# Sex-specific transcriptome dynamics of *Anopheles gambiae* during embryonic development

**DOI:** 10.1101/2024.12.24.630243

**Authors:** Agata Izabela Kalita, Eric Marois, Frank Rühle, Claudia Isabelle Keller Valsecchi

## Abstract

Malaria-transmitting mosquitoes are extremely sexually dimorphic in their anatomy and behaviour. Sex-specific gene expression in *Anopheles gambiae* is well-studied in adult stages, but its onset during embryogenesis, apart from sex-determination factors like *Yob*, remains largely unknown. Here, we report a comprehensive single-embryo transcriptome atlas of *A. gambiae* males and females to understand the earliest stages of establishing the sex-specific expression networks. Our dataset reveals embryonic RNA isoform diversity including a global shift towards distal alternative polyadenylation (APA) sites during the maternal-to-zygotic genome transition. Sex-biased gene expression and alternative splicing are limited during embryogenesis, with most sex-specific patterns emerging post-embryonically. X chromosome dosage compensation is established shortly after zygotic genome activation concomitant with direct binding of the master regulator protein SOA to X-linked promoters. Unlike the dosage compensation regulators in *Drosophila* or mammals, SOA operates in a one-step fashion, directly binding CA-rich promoter motifs without prioritizing certain gene groups over others. We propose that the *Anopheles* dosage compensation system represents an evolutionary endpoint of a gene-by-gene regulatory mechanism that evolved to a chromosome-wide scale.

## Introduction

In many species, males and females display differences in their physiology that are not limited to the gonads. For example, sex differences in adult insects comprise body size, ornamentation, coloration, and feeding behaviour, as only female, but not male mosquitoes are blood-feeding in order to produce eggs (Hopkins and Kopp 2021). This is relevant since the malaria-causing parasite *Plasmodium* is transmitted by a female mosquito after a previous blood meal taken on an infected individual. Understanding sex differences thus may have a big impact on the fight against malaria by inspiring new vector control strategies. Sexually dimorphic traits are determined by complex gene regulatory networks operating during the embryonic, larval and pupal developmental stages. The current understanding of the expression dynamics along *Anopheles* mosquito embryogenesis is based on a mixed-sex microarray dataset, which lacks information on sex-differences, non-coding transcripts, and splicing isoforms (Goltsev et al. 2009). Another study has investigated the transcriptome of male embryos in comparison with a mixed sex embryo population, limited to a single replicate per stage (Hall et al. 2016). Others (Papa et al. 2017; Hall et al. 2016) have studied sex-specific transcriptomes at post-embryonic stages (Rose et al. 2016; Jiang et al. 2015; Kalita et al. 2023). Thus, the sex-specific *Anopheles* gene regulation in embryos, especially at the stage where the embryo switches from maternal to zygotic transcription, is largely uncharacterized to date.

The initiation of sex-specific gene expression in the malaria mosquito *Anopheles gambiae* is conferred by the Y-linked maleness gene termed *Yob* (Krzywinska et al. 2016). *Yob* expression starts in early embryos at zygotic genome activation (ZGA) and triggers a largely unknown cascade of molecular events ultimately leading to the establishment of distinct phenotypes in males and females (Scali et al. 2005). Two recent studies (Kalita et al. 2023; Krzywinska et al. 2023) identified the *SOA/007* gene (hereafter *SOA*), encoding a male-specific DNA binding protein expressed after *Yob*, as the master regulator of *Anopheles* dosage compensation (DC). We showed that SOA protein associates specifically with the X chromosome. There, it mediates the upregulation of X chromosomal genes in XY males in order to match their expression levels in XX females (Kalita et al. 2023). How SOA binding is established during *Anopheles* development remains unknown to date. The striking process of collective gene regulation spanning an entire chromosome renders the study of the developmental onset and targeting principles of DC complexes particularly interesting.

The DC mechanism is well-characterized in another dipteran insect, the model organism *Drosophila melanogaster*. Here, this mechanism is established in a two-step fashion during embryogenesis (Samata et al. 2020; Prayitno et al. 2019; Lott et al. 2011). First, the master regulator of DC, the MSL2 protein, becomes expressed during the blastoderm stage, whereupon it binds to the X chromosome at around 300 high-affinity sites (HAS). In the second step, the multi-subunit MSL complex assembles and spreads to all X-linked genes. Through Histone H4 Lysine 16 acetylation along the entire X chromosome, the MSL complex mediates upregulation of expressed X-linked genes. Acetylation gradually increases over several hours during embryogenesis and some genes, e.g. developmental regulators and genes that are located further away from the HAS, get compensated with a delay. Notably, because of the property of 1) HAS-targeting and 2) spreading, the compensation mechanism can in principle act on any gene located on the X. Accordingly, when autosomal genes are artificially inserted on the X chromosome they can become targeted by the MSL complex and acquire DC (Gorchakov et al. 2009). However, in another dipteran, *Lucilia cuprina*, the DC mechanism cannot compensate transgenes and appears to be specific for genes native to the X chromosome (Linger, Belikoff, and Scott 2015). Although the DC mechanism in *L. cuprina* is not well understood, it suggests that DC in insects is not necessarily based on the same universal molecular mechanism.

In our characterization of SOA in mosquitoes at the pupal stage (∼10 days of development (Bayoh and Lindsay 2003)), we found that this protein associates directly with promoter regions of expressed genes, which is remarkably distinct from *Drosophila* MSL2. However, the dynamics of SOA binding, whether it initially targets HAS or if specific gene groups are particularly sensitive to its loss, are currently open questions.

Here, we use single-embryo RNA-seq to identify the sex-specific transcriptome and onset of DC mechanism along *Anopheles* embryogenesis. We characterize stage- and sex-specific RNA processing events and identify the expression onset of sexually dimorphic protein-coding and non-coding RNA transcripts. Genome-wide profiling of SOA by CUT&Tag in embryos and analysis of its loss reveal that 1) as in pupae, SOA directly localizes to promoters and 2) it readily establishes DC upon zygotic genome activation. Thus, we propose that the SOA pathway operates “gene-by-gene”, yet in a chromosome-wide fashion, as each X-linked promoter has had sufficient evolutionary time to accumulate SOA binding sites. Our atlas provides a framework to explore sex differences and X chromosome regulation in development and evolution and thereby, enables future functional analyses of sex-specific expression patterns and isoform usage.

## Results

### The developmental transcriptome of male and female *Anopheles* embryos

We isolated RNA from single *A. gambiae* embryos before and at eight different time points after ZGA as well as from neonate first-instar larvae (Fig. 1A). Because male and female embryos cannot be visually distinguished, we genotyped single embryos prior to generating libraries for RNA-seq (see methods and Fig. 1B, Fig. S1A). We then performed poly(A)+ RNA-seq followed by quality control (QC), including assessing the mapping rates, 3’-end bias indicative of degradation, library complexity or clustering (Supplemental Table S1). To the best of our knowledge, our atlas is the first RNA-seq dataset of single male and female embryos of any mosquito species. It represents a significant improvement over a microarray study on a mixed-sex *A. gambiae* population (Goltsev et al. 2009), which contains information on about 10’000 protein-coding genes (Fig. S1B) and a previous RNA-seq study (Hall et al. 2016), where only males in comparison with a mixed population (single replicate) were investigated.

**Figure 1.**
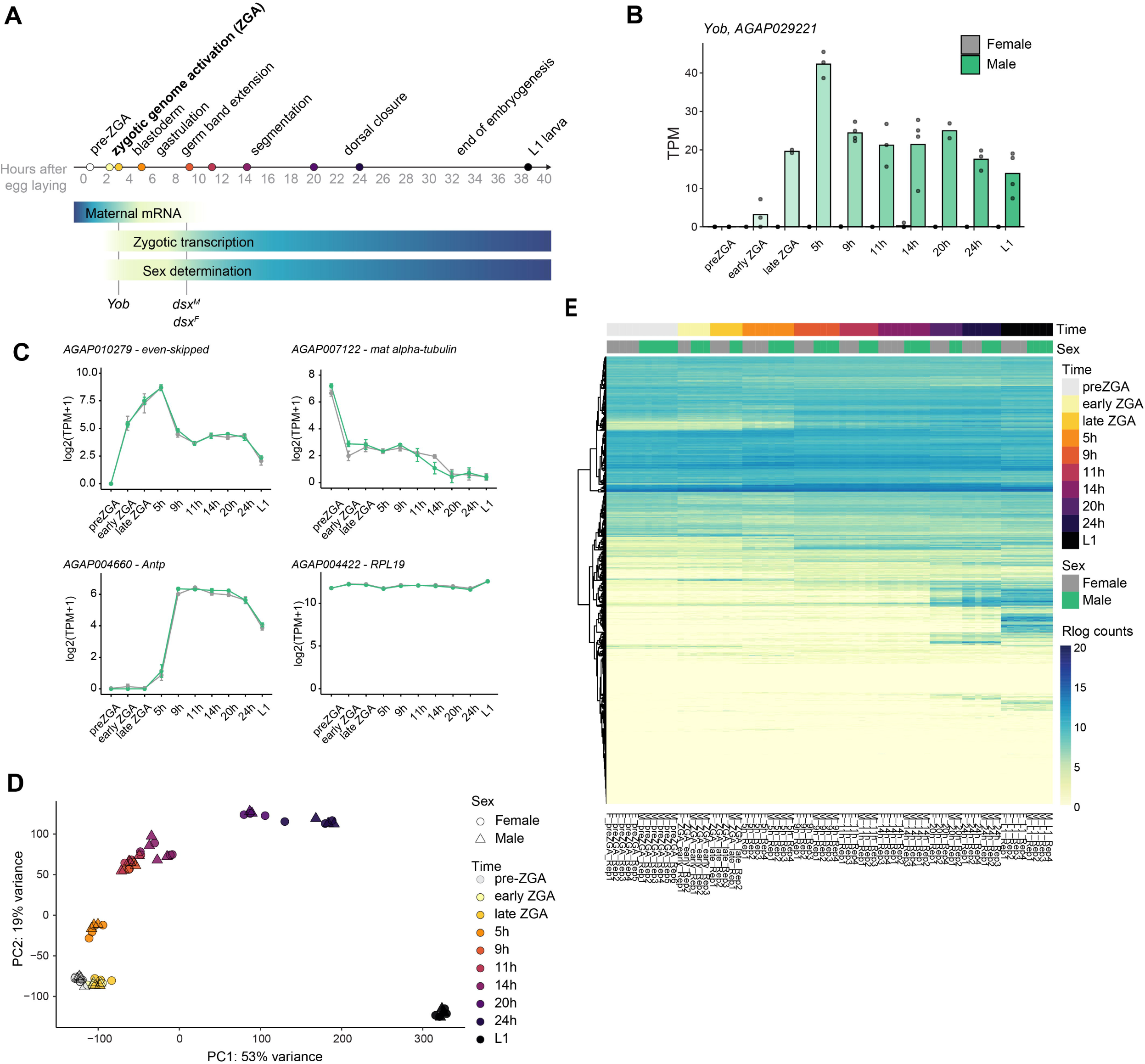
Single-embryo RNA-seq reveals developmental changes across embryogenesis. A) Scheme of *Anopheles gambiae* embryogenesis timing and respective developmental stages. The colored points on the timescale represent the times when samples were collected for RNA-seq. The gradient bar below depicts the approximate changes in the abundance of maternal versus zygotic transcripts, as well as the establishment of sex determination. The known sex determination cascade factors (*Yob* and *dsx*) and the start of their sex-specific expression/splicing is indicated below. B) Barplot representing the expression of *Yob* across embryogenesis in male versus female samples. The y-axis shows TPMs, individual points represent replicate values, while the height of the barplot is calculated as the mean of all replicates in the group. C) Expression patterns of the orthologues of known developmental regulators. The expression patterns in males are shown in green, for females in grey. D) A principal component (PC) analysis plot representing the PC1 and PC2 contribution to the variance in gene expression between samples. All expressed genes were used for determining the PC values. Male samples are represented as triangles and female samples as circles, with the color representing the developmental time point. E) Heatmap showing the developmental progression of gene expression. All genes are included, and the scale shows the regularized log-transformed expression values.

We mapped our data to a custom genome annotation (*n*=16,725 genes), which is based on the current AgamP4.14 Ensembl annotation, but additionally contains 3,171 putative long non-coding (lnc)RNA genes identified in (Jenkins, Waterhouse, and Muskavitch 2015). The additional genes included in our data display on average lower expression levels and higher variability across developmental stages compared with the gene set published in (Goltsev et al. 2009) (Fig. S1C-D).

To validate that our transcriptome accurately captures the intended developmental timepoints, we scrutinized genes with well-established dynamics in *Drosophila* and *Anopheles* (Goltsev et al. 2009, 2004). This confirmed the expected expression patterns of *RPL19* (constitutive), mat α-tubulin (maternal), *even-skipped* (early embryonic), *fushi tarazu* (*ftz*) and *hunchback* (segmentation, ZGA), as well as the Hox gene *Antennapedia* (gastrulation) (Fig. 1C, Fig. S1E). To further confirm that we have accurately captured the transition before and after ZGA, we investigated the expression patterns of *A. gambiae* orthologues of previously characterized *bona-fide* early zygotic *A. stephensi* genes (Wu et al. 2018), (Fig. S1F and methods). With some exceptions that were e.g. maternally deposited or not expressed at all, the *A. gambiae* orthologues showed a pronounced induction after ZGA (Fig. S1F).

When plotting the expression dynamics of all transcripts along the developmental time-points, we noticed that those are similar between males and females (Fig. 1D, 1E). A principal component analysis (PCA) showed that most of the variance in our dataset can be explained by differences between developmental stages rather than sexes (Fig. 1D). We thus focused first on analyses of developmental changes (for sex differences, see below) and used unsupervised clustering for gene classification according to their expression dynamics (16 clusters) (Fig. S2A). Those were then further grouped into 9 clusters based on whether they are stable or dynamic, and whether they are only maternal, only zygotic or both (Supplemental Table S2). We then investigated the groups for (1) regulatory patterns using motif enrichment analysis in promoters (400 bp upstream and 100 bp downstream of the annotated transcription start site) and (2) enriched gene functions using a gene ontology (SLIM-GO) enrichment analysis (Heinz et al. 2010) (Fig. S2B). Promoters of genes in Cluster B (exclusively early zygotic) are enriched in the Zelda binding motif, which is consistent with findings in *Drosophila* where Zelda is known to drive early zygotic transcription by binding these DNA motifs (Harrison et al. 2011). Trl/GAGA factor binding motif is enriched in Clusters C, D, and E (transcripts that are not maternally deposited and expressed in mid- and late embryos or larvae). Similar to *Drosophila* (X. Li et al. 2023), Trl/GAGA factor could establish faithful 3D organization for proper activation of these genes during embryogenesis. Other enriched motifs are typical of general transcription factors (TATA box or Initiator motif) and few clusters were also enriched for an E-box binding motif. The latter is noteworthy, because E-box transcription factors regulate circadian rhythm genes and oviposition in female *Anopheles* occurs preferentially at dusk. Therefore, those transcription factors could intrinsically ensure appropriate timing starting from egg laying to larval hatching.

The GO term analysis revealed expected patterns as e.g. “housekeeping” functions in the stable Cluster I, “muscle process” for the larval stage (Cluster E), “mitotic cell cycle” (mid embryo - Cluster G) or “ribosome biogenesis” (semi-stable, Cluster H). Cluster F, interestingly, is characterized by ’protein folding’ as a GO term specifically enriched in maternal and early embryonic stages, indicating that, prior to ZGA, maintaining a faithful proteome appears critical for supporting early developmental processes.

After having investigated global expression patterns, we turned our attention to individual genes. Besides the expected developmental genes (e.g. *hunchback*, see above), clustering also identified lncRNAs with dynamic expression patterns across embryogenesis. For example, the spliced lncRNA *merged.4615* in Cluster B is maternally deposited, but then becomes also actively and highly transcribed in the early embryo, reaching maximum expression in 5h embryos (Fig. S3A). Its expression pattern differs from the adjacent genes (*AGAP001238* and *AGAP001239*), suggesting active transcriptional regulation, and thus unlikely functioning by transcriptional interference with the neighboring genes. Tfsitescan prediction revealed a Grainy head binding motif in the upstream region of the gene (data not shown) and comparing the expression dynamics between the two implies that Grainy head could be a negative regulator of *merged.4615* (Fig. S3A). Based on synteny analysis, we found that this lncRNA is also highly expressed in *Aedes aegypti* embryos and *Anopheles albimanus* adults (Giraldo-Calderón et al. 2022). As expected for a lncRNA, *merged.4615* exhibited overall low sequence conservation, apart from two conserved stretches (25 nt at 88% identity and 23 nt at 65% identity across three mosquito species, Fig. S3B). This suggests that *merged.4615* could encode a precursor for a small RNA. Consistently, we observed two peaks with high read counts in publicly available small RNA sequencing experiments (Fig. S3C, (Biryukova, Ye, and Levashina 2014)). The peak heights decrease following Ago1 RNAi treatment in adult mosquitoes, indicating that these small RNAs are likely microRNAs.

Our study does not intend a functional exploration of these new transcripts and transcription factors, but these proof-of-concept analyses underscore the quality and usefulness of the first single-embryo transcriptome atlas of any mosquito species presented here.

### Developmental differences in alternative polyadenylation sites

Using RNA-seq can reveal isoform diversity generated through alternative splicing, transcription start site selection, and polyadenylation site variation. When investigating stage-specific processing events, we noticed that maternal transcripts (preZGA) often have shorter 3’UTRs compared to post-ZGA samples (Fig. 2A-C). To quantify and score these differences, we used APAtrap (Ye et al. 2018), revealing that 5,176 out of 13,100 expressed genes in any of our samples (>1 TPM) exhibit alternative polyadenylation (APA) sites (Supplemental Table S3). The analysis reveals a progressive shift in usage upon ZGA, with decreased proximal and increased distal APA usage (Fig. 2D). The most pronounced change occurs between pre-ZGA and early ZGA, with 1,245 genes showing significantly more distal APA usage, while only 94 show more proximal usage (p<0.01) (Fig. 2E). Distal APA usage results in elongated 3’UTRs, which affects 14% of all expressed genes at these stages (Fig. 2F). This suggests the existence of a global shift in 3’UTR site usage after ZGA. A GO term analysis of genes with significant APA change between preZGA to early ZGA samples revealed that these were enriched in biological processes related to chromatin remodeling, translation and protein folding (Fig. S4A, top). The molecular function of the genes was related to ATP and RNA binding, as well as protein kinase activity (Fig. S4A, bottom).

**Figure 2.**
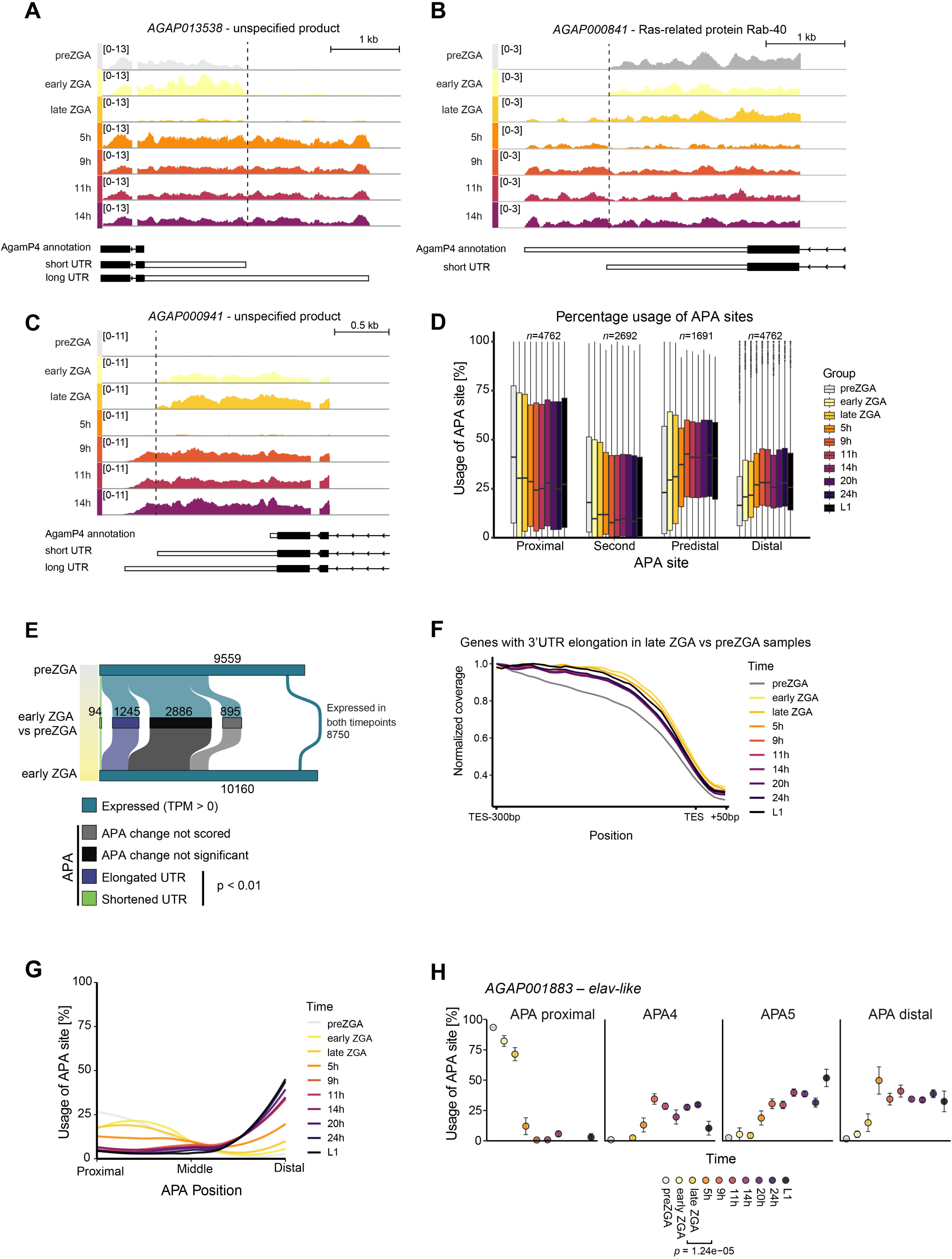
Transcripts gain a 3’UTR extension upon the zygotic genome activation A) Normalized coverage plot (CPM) of female samples across developmental timepoints for *AGAP013538.* The annotation scheme below shows the publicly available gene model (UTR in white outlined in black) and the most prevalent 3’UTR isoforms in embryos. B) as in (A) but for *AGAP000841*. C) as in (A) but for *AGAP000941*. D) Relative alternative polyadenylation (APA) usage across development. The *n* value above the plot indicates the number of plotted transcripts. The APA sites visualized are limited to four, i.e. proximal, second, predistal (second to last, excluding proximal and second) and distal. Some transcripts only have the proximal and distal sites annotated, resulting in different total transcript numbers for APA second and predistal. Only transcripts with more than one predicted polyadenylation site expressed in all timepoints are plotted. E) A scheme of changes in expression and APA usage from preZGA to early ZGA time point. In blue the number of genes that are expressed in respective stages. The middle bar of the plot shows the results of deAPA. In grey, genes where APA usage could not be scored because of filtering. In black, genes where APA usage was scored but there were no statistically significant changes in APA usage between the time points. In color: genes that show statistically significant (adjusted p-value < 0.01) change in APA usage: more distal (purple) or proximal (green) polyA sites in early ZGA samples as compared to preZGA samples F) A normalized coverage metaplot of genes with scored significant 3’UTR extension from preZGA to ZGA late embryos. A region surrounding the transcription end site (TES): 300 bp upstream and 50 bp downstream is depicted. The coverage is normalized to value at TES-300 bp as one for each transcript. Only female samples are plotted for brevity. G) Relative APA site usage from the most proximal to the most distal plotted together for the 23 genes that have been scored as significantly elongating their 3’UTRs and previously reported to have significant 3’UTR extension events in *Drosophila* neural development. H) Plot showing the % usage of *AGAP001883* (*elav-like*) APAs with adjusted *p*-value from deAPA. Only APA sites with relatively high usage are shown.

Previously, 3’UTR extension was reported for approximately 30 genes in the neuronal lineage along *Drosophila* embryogenesis (Hilgers et al. 2011). The majority of the *Anopheles* orthologues also show a 3’UTR extension in our dataset (23 significant by APAtrap, p<0.05), indicating that this process is conserved (Fig. 2G, 2H, S4B). This includes genes like *elav* and its paralogue, *fne*/*Rbp9*-like (Fig. S4B, S4C). In *Drosophila,* the RNA binding proteins encoded by *elav/fne* are positive regulators of 3’UTR extension by binding the proximal APA sites (Hilgers, Lemke, and Levine 2012; Wei et al. 2020; Carrasco et al. 2020). Besides the 23 genes with significant APA usage, we also found few examples where *Drosophila* and *Anopheles* showed different trends. The 3’UTR extension of *polo* (Pinto et al. 2011), or Hox cluster gene orthologs (*Ubx, abd-A, Abd-B,* and *Antp*) (Thomsen et al. 2010) was not scored in our *Anopheles* dataset (data not shown).

The prevalent 3’UTR extension upon ZGA observed here is consistent with recent discoveries in *Drosophila*, where analysis of 3’-seq datasets across tissues and developmental stages revealed the short 3’UTR as a feature of the maternal transcriptome and 3’UTR elongation as a zygotic signature (Lee et al. 2022). The conservation of 3’UTR extension during embryogenesis in *Drosophila* (Hilgers et al. 2011; Lee et al. 2022) and *Anopheles* (this study) suggests its functional importance, likely during the maternal-to-zygotic genome transition. A mechanistic exploration of this phenomenon will be best pursued in *Drosophila*, given its superior genetic and biochemical tools compared to *Anopheles*.

### Sex-biased gene expression and splicing along embryogenesis

#### Differential gene expression

Having analyzed the stage-specific transcriptome, we focused on the differences between sexes. For this, we first performed differential gene expression analysis between males and females (Supplemental Table S4, Fig. 3A). Excluding the late ZGA time-point (see next section on X chromosome DC), this revealed a total number of 111 sex-biased protein-coding genes (FDR<0.05), as well as 9 sex-biased putative lncRNAs (Jenkins, Waterhouse, and Muskavitch 2015). The sex-biased genes in embryos were significantly overrepresented on the X chromosome, especially among the female-biased transcripts, where approximately two thirds of the DE genes are located on the X (Fig. 3B). This is in line with earlier findings suggesting a “feminization” of the *Anopheles* X chromosome (Papa et al. 2017; Hahn and Lanzaro 2005), but could also be caused by inefficient DC in the initial stages of embryogenesis (see below). GO term analysis revealed an enrichment of biological processes such as “sex differentiation”, “RNA splicing” or “protein-DNA complex assembly” (Fig. 3C). Similarly, “channel regulator activity” or “protein-DNA complex” were enriched as GO molecular functions and GO cellular components, respectively (Fig. S5A). This is consistent with the view that sex-differentiation pathways are often exerted by a combination of sequence-specific transcription factors, splicing regulators and developmental regulatory proteins (e.g. membrane receptors) (Salz and Erickson 2010).

**Figure 3.**
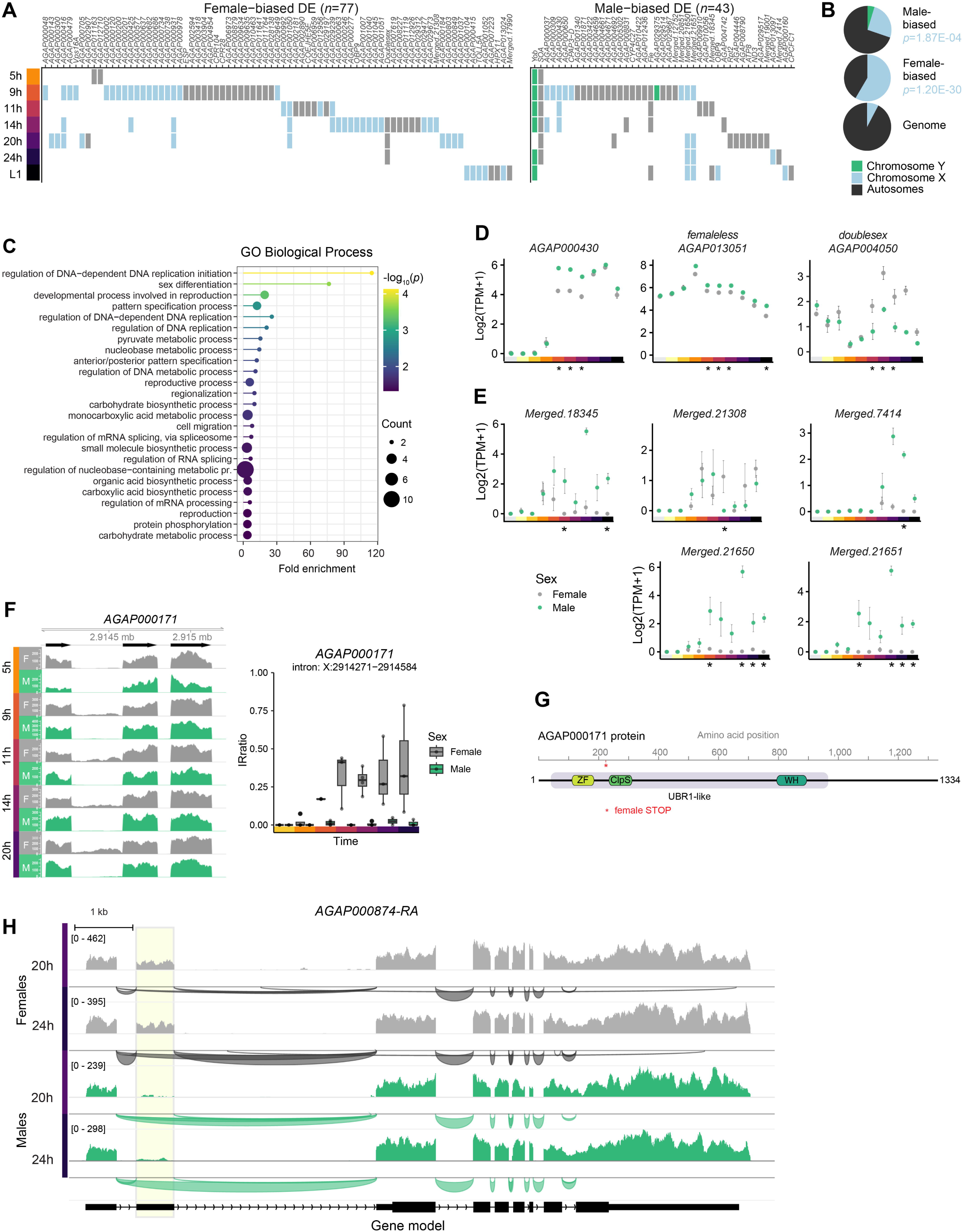
Sex-specific differences in gene expression and splicing. A) Tile plot of differentially expressed (DE) genes from 5 hour embryos onwards. The tile plot depicts the gene name on the x-axis, and the time on the y-axis. A filled tile indicates that a given gene is scored as differentially expressed at a given timepoint. The colors of the tiles represent the chromosomal location of the gene (grey: autosomal, green: Y chromosome, blue: X chromosome). B) Pie charts summarizing the chromosomal distribution of sex-biased genes, compared to the distribution of all annotated genes. The *p*-value is calculated using the Fisher’s test for overrepresentation (one-sided). C) A lollipop chart depicting the results of a Gene Ontology (GO) term analysis of sex-biased genes for the biological process ontology. The x-axis indicates the fold enrichment over random distribution. The color represents the -log10 (adjusted *p*- value), while the size of the circle is based on the number of genes in the set assigned to a given GO term. D) Plots showing expression in log2(TPM+1) of selected sex-biased genes across embryogenesis. The x-axis indicates the developmental time. The dots indicate the mean values of all the replicates, while the error bars show the standard error of the mean (SEM). The asterisk indicates timepoints at which the gene was scored as differentially expressed between the sexes by DEseq2 (FDR <0.05). E) as in (D) but for selected sex-biased non-coding genes across embryogenesis, F) Left: Coverage tracks of RNA-seq (all replicates merged in a given sex and timepoint). The screenshot shows intron 2 and surrounding exons of *AGAP000171.* The y-axis represents the number of reads in the bam file (not normalized). Right: Bar plot showing the sex-specific intron retention in *AGAP000171*, calculated with IRFinder. The y-axis represents the IRratio - the extent to which the intron is retained. Values from individual replicates are depicted over the barplot. G) InterProScan prediction of protein domains in AGAP000171. ZF - Zinc finger, UBR- type; ClpS - ClpS-homology domain; WH - Winged helix DNA-binding domain; UBR1-like - E3 ubiquitin-protein ligase UBR1-like protein family; STOP - stop codon resulting from the inclusion of intron 3. H) *AGAP000874* coverage and sashimi plots of RNA-seq with merged bam files generated from all replicates in a given sex and timepoint. The y-axis represents the number of reads in the bam file (not normalized), while the sashimi plot below shows the reads that span the exon-exon junctions.

Compared to the several hundreds of sex-biased genes present at adult stages (Papa et al. 2017), our embryonic dataset reveals a relatively small number of sex-biased genes. Notably, genes reported as fully sex-specific in adult somatic tissues (Papa et al. 2017) were either not expressed in embryos or were not sex-specific during embryogenesis (Fig. S5B), suggesting that most adult sexually dimorphic gene expression arises post-embryogenesis. Some genes identified as sex-biased during embryogenesis though remain biased in adults (Fig. S5C, S5D), likely reflecting common upstream regulators. These findings are not completely unexpected, since the morphological and behavioral differences of adult mosquitoes are dramatic compared to embryos, where males and females are phenotypically indistinguishable.

Among the sex-biased genes were the maleness gene *Yob* (*AGAP029221* (Krzywinska et al. 2016)), the DC master regulator *SOA* (*AGAP005748)* (Kalita et al. 2023), *Doublesex* (*AGAP004050)* (Scali et al. 2005) and *femaleless* (*Fle*, *AGAP013051*) (Krzywinska et al. 2021) (Fig. 3D). *Fle* has been proposed to be a regulator of DC and sex determination, responsible for establishing the female splicing pattern of *doublesex.* Interestingly, although *femaleless* plays a crucial role in female sexual development, we scored *femaleless* expression as male-biased (Fig. 3D, middle). A recent study, however, proposed a posttranscriptional regulation of *Fle* (Krzywinska et al. 2023), whereby the transcript but not the protein is expressed in both sexes. One example of a novel sex-biased gene is *AGAP000430*, which was scored as DE at 9 to 14 hours after egg laying (Fig. 3A, D). *AGAP000430* encodes a C2-domain containing membrane protein, whose *Drosophila* orthologue Synaptotagmin 14 is involved in nervous system development and synapse function. *AGAP000430* is also expressed in a male-biased fashion at the adult stage (Fig. S5E, (Papa et al. 2017)). It is possible that *AGAP000430* expression-bias contributes to the generation of male-specific neurons, for example to initiate male-specific behavioral patterns. Apart from protein-coding genes, there were also lncRNAs scored as differentially expressed between the sexes (Fig. 3E). Notably, despite extensive searches and *de novo* transcript discovery approaches, we were unable to find any X-linked lncRNA transcript akin to *roX1/2* or *Xist* in one of the two sexes.

#### Alternative splicing

Alternative splicing plays a critical role in sex determination and differentiation across development by enabling the generation of sex-specific transcripts from a shared genome. To investigate this, we conducted splicing analyses with rMATS (data not shown) and IRFfinder, which can identify differential intron retention events (Middleton et al. 2017). Most alternative splicing or intron retention events were misidentified due to overlapping transcripts, unannotated UTRs, or low transcript coverage, highlighting the challenges of studying alternative splicing in non-model organisms at understudied developmental stages. Compared to the relatively large extent of sex-specific splicing in adults (e.g. the ones conserved among four *Anopheles* species (Papa et al. 2017)) we found very few alternative splicing events in our dataset, suggesting that most of the adult patterns are established post-embryogenesis (data not shown).

However, careful examination of all statistically significant events revealed a few with potential biological relevance. The most striking example revealed by IRFfinder was *SOA* (*AGAP005748),* which we previously reported to produce two protein isoforms: a truncated, non-functional isoform in females, as well as a male full-length isoform capable of inducing DC (Kalita et al. 2023). Apart from *SOA*, we also found a previously unreported female-specific retention event in *AGAP000171* intron 3 (Fig. 3F) affecting a subset of the *AGAP000171* transcripts. The retained intron 3 contains an in-frame premature stop codon, resulting in a protein truncation after 224 amino acids (Fig. 3G). This is predicted to produce a protein lacking critical domains of the UBR-1-like E3 ubiquitin ligase encoded by *AGAP000171*. Comparison to publicly available datasets revealed that the female-specific intron retention remains substantial in adult females, especially in antennae (SCAN 2024; Giraldo-Calderón et al. 2022; Pitts et al. 2011) (data not shown).

Another example of a gene with sex-specific isoforms is *AGAP000874* (Fig. 3H). The gene is a duplication of *AGAP000875*, which encodes a protein predicted to be involved in transcription regulation. The gene is not expressed until 20 hours into embryogenesis, at which point exon 2 is spliced out in males, but not females. The second exon is part of the 5’UTR, indicating that the sex-specific splicing does not affect the coding potential of the transcript, but might result in changed translation efficiency or transcript stability.

Together, these findings highlight the minor yet noteworthy role of sex-biased gene expression and alternative splicing in establishing sex-specific differences during embryogenesis. The fact that most sex-specific splicing patterns emerge post-embryonically, likely reflects the increasing complexity and phenotypic manifestation of sex-specific functions in later developmental stages.

### The onset of DC during embryogenesis

One gene regulatory process relevant to sex differences is DC and we next investigated its establishment, particularly when zygotic transcription begins. Before Zygotic Genome Activation (ZGA), all transcripts are maternally deposited and therefore equally expressed between the sexes (no differentially expressed genes, median log2FC for the X chromosome 0.0735, Fig. 4A, B). Early at ZGA (around 3 hours after oviposition), expression is still balanced between the sexes and between chromosomes. Shortly thereafter (late ZGA, 3.5 hours) we detected a striking imbalance because males express X-linked genes only from a single copy, compared to the double dose in females and autosomal genes, respectively (Fig. 4B, C). When comparing males to females, this translates to a log2FC of -0.8859 for the X-linked genes, meaning that the level of X-linked genes in the males is 53.74% of the level in females. 189 out of 541 DE genes were X-linked (significant overrepresentation, *p*=1.80E-77, Fig. 4C). In 5 hour embryos, this imbalance is already partially compensated (log2FC -0.2044, 86.8%) with further fine-tuning later in development. This corresponds to the appearance of the master regulator of DC - the male-specific SOA isoform (Fig 4D), whose mechanistic and physiological roles we have dissected earlier in (Kalita et al. 2023). To investigate if there is a difference in the timing and extent of DC between different gene groups, we turned back to the developmental clusters (Fig. S2B). Comparing the log2FC between males and females revealed that X-linked genes in clusters with maternal contributions show less expression imbalance compared to genes in clusters consisting of purely zygotic genes, suggesting that maternal factors help mitigate DC imbalances during early development (Fig. 4E). Of note, the chromosomal distribution of genes within clusters is not random (Fig. 4F). Comparing the chromosomal distribution of all clustered genes, maternal and early-embryo clusters show a paucity of X-linked (e.g. Cluster B - early embryo: only 10 out of 290 genes are X-linked). Because DC is not yet established at this early developmental stage, X chromosomal location could cause male-female differences for an early embryonic gene expressed at this stage. In *Drosophila*, early developmental regulatory X-linked genes such as *giant, vnd, nullo, btd, tsg,* and *sog* are compensated in a non-canonical fashion despite the absence of MSL-H4K16ac (Lott et al. 2011). *Anopheles* apparently solves this issue in a different way by encoding at least some of these genes on autosomes (e.g. *giant, vnd)* (Fig. S5F). Others show distinct expression dynamics compared to *Drosophila* (Fig. S5G). They are either maternally deposited, or expressed later in development, when DC is established. Both solutions mitigate the risk of imbalance.

**Figure 4.**
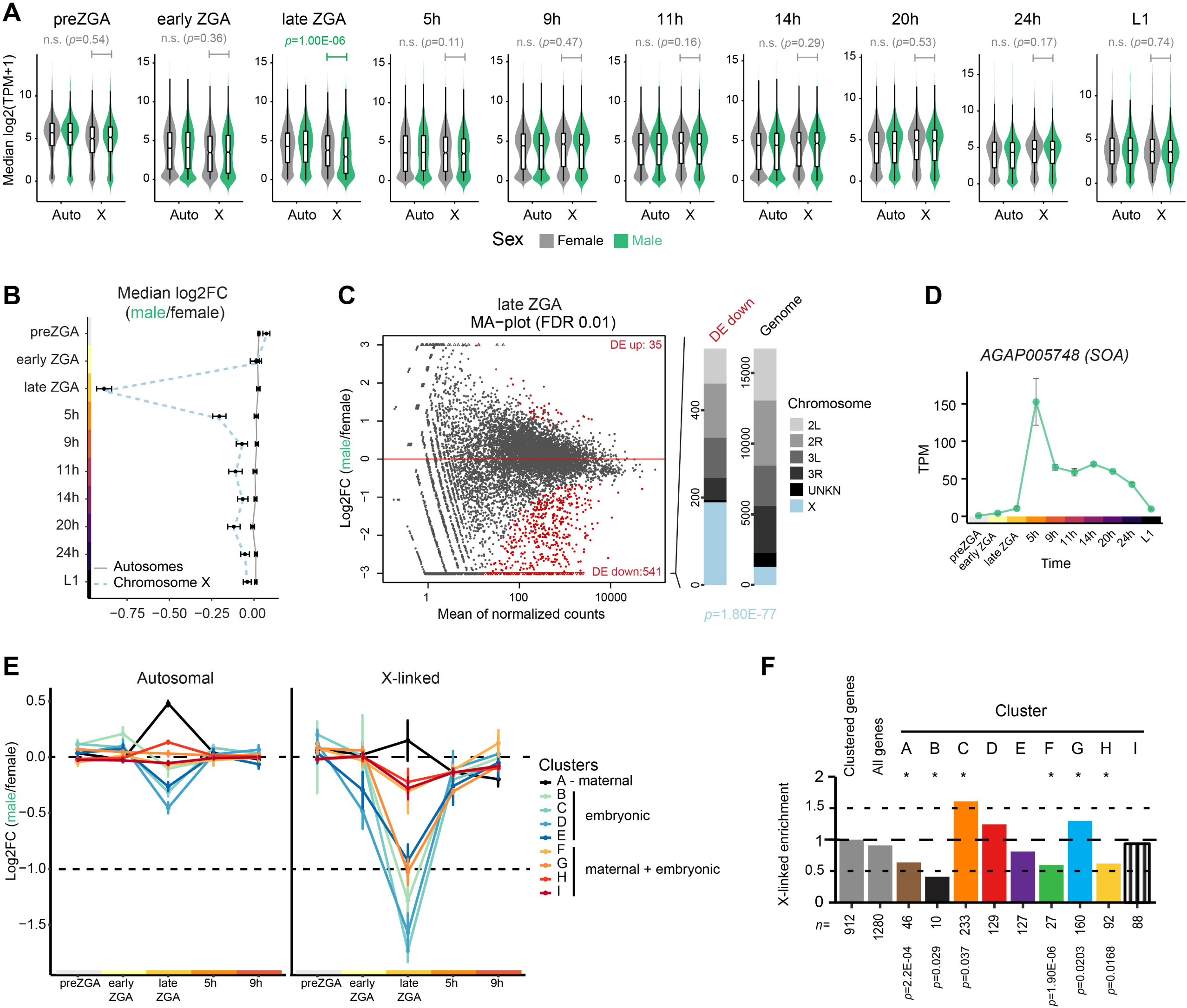
Sex chromosome dosage compensation along embryogenesis. A) Violin and box plot of the expression (log2TPM+1) of X-linked and autosomal genes in male and female embryos at different timepoints. The *p*-values above the plots are calculated with a two-sided Wilcoxon rank sum test with continuity correction. B) Plot of log2FC between males and females as calculated by DEseq2, grouped into X-linked and autosomal genes. The dots indicate the median value, with the error bars calculated as the standard error of the median. C) Left: MA plot of DEseq2 results for male to female comparison at late ZGA timepoint. The y-axis shows the log2FC value between males and females, the x-axis shows the mean of normalized counts. Differentially expressed genes (FDR<0.01) are colored in red, the remaining genes are in grey. Right: Chromosomal distribution of genes with significantly lower expression in males than in females, compared to the chromosomal distribution of all genes. The *p*-value below the plot is calculated with a one-sided Fisher’s test for overrepresentation. D) Plot showing expression (TPM) of *SOA* in the males across embryogenesis. The dots represent the mean values of all the replicates, while the error bars show the standard error of the mean (SEM). E) Plot of log2FC between males and females as calculated by DEseq2. X-linked genes are separated into previously assigned clusters (Fig. S2B). The dots indicate the median value, with the error bars calculated as the standard error of the median. F) Enrichment of X-linked genes in the developmental clusters (Fig. S2B) as compared to all clustered genes (mean TPM>5 at any timepoint). The numbers below the axis indicate how many X-linked genes were assigned to a given cluster. The asterisks above the plot mark the clusters with significant deviation from the expected distribution, as calculated with the Fisher’s test with exact *p*-values provided below.

We showed earlier at the pupal stage that the SOA protein binds the promoters of X-linked genes and its binding is necessary for achieving DC (Kalita et al. 2023). To examine the onset of DC, we analyzed the SOA binding dynamics during embryogenesis by generating SOA CUT&Tag datasets from pooled embryos across three developmental stages, using a homozygous *SOA* null mutant line (*SOA-KI*, extensively characterized in (Kalita et al. 2023)) as a negative control (Fig. S6A). Our earliest time-point captured 3-4 hour embryos, where - consistent with low SOA mRNA expression (Kalita et al. 2023) (Fig. 4D) - we did not observe the characteristic SOA binding pattern. Although few peaks were scored as statistically significant by DiffBind, we found that (1) they were not localized to the X chromosome, (2) they did not show the characteristic SOA CA-rich binding motif and (3) their coverage in later timepoints was similar between WT and *SOA-KI* mutant samples (Fig. 5A). We thus favor the idea that these peaks, rather than being early binding sites, reflect technical artifacts such as phantom peaks (Jain et al. 2015). In contrast, SOA binding at X-linked gene genes was prominent in 6-7 hour embryos (Fig. 5A, 5B), which is consistent with onset of DC accompanied by high expression of SOA transcript starting from 5 hour embryos (Fig. 4D). The SOA binding in 16-20 hour embryos showed a similar pattern to that observed in 6-7 hour embryos, albeit with a slightly weaker signal (Fig. 5A), likely reflecting a reduction in SOA expression levels following the critical window for DC establishment (Fig. 4D). As in pupae, most of the SOA peaks at these two timepoints were located at promoter regions (Fig. 5C).

**Figure 5.**
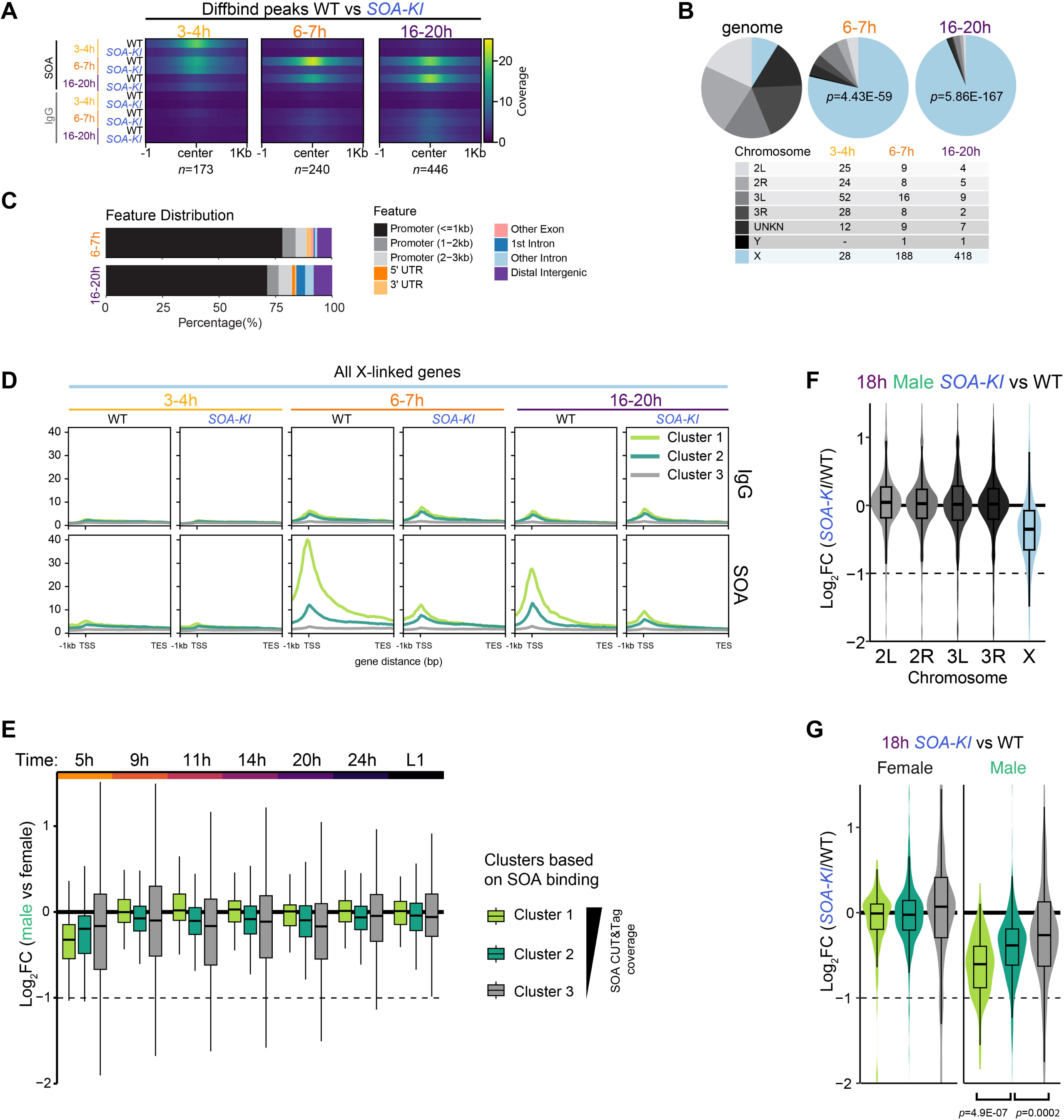
SOA binding across embryogenesis and its effect on gene expression. A) Heatmaps comparing mean SOA antibody and IgG CUT&Tag coverage in peak sets with significantly higher signal in WT compared to *SOA-KI* embryos (scored by DiffBind) in 3-4 hour embryos (left), 6-7 hour embryos (middle) and 16-20 hour embryos (right). Significant peaks from all chromosomal arms are shown (numbers of peaks in each group below the plots). The signal is shown ± 1 Kilobase (Kb) from the peak center. B) Chromosomal distribution of peaks with significantly higher signal (scored by DiffBind) in wild-type (WT) compared to *SOA-KI* embryos. The *p*-value is calculated with a one-sided Fisher’s test for overrepresentation of X-linked peaks. Table below the plots shows the peaks assigned to each chromosome in each set. UNKN, scaffolds that are not assigned to any chromosome. C) Bar plot of annotations of X-linked SOA peaks for genomic features. UTR, untranslated region. D) Metaplot of IgG control and SOA antibody CUT&Tag coverage at 3 random k-means clusters generated from all X-linked genes. TSS - transcription start site, TES - transcription end site. E) Box plots showing the log2FC across embryonic development on all X-linked genes grouped by unsupervised *k*-means clustering in 3 groups with strong, medium and weak SOA binding strength, see (D). F) Violin and box plots show the DESeq2-obtained log2FC in WT compared to homozygous *SOA-KI* mutant 18h male embryos. All expressed genes (mean read counts across all replicates > 0) are plotted. The genes are grouped by chromosomal location. Plot for female samples can be found in Fig. S6E. G) Violin and box plots show the DESeq2-obtained log_2_FC in 18h embryos (homozygous *SOA-KI* mutant compared to WT). All expressed X-linked genes (mean read counts across all replicates > 0) are plotted. The genes were separated by unsupervised *k*-means clustering in 3 groups with strong, medium and weak SOA binding strength, see (D). *p*-values obtained from two-sided Wilcoxon test with Bonferroni correction.

Next, we used *k*-means clustering to separate all X-linked genes into three groups based on their SOA binding strength (Fig. 5D). The genes with the strongest SOA binding by CUT&Tag were the most imbalanced in RNA-seq in 5 hour embryos (Fig. 5E). However, this group was fully compensated by 9 hours after oviposition, while establishment of DC was more gradual for the remaining two groups with weaker SOA binding. Like in the pupa stage (Kalita et al. 2023), the genes with strongest SOA binding in embryos displayed overall higher expression compared to genes with intermediate or weak SOA binding (Fig. S6B). To understand if SOA binding varies between X-linked genes with different developmental dynamics, we re-examined the expression clusters based on RNA-seq (Fig. S2B, S6C) and plotted the SOA CUT&Tag enrichment at their TSS (Fig. S6C). Genes with rather stable expression patterns (clusters G,H,I) exhibited the strongest SOA binding. Late embryonic and larval genes showed no SOA binding, which suggests that SOA binding co-occurs with the activation of a gene, rather than preceding it. Intriguingly, the few early zygotic X-linked genes had little SOA binding, possibly because of the very transient nature of their expression induction at approximately 5 hours into development (before the 6-7 hour timepoint sampled by CUT&Tag).

Overall, these data indicate that specificity of SOA to X-linked gene promoters is achieved immediately with the onset of the corresponding gene’s expression. This contrasts with the progressive binding by the MSL complex and DC establishment over different gene groups along *Drosophila* embryogenesis (Samata et al. 2020; Prayitno et al. 2019).

#### Inefficient DC causes sex-biased expression

Finally, we wanted to examine the extent of expression imbalance in embryos lacking SOA. For this purpose, we conducted RNA-seq on WT and *SOA-KI* embryos from both sexes (Fig. S6D). As expected, the expression of X-linked genes was substantially reduced in *SOA-KI* male embryos compared to wild-type (Fig. 5F, S6E, F) and similar to the pupa stage (Kalita et al. 2023), this effect depends on SOA binding strength (Fig. 5G).

Because female-biased genes tend to be overrepresented on the X-chromosome (Fig. 3A, B), and SOA binding appears to fine-tune RNA expression quite acutely (Fig. 5E), we wondered whether the female-biased expression of X-linked genes could be explained by inefficient DC. SOA binding, as assessed by CUT&Tag, is notably weaker at female-biased X-linked genes compared to unbiased genes (Fig. S6F). This is also reflected by the extent of imbalance scored after perturbation of the SOA pathway: The expression of X-linked female-biased genes is less affected in either *SOA-KI* mutant males or females ectopically expressing SOA (*SOA-R*) compared to unbiased genes. (Fig. S6G). This suggests that the female-bias of X-linked genes arises from weaker upregulation in males due to “resistance” to SOA-mediated DC.

## Discussion

Our single-embryo sex specific RNA-seq dataset provides significantly more information compared to previously published embryonic transcriptomes. First, we provide sex-specific data across multiple timepoints, increasing our understanding of the establishment of DC across development and the onset of sex-specific expression patterns. Additionally, profiling embryos by RNA-seq allowed us to understand the features of the embryonic transcriptome beyond the mere expression levels, for example identifying the differential usage of APA sites across development. Moreover, analyzing single embryos enabled a detailed understanding of rapid developmental transitions that are often overlooked in bulk collections of embryos. Consequently, this dataset offers a valuable resource for e.g. evolutionary studies, enabling direct comparisons of embryonic transcriptomes across insect species.

The cases of 3’UTR extension in our dataset include previously reported transcripts (Hilgers et al. 2011). However, apart from these striking extension events, in our analysis we have also observed more than a thousand genes having their 3’UTRs extended upon ZGA. This suggests this regulation mechanism to be a general ZGA feature, rather than specific to a small subset of neuronal regulators. Recently, the global shift in 3’UTR length upon maternal-to-zygotic (MTZ) transition has also been observed in *Drosophila (Lee et al. 2022)*.

Our data implies that this feature might be conserved through Dipterans or even more insect groups. Of note, dynamic changes in 3’UTR length have also been shown to play a role in zebrafish MZT (Y. Li et al. 2012; Mishima and Tomari 2016).

This 3’ UTR extension phenomenon may be linked to the rapid developmental changes occurring in the embryo. Although 3’UTRs are non-coding sequences, their length could affect transcript stability, translational efficiency and availability of miRNA binding sites. The increased 3’UTR length allows inclusion of additional binding sites for regulatory proteins and miRNAs that can control gene expression post-transcriptionally. This mode of gene regulation is more rapid compared to transcriptional regulation, which might be an especially important feature for embryonic development (Chen et al. 2014). The prediction of APA sites in our analysis relied solely on RNA-seq data. Such computational prediction can have its limitations; however they allow for insights into APA usage in absence of 3′-end-seq information in a species of interest. Improved annotation of mosquito 3’UTRs would aid in correct prediction of regulators acting on a given transcript (e.g. microRNA binding prediction or RNA binding protein motif search). Further investigation of this feature will require 3’-seq mapping to unambiguously annotate APA sites and spatially resolved transcriptomics to understand whether this phenomenon is more prevalent in certain tissues of *Anopheles* embryos.

Sequencing of male and female embryos separately allowed us to discover genes which are expressed in a sex-biased manner in early embryogenesis. Compared to adult mosquitoes (Papa et al. 2017), the sex bias in expression and splicing is modest in embryos, but includes key upstream regulators of sex-specific development (e.g. *Yob, SOA, doublesex, femaleless*). This reinforces the value of analyzing sex-specific gene expression during early developmental stages, where key regulatory factors can be identified without being obscured by the extensive number of sex-biased genes later in development (Kalita and Keller Valsecchi 2024). To get further insights into the gene-regulatory dynamics during the MZT and associated timing of DC in this species, which is apparently very rapid, it will be necessary to design experiments using crosses of diverging strains (Liang and Sharakhov 2019). In their hybrid offspring, any transcripts aligning to only the paternal genome must be of zygotic origin. Additionally, quantification of transcript abundance, e.g. by single-molecule RNA-FISH, can facilitate future investigations, especially because it provides an additional, spatial resolution that bulk RNA-seq cannot provide.

Analysis of our dataset together with SOA CUT&Tag binding profiles allowed us to gain deeper insights into the dynamic onset and mechanism of DC. There is no other, consistent sex-specific putative SOA co-factor in our dataset. This indicates that SOA may co-opt a conserved, non sex-specific machinery to regulate its target genes. We also did not find any evidence for a X-linked lncRNA akin to *Drosophila roX1/2* (Meller and Rattner 2002), Anolis lizard *MAYEX* (Tenorio et al. 2024) or mammalian *Xist* (Brockdorff et al. 1992), although it is formally possible that such a sex-specific transcript is non-polyadenylated (Pan, Darras, and Keller 2024), and thus, not captured by our dataset. However, our CUT&Tag data in embryos reveals no initial high affinity binding sites. Instead, the SOA binding pattern is established rapidly between 3-4 hours and 6-7 hours in embryogenesis and its direct association with X-linked promoters is the main predictor of DC efficiency. The onset of DC via promoter binding during *Anopheles* embryogenesis is rapid and precise. Because we consider it unlikely that a specific RNA species aiding recruitment is produced at each compensated gene individually, we favor the idea of a DNA-guided mechanism for each X-linked gene. This contrasts with the progressive establishment seen in *Drosophila (Samata et al. 2020; Prayitno et al. 2019)* that correlates with accumulation of roX1/2 RNA. Our findings imply that each promoter would need to evolve SOA binding elements, which remains to be proven using transgenic tools.

Independent of this experimental validation, which is challenging in non-model organisms such as *Anopheles*, this direct binding mechanism of SOA is further supported when analyzing distinct gene groups: early in development, maternal contributions appear to partially mitigate X-linked imbalances, evidenced by a lower imbalance in genes with maternal deposits compared to purely zygotic genes. Interestingly, maternally deposited and early zygotic genes seem to be depleted from the X chromosome, providing *Anopheles* a solution to mitigate imbalance due to X dosage differences before DC starts. This gene-specific role of SOA in inducing upregulation is also reflected in the group of female-biased X-linked genes, which exhibit SOA resistance in males. Thus, SOA shapes sex-specific gene regulation beyond DC.

More generally, an extensive discussion of the *Anopheles*-specific developmental patterning programs is provided in (Goltsev et al. 2007) but our data may further help to expand our understanding of insect embryo development beyond *Drosophila*. For instance, unlike *Drosophila*, mosquito embryos develop on water surfaces, exposing them to fluctuating temperatures and water conditions, which likely influence gene regulatory networks and developmental timing. Comparing these adaptations in *A. gambiae* and *D. melanogaster* can provide insights into how environmental factors shape developmental pathways and evolutionary differences between different insect embryos.

## Materials & Methods

### Mosquito rearing & staging

*A. gambiae* mosquitoes were maintained in standard insectary conditions (26–28°C, 75– 80% humidity, and 12-/12-h light/dark cycle). Egg development was induced by blood feeding and egg laying started by presenting a wet egg dish for approximately 20 minutes to the mosquito females. Individual eggs were collected at designated time points after presenting the egg dish (note that egg laying and fertilization in mosquitoes are synchronized). Because the dipteran development and transcriptome at zygotic genome activation (ZGA) undergoes very rapid changes, the 20 min collection time-window was not sufficient to appropriately stage ZGA. For the ZGA timepoint, we therefore sampled a total of 10 embryos, which were then allocated to early and late ZGA based on data from (Wu et al. 2018). For this, the *A. gambiae* orthologues were retrieved on Ensembl Biomart and the embryos were clustered based on the expression levels of the early ZGA genes. In addition, we manually inspected several early zygotic genes based on (Goltsev et al. 2009) and allocated the embryos to early and late timepoints, respectively.

#### Time course RNA isolation, library generation and sequencing

RNA was extracted using Trizol (Fisher Scientific, Cat. No.15-596-026) and the Direct-zol RNA MicroPrep Kit (Zymo Research, Cat. No. R2062). For preZGA samples, only the aqueous phase formed after phenol-chloroform separation was loaded on the column after mixing with 100% EtOH. The phenol-chloroform phase was used for DNA isolation following the manufacturers’ instruction manual (Fisher Scientific, Cat. No.15-596-026). RNA integrity was assessed with High Sensitivity RNA ScreenTape (Agilent, 5067-5579, 5067-5580). The library preparation was performed with Illumina’s Stranded mRNA Prep Ligation Kit following Stranded mRNA Prep Ligation ReferenceGuide (June 2020) (Document # 1000000124518 v00) using 13 PCR cycles. The samples were sequenced in paired-end mode, 2 x 75 on an Illumina HiSeq 4000 or a NextSeq500 (see Supplemental Table 1 for details).

### Genotyping

pre-ZGA embryos were genotyped by PCR on DNA isolated using the Trizol-chloroform method for RNA and DNA isolation from one lysate. PCR was performed with FastStart Universal SYBR Green Master (ROX) mix (Roche, 04913850001) using Y-linked repeats (S23 (Krzywinski et al. 2004)) as template and autosomal repeats (Ag404 (Krzywinski, Sangaré, and Besansky 2005)) as control. In brief, DNA and RNA were extracted simultaneously from single embryos collected before zygotic genome activation (ZGA). The DNA fraction was then subjected to quantitative PCR (qPCR), where a Y chromosomal satellite repeat was amplified allowing to identify males (Krzywinski et al. 2004). For embryos after the ZGA stage, the RNA levels of *Yob*, a Y-linked sex determination gene, were determined by subjecting 10% of the isolated RNA to reverse-transcription RT-qPCR (RNA-seq confirmation of correct sexing in Fig. 1B). post-ZGA samples were genotyped based on expression of Yg2 (i.e. Yob) mRNA by qPCR with Rp49 as a control. qPCR was performed on a Roche LightCycler II using FastStart Universal SYBR Green Master (04913914001; Roche) in a 7 μl reaction at 300 nM final primer concentration. Cycling conditions were as recommended by the manufacturer.

### Data processing and visualization

Reads were trimmed with cutadapt (1.18). To filter out rRNA reads, they were mapped to the ribosomal RNA sequences extracted from Ensembl AgamP4.14 genome using the Ensembl AgamP4 annotation (release 48) with STAR (v. 2.7.3a) with the following parameters: outFilterMultimapNmax 1000000 outFilterMismatchNoverLmax 0.04 outFilterMismatchNmax 999. Unmapped reads were used in downstream processing. Reads were mapped to the Ensembl AgamP4 genome using the Ensembl AgamP4 annotation (release 48) together with lncRNA annotation from (Jenkins, Waterhouse, and Muskavitch 2015) with STAR (v. 2.7.3a) using the following parameters: outFilterMismatchNoverLmax 0.04 outFilterMismatchNmax 999. Only uniquely mapped reads were used for downstream analysis. Coverage signal tracks (bigWigs) of primary alignments were generated using Samtools (v1.9) and DeepTools (v3.1.0). Primary alignments were assigned to features using subread 1.6.5 with the AgamP4 annotation (release 48) combined with lncRNA annotation (Jenkins, Waterhouse, and Muskavitch 2015) as a reference. Differential expression analysis was performed using DESeq2 (1.26.0) and only genes with FDR<0.05 in were considered as differentially expressed. To compare our transcriptome atlas to the microarray data generated by (Goltsev et al. 2009), we downloaded the expression data provided in Supplemental File 6 and converted the AGAP GeneIDs to the current annotation available on vectorbase.org (Giraldo-Calderón et al. 2022) (“Searches” Tab). Genome browser snapshots were generated with IGV (https://software.broadinstitute.org/software/igv/) (Robinson et al. 2011).

### ZGA orthologues

A stringent list of early zygotic transcripts described in *A. stephensi* was used to find their orthologues. Out of 71 genes, 49 have assigned orthologues in *A. gambiae*. Next, after visual inspection, genes with expression in preZGA samples were filtered out from the list (*AGAP006073, AGAP009974, AGAP002361, AGAP029670, AGAP007651, AGAP00369*).

Additionally, X-linked genes which would presumably differ in their expression between sexes upon ZGA due to lack of DC at this timepoint were excluded (*AGAP000819, AGAP000215, AGAP000014, AGAP000317, AGAP000714*), resulting in a final list of 38 autosomal genes.

### Clustering analysis

Gene expression was normalized to TPMs (Transcripts per Million). We calculated the mean gene expression for a given time point (combining both male and female samples). Next, only genes with mean expression per group of more than 5 TPMs at any time point were selected to avoid noise in our clustering analysis, resulting in 10’841 genes passing the filtering. The expression profiles were scaled from 0 to 1, where 1 is the highest expression of a given gene across all time points. We used the R package kohonen (3.0.12) (Wehrens and Kruisselbrink 2018) to construct self-organizing maps, resulting in expression profile clustering into 16 groups. Next, we inspected these clusters and assigned them a biologically relevant category based on the timing of the highest expression.

### HOMER motif enrichment analysis

The promoters of manually grouped clusters were analysed using the HOMER motif discovery tool. In brief, regions 400 bp upstream and 100 downstream of the annotated transcription start site were input to find enriched motifs of length 6,8, or 10 bp from the insect motif database. Homer 1 was used to only search through annotated, known motifs.

### APA trap

We analyzed all embryogenesis timecourse RNA-seq samples to identify the APA sites and their usage. First, we used bedtools (2.29.2) function genomeCoverageBed to generate the bedGraph files of our RNA-seq alignments. Next, we used the identifyDistal3UTR script from APAtrap (Ye et al. 2018) with bedGraphs of all the files and a bed file of gene locations generated based on the Ensembl AgamP4 annotation (release 48) as input. We used the default settings for genomes with short UTRs (50 window size and 5000 pre-extension 3’UTR size). Next, using a bed file with newly identified 3’UTR coordinates we ran predictAPA for all samples together, grouped by sex and time, using the following settings: coverage drop cutoff 0.2, coverage threshold: 20, APA min dis: 100, scan window size: 50. After inspection revealed little difference between the sexes, we assigned all the samples at a given time point to one group, regardless of sex. We then ran deAPA to score differential usage of APA sites between the timepoints, setting the minimum number of replicates showing APA change to 3. This allowed us to identify transcripts that change APA usage during development, which we identified based on the adjusted *p*-value < 0.01. In Figure 2G, we plot the smoothed average of APA usage for the 23 orthologues of genes previously identified in Drosophila melanogaster as elongating their 3’UTRs. Since genes have a different number of APA sites, the approximate APA usage across genes is calculated with a polynomial regression.

### CUT&Tag library generation and sequencing

Mixed-sex embryos from homozygous wild-type and *SOA-KI* loss-of-function lines were collected at the following timespans after egg laying: 3 to 4 hours, 6 to 7 hours, or 16 to 20 hours. Embryos were flash-frozen and stored at -80L. After thawing, embryos were homogenized in DPBS and filtered through a strainer. Cells were washed twice with PBS and resuspended in Wash Buffer. After DAPI staining, the cells were counted and adjusted to 100’000 cells per reaction (for 3-4 hours samples) and 400 000 cells per reaction (6-7 hours and 16-20 hours samples). The cells were bound to activated concanavalin A beads and incubated with the primary antibody (SOA or IgG; 1:100 dilution) overnight in Antibody Buffer. The cells on beads were rinsed with Dig-wash buffer and incubated at room temperature for 1h with the secondary antibody (Guinea Pig anti-Rabbit IgG, ABIN101961). After a rinse and a wash with Dig-wash buffer, 1:100 pA-Tn5 (prepared by the IMB Protein Production Core Facility) in Dig-300 buffer was added and incubated on a Nutator for 1 hour at room temperature. After a rinse and a wash with Dig-300 buffer, 150 μl of Tagmentation buffer containing MgCl_2_ was added and the samples were incubated at 37°C for 1h. Next, 5 μl of 0.5M EDTA, 1.5 μl of 10% SDS, and 1.25 μl of Proteinase K (20 mg/ml, Invitrogen, 10665795) were added and the samples were incubated for 1h at 55°C. After Proteinase K digestion, DNA was extracted using the DNA Clean&Concentrator-5 Zymo kit. Libraries were generated by PCR amplification (15 cycles) with NEBNext Ultra II Q5 Master Mix with custom primers. Amplified libraries were purified with SPRI magnetic beads and analysed using the DNA High Sensitivity Bioanalyzer. The libraries were pooled in equimolar ratios aiming at 10 Mio reads per library. The pool was sequenced on a NextSeq 2000 P3 (200 cycles) FC, PE for 2x111 cycles plus 2x8 cycles for the index reads. The analysis was performed using the NGSpipe2go ChIP-seq pipeline with modifications (“NGSpipe2go,” n.d.). Briefly, the reads were trimmed using cutadapt (v.4.6) to remove Illumina adapter sequences and subsequently mapped to the AgamP4 reference genome with bowtie2 (v.2.4.5). Multimapping and duplicate reads were removed. We then called peaks using macs2 (v.2.1.2) with the corresponding IgG samples as control, using the effective genome size of 300Mb. The resulting consensus peakset containing peaks called in at least 2 samples was used for downstream analysis with DiffBind (v.3.8.4) to identify sites that were significantly (FDRL<L0.05) differentially bound between wild-type and *SOA-KI* samples (results in Supplemental Table 5). Mean coverage of replicates was calculated with WiggleTools (v.1.2.8) and downstream visualization of differentially bound peaks (for example, heatmaps) were generated using deepTools (v.3.5.1).

### RNA-seq of *SOA-KI* embryos

RNA was isolated as described in “Time course RNA isolation, library generation and sequencing”. Genotyping was performed as described in “Genotyping” for post-ZGA samples. Two embryos of the same sex and genotype were pooled together prior to library generation, which were prepared with Illumina’s Stranded mRNA Prep Ligation Kit following Stranded mRNA Prep Ligation ReferenceGuide (April 2021) (Document 1000000124518 v02). Libraries were prepared with a starting amount of 4 ng and amplified with 13 PCR cycles. Two post PCR purification steps with AMPure XP Beads were performed. Libraries were pooled together in equimolar ratio aiming for at least 10 million reads per sample and sequenced on a NextSeq 2000 P3 (200 cycles) FC, PE for 2x108 cycles plus 2x10 cycles for the index reads and 1 dark cycle upfront R1 and R2. Reads were mapped to the AgamP4 genome using the Ensembl AgamP4 annotation (release 48) with STAR (v. 2.7.3a) using the following parameters: outFilterMismatchNoverLmax 0.04 outFilterMismatchNmax 999. Only uniquely mapped reads were used for downstream analysis. Coverage signal tracks (bigWigs) of primary alignments were generated using Samtools (v1.10) and DeepTools (v3.5.1). Primary alignments were assigned to features using subread 2.0.0 with the AgamP4 annotation (release 48) as a reference. Differential expression analysis was performed using DESeq2 (1.26.0) and only genes with FDR<0.05 were considered differentially expressed. One replicate per group (except *SOA-KI* males) was excluded because of significant deviation from the other samples. The exclusion was based on high Euclidean distances and clustering separately from all other samples in PCA plots (excluded samples: 24_M_WT_18h_Rep4, 18_F_WT_18h_Rep2, 28_F_SoaKI_18h_Rep4).

### Primers

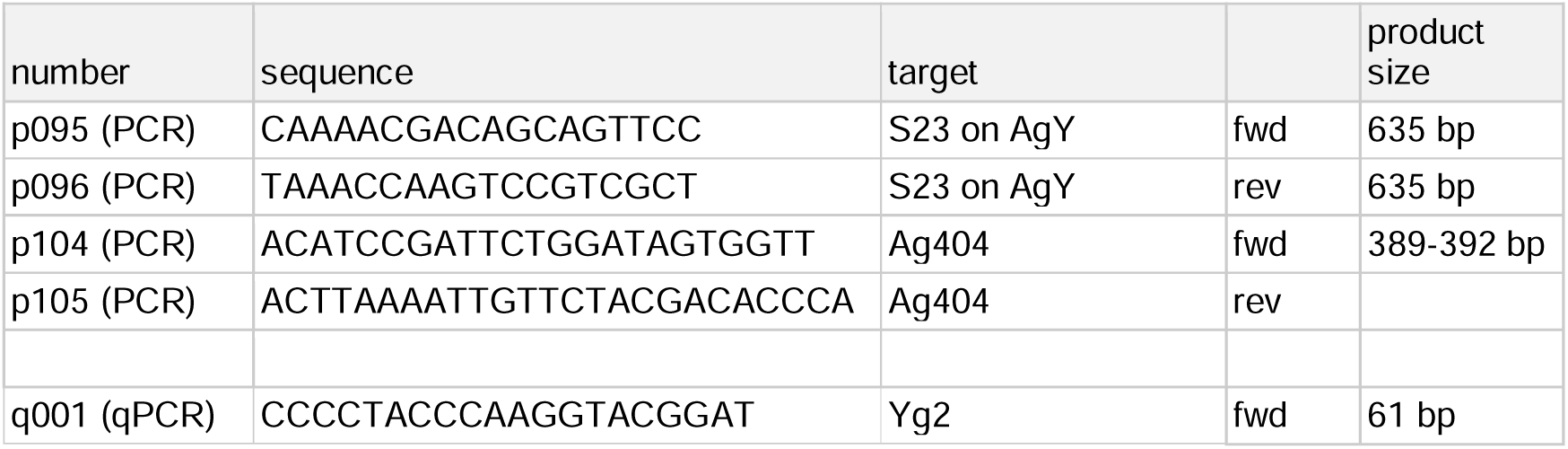
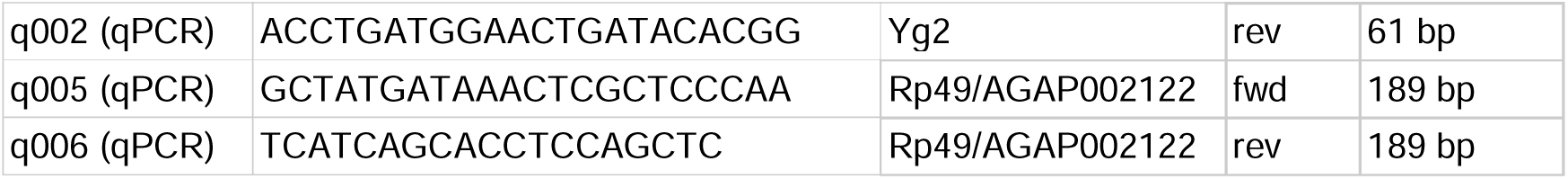

### Statistics, software and web resources

All statistics were calculated with R version 3.6.1 for the time course RNA-seq (DEseq2) analysis, later analyses were done with R version 4.2.2. In the box plots the line that divides the box into two parts represents the median, box bottom, and top edges represent interquartile ranges (IQR, 0.25th to 0.75th quartile [Q1-Q3]), whiskers represent Q1 − 1.5*IQR (bottom), Q3 + 1.5*IQR (top). Bar plots represent the mean ± SEM with overlaid data points representing independent experiments. Results were considered significant at FDR or p-value below 0.05, respectively. ns, not significant; NA, not analyzed; SEM, standard error of the mean. Unless otherwise indicated statistical tests were two-sided. Gene and protein sequences were retrieved from FlyBase (http://flybase.org) and VectorBase (https://www.vectorbase.org/). Lists of 1:1 orthologues were obtained via the Biomart tool on VectorBase. The GO term analysis was performed on VectorBase. No custom codes were used in this study.

### Data availability

The RNA-seq transcriptome atlas data is deposited under GSE210624. The SOA CUT&Tag and *SOA-KI* mutant RNA-seq data is deposited under GSE285042.

## Acknowledgements

The authors wish to thank all members of the Gene Dosage Lab for inputs on the project. They would also like to acknowledge Maya Wilkens for sharing her expertise on the clustering analysis. Support by the IMB Genomics Core Facility and the use of its NextSeq500 (funded by the Deutsche Forschungsgemeinschaft (DFG, German Research Foundation) – INST 247/870-1 FUGG) and NextSeq2000 is gratefully acknowledged. The authors would like to thank the Genomics Core Facility (GeneCore) of EMBL (Heidelberg) for supporting the embryogenesis transcriptome atlas in this project. AIK was supported by a Boehringer Ingelheim Foundation Fellowship. CIKV is supported by the Deutsche Forschungsgemeinschaft (DFG, German Research Foundation) - Individual Project Grant 513744403, Scientific Network Grant 531902894, GRK GenEvo 407023052, GRK 4R 491145305, Forschungsinitiative Rheinland-Pfalz (ReALity) and institutional funding from the IMB. Mosquito breeding and sample production was performed in the IBMC Insectarium facility with financial support from the University of Strasbourg, Inserm, CNRS and Agence Nationale de la Recherche through grants EQUIPEX ANR-11-EQPX-0022/ANR and GDaMO ANR-19-CE35-0007-01. The funders had no role in study design, data collection and analysis, decision to publish, or preparation of the manuscript.

## Author Contributions

The study was conceived by CIKV and AIK. EM performed mosquito rearing and collected starting material for RNA isolation and CUT&Tag. AIK performed all wet lab experiments. FR established pipelines and consulted on data analysis. AIK and CIKV performed data analyses and wrote the initial draft of the manuscript. All authors have revised the manuscript.

## Supplemental Files

Supplemental Table S1

Statistic related to RNA-seq: raw read counts from all the replicates, mapping statistics single-embryo RNA-seq, RNA-seq raw read counts for wild-type versus *SOA-KI*, mapping statistics *SOA-KI*

Supplemental Table S2

Clustering analysis results with scaled expression, GOterms clusters motifs

Supplemental Table S3

Alternative polyadenylation site analysis (APAtrap): bed file, APA results; deAPA statistics

Supplemental Table S4

DEseq2 results of male versus female embryos in all timepoints DEseq2 result of wild-type versus *SOA-KI* embryos

Supplemental Table S5

SOA CUT&Tag analysis: DiffBind results for wild-type compared to SOA-KI samples (4h, 7h, and 20h).

Supplemental File S1

PDF File showing expression dynamics along embryogenesis for each *Anopheles* gene plotted separately for males and females.

## Supplemental figure legends

**Figure S1.**
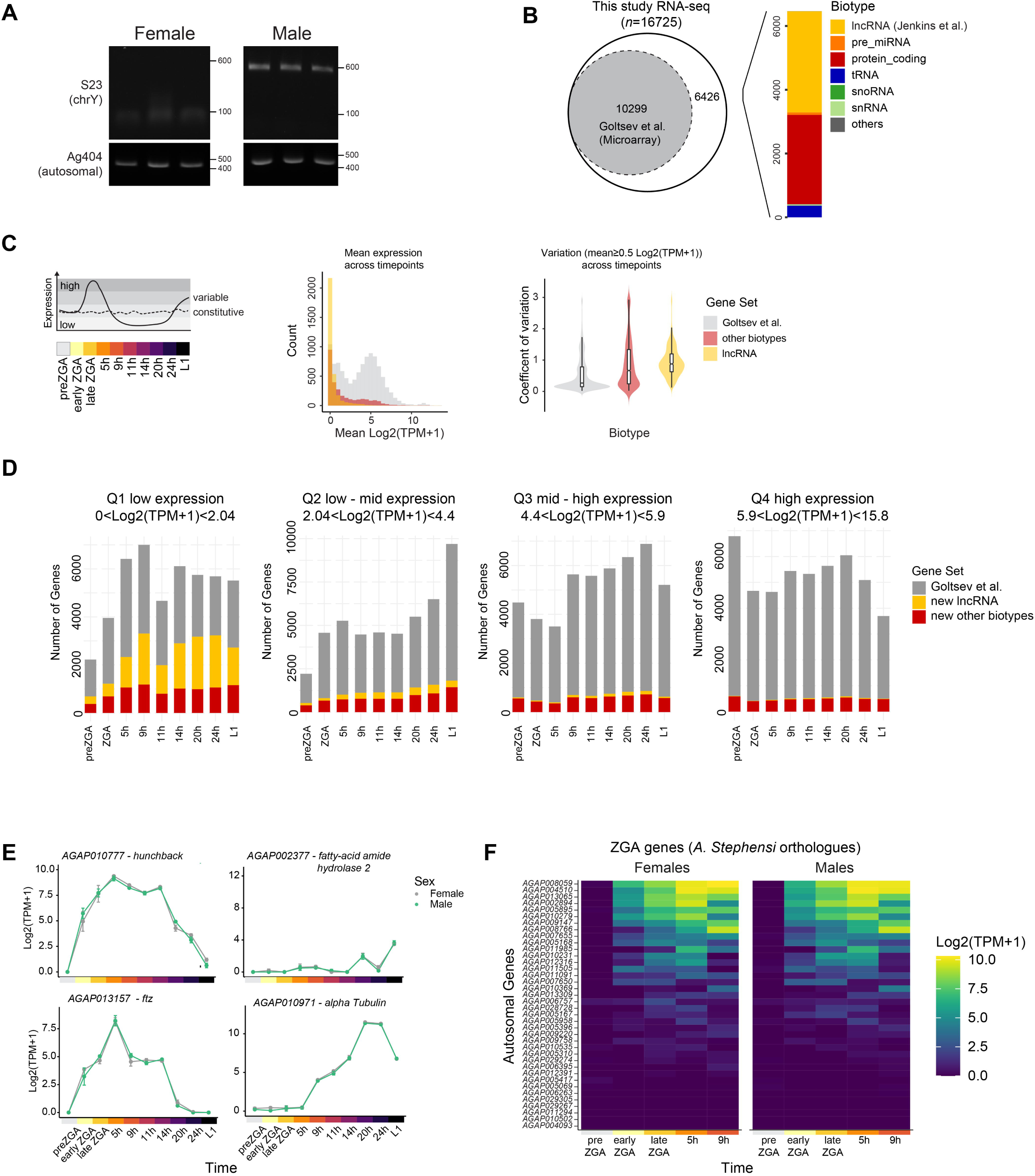
Embryogenesis transcriptome atlas sample characterization. A) Agarose gel showing the results of embryo genotyping based on PCR with Y-linked (top) and autosomal control (bottom) repeat-specific primers. B) Left: A Venn diagram comparing the numbers of genes measured in our study by RNA-seq to previously published microarray data (Goltsev et al. 2009). Right: bar plot showing the predicted biotype of genes in our RNA-seq that were not measured in the previously published microarray data. lncRNA - long non-coding RNA, snRNA - small nuclear RNA, snoRNA - small nucleolar RNA, tRNA - transfer RNA, miRNA - microRNA. C) Characterization of gene sets in the customs annotation. Left: schematic representation of variable and constitutive gene expression patterns. Middle: histogram showing the number of genes belonging to each gene set in a given mean expression (log2TPM+1) bin. Right: violin and bar plot showing the variation coefficient of different gene sets across embryogenesis. D) Stacked barplots showing numbers of genes in each gene set belonging to respective expression quartile across the timepoints studied. E) Plots showing expression in log2(TPM+1) of selected developmental genes across embryogenesis, plotted separately for male and female samples. The dots indicate the mean values of all the replicates, while the error bars show the standard error of the mean (SEM). F) Heatmap comparing the male and female expression levels of zygotic genes (gene names on y-axis) across early embryonic development (x-axis for timepoints). The tiles are colored based on expression levels in log2(TPM+1)

**Figure S2.**
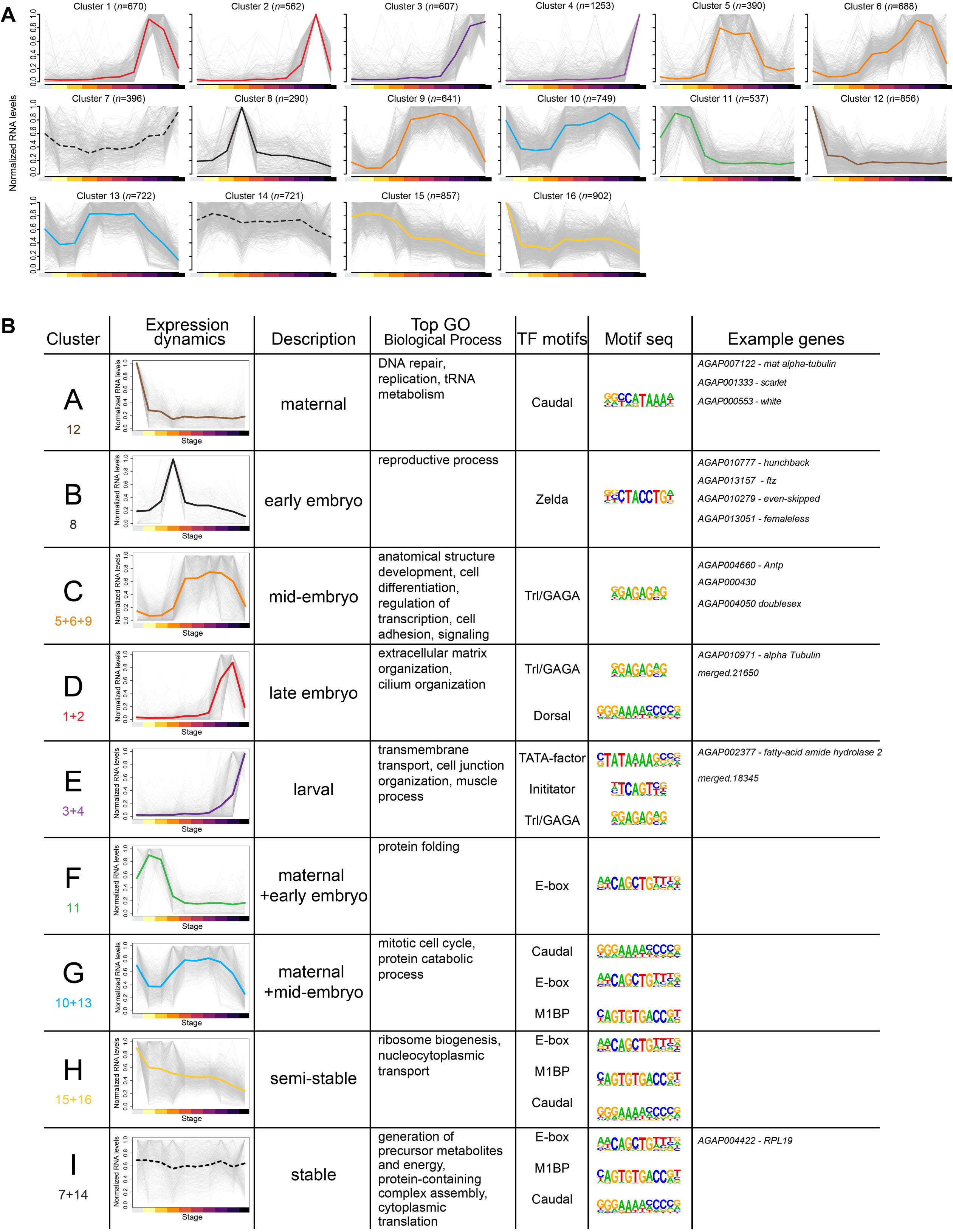
Self-organizing maps reveal clusters of genes with similar developmental dynamics. A) Results of clustering into 16 groups with self organizing maps for genes with the mean expression of at least 5 TPM at any developmental timepoint. *n* - number of genes assigned to the cluster, y-axis indicates expression scaled from 0 to 1 (where 1 indicating maximum mean expression). B) Manually curated clustering based on developmental dynamics. Below the new cluster name (A-I) are the cluster numbers from (A) that were merged to form the curated cluster. Top GO - top gene ontology terms significantly enriched in the curated cluster. TF motifs - names of transcription factors for which the known binding motifs were scored as significantly enriched in the promoters of genes belonging to a given cluster as scored by HOMER (v.1). Motif seq - the consensus motif sequence of the transcription factors described in previous column. Example genes belonging to the clusters are shown.

**Figure S3.**
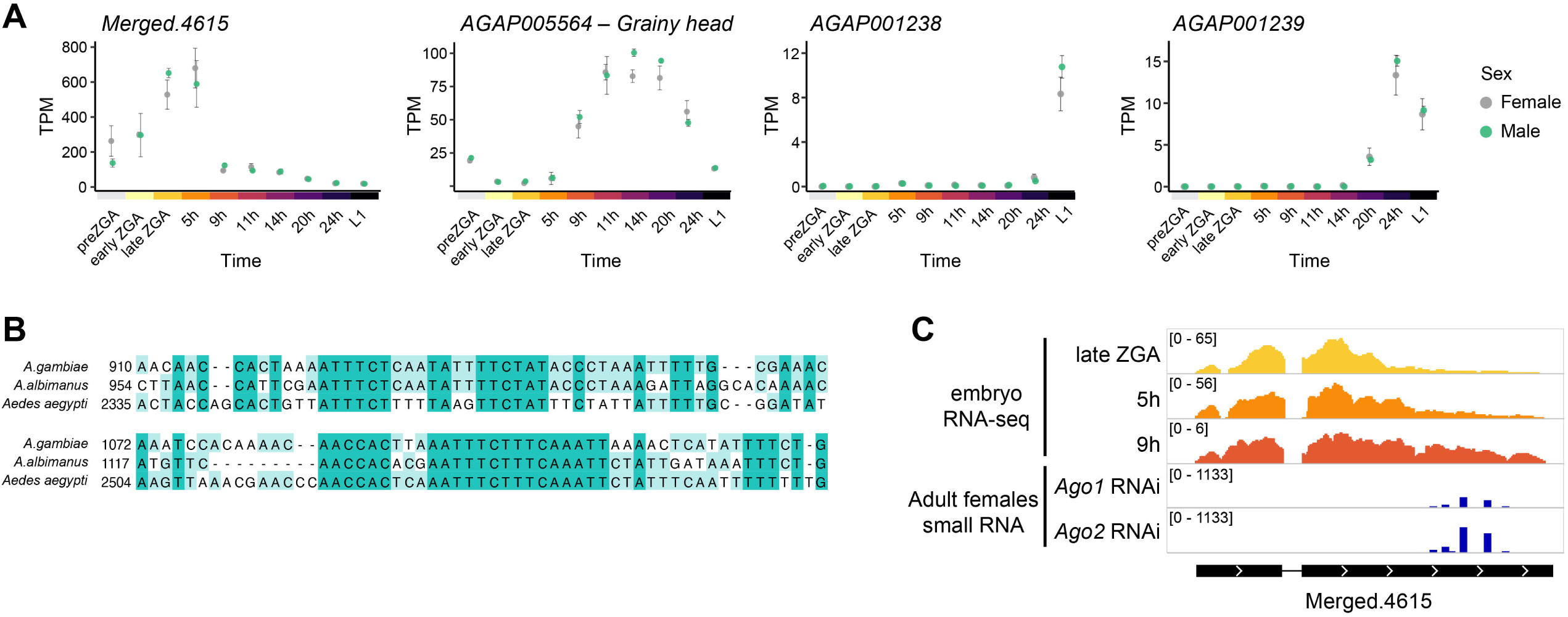
Characterization of a putative miRNA precursor. A) Plots showing expression of *Merged.4615* lncRNA, its putative regulator and neighboring genes across embryogenesis, plotted separately for male and female samples. The x-axis indicates the developmental time. The y-axis shows the TPM values. The dots indicate the mean values of all replicates, while the error bars show the standard error of the mean (SEM). B) Alignment of *Merged.4615* and its orthologues in *Anopheles albimanus* and *Aedes aegypti* identified based on synteny. Only a part of alignment that includes the highly conserved sequences is shown. The lncRNA can be found in the following genomic regions: *A. gambiae Merged.4615,* chromosome 2R: 1827987 - 1829487; *A. albimanus* scaffold CM024292 (chromosome 2): 2551012 - 2552335, annotated as *AALB20_031383.R62999; A. aegypti:* AaegL5 chromosome 3: 158079404 - 158083570. C) Coverage tracks showing the *Merged.4615* region from this RNA-seq (top) and published small RNA sequencing experiments (bottom). Analyzed and normalized coverage tracks published by (Biryukova, Ye, and Levashina 2014) were obtained from the Mosquito Small RNA Genomics database (Ma et al. 2021). *Ago2* RNAi is a negative control for *Ago1* RNAi treatment, since *Ago2* is involved in siRNA but not miRNA biosynthesis in *Anopheles*.

**Figure S4.**
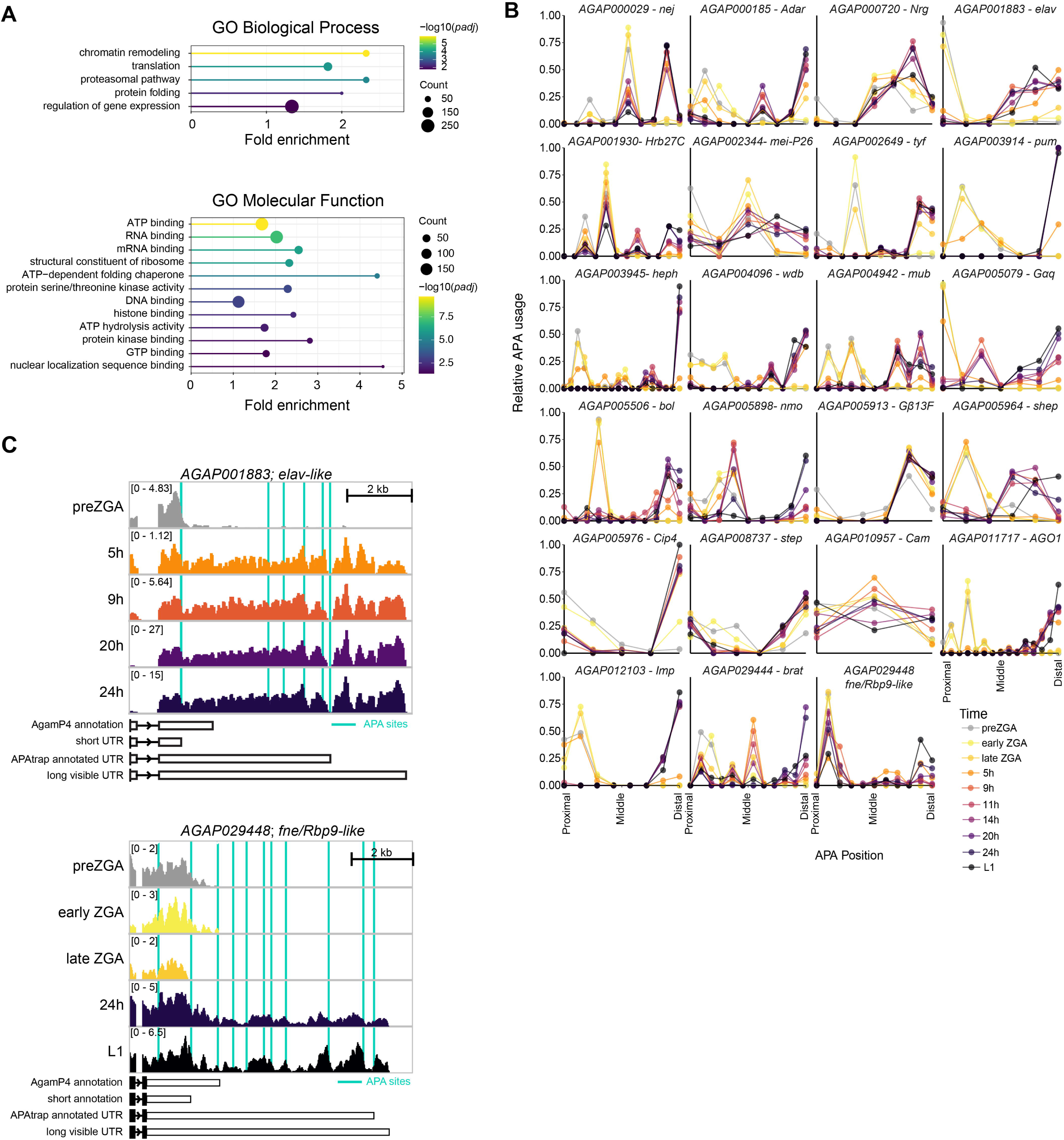
Conservation of 3’UTR extension between A. gambiae and D. melanogaster. A) A lollipop chart depicting the results of a Gene Ontology (GO) term analysis genes with 3’UTR elongation from preZGA to early ZGA stages. The x-axis indicates the fold enrichment over random distribution. The color represents the significance, while the size of the circle is based on the number of genes in the set assigned to a given GO term. Top: enriched terms in Biological Process gene ontology; Bottom: enriched terms in Molecular Function gene ontology; B) Relative APA usage for *A. gambiae* orthologues of genes previously reported to have 3’UTR extension in embryonic neural *D. melanogaster* tissues (Hilgers et al. 2011). The x-axis represents APA positions ordered from the most proximal to the most distal, while the y-axis shows the percentage of transcripts using a respective polyadenylation site. C) Coverage tracks of female RNA-seq embryo samples for the *elav* orthologue, *AGAP001883* (CPM-normalized). D) Coverage tracks of female RNA-seq embryo samples for the *fne/Rbp9* orthologue, *AGAP029448* (CPM-normalized) in selected timepoints.

**Figure S5.**
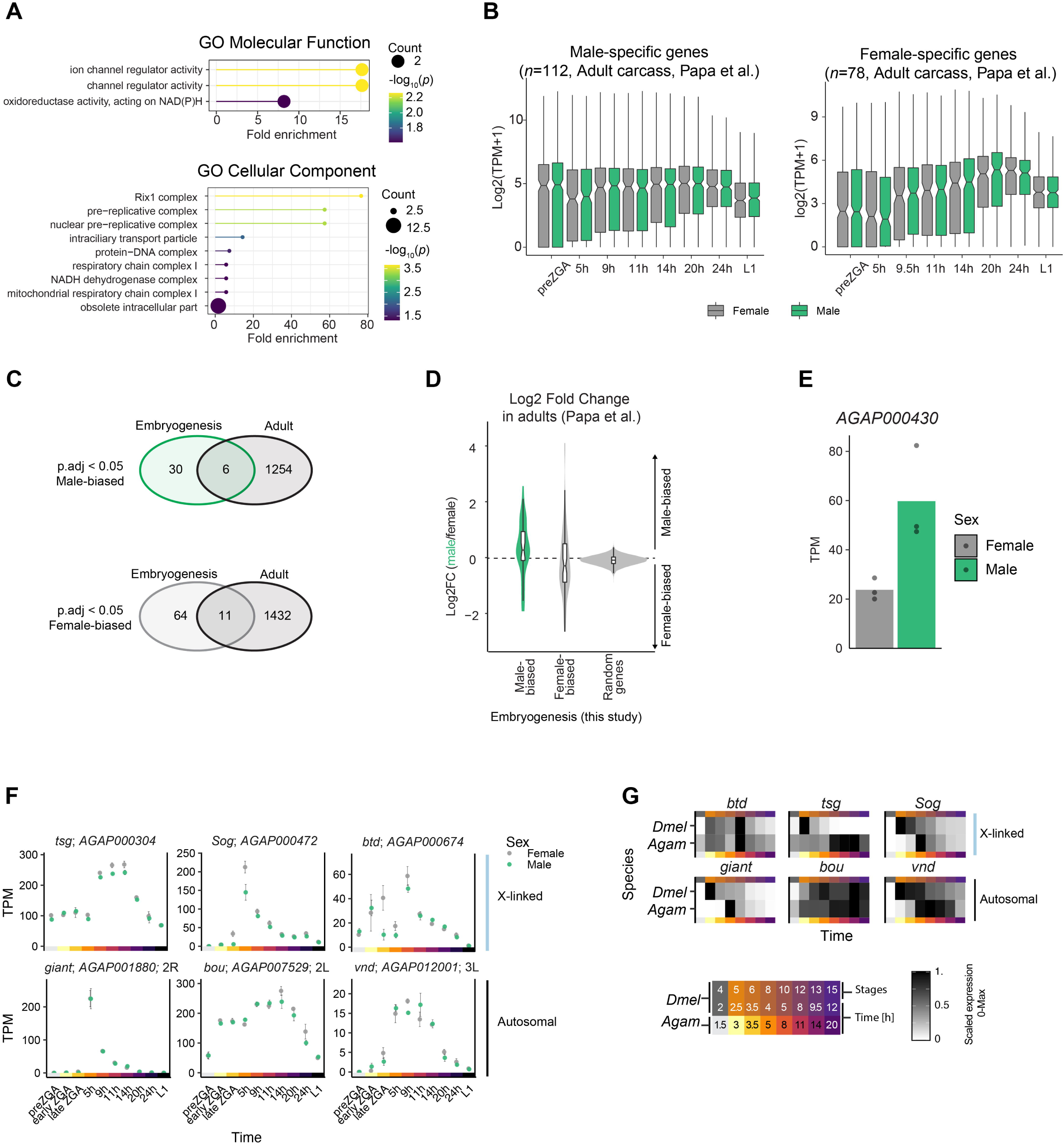
Comparison of sex-specificity and expression dynamics to previously published datasets in adult mosquitoes and *D. melanogaster* embryos. A) A lollipop chart depicting the results of a Gene Ontology (GO) term analysis of genes that are sex-biased in the embryo dataset the biological process (top) and cellular component (bottom) ontology. The x-axis indicates the fold enrichment over random distribution. The color represents the significance, while the size of the circle is based on the number of genes in the set assigned to a given GO term. B) Embryonic expression patterns in log2(TPM+1) of previously reported genes with sex-specific expression in adult mosquitoes (left: male-specific; right: female-specific). C) Venn diagrams showing overlaps between genes with sex-biased expression in embryogenesis (this study) and adult mosquitoes (Papa et al. 2017). Merged lncRNAs (7 in males, 2 in females) are not shown as they were not covered by the (Papa et al. 2017) study. D) Log2FC between adult males and females (Papa et al. 2017) for genes scored as sex-biased in embryogenesis dataset (this study). E) Expression levels in TPM of *AGAP000430* in adult male and female *A. gambiae* mosquitoes (Papa et al. 2017). F) Plots showing expression of the orthologues of important developmental regulators that are dosage compensated by a non-canonical mechanism in *D. melanogaster*. The expression in log2(TPM+1) is plotted across *A. gambiae* embryogenesis, separately for male and female samples. The dots indicate the mean values of all the replicates, while the error bars show the standard error of the mean (SEM) G) Comparison of embryonic expression patterns of important developmental regulators between *D. melanogaster* (*Dmel*) and *A. gambiae* (*Agam*) (Prayitno et al. 2019). The legend below shows the approximate times after oviposition/egg laying that are compared. The expression is scaled from zero to maximal expression levels across embryogenesis.

**Figure S6.**
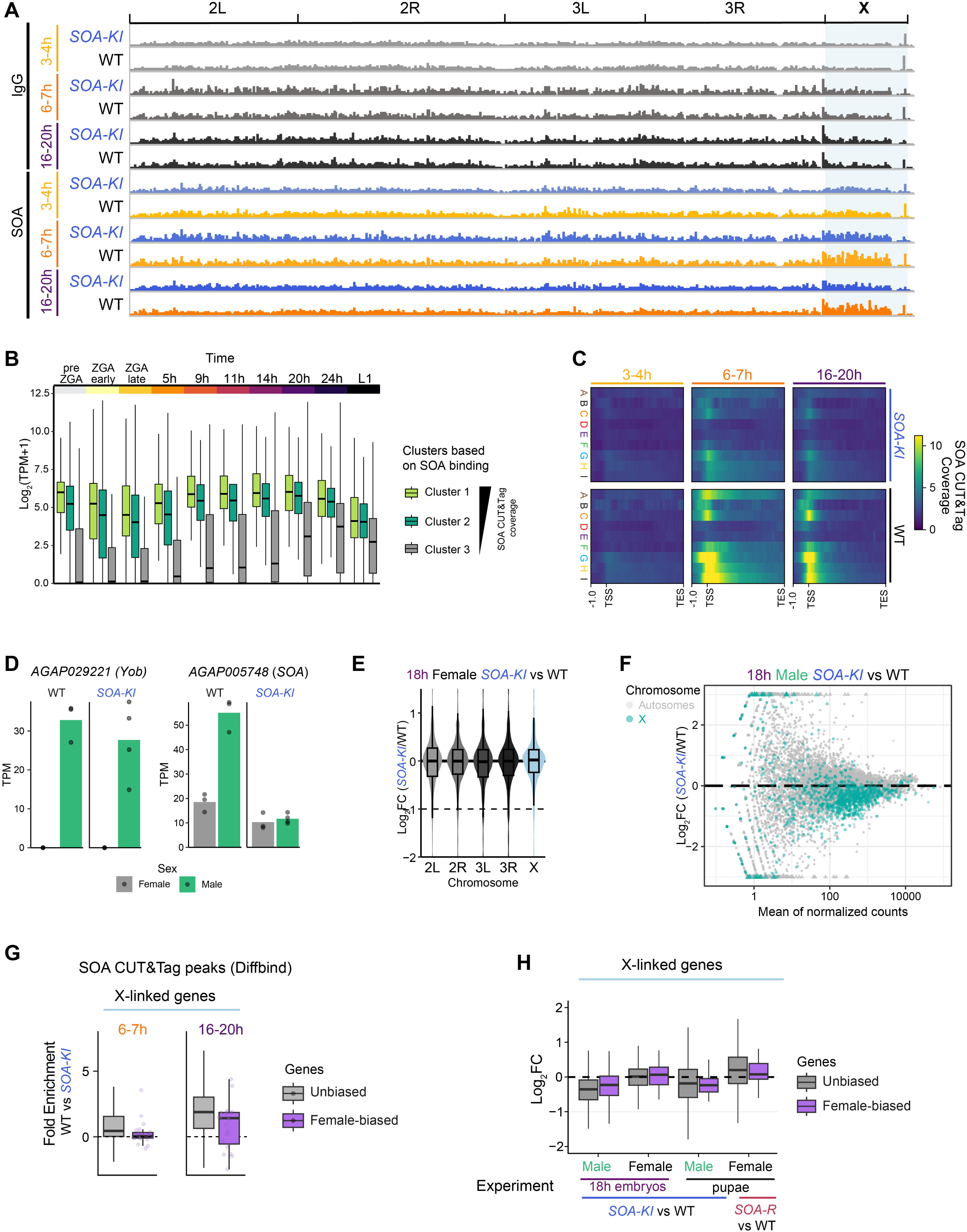
Characterization of SOA binding and effects of SOA loss in embryos. A) Embryogenesis SOA CUT&Tag coverage plot across all chromosomal arms. B) Embryonic expression in log2(TPM+1) of X-linked genes grouped into 3 clusters based on SOA binding strength (see Fig. 5D). C) Heatmaps comparing mean CUT&Tag coverage for X-linked genes belonging to different developmental clusters (Fig. S2B). The signal is shown from 1Kb upstream until the transcription start site (TSS) followed by the gene body until transcription end site (TES) scaled to 5 Kb. D) Expression levels of *Yob* and *SOA* in TPM confirming the sex and genotype of embryos used in *SOA-KI* RNA-seq. E) Violin and box plots show the DESeq2-obtained log2FC in WT compared to homozygous *SOA-KI* mutant 18h female embryos. All expressed genes (mean read counts across all replicates > 0) are plotted. The genes are grouped by chromosomal location. Plot for male samples can be found in Fig. 5F. F) MA plot of DEseq2 results for comparison of *SOA-KI* to wild-type male 18h embryos. The y-axis shows the log_2_FC between *SOA-KI* and WT, the x-axis shows the mean of normalized counts. X-linked genes are colored in blue, the remaining genes are in grey. G) Strength of SOA binding in embryo CUT&Tag experiments for X-linked genes with female-biased expression as compared to unbiased genes (defined by DESeq2, genes in Fig. 3A). The y-axis shows the fold difference in CUT&Tag signal between WT and *SOA-KI* samples, as scored by DiffBind. Because some genes may have multiple peaks scored, the peak with the highest fold value per gene was chosen for plotting. H) Log_2_FC in expression in various *SOA* mutant lines. X-linked genes with female-biased expression as compared to unbiased genes. Pupae datasets from (Kalita et al. 2023). *SOA-R* is a rescue allele, where the spliced coding sequence is inserted under the endogenous SOA promoter, leading to expression of full length isoform of SOA in females.

