## Supplemental File S1 for "Sex-specific transcriptome dynamics of *Anopheles gambiae* during embryonic development"

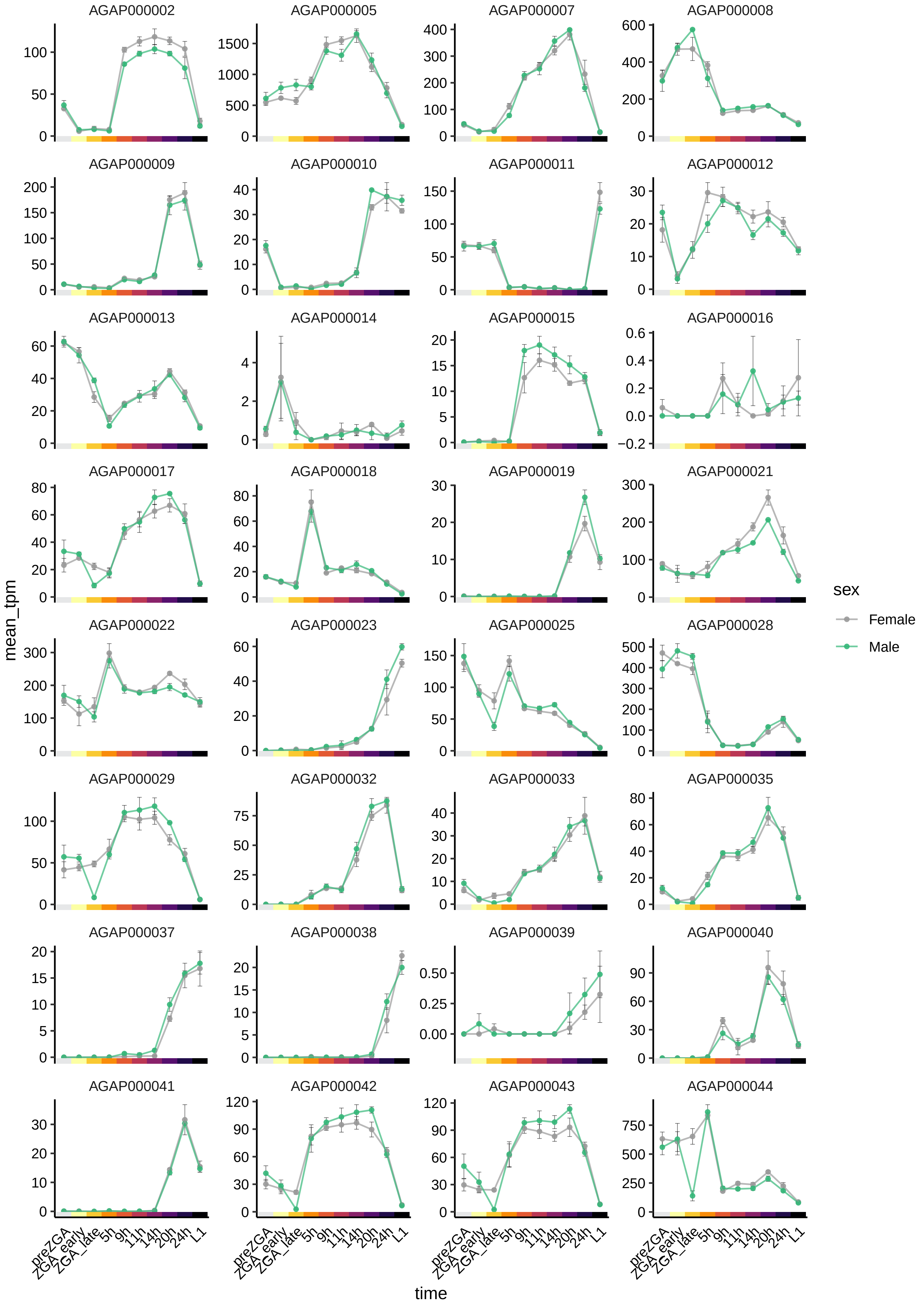

AGAP000002
AGAP000005
AGAP000007
AGAP000008
AGAP000009
AGAP000010
AGAP000011
AGAP000012
AGAP000013
AGAP000014
AGAP000015
AGAP000016
AGAP000017
AGAP000018
AGAP000019
AGAP000021
AGAP000022
AGAP000023
AGAP000025
AGAP000028
AGAP000029
AGAP000032
AGAP000033
AGAP000035
AGAP000037
AGAP000038
AGAP000039
AGAP000040
AGAP000041
AGAP000042
AGAP000043
AGAP000044
mean_tpm
time
preZGA
ZGA_early
ZGA_late
5h
9h
11h
14h
20h
24h
L1
sex
Female
Male

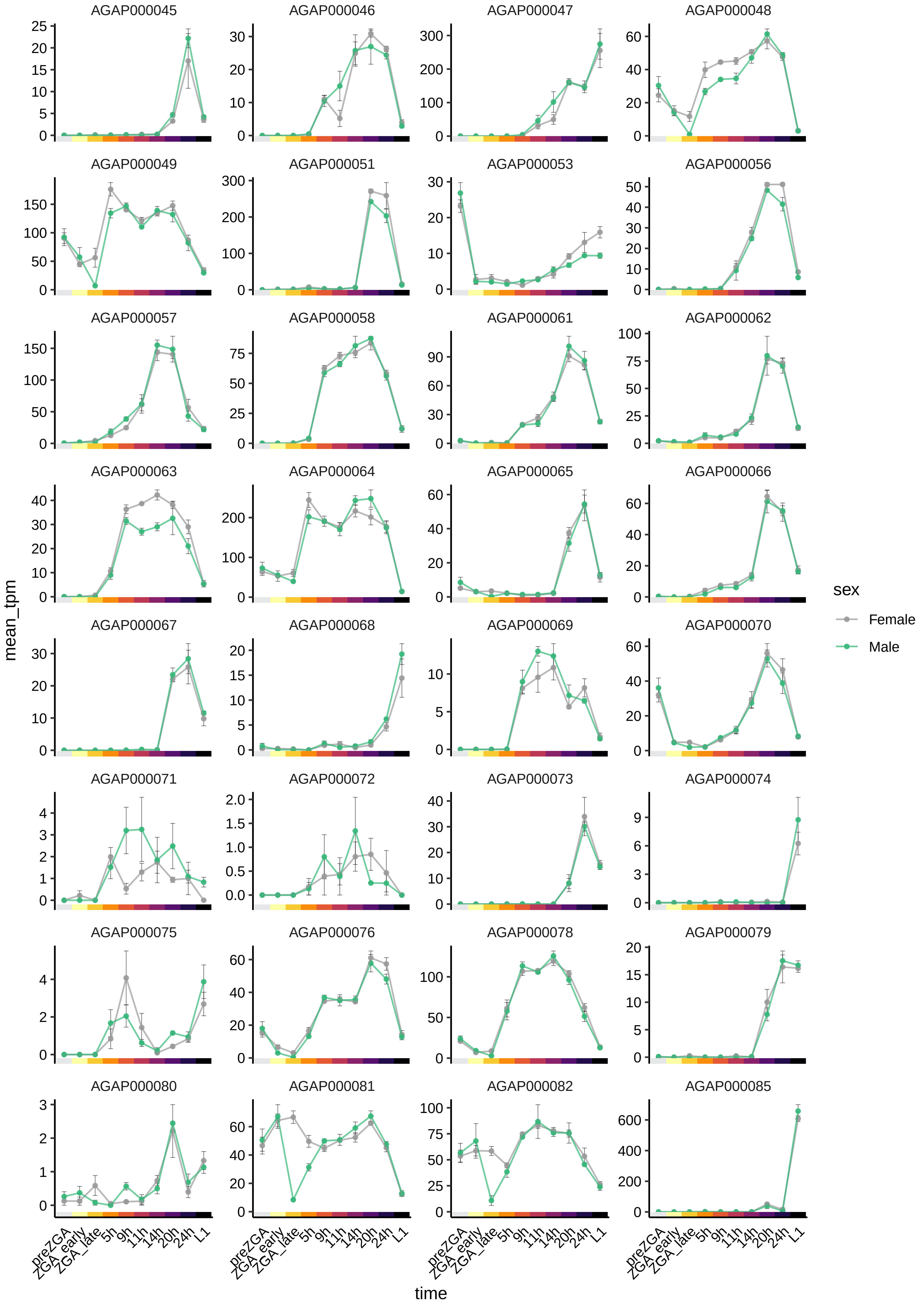
AGAP000045
AGAP000046
AGAP000047
AGAP000048
AGAP000049
AGAP000051
AGAP000053
AGAP000056
AGAP000057
AGAP000058
AGAP000061
AGAP000062
AGAP000063
AGAP000064
AGAP000065
AGAP000066
AGAP000067
AGAP000068
AGAP000069
AGAP000070
AGAP000071
AGAP000072
AGAP000073
AGAP000074
AGAP000075
AGAP000076
AGAP000078
AGAP000079
AGAP000080
AGAP000081
AGAP000082
AGAP000085
mean_tpm
time
preZGA
ZGA_early
ZGA_late
5h
9h
11h
14h
20h
24h
L1
sex
Female
Male

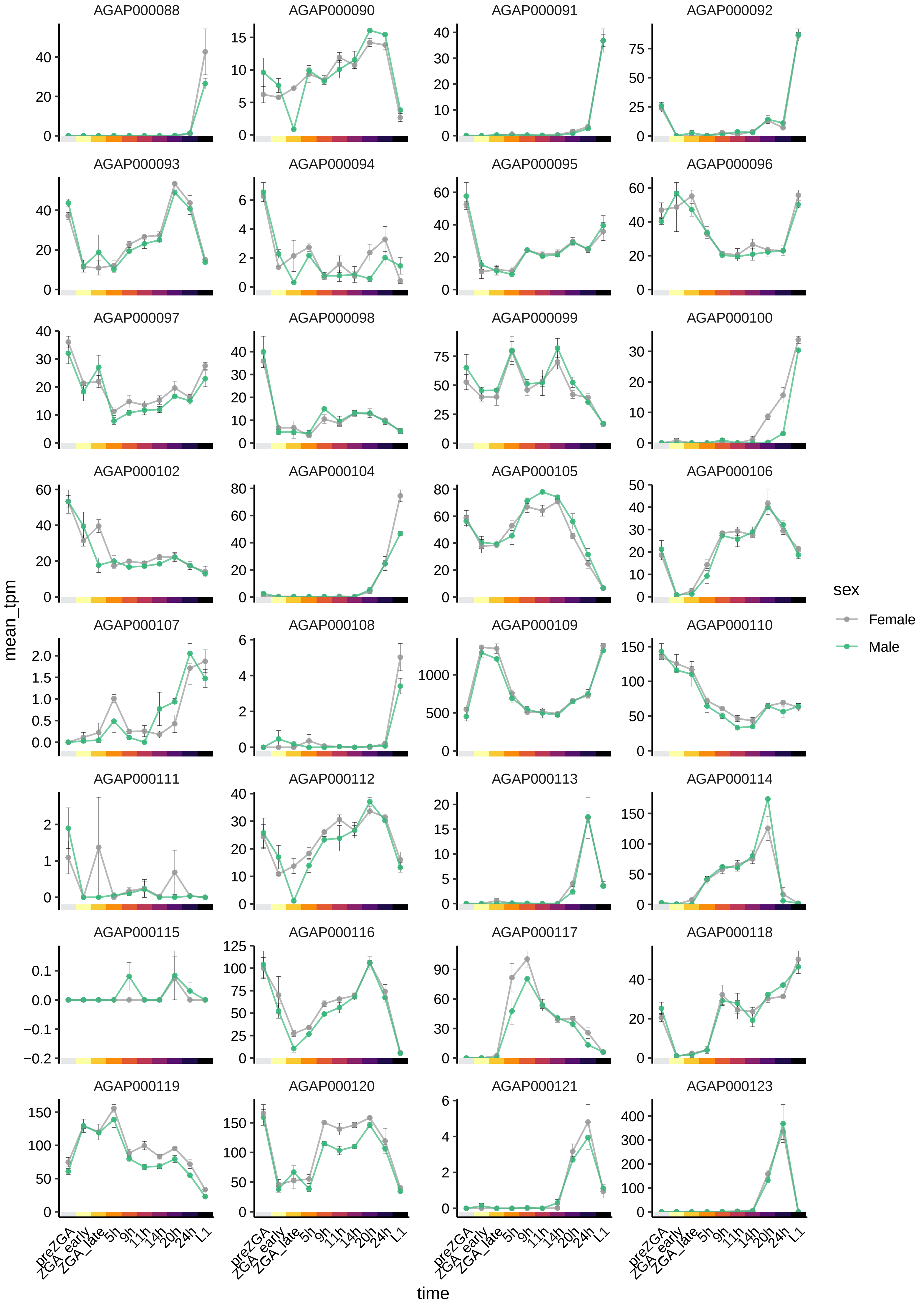

AGAP000088
AGAP000090
AGAP000091
AGAP000092
AGAP000093
AGAP000094
AGAP000095
AGAP000096
AGAP000097
AGAP000098
AGAP000099
AGAP000100
AGAP000102
AGAP000104
AGAP000105
AGAP000106
AGAP000107
AGAP000108
AGAP000109
AGAP000110
AGAP000111
AGAP000112
AGAP000113
AGAP000114
AGAP000115
AGAP000116
AGAP000117
AGAP000118
AGAP000119
AGAP000120
AGAP000121
AGAP000123
mean_tpm
time
preZGA
ZGA_early
ZGA_late
5h
9h
11h
14h
20h
24h
L1
sex
Female
Male

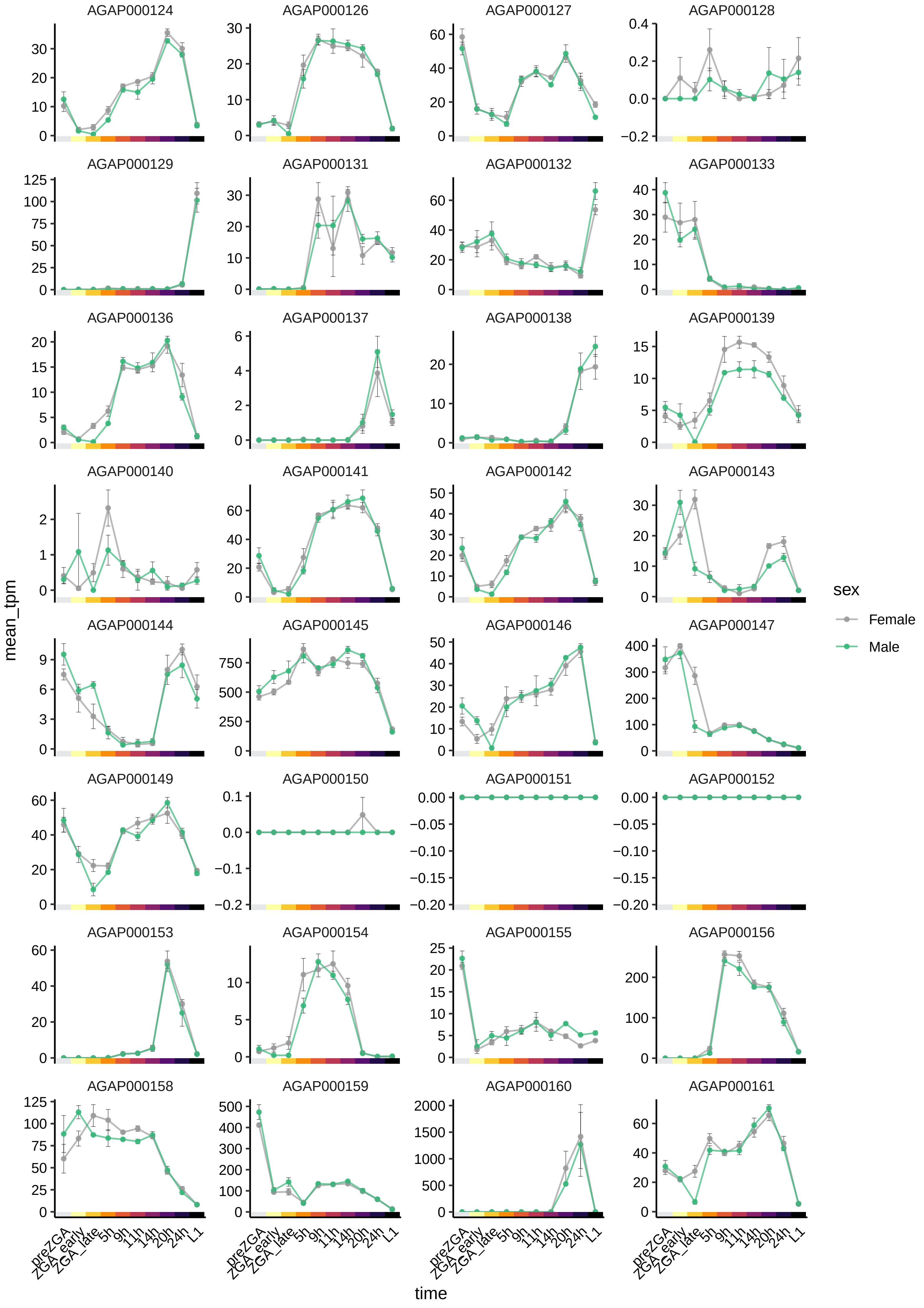
AGAP000124
AGAP000126
AGAP000127
AGAP000128
AGAP000129
AGAP000131
AGAP000132
AGAP000133
AGAP000136
AGAP000137
AGAP000138
AGAP000139
AGAP000140
AGAP000141
AGAP000142
AGAP000143
AGAP000144
AGAP000145
AGAP000146
AGAP000147
AGAP000149
AGAP000150
AGAP000151
AGAP000152
AGAP000153
AGAP000154
AGAP000155
AGAP000156
AGAP000158
AGAP000159
AGAP000160
AGAP000161
mean_tpm
preZGA
ZGA_early
ZGA_late
5h
9h
11h
14h
20h
24h
L1
time
sex
Female
Male

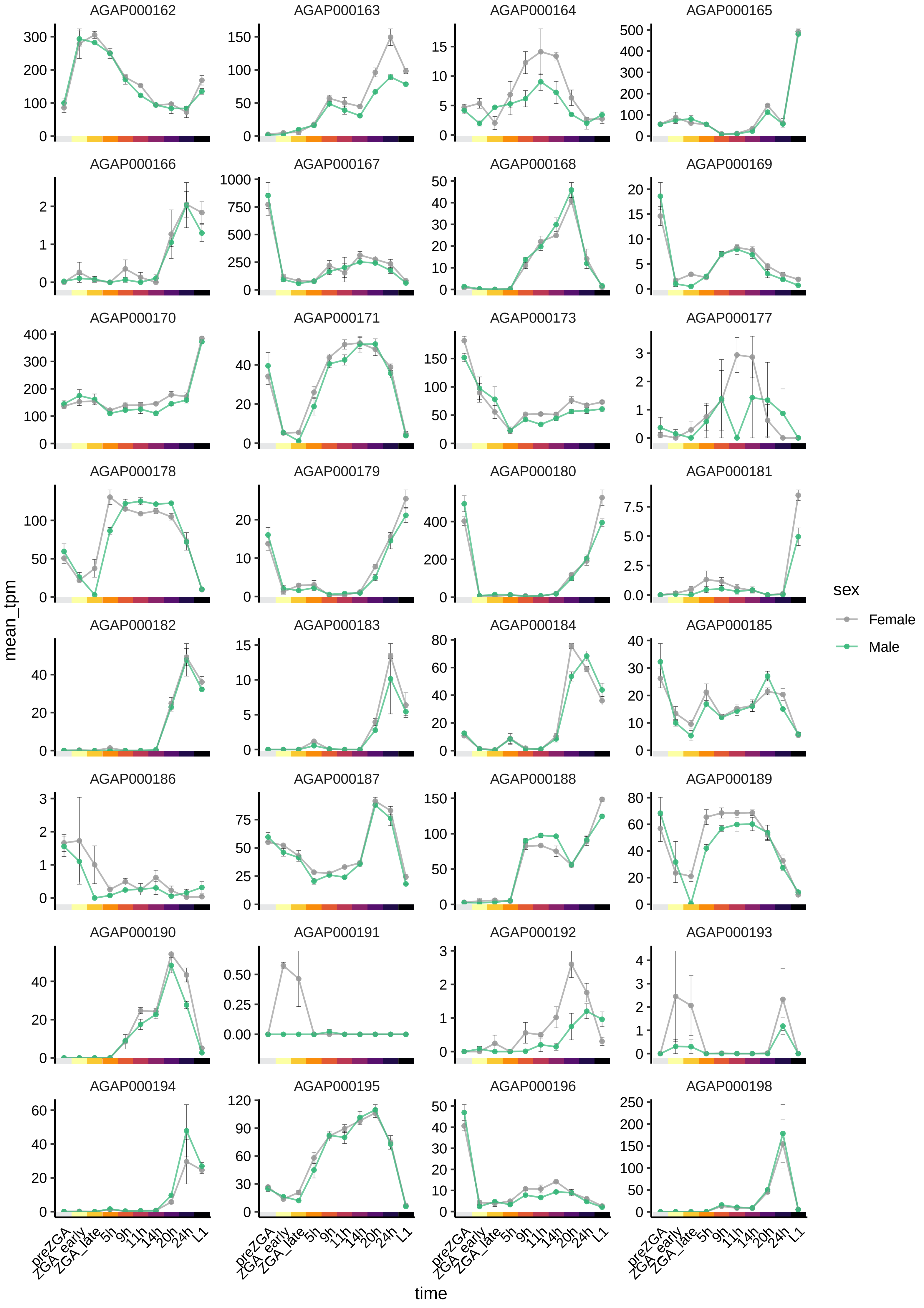

AGAP000162
AGAP000163
AGAP000164
AGAP000165
AGAP000166
AGAP000167
AGAP000168
AGAP000169
AGAP000170
AGAP000171
AGAP000173
AGAP000177
AGAP000178
AGAP000179
AGAP000180
AGAP000181
AGAP000182
AGAP000183
AGAP000184
AGAP000185
AGAP000186
AGAP000187
AGAP000188
AGAP000189
AGAP000190
AGAP000191
AGAP000192
AGAP000193
AGAP000194
AGAP000195
AGAP000196
AGAP000198
mean_tpm
time
preZGA
ZGA_early
ZGA_late
5h
9h
11h
14h
20h
24h
L1
sex
Female
Male

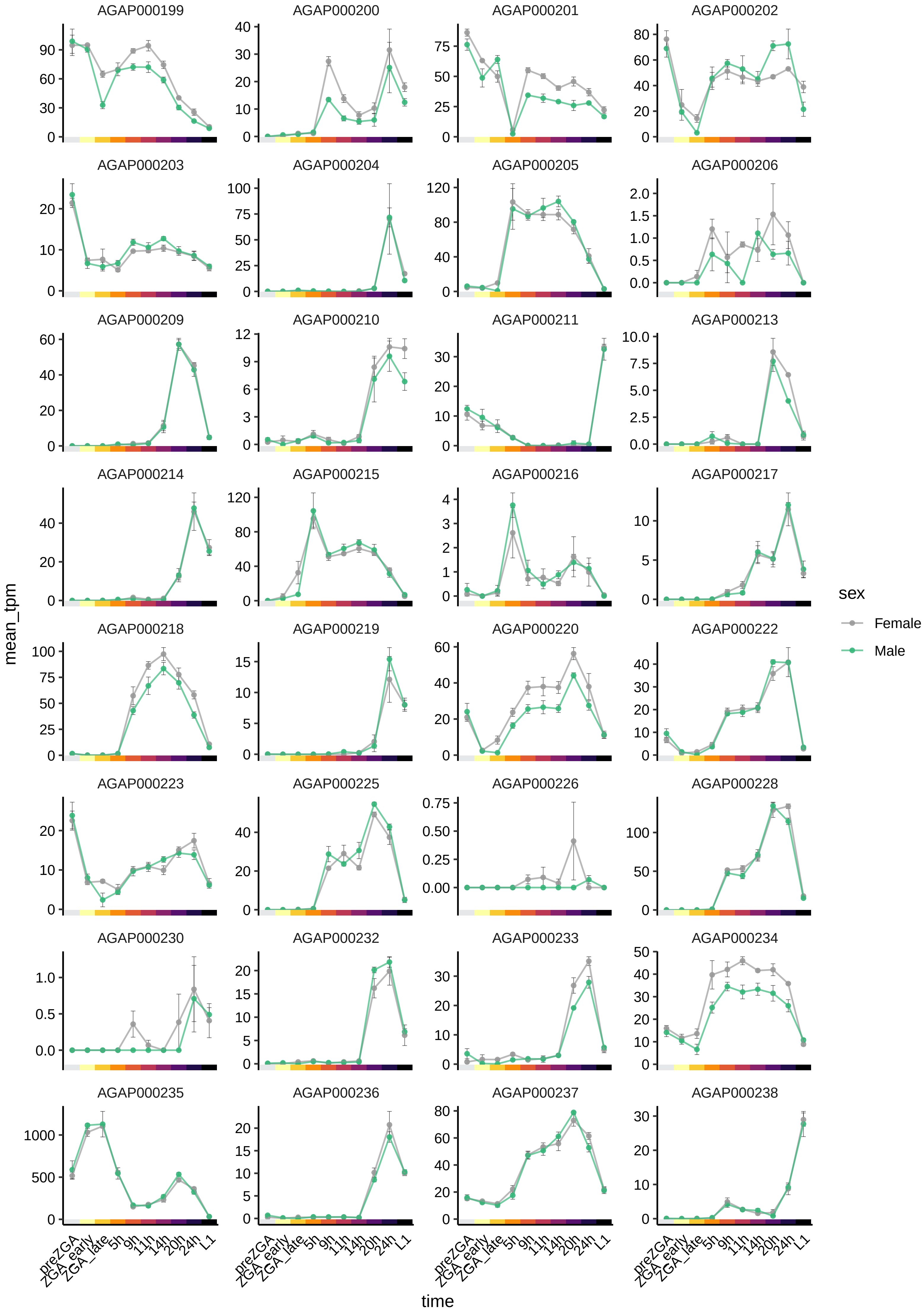

AGAP000199
AGAP000200
AGAP000201
AGAP000202
AGAP000203
AGAP000204
AGAP000205
AGAP000206
AGAP000209
AGAP000210
AGAP000211
AGAP000213
AGAP000214
AGAP000215
AGAP000216
AGAP000217
AGAP000218
AGAP000219
AGAP000220
AGAP000222
AGAP000223
AGAP000225
AGAP000226
AGAP000228
AGAP000230
AGAP000232
AGAP000233
AGAP000234
AGAP000235
AGAP000236
AGAP000237
AGAP000238
mean_tpm
time
preZGA
ZGA_early
ZGA_late
5h
9h
11h
14h
20h
24h
L1
sex
Female
Male

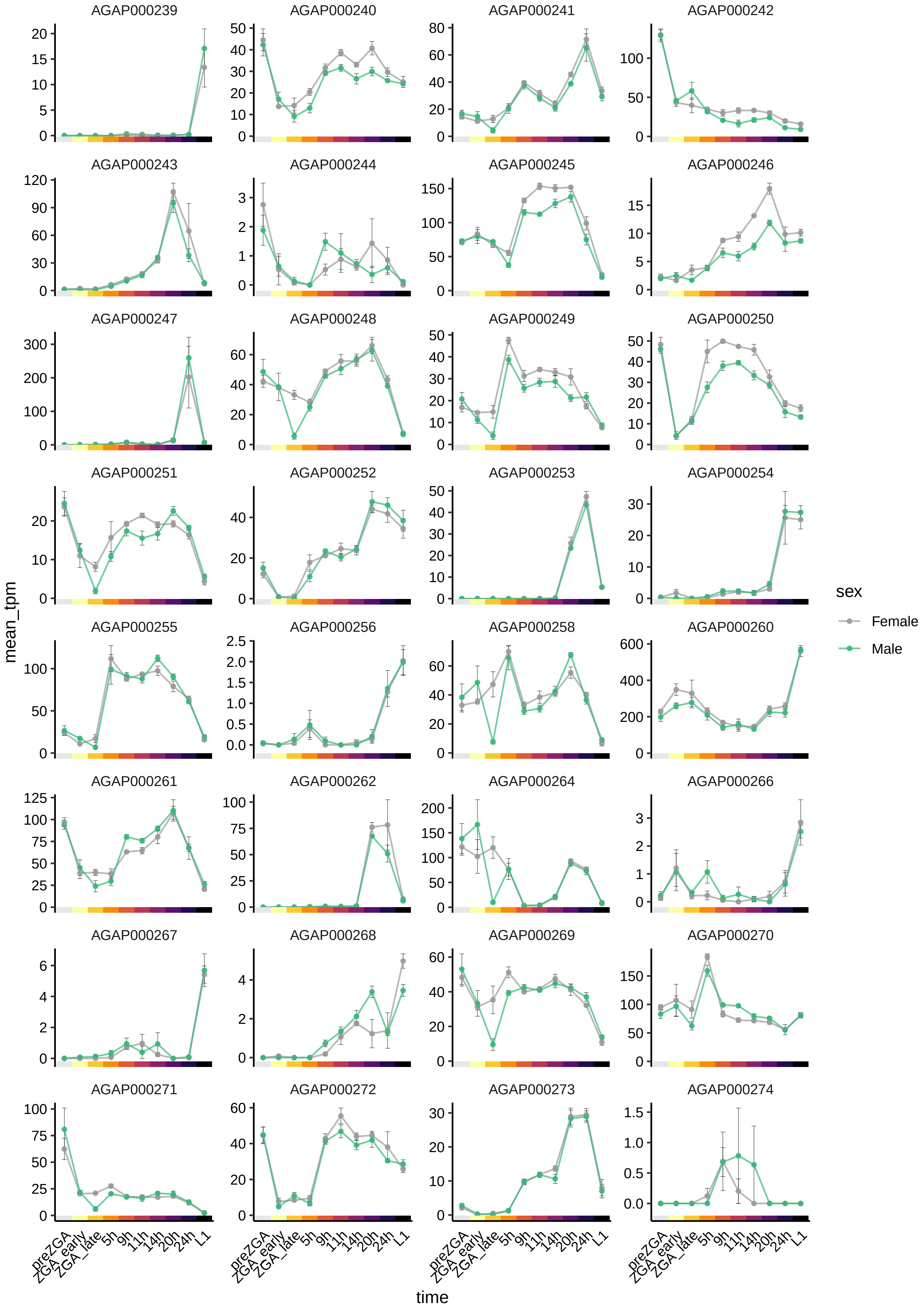

AGAP000239
AGAP000240
AGAP000241
AGAP000242
AGAP000243
AGAP000244
AGAP000245
AGAP000246
AGAP000247
AGAP000248
AGAP000249
AGAP000250
AGAP000251
AGAP000252
AGAP000253
AGAP000254
AGAP000255
AGAP000256
AGAP000258
AGAP000260
AGAP000261
AGAP000262
AGAP000264
AGAP000266
AGAP000267
AGAP000268
AGAP000269
AGAP000270
AGAP000271
AGAP000272
AGAP000273
AGAP000274
mean_tpm
time
preZGA
ZGA_early
ZGA_late
5h
9h
11h
14h
20h
24h
L1
sex
Female
Male

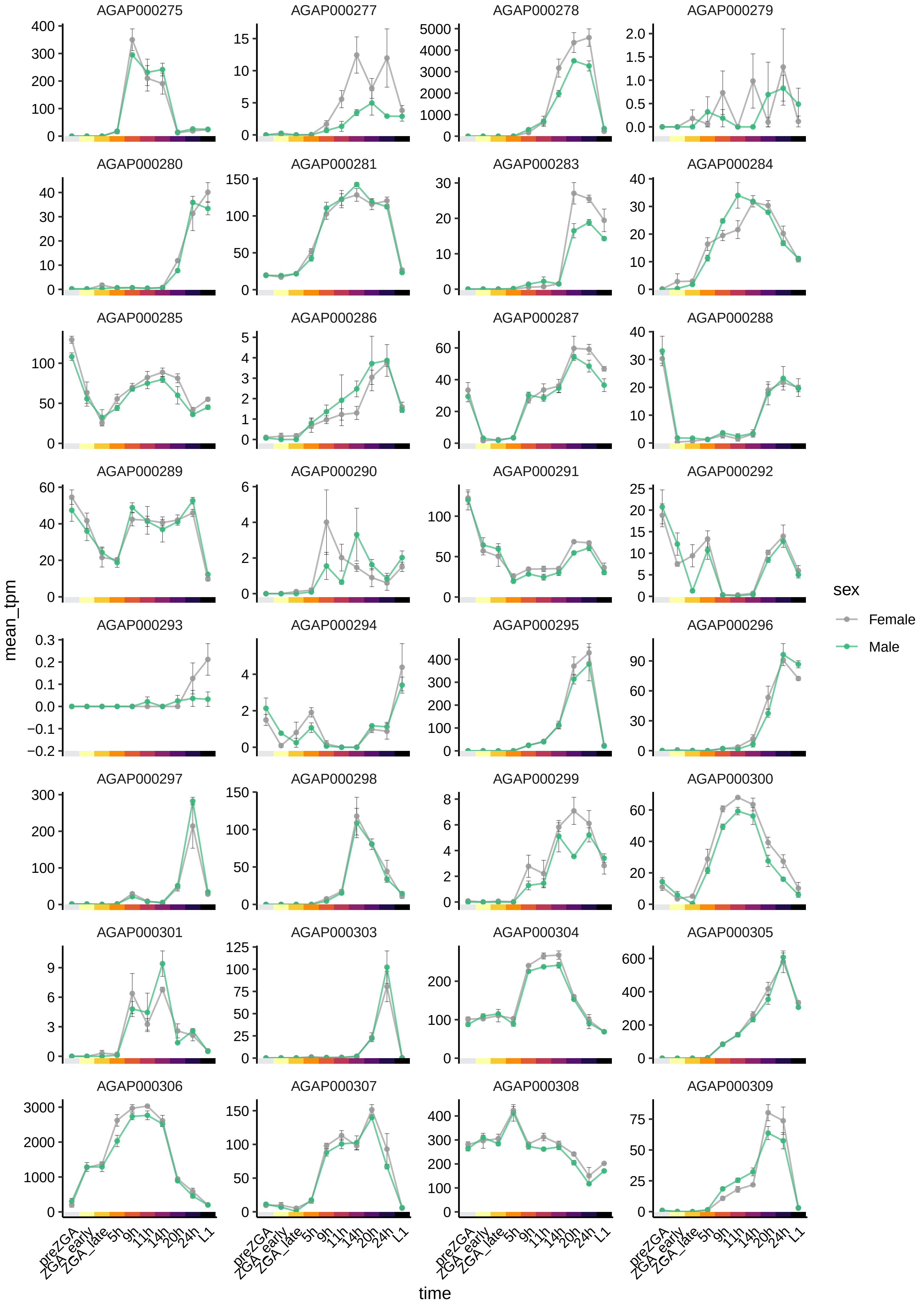
AGAP000275
AGAP000277
AGAP000278
AGAP000279
AGAP000280
AGAP000281
AGAP000283
AGAP000284
AGAP000285
AGAP000286
AGAP000287
AGAP000288
AGAP000289
AGAP000290
AGAP000291
AGAP000292
AGAP000293
AGAP000294
AGAP000295
AGAP000296
AGAP000297
AGAP000298
AGAP000299
AGAP000300
AGAP000301
AGAP000303
AGAP000304
AGAP000305
AGAP000306
AGAP000307
AGAP000308
AGAP000309
mean_tpm
time
preZGA
ZGA_early
ZGA_late
5h
9h
11h
14h
20h
24h
L1
sex
Female
Male

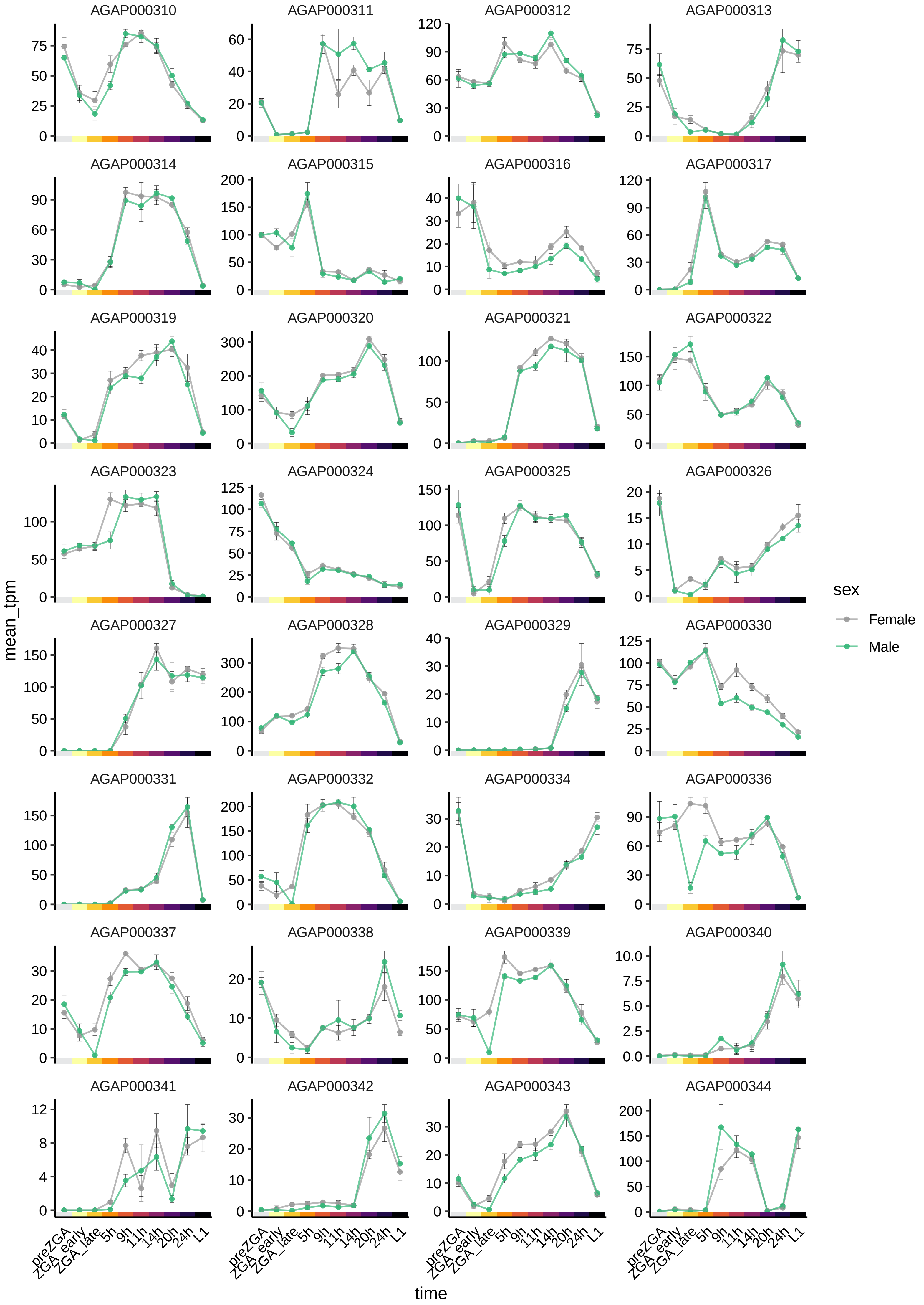

AGAP000310
AGAP000311
AGAP000312
AGAP000313
AGAP000314
AGAP000315
AGAP000316
AGAP000317
AGAP000319
AGAP000320
AGAP000321
AGAP000322
AGAP000323
AGAP000324
AGAP000325
AGAP000326
AGAP000327
AGAP000328
AGAP000329
AGAP000330
AGAP000331
AGAP000332
AGAP000334
AGAP000336
AGAP000337
AGAP000338
AGAP000339
AGAP000340
AGAP000341
AGAP000342
AGAP000343
AGAP000344
mean_tpm
time
preZGA
ZGA_early
ZGA_late
5h
9h
11h
14h
20h
24h
L1
sex
Female
Male

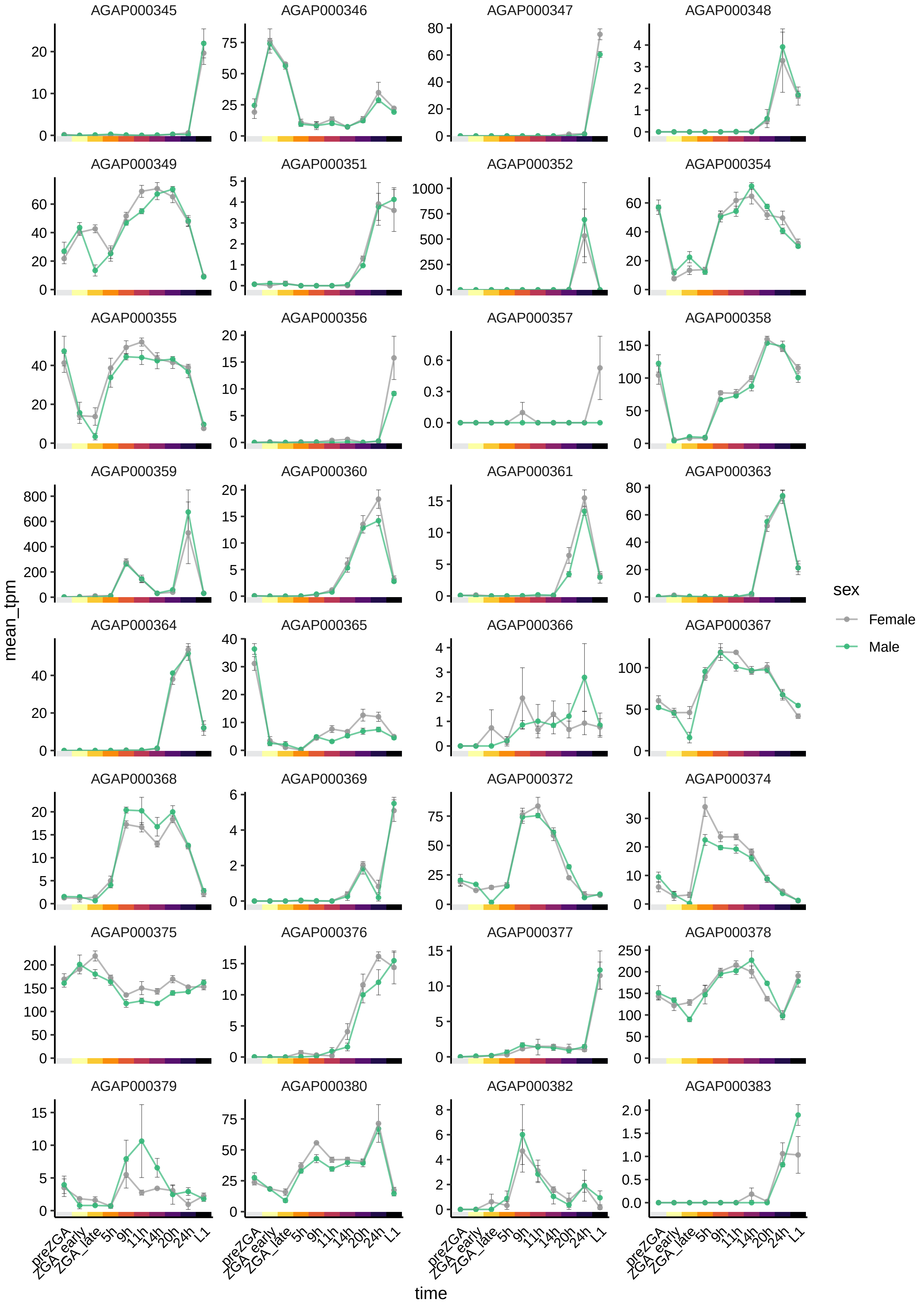
AGAP000345
AGAP000346
AGAP000347
AGAP000348
AGAP000349
AGAP000351
AGAP000352
AGAP000354
AGAP000355
AGAP000356
AGAP000357
AGAP000358
AGAP000359
AGAP000360
AGAP000361
AGAP000363
AGAP000364
AGAP000365
AGAP000366
AGAP000367
AGAP000368
AGAP000369
AGAP000372
AGAP000374
AGAP000375
AGAP000376
AGAP000377
AGAP000378
AGAP000379
AGAP000380
AGAP000382
AGAP000383
mean_tpm
time
preZGA
ZGA_early
ZGA_late
5h
9h
11h
14h
20h
24h
L1
sex
Female
Male

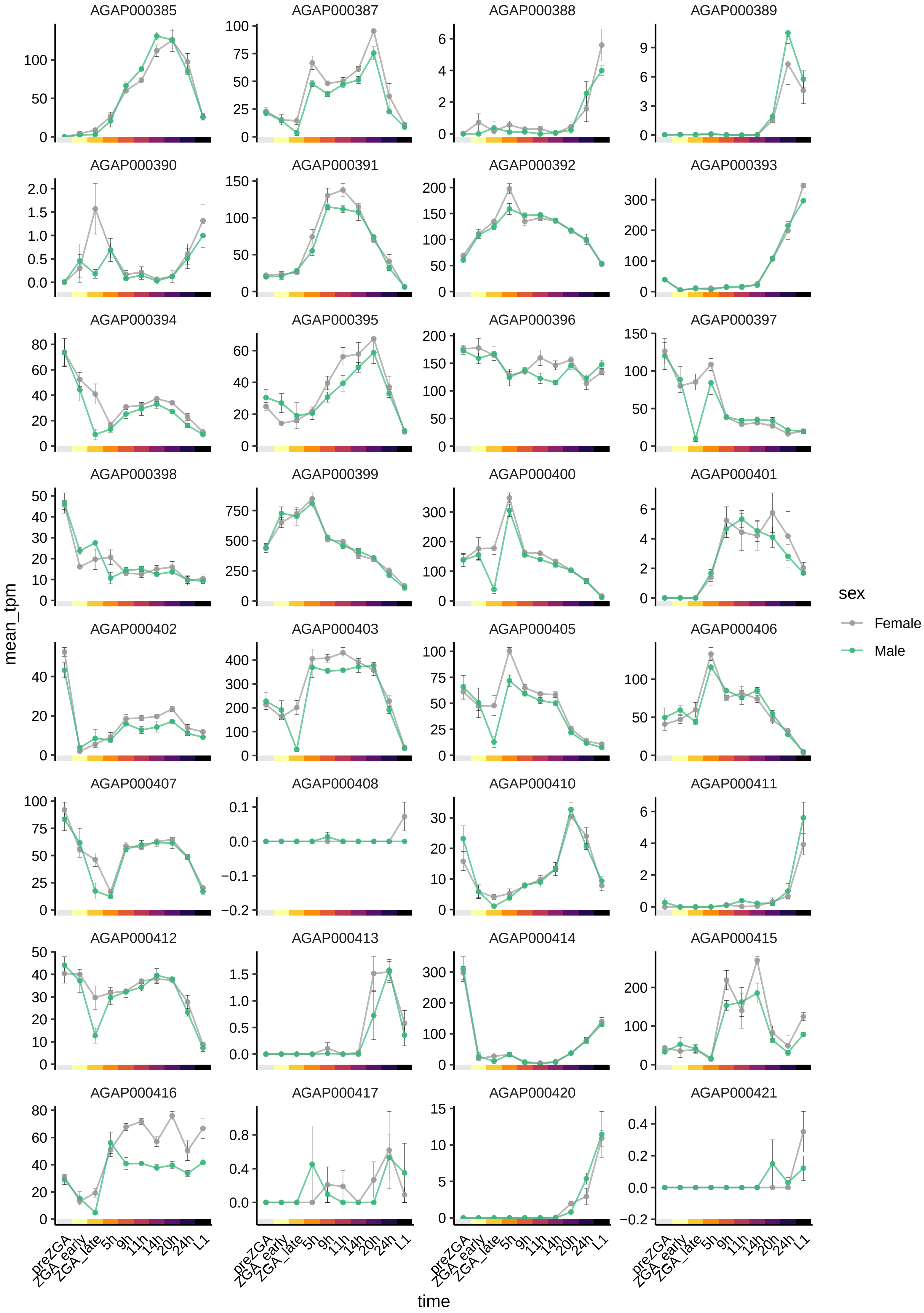

AGAP000385
AGAP000387
AGAP000388
AGAP000389
AGAP000390
AGAP000391
AGAP000392
AGAP000393
AGAP000394
AGAP000395
AGAP000396
AGAP000397
AGAP000398
AGAP000399
AGAP000400
AGAP000401
AGAP000402
AGAP000403
AGAP000405
AGAP000406
AGAP000407
AGAP000408
AGAP000410
AGAP000411
AGAP000412
AGAP000413
AGAP000414
AGAP000415
AGAP000416
AGAP000417
AGAP000420
AGAP000421
mean_tpm
time
preZGA
ZGA_early
ZGA_late
5h
9h
11h
14h
20h
24h
L1
sex
Female
Male

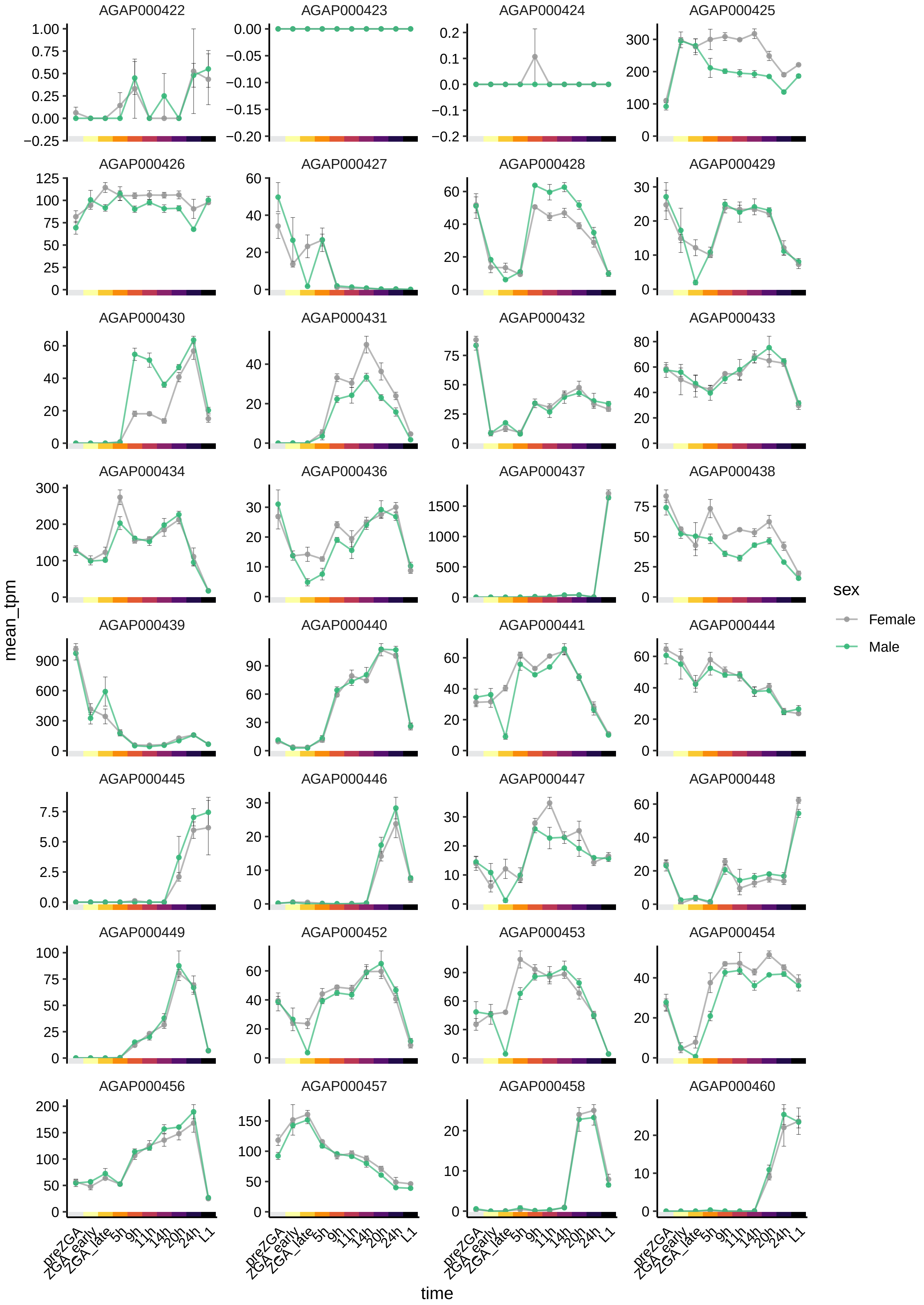
AGAP000422
AGAP000423
AGAP000424
AGAP000425
AGAP000426
AGAP000427
AGAP000428
AGAP000429
AGAP000430
AGAP000431
AGAP000432
AGAP000433
AGAP000434
AGAP000436
AGAP000437
AGAP000438
AGAP000439
AGAP000440
AGAP000441
AGAP000444
AGAP000445
AGAP000446
AGAP000447
AGAP000448
AGAP000449
AGAP000452
AGAP000453
AGAP000454
AGAP000456
AGAP000457
AGAP000458
AGAP000460
mean_tpm
time
preZGA
ZGA_early
ZGA_late
5h
9h
11h
14h
20h
24h
L1
sex
Female
Male

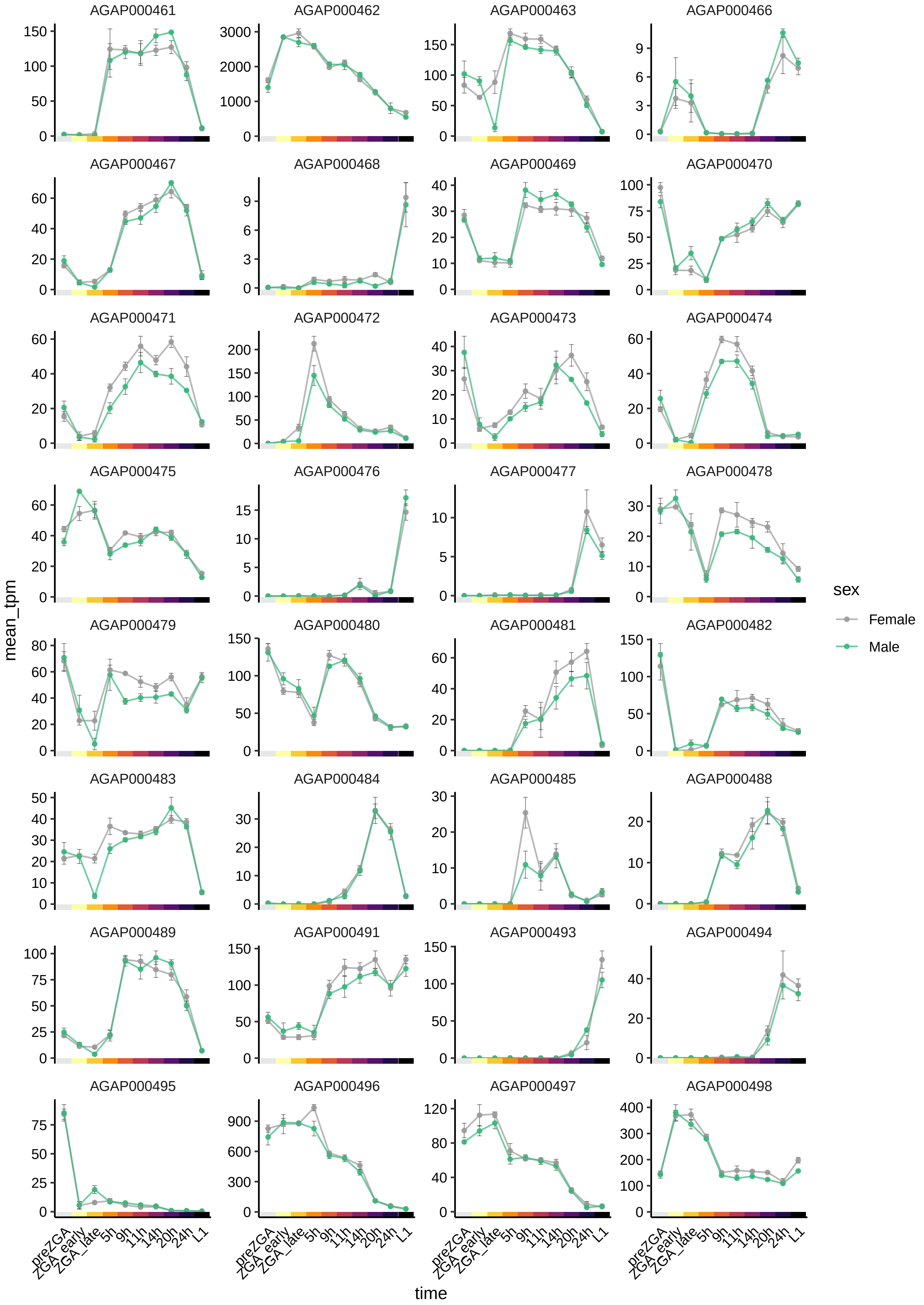
AGAP000461
AGAP000462
AGAP000463
AGAP000466
AGAP000467
AGAP000468
AGAP000469
AGAP000470
AGAP000471
AGAP000472
AGAP000473
AGAP000474
AGAP000475
AGAP000476
AGAP000477
AGAP000478
AGAP000479
AGAP000480
AGAP000481
AGAP000482
AGAP000483
AGAP000484
AGAP000485
AGAP000488
AGAP000489
AGAP000491
AGAP000493
AGAP000494
AGAP000495
AGAP000496
AGAP000497
AGAP000498
mean_tpm
time
preZGA
ZGA_early
ZGA_late
5h
9h
11h
14h
20h
24h
L1
sex
Female
Male

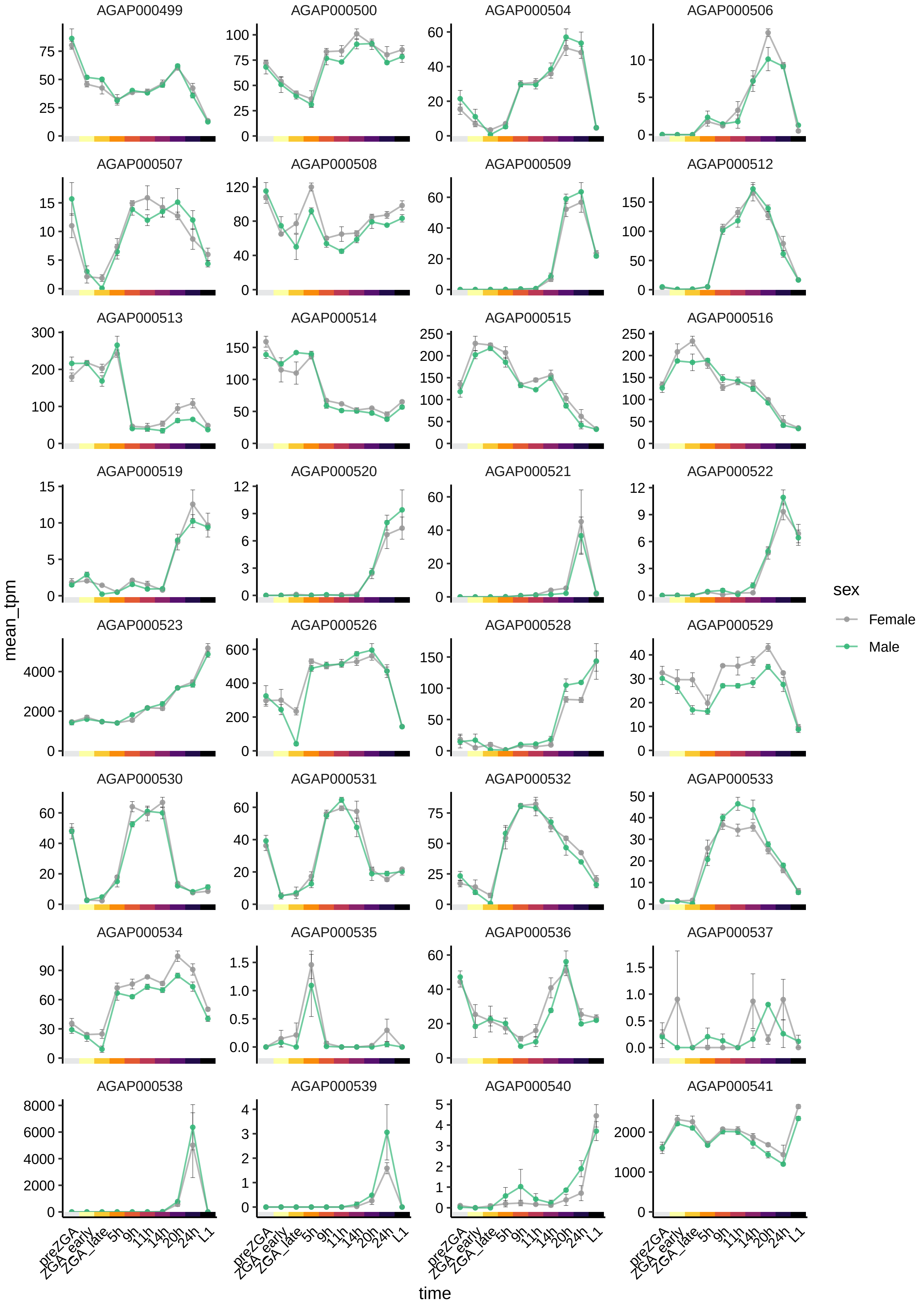

AGAP000499
AGAP000500
AGAP000504
AGAP000506
AGAP000507
AGAP000508
AGAP000509
AGAP000512
AGAP000513
AGAP000514
AGAP000515
AGAP000516
AGAP000519
AGAP000520
AGAP000521
AGAP000522
AGAP000523
AGAP000526
AGAP000528
AGAP000529
AGAP000530
AGAP000531
AGAP000532
AGAP000533
AGAP000534
AGAP000535
AGAP000536
AGAP000537
AGAP000538
AGAP000539
AGAP000540
AGAP000541
mean_tpm
time
preZGA
ZGA_early
ZGA_late
5h
9h
11h
14h
20h
24h
L1
sex
Female
Male

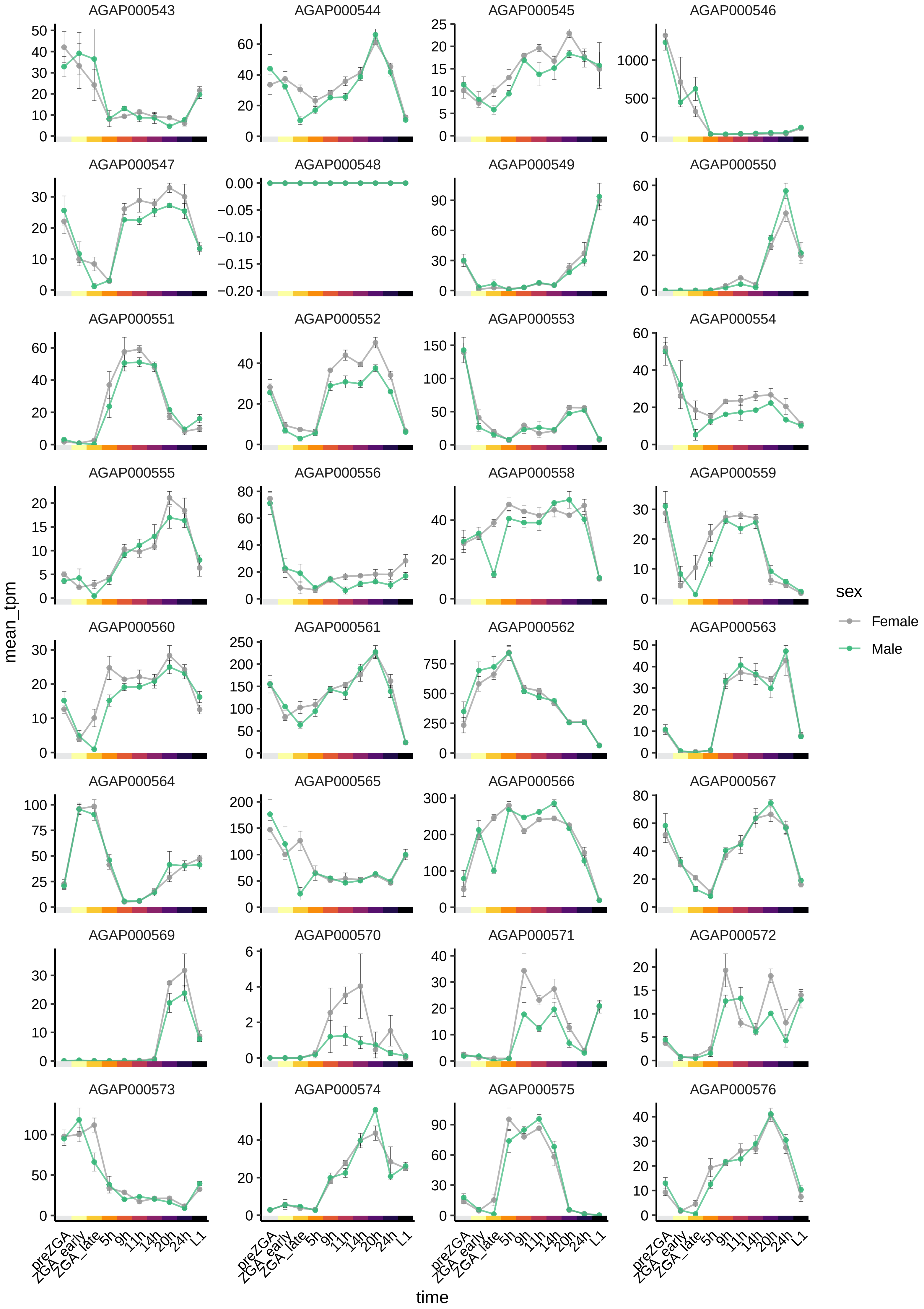

AGAP000543
AGAP000544
AGAP000545
AGAP000546
AGAP000547
AGAP000548
AGAP000549
AGAP000550
AGAP000551
AGAP000552
AGAP000553
AGAP000554
AGAP000555
AGAP000556
AGAP000558
AGAP000559
AGAP000560
AGAP000561
AGAP000562
AGAP000563
AGAP000564
AGAP000565
AGAP000566
AGAP000567
AGAP000569
AGAP000570
AGAP000571
AGAP000572
AGAP000573
AGAP000574
AGAP000575
AGAP000576
mean_tpm
time
preZGA
ZGA_early
ZGA_late
5h
9h
11h
14h
20h
24h
L1
sex
Female
Male

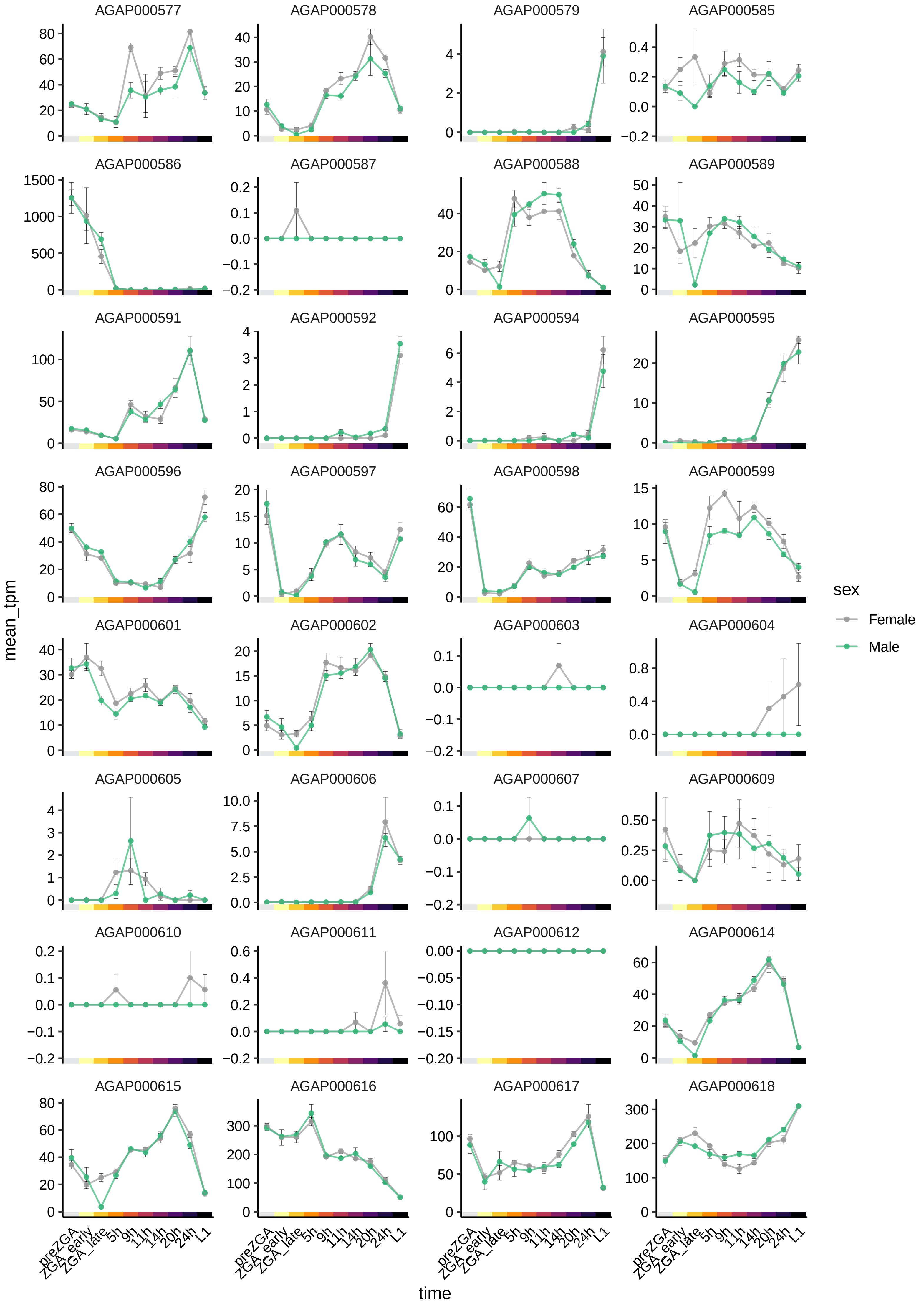

AGAP000577
AGAP000578
AGAP000579
AGAP000585
AGAP000586
AGAP000587
AGAP000588
AGAP000589
AGAP000591
AGAP000592
AGAP000594
AGAP000595
AGAP000596
AGAP000597
AGAP000598
AGAP000599
AGAP000601
AGAP000602
AGAP000603
AGAP000604
AGAP000605
AGAP000606
AGAP000607
AGAP000609
AGAP000610
AGAP000611
AGAP000612
AGAP000614
AGAP000615
AGAP000616
AGAP000617
AGAP000618
mean_tpm
time
preZGA
ZGA_early
ZGA_late
5h
9h
11h
14h
20h
24h
L1
sex
Female
Male

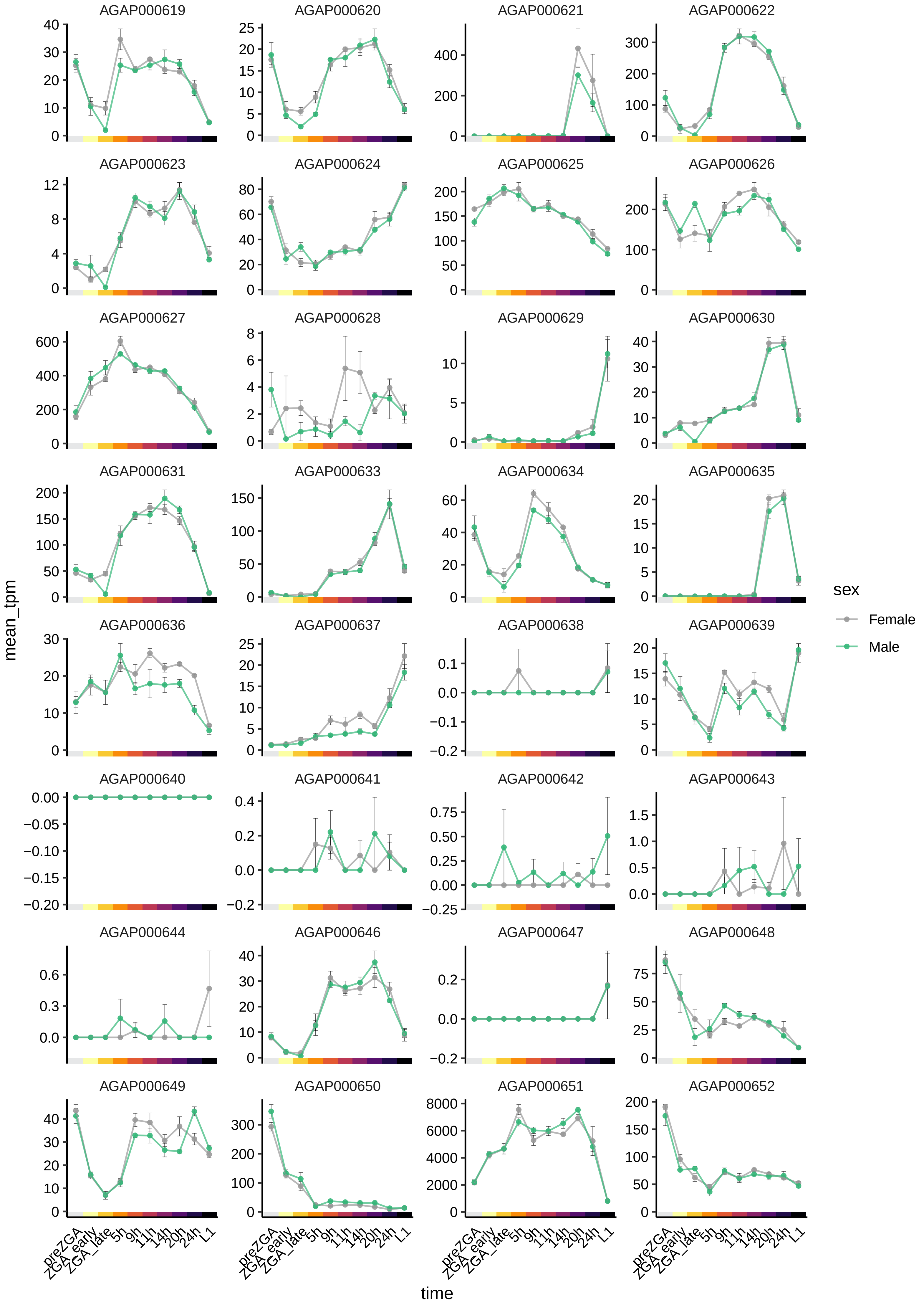
AGAP000619
AGAP000620
AGAP000621
AGAP000622
AGAP000623
AGAP000624
AGAP000625
AGAP000626
AGAP000627
AGAP000628
AGAP000629
AGAP000630
AGAP000631
AGAP000633
AGAP000634
AGAP000635
AGAP000636
AGAP000637
AGAP000638
AGAP000639
AGAP000640
AGAP000641
AGAP000642
AGAP000643
AGAP000644
AGAP000646
AGAP000647
AGAP000648
AGAP000649
AGAP000650
AGAP000651
AGAP000652
mean_tpm
time
preZGA
ZGA_early
ZGA_late
5h
9h
11h
14h
20h
24h
L1
sex
Female
Male

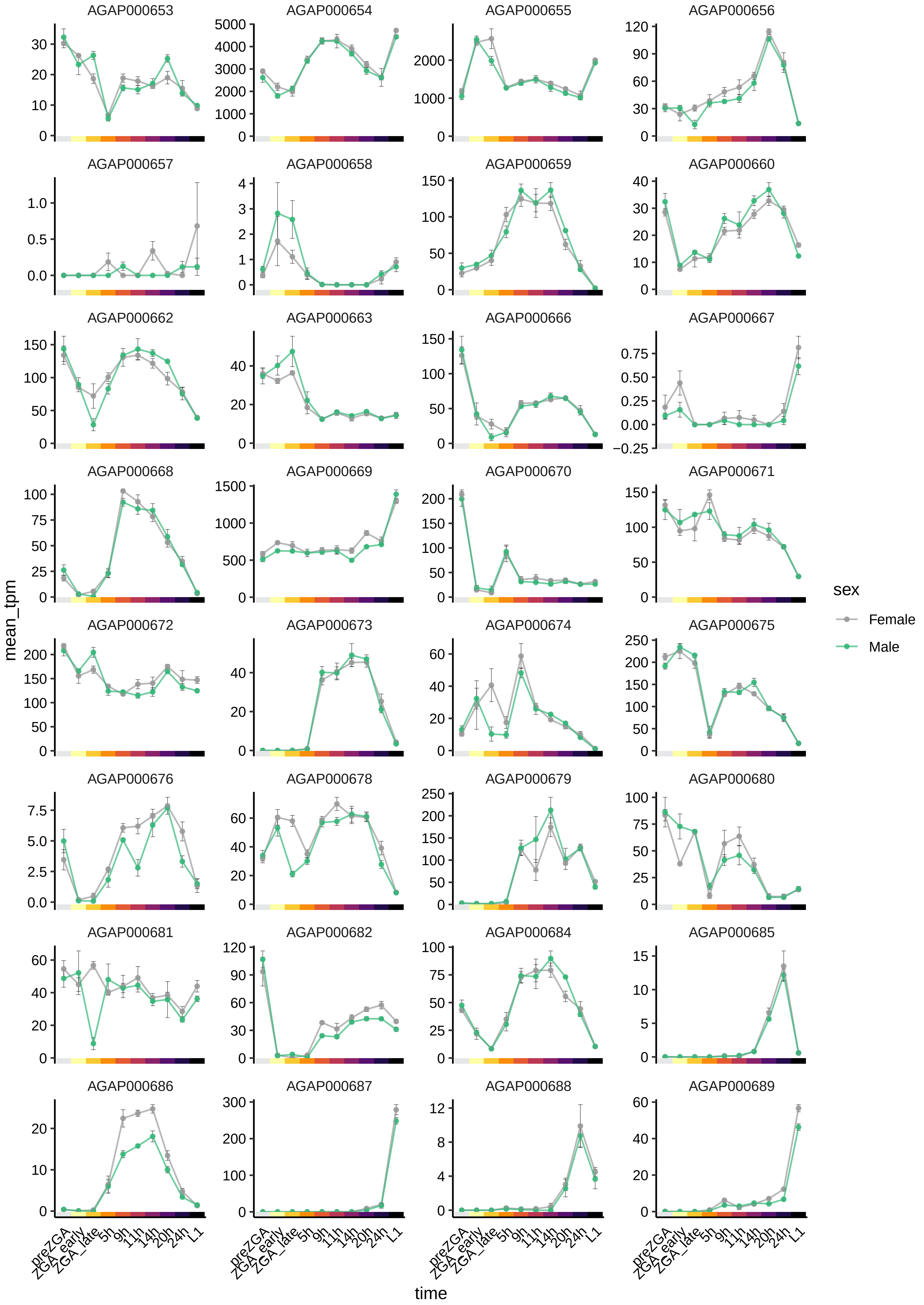

AGAP000653
AGAP000654
AGAP000655
AGAP000656
AGAP000657
AGAP000658
AGAP000659
AGAP000660
AGAP000662
AGAP000663
AGAP000666
AGAP000667
AGAP000668
AGAP000669
AGAP000670
AGAP000671
AGAP000672
AGAP000673
AGAP000674
AGAP000675
AGAP000676
AGAP000678
AGAP000679
AGAP000680
AGAP000681
AGAP000682
AGAP000684
AGAP000685
AGAP000686
AGAP000687
AGAP000688
AGAP000689
mean_tpm
time
preZGA
ZGA_early
ZGA_late
5h
9h
11h
14h
20h
24h
L1
sex
Female
Male

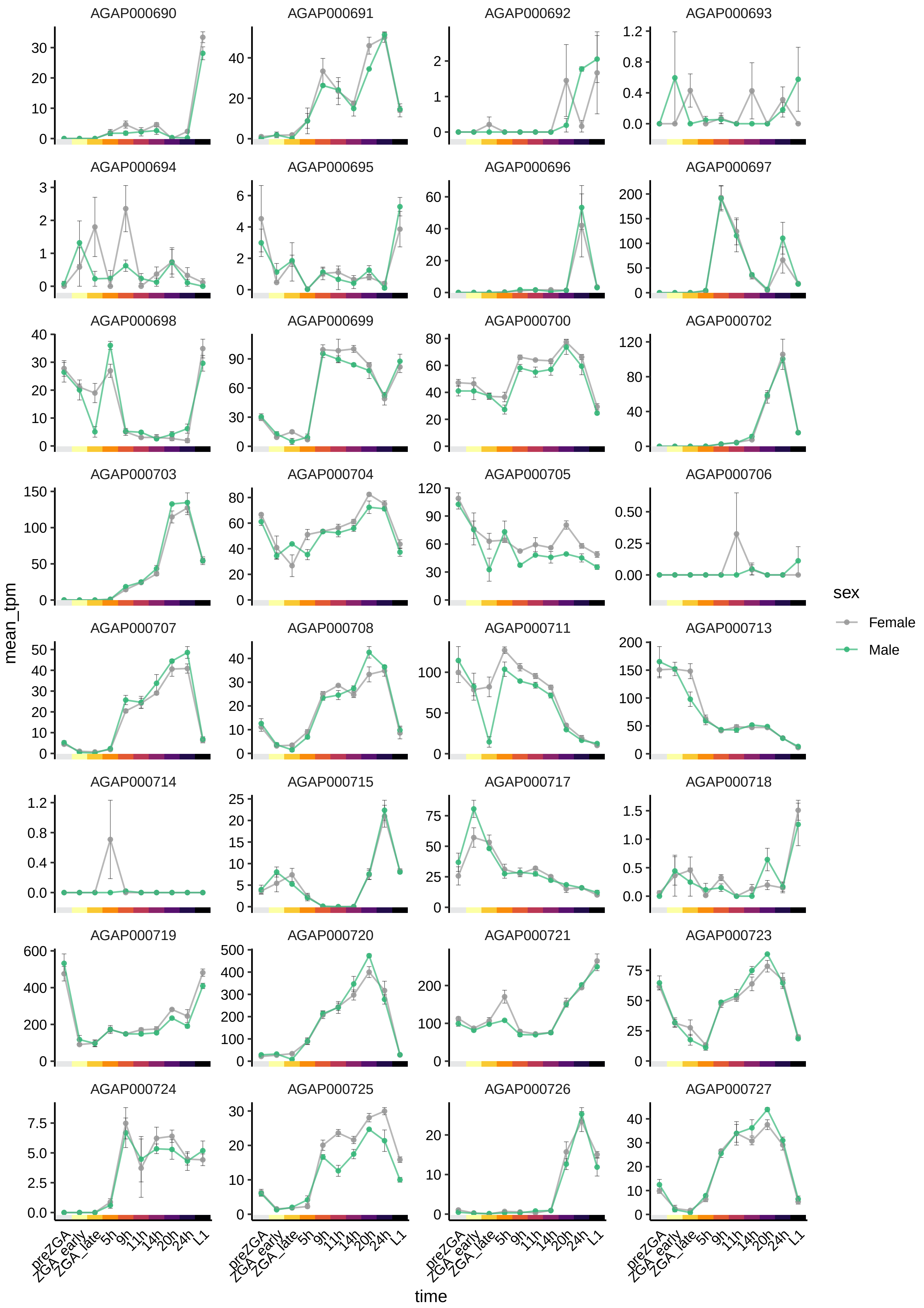
AGAP000690
AGAP000691
AGAP000692
AGAP000693
AGAP000694
AGAP000695
AGAP000696
AGAP000697
AGAP000698
AGAP000699
AGAP000700
AGAP000702
AGAP000703
AGAP000704
AGAP000705
AGAP000706
AGAP000707
AGAP000708
AGAP000711
AGAP000713
AGAP000714
AGAP000715
AGAP000717
AGAP000718
AGAP000719
AGAP000720
AGAP000721
AGAP000723
AGAP000724
AGAP000725
AGAP000726
AGAP000727
mean_tpm
time
preZGA
ZGA_early
ZGA_late
5h
9h
11h
14h
20h
24h
L1
sex
Female
Male

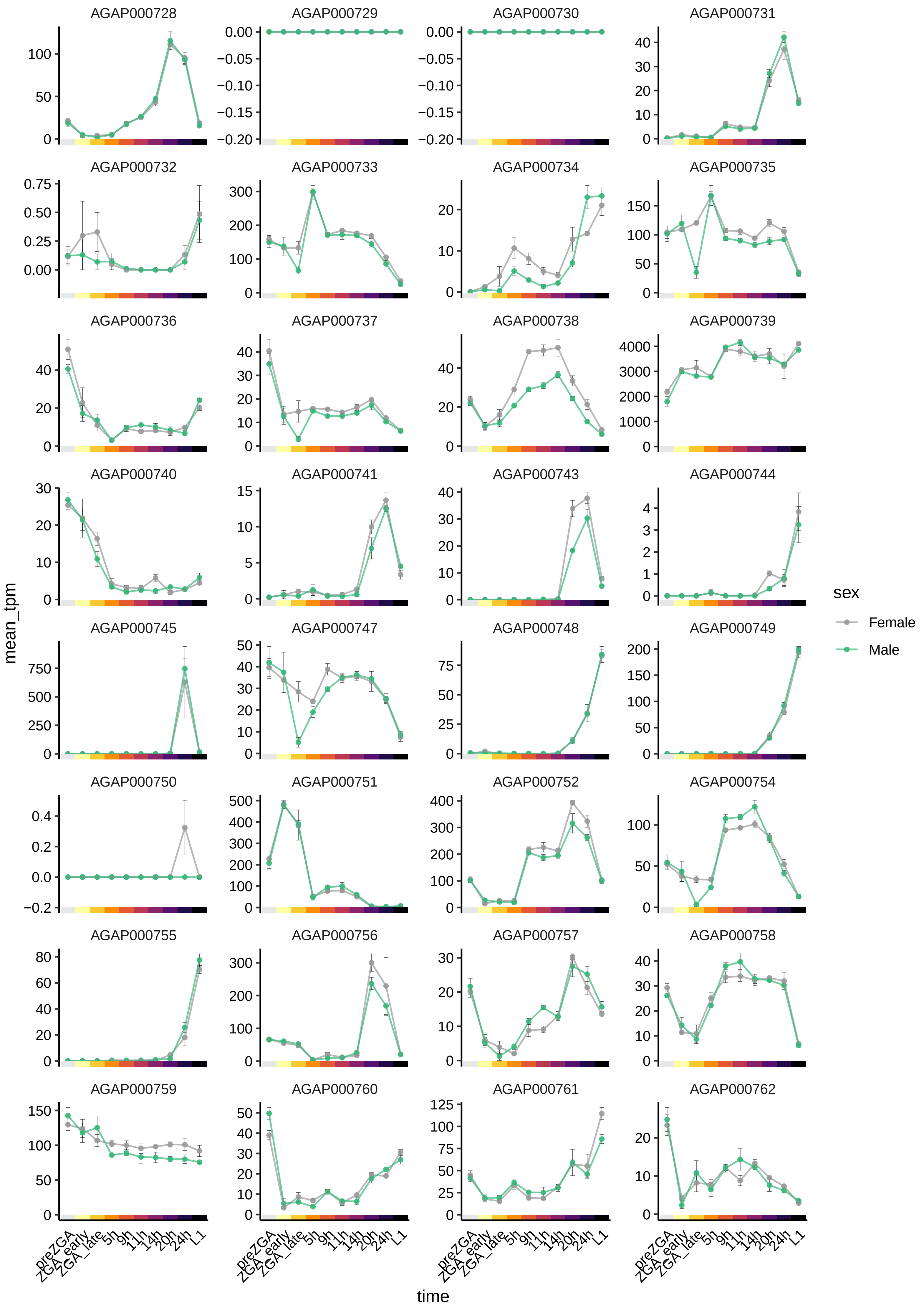

AGAP000728
AGAP000729
AGAP000730
AGAP000731
AGAP000732
AGAP000733
AGAP000734
AGAP000735
AGAP000736
AGAP000737
AGAP000738
AGAP000739
AGAP000740
AGAP000741
AGAP000743
AGAP000744
AGAP000745
AGAP000747
AGAP000748
AGAP000749
AGAP000750
AGAP000751
AGAP000752
AGAP000754
AGAP000755
AGAP000756
AGAP000757
AGAP000758
AGAP000759
AGAP000760
AGAP000761
AGAP000762
mean_tpm
time
preZGA
ZGA_early
ZGA_late
5h
9h
11h
14h
20h
24h
L1
sex
Female
Male

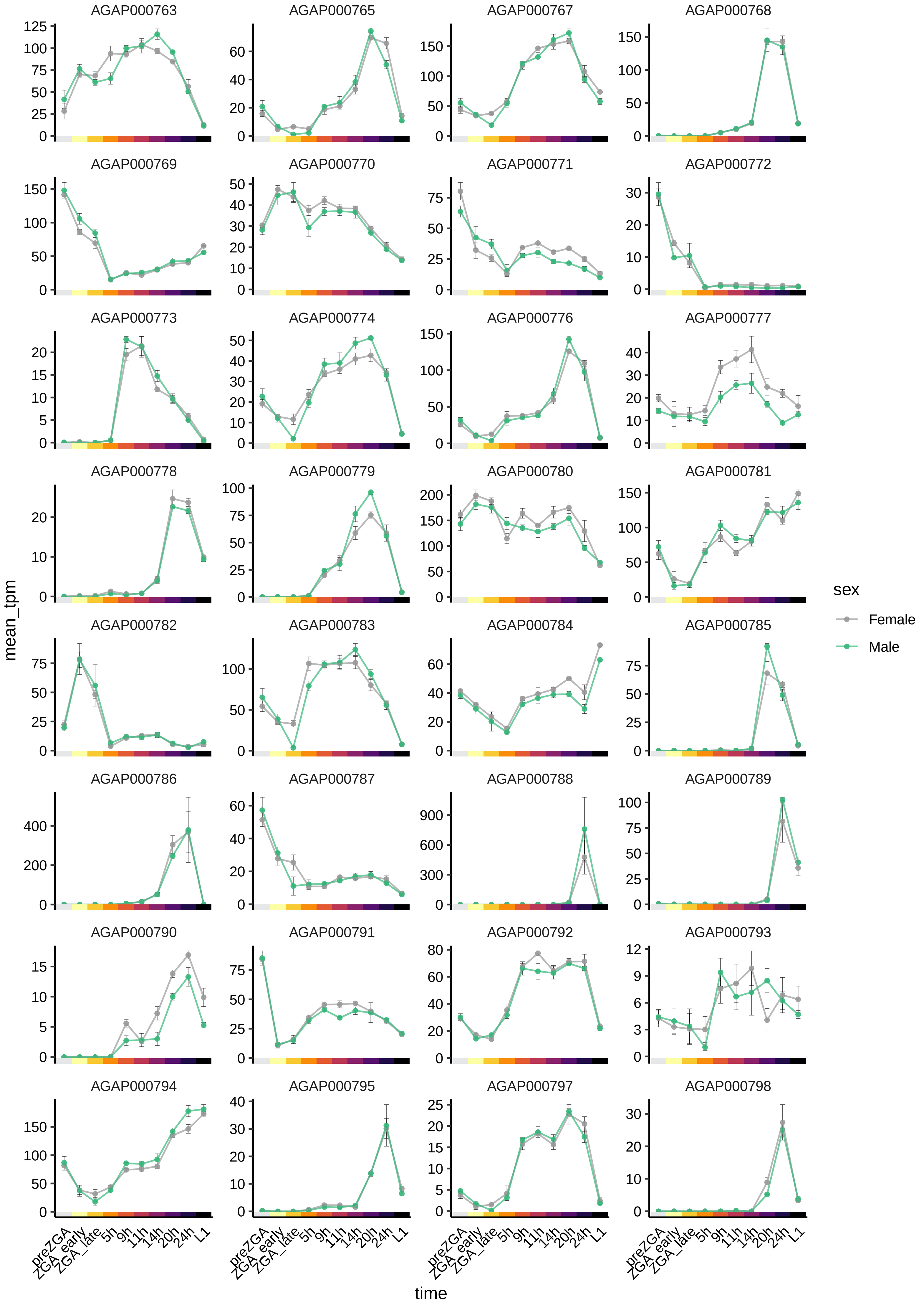

AGAP000763
AGAP000765
AGAP000767
AGAP000768
AGAP000769
AGAP000770
AGAP000771
AGAP000772
AGAP000773
AGAP000774
AGAP000776
AGAP000777
AGAP000778
AGAP000779
AGAP000780
AGAP000781
AGAP000782
AGAP000783
AGAP000784
AGAP000785
AGAP000786
AGAP000787
AGAP000788
AGAP000789
AGAP000790
AGAP000791
AGAP000792
AGAP000793
AGAP000794
AGAP000795
AGAP000797
AGAP000798
mean_tpm
time
preZGA
ZGA_early
ZGA_late
5h
9h
11h
14h
20h
24h
L1
sex
Female
Male

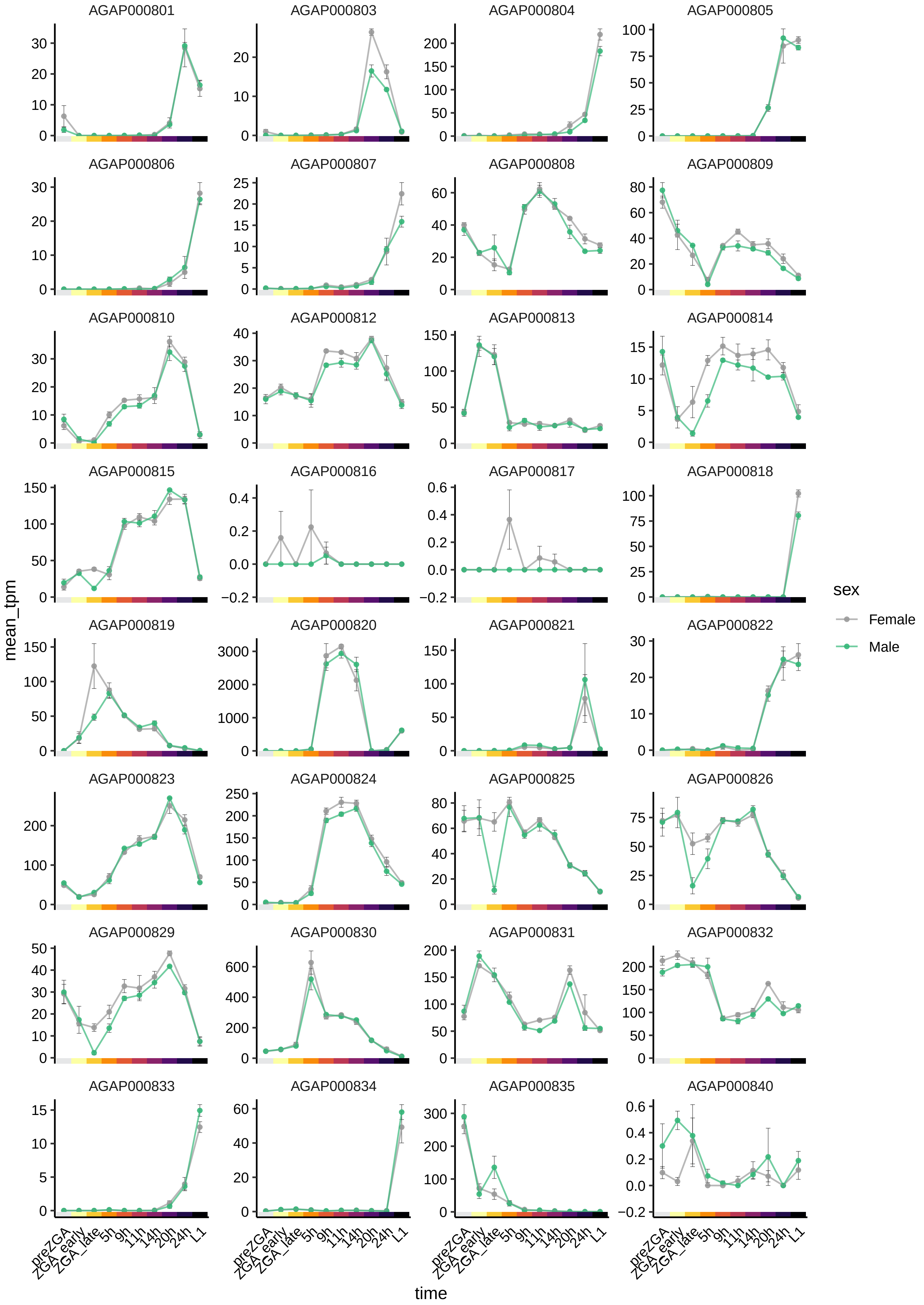
AGAP000801
AGAP000803
AGAP000804
AGAP000805
AGAP000806
AGAP000807
AGAP000808
AGAP000809
AGAP000810
AGAP000812
AGAP000813
AGAP000814
AGAP000815
AGAP000816
AGAP000817
AGAP000818
AGAP000819
AGAP000820
AGAP000821
AGAP000822
AGAP000823
AGAP000824
AGAP000825
AGAP000826
AGAP000829
AGAP000830
AGAP000831
AGAP000832
AGAP000833
AGAP000834
AGAP000835
AGAP000840
mean_tpm
time
preZGA
ZGA_early
ZGA_late
5h
9h
11h
14h
20h
24h
L1
sex
Female
Male

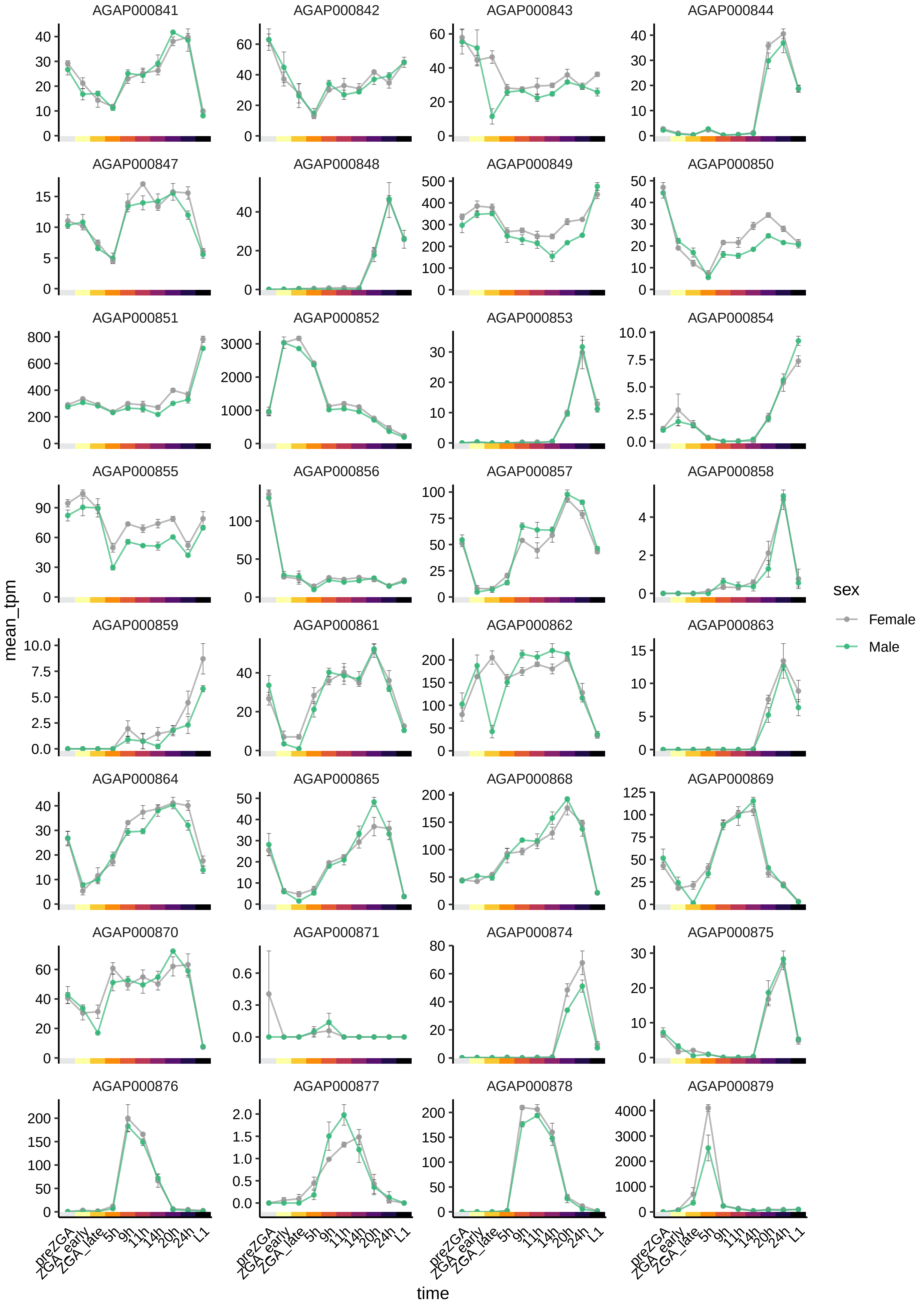

AGAP000841
AGAP000842
AGAP000843
AGAP000844
AGAP000847
AGAP000848
AGAP000849
AGAP000850
AGAP000851
AGAP000852
AGAP000853
AGAP000854
AGAP000855
AGAP000856
AGAP000857
AGAP000858
AGAP000859
AGAP000861
AGAP000862
AGAP000863
AGAP000864
AGAP000865
AGAP000868
AGAP000869
AGAP000870
AGAP000871
AGAP000874
AGAP000875
AGAP000876
AGAP000877
AGAP000878
AGAP000879
mean_tpm
time
preZGA
ZGA_early
ZGA_late
5h
9h
11h
14h
20h
24h
L1
sex
Female
Male

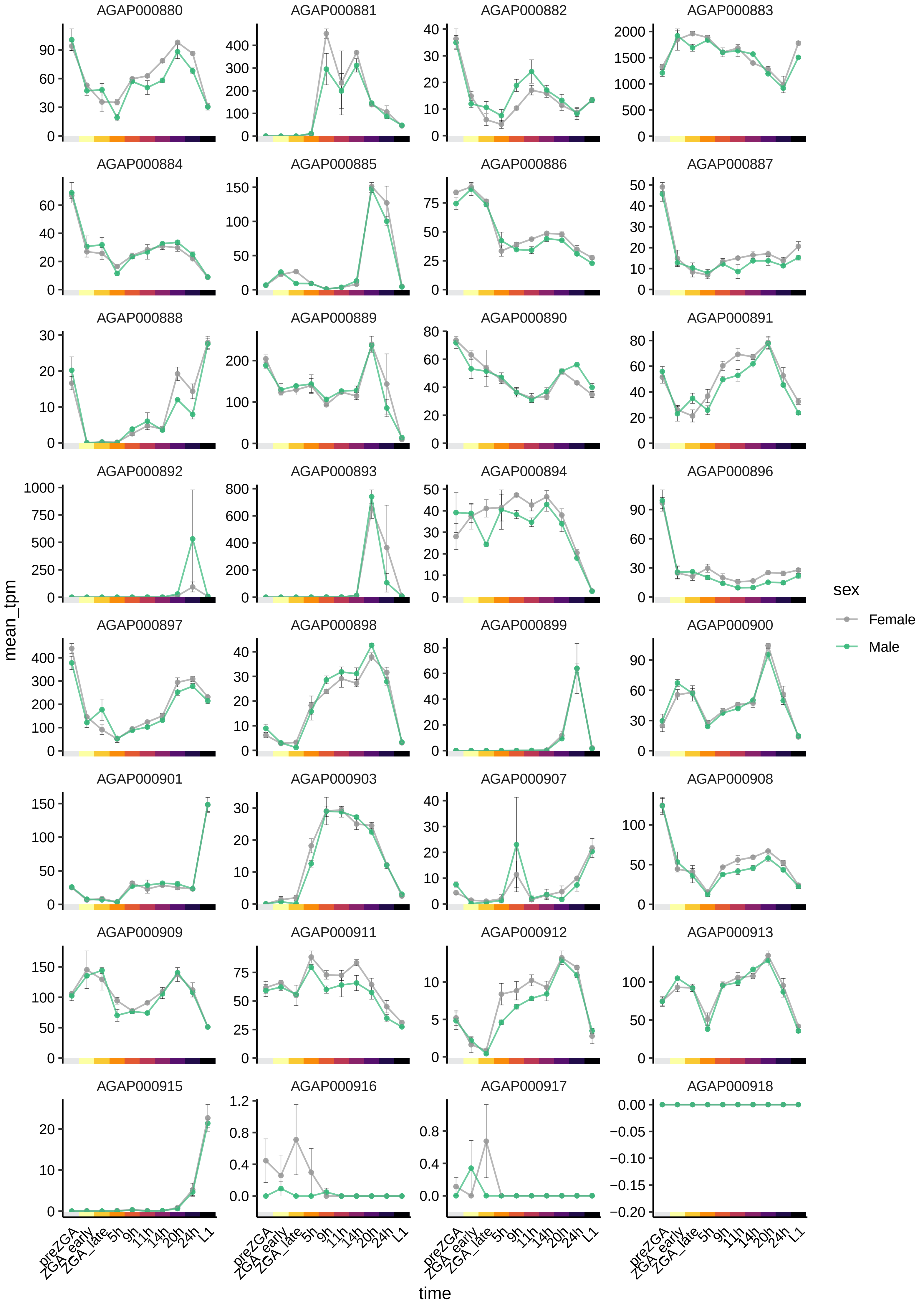

AGAP000880
AGAP000881
AGAP000882
AGAP000883
AGAP000884
AGAP000885
AGAP000886
AGAP000887
AGAP000888
AGAP000889
AGAP000890
AGAP000891
AGAP000892
AGAP000893
AGAP000894
AGAP000896
AGAP000897
AGAP000898
AGAP000899
AGAP000900
AGAP000901
AGAP000903
AGAP000907
AGAP000908
AGAP000909
AGAP000911
AGAP000912
AGAP000913
AGAP000915
AGAP000916
AGAP000917
AGAP000918
mean_tpm
time
preZGA
ZGA_early
ZGA_late
5h
9h
11h
14h
20h
24h
L1
sex
Female
Male

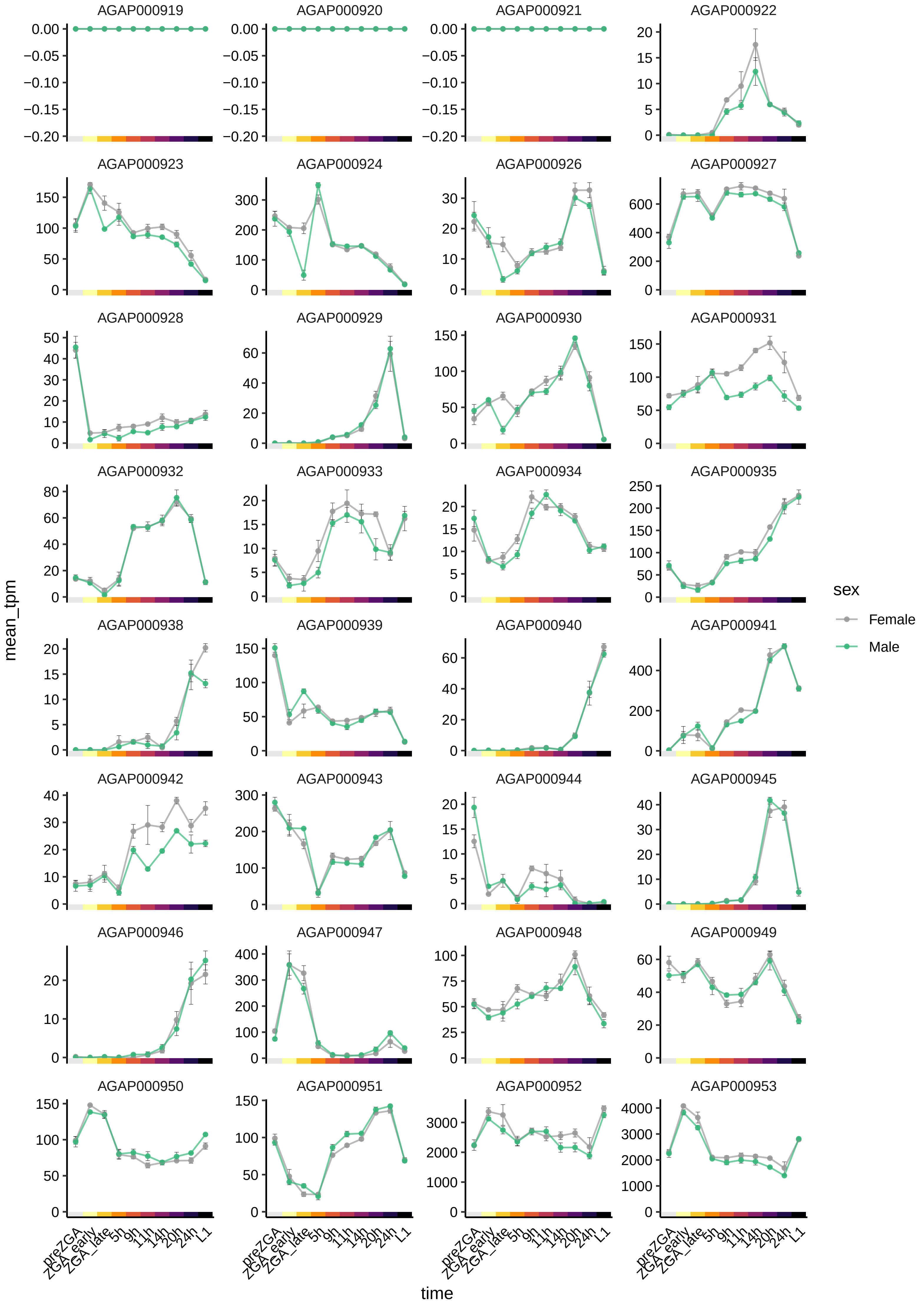

AGAP000919
AGAP000920
AGAP000921
AGAP000922
AGAP000923
AGAP000924
AGAP000926
AGAP000927
AGAP000928
AGAP000929
AGAP000930
AGAP000931
AGAP000932
AGAP000933
AGAP000934
AGAP000935
AGAP000938
AGAP000939
AGAP000940
AGAP000941
AGAP000942
AGAP000943
AGAP000944
AGAP000945
AGAP000946
AGAP000947
AGAP000948
AGAP000949
AGAP000950
AGAP000951
AGAP000952
AGAP000953
mean_tpm
time
preZGA
ZGA_early
ZGA_late
5h
9h
11h
14h
20h
24h
L1
sex
Female
Male

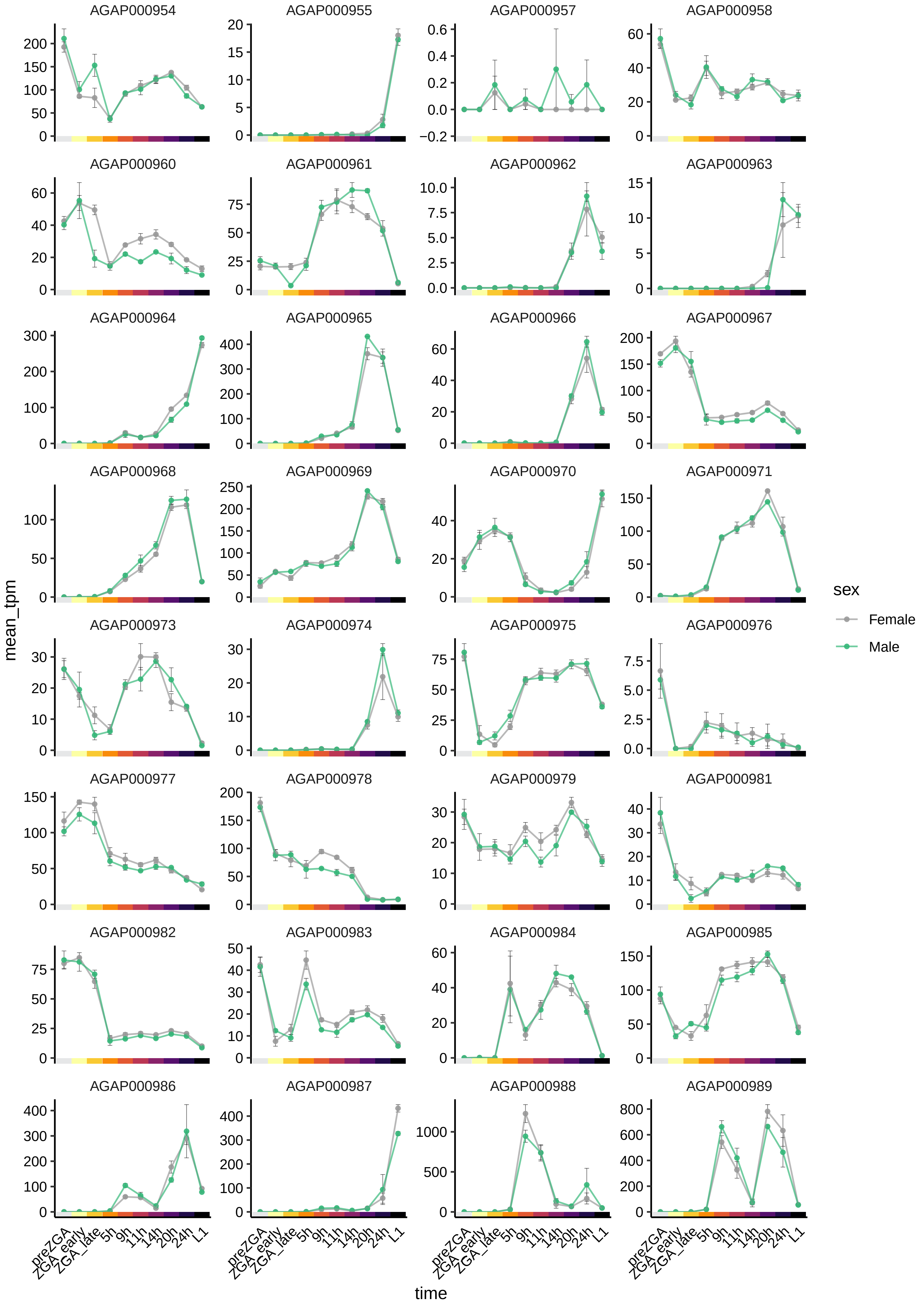
AGAP000954
AGAP000955
AGAP000957
AGAP000958
AGAP000960
AGAP000961
AGAP000962
AGAP000963
AGAP000964
AGAP000965
AGAP000966
AGAP000967
AGAP000968
AGAP000969
AGAP000970
AGAP000971
AGAP000973
AGAP000974
AGAP000975
AGAP000976
AGAP000977
AGAP000978
AGAP000979
AGAP000981
AGAP000982
AGAP000983
AGAP000984
AGAP000985
AGAP000986
AGAP000987
AGAP000988
AGAP000989
mean_tpm
preZGA
ZGA_early
ZGA_late
5h
9h
11h
14h
20h
24h
L1
time
sex
Female
Male

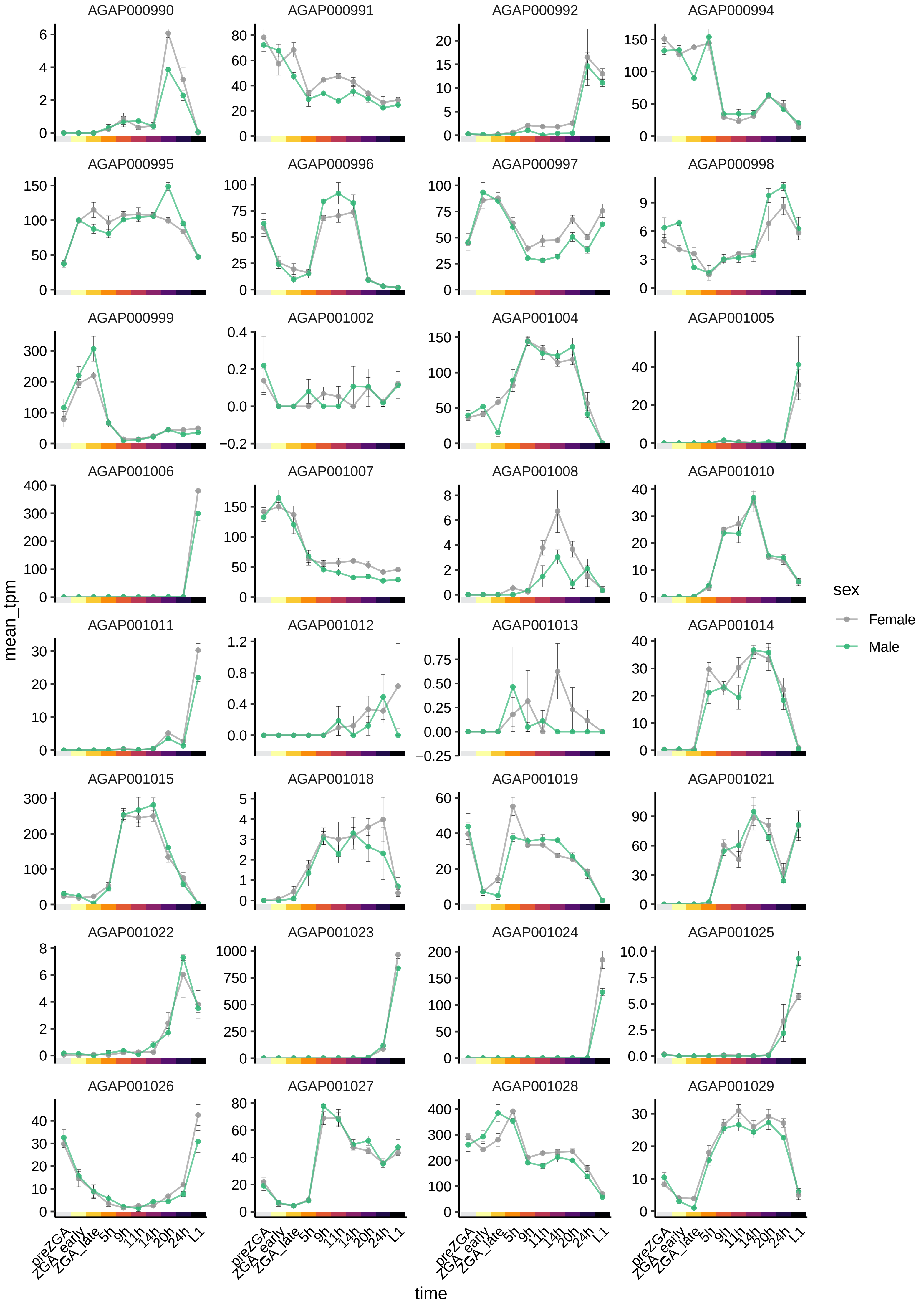

AGAP000990
AGAP000991
AGAP000992
AGAP000994
AGAP000995
AGAP000996
AGAP000997
AGAP000998
AGAP000999
AGAP001002
AGAP001004
AGAP001005
AGAP001006
AGAP001007
AGAP001008
AGAP001010
AGAP001011
AGAP001012
AGAP001013
AGAP001014
AGAP001015
AGAP001018
AGAP001019
AGAP001021
AGAP001022
AGAP001023
AGAP001024
AGAP001025
AGAP001026
AGAP001027
AGAP001028
AGAP001029
mean_tpm
time
preZGA
ZGA_early
ZGA_late
5h
9h
11h
14h
20h
24h
L1
sex
Female
Male

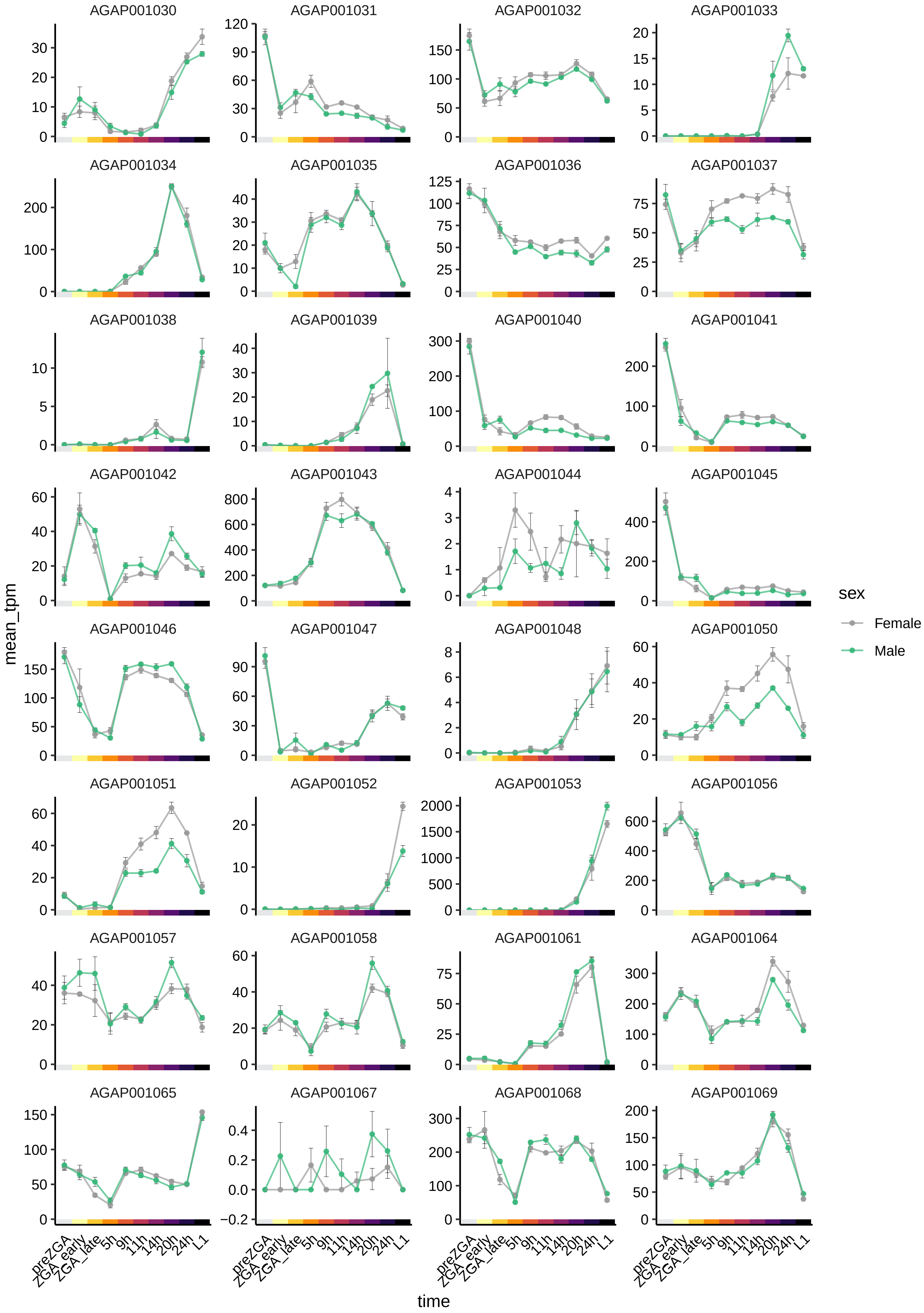

AGAP001030
AGAP001031
AGAP001032
AGAP001033
AGAP001034
AGAP001035
AGAP001036
AGAP001037
AGAP001038
AGAP001039
AGAP001040
AGAP001041
AGAP001042
AGAP001043
AGAP001044
AGAP001045
AGAP001046
AGAP001047
AGAP001048
AGAP001050
AGAP001051
AGAP001052
AGAP001053
AGAP001056
AGAP001057
AGAP001058
AGAP001061
AGAP001064
AGAP001065
AGAP001067
AGAP001068
AGAP001069
mean_tpm
time
preZGA
ZGA_early
ZGA_late
5h
9h
11h
14h
20h
24h
L1
sex
Female
Male

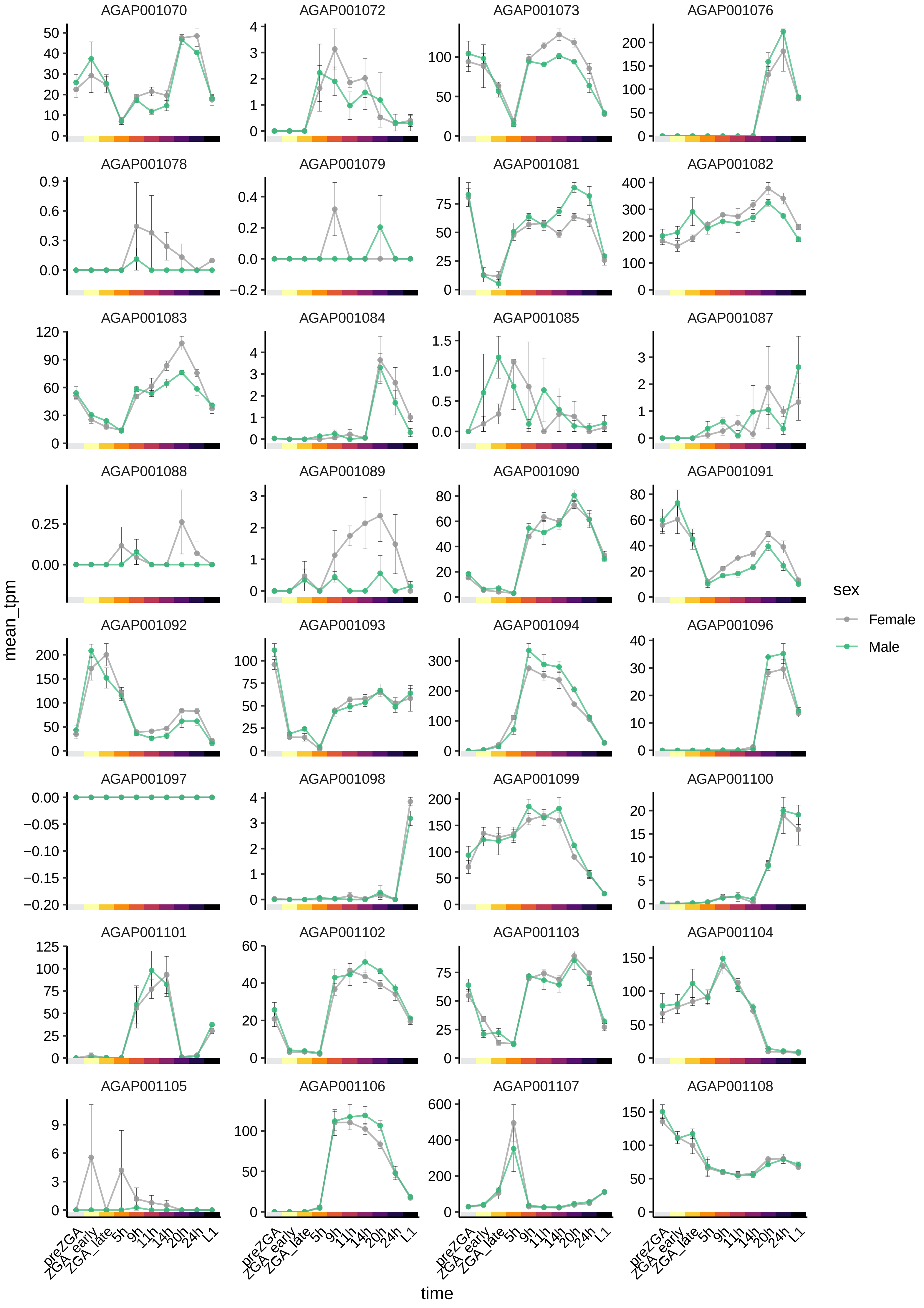
AGAP001070
AGAP001072
AGAP001073
AGAP001076
AGAP001078
AGAP001079
AGAP001081
AGAP001082
AGAP001083
AGAP001084
AGAP001085
AGAP001087
AGAP001088
AGAP001089
AGAP001090
AGAP001091
AGAP001092
AGAP001093
AGAP001094
AGAP001096
AGAP001097
AGAP001098
AGAP001099
AGAP001100
AGAP001101
AGAP001102
AGAP001103
AGAP001104
AGAP001105
AGAP001106
AGAP001107
AGAP001108
mean_tpm
time
preZGA
ZGA_early
ZGA_late
5h
9h
11h
14h
20h
24h
L1
sex
Female
Male

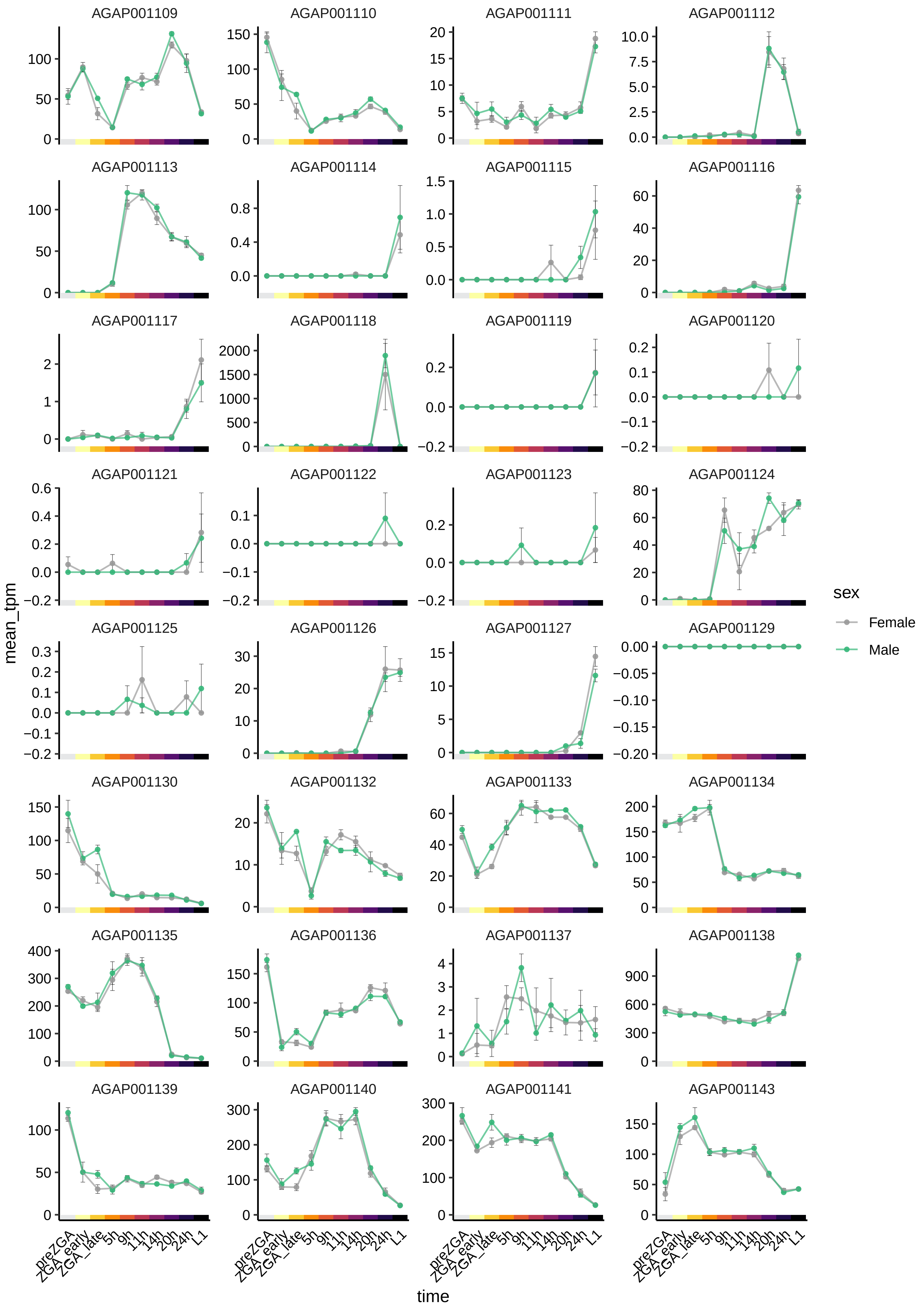

AGAP001109
AGAP001110
AGAP001111
AGAP001112
AGAP001113
AGAP001114
AGAP001115
AGAP001116
AGAP001117
AGAP001118
AGAP001119
AGAP001120
AGAP001121
AGAP001122
AGAP001123
AGAP001124
AGAP001125
AGAP001126
AGAP001127
AGAP001129
AGAP001130
AGAP001132
AGAP001133
AGAP001134
AGAP001135
AGAP001136
AGAP001137
AGAP001138
AGAP001139
AGAP001140
AGAP001141
AGAP001143
mean_tpm
time
preZGA
ZGA_early
ZGA_late
5h
9h
11h
14h
20h
24h
L1
sex
Female
Male

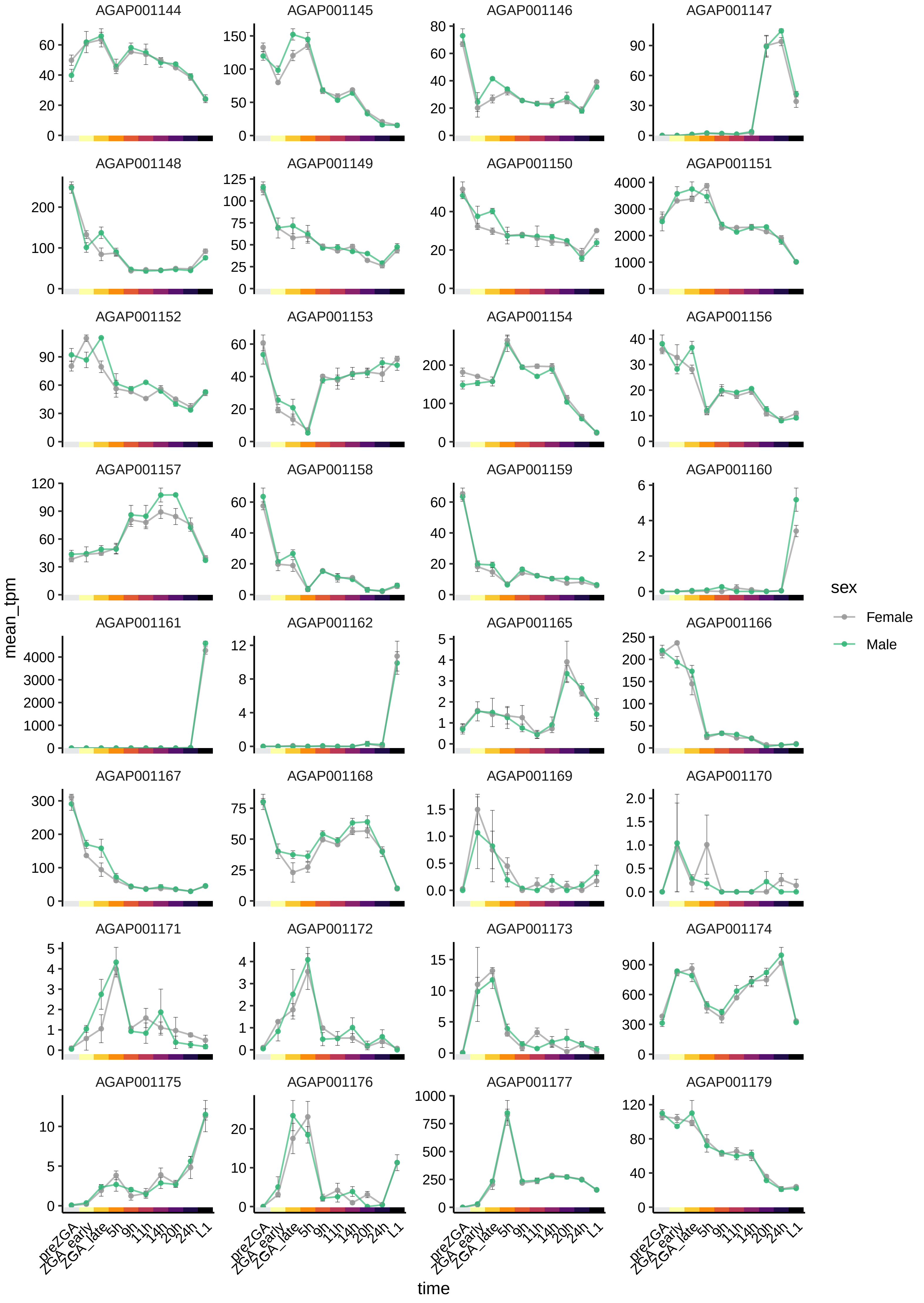

AGAP001144
AGAP001145
AGAP001146
AGAP001147
AGAP001148
AGAP001149
AGAP001150
AGAP001151
AGAP001152
AGAP001153
AGAP001154
AGAP001156
AGAP001157
AGAP001158
AGAP001159
AGAP001160
AGAP001161
AGAP001162
AGAP001165
AGAP001166
AGAP001167
AGAP001168
AGAP001169
AGAP001170
AGAP001171
AGAP001172
AGAP001173
AGAP001174
AGAP001175
AGAP001176
AGAP001177
AGAP001179
mean_tpm
time
preZGA
ZGA_early
ZGA_late
5h
9h
11h
14h
20h
24h
L1
sex
Female
Male

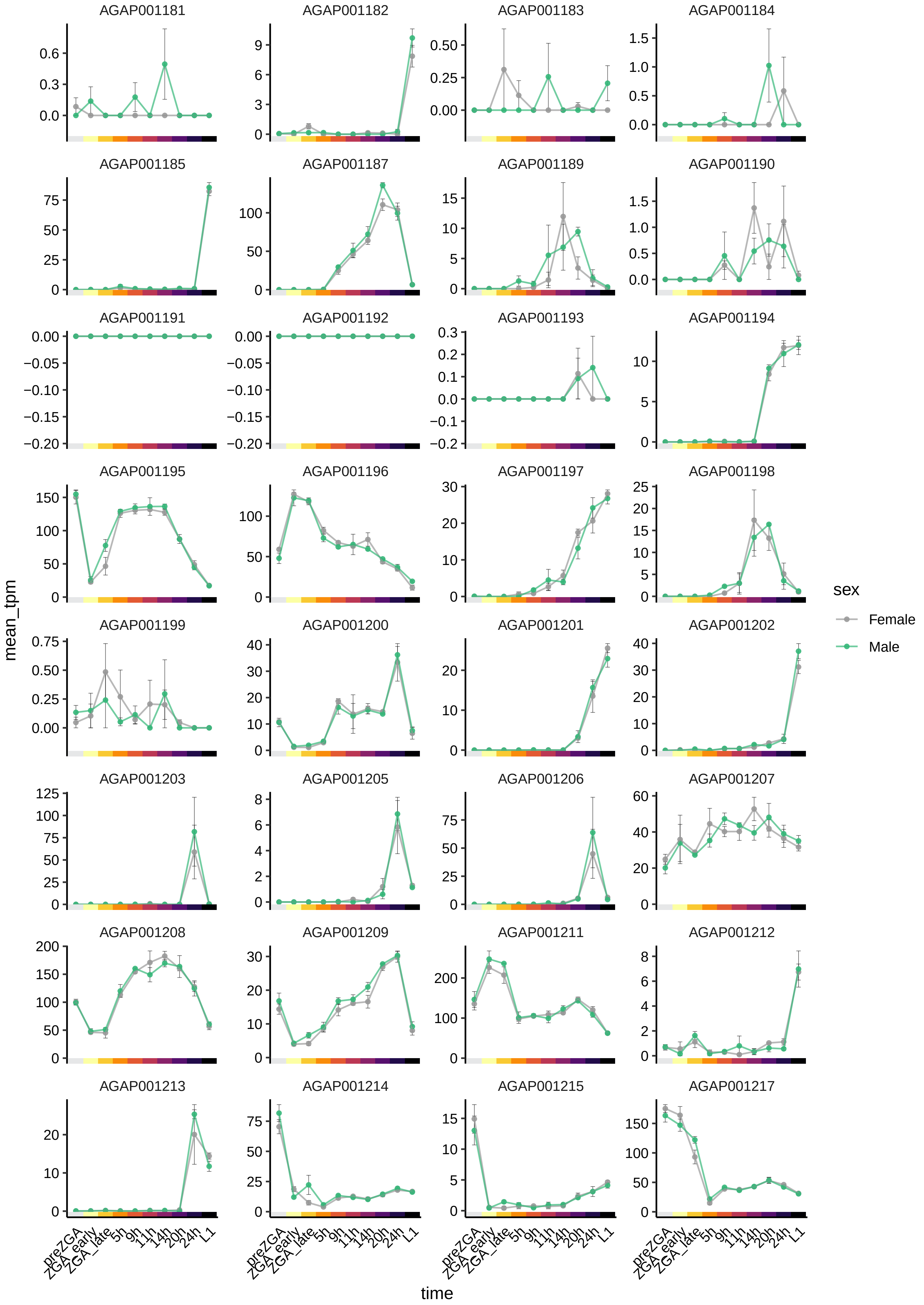
AGAP001181
AGAP001182
AGAP001183
AGAP001184
AGAP001185
AGAP001187
AGAP001189
AGAP001190
AGAP001191
AGAP001192
AGAP001193
AGAP001194
AGAP001195
AGAP001196
AGAP001197
AGAP001198
AGAP001199
AGAP001200
AGAP001201
AGAP001202
AGAP001203
AGAP001205
AGAP001206
AGAP001207
AGAP001208
AGAP001209
AGAP001211
AGAP001212
AGAP001213
AGAP001214
AGAP001215
AGAP001217
mean_tpm
time
preZGA
ZGA_early
ZGA_late
5h
9h
11h
14h
20h
24h
L1
sex
Female
Male

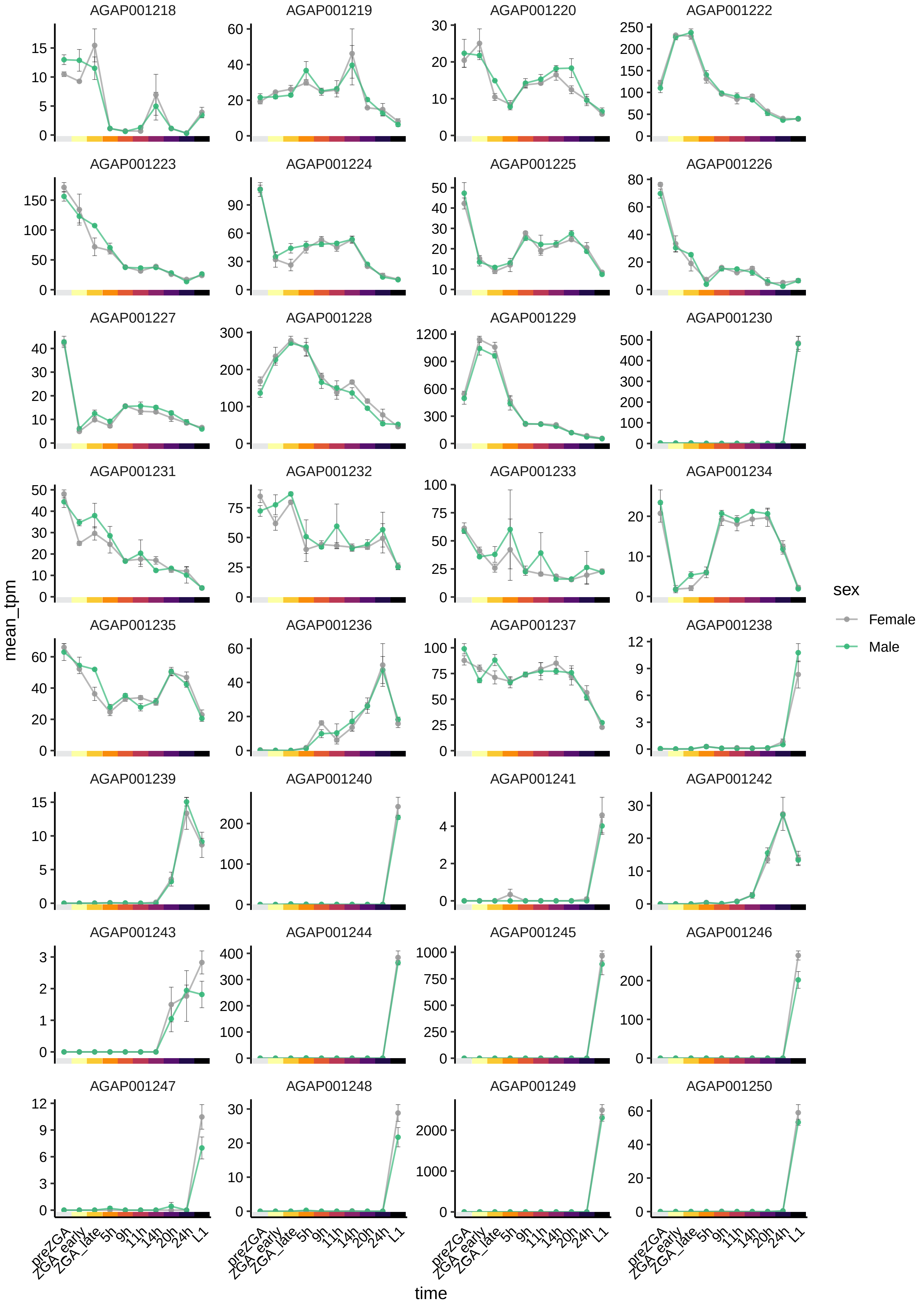

AGAP001218
AGAP001219
AGAP001220
AGAP001222
AGAP001223
AGAP001224
AGAP001225
AGAP001226
AGAP001227
AGAP001228
AGAP001229
AGAP001230
AGAP001231
AGAP001232
AGAP001233
AGAP001234
AGAP001235
AGAP001236
AGAP001237
AGAP001238
AGAP001239
AGAP001240
AGAP001241
AGAP001242
AGAP001243
AGAP001244
AGAP001245
AGAP001246
AGAP001247
AGAP001248
AGAP001249
AGAP001250
mean_tpm
time
preZGA
ZGA_early
ZGA_late
5h
9h
11h
14h
20h
24h
L1
sex
Female
Male

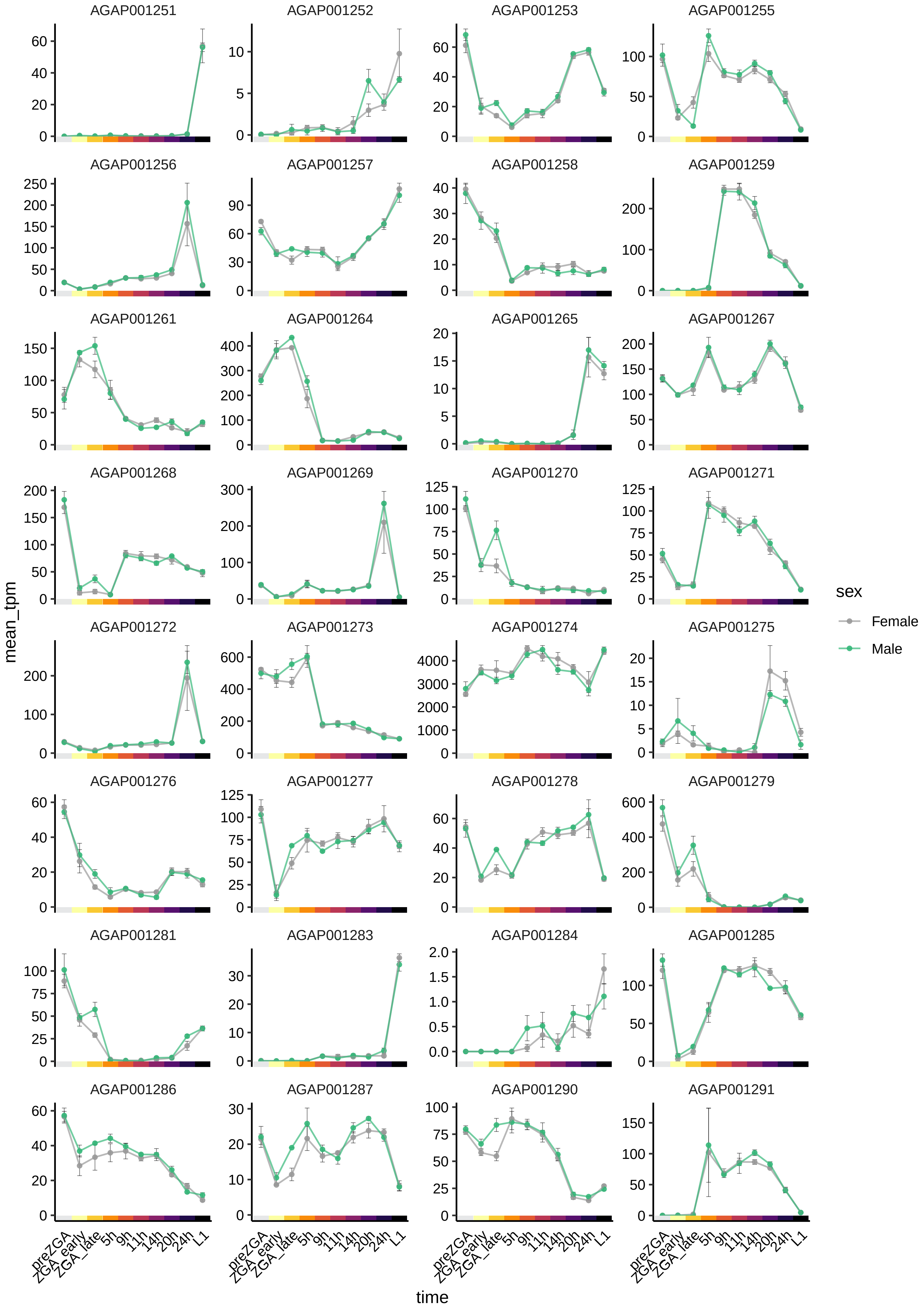

AGAP001251
AGAP001252
AGAP001253
AGAP001255
AGAP001256
AGAP001257
AGAP001258
AGAP001259
AGAP001261
AGAP001264
AGAP001265
AGAP001267
AGAP001268
AGAP001269
AGAP001270
AGAP001271
AGAP001272
AGAP001273
AGAP001274
AGAP001275
AGAP001276
AGAP001277
AGAP001278
AGAP001279
AGAP001281
AGAP001283
AGAP001284
AGAP001285
AGAP001286
AGAP001287
AGAP001290
AGAP001291
mean_tpm
time
preZGA
ZGA_early
ZGA_late
5h
9h
11h
14h
20h
24h
L1
sex
Female
Male

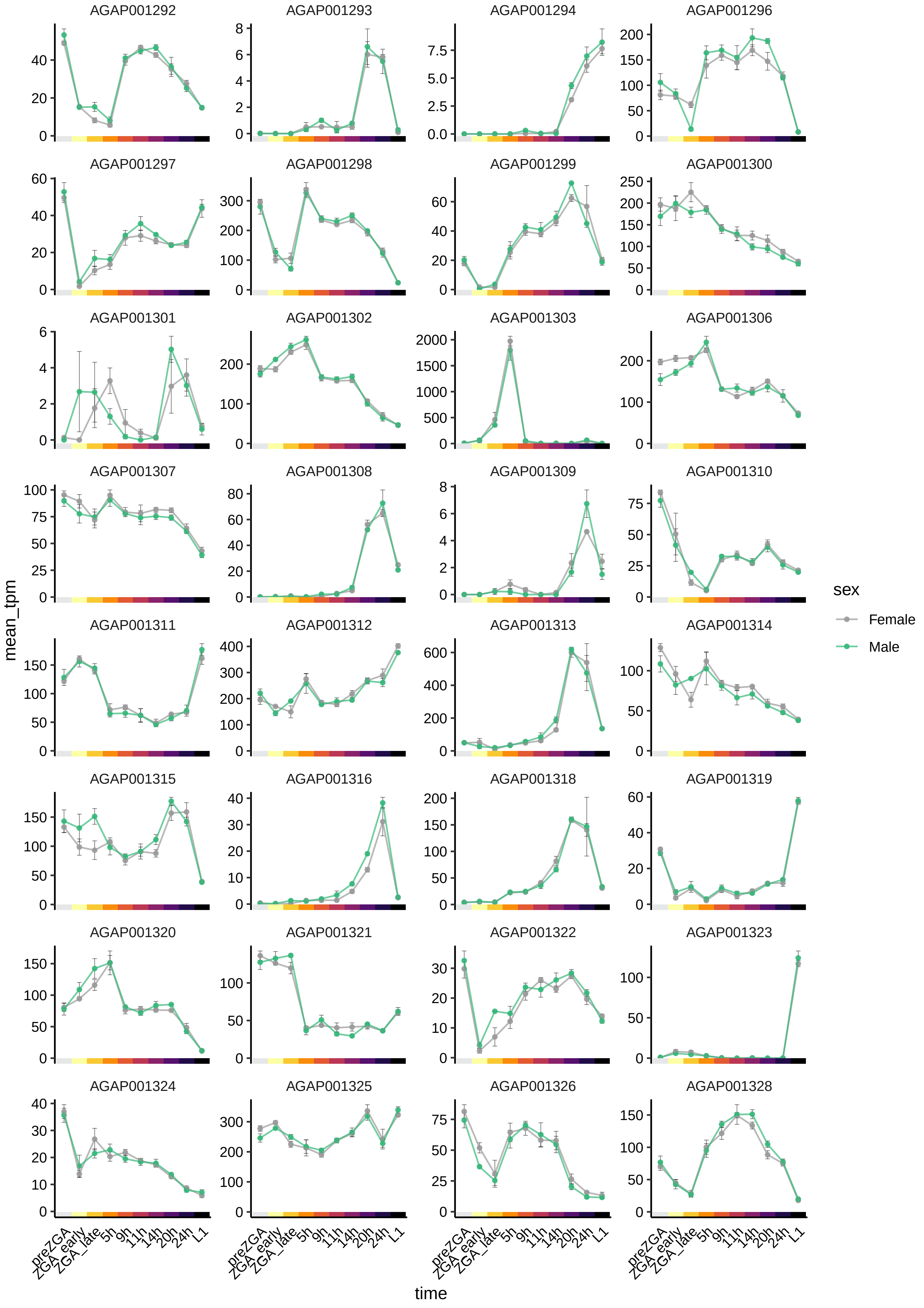
AGAP001292
AGAP001293
AGAP001294
AGAP001296
AGAP001297
AGAP001298
AGAP001299
AGAP001300
AGAP001301
AGAP001302
AGAP001303
AGAP001306
AGAP001307
AGAP001308
AGAP001309
AGAP001310
AGAP001311
AGAP001312
AGAP001313
AGAP001314
AGAP001315
AGAP001316
AGAP001318
AGAP001319
AGAP001320
AGAP001321
AGAP001322
AGAP001323
AGAP001324
AGAP001325
AGAP001326
AGAP001328
mean_tpm
time
preZGA
ZGA_early
ZGA_late
5h
9h
11h
14h
20h
24h
L1
sex
Female
Male

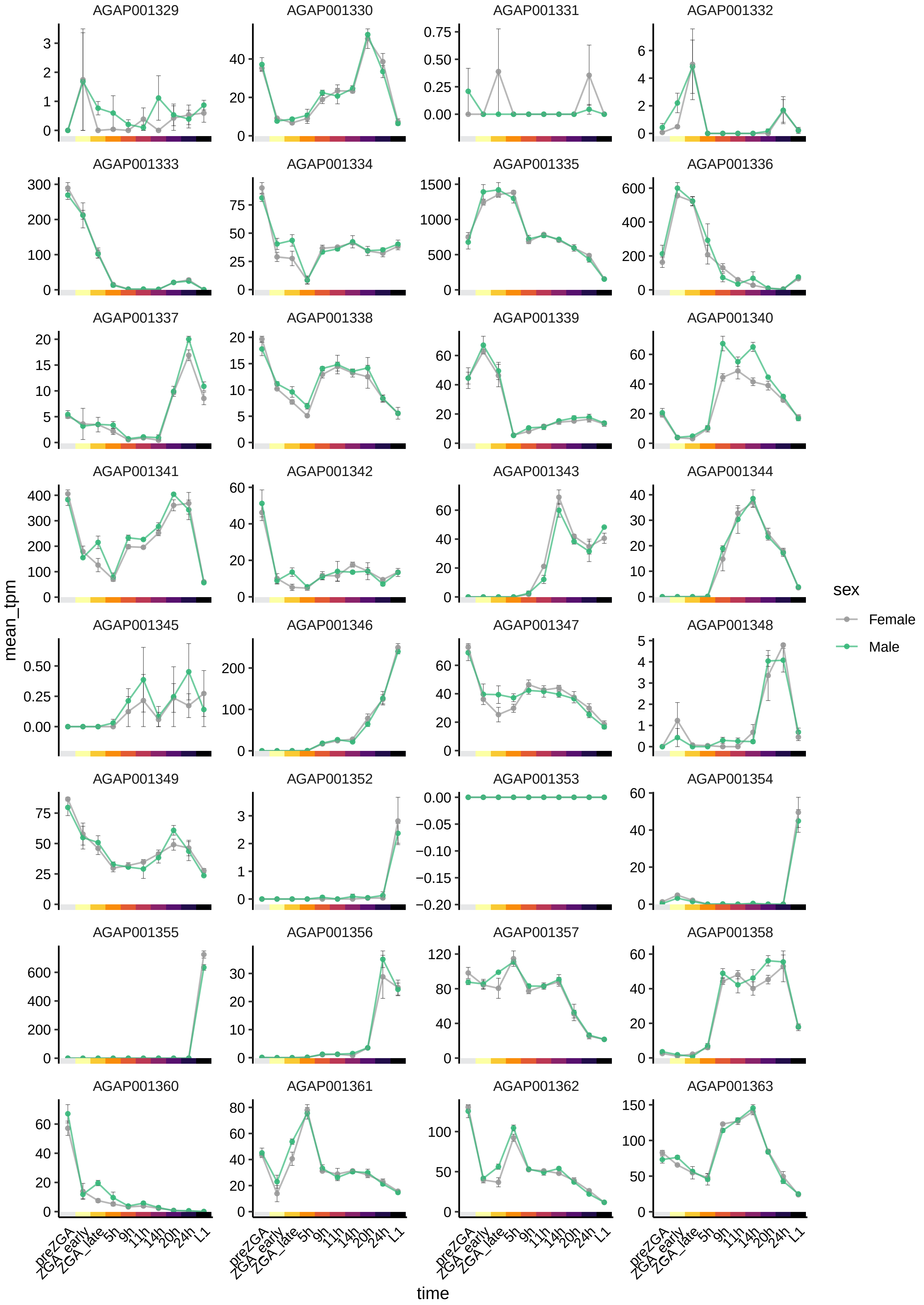
AGAP001329
AGAP001330
AGAP001331
AGAP001332
AGAP001333
AGAP001334
AGAP001335
AGAP001336
AGAP001337
AGAP001338
AGAP001339
AGAP001340
AGAP001341
AGAP001342
AGAP001343
AGAP001344
AGAP001345
AGAP001346
AGAP001347
AGAP001348
AGAP001349
AGAP001352
AGAP001353
AGAP001354
AGAP001355
AGAP001356
AGAP001357
AGAP001358
AGAP001360
AGAP001361
AGAP001362
AGAP001363
mean_tpm
time
preZGA
ZGA_early
ZGA_late
5h
9h
11h
14h
20h
24h
L1
sex
Female
Male

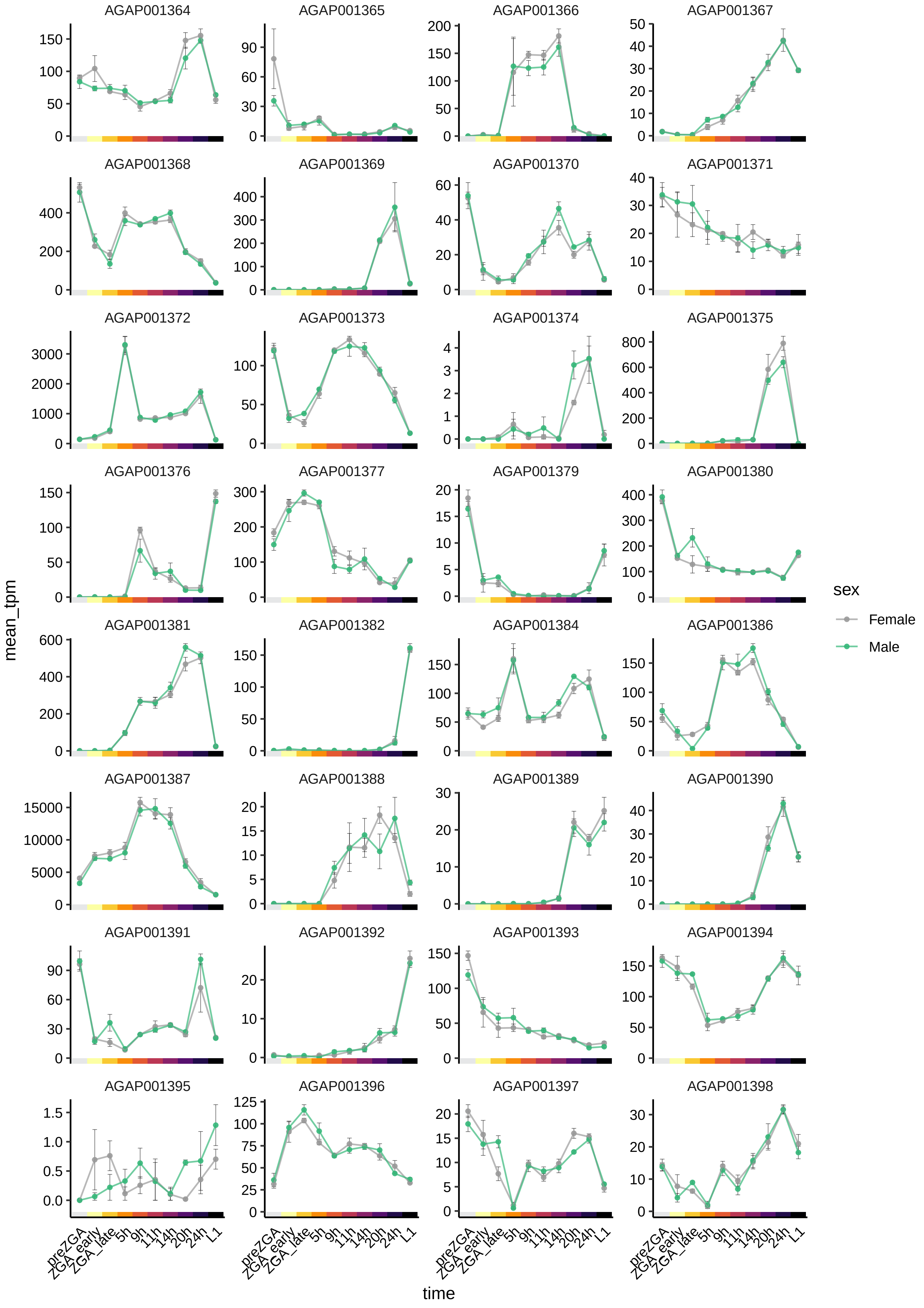

AGAP001364
AGAP001365
AGAP001366
AGAP001367
AGAP001368
AGAP001369
AGAP001370
AGAP001371
AGAP001372
AGAP001373
AGAP001374
AGAP001375
AGAP001376
AGAP001377
AGAP001379
AGAP001380
AGAP001381
AGAP001382
AGAP001384
AGAP001386
AGAP001387
AGAP001388
AGAP001389
AGAP001390
AGAP001391
AGAP001392
AGAP001393
AGAP001394
AGAP001395
AGAP001396
AGAP001397
AGAP001398
mean_tpm
time
preZGA
ZGA_early
ZGA_late
5h
9h
11h
14h
20h
24h
L1
sex
Female
Male

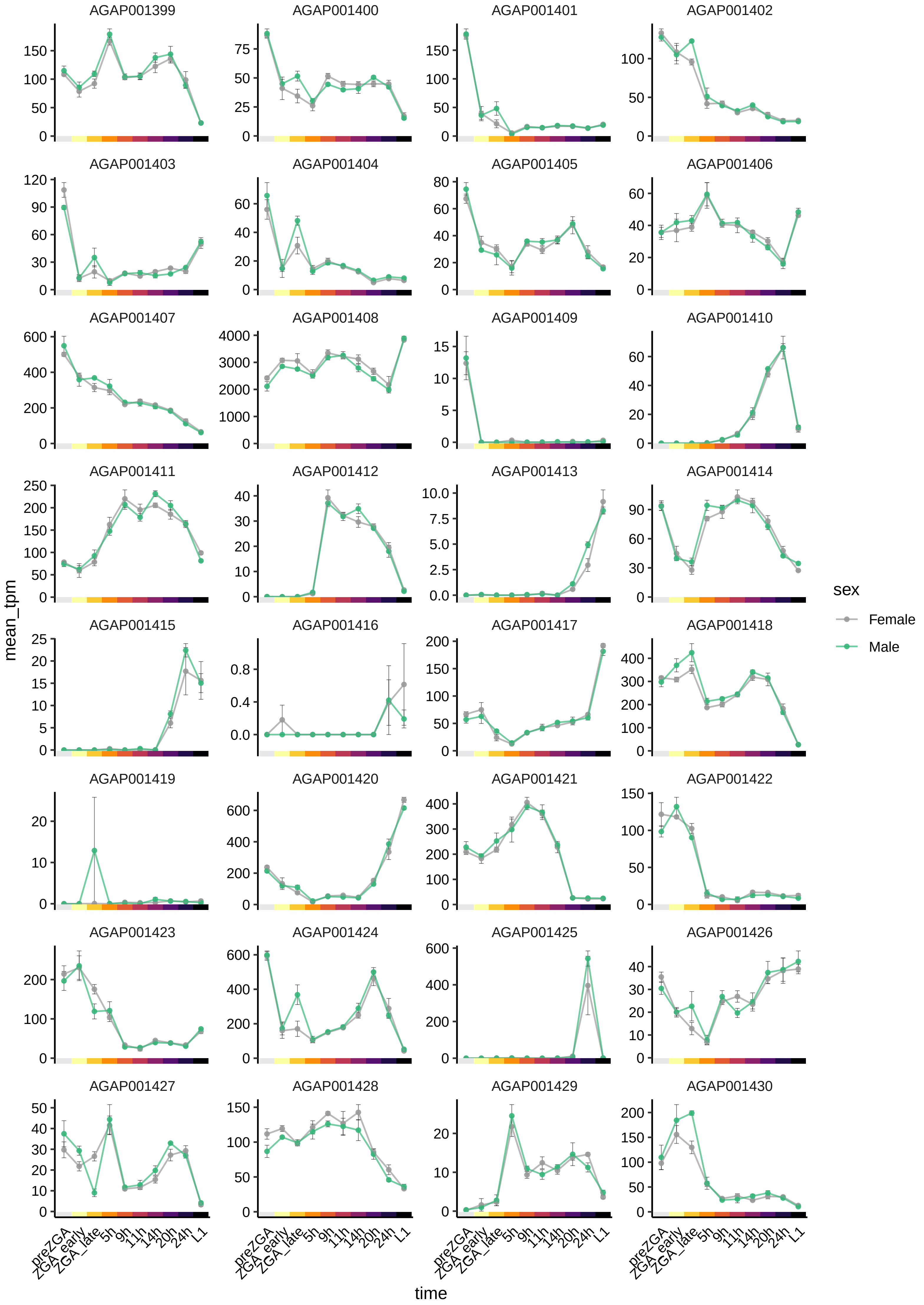
AGAP001399
AGAP001400
AGAP001401
AGAP001402
AGAP001403
AGAP001404
AGAP001405
AGAP001406
AGAP001407
AGAP001408
AGAP001409
AGAP001410
AGAP001411
AGAP001412
AGAP001413
AGAP001414
AGAP001415
AGAP001416
AGAP001417
AGAP001418
AGAP001419
AGAP001420
AGAP001421
AGAP001422
AGAP001423
AGAP001424
AGAP001425
AGAP001426
AGAP001427
AGAP001428
AGAP001429
AGAP001430
mean_tpm
time
preZGA
ZGA_early
ZGA_late
5h
9h
11h
14h
20h
24h
L1
sex
Female
Male

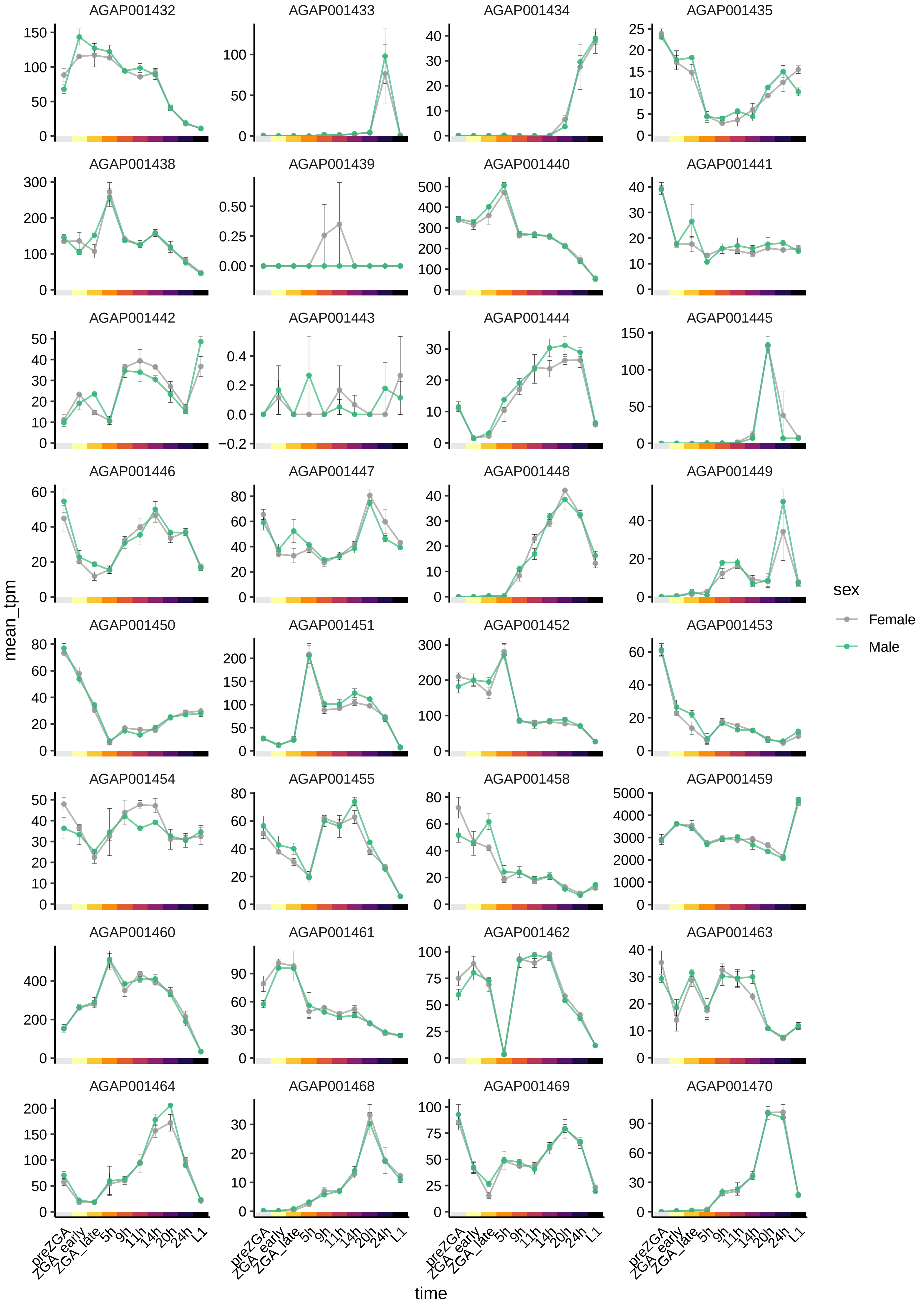

AGAP001432
AGAP001433
AGAP001434
AGAP001435
AGAP001438
AGAP001439
AGAP001440
AGAP001441
AGAP001442
AGAP001443
AGAP001444
AGAP001445
AGAP001446
AGAP001447
AGAP001448
AGAP001449
AGAP001450
AGAP001451
AGAP001452
AGAP001453
AGAP001454
AGAP001455
AGAP001458
AGAP001459
AGAP001460
AGAP001461
AGAP001462
AGAP001463
AGAP001464
AGAP001468
AGAP001469
AGAP001470
mean_tpm
time
preZGA
ZGA_early
ZGA_late
5h
9h
11h
14h
20h
24h
L1
sex
Female
Male

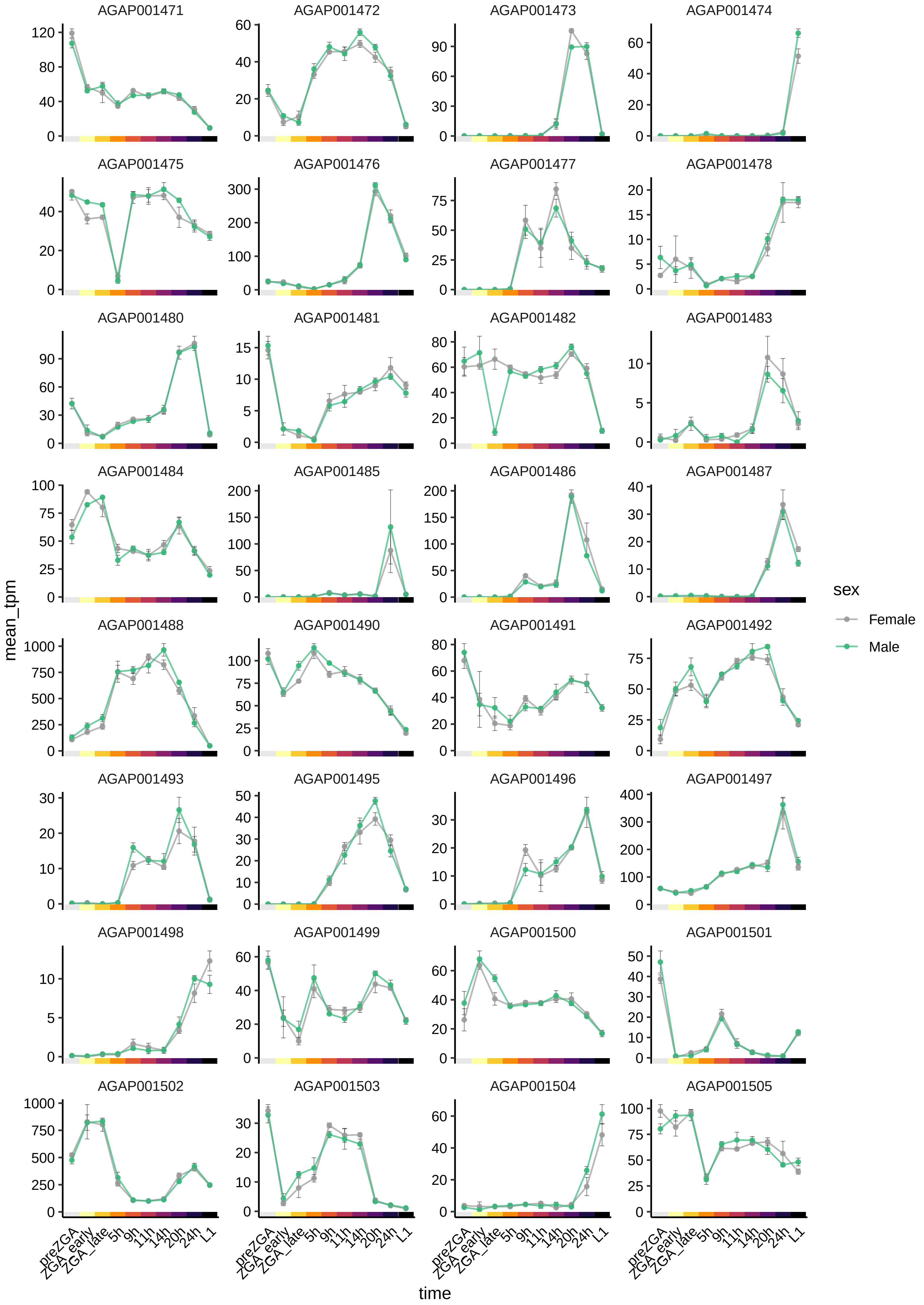

AGAP001471
AGAP001472
AGAP001473
AGAP001474
AGAP001475
AGAP001476
AGAP001477
AGAP001478
AGAP001480
AGAP001481
AGAP001482
AGAP001483
AGAP001484
AGAP001485
AGAP001486
AGAP001487
AGAP001488
AGAP001490
AGAP001491
AGAP001492
AGAP001493
AGAP001495
AGAP001496
AGAP001497
AGAP001498
AGAP001499
AGAP001500
AGAP001501
AGAP001502
AGAP001503
AGAP001504
AGAP001505
mean_tpm
time
preZGA
ZGA_early
ZGA_late
5h
9h
11h
14h
20h
24h
L1
sex
Female
Male

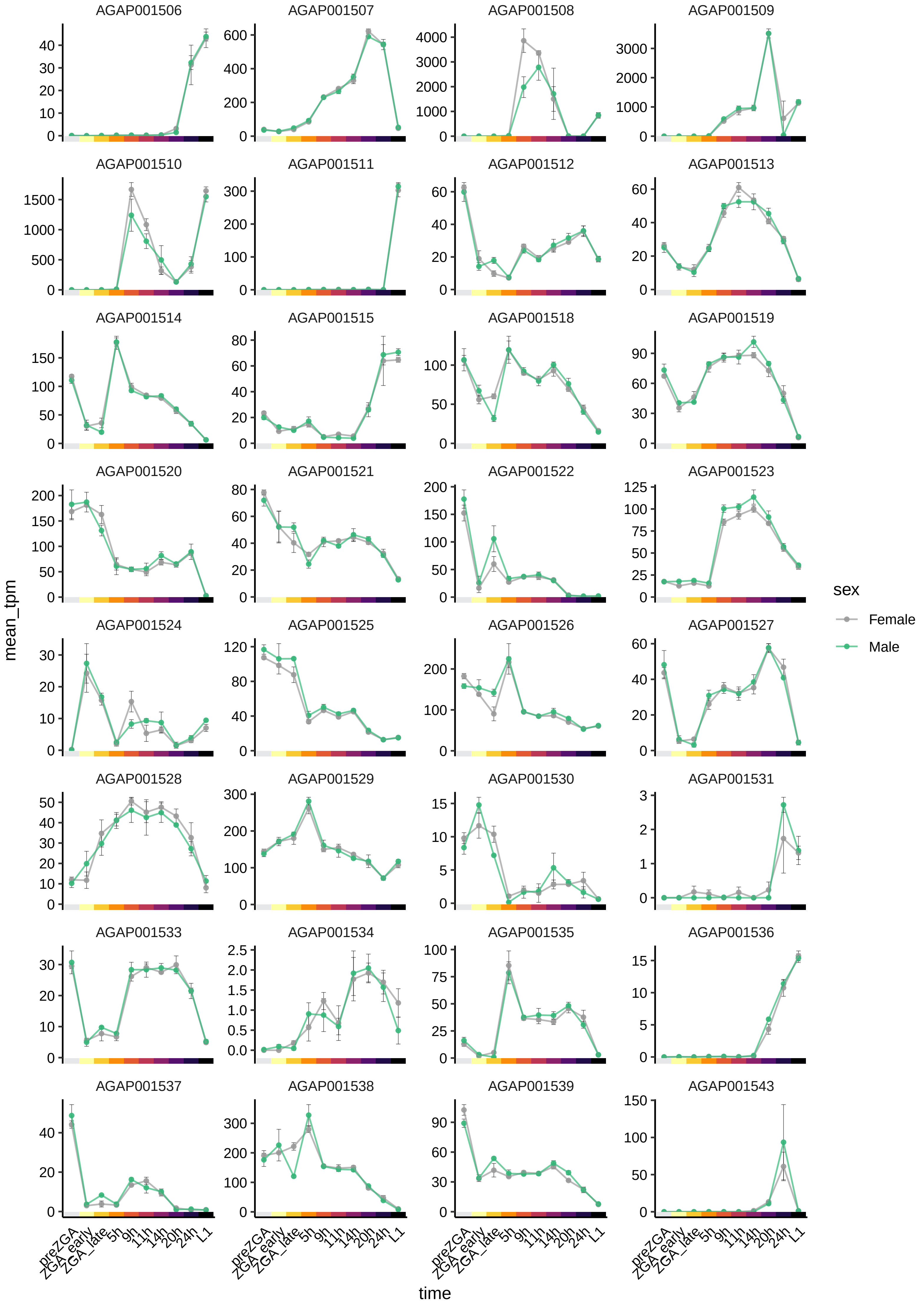
AGAP001506
AGAP001507
AGAP001508
AGAP001509
AGAP001510
AGAP001511
AGAP001512
AGAP001513
AGAP001514
AGAP001515
AGAP001518
AGAP001519
AGAP001520
AGAP001521
AGAP001522
AGAP001523
AGAP001524
AGAP001525
AGAP001526
AGAP001527
AGAP001528
AGAP001529
AGAP001530
AGAP001531
AGAP001533
AGAP001534
AGAP001535
AGAP001536
AGAP001537
AGAP001538
AGAP001539
AGAP001543
mean_tpm
time
preZGA
ZGA_early
ZGA_late
5h
9h
11h
14h
20h
24h
L1
sex
Female
Male

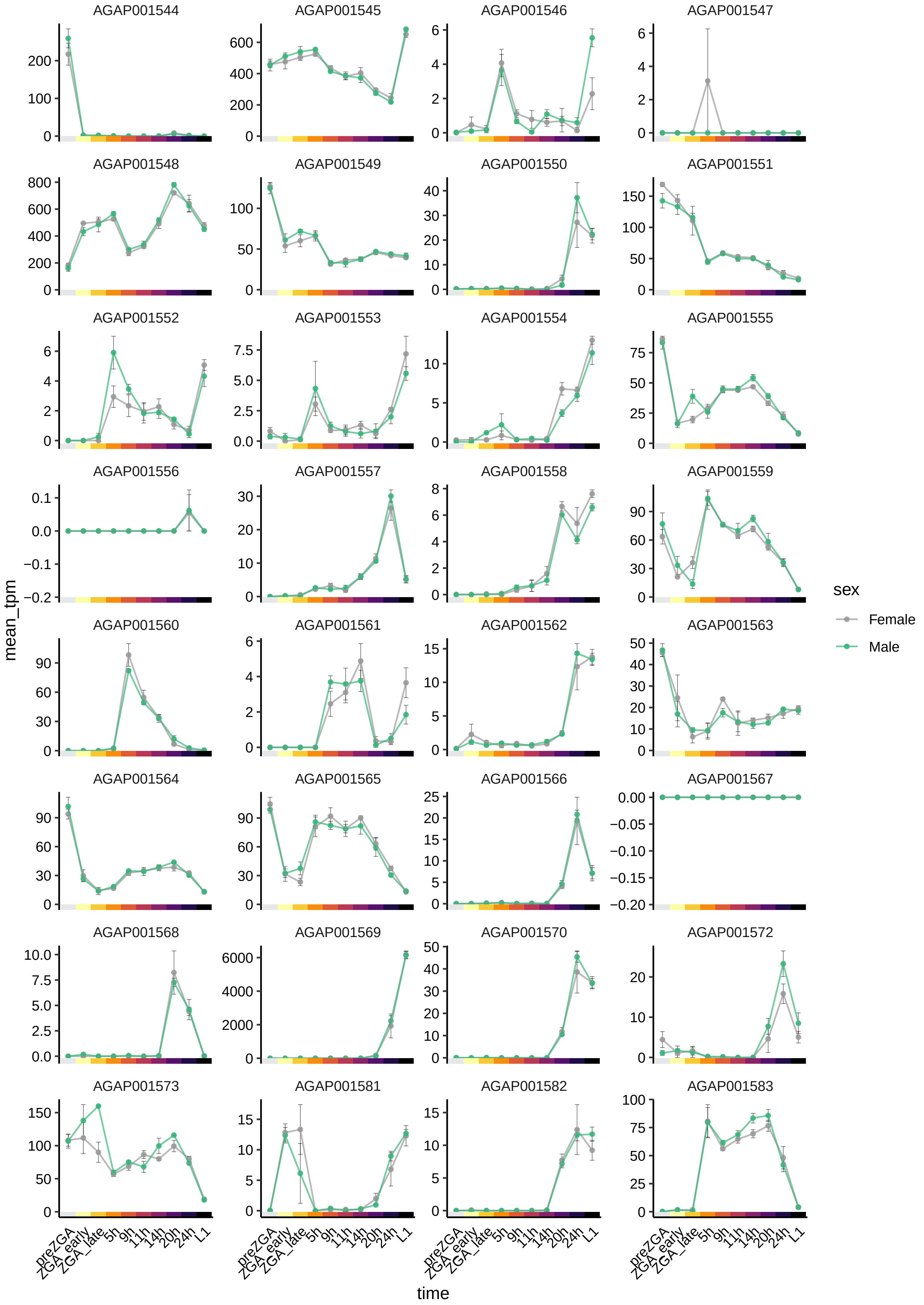
AGAP001544
AGAP001545
AGAP001546
AGAP001547
AGAP001548
AGAP001549
AGAP001550
AGAP001551
AGAP001552
AGAP001553
AGAP001554
AGAP001555
AGAP001556
AGAP001557
AGAP001558
AGAP001559
AGAP001560
AGAP001561
AGAP001562
AGAP001563
AGAP001564
AGAP001565
AGAP001566
AGAP001567
AGAP001568
AGAP001569
AGAP001570
AGAP001572
AGAP001573
AGAP001581
AGAP001582
AGAP001583
mean_tpm
time
preZGA
ZGA_early
ZGA_late
5h
9h
11h
14h
20h
24h
L1
sex
Female
Male

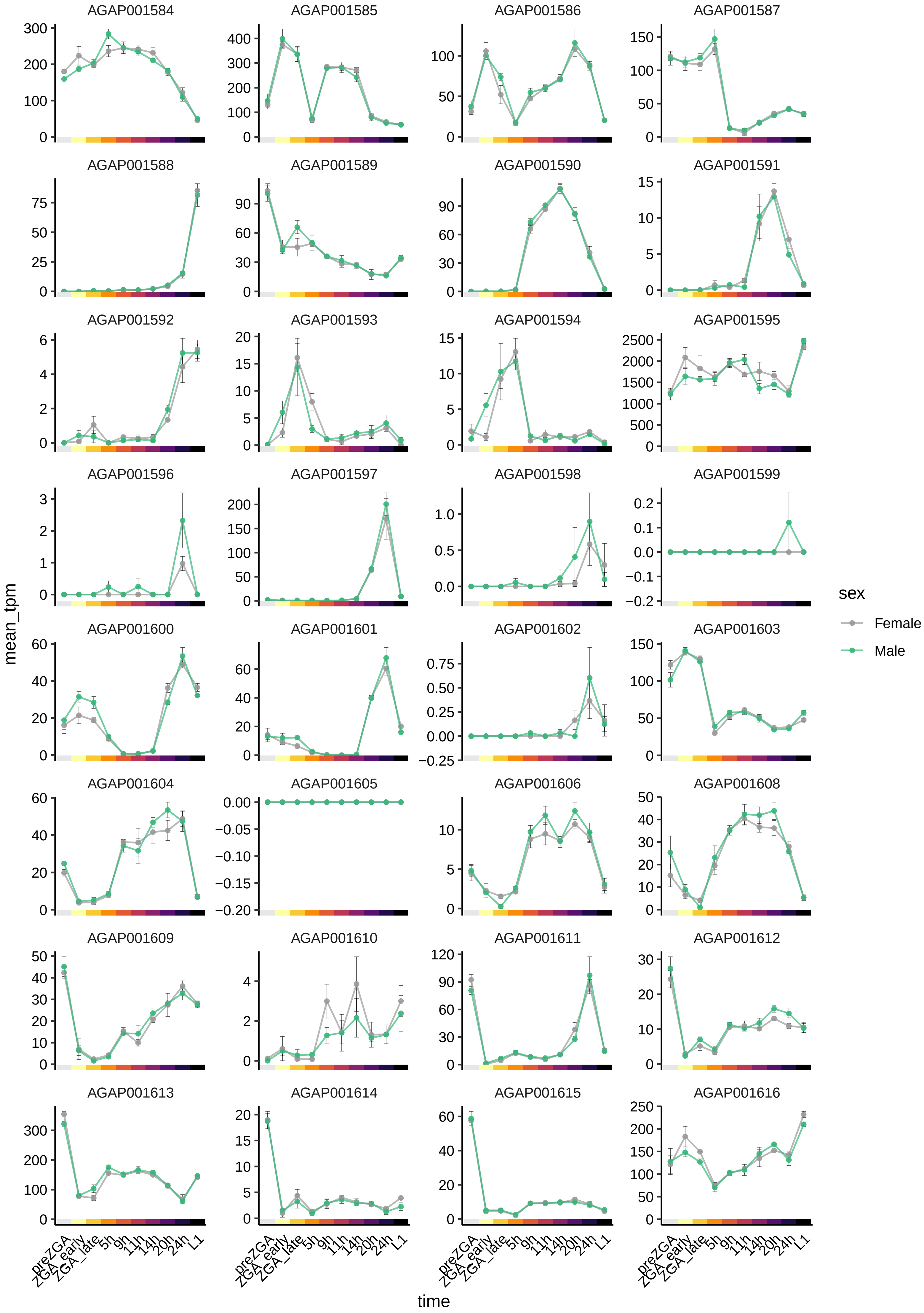

AGAP001584
AGAP001585
AGAP001586
AGAP001587
AGAP001588
AGAP001589
AGAP001590
AGAP001591
AGAP001592
AGAP001593
AGAP001594
AGAP001595
AGAP001596
AGAP001597
AGAP001598
AGAP001599
AGAP001600
AGAP001601
AGAP001602
AGAP001603
AGAP001604
AGAP001605
AGAP001606
AGAP001608
AGAP001609
AGAP001610
AGAP001611
AGAP001612
AGAP001613
AGAP001614
AGAP001615
AGAP001616
mean_tpm
time
preZGA
ZGA_early
ZGA_late
5h
9h
11h
14h
20h
24h
L1
sex
Female
Male

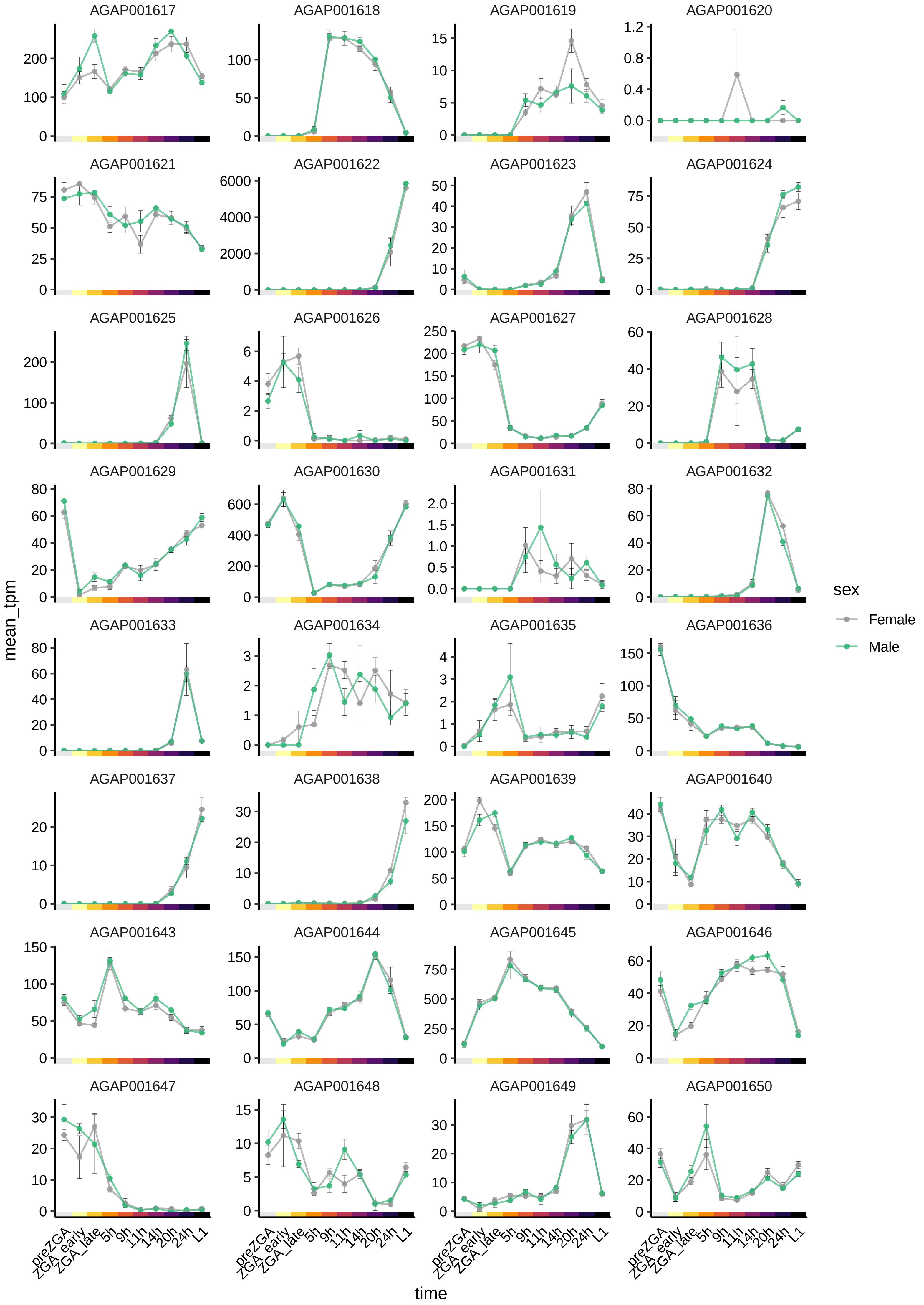

AGAP001617
AGAP001618
AGAP001619
AGAP001620
AGAP001621
AGAP001622
AGAP001623
AGAP001624
AGAP001625
AGAP001626
AGAP001627
AGAP001628
AGAP001629
AGAP001630
AGAP001631
AGAP001632
AGAP001633
AGAP001634
AGAP001635
AGAP001636
AGAP001637
AGAP001638
AGAP001639
AGAP001640
AGAP001643
AGAP001644
AGAP001645
AGAP001646
AGAP001647
AGAP001648
AGAP001649
AGAP001650
mean_tpm
time
preZGA
ZGA_early
ZGA_late
5h
9h
11h
14h
20h
24h
L1
sex
Female
Male

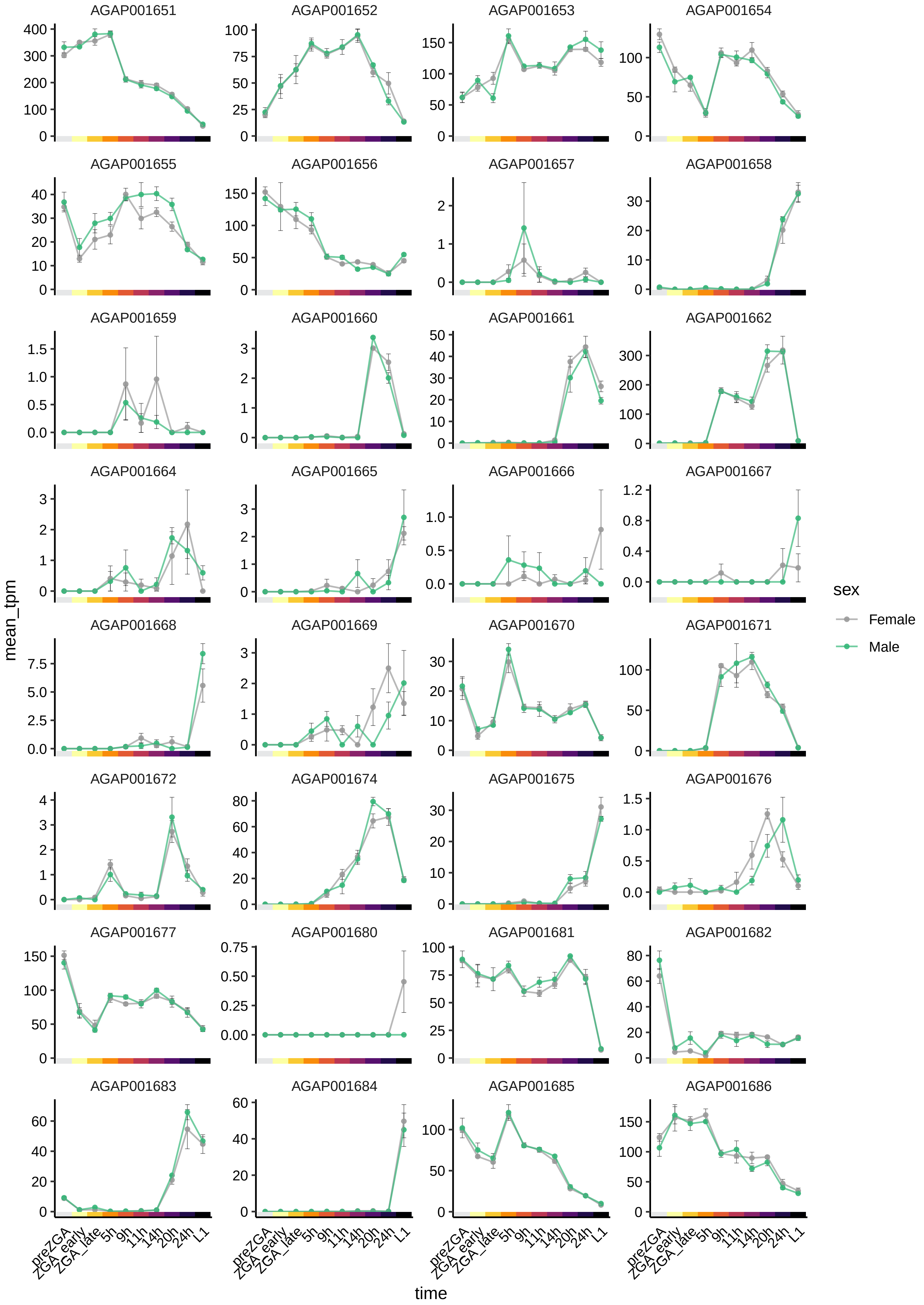

AGAP001651
AGAP001652
AGAP001653
AGAP001654
AGAP001655
AGAP001656
AGAP001657
AGAP001658
AGAP001659
AGAP001660
AGAP001661
AGAP001662
AGAP001664
AGAP001665
AGAP001666
AGAP001667
AGAP001668
AGAP001669
AGAP001670
AGAP001671
AGAP001672
AGAP001674
AGAP001675
AGAP001676
AGAP001677
AGAP001680
AGAP001681
AGAP001682
AGAP001683
AGAP001684
AGAP001685
AGAP001686
mean_tpm
time
preZGA
ZGA_early
ZGA_late
5h
9h
11h
14h
20h
24h
L1
sex
Female
Male

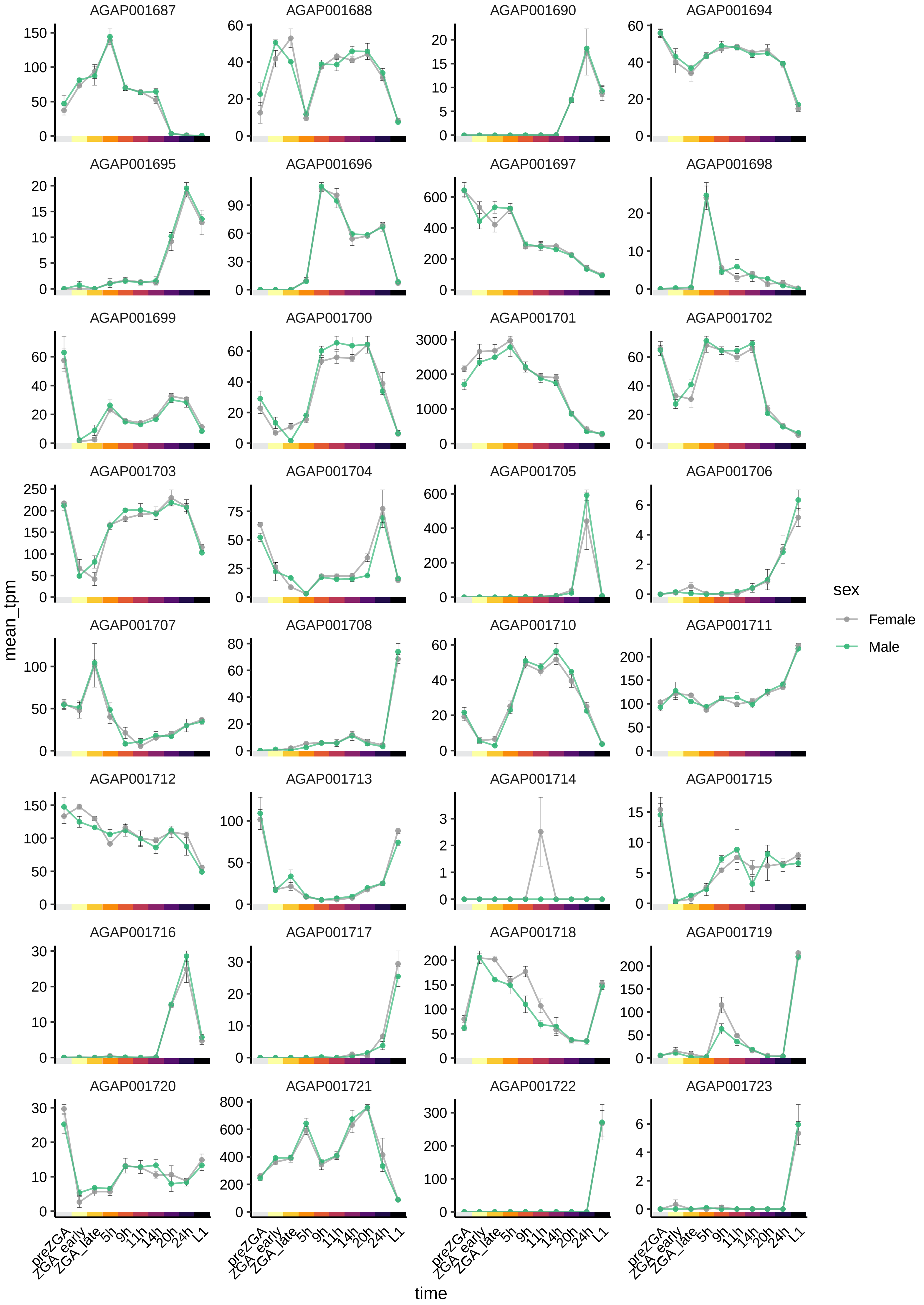

AGAP001687
AGAP001688
AGAP001690
AGAP001694
AGAP001695
AGAP001696
AGAP001697
AGAP001698
AGAP001699
AGAP001700
AGAP001701
AGAP001702
AGAP001703
AGAP001704
AGAP001705
AGAP001706
AGAP001707
AGAP001708
AGAP001710
AGAP001711
AGAP001712
AGAP001713
AGAP001714
AGAP001715
AGAP001716
AGAP001717
AGAP001718
AGAP001719
AGAP001720
AGAP001721
AGAP001722
AGAP001723
mean_tpm
time
preZGA
ZGA_early
ZGA_late
5h
9h
11h
14h
20h
24h
L1
sex
Female
Male

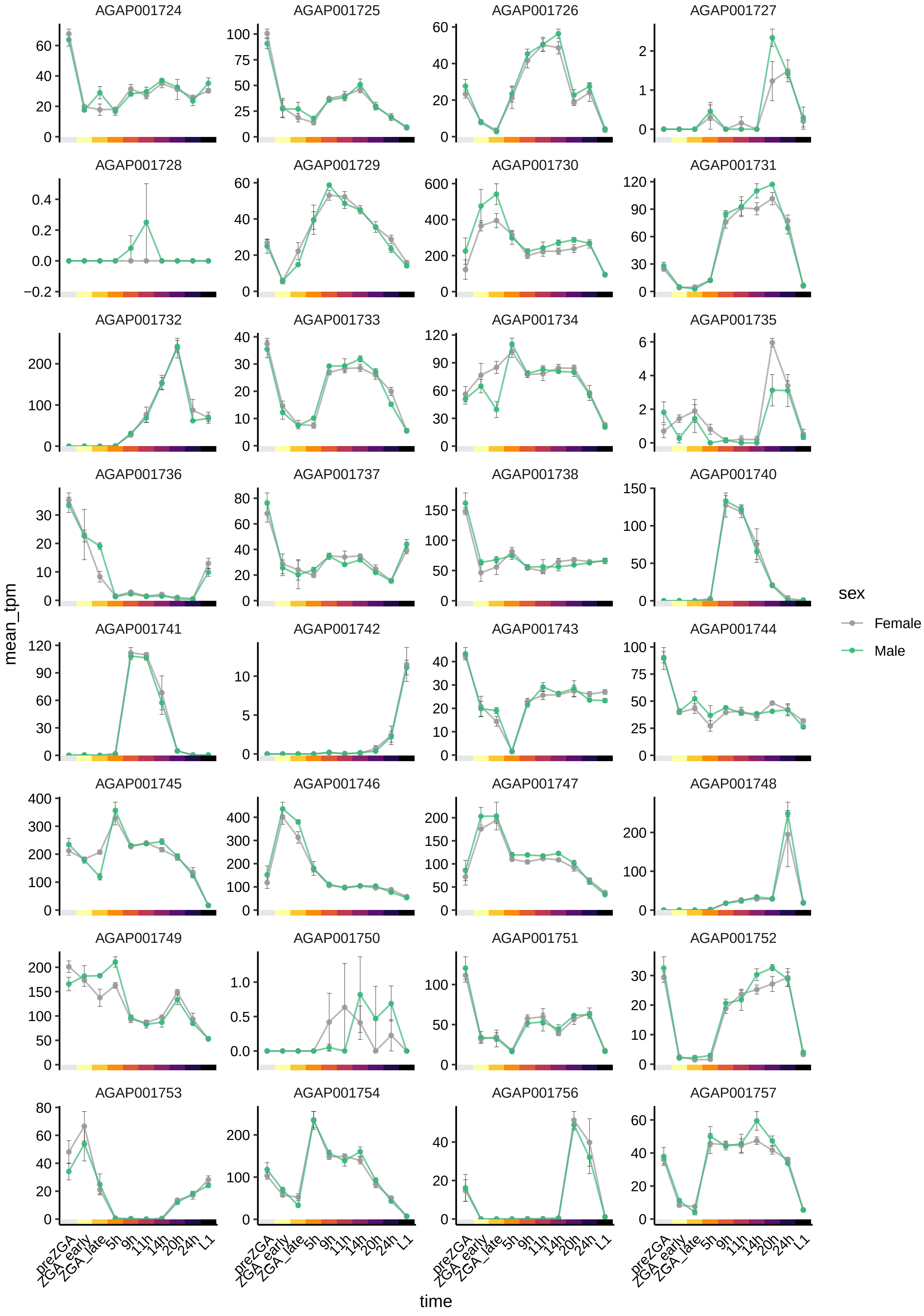
AGAP001724
AGAP001725
AGAP001726
AGAP001727
AGAP001728
AGAP001729
AGAP001730
AGAP001731
AGAP001732
AGAP001733
AGAP001734
AGAP001735
AGAP001736
AGAP001737
AGAP001738
AGAP001740
AGAP001741
AGAP001742
AGAP001743
AGAP001744
AGAP001745
AGAP001746
AGAP001747
AGAP001748
AGAP001749
AGAP001750
AGAP001751
AGAP001752
AGAP001753
AGAP001754
AGAP001756
AGAP001757
mean_tpm
time
preZGA
ZGA_early
ZGA_late
5h
9h
11h
14h
20h
24h
L1
sex
Female
Male

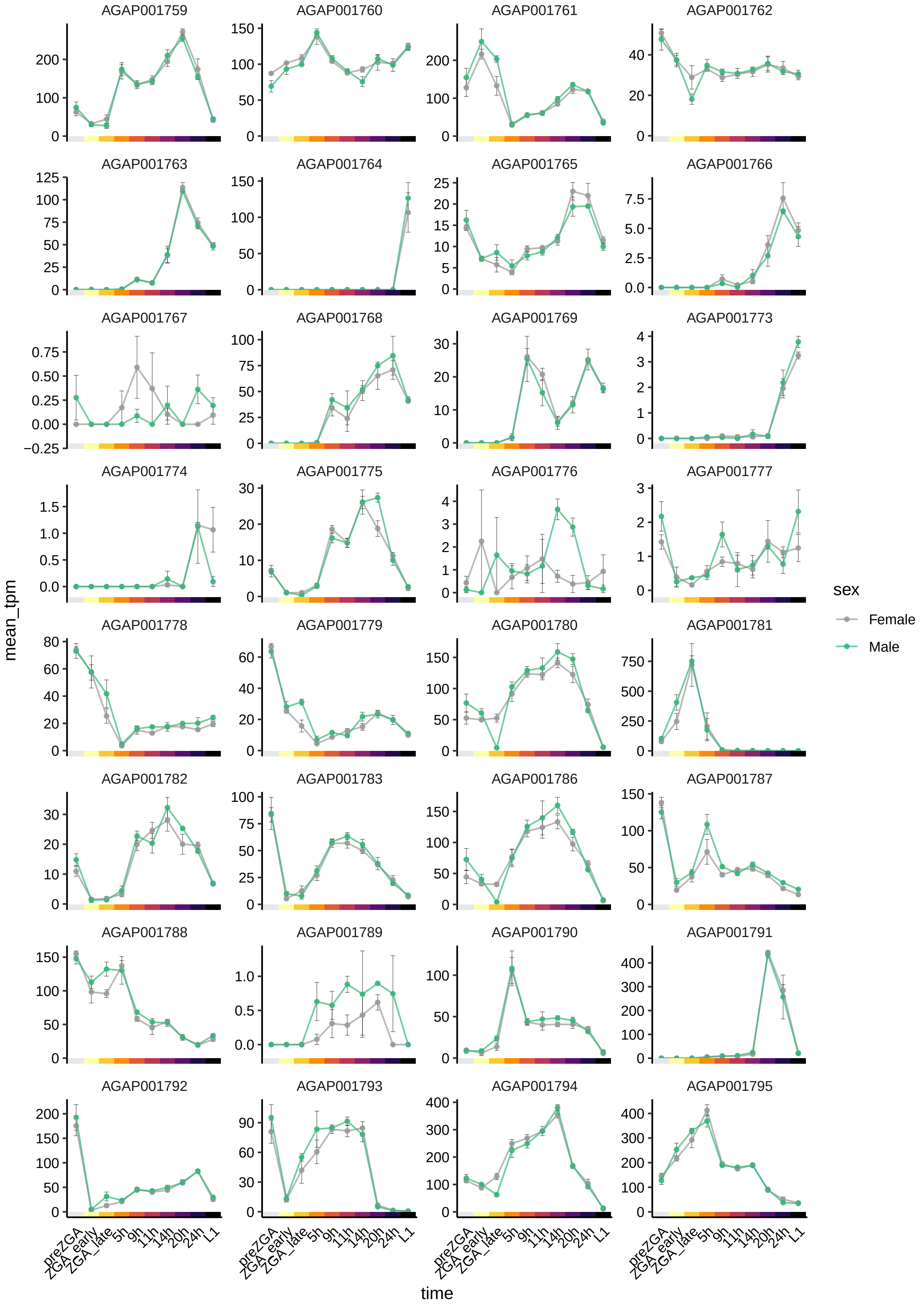
AGAP001759
AGAP001760
AGAP001761
AGAP001762
AGAP001763
AGAP001764
AGAP001765
AGAP001766
AGAP001767
AGAP001768
AGAP001769
AGAP001773
AGAP001774
AGAP001775
AGAP001776
AGAP001777
AGAP001778
AGAP001779
AGAP001780
AGAP001781
AGAP001782
AGAP001783
AGAP001786
AGAP001787
AGAP001788
AGAP001789
AGAP001790
AGAP001791
AGAP001792
AGAP001793
AGAP001794
AGAP001795
mean_tpm
time
preZGA
ZGA_early
ZGA_late
5h
9h
11h
14h
20h
24h
L1
sex
Female
Male

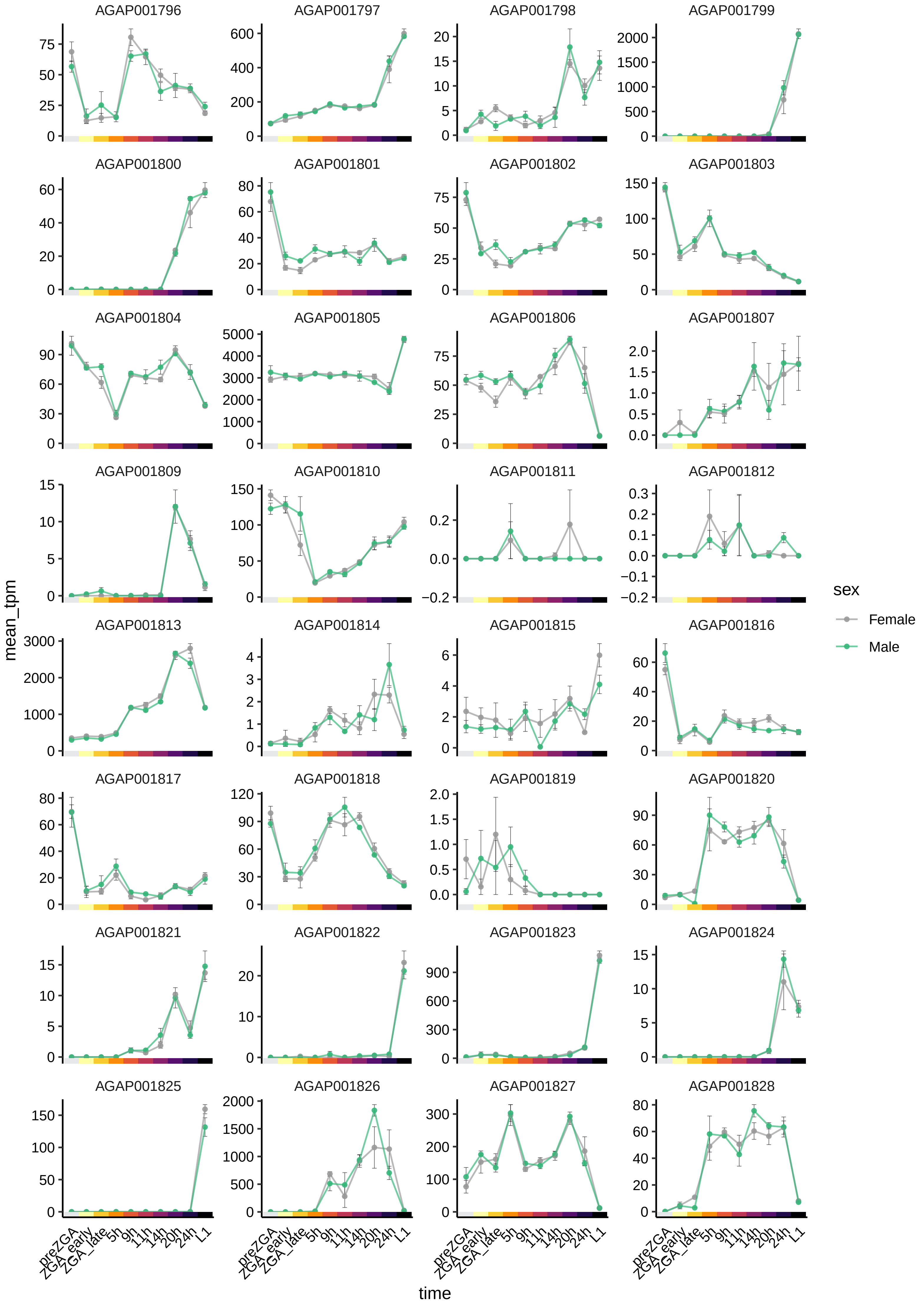

AGAP001796
AGAP001797
AGAP001798
AGAP001799
AGAP001800
AGAP001801
AGAP001802
AGAP001803
AGAP001804
AGAP001805
AGAP001806
AGAP001807
AGAP001809
AGAP001810
AGAP001811
AGAP001812
AGAP001813
AGAP001814
AGAP001815
AGAP001816
AGAP001817
AGAP001818
AGAP001819
AGAP001820
AGAP001821
AGAP001822
AGAP001823
AGAP001824
AGAP001825
AGAP001826
AGAP001827
AGAP001828
mean_tpm
time
preZGA
ZGA_early
ZGA_late
5h
9h
11h
14h
20h
24h
L1
sex
Female
Male

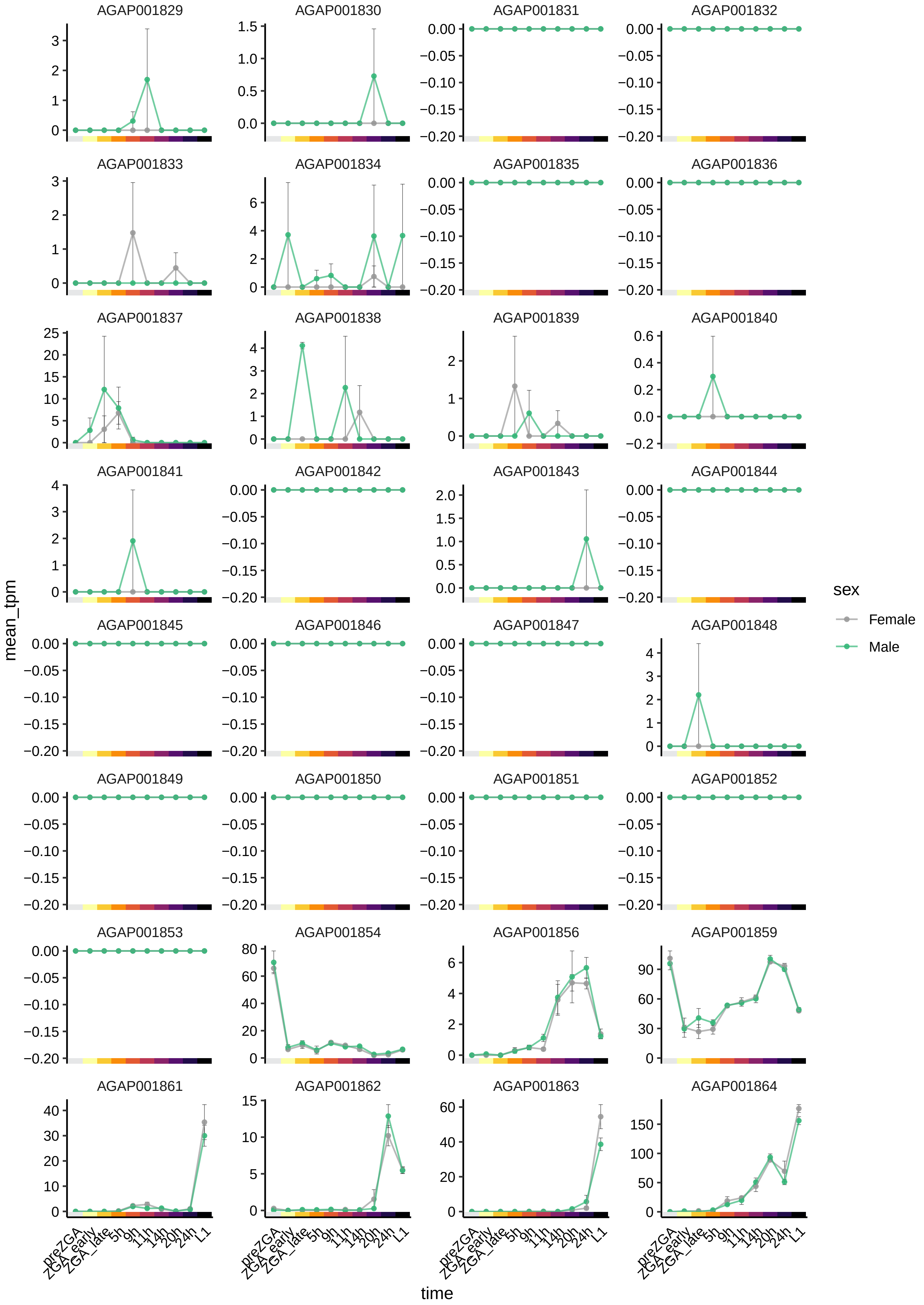
AGAP001829
AGAP001830
AGAP001831
AGAP001832
AGAP001833
AGAP001834
AGAP001835
AGAP001836
AGAP001837
AGAP001838
AGAP001839
AGAP001840
AGAP001841
AGAP001842
AGAP001843
AGAP001844
AGAP001845
AGAP001846
AGAP001847
AGAP001848
AGAP001849
AGAP001850
AGAP001851
AGAP001852
AGAP001853
AGAP001854
AGAP001856
AGAP001859
AGAP001861
AGAP001862
AGAP001863
AGAP001864
mean_tpm
time
preZGA
ZGA_early
ZGA_late
5h
9h
11h
14h
20h
24h
L1
sex
Female
Male

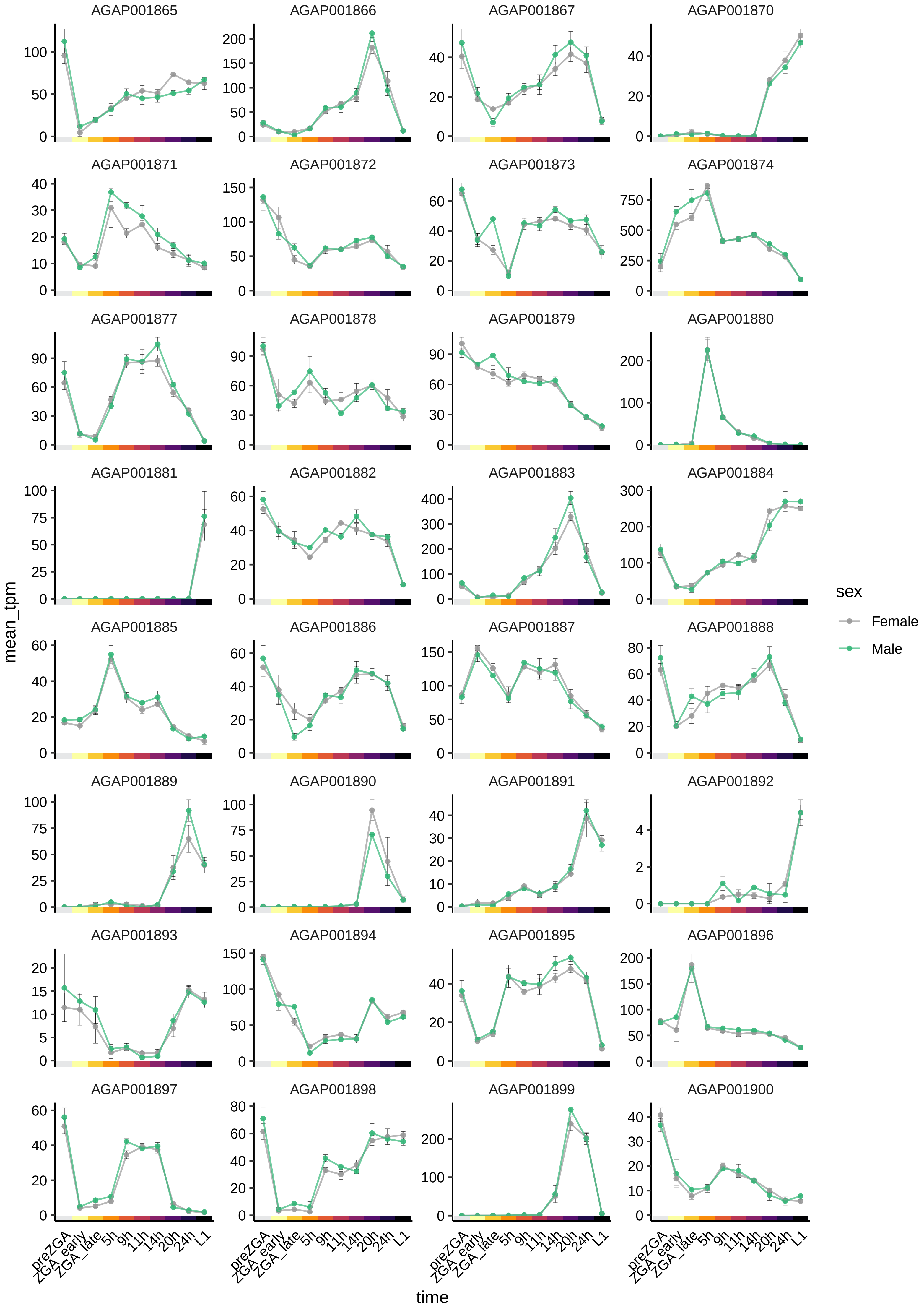

AGAP001865
AGAP001866
AGAP001867
AGAP001870
AGAP001871
AGAP001872
AGAP001873
AGAP001874
AGAP001877
AGAP001878
AGAP001879
AGAP001880
AGAP001881
AGAP001882
AGAP001883
AGAP001884
AGAP001885
AGAP001886
AGAP001887
AGAP001888
AGAP001889
AGAP001890
AGAP001891
AGAP001892
AGAP001893
AGAP001894
AGAP001895
AGAP001896
AGAP001897
AGAP001898
AGAP001899
AGAP001900
mean_tpm
time
preZGA
ZGA_early
ZGA_late
5h
9h
11h
14h
20h
24h
L1
sex
Female
Male

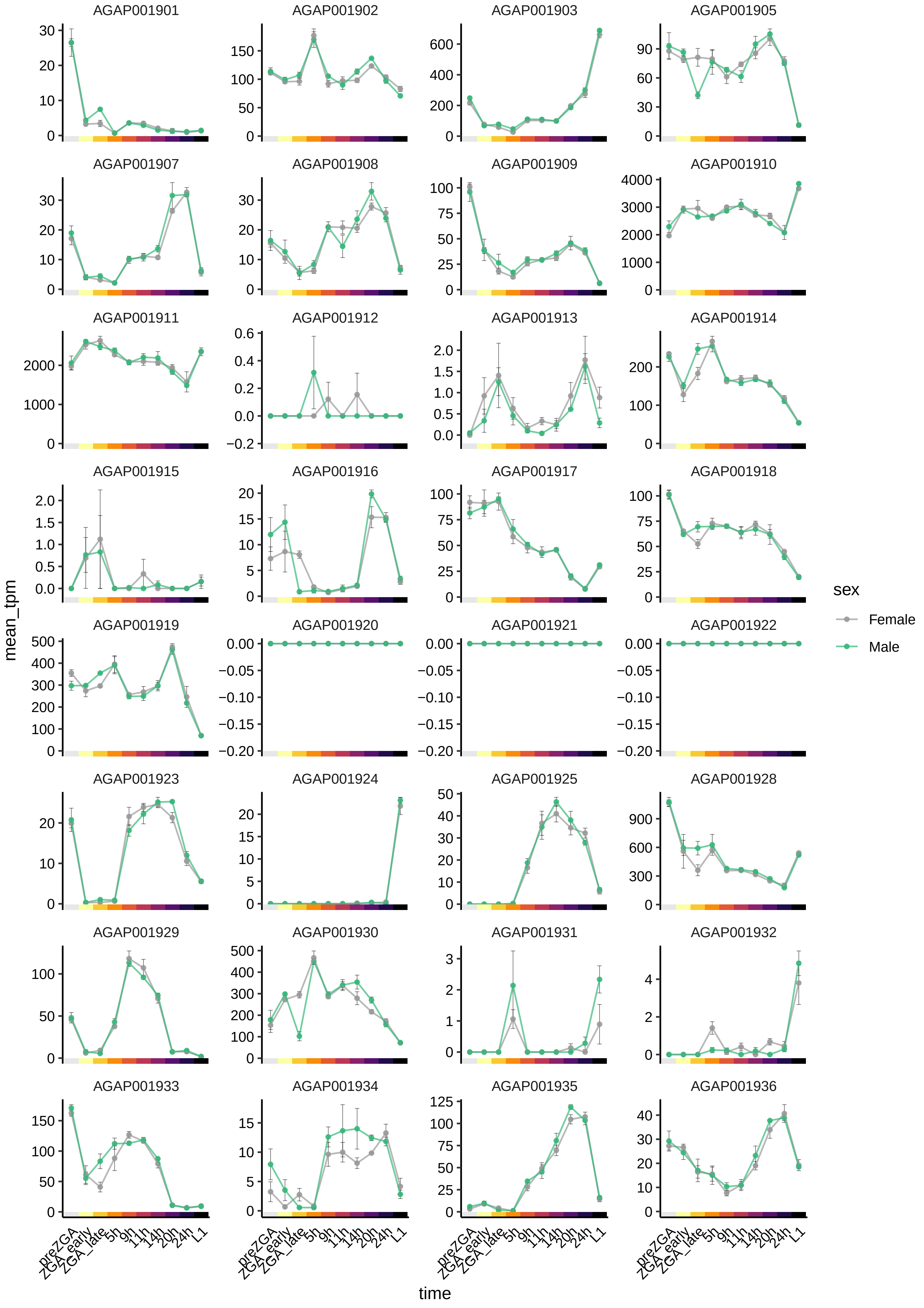
AGAP001901
AGAP001902
AGAP001903
AGAP001905
AGAP001907
AGAP001908
AGAP001909
AGAP001910
AGAP001911
AGAP001912
AGAP001913
AGAP001914
AGAP001915
AGAP001916
AGAP001917
AGAP001918
AGAP001919
AGAP001920
AGAP001921
AGAP001922
AGAP001923
AGAP001924
AGAP001925
AGAP001928
AGAP001929
AGAP001930
AGAP001931
AGAP001932
AGAP001933
AGAP001934
AGAP001935
AGAP001936
mean_tpm
time
preZGA
ZGA_early
ZGA_late
5h
9h
11h
14h
20h
24h
L1
sex
Female
Male

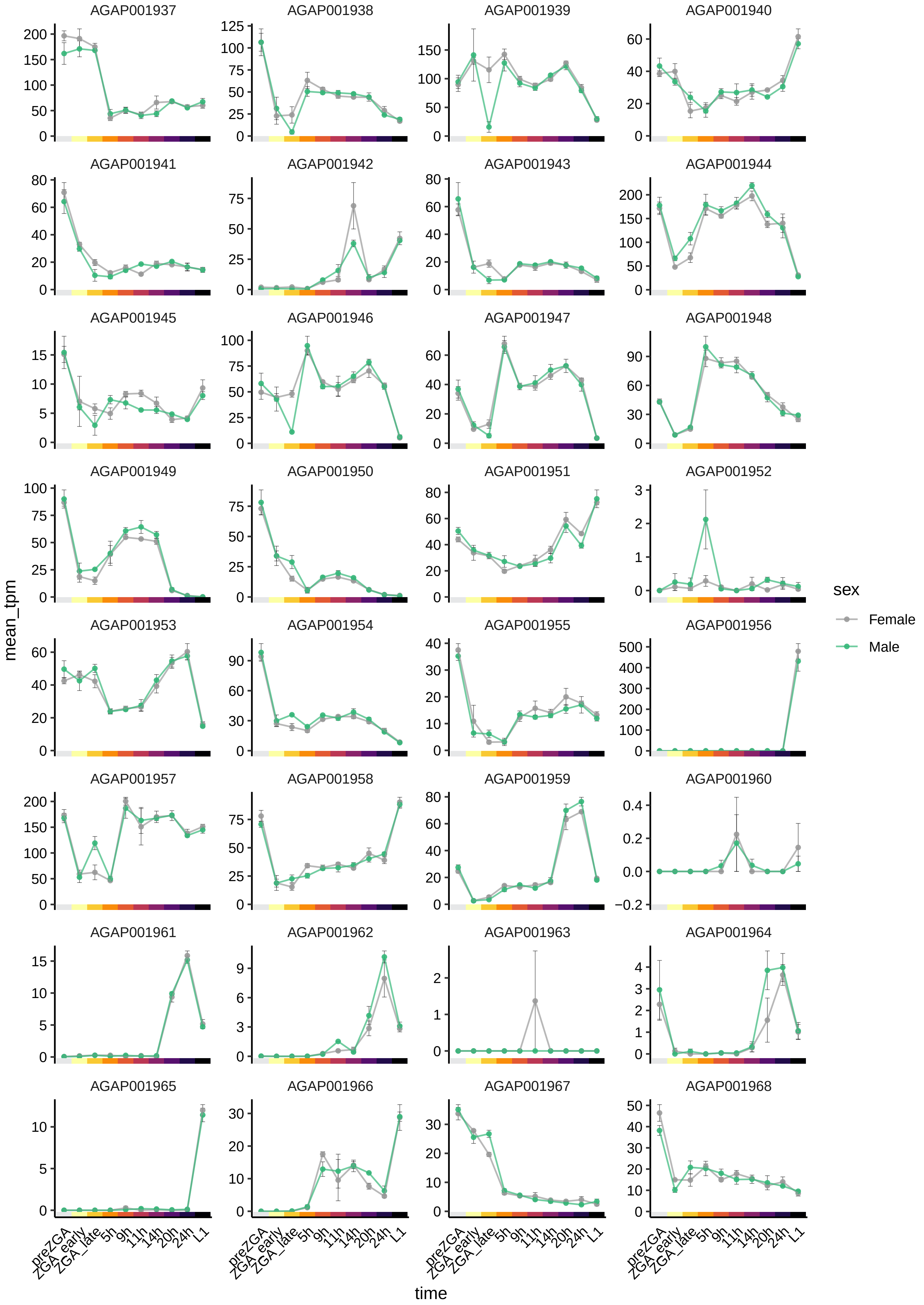

AGAP001937
AGAP001938
AGAP001939
AGAP001940
AGAP001941
AGAP001942
AGAP001943
AGAP001944
AGAP001945
AGAP001946
AGAP001947
AGAP001948
AGAP001949
AGAP001950
AGAP001951
AGAP001952
AGAP001953
AGAP001954
AGAP001955
AGAP001956
AGAP001957
AGAP001958
AGAP001959
AGAP001960
AGAP001961
AGAP001962
AGAP001963
AGAP001964
AGAP001965
AGAP001966
AGAP001967
AGAP001968
mean_tpm
time
preZGA
ZGA_early
ZGA_late
5h
9h
11h
14h
20h
24h
L1
sex
Female
Male

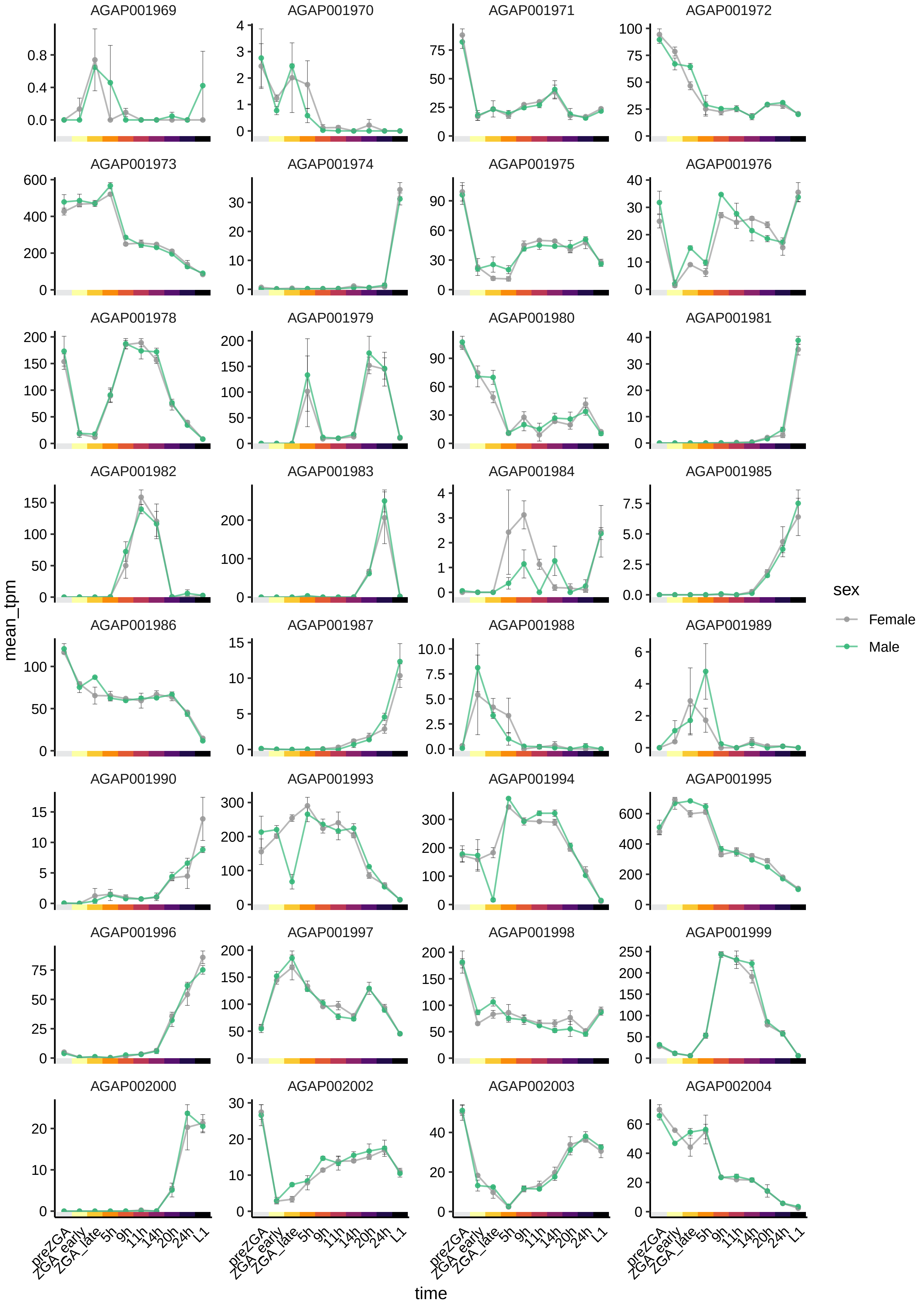
AGAP001969
AGAP001970
AGAP001971
AGAP001972
AGAP001973
AGAP001974
AGAP001975
AGAP001976
AGAP001978
AGAP001979
AGAP001980
AGAP001981
AGAP001982
AGAP001983
AGAP001984
AGAP001985
AGAP001986
AGAP001987
AGAP001988
AGAP001989
AGAP001990
AGAP001993
AGAP001994
AGAP001995
AGAP001996
AGAP001997
AGAP001998
AGAP001999
AGAP002000
AGAP002002
AGAP002003
AGAP002004
mean_tpm
time
preZGA
ZGA_early
ZGA_late
5h
9h
11h
14h
20h
24h
L1
sex
Female
Male

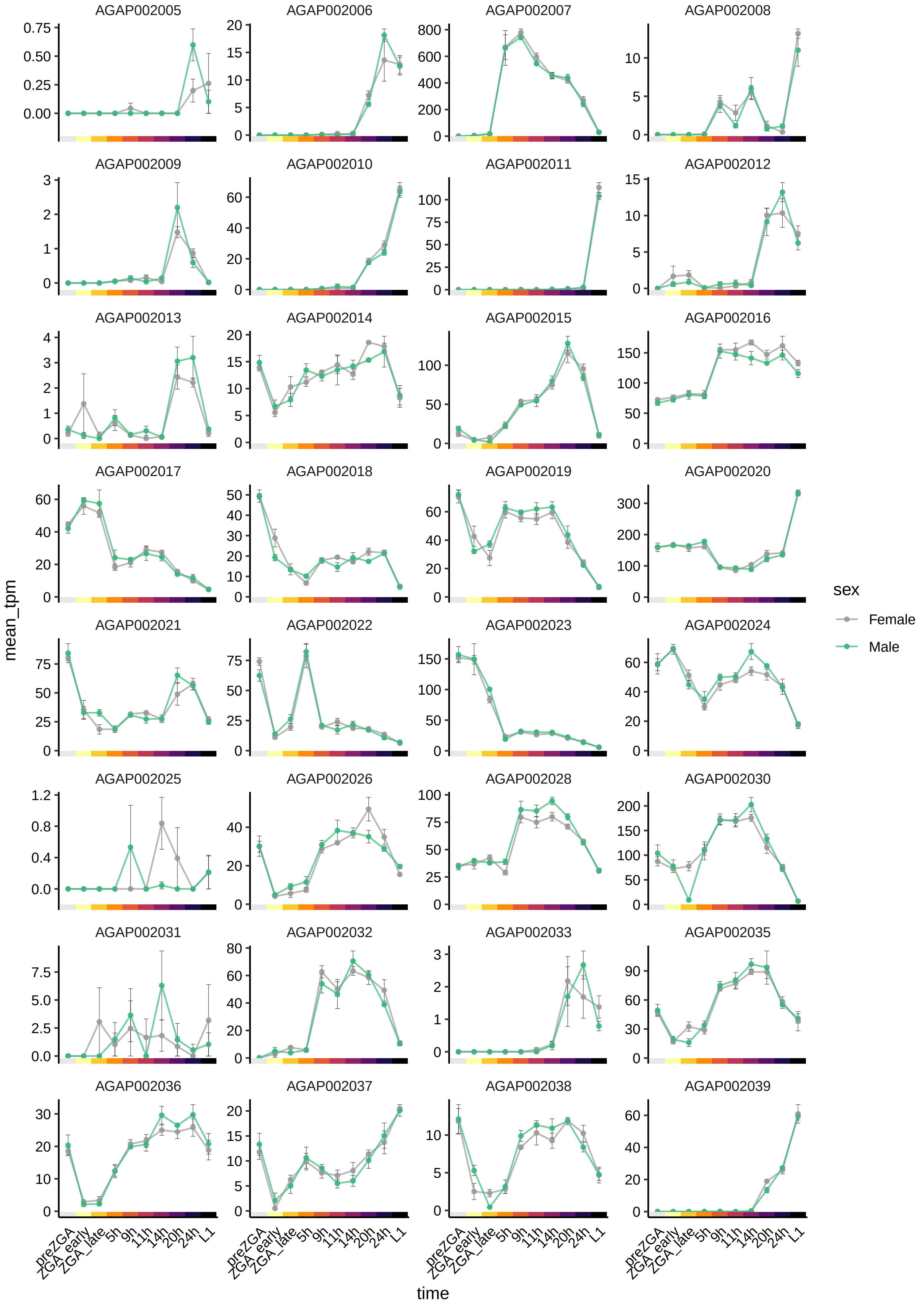

AGAP002005
AGAP002006
AGAP002007
AGAP002008
AGAP002009
AGAP002010
AGAP002011
AGAP002012
AGAP002013
AGAP002014
AGAP002015
AGAP002016
AGAP002017
AGAP002018
AGAP002019
AGAP002020
AGAP002021
AGAP002022
AGAP002023
AGAP002024
AGAP002025
AGAP002026
AGAP002028
AGAP002030
AGAP002031
AGAP002032
AGAP002033
AGAP002035
AGAP002036
AGAP002037
AGAP002038
AGAP002039
mean_tpm
time
preZGA
ZGA_early
ZGA_late
5h
9h
11h
14h
20h
24h
L1
sex
Female
Male

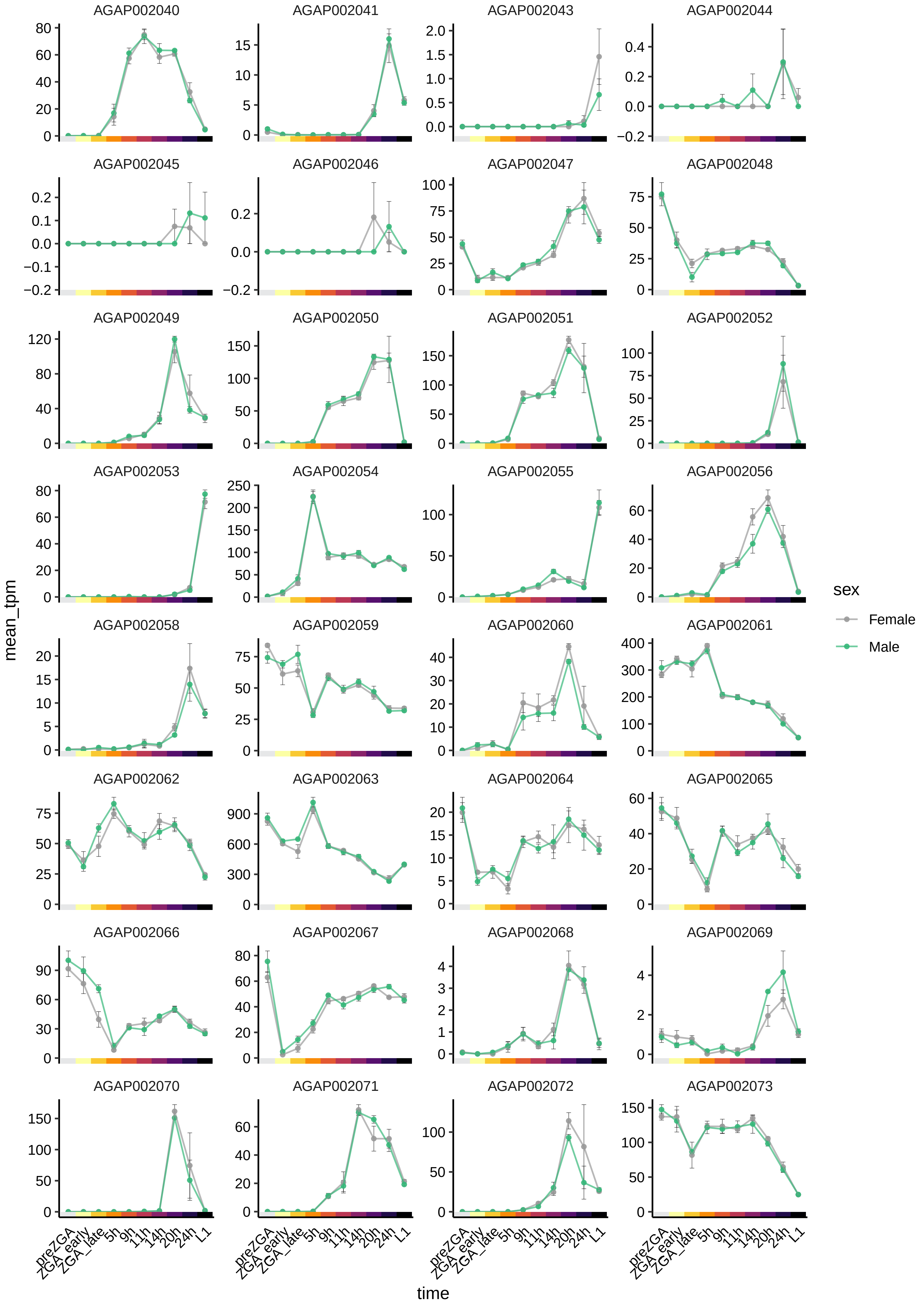

AGAP002040
AGAP002041
AGAP002043
AGAP002044
AGAP002045
AGAP002046
AGAP002047
AGAP002048
AGAP002049
AGAP002050
AGAP002051
AGAP002052
AGAP002053
AGAP002054
AGAP002055
AGAP002056
AGAP002058
AGAP002059
AGAP002060
AGAP002061
AGAP002062
AGAP002063
AGAP002064
AGAP002065
AGAP002066
AGAP002067
AGAP002068
AGAP002069
AGAP002070
AGAP002071
AGAP002072
AGAP002073
mean_tpm
time
preZGA
ZGA_early
ZGA_late
5h
9h
11h
14h
20h
24h
L1
sex
Female
Male

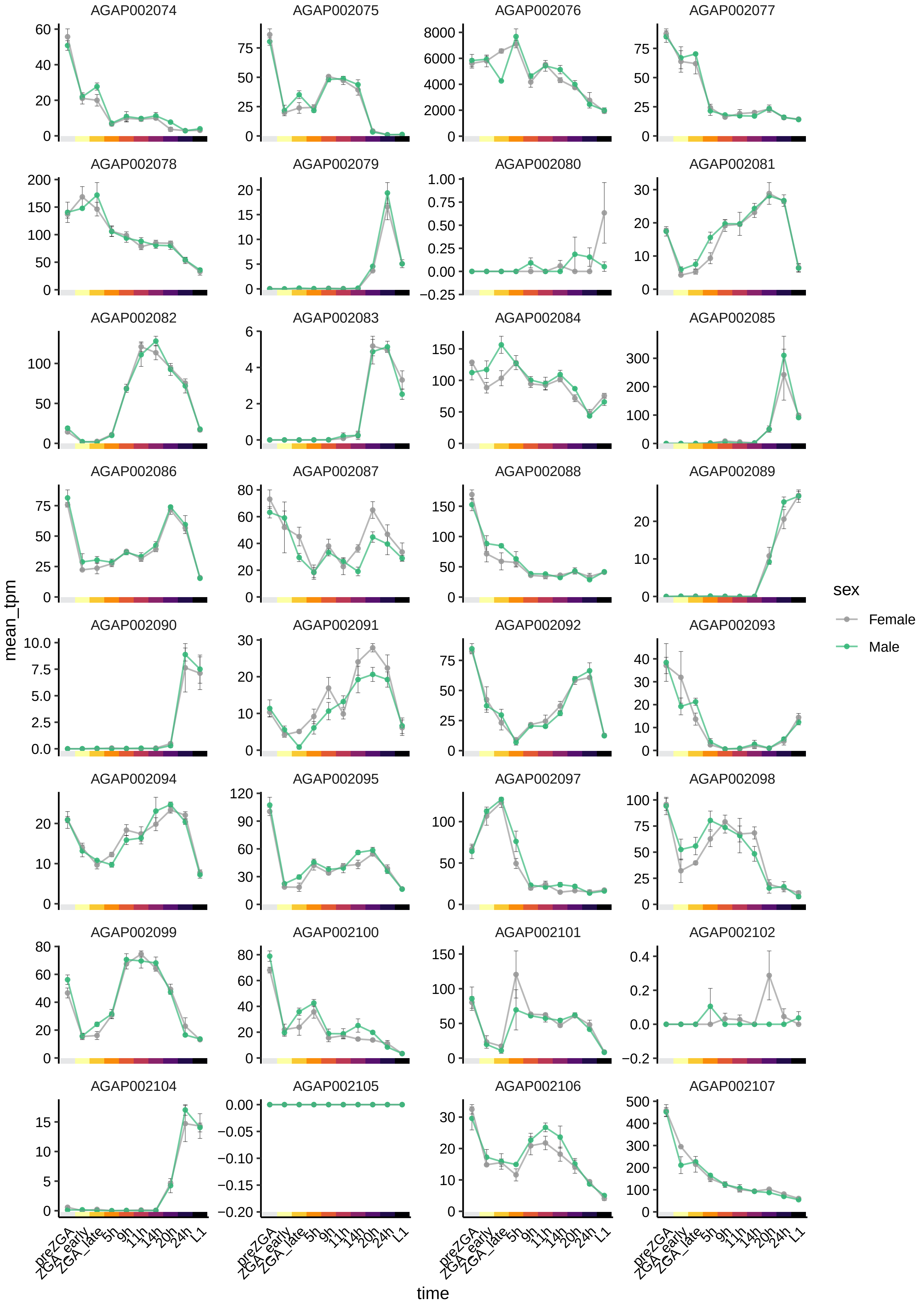
AGAP002074
AGAP002075
AGAP002076
AGAP002077
AGAP002078
AGAP002079
AGAP002080
AGAP002081
AGAP002082
AGAP002083
AGAP002084
AGAP002085
AGAP002086
AGAP002087
AGAP002088
AGAP002089
AGAP002090
AGAP002091
AGAP002092
AGAP002093
AGAP002094
AGAP002095
AGAP002097
AGAP002098
AGAP002099
AGAP002100
AGAP002101
AGAP002102
AGAP002104
AGAP002105
AGAP002106
AGAP002107
mean_tpm
time
preZGA
ZGA_early
ZGA_late
5h
9h
11h
14h
20h
24h
L1
sex
Female
Male

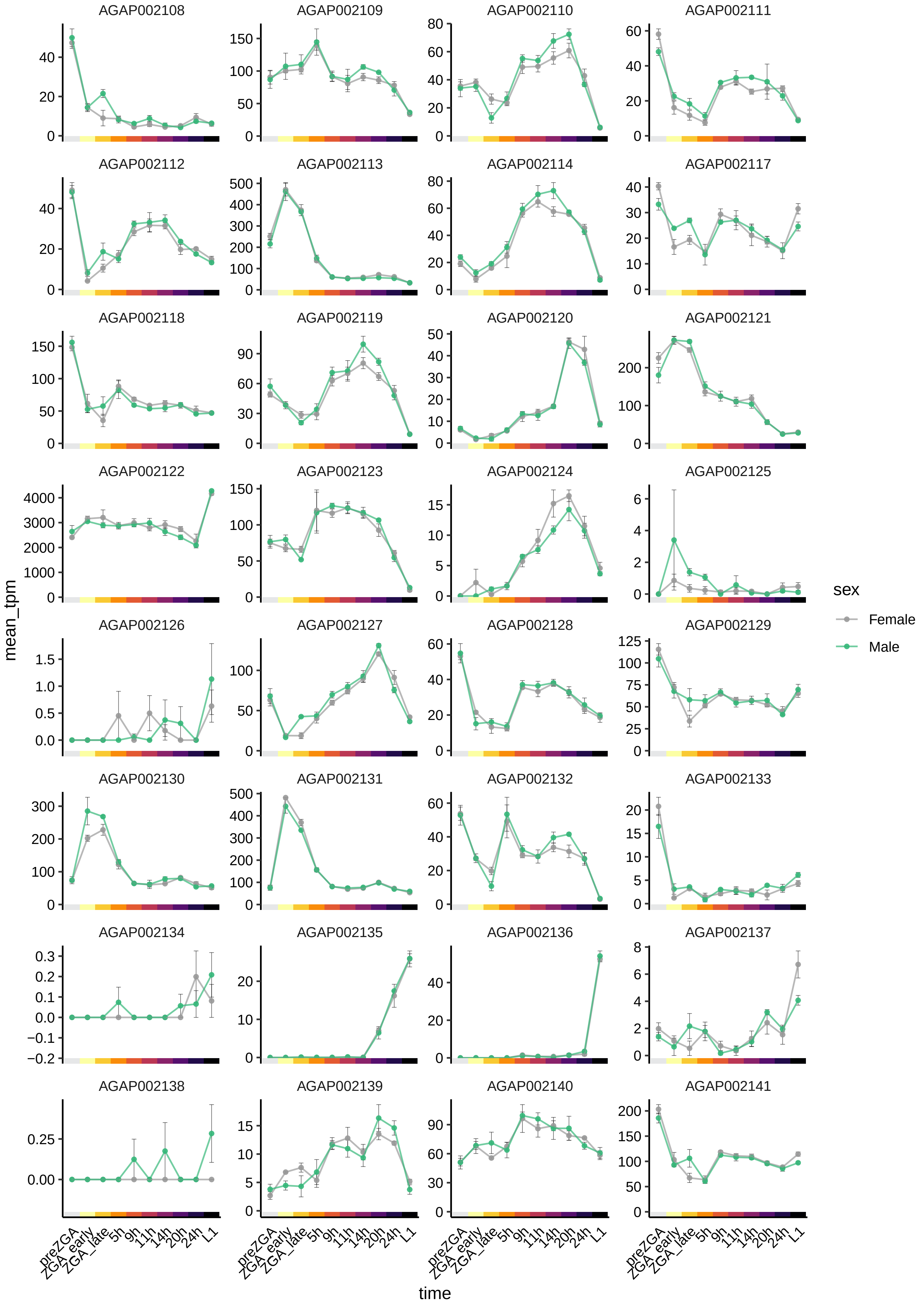

AGAP002108
AGAP002109
AGAP002110
AGAP002111
AGAP002112
AGAP002113
AGAP002114
AGAP002117
AGAP002118
AGAP002119
AGAP002120
AGAP002121
AGAP002122
AGAP002123
AGAP002124
AGAP002125
AGAP002126
AGAP002127
AGAP002128
AGAP002129
AGAP002130
AGAP002131
AGAP002132
AGAP002133
AGAP002134
AGAP002135
AGAP002136
AGAP002137
AGAP002138
AGAP002139
AGAP002140
AGAP002141
mean_tpm
time
preZGA
ZGA_early
ZGA_late
5h
9h
11h
14h
20h
24h
L1
sex
Female
Male

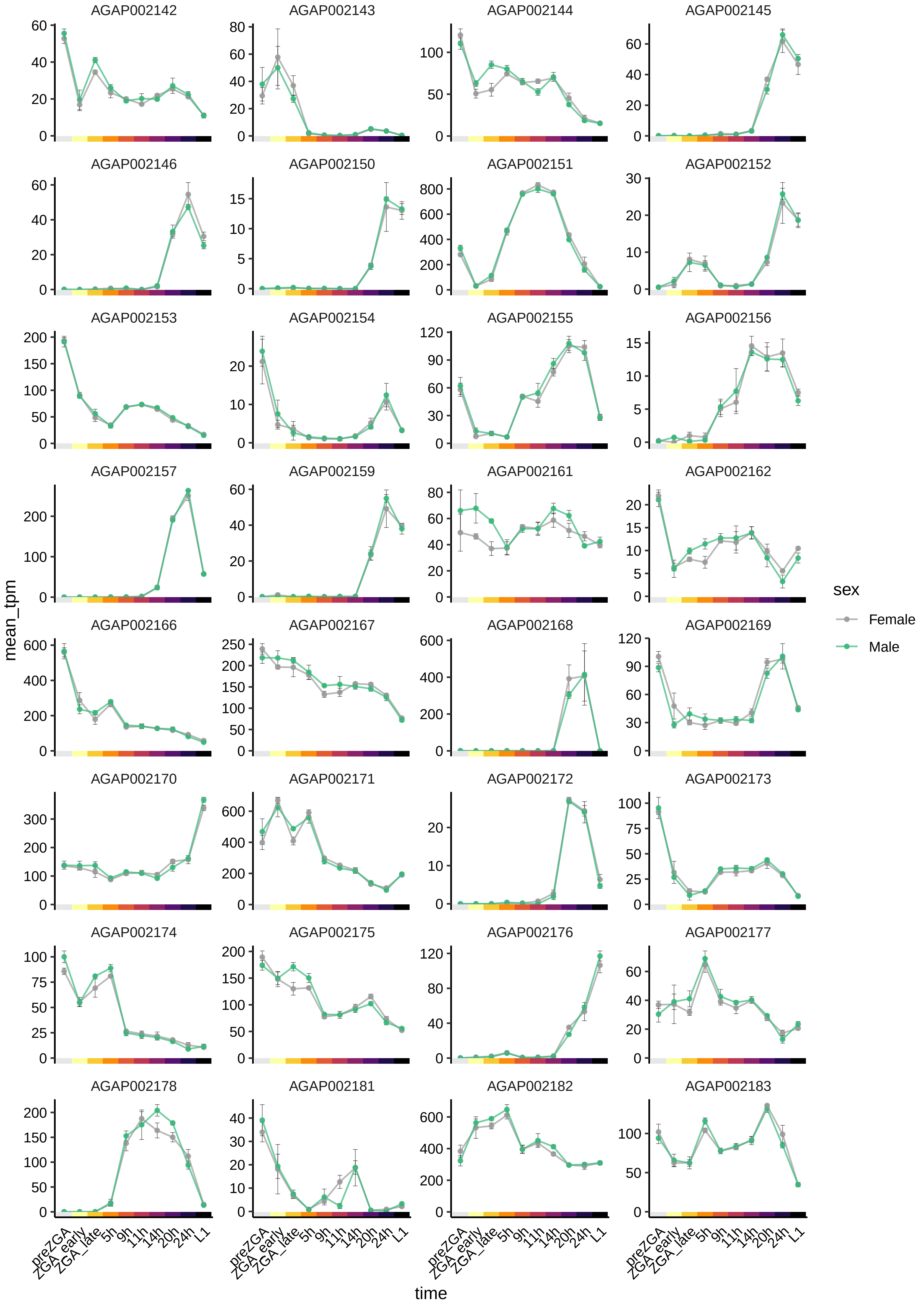

AGAP002142
AGAP002143
AGAP002144
AGAP002145
AGAP002146
AGAP002150
AGAP002151
AGAP002152
AGAP002153
AGAP002154
AGAP002155
AGAP002156
AGAP002157
AGAP002159
AGAP002161
AGAP002162
AGAP002166
AGAP002167
AGAP002168
AGAP002169
AGAP002170
AGAP002171
AGAP002172
AGAP002173
AGAP002174
AGAP002175
AGAP002176
AGAP002177
AGAP002178
AGAP002181
AGAP002182
AGAP002183
mean_tpm
time
preZGA
ZGA_early
ZGA_late
5h
9h
11h
14h
20h
24h
L1
sex
Female
Male

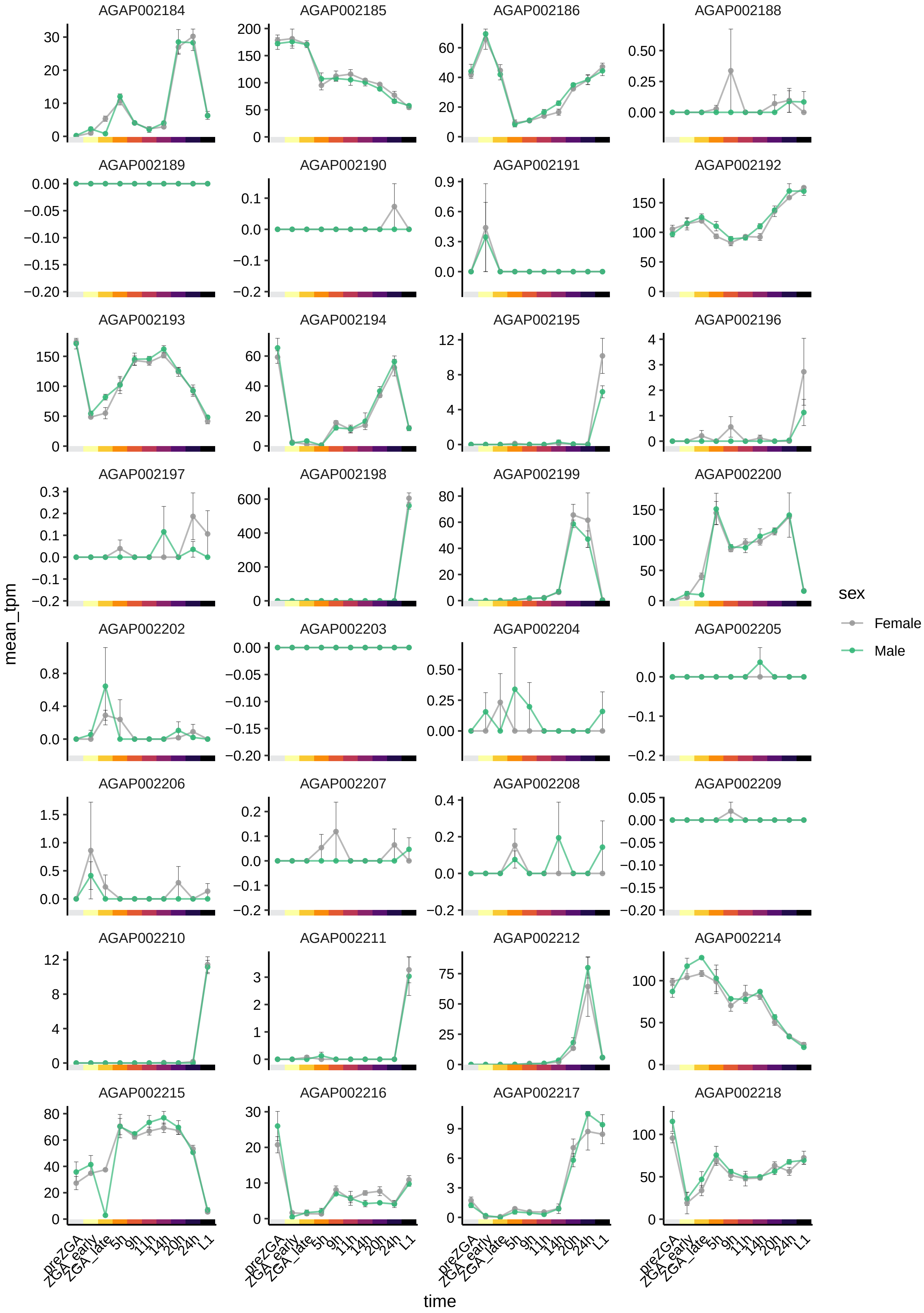
AGAP002184
AGAP002185
AGAP002186
AGAP002188
AGAP002189
AGAP002190
AGAP002191
AGAP002192
AGAP002193
AGAP002194
AGAP002195
AGAP002196
AGAP002197
AGAP002198
AGAP002199
AGAP002200
AGAP002202
AGAP002203
AGAP002204
AGAP002205
AGAP002206
AGAP002207
AGAP002208
AGAP002209
AGAP002210
AGAP002211
AGAP002212
AGAP002214
AGAP002215
AGAP002216
AGAP002217
AGAP002218
mean_tpm
time
preZGA
ZGA_early
ZGA_late
5h
9h
11h
14h
20h
24h
L1
sex
Female
Male

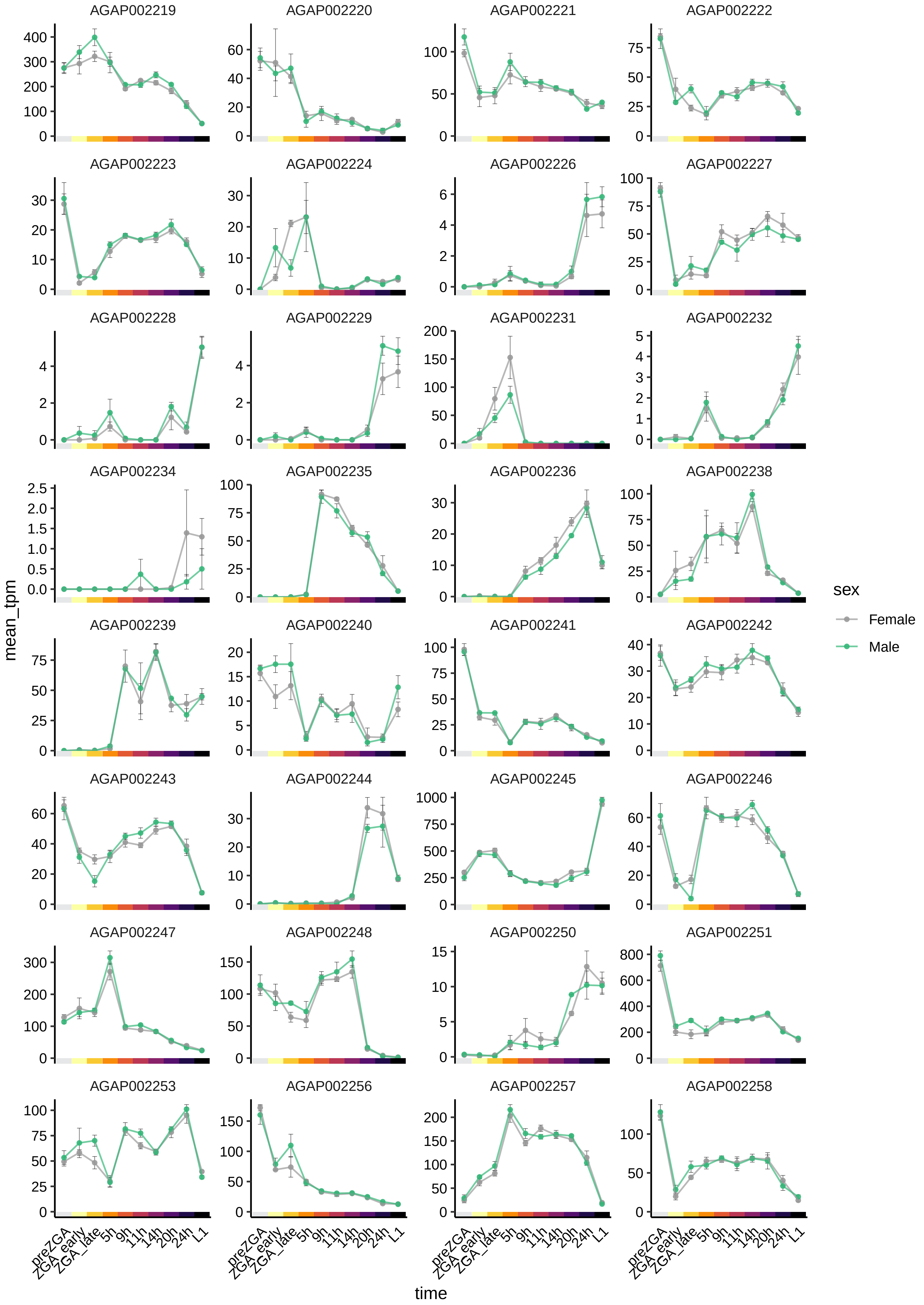
AGAP002219
AGAP002220
AGAP002221
AGAP002222
AGAP002223
AGAP002224
AGAP002226
AGAP002227
AGAP002228
AGAP002229
AGAP002231
AGAP002232
AGAP002234
AGAP002235
AGAP002236
AGAP002238
AGAP002239
AGAP002240
AGAP002241
AGAP002242
AGAP002243
AGAP002244
AGAP002245
AGAP002246
AGAP002247
AGAP002248
AGAP002250
AGAP002251
AGAP002253
AGAP002256
AGAP002257
AGAP002258
mean_tpm
time
preZGA
ZGA_early
ZGA_late
5h
9h
11h
14h
20h
24h
L1
sex
Female
Male

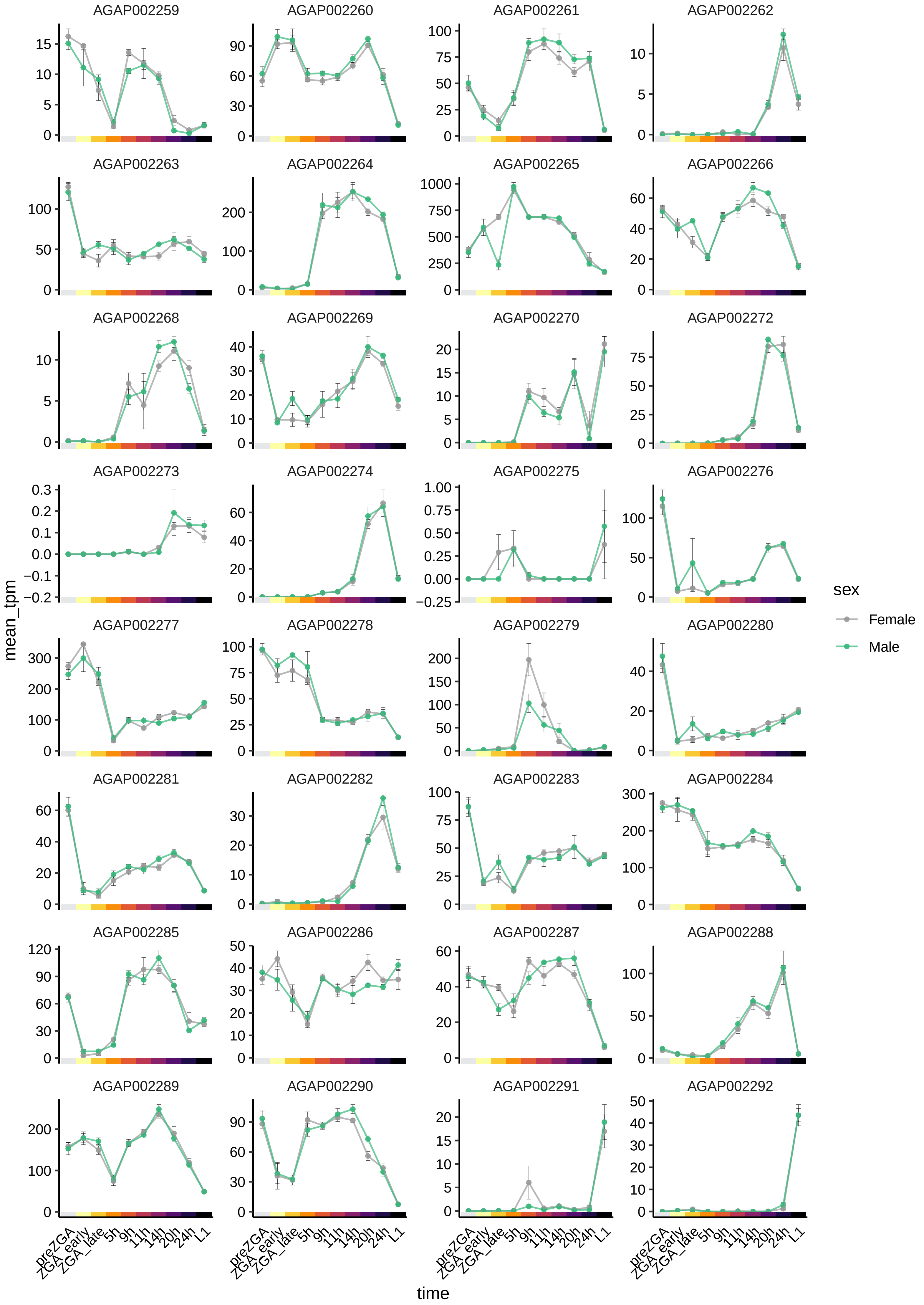
AGAP002259
AGAP002260
AGAP002261
AGAP002262
AGAP002263
AGAP002264
AGAP002265
AGAP002266
AGAP002268
AGAP002269
AGAP002270
AGAP002272
AGAP002273
AGAP002274
AGAP002275
AGAP002276
AGAP002277
AGAP002278
AGAP002279
AGAP002280
AGAP002281
AGAP002282
AGAP002283
AGAP002284
AGAP002285
AGAP002286
AGAP002287
AGAP002288
AGAP002289
AGAP002290
AGAP002291
AGAP002292
mean_tpm
time
preZGA
ZGA_early
ZGA_late
5h
9h
11h
14h
20h
24h
L1
sex
Female
Male

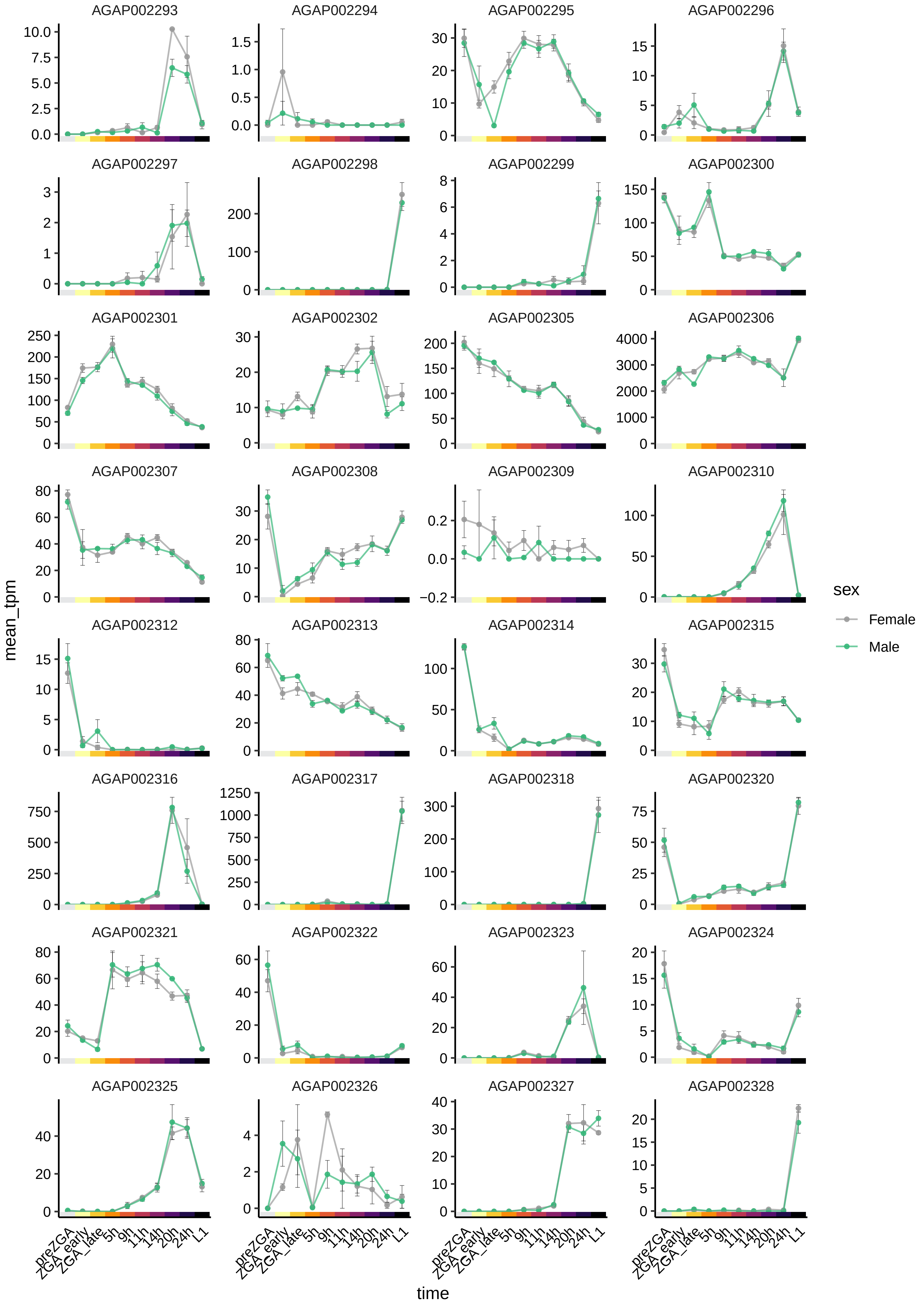
AGAP002293
AGAP002294
AGAP002295
AGAP002296
AGAP002297
AGAP002298
AGAP002299
AGAP002300
AGAP002301
AGAP002302
AGAP002305
AGAP002306
AGAP002307
AGAP002308
AGAP002309
AGAP002310
AGAP002312
AGAP002313
AGAP002314
AGAP002315
AGAP002316
AGAP002317
AGAP002318
AGAP002320
AGAP002321
AGAP002322
AGAP002323
AGAP002324
AGAP002325
AGAP002326
AGAP002327
AGAP002328
mean_tpm
time
preZGA
ZGA_early
ZGA_late
5h
9h
11h
14h
20h
24h
L1
sex
Female
Male

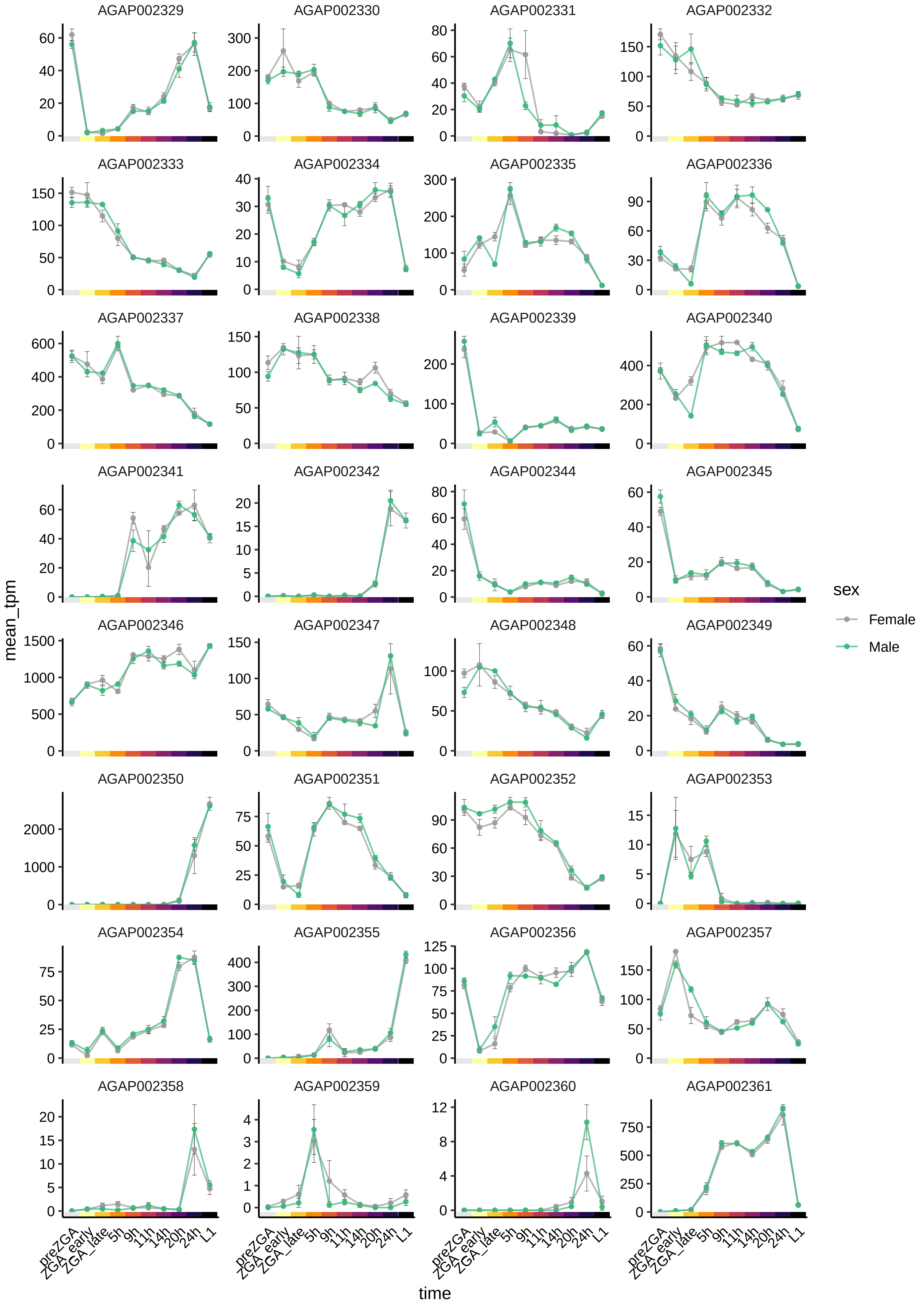

AGAP002329
AGAP002330
AGAP002331
AGAP002332
AGAP002333
AGAP002334
AGAP002335
AGAP002336
AGAP002337
AGAP002338
AGAP002339
AGAP002340
AGAP002341
AGAP002342
AGAP002344
AGAP002345
AGAP002346
AGAP002347
AGAP002348
AGAP002349
AGAP002350
AGAP002351
AGAP002352
AGAP002353
AGAP002354
AGAP002355
AGAP002356
AGAP002357
AGAP002358
AGAP002359
AGAP002360
AGAP002361
mean_tpm
time
preZGA
ZGA_early
ZGA_late
5h
9h
11h
14h
20h
24h
L1
sex
Female
Male

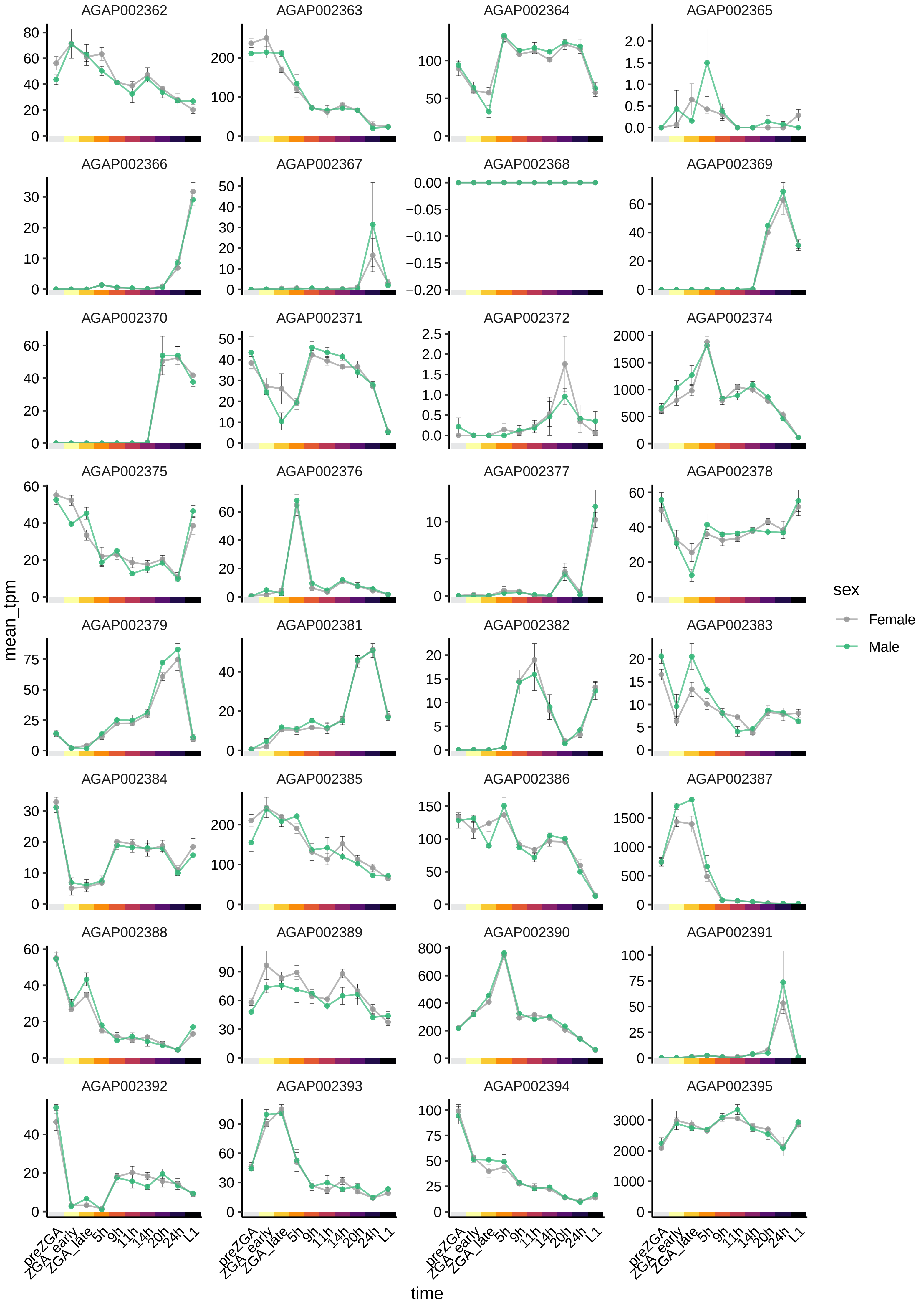
AGAP002362
AGAP002363
AGAP002364
AGAP002365
AGAP002366
AGAP002367
AGAP002368
AGAP002369
AGAP002370
AGAP002371
AGAP002372
AGAP002374
AGAP002375
AGAP002376
AGAP002377
AGAP002378
AGAP002379
AGAP002381
AGAP002382
AGAP002383
AGAP002384
AGAP002385
AGAP002386
AGAP002387
AGAP002388
AGAP002389
AGAP002390
AGAP002391
AGAP002392
AGAP002393
AGAP002394
AGAP002395
mean_tpm
time
preZGA
ZGA_early
ZGA_late
5h
9h
11h
14h
20h
24h
L1
sex
Female
Male

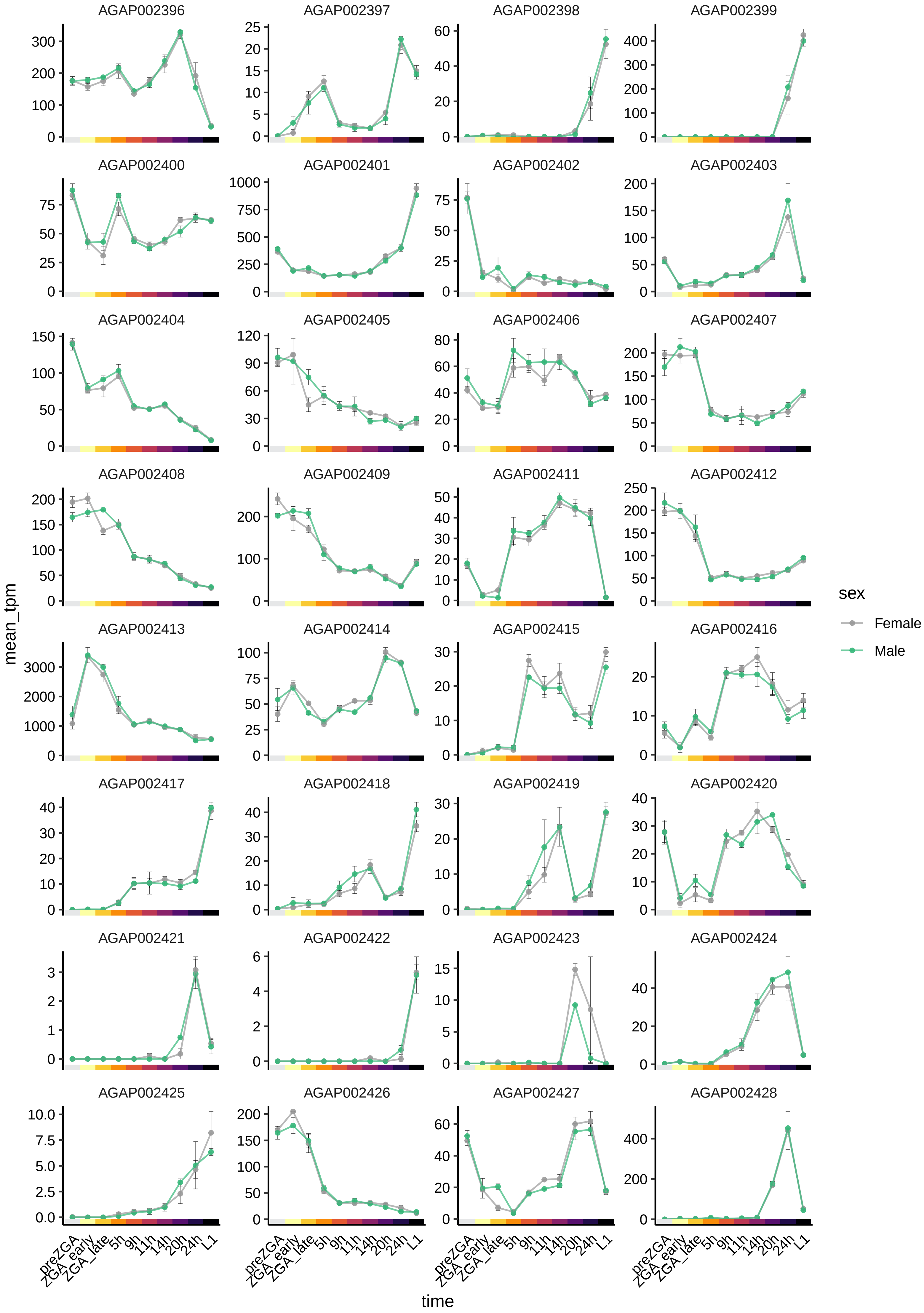

AGAP002396
AGAP002397
AGAP002398
AGAP002399
AGAP002400
AGAP002401
AGAP002402
AGAP002403
AGAP002404
AGAP002405
AGAP002406
AGAP002407
AGAP002408
AGAP002409
AGAP002411
AGAP002412
AGAP002413
AGAP002414
AGAP002415
AGAP002416
AGAP002417
AGAP002418
AGAP002419
AGAP002420
AGAP002421
AGAP002422
AGAP002423
AGAP002424
AGAP002425
AGAP002426
AGAP002427
AGAP002428
mean_tpm
time
preZGA
ZGA_early
ZGA_late
5h
9h
11h
14h
20h
24h
L1
sex
Female
Male

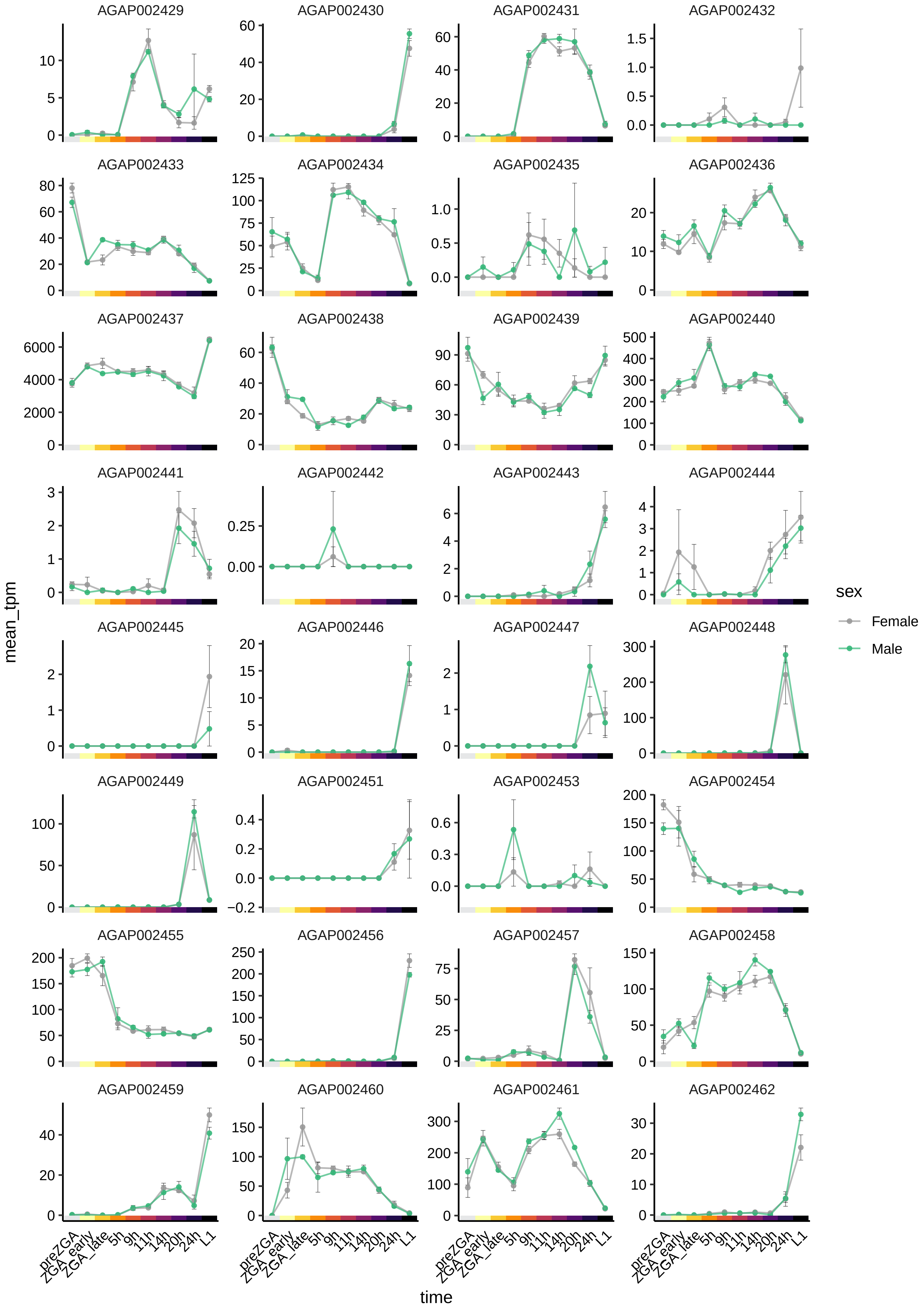

AGAP002429
AGAP002430
AGAP002431
AGAP002432
AGAP002433
AGAP002434
AGAP002435
AGAP002436
AGAP002437
AGAP002438
AGAP002439
AGAP002440
AGAP002441
AGAP002442
AGAP002443
AGAP002444
AGAP002445
AGAP002446
AGAP002447
AGAP002448
AGAP002449
AGAP002451
AGAP002453
AGAP002454
AGAP002455
AGAP002456
AGAP002457
AGAP002458
AGAP002459
AGAP002460
AGAP002461
AGAP002462
mean_tpm
time
preZGA
ZGA_early
ZGA_late
5h
9h
11h
14h
20h
24h
L1
sex
Female
Male

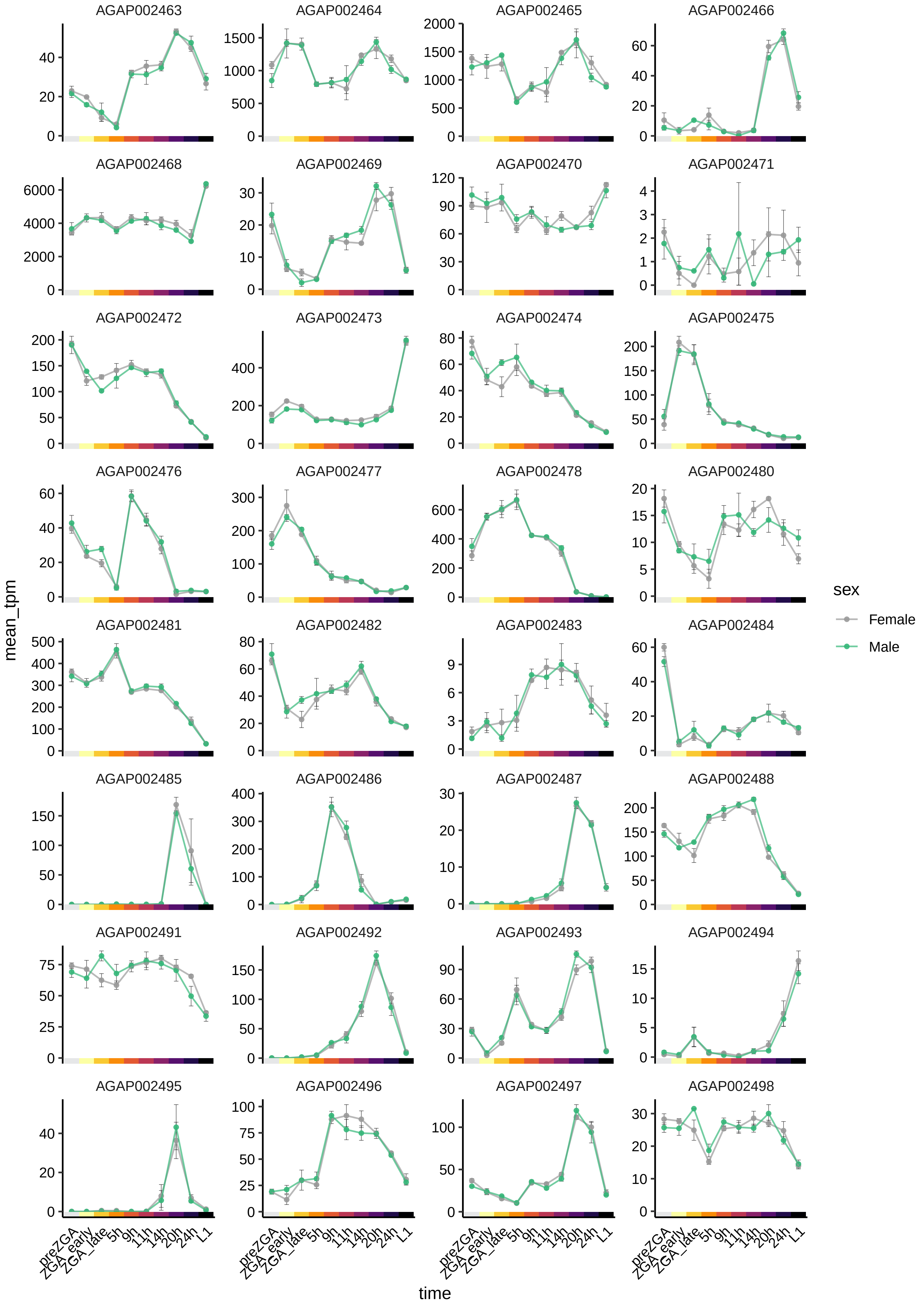

AGAP002463
AGAP002464
AGAP002465
AGAP002466
AGAP002468
AGAP002469
AGAP002470
AGAP002471
AGAP002472
AGAP002473
AGAP002474
AGAP002475
AGAP002476
AGAP002477
AGAP002478
AGAP002480
AGAP002481
AGAP002482
AGAP002483
AGAP002484
AGAP002485
AGAP002486
AGAP002487
AGAP002488
AGAP002491
AGAP002492
AGAP002493
AGAP002494
AGAP002495
AGAP002496
AGAP002497
AGAP002498
mean_tpm
time
preZGA
ZGA_early
ZGA_late
5h
9h
11h
14h
20h
24h
L1
sex
Female
Male

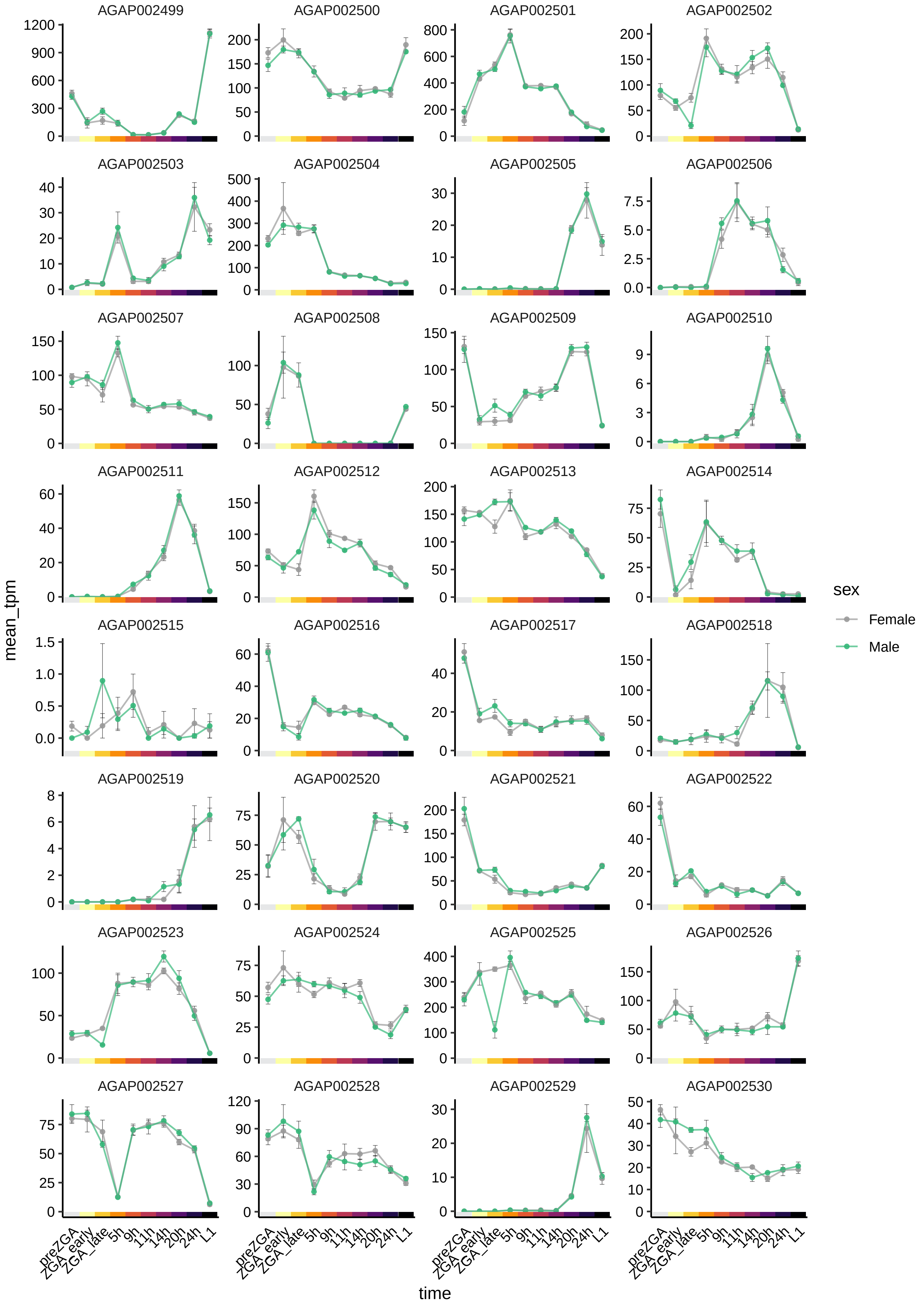

AGAP002499
AGAP002500
AGAP002501
AGAP002502
AGAP002503
AGAP002504
AGAP002505
AGAP002506
AGAP002507
AGAP002508
AGAP002509
AGAP002510
AGAP002511
AGAP002512
AGAP002513
AGAP002514
AGAP002515
AGAP002516
AGAP002517
AGAP002518
AGAP002519
AGAP002520
AGAP002521
AGAP002522
AGAP002523
AGAP002524
AGAP002525
AGAP002526
AGAP002527
AGAP002528
AGAP002529
AGAP002530
mean_tpm
time
preZGA
ZGA_early
ZGA_late
5h
9h
11h
14h
20h
24h
L1
sex
Female
Male

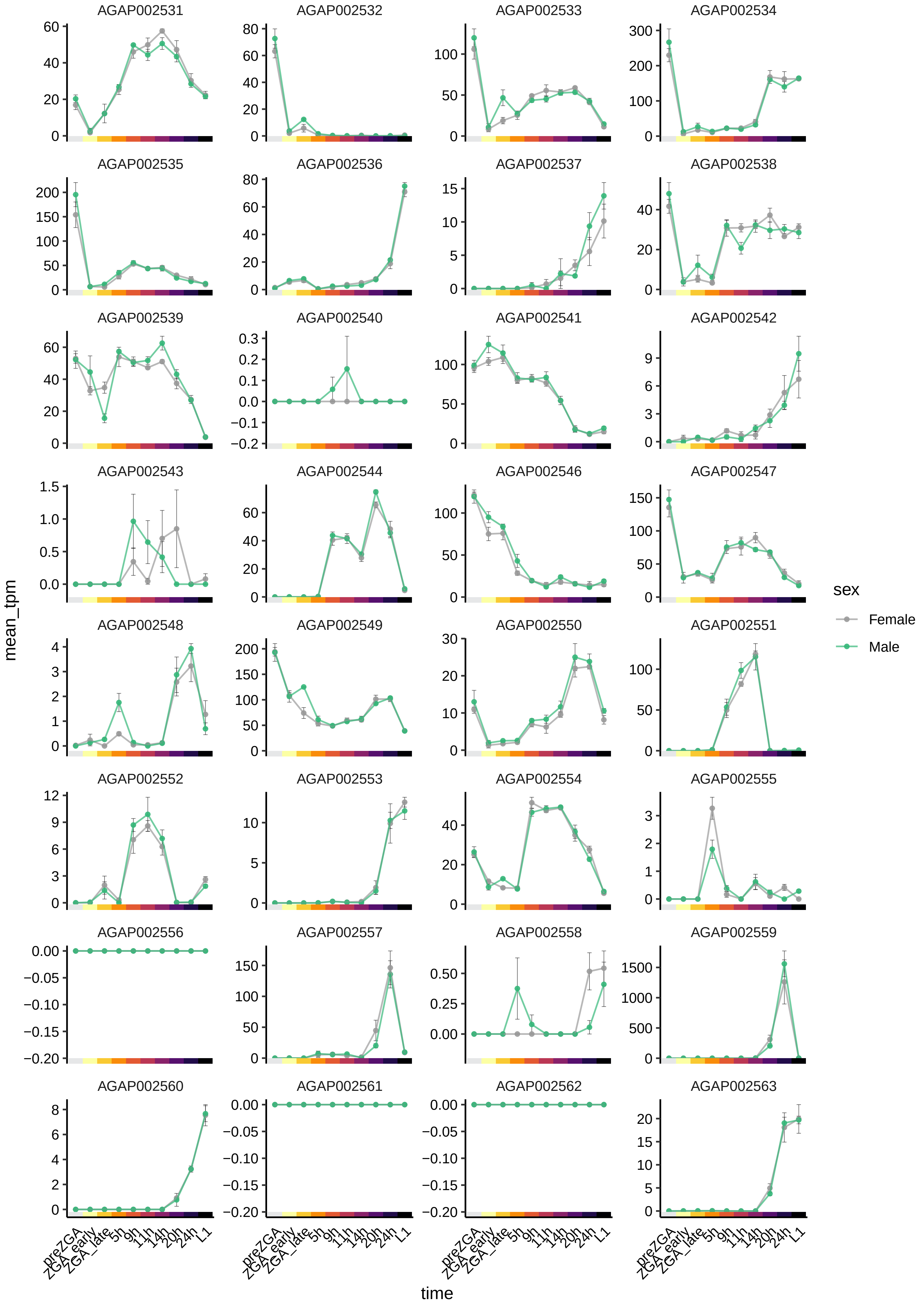
AGAP002531
AGAP002532
AGAP002533
AGAP002534
AGAP002535
AGAP002536
AGAP002537
AGAP002538
AGAP002539
AGAP002540
AGAP002541
AGAP002542
AGAP002543
AGAP002544
AGAP002546
AGAP002547
AGAP002548
AGAP002549
AGAP002550
AGAP002551
AGAP002552
AGAP002553
AGAP002554
AGAP002555
AGAP002556
AGAP002557
AGAP002558
AGAP002559
AGAP002560
AGAP002561
AGAP002562
AGAP002563
mean_tpm
time
preZGA
ZGA_early
ZGA_late
5h
9h
11h
14h
20h
24h
L1
sex
Female
Male

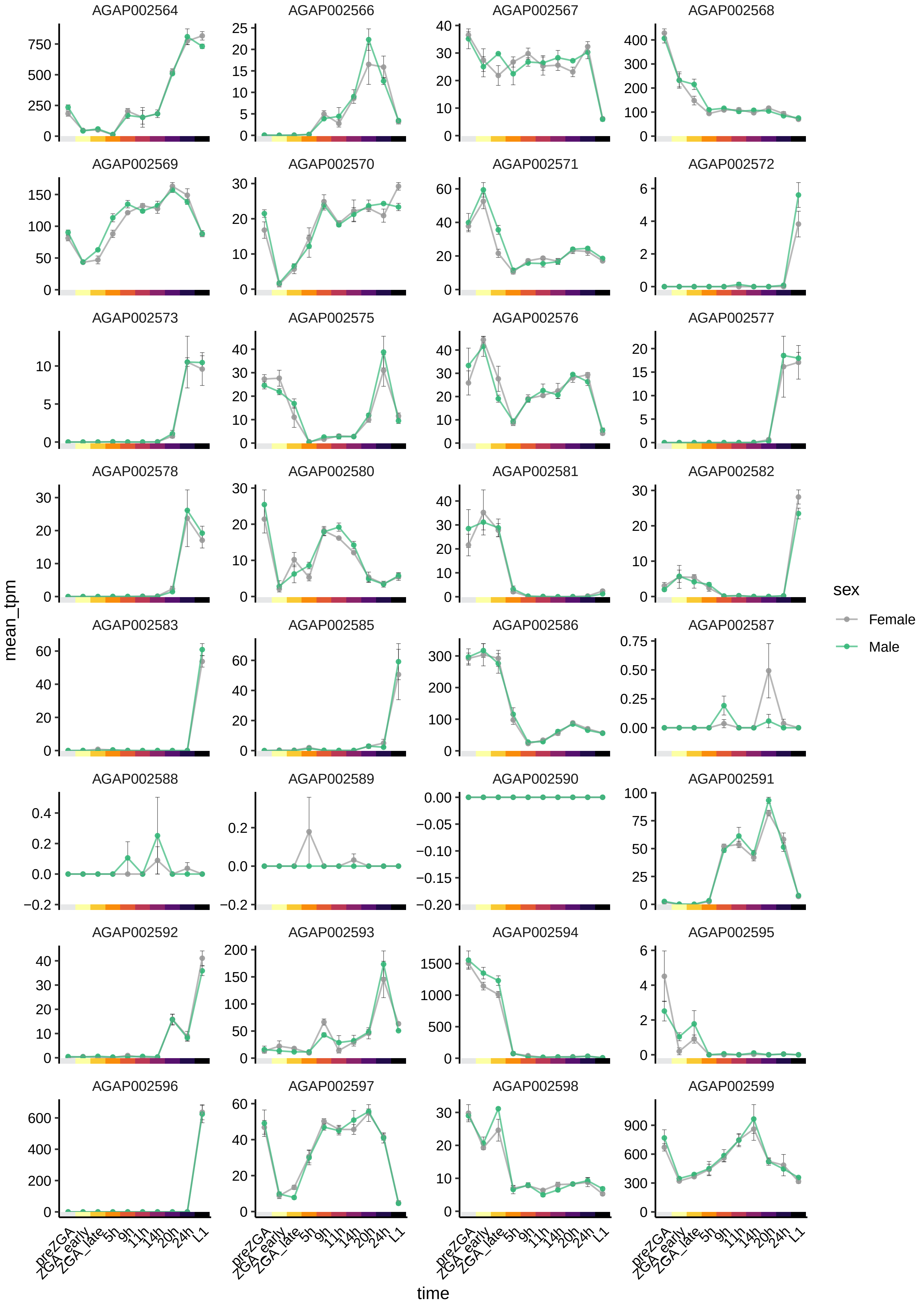

AGAP002564
AGAP002566
AGAP002567
AGAP002568
AGAP002569
AGAP002570
AGAP002571
AGAP002572
AGAP002573
AGAP002575
AGAP002576
AGAP002577
AGAP002578
AGAP002580
AGAP002581
AGAP002582
AGAP002583
AGAP002585
AGAP002586
AGAP002587
AGAP002588
AGAP002589
AGAP002590
AGAP002591
AGAP002592
AGAP002593
AGAP002594
AGAP002595
AGAP002596
AGAP002597
AGAP002598
AGAP002599
mean_tpm
time
preZGA
ZGA_early
ZGA_late
5h
9h
11h
14h
20h
24h
L1
sex
Female
Male

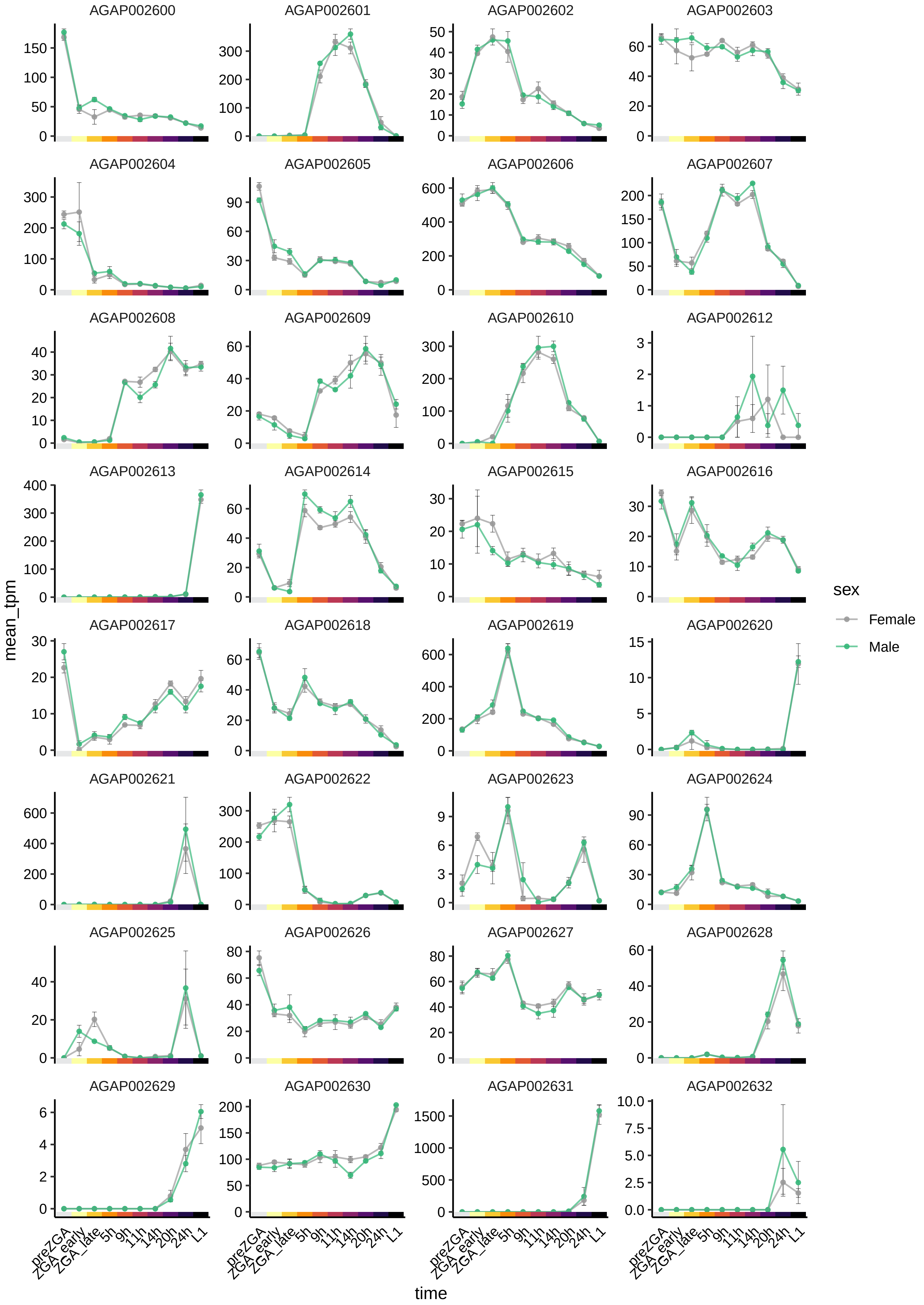

AGAP002600
AGAP002601
AGAP002602
AGAP002603
AGAP002604
AGAP002605
AGAP002606
AGAP002607
AGAP002608
AGAP002609
AGAP002610
AGAP002612
AGAP002613
AGAP002614
AGAP002615
AGAP002616
AGAP002617
AGAP002618
AGAP002619
AGAP002620
AGAP002621
AGAP002622
AGAP002623
AGAP002624
AGAP002625
AGAP002626
AGAP002627
AGAP002628
AGAP002629
AGAP002630
AGAP002631
AGAP002632
mean_tpm
time
preZGA
ZGA_early
ZGA_late
5h
9h
11h
14h
20h
24h
L1
sex
Female
Male

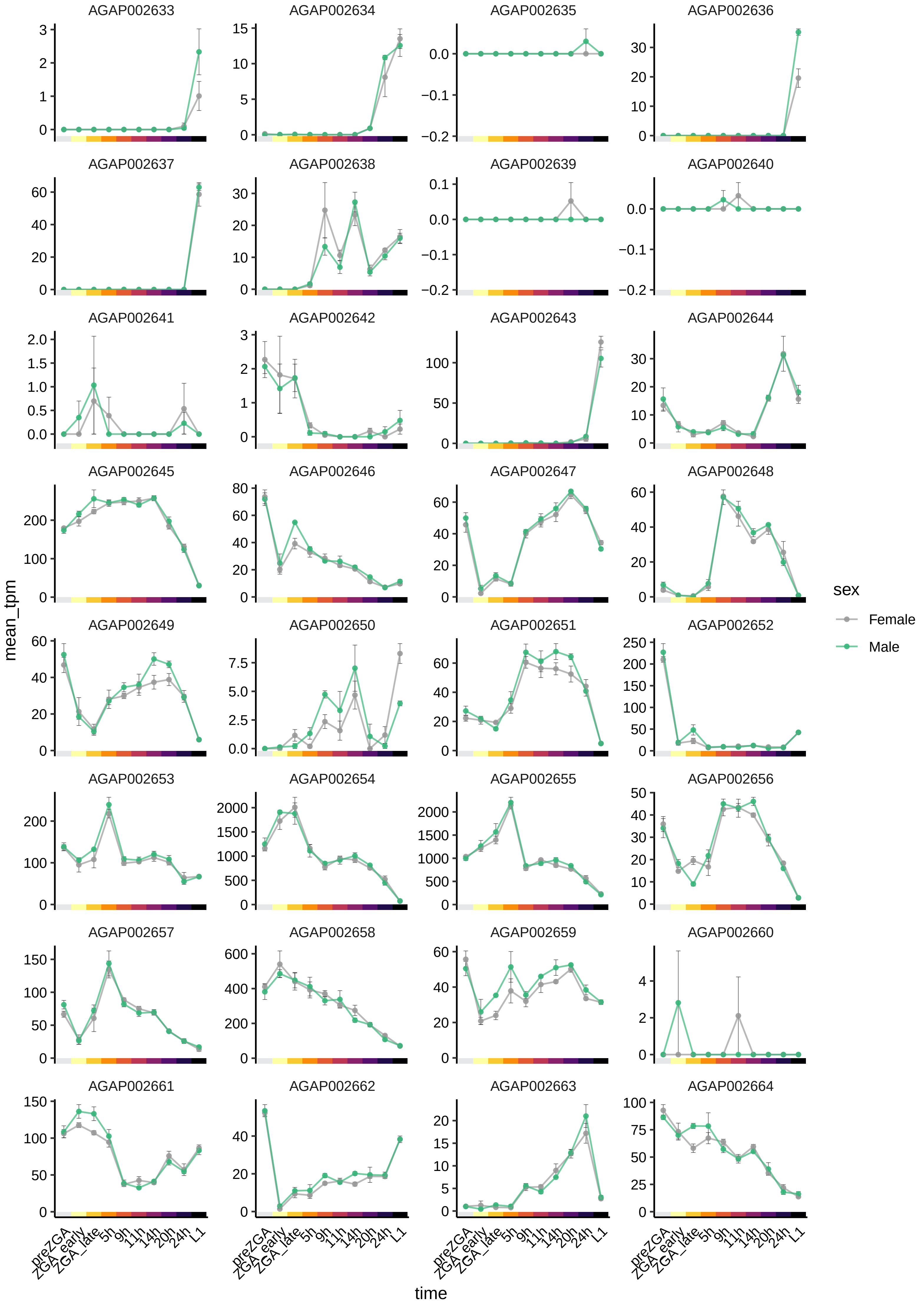

AGAP002633
AGAP002634
AGAP002635
AGAP002636
AGAP002637
AGAP002638
AGAP002639
AGAP002640
AGAP002641
AGAP002642
AGAP002643
AGAP002644
AGAP002645
AGAP002646
AGAP002647
AGAP002648
AGAP002649
AGAP002650
AGAP002651
AGAP002652
AGAP002653
AGAP002654
AGAP002655
AGAP002656
AGAP002657
AGAP002658
AGAP002659
AGAP002660
AGAP002661
AGAP002662
AGAP002663
AGAP002664
mean_tpm
time
preZGA
ZGA_early
ZGA_late
5h
9h
11h
14h
20h
24h
L1
sex
Female
Male

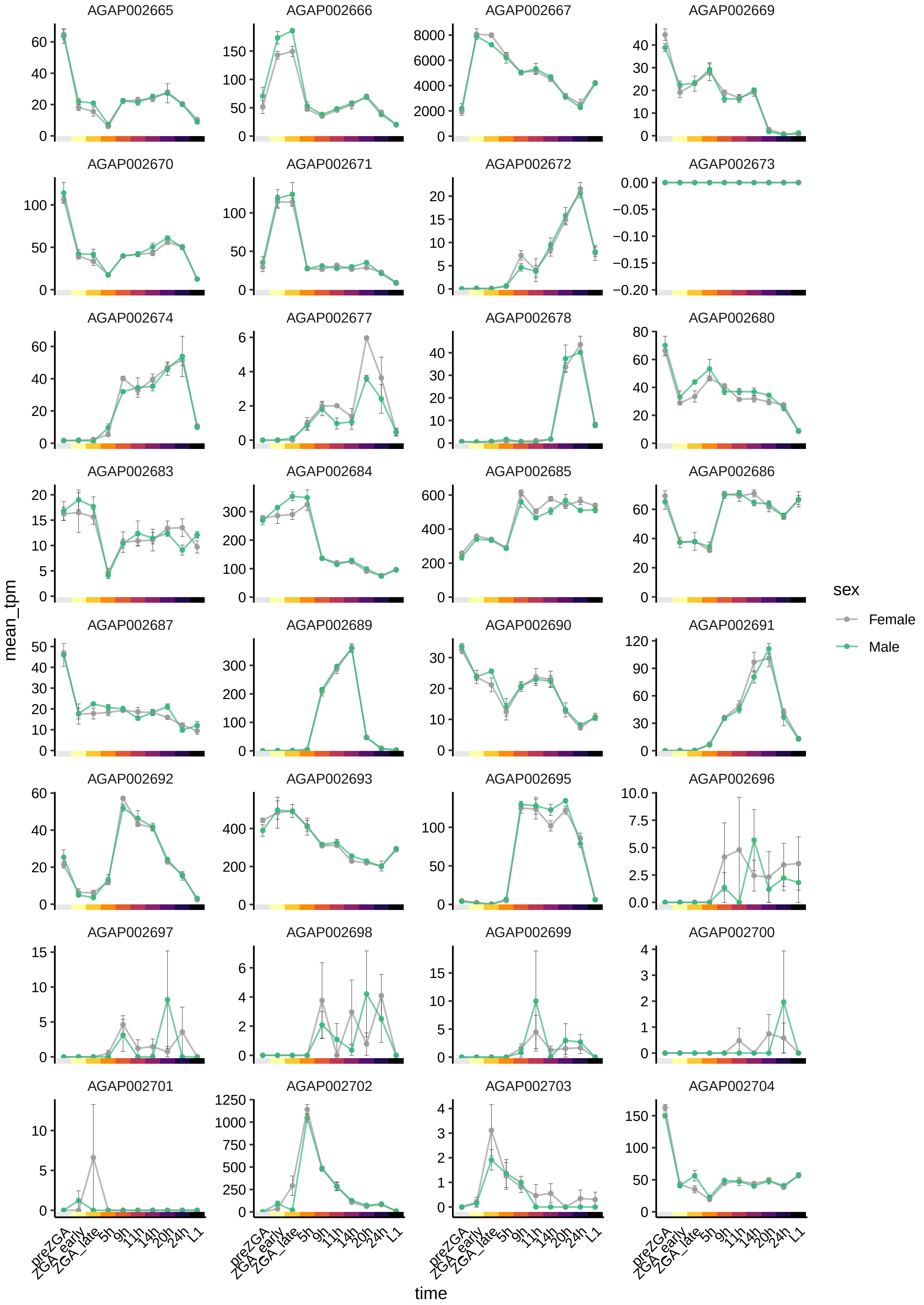
AGAP002665
AGAP002666
AGAP002667
AGAP002669
AGAP002670
AGAP002671
AGAP002672
AGAP002673
AGAP002674
AGAP002677
AGAP002678
AGAP002680
AGAP002683
AGAP002684
AGAP002685
AGAP002686
AGAP002687
AGAP002689
AGAP002690
AGAP002691
AGAP002692
AGAP002693
AGAP002695
AGAP002696
AGAP002697
AGAP002698
AGAP002699
AGAP002700
AGAP002701
AGAP002702
AGAP002703
AGAP002704
mean_tpm
time
preZGA
ZGA_early
ZGA_late
5h
9h
11h
14h
20h
24h
L1
sex
Female
Male

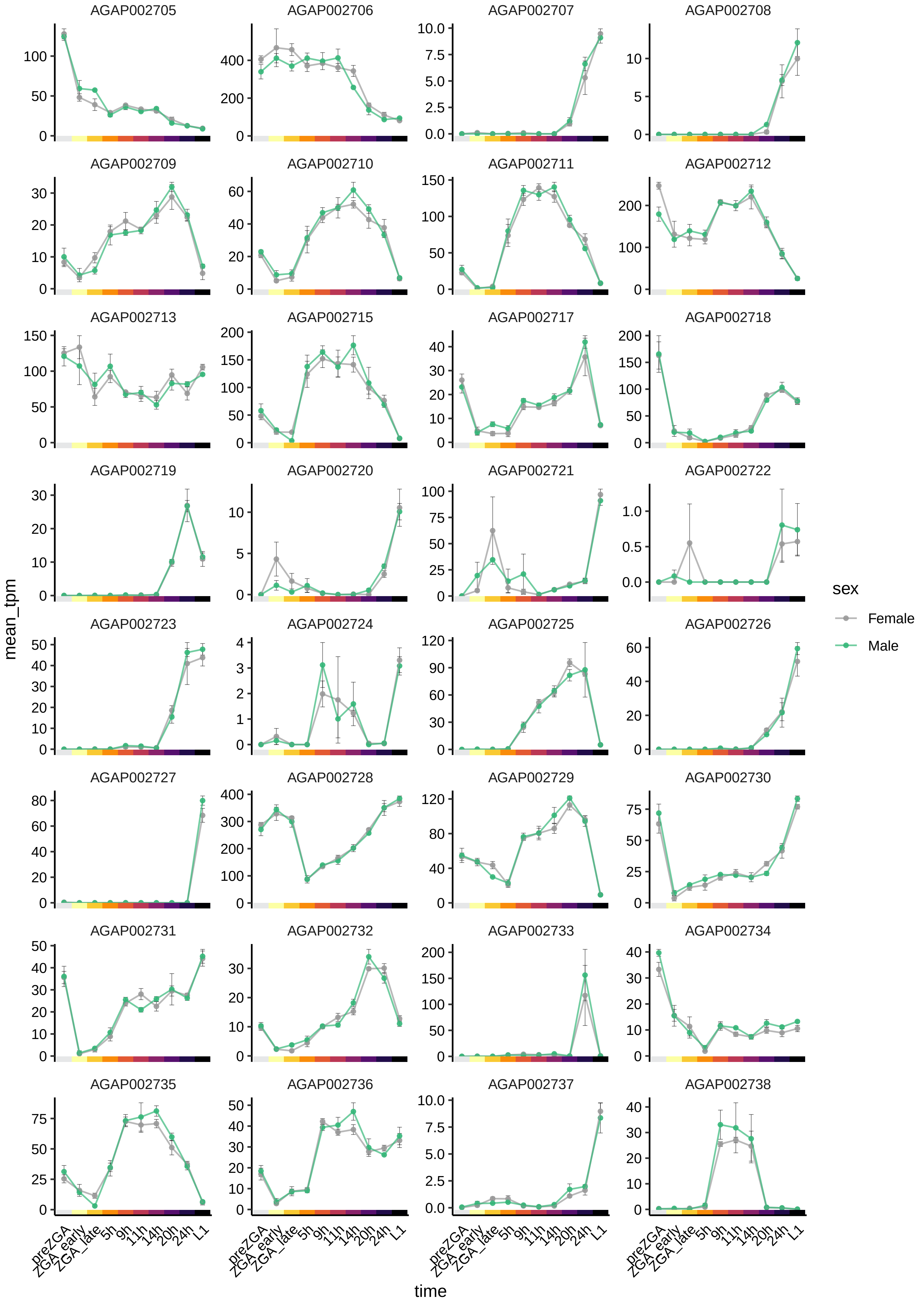
AGAP002705
AGAP002706
AGAP002707
AGAP002708
AGAP002709
AGAP002710
AGAP002711
AGAP002712
AGAP002713
AGAP002715
AGAP002717
AGAP002718
AGAP002719
AGAP002720
AGAP002721
AGAP002722
AGAP002723
AGAP002724
AGAP002725
AGAP002726
AGAP002727
AGAP002728
AGAP002729
AGAP002730
AGAP002731
AGAP002732
AGAP002733
AGAP002734
AGAP002735
AGAP002736
AGAP002737
AGAP002738
mean_tpm
preZGA
ZGA_early
ZGA_late
5h
9h
11h
14h
20h
24h
L1
time
sex
Female
Male

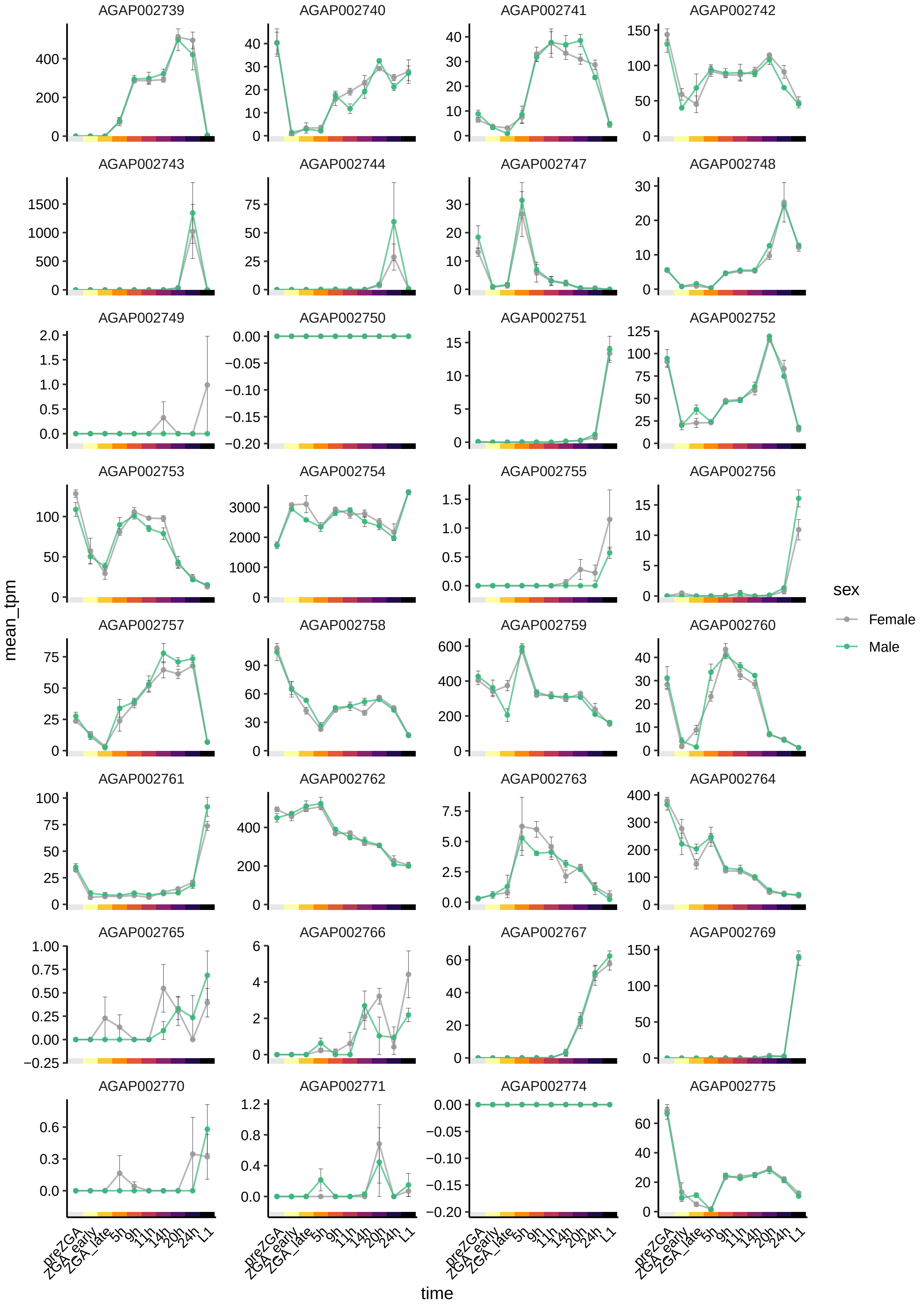

AGAP002739
AGAP002740
AGAP002741
AGAP002742
AGAP002743
AGAP002744
AGAP002747
AGAP002748
AGAP002749
AGAP002750
AGAP002751
AGAP002752
AGAP002753
AGAP002754
AGAP002755
AGAP002756
AGAP002757
AGAP002758
AGAP002759
AGAP002760
AGAP002761
AGAP002762
AGAP002763
AGAP002764
AGAP002765
AGAP002766
AGAP002767
AGAP002769
AGAP002770
AGAP002771
AGAP002774
AGAP002775
mean_tpm
time
preZGA
ZGA_early
ZGA_late
5h
9h
11h
14h
20h
24h
L1
sex
Female
Male

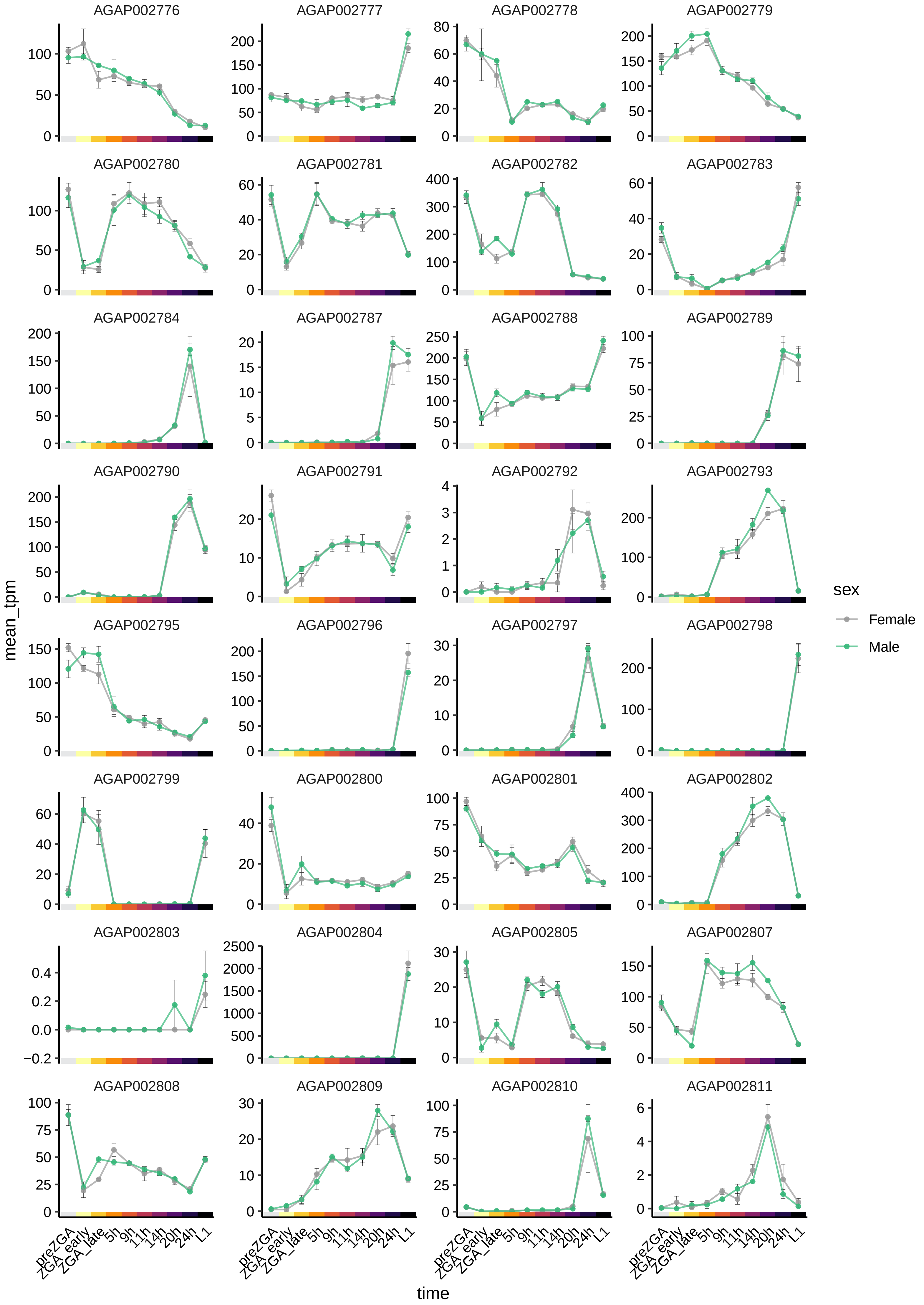

AGAP002776
AGAP002777
AGAP002778
AGAP002779
AGAP002780
AGAP002781
AGAP002782
AGAP002783
AGAP002784
AGAP002787
AGAP002788
AGAP002789
AGAP002790
AGAP002791
AGAP002792
AGAP002793
AGAP002795
AGAP002796
AGAP002797
AGAP002798
AGAP002799
AGAP002800
AGAP002801
AGAP002802
AGAP002803
AGAP002804
AGAP002805
AGAP002807
AGAP002808
AGAP002809
AGAP002810
AGAP002811
mean_tpm
time
preZGA
ZGA_early
ZGA_late
5h
9h
11h
14h
20h
24h
L1
sex
Female
Male

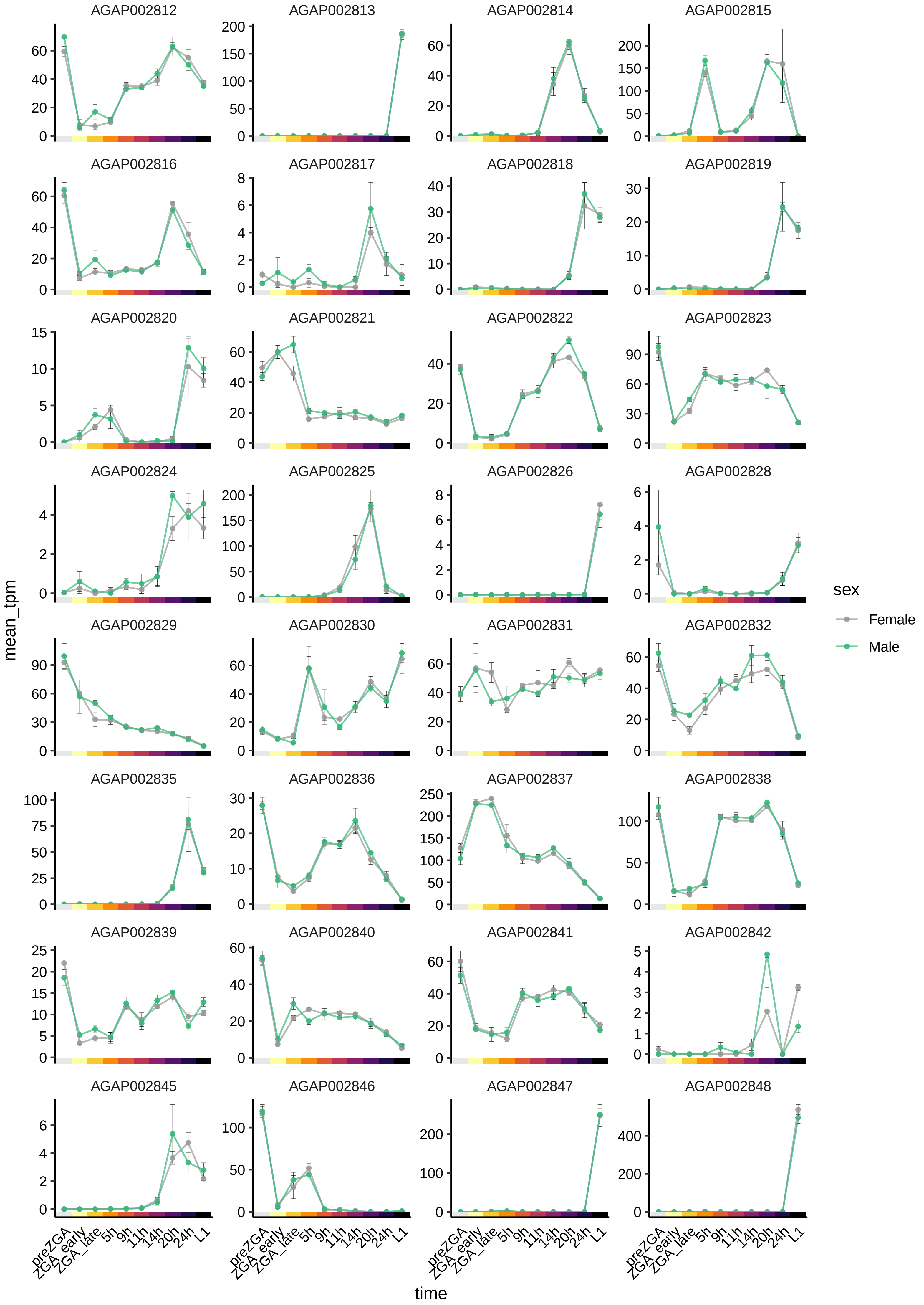
AGAP002812
AGAP002813
AGAP002814
AGAP002815
AGAP002816
AGAP002817
AGAP002818
AGAP002819
AGAP002820
AGAP002821
AGAP002822
AGAP002823
AGAP002824
AGAP002825
AGAP002826
AGAP002828
AGAP002829
AGAP002830
AGAP002831
AGAP002832
AGAP002835
AGAP002836
AGAP002837
AGAP002838
AGAP002839
AGAP002840
AGAP002841
AGAP002842
AGAP002845
AGAP002846
AGAP002847
AGAP002848
mean_tpm
time
preZGA
ZGA_early
ZGA_late
5h
9h
11h
14h
20h
24h
L1
sex
Female
Male

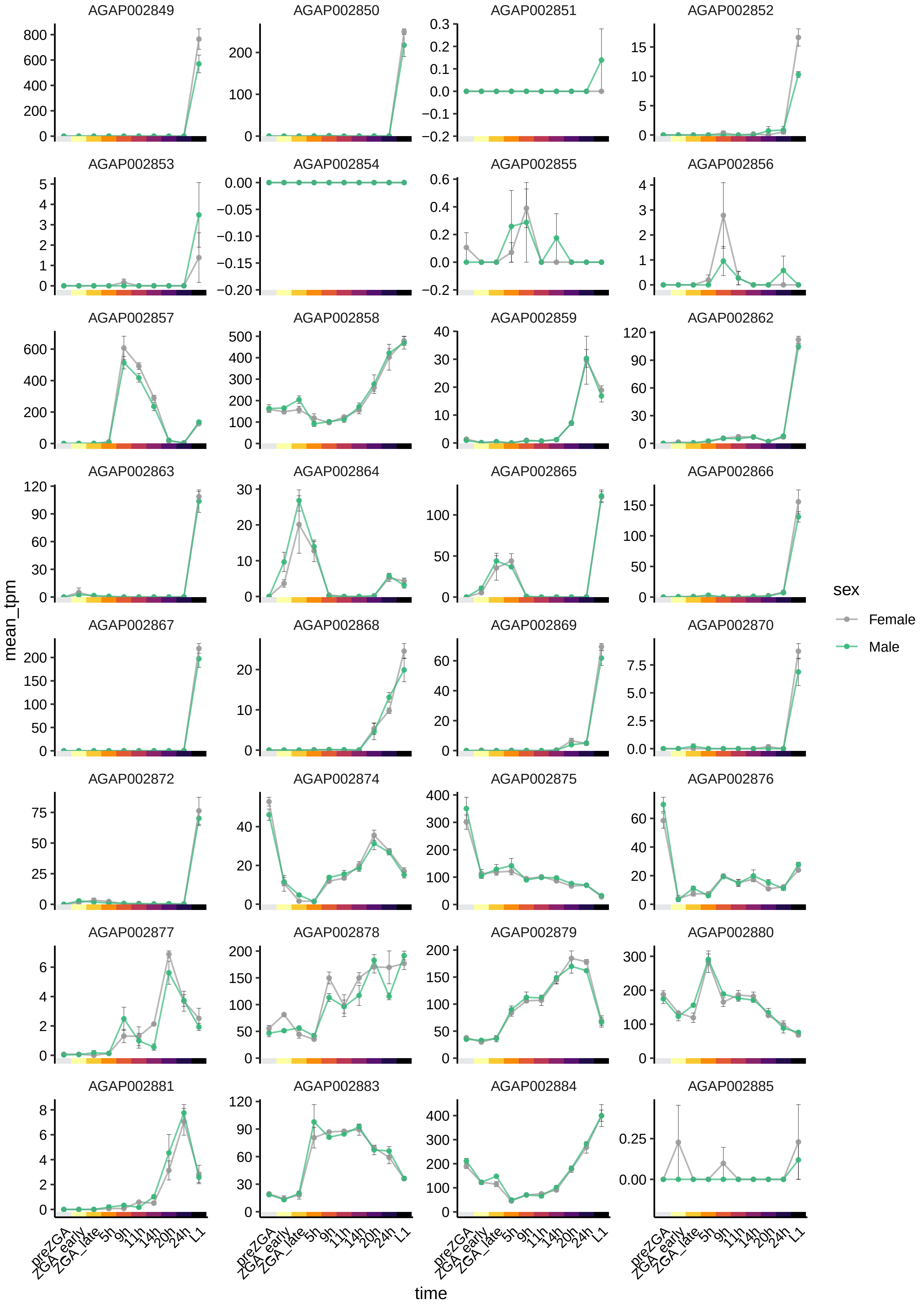

AGAP002849
AGAP002850
AGAP002851
AGAP002852
AGAP002853
AGAP002854
AGAP002855
AGAP002856
AGAP002857
AGAP002858
AGAP002859
AGAP002862
AGAP002863
AGAP002864
AGAP002865
AGAP002866
AGAP002867
AGAP002868
AGAP002869
AGAP002870
AGAP002872
AGAP002874
AGAP002875
AGAP002876
AGAP002877
AGAP002878
AGAP002879
AGAP002880
AGAP002881
AGAP002883
AGAP002884
AGAP002885
mean_tpm
time
preZGA
ZGA_early
ZGA_late
5h
9h
11h
14h
20h
24h
L1
sex
Female
Male

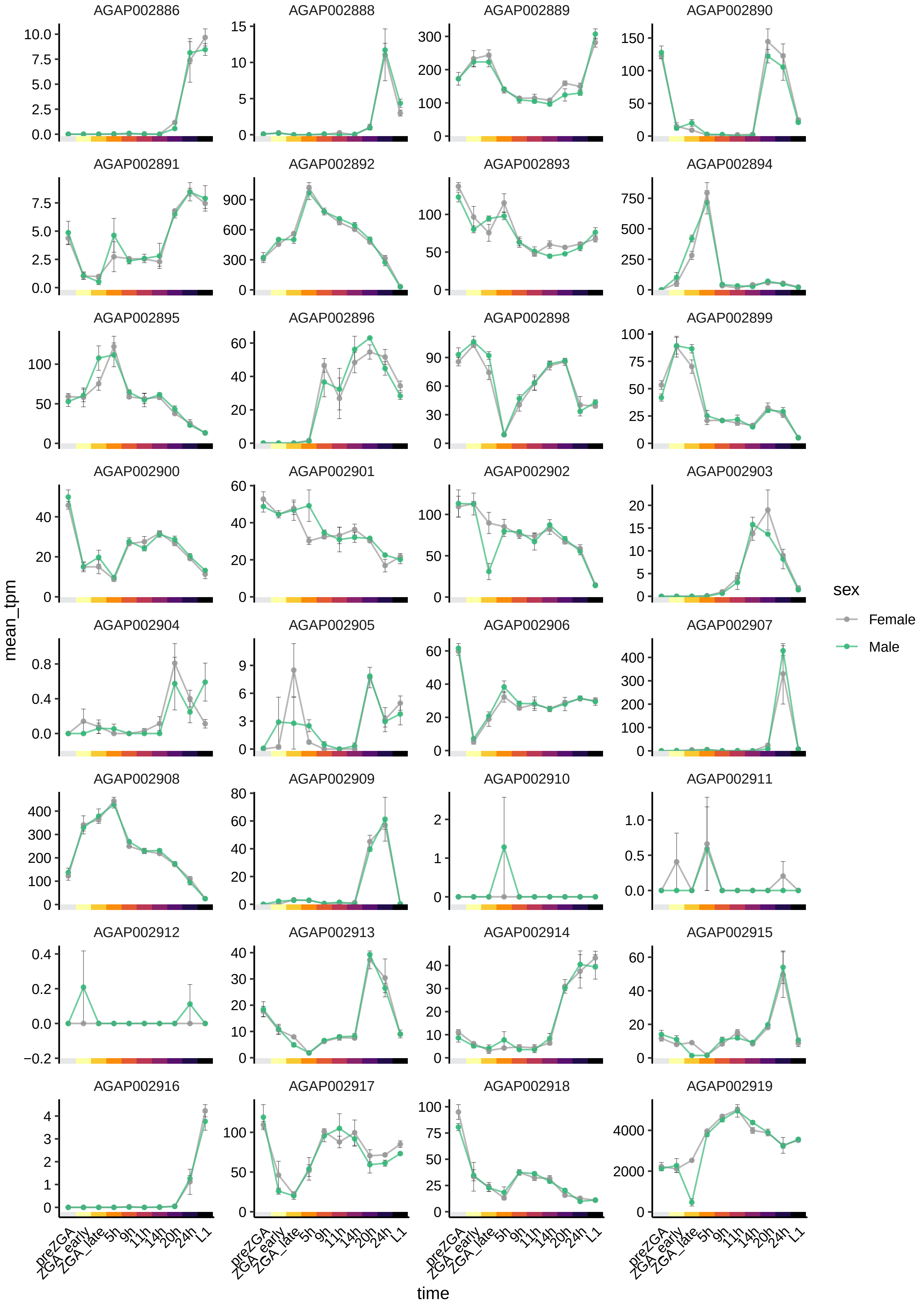

AGAP002886
AGAP002888
AGAP002889
AGAP002890
AGAP002891
AGAP002892
AGAP002893
AGAP002894
AGAP002895
AGAP002896
AGAP002898
AGAP002899
AGAP002900
AGAP002901
AGAP002902
AGAP002903
AGAP002904
AGAP002905
AGAP002906
AGAP002907
AGAP002908
AGAP002909
AGAP002910
AGAP002911
AGAP002912
AGAP002913
AGAP002914
AGAP002915
AGAP002916
AGAP002917
AGAP002918
AGAP002919
mean_tpm
time
preZGA
ZGA_early
ZGA_late
5h
9h
11h
14h
20h
24h
L1
sex
Female
Male

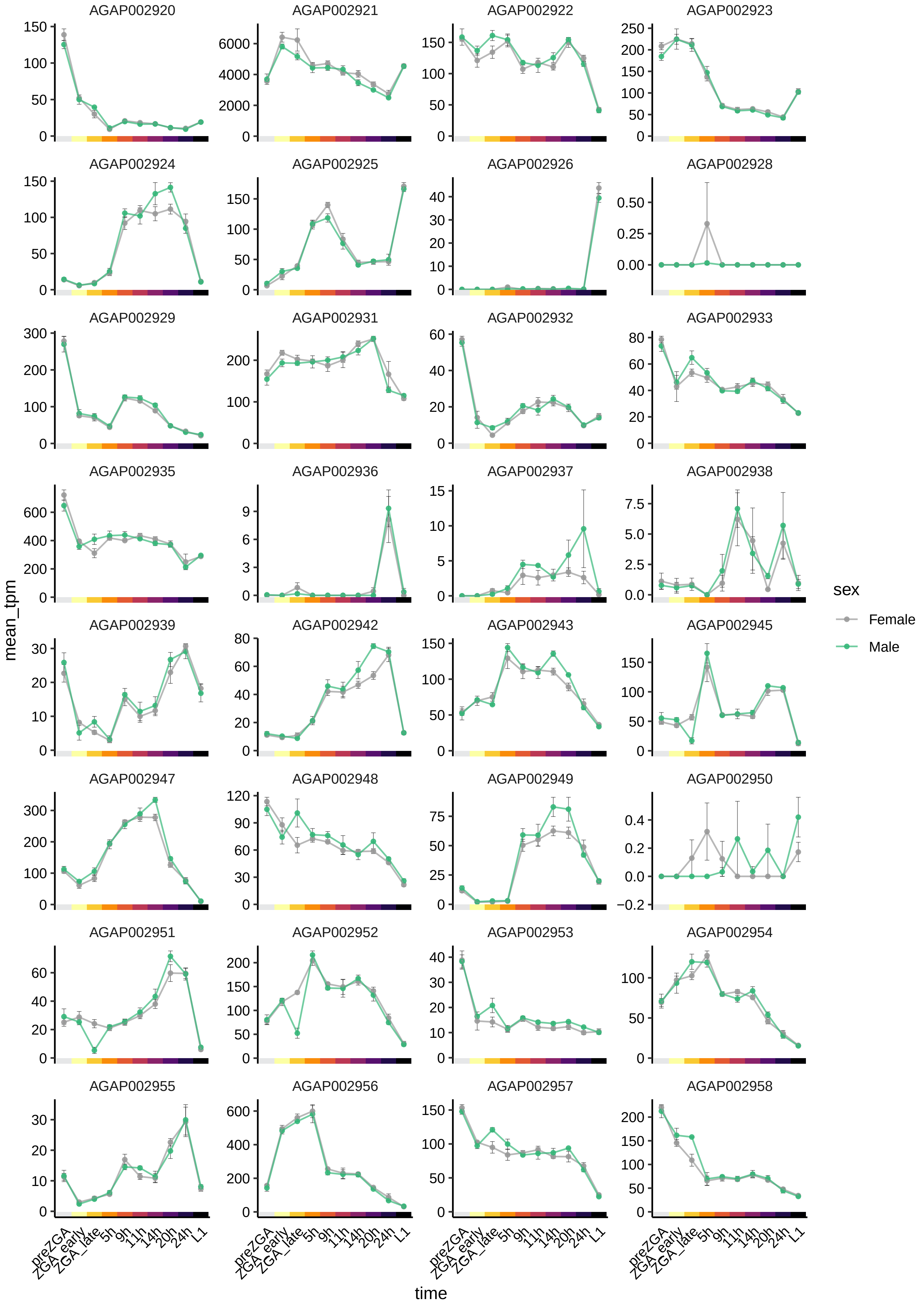

AGAP002920
AGAP002921
AGAP002922
AGAP002923
AGAP002924
AGAP002925
AGAP002926
AGAP002928
AGAP002929
AGAP002931
AGAP002932
AGAP002933
AGAP002935
AGAP002936
AGAP002937
AGAP002938
AGAP002939
AGAP002942
AGAP002943
AGAP002945
AGAP002947
AGAP002948
AGAP002949
AGAP002950
AGAP002951
AGAP002952
AGAP002953
AGAP002954
AGAP002955
AGAP002956
AGAP002957
AGAP002958
mean_tpm
time
preZGA
ZGA_early
ZGA_late
5h
9h
11h
14h
20h
24h
L1
sex
Female
Male

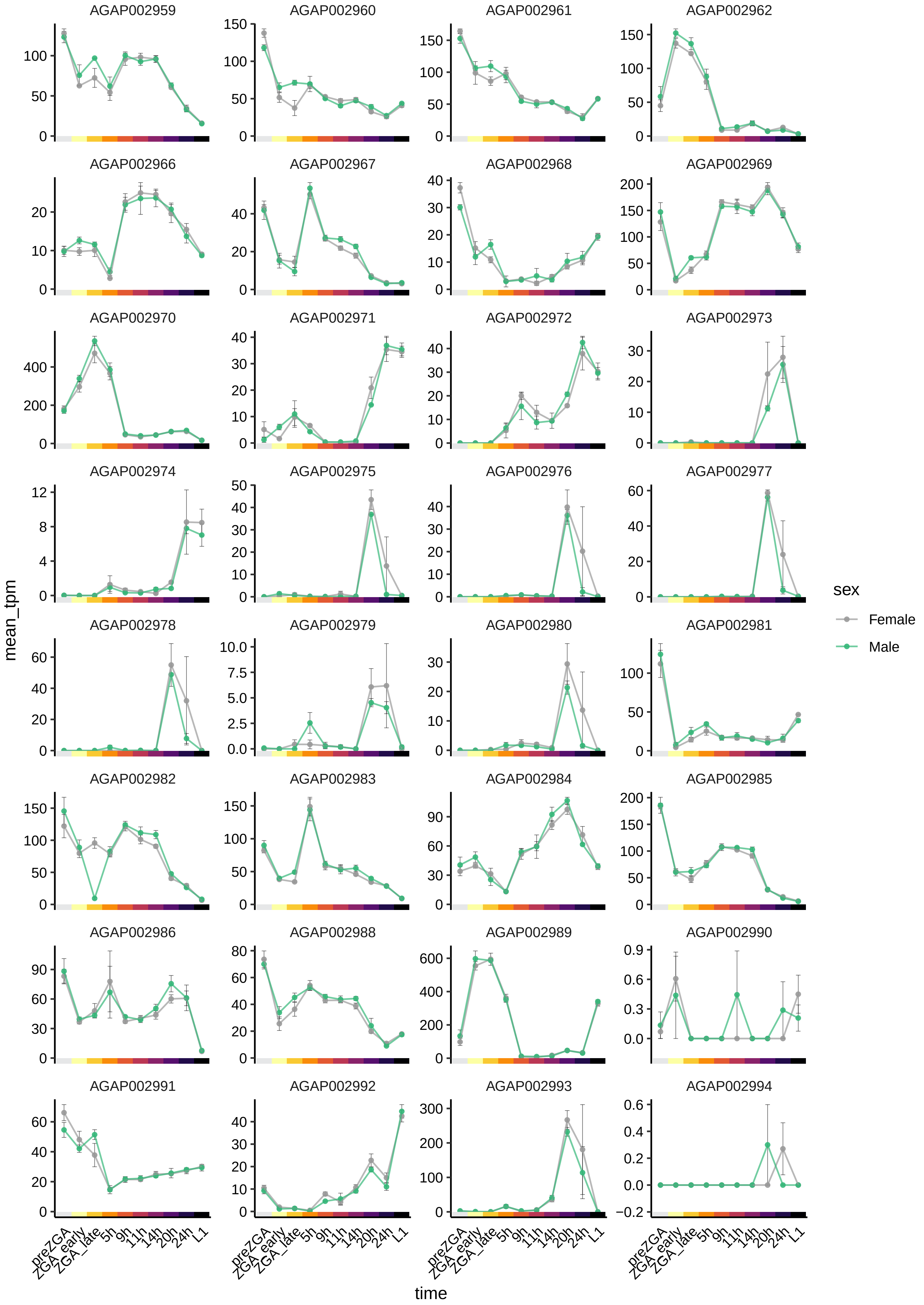

AGAP002959
AGAP002960
AGAP002961
AGAP002962
AGAP002966
AGAP002967
AGAP002968
AGAP002969
AGAP002970
AGAP002971
AGAP002972
AGAP002973
AGAP002974
AGAP002975
AGAP002976
AGAP002977
AGAP002978
AGAP002979
AGAP002980
AGAP002981
AGAP002982
AGAP002983
AGAP002984
AGAP002985
AGAP002986
AGAP002988
AGAP002989
AGAP002990
AGAP002991
AGAP002992
AGAP002993
AGAP002994
mean_tpm
time
preZGA
ZGA_early
ZGA_late
5h
9h
11h
14h
20h
24h
L1
sex
Female
Male

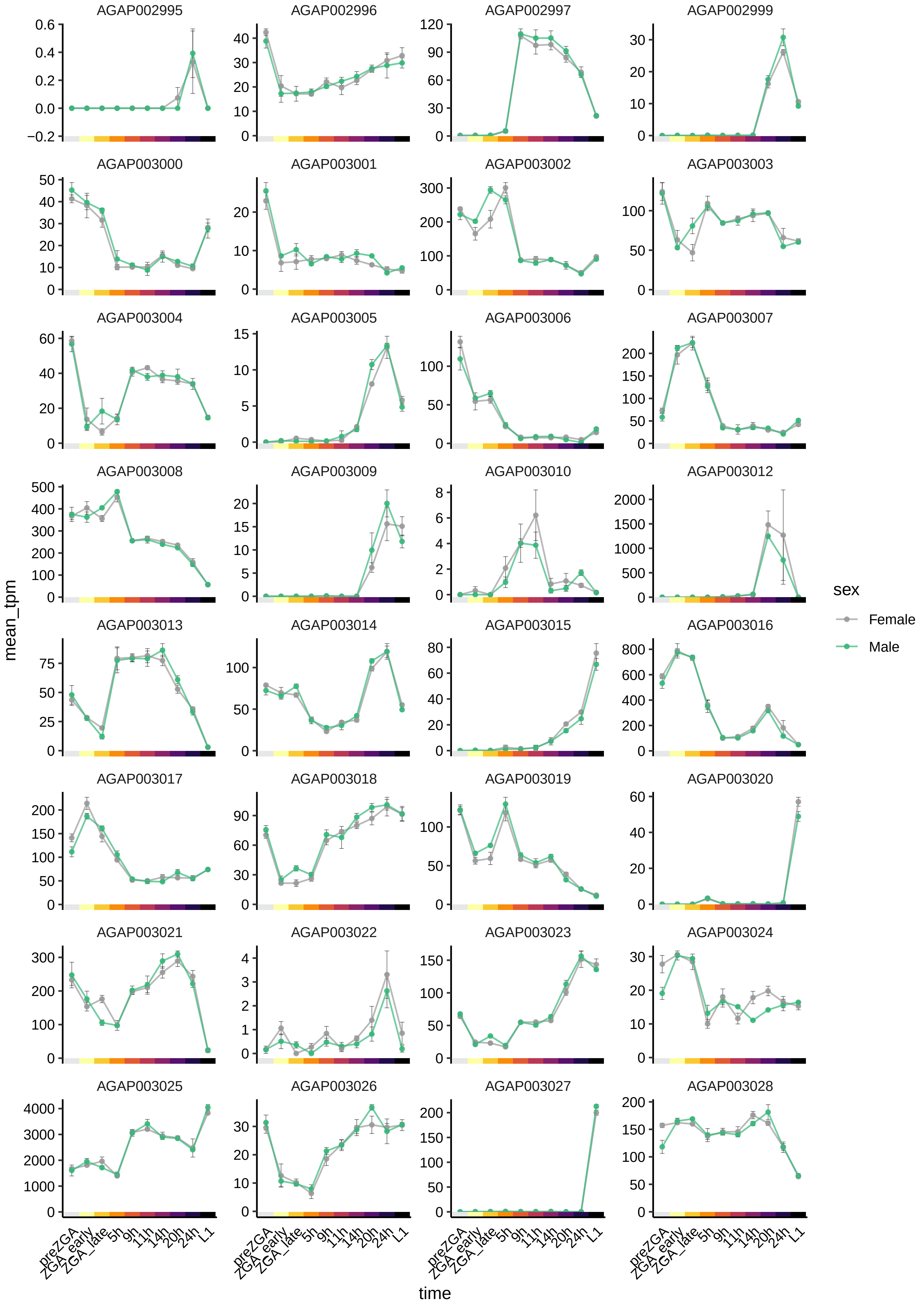
AGAP002995
AGAP002996
AGAP002997
AGAP002999
AGAP003000
AGAP003001
AGAP003002
AGAP003003
AGAP003004
AGAP003005
AGAP003006
AGAP003007
AGAP003008
AGAP003009
AGAP003010
AGAP003012
AGAP003013
AGAP003014
AGAP003015
AGAP003016
AGAP003017
AGAP003018
AGAP003019
AGAP003020
AGAP003021
AGAP003022
AGAP003023
AGAP003024
AGAP003025
AGAP003026
AGAP003027
AGAP003028
mean_tpm
time
preZGA
ZGA_early
ZGA_late
5h
9h
11h
14h
20h
24h
L1
sex
Female
Male

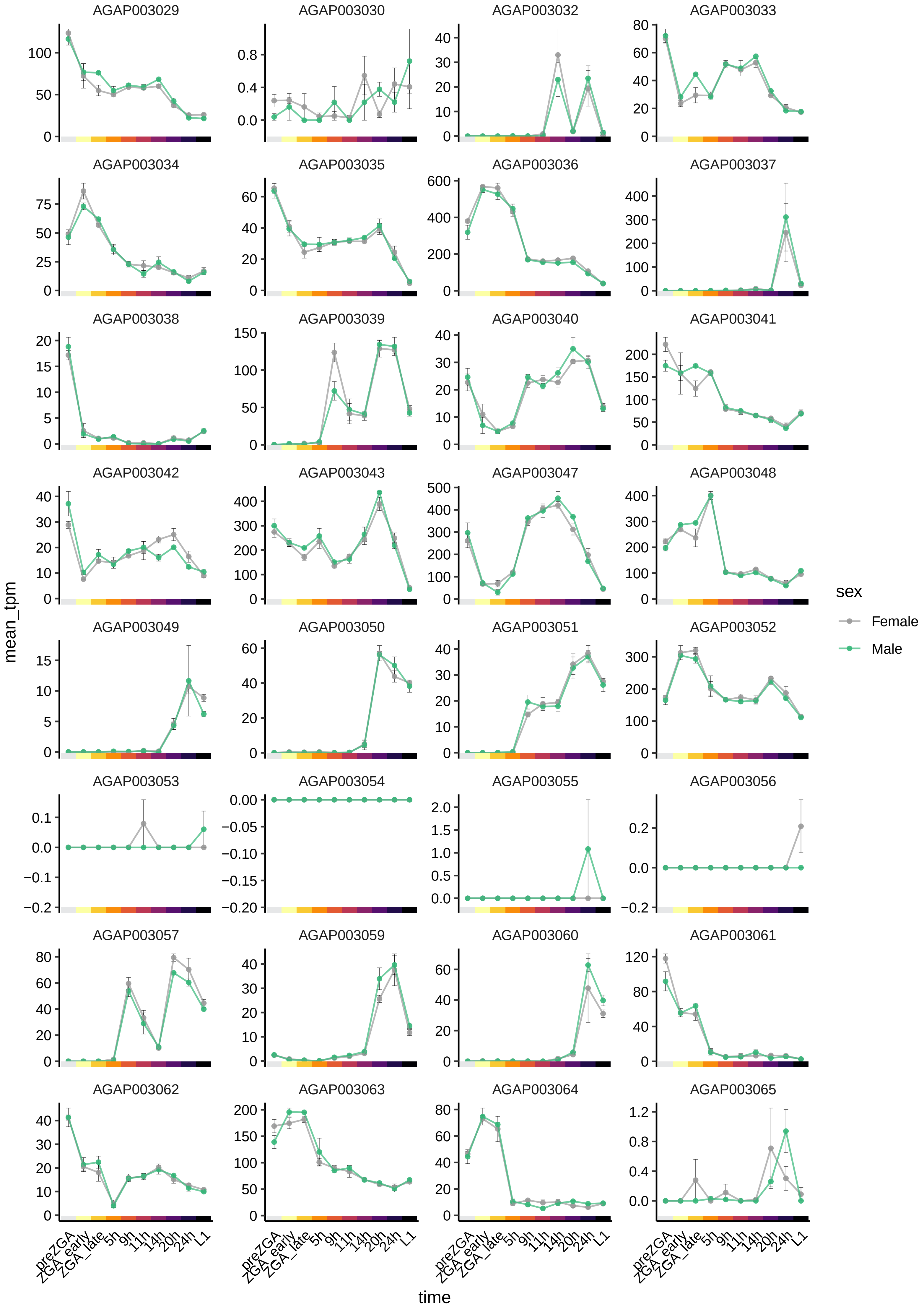

AGAP003029
AGAP003030
AGAP003032
AGAP003033
AGAP003034
AGAP003035
AGAP003036
AGAP003037
AGAP003038
AGAP003039
AGAP003040
AGAP003041
AGAP003042
AGAP003043
AGAP003047
AGAP003048
AGAP003049
AGAP003050
AGAP003051
AGAP003052
AGAP003053
AGAP003054
AGAP003055
AGAP003056
AGAP003057
AGAP003059
AGAP003060
AGAP003061
AGAP003062
AGAP003063
AGAP003064
AGAP003065
mean_tpm
time
preZGA
ZGA_early
ZGA_late
5h
9h
11h
14h
20h
24h
L1
sex
Female
Male

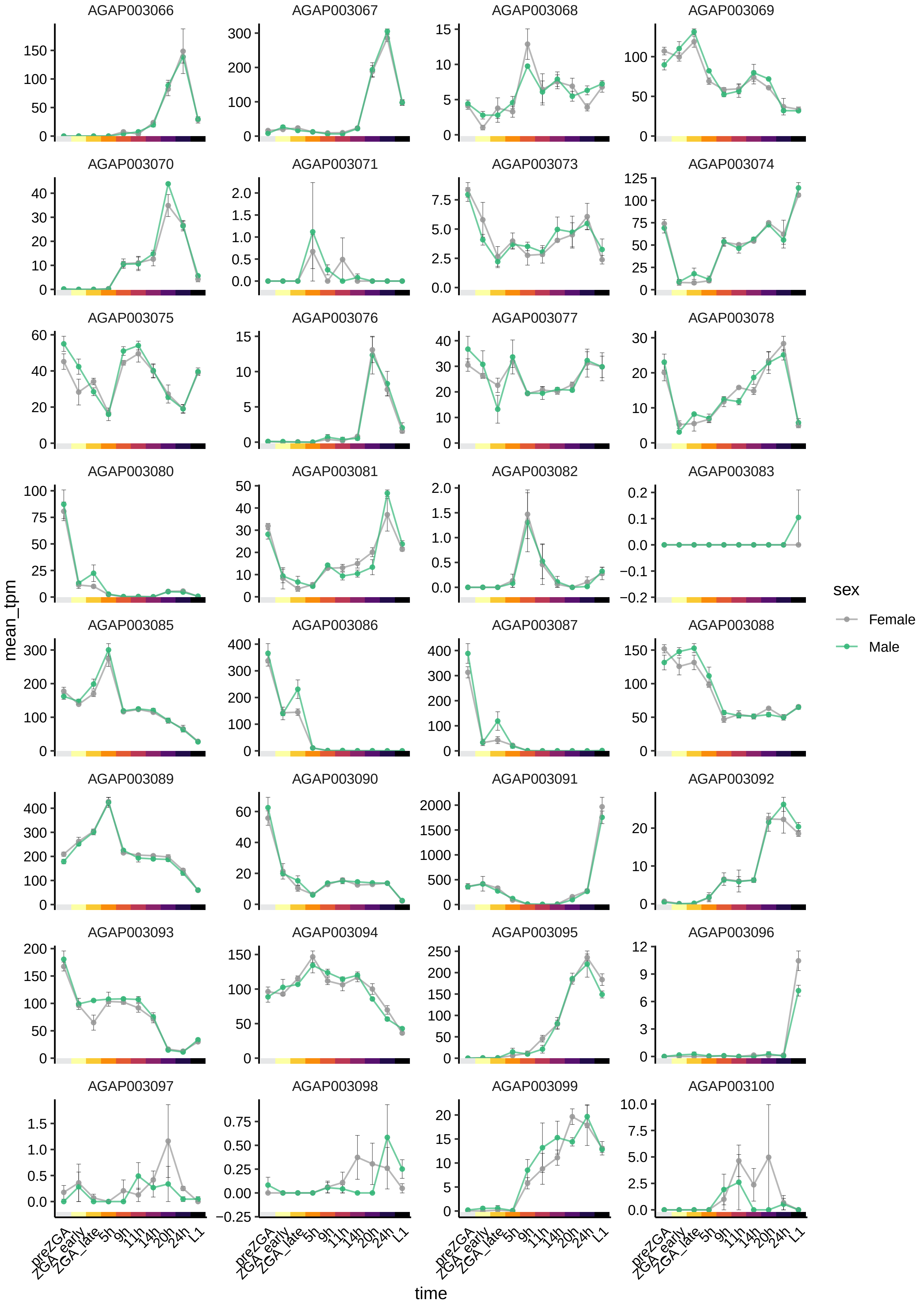
AGAP003066
AGAP003067
AGAP003068
AGAP003069
AGAP003070
AGAP003071
AGAP003073
AGAP003074
AGAP003075
AGAP003076
AGAP003077
AGAP003078
AGAP003080
AGAP003081
AGAP003082
AGAP003083
AGAP003085
AGAP003086
AGAP003087
AGAP003088
AGAP003089
AGAP003090
AGAP003091
AGAP003092
AGAP003093
AGAP003094
AGAP003095
AGAP003096
AGAP003097
AGAP003098
AGAP003099
AGAP003100
mean_tpm
time
preZGA
ZGA_early
ZGA_late
5h
9h
11h
14h
20h
24h
L1
sex
Female
Male

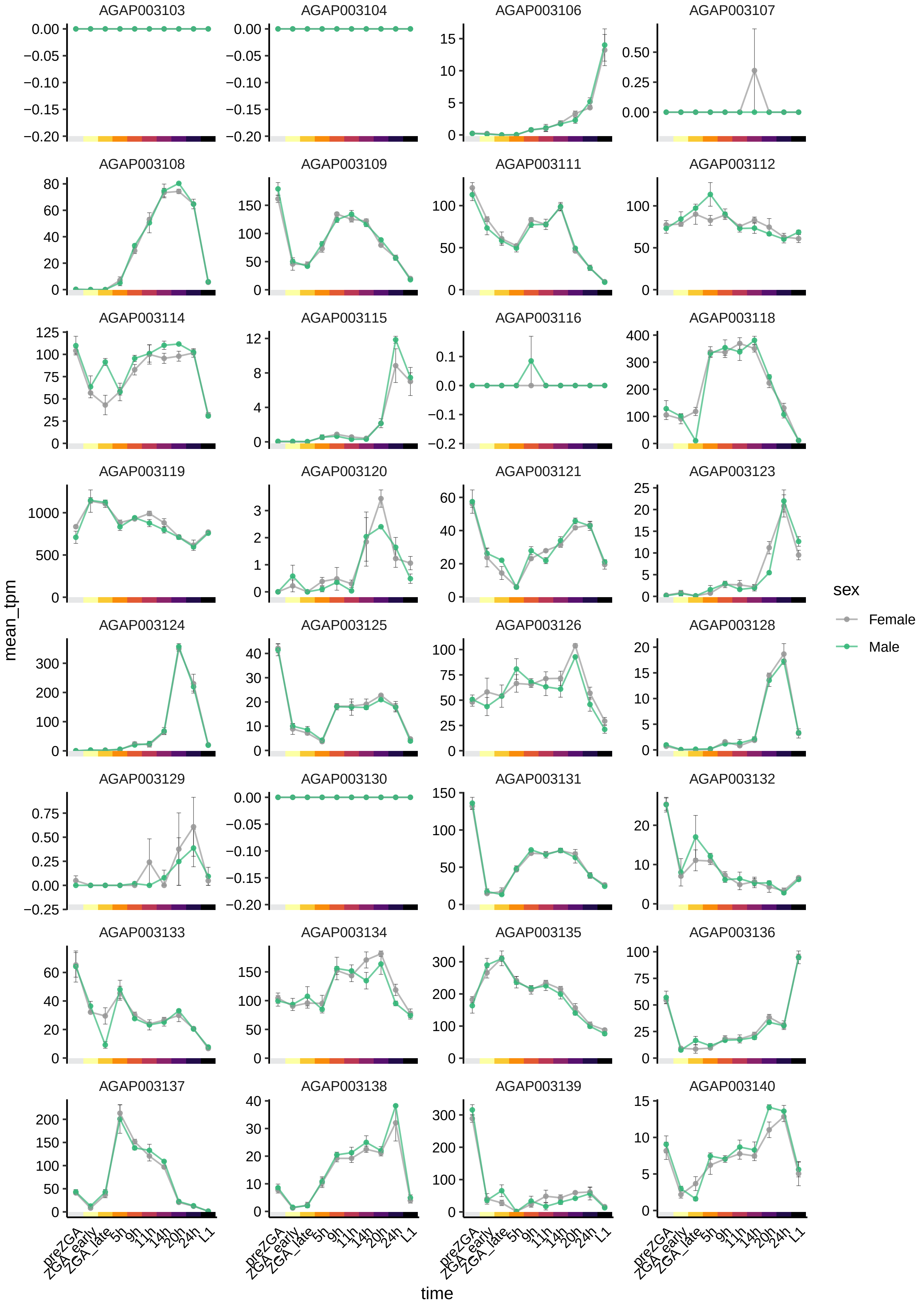
AGAP003103
AGAP003104
AGAP003106
AGAP003107
AGAP003108
AGAP003109
AGAP003111
AGAP003112
AGAP003114
AGAP003115
AGAP003116
AGAP003118
AGAP003119
AGAP003120
AGAP003121
AGAP003123
AGAP003124
AGAP003125
AGAP003126
AGAP003128
AGAP003129
AGAP003130
AGAP003131
AGAP003132
AGAP003133
AGAP003134
AGAP003135
AGAP003136
AGAP003137
AGAP003138
AGAP003139
AGAP003140
mean_tpm
time
preZGA
ZGA_early
ZGA_late
5h
9h
11h
14h
20h
24h
L1
sex
Female
Male

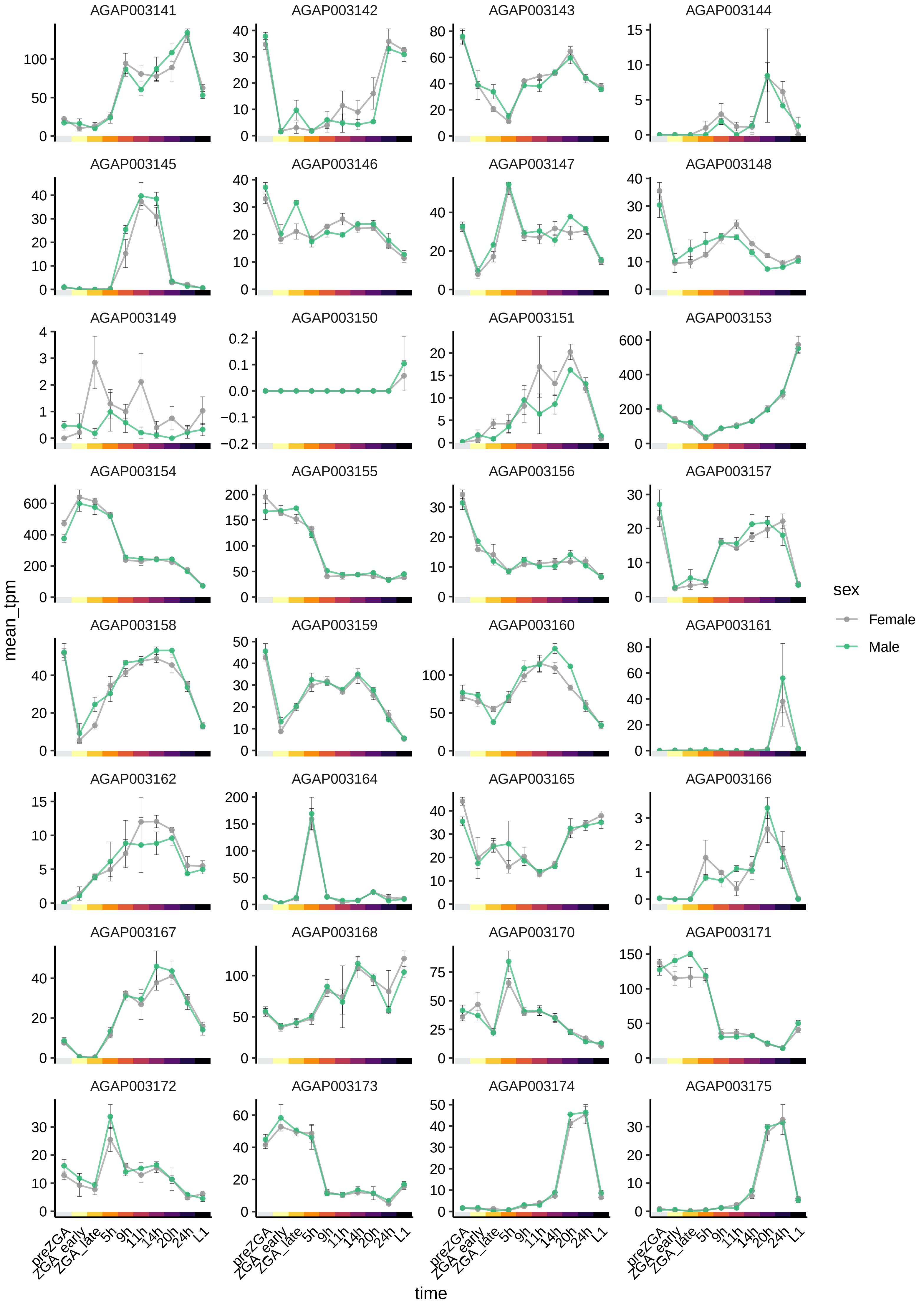

AGAP003141
AGAP003142
AGAP003143
AGAP003144
AGAP003145
AGAP003146
AGAP003147
AGAP003148
AGAP003149
AGAP003150
AGAP003151
AGAP003153
AGAP003154
AGAP003155
AGAP003156
AGAP003157
AGAP003158
AGAP003159
AGAP003160
AGAP003161
AGAP003162
AGAP003164
AGAP003165
AGAP003166
AGAP003167
AGAP003168
AGAP003170
AGAP003171
AGAP003172
AGAP003173
AGAP003174
AGAP003175
mean_tpm
time
preZGA
ZGA_early
ZGA_late
5h
9h
11h
14h
20h
24h
L1
sex
Female
Male

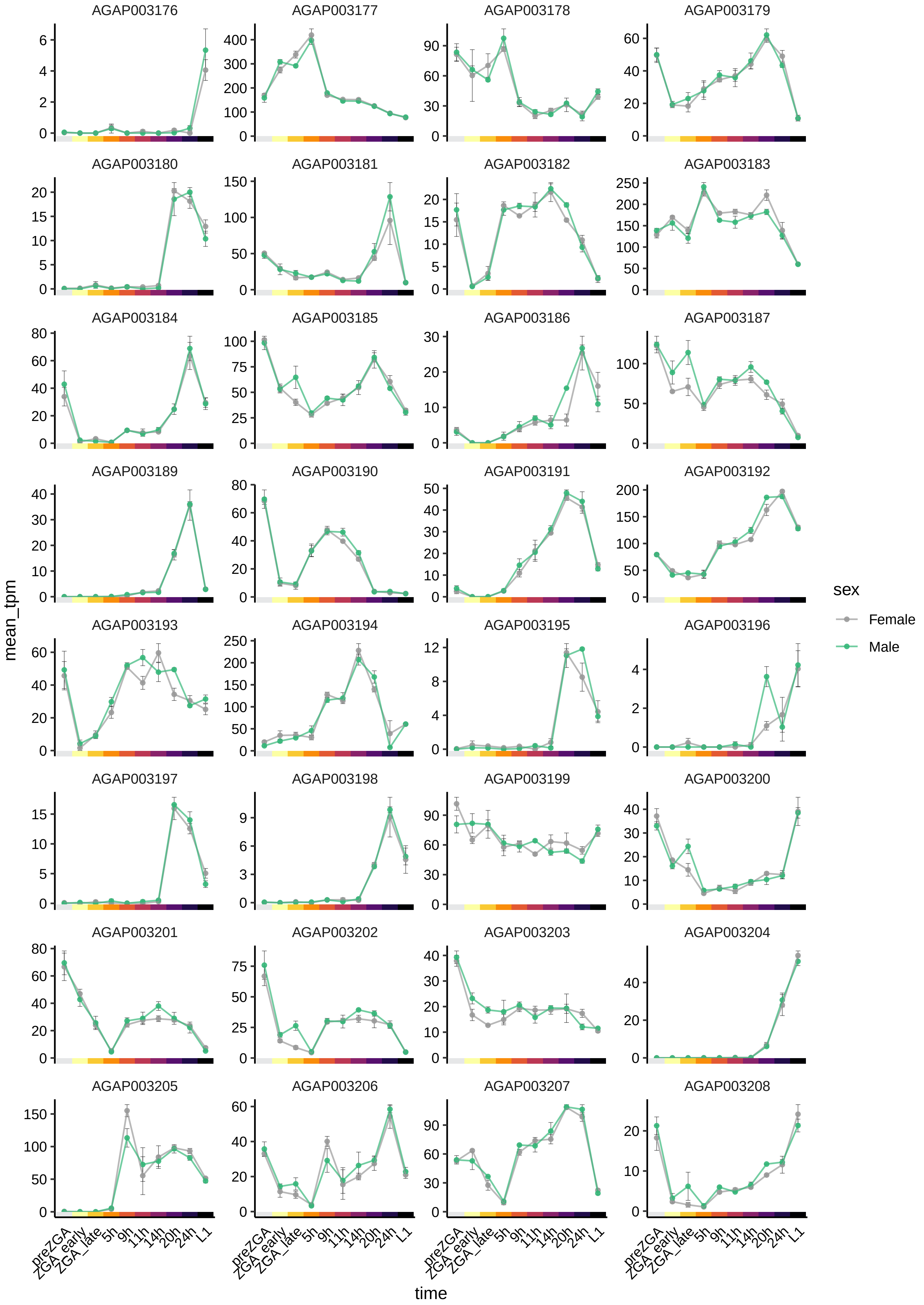

AGAP003176
AGAP003177
AGAP003178
AGAP003179
AGAP003180
AGAP003181
AGAP003182
AGAP003183
AGAP003184
AGAP003185
AGAP003186
AGAP003187
AGAP003189
AGAP003190
AGAP003191
AGAP003192
AGAP003193
AGAP003194
AGAP003195
AGAP003196
AGAP003197
AGAP003198
AGAP003199
AGAP003200
AGAP003201
AGAP003202
AGAP003203
AGAP003204
AGAP003205
AGAP003206
AGAP003207
AGAP003208
mean_tpm
time
preZGA
ZGA_early
ZGA_late
5h
9h
11h
14h
20h
24h
L1
sex
Female
Male

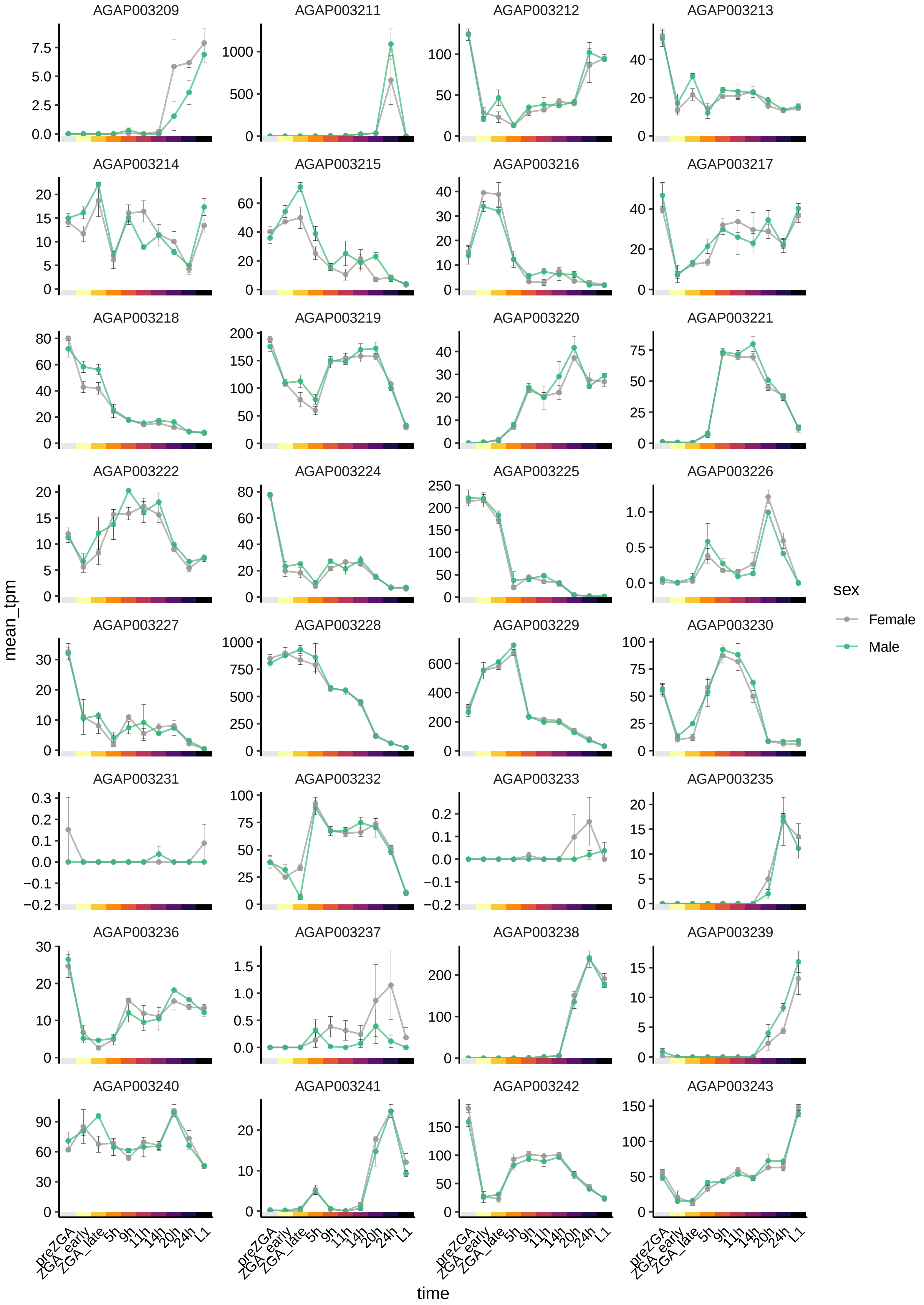

AGAP003209
AGAP003211
AGAP003212
AGAP003213
AGAP003214
AGAP003215
AGAP003216
AGAP003217
AGAP003218
AGAP003219
AGAP003220
AGAP003221
AGAP003222
AGAP003224
AGAP003225
AGAP003226
AGAP003227
AGAP003228
AGAP003229
AGAP003230
AGAP003231
AGAP003232
AGAP003233
AGAP003235
AGAP003236
AGAP003237
AGAP003238
AGAP003239
AGAP003240
AGAP003241
AGAP003242
AGAP003243
mean_tpm
time
preZGA
ZGA_early
ZGA_late
5h
9h
11h
14h
20h
24h
L1
sex
Female
Male

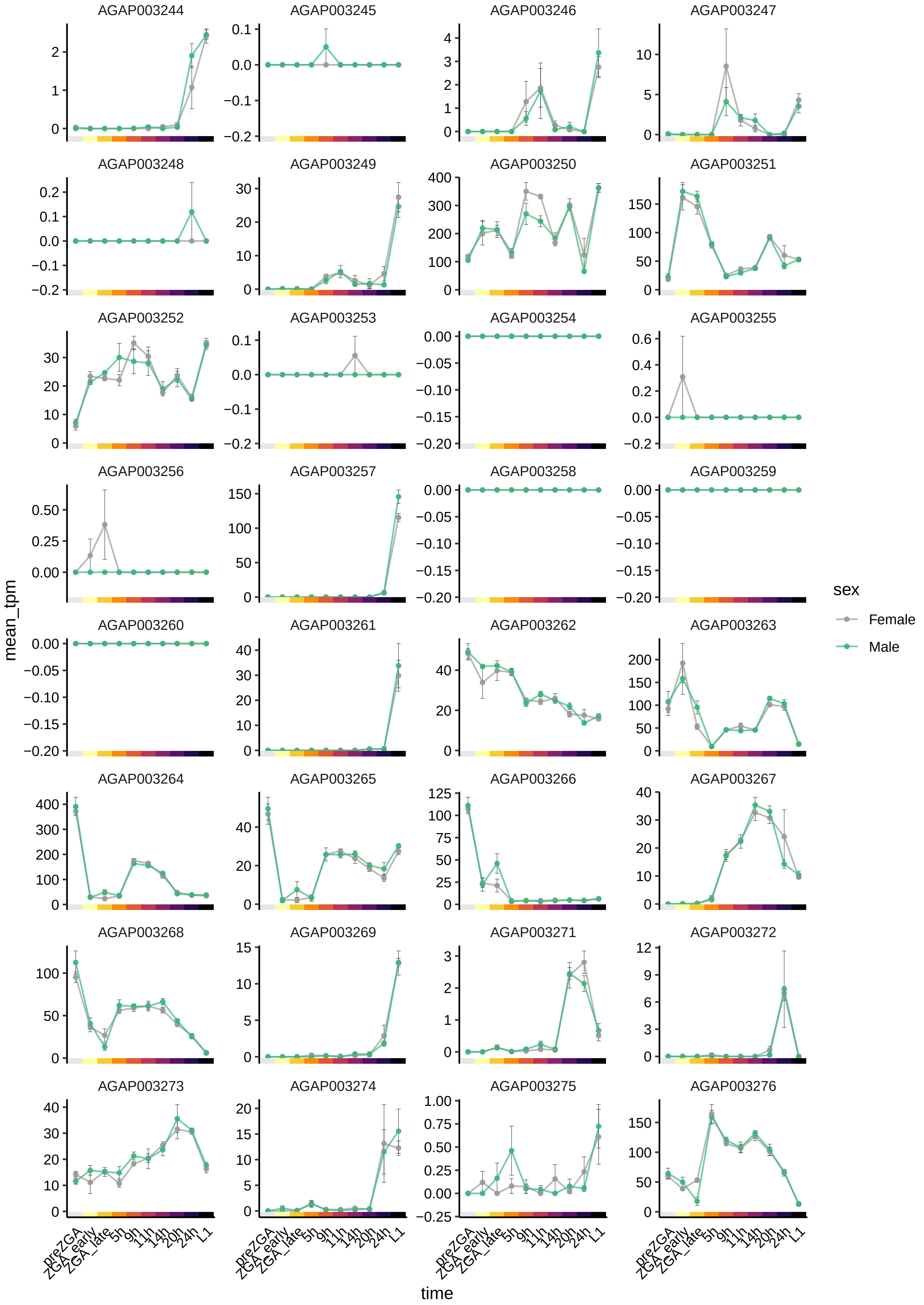
AGAP003244
AGAP003245
AGAP003246
AGAP003247
AGAP003248
AGAP003249
AGAP003250
AGAP003251
AGAP003252
AGAP003253
AGAP003254
AGAP003255
AGAP003256
AGAP003257
AGAP003258
AGAP003259
AGAP003260
AGAP003261
AGAP003262
AGAP003263
AGAP003264
AGAP003265
AGAP003266
AGAP003267
AGAP003268
AGAP003269
AGAP003271
AGAP003272
AGAP003273
AGAP003274
AGAP003275
AGAP003276
mean_tpm
time
preZGA
ZGA_early
ZGA_late
5h
9h
11h
14h
20h
24h
L1
sex
Female
Male

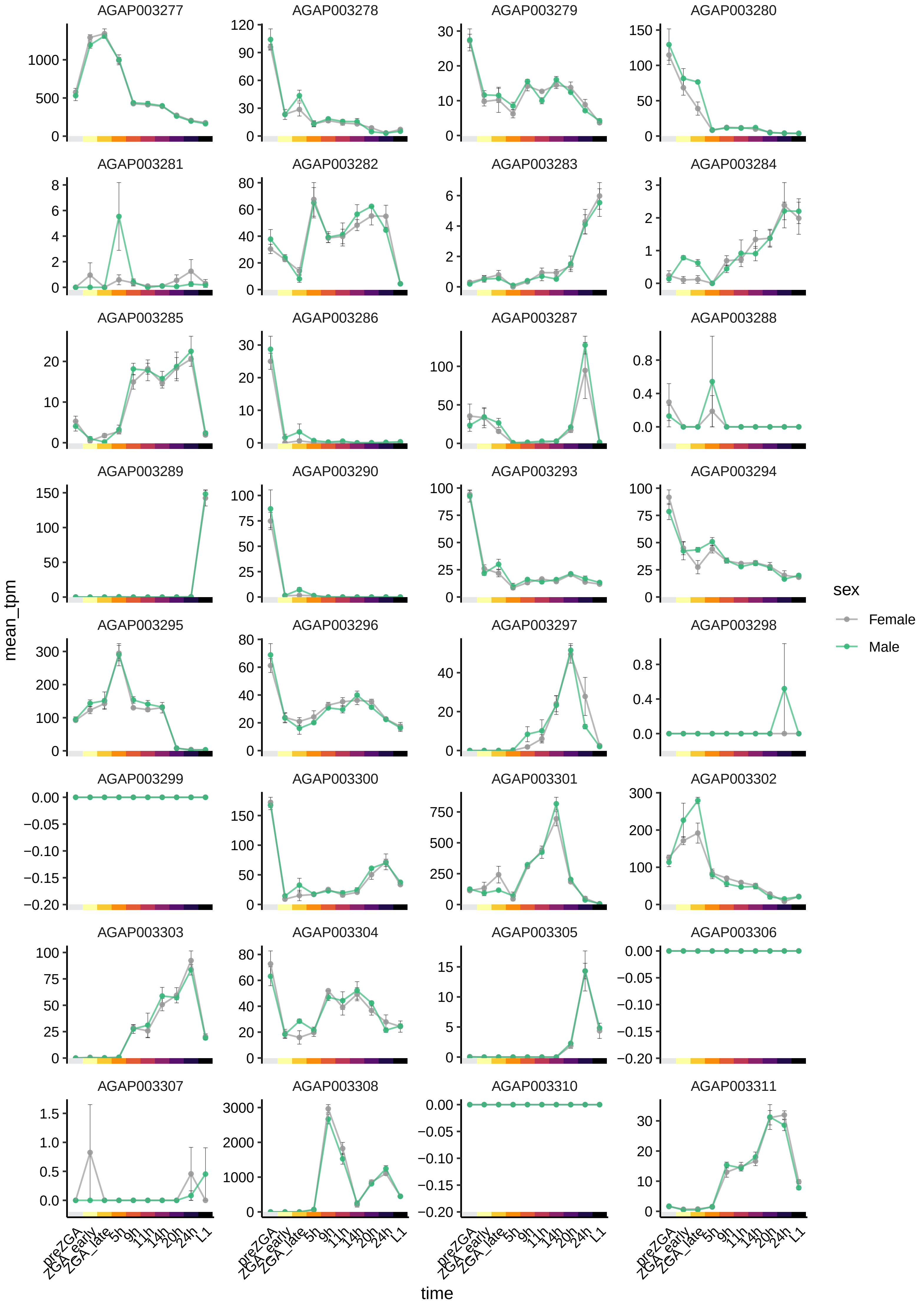

AGAP003277
AGAP003278
AGAP003279
AGAP003280
AGAP003281
AGAP003282
AGAP003283
AGAP003284
AGAP003285
AGAP003286
AGAP003287
AGAP003288
AGAP003289
AGAP003290
AGAP003293
AGAP003294
AGAP003295
AGAP003296
AGAP003297
AGAP003298
AGAP003299
AGAP003300
AGAP003301
AGAP003302
AGAP003303
AGAP003304
AGAP003305
AGAP003306
AGAP003307
AGAP003308
AGAP003310
AGAP003311
mean_tpm
time
preZGA
ZGA_early
ZGA_late
5h
9h
11h
14h
20h
24h
L1
sex
Female
Male

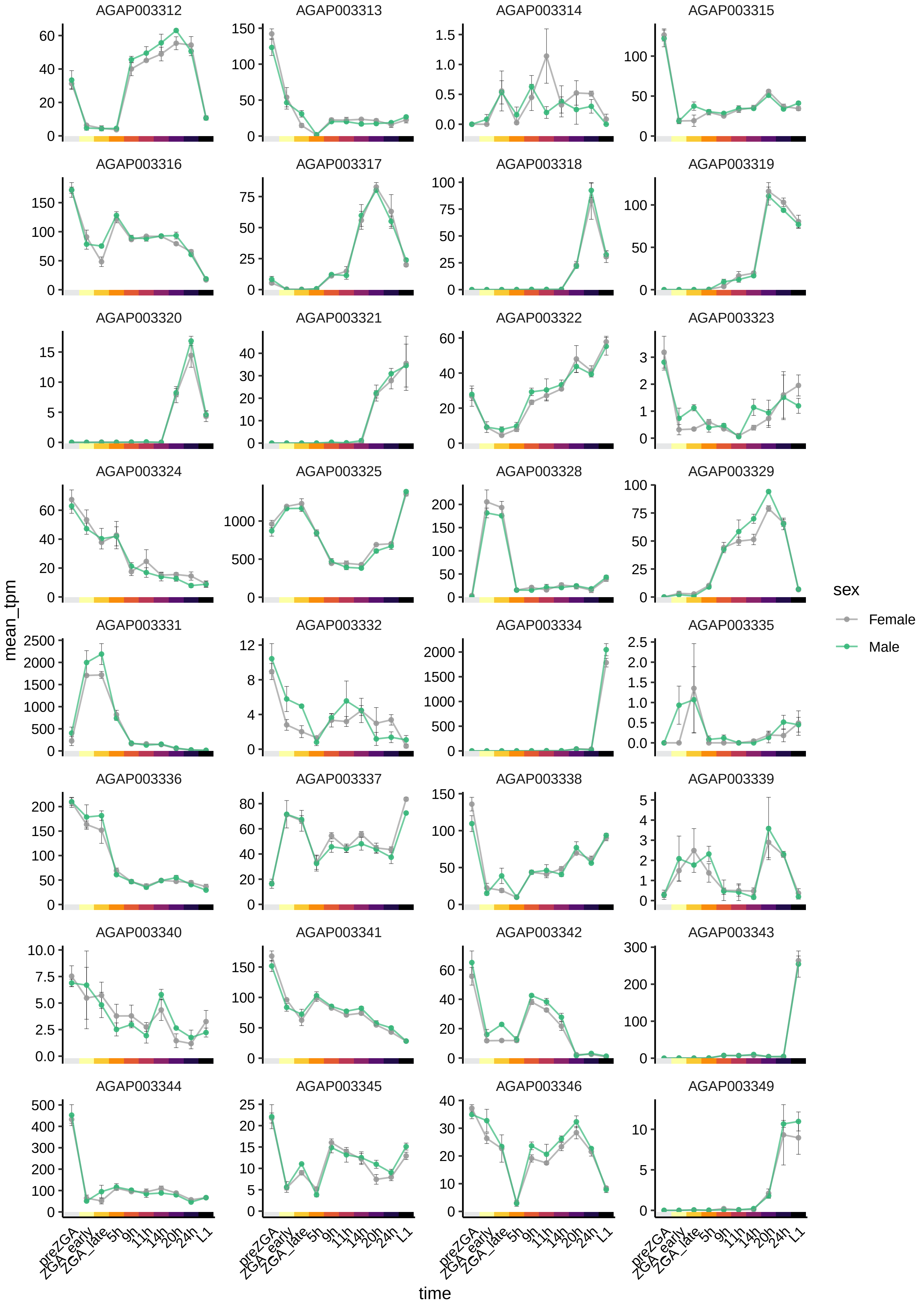
AGAP003312
AGAP003313
AGAP003314
AGAP003315
AGAP003316
AGAP003317
AGAP003318
AGAP003319
AGAP003320
AGAP003321
AGAP003322
AGAP003323
AGAP003324
AGAP003325
AGAP003328
AGAP003329
AGAP003331
AGAP003332
AGAP003334
AGAP003335
AGAP003336
AGAP003337
AGAP003338
AGAP003339
AGAP003340
AGAP003341
AGAP003342
AGAP003343
AGAP003344
AGAP003345
AGAP003346
AGAP003349
mean_tpm
time
preZGA
ZGA_early
ZGA_late
5h
9h
11h
14h
20h
24h
L1
sex
Female
Male

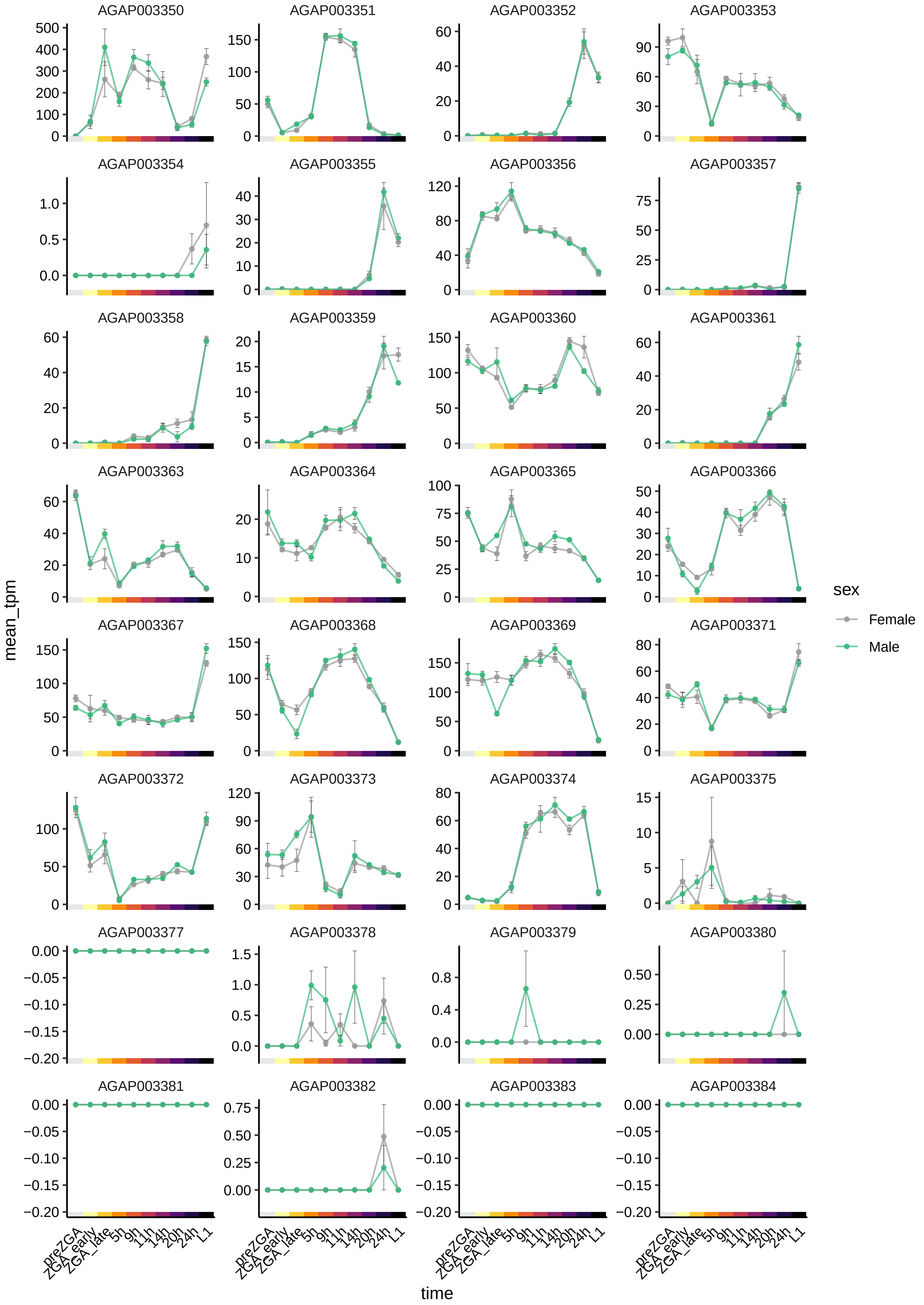
AGAP003350
AGAP003351
AGAP003352
AGAP003353
AGAP003354
AGAP003355
AGAP003356
AGAP003357
AGAP003358
AGAP003359
AGAP003360
AGAP003361
AGAP003363
AGAP003364
AGAP003365
AGAP003366
AGAP003367
AGAP003368
AGAP003369
AGAP003371
AGAP003372
AGAP003373
AGAP003374
AGAP003375
AGAP003377
AGAP003378
AGAP003379
AGAP003380
AGAP003381
AGAP003382
AGAP003383
AGAP003384
mean_tpm
time
preZGA
ZGA_early
ZGA_late
5h
9h
11h
14h
20h
24h
L1
sex
Female
Male

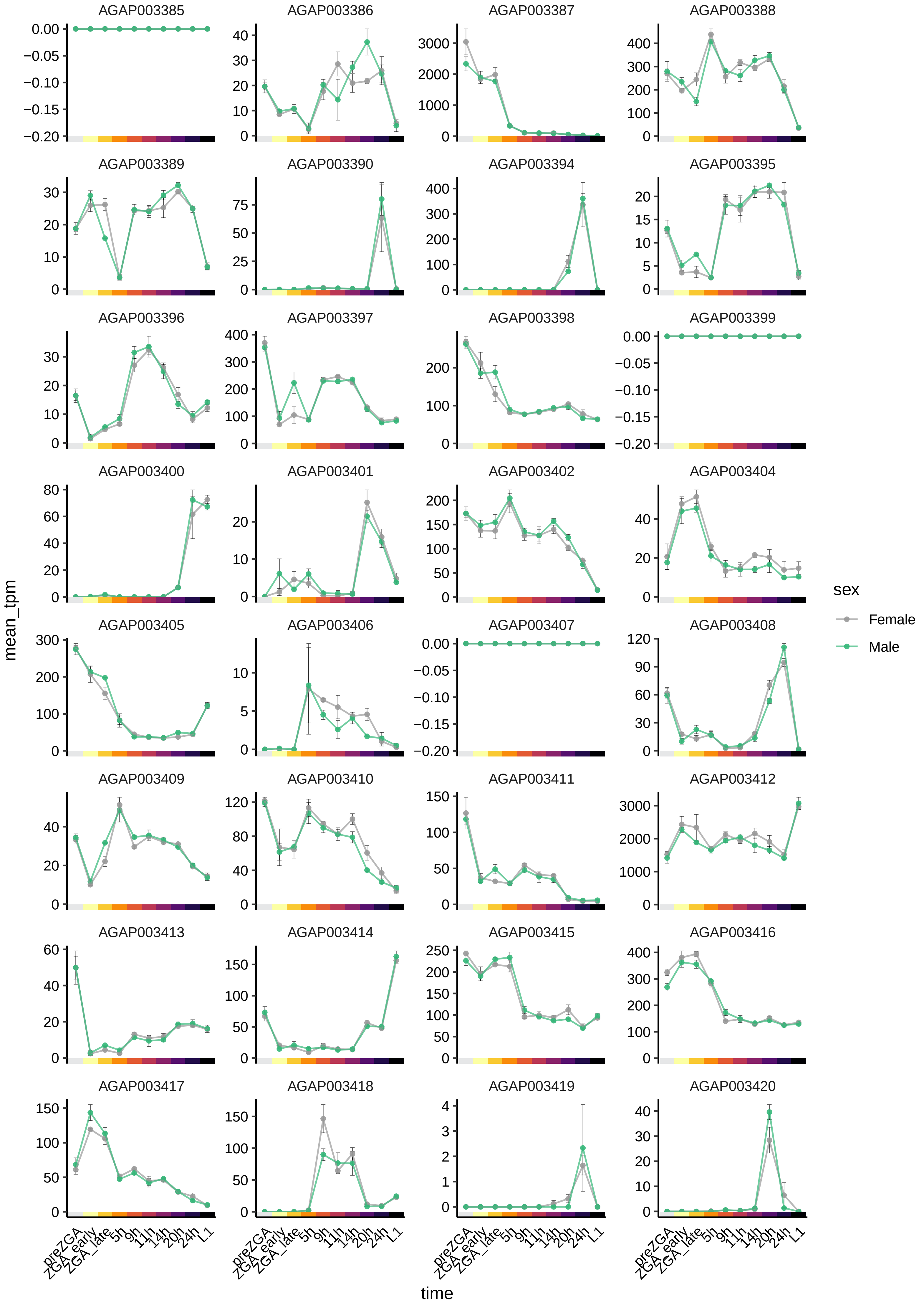

AGAP003385
AGAP003386
AGAP003387
AGAP003388
AGAP003389
AGAP003390
AGAP003394
AGAP003395
AGAP003396
AGAP003397
AGAP003398
AGAP003399
AGAP003400
AGAP003401
AGAP003402
AGAP003404
AGAP003405
AGAP003406
AGAP003407
AGAP003408
AGAP003409
AGAP003410
AGAP003411
AGAP003412
AGAP003413
AGAP003414
AGAP003415
AGAP003416
AGAP003417
AGAP003418
AGAP003419
AGAP003420
mean_tpm
time
preZGA
ZGA_early
ZGA_late
5h
9h
11h
14h
20h
24h
L1
sex
Female
Male

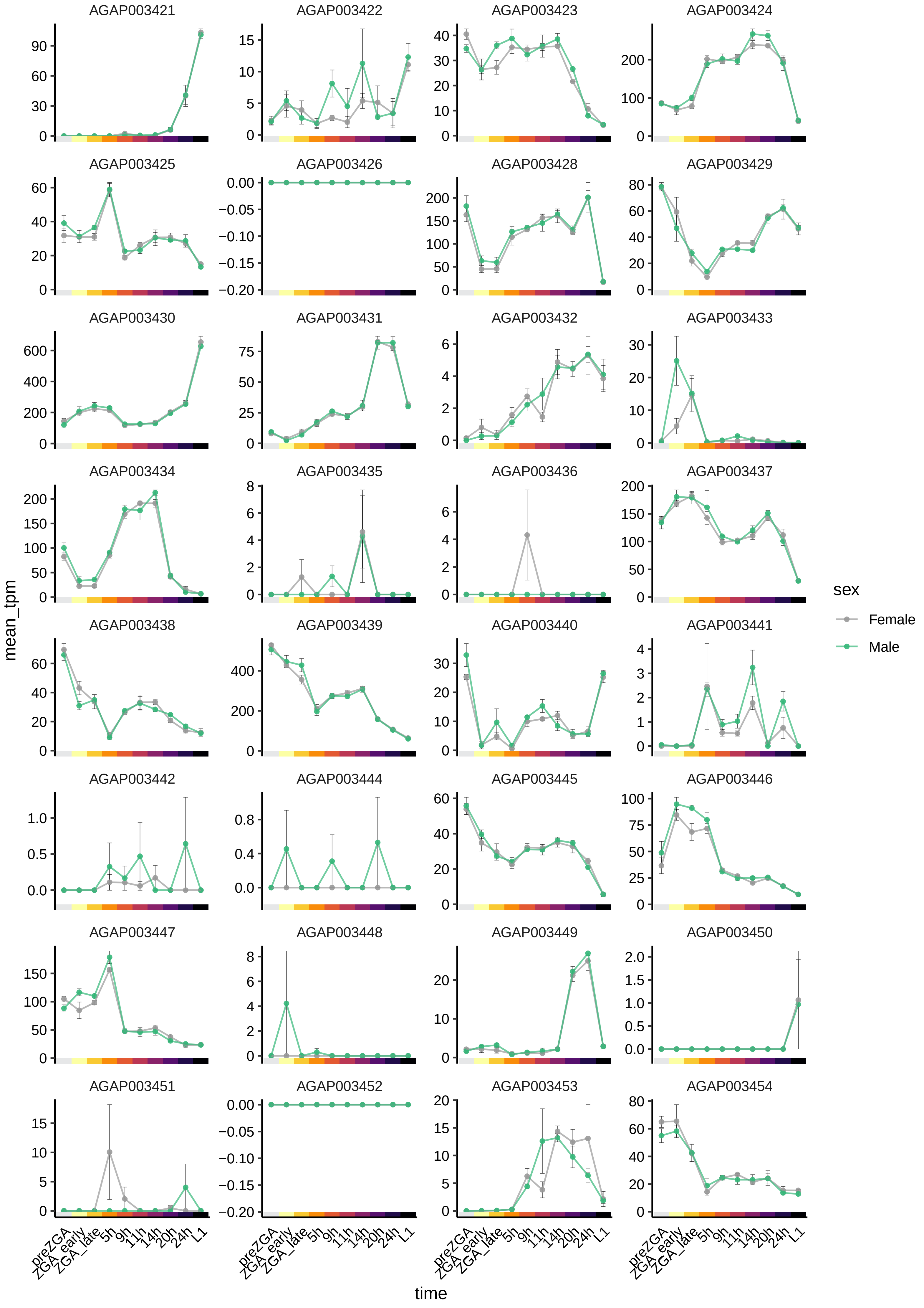
AGAP003421
AGAP003422
AGAP003423
AGAP003424
AGAP003425
AGAP003426
AGAP003428
AGAP003429
AGAP003430
AGAP003431
AGAP003432
AGAP003433
AGAP003434
AGAP003435
AGAP003436
AGAP003437
AGAP003438
AGAP003439
AGAP003440
AGAP003441
AGAP003442
AGAP003444
AGAP003445
AGAP003446
AGAP003447
AGAP003448
AGAP003449
AGAP003450
AGAP003451
AGAP003452
AGAP003453
AGAP003454
mean_tpm
time
preZGA
ZGA_early
ZGA_late
5h
9h
11h
14h
20h
24h
L1
sex
Female
Male

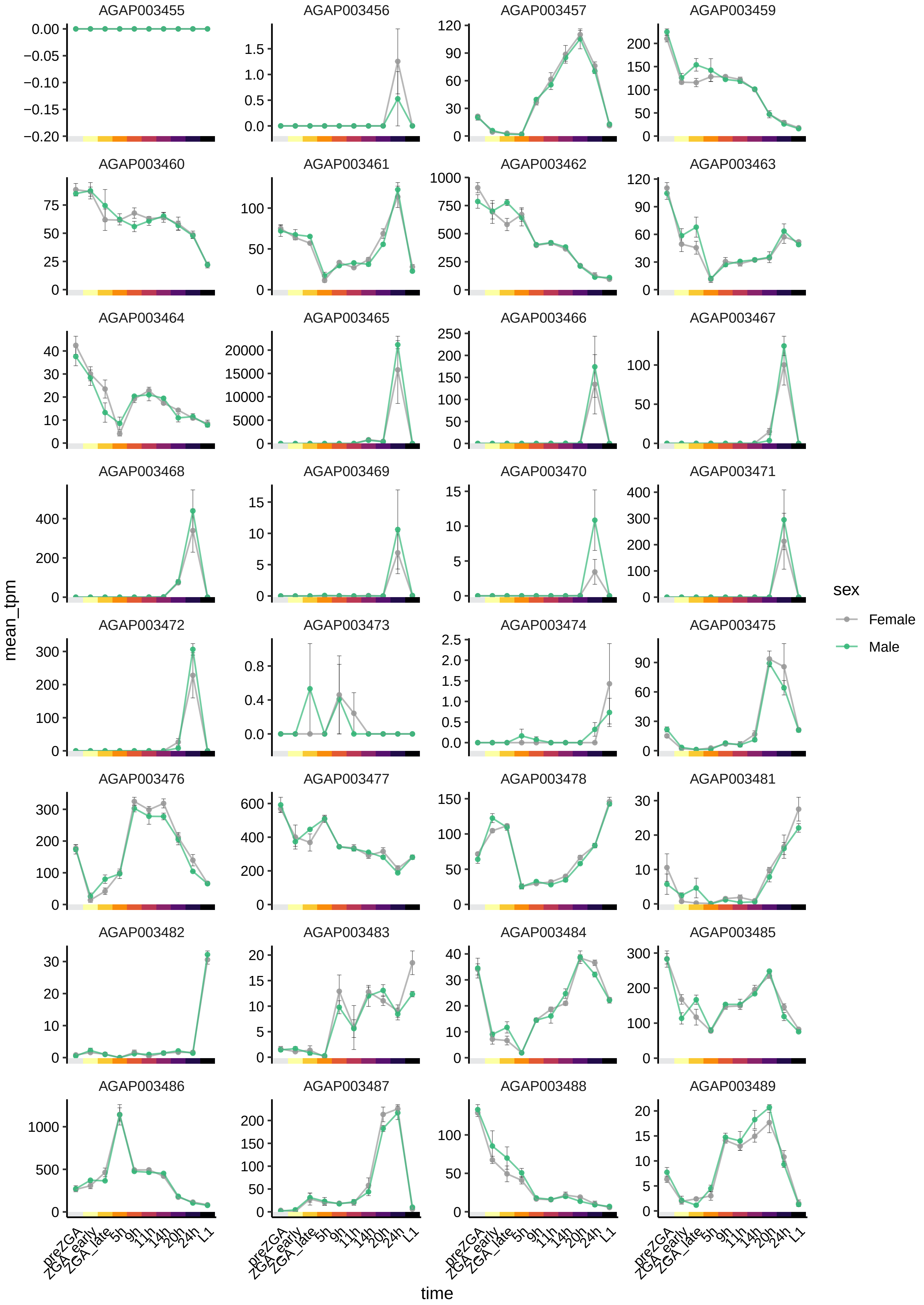

AGAP003455
AGAP003456
AGAP003457
AGAP003459
AGAP003460
AGAP003461
AGAP003462
AGAP003463
AGAP003464
AGAP003465
AGAP003466
AGAP003467
AGAP003468
AGAP003469
AGAP003470
AGAP003471
AGAP003472
AGAP003473
AGAP003474
AGAP003475
AGAP003476
AGAP003477
AGAP003478
AGAP003481
AGAP003482
AGAP003483
AGAP003484
AGAP003485
AGAP003486
AGAP003487
AGAP003488
AGAP003489
mean_tpm
time
preZGA
ZGA_early
ZGA_late
5h
9h
11h
14h
20h
24h
L1
sex
Female
Male

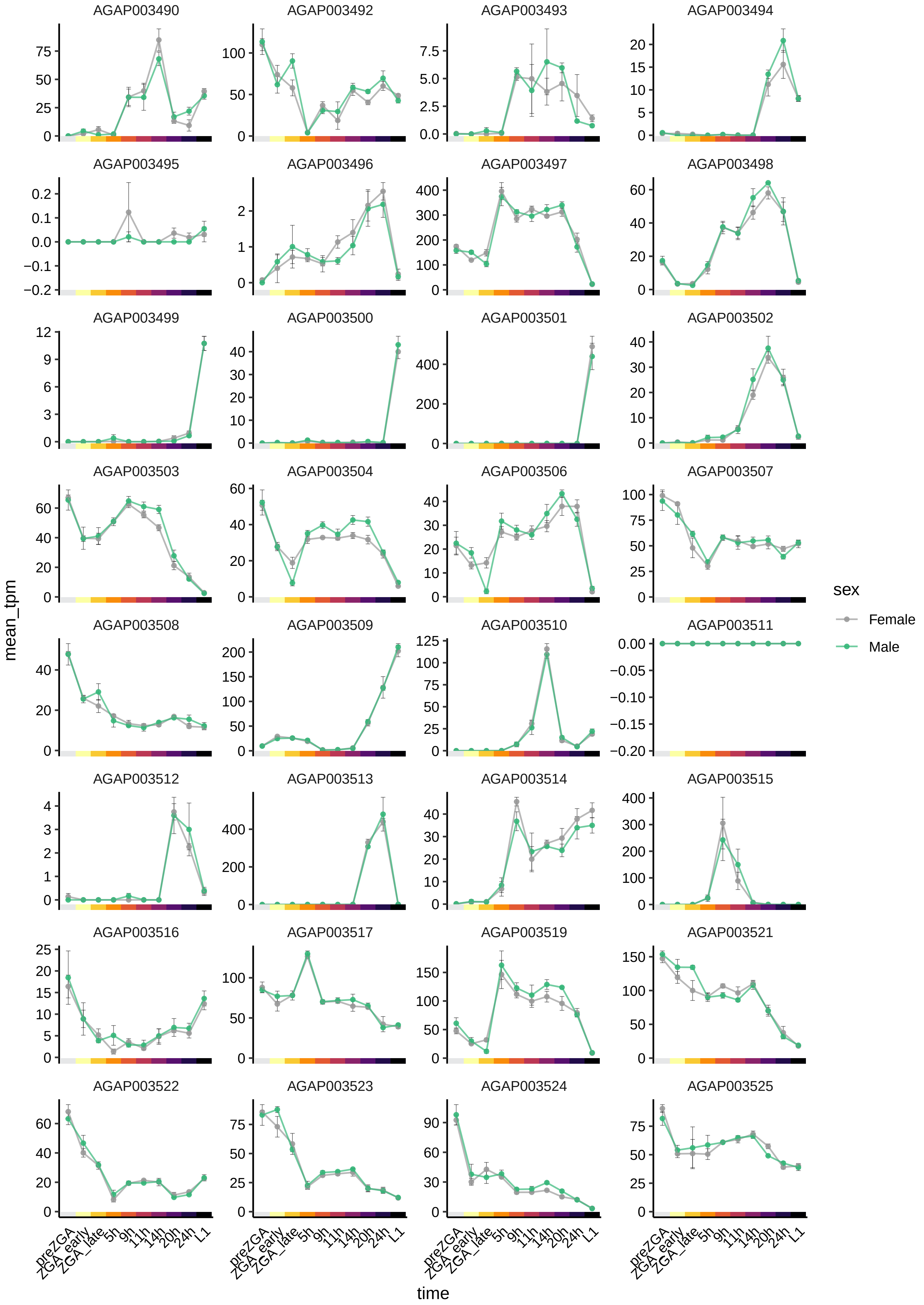

AGAP003490
AGAP003492
AGAP003493
AGAP003494
AGAP003495
AGAP003496
AGAP003497
AGAP003498
AGAP003499
AGAP003500
AGAP003501
AGAP003502
AGAP003503
AGAP003504
AGAP003506
AGAP003507
AGAP003508
AGAP003509
AGAP003510
AGAP003511
AGAP003512
AGAP003513
AGAP003514
AGAP003515
AGAP003516
AGAP003517
AGAP003519
AGAP003521
AGAP003522
AGAP003523
AGAP003524
AGAP003525
mean_tpm
time
preZGA
ZGA_early
ZGA_late
5h
9h
11h
14h
20h
24h
L1
sex
Female
Male

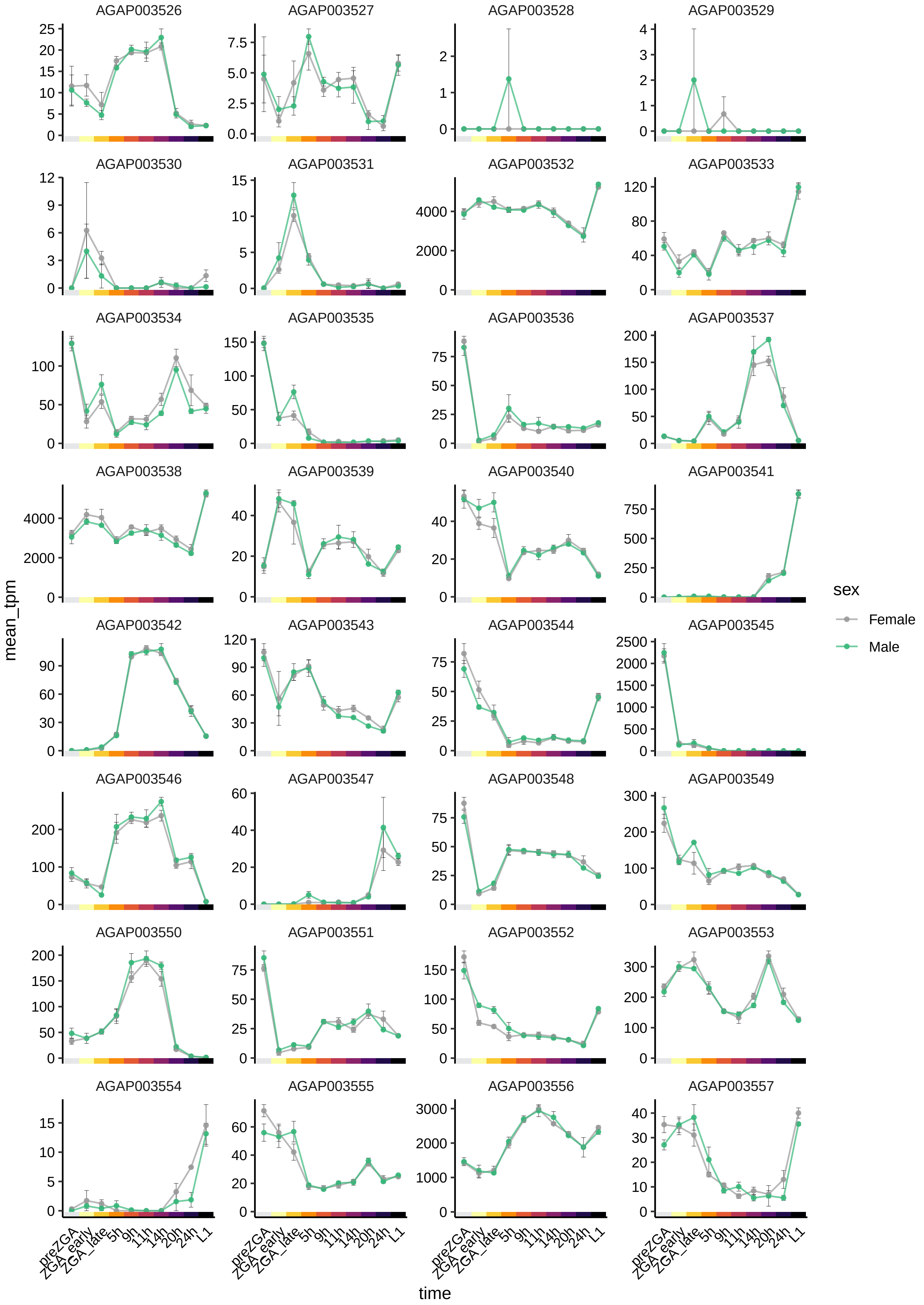
AGAP003526
AGAP003527
AGAP003528
AGAP003529
AGAP003530
AGAP003531
AGAP003532
AGAP003533
AGAP003534
AGAP003535
AGAP003536
AGAP003537
AGAP003538
AGAP003539
AGAP003540
AGAP003541
AGAP003542
AGAP003543
AGAP003544
AGAP003545
AGAP003546
AGAP003547
AGAP003548
AGAP003549
AGAP003550
AGAP003551
AGAP003552
AGAP003553
AGAP003554
AGAP003555
AGAP003556
AGAP003557
mean_tpm
time
preZGA
ZGA_early
ZGA_late
5h
9h
11h
14h
20h
24h
L1
sex
Female
Male

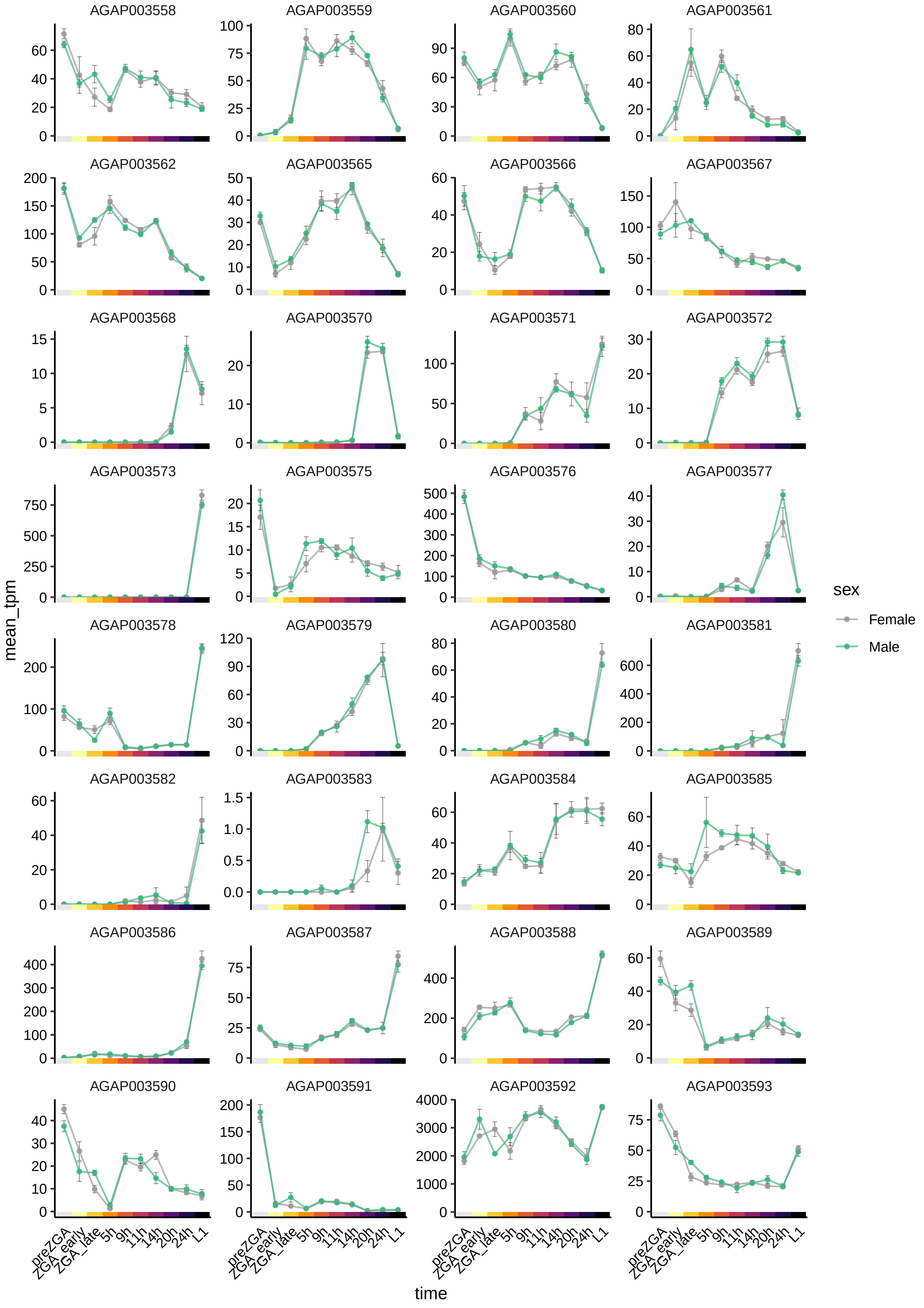

AGAP003558
AGAP003559
AGAP003560
AGAP003561
AGAP003562
AGAP003565
AGAP003566
AGAP003567
AGAP003568
AGAP003570
AGAP003571
AGAP003572
AGAP003573
AGAP003575
AGAP003576
AGAP003577
AGAP003578
AGAP003579
AGAP003580
AGAP003581
AGAP003582
AGAP003583
AGAP003584
AGAP003585
AGAP003586
AGAP003587
AGAP003588
AGAP003589
AGAP003590
AGAP003591
AGAP003592
AGAP003593
mean_tpm
time
preZGA
ZGA_early
ZGA_late
5h
9h
11h
14h
20h
24h
L1
sex
Female
Male

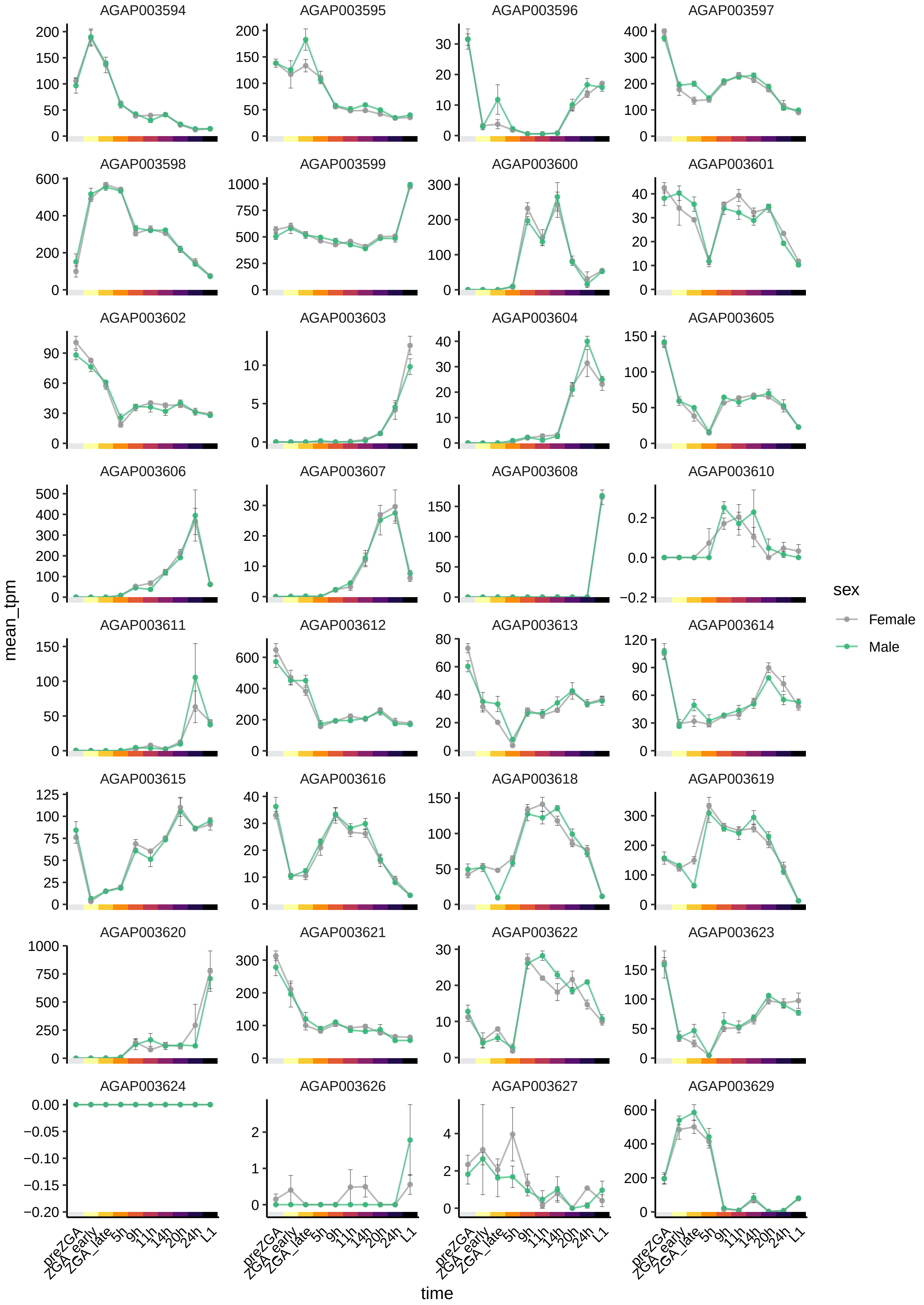

AGAP003594
AGAP003595
AGAP003596
AGAP003597
AGAP003598
AGAP003599
AGAP003600
AGAP003601
AGAP003602
AGAP003603
AGAP003604
AGAP003605
AGAP003606
AGAP003607
AGAP003608
AGAP003610
AGAP003611
AGAP003612
AGAP003613
AGAP003614
AGAP003615
AGAP003616
AGAP003618
AGAP003619
AGAP003620
AGAP003621
AGAP003622
AGAP003623
AGAP003624
AGAP003626
AGAP003627
AGAP003629
mean_tpm
time
preZGA
ZGA_early
ZGA_late
5h
9h
11h
14h
20h
24h
L1
sex
Female
Male

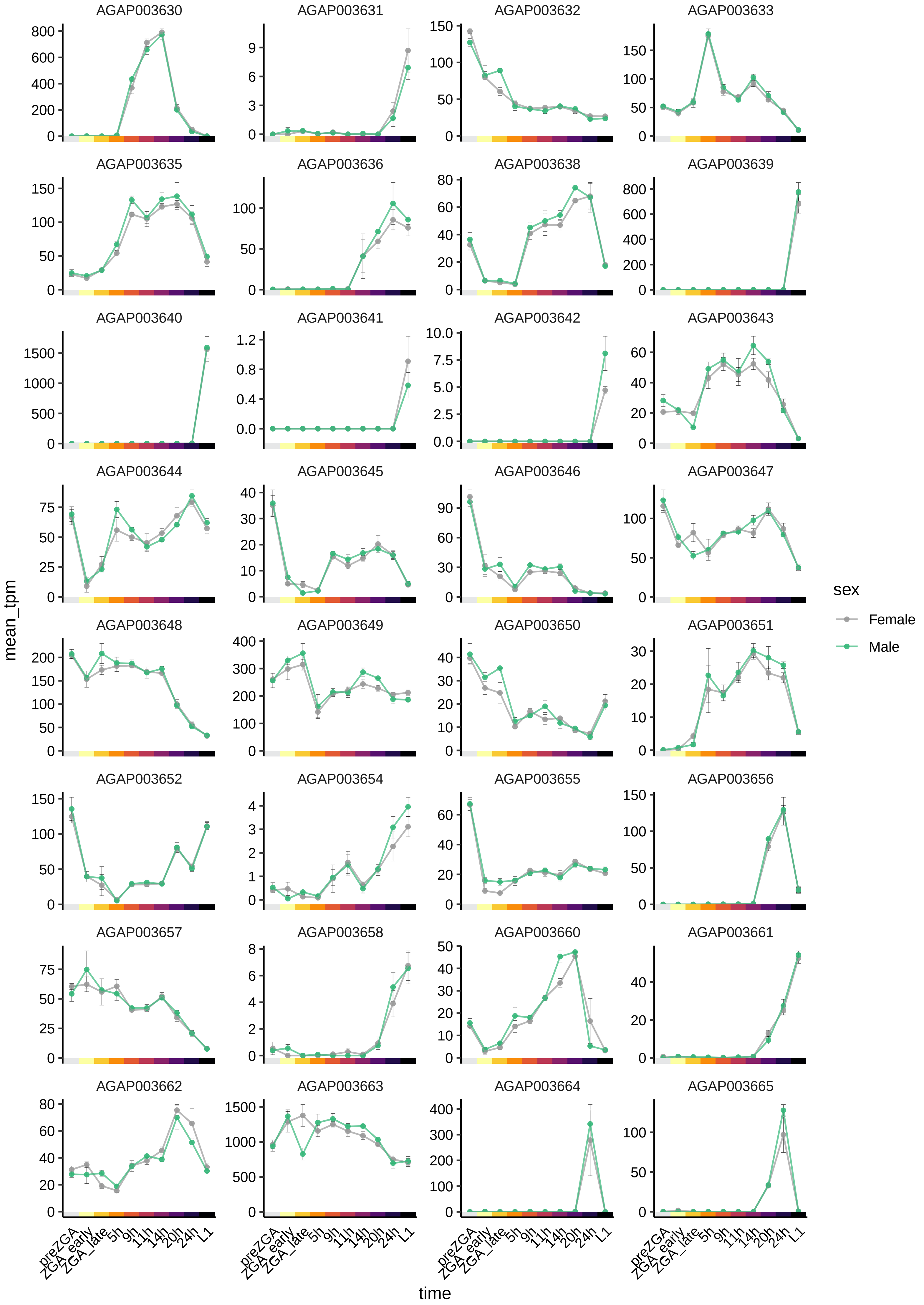
AGAP003630
AGAP003631
AGAP003632
AGAP003633
AGAP003635
AGAP003636
AGAP003638
AGAP003639
AGAP003640
AGAP003641
AGAP003642
AGAP003643
AGAP003644
AGAP003645
AGAP003646
AGAP003647
AGAP003648
AGAP003649
AGAP003650
AGAP003651
AGAP003652
AGAP003654
AGAP003655
AGAP003656
AGAP003657
AGAP003658
AGAP003660
AGAP003661
AGAP003662
AGAP003663
AGAP003664
AGAP003665
mean_tpm
time
preZGA
ZGA_early
ZGA_late
5h
9h
11h
14h
20h
24h
L1
sex
Female
Male

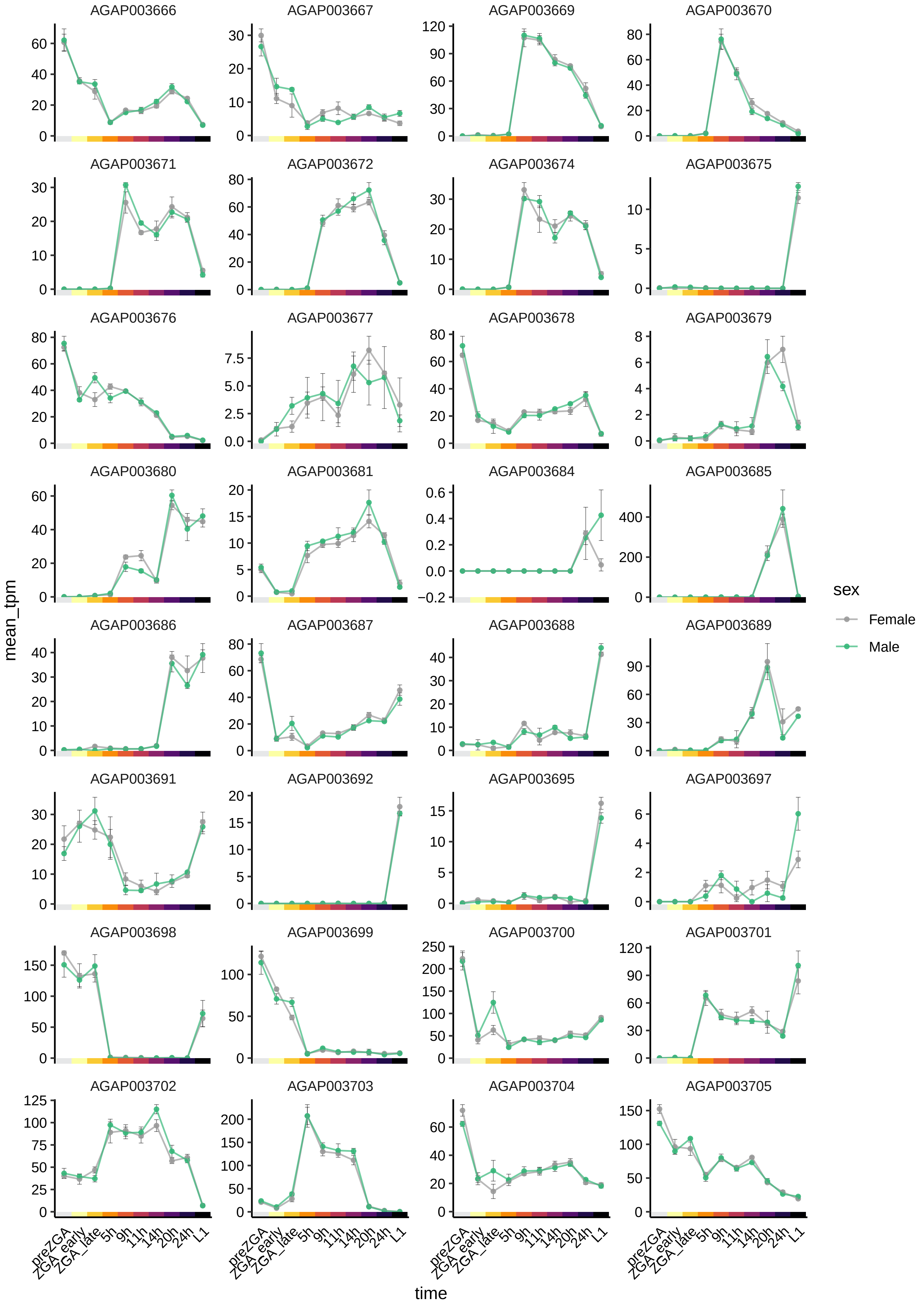

AGAP003666
AGAP003667
AGAP003669
AGAP003670
AGAP003671
AGAP003672
AGAP003674
AGAP003675
AGAP003676
AGAP003677
AGAP003678
AGAP003679
AGAP003680
AGAP003681
AGAP003684
AGAP003685
AGAP003686
AGAP003687
AGAP003688
AGAP003689
AGAP003691
AGAP003692
AGAP003695
AGAP003697
AGAP003698
AGAP003699
AGAP003700
AGAP003701
AGAP003702
AGAP003703
AGAP003704
AGAP003705
mean_tpm
time
preZGA
ZGA_early
ZGA_late
5h
9h
11h
14h
20h
24h
L1
sex
Female
Male

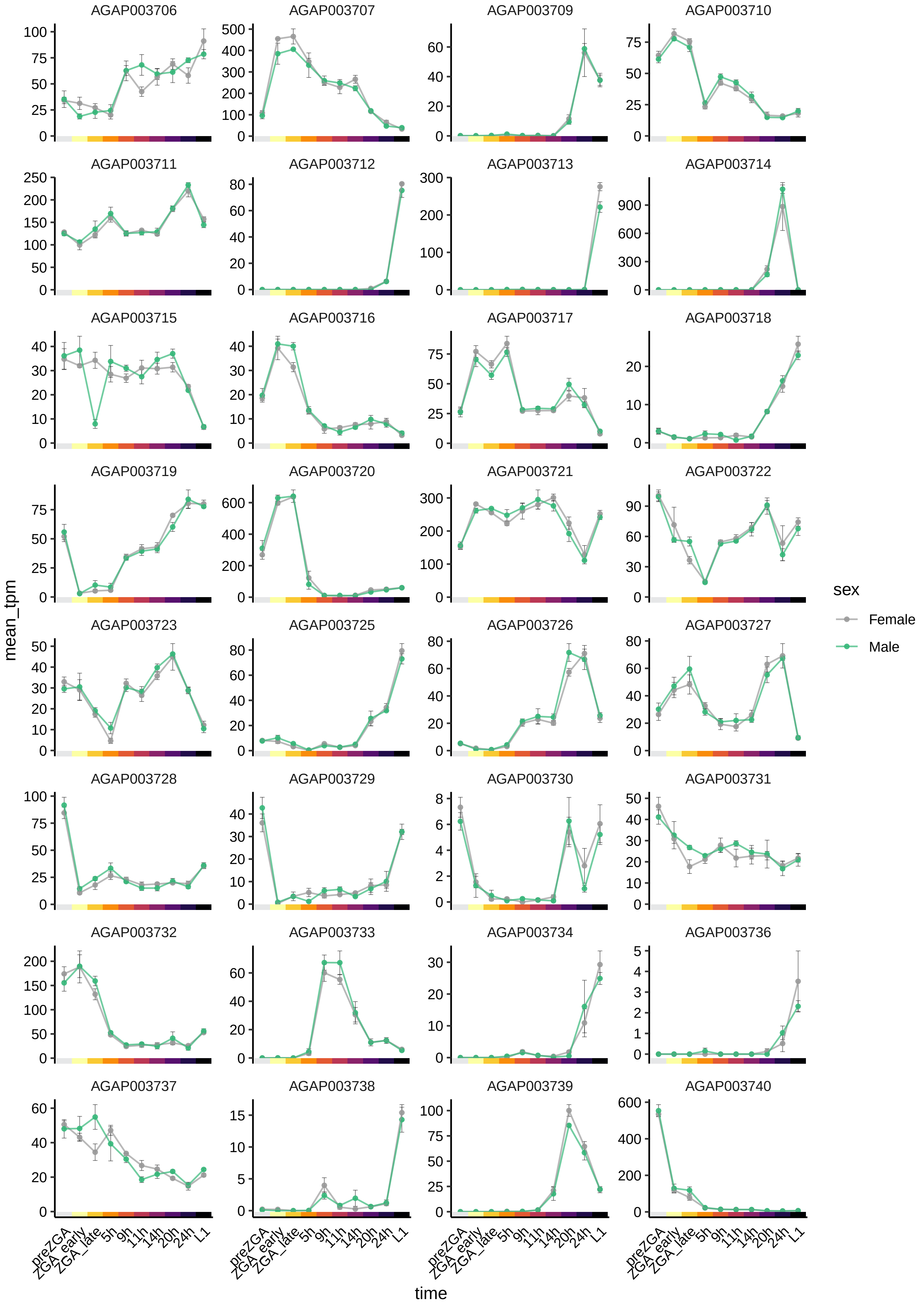

AGAP003706
AGAP003707
AGAP003709
AGAP003710
AGAP003711
AGAP003712
AGAP003713
AGAP003714
AGAP003715
AGAP003716
AGAP003717
AGAP003718
AGAP003719
AGAP003720
AGAP003721
AGAP003722
AGAP003723
AGAP003725
AGAP003726
AGAP003727
AGAP003728
AGAP003729
AGAP003730
AGAP003731
AGAP003732
AGAP003733
AGAP003734
AGAP003736
AGAP003737
AGAP003738
AGAP003739
AGAP003740
mean_tpm
time
preZGA
ZGA_early
ZGA_late
5h
9h
11h
14h
20h
24h
L1
sex
Female
Male

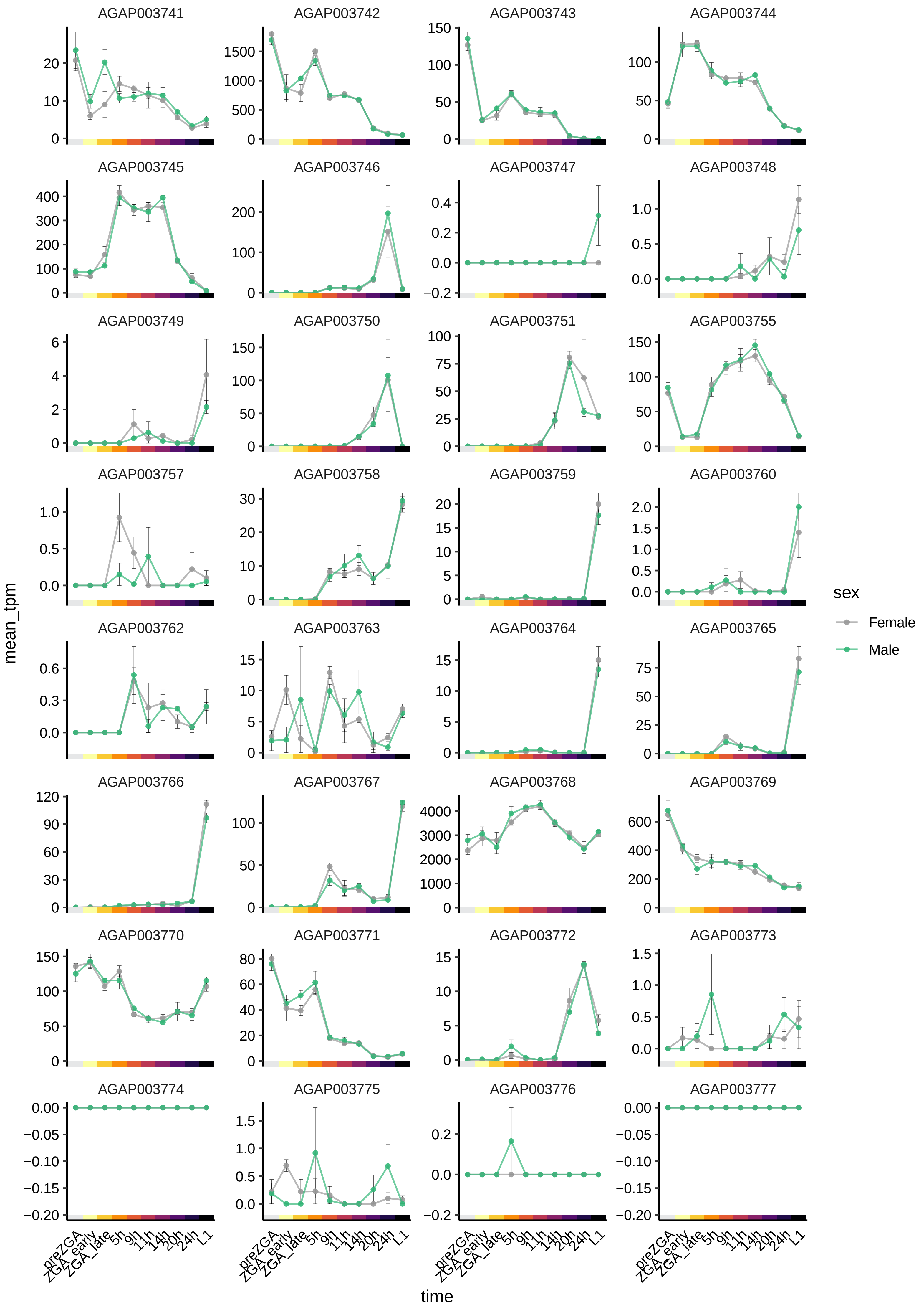
AGAP003741
AGAP003742
AGAP003743
AGAP003744
AGAP003745
AGAP003746
AGAP003747
AGAP003748
AGAP003749
AGAP003750
AGAP003751
AGAP003755
AGAP003757
AGAP003758
AGAP003759
AGAP003760
AGAP003762
AGAP003763
AGAP003764
AGAP003765
AGAP003766
AGAP003767
AGAP003768
AGAP003769
AGAP003770
AGAP003771
AGAP003772
AGAP003773
AGAP003774
AGAP003775
AGAP003776
AGAP003777
mean_tpm
time
preZGA
ZGA_early
ZGA_late
5h
9h
11h
14h
20h
24h
L1
sex
Female
Male

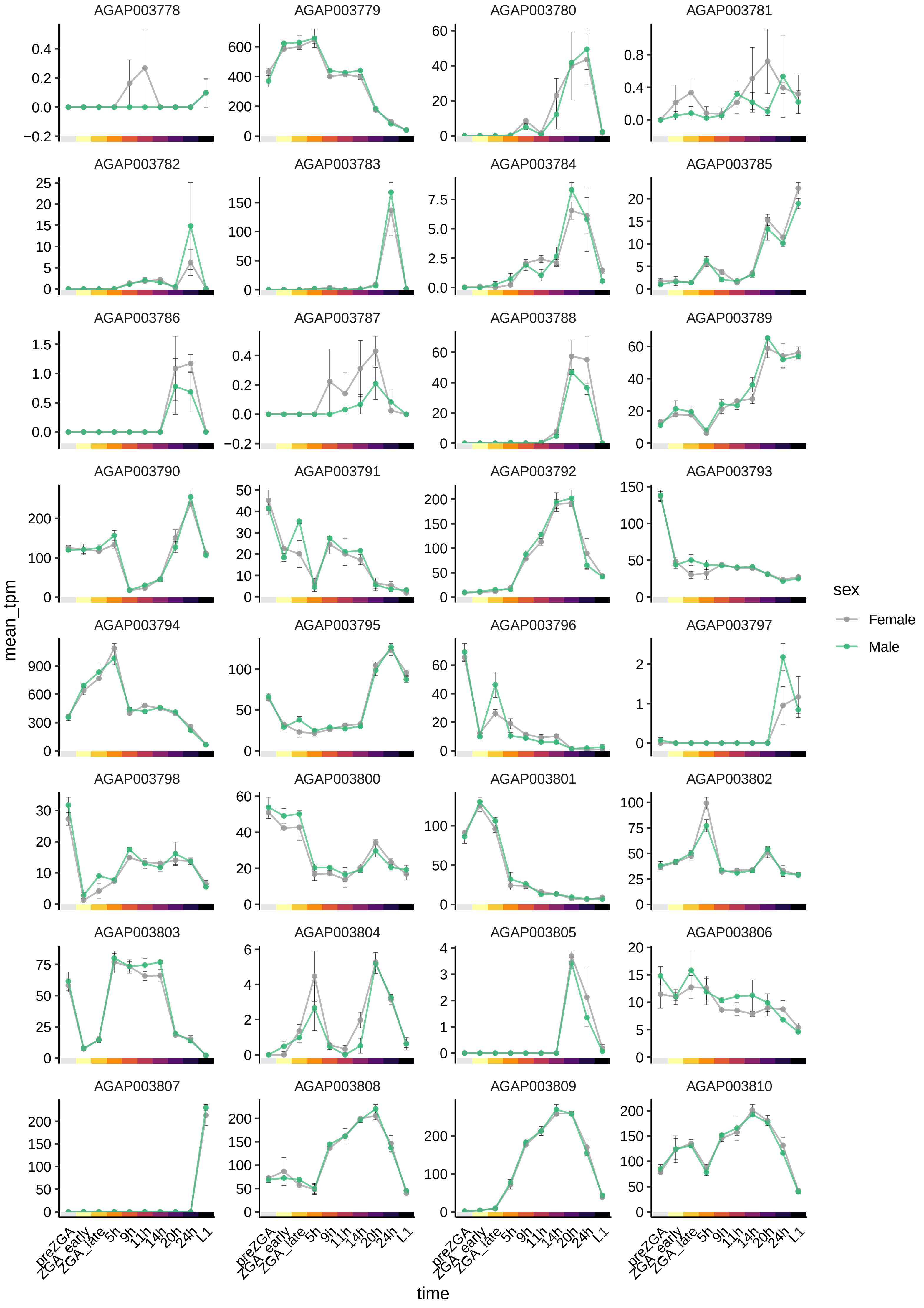
AGAP003778
AGAP003779
AGAP003780
AGAP003781
AGAP003782
AGAP003783
AGAP003784
AGAP003785
AGAP003786
AGAP003787
AGAP003788
AGAP003789
AGAP003790
AGAP003791
AGAP003792
AGAP003793
AGAP003794
AGAP003795
AGAP003796
AGAP003797
AGAP003798
AGAP003800
AGAP003801
AGAP003802
AGAP003803
AGAP003804
AGAP003805
AGAP003806
AGAP003807
AGAP003808
AGAP003809
AGAP003810
mean_tpm
time
preZGA
ZGA_early
ZGA_late
5h
9h
11h
14h
20h
24h
L1
sex
Female
Male

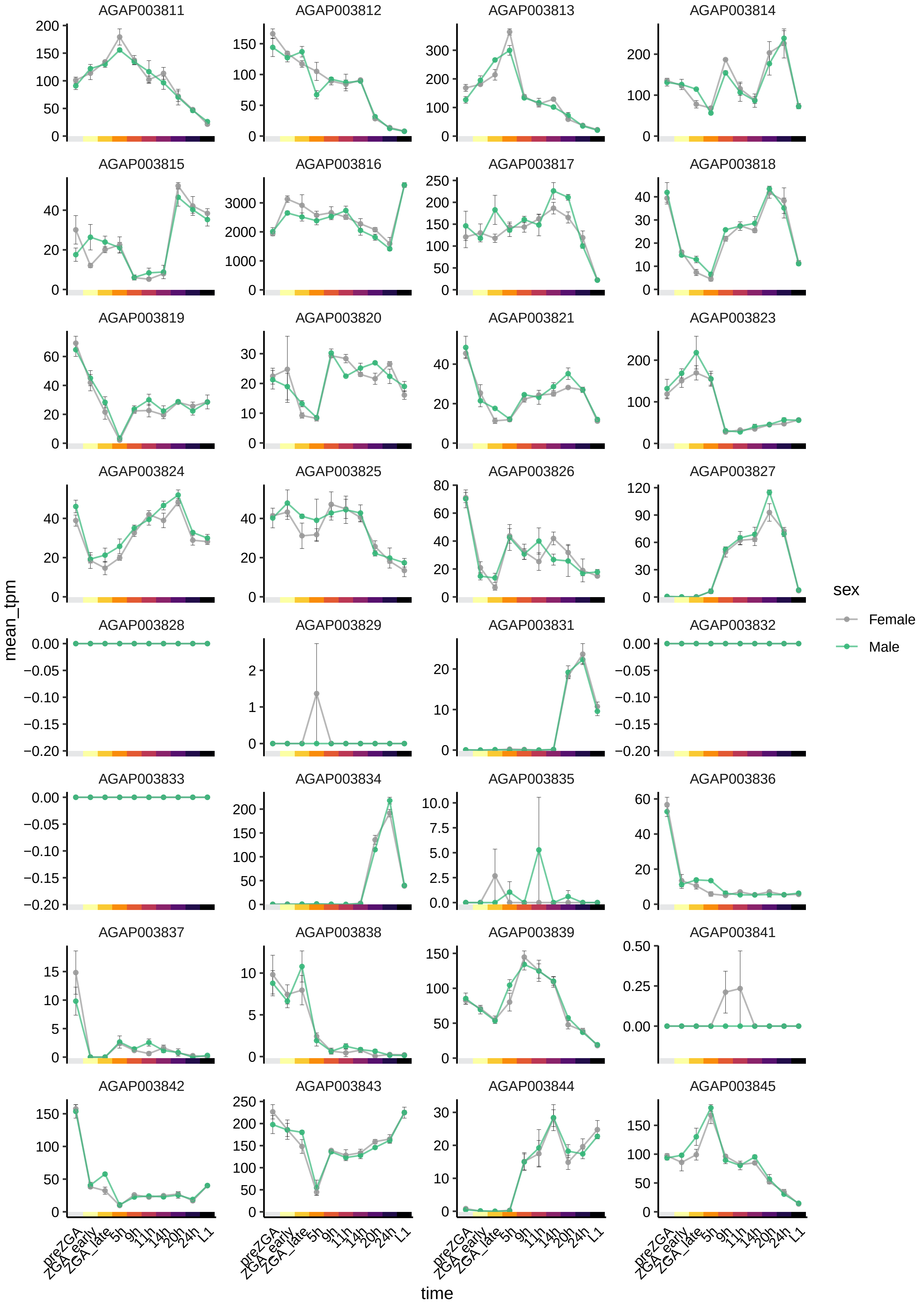

AGAP003811
AGAP003812
AGAP003813
AGAP003814
AGAP003815
AGAP003816
AGAP003817
AGAP003818
AGAP003819
AGAP003820
AGAP003821
AGAP003823
AGAP003824
AGAP003825
AGAP003826
AGAP003827
AGAP003828
AGAP003829
AGAP003831
AGAP003832
AGAP003833
AGAP003834
AGAP003835
AGAP003836
AGAP003837
AGAP003838
AGAP003839
AGAP003841
AGAP003842
AGAP003843
AGAP003844
AGAP003845
mean_tpm
time
preZGA
ZGA_early
ZGA_late
5h
9h
11h
14h
20h
24h
L1
sex
Female
Male

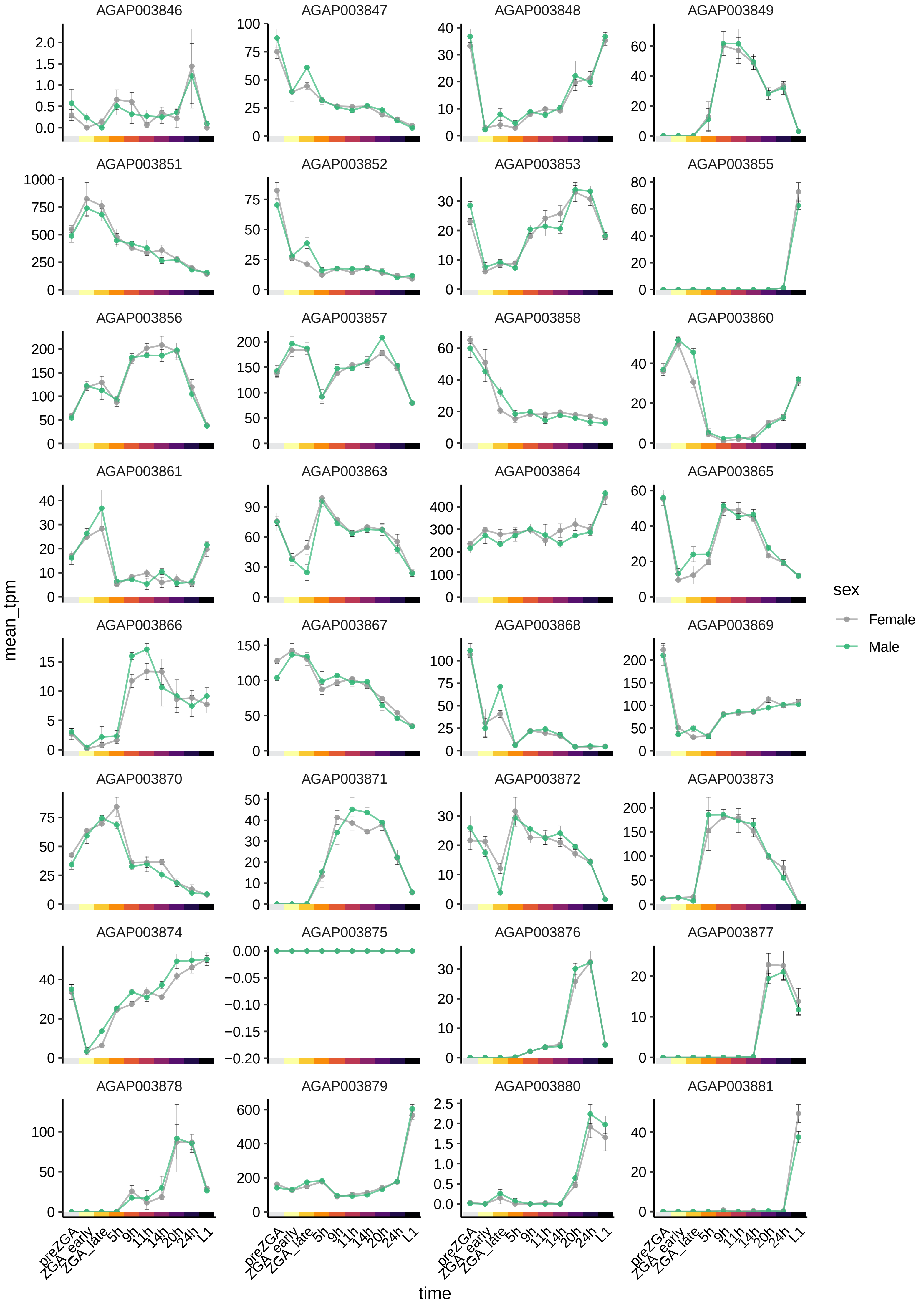
AGAP003846
AGAP003847
AGAP003848
AGAP003849
AGAP003851
AGAP003852
AGAP003853
AGAP003855
AGAP003856
AGAP003857
AGAP003858
AGAP003860
AGAP003861
AGAP003863
AGAP003864
AGAP003865
AGAP003866
AGAP003867
AGAP003868
AGAP003869
AGAP003870
AGAP003871
AGAP003872
AGAP003873
AGAP003874
AGAP003875
AGAP003876
AGAP003877
AGAP003878
AGAP003879
AGAP003880
AGAP003881
mean_tpm
time
preZGA
ZGA_early
ZGA_late
5h
9h
11h
14h
20h
24h
L1
sex
Female
Male

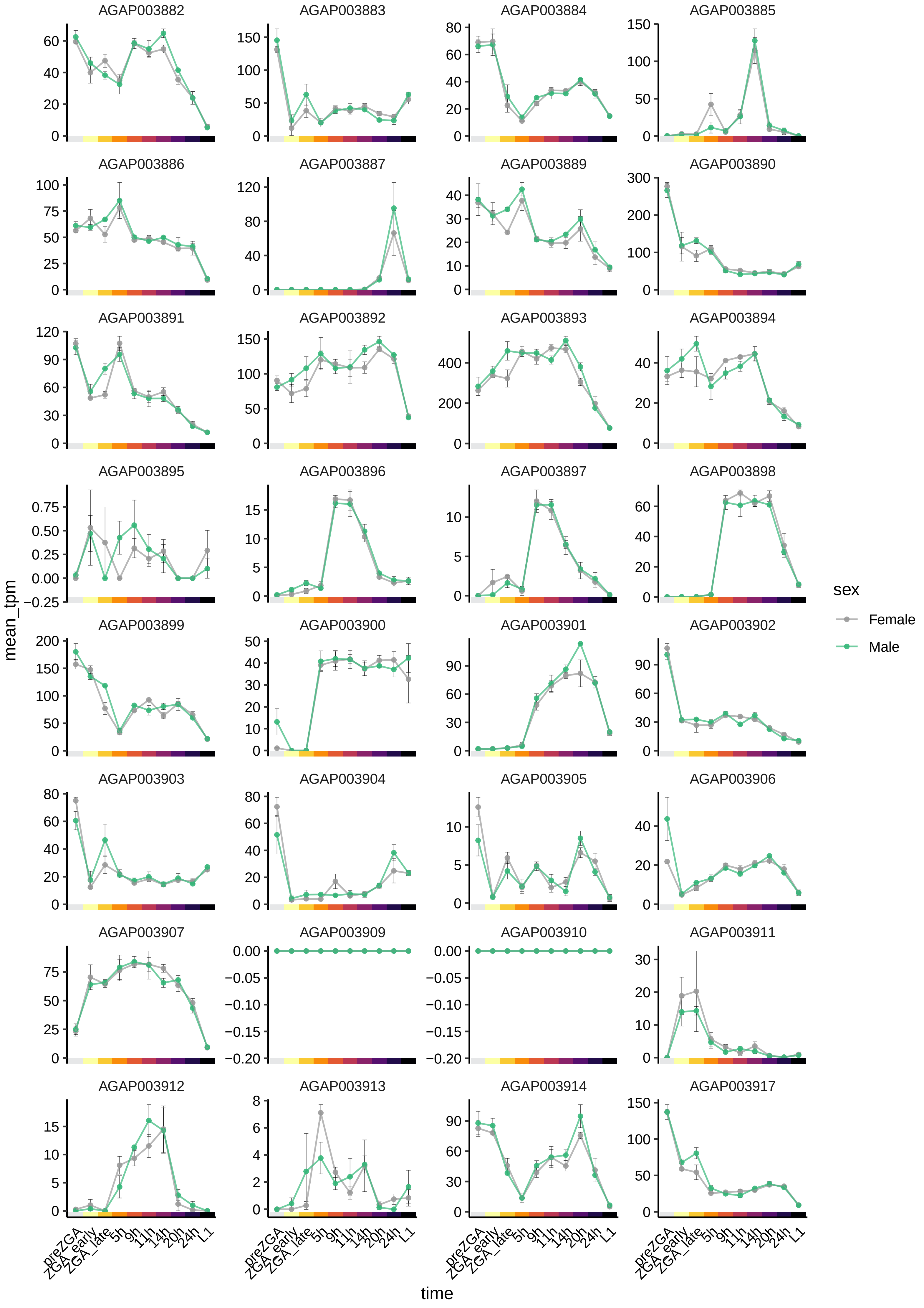
AGAP003882
AGAP003883
AGAP003884
AGAP003885
AGAP003886
AGAP003887
AGAP003889
AGAP003890
AGAP003891
AGAP003892
AGAP003893
AGAP003894
AGAP003895
AGAP003896
AGAP003897
AGAP003898
AGAP003899
AGAP003900
AGAP003901
AGAP003902
AGAP003903
AGAP003904
AGAP003905
AGAP003906
AGAP003907
AGAP003909
AGAP003910
AGAP003911
AGAP003912
AGAP003913
AGAP003914
AGAP003917
mean_tpm
time
preZGA
ZGA_early
ZGA_late
5h
9h
11h
14h
20h
24h
L1
sex
Female
Male

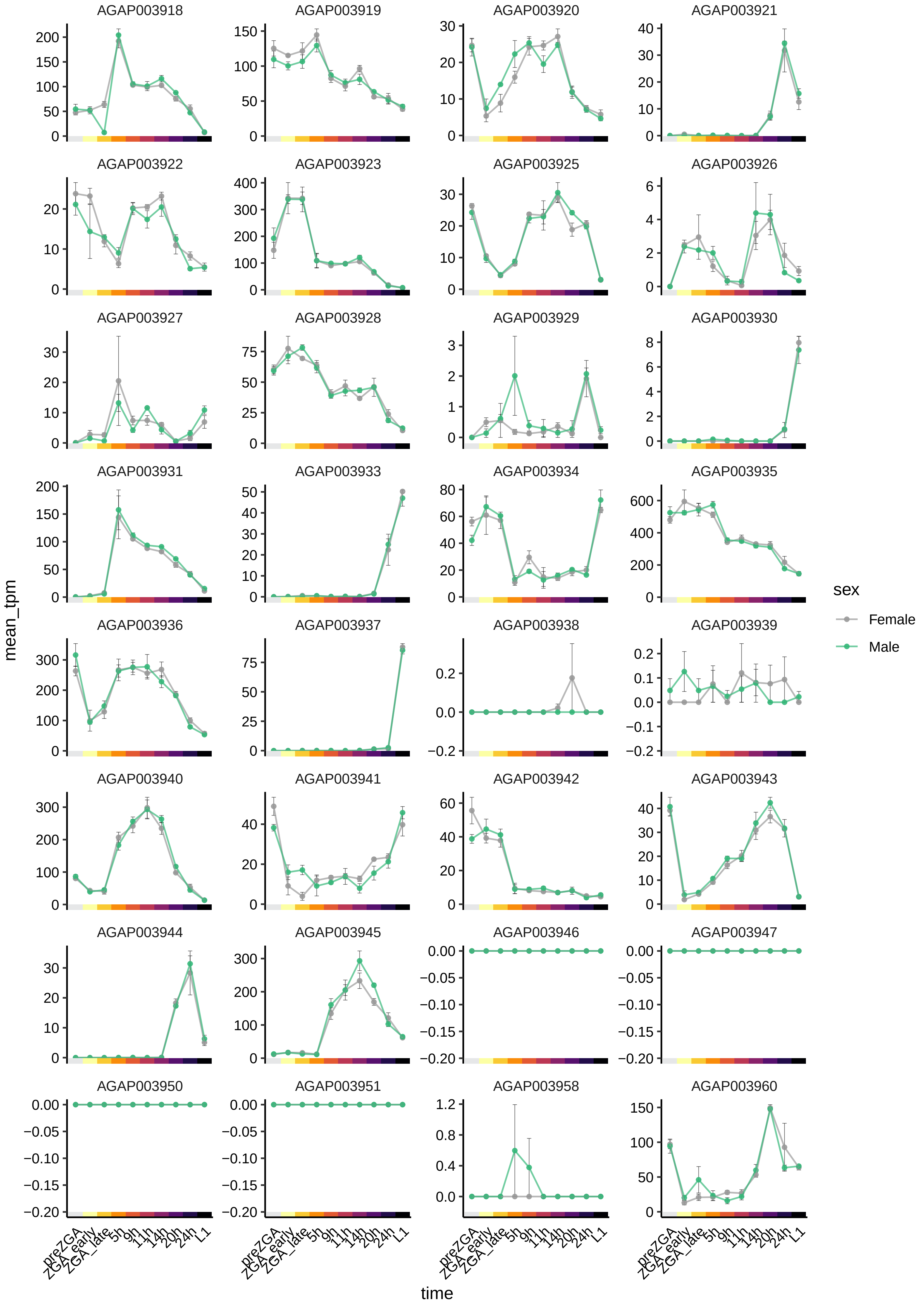

AGAP003918
AGAP003919
AGAP003920
AGAP003921
AGAP003922
AGAP003923
AGAP003925
AGAP003926
AGAP003927
AGAP003928
AGAP003929
AGAP003930
AGAP003931
AGAP003933
AGAP003934
AGAP003935
AGAP003936
AGAP003937
AGAP003938
AGAP003939
AGAP003940
AGAP003941
AGAP003942
AGAP003943
AGAP003944
AGAP003945
AGAP003946
AGAP003947
AGAP003950
AGAP003951
AGAP003958
AGAP003960
mean_tpm
time
preZGA
ZGA_early
ZGA_late
5h
9h
11h
14h
20h
24h
L1
sex
Female
Male

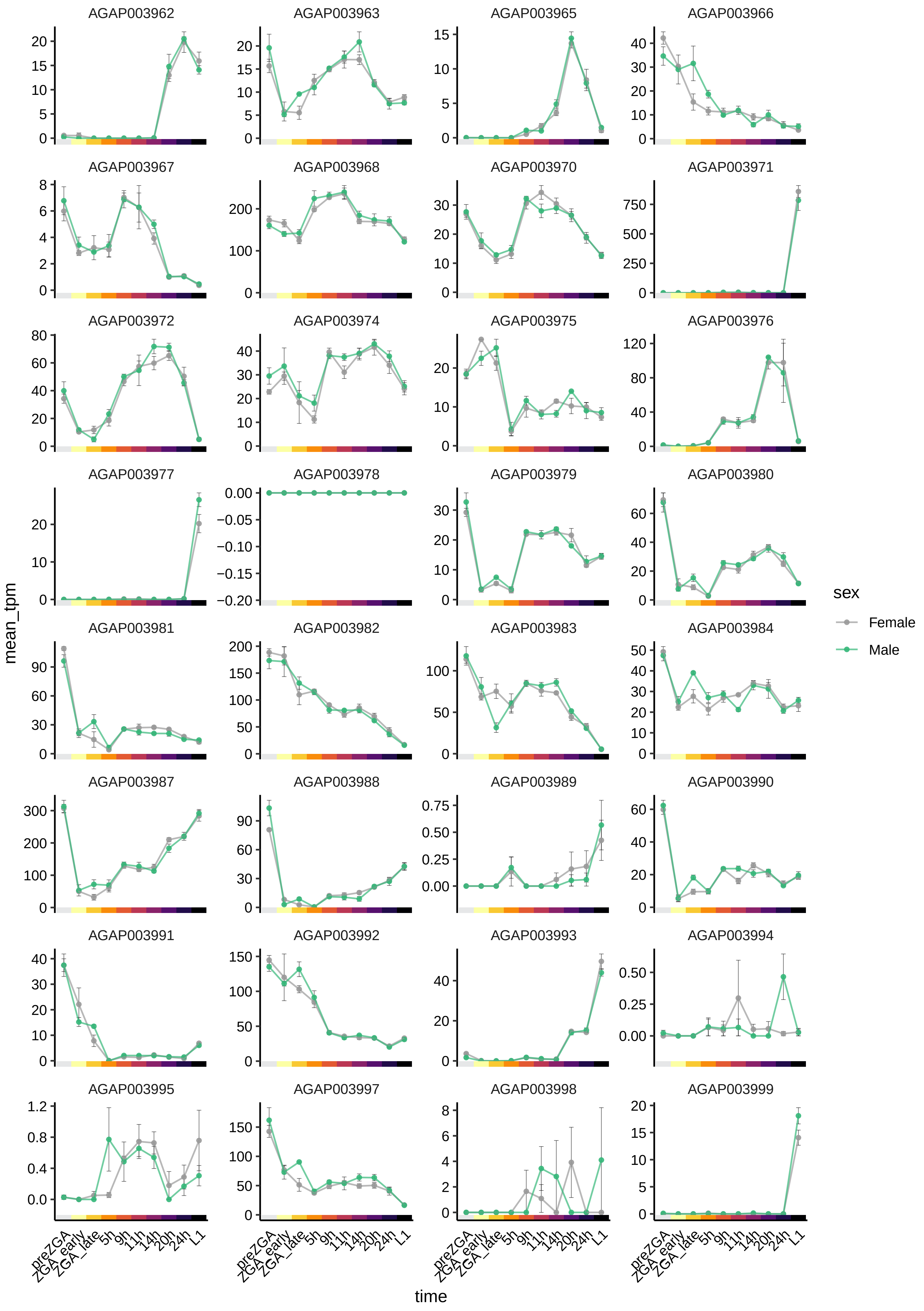
AGAP003962
AGAP003963
AGAP003965
AGAP003966
AGAP003967
AGAP003968
AGAP003970
AGAP003971
AGAP003972
AGAP003974
AGAP003975
AGAP003976
AGAP003977
AGAP003978
AGAP003979
AGAP003980
AGAP003981
AGAP003982
AGAP003983
AGAP003984
AGAP003987
AGAP003988
AGAP003989
AGAP003990
AGAP003991
AGAP003992
AGAP003993
AGAP003994
AGAP003995
AGAP003997
AGAP003998
AGAP003999
mean_tpm
time
preZGA
ZGA_early
ZGA_late
5h
9h
11h
14h
20h
24h
L1
sex
Female
Male

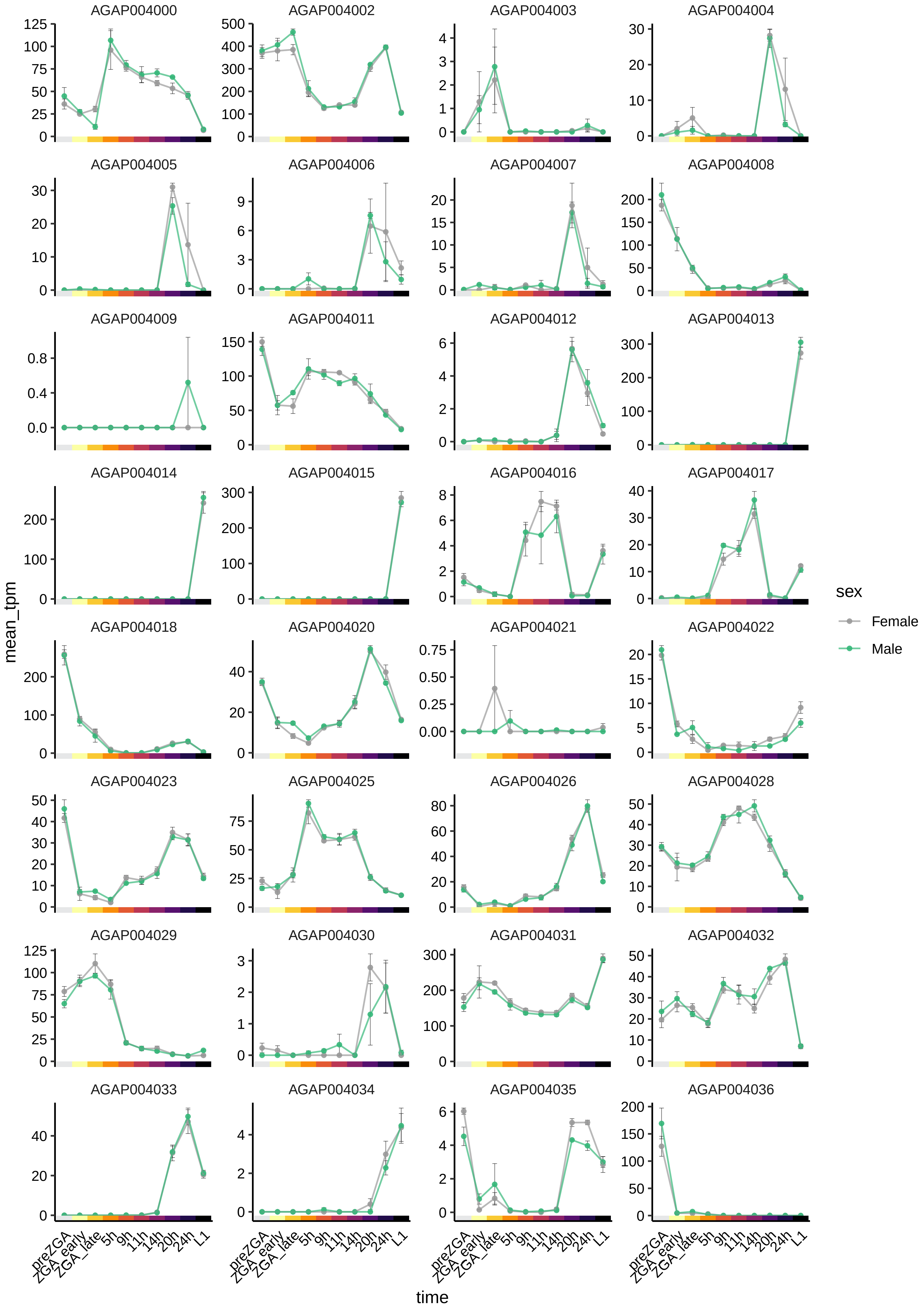

AGAP004000
AGAP004002
AGAP004003
AGAP004004
AGAP004005
AGAP004006
AGAP004007
AGAP004008
AGAP004009
AGAP004011
AGAP004012
AGAP004013
AGAP004014
AGAP004015
AGAP004016
AGAP004017
AGAP004018
AGAP004020
AGAP004021
AGAP004022
AGAP004023
AGAP004025
AGAP004026
AGAP004028
AGAP004029
AGAP004030
AGAP004031
AGAP004032
AGAP004033
AGAP004034
AGAP004035
AGAP004036
mean_tpm
time
preZGA
ZGA_early
ZGA_late
5h
9h
11h
14h
20h
24h
L1
sex
Female
Male

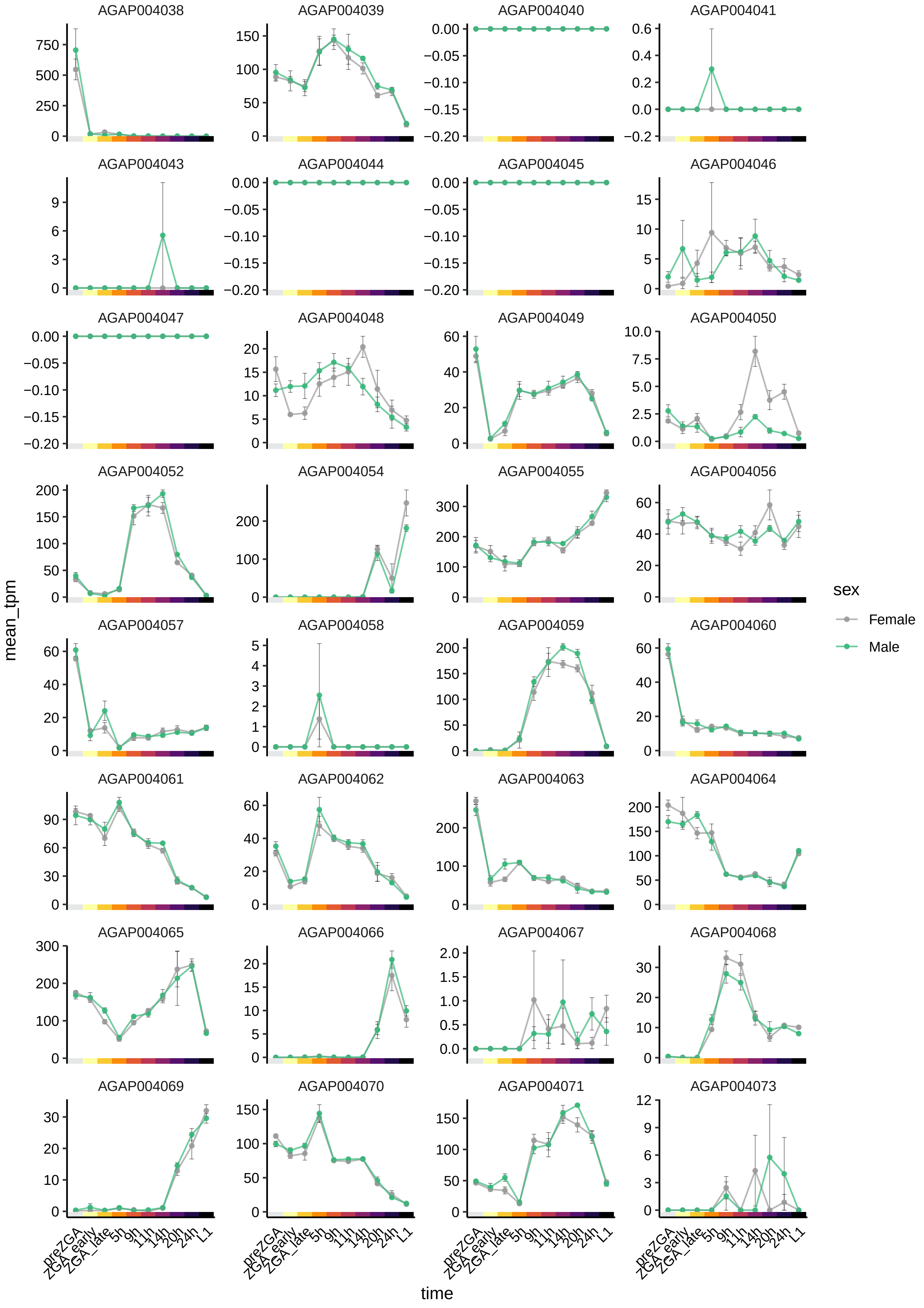

AGAP004038
AGAP004039
AGAP004040
AGAP004041
AGAP004043
AGAP004044
AGAP004045
AGAP004046
AGAP004047
AGAP004048
AGAP004049
AGAP004050
AGAP004052
AGAP004054
AGAP004055
AGAP004056
AGAP004057
AGAP004058
AGAP004059
AGAP004060
AGAP004061
AGAP004062
AGAP004063
AGAP004064
AGAP004065
AGAP004066
AGAP004067
AGAP004068
AGAP004069
AGAP004070
AGAP004071
AGAP004073
mean_tpm
time
preZGA
ZGA_early
ZGA_late
5h
9h
11h
14h
20h
24h
L1
sex
Female
Male

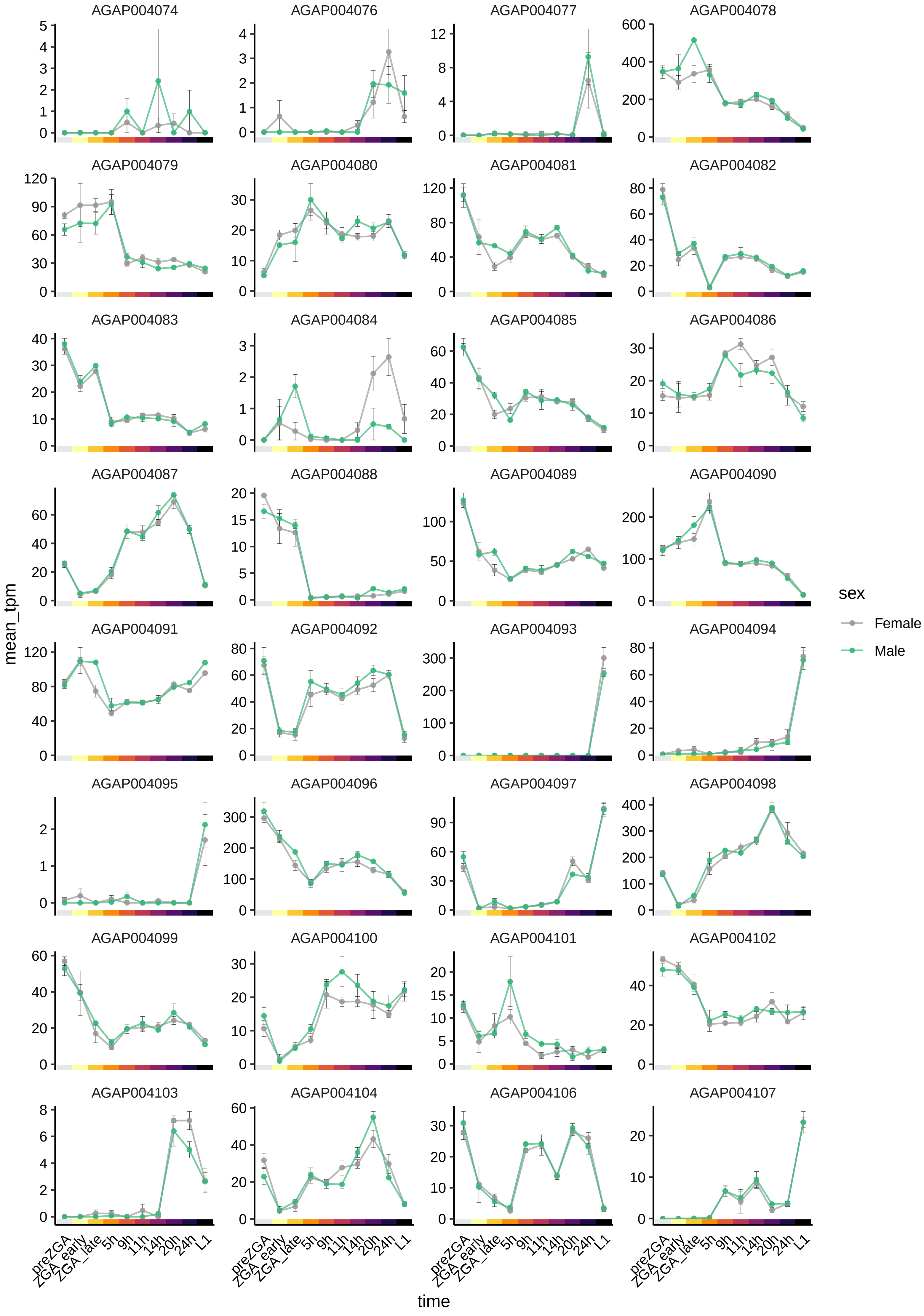

AGAP004074
AGAP004076
AGAP004077
AGAP004078
AGAP004079
AGAP004080
AGAP004081
AGAP004082
AGAP004083
AGAP004084
AGAP004085
AGAP004086
AGAP004087
AGAP004088
AGAP004089
AGAP004090
AGAP004091
AGAP004092
AGAP004093
AGAP004094
AGAP004095
AGAP004096
AGAP004097
AGAP004098
AGAP004099
AGAP004100
AGAP004101
AGAP004102
AGAP004103
AGAP004104
AGAP004106
AGAP004107
mean_tpm
time
preZGA
ZGA_early
ZGA_late
5h
9h
11h
14h
20h
24h
L1
sex
Female
Male

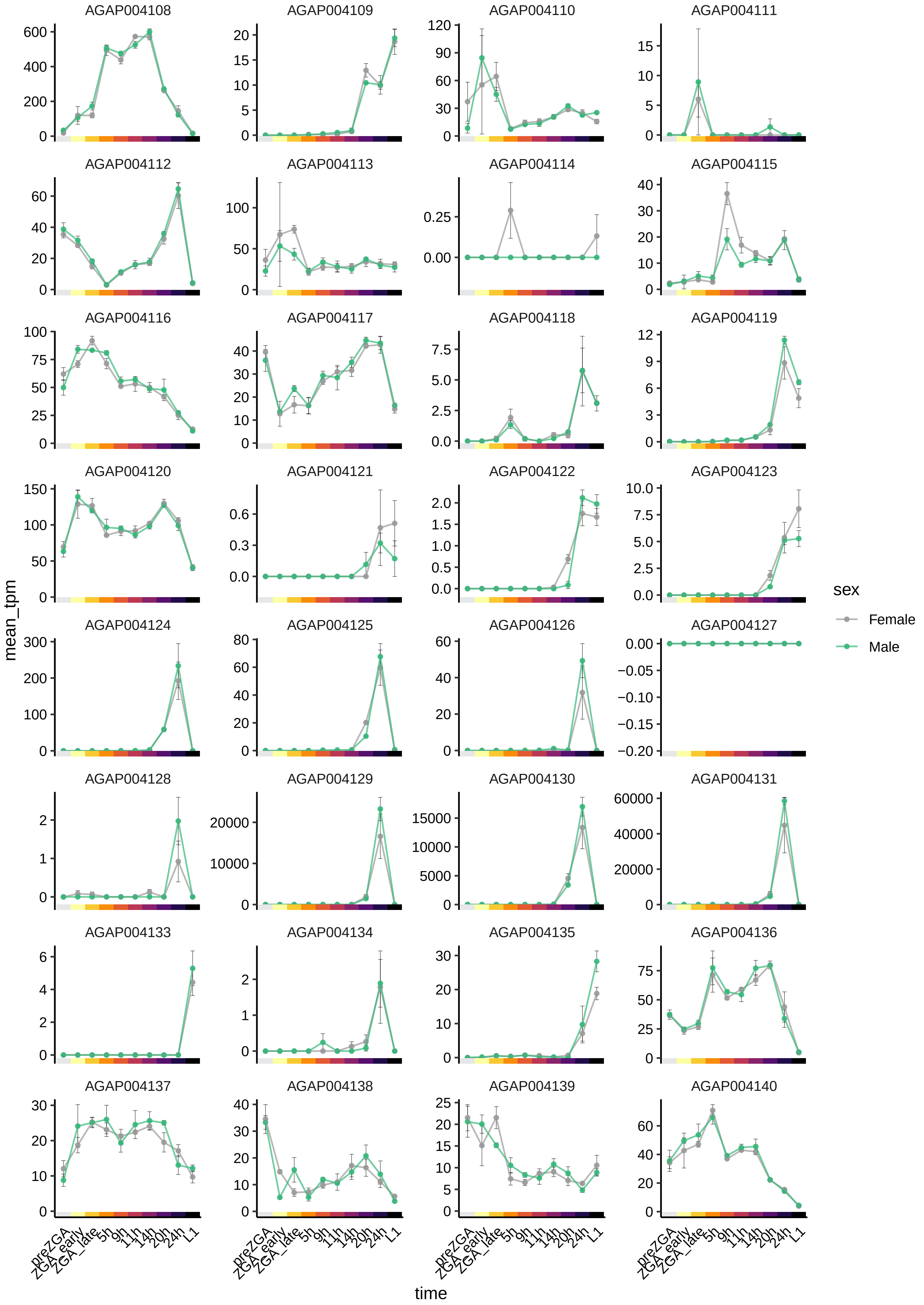
AGAP004108
AGAP004109
AGAP004110
AGAP004111
AGAP004112
AGAP004113
AGAP004114
AGAP004115
AGAP004116
AGAP004117
AGAP004118
AGAP004119
AGAP004120
AGAP004121
AGAP004122
AGAP004123
AGAP004124
AGAP004125
AGAP004126
AGAP004127
AGAP004128
AGAP004129
AGAP004130
AGAP004131
AGAP004133
AGAP004134
AGAP004135
AGAP004136
AGAP004137
AGAP004138
AGAP004139
AGAP004140
mean_tpm
time
preZGA
ZGA_early
ZGA_late
5h
9h
11h
14h
20h
24h
L1
sex
Female
Male

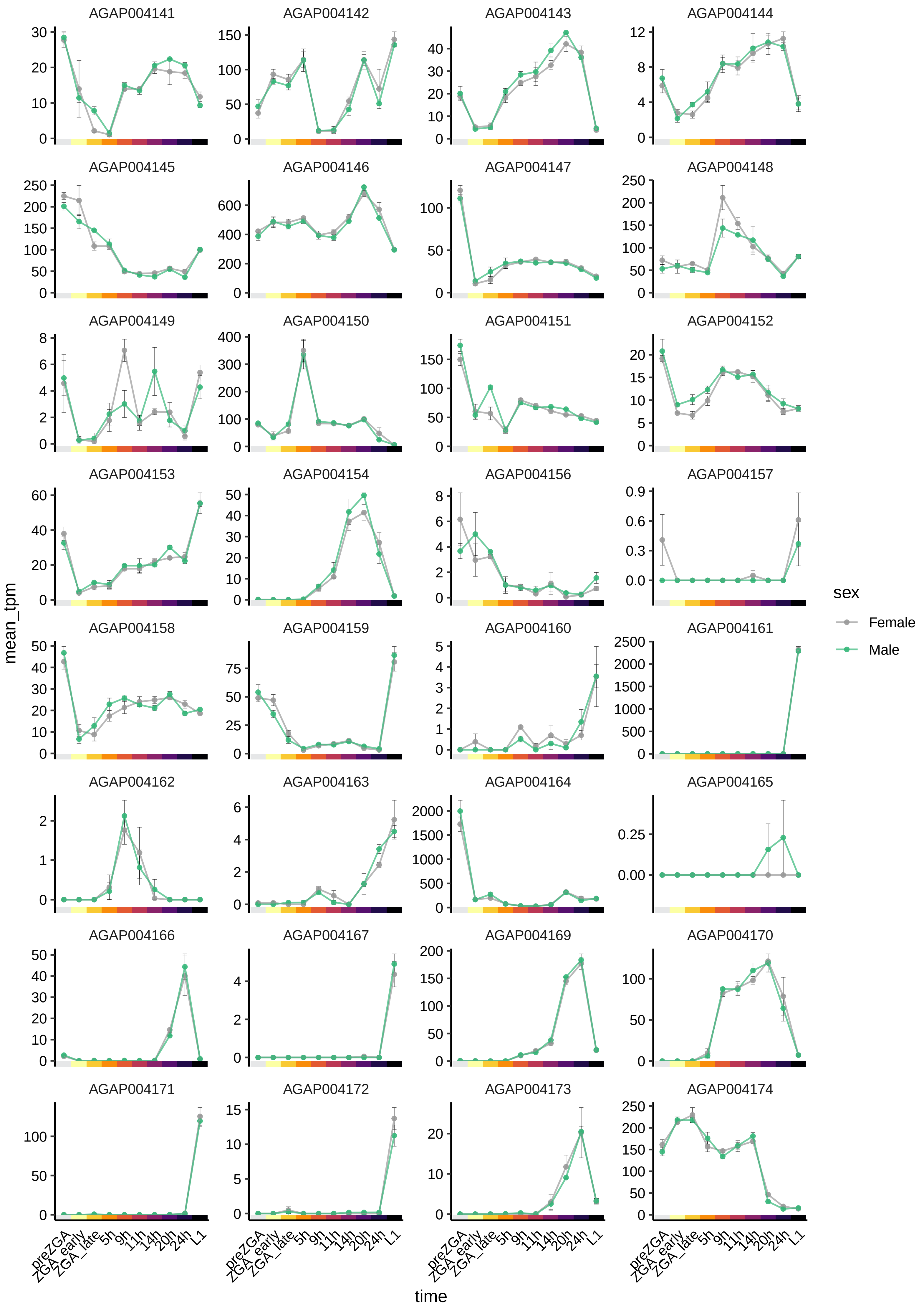

AGAP004141
AGAP004142
AGAP004143
AGAP004144
AGAP004145
AGAP004146
AGAP004147
AGAP004148
AGAP004149
AGAP004150
AGAP004151
AGAP004152
AGAP004153
AGAP004154
AGAP004156
AGAP004157
AGAP004158
AGAP004159
AGAP004160
AGAP004161
AGAP004162
AGAP004163
AGAP004164
AGAP004165
AGAP004166
AGAP004167
AGAP004169
AGAP004170
AGAP004171
AGAP004172
AGAP004173
AGAP004174
mean_tpm
time
preZGA
ZGA_early
ZGA_late
5h
9h
11h
14h
20h
24h
L1
sex
Female
Male

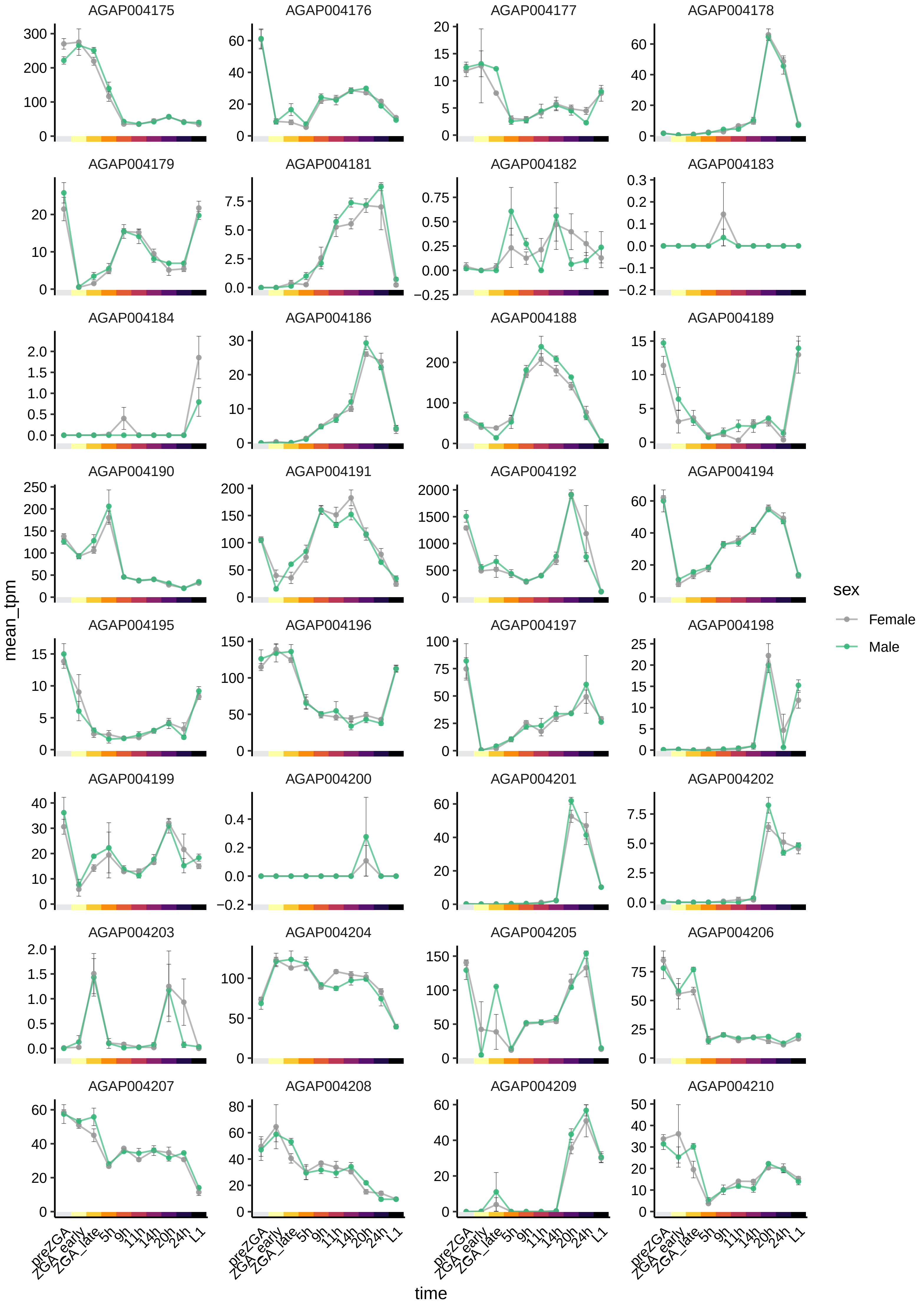

AGAP004175
AGAP004176
AGAP004177
AGAP004178
AGAP004179
AGAP004181
AGAP004182
AGAP004183
AGAP004184
AGAP004186
AGAP004188
AGAP004189
AGAP004190
AGAP004191
AGAP004192
AGAP004194
AGAP004195
AGAP004196
AGAP004197
AGAP004198
AGAP004199
AGAP004200
AGAP004201
AGAP004202
AGAP004203
AGAP004204
AGAP004205
AGAP004206
AGAP004207
AGAP004208
AGAP004209
AGAP004210
mean_tpm
time
preZGA
ZGA_early
ZGA_late
5h
9h
11h
14h
20h
24h
L1
sex
Female
Male

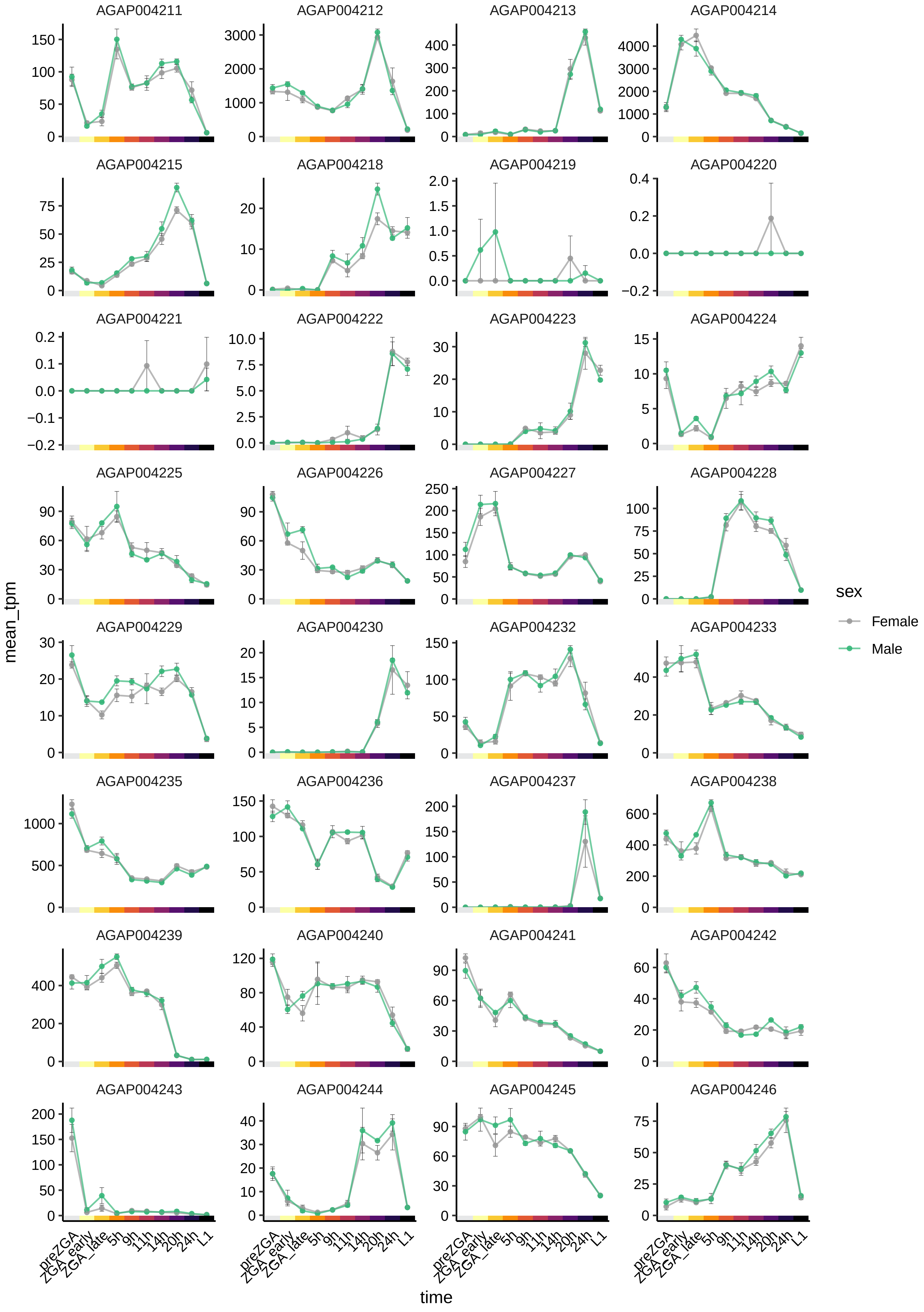

AGAP004211
AGAP004212
AGAP004213
AGAP004214
AGAP004215
AGAP004218
AGAP004219
AGAP004220
AGAP004221
AGAP004222
AGAP004223
AGAP004224
AGAP004225
AGAP004226
AGAP004227
AGAP004228
AGAP004229
AGAP004230
AGAP004232
AGAP004233
AGAP004235
AGAP004236
AGAP004237
AGAP004238
AGAP004239
AGAP004240
AGAP004241
AGAP004242
AGAP004243
AGAP004244
AGAP004245
AGAP004246
mean_tpm
time
preZGA
ZGA_early
ZGA_late
5h
9h
11h
14h
20h
24h
L1
sex
Female
Male

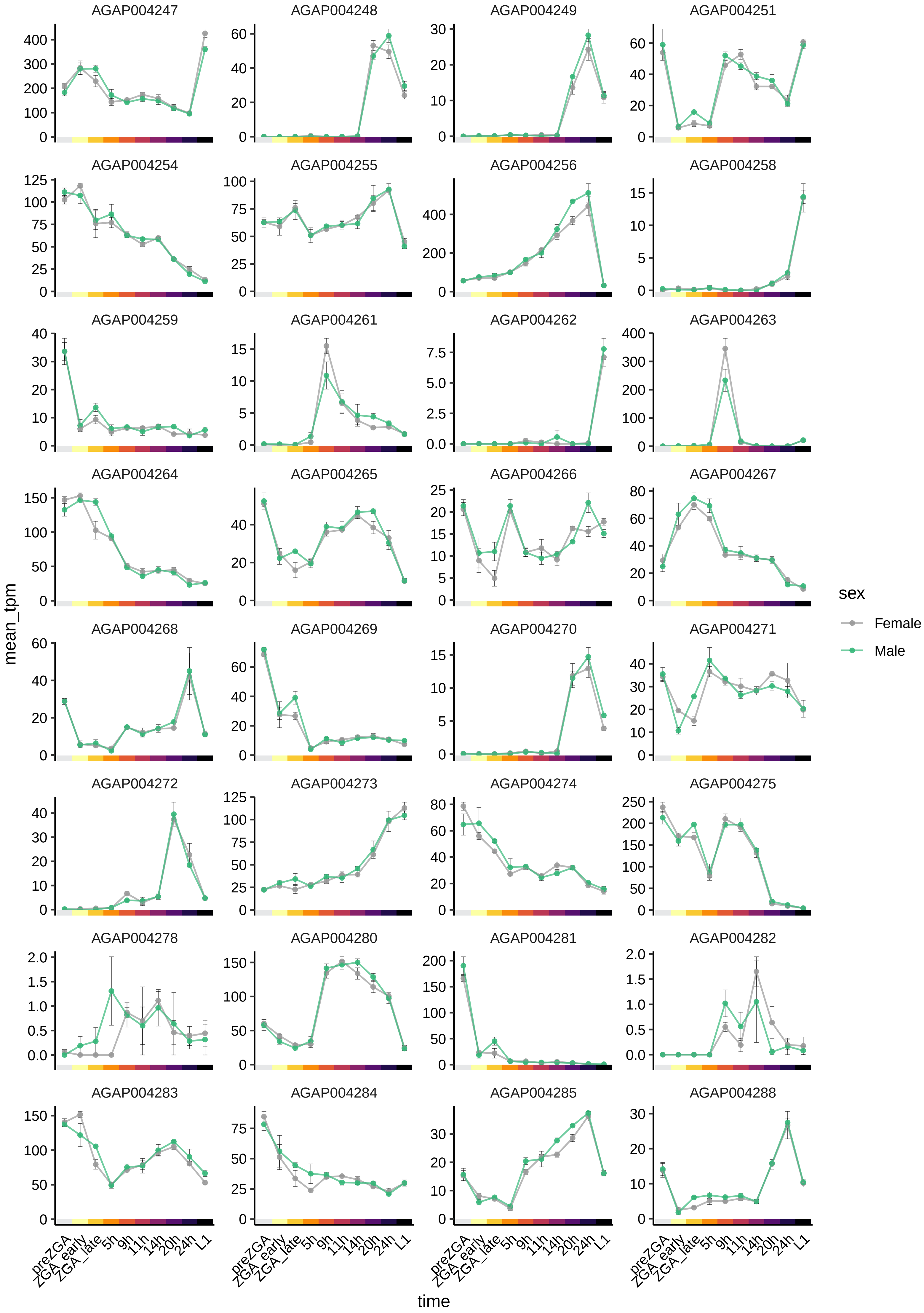

AGAP004247
AGAP004248
AGAP004249
AGAP004251
AGAP004254
AGAP004255
AGAP004256
AGAP004258
AGAP004259
AGAP004261
AGAP004262
AGAP004263
AGAP004264
AGAP004265
AGAP004266
AGAP004267
AGAP004268
AGAP004269
AGAP004270
AGAP004271
AGAP004272
AGAP004273
AGAP004274
AGAP004275
AGAP004278
AGAP004280
AGAP004281
AGAP004282
AGAP004283
AGAP004284
AGAP004285
AGAP004288
mean_tpm
time
preZGA
ZGA_early
ZGA_late
5h
9h
11h
14h
20h
24h
L1
sex
Female
Male

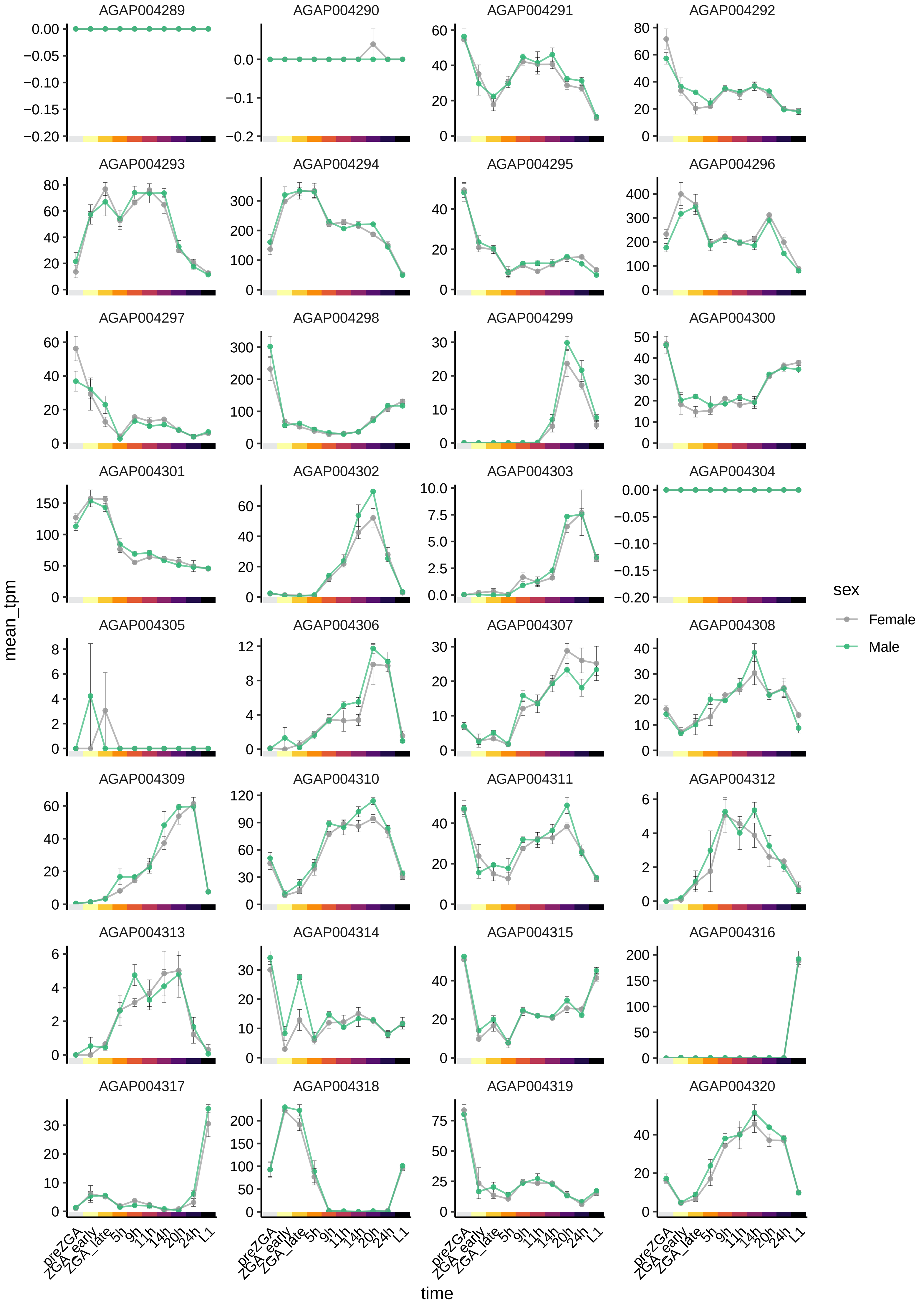

AGAP004289
AGAP004290
AGAP004291
AGAP004292
AGAP004293
AGAP004294
AGAP004295
AGAP004296
AGAP004297
AGAP004298
AGAP004299
AGAP004300
AGAP004301
AGAP004302
AGAP004303
AGAP004304
AGAP004305
AGAP004306
AGAP004307
AGAP004308
AGAP004309
AGAP004310
AGAP004311
AGAP004312
AGAP004313
AGAP004314
AGAP004315
AGAP004316
AGAP004317
AGAP004318
AGAP004319
AGAP004320
mean_tpm
time
preZGA
ZGA_early
ZGA_late
5h
9h
11h
14h
20h
24h
L1
sex
Female
Male

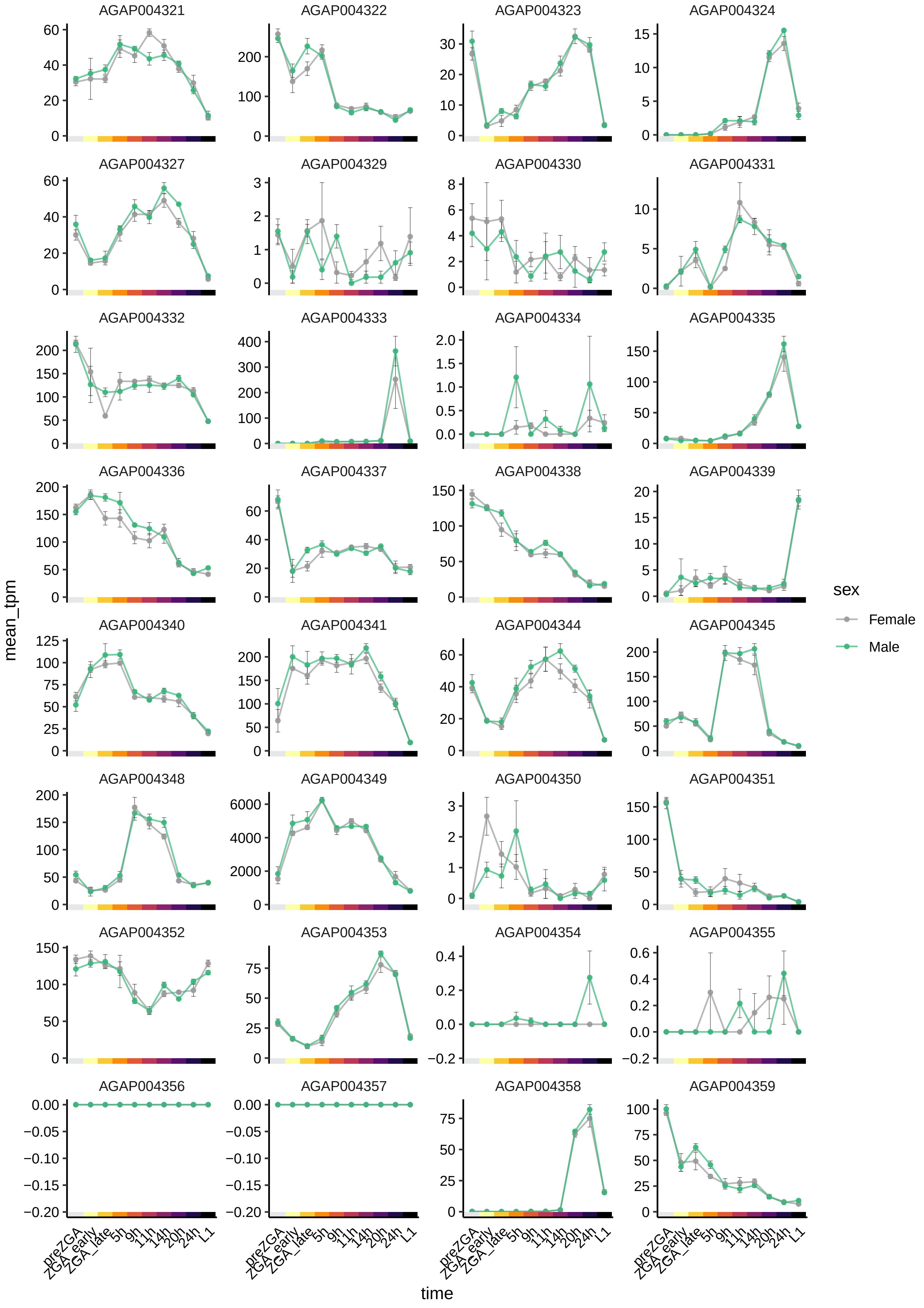

AGAP004321
AGAP004322
AGAP004323
AGAP004324
AGAP004327
AGAP004329
AGAP004330
AGAP004331
AGAP004332
AGAP004333
AGAP004334
AGAP004335
AGAP004336
AGAP004337
AGAP004338
AGAP004339
AGAP004340
AGAP004341
AGAP004344
AGAP004345
AGAP004348
AGAP004349
AGAP004350
AGAP004351
AGAP004352
AGAP004353
AGAP004354
AGAP004355
AGAP004356
AGAP004357
AGAP004358
AGAP004359
mean_tpm
time
preZGA
ZGA_early
ZGA_late
5h
9h
11h
14h
20h
24h
L1
sex
Female
Male

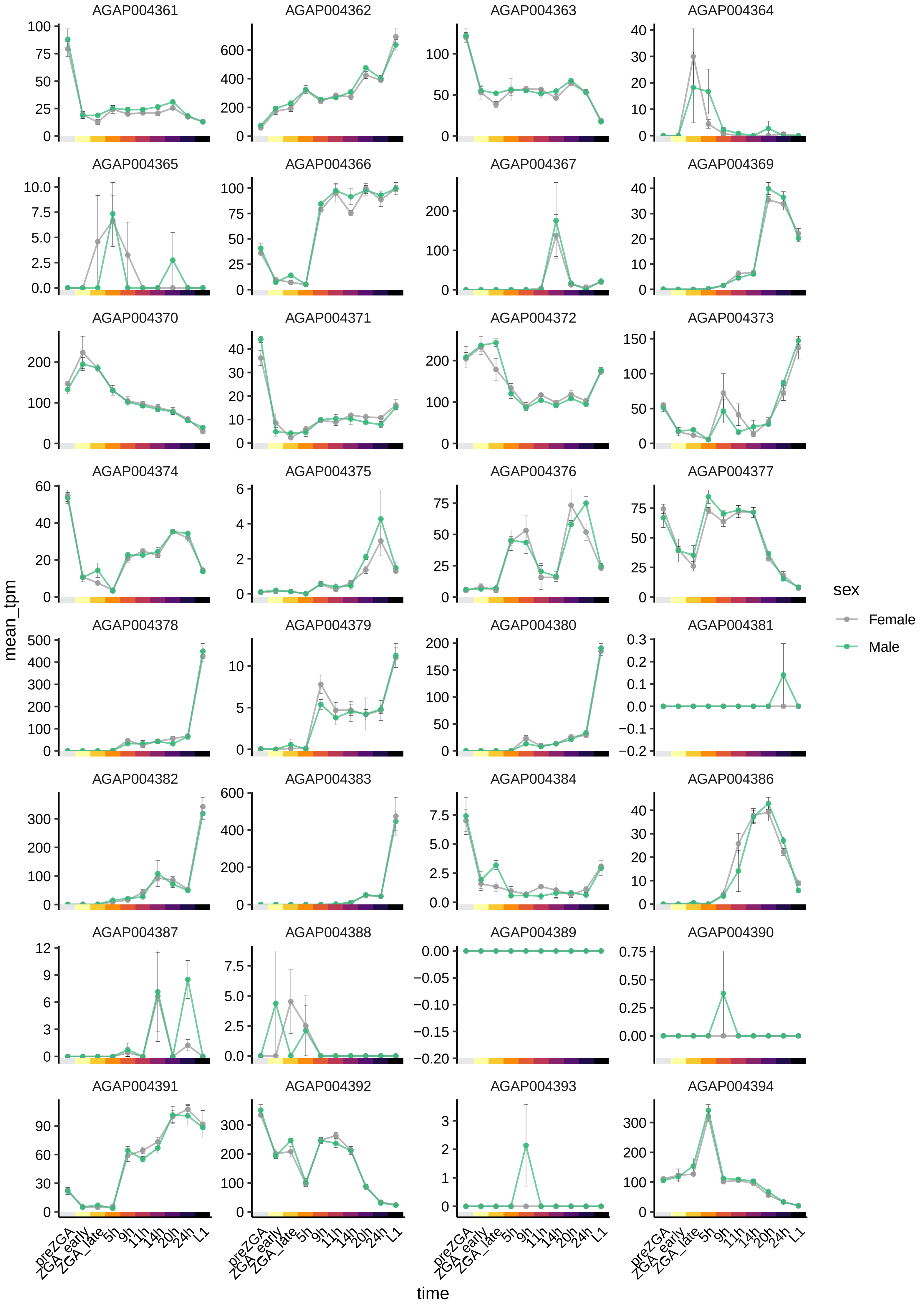

AGAP004361
AGAP004362
AGAP004363
AGAP004364
AGAP004365
AGAP004366
AGAP004367
AGAP004369
AGAP004370
AGAP004371
AGAP004372
AGAP004373
AGAP004374
AGAP004375
AGAP004376
AGAP004377
AGAP004378
AGAP004379
AGAP004380
AGAP004381
AGAP004382
AGAP004383
AGAP004384
AGAP004386
AGAP004387
AGAP004388
AGAP004389
AGAP004390
AGAP004391
AGAP004392
AGAP004393
AGAP004394
mean_tpm
time
preZGA
ZGA_early
ZGA_late
5h
9h
11h
14h
20h
24h
L1
sex
Female
Male

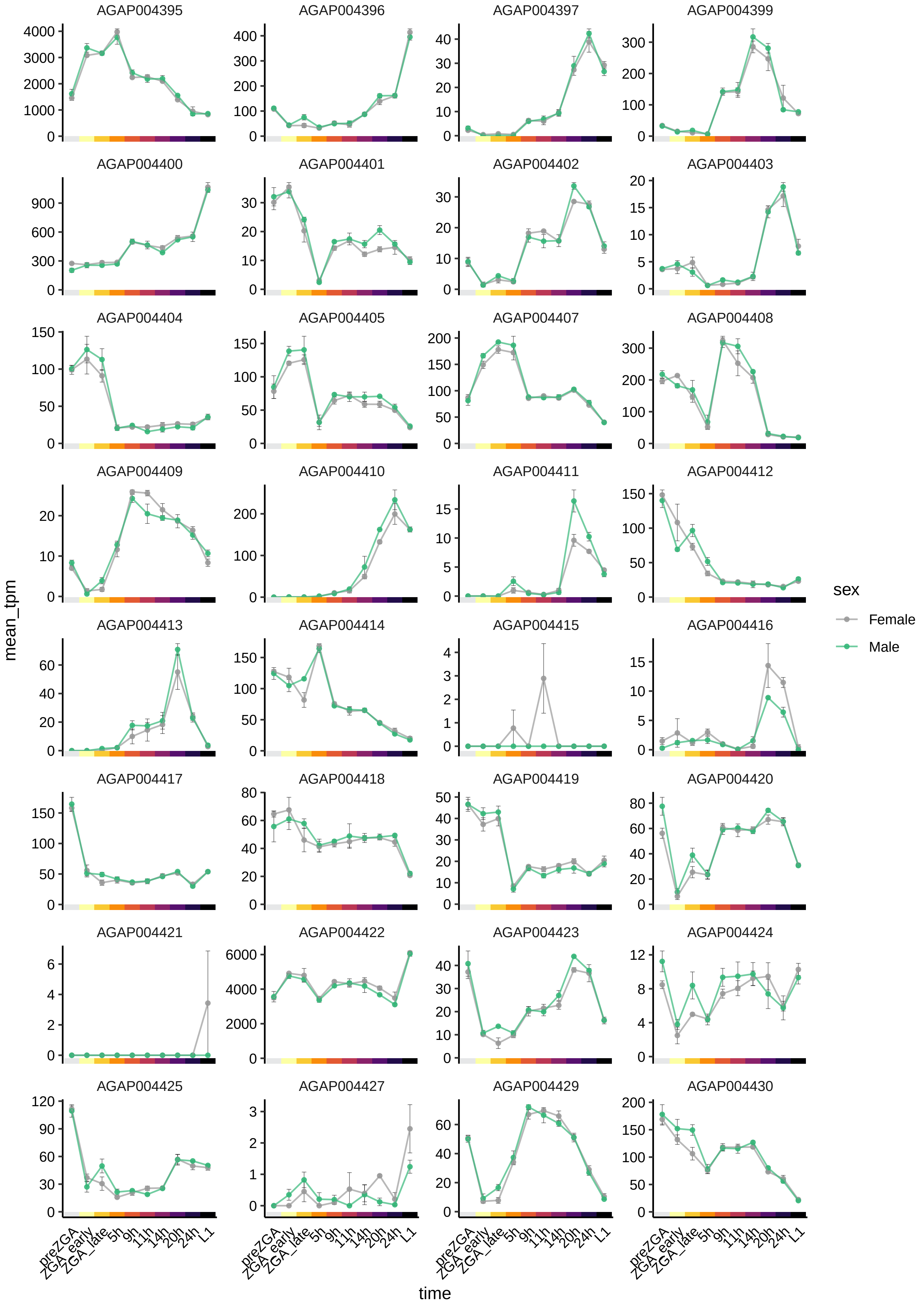

AGAP004395
AGAP004396
AGAP004397
AGAP004399
AGAP004400
AGAP004401
AGAP004402
AGAP004403
AGAP004404
AGAP004405
AGAP004407
AGAP004408
AGAP004409
AGAP004410
AGAP004411
AGAP004412
AGAP004413
AGAP004414
AGAP004415
AGAP004416
AGAP004417
AGAP004418
AGAP004419
AGAP004420
AGAP004421
AGAP004422
AGAP004423
AGAP004424
AGAP004425
AGAP004427
AGAP004429
AGAP004430
mean_tpm
time
preZGA
ZGA_early
ZGA_late
5h
9h
11h
14h
20h
24h
L1
sex
Female
Male

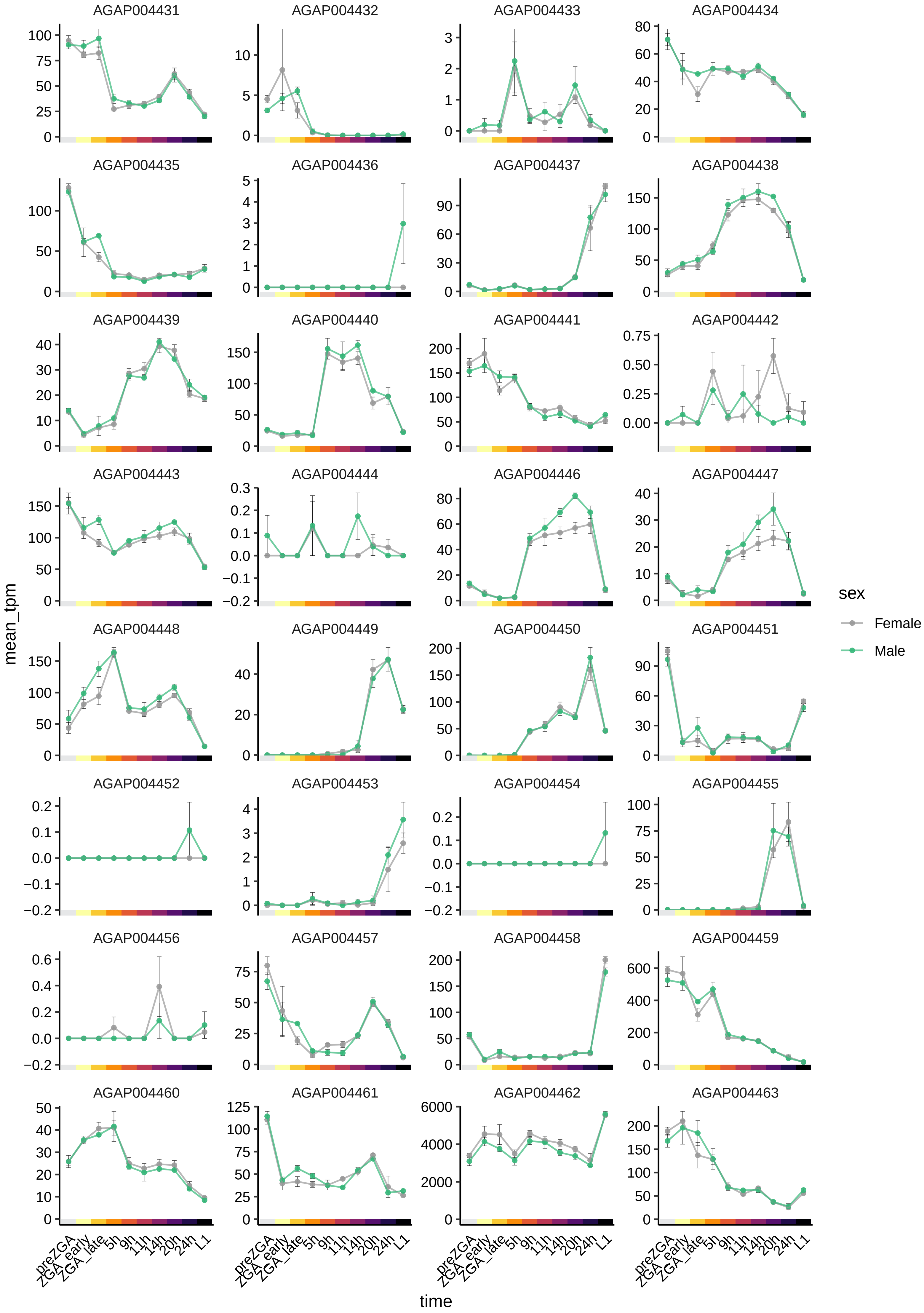
AGAP004431
AGAP004432
AGAP004433
AGAP004434
AGAP004435
AGAP004436
AGAP004437
AGAP004438
AGAP004439
AGAP004440
AGAP004441
AGAP004442
AGAP004443
AGAP004444
AGAP004446
AGAP004447
AGAP004448
AGAP004449
AGAP004450
AGAP004451
AGAP004452
AGAP004453
AGAP004454
AGAP004455
AGAP004456
AGAP004457
AGAP004458
AGAP004459
AGAP004460
AGAP004461
AGAP004462
AGAP004463
mean_tpm
time
preZGA
ZGA_early
ZGA_late
5h
9h
11h
14h
20h
24h
L1
sex
Female
Male

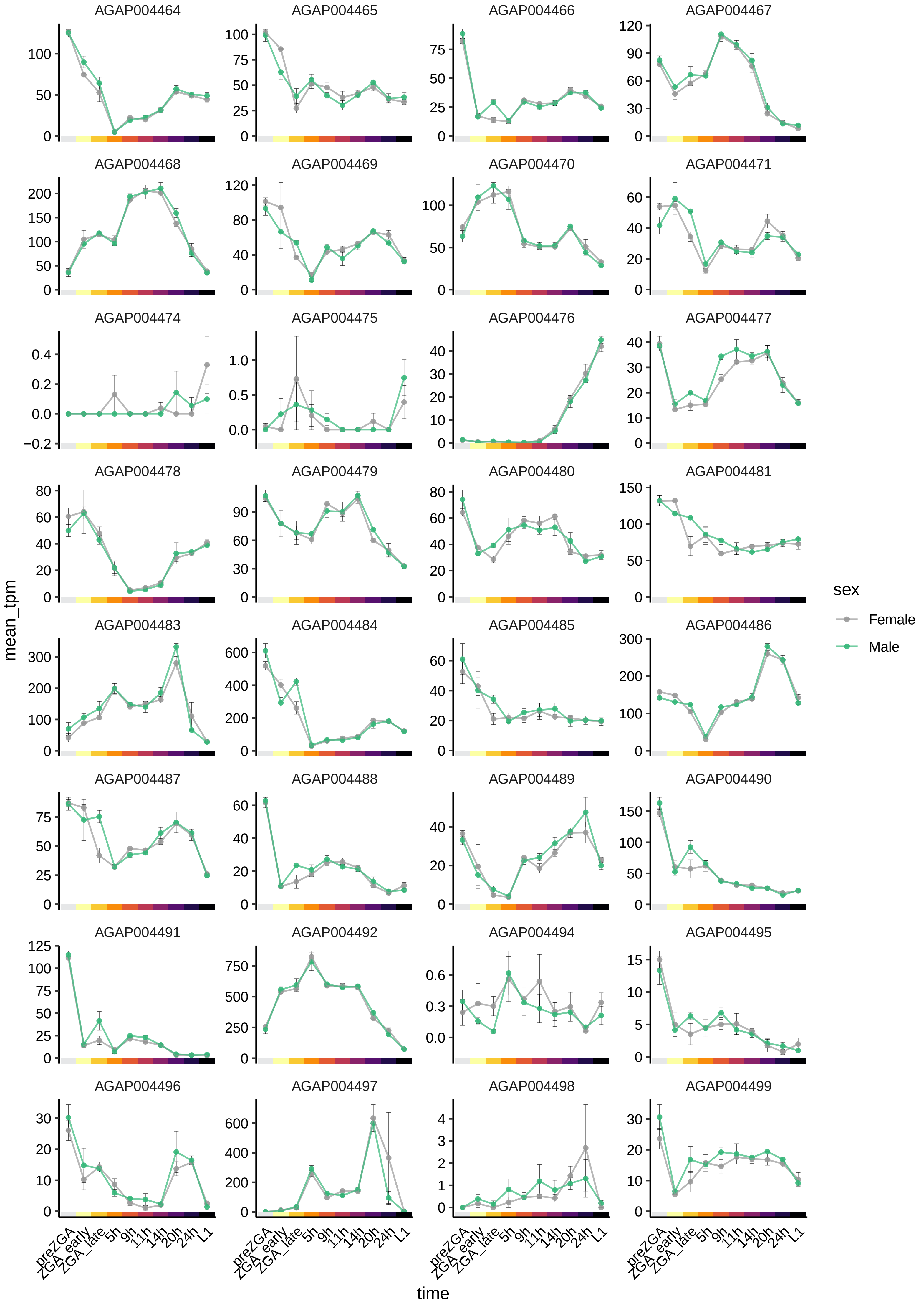
AGAP004464
AGAP004465
AGAP004466
AGAP004467
AGAP004468
AGAP004469
AGAP004470
AGAP004471
AGAP004474
AGAP004475
AGAP004476
AGAP004477
AGAP004478
AGAP004479
AGAP004480
AGAP004481
AGAP004483
AGAP004484
AGAP004485
AGAP004486
AGAP004487
AGAP004488
AGAP004489
AGAP004490
AGAP004491
AGAP004492
AGAP004494
AGAP004495
AGAP004496
AGAP004497
AGAP004498
AGAP004499
mean_tpm
time
preZGA
ZGA_early
ZGA_late
5h
9h
11h
14h
20h
24h
L1
sex
Female
Male

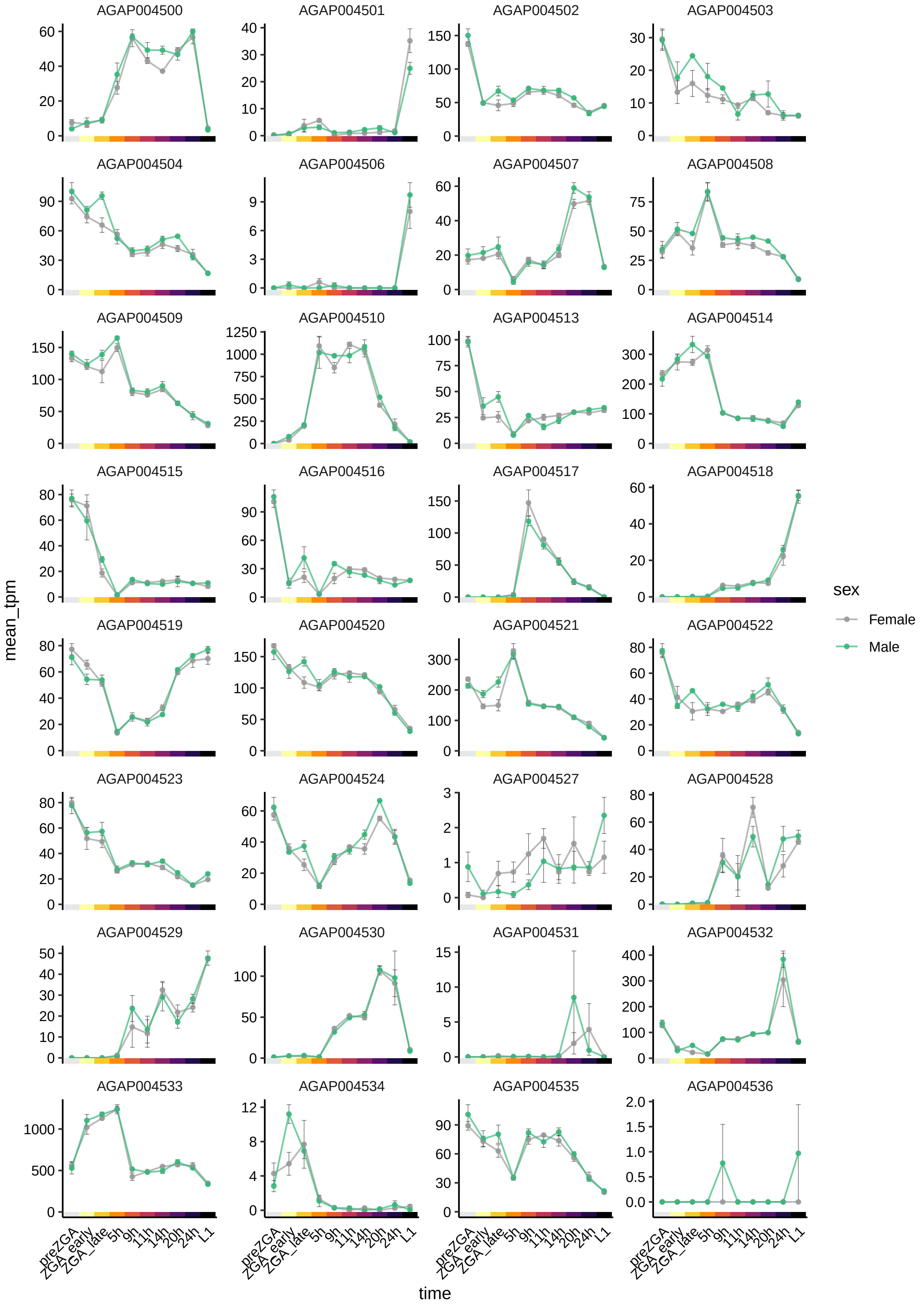

AGAP004500
AGAP004501
AGAP004502
AGAP004503
AGAP004504
AGAP004506
AGAP004507
AGAP004508
AGAP004509
AGAP004510
AGAP004513
AGAP004514
AGAP004515
AGAP004516
AGAP004517
AGAP004518
AGAP004519
AGAP004520
AGAP004521
AGAP004522
AGAP004523
AGAP004524
AGAP004527
AGAP004528
AGAP004529
AGAP004530
AGAP004531
AGAP004532
AGAP004533
AGAP004534
AGAP004535
AGAP004536
mean_tpm
time
preZGA
ZGA_early
ZGA_late
5h
9h
11h
14h
20h
24h
L1
sex
Female
Male

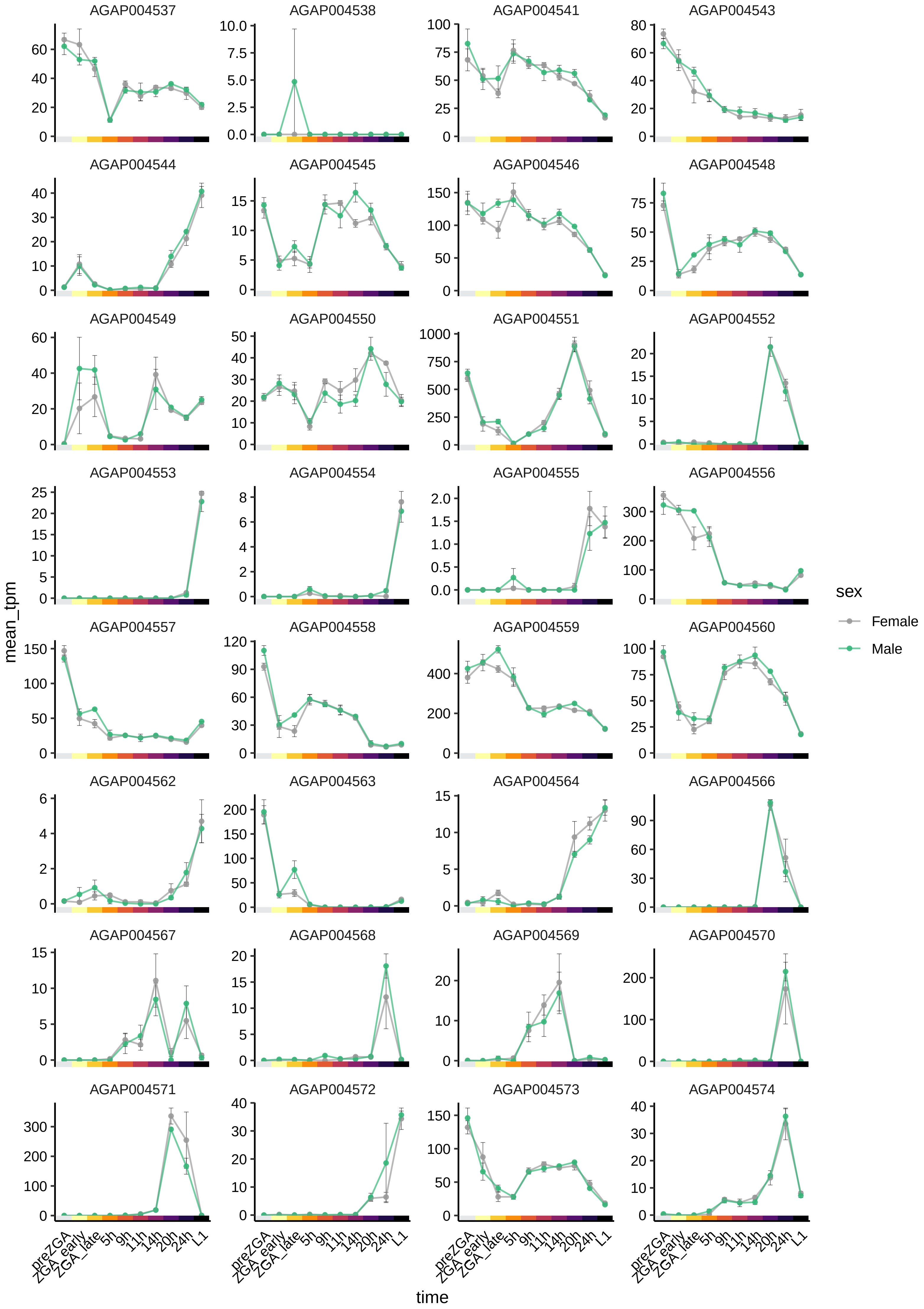
AGAP004537
AGAP004538
AGAP004541
AGAP004543
AGAP004544
AGAP004545
AGAP004546
AGAP004548
AGAP004549
AGAP004550
AGAP004551
AGAP004552
AGAP004553
AGAP004554
AGAP004555
AGAP004556
AGAP004557
AGAP004558
AGAP004559
AGAP004560
AGAP004562
AGAP004563
AGAP004564
AGAP004566
AGAP004567
AGAP004568
AGAP004569
AGAP004570
AGAP004571
AGAP004572
AGAP004573
AGAP004574
mean_tpm
time
preZGA
ZGA_early
ZGA_late
5h
9h
11h
14h
20h
24h
L1
sex
Female
Male

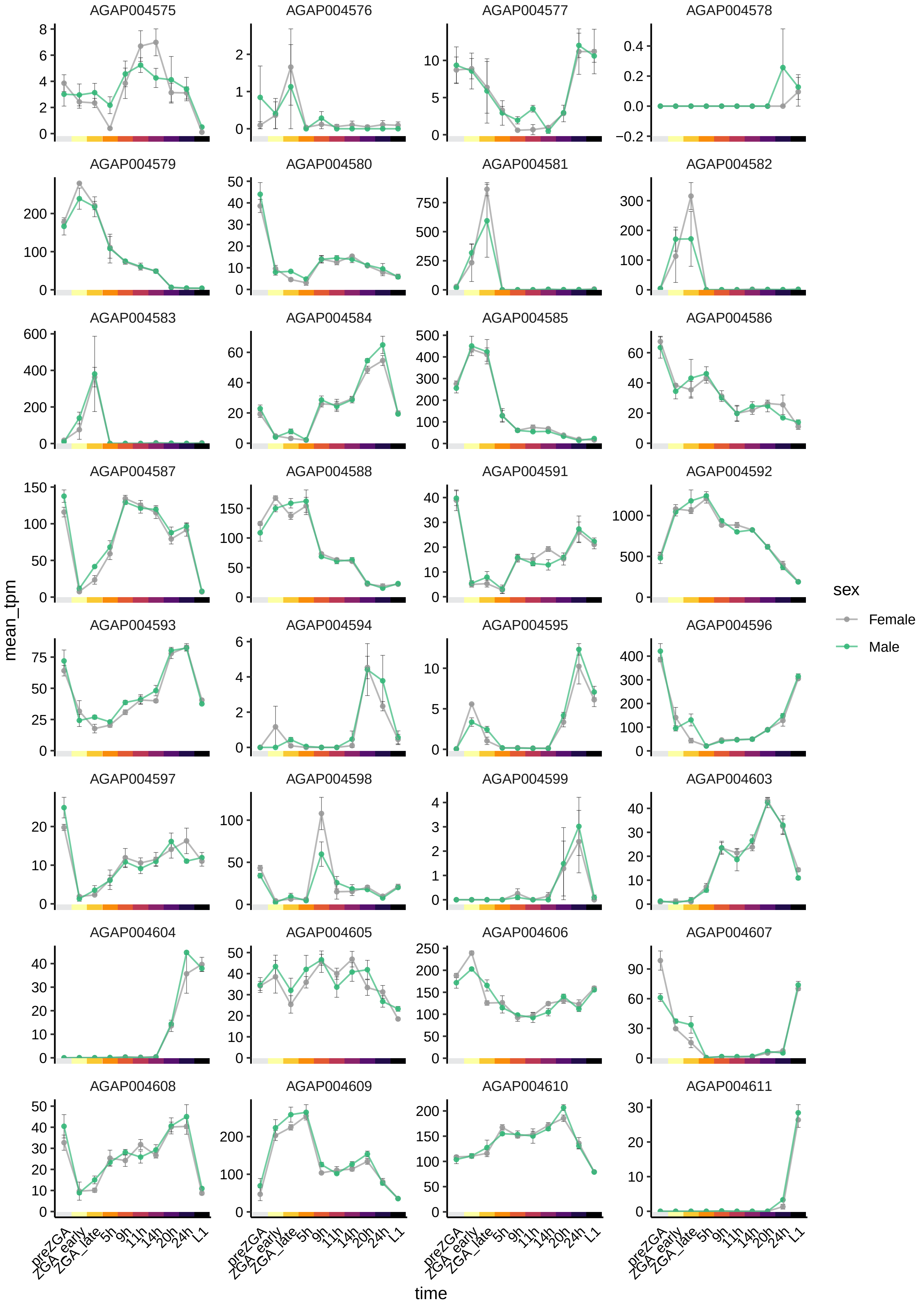

AGAP004575
AGAP004576
AGAP004577
AGAP004578
AGAP004579
AGAP004580
AGAP004581
AGAP004582
AGAP004583
AGAP004584
AGAP004585
AGAP004586
AGAP004587
AGAP004588
AGAP004591
AGAP004592
AGAP004593
AGAP004594
AGAP004595
AGAP004596
AGAP004597
AGAP004598
AGAP004599
AGAP004603
AGAP004604
AGAP004605
AGAP004606
AGAP004607
AGAP004608
AGAP004609
AGAP004610
AGAP004611
mean_tpm
time
preZGA
ZGA_early
ZGA_late
5h
9h
11h
14h
20h
24h
L1
sex
Female
Male

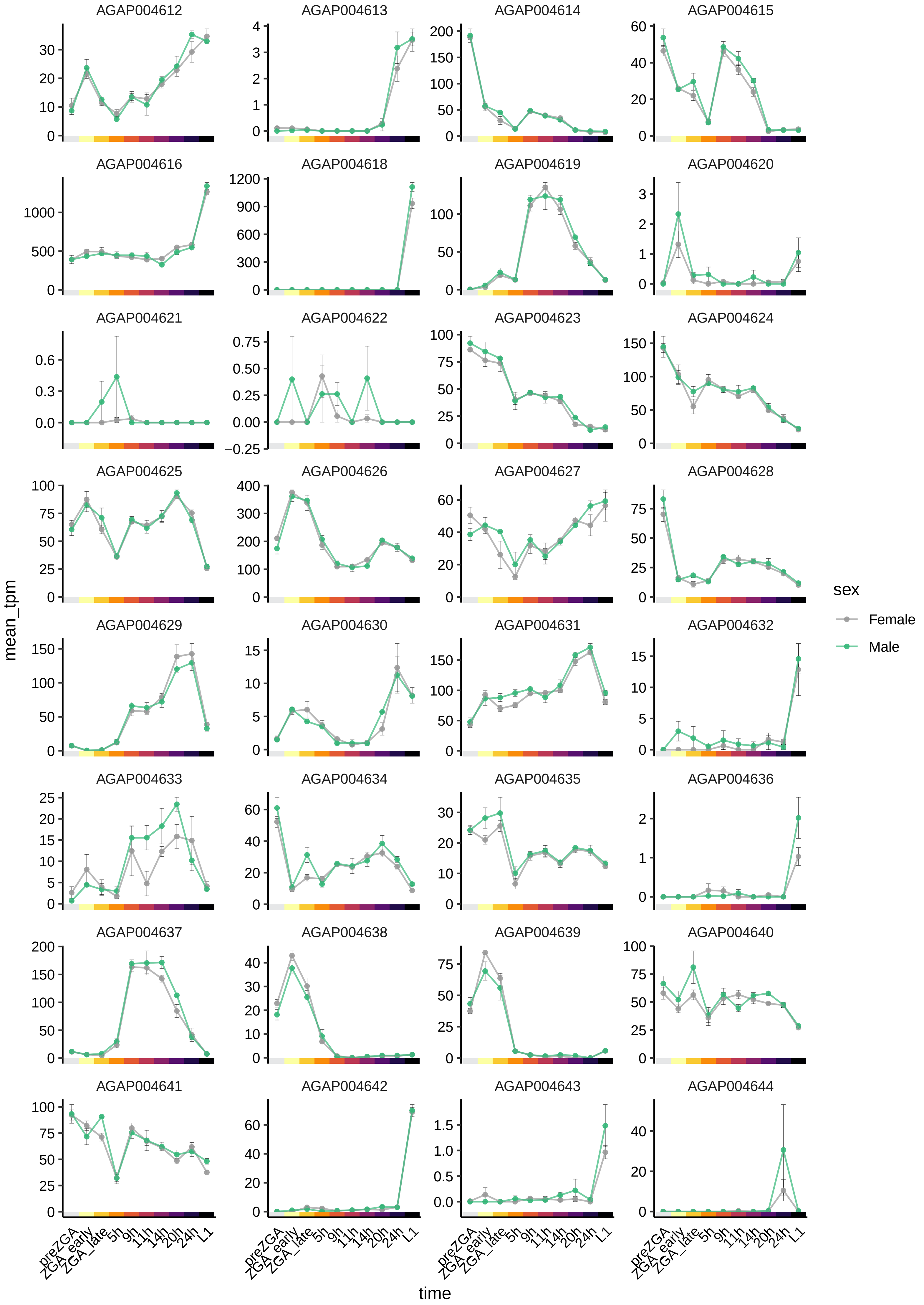

AGAP004612
AGAP004613
AGAP004614
AGAP004615
AGAP004616
AGAP004618
AGAP004619
AGAP004620
AGAP004621
AGAP004622
AGAP004623
AGAP004624
AGAP004625
AGAP004626
AGAP004627
AGAP004628
AGAP004629
AGAP004630
AGAP004631
AGAP004632
AGAP004633
AGAP004634
AGAP004635
AGAP004636
AGAP004637
AGAP004638
AGAP004639
AGAP004640
AGAP004641
AGAP004642
AGAP004643
AGAP004644
mean_tpm
time
preZGA
ZGA_early
ZGA_late
5h
9h
11h
14h
20h
24h
L1
sex
Female
Male

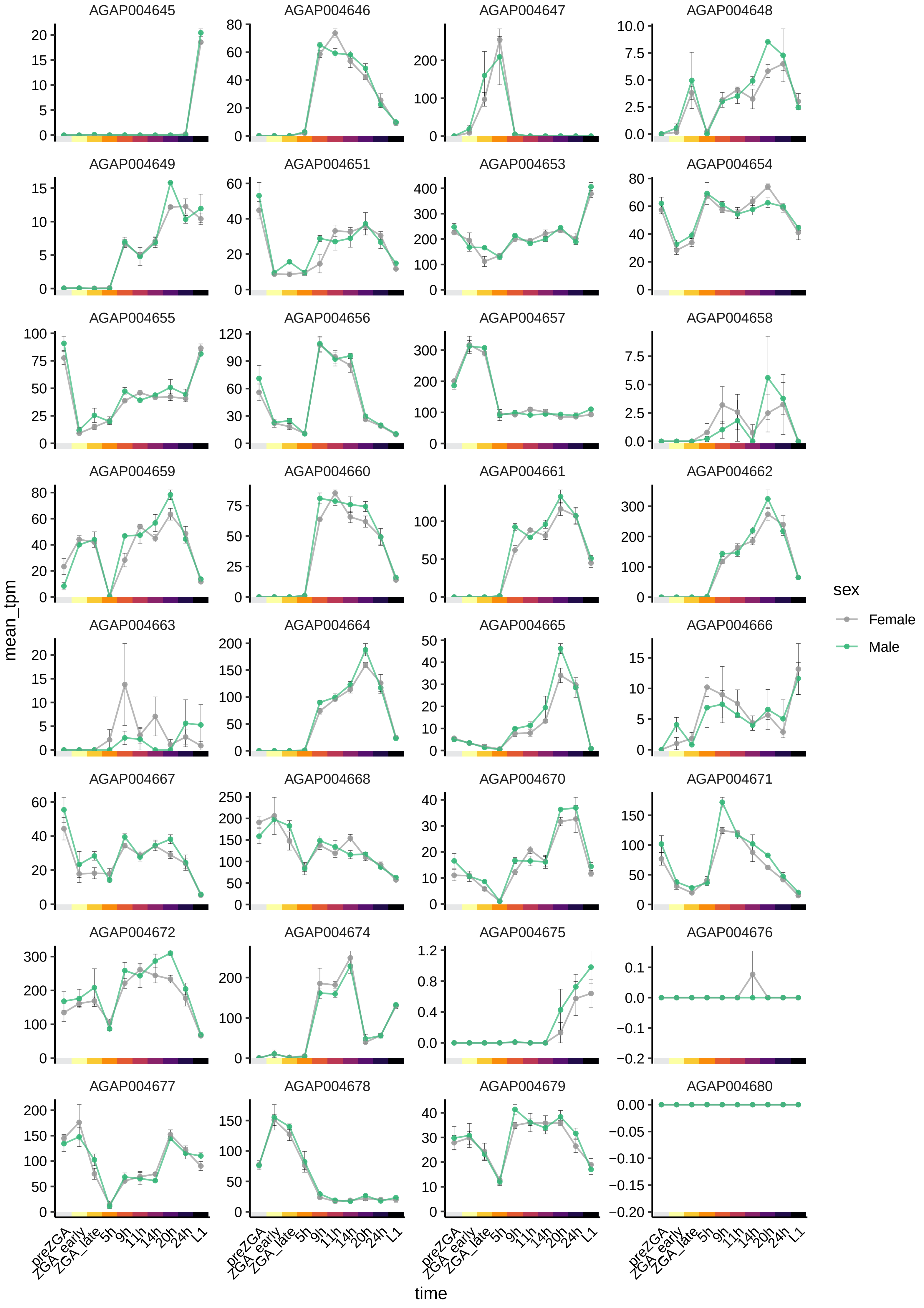
AGAP004645
AGAP004646
AGAP004647
AGAP004648
AGAP004649
AGAP004651
AGAP004653
AGAP004654
AGAP004655
AGAP004656
AGAP004657
AGAP004658
AGAP004659
AGAP004660
AGAP004661
AGAP004662
AGAP004663
AGAP004664
AGAP004665
AGAP004666
AGAP004667
AGAP004668
AGAP004670
AGAP004671
AGAP004672
AGAP004674
AGAP004675
AGAP004676
AGAP004677
AGAP004678
AGAP004679
AGAP004680
mean_tpm
time
preZGA
ZGA_early
ZGA_late
5h
9h
11h
14h
20h
24h
L1
sex
Female
Male

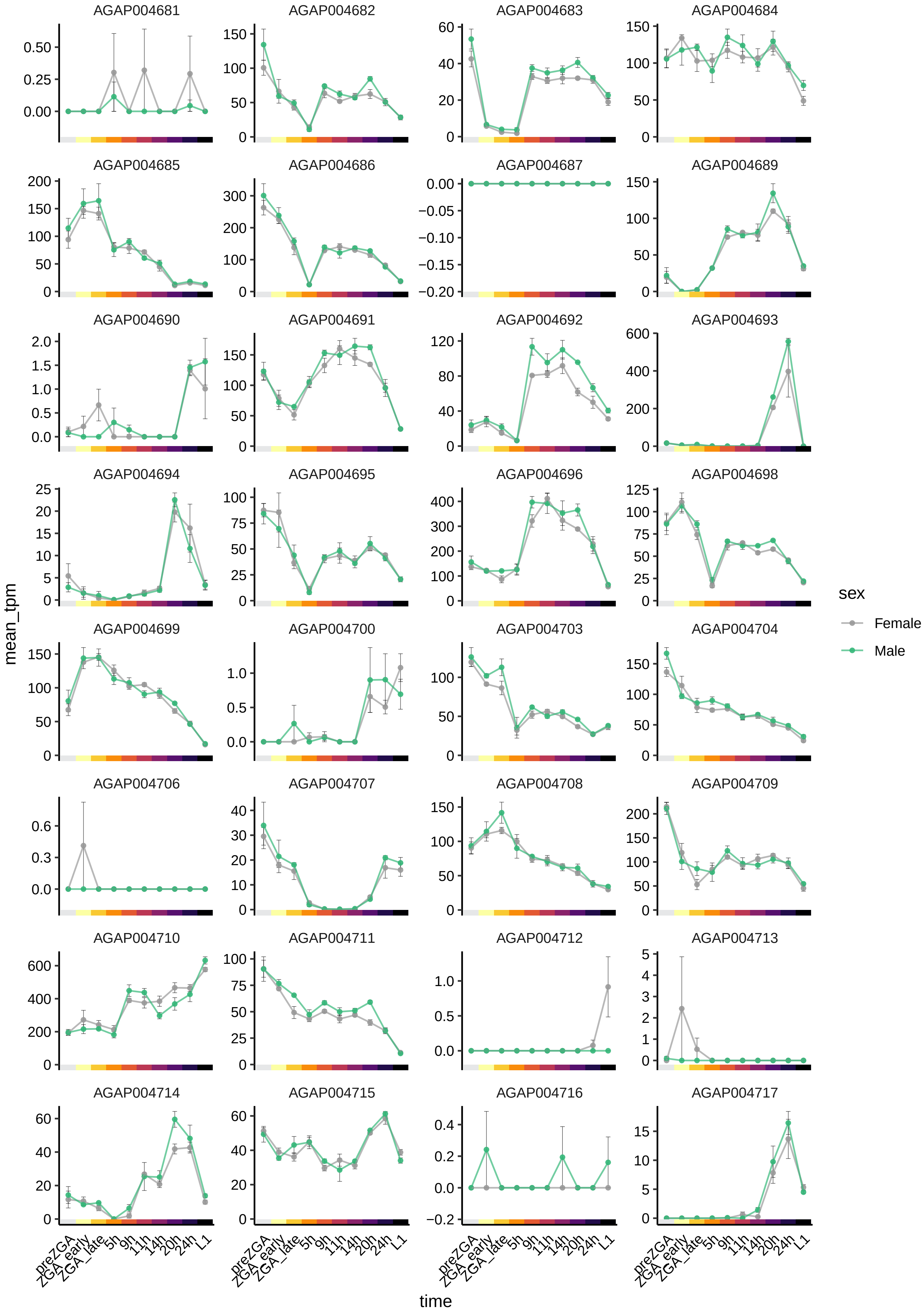
AGAP004681
AGAP004682
AGAP004683
AGAP004684
AGAP004685
AGAP004686
AGAP004687
AGAP004689
AGAP004690
AGAP004691
AGAP004692
AGAP004693
AGAP004694
AGAP004695
AGAP004696
AGAP004698
AGAP004699
AGAP004700
AGAP004703
AGAP004704
AGAP004706
AGAP004707
AGAP004708
AGAP004709
AGAP004710
AGAP004711
AGAP004712
AGAP004713
AGAP004714
AGAP004715
AGAP004716
AGAP004717
mean_tpm
time
preZGA
ZGA_early
ZGA_late
5h
9h
11h
14h
20h
24h
L1
sex
Female
Male

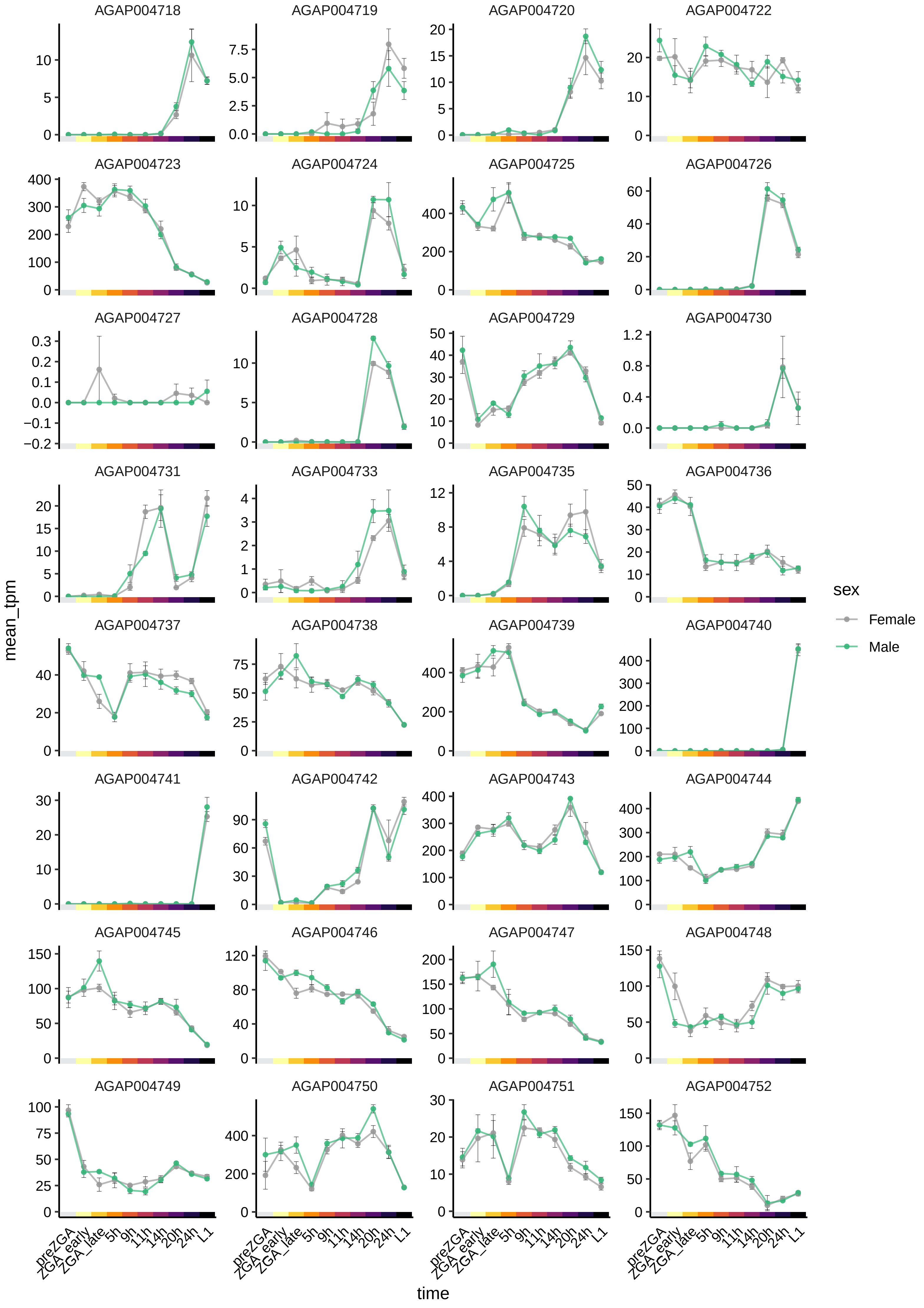

AGAP004718
AGAP004719
AGAP004720
AGAP004722
AGAP004723
AGAP004724
AGAP004725
AGAP004726
AGAP004727
AGAP004728
AGAP004729
AGAP004730
AGAP004731
AGAP004733
AGAP004735
AGAP004736
AGAP004737
AGAP004738
AGAP004739
AGAP004740
AGAP004741
AGAP004742
AGAP004743
AGAP004744
AGAP004745
AGAP004746
AGAP004747
AGAP004748
AGAP004749
AGAP004750
AGAP004751
AGAP004752
mean_tpm
time
preZGA
ZGA_early
ZGA_late
5h
9h
11h
14h
20h
24h
L1
sex
Female
Male

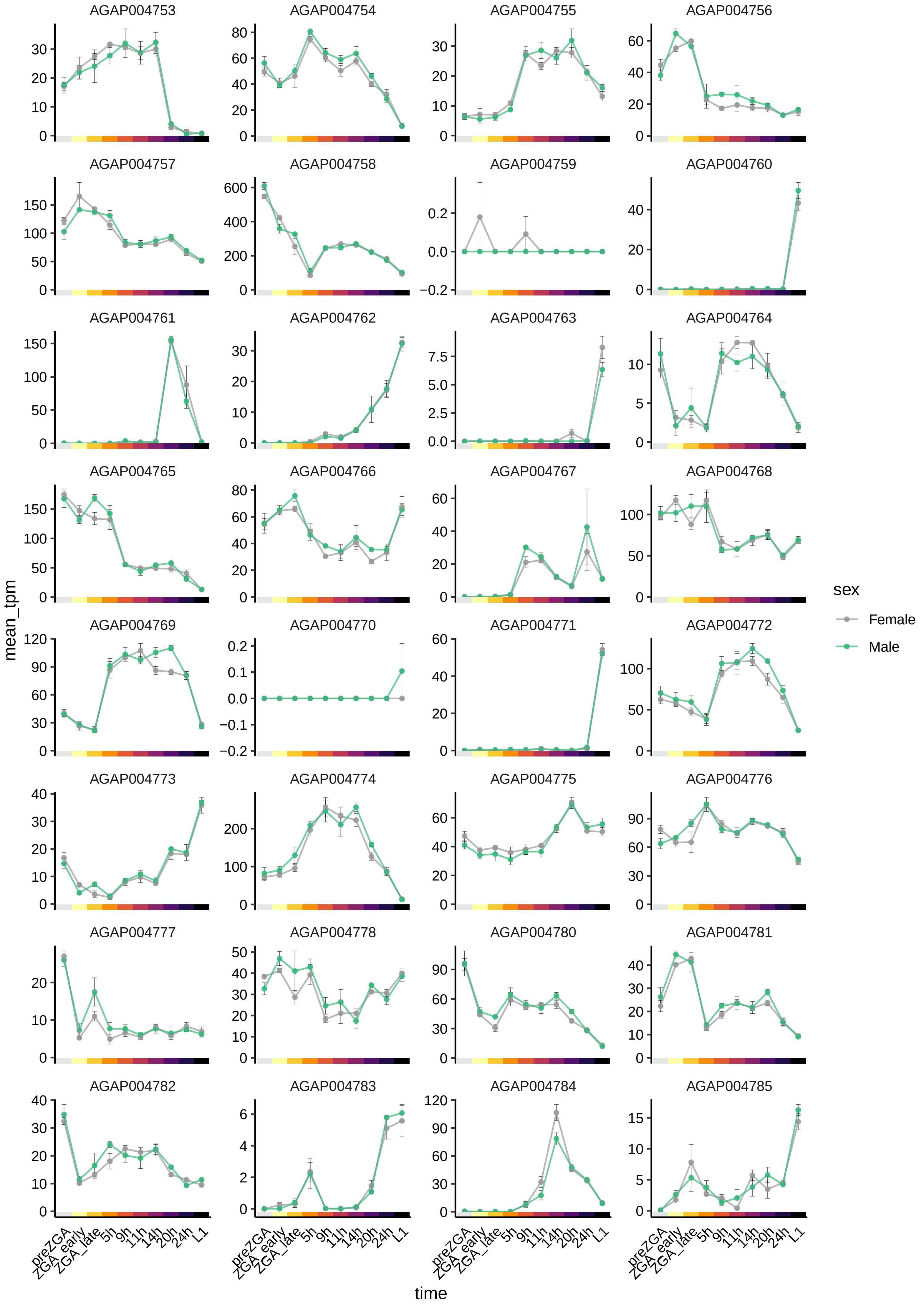
AGAP004753
AGAP004754
AGAP004755
AGAP004756
AGAP004757
AGAP004758
AGAP004759
AGAP004760
AGAP004761
AGAP004762
AGAP004763
AGAP004764
AGAP004765
AGAP004766
AGAP004767
AGAP004768
AGAP004769
AGAP004770
AGAP004771
AGAP004772
AGAP004773
AGAP004774
AGAP004775
AGAP004776
AGAP004777
AGAP004778
AGAP004780
AGAP004781
AGAP004782
AGAP004783
AGAP004784
AGAP004785
mean_tpm
time
preZGA
ZGA_early
ZGA_late
5h
9h
11h
14h
20h
24h
L1
sex
Female
Male

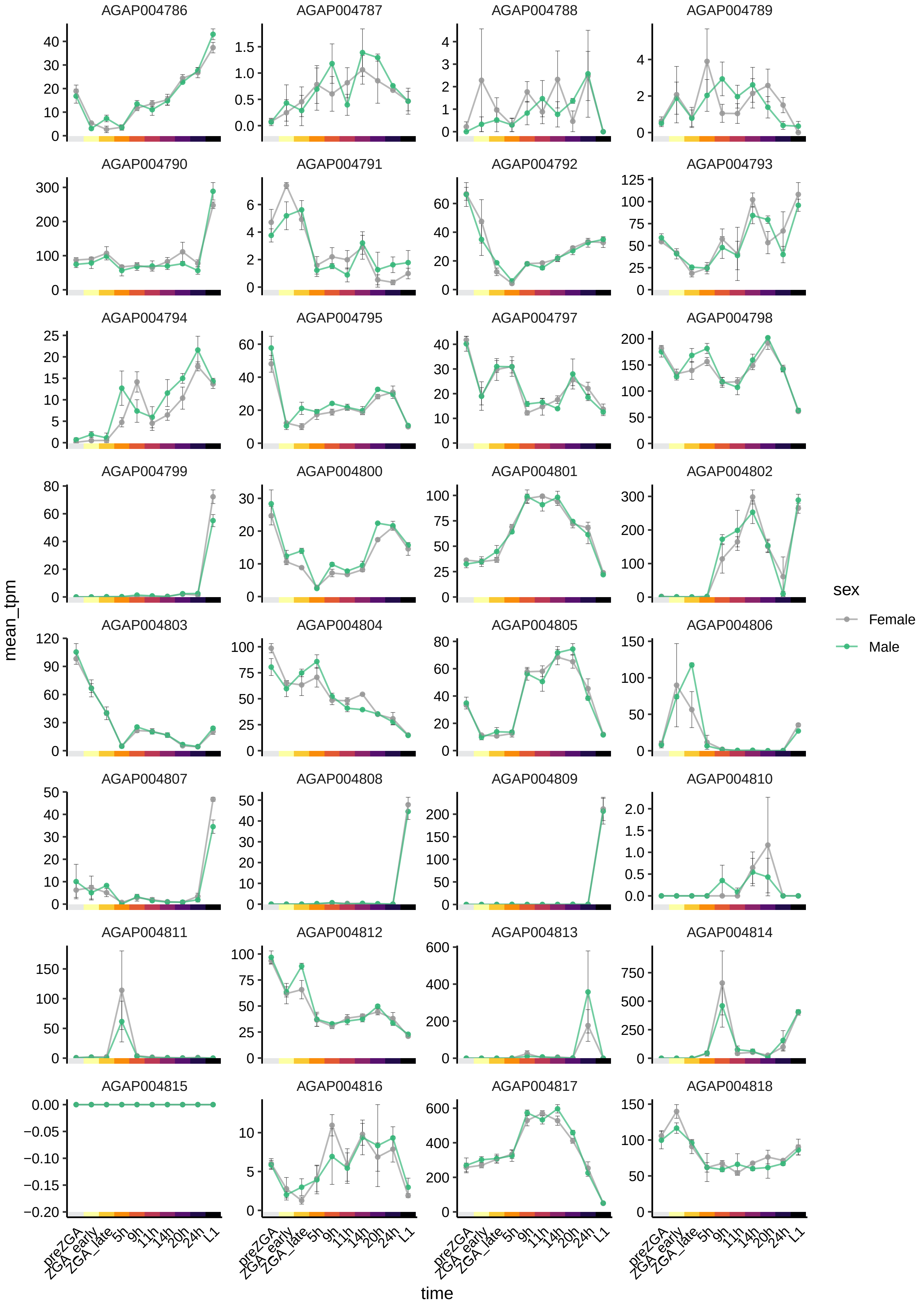

AGAP004786
AGAP004787
AGAP004788
AGAP004789
AGAP004790
AGAP004791
AGAP004792
AGAP004793
AGAP004794
AGAP004795
AGAP004797
AGAP004798
AGAP004799
AGAP004800
AGAP004801
AGAP004802
AGAP004803
AGAP004804
AGAP004805
AGAP004806
AGAP004807
AGAP004808
AGAP004809
AGAP004810
AGAP004811
AGAP004812
AGAP004813
AGAP004814
AGAP004815
AGAP004816
AGAP004817
AGAP004818
mean_tpm
time
preZGA
ZGA_early
ZGA_late
5h
9h
11h
14h
20h
24h
L1
sex
Female
Male

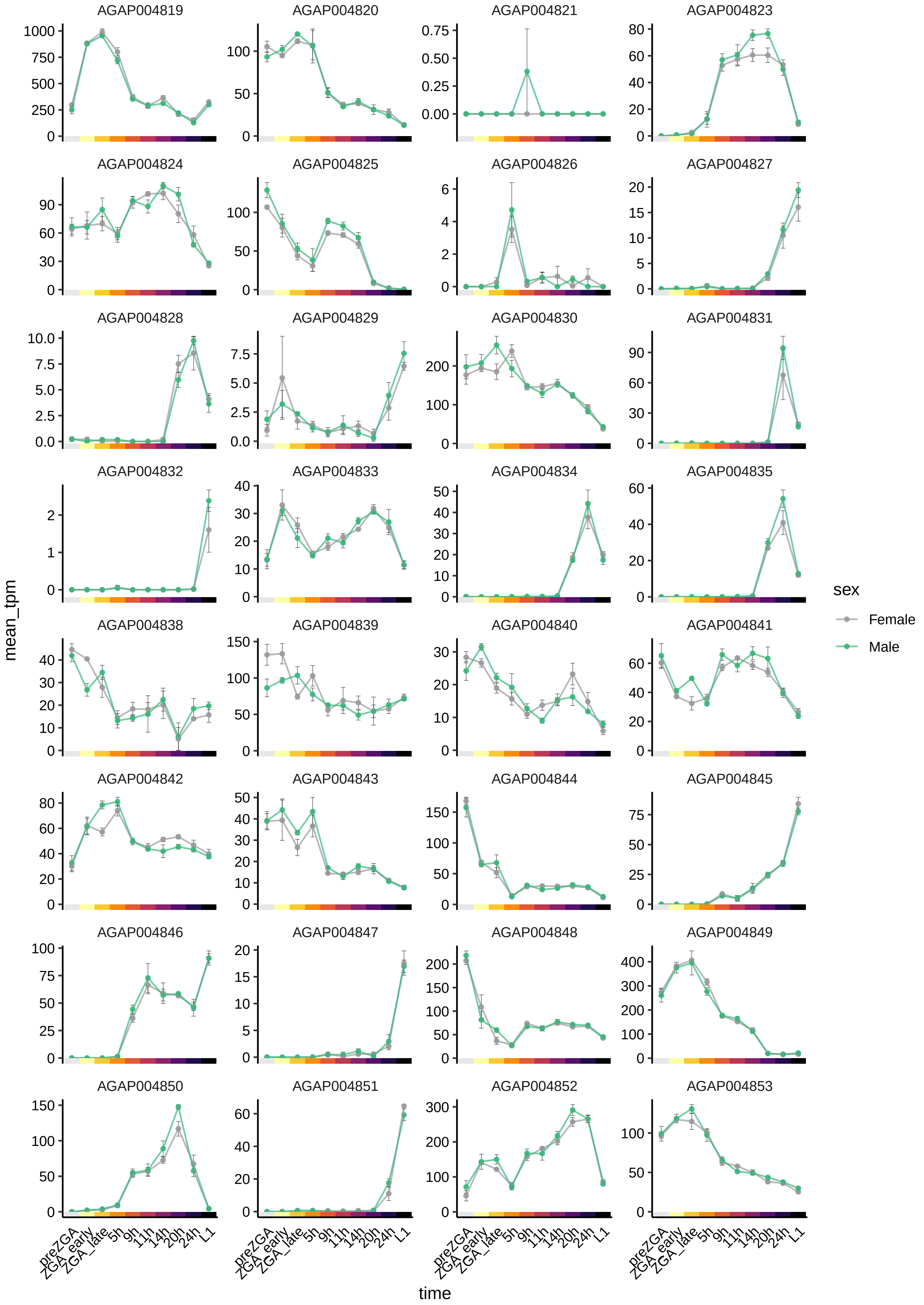

AGAP004819
AGAP004820
AGAP004821
AGAP004823
AGAP004824
AGAP004825
AGAP004826
AGAP004827
AGAP004828
AGAP004829
AGAP004830
AGAP004831
AGAP004832
AGAP004833
AGAP004834
AGAP004835
AGAP004838
AGAP004839
AGAP004840
AGAP004841
AGAP004842
AGAP004843
AGAP004844
AGAP004845
AGAP004846
AGAP004847
AGAP004848
AGAP004849
AGAP004850
AGAP004851
AGAP004852
AGAP004853
mean_tpm
time
preZGA
ZGA_early
ZGA_late
5h
9h
11h
14h
20h
24h
L1
sex
Female
Male

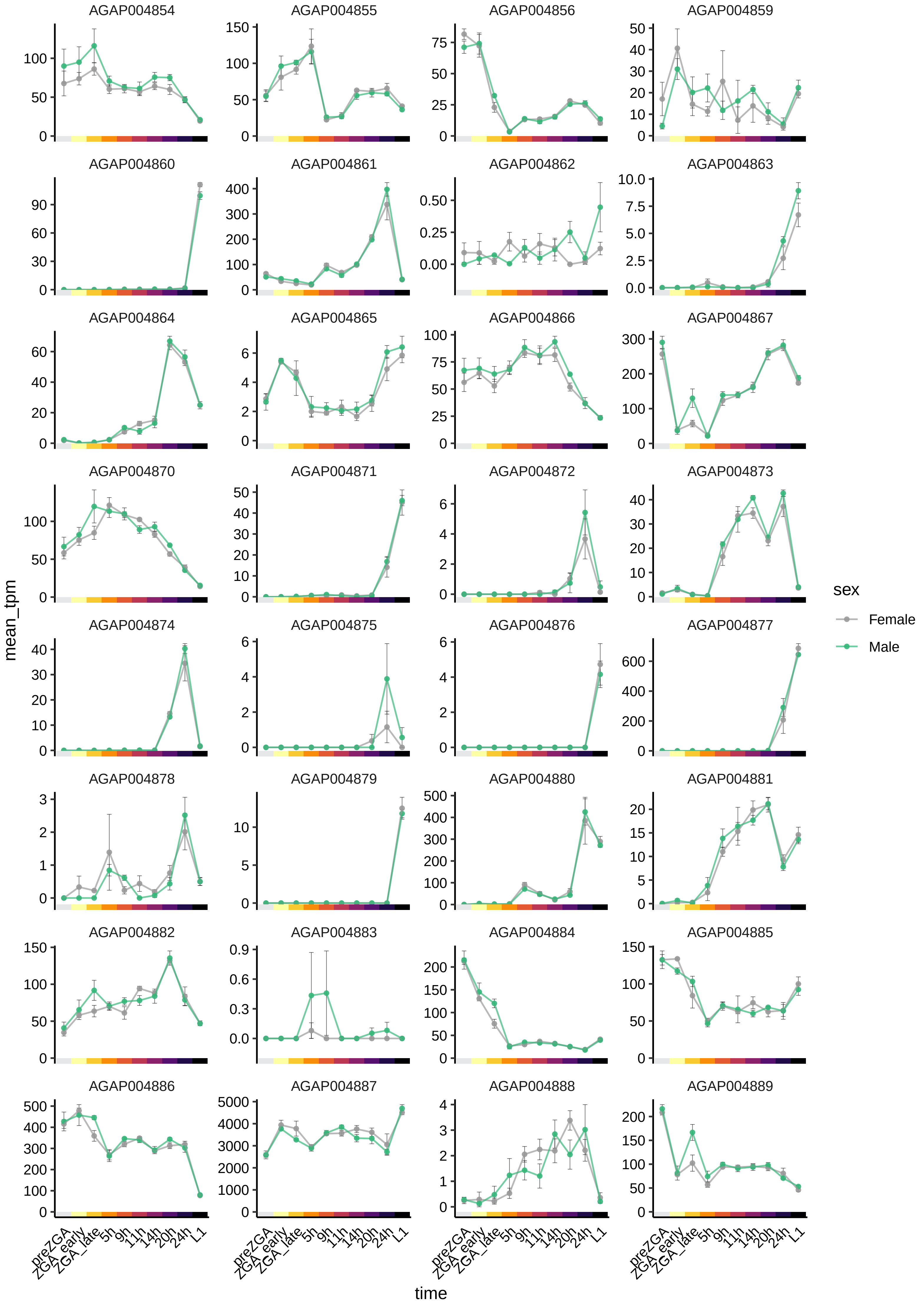
AGAP004854
AGAP004855
AGAP004856
AGAP004859
AGAP004860
AGAP004861
AGAP004862
AGAP004863
AGAP004864
AGAP004865
AGAP004866
AGAP004867
AGAP004870
AGAP004871
AGAP004872
AGAP004873
AGAP004874
AGAP004875
AGAP004876
AGAP004877
AGAP004878
AGAP004879
AGAP004880
AGAP004881
AGAP004882
AGAP004883
AGAP004884
AGAP004885
AGAP004886
AGAP004887
AGAP004888
AGAP004889
mean_tpm
time
preZGA
ZGA_early
ZGA_late
5h
9h
11h
14h
20h
24h
L1
sex
Female
Male

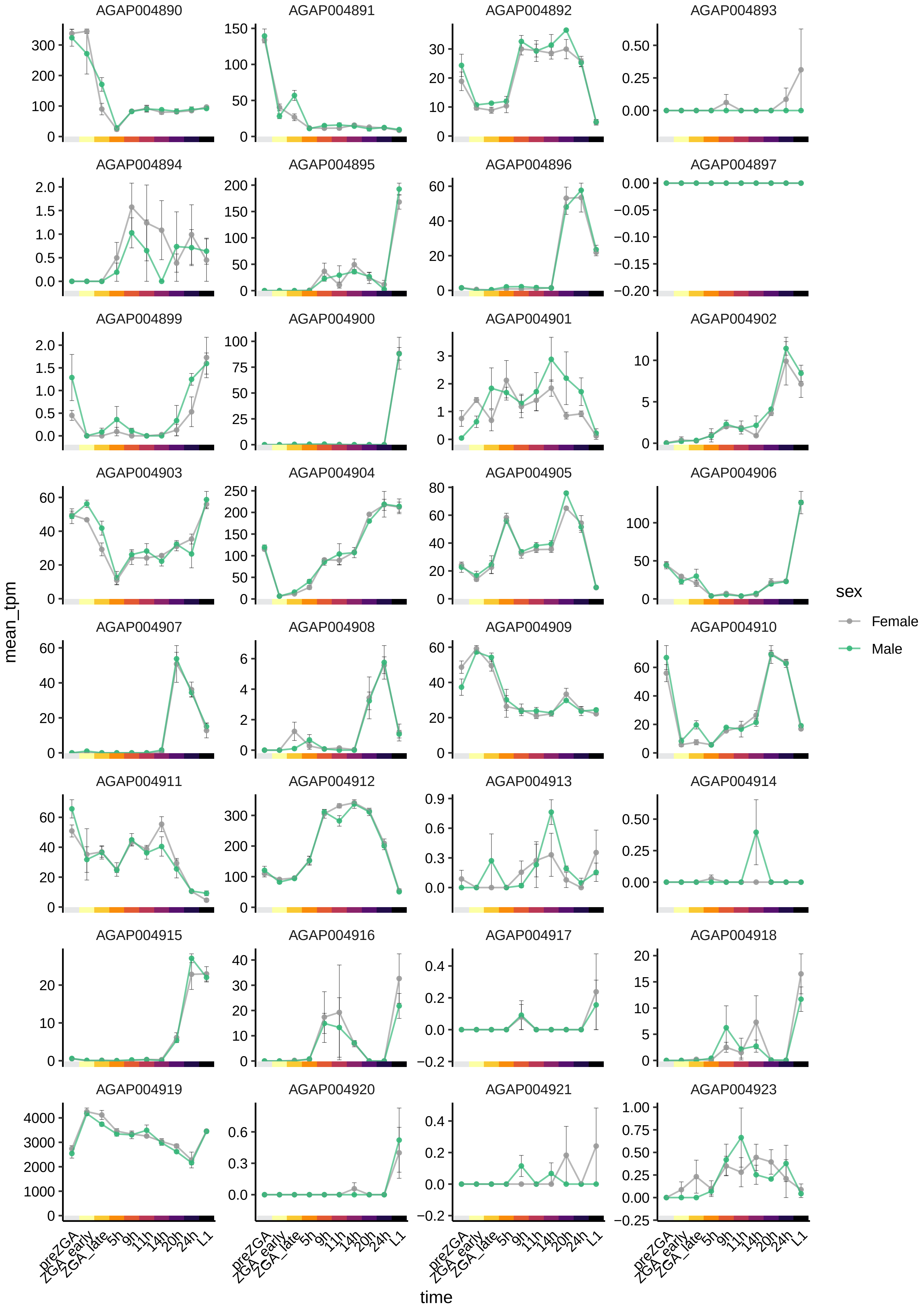

AGAP004890
AGAP004891
AGAP004892
AGAP004893
AGAP004894
AGAP004895
AGAP004896
AGAP004897
AGAP004899
AGAP004900
AGAP004901
AGAP004902
AGAP004903
AGAP004904
AGAP004905
AGAP004906
AGAP004907
AGAP004908
AGAP004909
AGAP004910
AGAP004911
AGAP004912
AGAP004913
AGAP004914
AGAP004915
AGAP004916
AGAP004917
AGAP004918
AGAP004919
AGAP004920
AGAP004921
AGAP004923
mean_tpm
time
preZGA
ZGA_early
ZGA_late
5h
9h
11h
14h
20h
24h
L1
sex
Female
Male

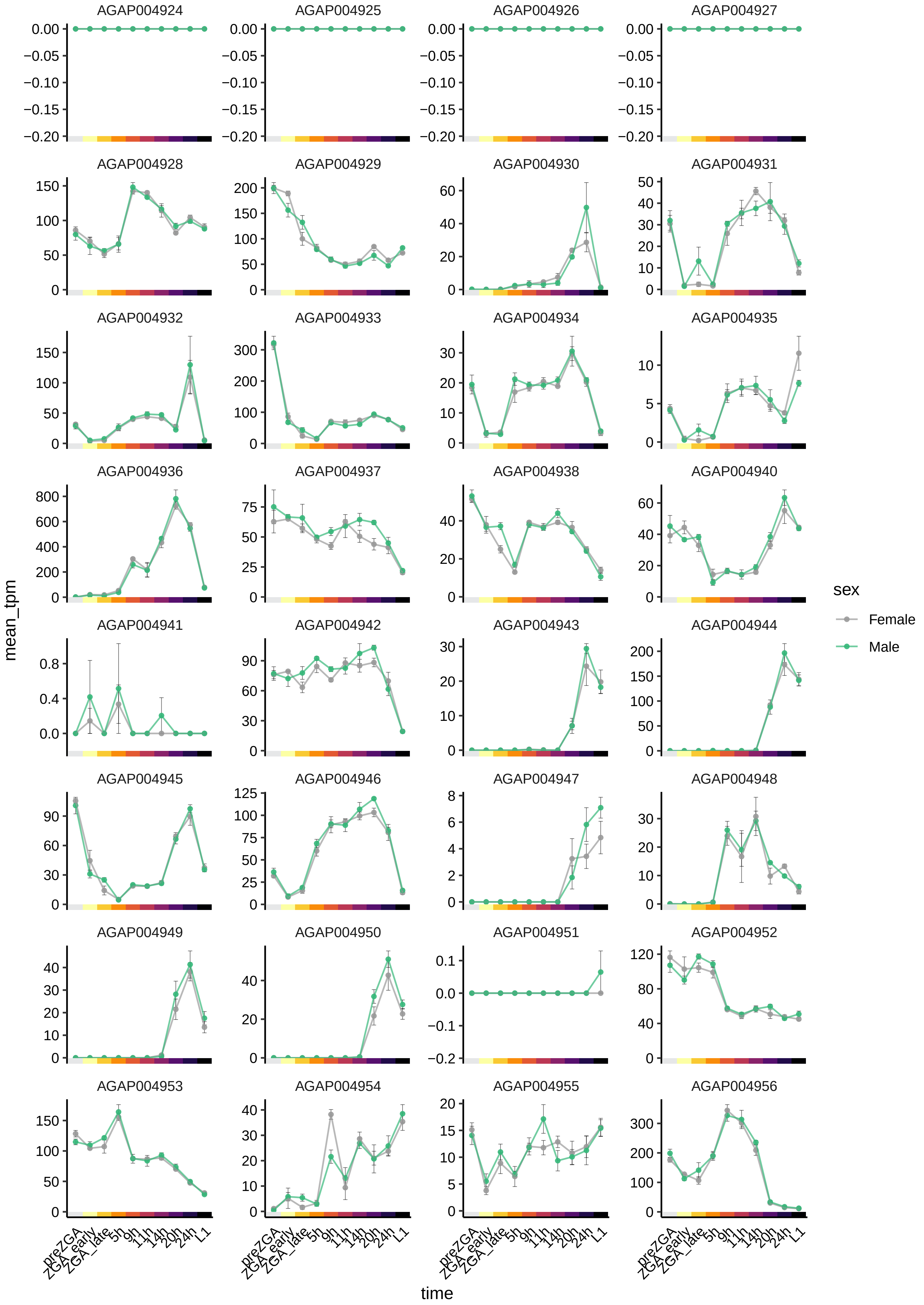
AGAP004924
AGAP004925
AGAP004926
AGAP004927
AGAP004928
AGAP004929
AGAP004930
AGAP004931
AGAP004932
AGAP004933
AGAP004934
AGAP004935
AGAP004936
AGAP004937
AGAP004938
AGAP004940
AGAP004941
AGAP004942
AGAP004943
AGAP004944
AGAP004945
AGAP004946
AGAP004947
AGAP004948
AGAP004949
AGAP004950
AGAP004951
AGAP004952
AGAP004953
AGAP004954
AGAP004955
AGAP004956
mean_tpm
time
preZGA
ZGA_early
ZGA_late
5h
9h
11h
14h
20h
24h
L1
sex
Female
Male

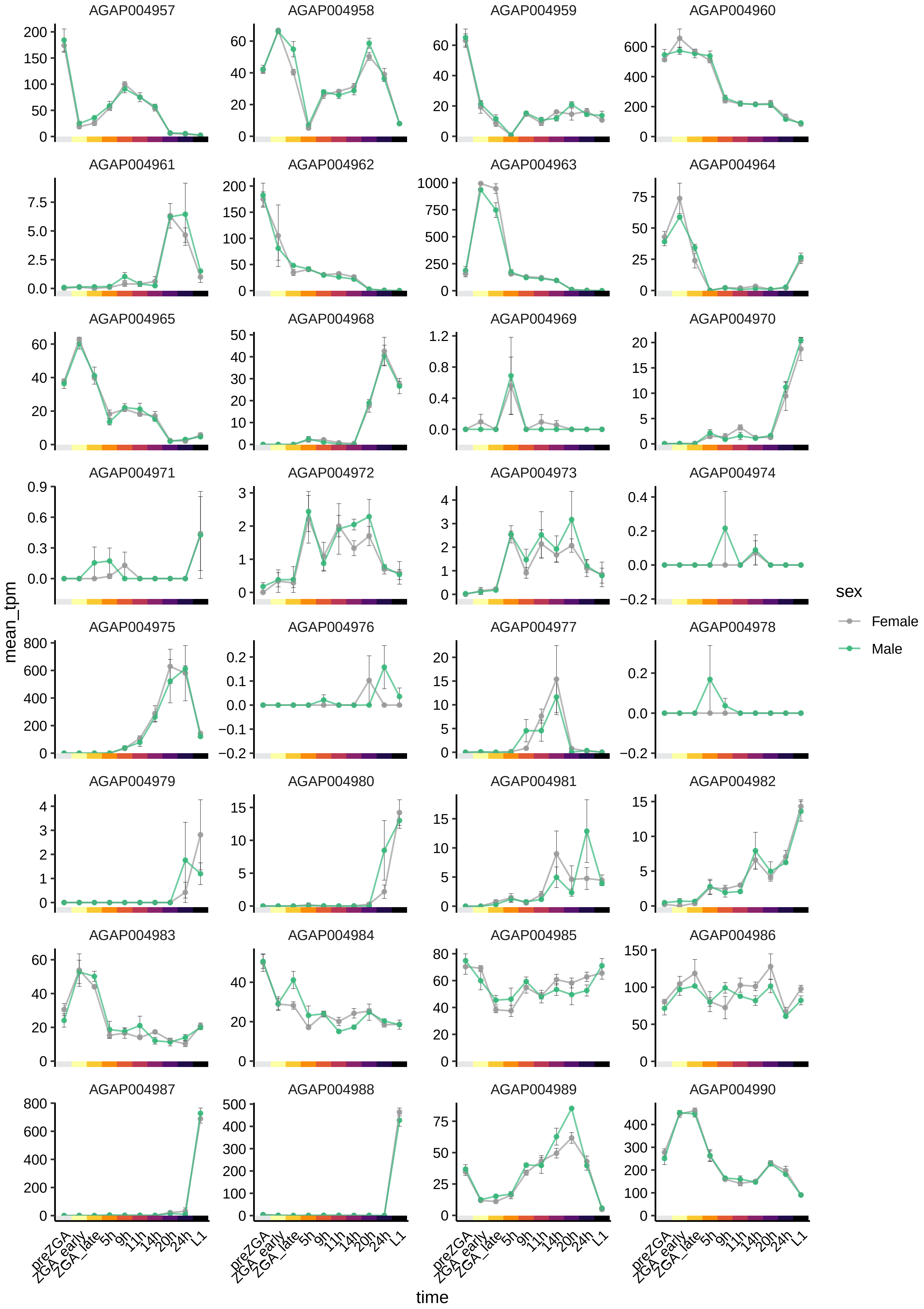

AGAP004957
AGAP004958
AGAP004959
AGAP004960
AGAP004961
AGAP004962
AGAP004963
AGAP004964
AGAP004965
AGAP004968
AGAP004969
AGAP004970
AGAP004971
AGAP004972
AGAP004973
AGAP004974
AGAP004975
AGAP004976
AGAP004977
AGAP004978
AGAP004979
AGAP004980
AGAP004981
AGAP004982
AGAP004983
AGAP004984
AGAP004985
AGAP004986
AGAP004987
AGAP004988
AGAP004989
AGAP004990
mean_tpm
time
preZGA
ZGA_early
ZGA_late
5h
9h
11h
14h
20h
24h
L1
sex
Female
Male

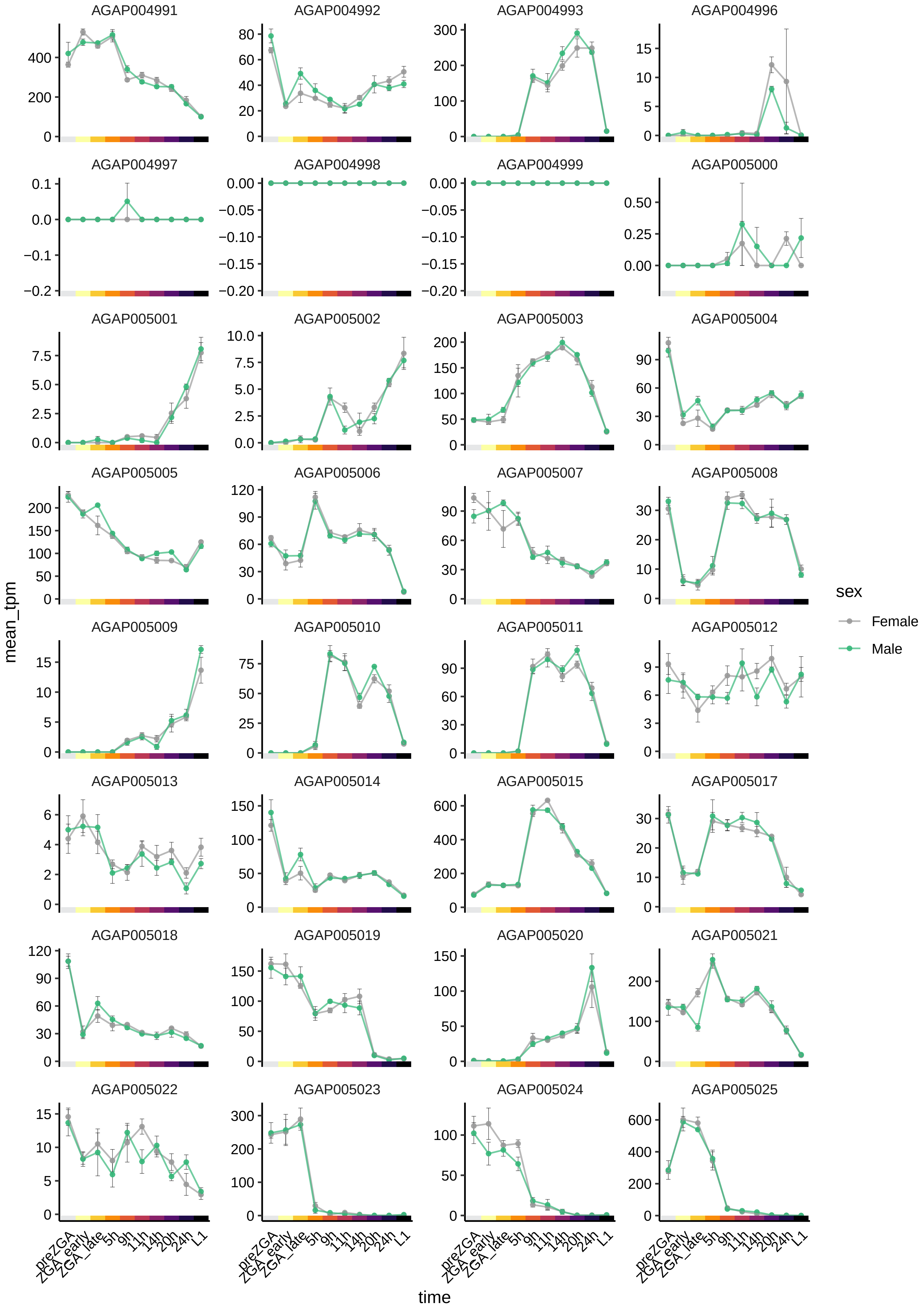

AGAP004991
AGAP004992
AGAP004993
AGAP004996
AGAP004997
AGAP004998
AGAP004999
AGAP005000
AGAP005001
AGAP005002
AGAP005003
AGAP005004
AGAP005005
AGAP005006
AGAP005007
AGAP005008
AGAP005009
AGAP005010
AGAP005011
AGAP005012
AGAP005013
AGAP005014
AGAP005015
AGAP005017
AGAP005018
AGAP005019
AGAP005020
AGAP005021
AGAP005022
AGAP005023
AGAP005024
AGAP005025
mean_tpm
time
preZGA
ZGA_early
ZGA_late
5h
9h
11h
14h
20h
24h
L1
sex
Female
Male

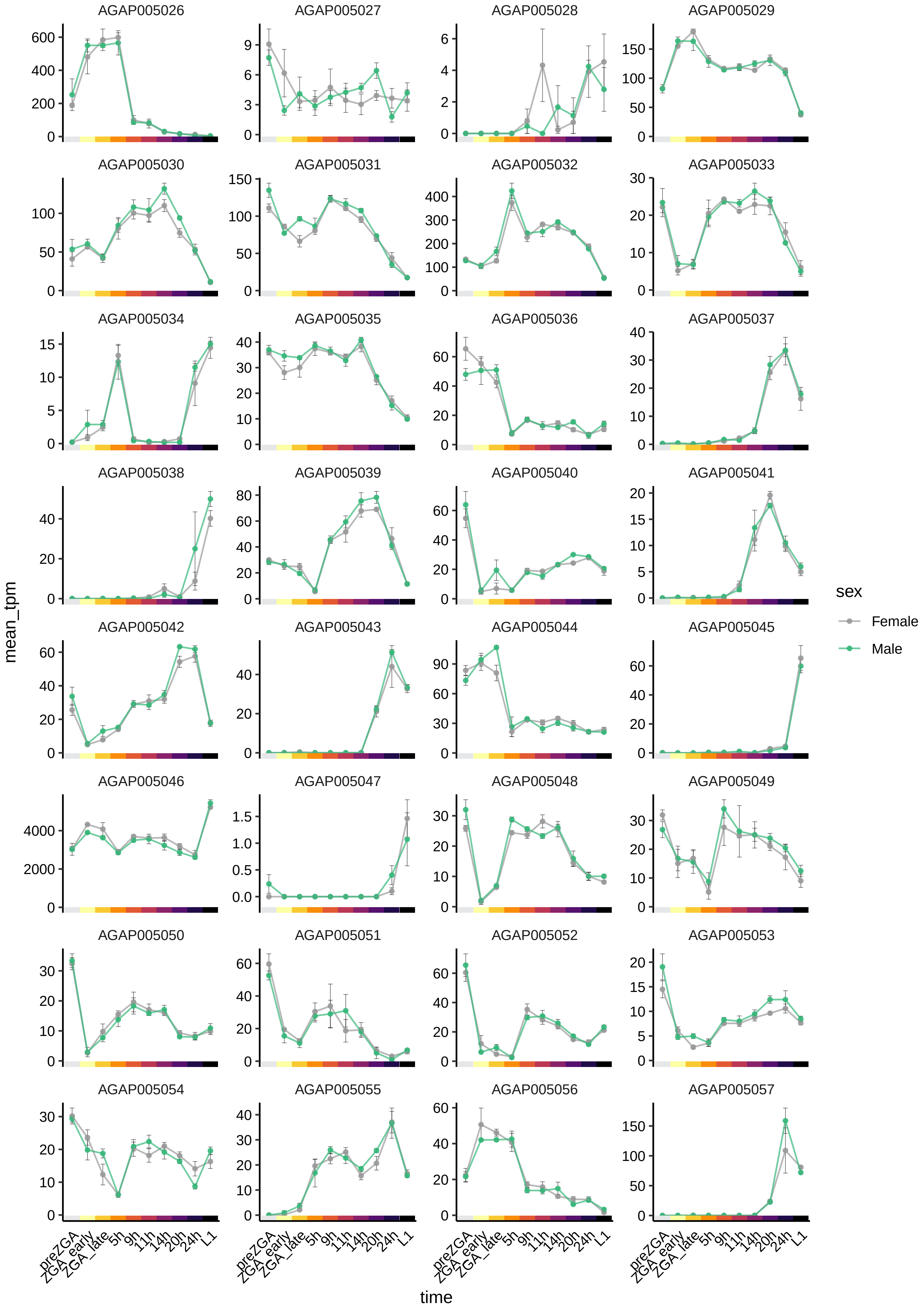
AGAP005026
AGAP005027
AGAP005028
AGAP005029
AGAP005030
AGAP005031
AGAP005032
AGAP005033
AGAP005034
AGAP005035
AGAP005036
AGAP005037
AGAP005038
AGAP005039
AGAP005040
AGAP005041
AGAP005042
AGAP005043
AGAP005044
AGAP005045
AGAP005046
AGAP005047
AGAP005048
AGAP005049
AGAP005050
AGAP005051
AGAP005052
AGAP005053
AGAP005054
AGAP005055
AGAP005056
AGAP005057
mean_tpm
time
preZGA
ZGA_early
ZGA_late
5h
9h
11h
14h
20h
24h
L1
sex
Female
Male

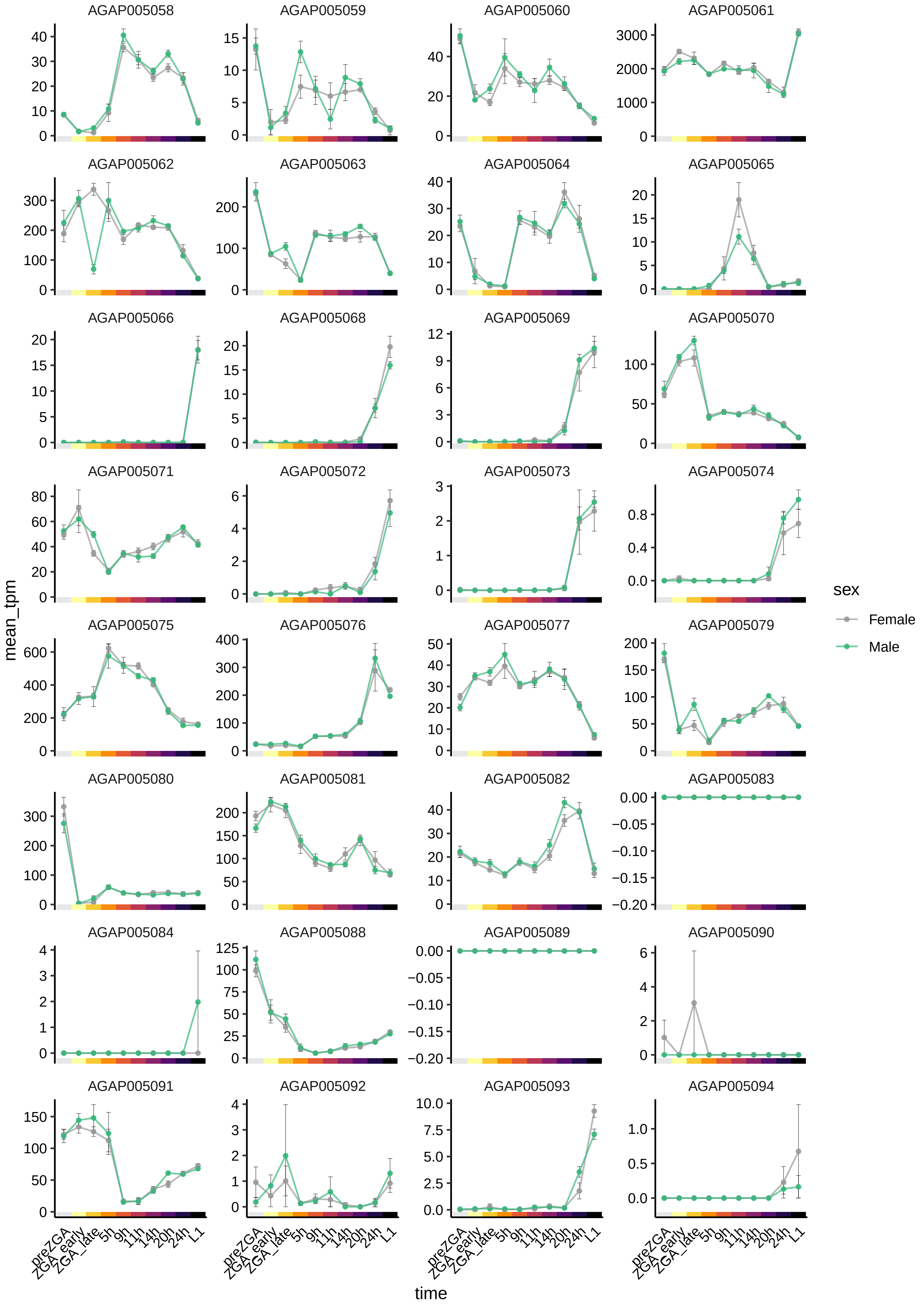

AGAP005058
AGAP005059
AGAP005060
AGAP005061
AGAP005062
AGAP005063
AGAP005064
AGAP005065
AGAP005066
AGAP005068
AGAP005069
AGAP005070
AGAP005071
AGAP005072
AGAP005073
AGAP005074
AGAP005075
AGAP005076
AGAP005077
AGAP005079
AGAP005080
AGAP005081
AGAP005082
AGAP005083
AGAP005084
AGAP005088
AGAP005089
AGAP005090
AGAP005091
AGAP005092
AGAP005093
AGAP005094
mean_tpm
time
preZGA
ZGA_early
ZGA_late
5h
9h
11h
14h
20h
24h
L1
sex
Female
Male

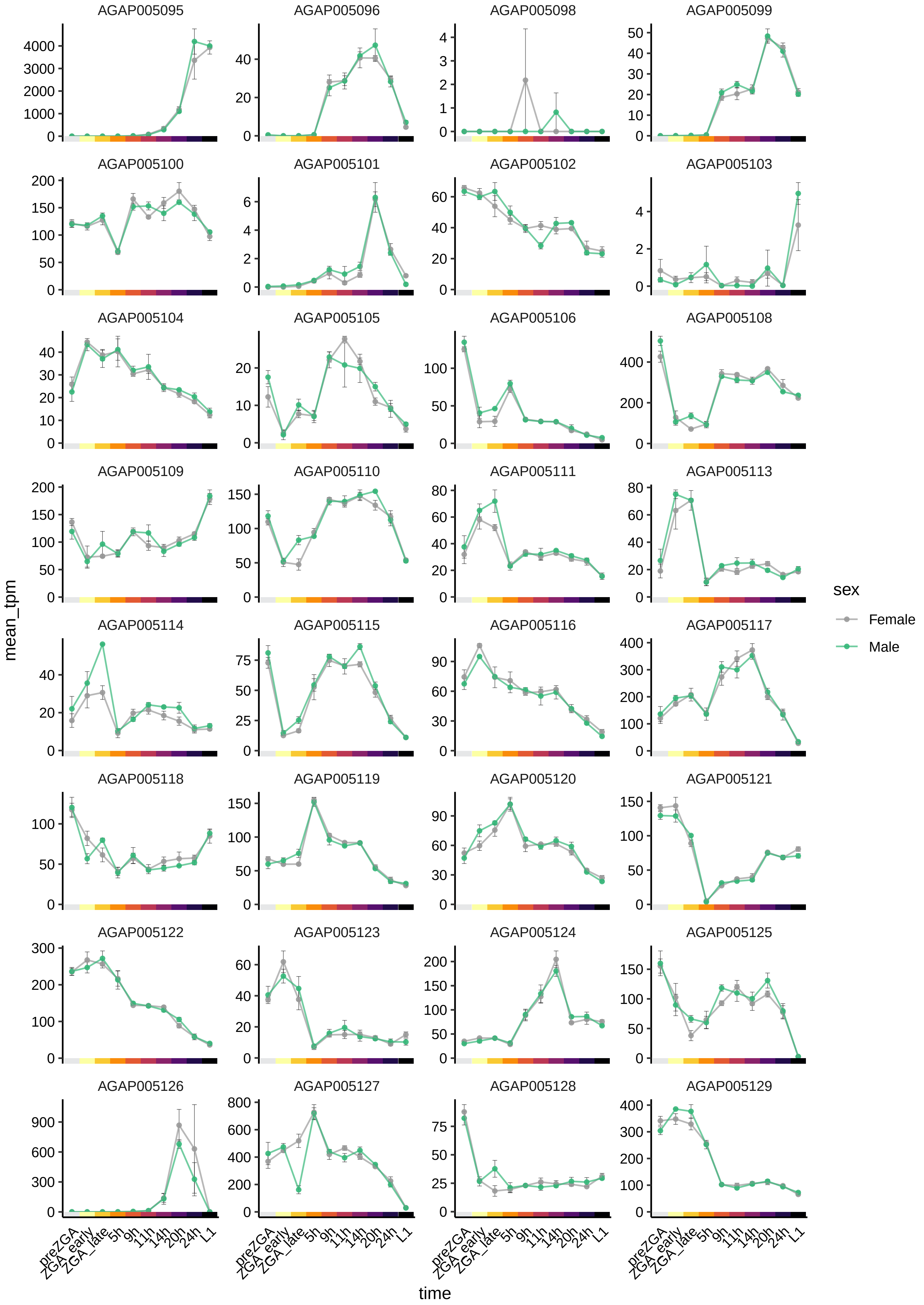

AGAP005095
AGAP005096
AGAP005098
AGAP005099
AGAP005100
AGAP005101
AGAP005102
AGAP005103
AGAP005104
AGAP005105
AGAP005106
AGAP005108
AGAP005109
AGAP005110
AGAP005111
AGAP005113
AGAP005114
AGAP005115
AGAP005116
AGAP005117
AGAP005118
AGAP005119
AGAP005120
AGAP005121
AGAP005122
AGAP005123
AGAP005124
AGAP005125
AGAP005126
AGAP005127
AGAP005128
AGAP005129
mean_tpm
time
preZGA
ZGA_early
ZGA_late
5h
9h
11h
14h
20h
24h
L1
sex
Female
Male

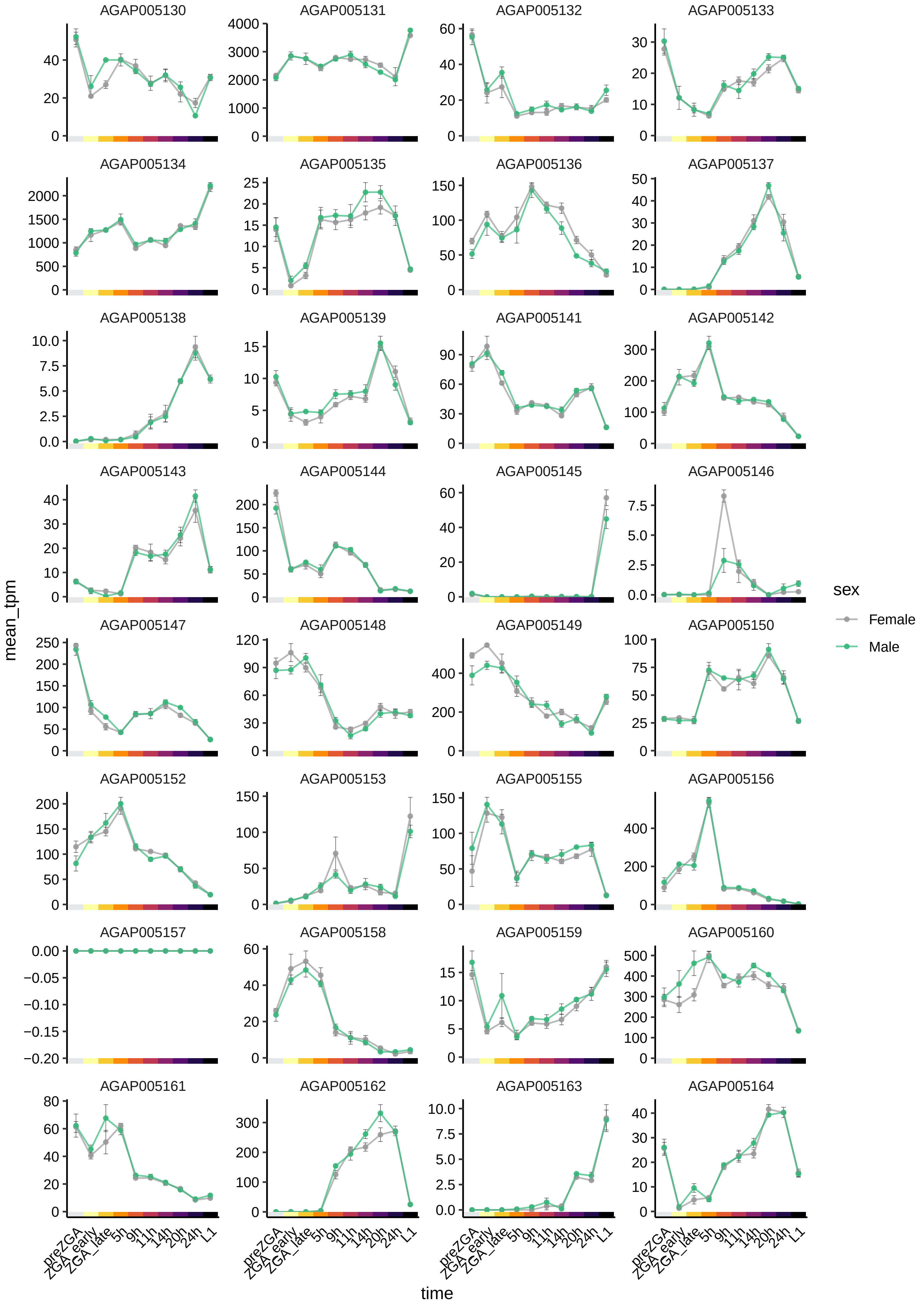

AGAP005130
AGAP005131
AGAP005132
AGAP005133
AGAP005134
AGAP005135
AGAP005136
AGAP005137
AGAP005138
AGAP005139
AGAP005141
AGAP005142
AGAP005143
AGAP005144
AGAP005145
AGAP005146
AGAP005147
AGAP005148
AGAP005149
AGAP005150
AGAP005152
AGAP005153
AGAP005155
AGAP005156
AGAP005157
AGAP005158
AGAP005159
AGAP005160
AGAP005161
AGAP005162
AGAP005163
AGAP005164
mean_tpm
time
preZGA
ZGA_early
ZGA_late
5h
9h
11h
14h
20h
24h
L1
sex
Female
Male

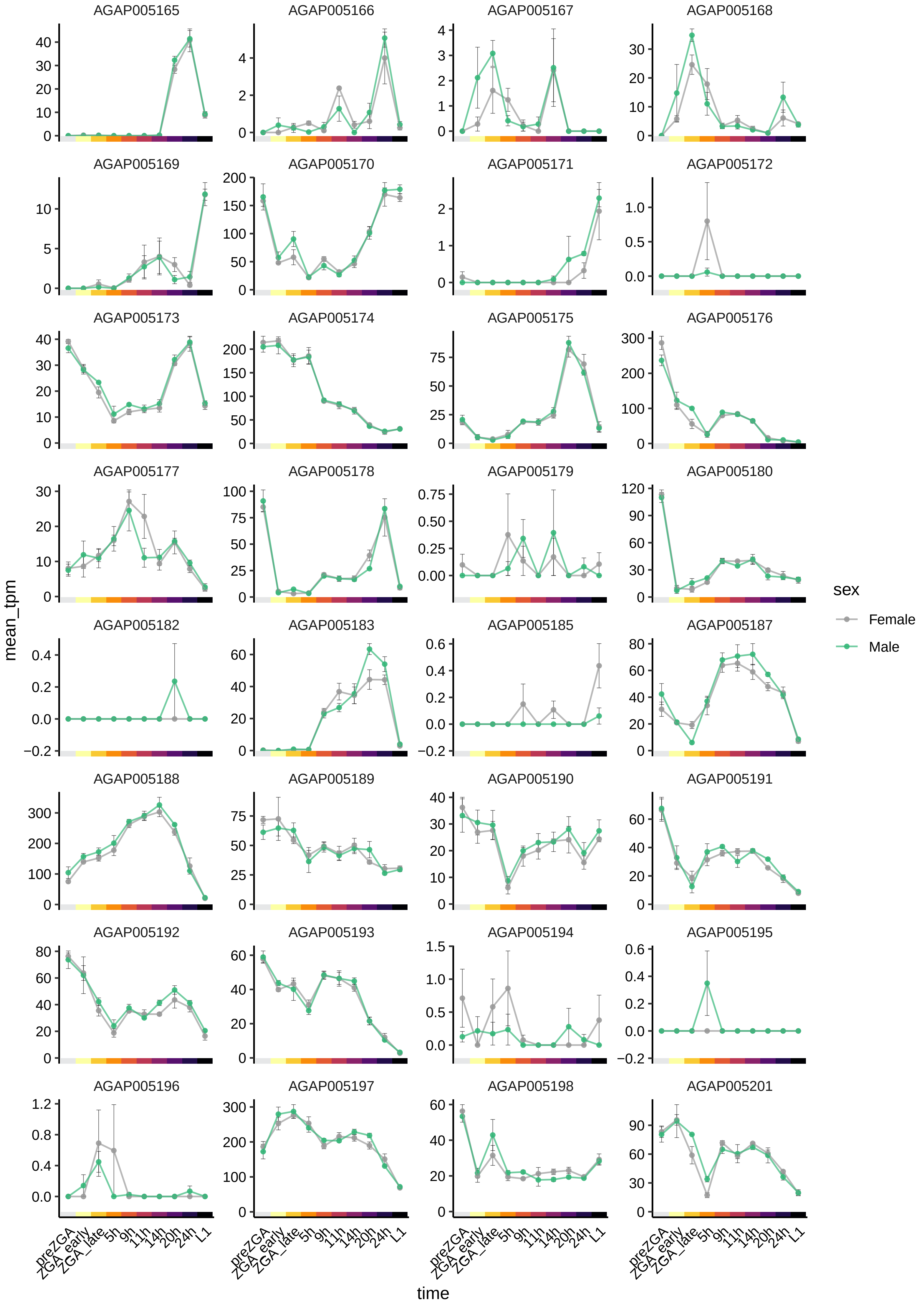
AGAP005165
AGAP005166
AGAP005167
AGAP005168
AGAP005169
AGAP005170
AGAP005171
AGAP005172
AGAP005173
AGAP005174
AGAP005175
AGAP005176
AGAP005177
AGAP005178
AGAP005179
AGAP005180
AGAP005182
AGAP005183
AGAP005185
AGAP005187
AGAP005188
AGAP005189
AGAP005190
AGAP005191
AGAP005192
AGAP005193
AGAP005194
AGAP005195
AGAP005196
AGAP005197
AGAP005198
AGAP005201
mean_tpm
time
preZGA
ZGA_early
ZGA_late
5h
9h
11h
14h
20h
24h
L1
sex
Female
Male

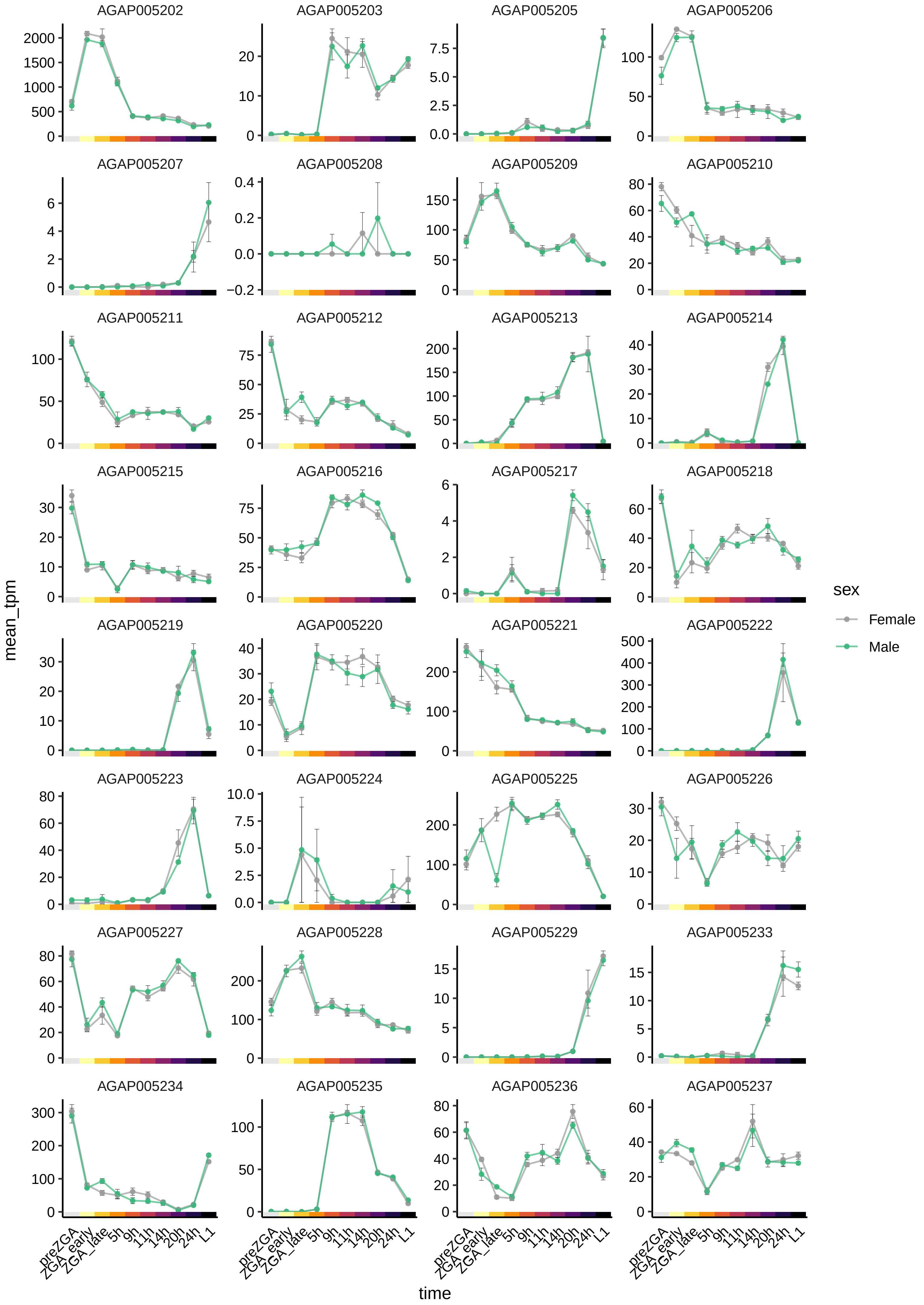
AGAP005202
AGAP005203
AGAP005205
AGAP005206
AGAP005207
AGAP005208
AGAP005209
AGAP005210
AGAP005211
AGAP005212
AGAP005213
AGAP005214
AGAP005215
AGAP005216
AGAP005217
AGAP005218
AGAP005219
AGAP005220
AGAP005221
AGAP005222
AGAP005223
AGAP005224
AGAP005225
AGAP005226
AGAP005227
AGAP005228
AGAP005229
AGAP005233
AGAP005234
AGAP005235
AGAP005236
AGAP005237
mean_tpm
time
preZGA
ZGA_early
ZGA_late
5h
9h
11h
14h
20h
24h
L1
sex
Female
Male

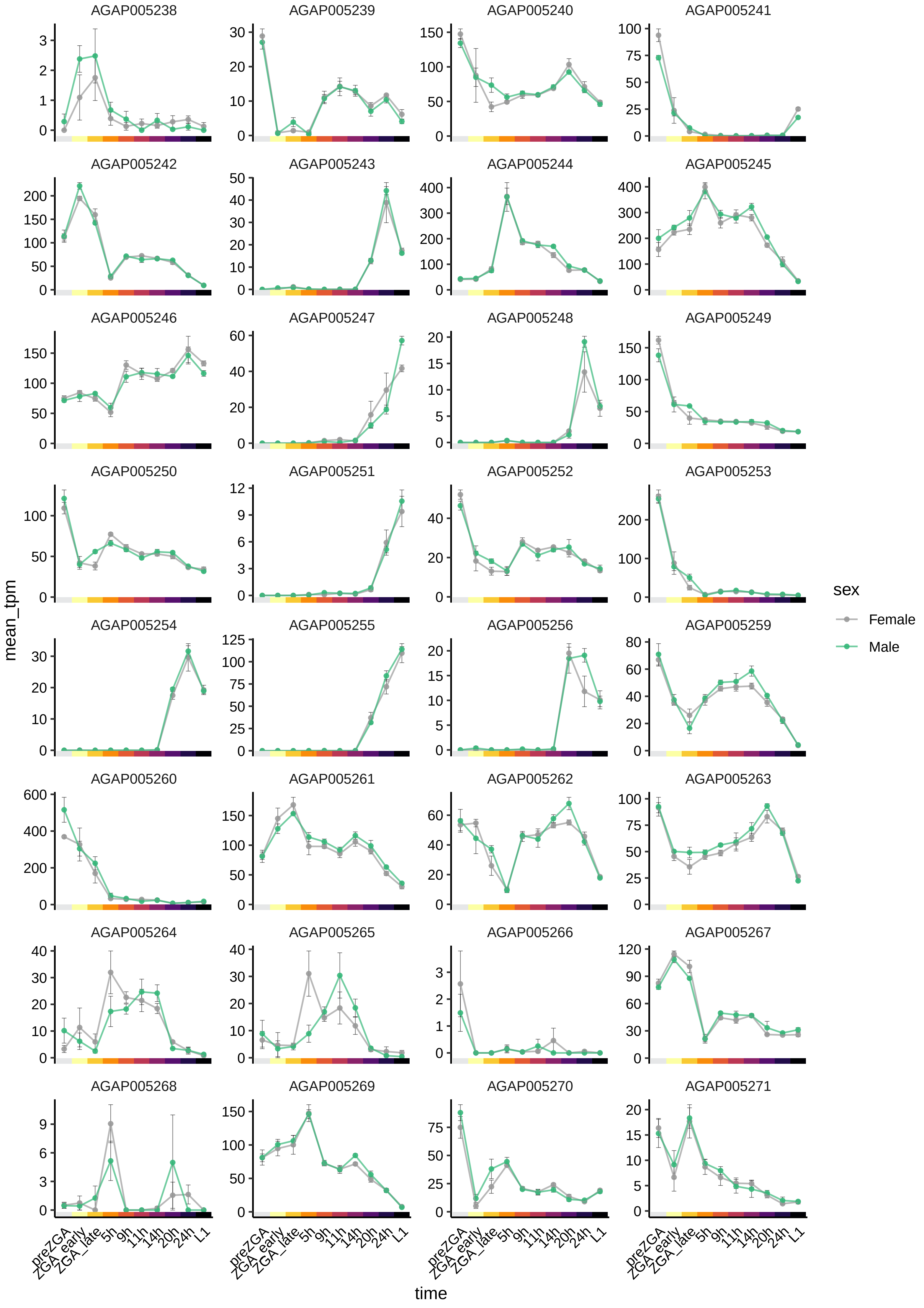

AGAP005238
AGAP005239
AGAP005240
AGAP005241
AGAP005242
AGAP005243
AGAP005244
AGAP005245
AGAP005246
AGAP005247
AGAP005248
AGAP005249
AGAP005250
AGAP005251
AGAP005252
AGAP005253
AGAP005254
AGAP005255
AGAP005256
AGAP005259
AGAP005260
AGAP005261
AGAP005262
AGAP005263
AGAP005264
AGAP005265
AGAP005266
AGAP005267
AGAP005268
AGAP005269
AGAP005270
AGAP005271
mean_tpm
time
preZGA
ZGA_early
ZGA_late
5h
9h
11h
14h
20h
24h
L1
sex
Female
Male

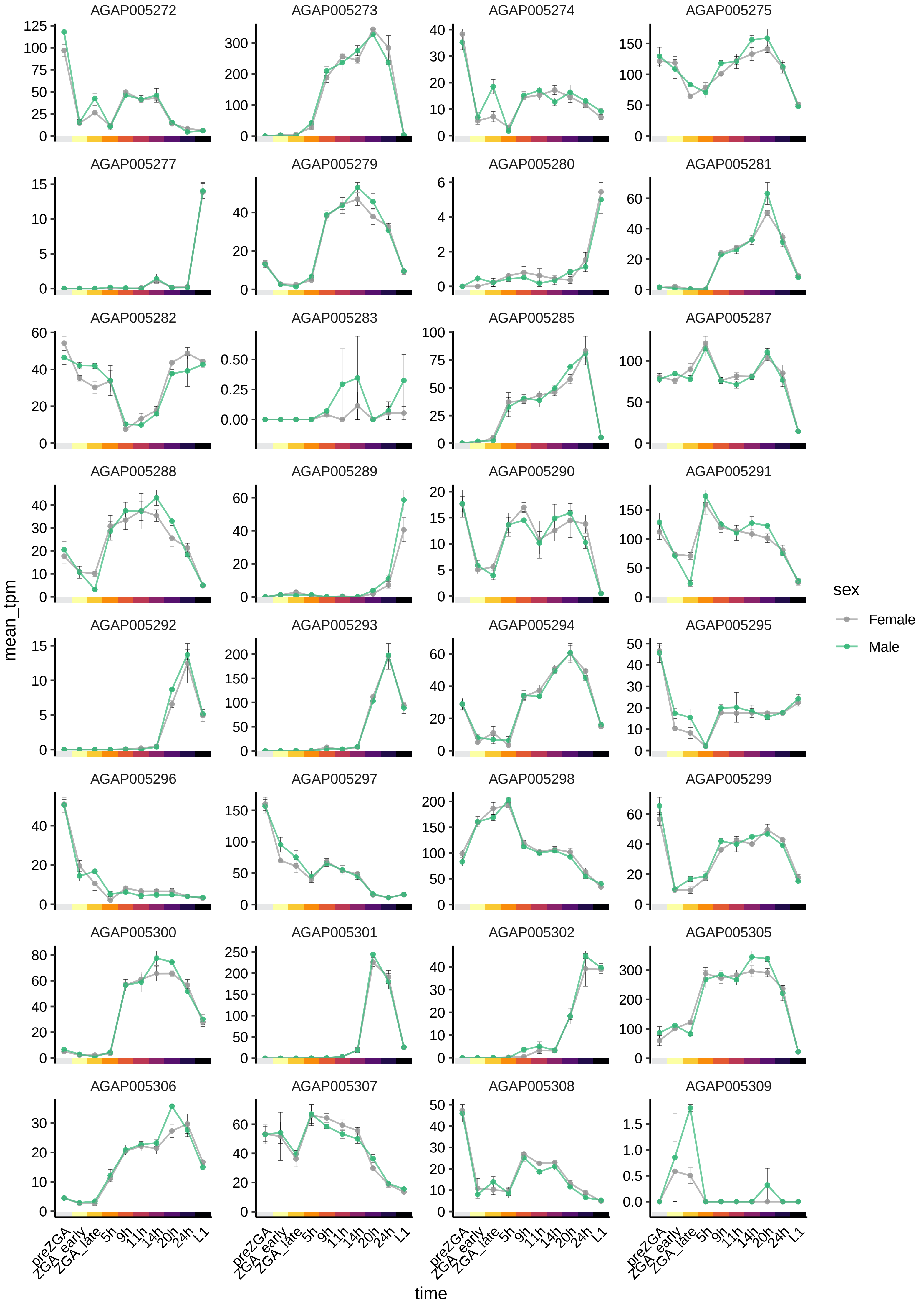
AGAP005272
AGAP005273
AGAP005274
AGAP005275
AGAP005277
AGAP005279
AGAP005280
AGAP005281
AGAP005282
AGAP005283
AGAP005285
AGAP005287
AGAP005288
AGAP005289
AGAP005290
AGAP005291
AGAP005292
AGAP005293
AGAP005294
AGAP005295
AGAP005296
AGAP005297
AGAP005298
AGAP005299
AGAP005300
AGAP005301
AGAP005302
AGAP005305
AGAP005306
AGAP005307
AGAP005308
AGAP005309
mean_tpm
time
preZGA
ZGA_early
ZGA_late
5h
9h
11h
14h
20h
24h
L1
sex
Female
Male

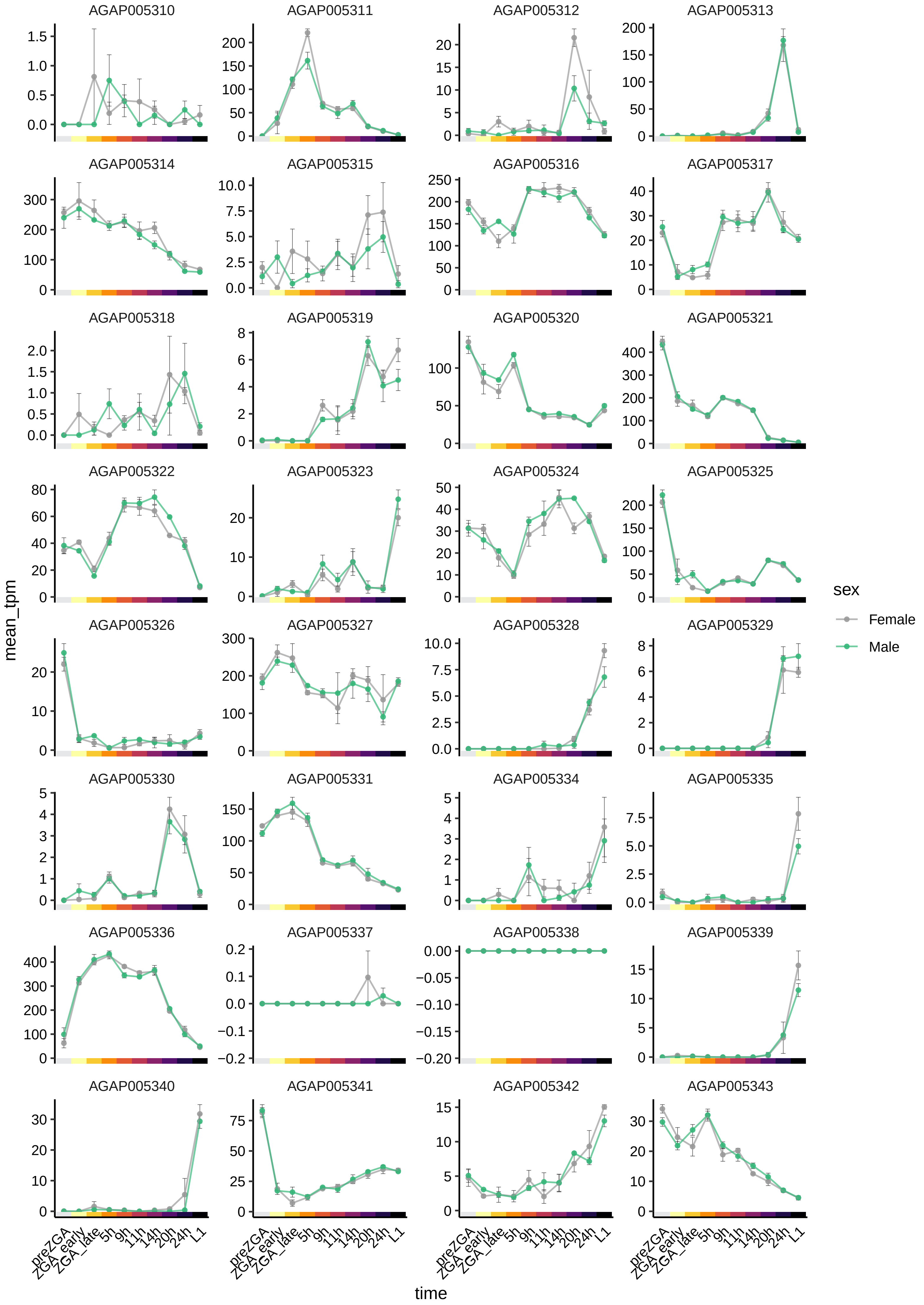
AGAP005310
AGAP005311
AGAP005312
AGAP005313
AGAP005314
AGAP005315
AGAP005316
AGAP005317
AGAP005318
AGAP005319
AGAP005320
AGAP005321
AGAP005322
AGAP005323
AGAP005324
AGAP005325
AGAP005326
AGAP005327
AGAP005328
AGAP005329
AGAP005330
AGAP005331
AGAP005334
AGAP005335
AGAP005336
AGAP005337
AGAP005338
AGAP005339
AGAP005340
AGAP005341
AGAP005342
AGAP005343
mean_tpm
time
preZGA
ZGA_early
ZGA_late
5h
9h
11h
14h
20h
24h
L1
sex
Female
Male

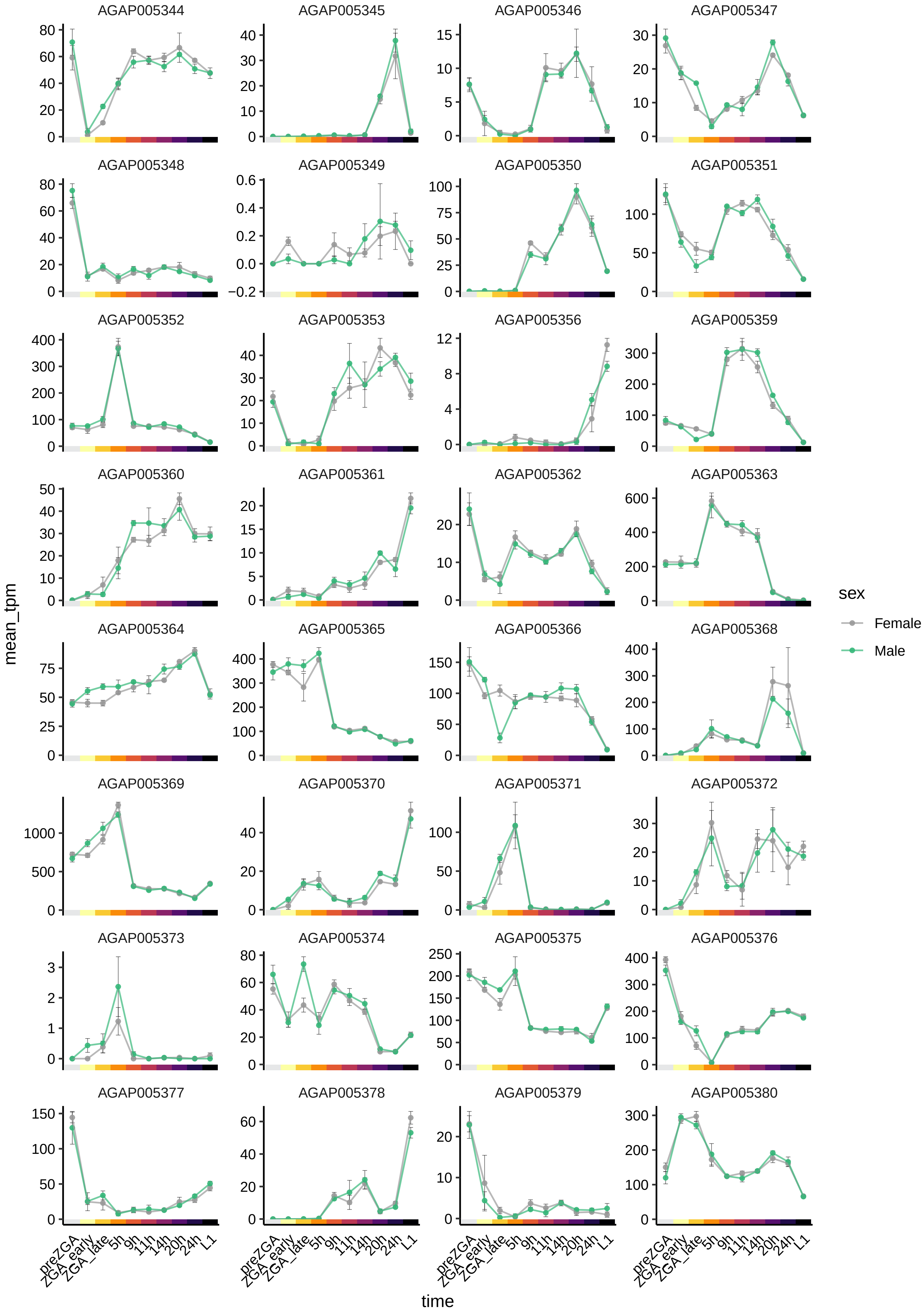

AGAP005344
AGAP005345
AGAP005346
AGAP005347
AGAP005348
AGAP005349
AGAP005350
AGAP005351
AGAP005352
AGAP005353
AGAP005356
AGAP005359
AGAP005360
AGAP005361
AGAP005362
AGAP005363
AGAP005364
AGAP005365
AGAP005366
AGAP005368
AGAP005369
AGAP005370
AGAP005371
AGAP005372
AGAP005373
AGAP005374
AGAP005375
AGAP005376
AGAP005377
AGAP005378
AGAP005379
AGAP005380
mean_tpm
time
preZGA
ZGA_early
ZGA_late
5h
9h
11h
14h
20h
24h
L1
sex
Female
Male

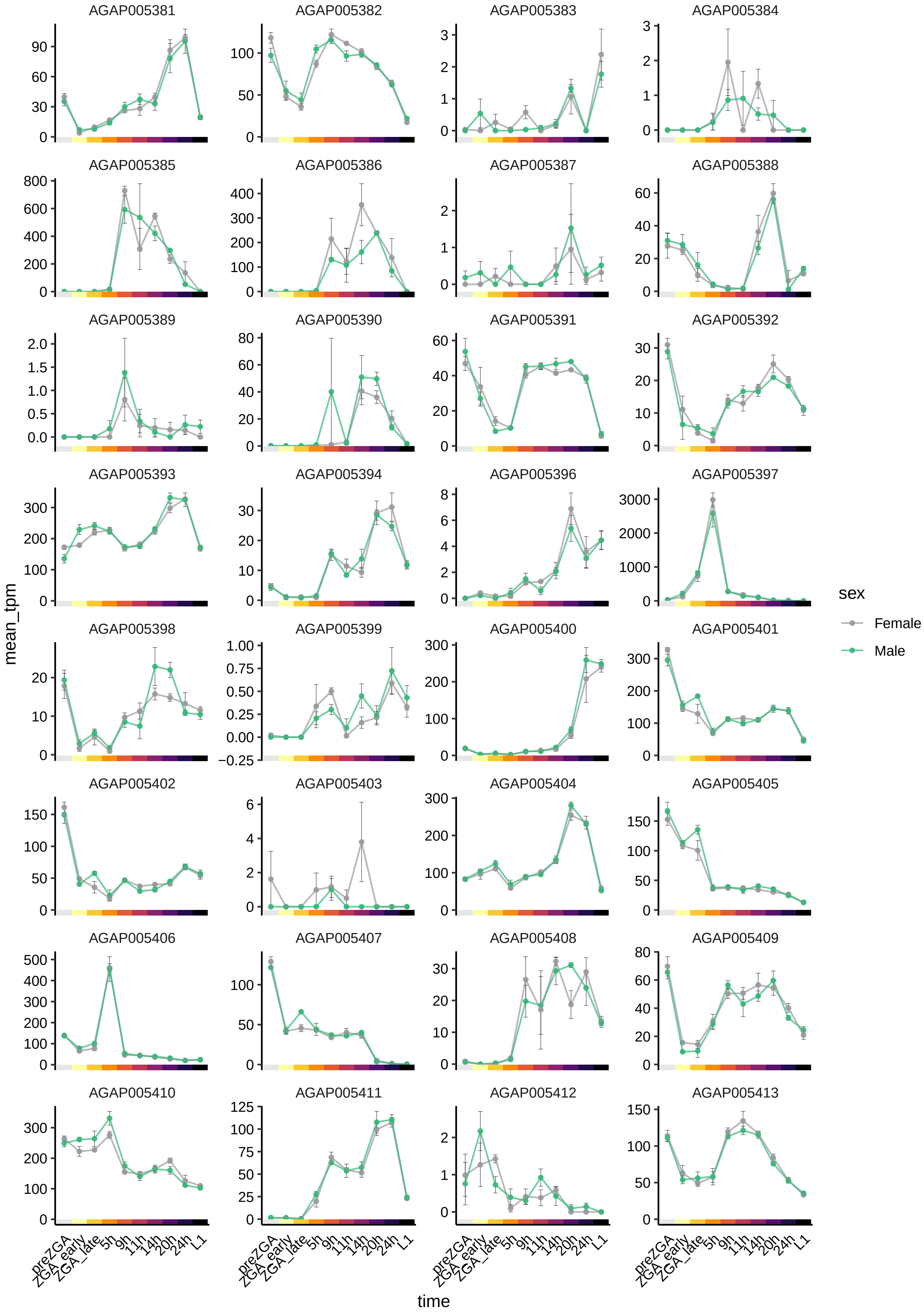

AGAP005381
AGAP005382
AGAP005383
AGAP005384
AGAP005385
AGAP005386
AGAP005387
AGAP005388
AGAP005389
AGAP005390
AGAP005391
AGAP005392
AGAP005393
AGAP005394
AGAP005396
AGAP005397
AGAP005398
AGAP005399
AGAP005400
AGAP005401
AGAP005402
AGAP005403
AGAP005404
AGAP005405
AGAP005406
AGAP005407
AGAP005408
AGAP005409
AGAP005410
AGAP005411
AGAP005412
AGAP005413
mean_tpm
time
preZGA
ZGA_early
ZGA_late
5h
9h
11h
14h
20h
24h
L1
sex
Female
Male

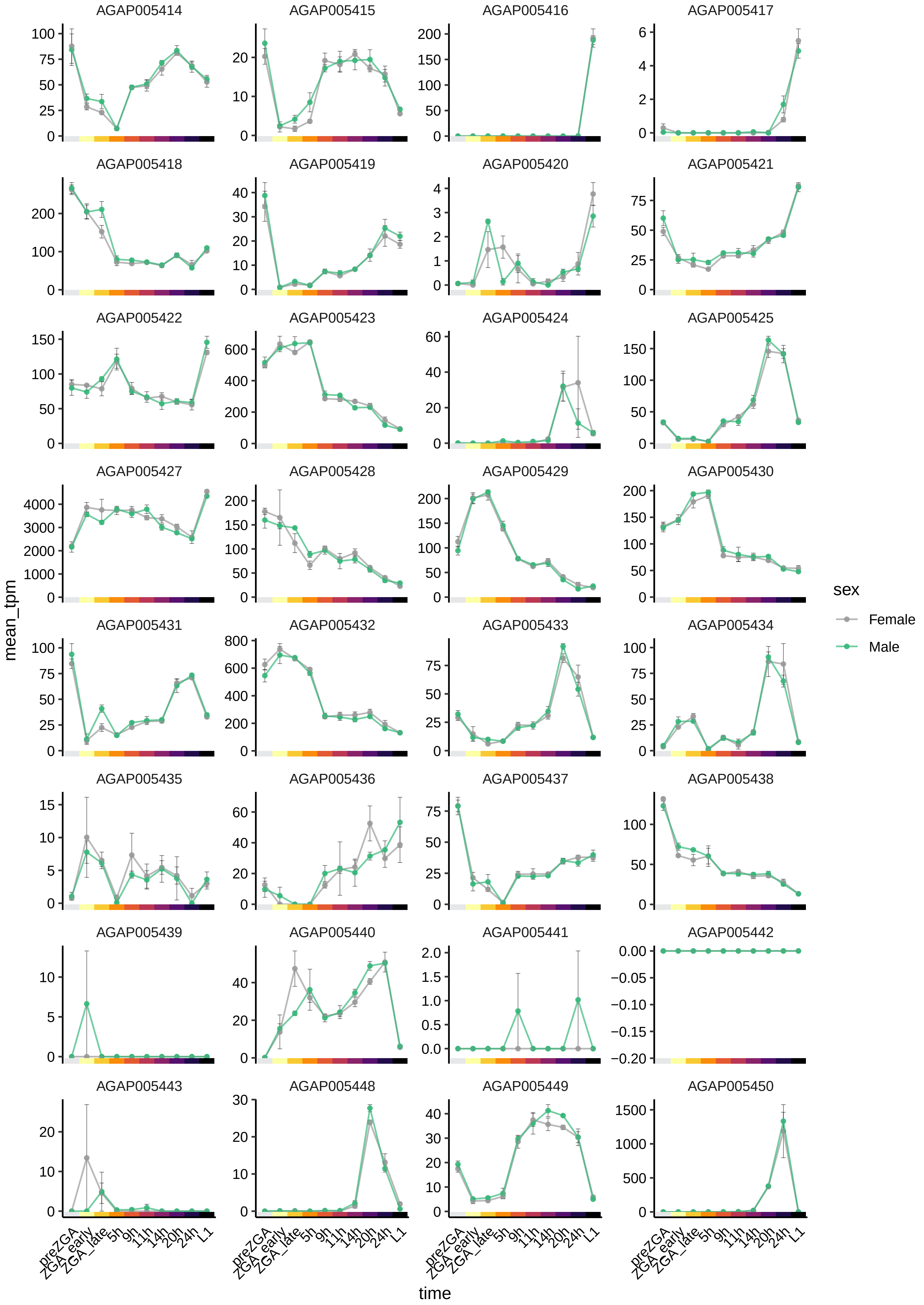
AGAP005414
AGAP005415
AGAP005416
AGAP005417
AGAP005418
AGAP005419
AGAP005420
AGAP005421
AGAP005422
AGAP005423
AGAP005424
AGAP005425
AGAP005427
AGAP005428
AGAP005429
AGAP005430
AGAP005431
AGAP005432
AGAP005433
AGAP005434
AGAP005435
AGAP005436
AGAP005437
AGAP005438
AGAP005439
AGAP005440
AGAP005441
AGAP005442
AGAP005443
AGAP005448
AGAP005449
AGAP005450
mean_tpm
time
preZGA
ZGA_early
ZGA_late
5h
9h
11h
14h
20h
24h
L1
sex
Female
Male

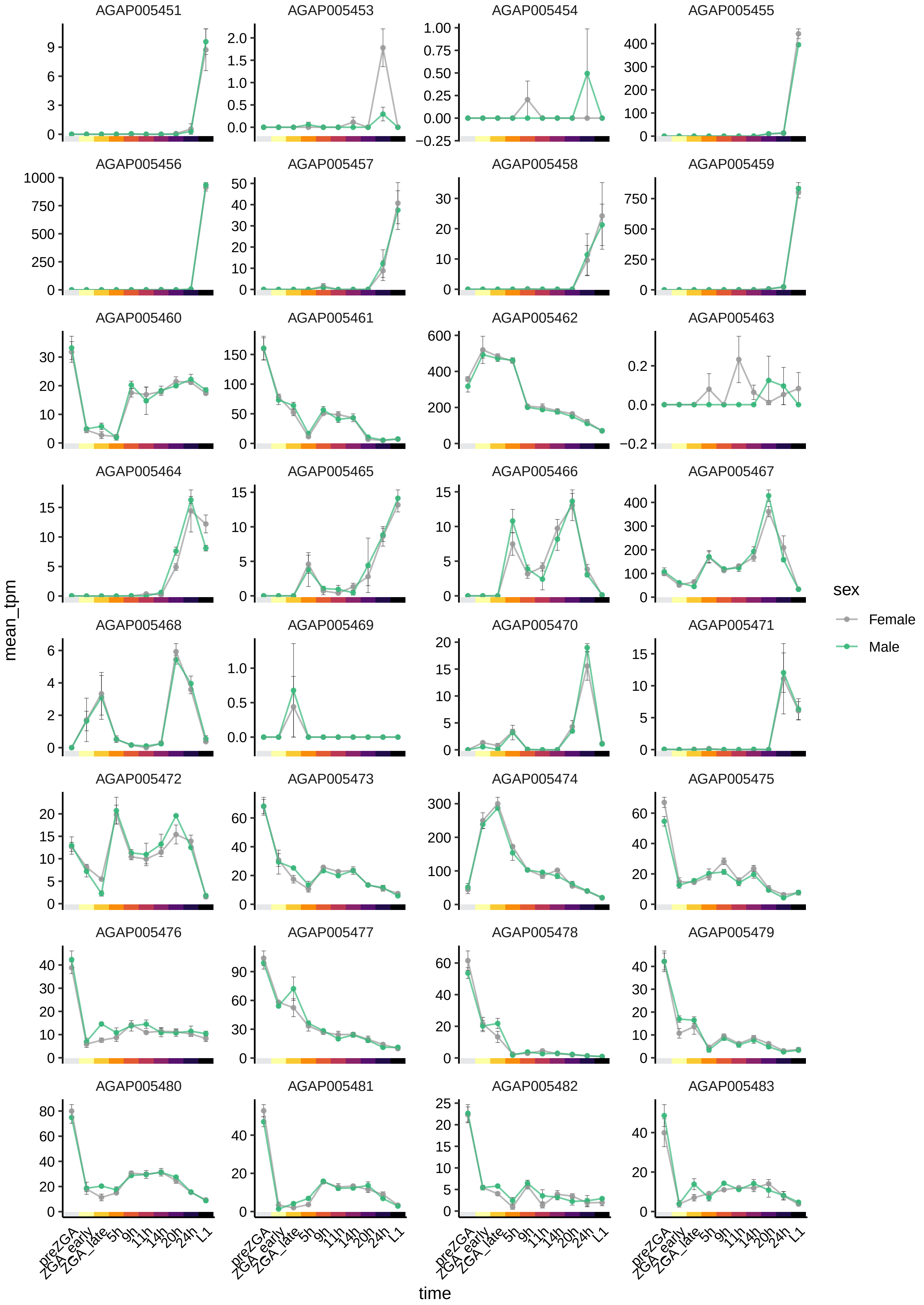

AGAP005451
AGAP005453
AGAP005454
AGAP005455
AGAP005456
AGAP005457
AGAP005458
AGAP005459
AGAP005460
AGAP005461
AGAP005462
AGAP005463
AGAP005464
AGAP005465
AGAP005466
AGAP005467
AGAP005468
AGAP005469
AGAP005470
AGAP005471
AGAP005472
AGAP005473
AGAP005474
AGAP005475
AGAP005476
AGAP005477
AGAP005478
AGAP005479
AGAP005480
AGAP005481
AGAP005482
AGAP005483
mean_tpm
time
preZGA
ZGA_early
ZGA_late
5h
9h
11h
14h
20h
24h
L1
sex
Female
Male

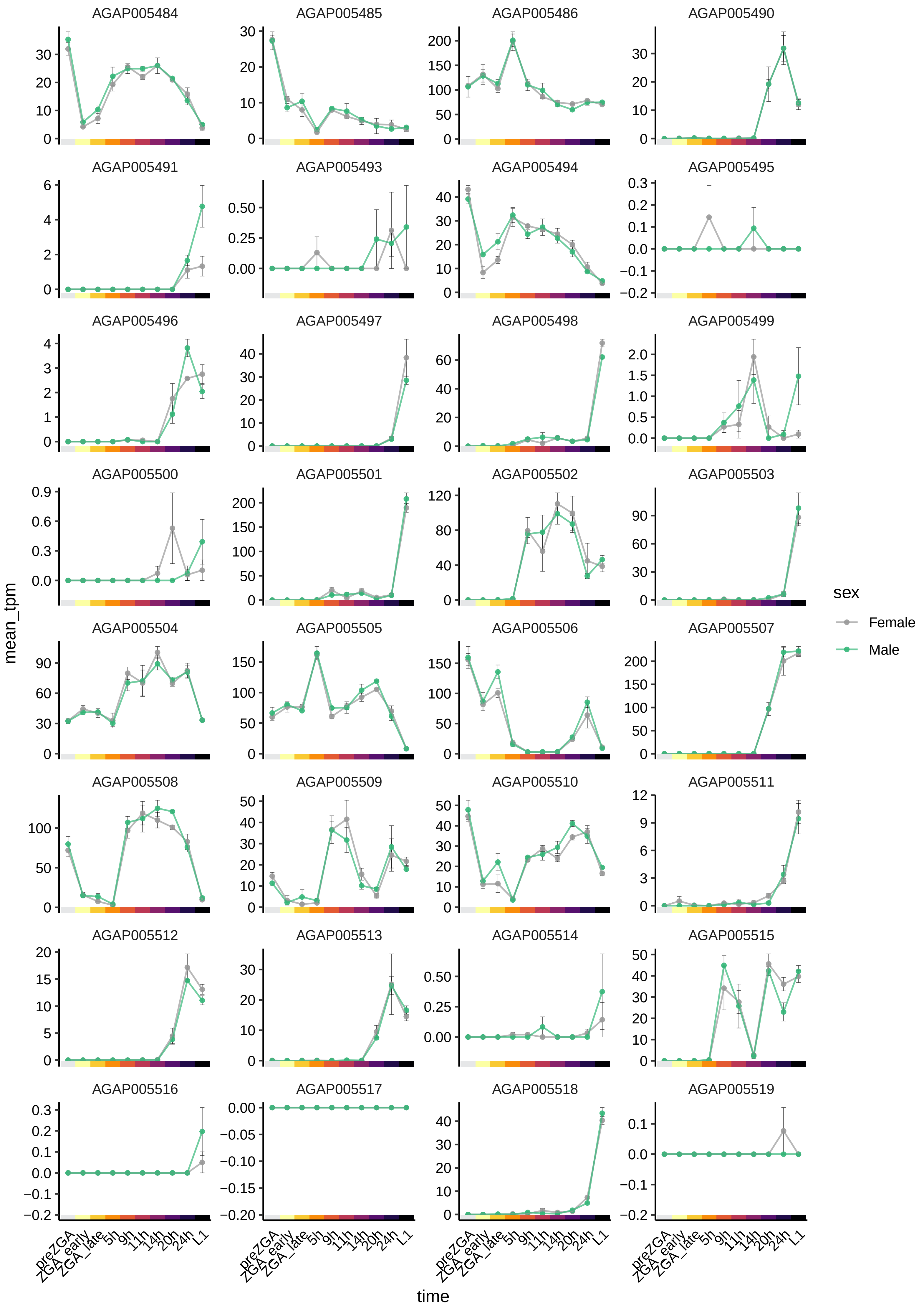
AGAP005484
AGAP005485
AGAP005486
AGAP005490
AGAP005491
AGAP005493
AGAP005494
AGAP005495
AGAP005496
AGAP005497
AGAP005498
AGAP005499
AGAP005500
AGAP005501
AGAP005502
AGAP005503
AGAP005504
AGAP005505
AGAP005506
AGAP005507
AGAP005508
AGAP005509
AGAP005510
AGAP005511
AGAP005512
AGAP005513
AGAP005514
AGAP005515
AGAP005516
AGAP005517
AGAP005518
AGAP005519
mean_tpm
time
preZGA
ZGA_early
ZGA_late
5h
9h
11h
14h
20h
24h
L1
sex
Female
Male

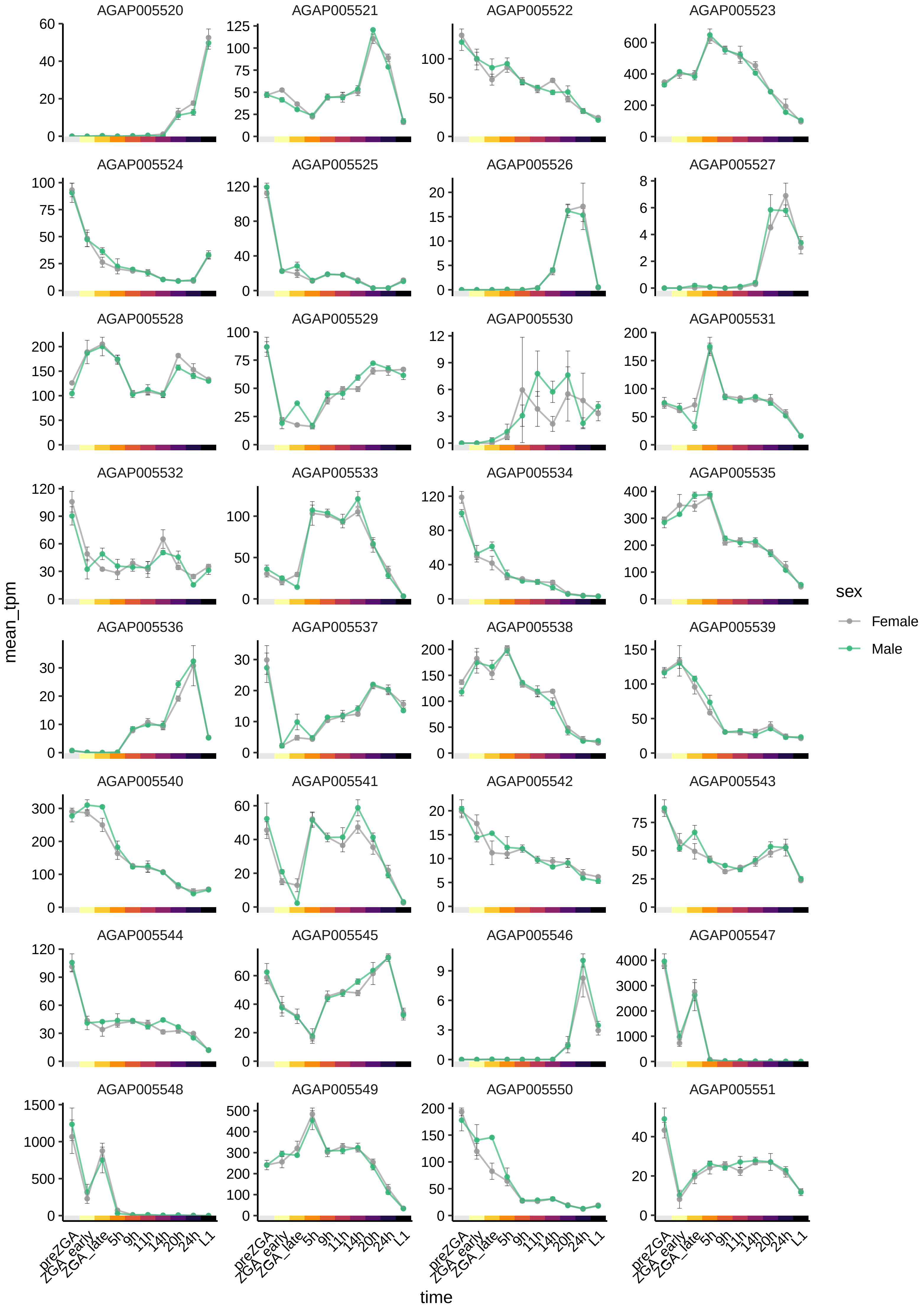
AGAP005520
AGAP005521
AGAP005522
AGAP005523
AGAP005524
AGAP005525
AGAP005526
AGAP005527
AGAP005528
AGAP005529
AGAP005530
AGAP005531
AGAP005532
AGAP005533
AGAP005534
AGAP005535
AGAP005536
AGAP005537
AGAP005538
AGAP005539
AGAP005540
AGAP005541
AGAP005542
AGAP005543
AGAP005544
AGAP005545
AGAP005546
AGAP005547
AGAP005548
AGAP005549
AGAP005550
AGAP005551
mean_tpm
time
preZGA
ZGA_early
ZGA_late
5h
9h
11h
14h
20h
24h
L1
sex
Female
Male

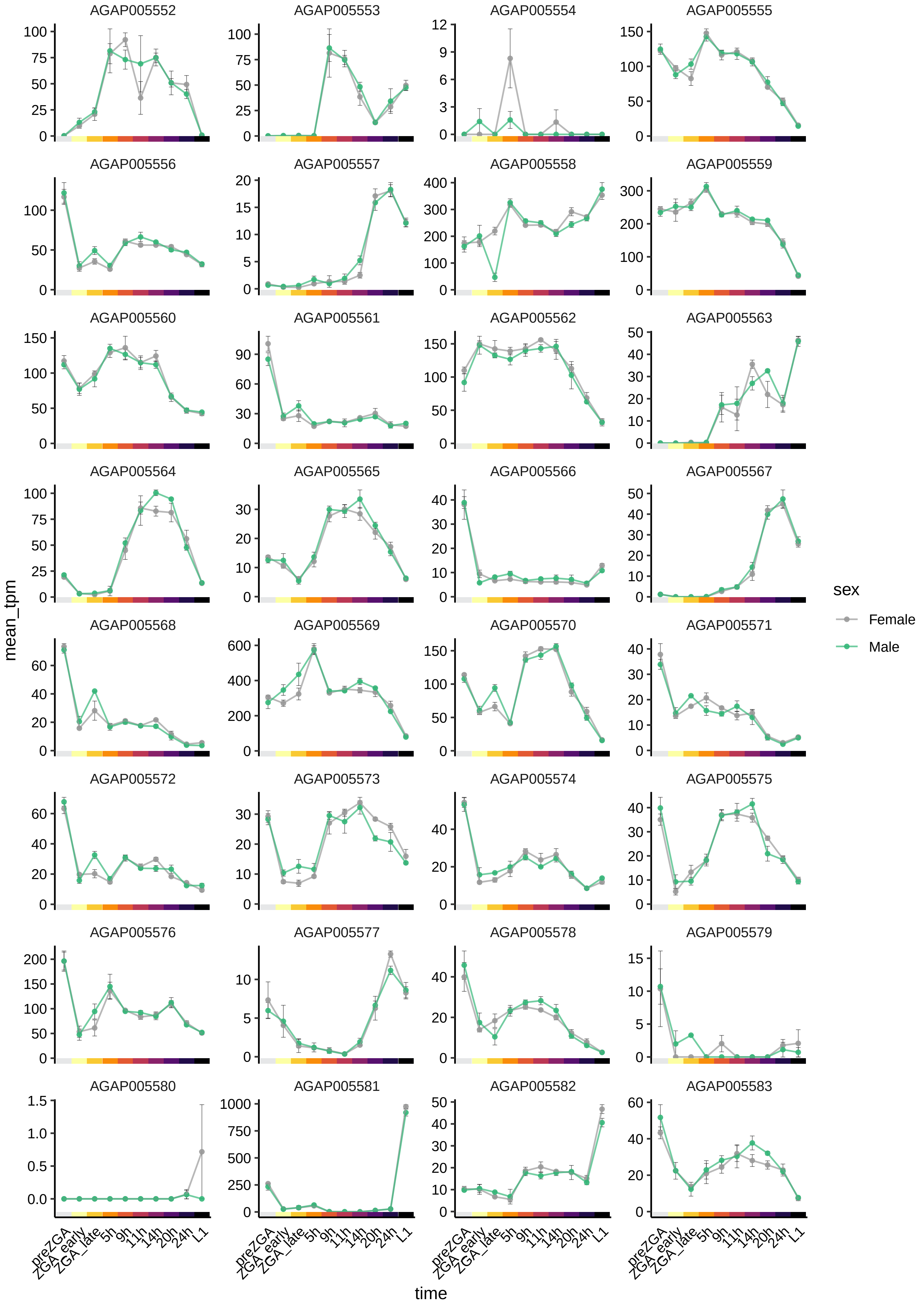

AGAP005552
AGAP005553
AGAP005554
AGAP005555
AGAP005556
AGAP005557
AGAP005558
AGAP005559
AGAP005560
AGAP005561
AGAP005562
AGAP005563
AGAP005564
AGAP005565
AGAP005566
AGAP005567
AGAP005568
AGAP005569
AGAP005570
AGAP005571
AGAP005572
AGAP005573
AGAP005574
AGAP005575
AGAP005576
AGAP005577
AGAP005578
AGAP005579
AGAP005580
AGAP005581
AGAP005582
AGAP005583
mean_tpm
time
preZGA
ZGA_early
ZGA_late
5h
9h
11h
14h
20h
24h
L1
sex
Female
Male

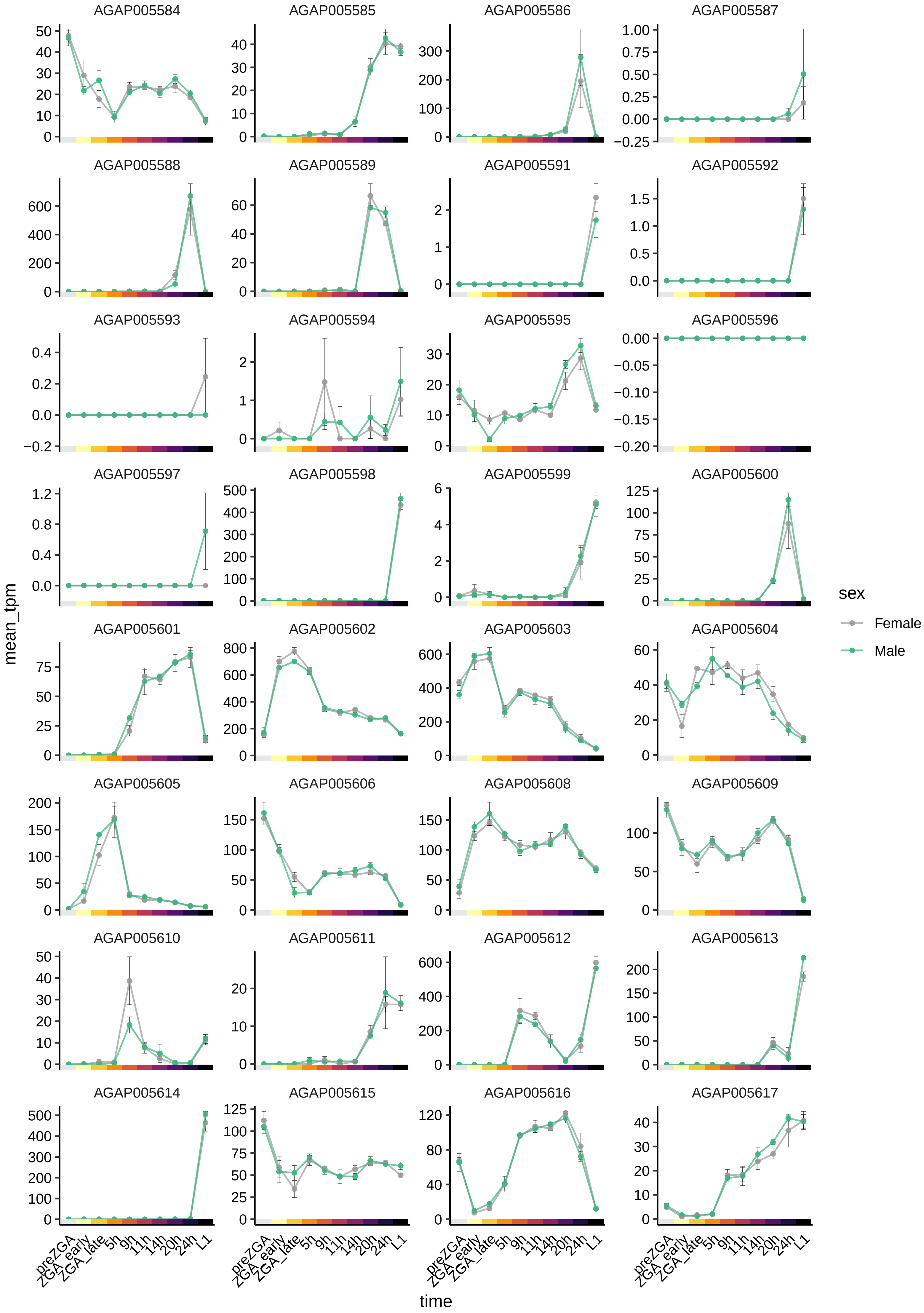

AGAP005584
AGAP005585
AGAP005586
AGAP005587
AGAP005588
AGAP005589
AGAP005591
AGAP005592
AGAP005593
AGAP005594
AGAP005595
AGAP005596
AGAP005597
AGAP005598
AGAP005599
AGAP005600
AGAP005601
AGAP005602
AGAP005603
AGAP005604
AGAP005605
AGAP005606
AGAP005608
AGAP005609
AGAP005610
AGAP005611
AGAP005612
AGAP005613
AGAP005614
AGAP005615
AGAP005616
AGAP005617
mean_tpm
time
preZGA
ZGA_early
ZGA_late
5h
9h
11h
14h
20h
24h
L1
sex
Female
Male

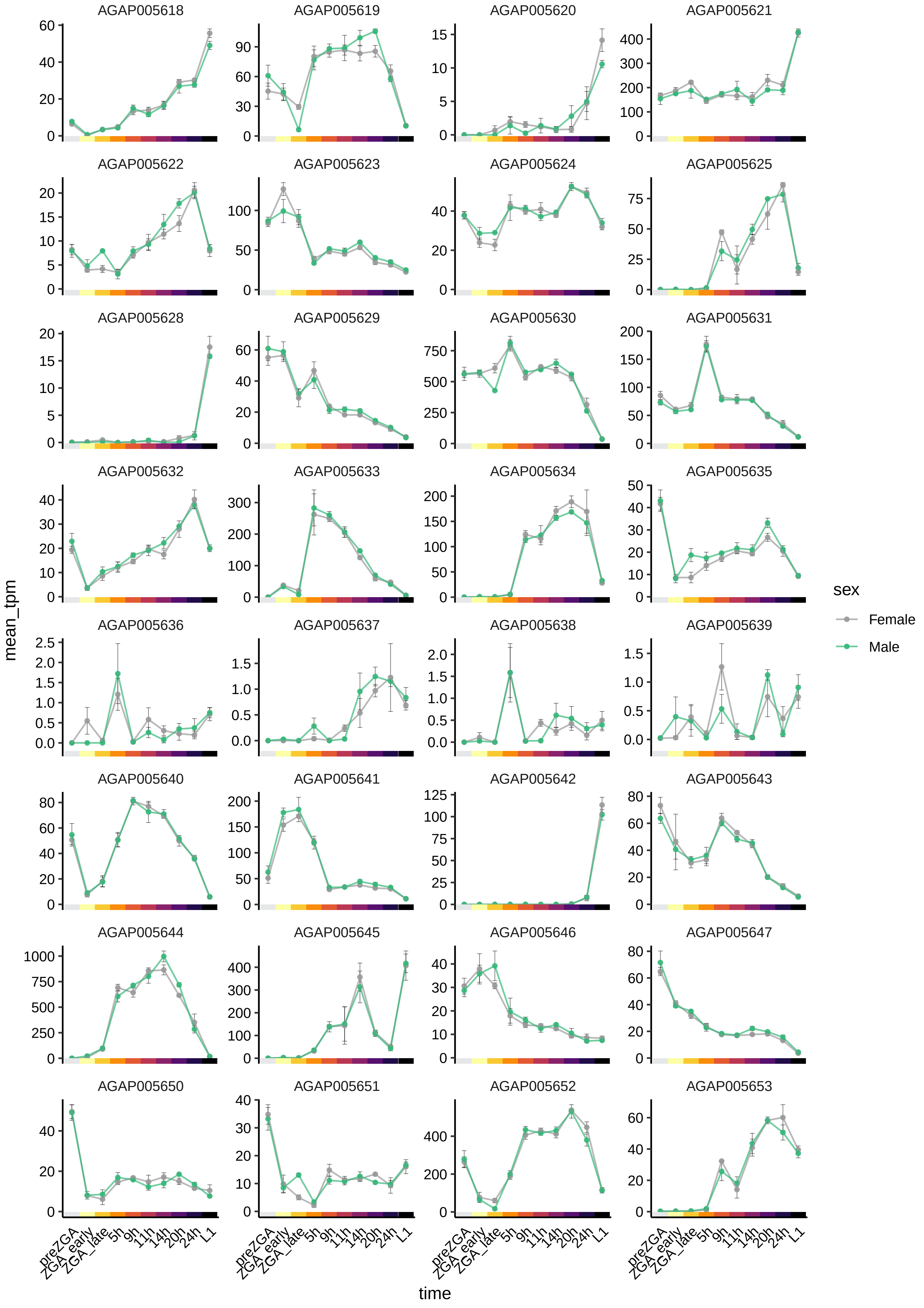

AGAP005618
AGAP005619
AGAP005620
AGAP005621
AGAP005622
AGAP005623
AGAP005624
AGAP005625
AGAP005628
AGAP005629
AGAP005630
AGAP005631
AGAP005632
AGAP005633
AGAP005634
AGAP005635
AGAP005636
AGAP005637
AGAP005638
AGAP005639
AGAP005640
AGAP005641
AGAP005642
AGAP005643
AGAP005644
AGAP005645
AGAP005646
AGAP005647
AGAP005650
AGAP005651
AGAP005652
AGAP005653
mean_tpm
time
preZGA
ZGA_early
ZGA_late
5h
9h
11h
14h
20h
24h
L1
sex
Female
Male

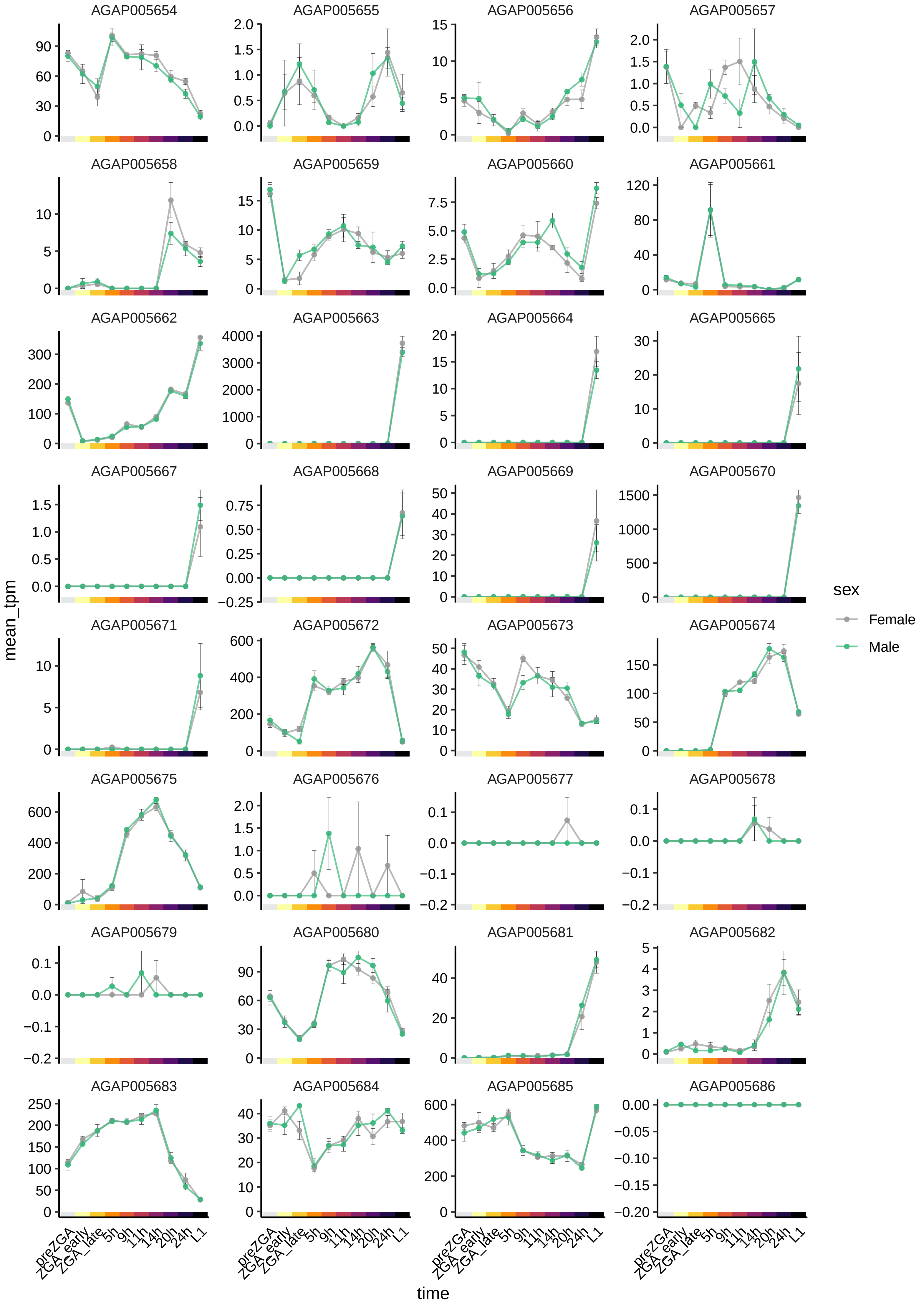
AGAP005654
AGAP005655
AGAP005656
AGAP005657
AGAP005658
AGAP005659
AGAP005660
AGAP005661
AGAP005662
AGAP005663
AGAP005664
AGAP005665
AGAP005667
AGAP005668
AGAP005669
AGAP005670
AGAP005671
AGAP005672
AGAP005673
AGAP005674
AGAP005675
AGAP005676
AGAP005677
AGAP005678
AGAP005679
AGAP005680
AGAP005681
AGAP005682
AGAP005683
AGAP005684
AGAP005685
AGAP005686
mean_tpm
time
preZGA
ZGA_early
ZGA_late
5h
9h
11h
14h
20h
24h
L1
sex
Female
Male

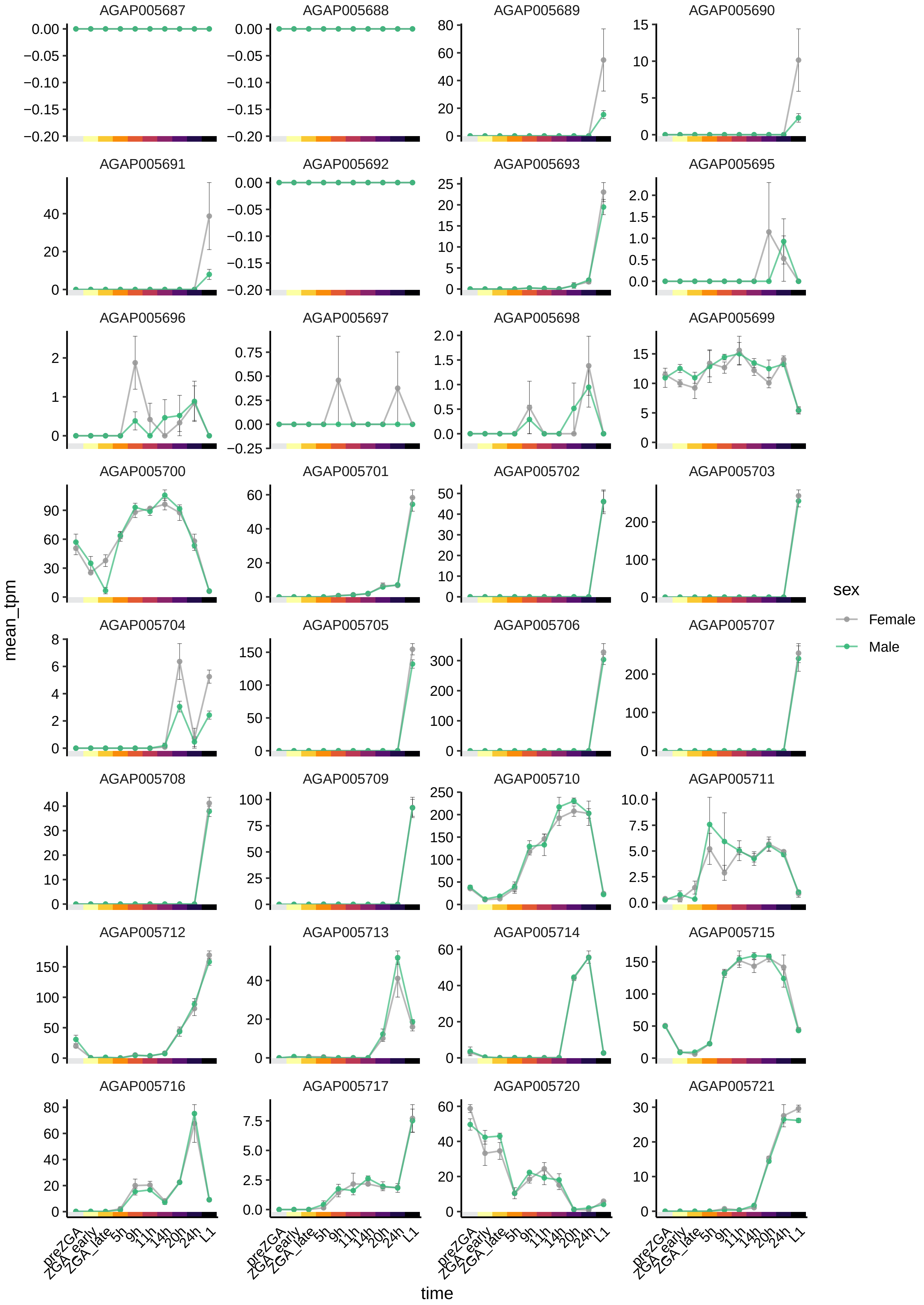
AGAP005687
AGAP005688
AGAP005689
AGAP005690
AGAP005691
AGAP005692
AGAP005693
AGAP005695
AGAP005696
AGAP005697
AGAP005698
AGAP005699
AGAP005700
AGAP005701
AGAP005702
AGAP005703
AGAP005704
AGAP005705
AGAP005706
AGAP005707
AGAP005708
AGAP005709
AGAP005710
AGAP005711
AGAP005712
AGAP005713
AGAP005714
AGAP005715
AGAP005716
AGAP005717
AGAP005720
AGAP005721
mean_tpm
time
preZGA
ZGA_early
ZGA_late
5h
9h
11h
14h
20h
24h
L1
sex
Female
Male

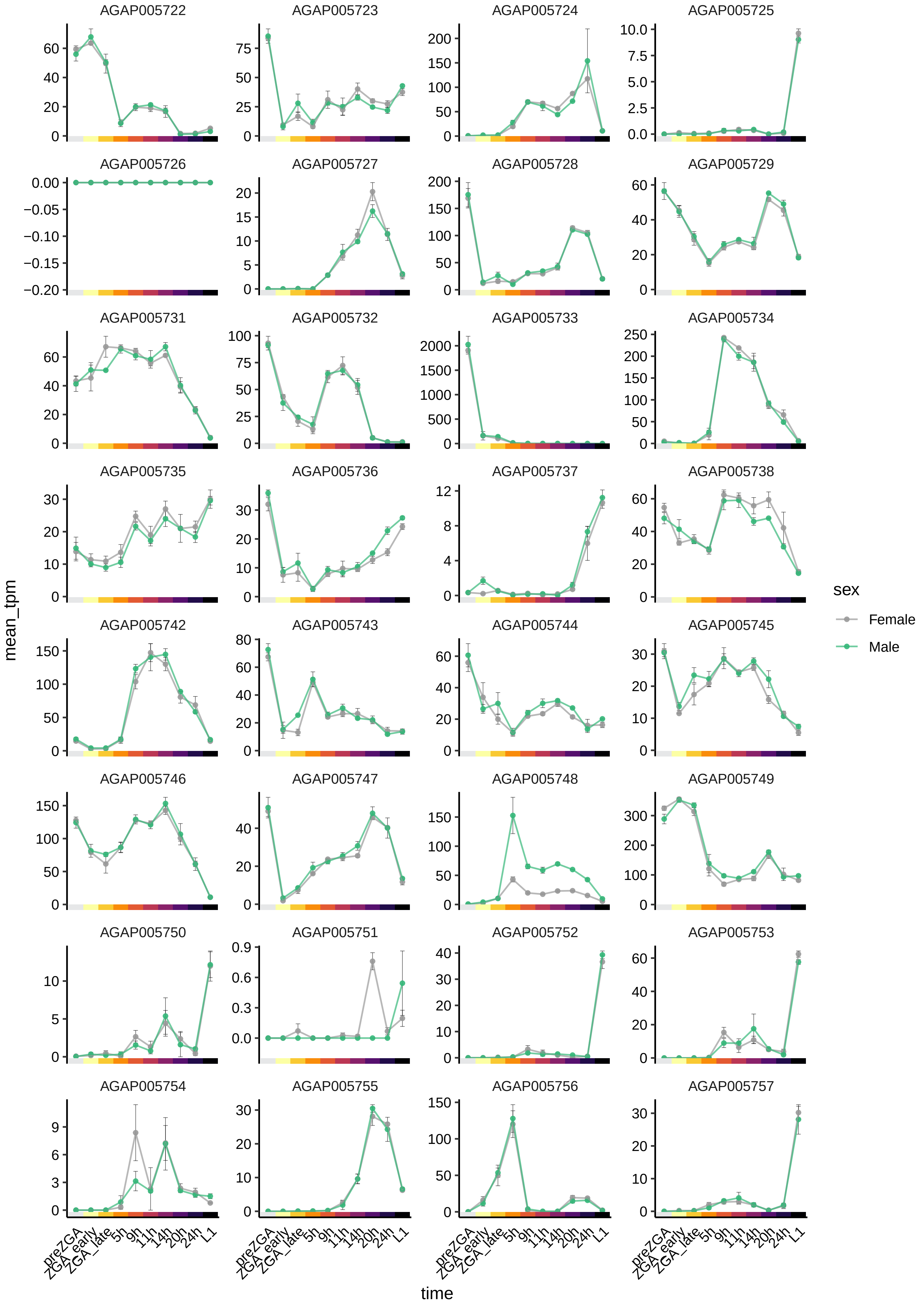
AGAP005722
AGAP005723
AGAP005724
AGAP005725
AGAP005726
AGAP005727
AGAP005728
AGAP005729
AGAP005731
AGAP005732
AGAP005733
AGAP005734
AGAP005735
AGAP005736
AGAP005737
AGAP005738
AGAP005742
AGAP005743
AGAP005744
AGAP005745
AGAP005746
AGAP005747
AGAP005748
AGAP005749
AGAP005750
AGAP005751
AGAP005752
AGAP005753
AGAP005754
AGAP005755
AGAP005756
AGAP005757
mean_tpm
time
preZGA
ZGA_early
ZGA_late
5h
9h
11h
14h
20h
24h
L1
sex
Female
Male

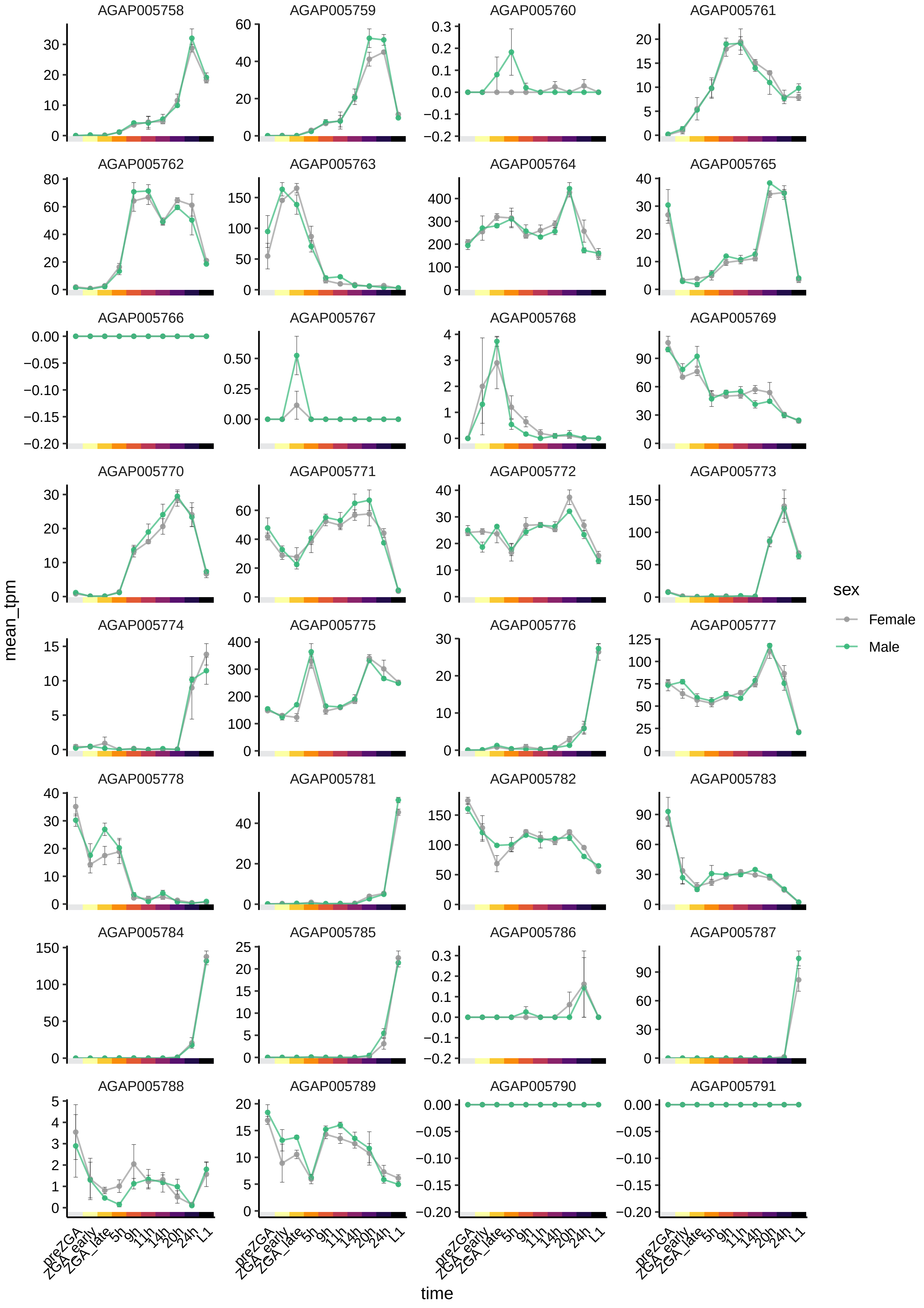

AGAP005758
AGAP005759
AGAP005760
AGAP005761
AGAP005762
AGAP005763
AGAP005764
AGAP005765
AGAP005766
AGAP005767
AGAP005768
AGAP005769
AGAP005770
AGAP005771
AGAP005772
AGAP005773
AGAP005774
AGAP005775
AGAP005776
AGAP005777
AGAP005778
AGAP005781
AGAP005782
AGAP005783
AGAP005784
AGAP005785
AGAP005786
AGAP005787
AGAP005788
AGAP005789
AGAP005790
AGAP005791
mean_tpm
time
preZGA
ZGA_early
ZGA_late
5h
9h
11h
14h
20h
24h
L1
sex
Female
Male

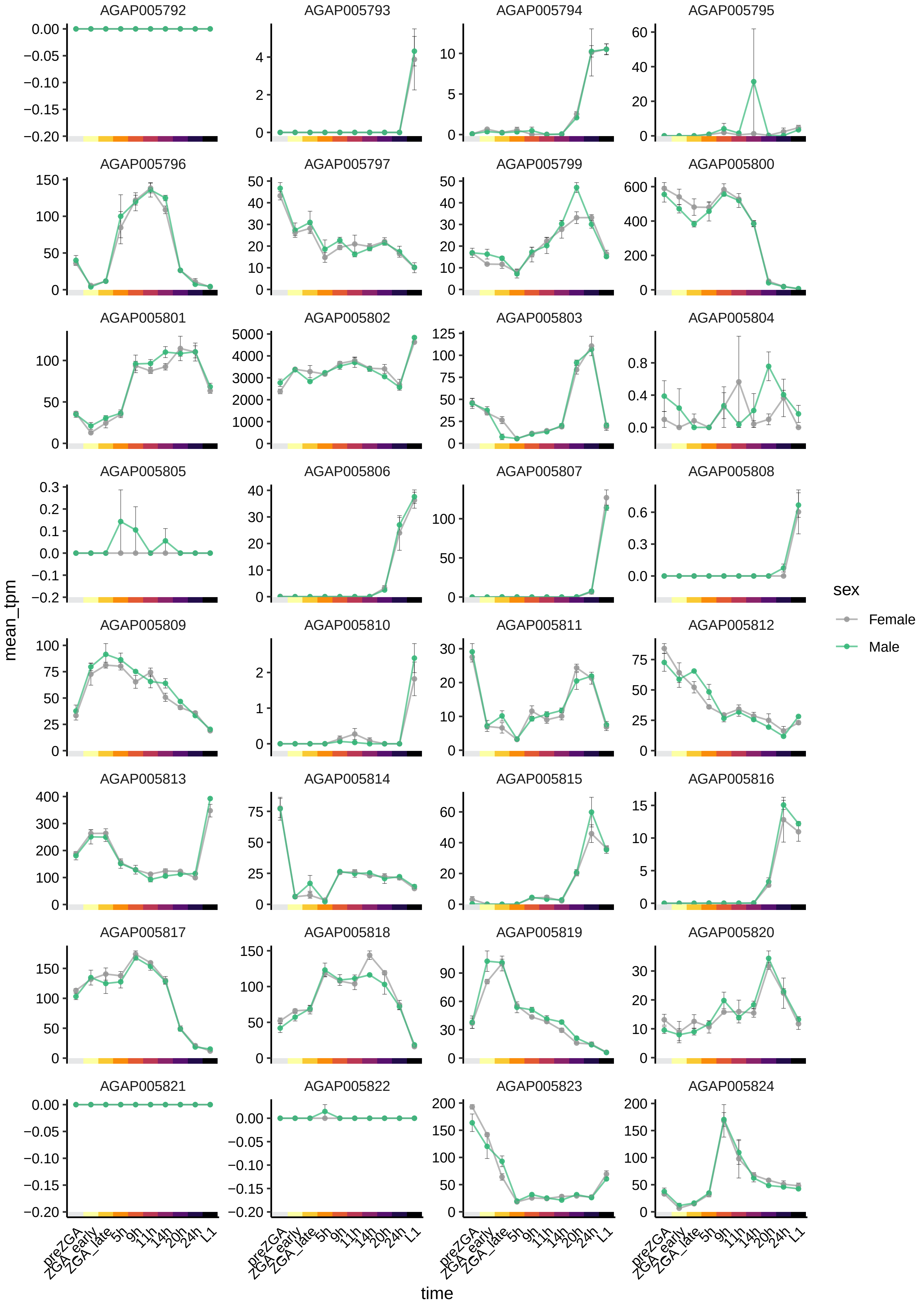

AGAP005792
AGAP005793
AGAP005794
AGAP005795
AGAP005796
AGAP005797
AGAP005799
AGAP005800
AGAP005801
AGAP005802
AGAP005803
AGAP005804
AGAP005805
AGAP005806
AGAP005807
AGAP005808
AGAP005809
AGAP005810
AGAP005811
AGAP005812
AGAP005813
AGAP005814
AGAP005815
AGAP005816
AGAP005817
AGAP005818
AGAP005819
AGAP005820
AGAP005821
AGAP005822
AGAP005823
AGAP005824
mean_tpm
time
preZGA
ZGA_early
ZGA_late
5h
9h
11h
14h
20h
24h
L1
sex
Female
Male

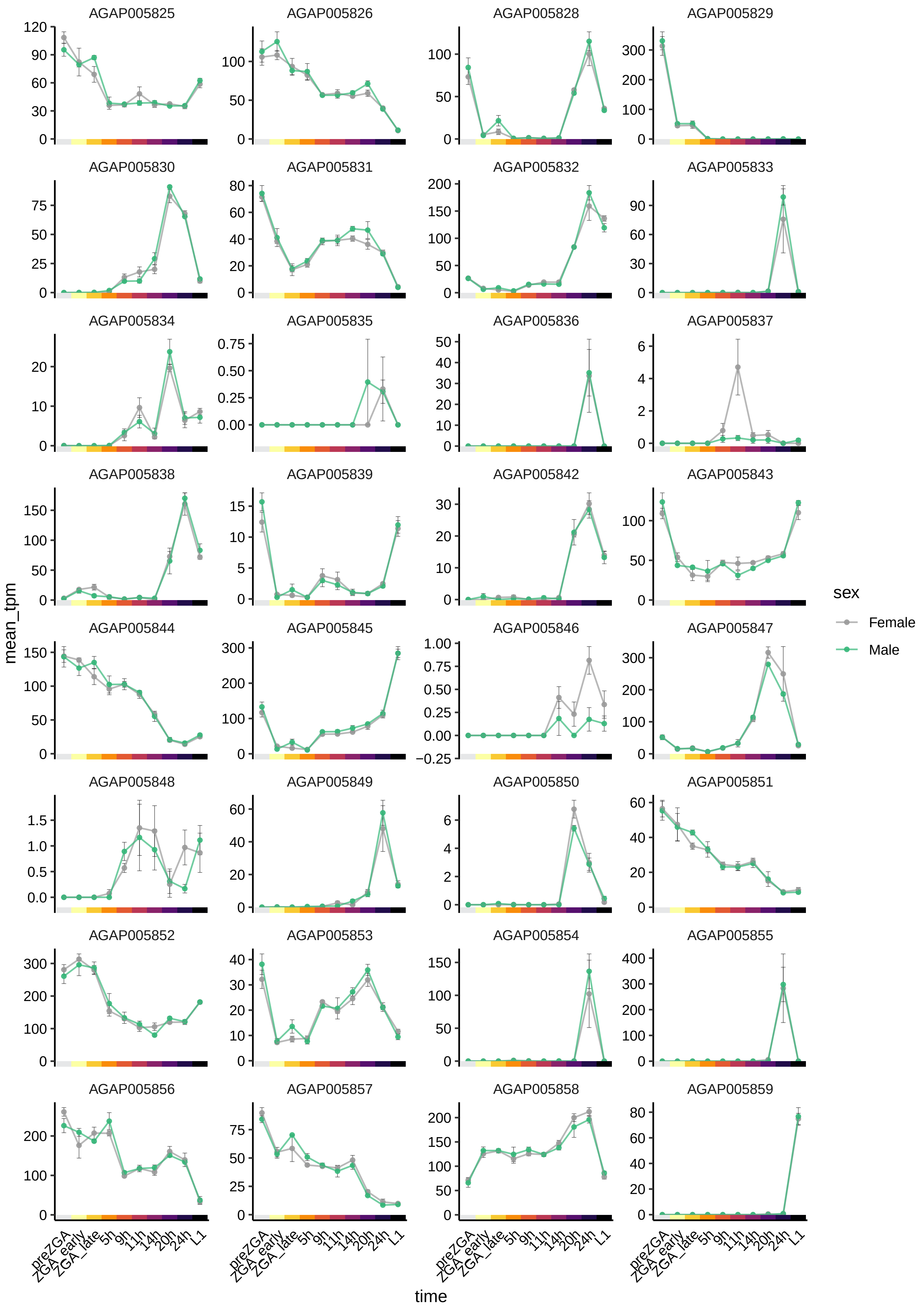
AGAP005825
AGAP005826
AGAP005828
AGAP005829
AGAP005830
AGAP005831
AGAP005832
AGAP005833
AGAP005834
AGAP005835
AGAP005836
AGAP005837
AGAP005838
AGAP005839
AGAP005842
AGAP005843
AGAP005844
AGAP005845
AGAP005846
AGAP005847
AGAP005848
AGAP005849
AGAP005850
AGAP005851
AGAP005852
AGAP005853
AGAP005854
AGAP005855
AGAP005856
AGAP005857
AGAP005858
AGAP005859
mean_tpm
time
preZGA
ZGA_early
ZGA_late
5h
9h
11h
14h
20h
24h
L1
sex
Female
Male

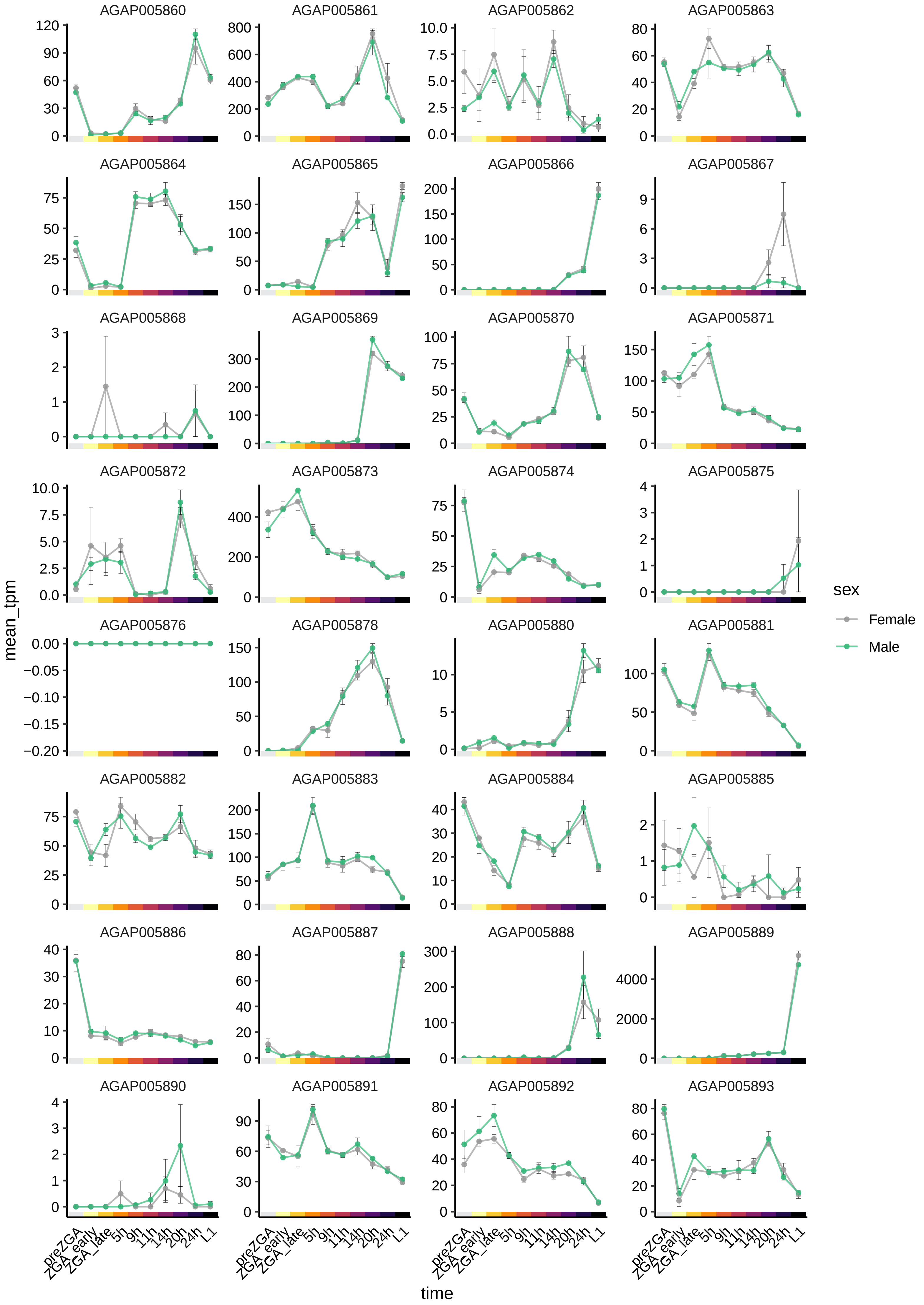
AGAP005860
AGAP005861
AGAP005862
AGAP005863
AGAP005864
AGAP005865
AGAP005866
AGAP005867
AGAP005868
AGAP005869
AGAP005870
AGAP005871
AGAP005872
AGAP005873
AGAP005874
AGAP005875
AGAP005876
AGAP005878
AGAP005880
AGAP005881
AGAP005882
AGAP005883
AGAP005884
AGAP005885
AGAP005886
AGAP005887
AGAP005888
AGAP005889
AGAP005890
AGAP005891
AGAP005892
AGAP005893
mean_tpm
time
preZGA
ZGA_early
ZGA_late
5h
9h
11h
14h
20h
24h
L1
sex
Female
Male

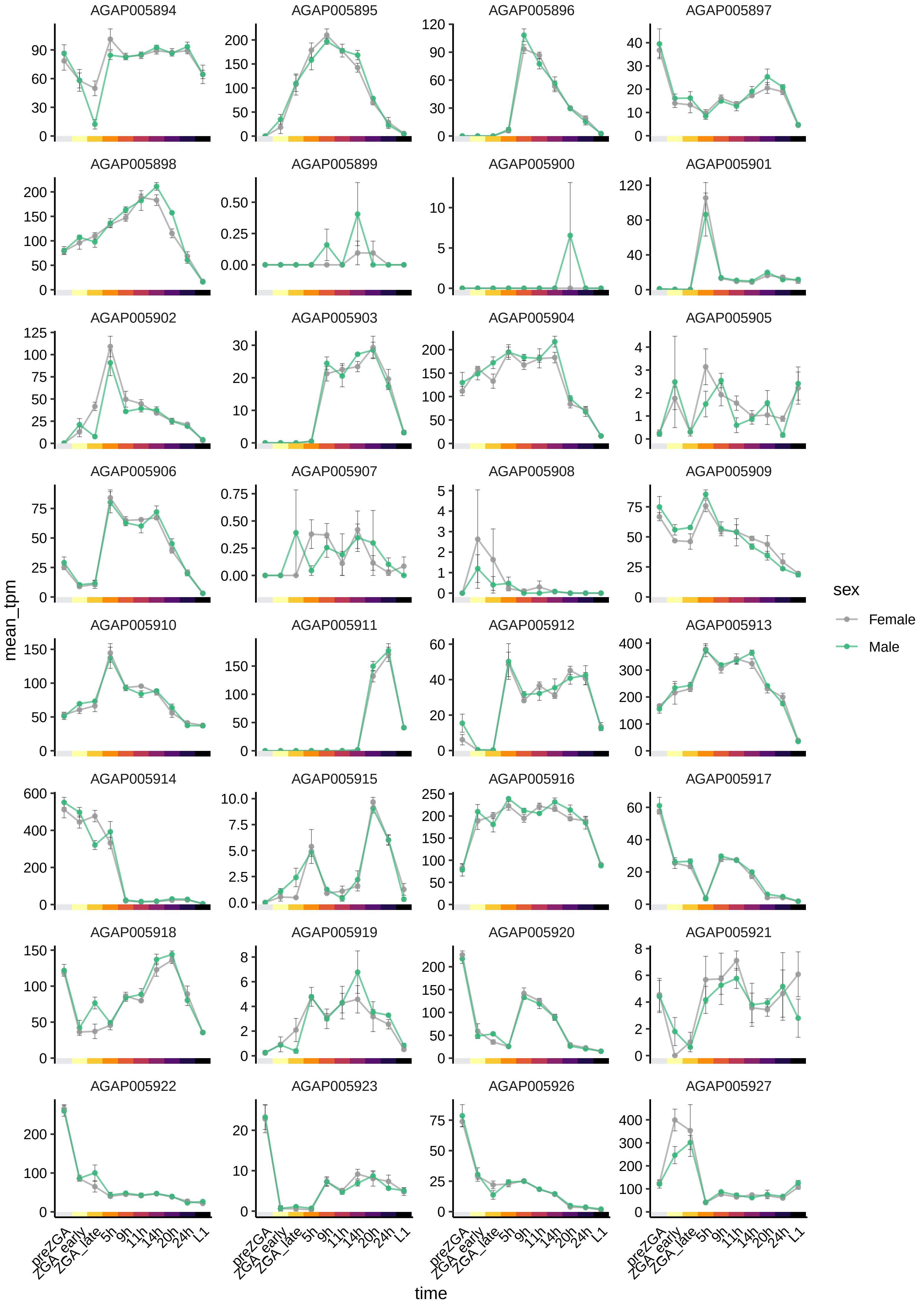

AGAP005894
AGAP005895
AGAP005896
AGAP005897
AGAP005898
AGAP005899
AGAP005900
AGAP005901
AGAP005902
AGAP005903
AGAP005904
AGAP005905
AGAP005906
AGAP005907
AGAP005908
AGAP005909
AGAP005910
AGAP005911
AGAP005912
AGAP005913
AGAP005914
AGAP005915
AGAP005916
AGAP005917
AGAP005918
AGAP005919
AGAP005920
AGAP005921
AGAP005922
AGAP005923
AGAP005926
AGAP005927
mean_tpm
time
preZGA
ZGA_early
ZGA_late
5h
9h
11h
14h
20h
24h
L1
sex
Female
Male

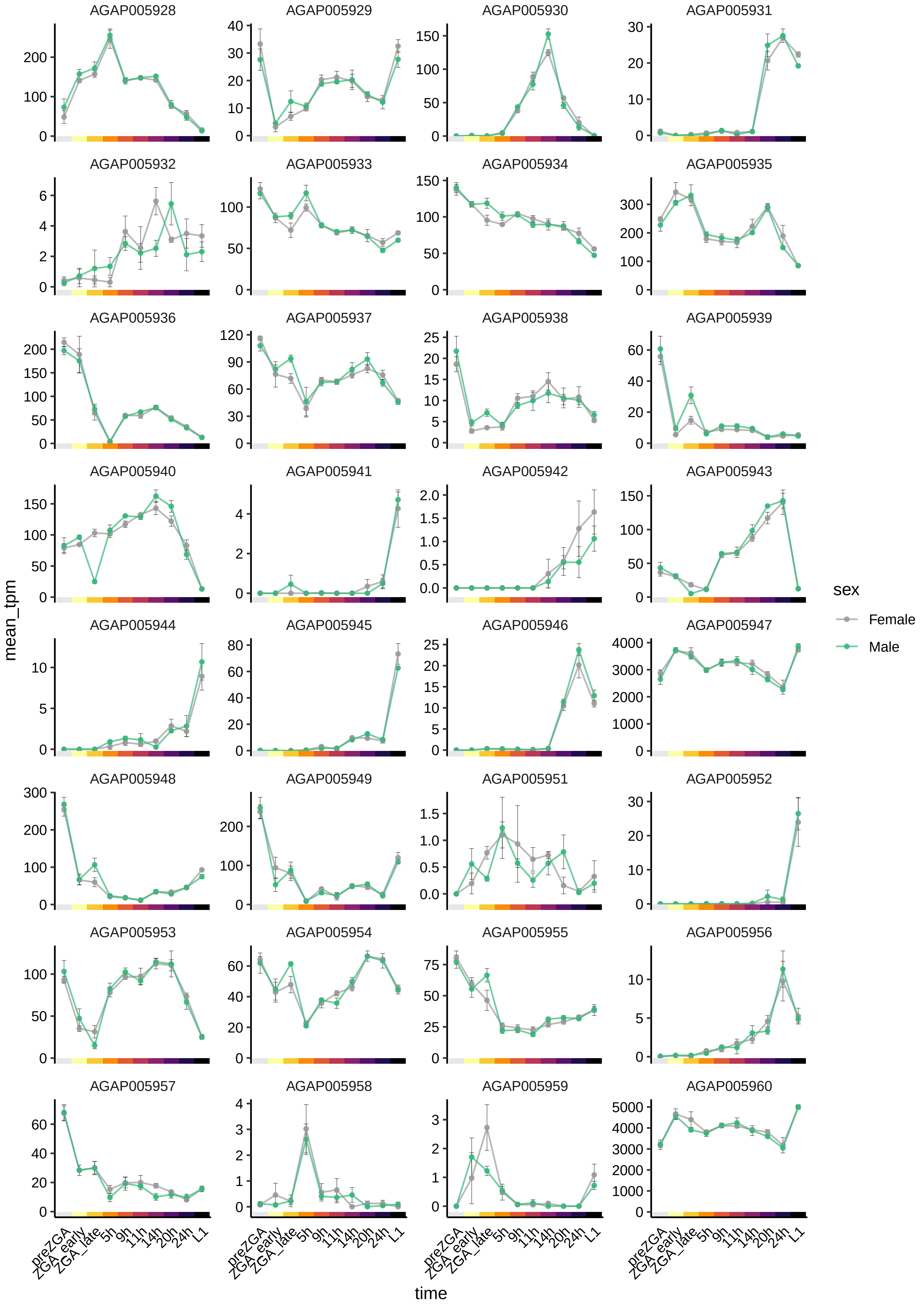

AGAP005928
AGAP005929
AGAP005930
AGAP005931
AGAP005932
AGAP005933
AGAP005934
AGAP005935
AGAP005936
AGAP005937
AGAP005938
AGAP005939
AGAP005940
AGAP005941
AGAP005942
AGAP005943
AGAP005944
AGAP005945
AGAP005946
AGAP005947
AGAP005948
AGAP005949
AGAP005951
AGAP005952
AGAP005953
AGAP005954
AGAP005955
AGAP005956
AGAP005957
AGAP005958
AGAP005959
AGAP005960
mean_tpm
time
preZGA
ZGA_early
ZGA_late
5h
9h
11h
14h
20h
24h
L1
sex
Female
Male

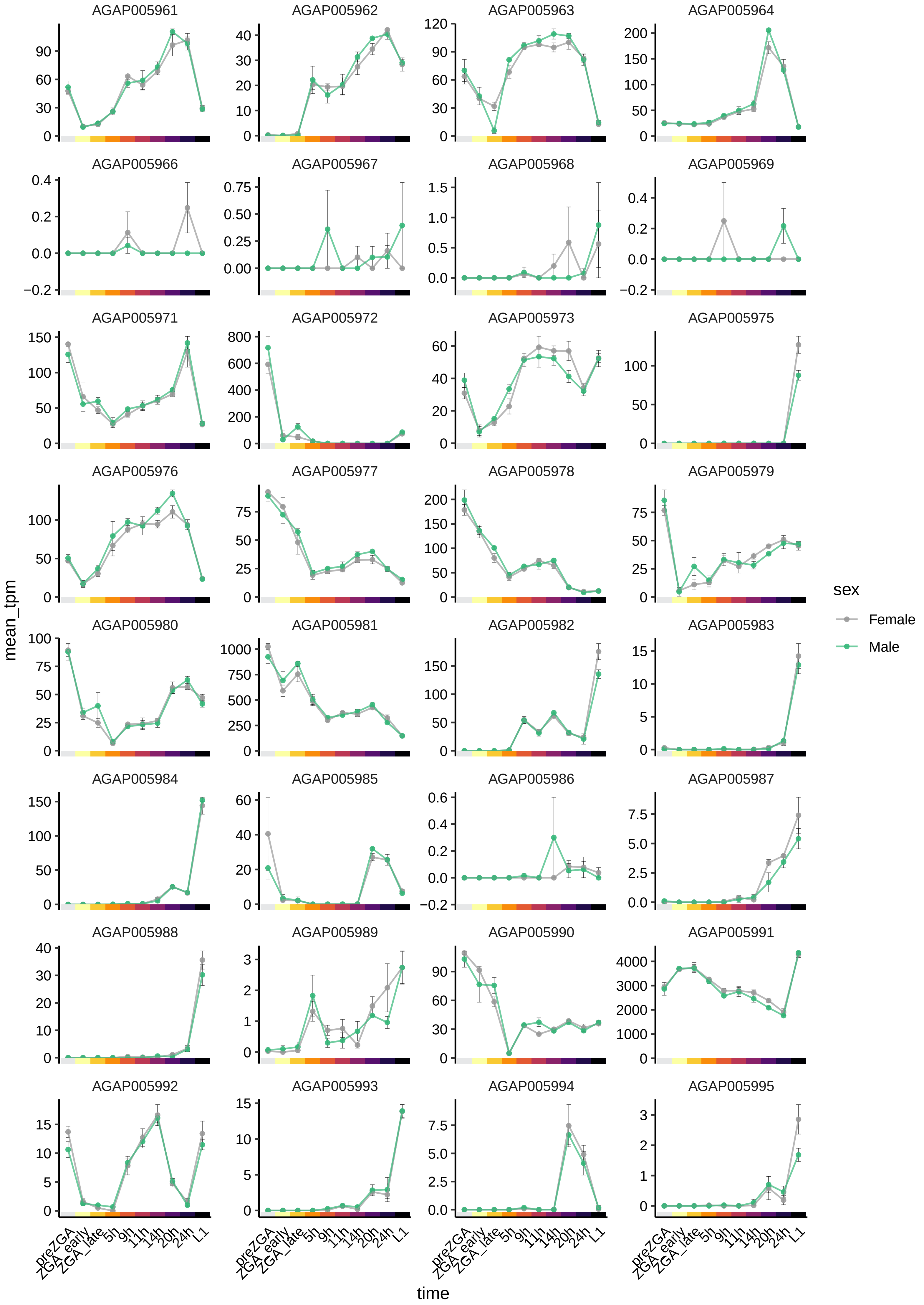

AGAP005961
AGAP005962
AGAP005963
AGAP005964
AGAP005966
AGAP005967
AGAP005968
AGAP005969
AGAP005971
AGAP005972
AGAP005973
AGAP005975
AGAP005976
AGAP005977
AGAP005978
AGAP005979
AGAP005980
AGAP005981
AGAP005982
AGAP005983
AGAP005984
AGAP005985
AGAP005986
AGAP005987
AGAP005988
AGAP005989
AGAP005990
AGAP005991
AGAP005992
AGAP005993
AGAP005994
AGAP005995
mean_tpm
time
preZGA
ZGA_early
ZGA_late
5h
9h
11h
14h
20h
24h
L1
sex
Female
Male

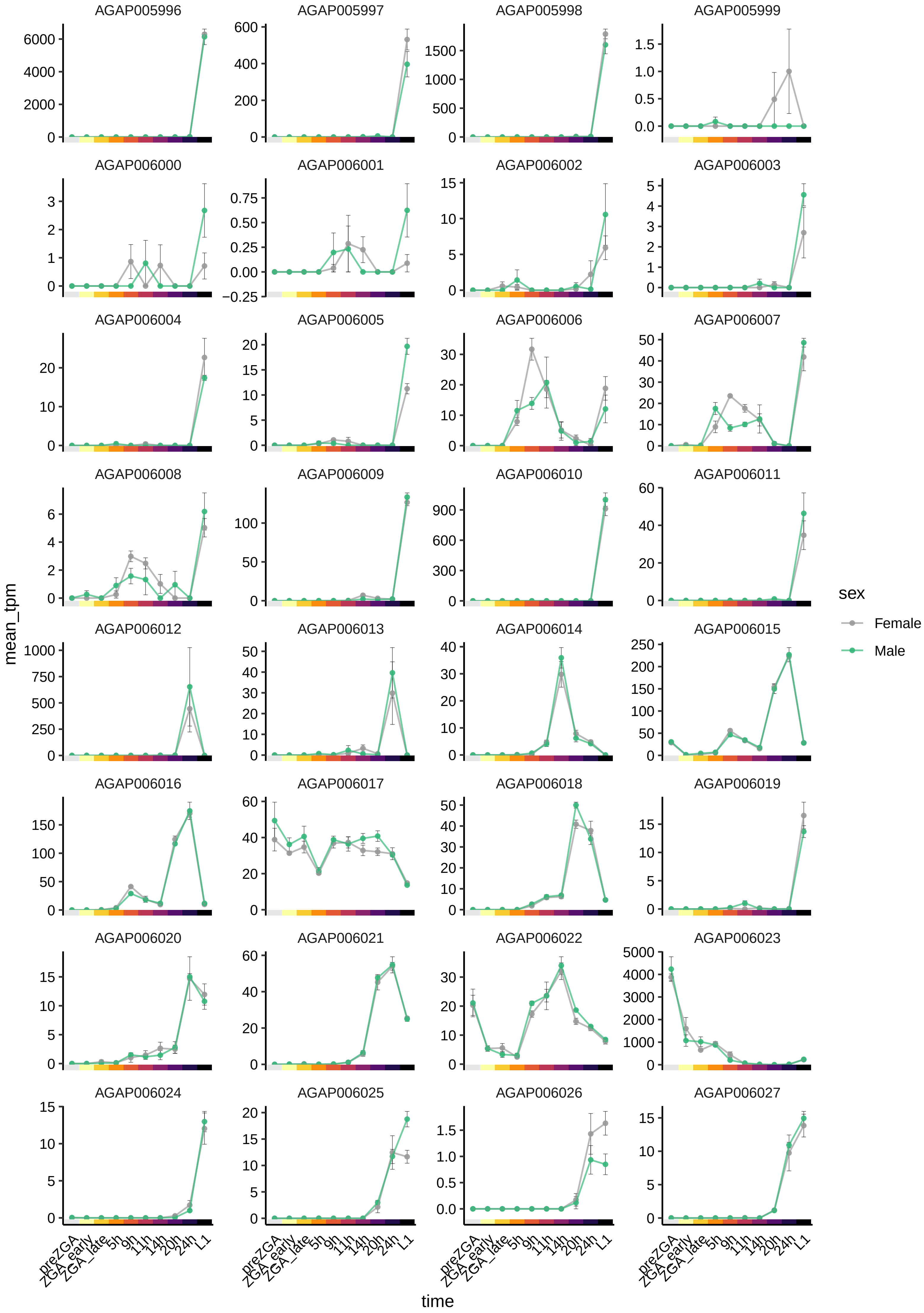

AGAP005996
AGAP005997
AGAP005998
AGAP005999
AGAP006000
AGAP006001
AGAP006002
AGAP006003
AGAP006004
AGAP006005
AGAP006006
AGAP006007
AGAP006008
AGAP006009
AGAP006010
AGAP006011
AGAP006012
AGAP006013
AGAP006014
AGAP006015
AGAP006016
AGAP006017
AGAP006018
AGAP006019
AGAP006020
AGAP006021
AGAP006022
AGAP006023
AGAP006024
AGAP006025
AGAP006026
AGAP006027
mean_tpm
time
preZGA
ZGA_early
ZGA_late
5h
9h
11h
14h
20h
24h
L1
sex
Female
Male

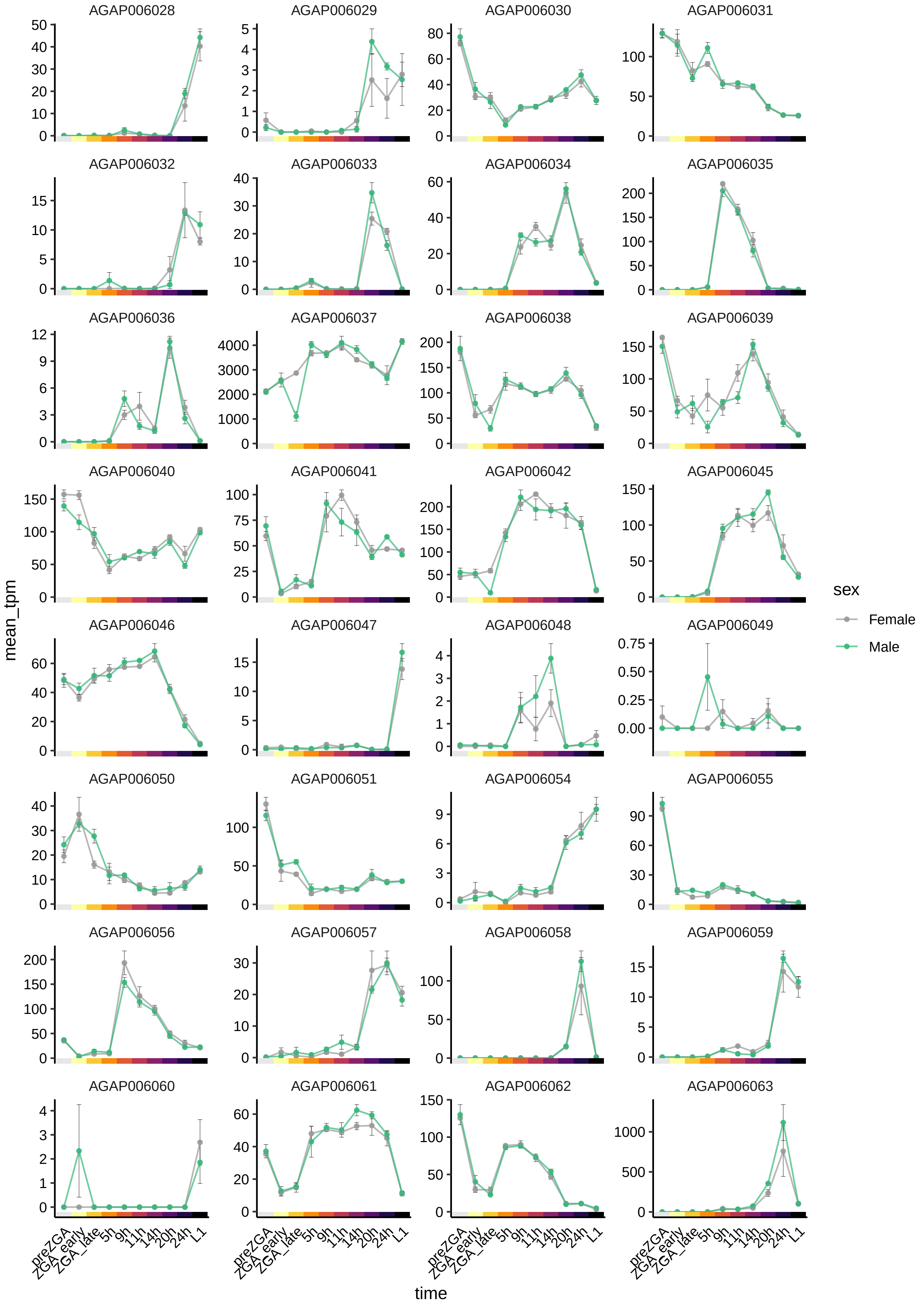

AGAP006028
AGAP006029
AGAP006030
AGAP006031
AGAP006032
AGAP006033
AGAP006034
AGAP006035
AGAP006036
AGAP006037
AGAP006038
AGAP006039
AGAP006040
AGAP006041
AGAP006042
AGAP006045
AGAP006046
AGAP006047
AGAP006048
AGAP006049
AGAP006050
AGAP006051
AGAP006054
AGAP006055
AGAP006056
AGAP006057
AGAP006058
AGAP006059
AGAP006060
AGAP006061
AGAP006062
AGAP006063
mean_tpm
time
preZGA
ZGA_early
ZGA_late
5h
9h
11h
14h
20h
24h
L1
sex
Female
Male

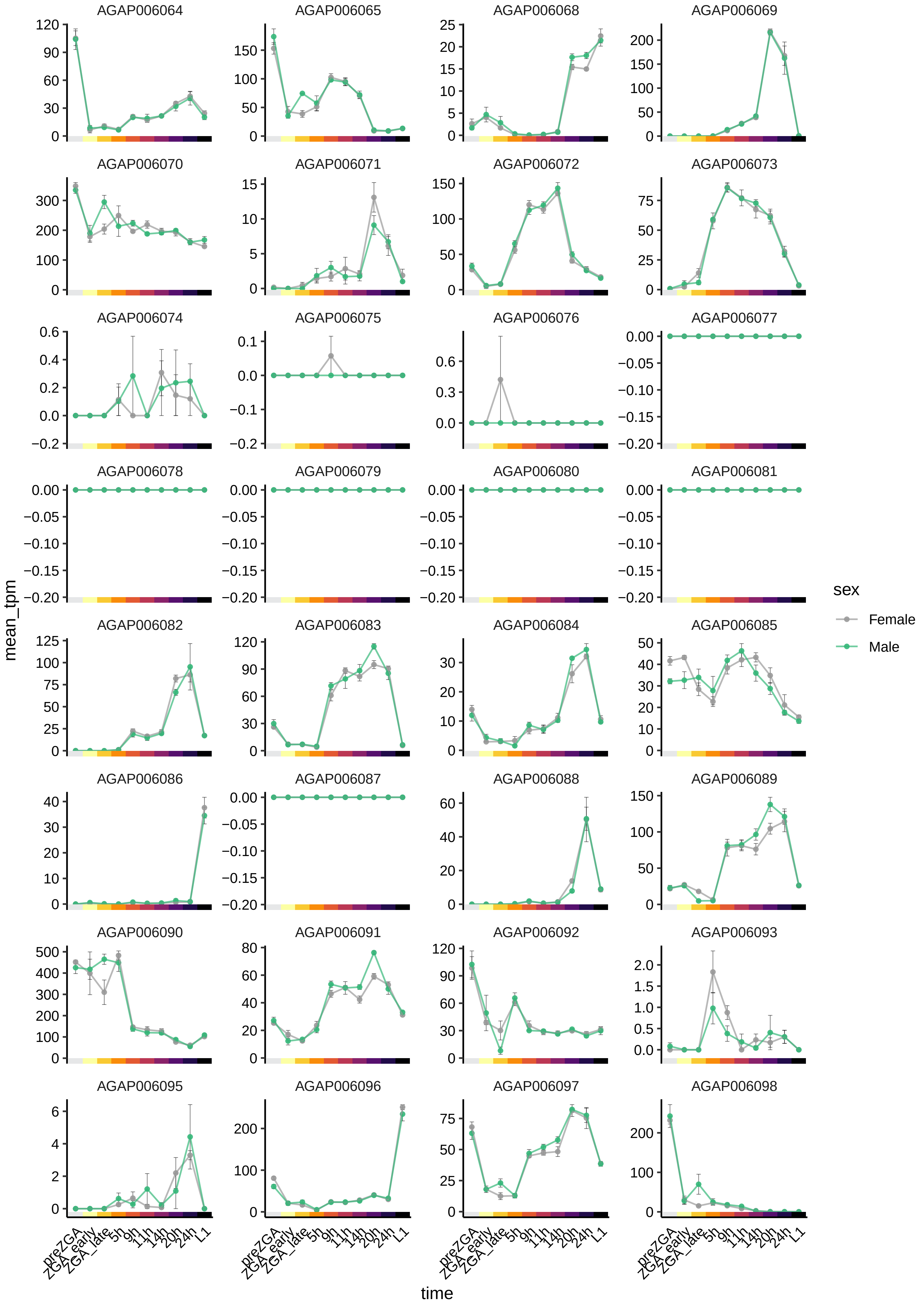
AGAP006064
AGAP006065
AGAP006068
AGAP006069
AGAP006070
AGAP006071
AGAP006072
AGAP006073
AGAP006074
AGAP006075
AGAP006076
AGAP006077
AGAP006078
AGAP006079
AGAP006080
AGAP006081
AGAP006082
AGAP006083
AGAP006084
AGAP006085
AGAP006086
AGAP006087
AGAP006088
AGAP006089
AGAP006090
AGAP006091
AGAP006092
AGAP006093
AGAP006095
AGAP006096
AGAP006097
AGAP006098
mean_tpm
time
preZGA
ZGA_early
ZGA_late
5h
9h
11h
14h
20h
24h
L1
sex
Female
Male

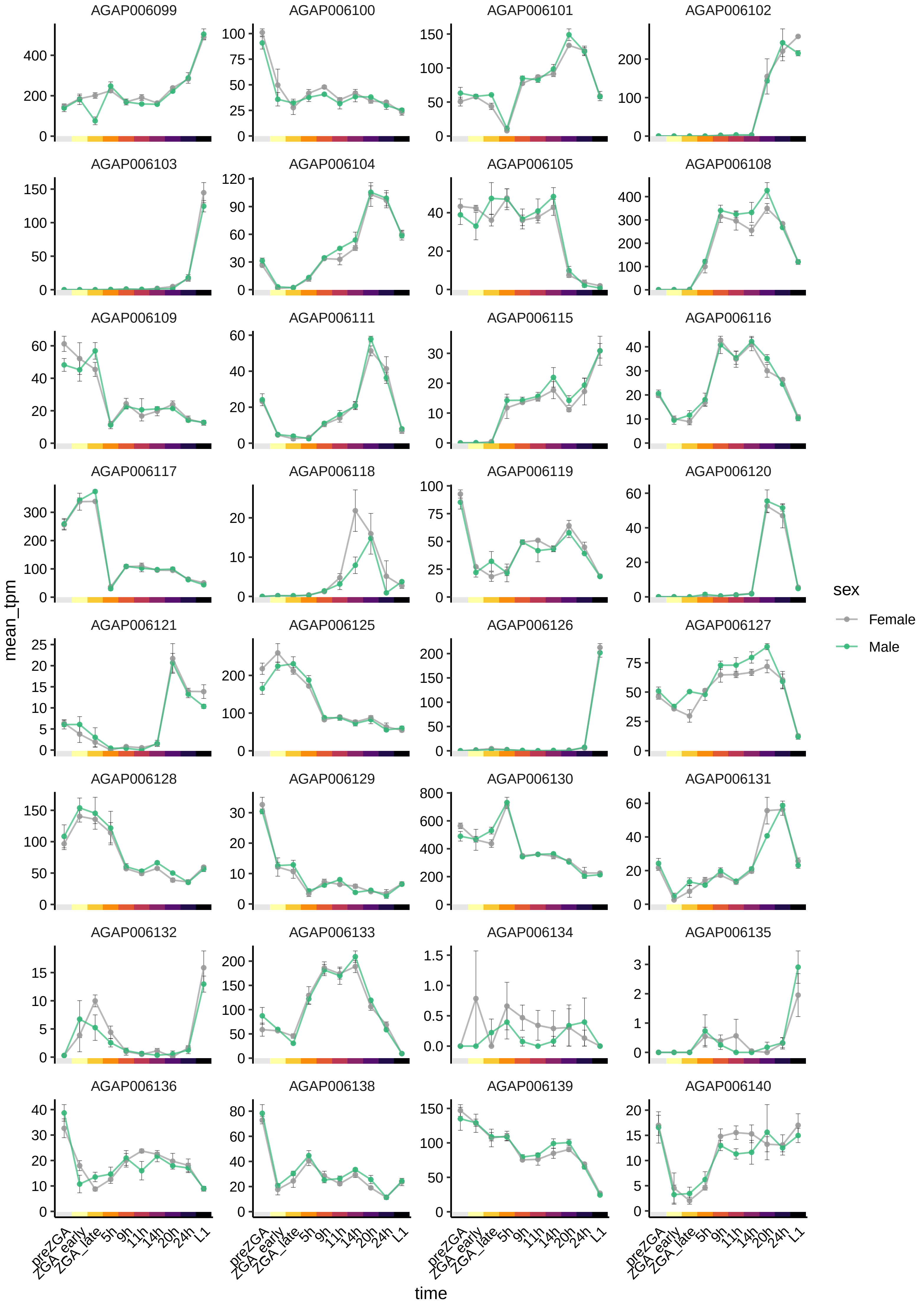
AGAP006099
AGAP006100
AGAP006101
AGAP006102
AGAP006103
AGAP006104
AGAP006105
AGAP006108
AGAP006109
AGAP006111
AGAP006115
AGAP006116
AGAP006117
AGAP006118
AGAP006119
AGAP006120
AGAP006121
AGAP006125
AGAP006126
AGAP006127
AGAP006128
AGAP006129
AGAP006130
AGAP006131
AGAP006132
AGAP006133
AGAP006134
AGAP006135
AGAP006136
AGAP006138
AGAP006139
AGAP006140
mean_tpm
time
preZGA
ZGA_early
ZGA_late
5h
9h
11h
14h
20h
24h
L1
sex
Female
Male

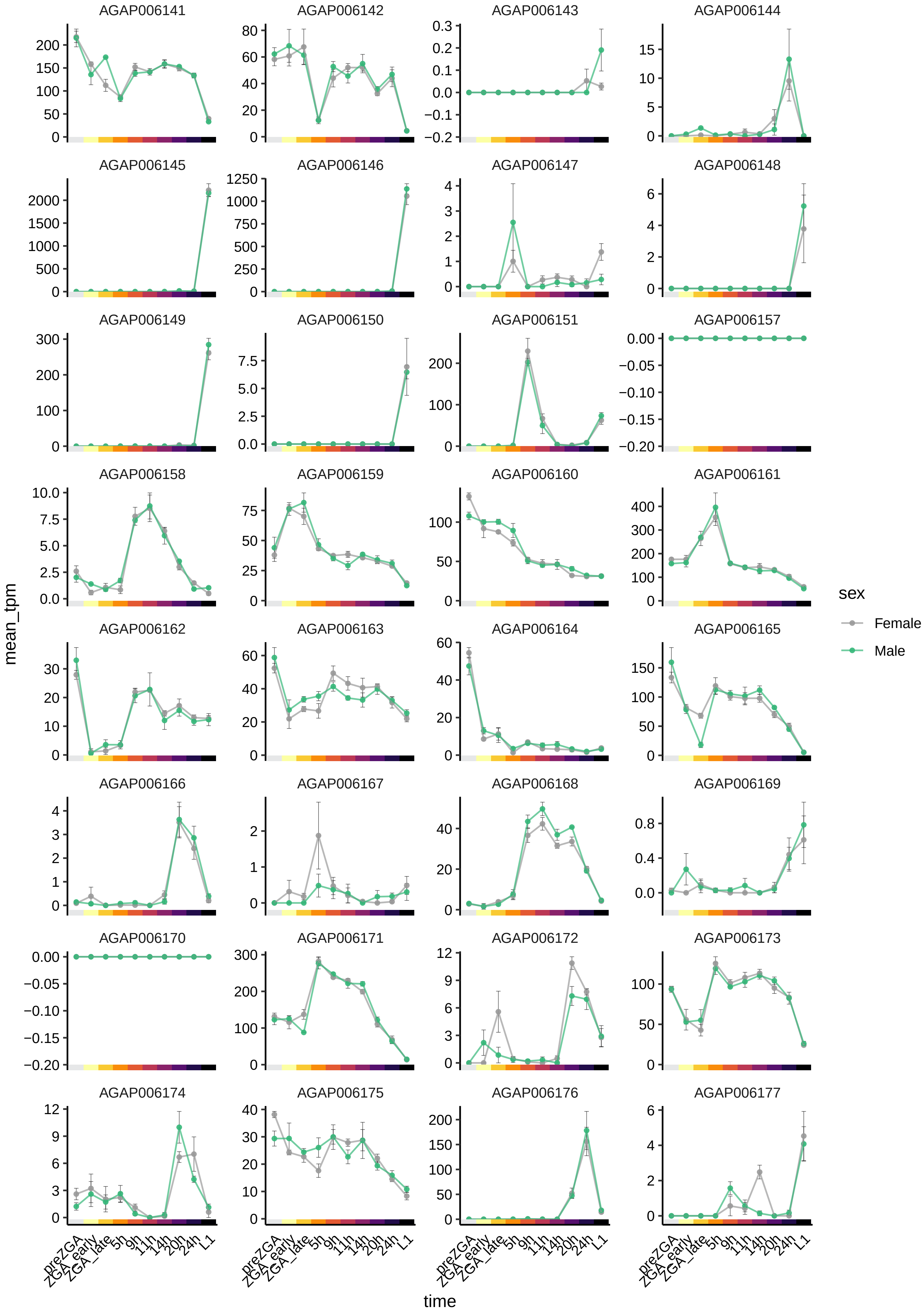
AGAP006141
AGAP006142
AGAP006143
AGAP006144
AGAP006145
AGAP006146
AGAP006147
AGAP006148
AGAP006149
AGAP006150
AGAP006151
AGAP006157
AGAP006158
AGAP006159
AGAP006160
AGAP006161
AGAP006162
AGAP006163
AGAP006164
AGAP006165
AGAP006166
AGAP006167
AGAP006168
AGAP006169
AGAP006170
AGAP006171
AGAP006172
AGAP006173
AGAP006174
AGAP006175
AGAP006176
AGAP006177
mean_tpm
time
preZGA
ZGA_early
ZGA_late
5h
9h
11h
14h
20h
24h
L1
sex
Female
Male

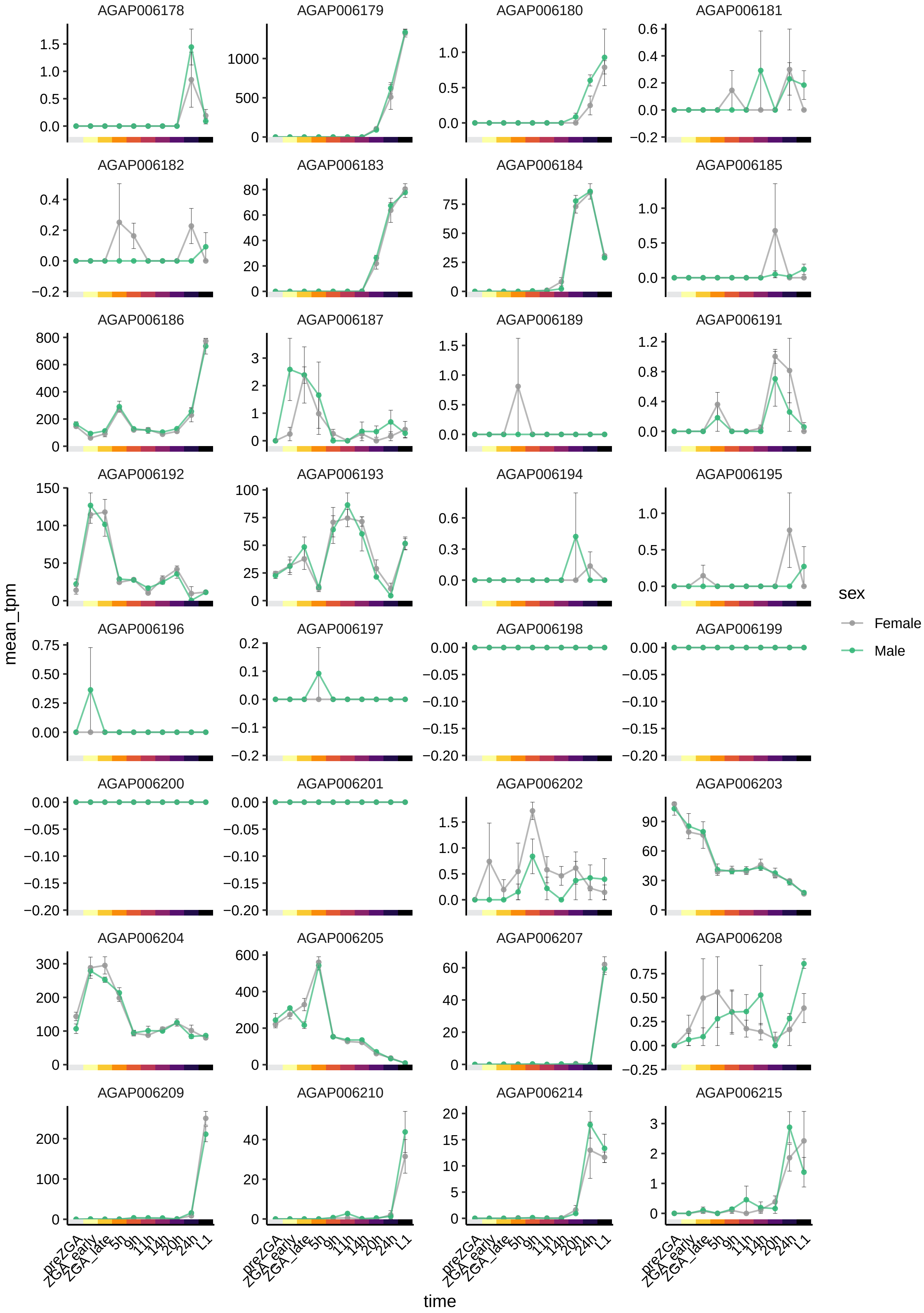
AGAP006178
AGAP006179
AGAP006180
AGAP006181
AGAP006182
AGAP006183
AGAP006184
AGAP006185
AGAP006186
AGAP006187
AGAP006189
AGAP006191
AGAP006192
AGAP006193
AGAP006194
AGAP006195
AGAP006196
AGAP006197
AGAP006198
AGAP006199
AGAP006200
AGAP006201
AGAP006202
AGAP006203
AGAP006204
AGAP006205
AGAP006207
AGAP006208
AGAP006209
AGAP006210
AGAP006214
AGAP006215
mean_tpm
time
preZGA
ZGA_early
ZGA_late
5h
9h
11h
14h
20h
24h
L1
sex
Female
Male

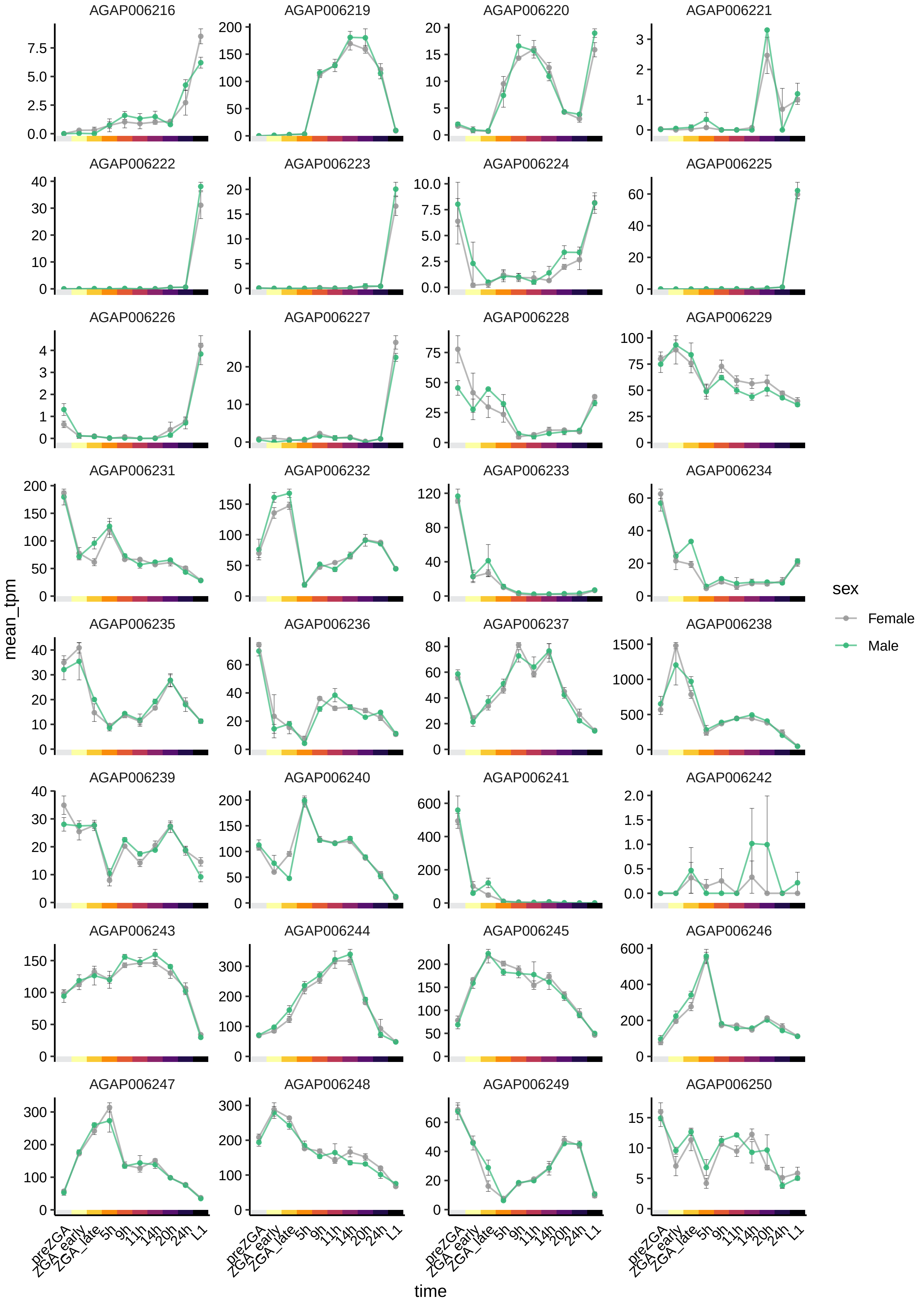
AGAP006216
AGAP006219
AGAP006220
AGAP006221
AGAP006222
AGAP006223
AGAP006224
AGAP006225
AGAP006226
AGAP006227
AGAP006228
AGAP006229
AGAP006231
AGAP006232
AGAP006233
AGAP006234
AGAP006235
AGAP006236
AGAP006237
AGAP006238
AGAP006239
AGAP006240
AGAP006241
AGAP006242
AGAP006243
AGAP006244
AGAP006245
AGAP006246
AGAP006247
AGAP006248
AGAP006249
AGAP006250
mean_tpm
time
preZGA
ZGA_early
ZGA_late
5h
9h
11h
14h
20h
24h
L1
sex
Female
Male

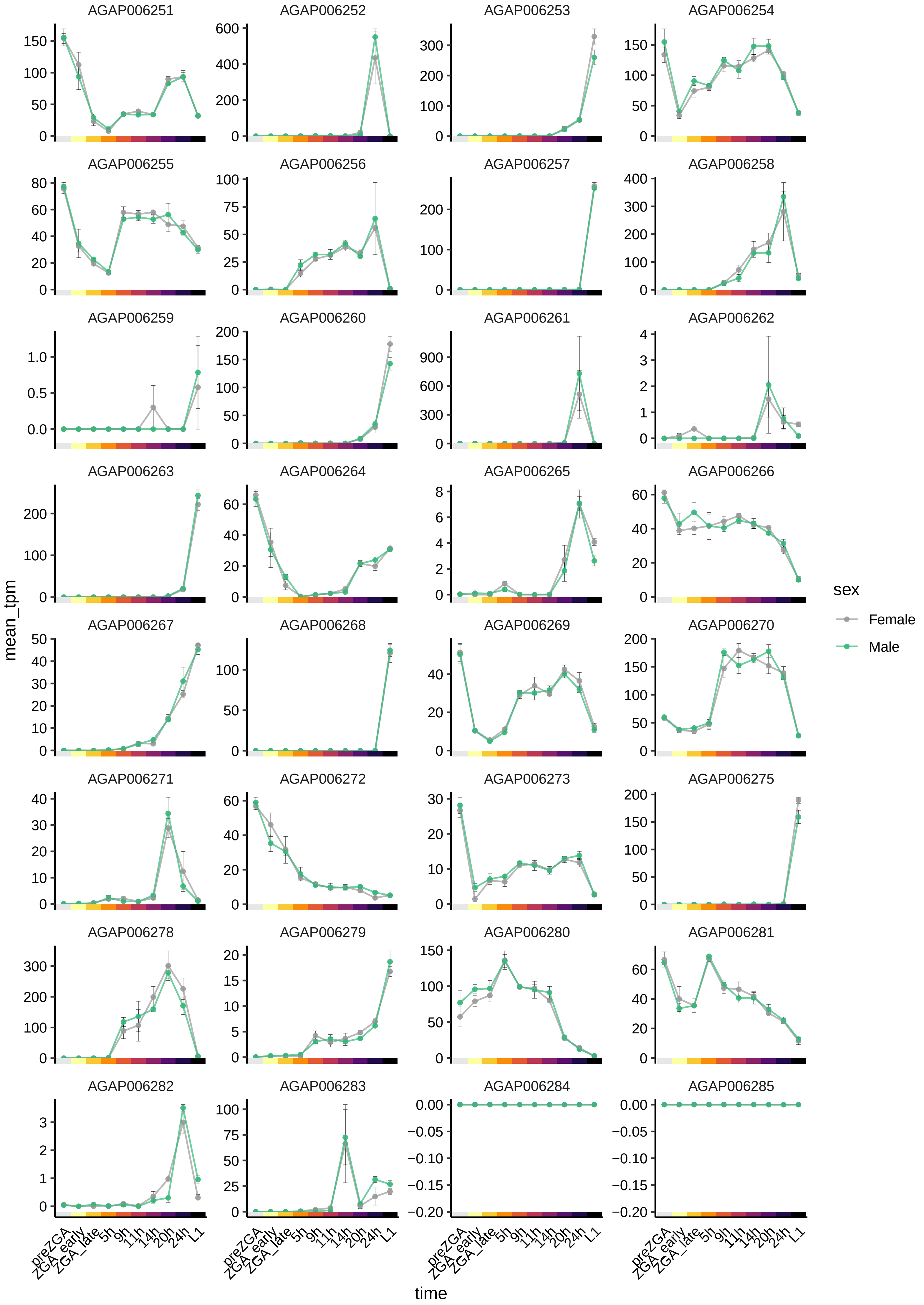
AGAP006251
AGAP006252
AGAP006253
AGAP006254
AGAP006255
AGAP006256
AGAP006257
AGAP006258
AGAP006259
AGAP006260
AGAP006261
AGAP006262
AGAP006263
AGAP006264
AGAP006265
AGAP006266
AGAP006267
AGAP006268
AGAP006269
AGAP006270
AGAP006271
AGAP006272
AGAP006273
AGAP006275
AGAP006278
AGAP006279
AGAP006280
AGAP006281
AGAP006282
AGAP006283
AGAP006284
AGAP006285
mean_tpm
time
preZGA
ZGA_early
ZGA_late
5h
9h
11h
14h
20h
24h
L1
sex
Female
Male

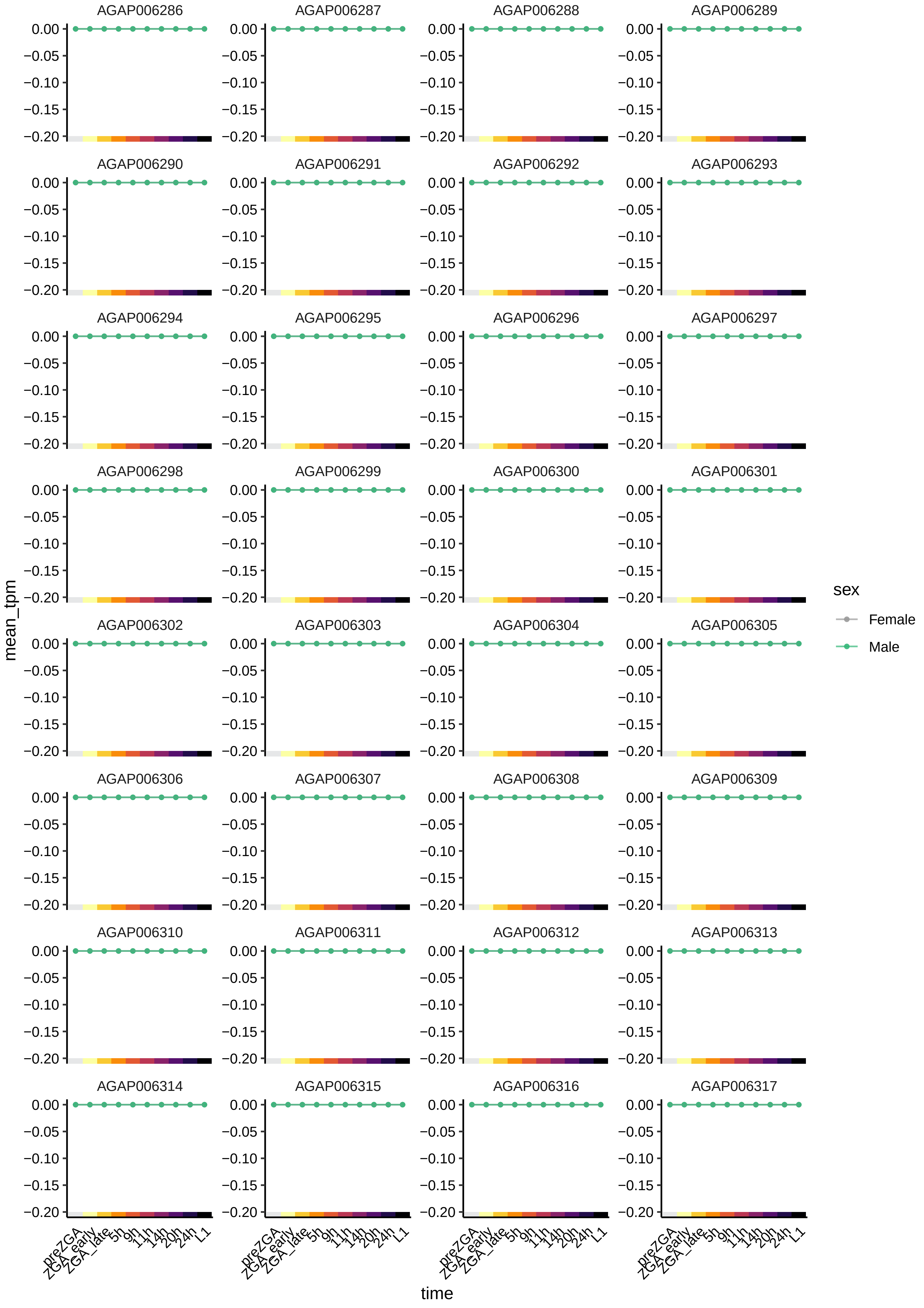

AGAP006286
AGAP006287
AGAP006288
AGAP006289
AGAP006290
AGAP006291
AGAP006292
AGAP006293
AGAP006294
AGAP006295
AGAP006296
AGAP006297
AGAP006298
AGAP006299
AGAP006300
AGAP006301
AGAP006302
AGAP006303
AGAP006304
AGAP006305
AGAP006306
AGAP006307
AGAP006308
AGAP006309
AGAP006310
AGAP006311
AGAP006312
AGAP006313
AGAP006314
AGAP006315
AGAP006316
AGAP006317
0.00
−0.05
−0.10
−0.15
−0.20
mean_tpm
preZGA
ZGA_early
ZGA_late
5h
9h
11h
14h
20h
24h
L1
time
sex
Female
Male

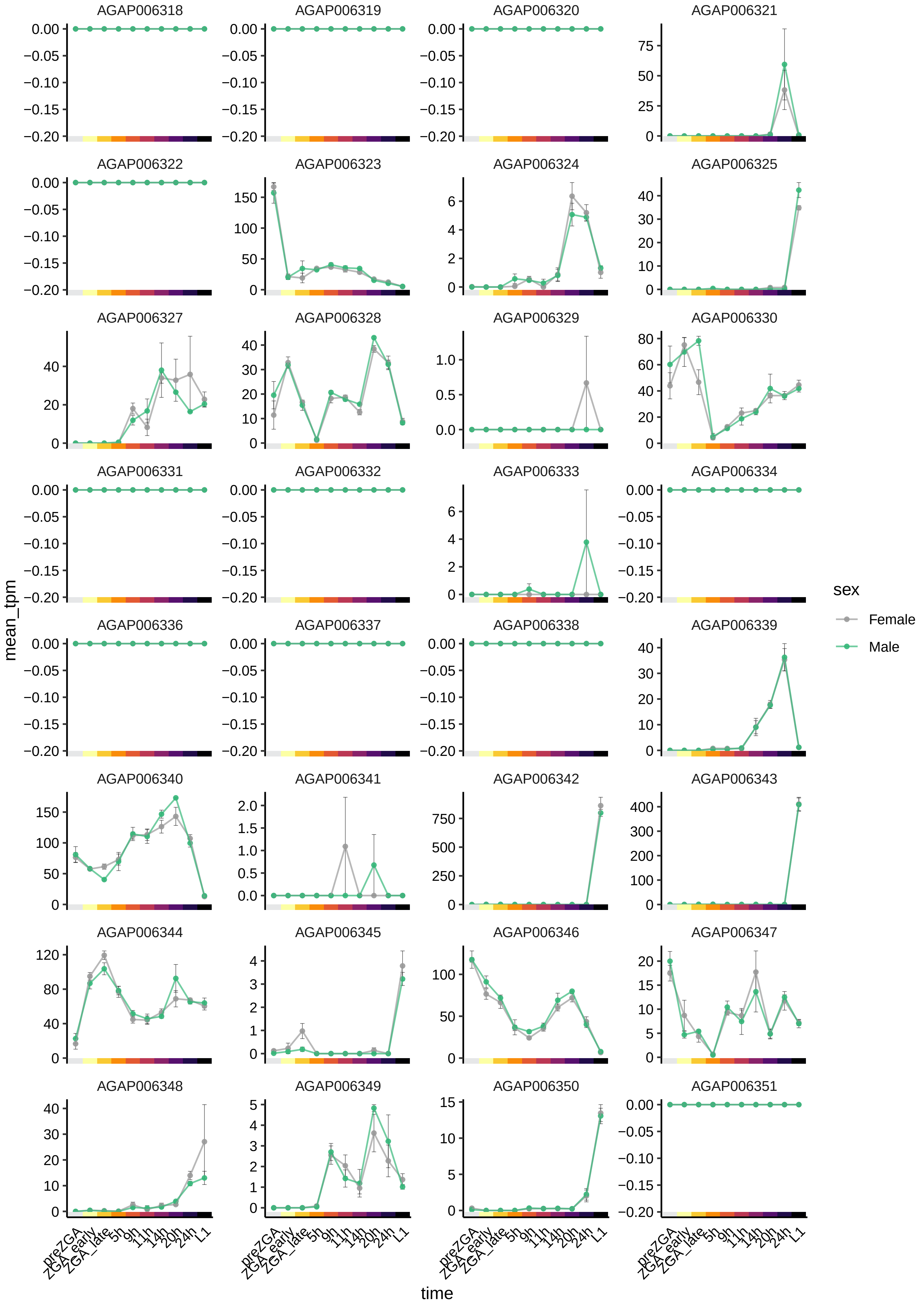

AGAP006318
AGAP006319
AGAP006320
AGAP006321
AGAP006322
AGAP006323
AGAP006324
AGAP006325
AGAP006327
AGAP006328
AGAP006329
AGAP006330
AGAP006331
AGAP006332
AGAP006333
AGAP006334
AGAP006336
AGAP006337
AGAP006338
AGAP006339
AGAP006340
AGAP006341
AGAP006342
AGAP006343
AGAP006344
AGAP006345
AGAP006346
AGAP006347
AGAP006348
AGAP006349
AGAP006350
AGAP006351
mean_tpm
time
preZGA
ZGA_early
ZGA_late
5h
9h
11h
14h
20h
24h
L1
sex
Female
Male

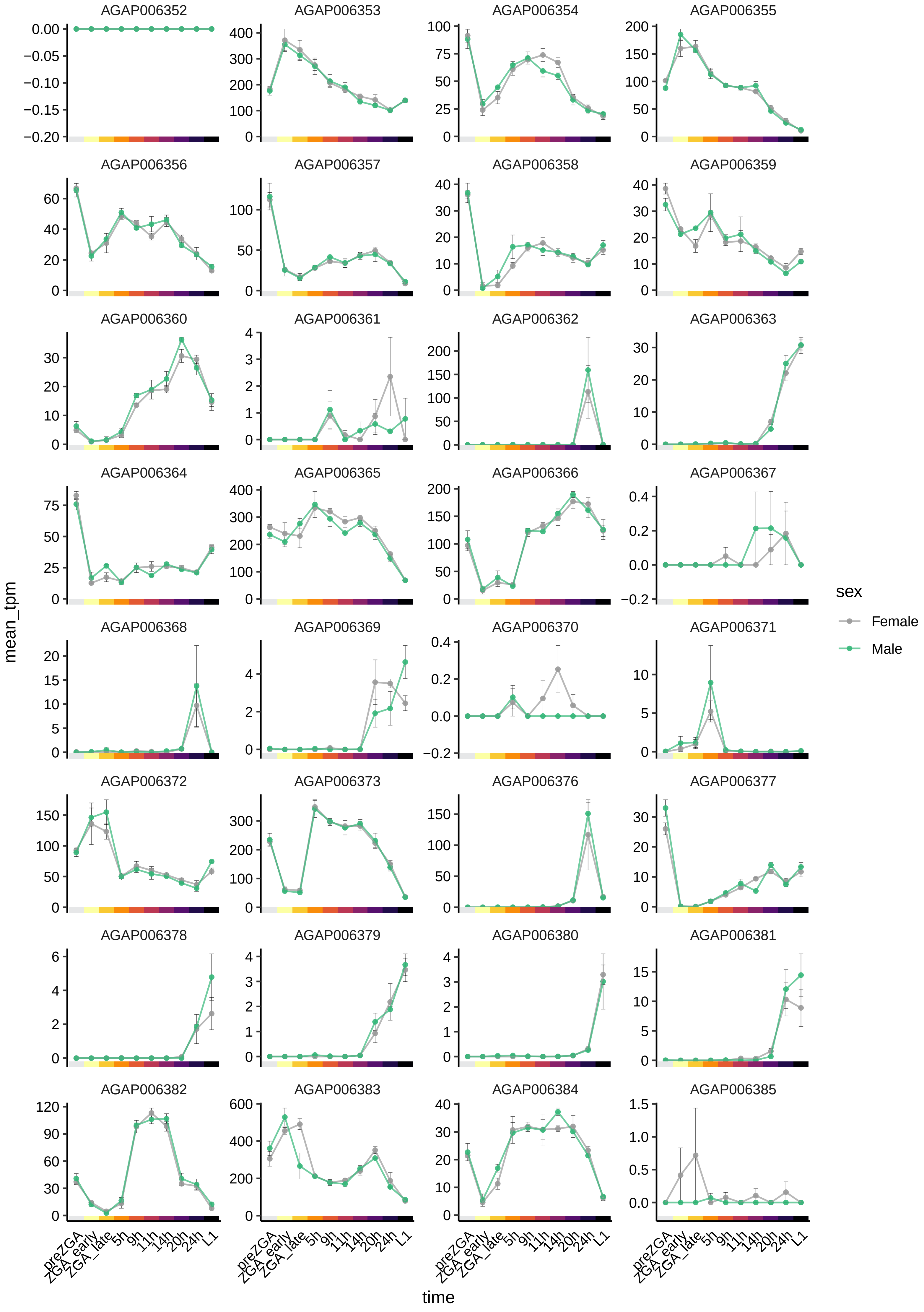

AGAP006352
AGAP006353
AGAP006354
AGAP006355
AGAP006356
AGAP006357
AGAP006358
AGAP006359
AGAP006360
AGAP006361
AGAP006362
AGAP006363
AGAP006364
AGAP006365
AGAP006366
AGAP006367
AGAP006368
AGAP006369
AGAP006370
AGAP006371
AGAP006372
AGAP006373
AGAP006376
AGAP006377
AGAP006378
AGAP006379
AGAP006380
AGAP006381
AGAP006382
AGAP006383
AGAP006384
AGAP006385
mean_tpm
time
preZGA
ZGA_early
ZGA_late
5h
9h
11h
14h
20h
24h
L1
sex
Female
Male

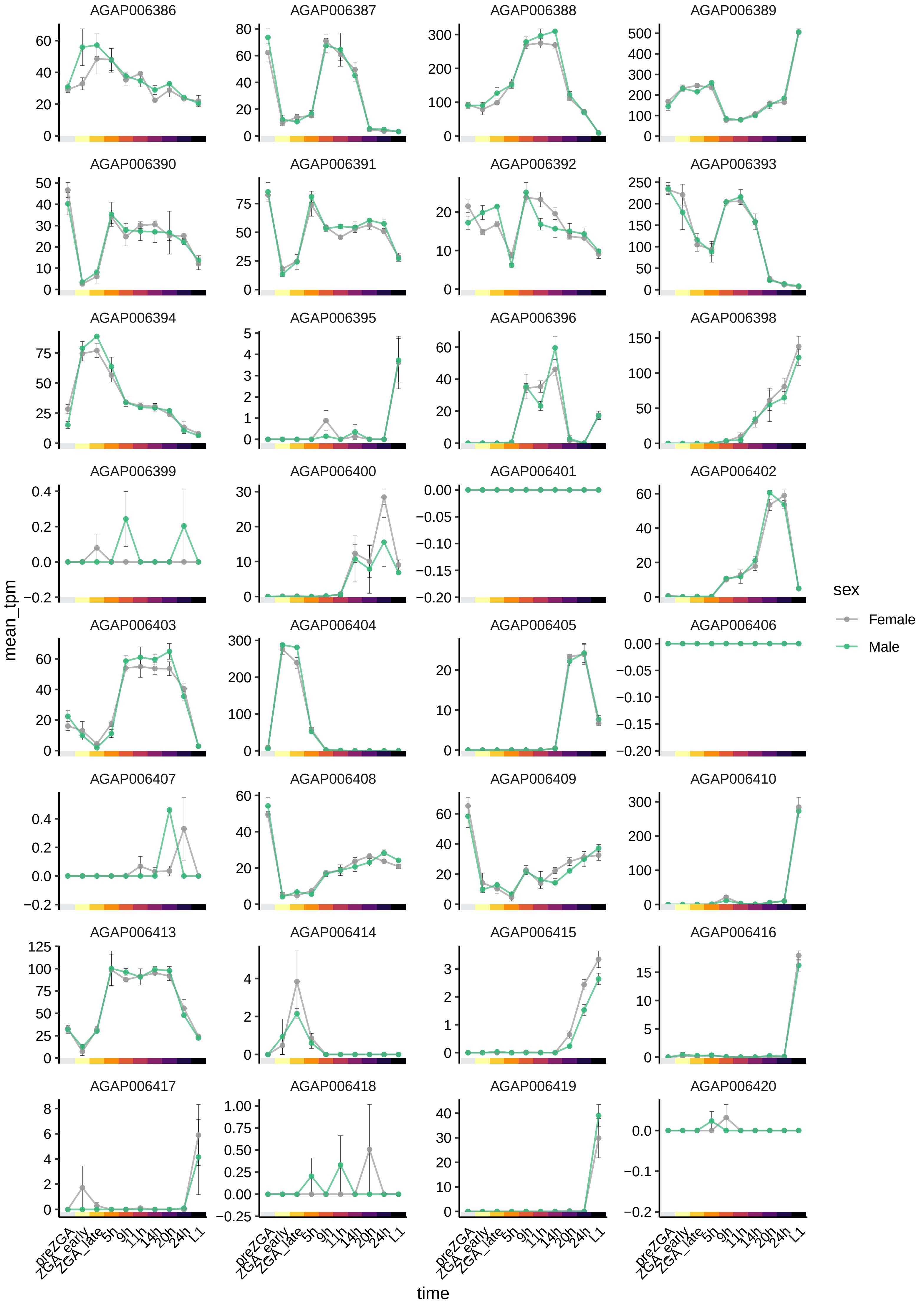

AGAP006386
AGAP006387
AGAP006388
AGAP006389
AGAP006390
AGAP006391
AGAP006392
AGAP006393
AGAP006394
AGAP006395
AGAP006396
AGAP006398
AGAP006399
AGAP006400
AGAP006401
AGAP006402
AGAP006403
AGAP006404
AGAP006405
AGAP006406
AGAP006407
AGAP006408
AGAP006409
AGAP006410
AGAP006413
AGAP006414
AGAP006415
AGAP006416
AGAP006417
AGAP006418
AGAP006419
AGAP006420
mean_tpm
time
preZGA
ZGA_early
ZGA_late
5h
9h
11h
14h
20h
24h
L1
sex
Female
Male

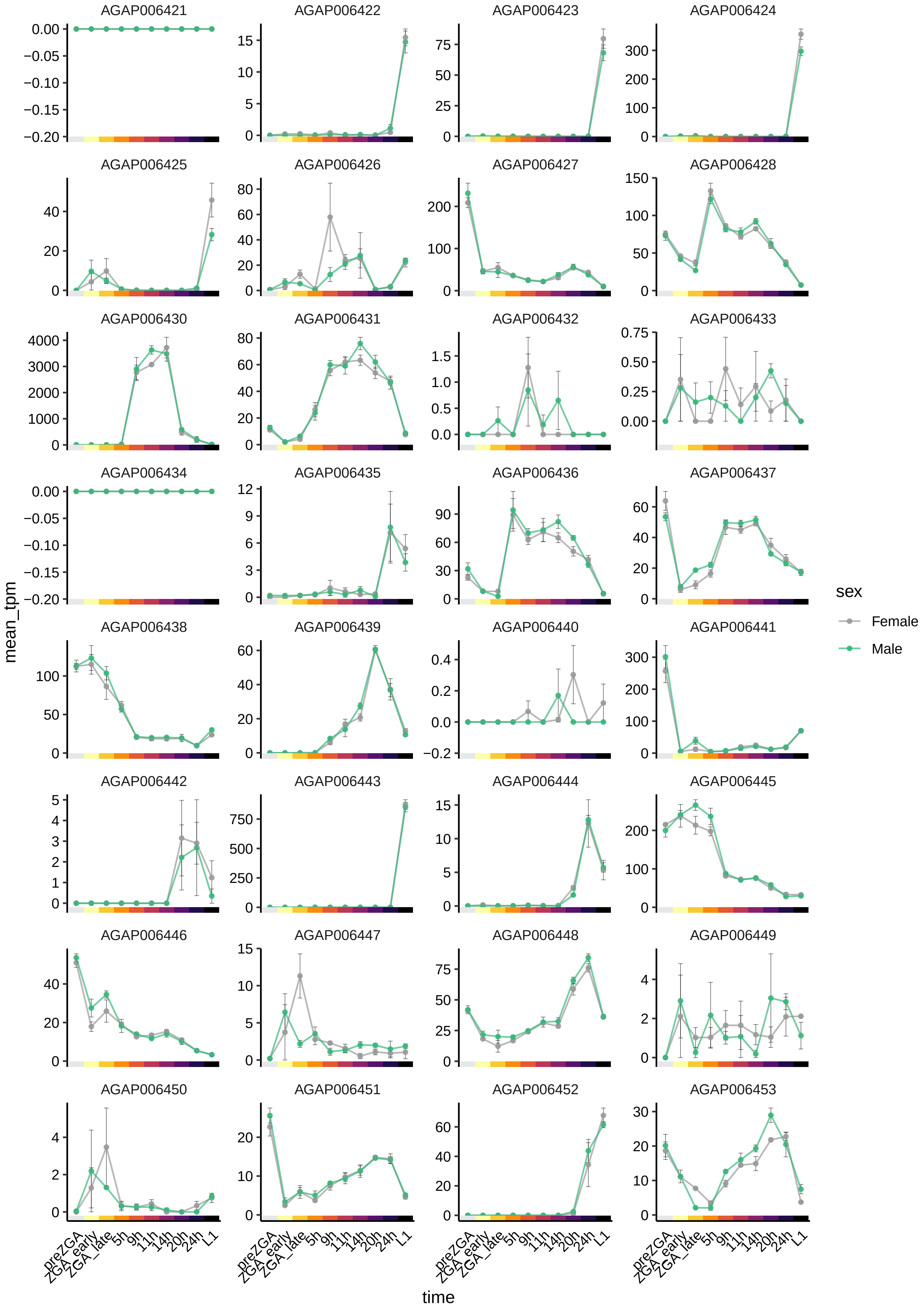
AGAP006421
AGAP006422
AGAP006423
AGAP006424
AGAP006425
AGAP006426
AGAP006427
AGAP006428
AGAP006430
AGAP006431
AGAP006432
AGAP006433
AGAP006434
AGAP006435
AGAP006436
AGAP006437
AGAP006438
AGAP006439
AGAP006440
AGAP006441
AGAP006442
AGAP006443
AGAP006444
AGAP006445
AGAP006446
AGAP006447
AGAP006448
AGAP006449
AGAP006450
AGAP006451
AGAP006452
AGAP006453
mean_tpm
time
preZGA
ZGA_early
ZGA_late
5h
9h
11h
14h
20h
24h
L1
sex
Female
Male

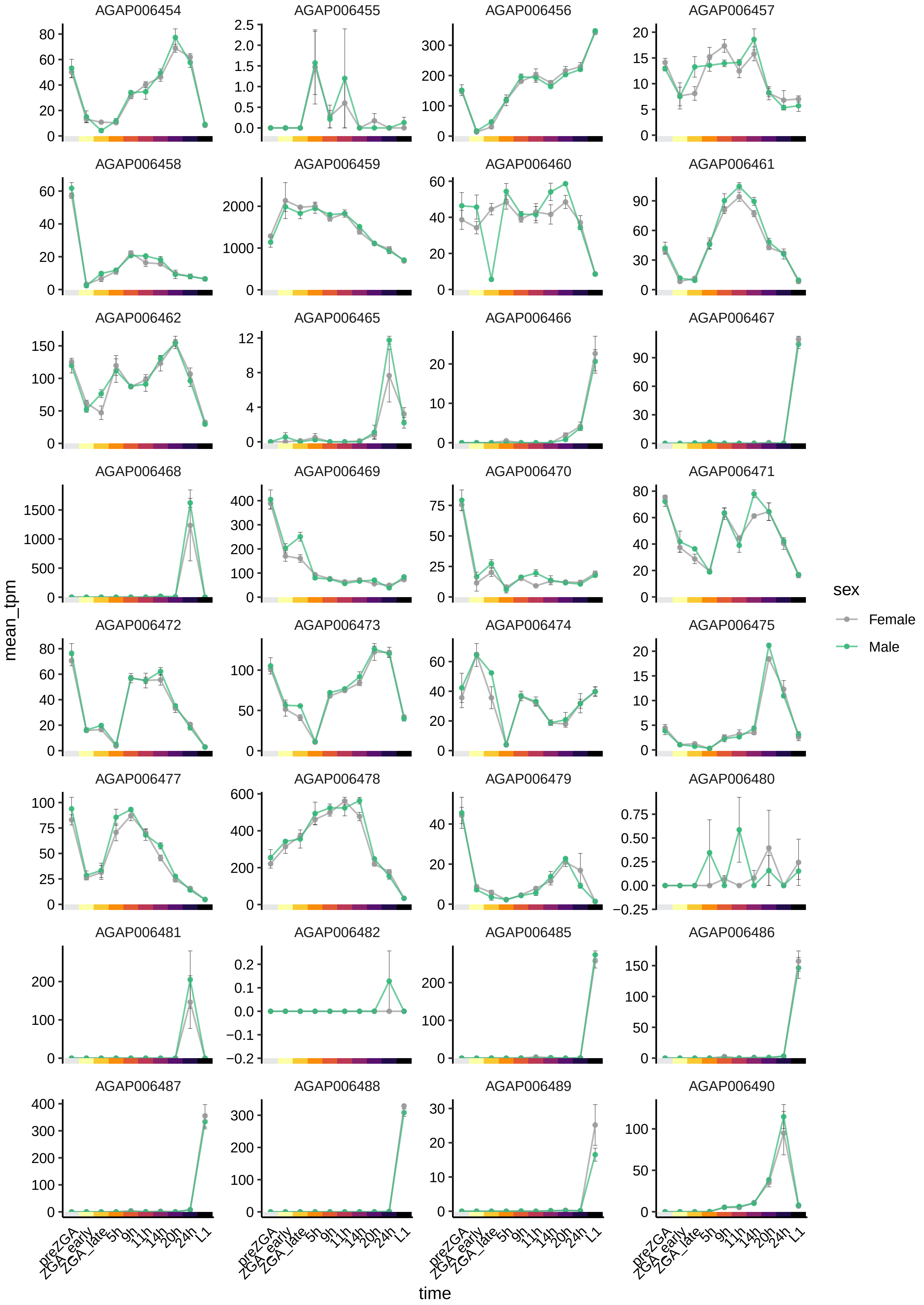

AGAP006454
AGAP006455
AGAP006456
AGAP006457
AGAP006458
AGAP006459
AGAP006460
AGAP006461
AGAP006462
AGAP006465
AGAP006466
AGAP006467
AGAP006468
AGAP006469
AGAP006470
AGAP006471
AGAP006472
AGAP006473
AGAP006474
AGAP006475
AGAP006477
AGAP006478
AGAP006479
AGAP006480
AGAP006481
AGAP006482
AGAP006485
AGAP006486
AGAP006487
AGAP006488
AGAP006489
AGAP006490
mean_tpm
time
preZGA
ZGA_early
ZGA_late
5h
9h
11h
14h
20h
24h
L1
sex
Female
Male

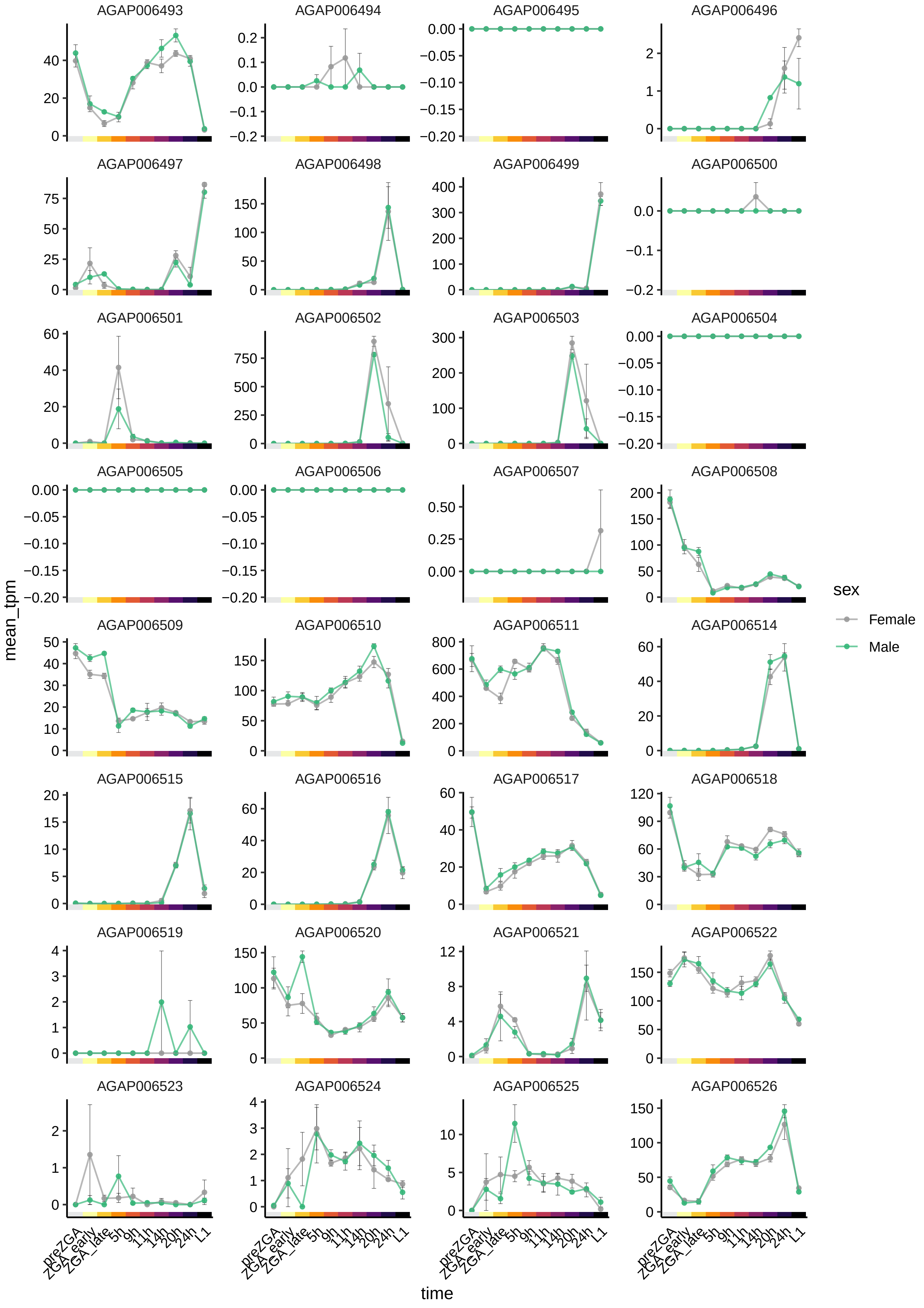

AGAP006493
AGAP006494
AGAP006495
AGAP006496
AGAP006497
AGAP006498
AGAP006499
AGAP006500
AGAP006501
AGAP006502
AGAP006503
AGAP006504
AGAP006505
AGAP006506
AGAP006507
AGAP006508
AGAP006509
AGAP006510
AGAP006511
AGAP006514
AGAP006515
AGAP006516
AGAP006517
AGAP006518
AGAP006519
AGAP006520
AGAP006521
AGAP006522
AGAP006523
AGAP006524
AGAP006525
AGAP006526
mean_tpm
time
preZGA
ZGA_early
ZGA_late
5h
9h
11h
14h
20h
24h
L1
sex
Female
Male

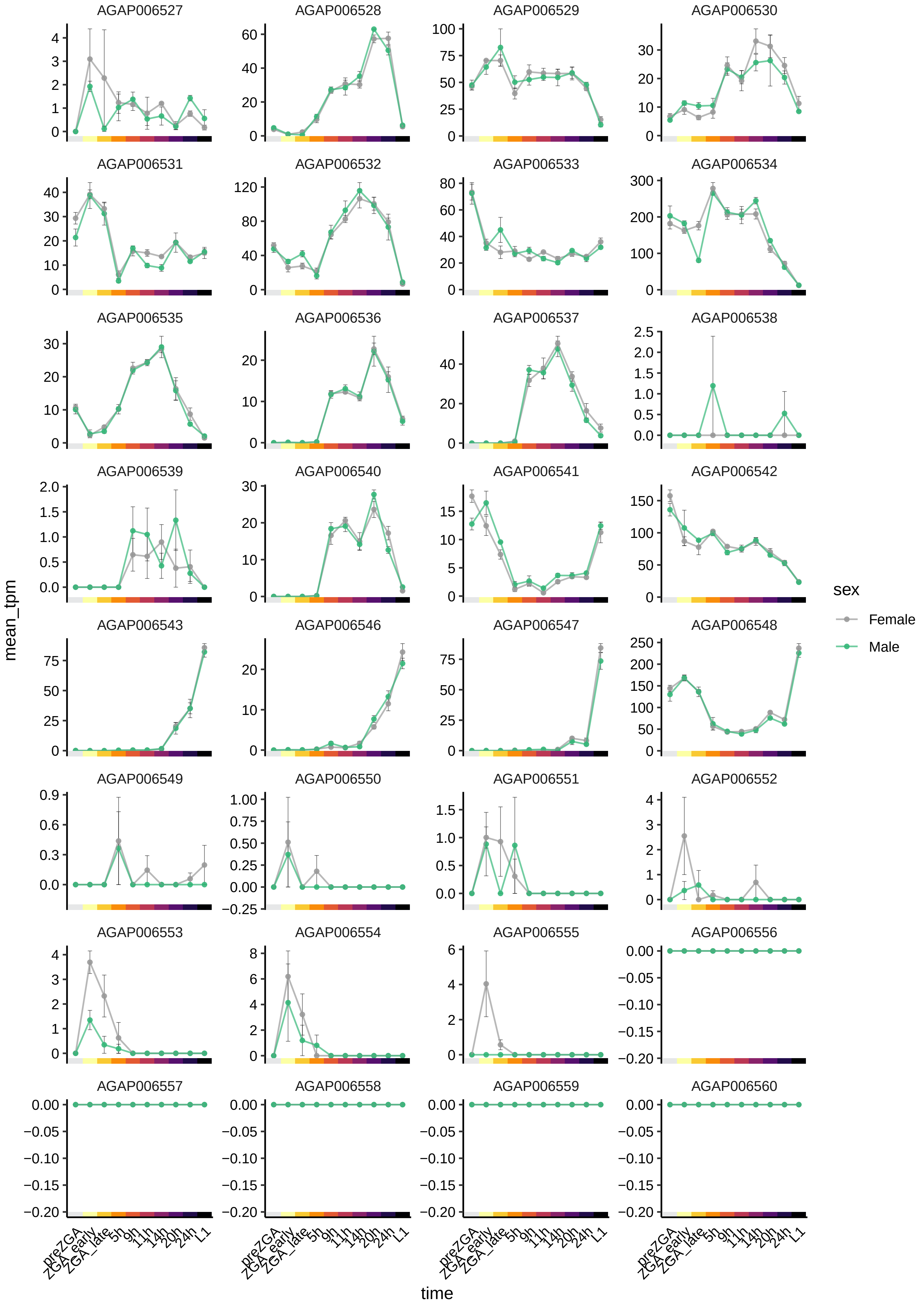
AGAP006527
AGAP006528
AGAP006529
AGAP006530
AGAP006531
AGAP006532
AGAP006533
AGAP006534
AGAP006535
AGAP006536
AGAP006537
AGAP006538
AGAP006539
AGAP006540
AGAP006541
AGAP006542
AGAP006543
AGAP006546
AGAP006547
AGAP006548
AGAP006549
AGAP006550
AGAP006551
AGAP006552
AGAP006553
AGAP006554
AGAP006555
AGAP006556
AGAP006557
AGAP006558
AGAP006559
AGAP006560
mean_tpm
time
preZGA
ZGA_early
ZGA_late
5h
9h
11h
14h
20h
24h
L1
sex
Female
Male

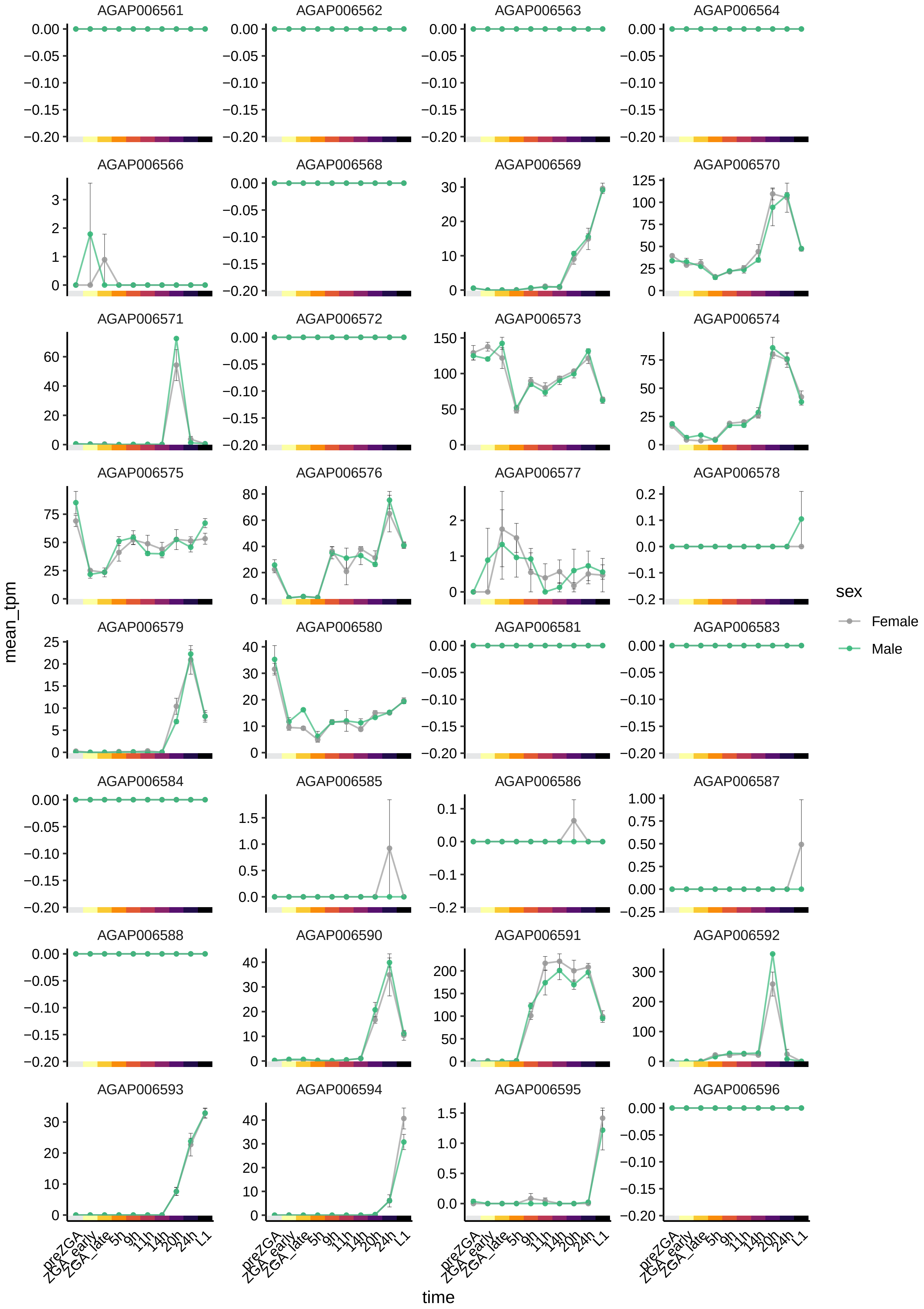

AGAP006561
AGAP006562
AGAP006563
AGAP006564
AGAP006566
AGAP006568
AGAP006569
AGAP006570
AGAP006571
AGAP006572
AGAP006573
AGAP006574
AGAP006575
AGAP006576
AGAP006577
AGAP006578
AGAP006579
AGAP006580
AGAP006581
AGAP006583
AGAP006584
AGAP006585
AGAP006586
AGAP006587
AGAP006588
AGAP006590
AGAP006591
AGAP006592
AGAP006593
AGAP006594
AGAP006595
AGAP006596
mean_tpm
time
preZGA
ZGA_early
ZGA_late
5h
9h
11h
14h
20h
24h
L1
sex
Female
Male

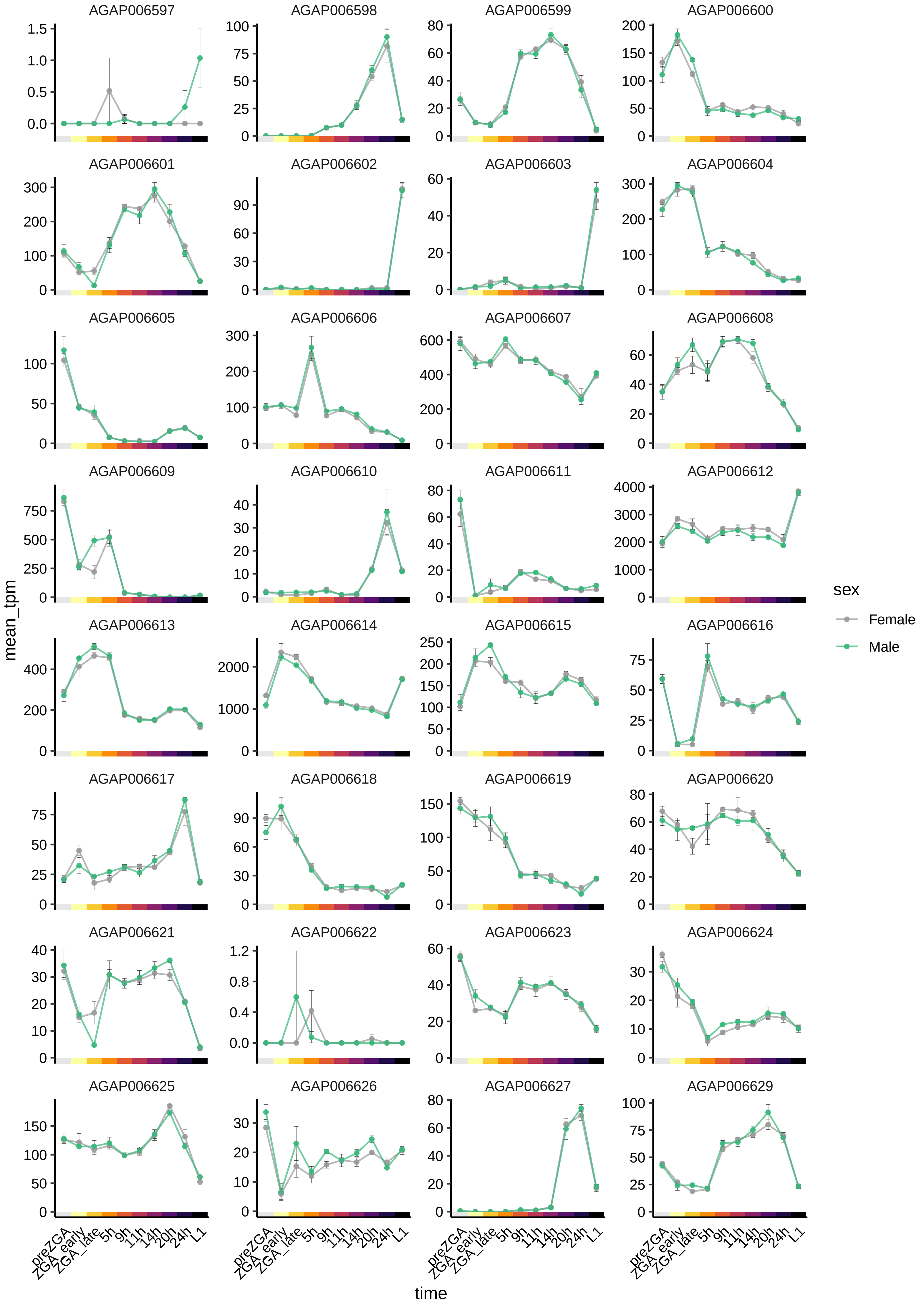
AGAP006597
AGAP006598
AGAP006599
AGAP006600
AGAP006601
AGAP006602
AGAP006603
AGAP006604
AGAP006605
AGAP006606
AGAP006607
AGAP006608
AGAP006609
AGAP006610
AGAP006611
AGAP006612
AGAP006613
AGAP006614
AGAP006615
AGAP006616
AGAP006617
AGAP006618
AGAP006619
AGAP006620
AGAP006621
AGAP006622
AGAP006623
AGAP006624
AGAP006625
AGAP006626
AGAP006627
AGAP006629
mean_tpm
time
preZGA
ZGA_early
ZGA_late
5h
9h
11h
14h
20h
24h
L1
sex
Female
Male

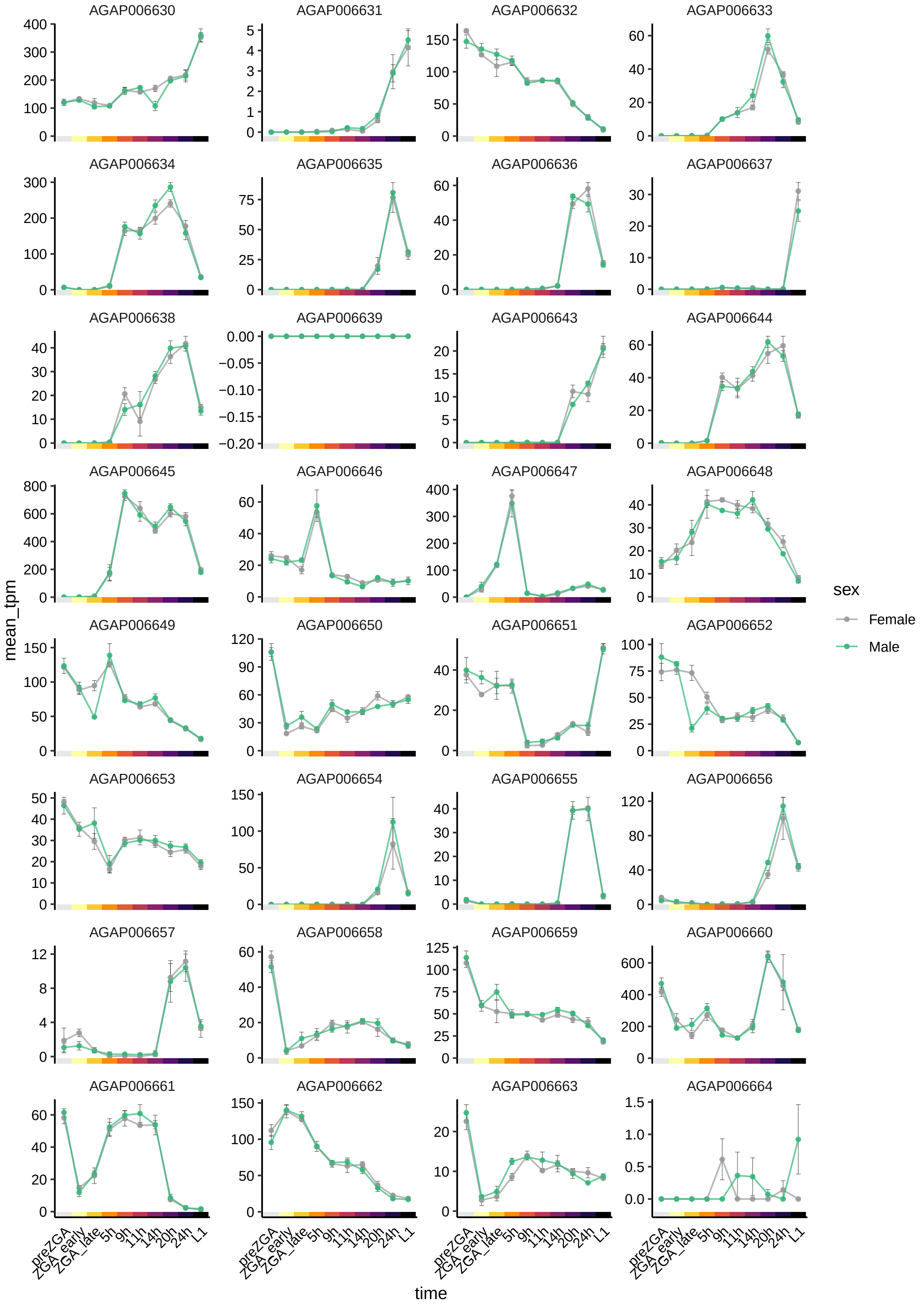
AGAP006630
AGAP006631
AGAP006632
AGAP006633
AGAP006634
AGAP006635
AGAP006636
AGAP006637
AGAP006638
AGAP006639
AGAP006643
AGAP006644
AGAP006645
AGAP006646
AGAP006647
AGAP006648
AGAP006649
AGAP006650
AGAP006651
AGAP006652
AGAP006653
AGAP006654
AGAP006655
AGAP006656
AGAP006657
AGAP006658
AGAP006659
AGAP006660
AGAP006661
AGAP006662
AGAP006663
AGAP006664
mean_tpm
time
preZGA
ZGA_early
ZGA_late
5h
9h
11h
14h
20h
24h
L1
sex
Female
Male

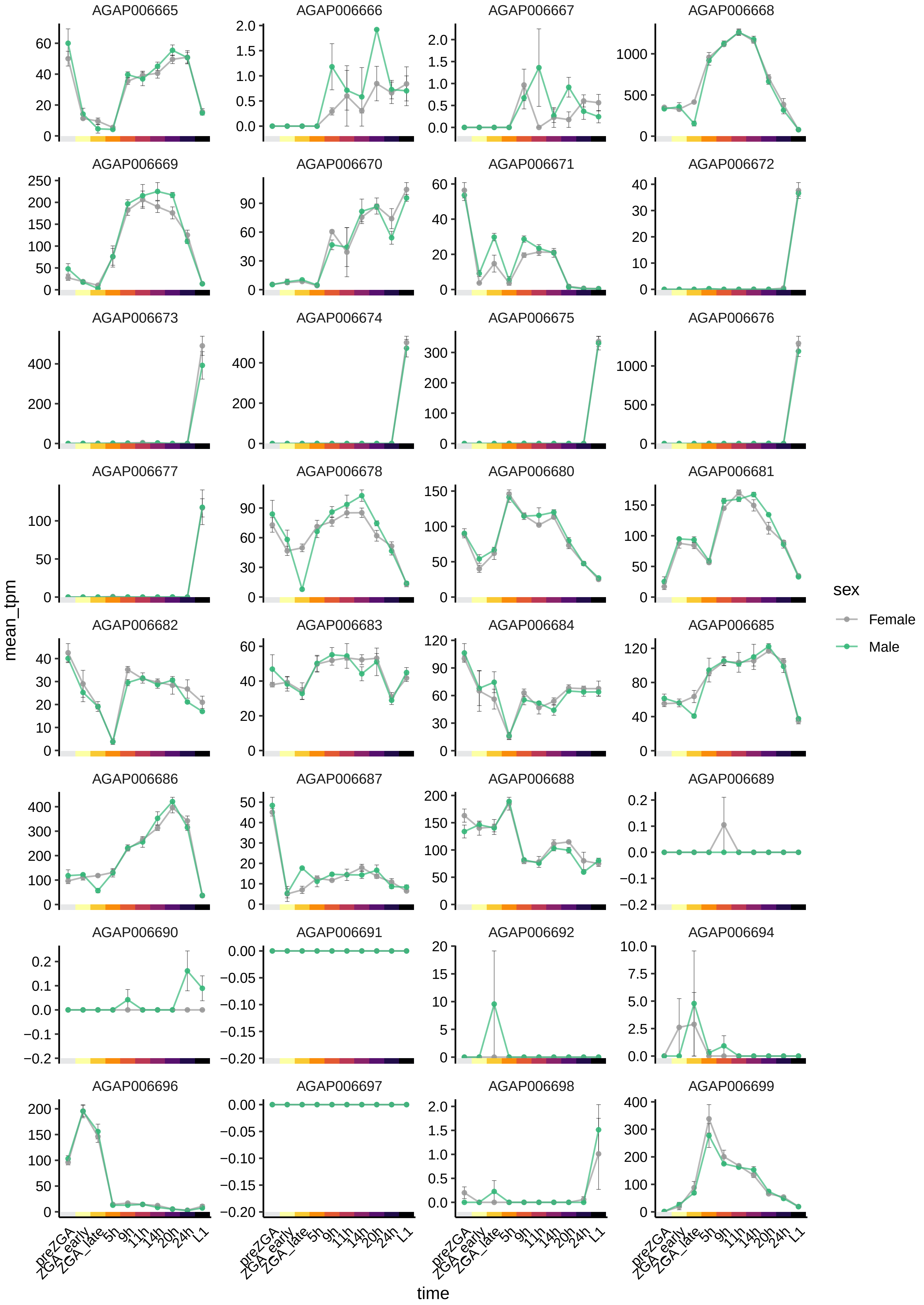

AGAP006665
AGAP006666
AGAP006667
AGAP006668
AGAP006669
AGAP006670
AGAP006671
AGAP006672
AGAP006673
AGAP006674
AGAP006675
AGAP006676
AGAP006677
AGAP006678
AGAP006680
AGAP006681
AGAP006682
AGAP006683
AGAP006684
AGAP006685
AGAP006686
AGAP006687
AGAP006688
AGAP006689
AGAP006690
AGAP006691
AGAP006692
AGAP006694
AGAP006696
AGAP006697
AGAP006698
AGAP006699
mean_tpm
time
preZGA
ZGA_early
ZGA_late
5h
9h
11h
14h
20h
24h
L1
sex
Female
Male

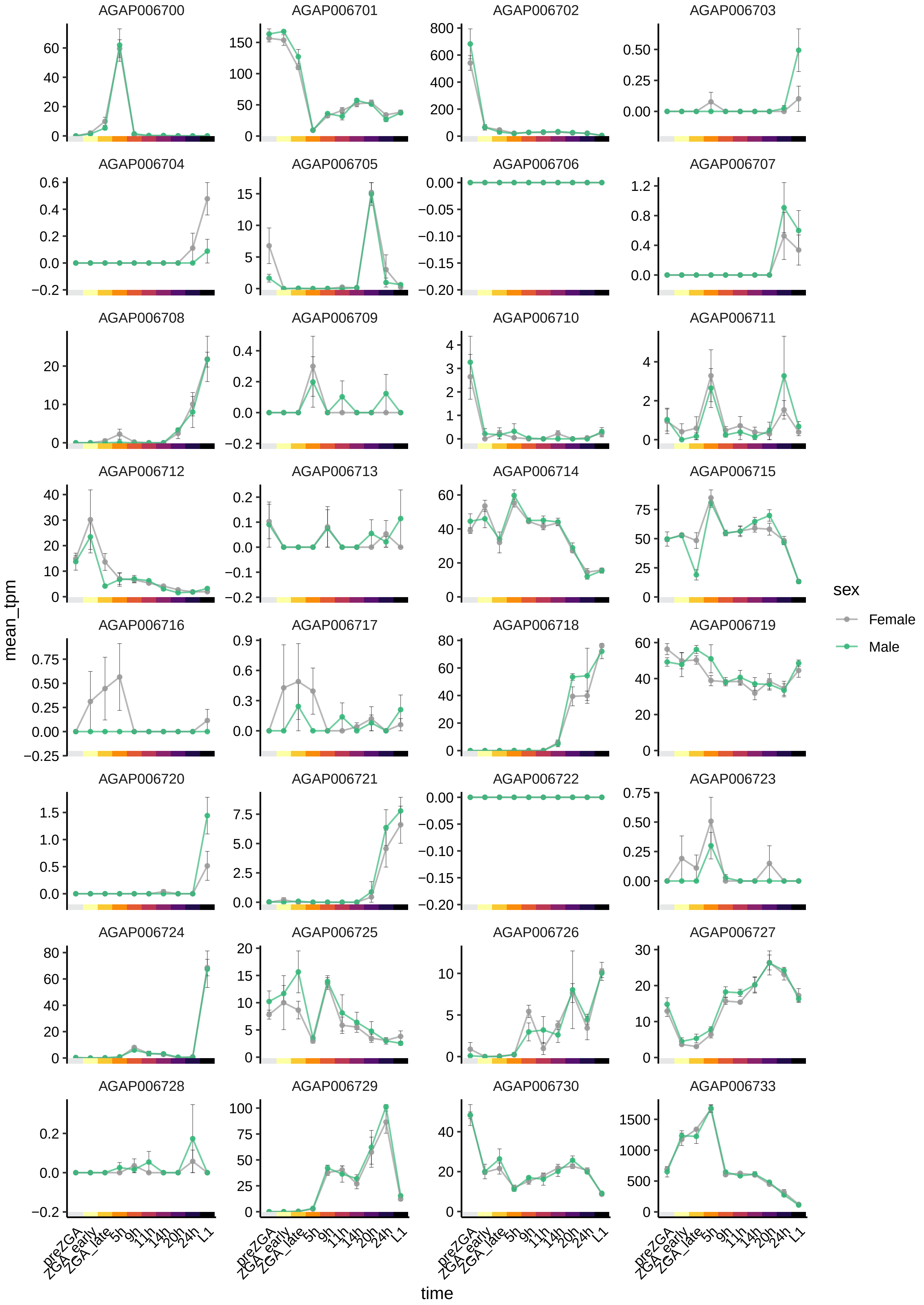

AGAP006700
AGAP006701
AGAP006702
AGAP006703
AGAP006704
AGAP006705
AGAP006706
AGAP006707
AGAP006708
AGAP006709
AGAP006710
AGAP006711
AGAP006712
AGAP006713
AGAP006714
AGAP006715
AGAP006716
AGAP006717
AGAP006718
AGAP006719
AGAP006720
AGAP006721
AGAP006722
AGAP006723
AGAP006724
AGAP006725
AGAP006726
AGAP006727
AGAP006728
AGAP006729
AGAP006730
AGAP006733
mean_tpm
time
preZGA
ZGA_early
ZGA_late
5h
9h
11h
14h
20h
24h
L1
sex
Female
Male

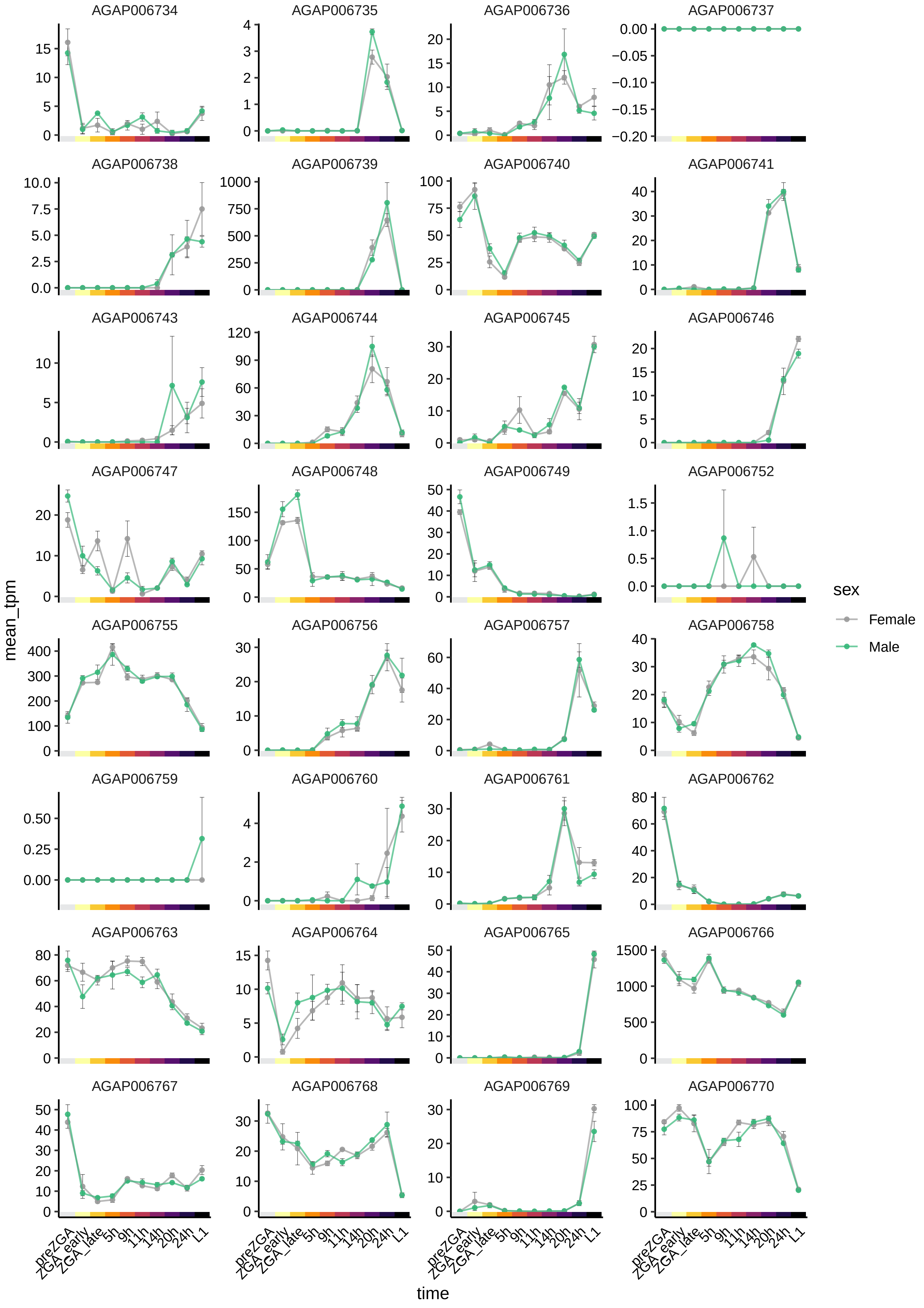
AGAP006734
AGAP006735
AGAP006736
AGAP006737
AGAP006738
AGAP006739
AGAP006740
AGAP006741
AGAP006743
AGAP006744
AGAP006745
AGAP006746
AGAP006747
AGAP006748
AGAP006749
AGAP006752
AGAP006755
AGAP006756
AGAP006757
AGAP006758
AGAP006759
AGAP006760
AGAP006761
AGAP006762
AGAP006763
AGAP006764
AGAP006765
AGAP006766
AGAP006767
AGAP006768
AGAP006769
AGAP006770
mean_tpm
time
preZGA
ZGA_early
ZGA_late
5h
9h
11h
14h
20h
24h
L1
sex
Female
Male

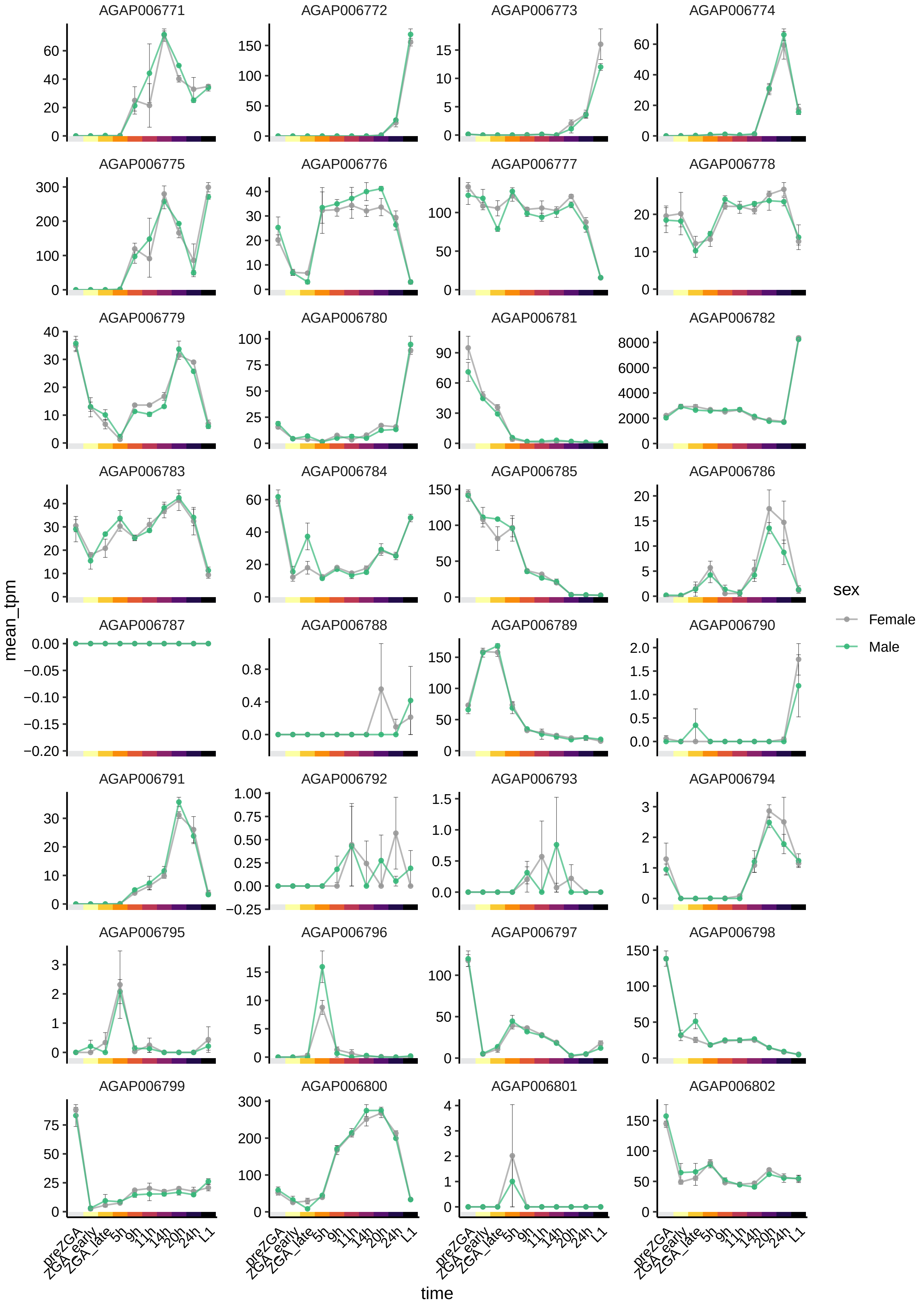

AGAP006771
AGAP006772
AGAP006773
AGAP006774
AGAP006775
AGAP006776
AGAP006777
AGAP006778
AGAP006779
AGAP006780
AGAP006781
AGAP006782
AGAP006783
AGAP006784
AGAP006785
AGAP006786
AGAP006787
AGAP006788
AGAP006789
AGAP006790
AGAP006791
AGAP006792
AGAP006793
AGAP006794
AGAP006795
AGAP006796
AGAP006797
AGAP006798
AGAP006799
AGAP006800
AGAP006801
AGAP006802
mean_tpm
time
preZGA
ZGA_early
ZGA_late
5h
9h
11h
14h
20h
24h
L1
sex
Female
Male

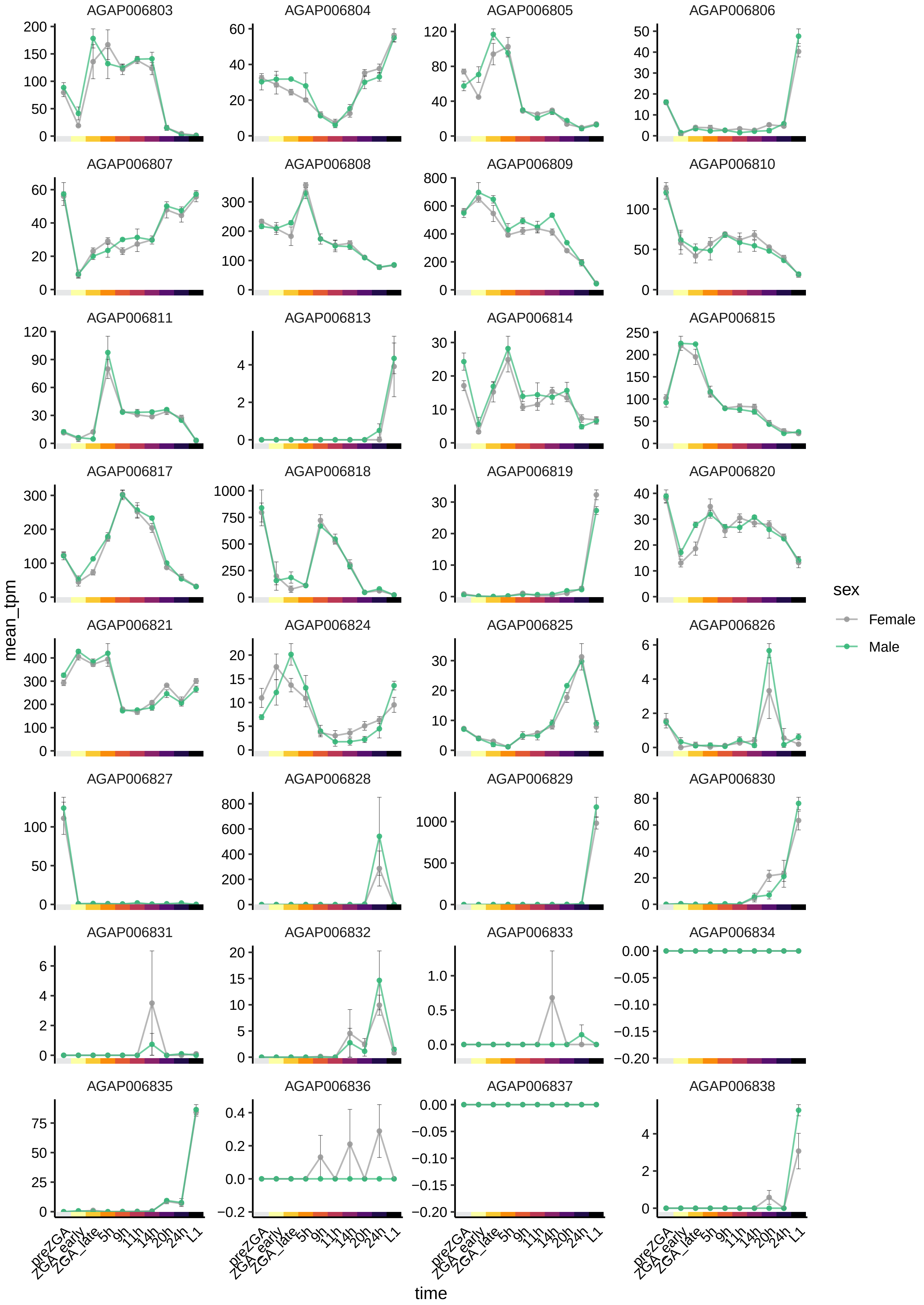
AGAP006803
AGAP006804
AGAP006805
AGAP006806
AGAP006807
AGAP006808
AGAP006809
AGAP006810
AGAP006811
AGAP006813
AGAP006814
AGAP006815
AGAP006817
AGAP006818
AGAP006819
AGAP006820
AGAP006821
AGAP006824
AGAP006825
AGAP006826
AGAP006827
AGAP006828
AGAP006829
AGAP006830
AGAP006831
AGAP006832
AGAP006833
AGAP006834
AGAP006835
AGAP006836
AGAP006837
AGAP006838
mean_tpm
time
preZGA
ZGA_early
ZGA_late
5h
9h
11h
14h
20h
24h
L1
sex
Female
Male

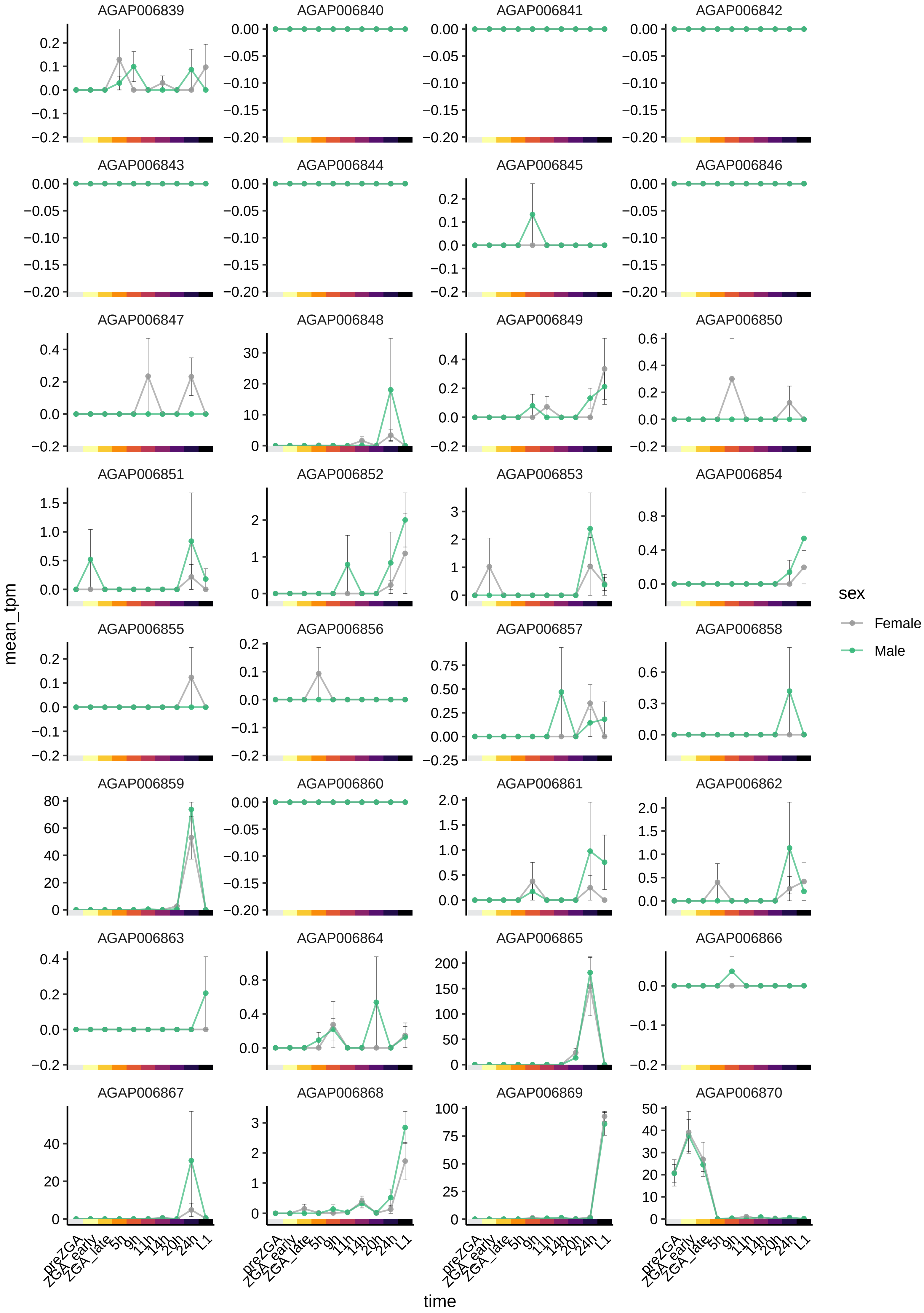
AGAP006839
AGAP006840
AGAP006841
AGAP006842
AGAP006843
AGAP006844
AGAP006845
AGAP006846
AGAP006847
AGAP006848
AGAP006849
AGAP006850
AGAP006851
AGAP006852
AGAP006853
AGAP006854
AGAP006855
AGAP006856
AGAP006857
AGAP006858
AGAP006859
AGAP006860
AGAP006861
AGAP006862
AGAP006863
AGAP006864
AGAP006865
AGAP006866
AGAP006867
AGAP006868
AGAP006869
AGAP006870
mean_tpm
time
preZGA
ZGA_early
ZGA_late
5h
9h
11h
14h
20h
24h
L1
sex
Female
Male

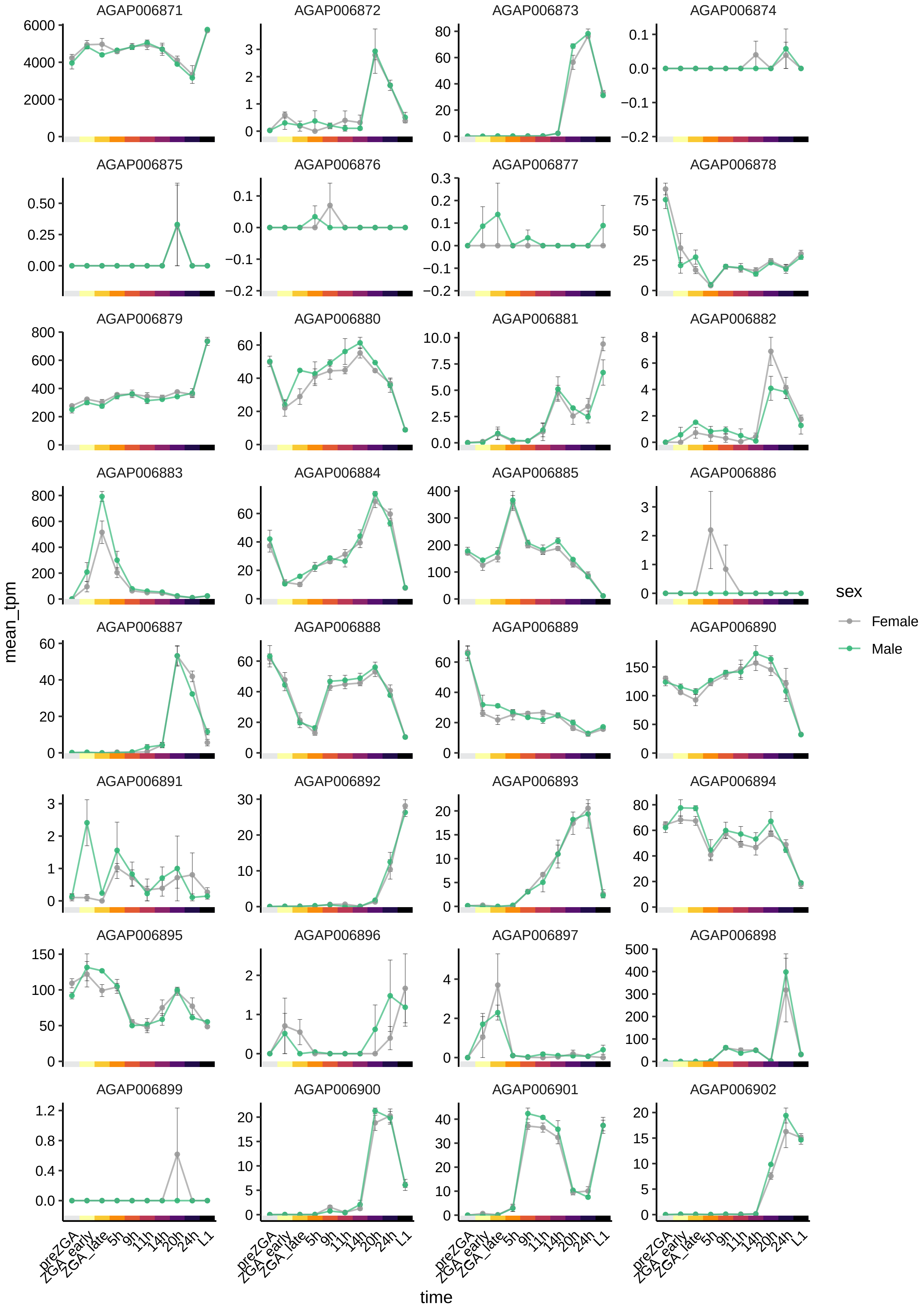

AGAP006871
AGAP006872
AGAP006873
AGAP006874
AGAP006875
AGAP006876
AGAP006877
AGAP006878
AGAP006879
AGAP006880
AGAP006881
AGAP006882
AGAP006883
AGAP006884
AGAP006885
AGAP006886
AGAP006887
AGAP006888
AGAP006889
AGAP006890
AGAP006891
AGAP006892
AGAP006893
AGAP006894
AGAP006895
AGAP006896
AGAP006897
AGAP006898
AGAP006899
AGAP006900
AGAP006901
AGAP006902
mean_tpm
time
preZGA
ZGA_early
ZGA_late
5h
9h
11h
14h
20h
24h
L1
sex
Female
Male

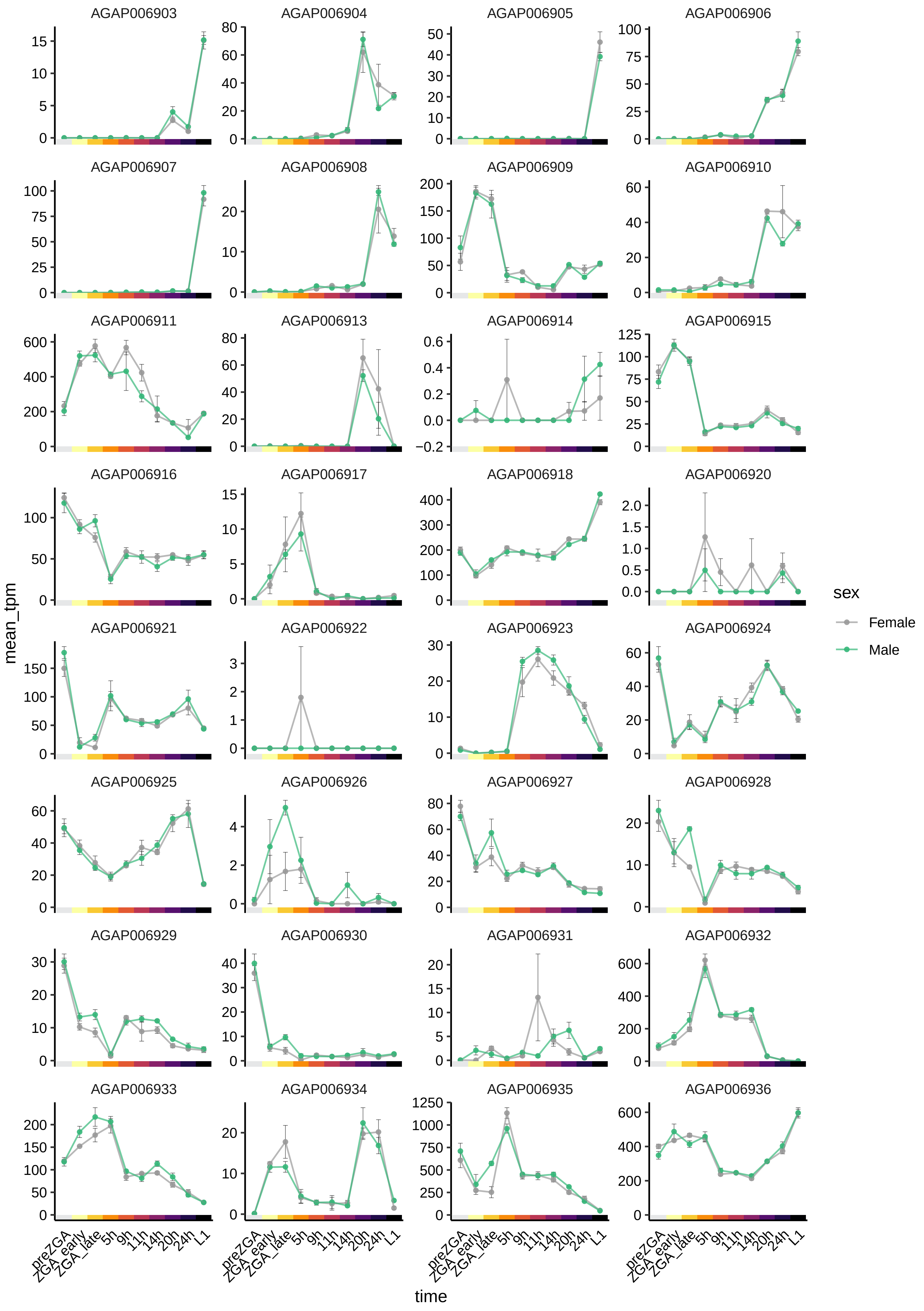

AGAP006903
AGAP006904
AGAP006905
AGAP006906
AGAP006907
AGAP006908
AGAP006909
AGAP006910
AGAP006911
AGAP006913
AGAP006914
AGAP006915
AGAP006916
AGAP006917
AGAP006918
AGAP006920
AGAP006921
AGAP006922
AGAP006923
AGAP006924
AGAP006925
AGAP006926
AGAP006927
AGAP006928
AGAP006929
AGAP006930
AGAP006931
AGAP006932
AGAP006933
AGAP006934
AGAP006935
AGAP006936
mean_tpm
time
preZGA
ZGA_early
ZGA_late
5h
9h
11h
14h
20h
24h
L1
sex
Female
Male

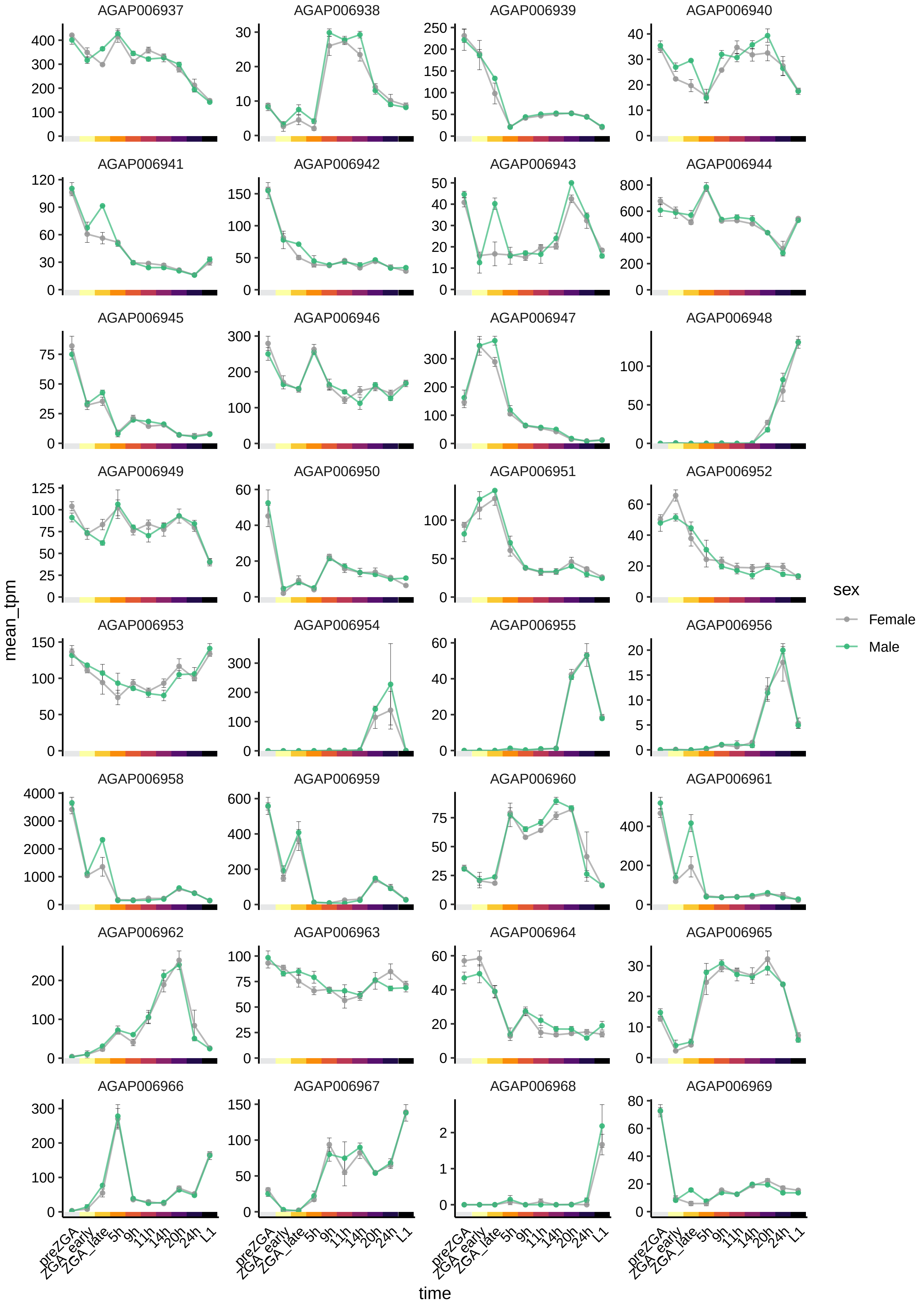
AGAP006937
AGAP006938
AGAP006939
AGAP006940
AGAP006941
AGAP006942
AGAP006943
AGAP006944
AGAP006945
AGAP006946
AGAP006947
AGAP006948
AGAP006949
AGAP006950
AGAP006951
AGAP006952
AGAP006953
AGAP006954
AGAP006955
AGAP006956
AGAP006958
AGAP006959
AGAP006960
AGAP006961
AGAP006962
AGAP006963
AGAP006964
AGAP006965
AGAP006966
AGAP006967
AGAP006968
AGAP006969
mean_tpm
time
preZGA
ZGA_early
ZGA_late
5h
9h
11h
14h
20h
24h
L1
sex
Female
Male

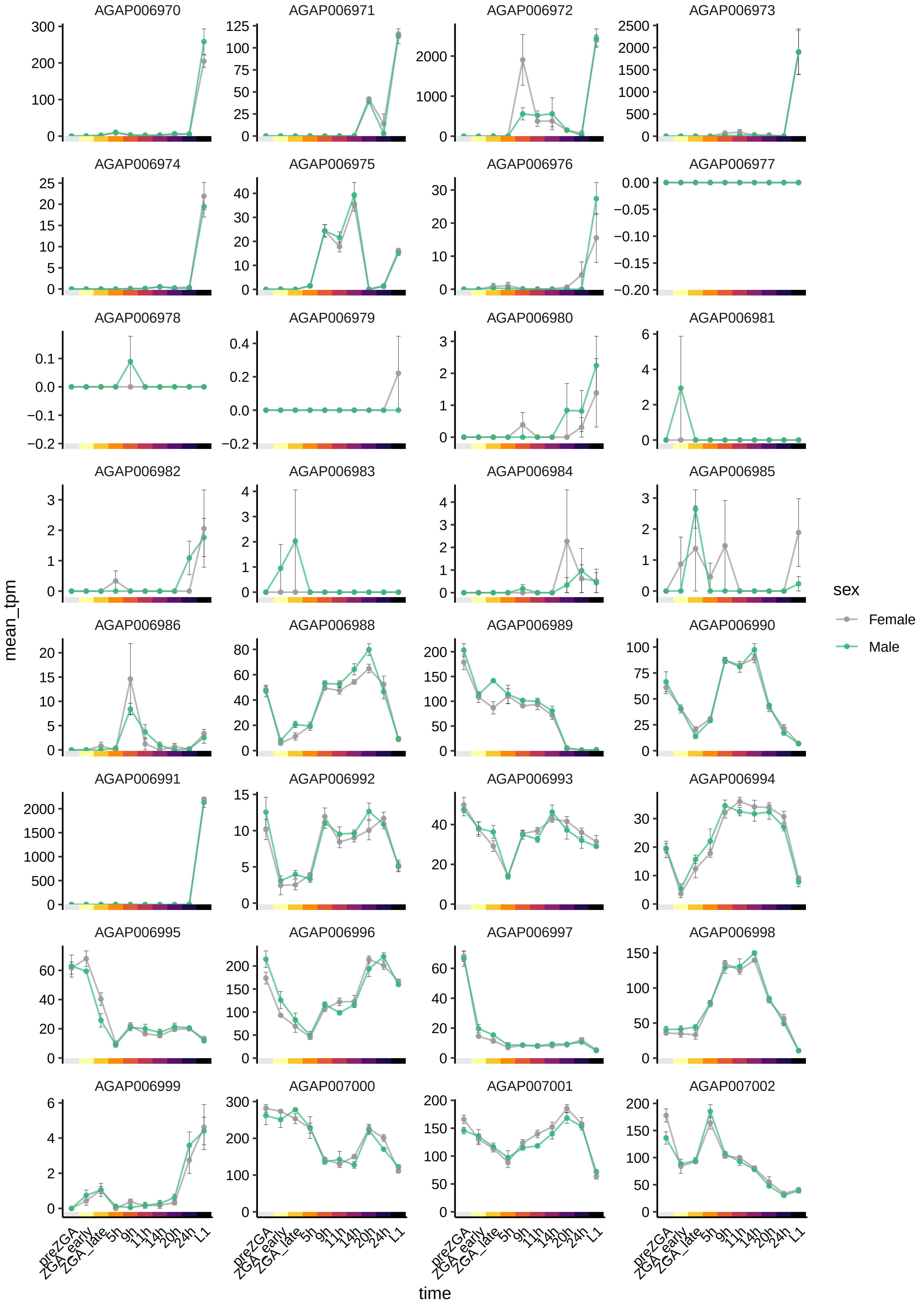

AGAP006970
AGAP006971
AGAP006972
AGAP006973
AGAP006974
AGAP006975
AGAP006976
AGAP006977
AGAP006978
AGAP006979
AGAP006980
AGAP006981
AGAP006982
AGAP006983
AGAP006984
AGAP006985
AGAP006986
AGAP006988
AGAP006989
AGAP006990
AGAP006991
AGAP006992
AGAP006993
AGAP006994
AGAP006995
AGAP006996
AGAP006997
AGAP006998
AGAP006999
AGAP007000
AGAP007001
AGAP007002
mean_tpm
time
preZGA
ZGA_early
ZGA_late
5h
9h
11h
14h
20h
24h
L1
sex
Female
Male

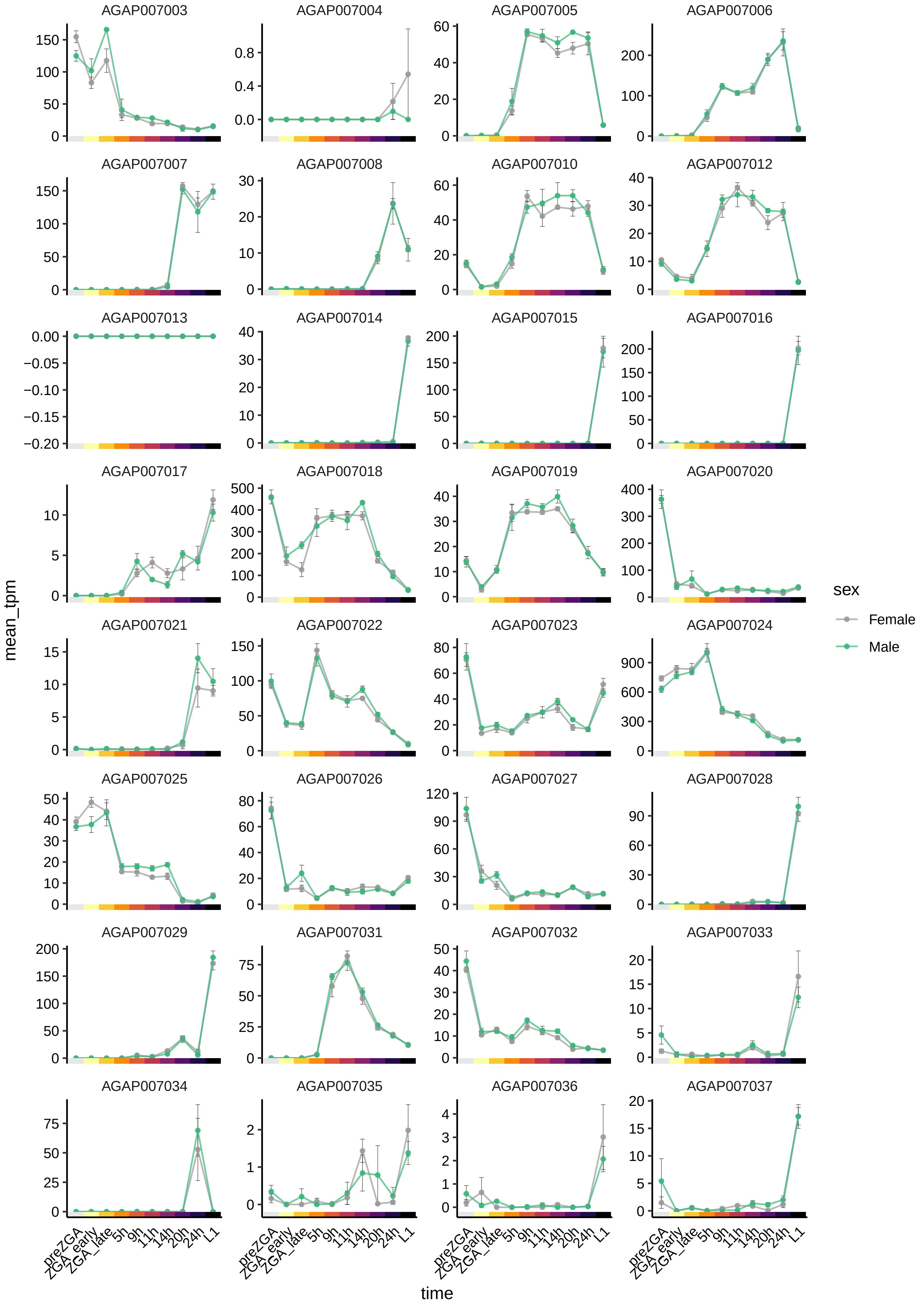
AGAP007003
AGAP007004
AGAP007005
AGAP007006
AGAP007007
AGAP007008
AGAP007010
AGAP007012
AGAP007013
AGAP007014
AGAP007015
AGAP007016
AGAP007017
AGAP007018
AGAP007019
AGAP007020
AGAP007021
AGAP007022
AGAP007023
AGAP007024
AGAP007025
AGAP007026
AGAP007027
AGAP007028
AGAP007029
AGAP007031
AGAP007032
AGAP007033
AGAP007034
AGAP007035
AGAP007036
AGAP007037
mean_tpm
time
preZGA
ZGA_early
ZGA_late
5h
9h
11h
14h
20h
24h
L1
sex
Female
Male

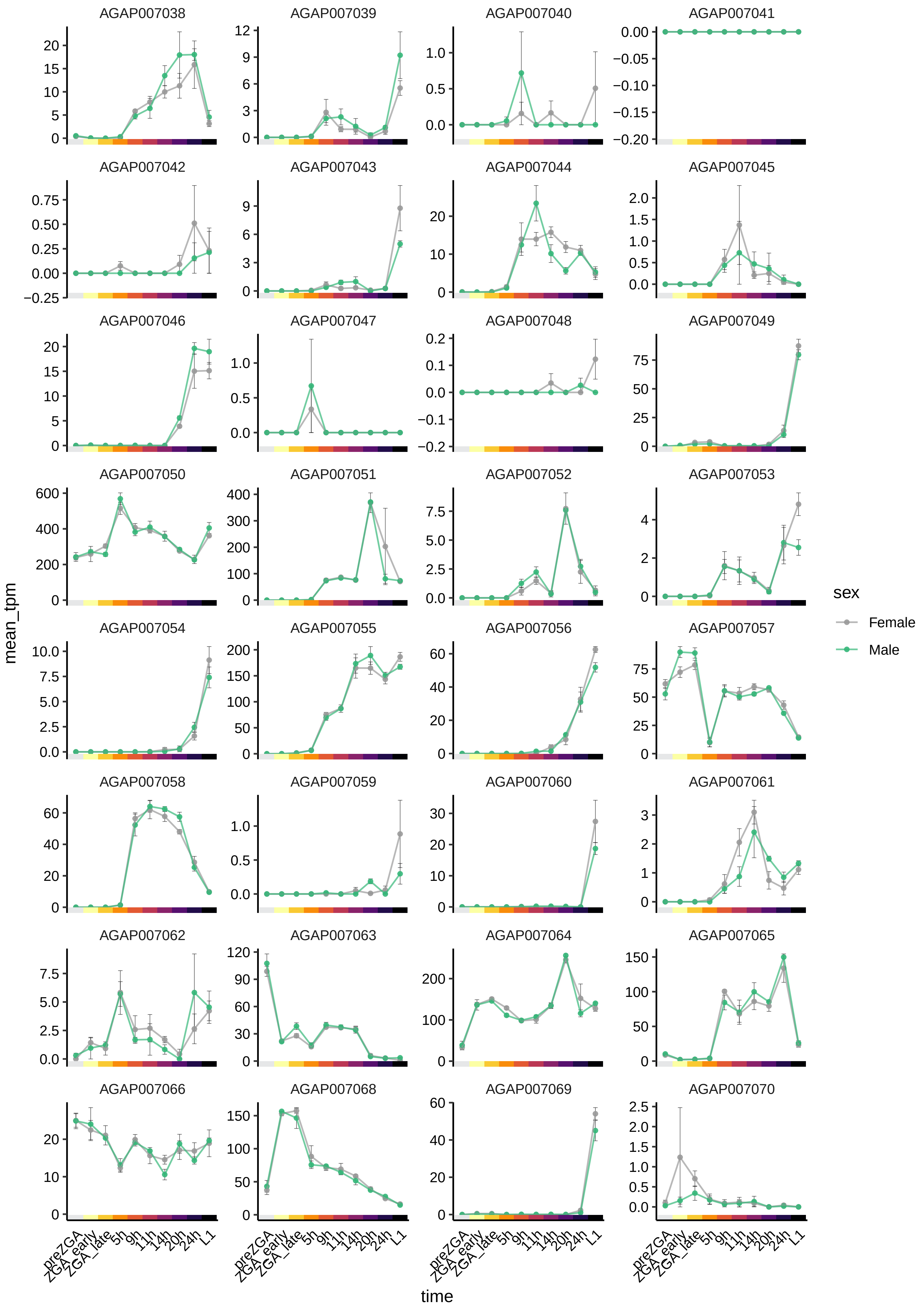

AGAP007038
AGAP007039
AGAP007040
AGAP007041
AGAP007042
AGAP007043
AGAP007044
AGAP007045
AGAP007046
AGAP007047
AGAP007048
AGAP007049
AGAP007050
AGAP007051
AGAP007052
AGAP007053
AGAP007054
AGAP007055
AGAP007056
AGAP007057
AGAP007058
AGAP007059
AGAP007060
AGAP007061
AGAP007062
AGAP007063
AGAP007064
AGAP007065
AGAP007066
AGAP007068
AGAP007069
AGAP007070
mean_tpm
time
preZGA
ZGA_early
ZGA_late
5h
9h
11h
14h
20h
24h
L1
sex
Female
Male

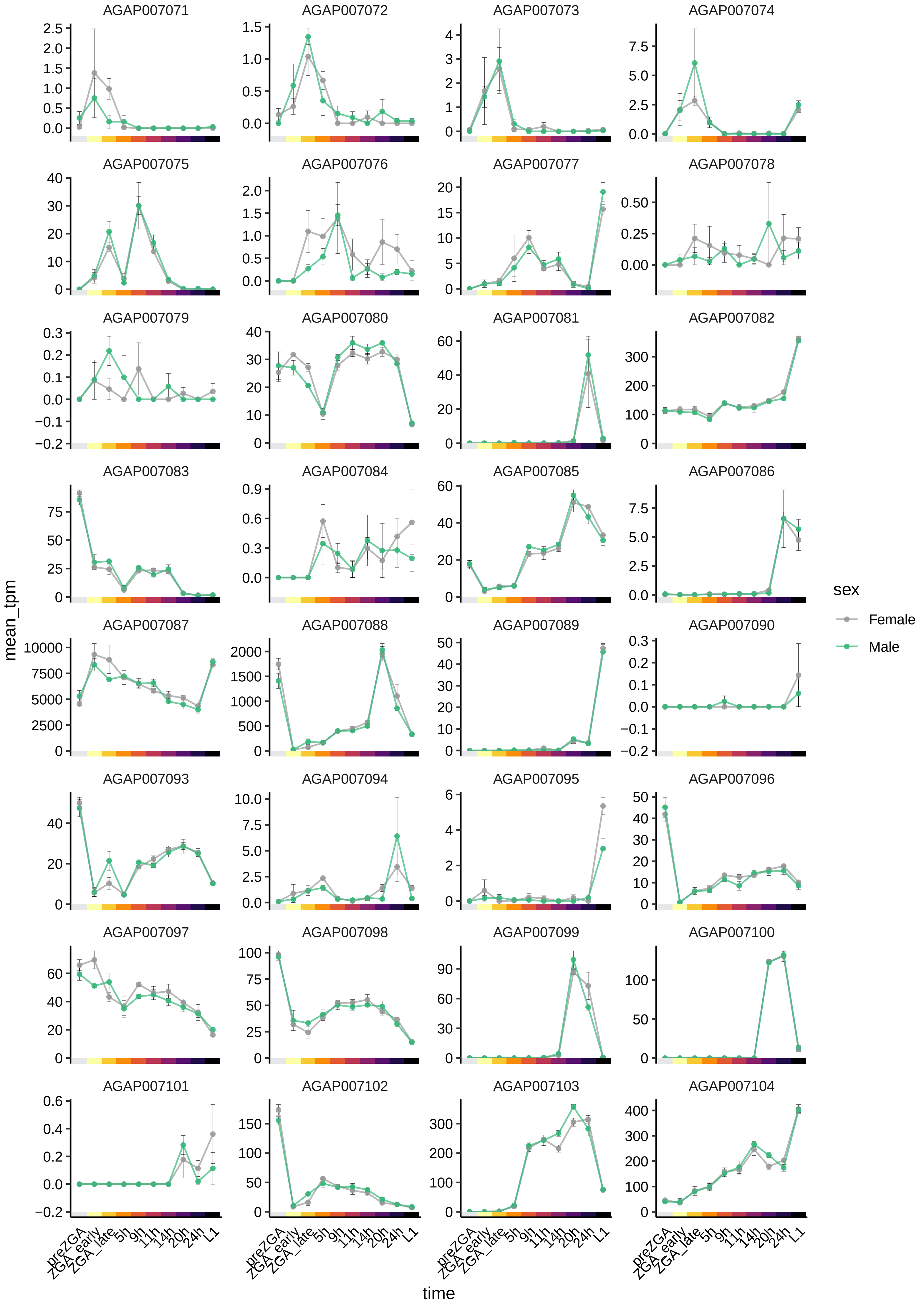
AGAP007071
AGAP007072
AGAP007073
AGAP007074
AGAP007075
AGAP007076
AGAP007077
AGAP007078
AGAP007079
AGAP007080
AGAP007081
AGAP007082
AGAP007083
AGAP007084
AGAP007085
AGAP007086
AGAP007087
AGAP007088
AGAP007089
AGAP007090
AGAP007093
AGAP007094
AGAP007095
AGAP007096
AGAP007097
AGAP007098
AGAP007099
AGAP007100
AGAP007101
AGAP007102
AGAP007103
AGAP007104
mean_tpm
time
preZGA
ZGA_early
ZGA_late
5h
9h
11h
14h
20h
24h
L1
sex
Female
Male

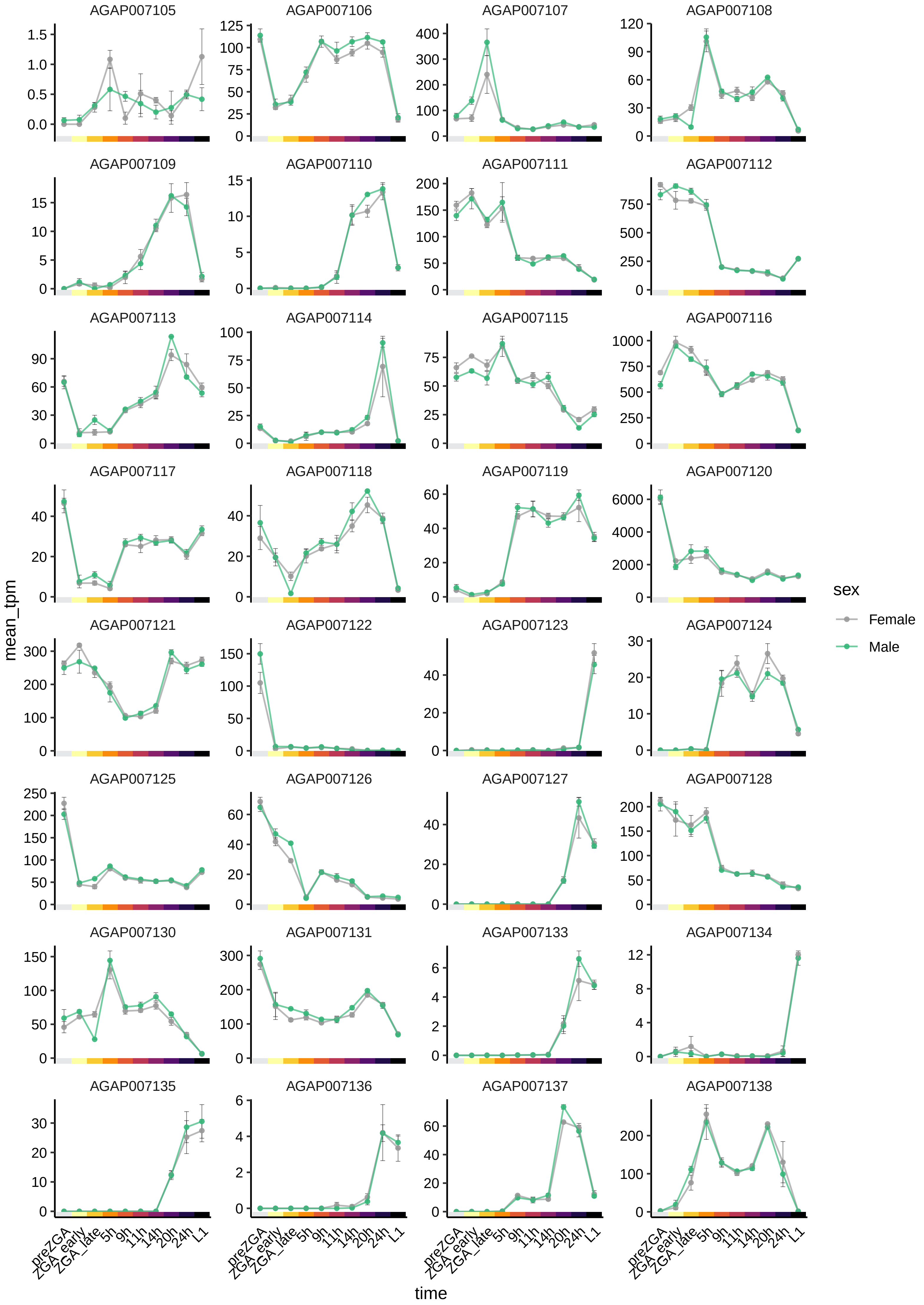

AGAP007105
AGAP007106
AGAP007107
AGAP007108
AGAP007109
AGAP007110
AGAP007111
AGAP007112
AGAP007113
AGAP007114
AGAP007115
AGAP007116
AGAP007117
AGAP007118
AGAP007119
AGAP007120
AGAP007121
AGAP007122
AGAP007123
AGAP007124
AGAP007125
AGAP007126
AGAP007127
AGAP007128
AGAP007130
AGAP007131
AGAP007133
AGAP007134
AGAP007135
AGAP007136
AGAP007137
AGAP007138
mean_tpm
time
preZGA
ZGA_early
ZGA_late
5h
9h
11h
14h
20h
24h
L1
sex
Female
Male

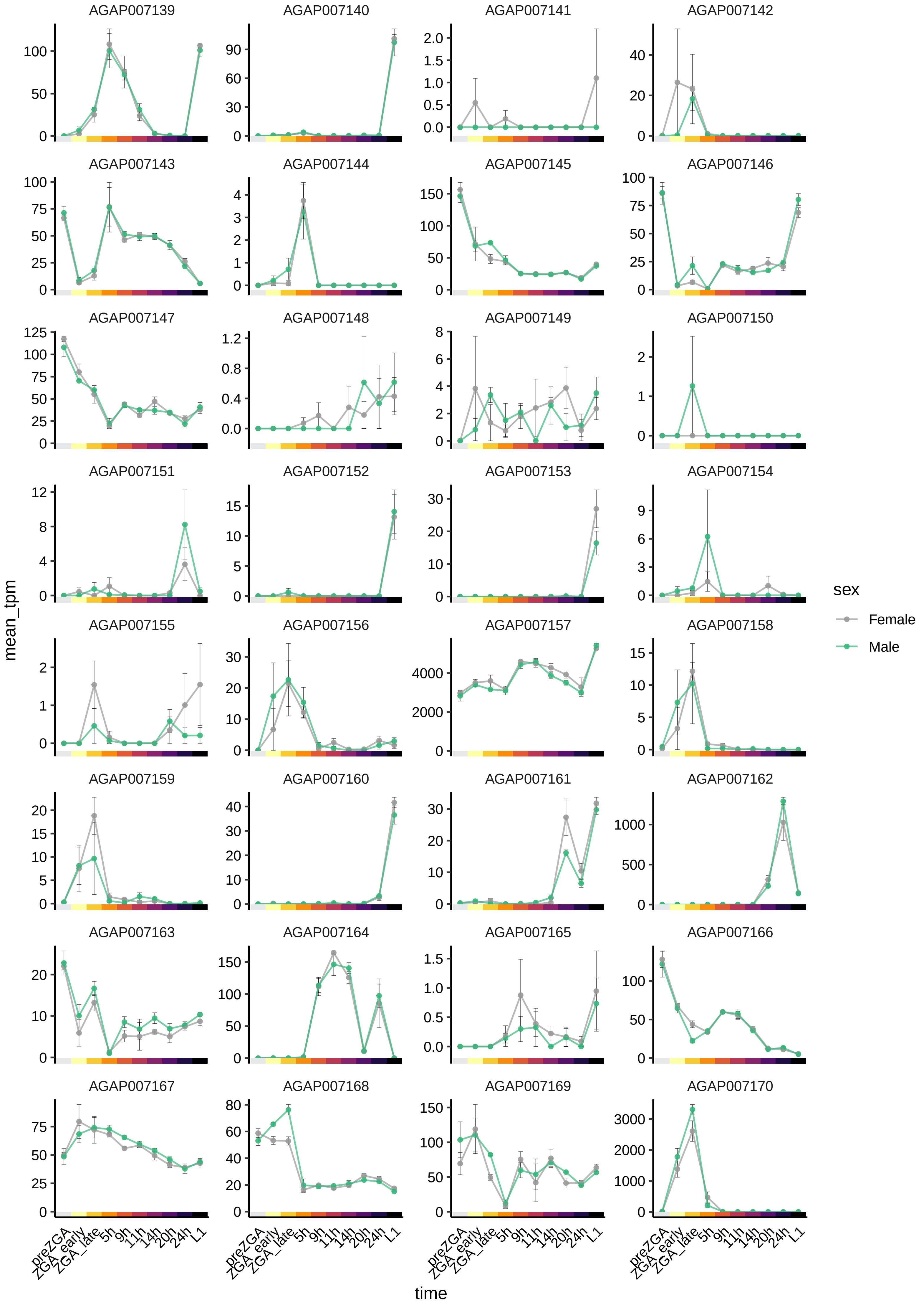
AGAP007139
AGAP007140
AGAP007141
AGAP007142
AGAP007143
AGAP007144
AGAP007145
AGAP007146
AGAP007147
AGAP007148
AGAP007149
AGAP007150
AGAP007151
AGAP007152
AGAP007153
AGAP007154
AGAP007155
AGAP007156
AGAP007157
AGAP007158
AGAP007159
AGAP007160
AGAP007161
AGAP007162
AGAP007163
AGAP007164
AGAP007165
AGAP007166
AGAP007167
AGAP007168
AGAP007169
AGAP007170
mean_tpm
preZGA
ZGA_early
ZGA_late
5h
9h
11h
14h
20h
24h
L1
time
sex
Female
Male

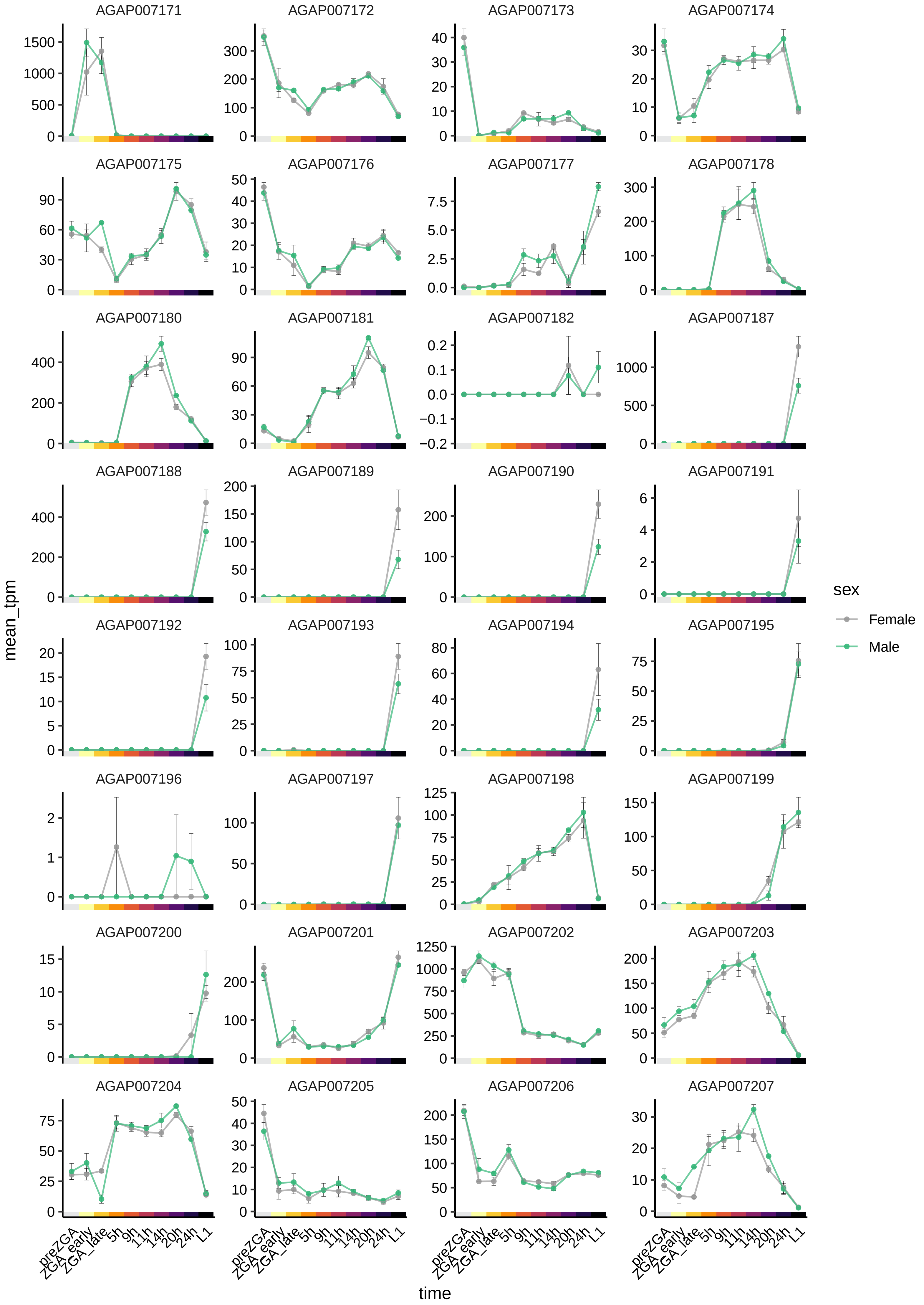

AGAP007171
AGAP007172
AGAP007173
AGAP007174
AGAP007175
AGAP007176
AGAP007177
AGAP007178
AGAP007180
AGAP007181
AGAP007182
AGAP007187
AGAP007188
AGAP007189
AGAP007190
AGAP007191
AGAP007192
AGAP007193
AGAP007194
AGAP007195
AGAP007196
AGAP007197
AGAP007198
AGAP007199
AGAP007200
AGAP007201
AGAP007202
AGAP007203
AGAP007204
AGAP007205
AGAP007206
AGAP007207
mean_tpm
time
preZGA
ZGA_early
ZGA_late
5h
9h
11h
14h
20h
24h
L1
sex
Female
Male

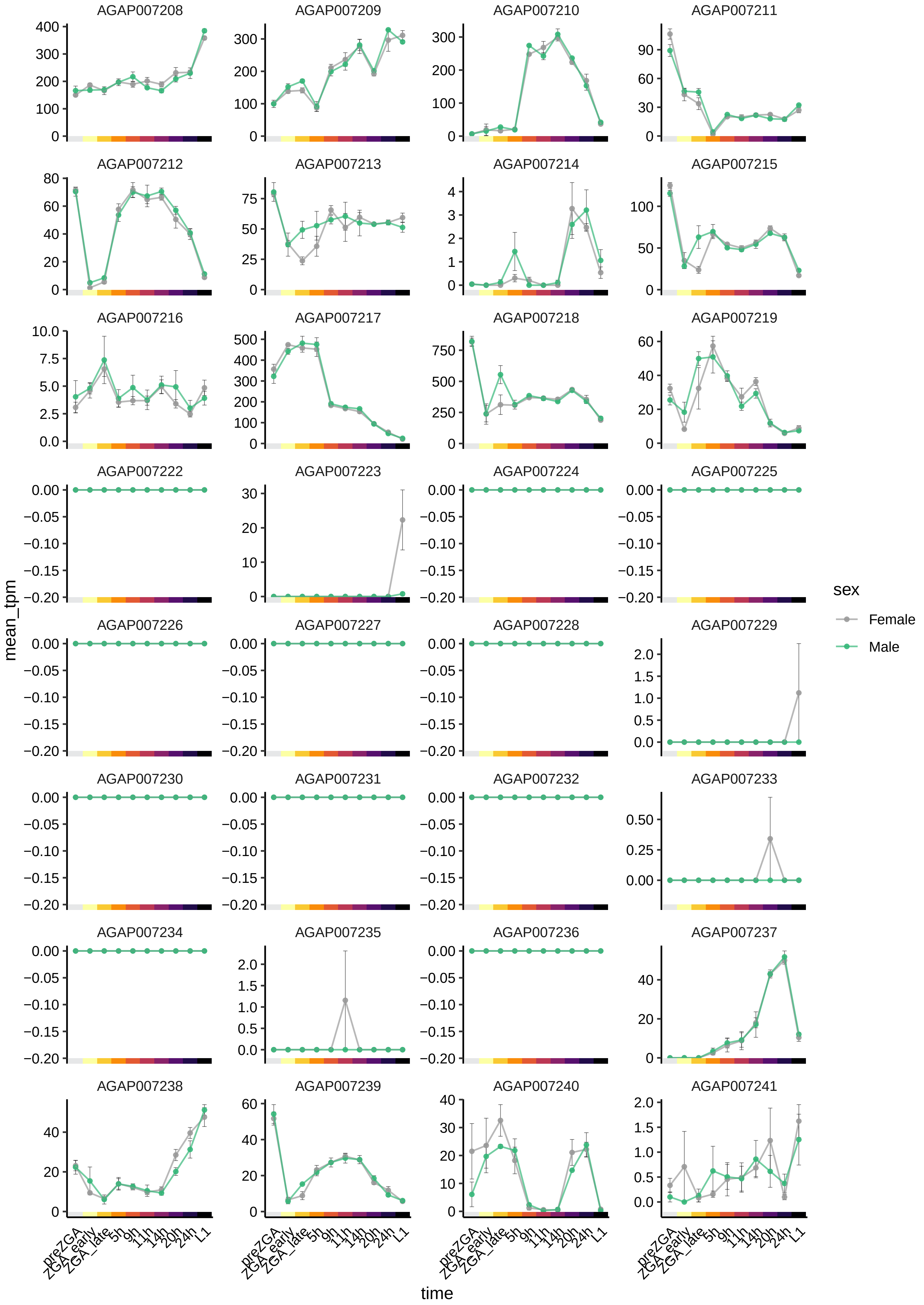

AGAP007208
AGAP007209
AGAP007210
AGAP007211
AGAP007212
AGAP007213
AGAP007214
AGAP007215
AGAP007216
AGAP007217
AGAP007218
AGAP007219
AGAP007222
AGAP007223
AGAP007224
AGAP007225
AGAP007226
AGAP007227
AGAP007228
AGAP007229
AGAP007230
AGAP007231
AGAP007232
AGAP007233
AGAP007234
AGAP007235
AGAP007236
AGAP007237
AGAP007238
AGAP007239
AGAP007240
AGAP007241
mean_tpm
time
preZGA
ZGA_early
ZGA_late
5h
9h
11h
14h
20h
24h
L1
sex
Female
Male

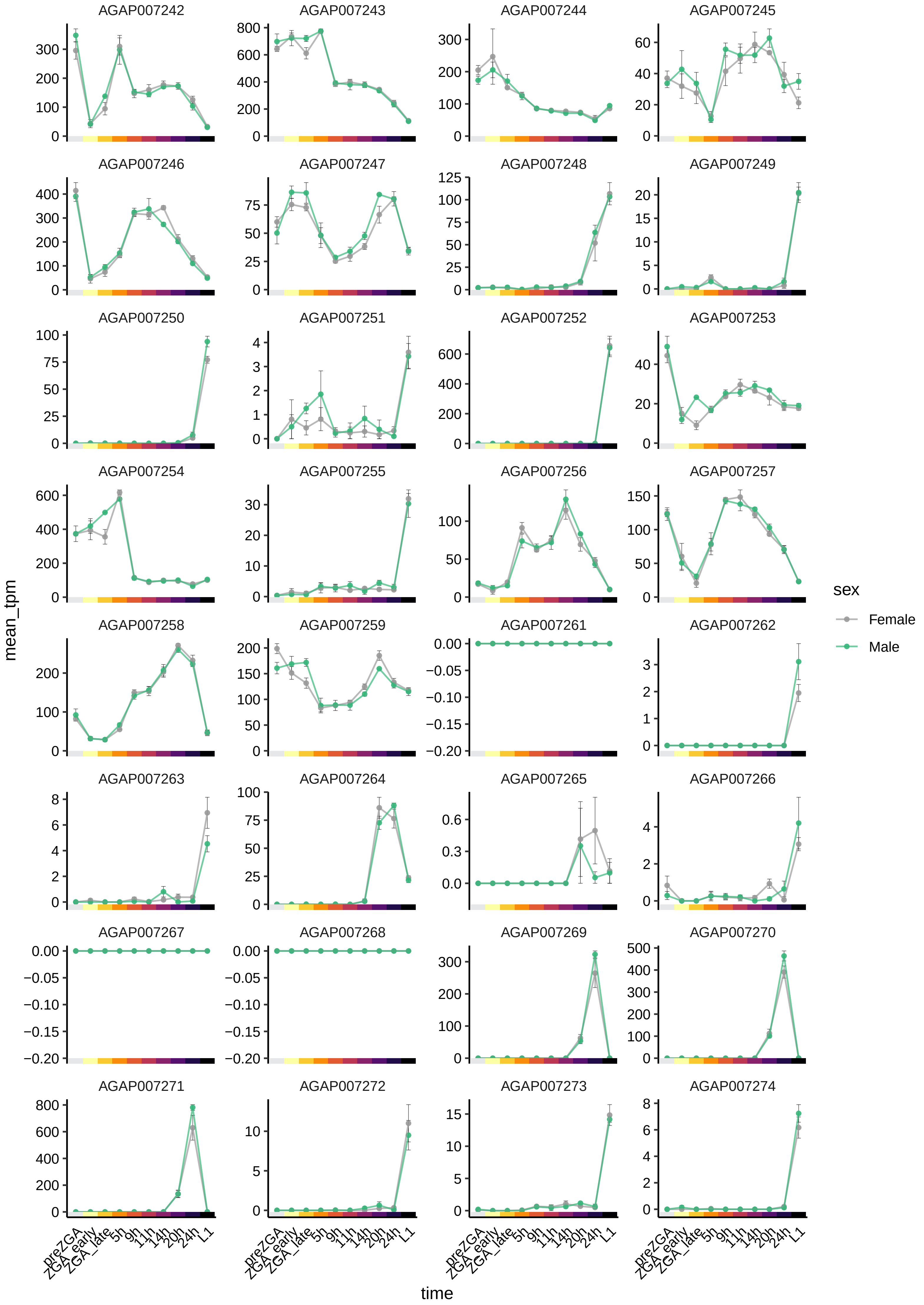

AGAP007242
AGAP007243
AGAP007244
AGAP007245
AGAP007246
AGAP007247
AGAP007248
AGAP007249
AGAP007250
AGAP007251
AGAP007252
AGAP007253
AGAP007254
AGAP007255
AGAP007256
AGAP007257
AGAP007258
AGAP007259
AGAP007261
AGAP007262
AGAP007263
AGAP007264
AGAP007265
AGAP007266
AGAP007267
AGAP007268
AGAP007269
AGAP007270
AGAP007271
AGAP007272
AGAP007273
AGAP007274
mean_tpm
time
preZGA
ZGA_early
ZGA_late
5h
9h
11h
14h
20h
24h
L1
sex
Female
Male

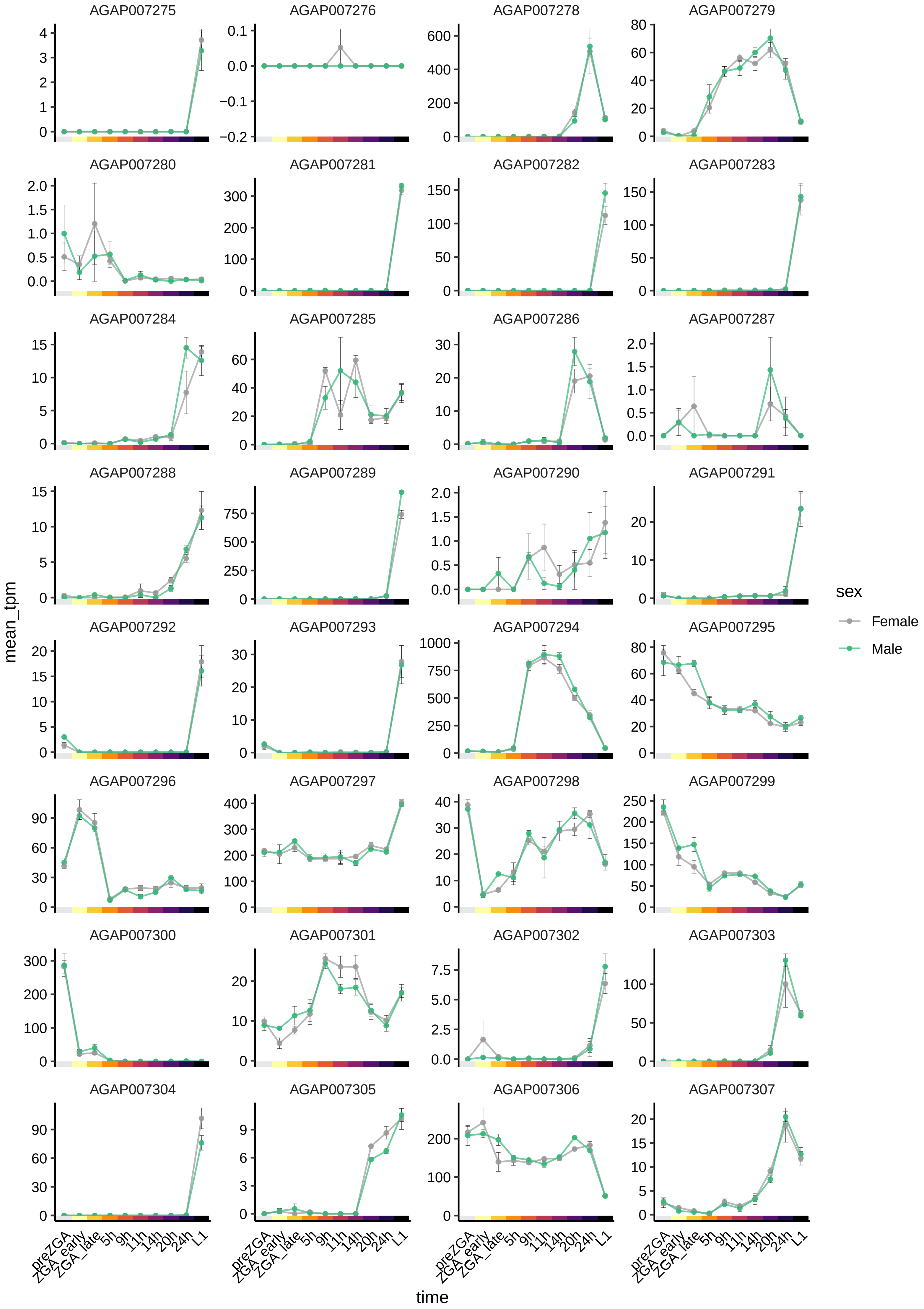

AGAP007275
AGAP007276
AGAP007278
AGAP007279
AGAP007280
AGAP007281
AGAP007282
AGAP007283
AGAP007284
AGAP007285
AGAP007286
AGAP007287
AGAP007288
AGAP007289
AGAP007290
AGAP007291
AGAP007292
AGAP007293
AGAP007294
AGAP007295
AGAP007296
AGAP007297
AGAP007298
AGAP007299
AGAP007300
AGAP007301
AGAP007302
AGAP007303
AGAP007304
AGAP007305
AGAP007306
AGAP007307
mean_tpm
time
preZGA
ZGA_early
ZGA_late
5h
9h
11h
14h
20h
24h
L1
sex
Female
Male

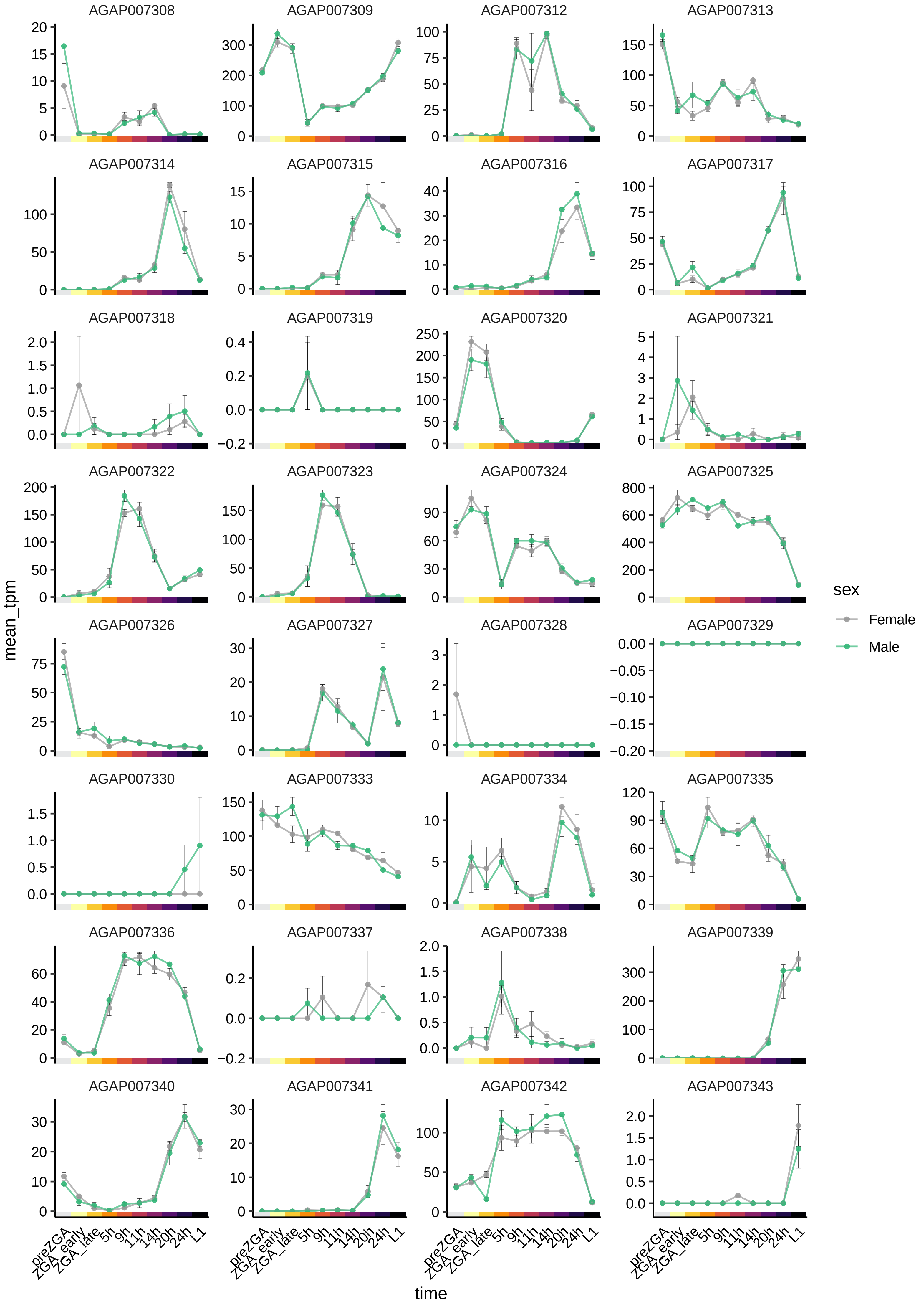
AGAP007308
AGAP007309
AGAP007312
AGAP007313
AGAP007314
AGAP007315
AGAP007316
AGAP007317
AGAP007318
AGAP007319
AGAP007320
AGAP007321
AGAP007322
AGAP007323
AGAP007324
AGAP007325
AGAP007326
AGAP007327
AGAP007328
AGAP007329
AGAP007330
AGAP007333
AGAP007334
AGAP007335
AGAP007336
AGAP007337
AGAP007338
AGAP007339
AGAP007340
AGAP007341
AGAP007342
AGAP007343
mean_tpm
time
preZGA
ZGA_early
ZGA_late
5h
9h
11h
14h
20h
24h
L1
sex
Female
Male

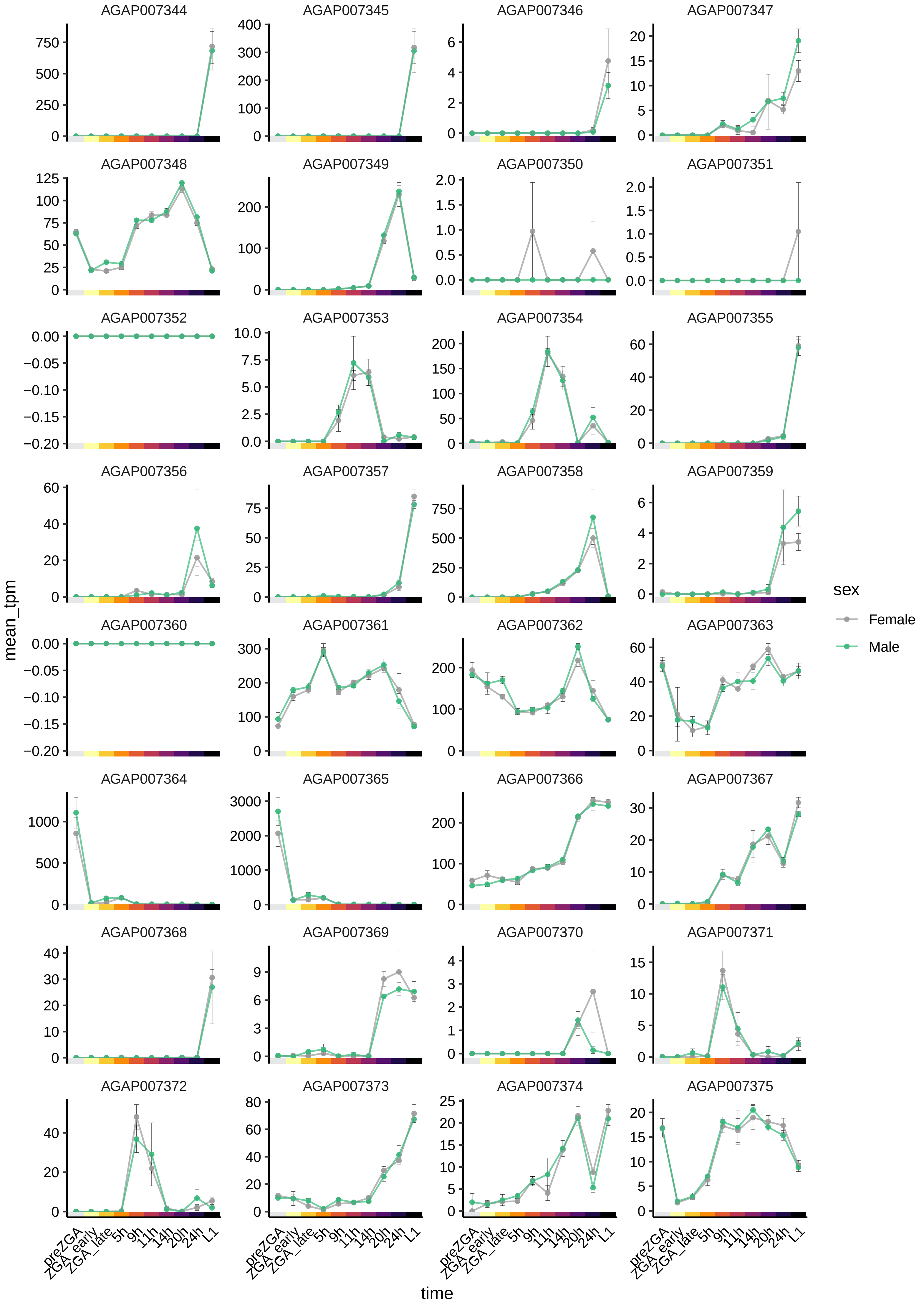

AGAP007344
AGAP007345
AGAP007346
AGAP007347
AGAP007348
AGAP007349
AGAP007350
AGAP007351
AGAP007352
AGAP007353
AGAP007354
AGAP007355
AGAP007356
AGAP007357
AGAP007358
AGAP007359
AGAP007360
AGAP007361
AGAP007362
AGAP007363
AGAP007364
AGAP007365
AGAP007366
AGAP007367
AGAP007368
AGAP007369
AGAP007370
AGAP007371
AGAP007372
AGAP007373
AGAP007374
AGAP007375
mean_tpm
time
preZGA
ZGA_early
ZGA_late
5h
9h
11h
14h
20h
24h
L1
sex
Female
Male

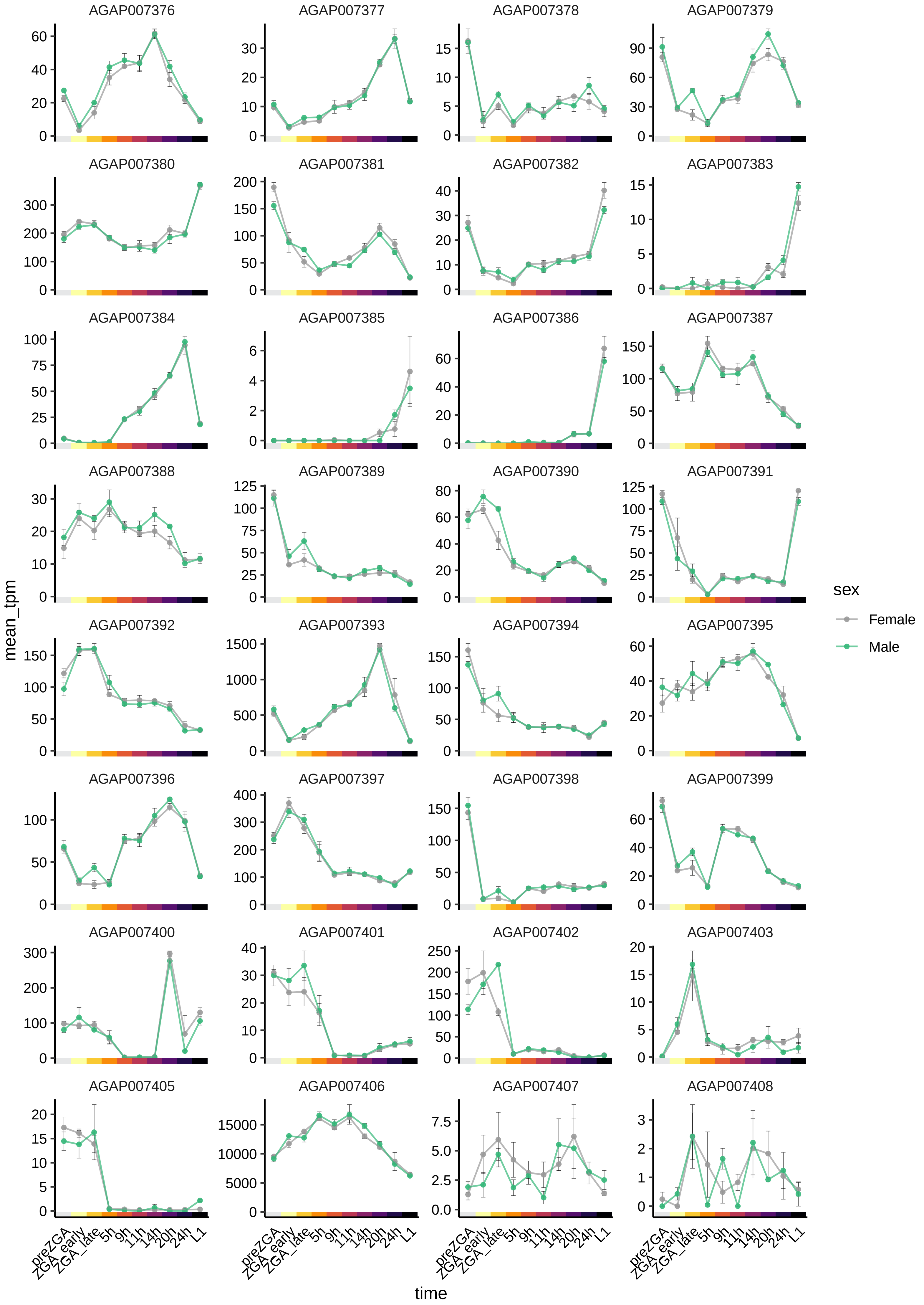
AGAP007376
AGAP007377
AGAP007378
AGAP007379
AGAP007380
AGAP007381
AGAP007382
AGAP007383
AGAP007384
AGAP007385
AGAP007386
AGAP007387
AGAP007388
AGAP007389
AGAP007390
AGAP007391
AGAP007392
AGAP007393
AGAP007394
AGAP007395
AGAP007396
AGAP007397
AGAP007398
AGAP007399
AGAP007400
AGAP007401
AGAP007402
AGAP007403
AGAP007405
AGAP007406
AGAP007407
AGAP007408
mean_tpm
time
preZGA
ZGA_early
ZGA_late
5h
9h
11h
14h
20h
24h
L1
sex
Female
Male

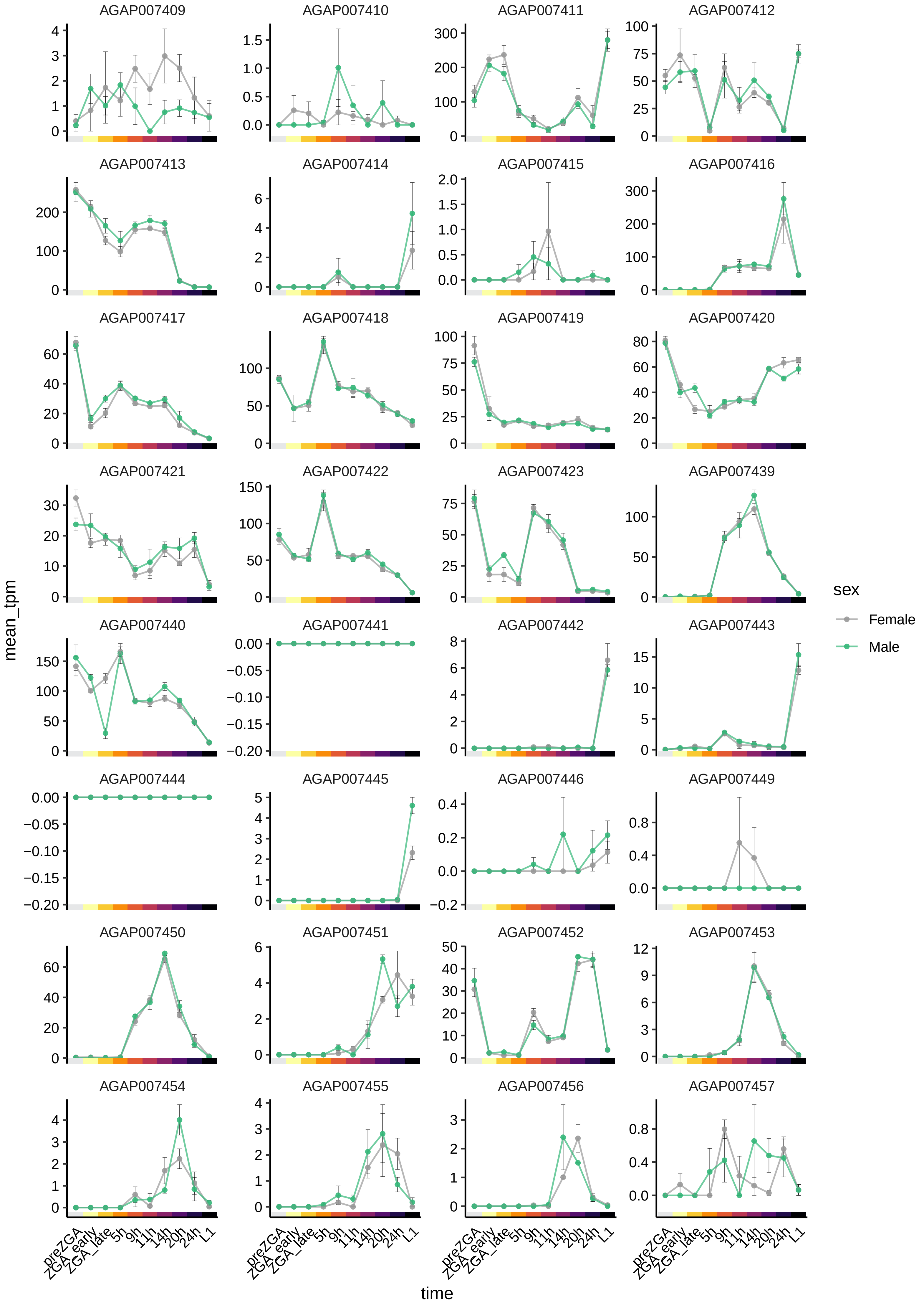

AGAP007409
AGAP007410
AGAP007411
AGAP007412
AGAP007413
AGAP007414
AGAP007415
AGAP007416
AGAP007417
AGAP007418
AGAP007419
AGAP007420
AGAP007421
AGAP007422
AGAP007423
AGAP007439
AGAP007440
AGAP007441
AGAP007442
AGAP007443
AGAP007444
AGAP007445
AGAP007446
AGAP007449
AGAP007450
AGAP007451
AGAP007452
AGAP007453
AGAP007454
AGAP007455
AGAP007456
AGAP007457
mean_tpm
time
preZGA
ZGA_early
ZGA_late
5h
9h
11h
14h
20h
24h
L1
sex
Female
Male

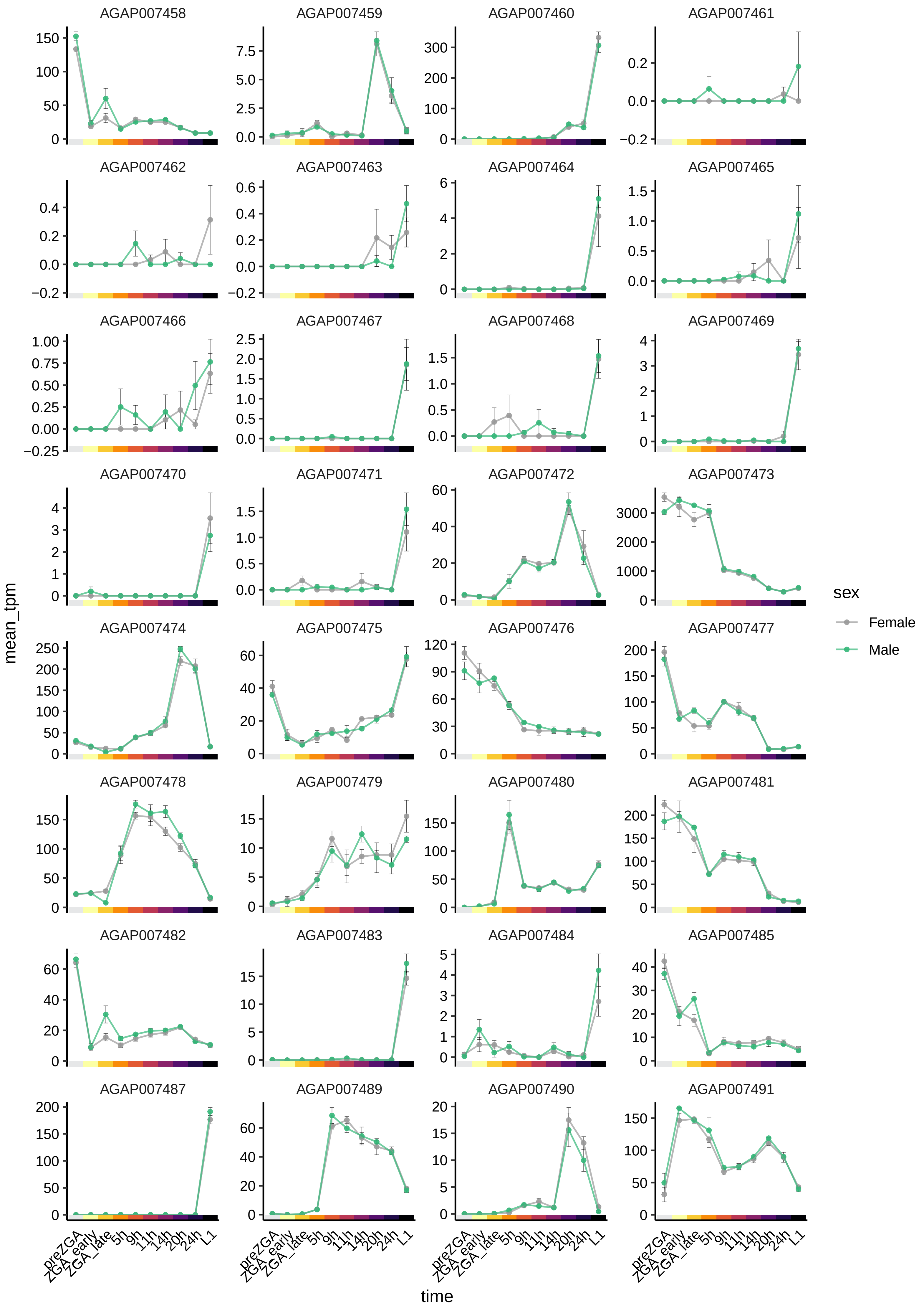

AGAP007458
AGAP007459
AGAP007460
AGAP007461
AGAP007462
AGAP007463
AGAP007464
AGAP007465
AGAP007466
AGAP007467
AGAP007468
AGAP007469
AGAP007470
AGAP007471
AGAP007472
AGAP007473
AGAP007474
AGAP007475
AGAP007476
AGAP007477
AGAP007478
AGAP007479
AGAP007480
AGAP007481
AGAP007482
AGAP007483
AGAP007484
AGAP007485
AGAP007487
AGAP007489
AGAP007490
AGAP007491
mean_tpm
time
preZGA
ZGA_early
ZGA_late
5h
9h
11h
14h
20h
24h
L1
sex
Female
Male

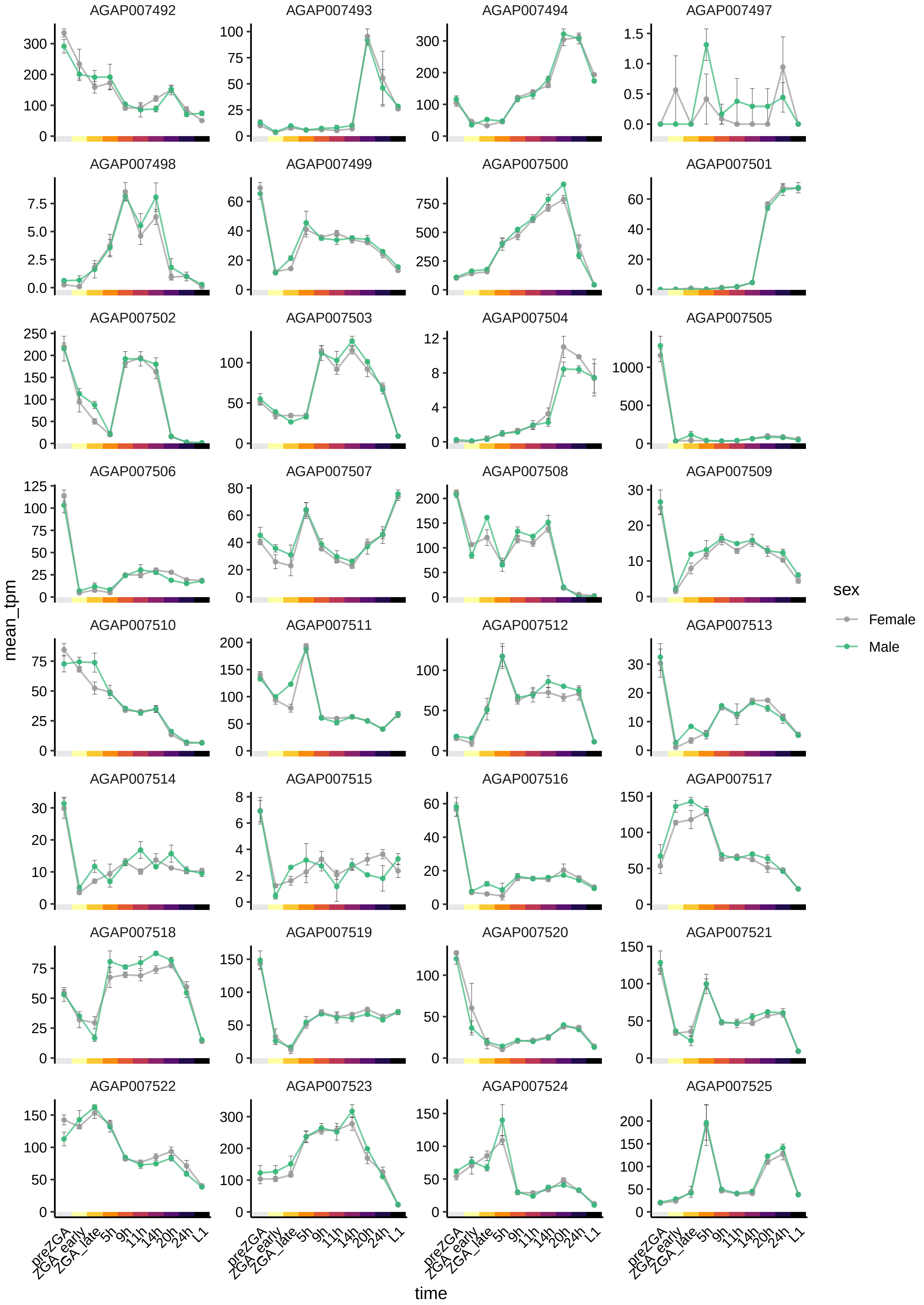

AGAP007492
AGAP007493
AGAP007494
AGAP007497
AGAP007498
AGAP007499
AGAP007500
AGAP007501
AGAP007502
AGAP007503
AGAP007504
AGAP007505
AGAP007506
AGAP007507
AGAP007508
AGAP007509
AGAP007510
AGAP007511
AGAP007512
AGAP007513
AGAP007514
AGAP007515
AGAP007516
AGAP007517
AGAP007518
AGAP007519
AGAP007520
AGAP007521
AGAP007522
AGAP007523
AGAP007524
AGAP007525
mean_tpm
time
preZGA
ZGA_early
ZGA_late
5h
9h
11h
14h
20h
24h
L1
sex
Female
Male

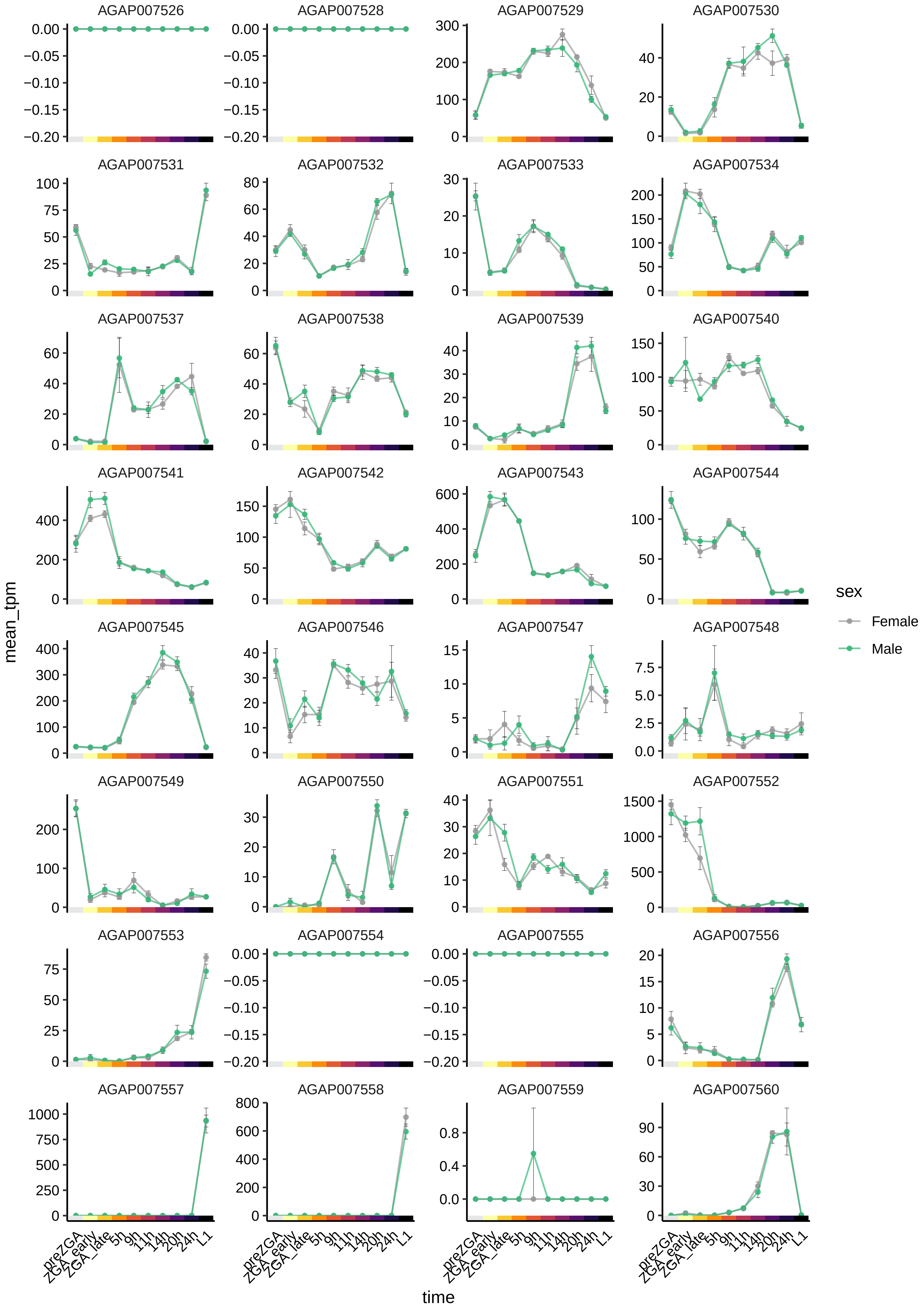
AGAP007526
AGAP007528
AGAP007529
AGAP007530
AGAP007531
AGAP007532
AGAP007533
AGAP007534
AGAP007537
AGAP007538
AGAP007539
AGAP007540
AGAP007541
AGAP007542
AGAP007543
AGAP007544
AGAP007545
AGAP007546
AGAP007547
AGAP007548
AGAP007549
AGAP007550
AGAP007551
AGAP007552
AGAP007553
AGAP007554
AGAP007555
AGAP007556
AGAP007557
AGAP007558
AGAP007559
AGAP007560
mean_tpm
time
preZGA
ZGA_early
ZGA_late
5h
9h
11h
14h
20h
24h
L1
sex
Female
Male

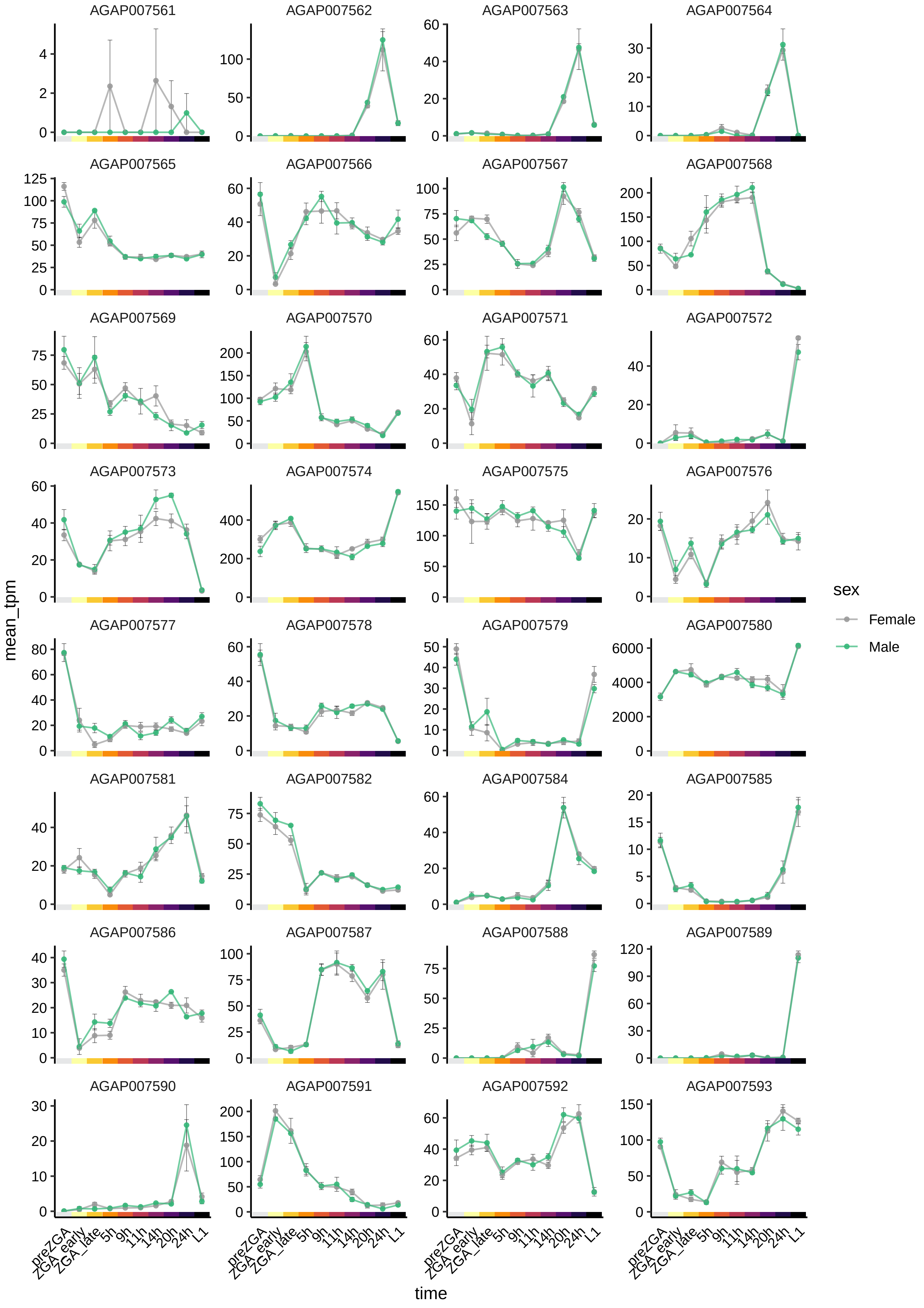
AGAP007561
AGAP007562
AGAP007563
AGAP007564
AGAP007565
AGAP007566
AGAP007567
AGAP007568
AGAP007569
AGAP007570
AGAP007571
AGAP007572
AGAP007573
AGAP007574
AGAP007575
AGAP007576
AGAP007577
AGAP007578
AGAP007579
AGAP007580
AGAP007581
AGAP007582
AGAP007584
AGAP007585
AGAP007586
AGAP007587
AGAP007588
AGAP007589
AGAP007590
AGAP007591
AGAP007592
AGAP007593
mean_tpm
time
preZGA
ZGA_early
ZGA_late
5h
9h
11h
14h
20h
24h
L1
sex
Female
Male

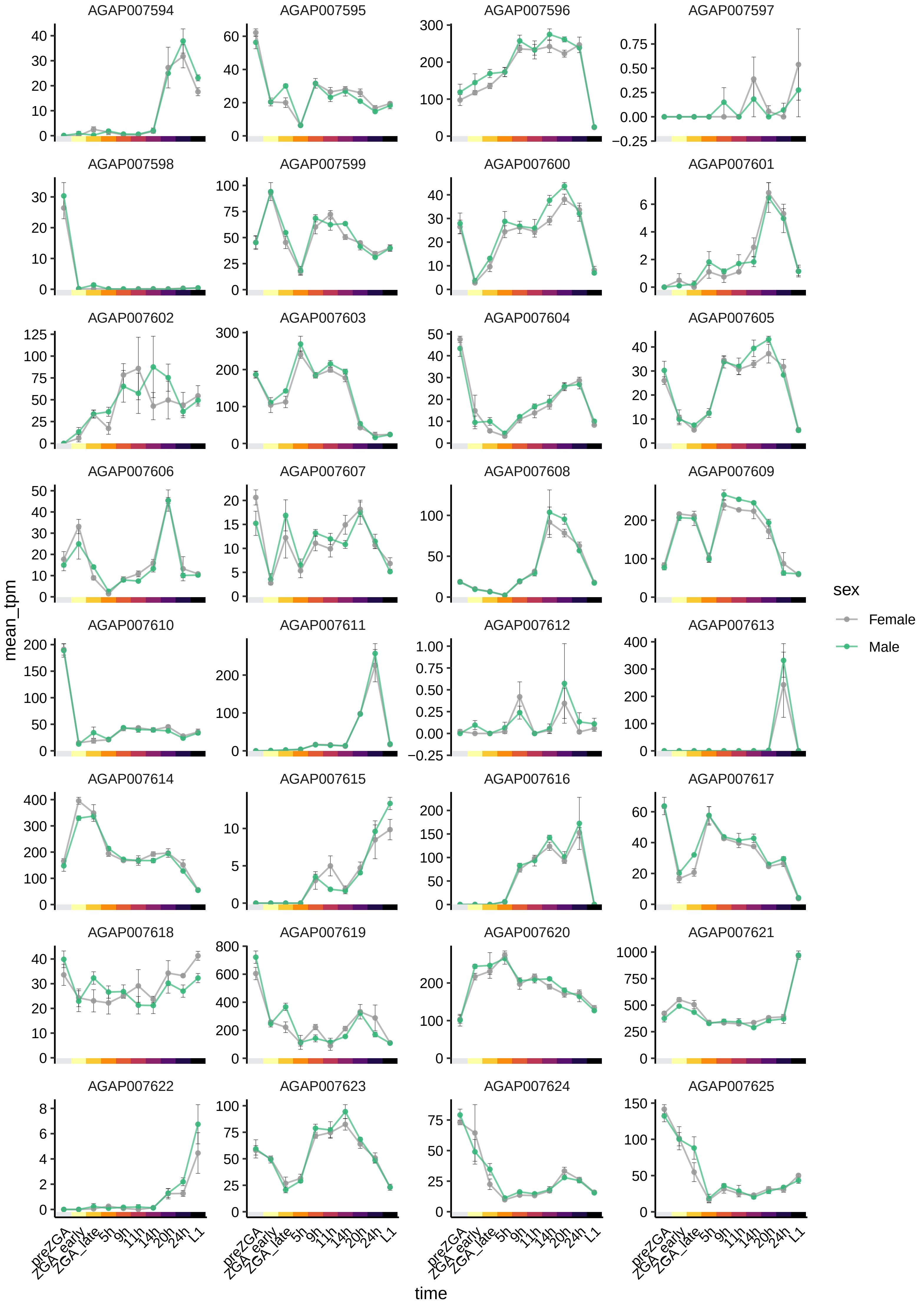
AGAP007594
AGAP007595
AGAP007596
AGAP007597
AGAP007598
AGAP007599
AGAP007600
AGAP007601
AGAP007602
AGAP007603
AGAP007604
AGAP007605
AGAP007606
AGAP007607
AGAP007608
AGAP007609
AGAP007610
AGAP007611
AGAP007612
AGAP007613
AGAP007614
AGAP007615
AGAP007616
AGAP007617
AGAP007618
AGAP007619
AGAP007620
AGAP007621
AGAP007622
AGAP007623
AGAP007624
AGAP007625
mean_tpm
time
preZGA
ZGA_early
ZGA_late
5h
9h
11h
14h
20h
24h
L1
sex
Female
Male

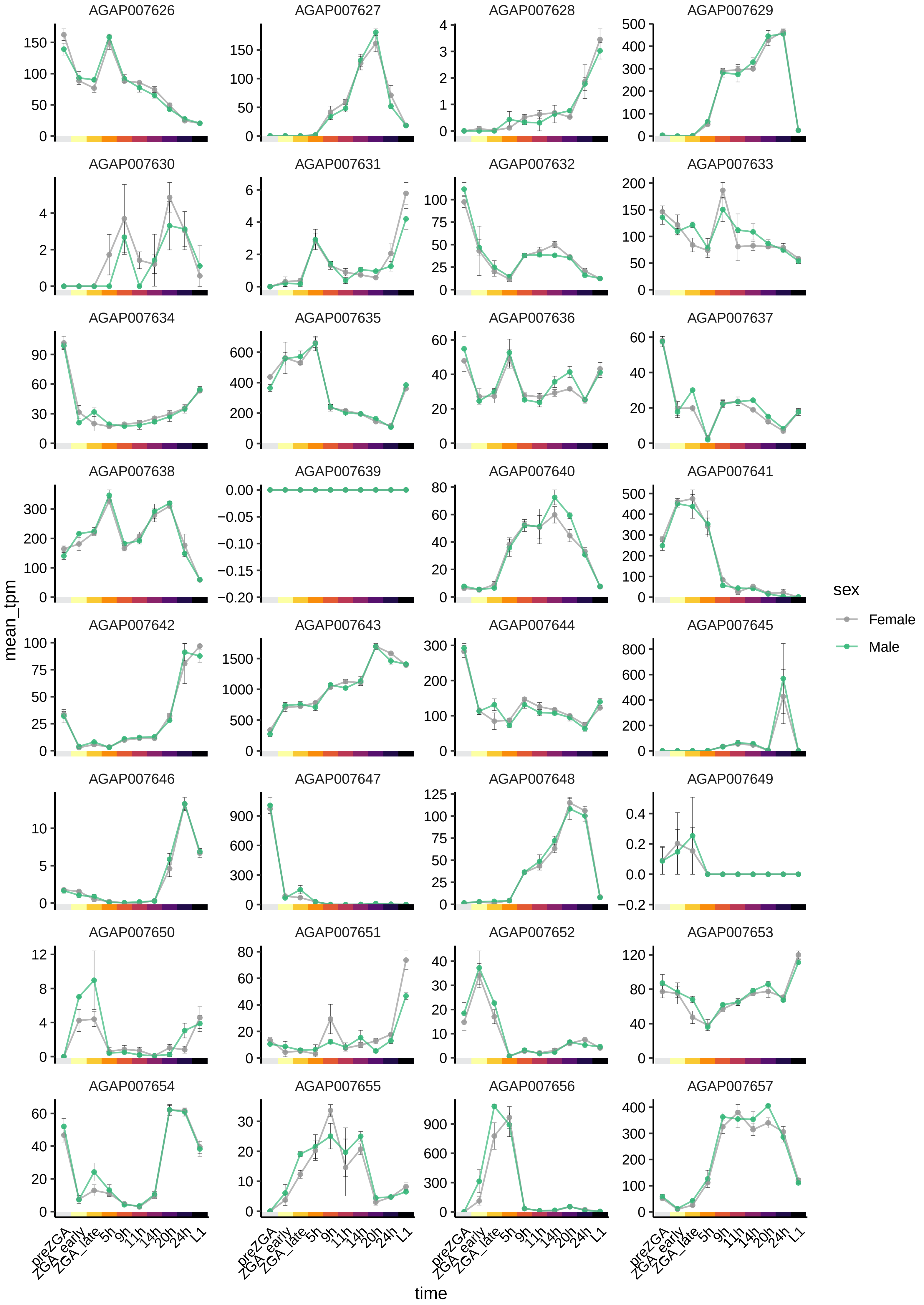

AGAP007626
AGAP007627
AGAP007628
AGAP007629
AGAP007630
AGAP007631
AGAP007632
AGAP007633
AGAP007634
AGAP007635
AGAP007636
AGAP007637
AGAP007638
AGAP007639
AGAP007640
AGAP007641
AGAP007642
AGAP007643
AGAP007644
AGAP007645
AGAP007646
AGAP007647
AGAP007648
AGAP007649
AGAP007650
AGAP007651
AGAP007652
AGAP007653
AGAP007654
AGAP007655
AGAP007656
AGAP007657
mean_tpm
time
preZGA
ZGA_early
ZGA_late
5h
9h
11h
14h
20h
24h
L1
sex
Female
Male

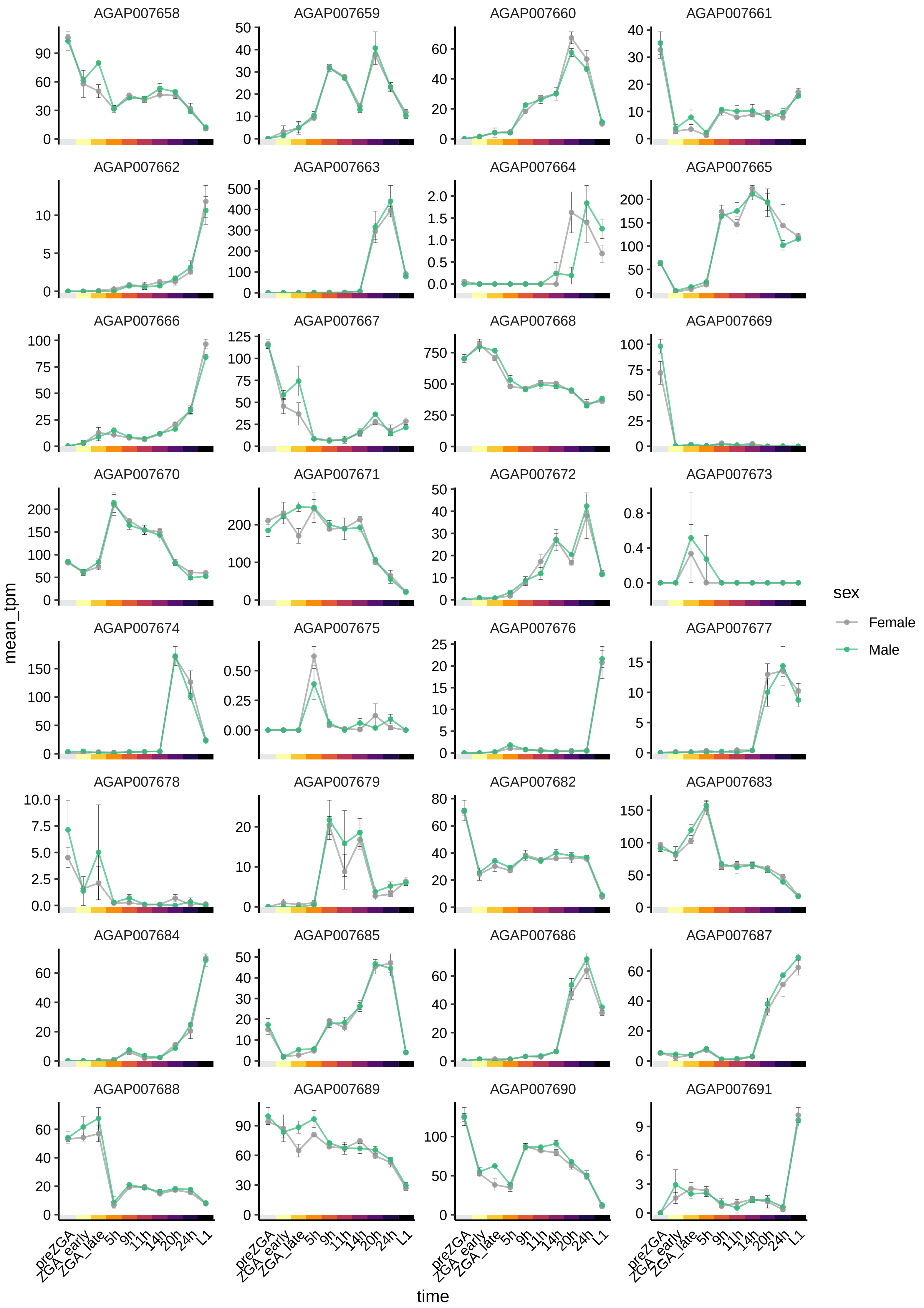

AGAP007658
AGAP007659
AGAP007660
AGAP007661
AGAP007662
AGAP007663
AGAP007664
AGAP007665
AGAP007666
AGAP007667
AGAP007668
AGAP007669
AGAP007670
AGAP007671
AGAP007672
AGAP007673
AGAP007674
AGAP007675
AGAP007676
AGAP007677
AGAP007678
AGAP007679
AGAP007682
AGAP007683
AGAP007684
AGAP007685
AGAP007686
AGAP007687
AGAP007688
AGAP007689
AGAP007690
AGAP007691
mean_tpm
time
preZGA
ZGA_early
ZGA_late
5h
9h
11h
14h
20h
24h
L1
sex
Female
Male

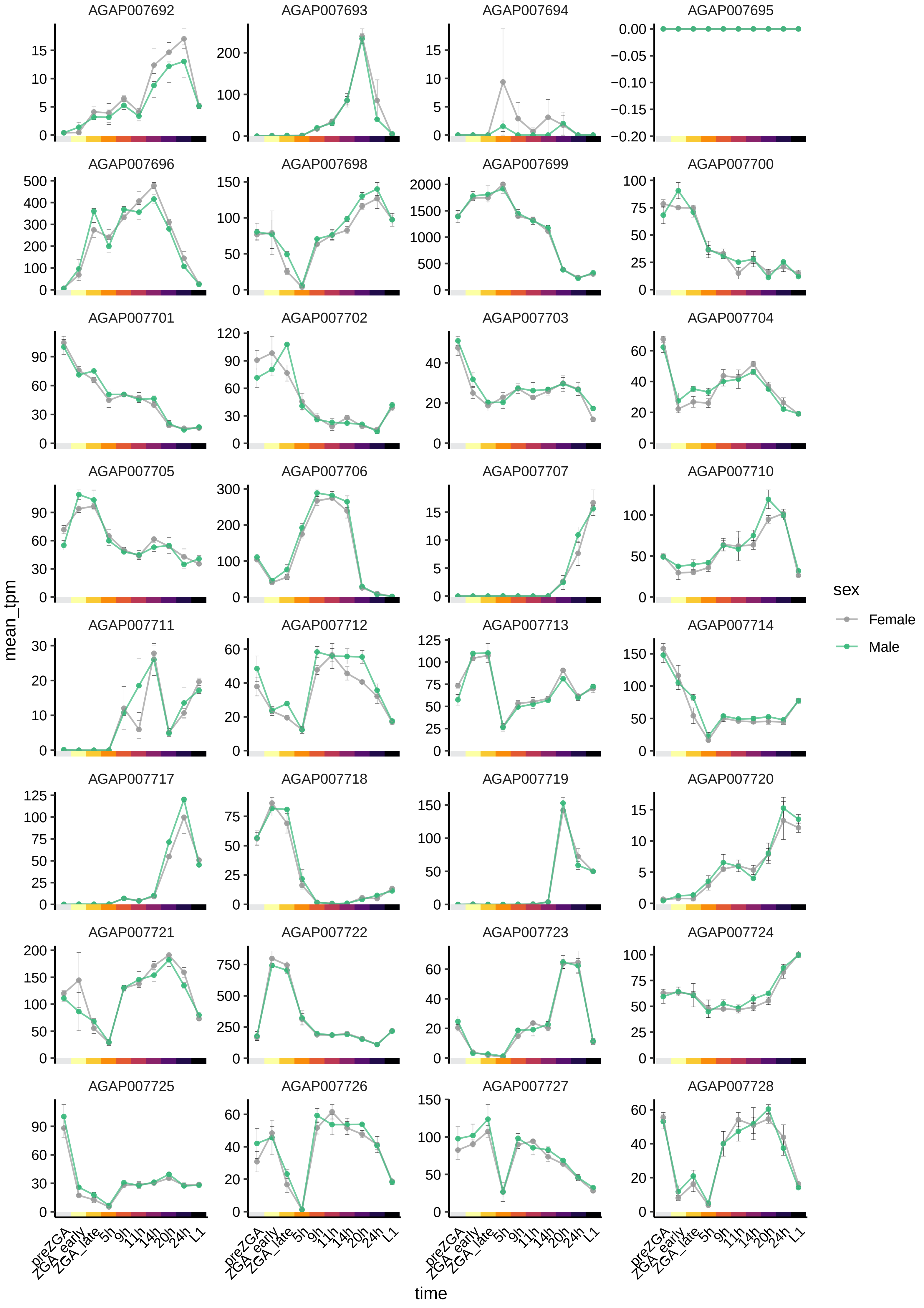
AGAP007692
AGAP007693
AGAP007694
AGAP007695
AGAP007696
AGAP007698
AGAP007699
AGAP007700
AGAP007701
AGAP007702
AGAP007703
AGAP007704
AGAP007705
AGAP007706
AGAP007707
AGAP007710
AGAP007711
AGAP007712
AGAP007713
AGAP007714
AGAP007717
AGAP007718
AGAP007719
AGAP007720
AGAP007721
AGAP007722
AGAP007723
AGAP007724
AGAP007725
AGAP007726
AGAP007727
AGAP007728
mean_tpm
time
preZGA
ZGA_early
ZGA_late
5h
9h
11h
14h
20h
24h
L1
sex
Female
Male

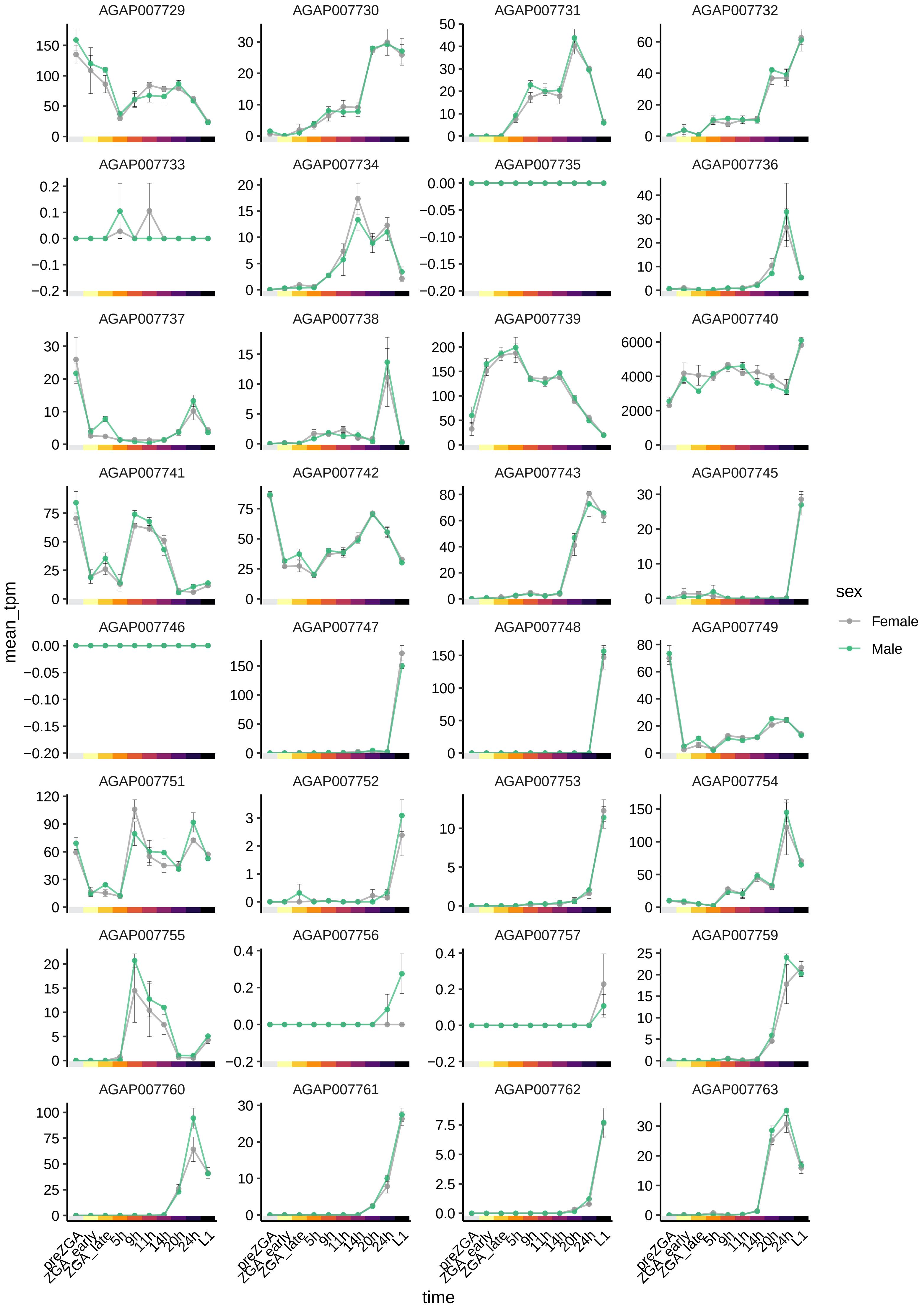
AGAP007729
AGAP007730
AGAP007731
AGAP007732
AGAP007733
AGAP007734
AGAP007735
AGAP007736
AGAP007737
AGAP007738
AGAP007739
AGAP007740
AGAP007741
AGAP007742
AGAP007743
AGAP007745
AGAP007746
AGAP007747
AGAP007748
AGAP007749
AGAP007751
AGAP007752
AGAP007753
AGAP007754
AGAP007755
AGAP007756
AGAP007757
AGAP007759
AGAP007760
AGAP007761
AGAP007762
AGAP007763
mean_tpm
time
preZGA
ZGA_early
ZGA_late
5h
9h
11h
14h
20h
24h
L1
sex
Female
Male

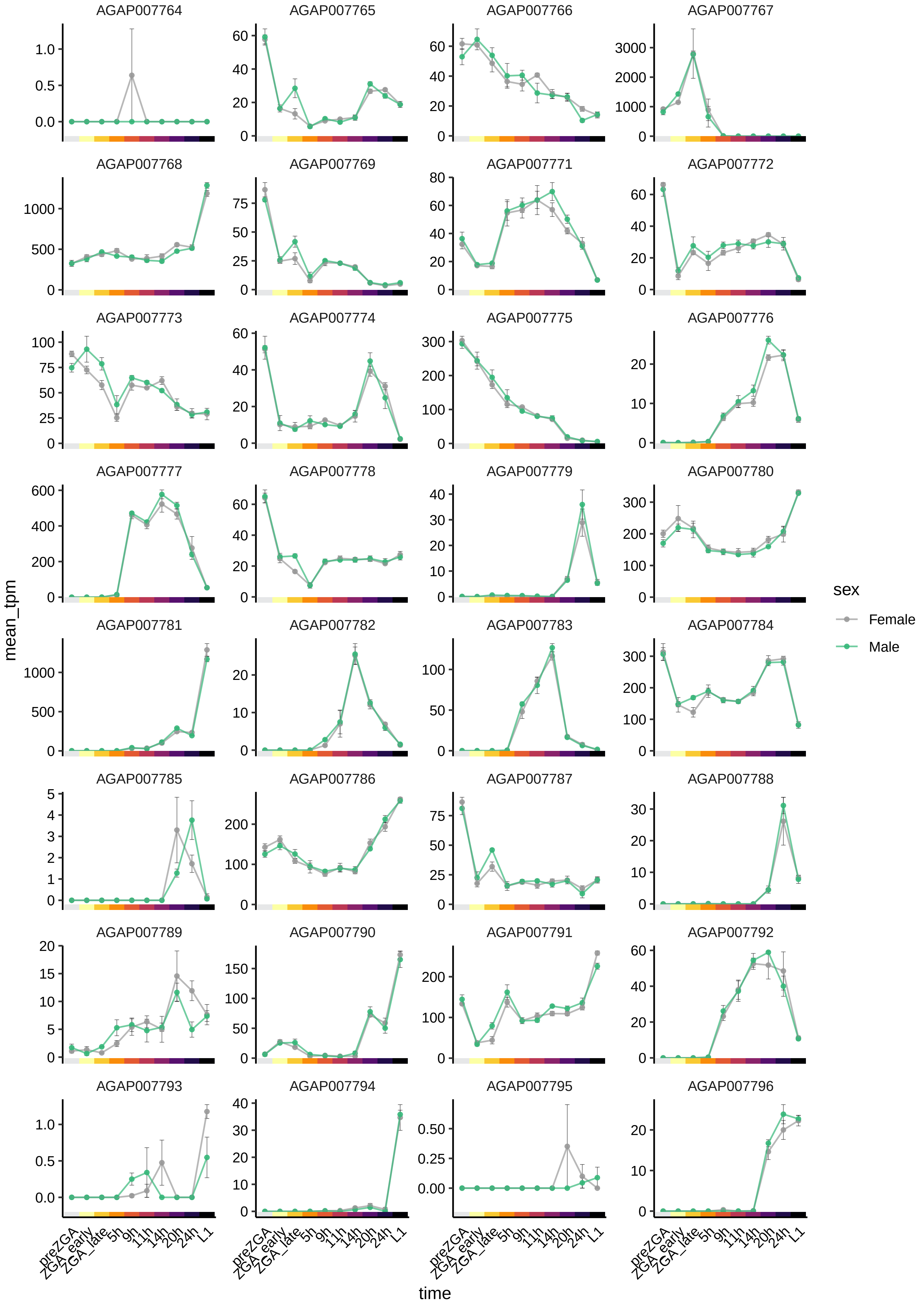
AGAP007764
AGAP007765
AGAP007766
AGAP007767
AGAP007768
AGAP007769
AGAP007771
AGAP007772
AGAP007773
AGAP007774
AGAP007775
AGAP007776
AGAP007777
AGAP007778
AGAP007779
AGAP007780
AGAP007781
AGAP007782
AGAP007783
AGAP007784
AGAP007785
AGAP007786
AGAP007787
AGAP007788
AGAP007789
AGAP007790
AGAP007791
AGAP007792
AGAP007793
AGAP007794
AGAP007795
AGAP007796
mean_tpm
time
preZGA
ZGA_early
ZGA_late
5h
9h
11h
14h
20h
24h
L1
sex
Female
Male

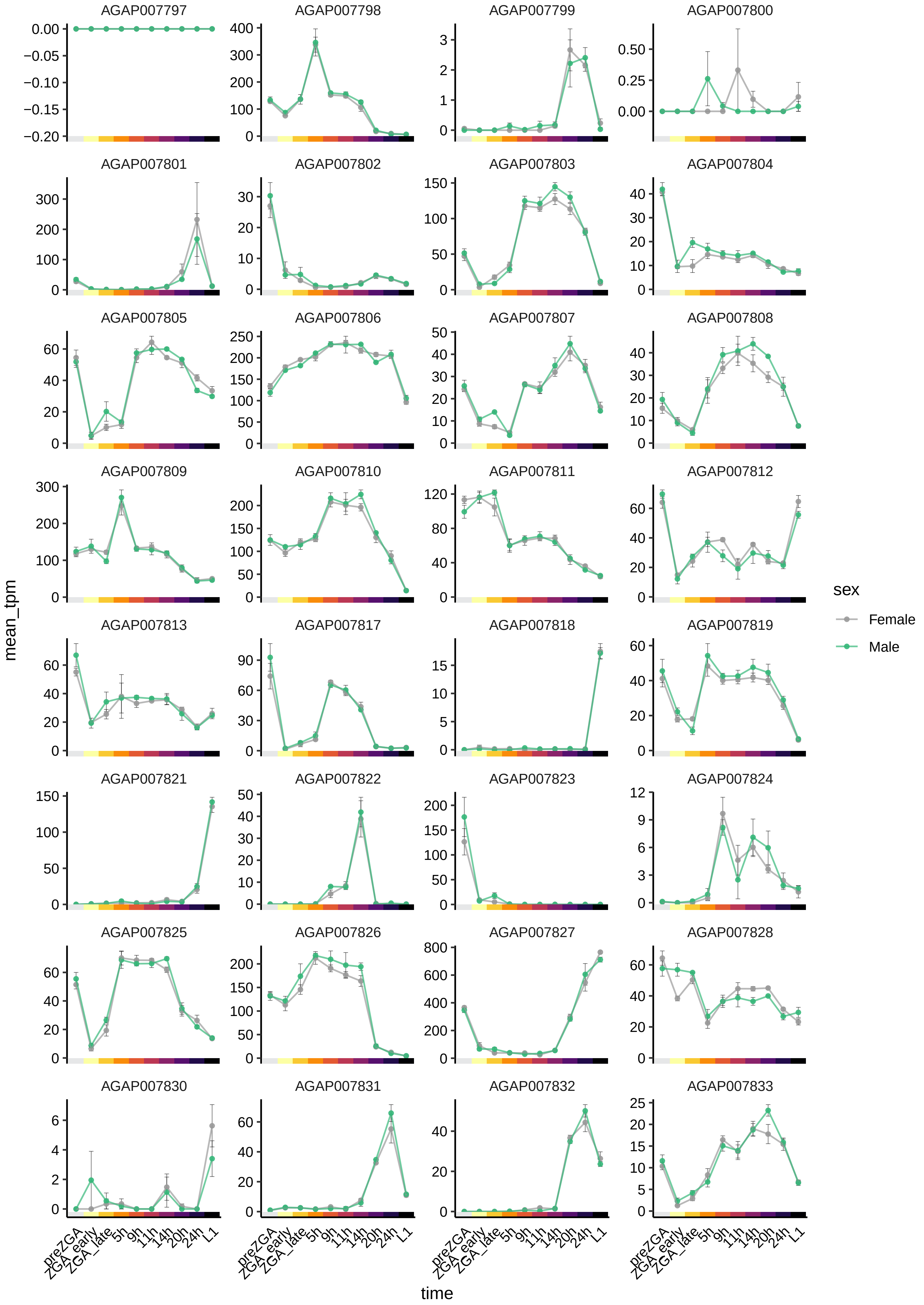

AGAP007797
AGAP007798
AGAP007799
AGAP007800
AGAP007801
AGAP007802
AGAP007803
AGAP007804
AGAP007805
AGAP007806
AGAP007807
AGAP007808
AGAP007809
AGAP007810
AGAP007811
AGAP007812
AGAP007813
AGAP007817
AGAP007818
AGAP007819
AGAP007821
AGAP007822
AGAP007823
AGAP007824
AGAP007825
AGAP007826
AGAP007827
AGAP007828
AGAP007830
AGAP007831
AGAP007832
AGAP007833
mean_tpm
time
preZGA
ZGA_early
ZGA_late
5h
9h
11h
14h
20h
24h
L1
sex
Female
Male

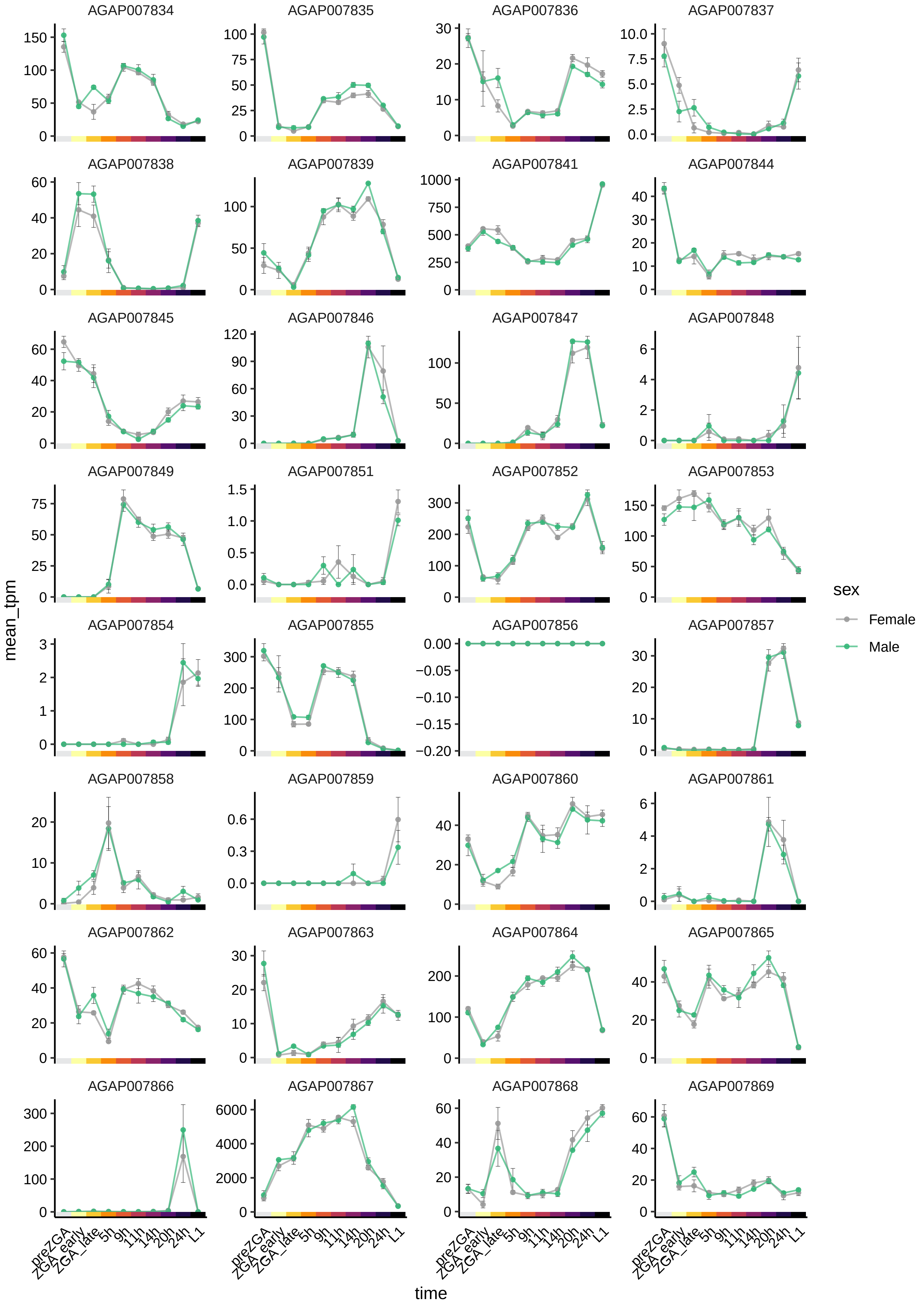

AGAP007834
AGAP007835
AGAP007836
AGAP007837
AGAP007838
AGAP007839
AGAP007841
AGAP007844
AGAP007845
AGAP007846
AGAP007847
AGAP007848
AGAP007849
AGAP007851
AGAP007852
AGAP007853
AGAP007854
AGAP007855
AGAP007856
AGAP007857
AGAP007858
AGAP007859
AGAP007860
AGAP007861
AGAP007862
AGAP007863
AGAP007864
AGAP007865
AGAP007866
AGAP007867
AGAP007868
AGAP007869
mean_tpm
time
preZGA
ZGA_early
ZGA_late
5h
9h
11h
14h
20h
24h
L1
sex
Female
Male

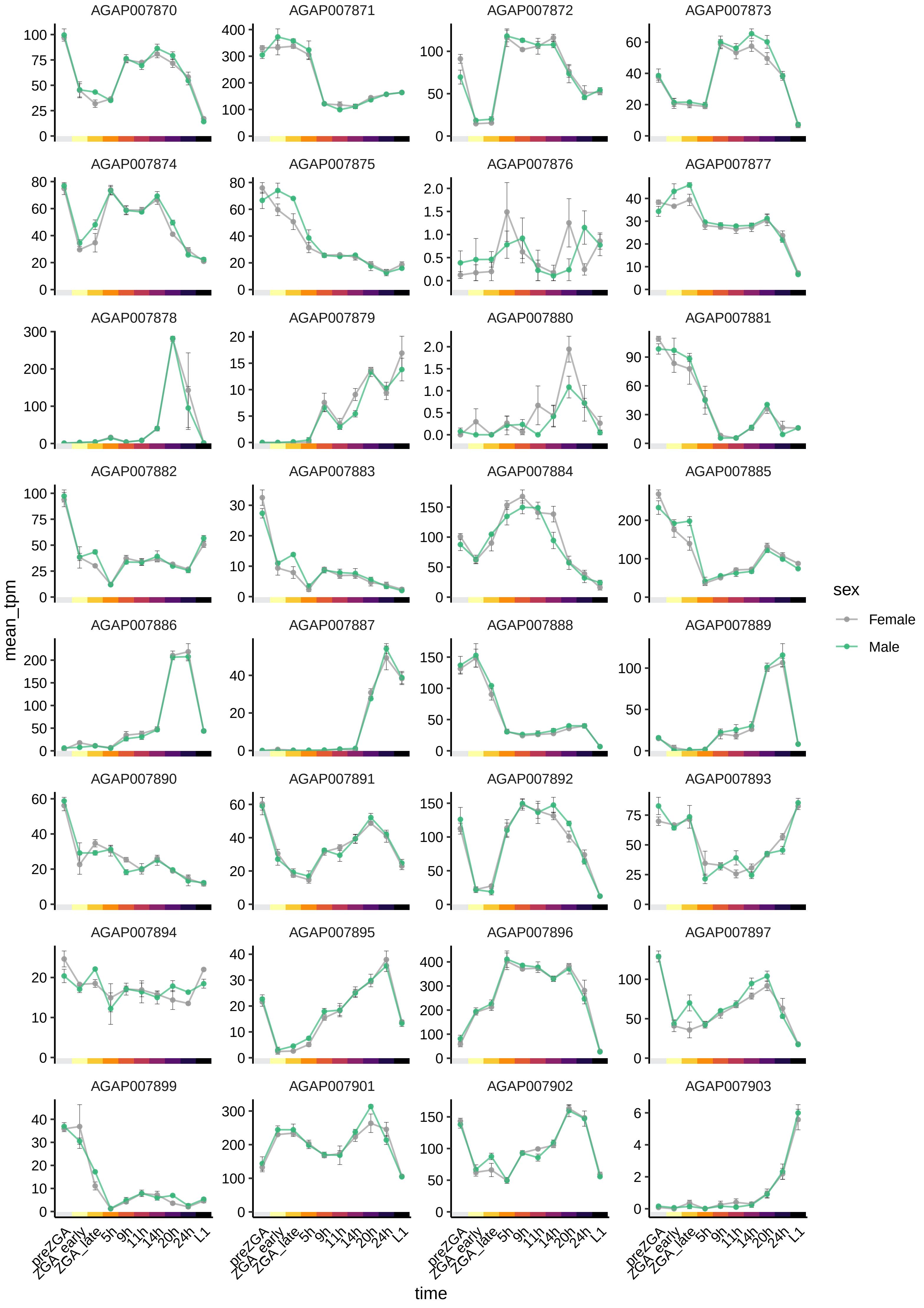
AGAP007870
AGAP007871
AGAP007872
AGAP007873
AGAP007874
AGAP007875
AGAP007876
AGAP007877
AGAP007878
AGAP007879
AGAP007880
AGAP007881
AGAP007882
AGAP007883
AGAP007884
AGAP007885
AGAP007886
AGAP007887
AGAP007888
AGAP007889
AGAP007890
AGAP007891
AGAP007892
AGAP007893
AGAP007894
AGAP007895
AGAP007896
AGAP007897
AGAP007899
AGAP007901
AGAP007902
AGAP007903
mean_tpm
time
preZGA
ZGA_early
ZGA_late
5h
9h
11h
14h
20h
24h
L1
sex
Female
Male

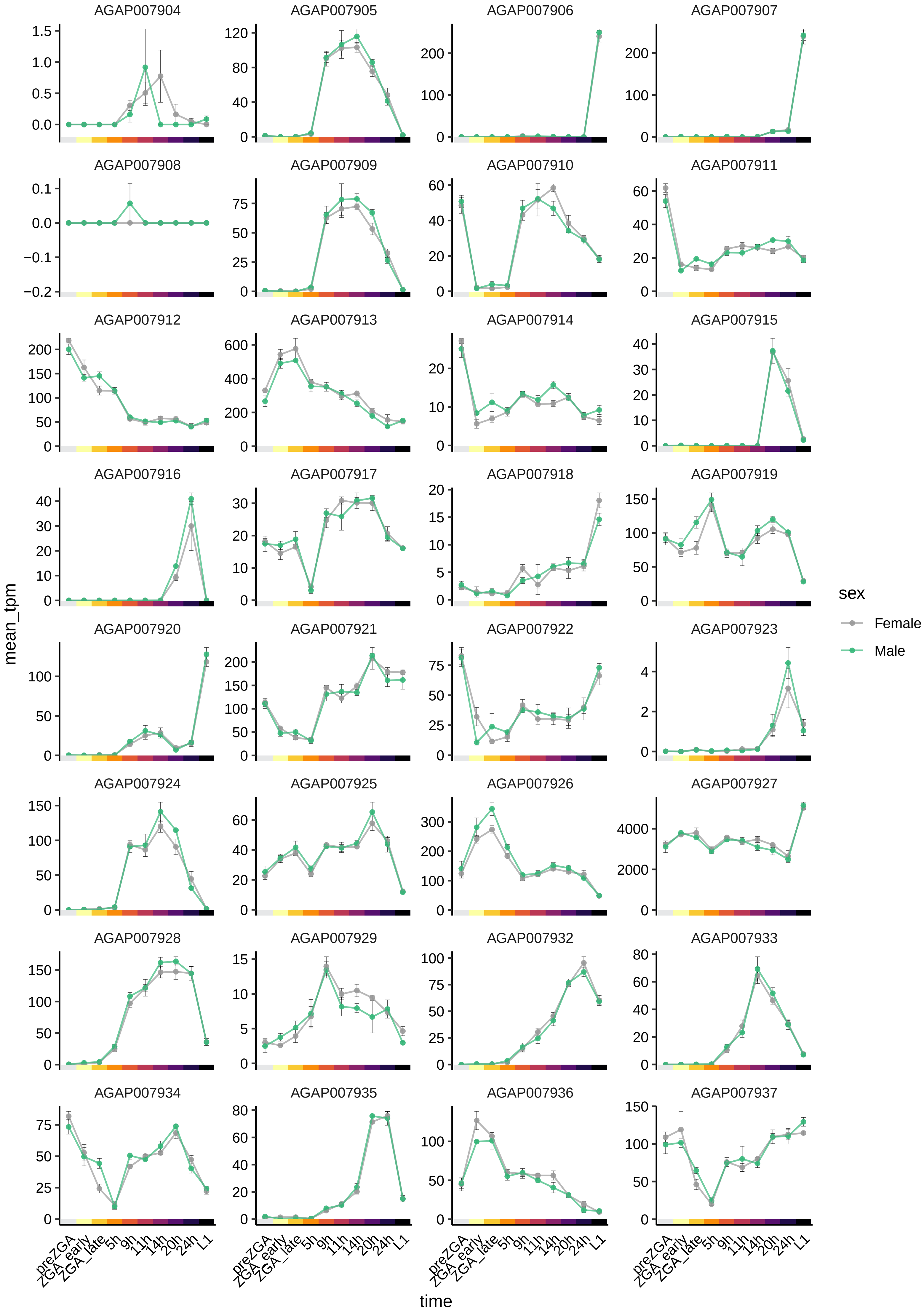
AGAP007904
AGAP007905
AGAP007906
AGAP007907
AGAP007908
AGAP007909
AGAP007910
AGAP007911
AGAP007912
AGAP007913
AGAP007914
AGAP007915
AGAP007916
AGAP007917
AGAP007918
AGAP007919
AGAP007920
AGAP007921
AGAP007922
AGAP007923
AGAP007924
AGAP007925
AGAP007926
AGAP007927
AGAP007928
AGAP007929
AGAP007932
AGAP007933
AGAP007934
AGAP007935
AGAP007936
AGAP007937
mean_tpm
time
preZGA
ZGA_early
ZGA_late
5h
9h
11h
14h
20h
24h
L1
sex
Female
Male

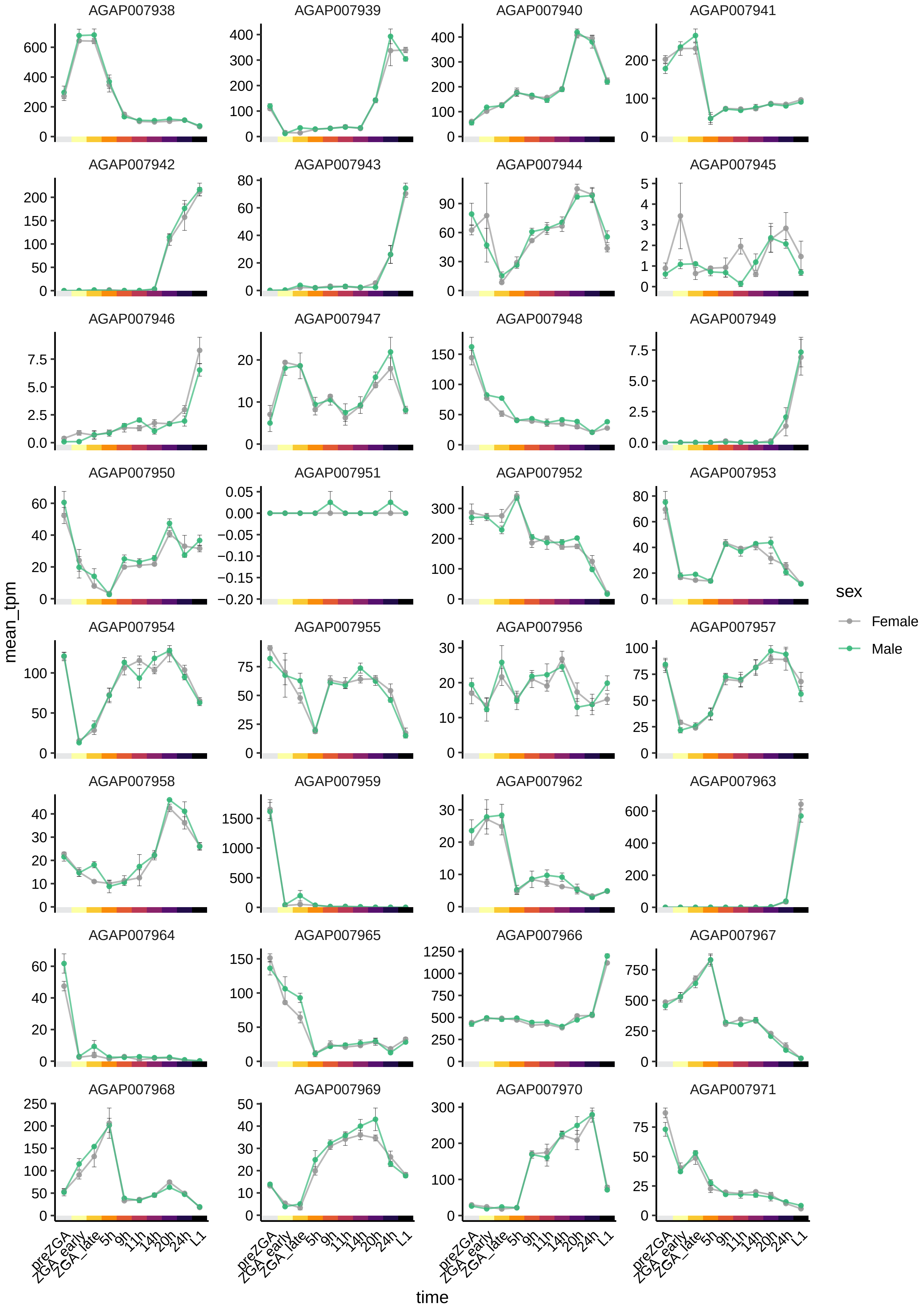

AGAP007938
AGAP007939
AGAP007940
AGAP007941
AGAP007942
AGAP007943
AGAP007944
AGAP007945
AGAP007946
AGAP007947
AGAP007948
AGAP007949
AGAP007950
AGAP007951
AGAP007952
AGAP007953
AGAP007954
AGAP007955
AGAP007956
AGAP007957
AGAP007958
AGAP007959
AGAP007962
AGAP007963
AGAP007964
AGAP007965
AGAP007966
AGAP007967
AGAP007968
AGAP007969
AGAP007970
AGAP007971
mean_tpm
time
preZGA
ZGA_early
ZGA_late
5h
9h
11h
14h
20h
24h
L1
sex
Female
Male

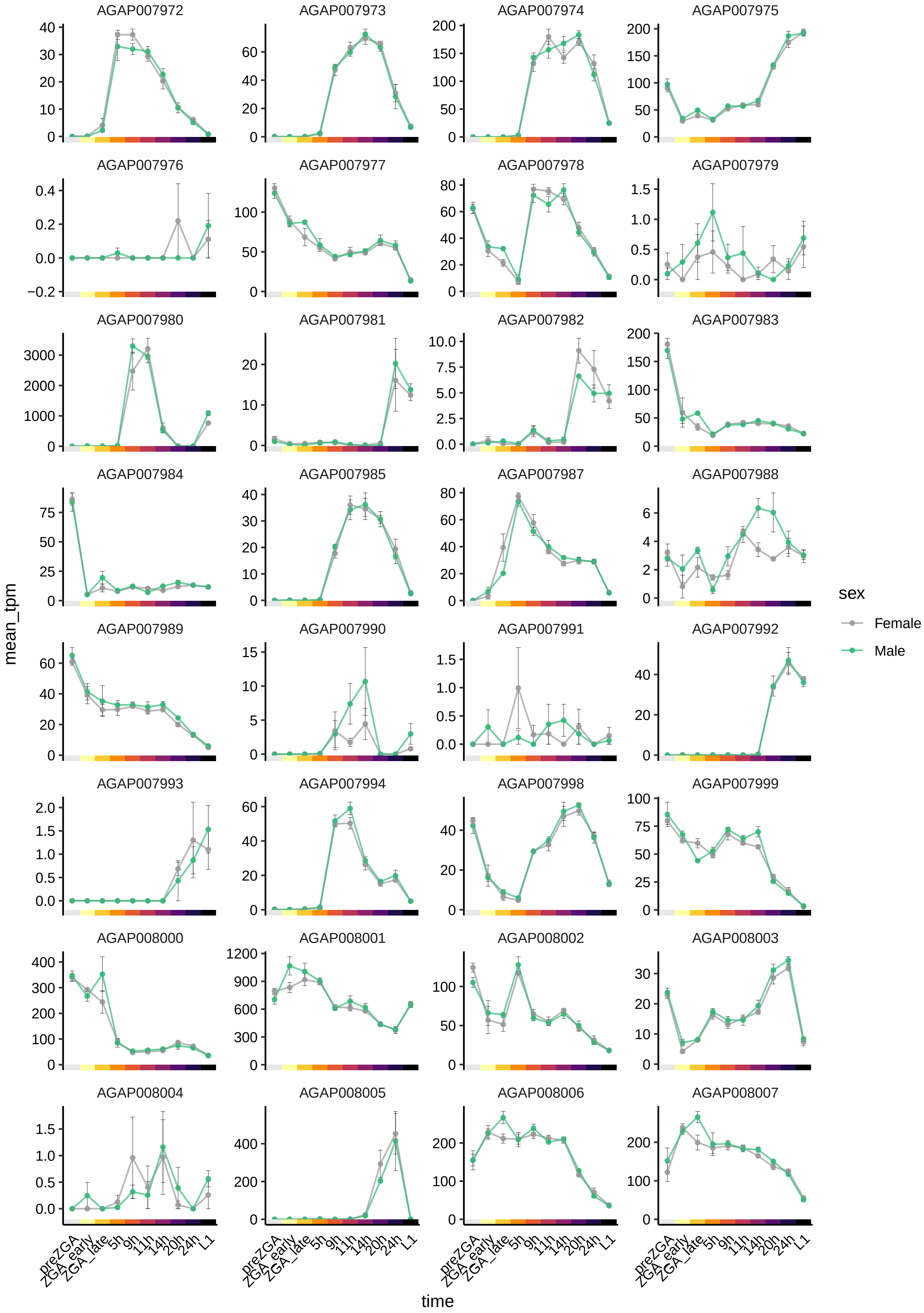
AGAP007972
AGAP007973
AGAP007974
AGAP007975
AGAP007976
AGAP007977
AGAP007978
AGAP007979
AGAP007980
AGAP007981
AGAP007982
AGAP007983
AGAP007984
AGAP007985
AGAP007987
AGAP007988
AGAP007989
AGAP007990
AGAP007991
AGAP007992
AGAP007993
AGAP007994
AGAP007998
AGAP007999
AGAP008000
AGAP008001
AGAP008002
AGAP008003
AGAP008004
AGAP008005
AGAP008006
AGAP008007
mean_tpm
time
preZGA
ZGA_early
ZGA_late
5h
9h
11h
14h
20h
24h
L1
sex
Female
Male

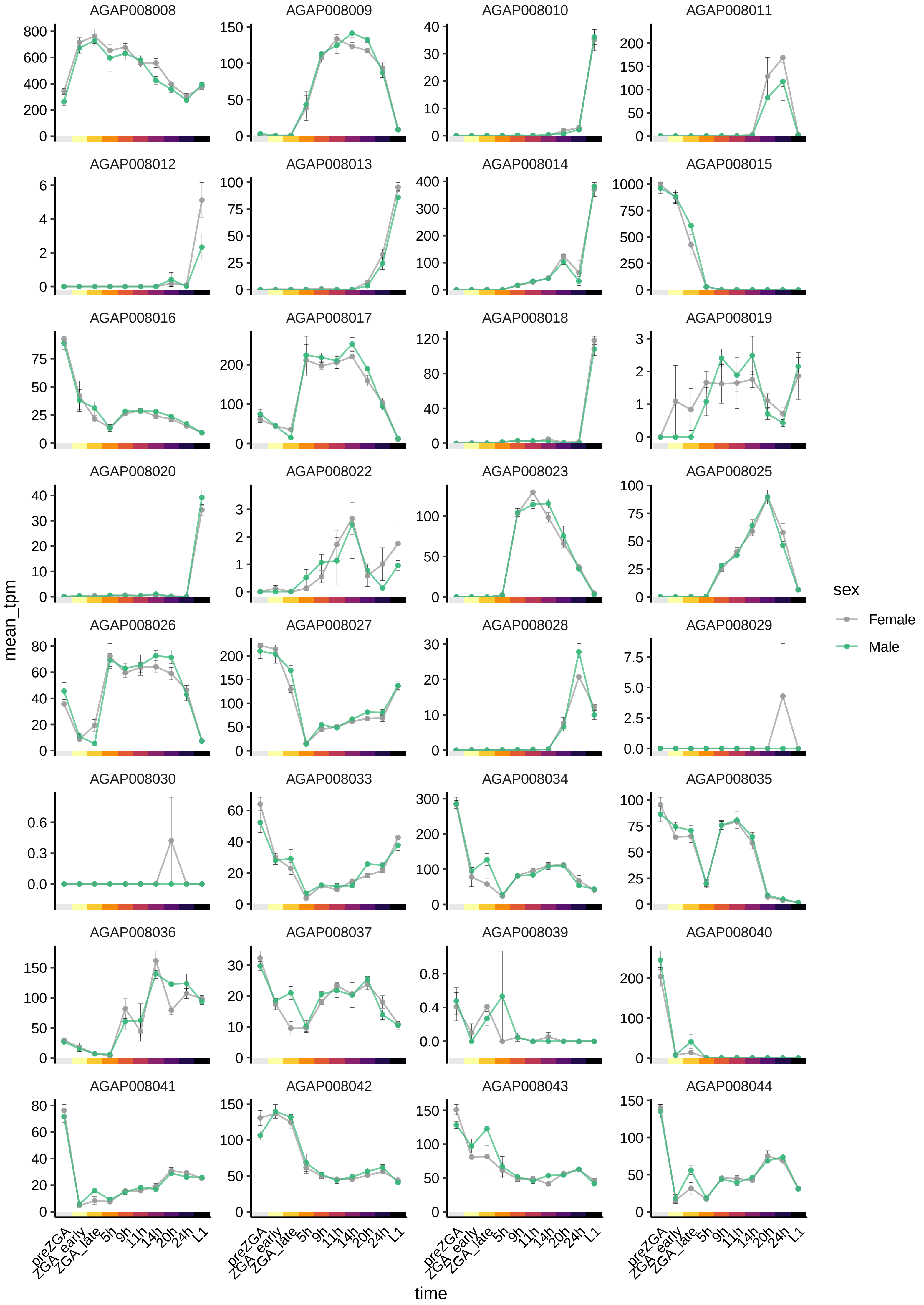

AGAP008008
AGAP008009
AGAP008010
AGAP008011
AGAP008012
AGAP008013
AGAP008014
AGAP008015
AGAP008016
AGAP008017
AGAP008018
AGAP008019
AGAP008020
AGAP008022
AGAP008023
AGAP008025
AGAP008026
AGAP008027
AGAP008028
AGAP008029
AGAP008030
AGAP008033
AGAP008034
AGAP008035
AGAP008036
AGAP008037
AGAP008039
AGAP008040
AGAP008041
AGAP008042
AGAP008043
AGAP008044
mean_tpm
time
preZGA
ZGA_early
ZGA_late
5h
9h
11h
14h
20h
24h
L1
sex
Female
Male

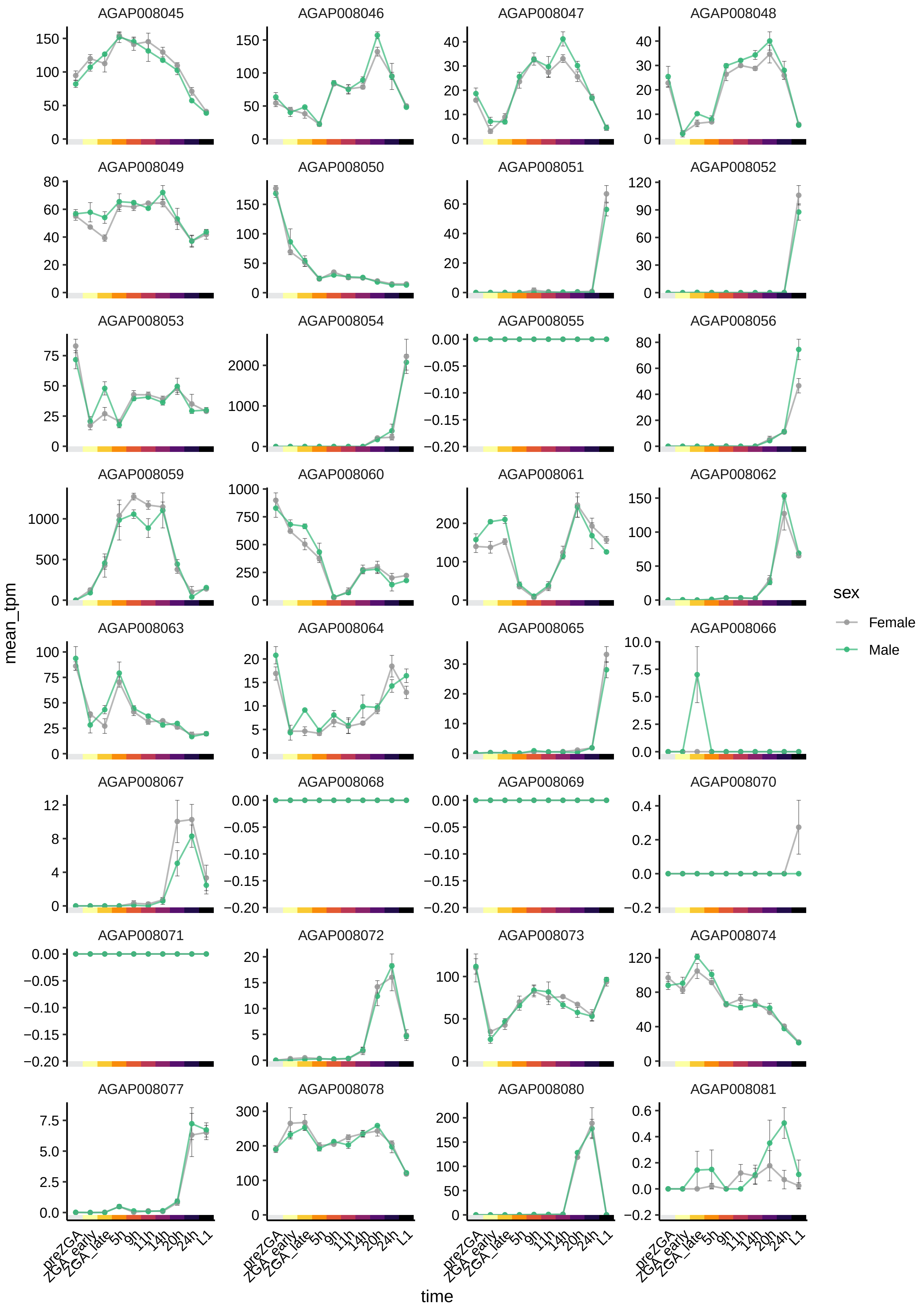
AGAP008045
AGAP008046
AGAP008047
AGAP008048
AGAP008049
AGAP008050
AGAP008051
AGAP008052
AGAP008053
AGAP008054
AGAP008055
AGAP008056
AGAP008059
AGAP008060
AGAP008061
AGAP008062
AGAP008063
AGAP008064
AGAP008065
AGAP008066
AGAP008067
AGAP008068
AGAP008069
AGAP008070
AGAP008071
AGAP008072
AGAP008073
AGAP008074
AGAP008077
AGAP008078
AGAP008080
AGAP008081
mean_tpm
preZGA
ZGA_early
ZGA_late
5h
9h
11h
14h
20h
24h
L1
time
sex
Female
Male

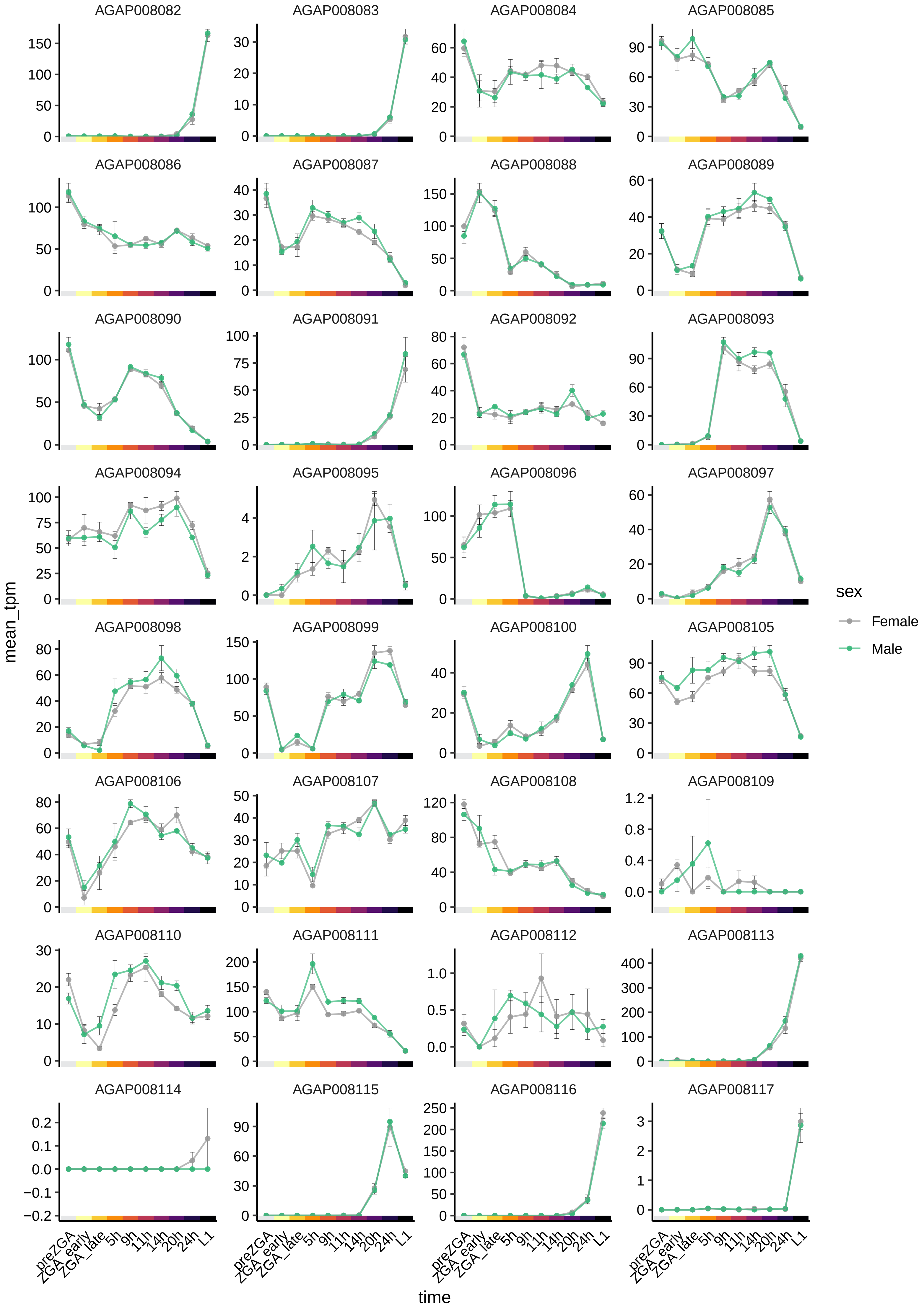
AGAP008082
AGAP008083
AGAP008084
AGAP008085
AGAP008086
AGAP008087
AGAP008088
AGAP008089
AGAP008090
AGAP008091
AGAP008092
AGAP008093
AGAP008094
AGAP008095
AGAP008096
AGAP008097
AGAP008098
AGAP008099
AGAP008100
AGAP008105
AGAP008106
AGAP008107
AGAP008108
AGAP008109
AGAP008110
AGAP008111
AGAP008112
AGAP008113
AGAP008114
AGAP008115
AGAP008116
AGAP008117
mean_tpm
time
preZGA
ZGA_early
ZGA_late
5h
9h
11h
14h
20h
24h
L1
sex
Female
Male

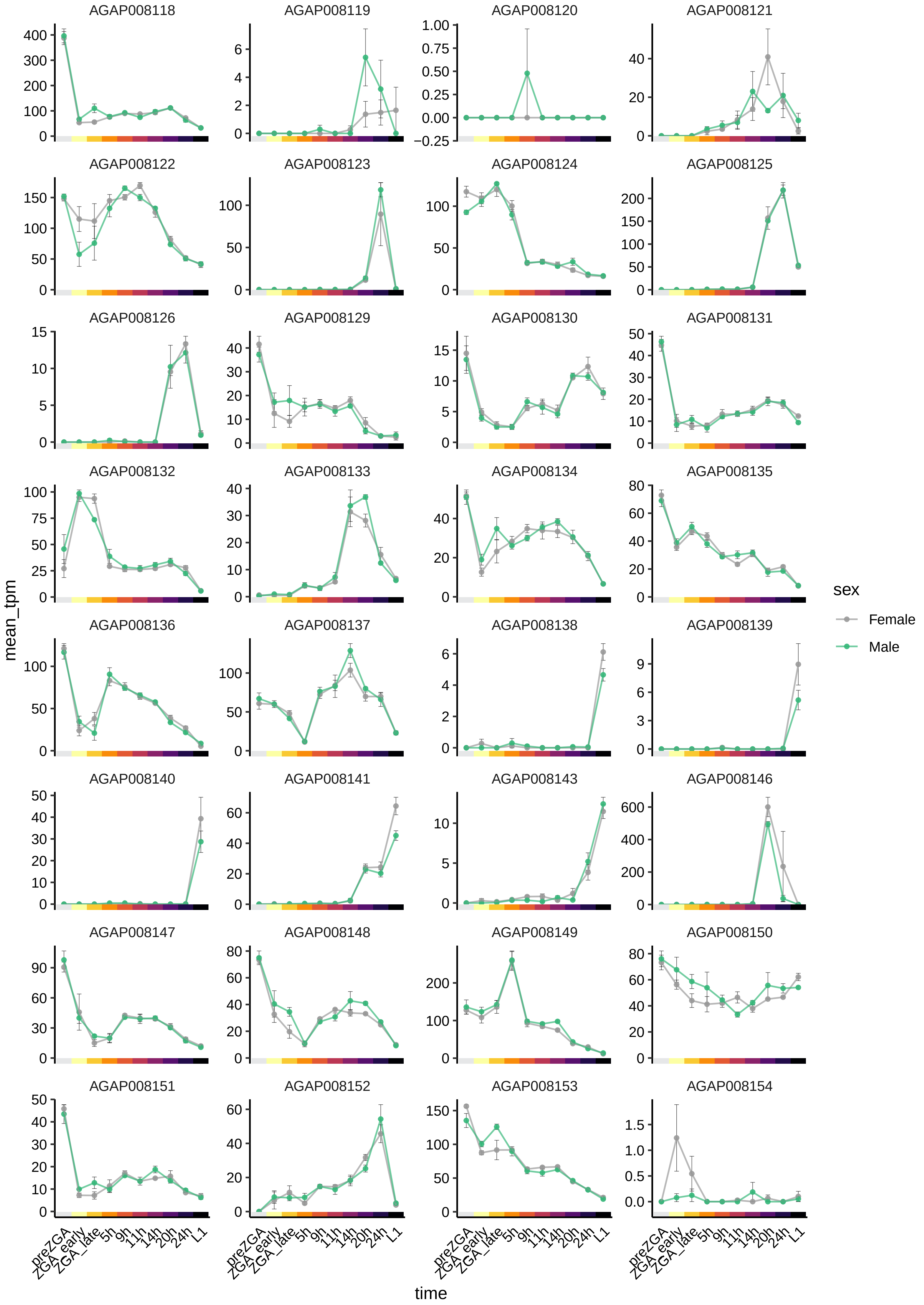
AGAP008118
AGAP008119
AGAP008120
AGAP008121
AGAP008122
AGAP008123
AGAP008124
AGAP008125
AGAP008126
AGAP008129
AGAP008130
AGAP008131
AGAP008132
AGAP008133
AGAP008134
AGAP008135
AGAP008136
AGAP008137
AGAP008138
AGAP008139
AGAP008140
AGAP008141
AGAP008143
AGAP008146
AGAP008147
AGAP008148
AGAP008149
AGAP008150
AGAP008151
AGAP008152
AGAP008153
AGAP008154
mean_tpm
time
preZGA
ZGA_early
ZGA_late
5h
9h
11h
14h
20h
24h
L1
sex
Female
Male

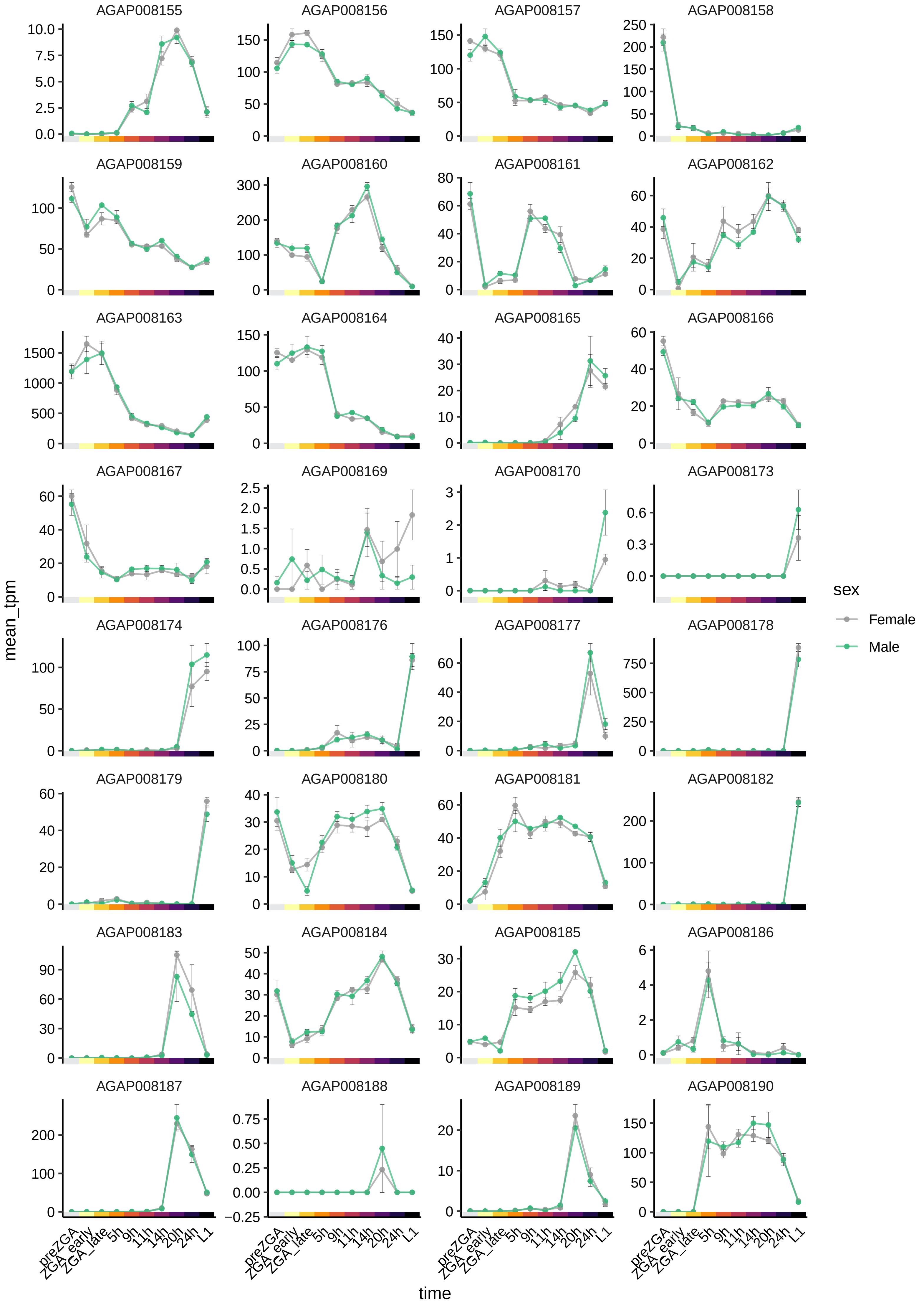

AGAP008155
AGAP008156
AGAP008157
AGAP008158
AGAP008159
AGAP008160
AGAP008161
AGAP008162
AGAP008163
AGAP008164
AGAP008165
AGAP008166
AGAP008167
AGAP008169
AGAP008170
AGAP008173
AGAP008174
AGAP008176
AGAP008177
AGAP008178
AGAP008179
AGAP008180
AGAP008181
AGAP008182
AGAP008183
AGAP008184
AGAP008185
AGAP008186
AGAP008187
AGAP008188
AGAP008189
AGAP008190
mean_tpm
time
preZGA
ZGA_early
ZGA_late
5h
9h
11h
14h
20h
24h
L1
sex
Female
Male

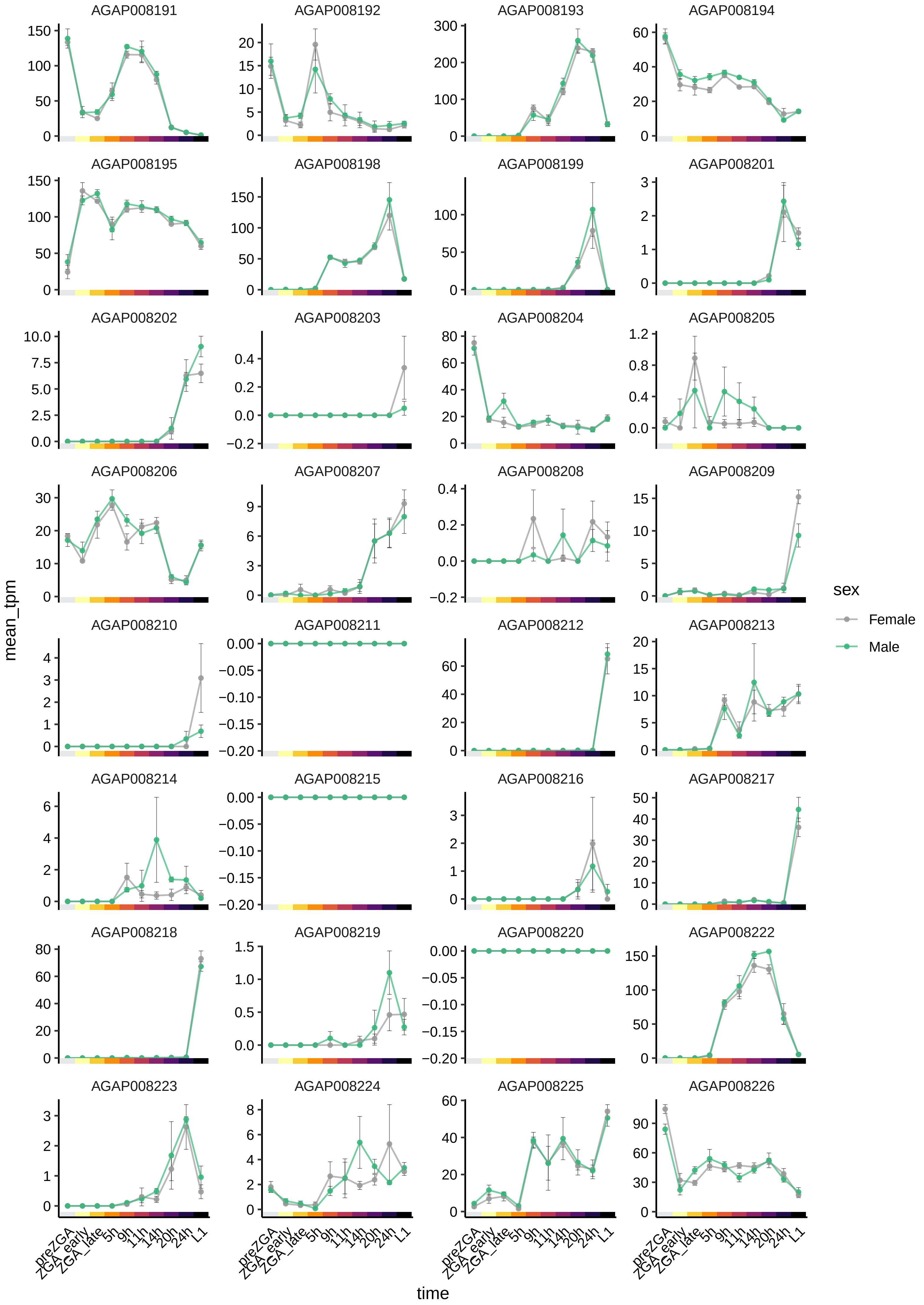

AGAP008191
AGAP008192
AGAP008193
AGAP008194
AGAP008195
AGAP008198
AGAP008199
AGAP008201
AGAP008202
AGAP008203
AGAP008204
AGAP008205
AGAP008206
AGAP008207
AGAP008208
AGAP008209
AGAP008210
AGAP008211
AGAP008212
AGAP008213
AGAP008214
AGAP008215
AGAP008216
AGAP008217
AGAP008218
AGAP008219
AGAP008220
AGAP008222
AGAP008223
AGAP008224
AGAP008225
AGAP008226
mean_tpm
time
preZGA
ZGA_early
ZGA_late
5h
9h
11h
14h
20h
24h
L1
sex
Female
Male

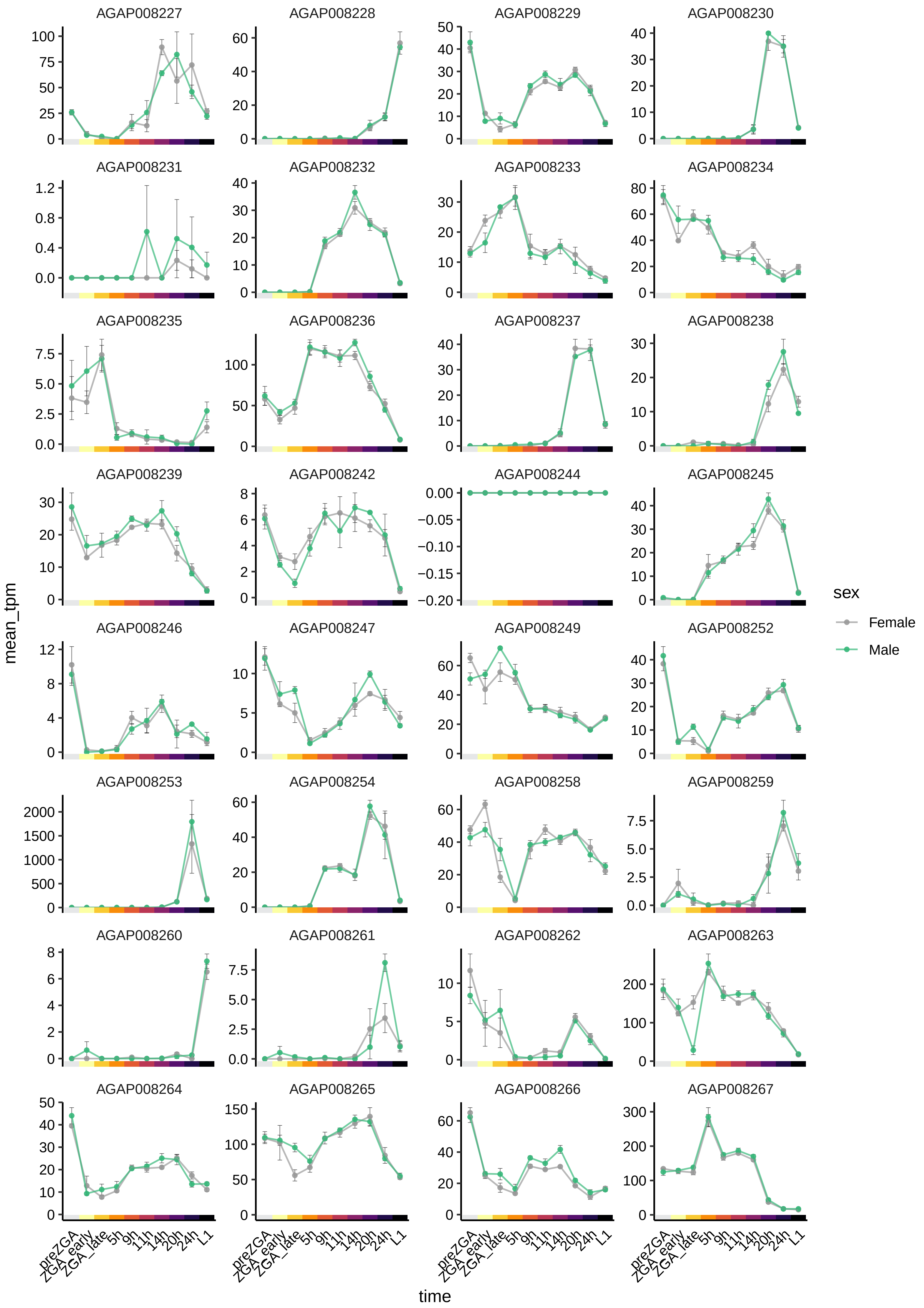
AGAP008227
AGAP008228
AGAP008229
AGAP008230
AGAP008231
AGAP008232
AGAP008233
AGAP008234
AGAP008235
AGAP008236
AGAP008237
AGAP008238
AGAP008239
AGAP008242
AGAP008244
AGAP008245
AGAP008246
AGAP008247
AGAP008249
AGAP008252
AGAP008253
AGAP008254
AGAP008258
AGAP008259
AGAP008260
AGAP008261
AGAP008262
AGAP008263
AGAP008264
AGAP008265
AGAP008266
AGAP008267
mean_tpm
time
preZGA
ZGA_early
ZGA_late
5h
9h
11h
14h
20h
24h
L1
sex
Female
Male

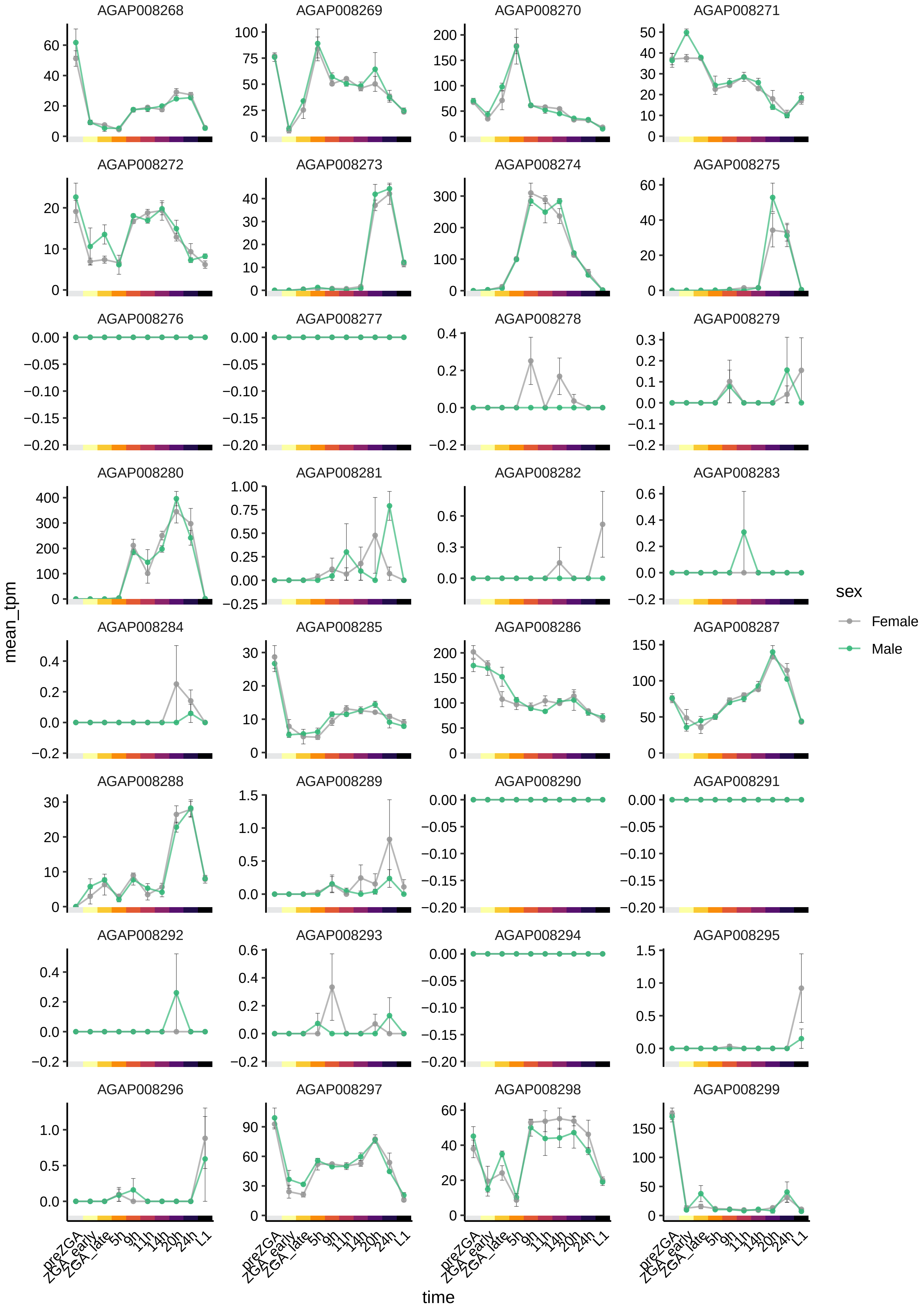

AGAP008268
AGAP008269
AGAP008270
AGAP008271
AGAP008272
AGAP008273
AGAP008274
AGAP008275
AGAP008276
AGAP008277
AGAP008278
AGAP008279
AGAP008280
AGAP008281
AGAP008282
AGAP008283
AGAP008284
AGAP008285
AGAP008286
AGAP008287
AGAP008288
AGAP008289
AGAP008290
AGAP008291
AGAP008292
AGAP008293
AGAP008294
AGAP008295
AGAP008296
AGAP008297
AGAP008298
AGAP008299
mean_tpm
time
preZGA
ZGA_early
ZGA_late
5h
9h
11h
14h
20h
24h
L1
sex
Female
Male

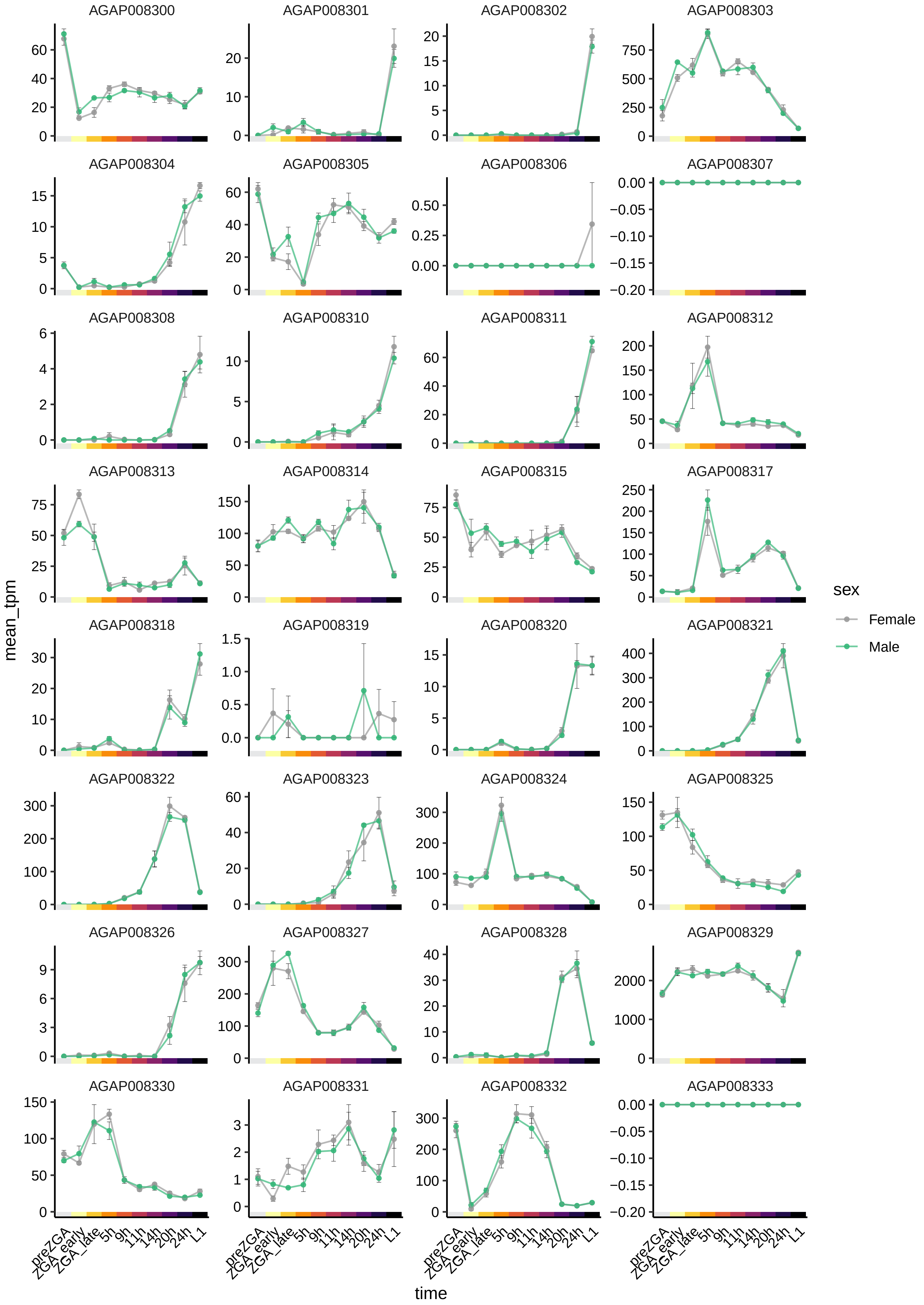
AGAP008300
AGAP008301
AGAP008302
AGAP008303
AGAP008304
AGAP008305
AGAP008306
AGAP008307
AGAP008308
AGAP008310
AGAP008311
AGAP008312
AGAP008313
AGAP008314
AGAP008315
AGAP008317
AGAP008318
AGAP008319
AGAP008320
AGAP008321
AGAP008322
AGAP008323
AGAP008324
AGAP008325
AGAP008326
AGAP008327
AGAP008328
AGAP008329
AGAP008330
AGAP008331
AGAP008332
AGAP008333
mean_tpm
time
preZGA
ZGA_early
ZGA_late
5h
9h
11h
14h
20h
24h
L1
sex
Female
Male

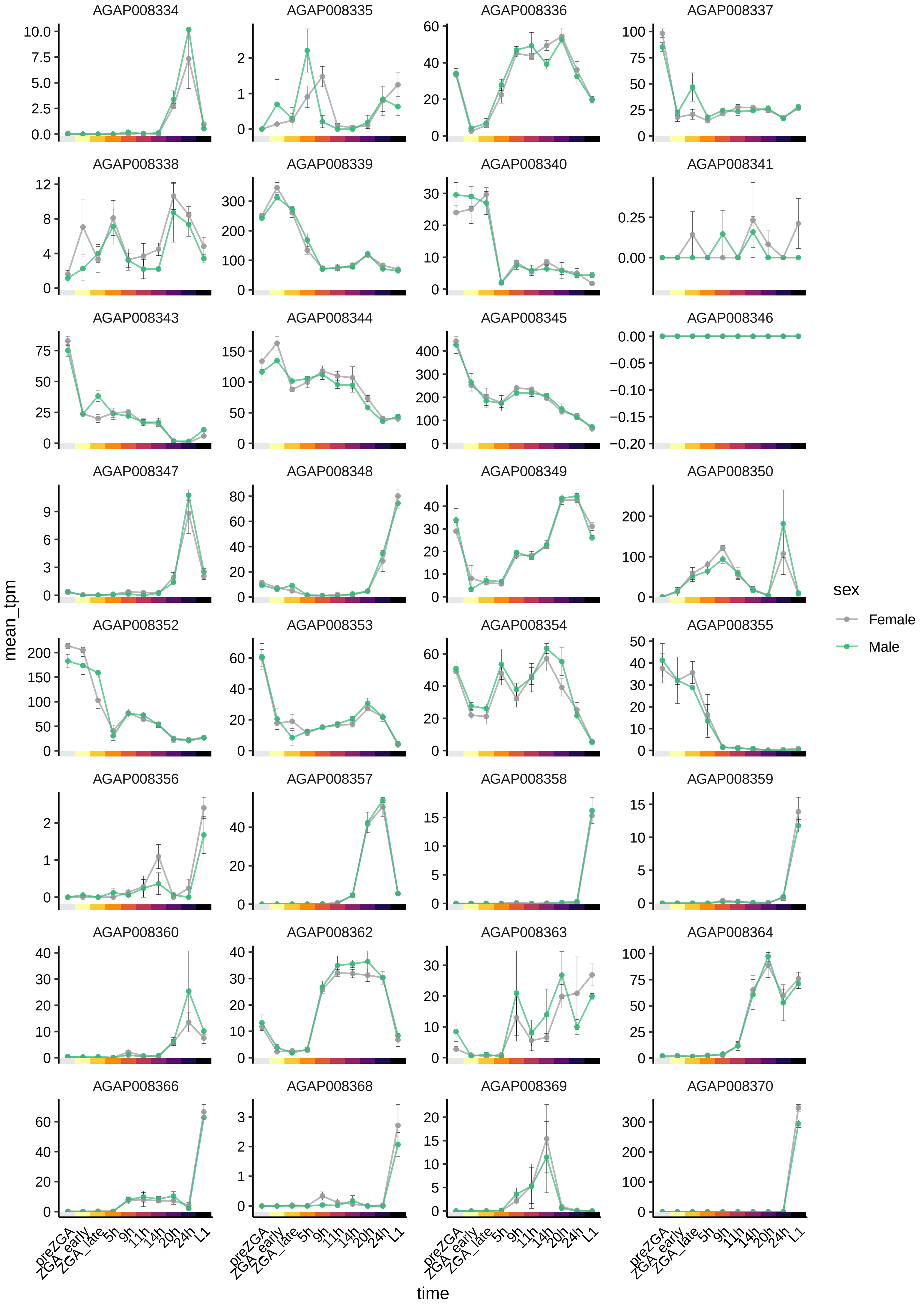
AGAP008334
AGAP008335
AGAP008336
AGAP008337
AGAP008338
AGAP008339
AGAP008340
AGAP008341
AGAP008343
AGAP008344
AGAP008345
AGAP008346
AGAP008347
AGAP008348
AGAP008349
AGAP008350
AGAP008352
AGAP008353
AGAP008354
AGAP008355
AGAP008356
AGAP008357
AGAP008358
AGAP008359
AGAP008360
AGAP008362
AGAP008363
AGAP008364
AGAP008366
AGAP008368
AGAP008369
AGAP008370
mean_tpm
time
preZGA
ZGA_early
ZGA_late
5h
9h
11h
14h
20h
24h
L1
sex
Female
Male

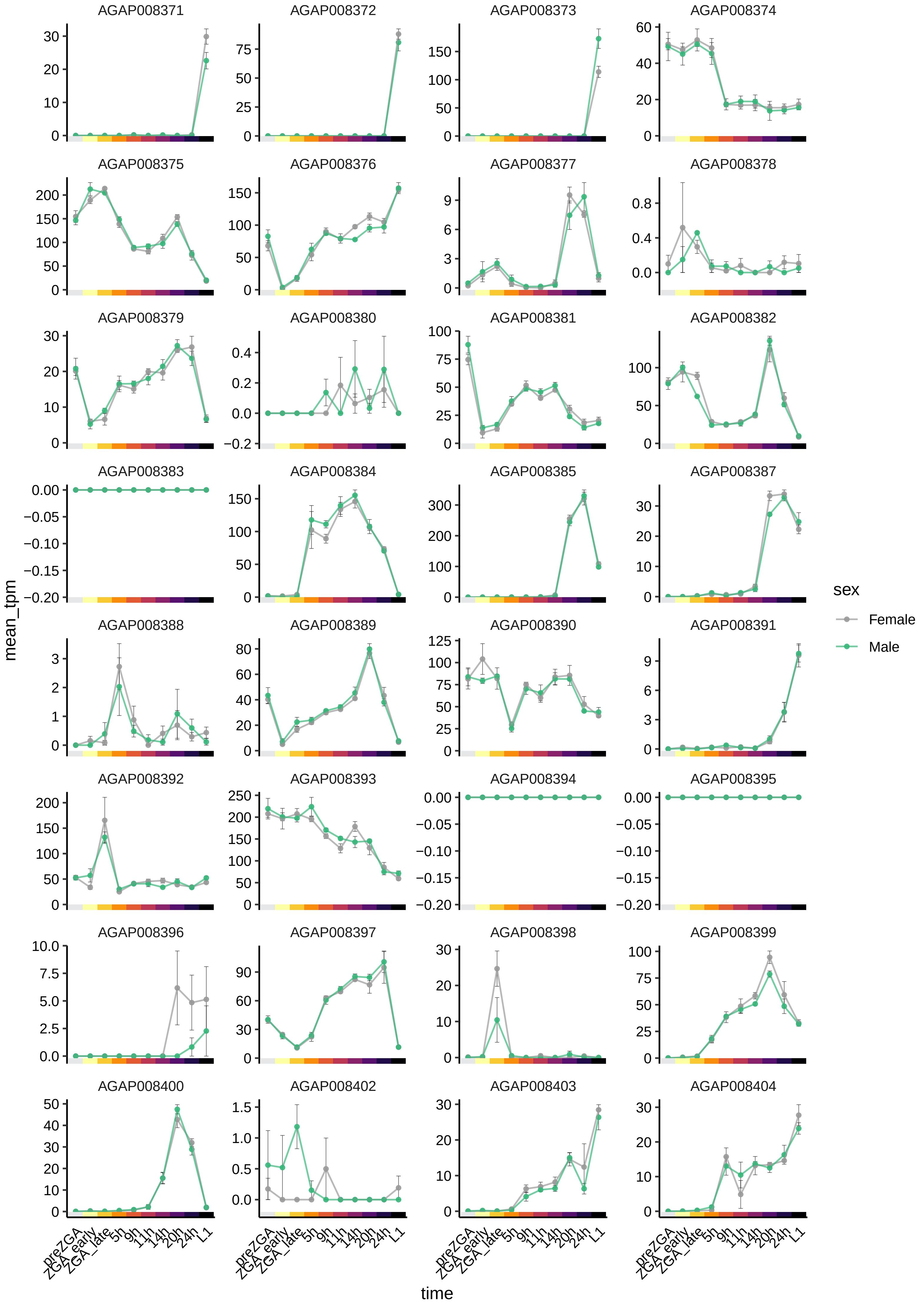

AGAP008371
AGAP008372
AGAP008373
AGAP008374
AGAP008375
AGAP008376
AGAP008377
AGAP008378
AGAP008379
AGAP008380
AGAP008381
AGAP008382
AGAP008383
AGAP008384
AGAP008385
AGAP008387
AGAP008388
AGAP008389
AGAP008390
AGAP008391
AGAP008392
AGAP008393
AGAP008394
AGAP008395
AGAP008396
AGAP008397
AGAP008398
AGAP008399
AGAP008400
AGAP008402
AGAP008403
AGAP008404
mean_tpm
time
preZGA
ZGA_early
ZGA_late
5h
9h
11h
14h
20h
24h
L1
sex
Female
Male

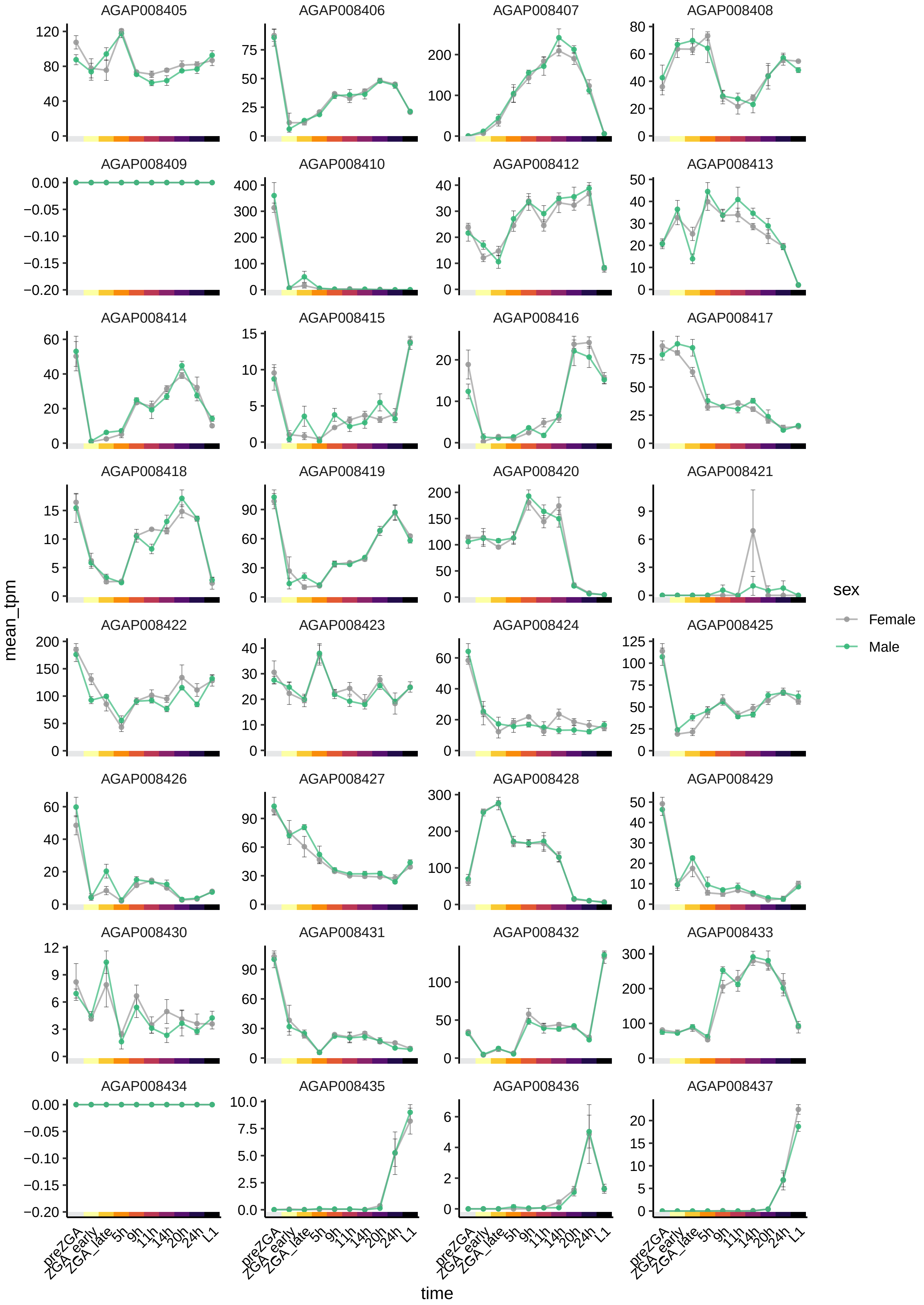
AGAP008405
AGAP008406
AGAP008407
AGAP008408
AGAP008409
AGAP008410
AGAP008412
AGAP008413
AGAP008414
AGAP008415
AGAP008416
AGAP008417
AGAP008418
AGAP008419
AGAP008420
AGAP008421
AGAP008422
AGAP008423
AGAP008424
AGAP008425
AGAP008426
AGAP008427
AGAP008428
AGAP008429
AGAP008430
AGAP008431
AGAP008432
AGAP008433
AGAP008434
AGAP008435
AGAP008436
AGAP008437
mean_tpm
time
preZGA
ZGA_early
ZGA_late
5h
9h
11h
14h
20h
24h
L1
sex
Female
Male

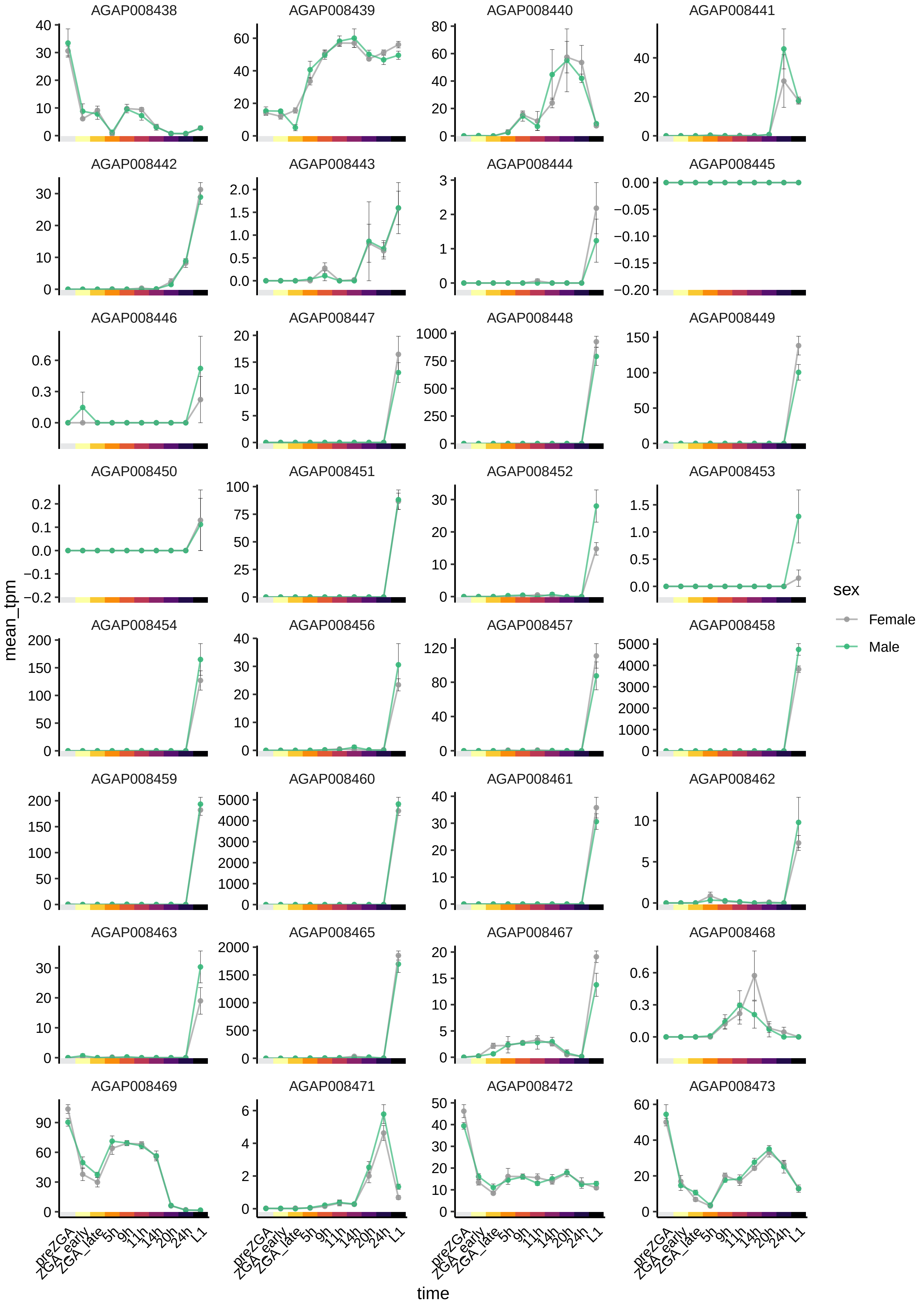

AGAP008438
AGAP008439
AGAP008440
AGAP008441
AGAP008442
AGAP008443
AGAP008444
AGAP008445
AGAP008446
AGAP008447
AGAP008448
AGAP008449
AGAP008450
AGAP008451
AGAP008452
AGAP008453
AGAP008454
AGAP008456
AGAP008457
AGAP008458
AGAP008459
AGAP008460
AGAP008461
AGAP008462
AGAP008463
AGAP008465
AGAP008467
AGAP008468
AGAP008469
AGAP008471
AGAP008472
AGAP008473
mean_tpm
time
preZGA
ZGA_early
ZGA_late
5h
9h
11h
14h
20h
24h
L1
sex
Female
Male

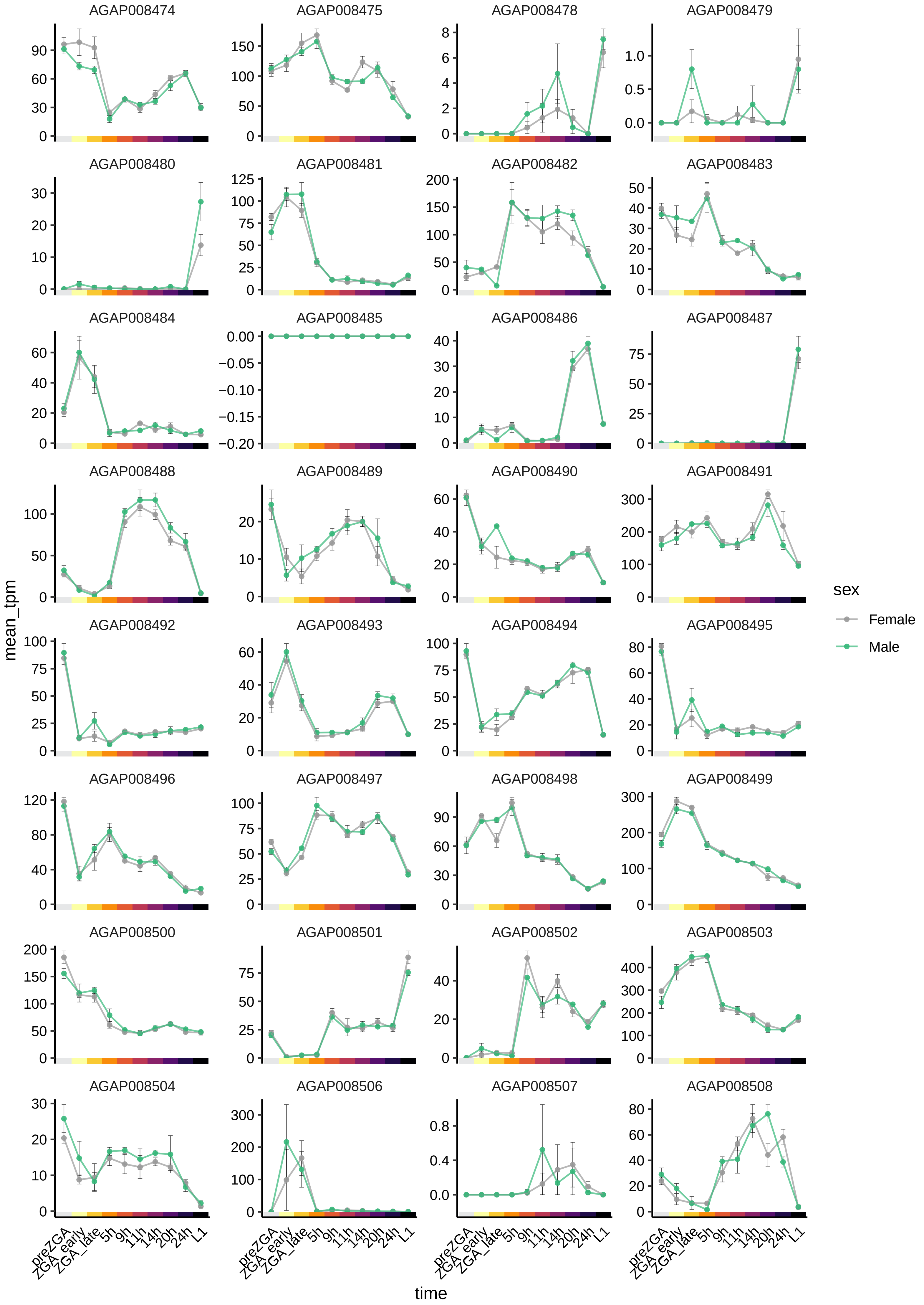
AGAP008474
AGAP008475
AGAP008478
AGAP008479
AGAP008480
AGAP008481
AGAP008482
AGAP008483
AGAP008484
AGAP008485
AGAP008486
AGAP008487
AGAP008488
AGAP008489
AGAP008490
AGAP008491
AGAP008492
AGAP008493
AGAP008494
AGAP008495
AGAP008496
AGAP008497
AGAP008498
AGAP008499
AGAP008500
AGAP008501
AGAP008502
AGAP008503
AGAP008504
AGAP008506
AGAP008507
AGAP008508
mean_tpm
time
preZGA
ZGA_early
ZGA_late
5h
9h
11h
14h
20h
24h
L1
sex
Female
Male

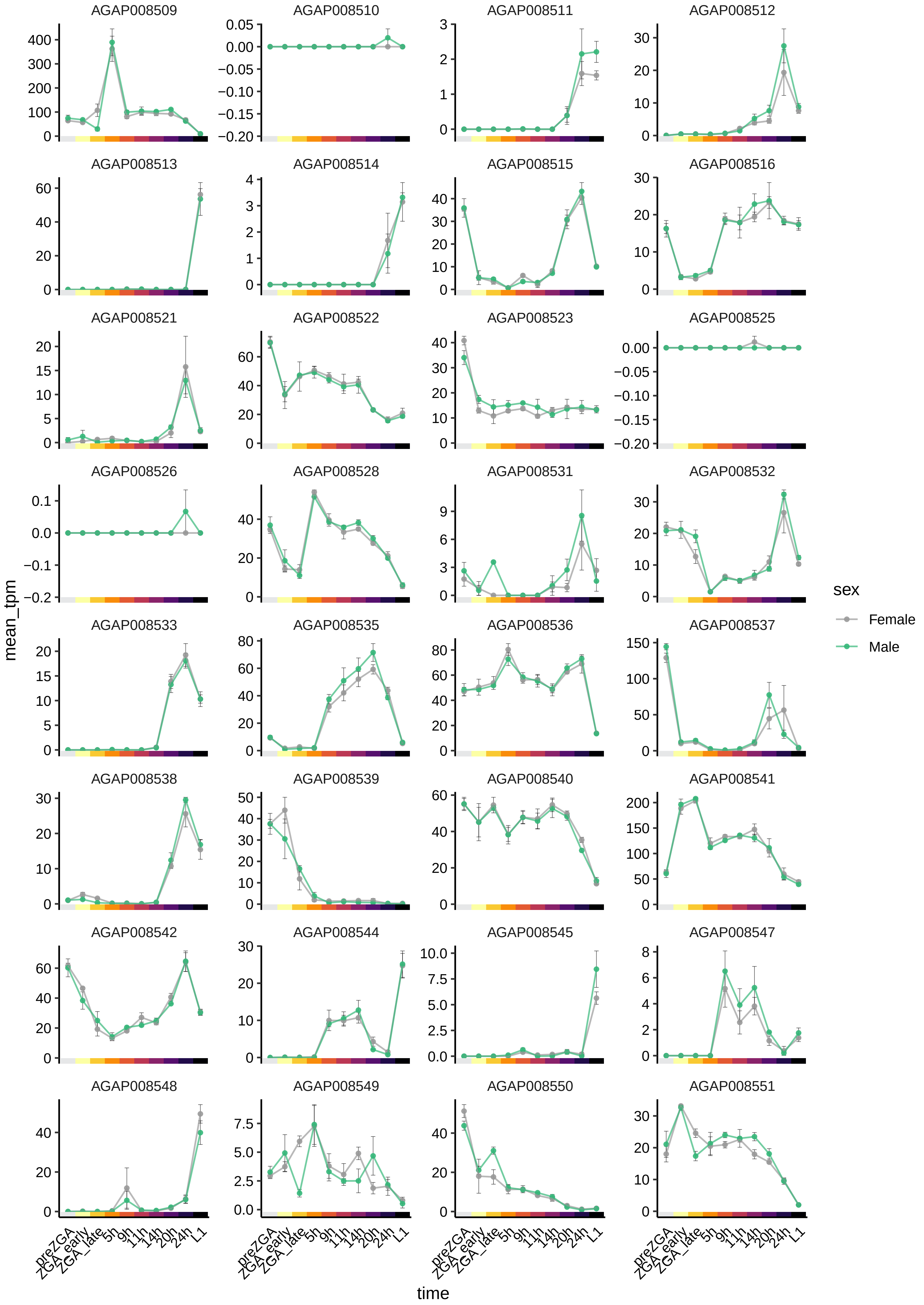
AGAP008509
AGAP008510
AGAP008511
AGAP008512
AGAP008513
AGAP008514
AGAP008515
AGAP008516
AGAP008521
AGAP008522
AGAP008523
AGAP008525
AGAP008526
AGAP008528
AGAP008531
AGAP008532
AGAP008533
AGAP008535
AGAP008536
AGAP008537
AGAP008538
AGAP008539
AGAP008540
AGAP008541
AGAP008542
AGAP008544
AGAP008545
AGAP008547
AGAP008548
AGAP008549
AGAP008550
AGAP008551
mean_tpm
time
preZGA
ZGA_early
ZGA_late
5h
9h
11h
14h
20h
24h
L1
sex
Female
Male

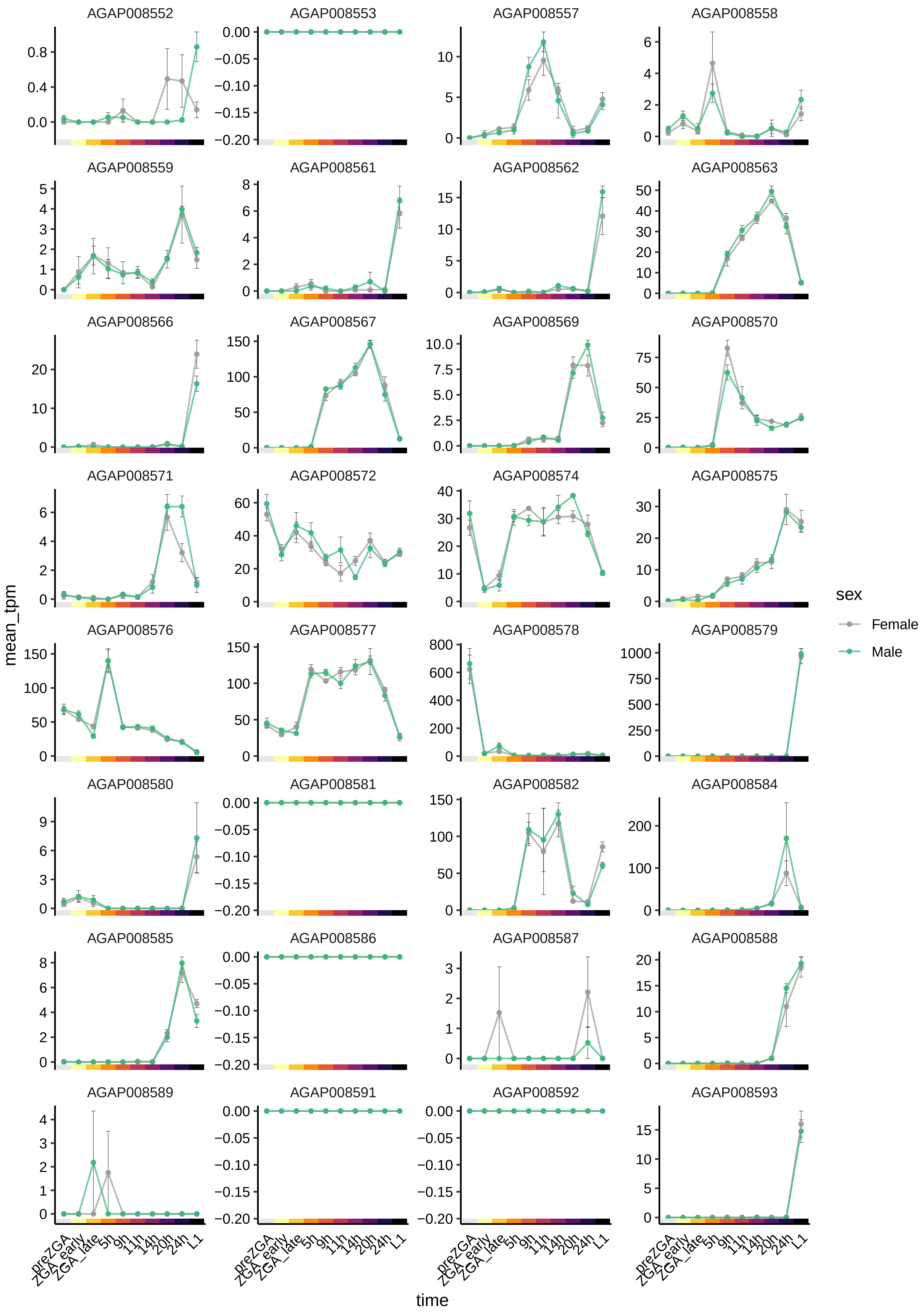

AGAP008552
AGAP008553
AGAP008557
AGAP008558
AGAP008559
AGAP008561
AGAP008562
AGAP008563
AGAP008566
AGAP008567
AGAP008569
AGAP008570
AGAP008571
AGAP008572
AGAP008574
AGAP008575
AGAP008576
AGAP008577
AGAP008578
AGAP008579
AGAP008580
AGAP008581
AGAP008582
AGAP008584
AGAP008585
AGAP008586
AGAP008587
AGAP008588
AGAP008589
AGAP008591
AGAP008592
AGAP008593
mean_tpm
time
preZGA
ZGA_early
ZGA_late
5h
9h
11h
14h
20h
24h
L1
sex
Female
Male

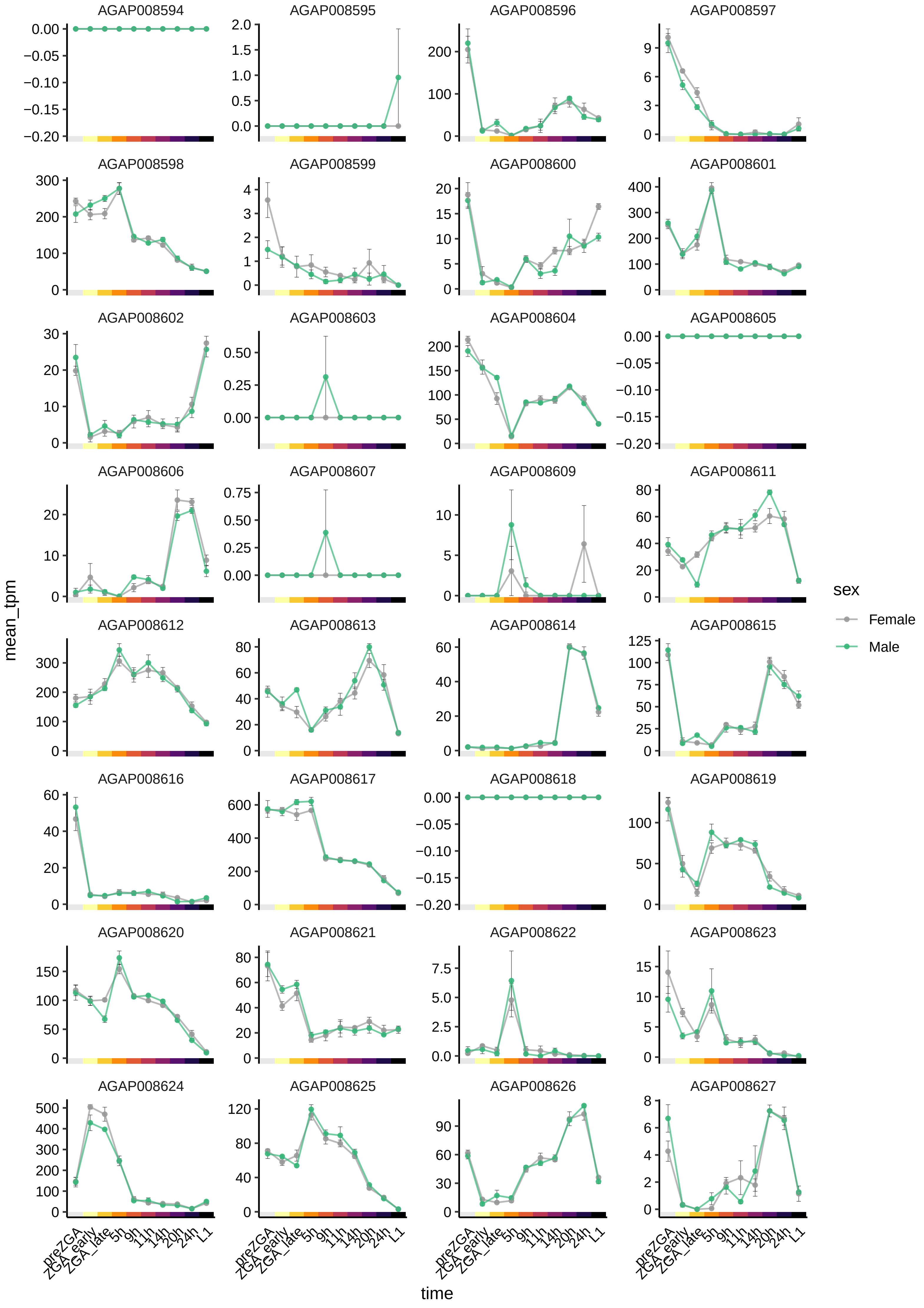
AGAP008594
AGAP008595
AGAP008596
AGAP008597
AGAP008598
AGAP008599
AGAP008600
AGAP008601
AGAP008602
AGAP008603
AGAP008604
AGAP008605
AGAP008606
AGAP008607
AGAP008609
AGAP008611
AGAP008612
AGAP008613
AGAP008614
AGAP008615
AGAP008616
AGAP008617
AGAP008618
AGAP008619
AGAP008620
AGAP008621
AGAP008622
AGAP008623
AGAP008624
AGAP008625
AGAP008626
AGAP008627
mean_tpm
time
preZGA
ZGA_early
ZGA_late
5h
9h
11h
14h
20h
24h
L1
sex
Female
Male

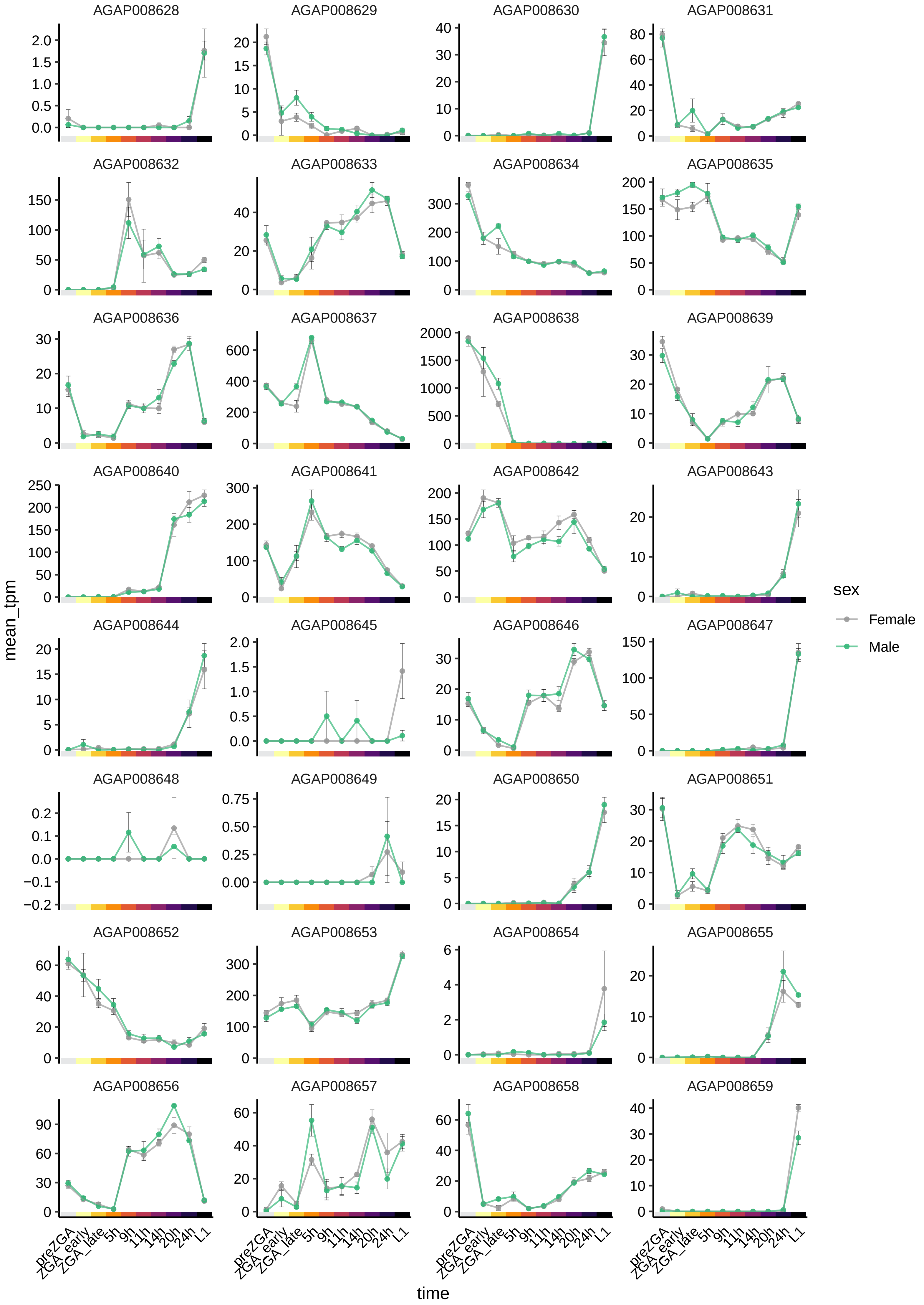

AGAP008628
AGAP008629
AGAP008630
AGAP008631
AGAP008632
AGAP008633
AGAP008634
AGAP008635
AGAP008636
AGAP008637
AGAP008638
AGAP008639
AGAP008640
AGAP008641
AGAP008642
AGAP008643
AGAP008644
AGAP008645
AGAP008646
AGAP008647
AGAP008648
AGAP008649
AGAP008650
AGAP008651
AGAP008652
AGAP008653
AGAP008654
AGAP008655
AGAP008656
AGAP008657
AGAP008658
AGAP008659
mean_tpm
time
preZGA
ZGA_early
ZGA_late
5h
9h
11h
14h
20h
24h
L1
sex
Female
Male

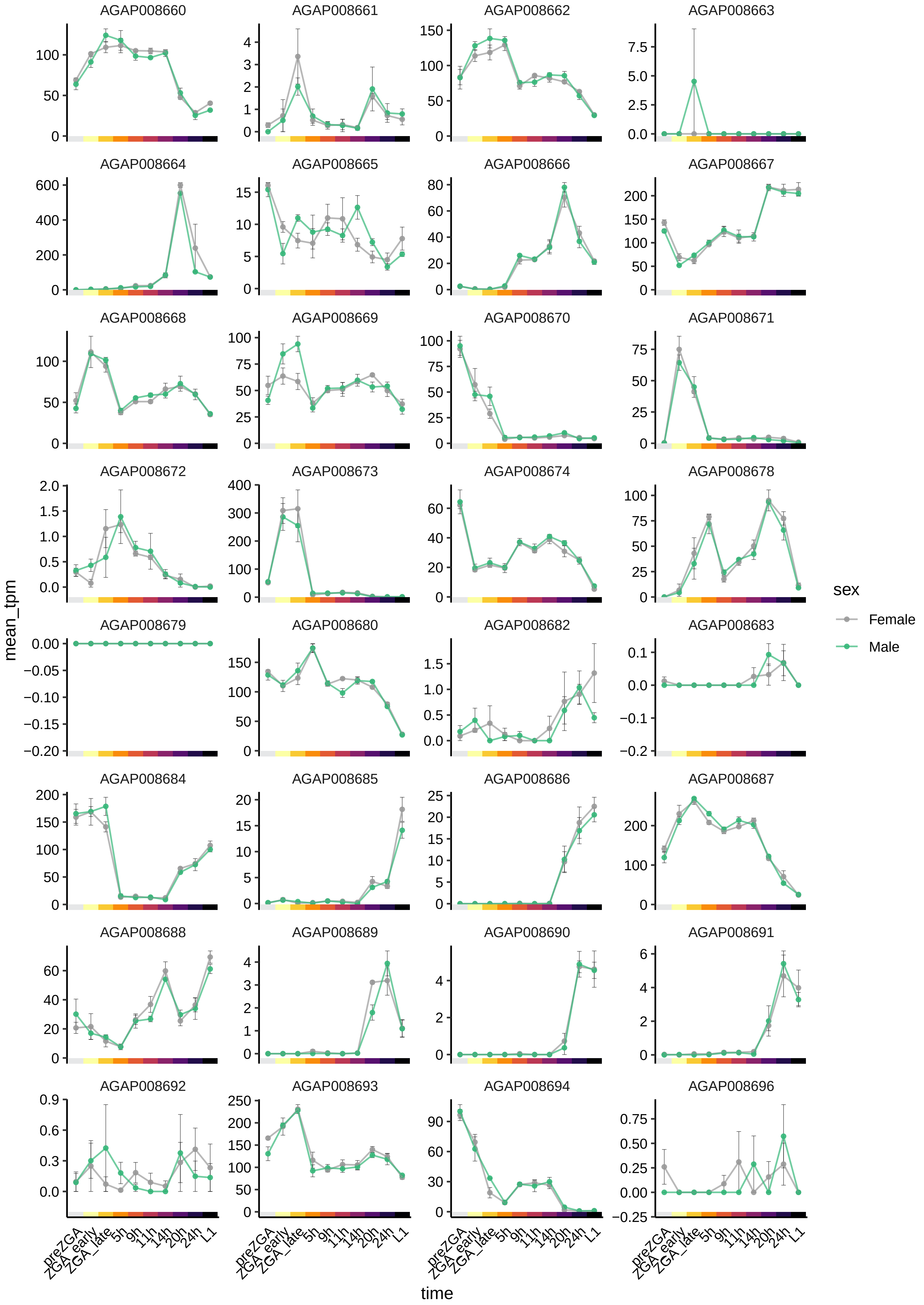
AGAP008660
AGAP008661
AGAP008662
AGAP008663
AGAP008664
AGAP008665
AGAP008666
AGAP008667
AGAP008668
AGAP008669
AGAP008670
AGAP008671
AGAP008672
AGAP008673
AGAP008674
AGAP008678
AGAP008679
AGAP008680
AGAP008682
AGAP008683
AGAP008684
AGAP008685
AGAP008686
AGAP008687
AGAP008688
AGAP008689
AGAP008690
AGAP008691
AGAP008692
AGAP008693
AGAP008694
AGAP008696
mean_tpm
time
preZGA
ZGA_early
ZGA_late
5h
9h
11h
14h
20h
24h
L1
sex
Female
Male

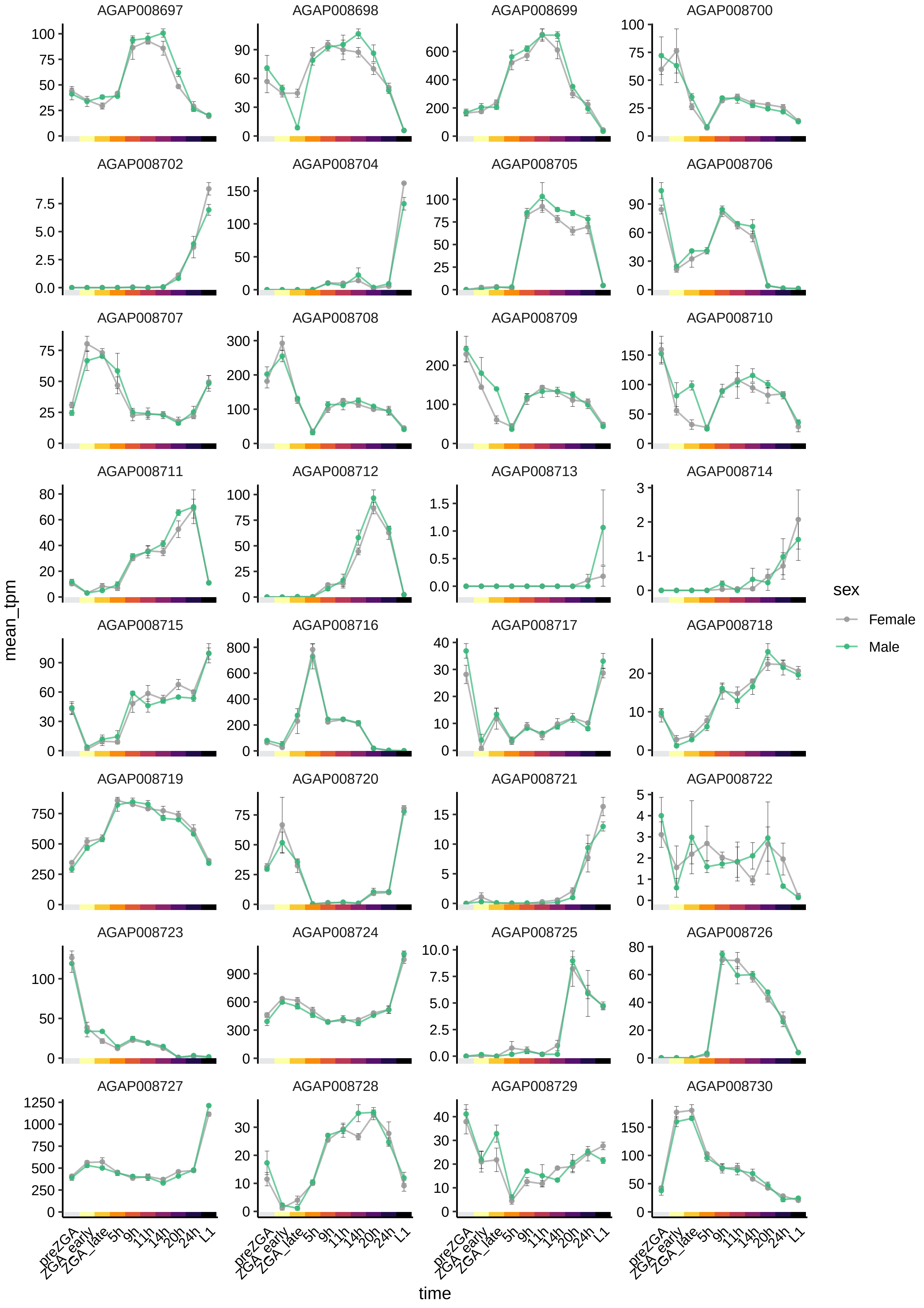
AGAP008697
AGAP008698
AGAP008699
AGAP008700
AGAP008702
AGAP008704
AGAP008705
AGAP008706
AGAP008707
AGAP008708
AGAP008709
AGAP008710
AGAP008711
AGAP008712
AGAP008713
AGAP008714
AGAP008715
AGAP008716
AGAP008717
AGAP008718
AGAP008719
AGAP008720
AGAP008721
AGAP008722
AGAP008723
AGAP008724
AGAP008725
AGAP008726
AGAP008727
AGAP008728
AGAP008729
AGAP008730
mean_tpm
time
preZGA
ZGA_early
ZGA_late
5h
9h
11h
14h
20h
24h
L1
sex
Female
Male

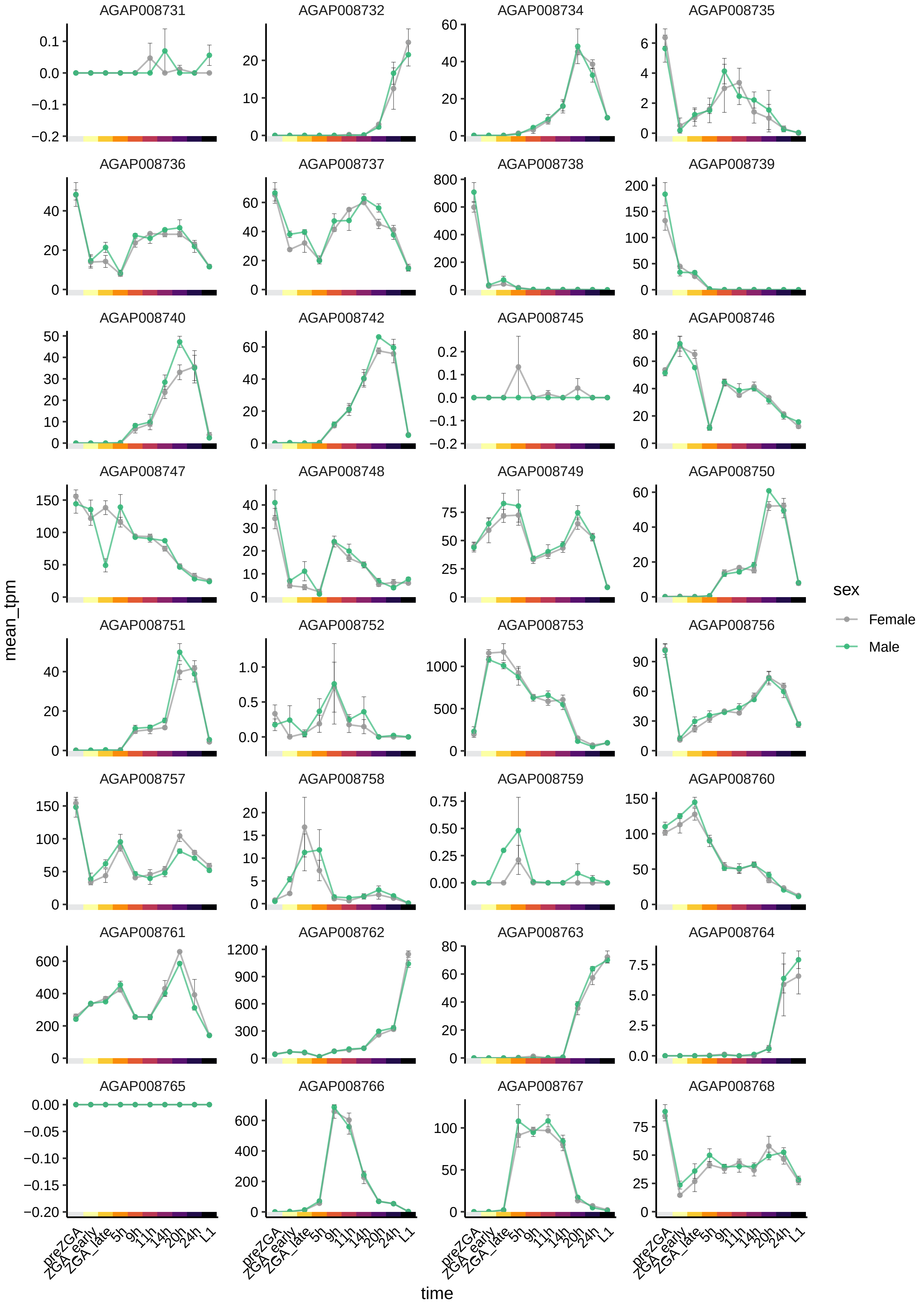

AGAP008731
AGAP008732
AGAP008734
AGAP008735
AGAP008736
AGAP008737
AGAP008738
AGAP008739
AGAP008740
AGAP008742
AGAP008745
AGAP008746
AGAP008747
AGAP008748
AGAP008749
AGAP008750
AGAP008751
AGAP008752
AGAP008753
AGAP008756
AGAP008757
AGAP008758
AGAP008759
AGAP008760
AGAP008761
AGAP008762
AGAP008763
AGAP008764
AGAP008765
AGAP008766
AGAP008767
AGAP008768
mean_tpm
time
preZGA
ZGA_early
ZGA_late
5h
9h
11h
14h
20h
24h
L1
sex
Female
Male

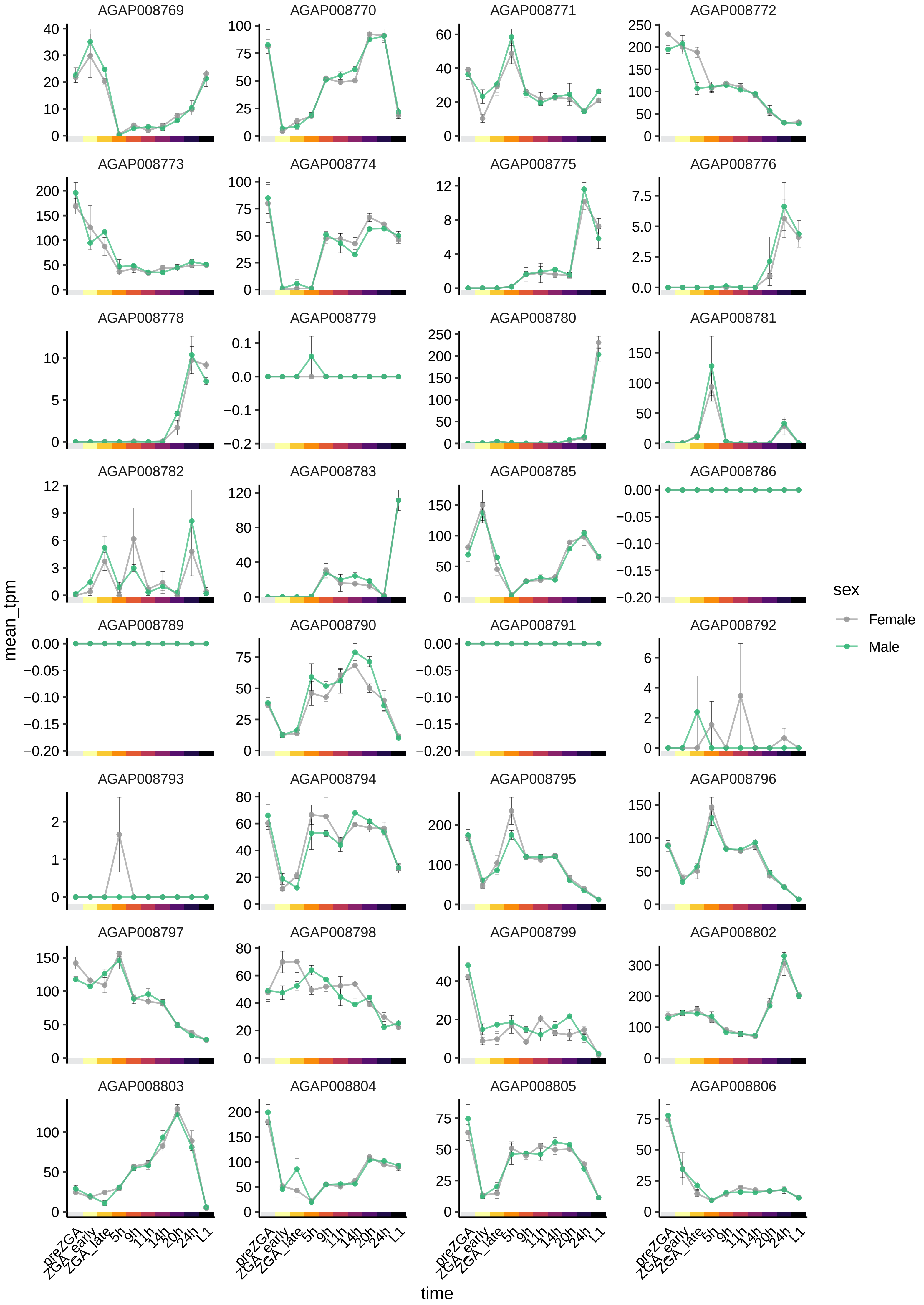

AGAP008769
AGAP008770
AGAP008771
AGAP008772
AGAP008773
AGAP008774
AGAP008775
AGAP008776
AGAP008778
AGAP008779
AGAP008780
AGAP008781
AGAP008782
AGAP008783
AGAP008785
AGAP008786
AGAP008789
AGAP008790
AGAP008791
AGAP008792
AGAP008793
AGAP008794
AGAP008795
AGAP008796
AGAP008797
AGAP008798
AGAP008799
AGAP008802
AGAP008803
AGAP008804
AGAP008805
AGAP008806
mean_tpm
time
preZGA
ZGA_early
ZGA_late
5h
9h
11h
14h
20h
24h
L1
sex
Female
Male

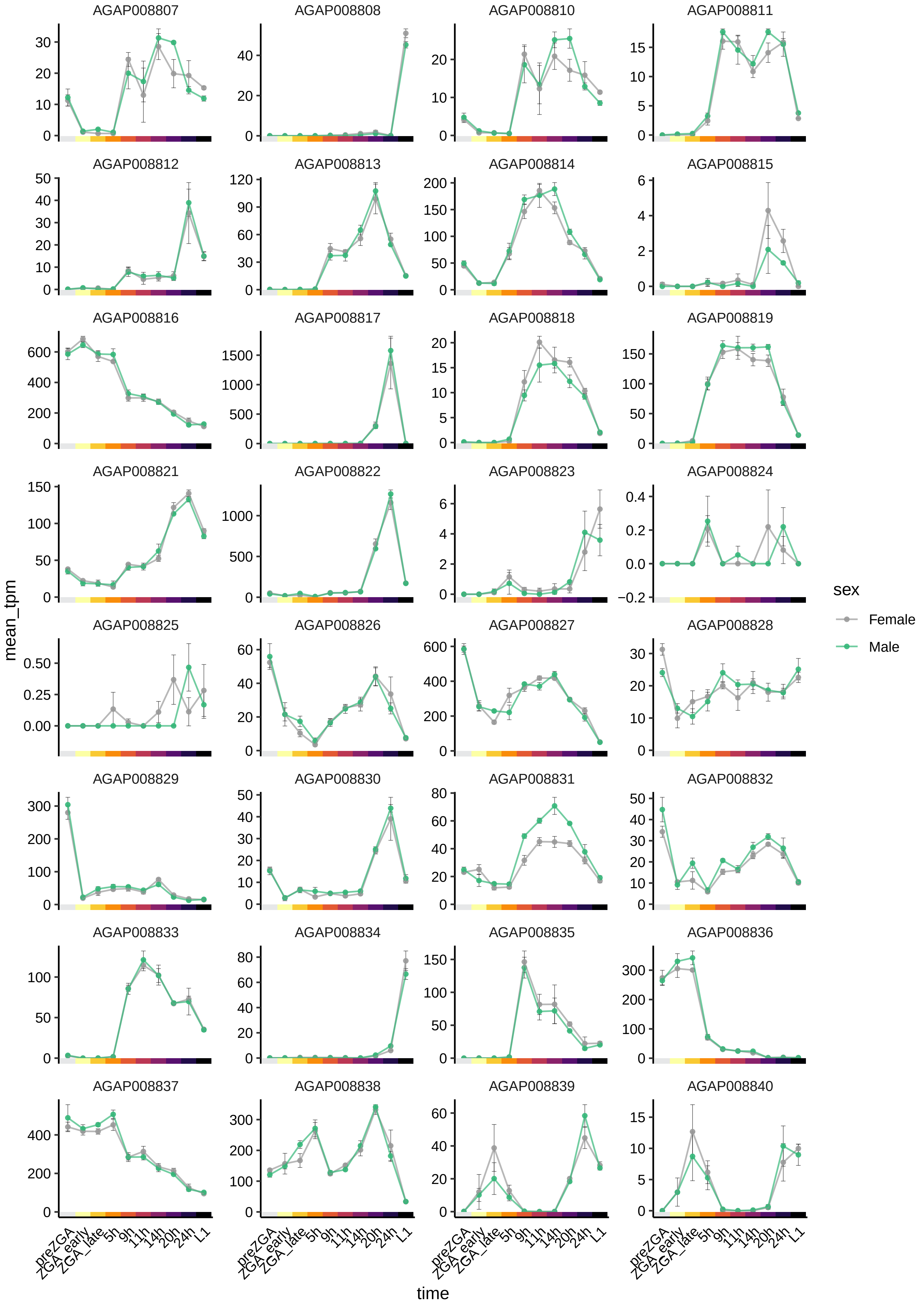

AGAP008807
AGAP008808
AGAP008810
AGAP008811
AGAP008812
AGAP008813
AGAP008814
AGAP008815
AGAP008816
AGAP008817
AGAP008818
AGAP008819
AGAP008821
AGAP008822
AGAP008823
AGAP008824
AGAP008825
AGAP008826
AGAP008827
AGAP008828
AGAP008829
AGAP008830
AGAP008831
AGAP008832
AGAP008833
AGAP008834
AGAP008835
AGAP008836
AGAP008837
AGAP008838
AGAP008839
AGAP008840
mean_tpm
time
preZGA
ZGA_early
ZGA_late
5h
9h
11h
14h
20h
24h
L1
sex
Female
Male

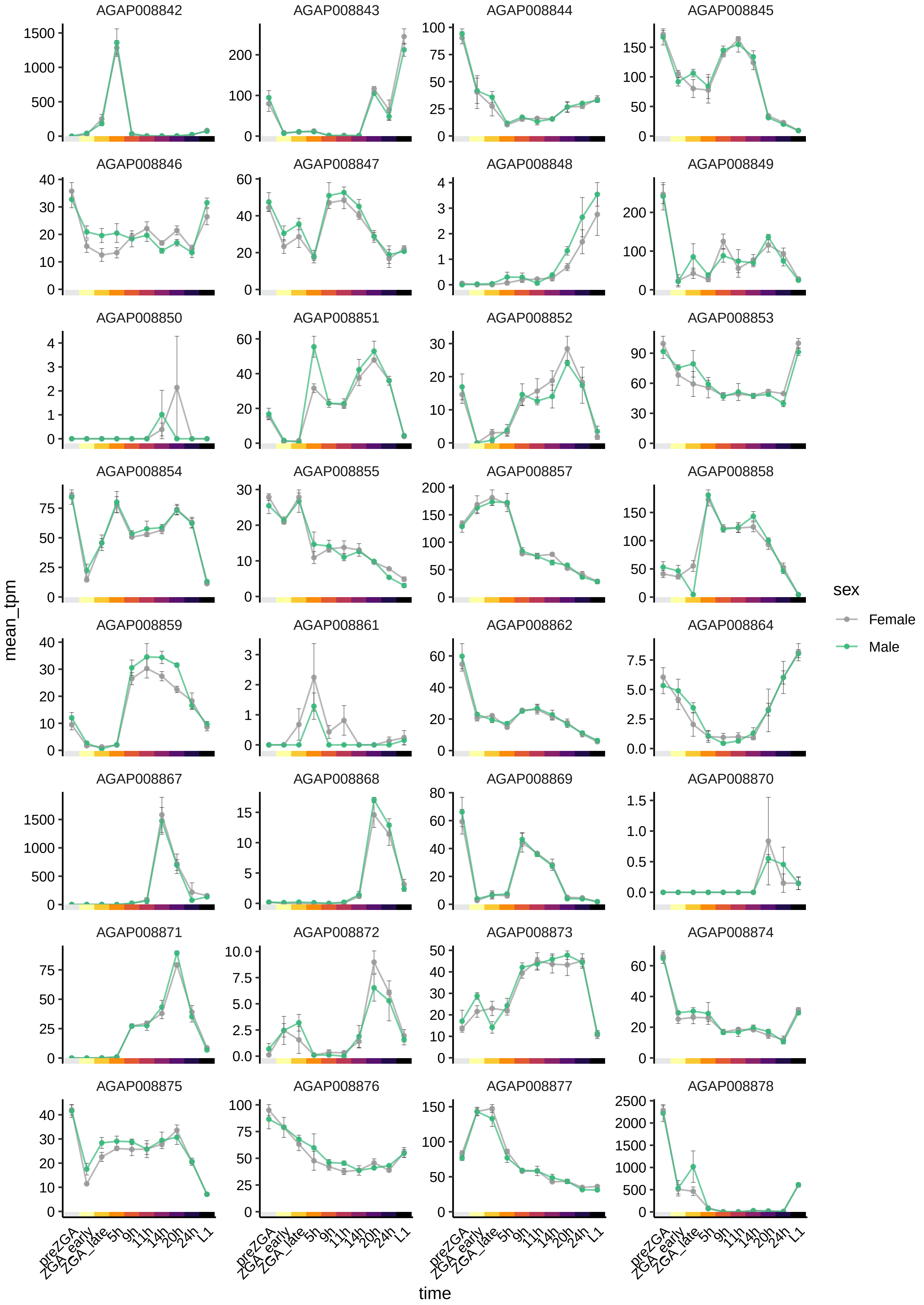
AGAP008842
AGAP008843
AGAP008844
AGAP008845
AGAP008846
AGAP008847
AGAP008848
AGAP008849
AGAP008850
AGAP008851
AGAP008852
AGAP008853
AGAP008854
AGAP008855
AGAP008857
AGAP008858
AGAP008859
AGAP008861
AGAP008862
AGAP008864
AGAP008867
AGAP008868
AGAP008869
AGAP008870
AGAP008871
AGAP008872
AGAP008873
AGAP008874
AGAP008875
AGAP008876
AGAP008877
AGAP008878
mean_tpm
time
preZGA
ZGA_early
ZGA_late
5h
9h
11h
14h
20h
24h
L1
sex
Female
Male

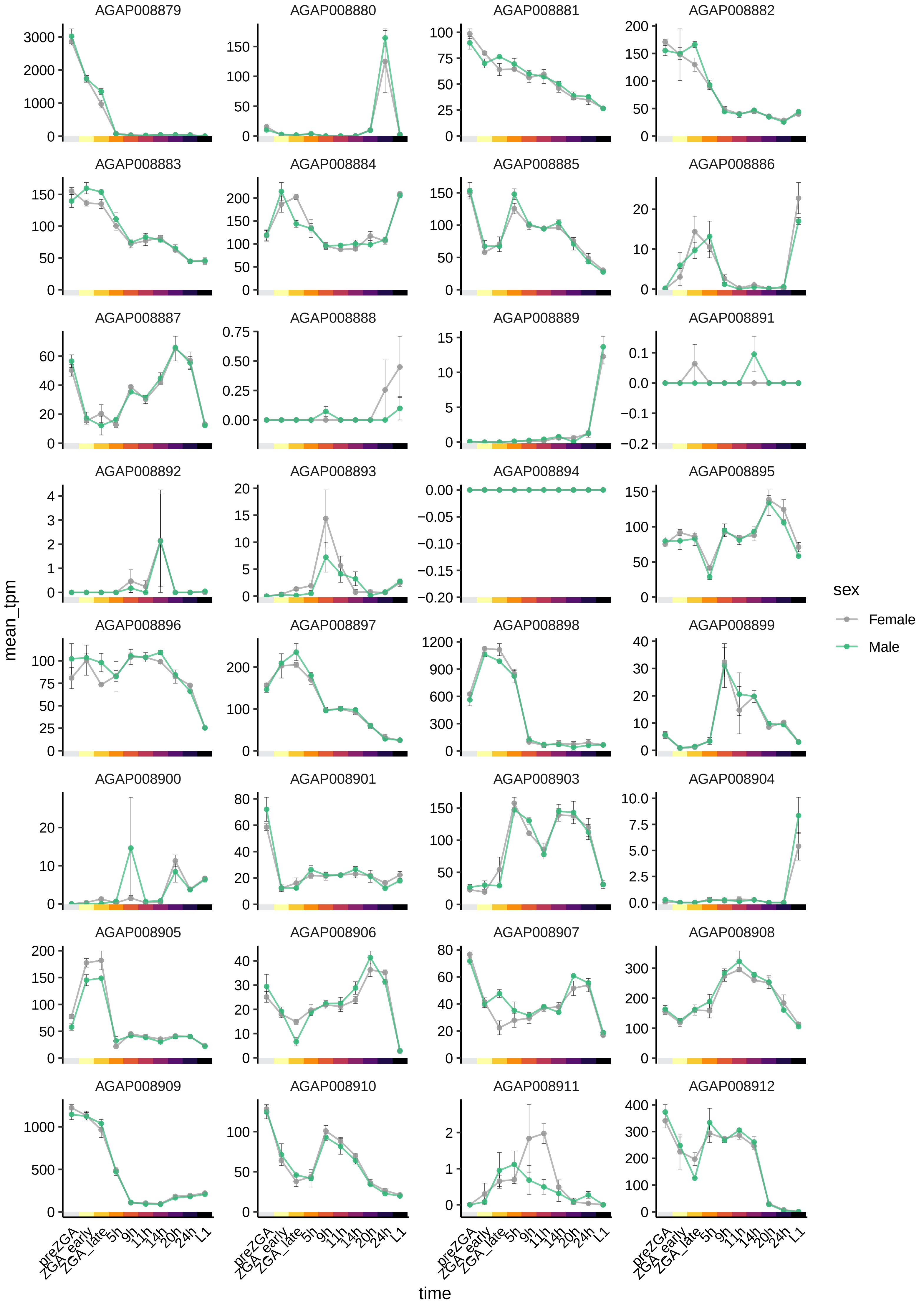

AGAP008879
AGAP008880
AGAP008881
AGAP008882
AGAP008883
AGAP008884
AGAP008885
AGAP008886
AGAP008887
AGAP008888
AGAP008889
AGAP008891
AGAP008892
AGAP008893
AGAP008894
AGAP008895
AGAP008896
AGAP008897
AGAP008898
AGAP008899
AGAP008900
AGAP008901
AGAP008903
AGAP008904
AGAP008905
AGAP008906
AGAP008907
AGAP008908
AGAP008909
AGAP008910
AGAP008911
AGAP008912
mean_tpm
time
preZGA
ZGA_early
ZGA_late
5h
9h
11h
14h
20h
24h
L1
sex
Female
Male

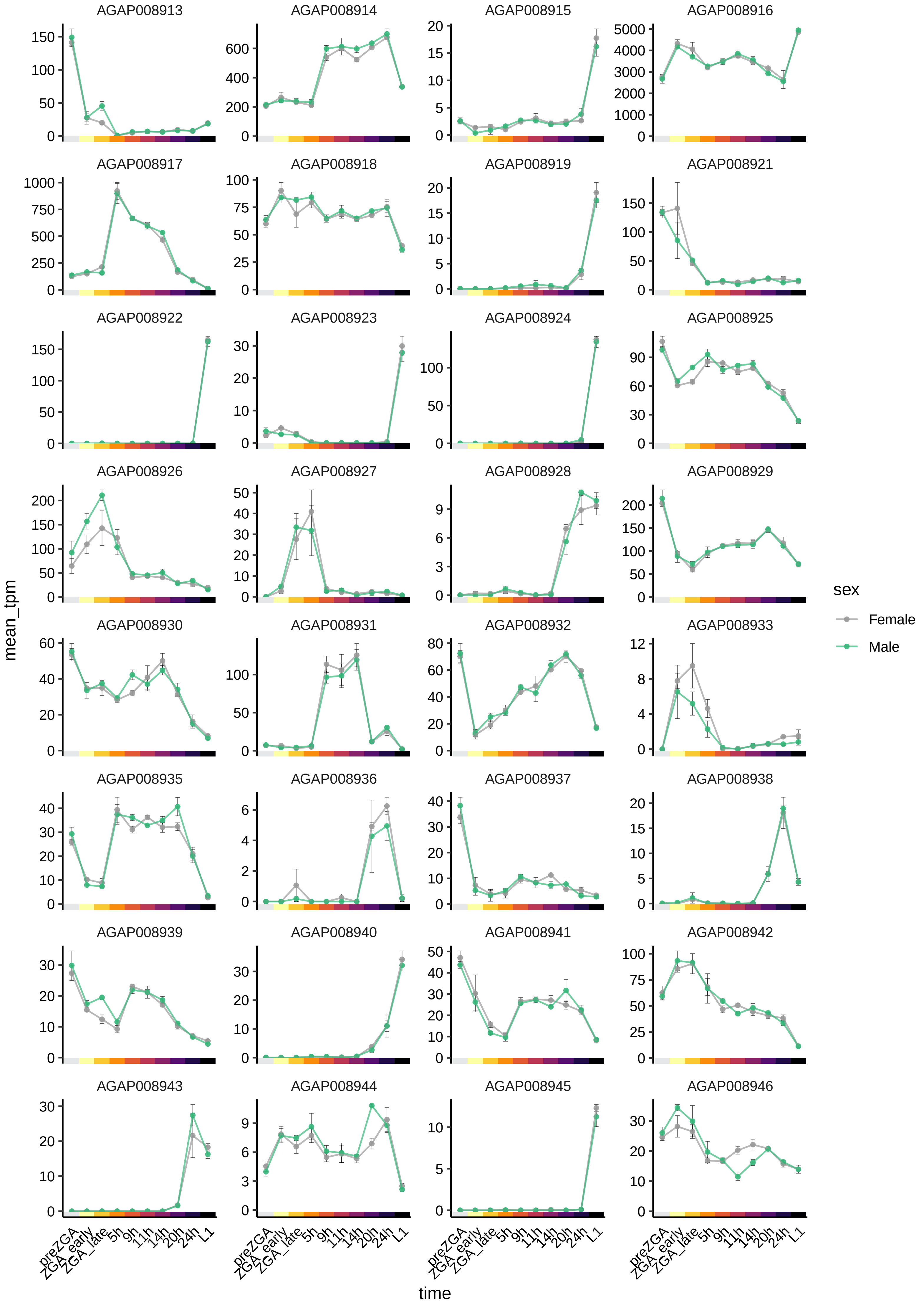

AGAP008913
AGAP008914
AGAP008915
AGAP008916
AGAP008917
AGAP008918
AGAP008919
AGAP008921
AGAP008922
AGAP008923
AGAP008924
AGAP008925
AGAP008926
AGAP008927
AGAP008928
AGAP008929
AGAP008930
AGAP008931
AGAP008932
AGAP008933
AGAP008935
AGAP008936
AGAP008937
AGAP008938
AGAP008939
AGAP008940
AGAP008941
AGAP008942
AGAP008943
AGAP008944
AGAP008945
AGAP008946
mean_tpm
time
preZGA
ZGA_early
ZGA_late
5h
9h
11h
14h
20h
24h
L1
sex
Female
Male

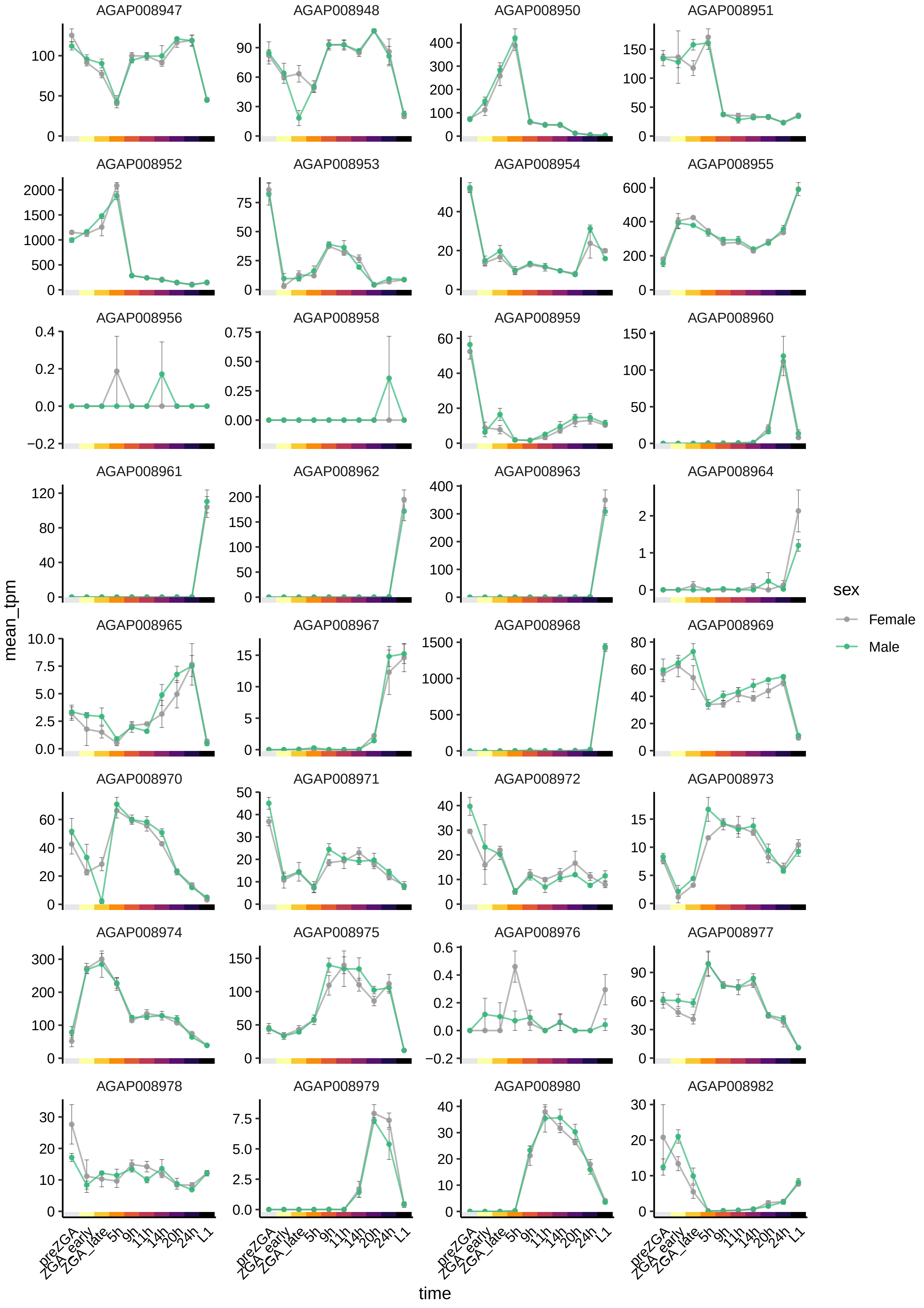

AGAP008947
AGAP008948
AGAP008950
AGAP008951
AGAP008952
AGAP008953
AGAP008954
AGAP008955
AGAP008956
AGAP008958
AGAP008959
AGAP008960
AGAP008961
AGAP008962
AGAP008963
AGAP008964
AGAP008965
AGAP008967
AGAP008968
AGAP008969
AGAP008970
AGAP008971
AGAP008972
AGAP008973
AGAP008974
AGAP008975
AGAP008976
AGAP008977
AGAP008978
AGAP008979
AGAP008980
AGAP008982
mean_tpm
time
preZGA
ZGA_early
ZGA_late
5h
9h
11h
14h
20h
24h
L1
sex
Female
Male

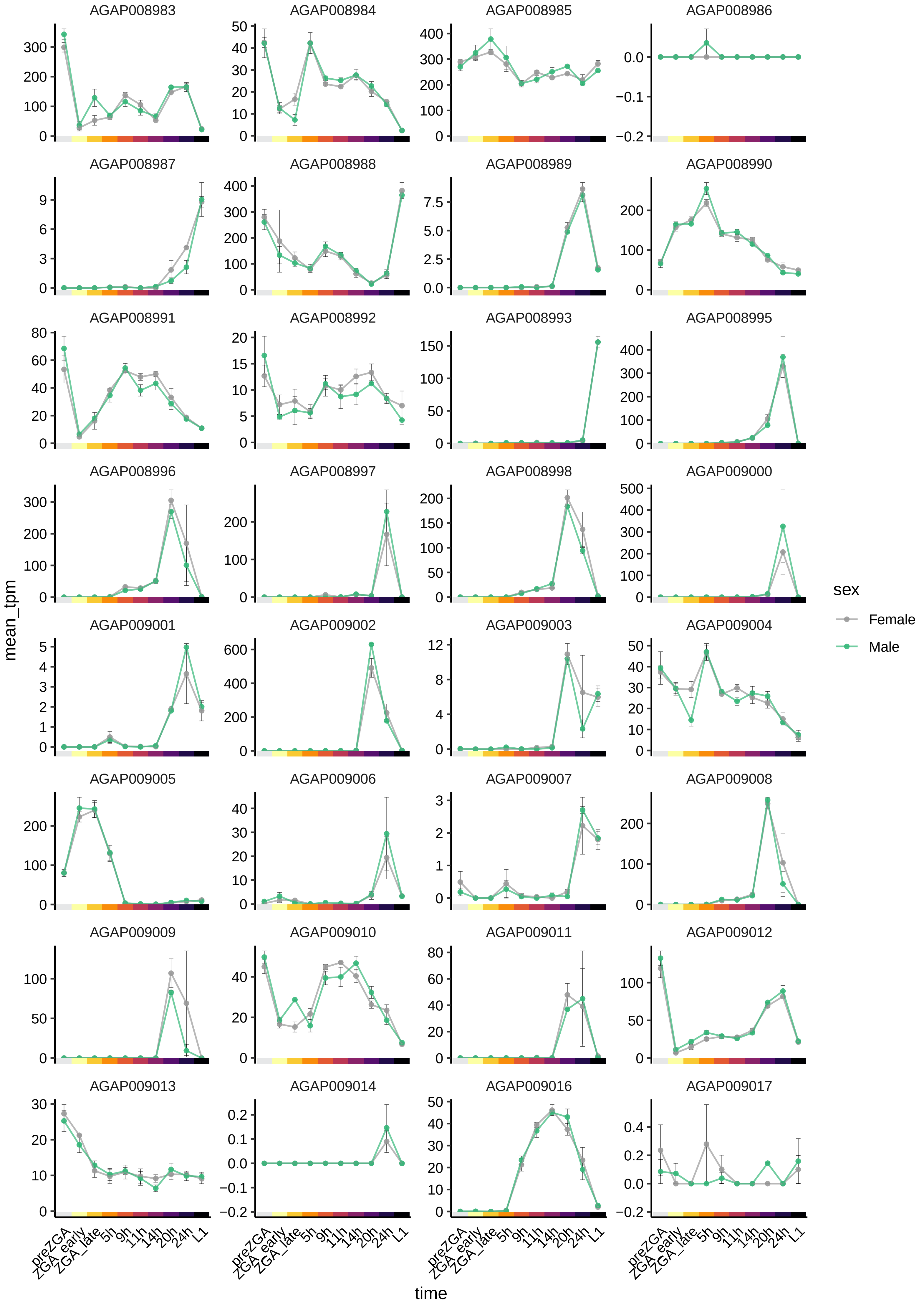

AGAP008983
AGAP008984
AGAP008985
AGAP008986
AGAP008987
AGAP008988
AGAP008989
AGAP008990
AGAP008991
AGAP008992
AGAP008993
AGAP008995
AGAP008996
AGAP008997
AGAP008998
AGAP009000
AGAP009001
AGAP009002
AGAP009003
AGAP009004
AGAP009005
AGAP009006
AGAP009007
AGAP009008
AGAP009009
AGAP009010
AGAP009011
AGAP009012
AGAP009013
AGAP009014
AGAP009016
AGAP009017
mean_tpm
time
preZGA
ZGA_early
ZGA_late
5h
9h
11h
14h
20h
24h
L1
sex
Female
Male

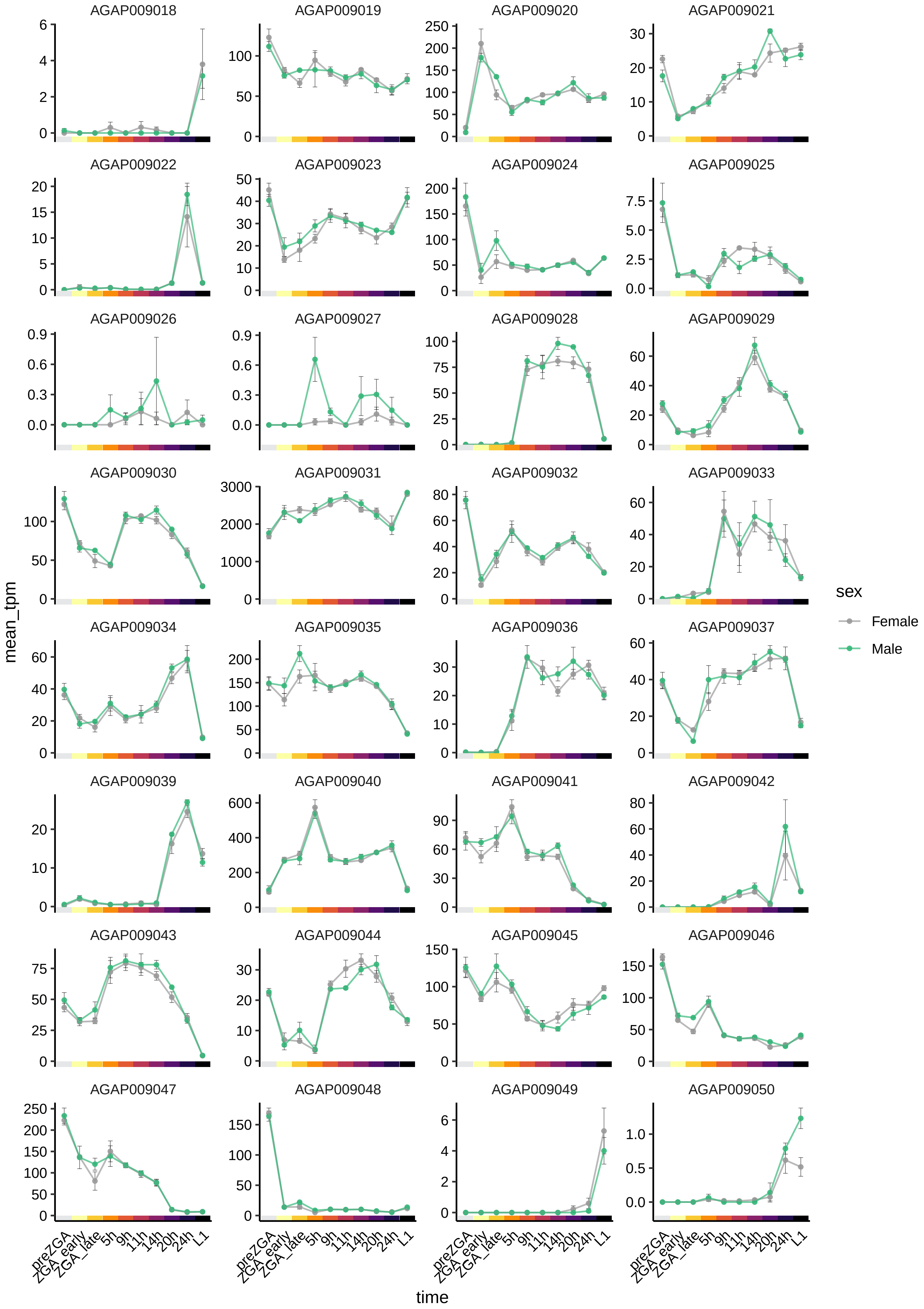
AGAP009018
AGAP009019
AGAP009020
AGAP009021
AGAP009022
AGAP009023
AGAP009024
AGAP009025
AGAP009026
AGAP009027
AGAP009028
AGAP009029
AGAP009030
AGAP009031
AGAP009032
AGAP009033
AGAP009034
AGAP009035
AGAP009036
AGAP009037
AGAP009039
AGAP009040
AGAP009041
AGAP009042
AGAP009043
AGAP009044
AGAP009045
AGAP009046
AGAP009047
AGAP009048
AGAP009049
AGAP009050
mean_tpm
time
preZGA
ZGA_early
ZGA_late
5h
9h
11h
14h
20h
24h
L1
sex
Female
Male

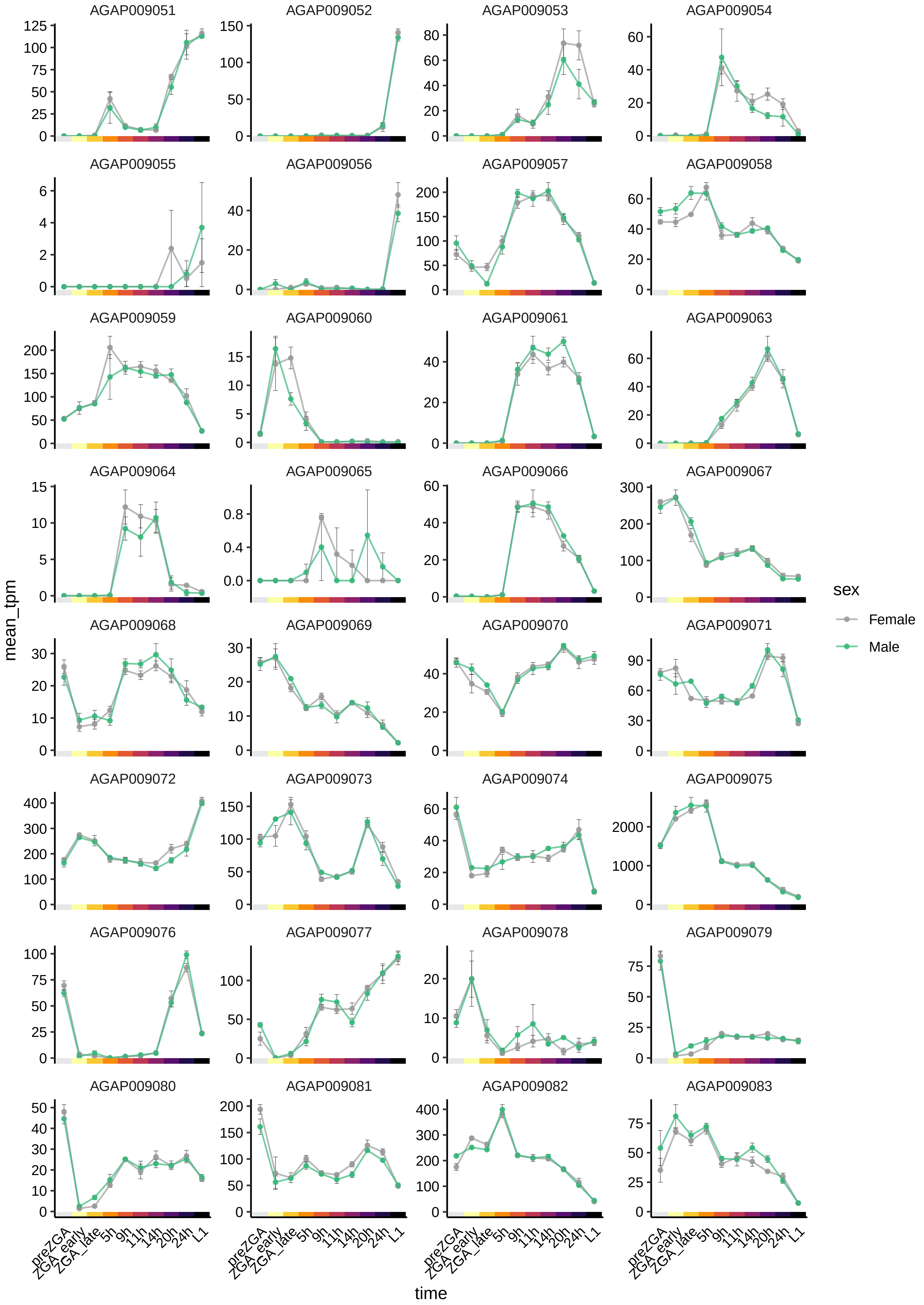
AGAP009051
AGAP009052
AGAP009053
AGAP009054
AGAP009055
AGAP009056
AGAP009057
AGAP009058
AGAP009059
AGAP009060
AGAP009061
AGAP009063
AGAP009064
AGAP009065
AGAP009066
AGAP009067
AGAP009068
AGAP009069
AGAP009070
AGAP009071
AGAP009072
AGAP009073
AGAP009074
AGAP009075
AGAP009076
AGAP009077
AGAP009078
AGAP009079
AGAP009080
AGAP009081
AGAP009082
AGAP009083
mean_tpm
time
preZGA
ZGA_early
ZGA_late
5h
9h
11h
14h
20h
24h
L1
sex
Female
Male

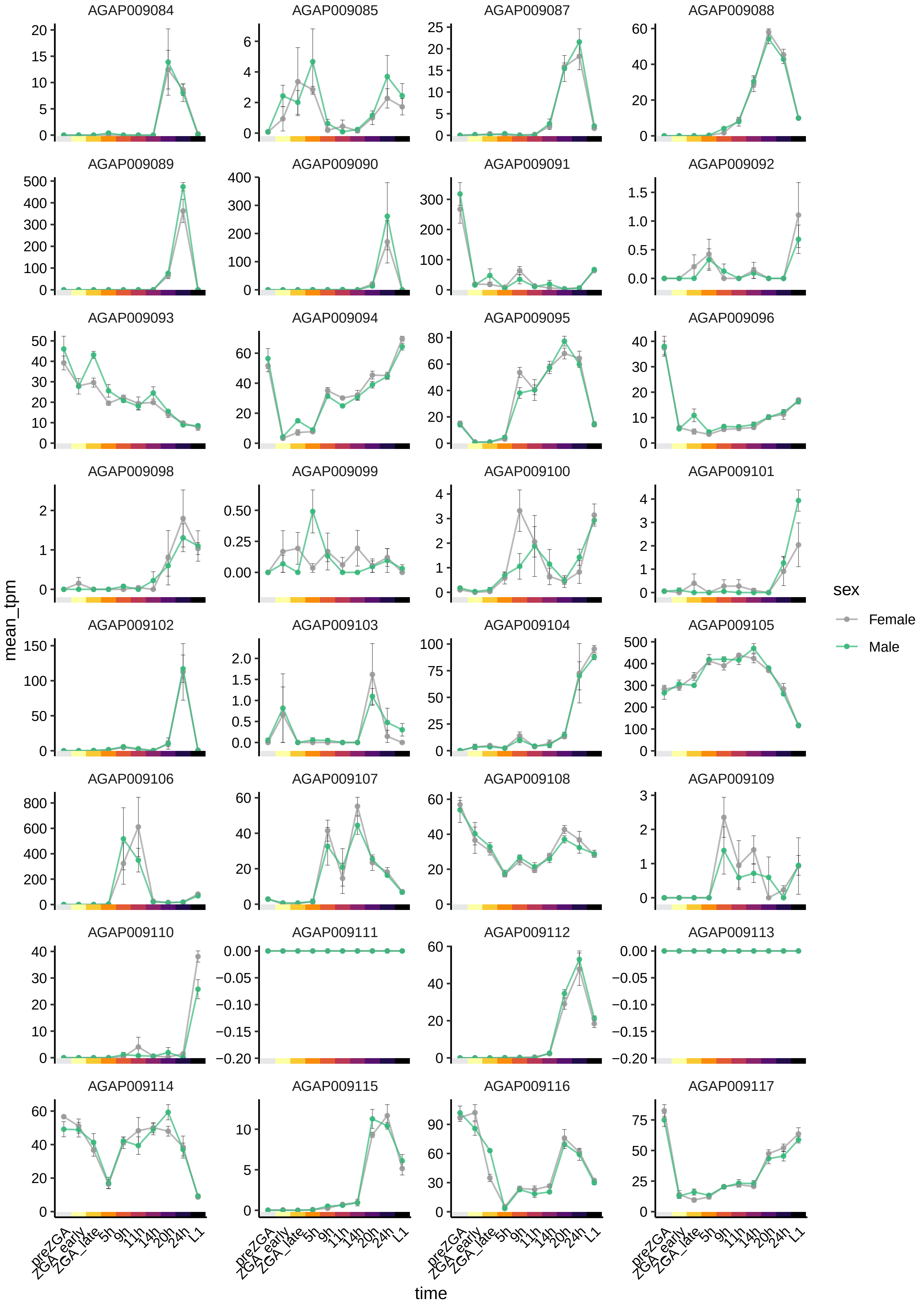

AGAP009084
AGAP009085
AGAP009087
AGAP009088
AGAP009089
AGAP009090
AGAP009091
AGAP009092
AGAP009093
AGAP009094
AGAP009095
AGAP009096
AGAP009098
AGAP009099
AGAP009100
AGAP009101
AGAP009102
AGAP009103
AGAP009104
AGAP009105
AGAP009106
AGAP009107
AGAP009108
AGAP009109
AGAP009110
AGAP009111
AGAP009112
AGAP009113
AGAP009114
AGAP009115
AGAP009116
AGAP009117
mean_tpm
time
preZGA
ZGA_early
ZGA_late
5h
9h
11h
14h
20h
24h
L1
sex
Female
Male

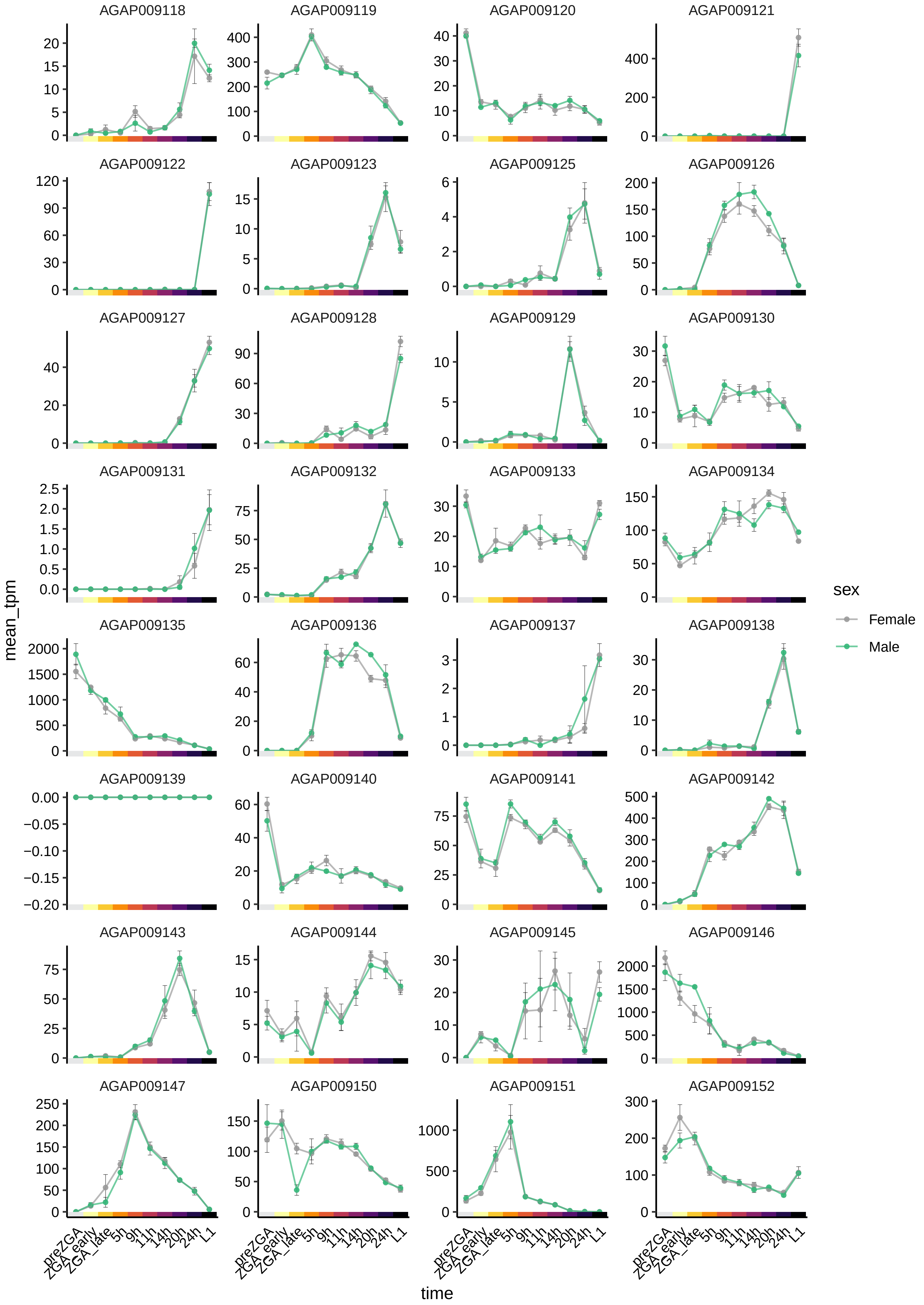

AGAP009118
AGAP009119
AGAP009120
AGAP009121
AGAP009122
AGAP009123
AGAP009125
AGAP009126
AGAP009127
AGAP009128
AGAP009129
AGAP009130
AGAP009131
AGAP009132
AGAP009133
AGAP009134
AGAP009135
AGAP009136
AGAP009137
AGAP009138
AGAP009139
AGAP009140
AGAP009141
AGAP009142
AGAP009143
AGAP009144
AGAP009145
AGAP009146
AGAP009147
AGAP009150
AGAP009151
AGAP009152
mean_tpm
time
preZGA
ZGA_early
ZGA_late
5h
9h
11h
14h
20h
24h
L1
sex
Female
Male

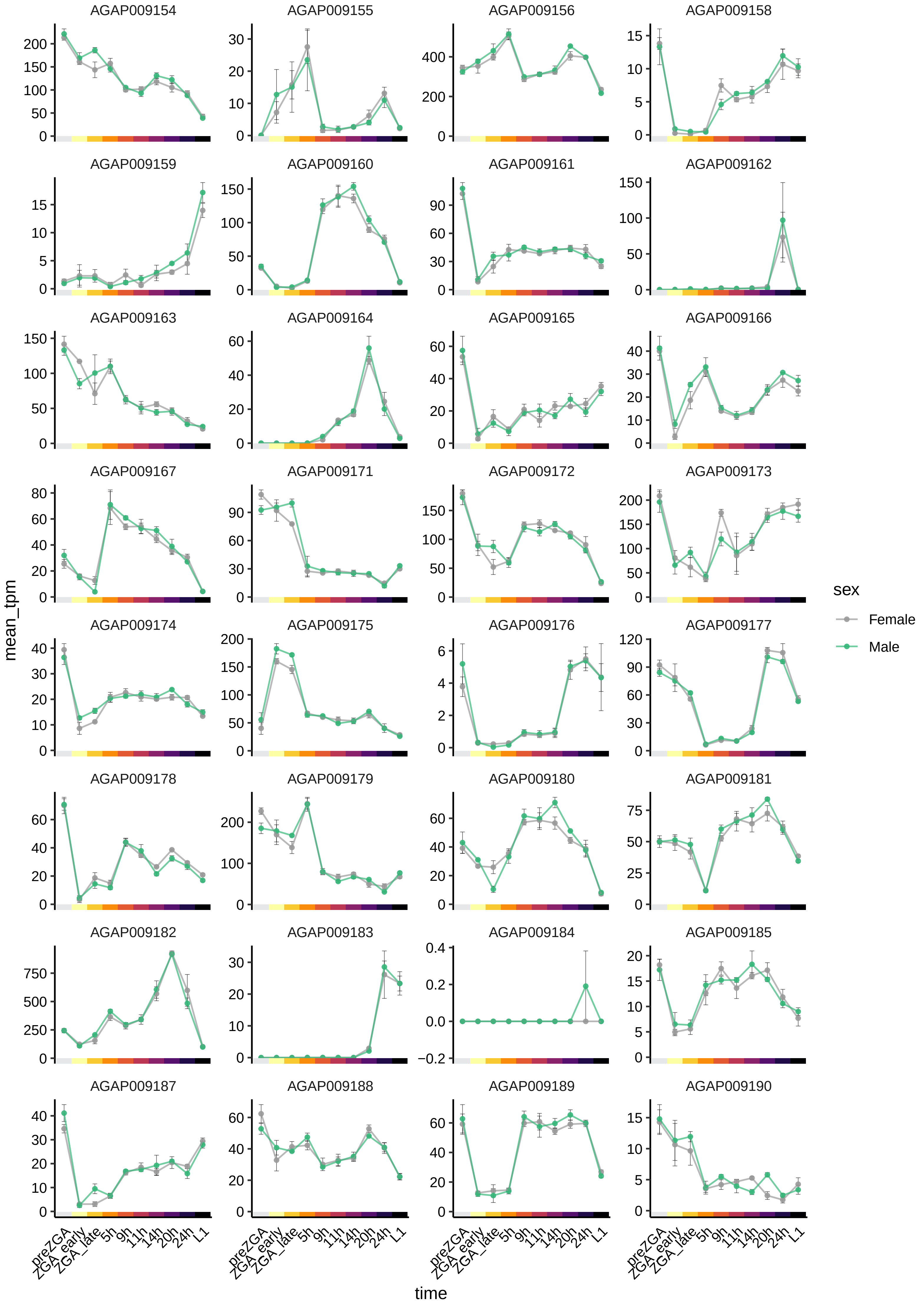

AGAP009154
AGAP009155
AGAP009156
AGAP009158
AGAP009159
AGAP009160
AGAP009161
AGAP009162
AGAP009163
AGAP009164
AGAP009165
AGAP009166
AGAP009167
AGAP009171
AGAP009172
AGAP009173
AGAP009174
AGAP009175
AGAP009176
AGAP009177
AGAP009178
AGAP009179
AGAP009180
AGAP009181
AGAP009182
AGAP009183
AGAP009184
AGAP009185
AGAP009187
AGAP009188
AGAP009189
AGAP009190
mean_tpm
time
preZGA
ZGA_early
ZGA_late
5h
9h
11h
14h
20h
24h
L1
sex
Female
Male

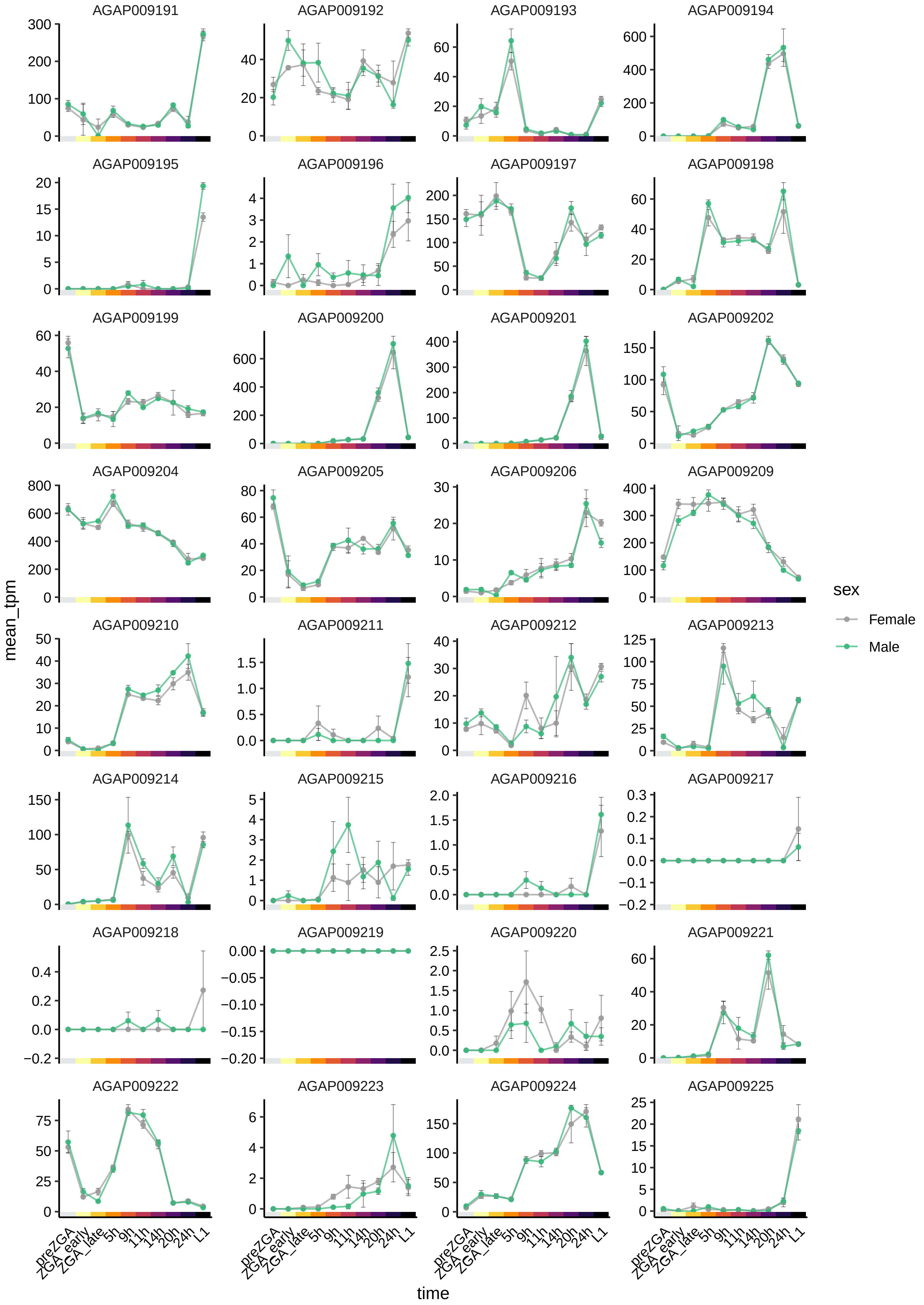
AGAP009191
AGAP009192
AGAP009193
AGAP009194
AGAP009195
AGAP009196
AGAP009197
AGAP009198
AGAP009199
AGAP009200
AGAP009201
AGAP009202
AGAP009204
AGAP009205
AGAP009206
AGAP009209
AGAP009210
AGAP009211
AGAP009212
AGAP009213
AGAP009214
AGAP009215
AGAP009216
AGAP009217
AGAP009218
AGAP009219
AGAP009220
AGAP009221
AGAP009222
AGAP009223
AGAP009224
AGAP009225
mean_tpm
time
preZGA
ZGA_early
ZGA_late
5h
9h
11h
14h
20h
24h
L1
sex
Female
Male

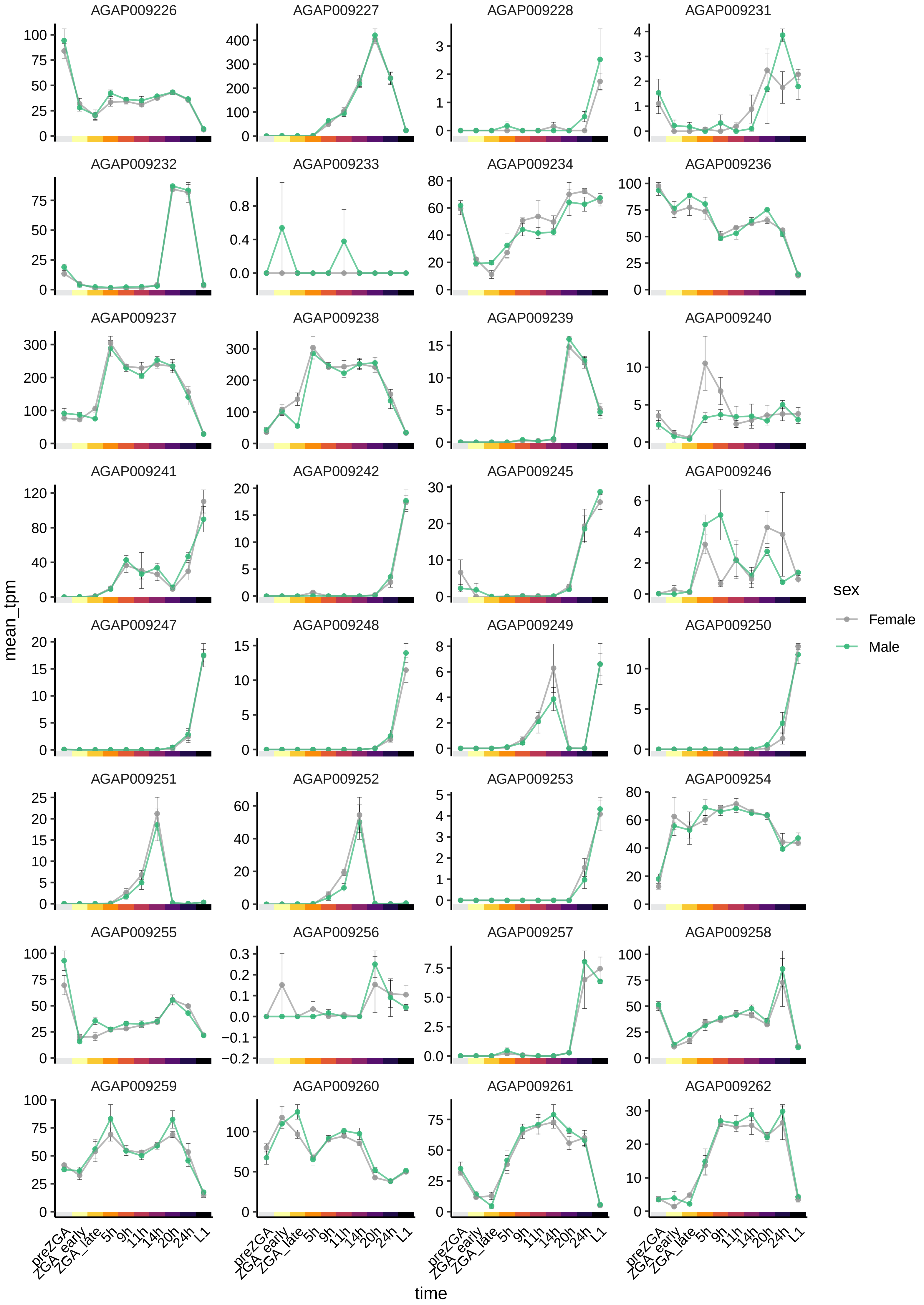
AGAP009226
AGAP009227
AGAP009228
AGAP009231
AGAP009232
AGAP009233
AGAP009234
AGAP009236
AGAP009237
AGAP009238
AGAP009239
AGAP009240
AGAP009241
AGAP009242
AGAP009245
AGAP009246
AGAP009247
AGAP009248
AGAP009249
AGAP009250
AGAP009251
AGAP009252
AGAP009253
AGAP009254
AGAP009255
AGAP009256
AGAP009257
AGAP009258
AGAP009259
AGAP009260
AGAP009261
AGAP009262
mean_tpm
time
preZGA
ZGA_early
ZGA_late
5h
9h
11h
14h
20h
24h
L1
sex
Female
Male

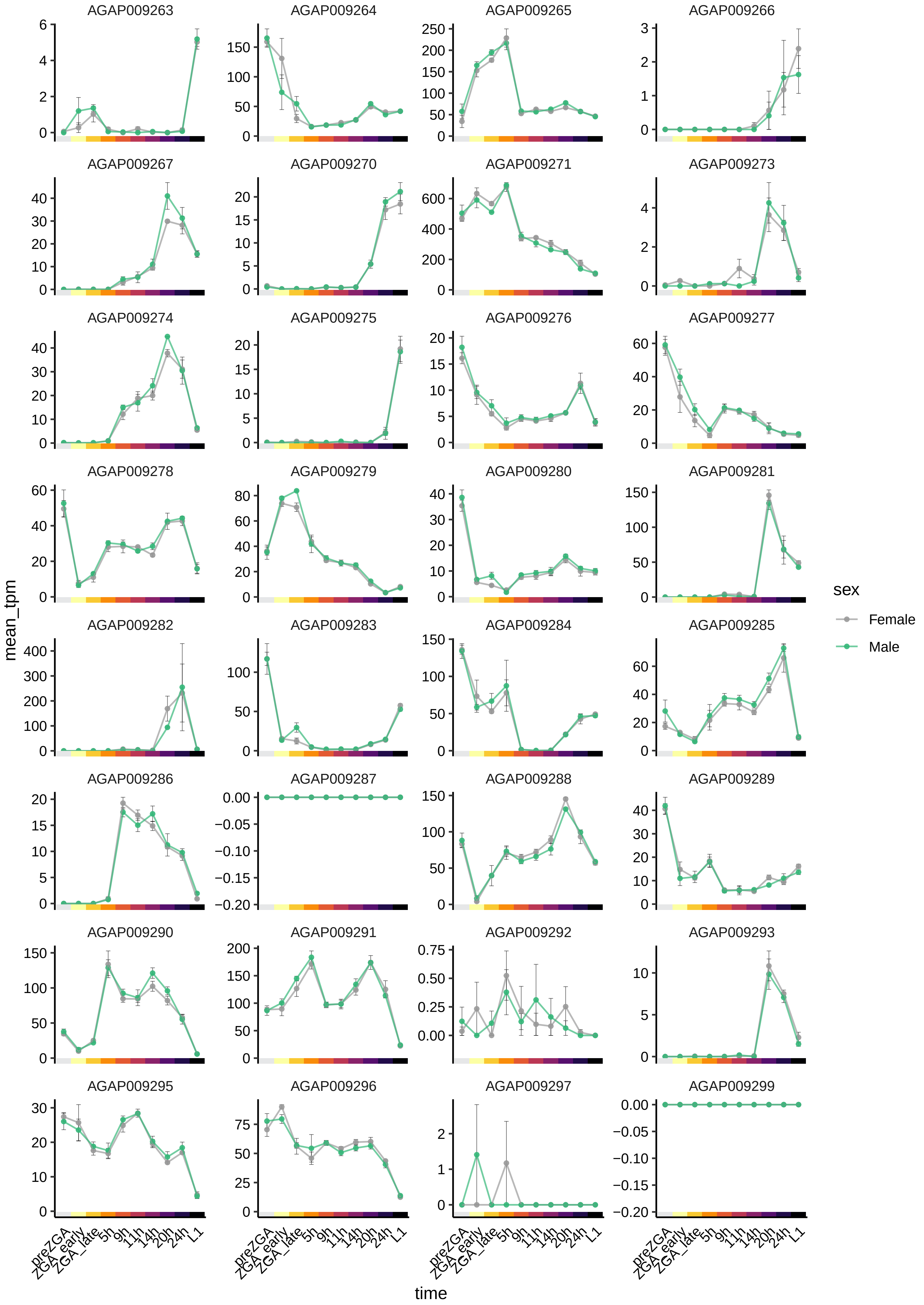
AGAP009263
AGAP009264
AGAP009265
AGAP009266
AGAP009267
AGAP009270
AGAP009271
AGAP009273
AGAP009274
AGAP009275
AGAP009276
AGAP009277
AGAP009278
AGAP009279
AGAP009280
AGAP009281
AGAP009282
AGAP009283
AGAP009284
AGAP009285
AGAP009286
AGAP009287
AGAP009288
AGAP009289
AGAP009290
AGAP009291
AGAP009292
AGAP009293
AGAP009295
AGAP009296
AGAP009297
AGAP009299
mean_tpm
time
preZGA
ZGA_early
ZGA_late
5h
9h
11h
14h
20h
24h
L1
sex
Female
Male

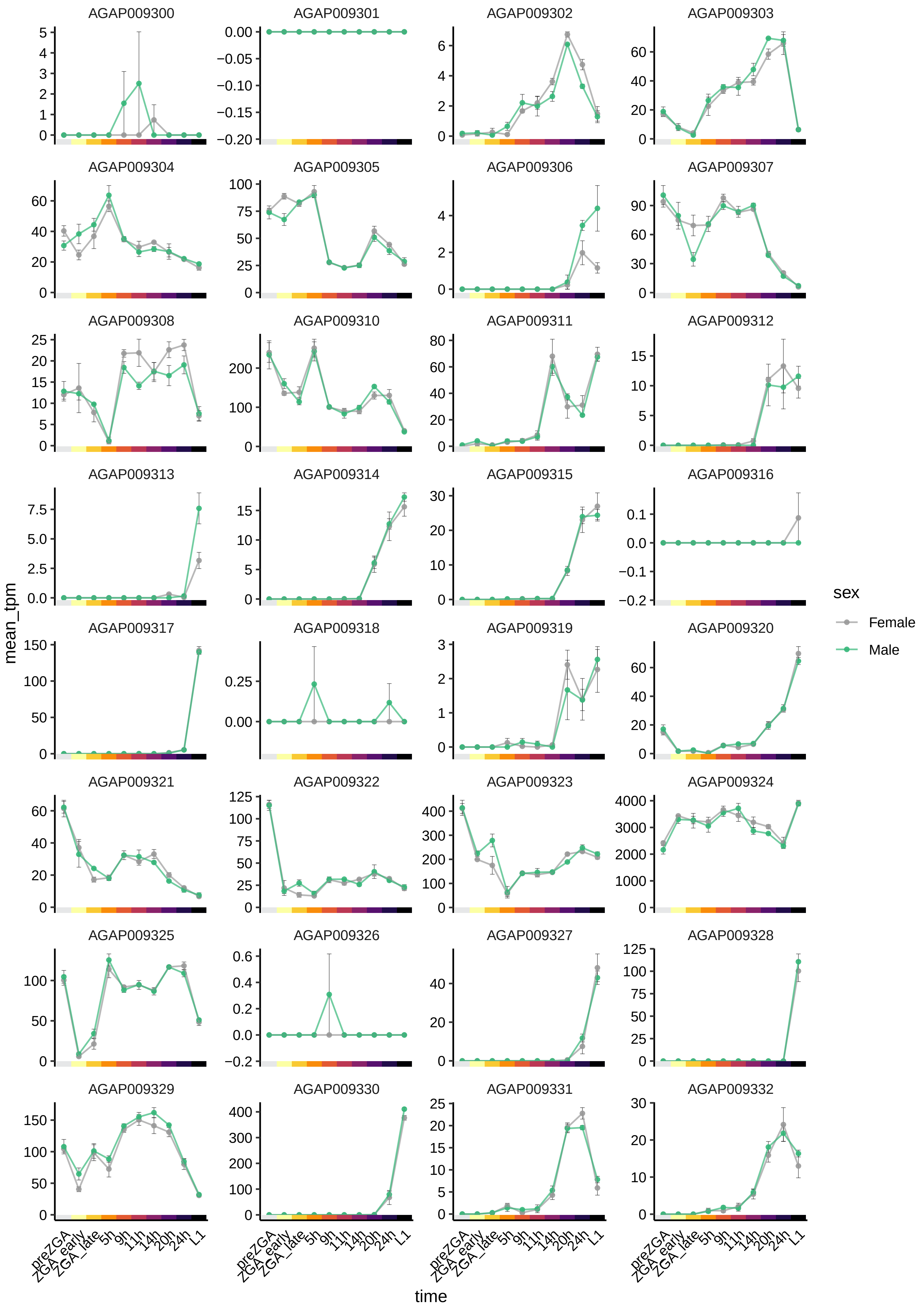
AGAP009300
AGAP009301
AGAP009302
AGAP009303
AGAP009304
AGAP009305
AGAP009306
AGAP009307
AGAP009308
AGAP009310
AGAP009311
AGAP009312
AGAP009313
AGAP009314
AGAP009315
AGAP009316
AGAP009317
AGAP009318
AGAP009319
AGAP009320
AGAP009321
AGAP009322
AGAP009323
AGAP009324
AGAP009325
AGAP009326
AGAP009327
AGAP009328
AGAP009329
AGAP009330
AGAP009331
AGAP009332
mean_tpm
time
preZGA
ZGA_early
ZGA_late
5h
9h
11h
14h
20h
24h
L1
sex
Female
Male

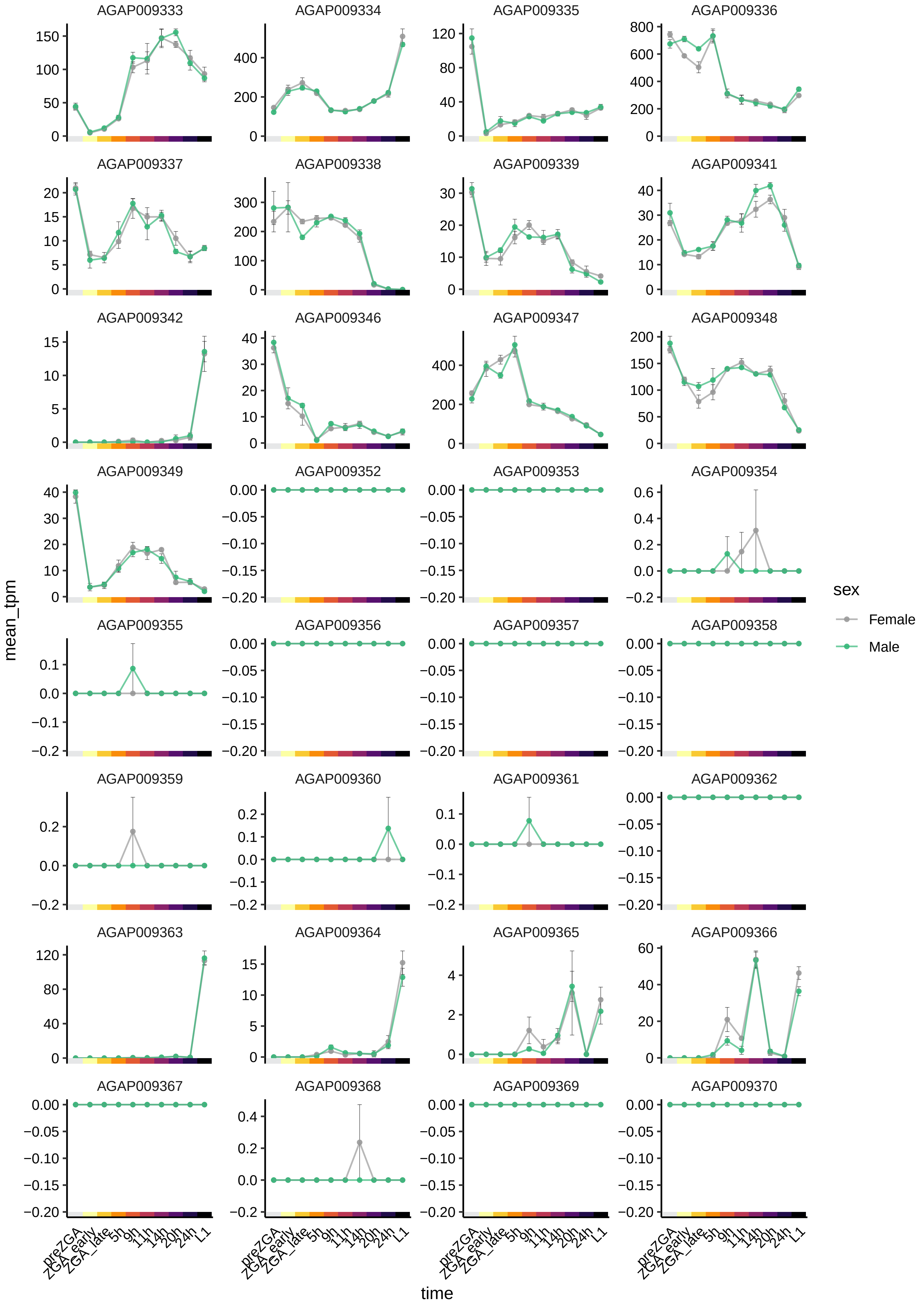

AGAP009333
AGAP009334
AGAP009335
AGAP009336
AGAP009337
AGAP009338
AGAP009339
AGAP009341
AGAP009342
AGAP009346
AGAP009347
AGAP009348
AGAP009349
AGAP009352
AGAP009353
AGAP009354
AGAP009355
AGAP009356
AGAP009357
AGAP009358
AGAP009359
AGAP009360
AGAP009361
AGAP009362
AGAP009363
AGAP009364
AGAP009365
AGAP009366
AGAP009367
AGAP009368
AGAP009369
AGAP009370
mean_tpm
time
preZGA
ZGA_early
ZGA_late
5h
9h
11h
14h
20h
24h
L1
sex
Female
Male

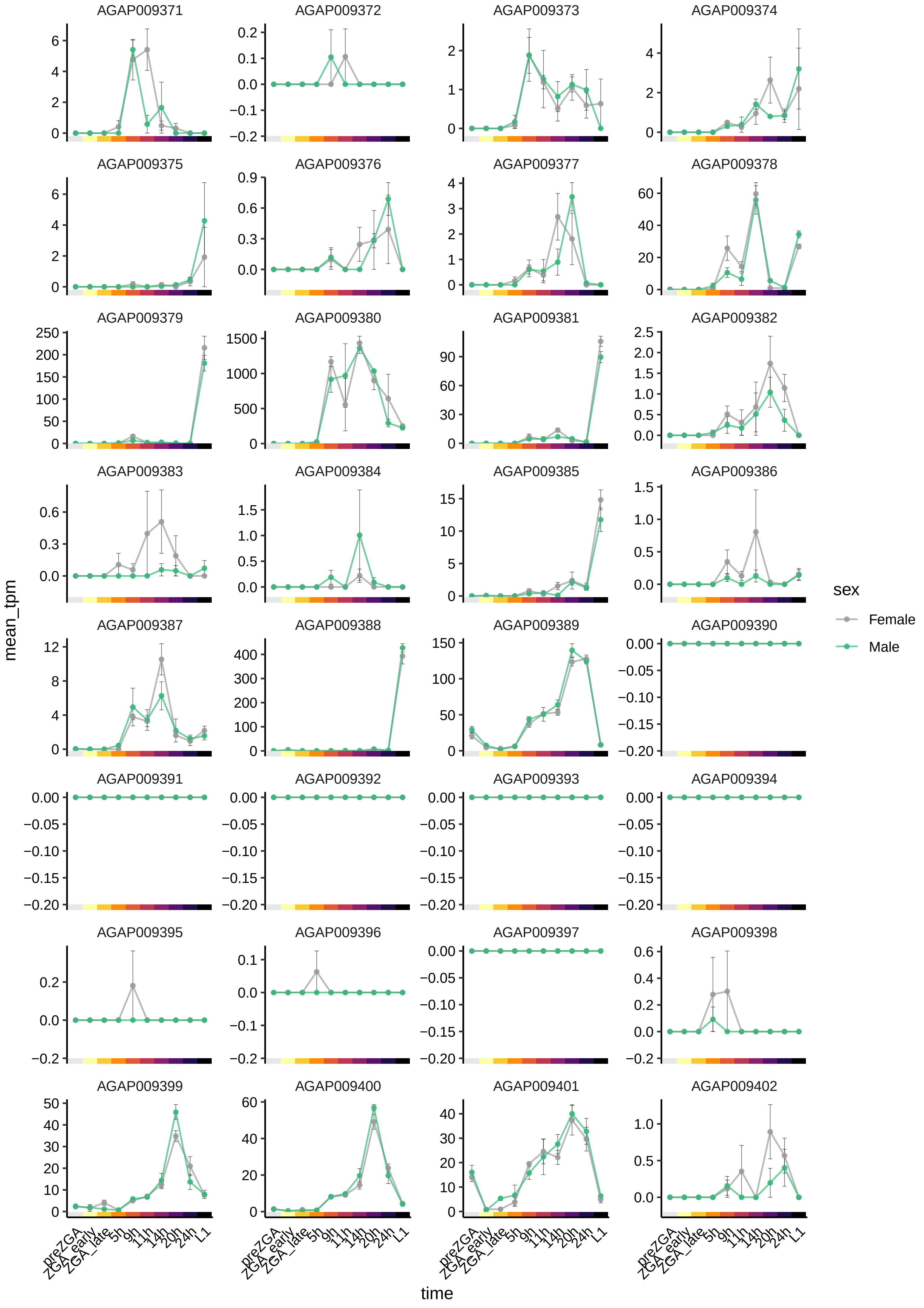

AGAP009371
AGAP009372
AGAP009373
AGAP009374
AGAP009375
AGAP009376
AGAP009377
AGAP009378
AGAP009379
AGAP009380
AGAP009381
AGAP009382
AGAP009383
AGAP009384
AGAP009385
AGAP009386
AGAP009387
AGAP009388
AGAP009389
AGAP009390
AGAP009391
AGAP009392
AGAP009393
AGAP009394
AGAP009395
AGAP009396
AGAP009397
AGAP009398
AGAP009399
AGAP009400
AGAP009401
AGAP009402
mean_tpm
time
preZGA
ZGA_early
ZGA_late
5h
9h
11h
14h
20h
24h
L1
sex
Female
Male

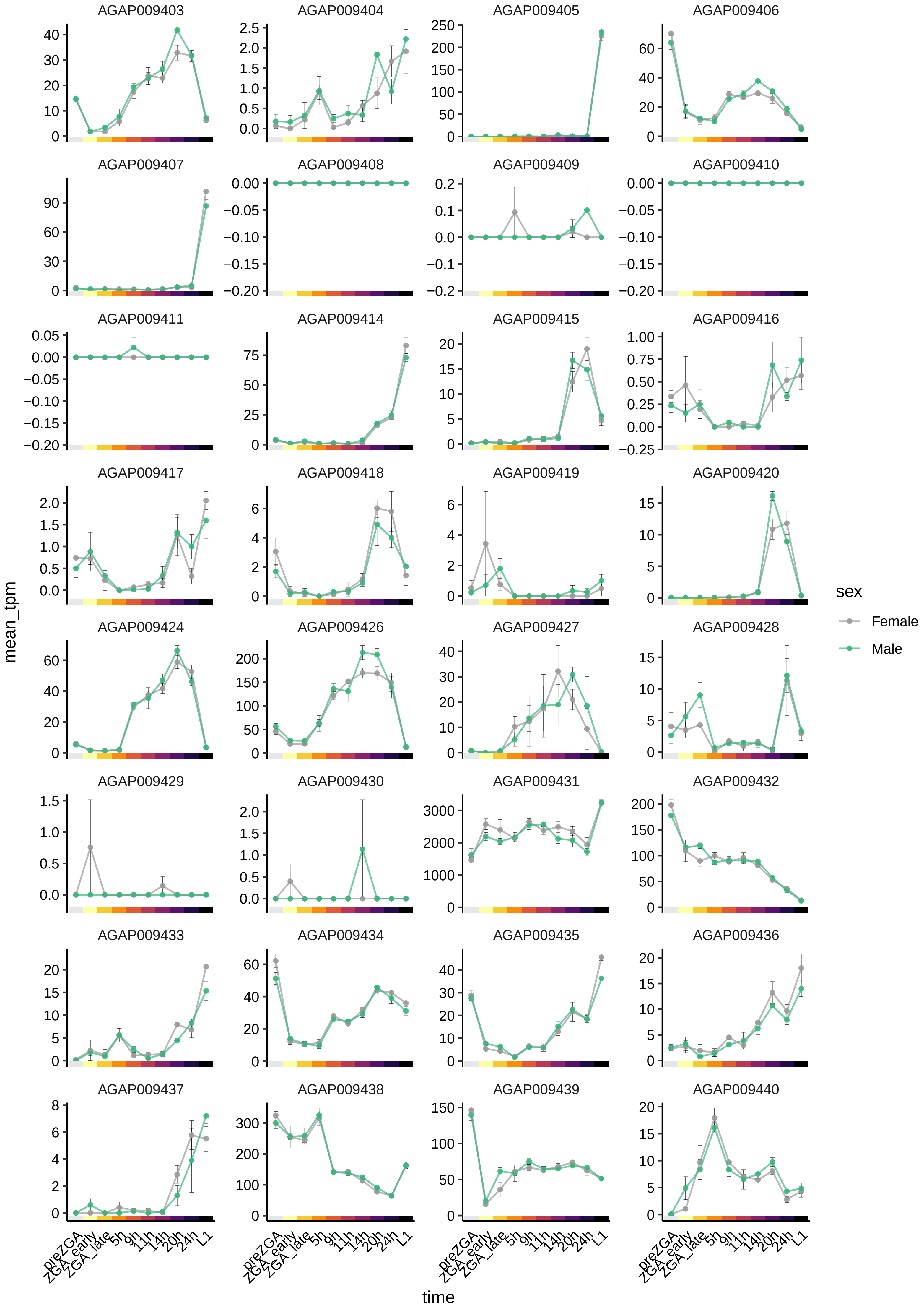

AGAP009403
AGAP009404
AGAP009405
AGAP009406
AGAP009407
AGAP009408
AGAP009409
AGAP009410
AGAP009411
AGAP009414
AGAP009415
AGAP009416
AGAP009417
AGAP009418
AGAP009419
AGAP009420
AGAP009424
AGAP009426
AGAP009427
AGAP009428
AGAP009429
AGAP009430
AGAP009431
AGAP009432
AGAP009433
AGAP009434
AGAP009435
AGAP009436
AGAP009437
AGAP009438
AGAP009439
AGAP009440
mean_tpm
time
preZGA
ZGA_early
ZGA_late
5h
9h
11h
14h
20h
24h
L1
sex
Female
Male

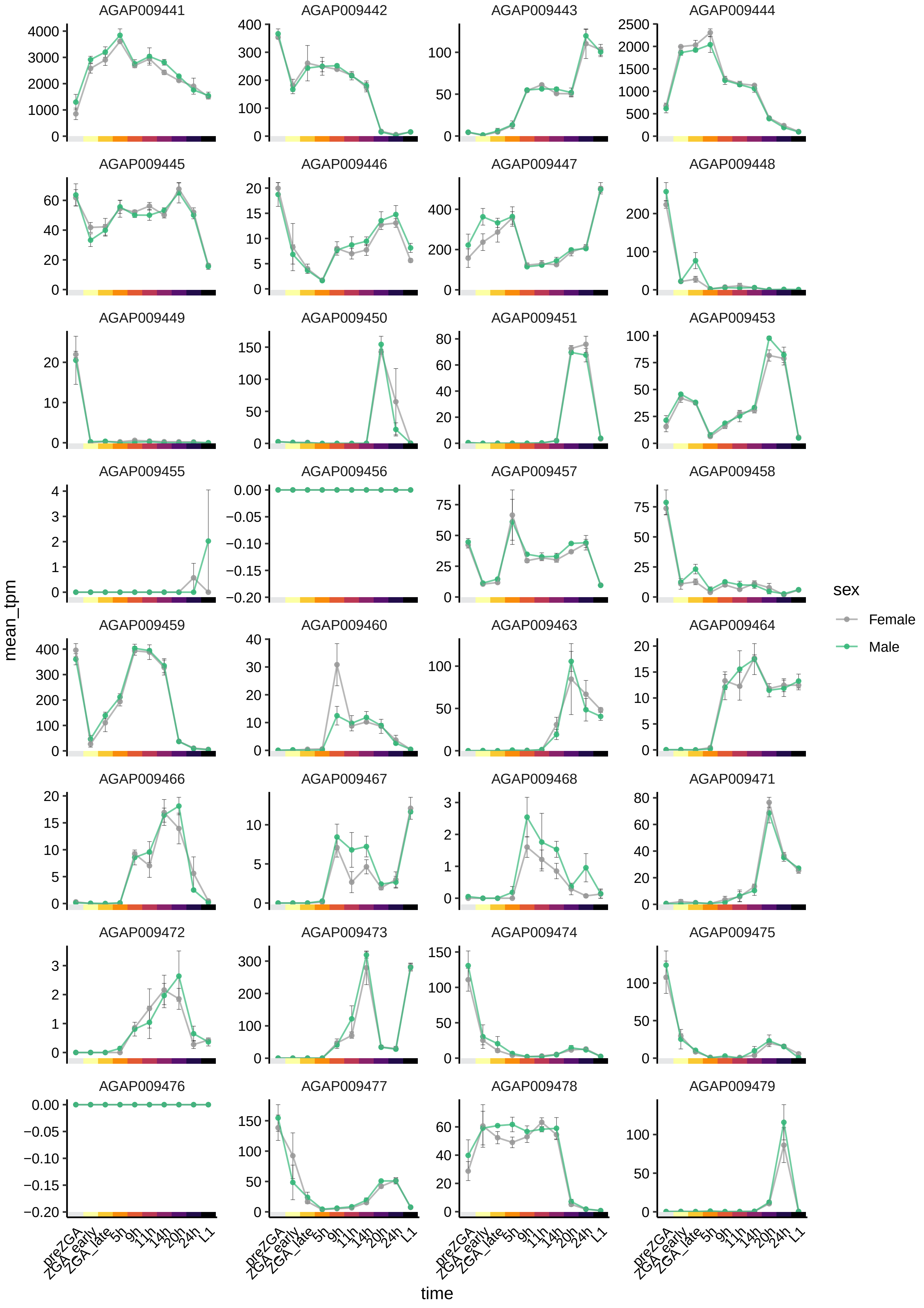
AGAP009441
AGAP009442
AGAP009443
AGAP009444
AGAP009445
AGAP009446
AGAP009447
AGAP009448
AGAP009449
AGAP009450
AGAP009451
AGAP009453
AGAP009455
AGAP009456
AGAP009457
AGAP009458
AGAP009459
AGAP009460
AGAP009463
AGAP009464
AGAP009466
AGAP009467
AGAP009468
AGAP009471
AGAP009472
AGAP009473
AGAP009474
AGAP009475
AGAP009476
AGAP009477
AGAP009478
AGAP009479
mean_tpm
time
preZGA
ZGA_early
ZGA_late
5h
9h
11h
14h
20h
24h
L1
sex
Female
Male

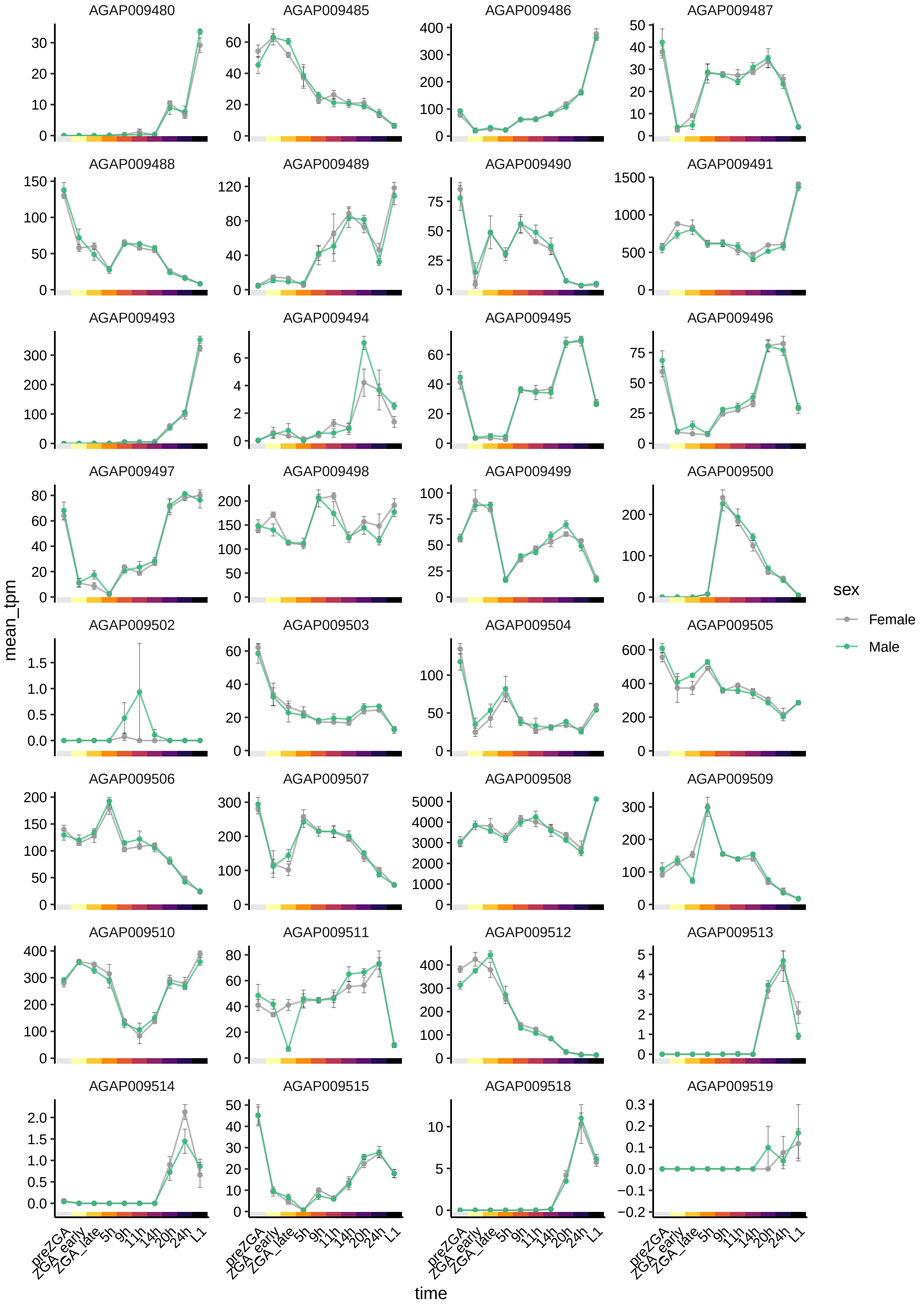

AGAP009480
AGAP009485
AGAP009486
AGAP009487
AGAP009488
AGAP009489
AGAP009490
AGAP009491
AGAP009493
AGAP009494
AGAP009495
AGAP009496
AGAP009497
AGAP009498
AGAP009499
AGAP009500
AGAP009502
AGAP009503
AGAP009504
AGAP009505
AGAP009506
AGAP009507
AGAP009508
AGAP009509
AGAP009510
AGAP009511
AGAP009512
AGAP009513
AGAP009514
AGAP009515
AGAP009518
AGAP009519
mean_tpm
time
preZGA
ZGA_early
ZGA_late
5h
9h
11h
14h
20h
24h
L1
sex
Female
Male

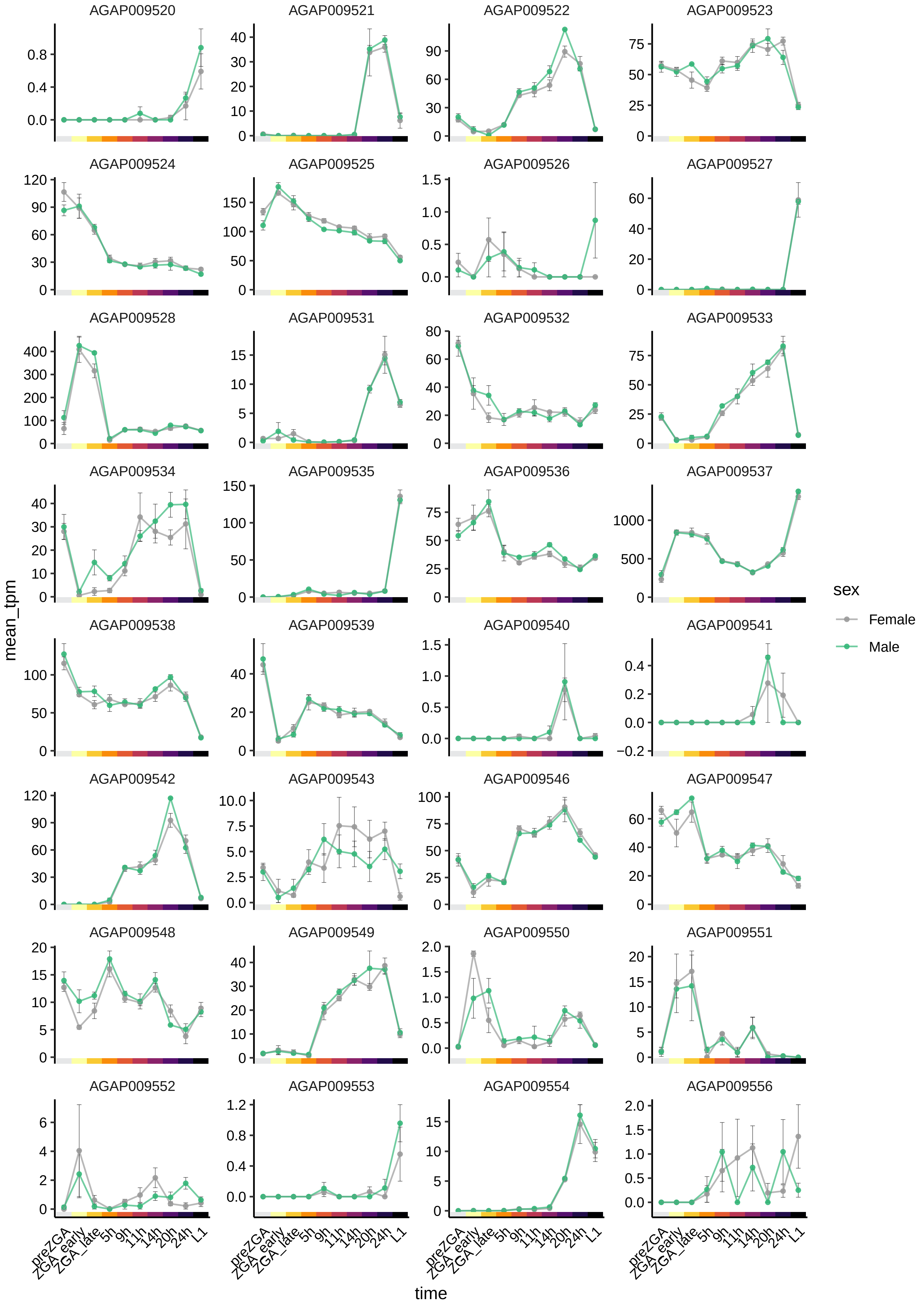

AGAP009520
AGAP009521
AGAP009522
AGAP009523
AGAP009524
AGAP009525
AGAP009526
AGAP009527
AGAP009528
AGAP009531
AGAP009532
AGAP009533
AGAP009534
AGAP009535
AGAP009536
AGAP009537
AGAP009538
AGAP009539
AGAP009540
AGAP009541
AGAP009542
AGAP009543
AGAP009546
AGAP009547
AGAP009548
AGAP009549
AGAP009550
AGAP009551
AGAP009552
AGAP009553
AGAP009554
AGAP009556
mean_tpm
time
preZGA
ZGA_early
ZGA_late
5h
9h
11h
14h
20h
24h
L1
sex
Female
Male

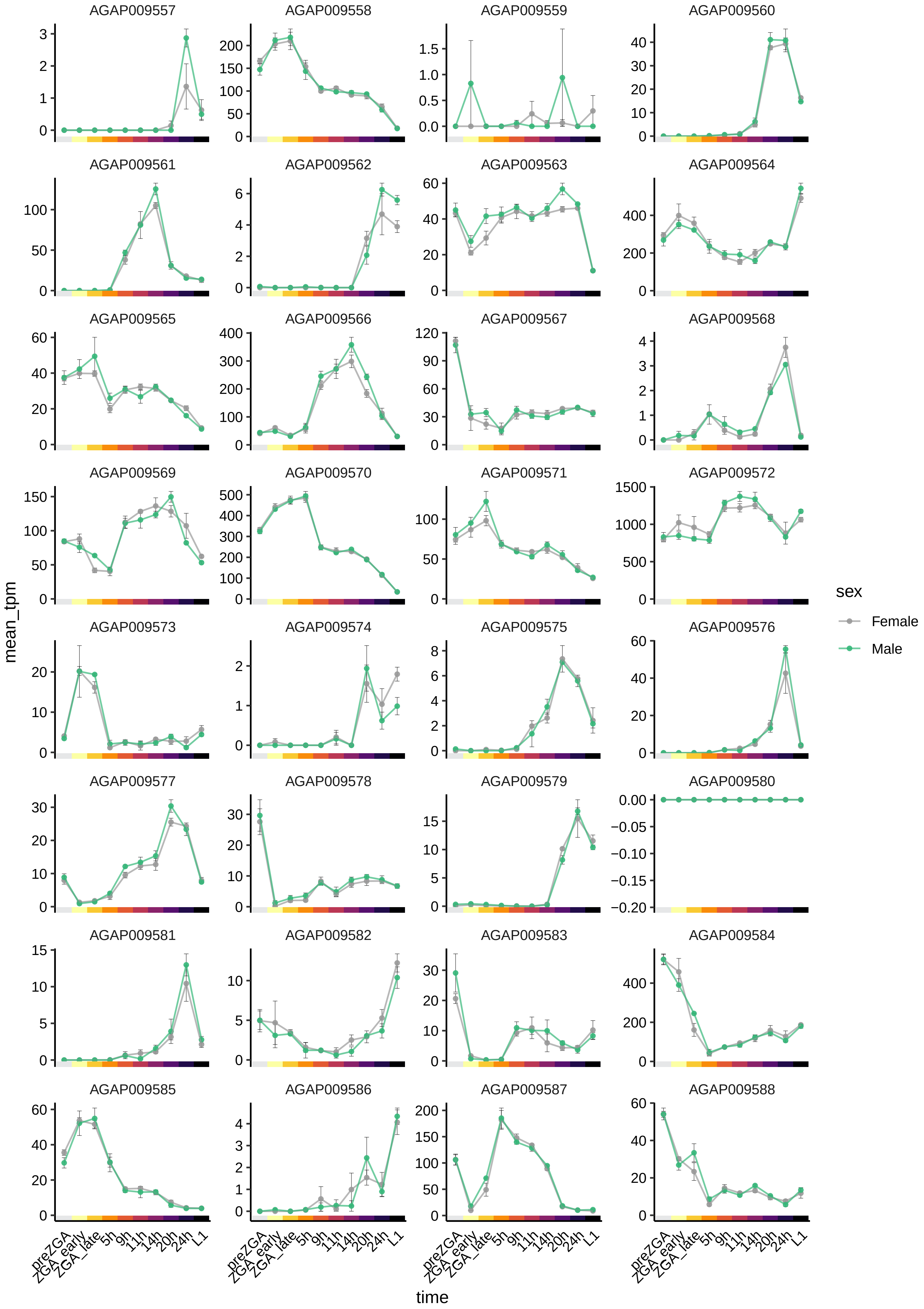
AGAP009557
AGAP009558
AGAP009559
AGAP009560
AGAP009561
AGAP009562
AGAP009563
AGAP009564
AGAP009565
AGAP009566
AGAP009567
AGAP009568
AGAP009569
AGAP009570
AGAP009571
AGAP009572
AGAP009573
AGAP009574
AGAP009575
AGAP009576
AGAP009577
AGAP009578
AGAP009579
AGAP009580
AGAP009581
AGAP009582
AGAP009583
AGAP009584
AGAP009585
AGAP009586
AGAP009587
AGAP009588
mean_tpm
time
preZGA
ZGA_early
ZGA_late
5h
9h
11h
14h
20h
24h
L1
sex
Female
Male

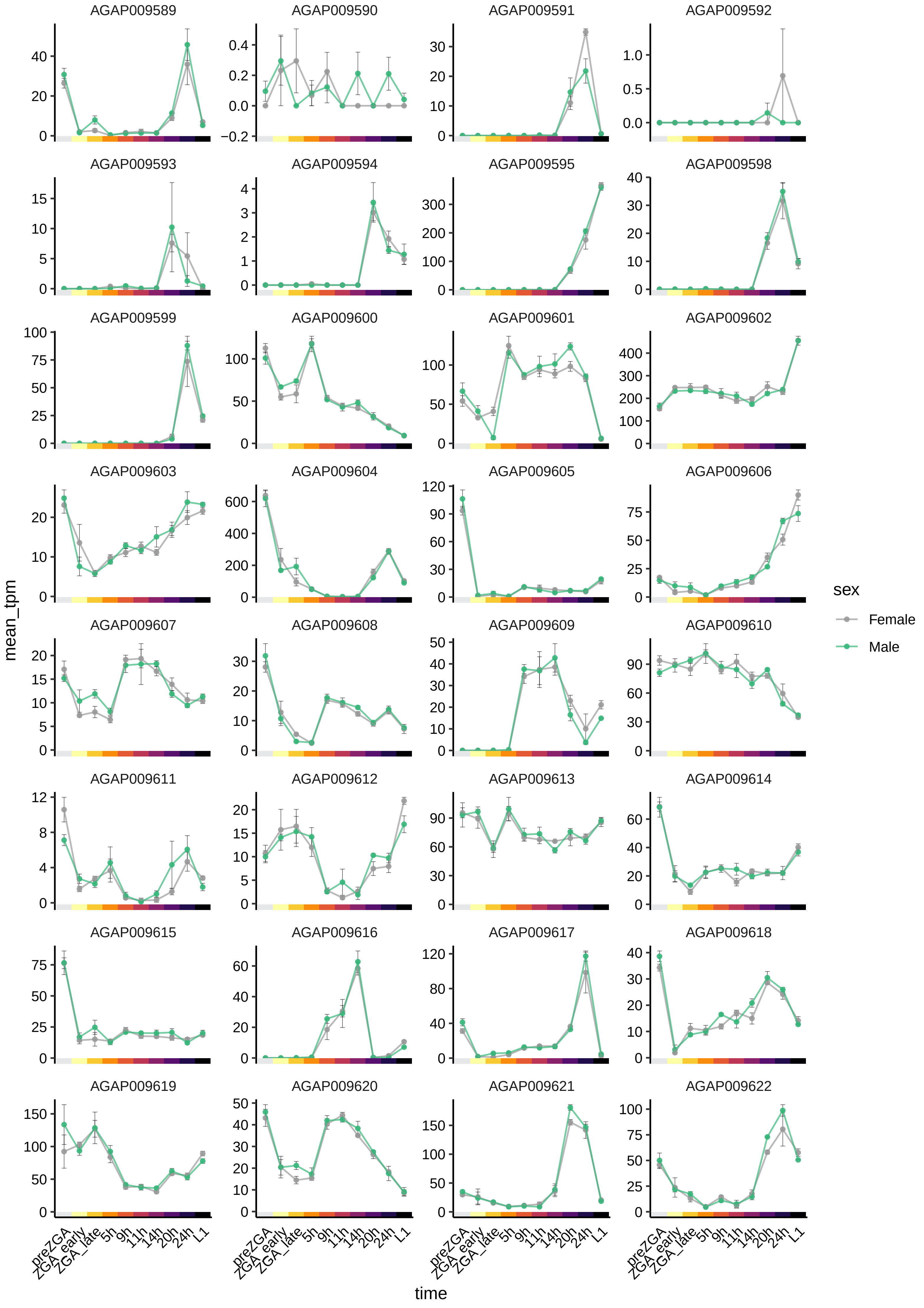
AGAP009589
AGAP009590
AGAP009591
AGAP009592
AGAP009593
AGAP009594
AGAP009595
AGAP009598
AGAP009599
AGAP009600
AGAP009601
AGAP009602
AGAP009603
AGAP009604
AGAP009605
AGAP009606
AGAP009607
AGAP009608
AGAP009609
AGAP009610
AGAP009611
AGAP009612
AGAP009613
AGAP009614
AGAP009615
AGAP009616
AGAP009617
AGAP009618
AGAP009619
AGAP009620
AGAP009621
AGAP009622
mean_tpm
time
preZGA
ZGA_early
ZGA_late
5h
9h
11h
14h
20h
24h
L1
sex
Female
Male

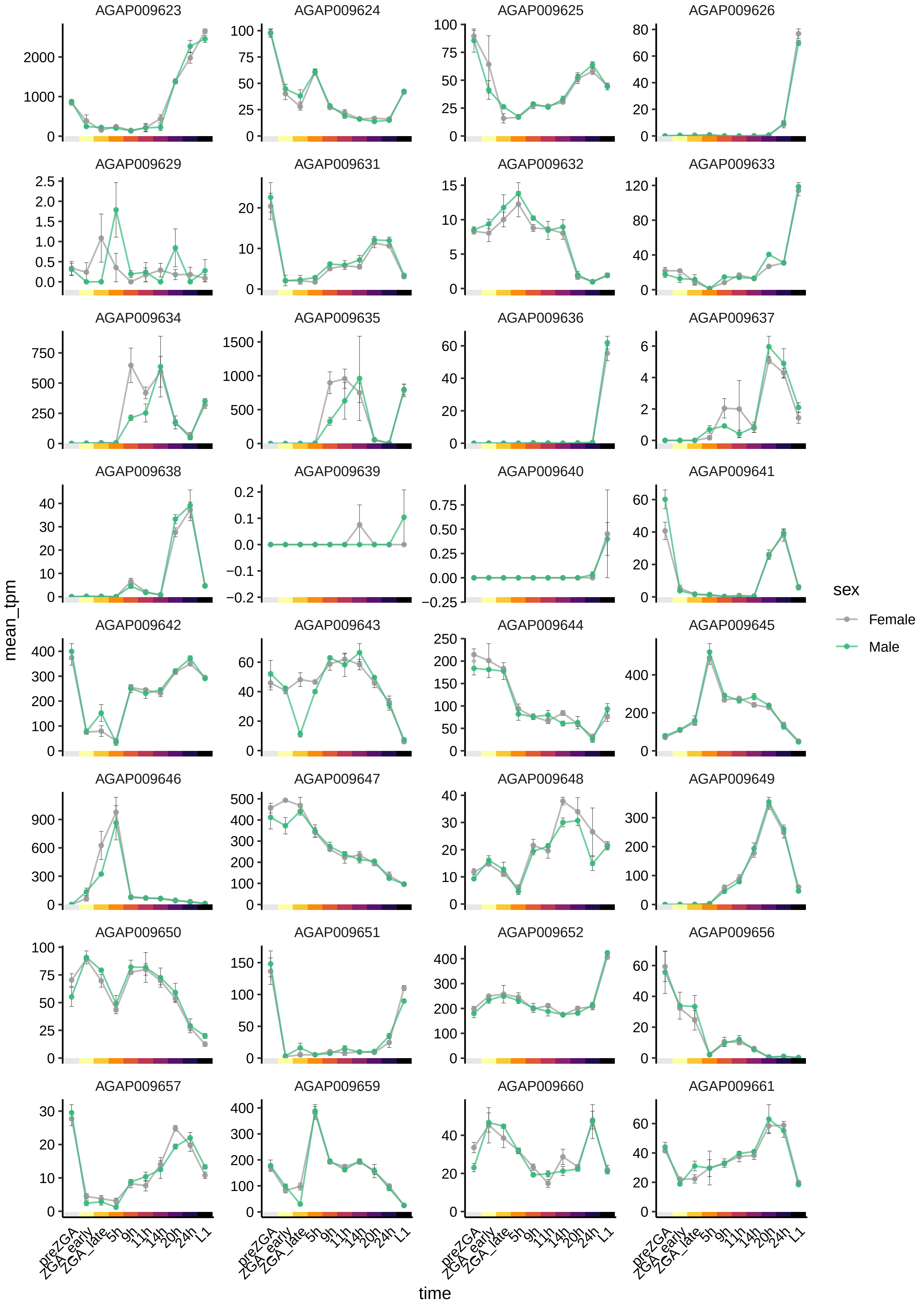
AGAP009623
AGAP009624
AGAP009625
AGAP009626
AGAP009629
AGAP009631
AGAP009632
AGAP009633
AGAP009634
AGAP009635
AGAP009636
AGAP009637
AGAP009638
AGAP009639
AGAP009640
AGAP009641
AGAP009642
AGAP009643
AGAP009644
AGAP009645
AGAP009646
AGAP009647
AGAP009648
AGAP009649
AGAP009650
AGAP009651
AGAP009652
AGAP009656
AGAP009657
AGAP009659
AGAP009660
AGAP009661
mean_tpm
time
preZGA
ZGA_early
ZGA_late
5h
9h
11h
14h
20h
24h
L1
sex
Female
Male

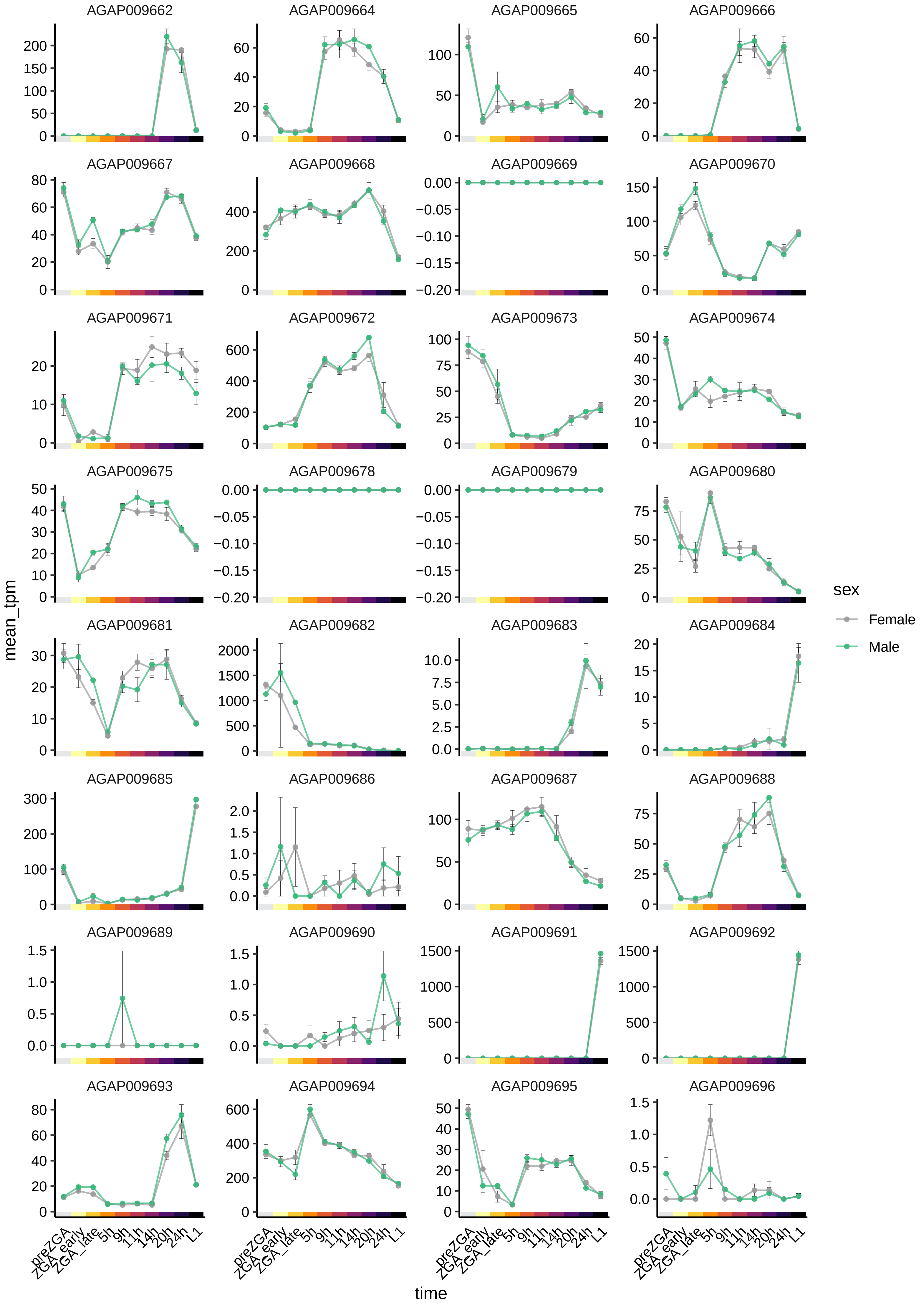
AGAP009662
AGAP009664
AGAP009665
AGAP009666
AGAP009667
AGAP009668
AGAP009669
AGAP009670
AGAP009671
AGAP009672
AGAP009673
AGAP009674
AGAP009675
AGAP009678
AGAP009679
AGAP009680
AGAP009681
AGAP009682
AGAP009683
AGAP009684
AGAP009685
AGAP009686
AGAP009687
AGAP009688
AGAP009689
AGAP009690
AGAP009691
AGAP009692
AGAP009693
AGAP009694
AGAP009695
AGAP009696
mean_tpm
time
preZGA
ZGA_early
ZGA_late
5h
9h
11h
14h
20h
24h
L1
sex
Female
Male

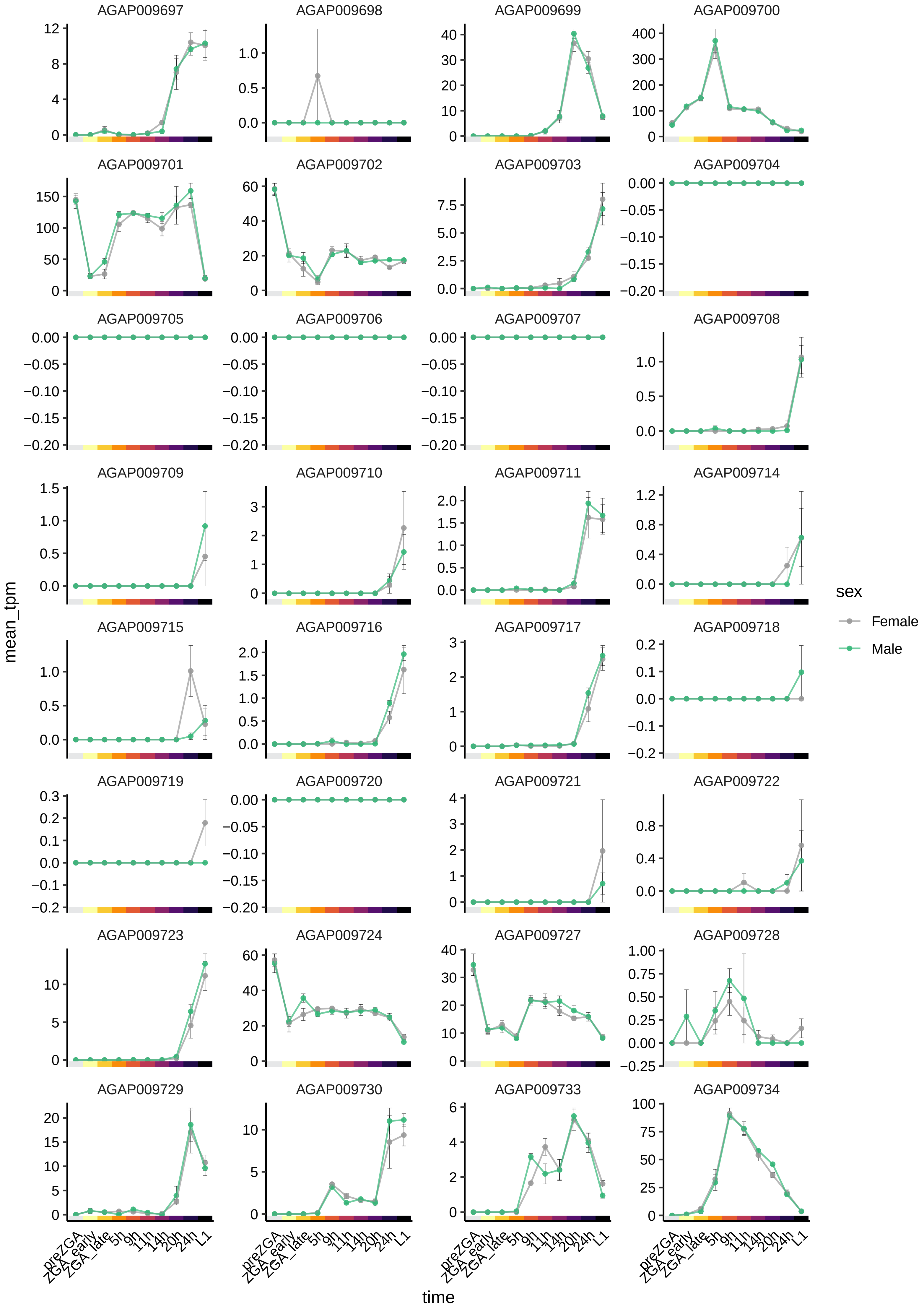

AGAP009697
AGAP009698
AGAP009699
AGAP009700
AGAP009701
AGAP009702
AGAP009703
AGAP009704
AGAP009705
AGAP009706
AGAP009707
AGAP009708
AGAP009709
AGAP009710
AGAP009711
AGAP009714
AGAP009715
AGAP009716
AGAP009717
AGAP009718
AGAP009719
AGAP009720
AGAP009721
AGAP009722
AGAP009723
AGAP009724
AGAP009727
AGAP009728
AGAP009729
AGAP009730
AGAP009733
AGAP009734
mean_tpm
time
preZGA
ZGA_early
ZGA_late
5h
9h
11h
14h
20h
24h
L1
sex
Female
Male

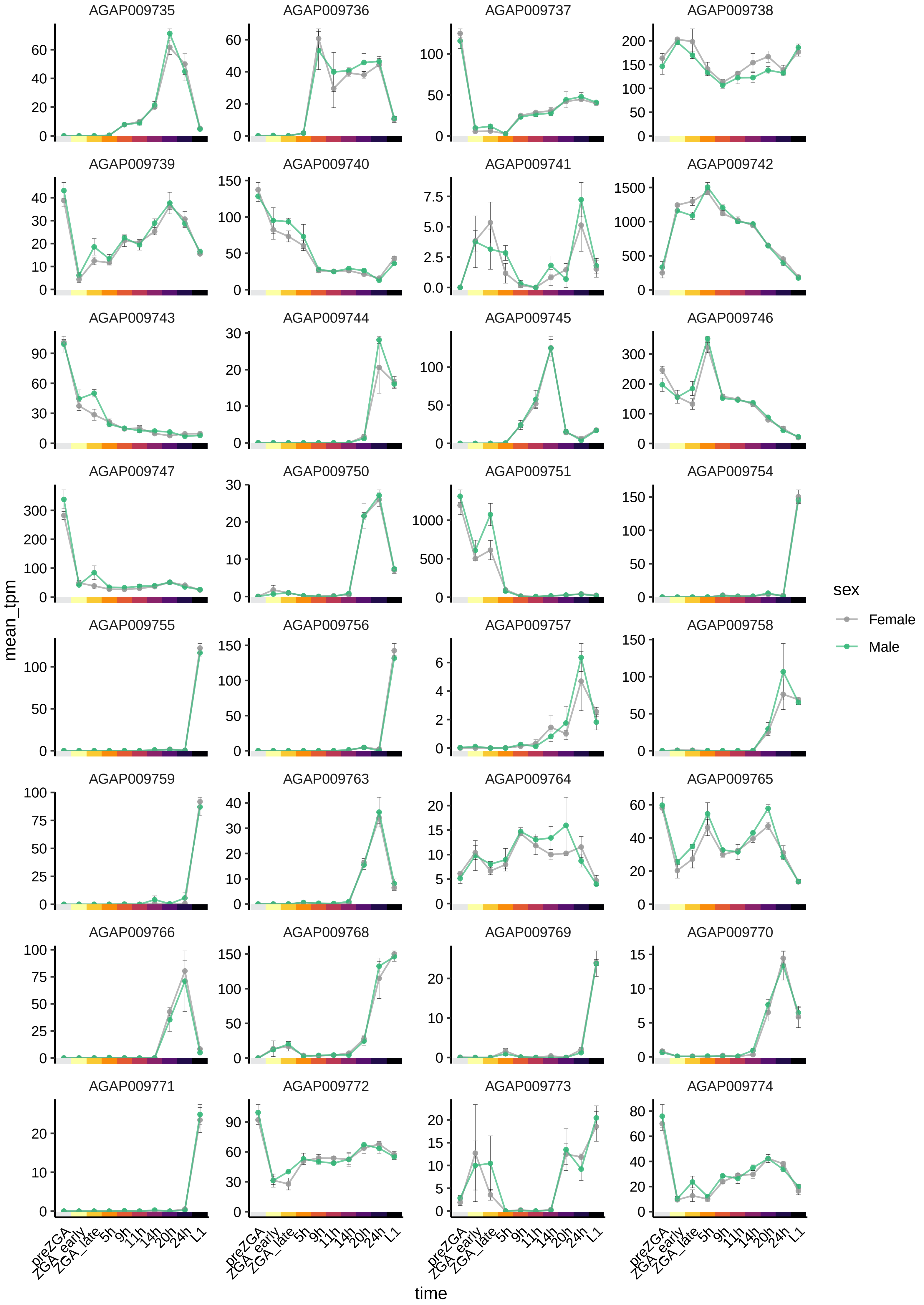

AGAP009735
AGAP009736
AGAP009737
AGAP009738
AGAP009739
AGAP009740
AGAP009741
AGAP009742
AGAP009743
AGAP009744
AGAP009745
AGAP009746
AGAP009747
AGAP009750
AGAP009751
AGAP009754
AGAP009755
AGAP009756
AGAP009757
AGAP009758
AGAP009759
AGAP009763
AGAP009764
AGAP009765
AGAP009766
AGAP009768
AGAP009769
AGAP009770
AGAP009771
AGAP009772
AGAP009773
AGAP009774
mean_tpm
time
preZGA
ZGA_early
ZGA_late
5h
9h
11h
14h
20h
24h
L1
sex
Female
Male

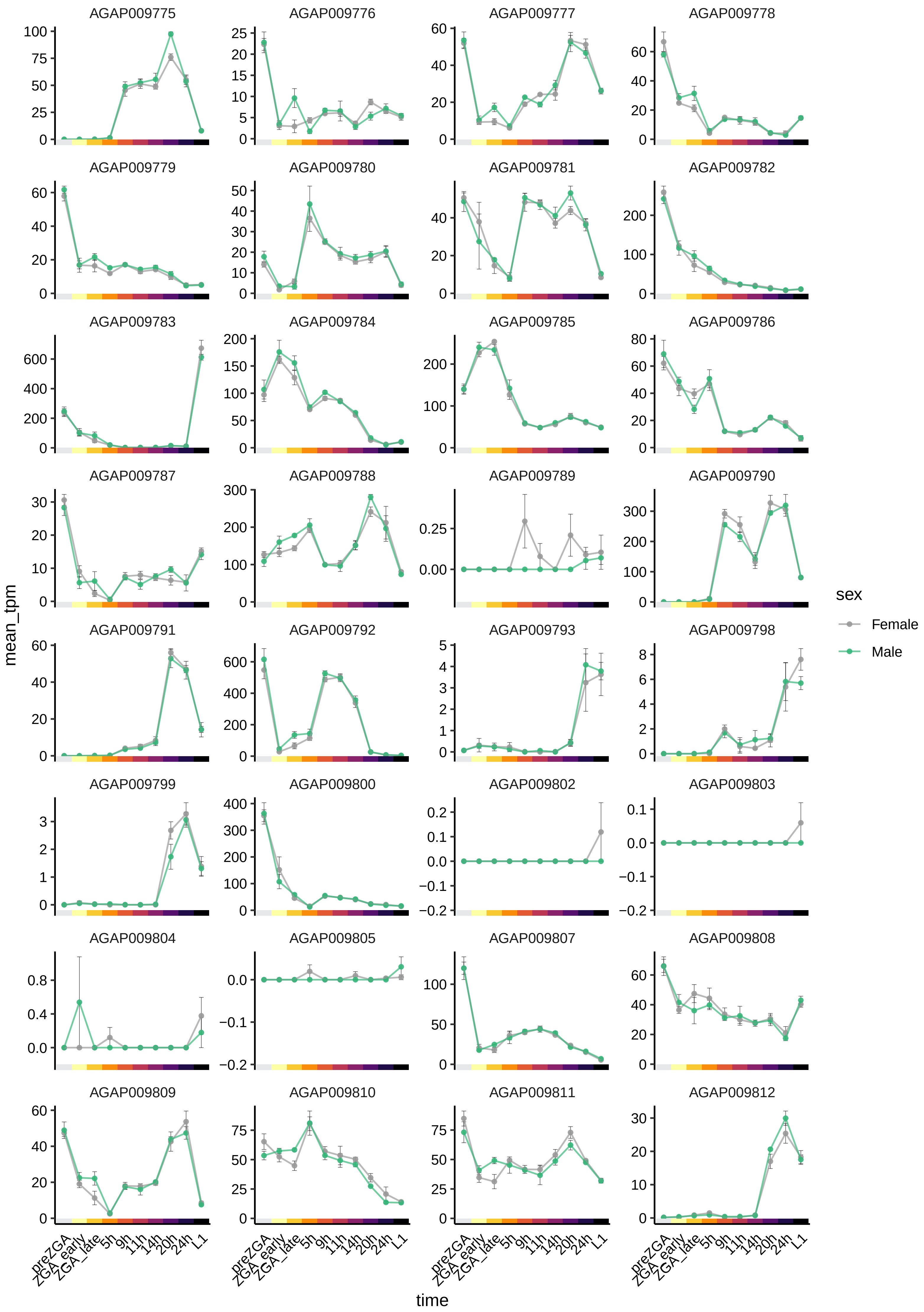

AGAP009775
AGAP009776
AGAP009777
AGAP009778
AGAP009779
AGAP009780
AGAP009781
AGAP009782
AGAP009783
AGAP009784
AGAP009785
AGAP009786
AGAP009787
AGAP009788
AGAP009789
AGAP009790
AGAP009791
AGAP009792
AGAP009793
AGAP009798
AGAP009799
AGAP009800
AGAP009802
AGAP009803
AGAP009804
AGAP009805
AGAP009807
AGAP009808
AGAP009809
AGAP009810
AGAP009811
AGAP009812
mean_tpm
time
preZGA
ZGA_early
ZGA_late
5h
9h
11h
14h
20h
24h
L1
sex
Female
Male

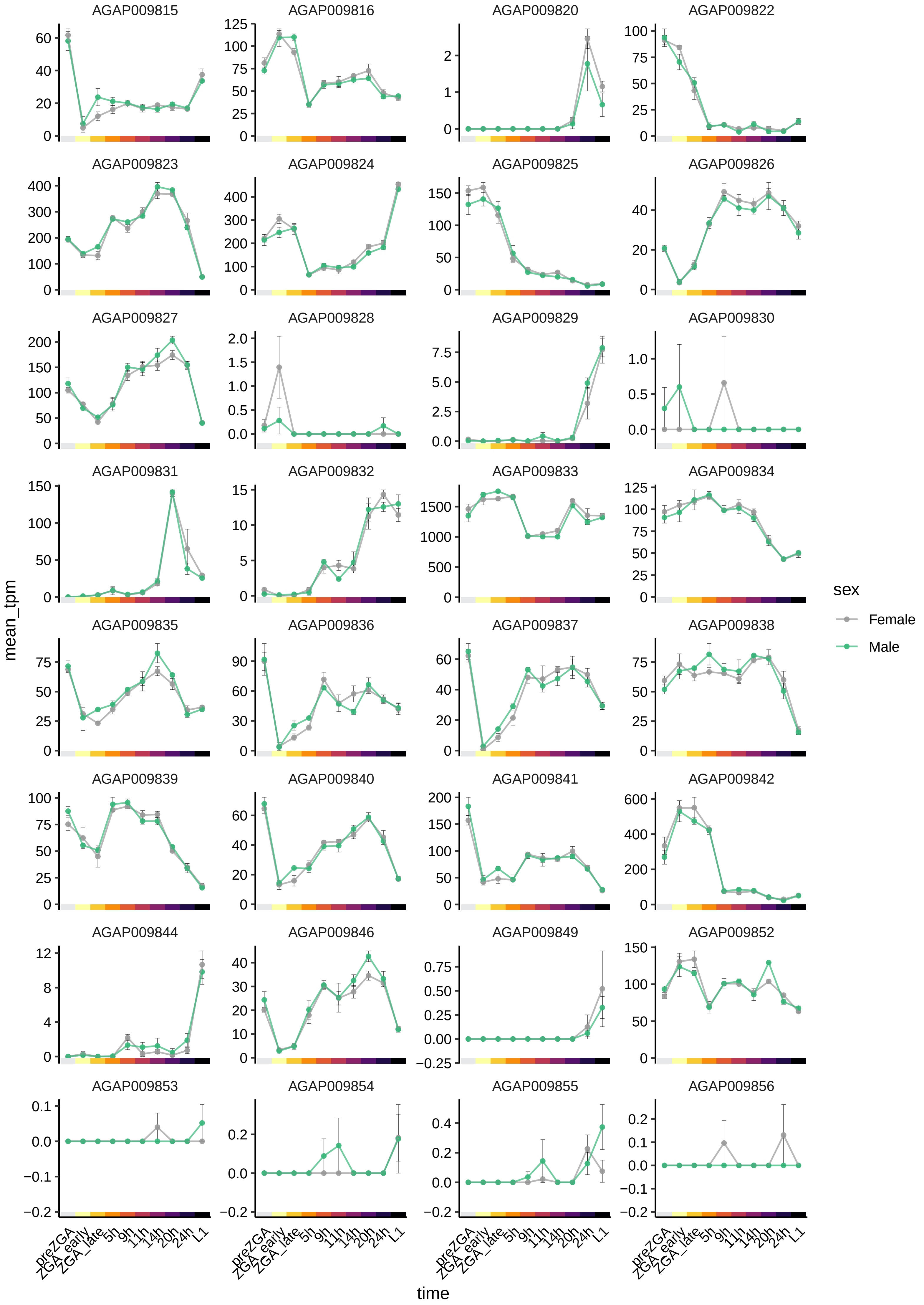

AGAP009815
AGAP009816
AGAP009820
AGAP009822
AGAP009823
AGAP009824
AGAP009825
AGAP009826
AGAP009827
AGAP009828
AGAP009829
AGAP009830
AGAP009831
AGAP009832
AGAP009833
AGAP009834
AGAP009835
AGAP009836
AGAP009837
AGAP009838
AGAP009839
AGAP009840
AGAP009841
AGAP009842
AGAP009844
AGAP009846
AGAP009849
AGAP009852
AGAP009853
AGAP009854
AGAP009855
AGAP009856
mean_tpm
time
preZGA
ZGA_early
ZGA_late
5h
9h
11h
14h
20h
24h
L1
sex
Female
Male

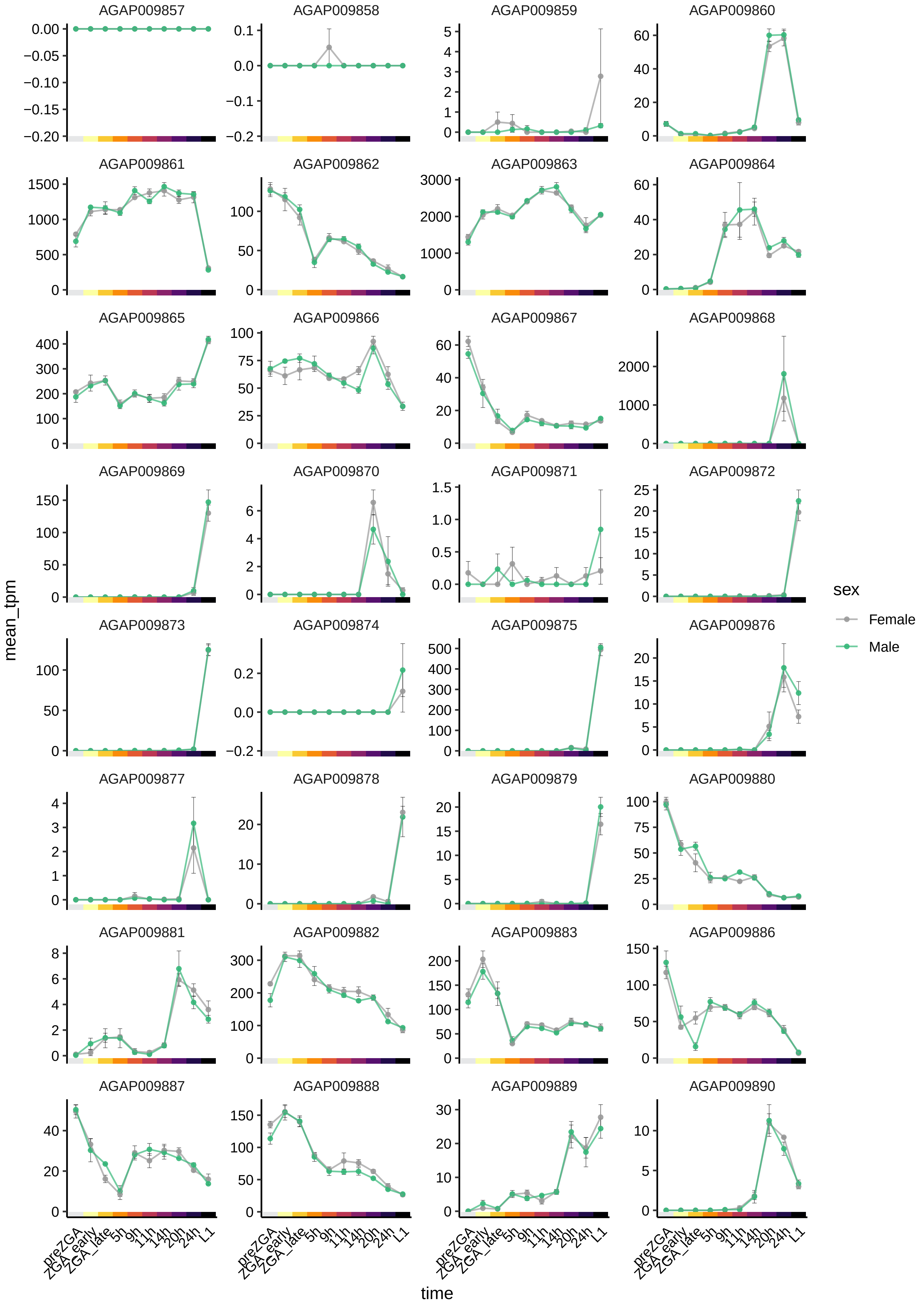

AGAP009857
AGAP009858
AGAP009859
AGAP009860
AGAP009861
AGAP009862
AGAP009863
AGAP009864
AGAP009865
AGAP009866
AGAP009867
AGAP009868
AGAP009869
AGAP009870
AGAP009871
AGAP009872
AGAP009873
AGAP009874
AGAP009875
AGAP009876
AGAP009877
AGAP009878
AGAP009879
AGAP009880
AGAP009881
AGAP009882
AGAP009883
AGAP009886
AGAP009887
AGAP009888
AGAP009889
AGAP009890
mean_tpm
time
preZGA
ZGA_early
ZGA_late
5h
9h
11h
14h
20h
24h
L1
sex
Female
Male

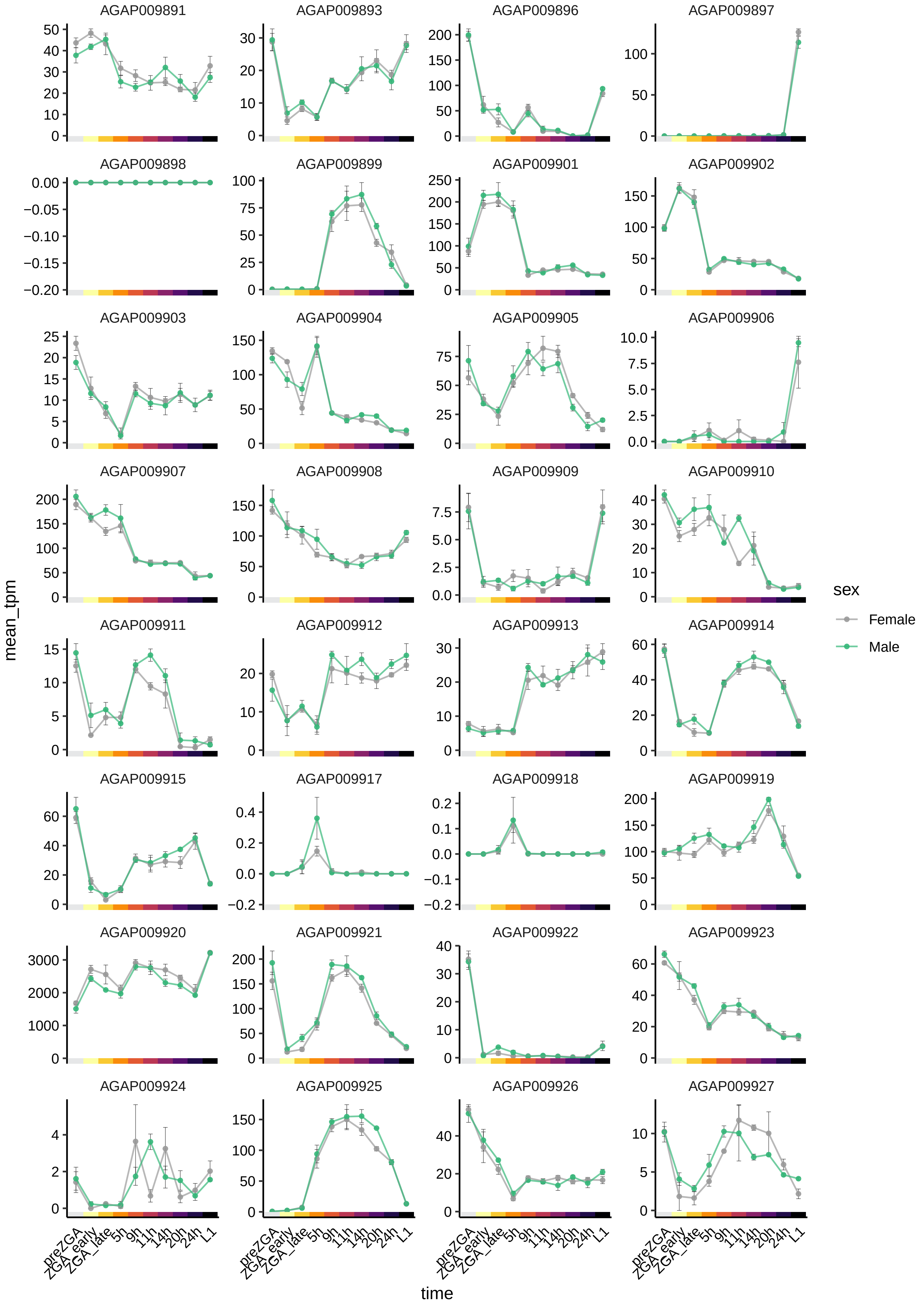
AGAP009891
AGAP009893
AGAP009896
AGAP009897
AGAP009898
AGAP009899
AGAP009901
AGAP009902
AGAP009903
AGAP009904
AGAP009905
AGAP009906
AGAP009907
AGAP009908
AGAP009909
AGAP009910
AGAP009911
AGAP009912
AGAP009913
AGAP009914
AGAP009915
AGAP009917
AGAP009918
AGAP009919
AGAP009920
AGAP009921
AGAP009922
AGAP009923
AGAP009924
AGAP009925
AGAP009926
AGAP009927
mean_tpm
time
preZGA
ZGA_early
ZGA_late
5h
9h
11h
14h
20h
24h
L1
sex
Female
Male

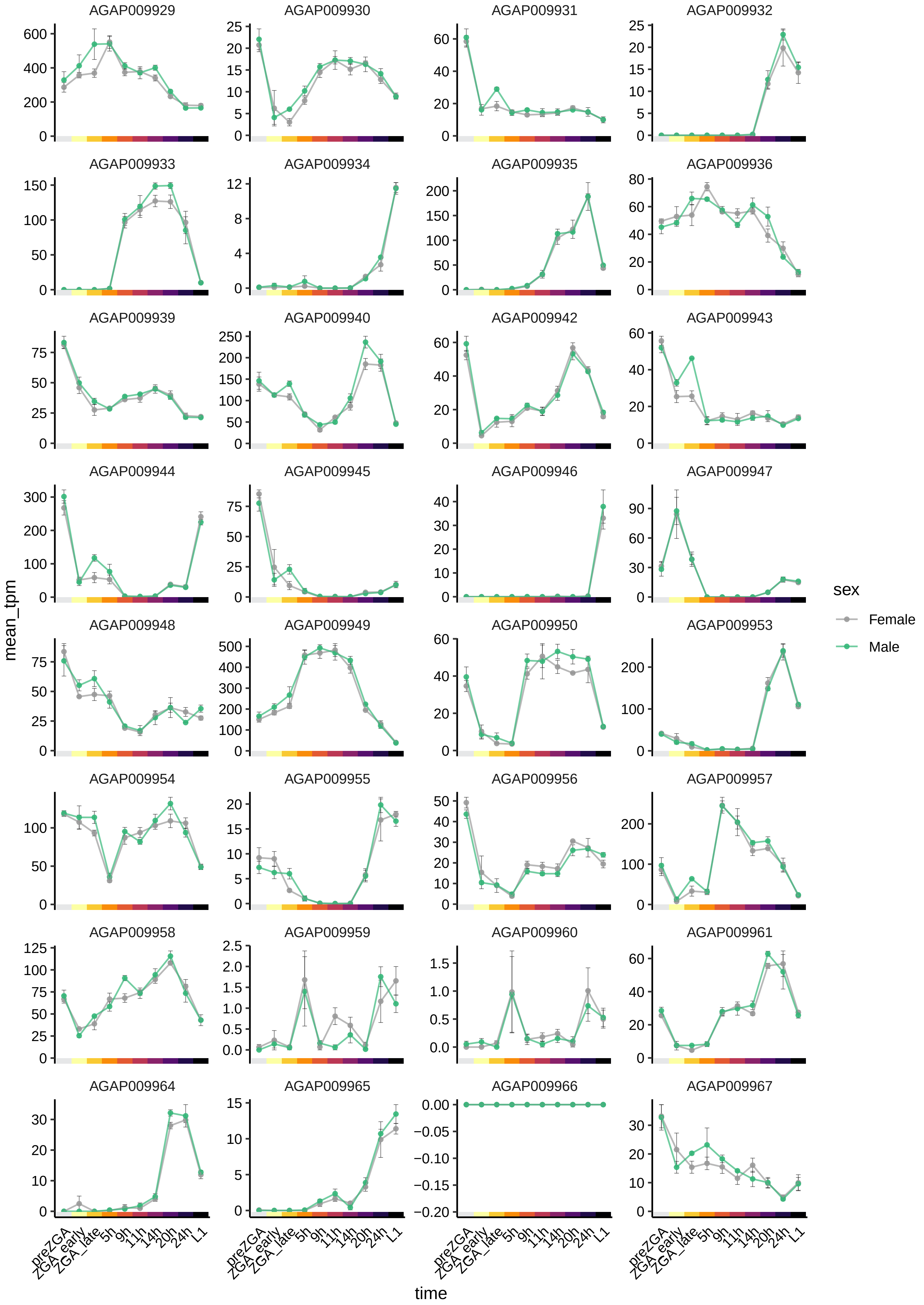
AGAP009929
AGAP009930
AGAP009931
AGAP009932
AGAP009933
AGAP009934
AGAP009935
AGAP009936
AGAP009939
AGAP009940
AGAP009942
AGAP009943
AGAP009944
AGAP009945
AGAP009946
AGAP009947
AGAP009948
AGAP009949
AGAP009950
AGAP009953
AGAP009954
AGAP009955
AGAP009956
AGAP009957
AGAP009958
AGAP009959
AGAP009960
AGAP009961
AGAP009964
AGAP009965
AGAP009966
AGAP009967
mean_tpm
time
preZGA
ZGA_early
ZGA_late
5h
9h
11h
14h
20h
24h
L1
sex
Female
Male

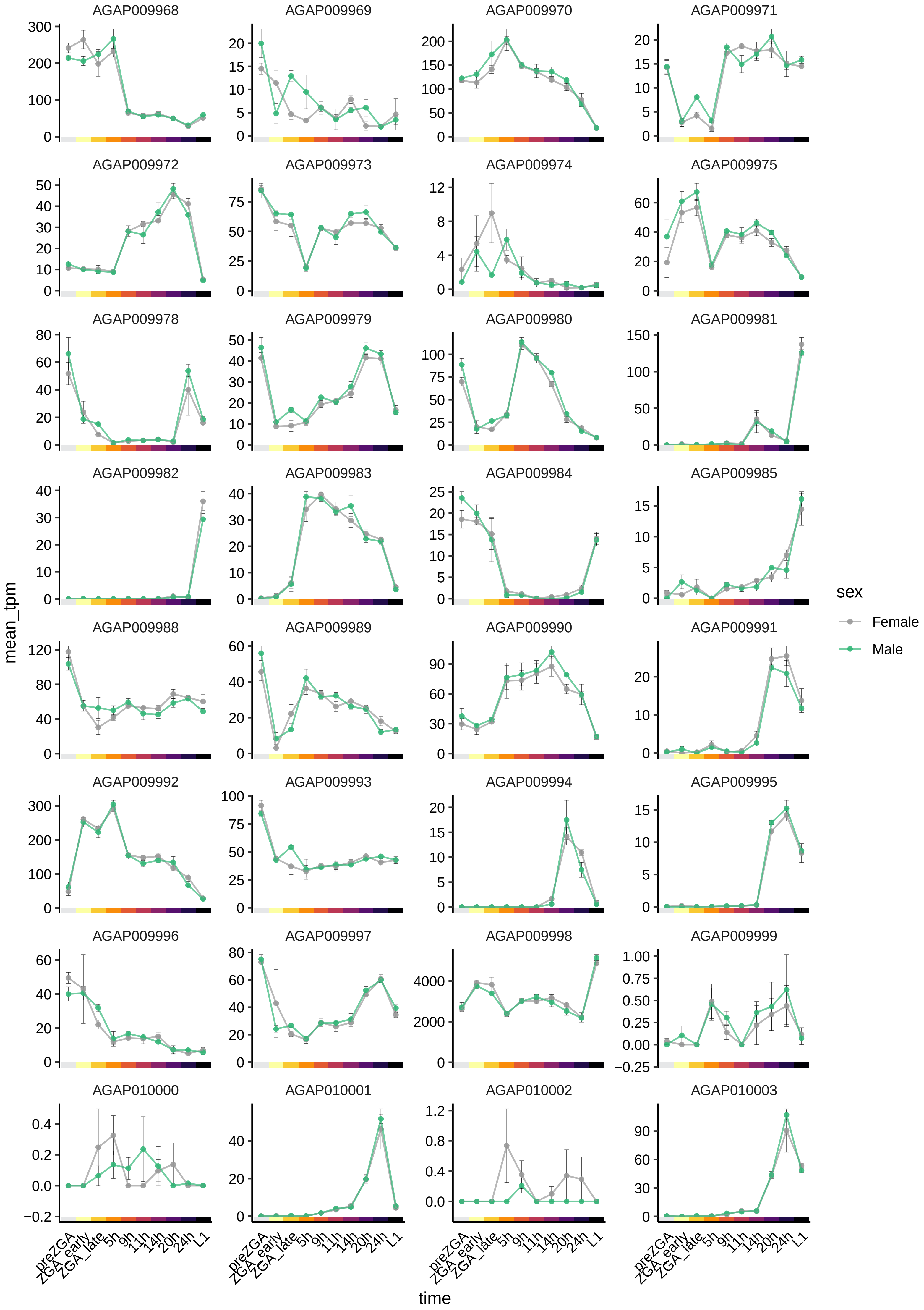

AGAP009968
AGAP009969
AGAP009970
AGAP009971
AGAP009972
AGAP009973
AGAP009974
AGAP009975
AGAP009978
AGAP009979
AGAP009980
AGAP009981
AGAP009982
AGAP009983
AGAP009984
AGAP009985
AGAP009988
AGAP009989
AGAP009990
AGAP009991
AGAP009992
AGAP009993
AGAP009994
AGAP009995
AGAP009996
AGAP009997
AGAP009998
AGAP009999
AGAP010000
AGAP010001
AGAP010002
AGAP010003
mean_tpm
time
preZGA
ZGA_early
ZGA_late
5h
9h
11h
14h
20h
24h
L1
sex
Female
Male

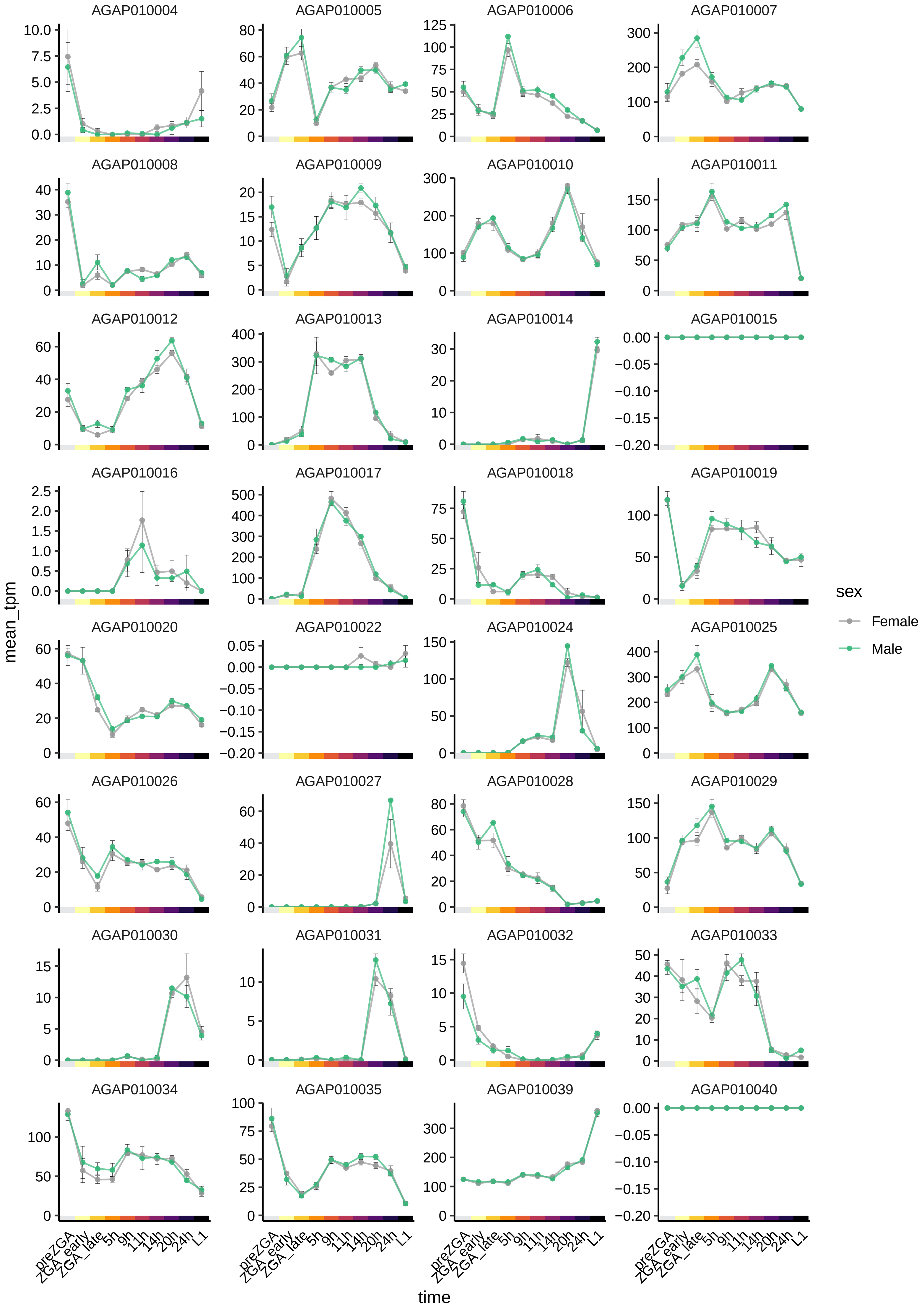
AGAP010004
AGAP010005
AGAP010006
AGAP010007
AGAP010008
AGAP010009
AGAP010010
AGAP010011
AGAP010012
AGAP010013
AGAP010014
AGAP010015
AGAP010016
AGAP010017
AGAP010018
AGAP010019
AGAP010020
AGAP010022
AGAP010024
AGAP010025
AGAP010026
AGAP010027
AGAP010028
AGAP010029
AGAP010030
AGAP010031
AGAP010032
AGAP010033
AGAP010034
AGAP010035
AGAP010039
AGAP010040
mean_tpm
time
preZGA
ZGA_early
ZGA_late
5h
9h
11h
14h
20h
24h
L1
sex
Female
Male

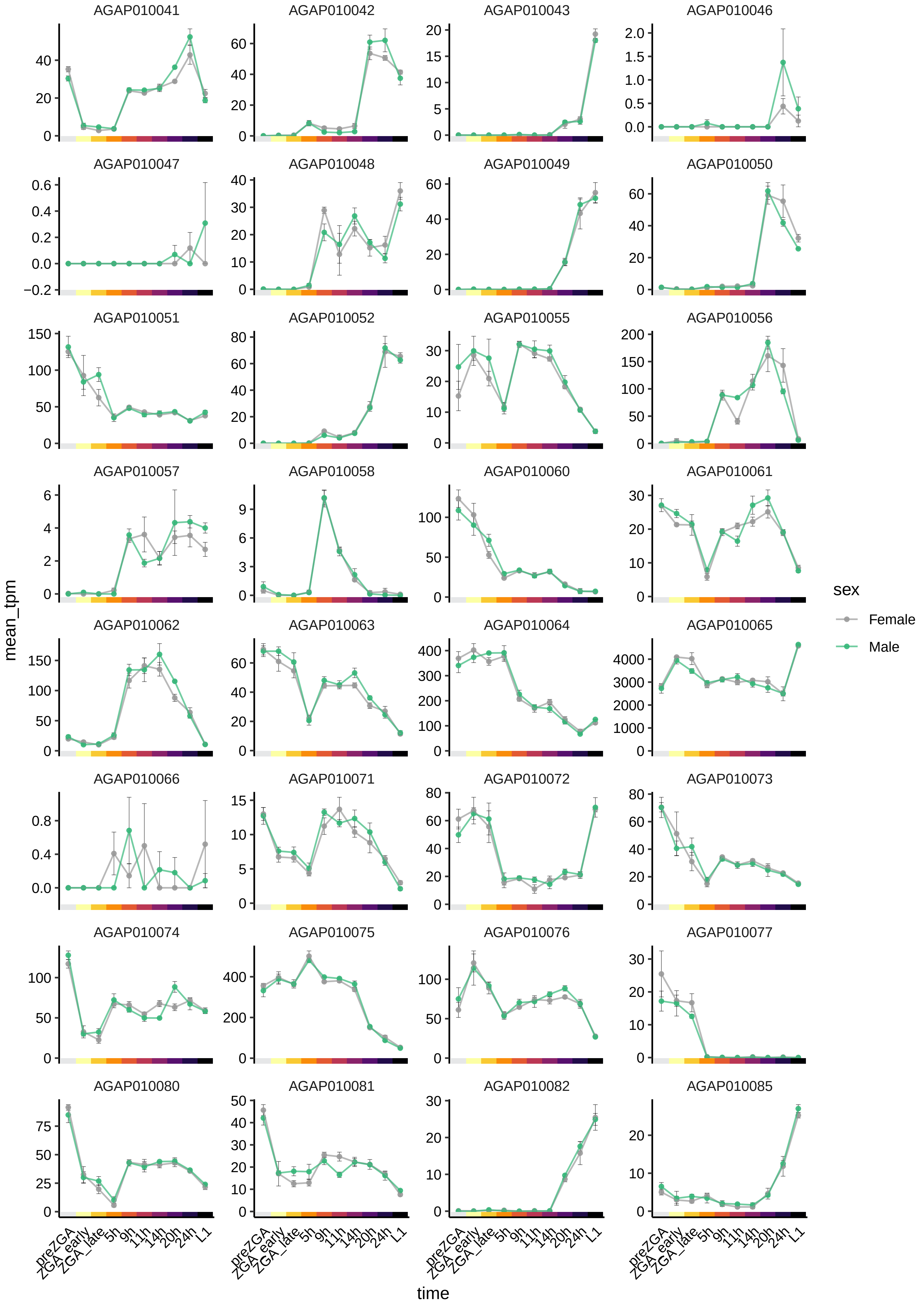

AGAP010041
AGAP010042
AGAP010043
AGAP010046
AGAP010047
AGAP010048
AGAP010049
AGAP010050
AGAP010051
AGAP010052
AGAP010055
AGAP010056
AGAP010057
AGAP010058
AGAP010060
AGAP010061
AGAP010062
AGAP010063
AGAP010064
AGAP010065
AGAP010066
AGAP010071
AGAP010072
AGAP010073
AGAP010074
AGAP010075
AGAP010076
AGAP010077
AGAP010080
AGAP010081
AGAP010082
AGAP010085
mean_tpm
time
preZGA
ZGA_early
ZGA_late
5h
9h
11h
14h
20h
24h
L1
sex
Female
Male

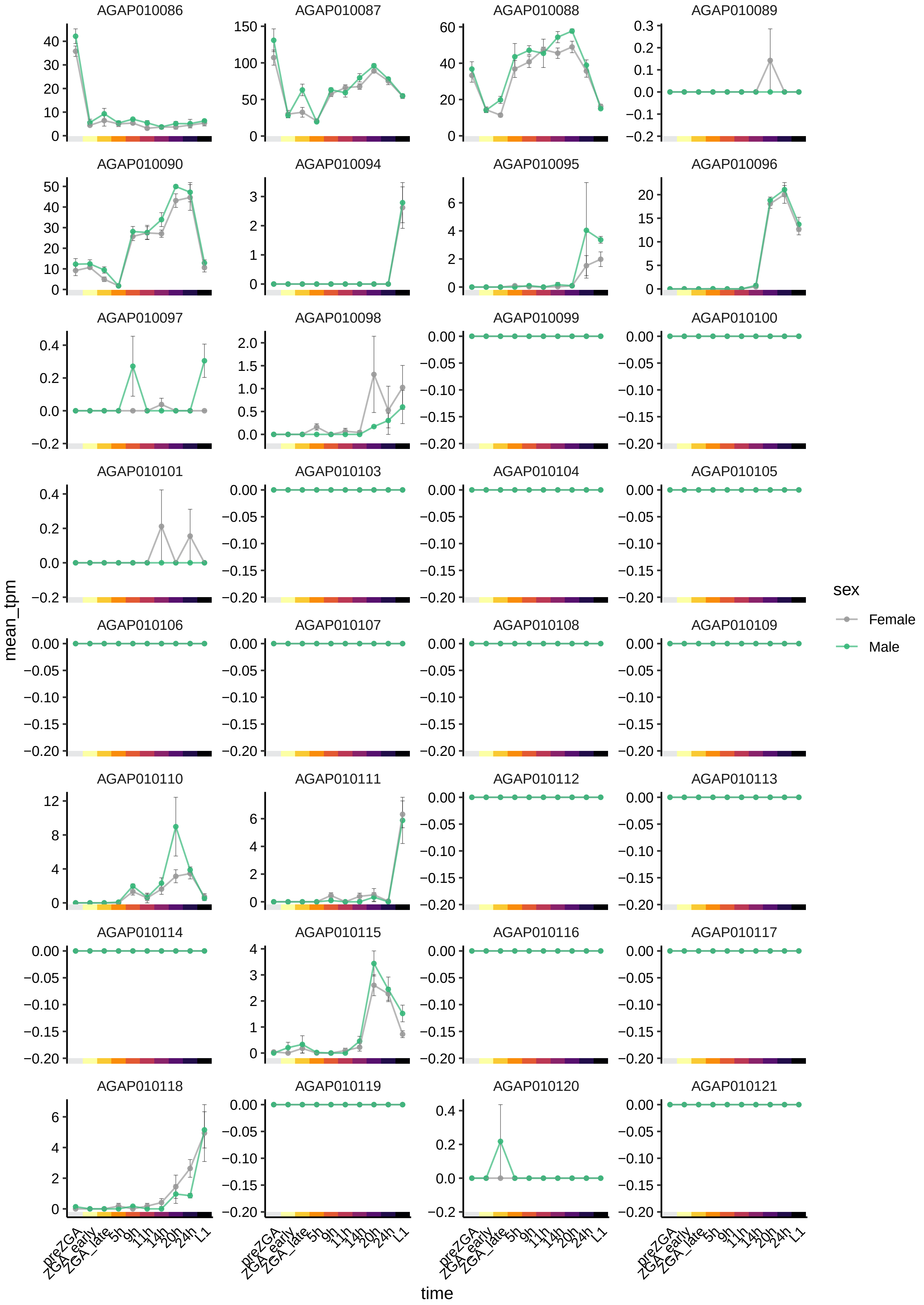

AGAP010086
AGAP010087
AGAP010088
AGAP010089
AGAP010090
AGAP010094
AGAP010095
AGAP010096
AGAP010097
AGAP010098
AGAP010099
AGAP010100
AGAP010101
AGAP010103
AGAP010104
AGAP010105
AGAP010106
AGAP010107
AGAP010108
AGAP010109
AGAP010110
AGAP010111
AGAP010112
AGAP010113
AGAP010114
AGAP010115
AGAP010116
AGAP010117
AGAP010118
AGAP010119
AGAP010120
AGAP010121
mean_tpm
time
preZGA
ZGA_early
ZGA_late
5h
9h
11h
14h
20h
24h
L1
sex
Female
Male

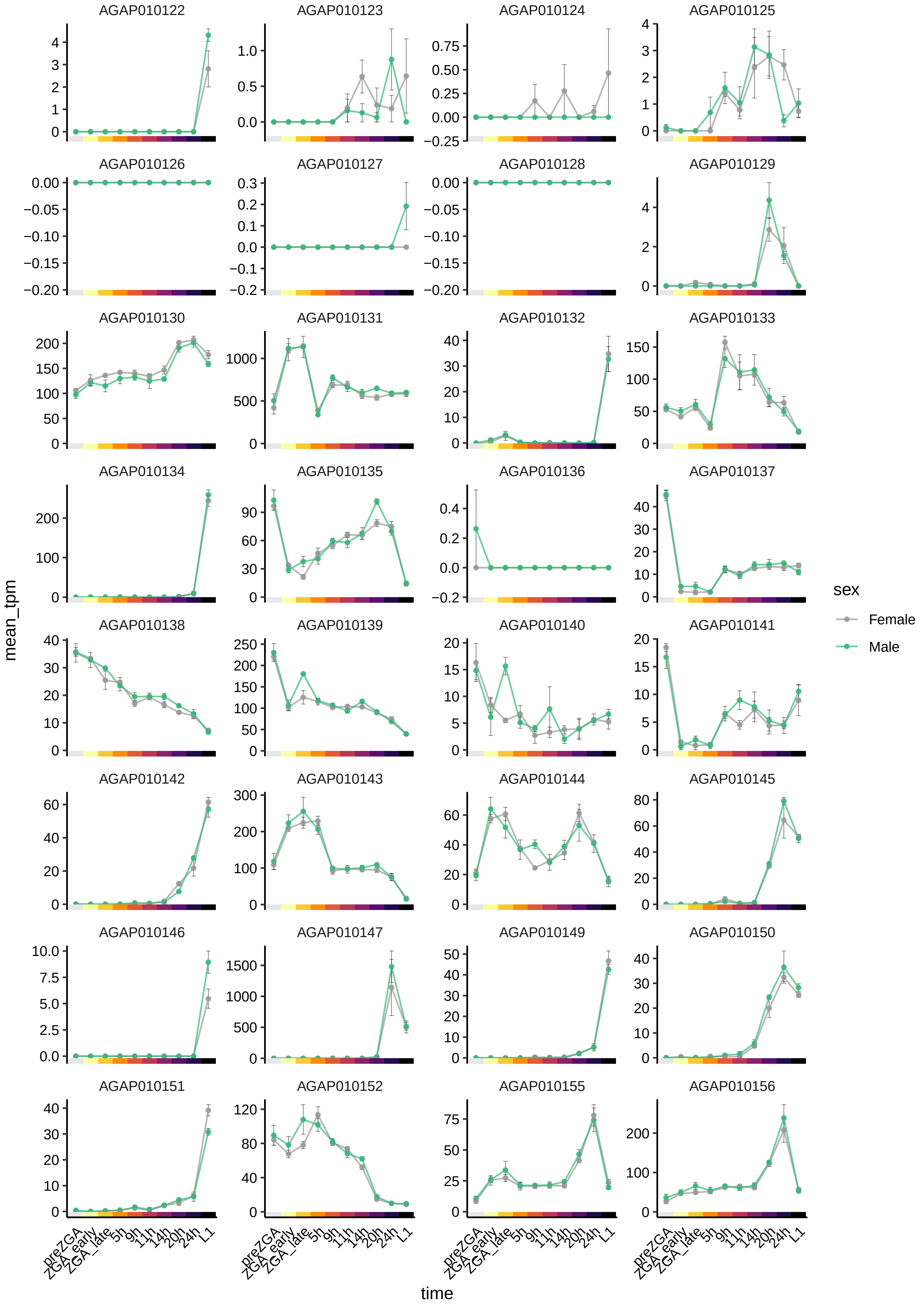

AGAP010122
AGAP010123
AGAP010124
AGAP010125
AGAP010126
AGAP010127
AGAP010128
AGAP010129
AGAP010130
AGAP010131
AGAP010132
AGAP010133
AGAP010134
AGAP010135
AGAP010136
AGAP010137
AGAP010138
AGAP010139
AGAP010140
AGAP010141
AGAP010142
AGAP010143
AGAP010144
AGAP010145
AGAP010146
AGAP010147
AGAP010149
AGAP010150
AGAP010151
AGAP010152
AGAP010155
AGAP010156
mean_tpm
time
preZGA
ZGA_early
ZGA_late
5h
9h
11h
14h
20h
24h
L1
sex
Female
Male

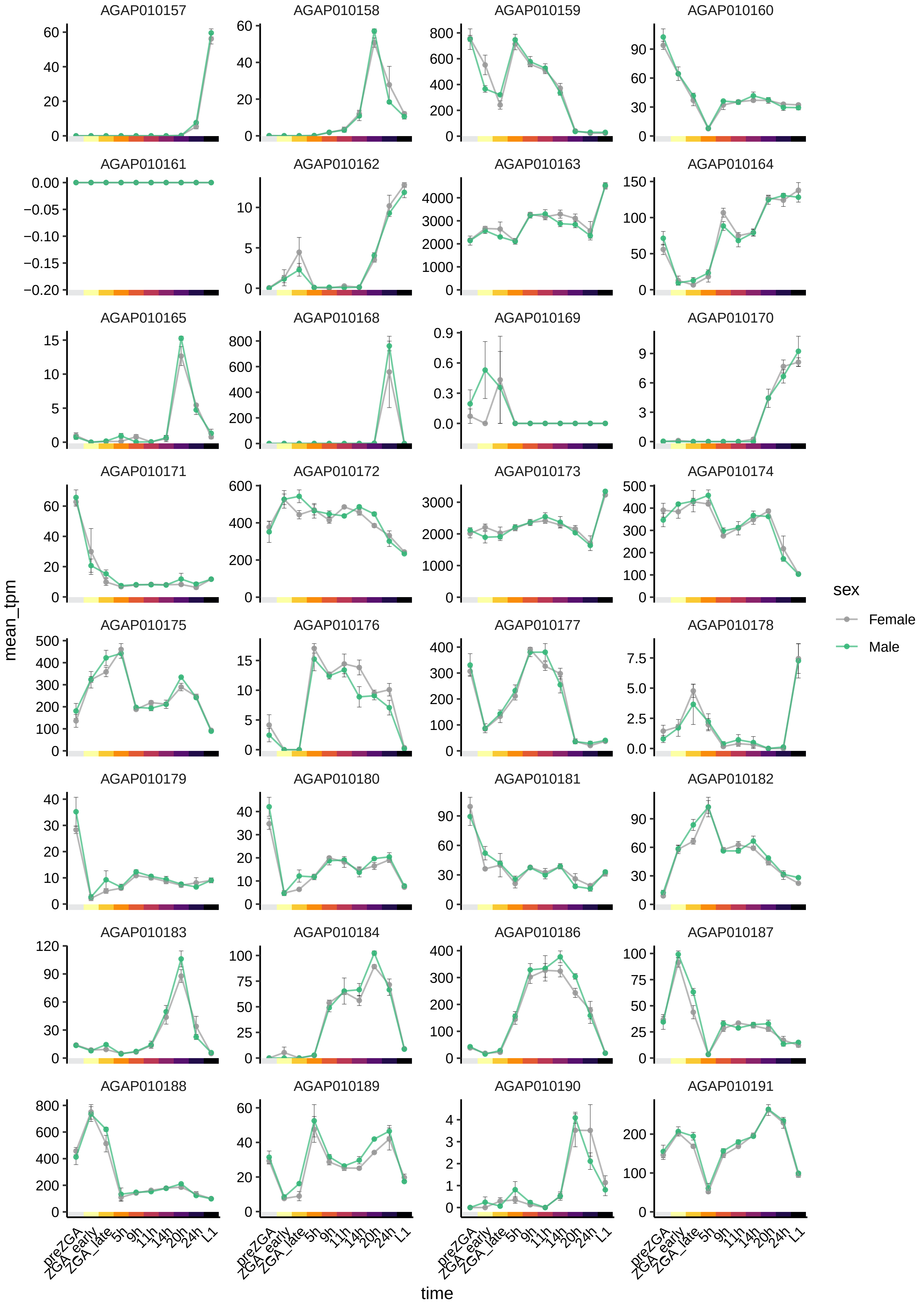
AGAP010157
AGAP010158
AGAP010159
AGAP010160
AGAP010161
AGAP010162
AGAP010163
AGAP010164
AGAP010165
AGAP010168
AGAP010169
AGAP010170
AGAP010171
AGAP010172
AGAP010173
AGAP010174
AGAP010175
AGAP010176
AGAP010177
AGAP010178
AGAP010179
AGAP010180
AGAP010181
AGAP010182
AGAP010183
AGAP010184
AGAP010186
AGAP010187
AGAP010188
AGAP010189
AGAP010190
AGAP010191
mean_tpm
time
preZGA
ZGA_early
ZGA_late
5h
9h
11h
14h
20h
24h
L1
sex
Female
Male

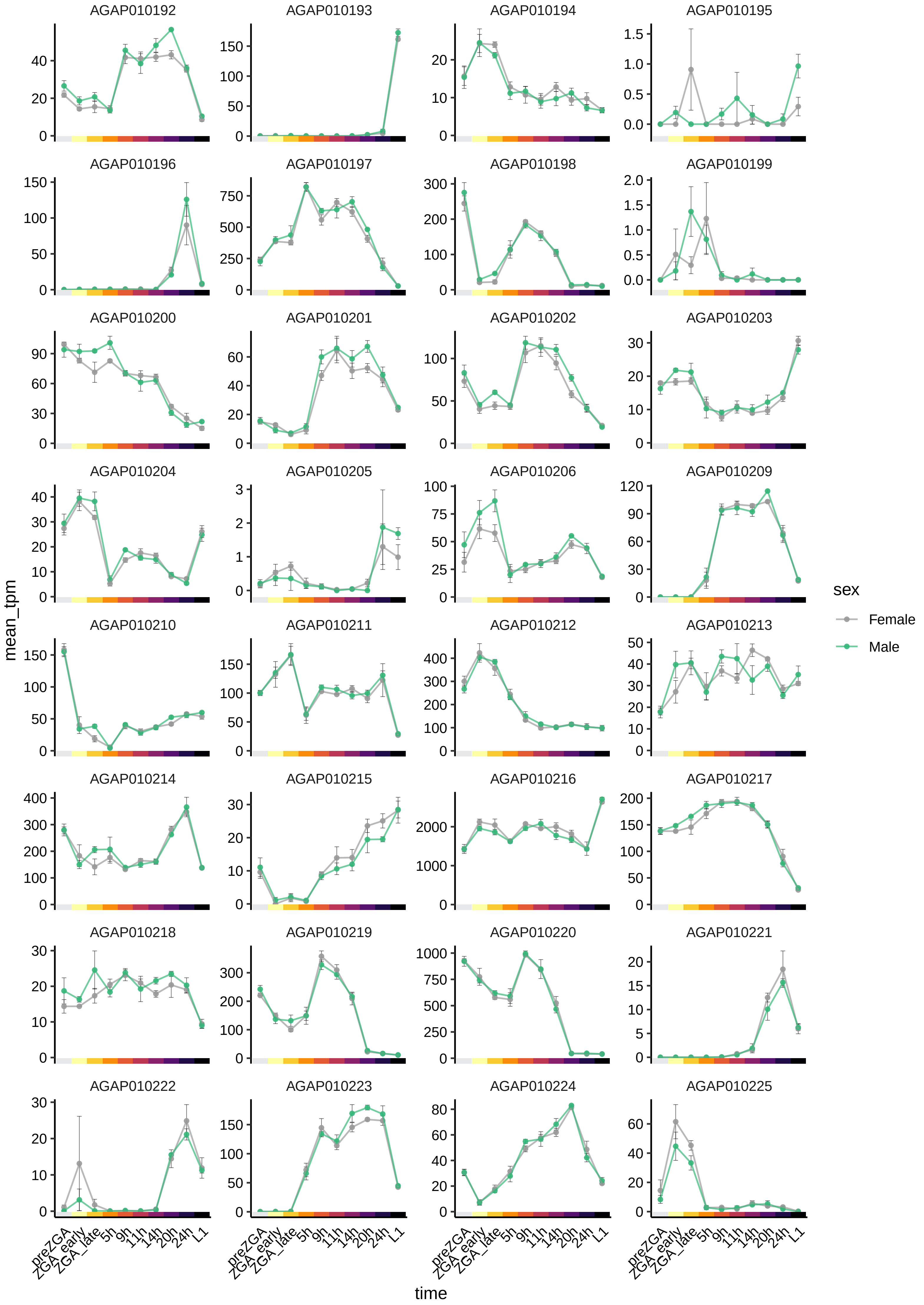

AGAP010192
AGAP010193
AGAP010194
AGAP010195
AGAP010196
AGAP010197
AGAP010198
AGAP010199
AGAP010200
AGAP010201
AGAP010202
AGAP010203
AGAP010204
AGAP010205
AGAP010206
AGAP010209
AGAP010210
AGAP010211
AGAP010212
AGAP010213
AGAP010214
AGAP010215
AGAP010216
AGAP010217
AGAP010218
AGAP010219
AGAP010220
AGAP010221
AGAP010222
AGAP010223
AGAP010224
AGAP010225
mean_tpm
time
preZGA
ZGA_early
ZGA_late
5h
9h
11h
14h
20h
24h
L1
sex
Female
Male

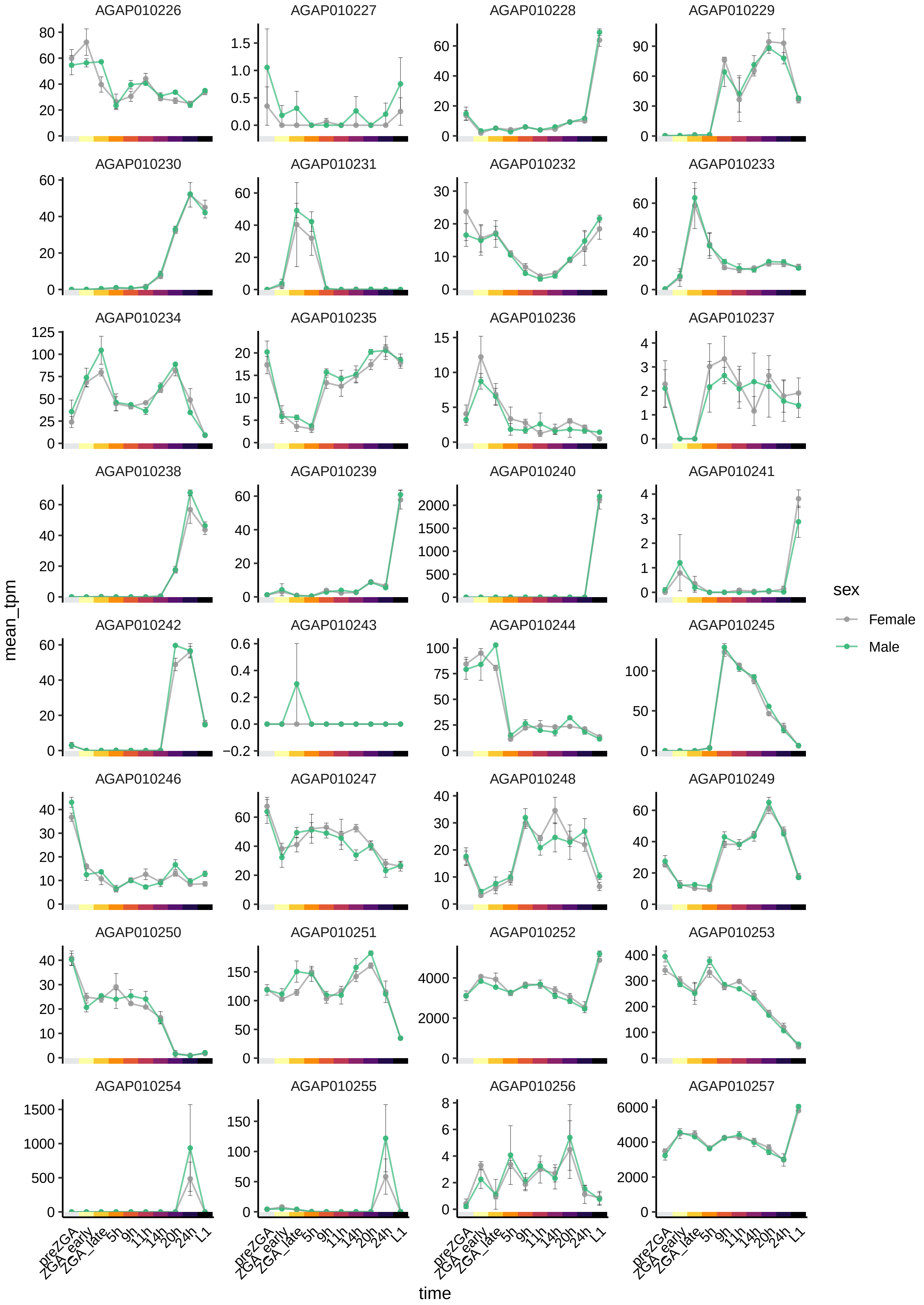
AGAP010226
AGAP010227
AGAP010228
AGAP010229
AGAP010230
AGAP010231
AGAP010232
AGAP010233
AGAP010234
AGAP010235
AGAP010236
AGAP010237
AGAP010238
AGAP010239
AGAP010240
AGAP010241
AGAP010242
AGAP010243
AGAP010244
AGAP010245
AGAP010246
AGAP010247
AGAP010248
AGAP010249
AGAP010250
AGAP010251
AGAP010252
AGAP010253
AGAP010254
AGAP010255
AGAP010256
AGAP010257
mean_tpm
time
preZGA
ZGA_early
ZGA_late
5h
9h
11h
14h
20h
24h
L1
sex
Female
Male

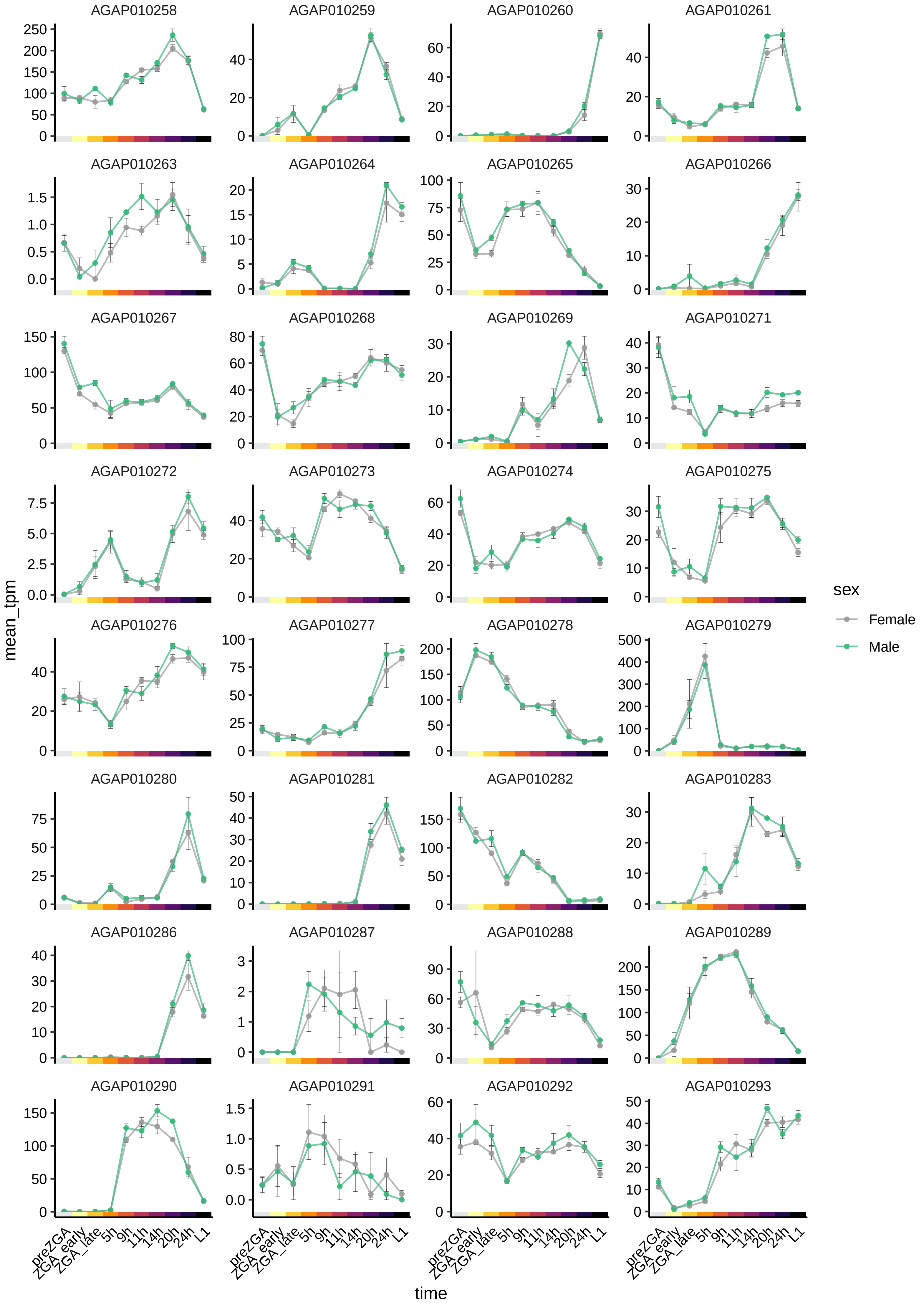
AGAP010258
AGAP010259
AGAP010260
AGAP010261
AGAP010263
AGAP010264
AGAP010265
AGAP010266
AGAP010267
AGAP010268
AGAP010269
AGAP010271
AGAP010272
AGAP010273
AGAP010274
AGAP010275
AGAP010276
AGAP010277
AGAP010278
AGAP010279
AGAP010280
AGAP010281
AGAP010282
AGAP010283
AGAP010286
AGAP010287
AGAP010288
AGAP010289
AGAP010290
AGAP010291
AGAP010292
AGAP010293
mean_tpm
time
preZGA
ZGA_early
ZGA_late
5h
9h
11h
14h
20h
24h
L1
sex
Female
Male

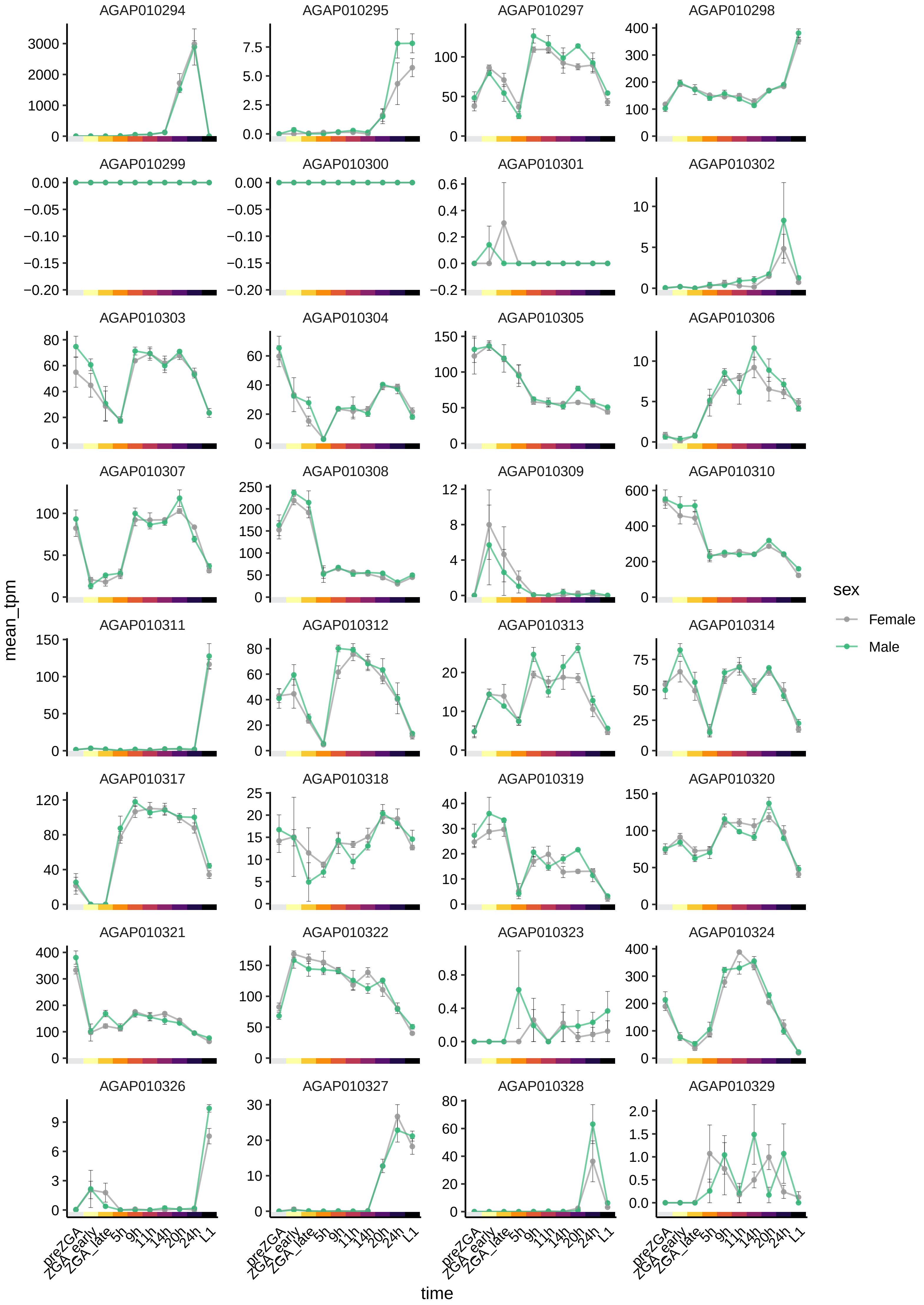

AGAP010294
AGAP010295
AGAP010297
AGAP010298
AGAP010299
AGAP010300
AGAP010301
AGAP010302
AGAP010303
AGAP010304
AGAP010305
AGAP010306
AGAP010307
AGAP010308
AGAP010309
AGAP010310
AGAP010311
AGAP010312
AGAP010313
AGAP010314
AGAP010317
AGAP010318
AGAP010319
AGAP010320
AGAP010321
AGAP010322
AGAP010323
AGAP010324
AGAP010326
AGAP010327
AGAP010328
AGAP010329
mean_tpm
time
preZGA
ZGA_early
ZGA_late
5h
9h
11h
14h
20h
24h
L1
sex
Female
Male

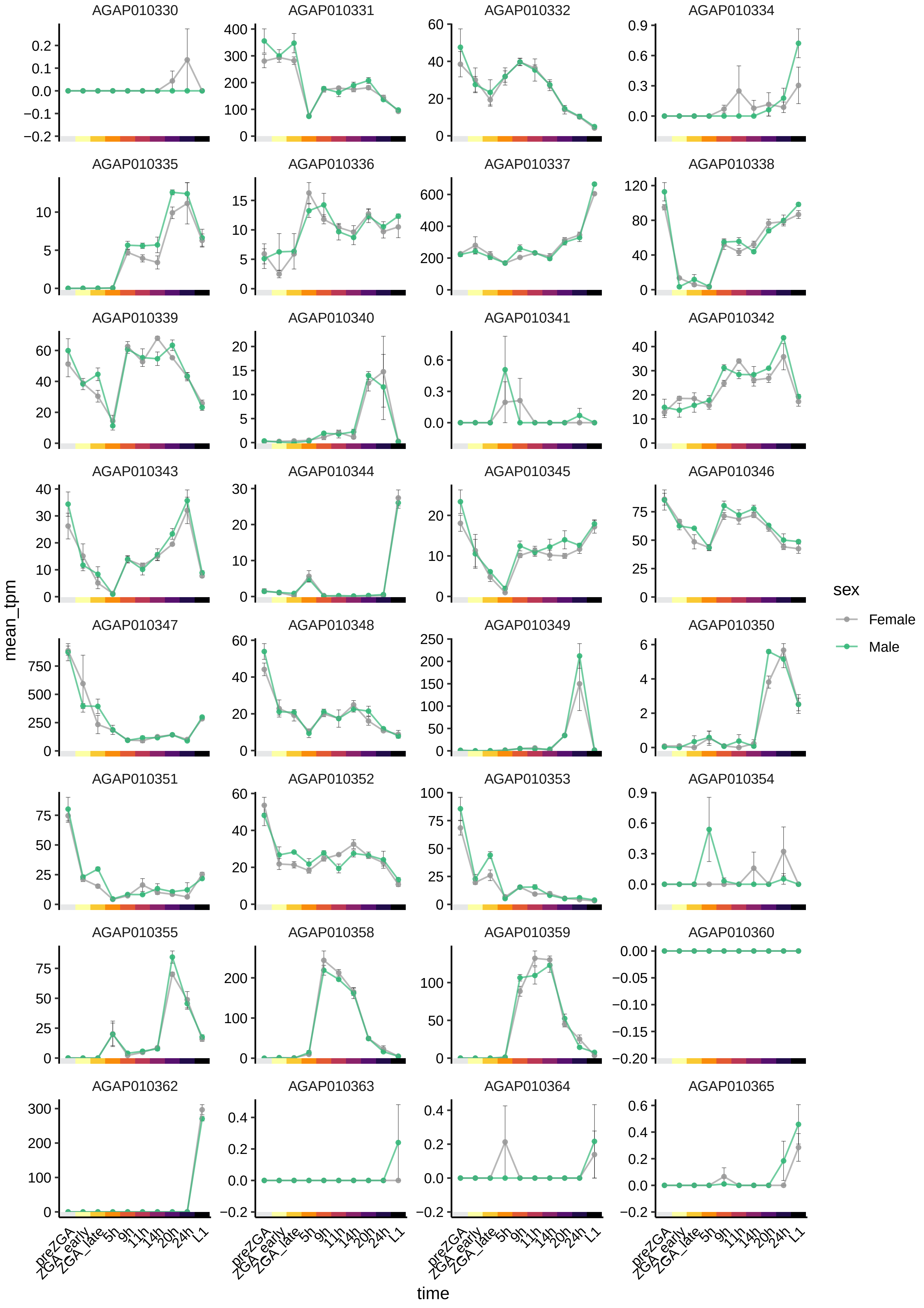
AGAP010330
AGAP010331
AGAP010332
AGAP010334
AGAP010335
AGAP010336
AGAP010337
AGAP010338
AGAP010339
AGAP010340
AGAP010341
AGAP010342
AGAP010343
AGAP010344
AGAP010345
AGAP010346
AGAP010347
AGAP010348
AGAP010349
AGAP010350
AGAP010351
AGAP010352
AGAP010353
AGAP010354
AGAP010355
AGAP010358
AGAP010359
AGAP010360
AGAP010362
AGAP010363
AGAP010364
AGAP010365
mean_tpm
time
preZGA
ZGA_early
ZGA_late
5h
9h
11h
14h
20h
24h
L1
sex
Female
Male

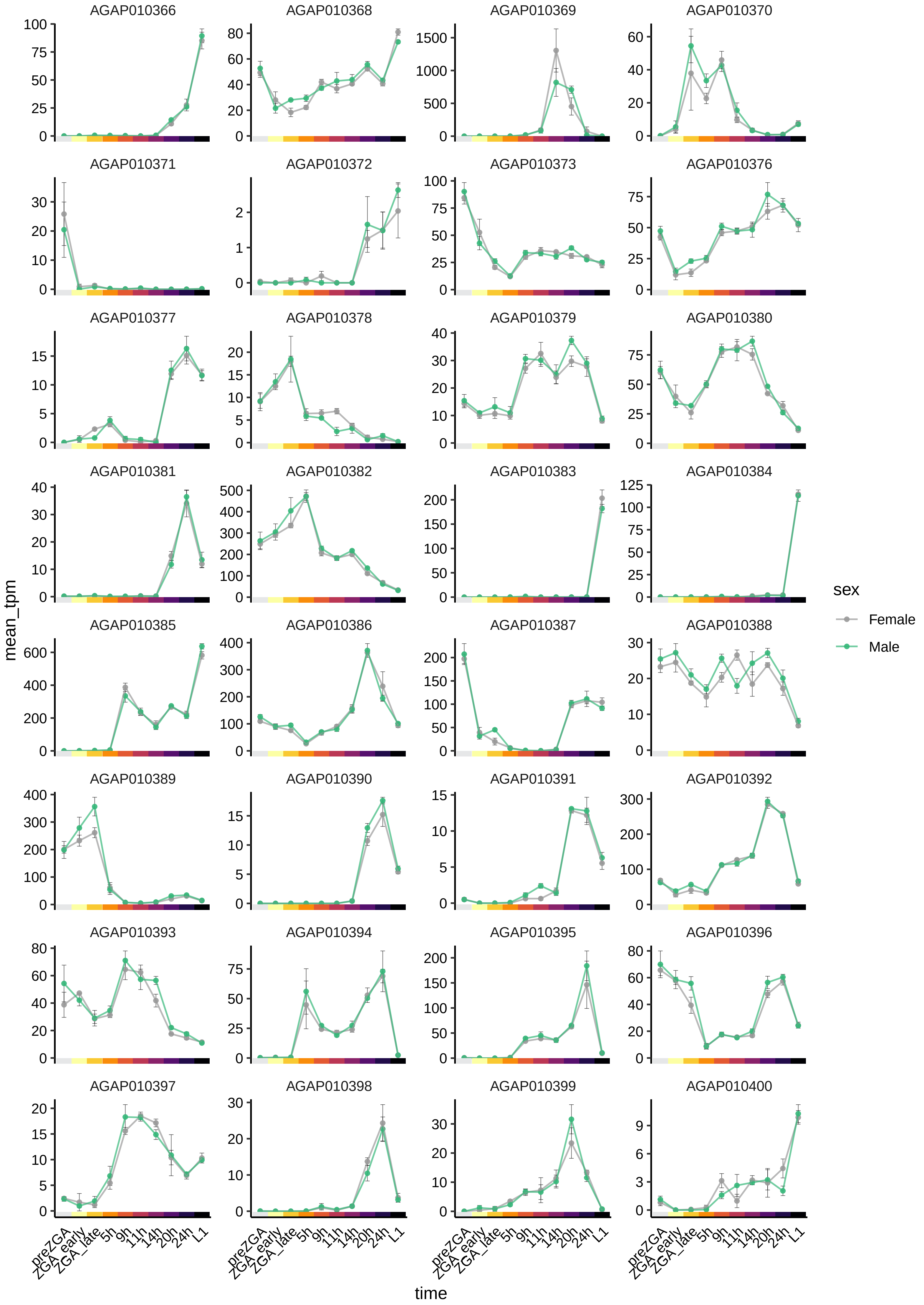

AGAP010366
AGAP010368
AGAP010369
AGAP010370
AGAP010371
AGAP010372
AGAP010373
AGAP010376
AGAP010377
AGAP010378
AGAP010379
AGAP010380
AGAP010381
AGAP010382
AGAP010383
AGAP010384
AGAP010385
AGAP010386
AGAP010387
AGAP010388
AGAP010389
AGAP010390
AGAP010391
AGAP010392
AGAP010393
AGAP010394
AGAP010395
AGAP010396
AGAP010397
AGAP010398
AGAP010399
AGAP010400
mean_tpm
time
preZGA
ZGA_early
ZGA_late
5h
9h
11h
14h
20h
24h
L1
sex
Female
Male

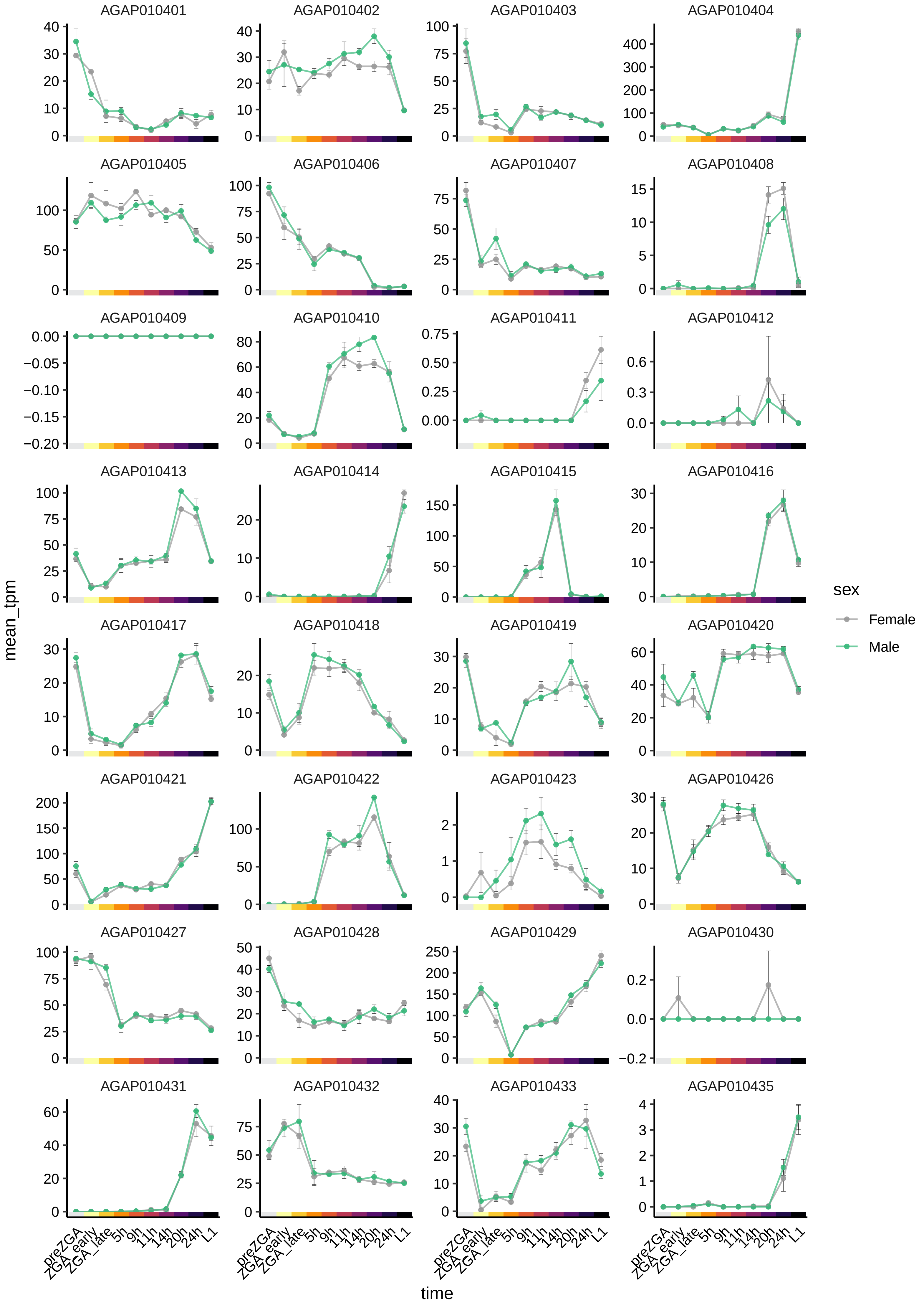
AGAP010401
AGAP010402
AGAP010403
AGAP010404
AGAP010405
AGAP010406
AGAP010407
AGAP010408
AGAP010409
AGAP010410
AGAP010411
AGAP010412
AGAP010413
AGAP010414
AGAP010415
AGAP010416
AGAP010417
AGAP010418
AGAP010419
AGAP010420
AGAP010421
AGAP010422
AGAP010423
AGAP010426
AGAP010427
AGAP010428
AGAP010429
AGAP010430
AGAP010431
AGAP010432
AGAP010433
AGAP010435
mean_tpm
time
preZGA
ZGA_early
ZGA_late
5h
9h
11h
14h
20h
24h
L1
sex
Female
Male

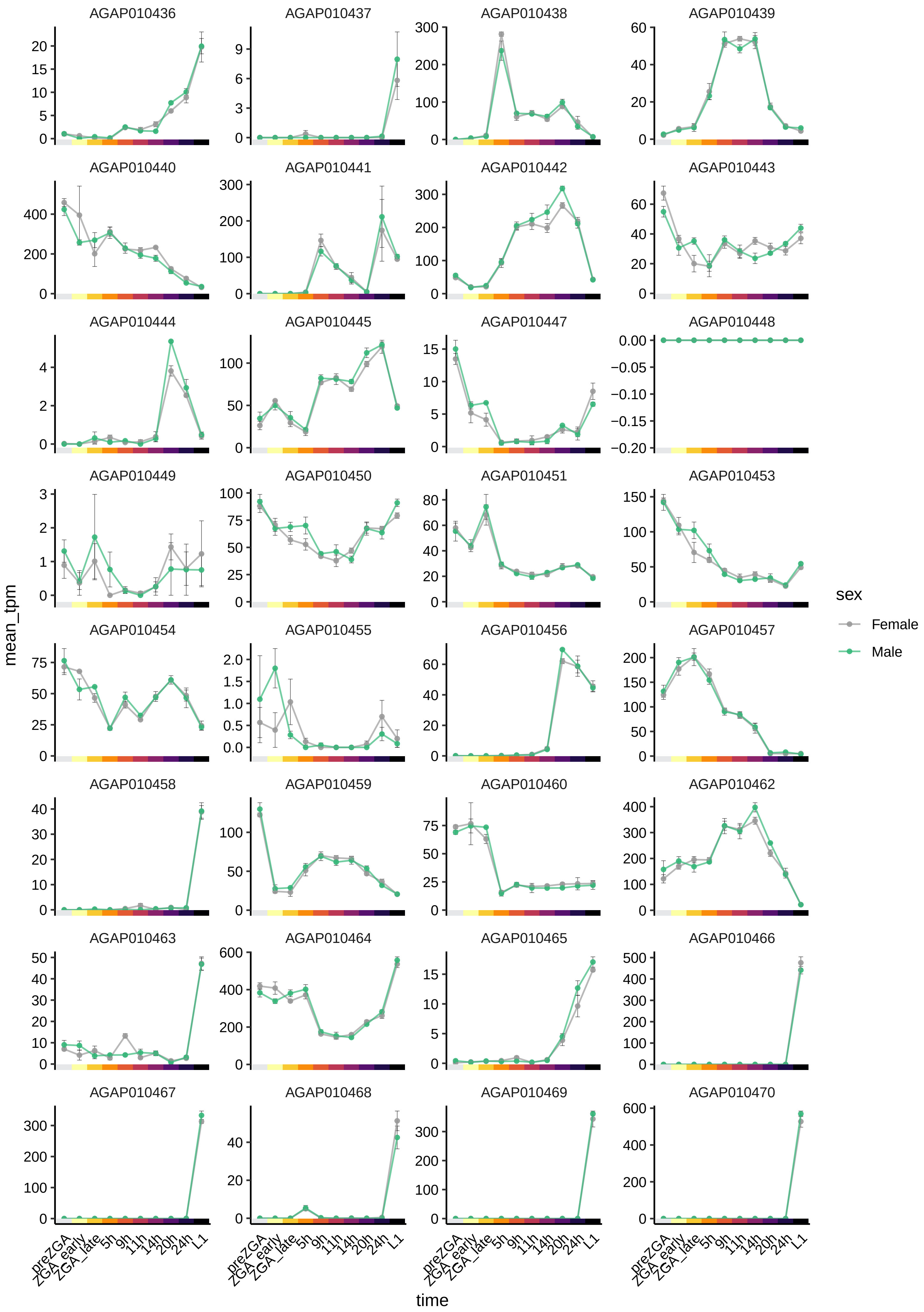
AGAP010436
AGAP010437
AGAP010438
AGAP010439
AGAP010440
AGAP010441
AGAP010442
AGAP010443
AGAP010444
AGAP010445
AGAP010447
AGAP010448
AGAP010449
AGAP010450
AGAP010451
AGAP010453
AGAP010454
AGAP010455
AGAP010456
AGAP010457
AGAP010458
AGAP010459
AGAP010460
AGAP010462
AGAP010463
AGAP010464
AGAP010465
AGAP010466
AGAP010467
AGAP010468
AGAP010469
AGAP010470
mean_tpm
time
preZGA
ZGA_early
ZGA_late
5h
9h
11h
14h
20h
24h
L1
sex
Female
Male

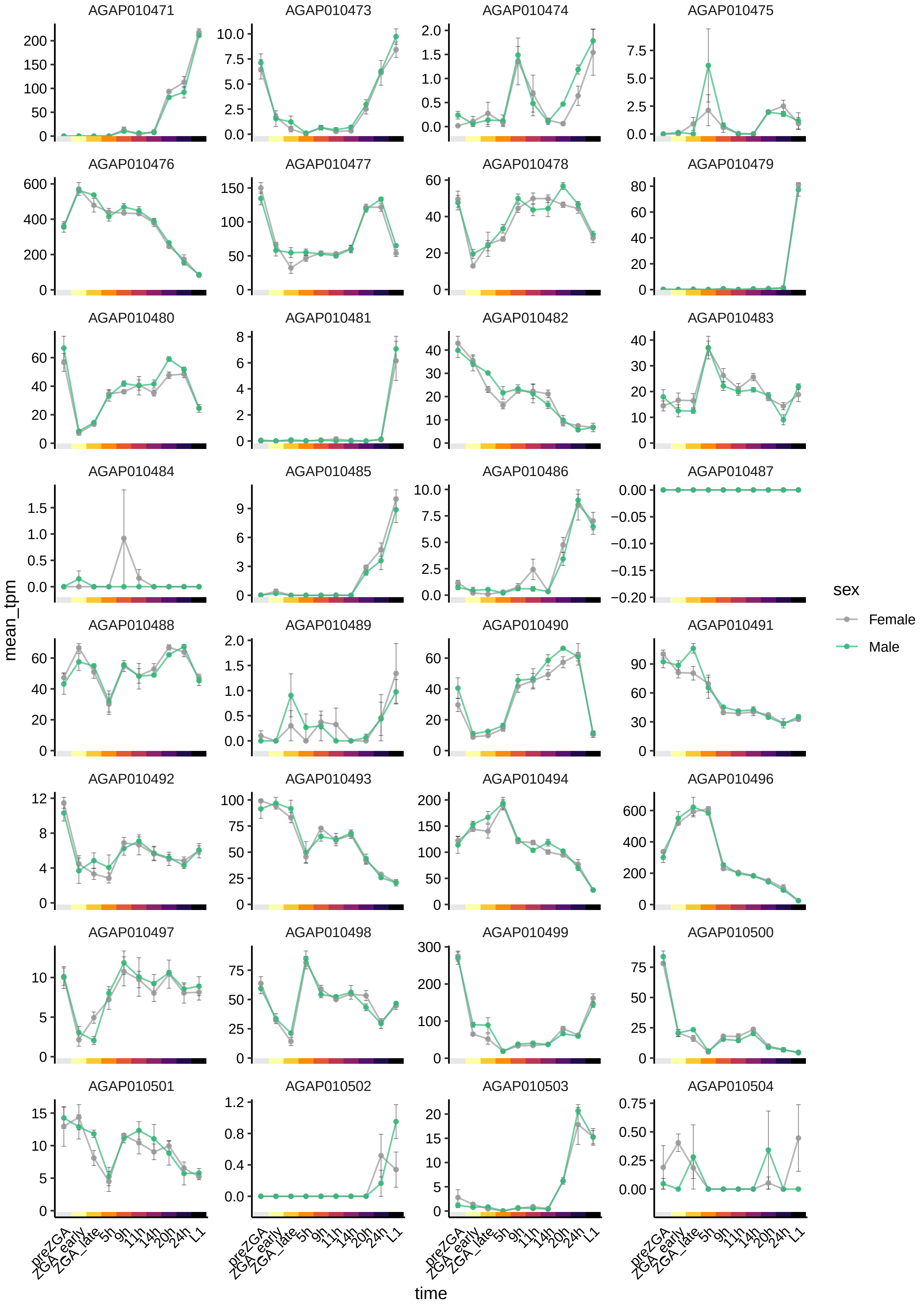

AGAP010471
AGAP010473
AGAP010474
AGAP010475
AGAP010476
AGAP010477
AGAP010478
AGAP010479
AGAP010480
AGAP010481
AGAP010482
AGAP010483
AGAP010484
AGAP010485
AGAP010486
AGAP010487
AGAP010488
AGAP010489
AGAP010490
AGAP010491
AGAP010492
AGAP010493
AGAP010494
AGAP010496
AGAP010497
AGAP010498
AGAP010499
AGAP010500
AGAP010501
AGAP010502
AGAP010503
AGAP010504
mean_tpm
time
preZGA
ZGA_early
ZGA_late
5h
9h
11h
14h
20h
24h
L1
sex
Female
Male

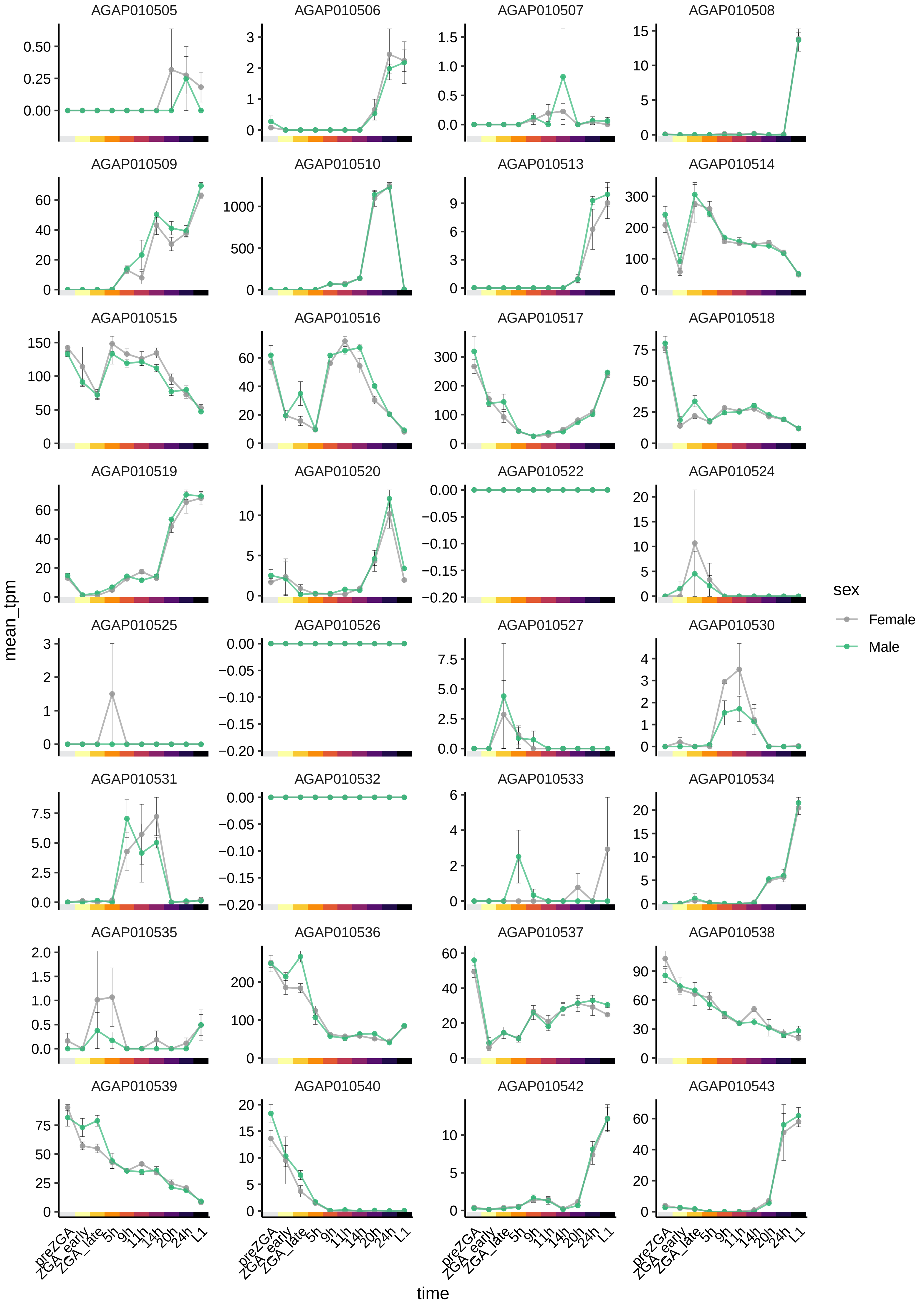
AGAP010505
AGAP010506
AGAP010507
AGAP010508
AGAP010509
AGAP010510
AGAP010513
AGAP010514
AGAP010515
AGAP010516
AGAP010517
AGAP010518
AGAP010519
AGAP010520
AGAP010522
AGAP010524
AGAP010525
AGAP010526
AGAP010527
AGAP010530
AGAP010531
AGAP010532
AGAP010533
AGAP010534
AGAP010535
AGAP010536
AGAP010537
AGAP010538
AGAP010539
AGAP010540
AGAP010542
AGAP010543
mean_tpm
time
preZGA
ZGA_early
ZGA_late
5h
9h
11h
14h
20h
24h
L1
sex
Female
Male

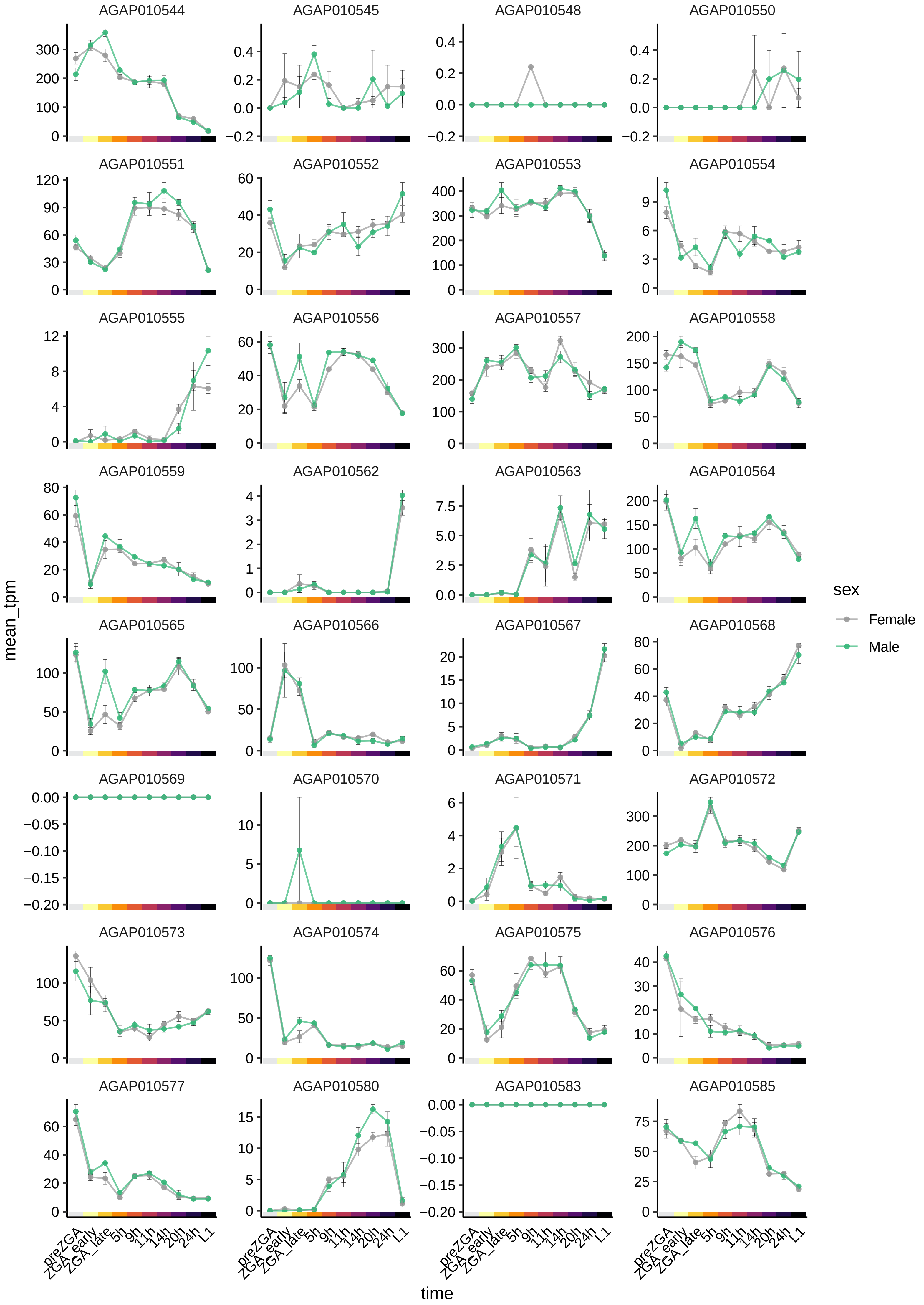

AGAP010544
AGAP010545
AGAP010548
AGAP010550
AGAP010551
AGAP010552
AGAP010553
AGAP010554
AGAP010555
AGAP010556
AGAP010557
AGAP010558
AGAP010559
AGAP010562
AGAP010563
AGAP010564
AGAP010565
AGAP010566
AGAP010567
AGAP010568
AGAP010569
AGAP010570
AGAP010571
AGAP010572
AGAP010573
AGAP010574
AGAP010575
AGAP010576
AGAP010577
AGAP010580
AGAP010583
AGAP010585
mean_tpm
time
preZGA
ZGA_early
ZGA_late
5h
9h
11h
14h
20h
24h
L1
sex
Female
Male

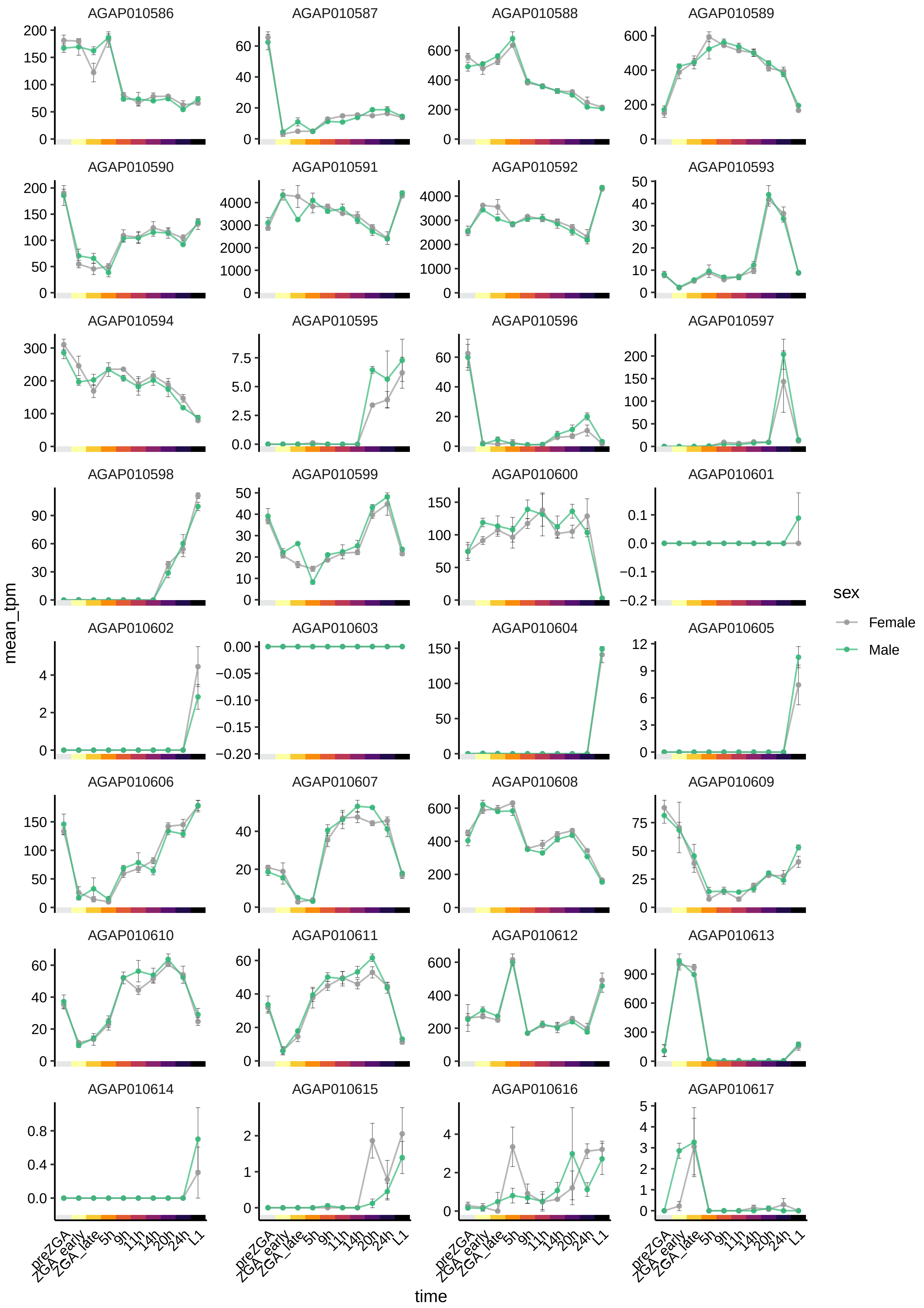

AGAP010586
AGAP010587
AGAP010588
AGAP010589
AGAP010590
AGAP010591
AGAP010592
AGAP010593
AGAP010594
AGAP010595
AGAP010596
AGAP010597
AGAP010598
AGAP010599
AGAP010600
AGAP010601
AGAP010602
AGAP010603
AGAP010604
AGAP010605
AGAP010606
AGAP010607
AGAP010608
AGAP010609
AGAP010610
AGAP010611
AGAP010612
AGAP010613
AGAP010614
AGAP010615
AGAP010616
AGAP010617
mean_tpm
time
preZGA
ZGA_early
ZGA_late
5h
9h
11h
14h
20h
24h
L1
sex
Female
Male

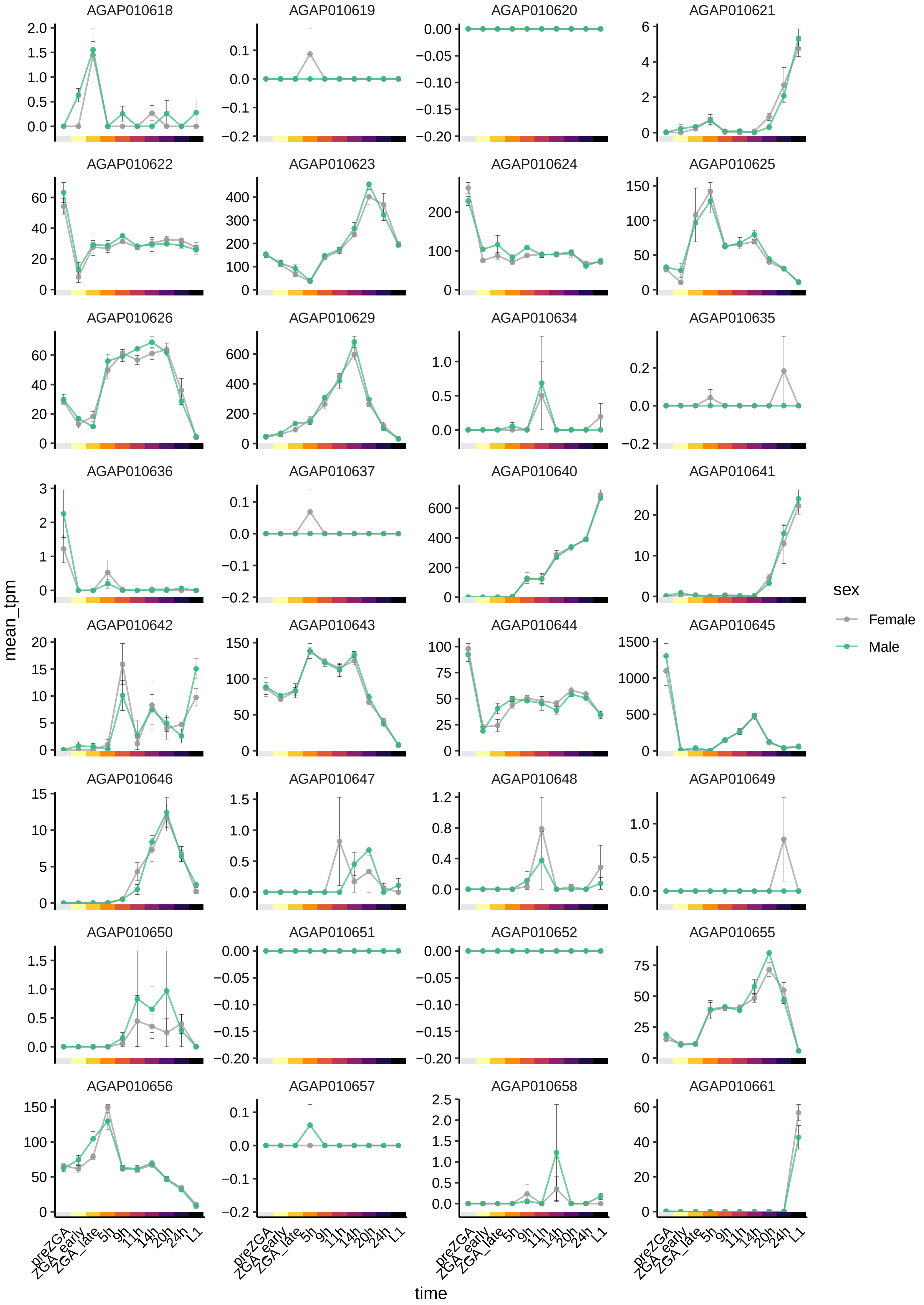

AGAP010618
AGAP010619
AGAP010620
AGAP010621
AGAP010622
AGAP010623
AGAP010624
AGAP010625
AGAP010626
AGAP010629
AGAP010634
AGAP010635
AGAP010636
AGAP010637
AGAP010640
AGAP010641
AGAP010642
AGAP010643
AGAP010644
AGAP010645
AGAP010646
AGAP010647
AGAP010648
AGAP010649
AGAP010650
AGAP010651
AGAP010652
AGAP010655
AGAP010656
AGAP010657
AGAP010658
AGAP010661
mean_tpm
time
preZGA
ZGA_early
ZGA_late
5h
9h
11h
14h
20h
24h
L1
sex
Female
Male

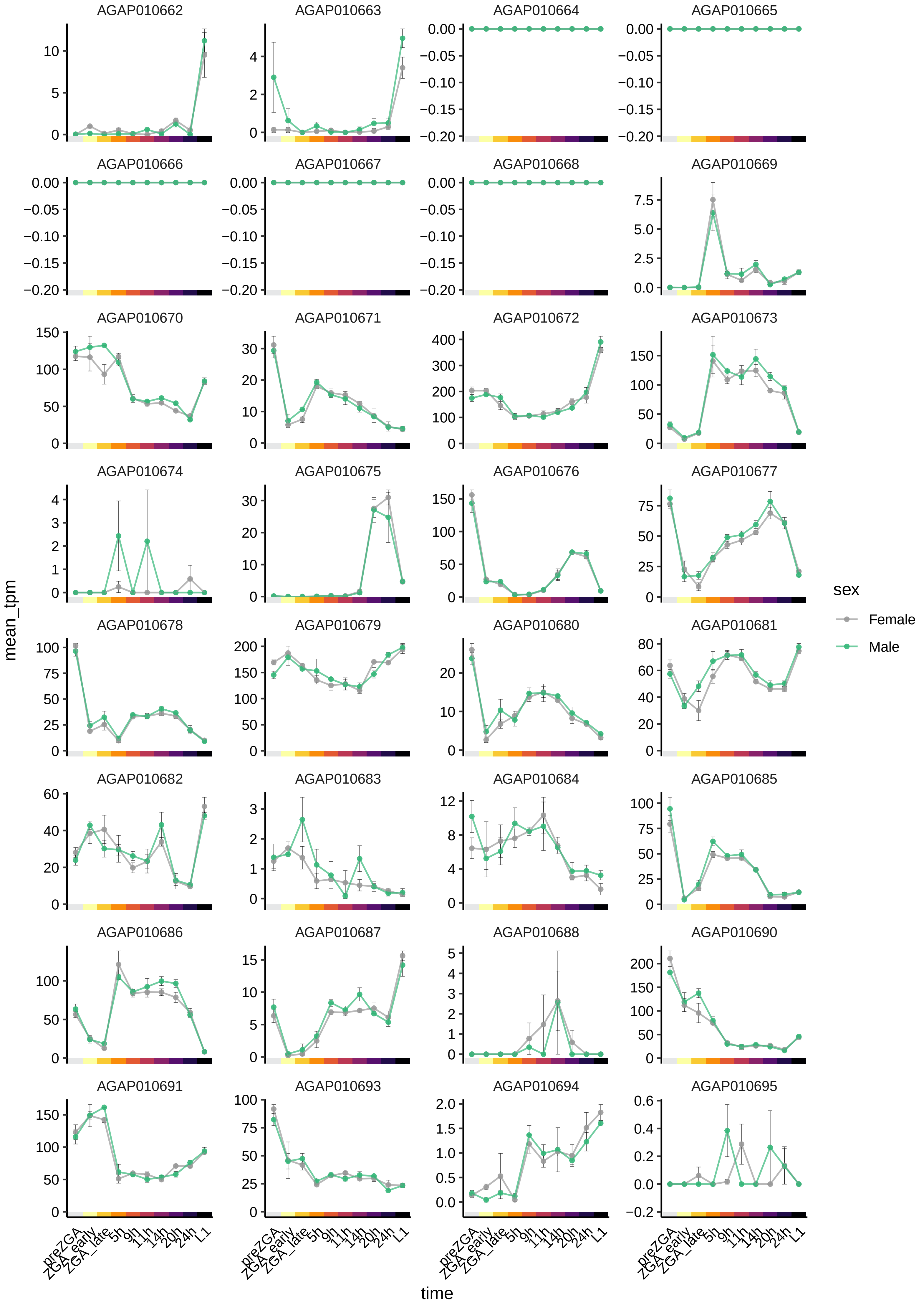

AGAP010662
AGAP010663
AGAP010664
AGAP010665
AGAP010666
AGAP010667
AGAP010668
AGAP010669
AGAP010670
AGAP010671
AGAP010672
AGAP010673
AGAP010674
AGAP010675
AGAP010676
AGAP010677
AGAP010678
AGAP010679
AGAP010680
AGAP010681
AGAP010682
AGAP010683
AGAP010684
AGAP010685
AGAP010686
AGAP010687
AGAP010688
AGAP010690
AGAP010691
AGAP010693
AGAP010694
AGAP010695
mean_tpm
time
preZGA
ZGA_early
ZGA_late
5h
9h
11h
14h
20h
24h
L1
sex
Female
Male

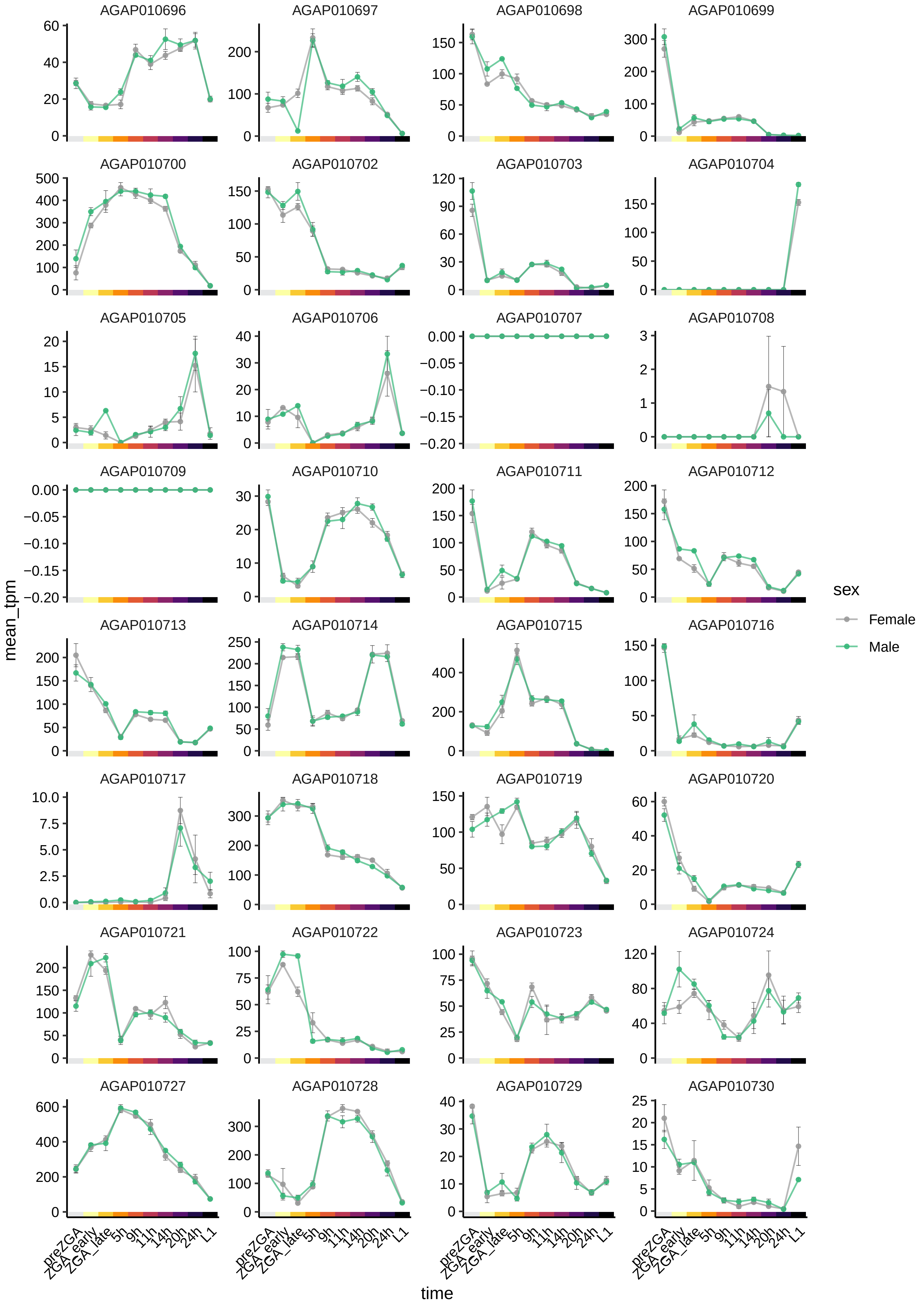
AGAP010696
AGAP010697
AGAP010698
AGAP010699
AGAP010700
AGAP010702
AGAP010703
AGAP010704
AGAP010705
AGAP010706
AGAP010707
AGAP010708
AGAP010709
AGAP010710
AGAP010711
AGAP010712
AGAP010713
AGAP010714
AGAP010715
AGAP010716
AGAP010717
AGAP010718
AGAP010719
AGAP010720
AGAP010721
AGAP010722
AGAP010723
AGAP010724
AGAP010727
AGAP010728
AGAP010729
AGAP010730
mean_tpm
time
preZGA
ZGA_early
ZGA_late
5h
9h
11h
14h
20h
24h
L1
sex
Female
Male

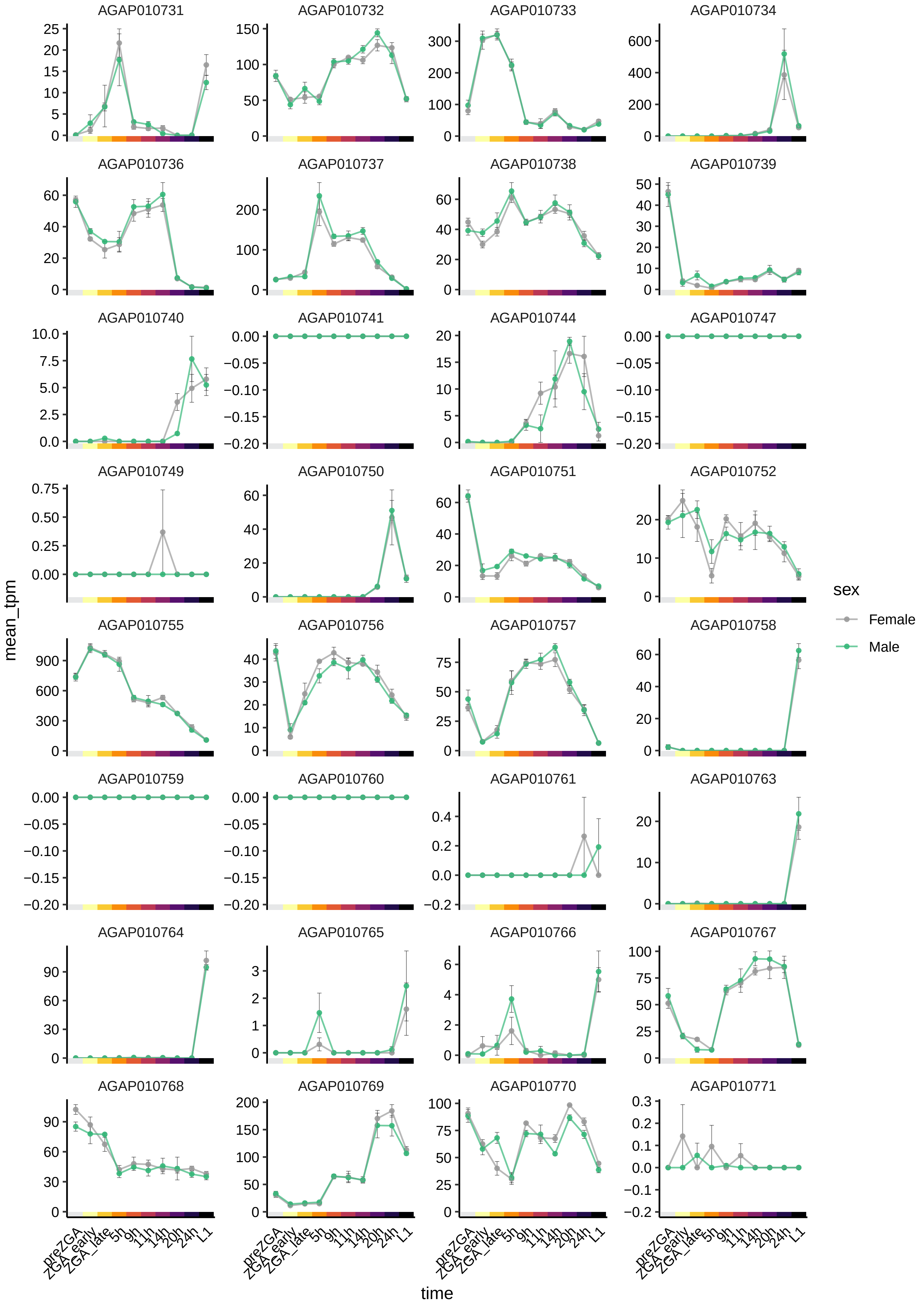

AGAP010731
AGAP010732
AGAP010733
AGAP010734
AGAP010736
AGAP010737
AGAP010738
AGAP010739
AGAP010740
AGAP010741
AGAP010744
AGAP010747
AGAP010749
AGAP010750
AGAP010751
AGAP010752
AGAP010755
AGAP010756
AGAP010757
AGAP010758
AGAP010759
AGAP010760
AGAP010761
AGAP010763
AGAP010764
AGAP010765
AGAP010766
AGAP010767
AGAP010768
AGAP010769
AGAP010770
AGAP010771
mean_tpm
time
preZGA
ZGA_early
ZGA_late
5h
9h
11h
14h
20h
24h
L1
sex
Female
Male

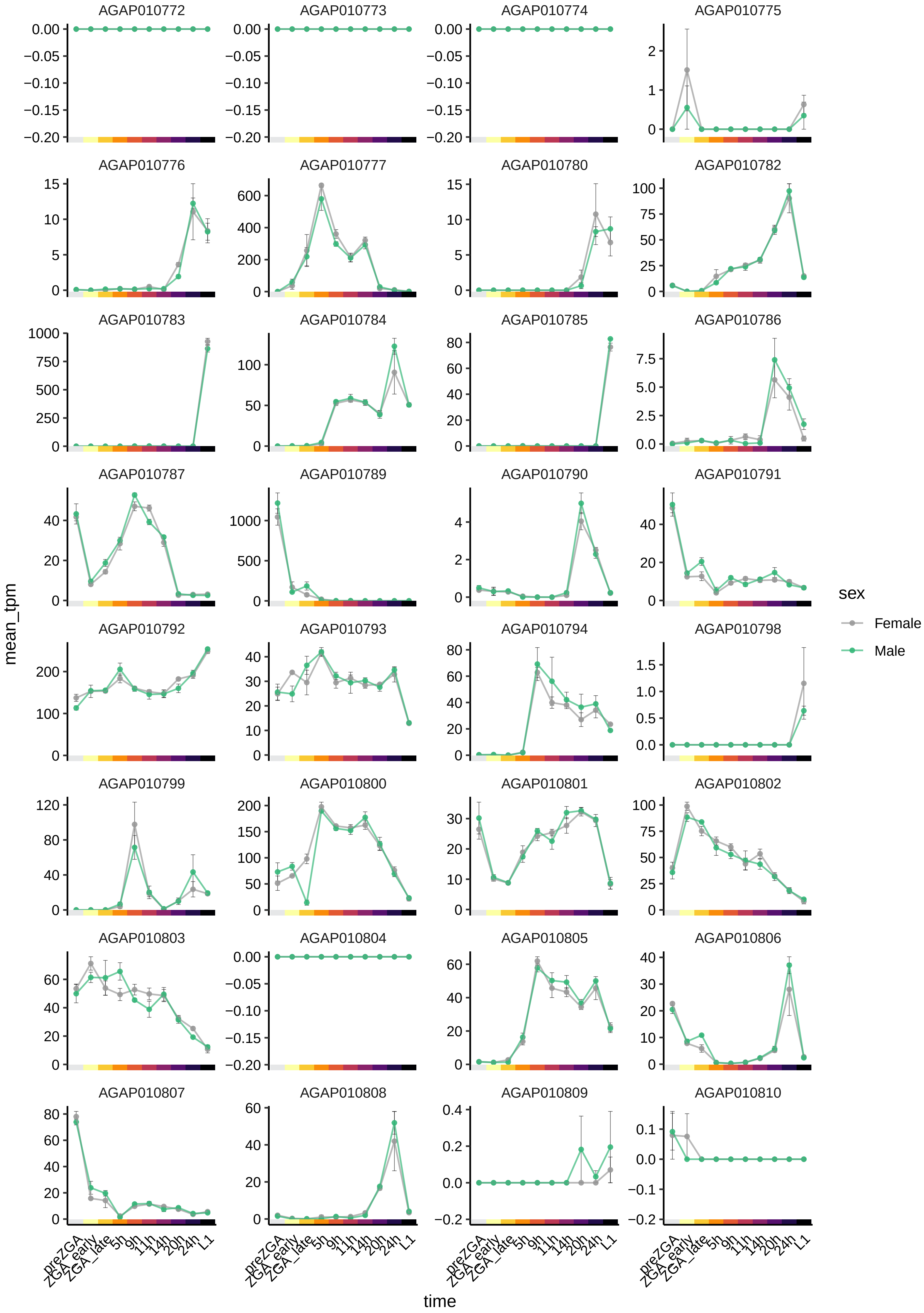
AGAP010772
AGAP010773
AGAP010774
AGAP010775
AGAP010776
AGAP010777
AGAP010780
AGAP010782
AGAP010783
AGAP010784
AGAP010785
AGAP010786
AGAP010787
AGAP010789
AGAP010790
AGAP010791
AGAP010792
AGAP010793
AGAP010794
AGAP010798
AGAP010799
AGAP010800
AGAP010801
AGAP010802
AGAP010803
AGAP010804
AGAP010805
AGAP010806
AGAP010807
AGAP010808
AGAP010809
AGAP010810
mean_tpm
preZGA
ZGA_early
ZGA_late
5h
9h
11h
14h
20h
24h
L1
time
sex
Female
Male

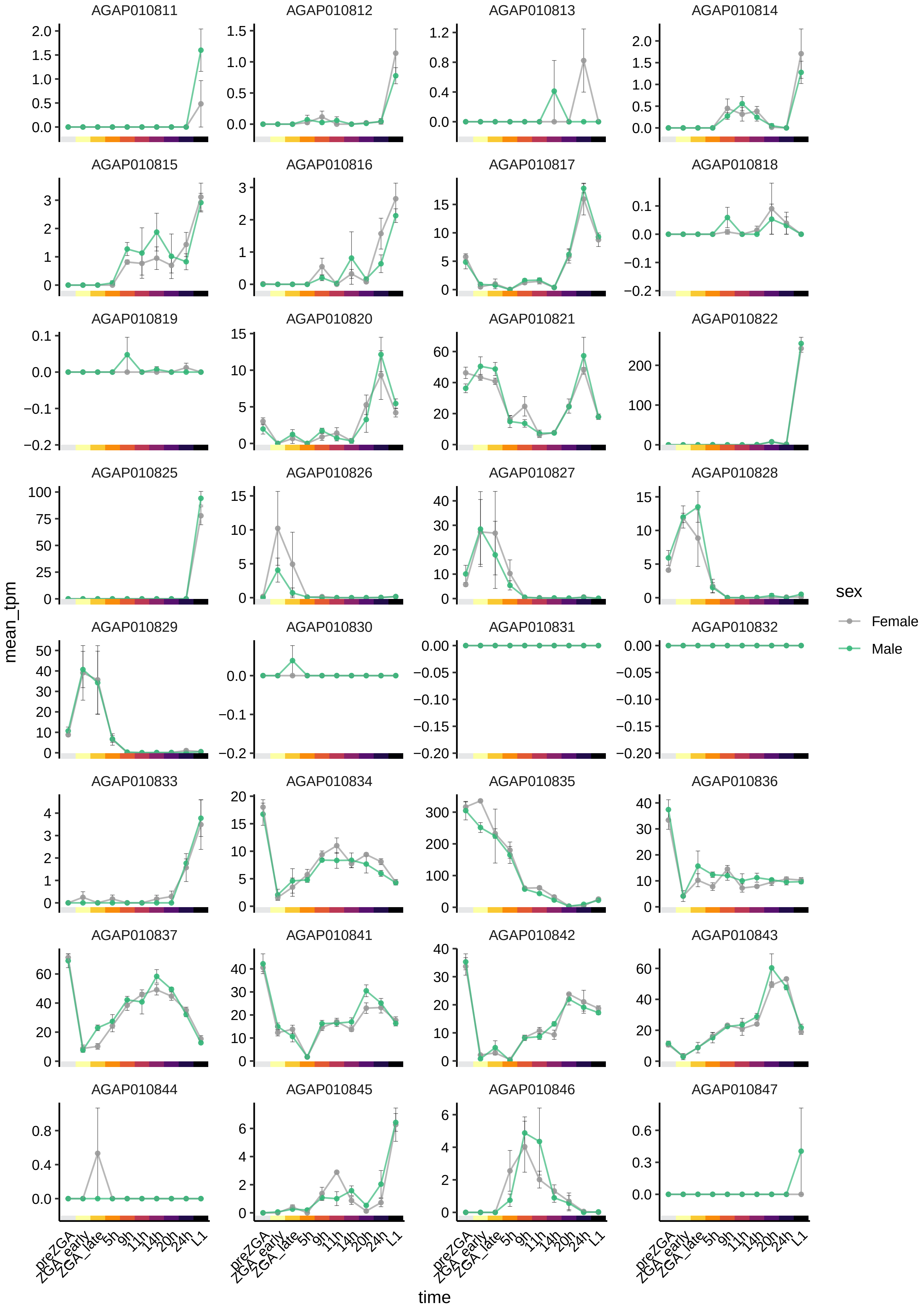

AGAP010811
AGAP010812
AGAP010813
AGAP010814
AGAP010815
AGAP010816
AGAP010817
AGAP010818
AGAP010819
AGAP010820
AGAP010821
AGAP010822
AGAP010825
AGAP010826
AGAP010827
AGAP010828
AGAP010829
AGAP010830
AGAP010831
AGAP010832
AGAP010833
AGAP010834
AGAP010835
AGAP010836
AGAP010837
AGAP010841
AGAP010842
AGAP010843
AGAP010844
AGAP010845
AGAP010846
AGAP010847
mean_tpm
time
preZGA
ZGA_early
ZGA_late
5h
9h
11h
14h
20h
24h
L1
sex
Female
Male

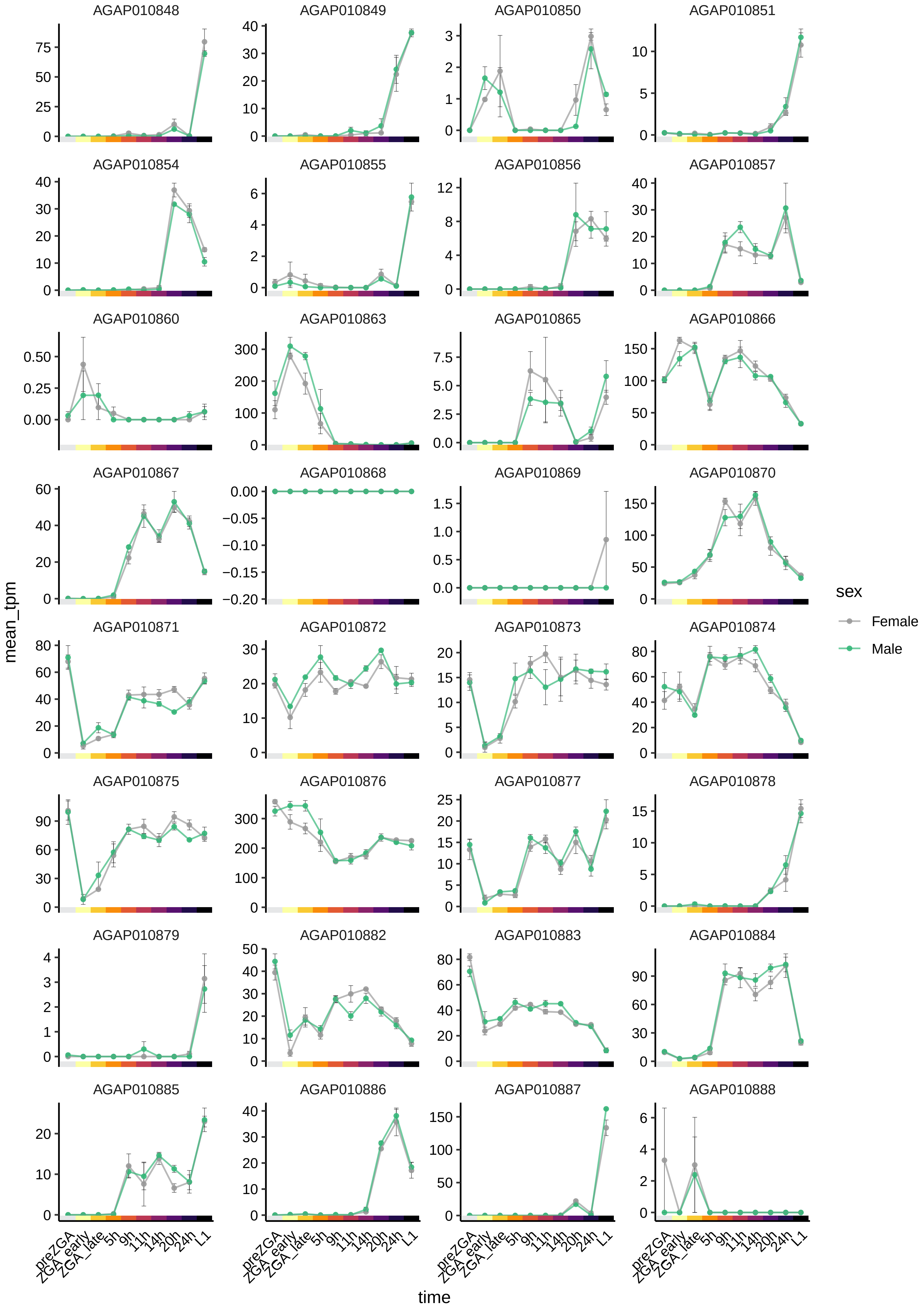

AGAP010848
AGAP010849
AGAP010850
AGAP010851
AGAP010854
AGAP010855
AGAP010856
AGAP010857
AGAP010860
AGAP010863
AGAP010865
AGAP010866
AGAP010867
AGAP010868
AGAP010869
AGAP010870
AGAP010871
AGAP010872
AGAP010873
AGAP010874
AGAP010875
AGAP010876
AGAP010877
AGAP010878
AGAP010879
AGAP010882
AGAP010883
AGAP010884
AGAP010885
AGAP010886
AGAP010887
AGAP010888
mean_tpm
time
preZGA
ZGA_early
ZGA_late
5h
9h
11h
14h
20h
24h
L1
sex
Female
Male

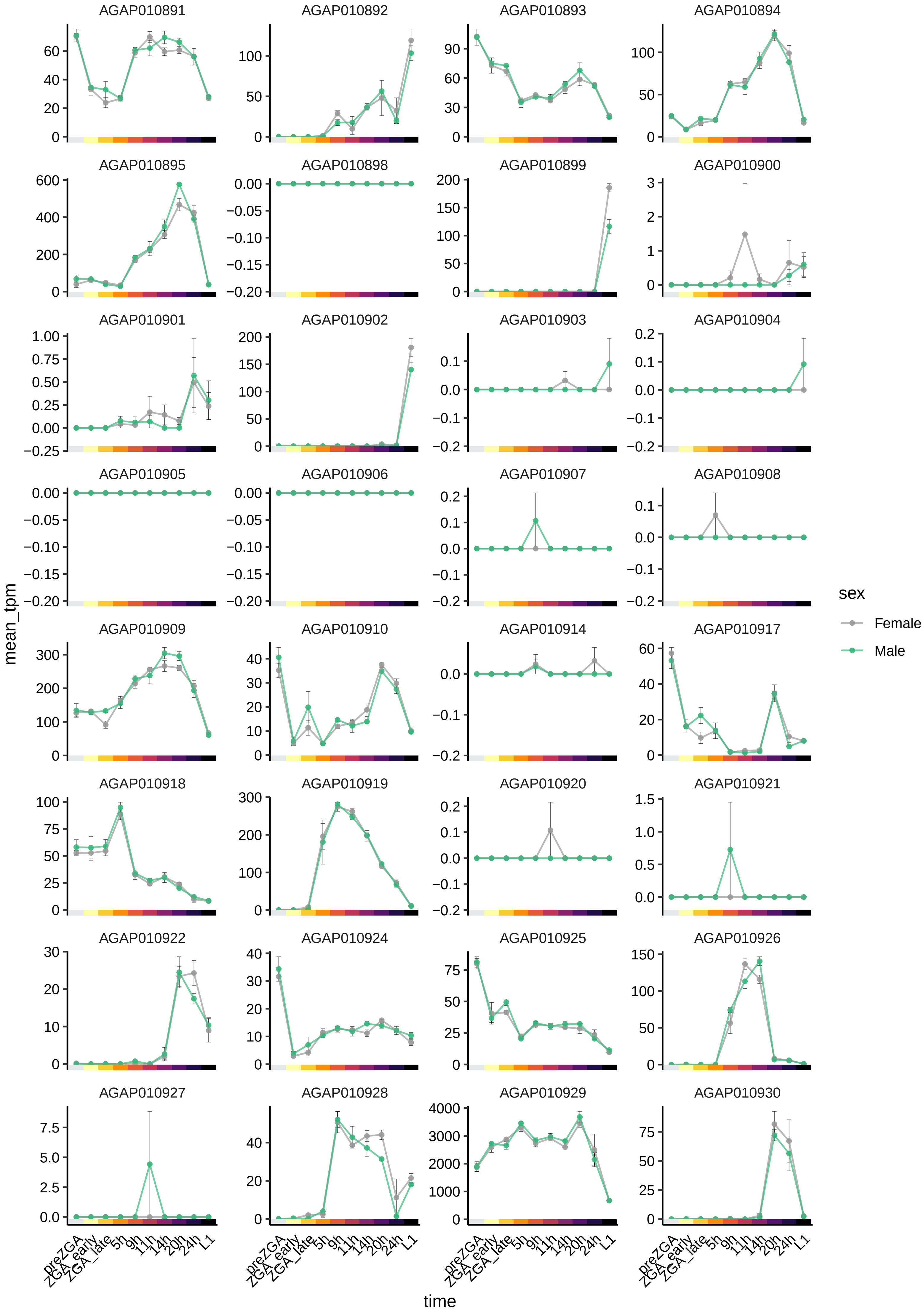
AGAP010891
AGAP010892
AGAP010893
AGAP010894
AGAP010895
AGAP010898
AGAP010899
AGAP010900
AGAP010901
AGAP010902
AGAP010903
AGAP010904
AGAP010905
AGAP010906
AGAP010907
AGAP010908
AGAP010909
AGAP010910
AGAP010914
AGAP010917
AGAP010918
AGAP010919
AGAP010920
AGAP010921
AGAP010922
AGAP010924
AGAP010925
AGAP010926
AGAP010927
AGAP010928
AGAP010929
AGAP010930
mean_tpm
time
preZGA
ZGA_early
ZGA_late
5h
9h
11h
14h
20h
24h
L1
sex
Female
Male

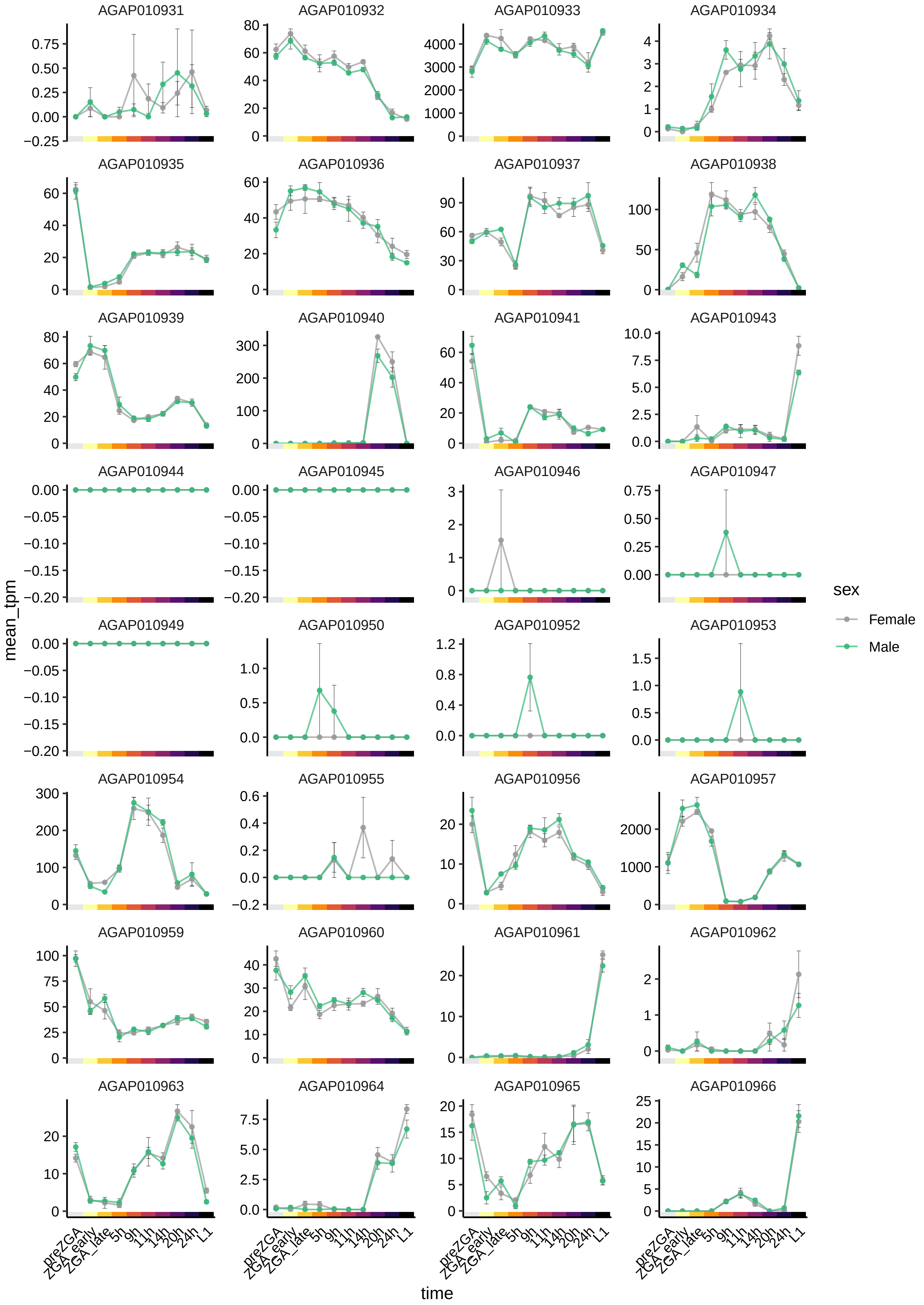

AGAP010931
AGAP010932
AGAP010933
AGAP010934
AGAP010935
AGAP010936
AGAP010937
AGAP010938
AGAP010939
AGAP010940
AGAP010941
AGAP010943
AGAP010944
AGAP010945
AGAP010946
AGAP010947
AGAP010949
AGAP010950
AGAP010952
AGAP010953
AGAP010954
AGAP010955
AGAP010956
AGAP010957
AGAP010959
AGAP010960
AGAP010961
AGAP010962
AGAP010963
AGAP010964
AGAP010965
AGAP010966
mean_tpm
time
preZGA
ZGA_early
ZGA_late
5h
9h
11h
14h
20h
24h
L1
sex
Female
Male

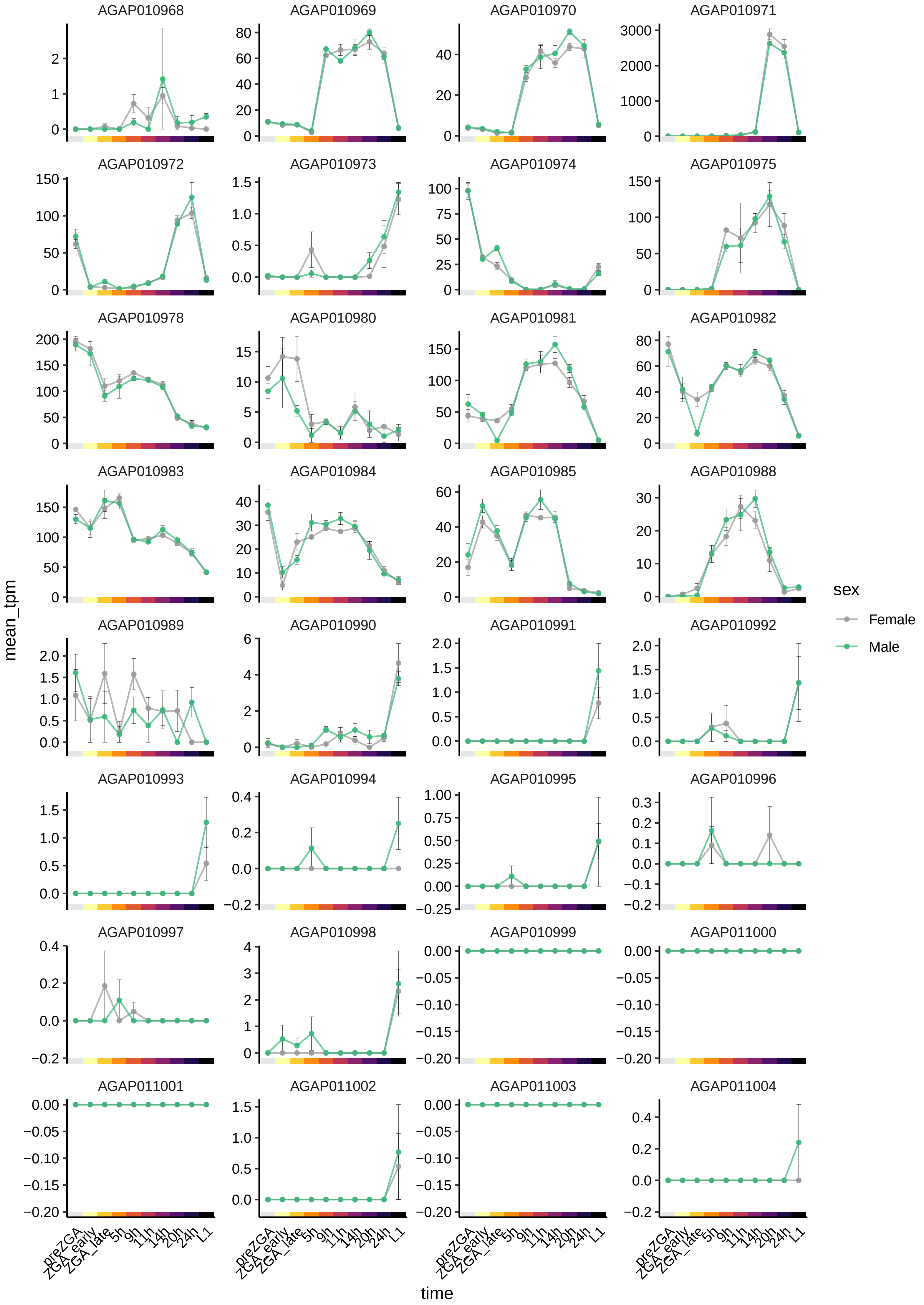
AGAP010968
AGAP010969
AGAP010970
AGAP010971
AGAP010972
AGAP010973
AGAP010974
AGAP010975
AGAP010978
AGAP010980
AGAP010981
AGAP010982
AGAP010983
AGAP010984
AGAP010985
AGAP010988
AGAP010989
AGAP010990
AGAP010991
AGAP010992
AGAP010993
AGAP010994
AGAP010995
AGAP010996
AGAP010997
AGAP010998
AGAP010999
AGAP011000
AGAP011001
AGAP011002
AGAP011003
AGAP011004
mean_tpm
time
preZGA
ZGA_early
ZGA_late
5h
9h
11h
14h
20h
24h
L1
sex
Female
Male

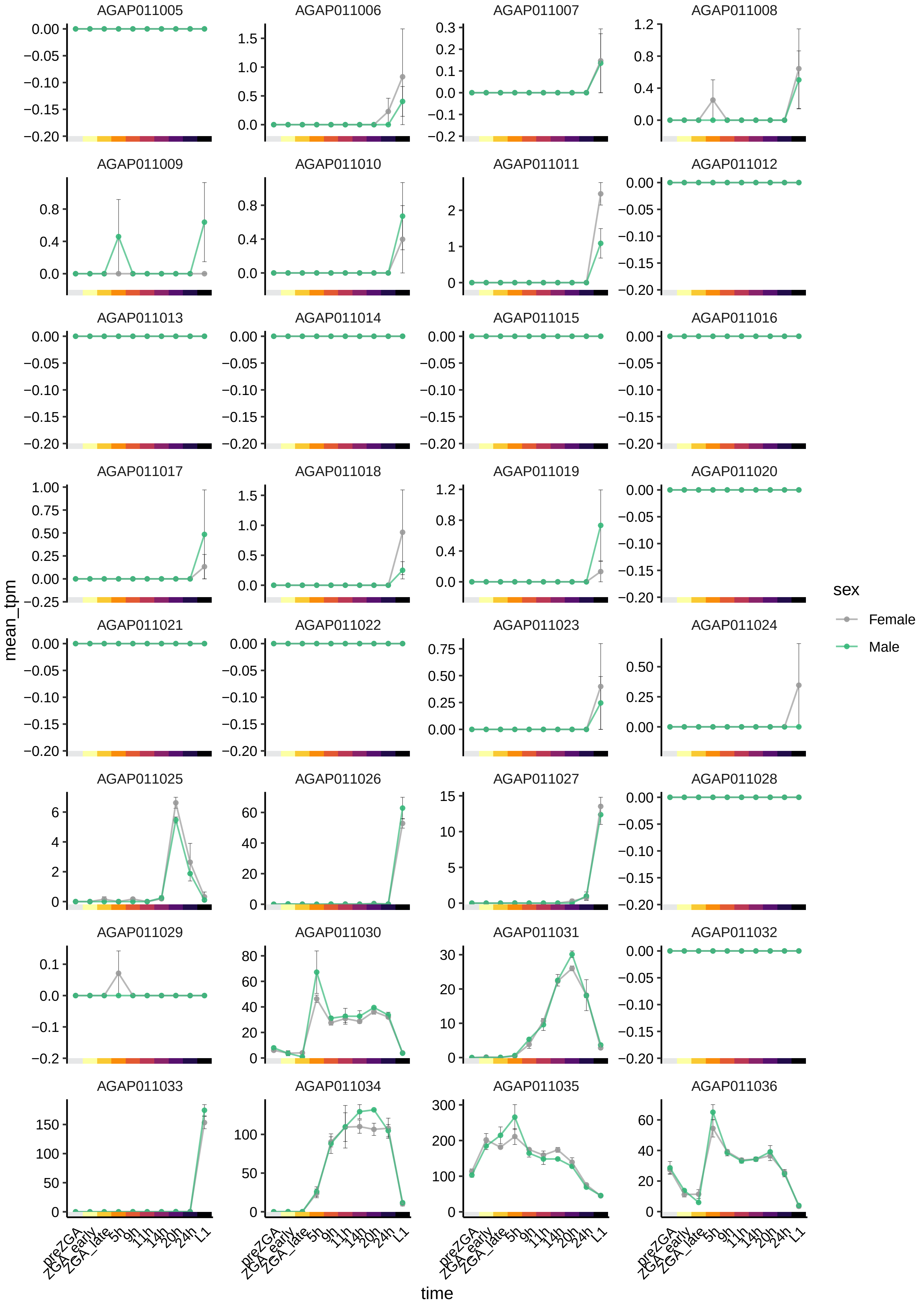

AGAP011005
AGAP011006
AGAP011007
AGAP011008
AGAP011009
AGAP011010
AGAP011011
AGAP011012
AGAP011013
AGAP011014
AGAP011015
AGAP011016
AGAP011017
AGAP011018
AGAP011019
AGAP011020
AGAP011021
AGAP011022
AGAP011023
AGAP011024
AGAP011025
AGAP011026
AGAP011027
AGAP011028
AGAP011029
AGAP011030
AGAP011031
AGAP011032
AGAP011033
AGAP011034
AGAP011035
AGAP011036
mean_tpm
time
preZGA
ZGA_early
ZGA_late
5h
9h
11h
14h
20h
24h
L1
sex
Female
Male

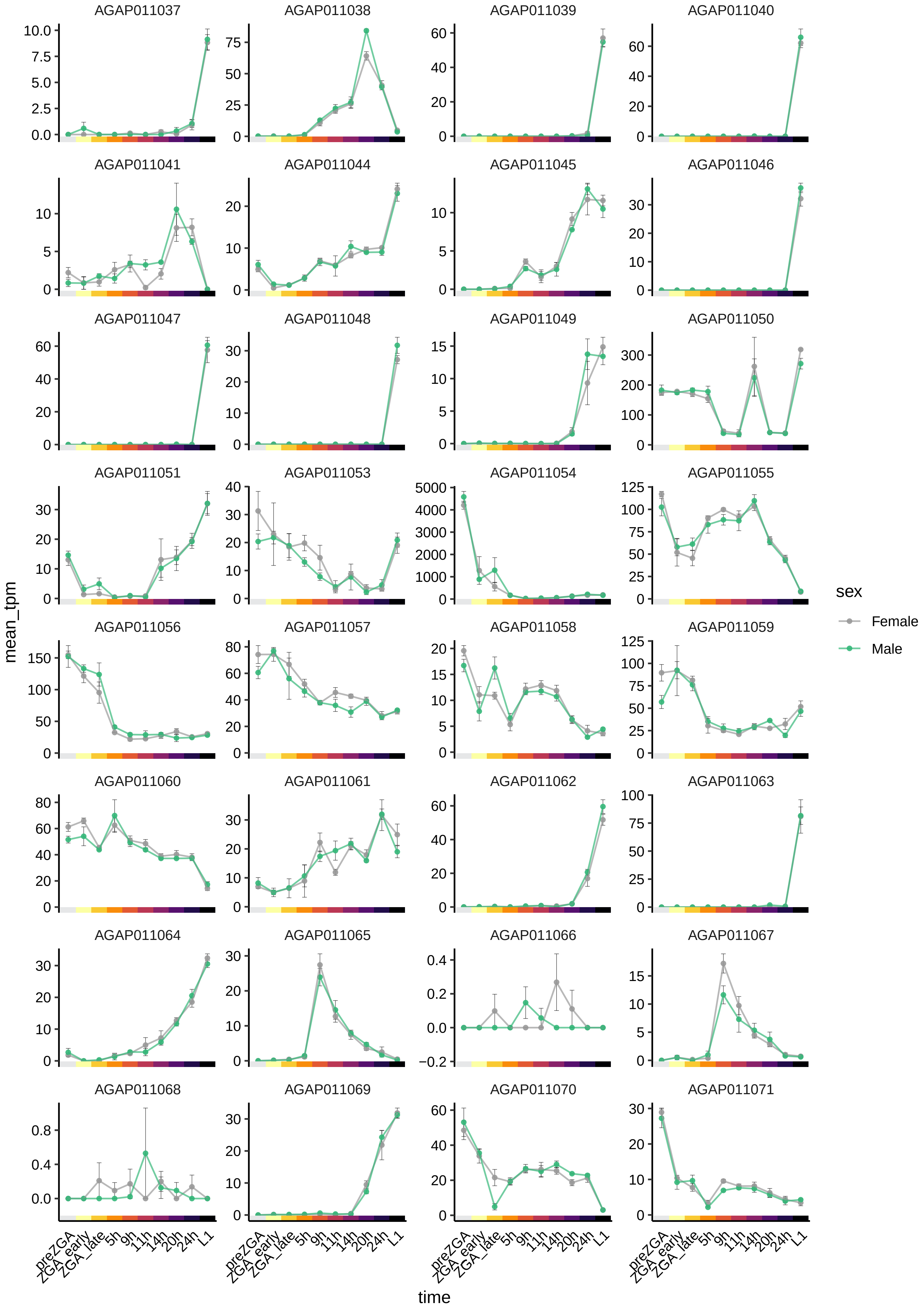

AGAP011037
AGAP011038
AGAP011039
AGAP011040
AGAP011041
AGAP011044
AGAP011045
AGAP011046
AGAP011047
AGAP011048
AGAP011049
AGAP011050
AGAP011051
AGAP011053
AGAP011054
AGAP011055
AGAP011056
AGAP011057
AGAP011058
AGAP011059
AGAP011060
AGAP011061
AGAP011062
AGAP011063
AGAP011064
AGAP011065
AGAP011066
AGAP011067
AGAP011068
AGAP011069
AGAP011070
AGAP011071
mean_tpm
time
preZGA
ZGA_early
ZGA_late
5h
9h
11h
14h
20h
24h
L1
sex
Female
Male

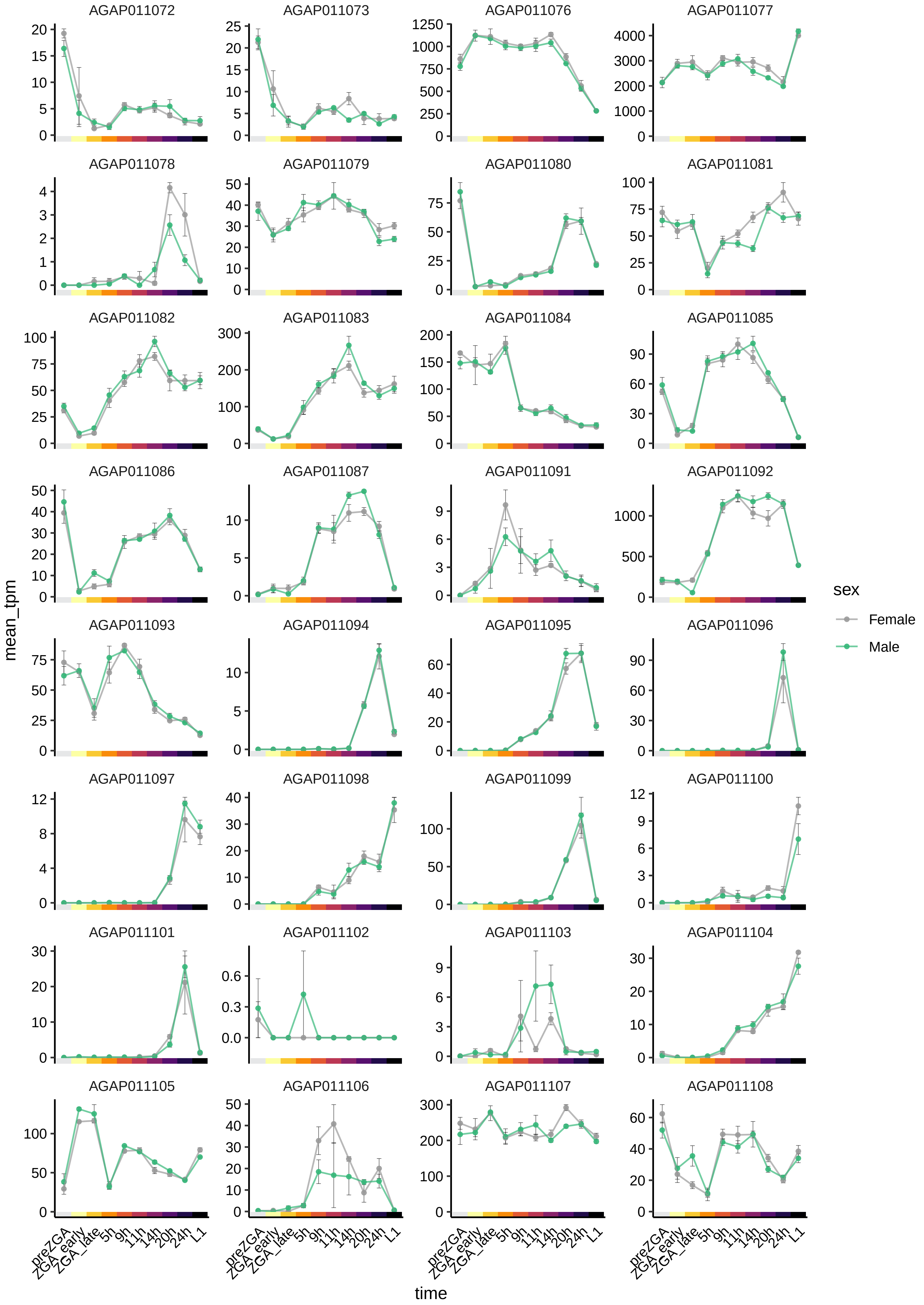

AGAP011072
AGAP011073
AGAP011076
AGAP011077
AGAP011078
AGAP011079
AGAP011080
AGAP011081
AGAP011082
AGAP011083
AGAP011084
AGAP011085
AGAP011086
AGAP011087
AGAP011091
AGAP011092
AGAP011093
AGAP011094
AGAP011095
AGAP011096
AGAP011097
AGAP011098
AGAP011099
AGAP011100
AGAP011101
AGAP011102
AGAP011103
AGAP011104
AGAP011105
AGAP011106
AGAP011107
AGAP011108
mean_tpm
time
preZGA
ZGA_early
ZGA_late
5h
9h
11h
14h
20h
24h
L1
sex
Female
Male

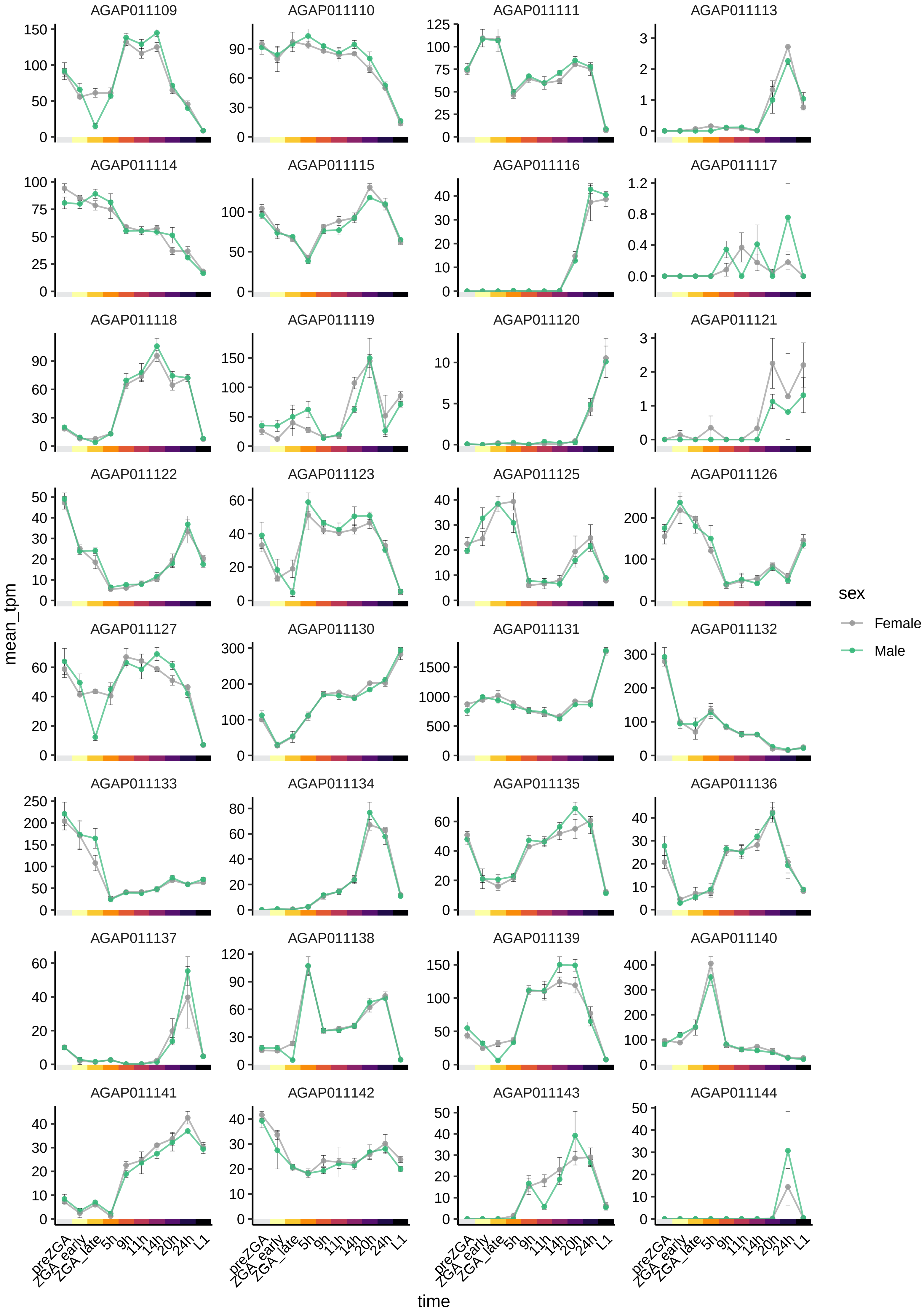

AGAP011109
AGAP011110
AGAP011111
AGAP011113
AGAP011114
AGAP011115
AGAP011116
AGAP011117
AGAP011118
AGAP011119
AGAP011120
AGAP011121
AGAP011122
AGAP011123
AGAP011125
AGAP011126
AGAP011127
AGAP011130
AGAP011131
AGAP011132
AGAP011133
AGAP011134
AGAP011135
AGAP011136
AGAP011137
AGAP011138
AGAP011139
AGAP011140
AGAP011141
AGAP011142
AGAP011143
AGAP011144
mean_tpm
time
preZGA
ZGA_early
ZGA_late
5h
9h
11h
14h
20h
24h
L1
sex
Female
Male

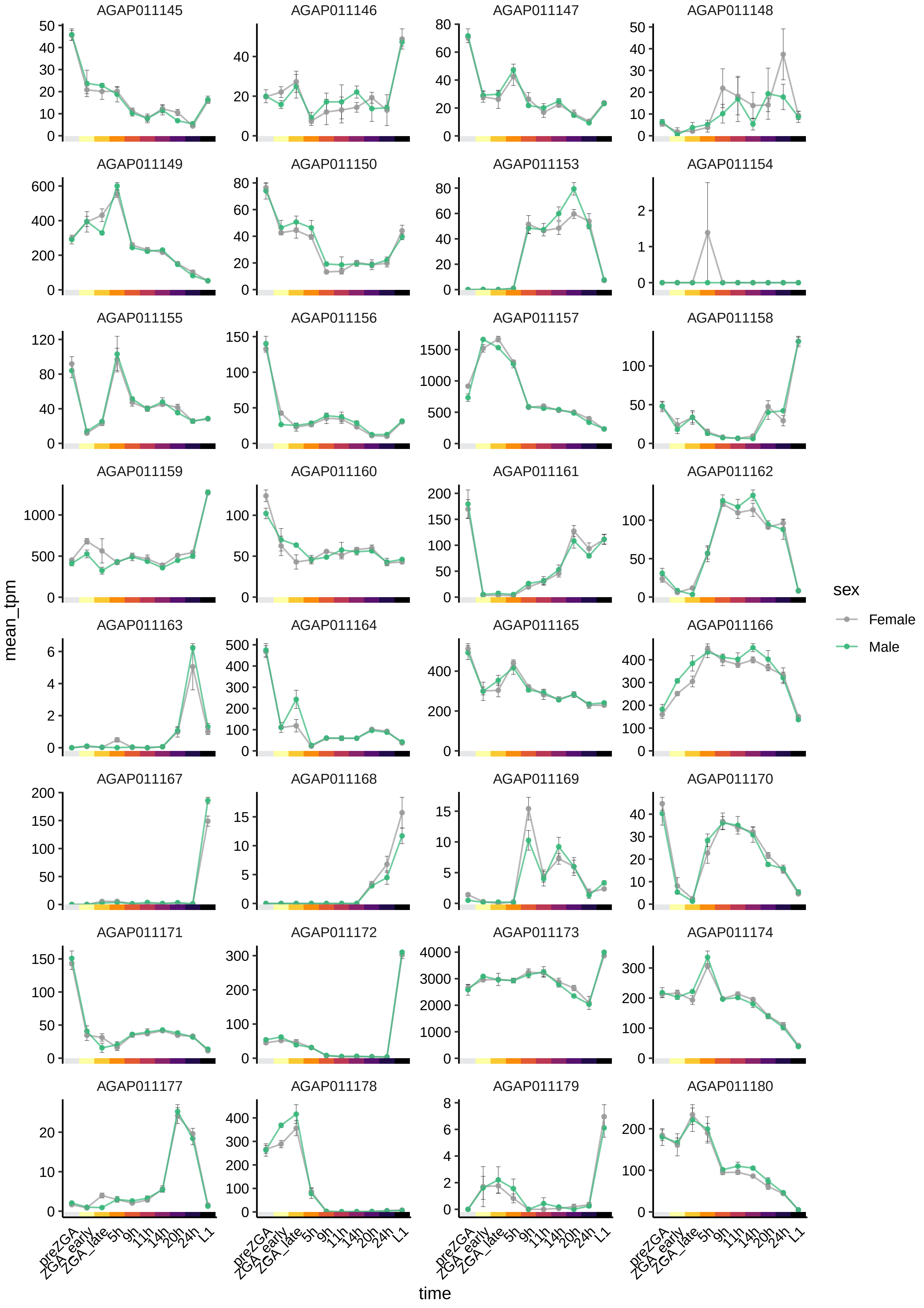

AGAP011145
AGAP011146
AGAP011147
AGAP011148
AGAP011149
AGAP011150
AGAP011153
AGAP011154
AGAP011155
AGAP011156
AGAP011157
AGAP011158
AGAP011159
AGAP011160
AGAP011161
AGAP011162
AGAP011163
AGAP011164
AGAP011165
AGAP011166
AGAP011167
AGAP011168
AGAP011169
AGAP011170
AGAP011171
AGAP011172
AGAP011173
AGAP011174
AGAP011177
AGAP011178
AGAP011179
AGAP011180
mean_tpm
time
preZGA
ZGA_early
ZGA_late
5h
9h
11h
14h
20h
24h
L1
sex
Female
Male

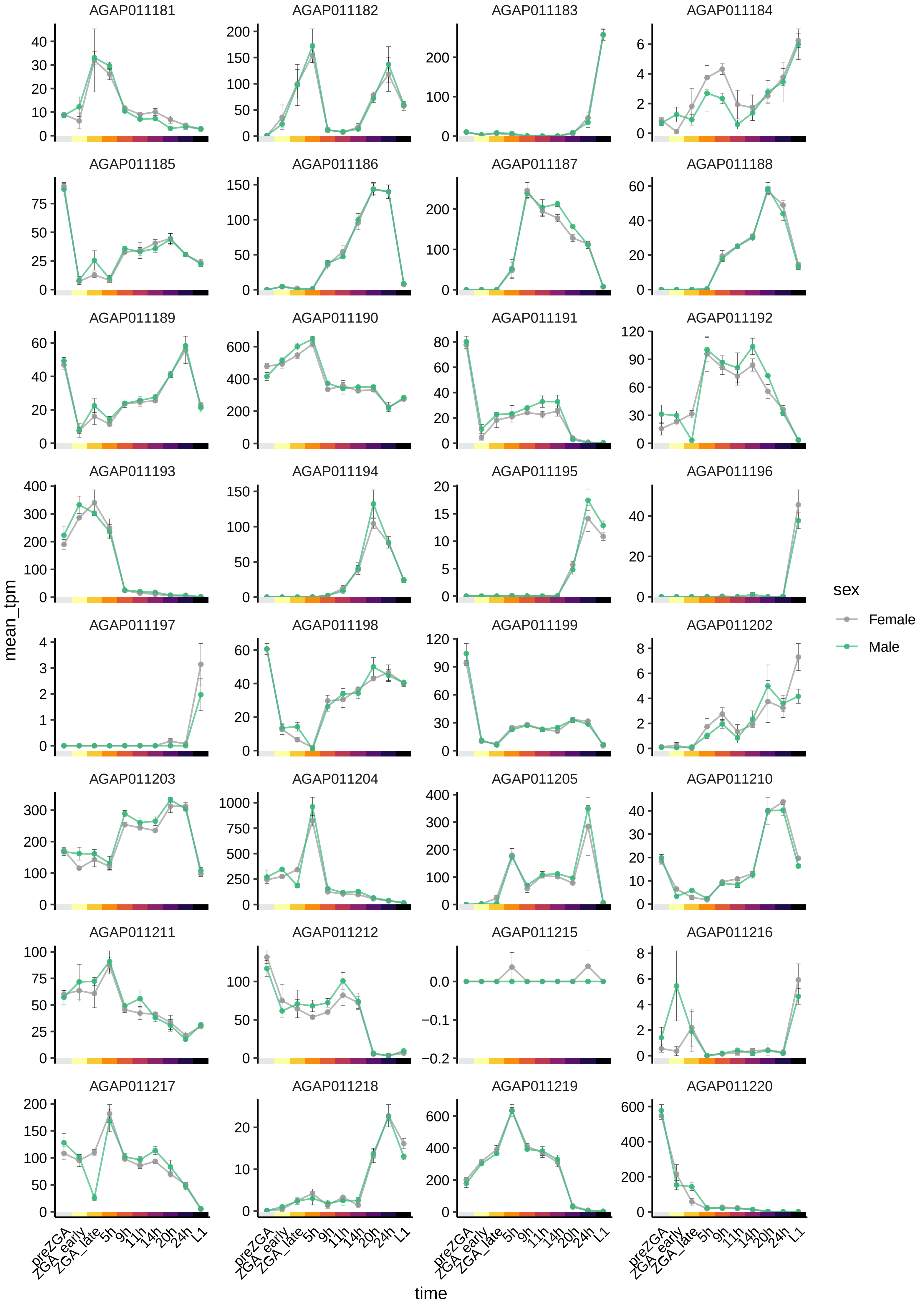

AGAP011181
AGAP011182
AGAP011183
AGAP011184
AGAP011185
AGAP011186
AGAP011187
AGAP011188
AGAP011189
AGAP011190
AGAP011191
AGAP011192
AGAP011193
AGAP011194
AGAP011195
AGAP011196
AGAP011197
AGAP011198
AGAP011199
AGAP011202
AGAP011203
AGAP011204
AGAP011205
AGAP011210
AGAP011211
AGAP011212
AGAP011215
AGAP011216
AGAP011217
AGAP011218
AGAP011219
AGAP011220
mean_tpm
time
preZGA
ZGA_early
ZGA_late
5h
9h
11h
14h
20h
24h
L1
sex
Female
Male

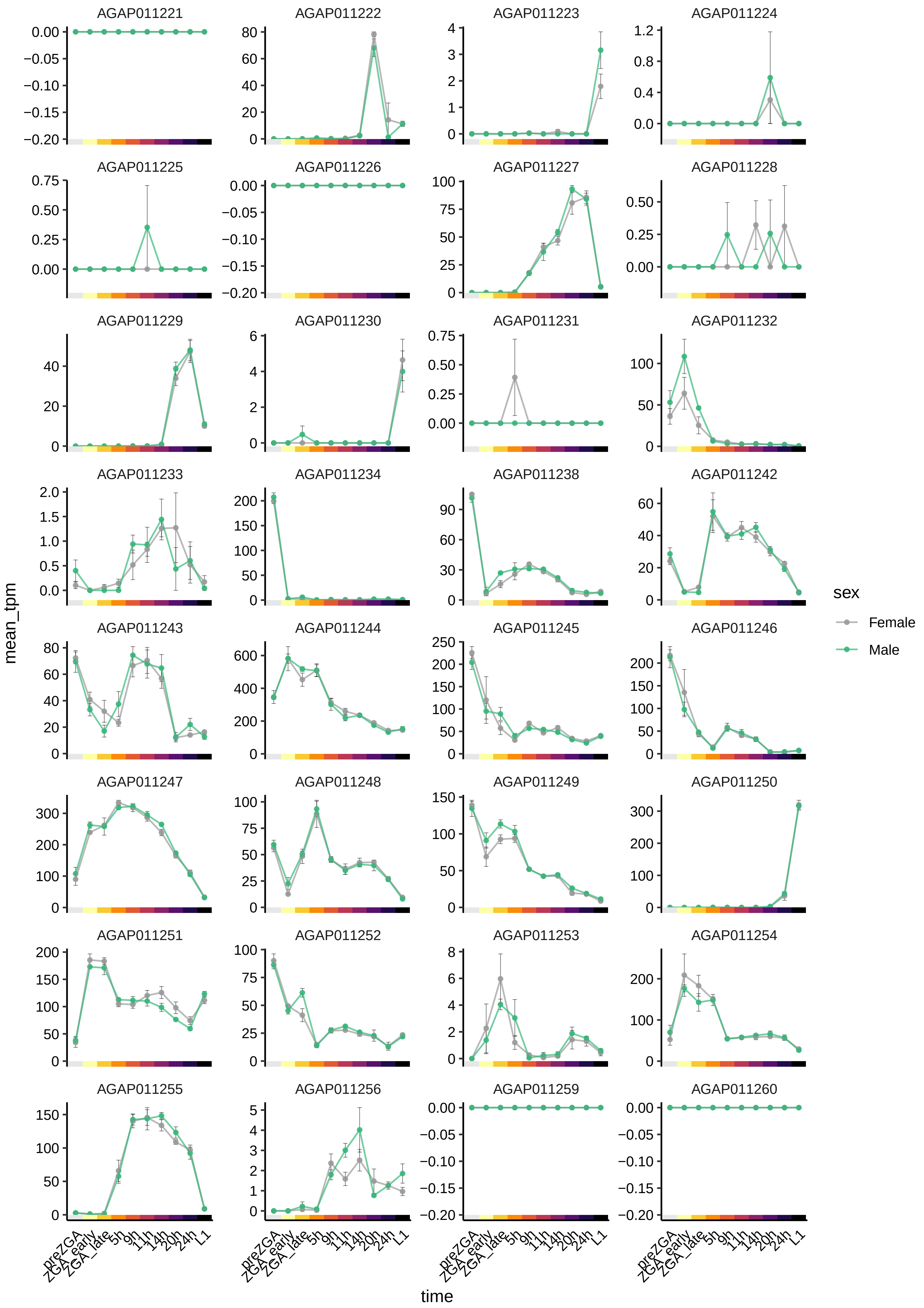
AGAP011221
AGAP011222
AGAP011223
AGAP011224
AGAP011225
AGAP011226
AGAP011227
AGAP011228
AGAP011229
AGAP011230
AGAP011231
AGAP011232
AGAP011233
AGAP011234
AGAP011238
AGAP011242
AGAP011243
AGAP011244
AGAP011245
AGAP011246
AGAP011247
AGAP011248
AGAP011249
AGAP011250
AGAP011251
AGAP011252
AGAP011253
AGAP011254
AGAP011255
AGAP011256
AGAP011259
AGAP011260
mean_tpm
time
preZGA
ZGA_early
ZGA_late
5h
9h
11h
14h
20h
24h
L1
sex
Female
Male

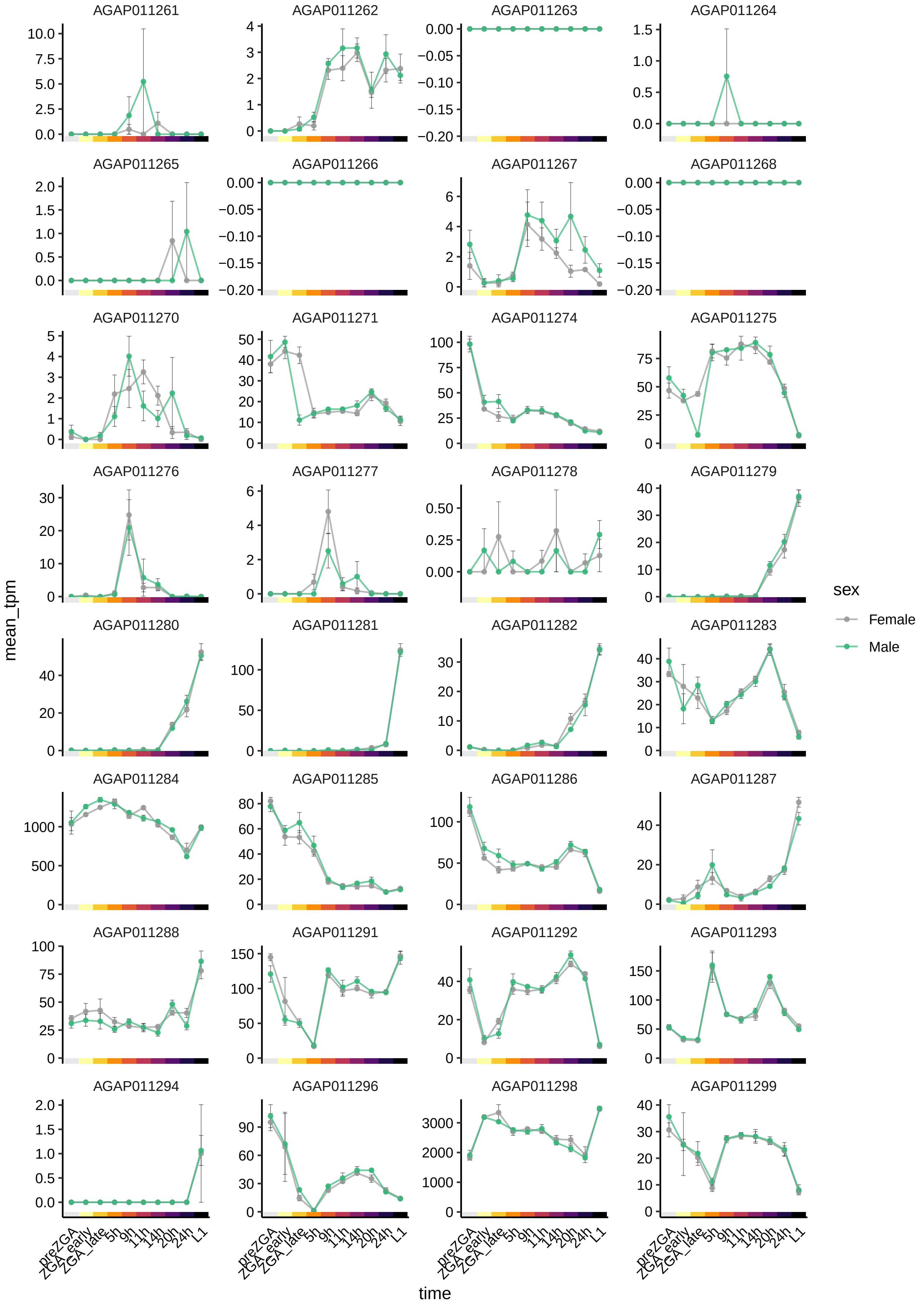

AGAP011261
AGAP011262
AGAP011263
AGAP011264
AGAP011265
AGAP011266
AGAP011267
AGAP011268
AGAP011270
AGAP011271
AGAP011274
AGAP011275
AGAP011276
AGAP011277
AGAP011278
AGAP011279
AGAP011280
AGAP011281
AGAP011282
AGAP011283
AGAP011284
AGAP011285
AGAP011286
AGAP011287
AGAP011288
AGAP011291
AGAP011292
AGAP011293
AGAP011294
AGAP011296
AGAP011298
AGAP011299
mean_tpm
time
preZGA
ZGA_early
ZGA_late
5h
9h
11h
14h
20h
24h
L1
sex
Female
Male

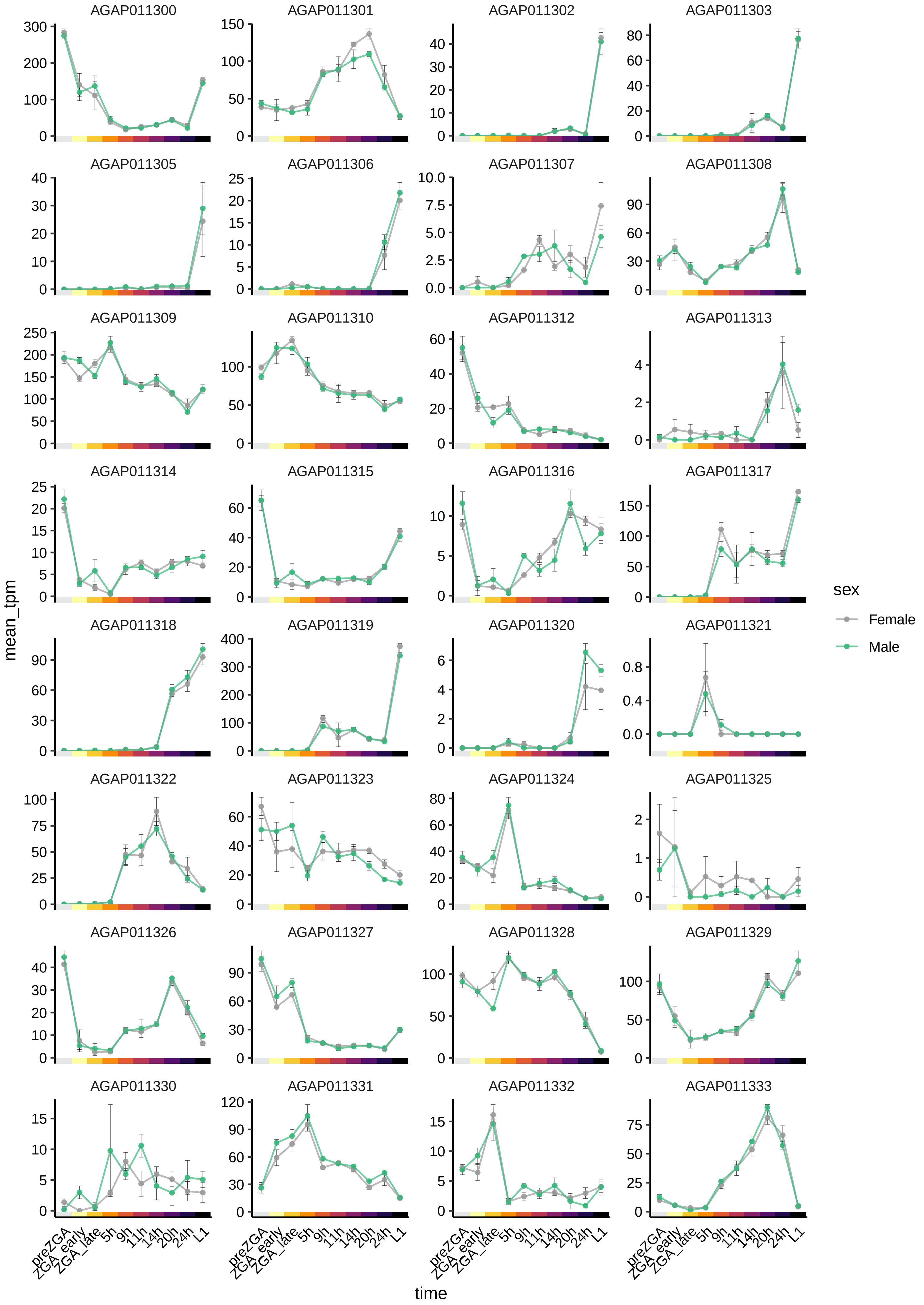

AGAP011300
AGAP011301
AGAP011302
AGAP011303
AGAP011305
AGAP011306
AGAP011307
AGAP011308
AGAP011309
AGAP011310
AGAP011312
AGAP011313
AGAP011314
AGAP011315
AGAP011316
AGAP011317
AGAP011318
AGAP011319
AGAP011320
AGAP011321
AGAP011322
AGAP011323
AGAP011324
AGAP011325
AGAP011326
AGAP011327
AGAP011328
AGAP011329
AGAP011330
AGAP011331
AGAP011332
AGAP011333
mean_tpm
time
preZGA
ZGA_early
ZGA_late
5h
9h
11h
14h
20h
24h
L1
sex
Female
Male

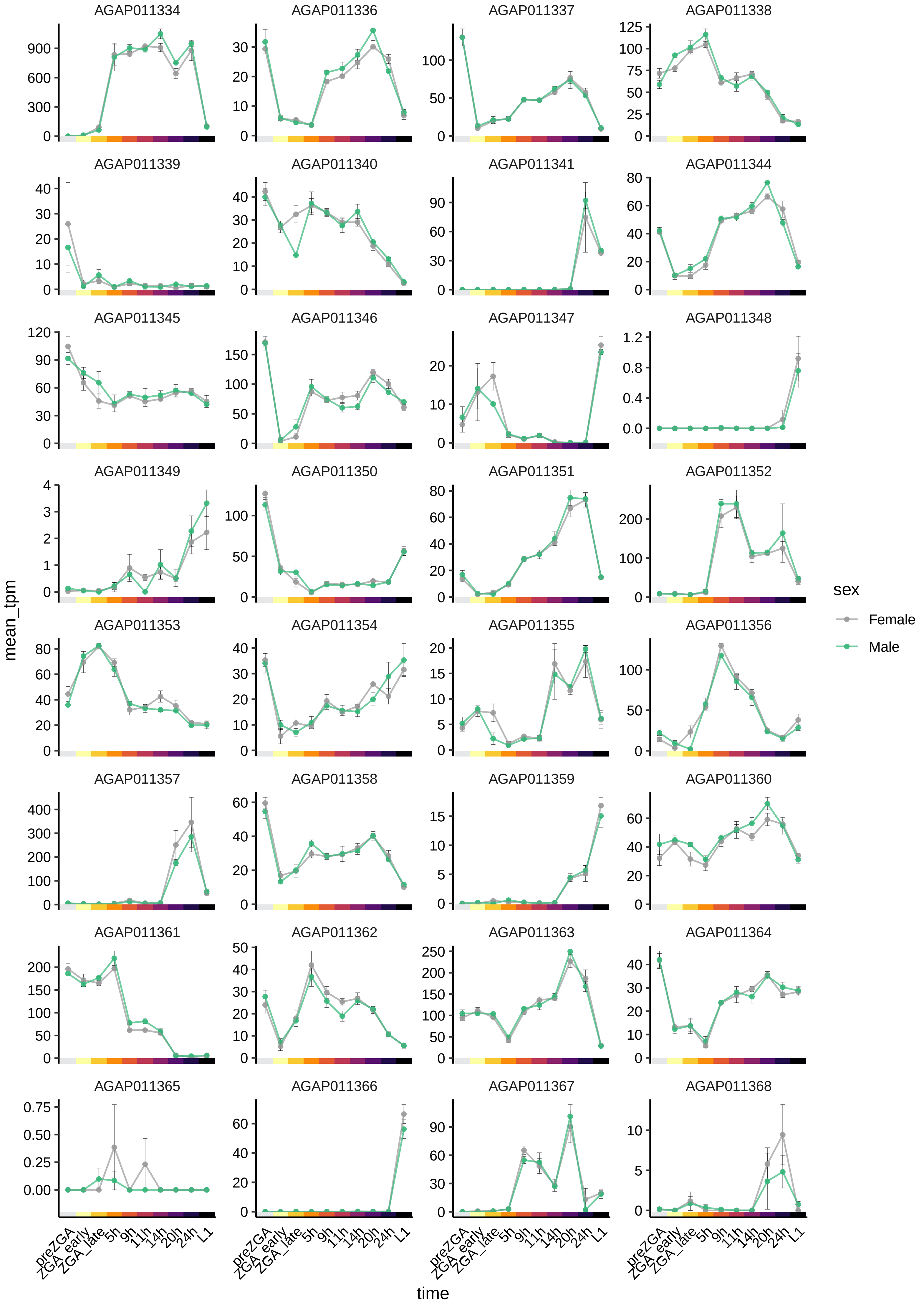
AGAP011334
AGAP011336
AGAP011337
AGAP011338
AGAP011339
AGAP011340
AGAP011341
AGAP011344
AGAP011345
AGAP011346
AGAP011347
AGAP011348
AGAP011349
AGAP011350
AGAP011351
AGAP011352
AGAP011353
AGAP011354
AGAP011355
AGAP011356
AGAP011357
AGAP011358
AGAP011359
AGAP011360
AGAP011361
AGAP011362
AGAP011363
AGAP011364
AGAP011365
AGAP011366
AGAP011367
AGAP011368
mean_tpm
time
preZGA
ZGA_early
ZGA_late
5h
9h
11h
14h
20h
24h
L1
sex
Female
Male

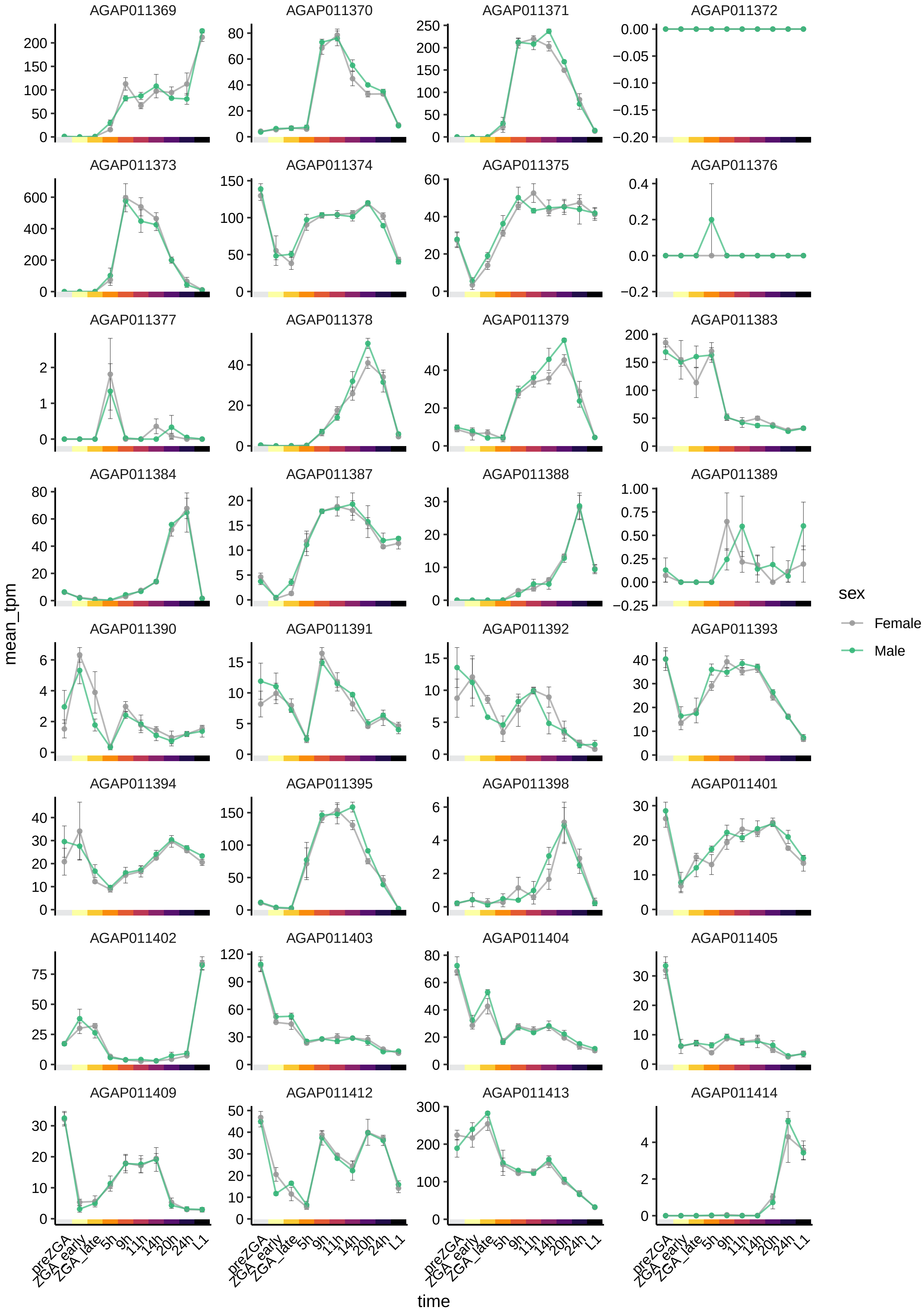

AGAP011369
AGAP011370
AGAP011371
AGAP011372
AGAP011373
AGAP011374
AGAP011375
AGAP011376
AGAP011377
AGAP011378
AGAP011379
AGAP011383
AGAP011384
AGAP011387
AGAP011388
AGAP011389
AGAP011390
AGAP011391
AGAP011392
AGAP011393
AGAP011394
AGAP011395
AGAP011398
AGAP011401
AGAP011402
AGAP011403
AGAP011404
AGAP011405
AGAP011409
AGAP011412
AGAP011413
AGAP011414
mean_tpm
time
preZGA
ZGA_early
ZGA_late
5h
9h
11h
14h
20h
24h
L1
sex
Female
Male

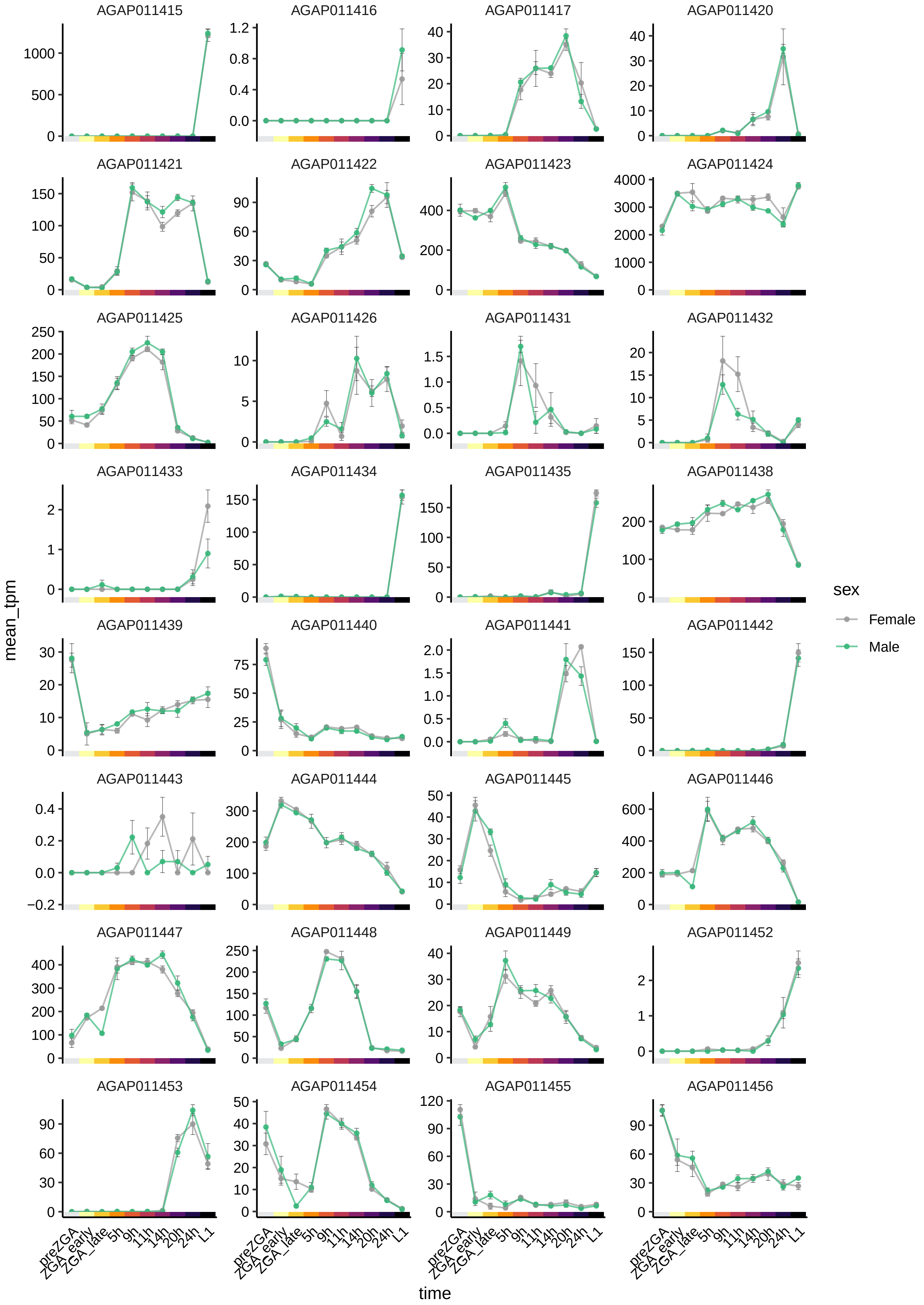

AGAP011415
AGAP011416
AGAP011417
AGAP011420
AGAP011421
AGAP011422
AGAP011423
AGAP011424
AGAP011425
AGAP011426
AGAP011431
AGAP011432
AGAP011433
AGAP011434
AGAP011435
AGAP011438
AGAP011439
AGAP011440
AGAP011441
AGAP011442
AGAP011443
AGAP011444
AGAP011445
AGAP011446
AGAP011447
AGAP011448
AGAP011449
AGAP011452
AGAP011453
AGAP011454
AGAP011455
AGAP011456
mean_tpm
time
preZGA
ZGA_early
ZGA_late
5h
9h
11h
14h
20h
24h
L1
sex
Female
Male

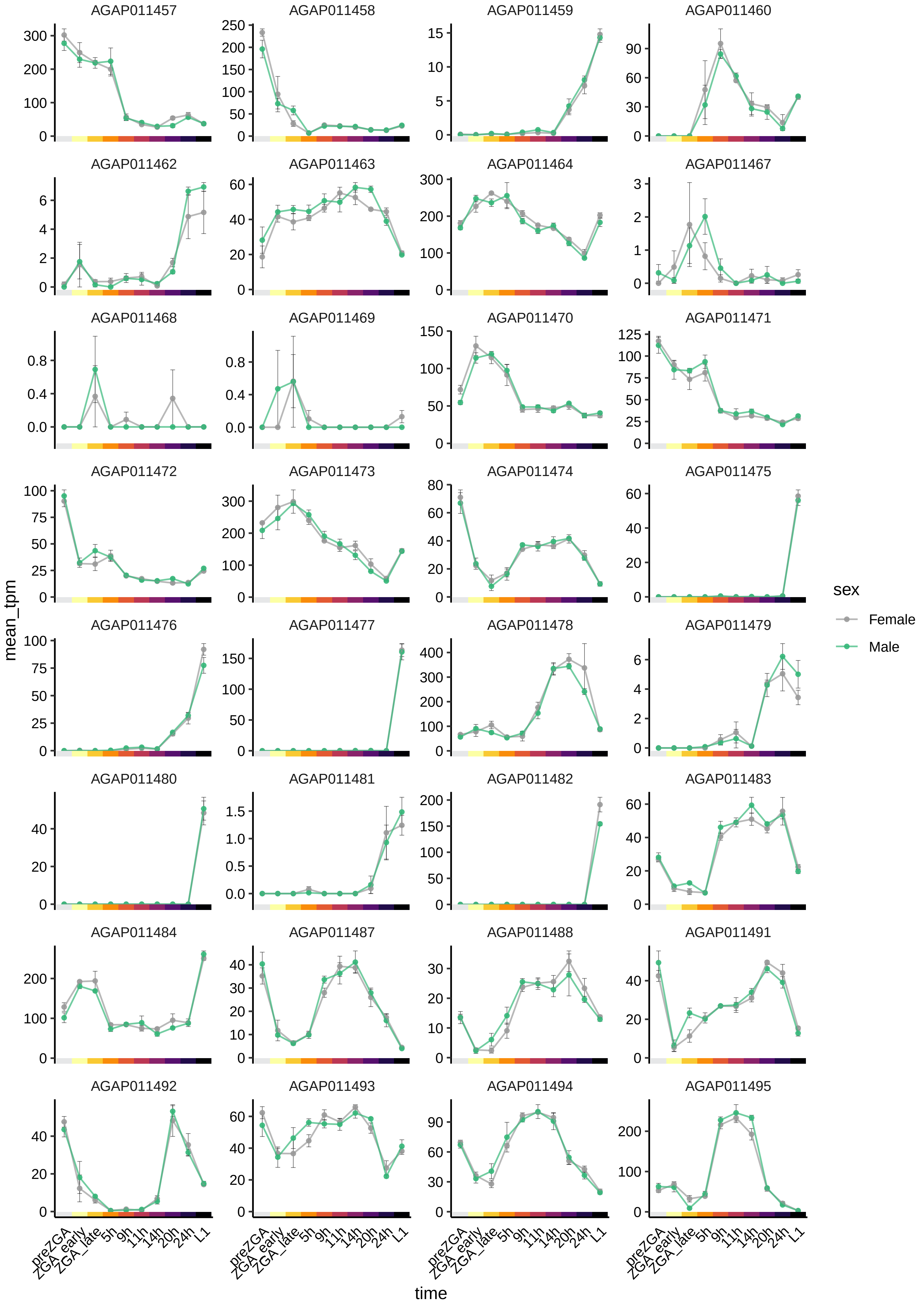

AGAP011457
AGAP011458
AGAP011459
AGAP011460
AGAP011462
AGAP011463
AGAP011464
AGAP011467
AGAP011468
AGAP011469
AGAP011470
AGAP011471
AGAP011472
AGAP011473
AGAP011474
AGAP011475
AGAP011476
AGAP011477
AGAP011478
AGAP011479
AGAP011480
AGAP011481
AGAP011482
AGAP011483
AGAP011484
AGAP011487
AGAP011488
AGAP011491
AGAP011492
AGAP011493
AGAP011494
AGAP011495
mean_tpm
time
preZGA
ZGA_early
ZGA_late
5h
9h
11h
14h
20h
24h
L1
sex
Female
Male

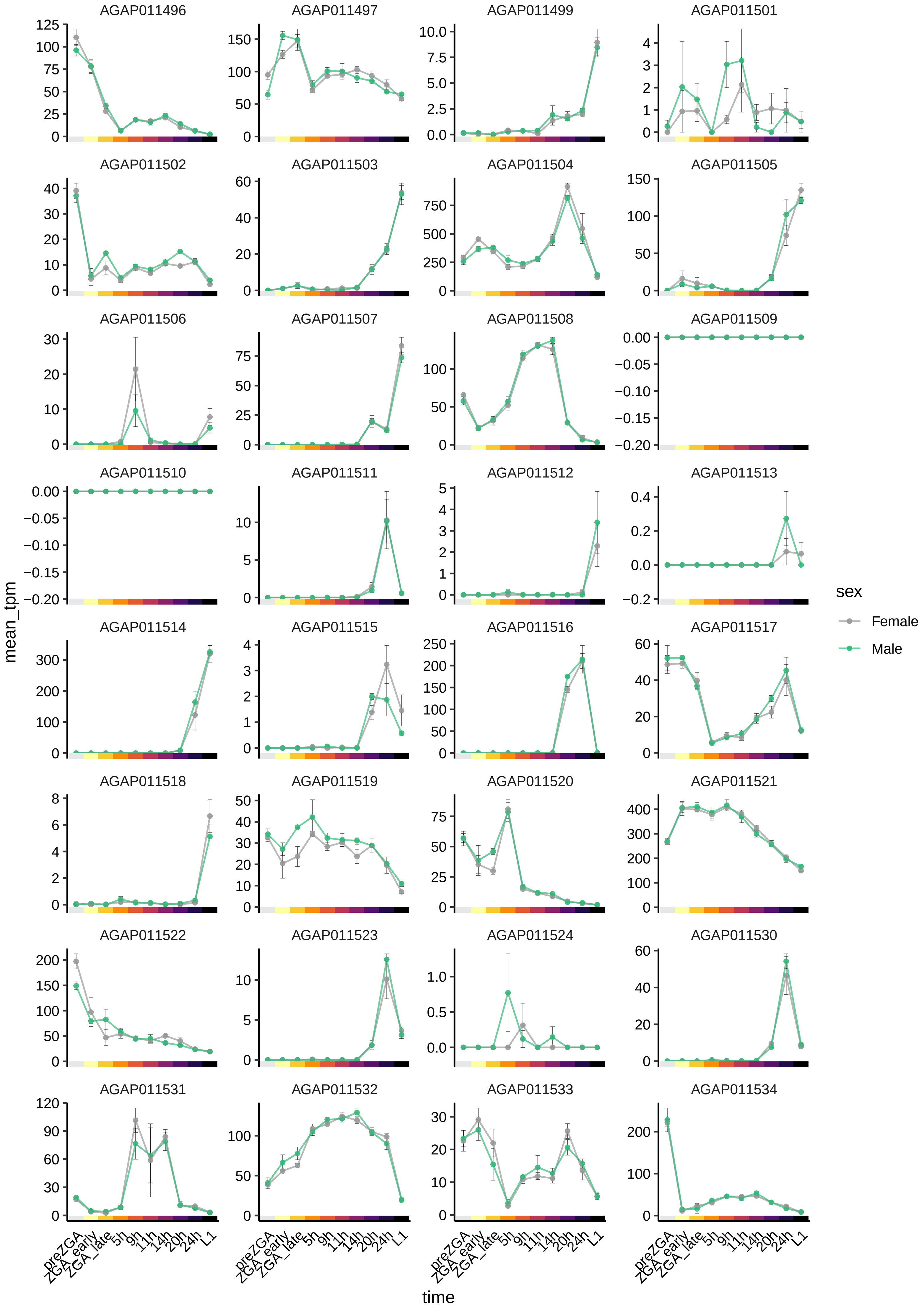
AGAP011496
AGAP011497
AGAP011499
AGAP011501
AGAP011502
AGAP011503
AGAP011504
AGAP011505
AGAP011506
AGAP011507
AGAP011508
AGAP011509
AGAP011510
AGAP011511
AGAP011512
AGAP011513
AGAP011514
AGAP011515
AGAP011516
AGAP011517
AGAP011518
AGAP011519
AGAP011520
AGAP011521
AGAP011522
AGAP011523
AGAP011524
AGAP011530
AGAP011531
AGAP011532
AGAP011533
AGAP011534
mean_tpm
time
preZGA
ZGA_early
ZGA_late
5h
9h
11h
14h
20h
24h
L1
sex
Female
Male

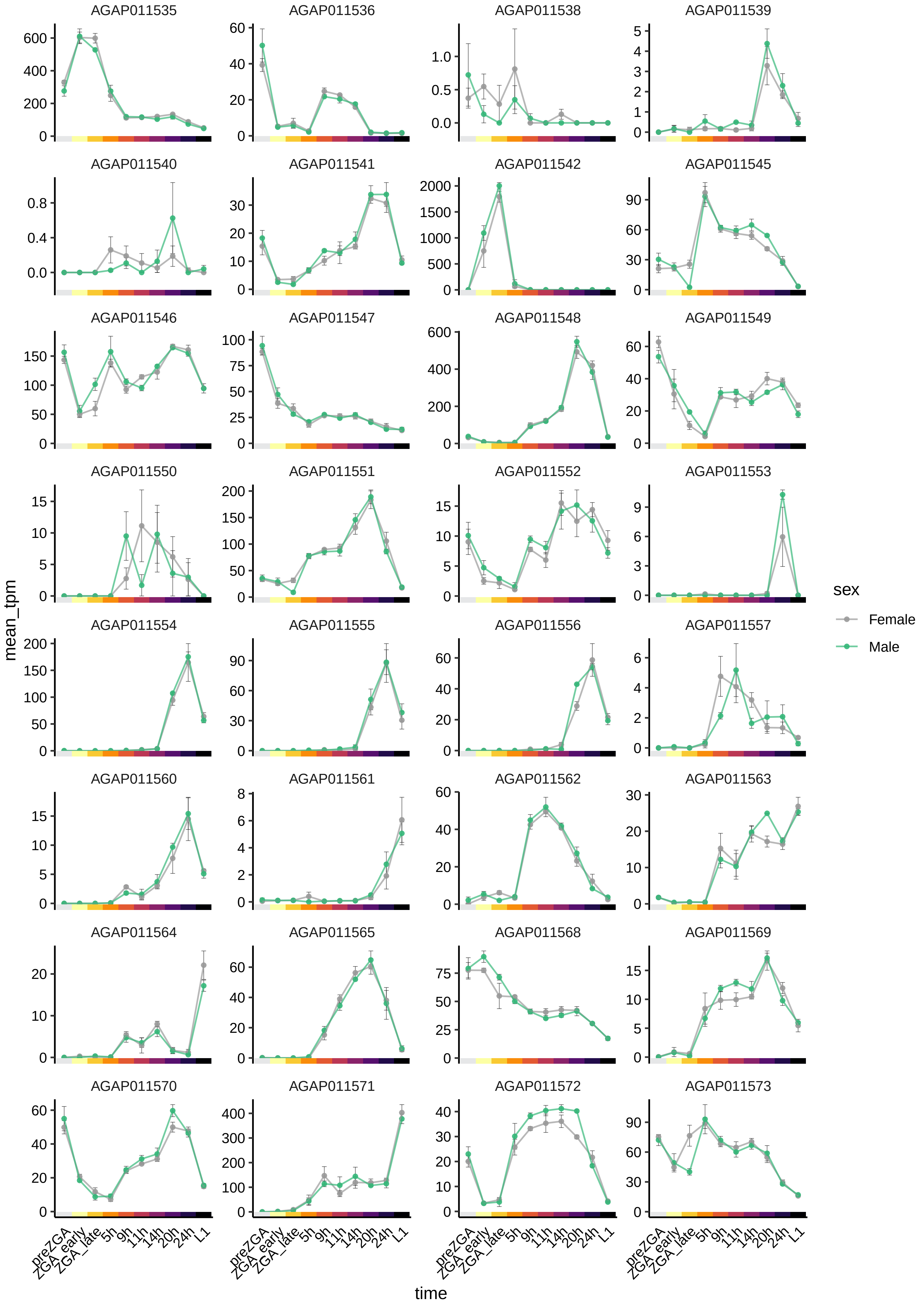

AGAP011535
AGAP011536
AGAP011538
AGAP011539
AGAP011540
AGAP011541
AGAP011542
AGAP011545
AGAP011546
AGAP011547
AGAP011548
AGAP011549
AGAP011550
AGAP011551
AGAP011552
AGAP011553
AGAP011554
AGAP011555
AGAP011556
AGAP011557
AGAP011560
AGAP011561
AGAP011562
AGAP011563
AGAP011564
AGAP011565
AGAP011568
AGAP011569
AGAP011570
AGAP011571
AGAP011572
AGAP011573
mean_tpm
time
preZGA
ZGA_early
ZGA_late
5h
9h
11h
14h
20h
24h
L1
sex
Female
Male

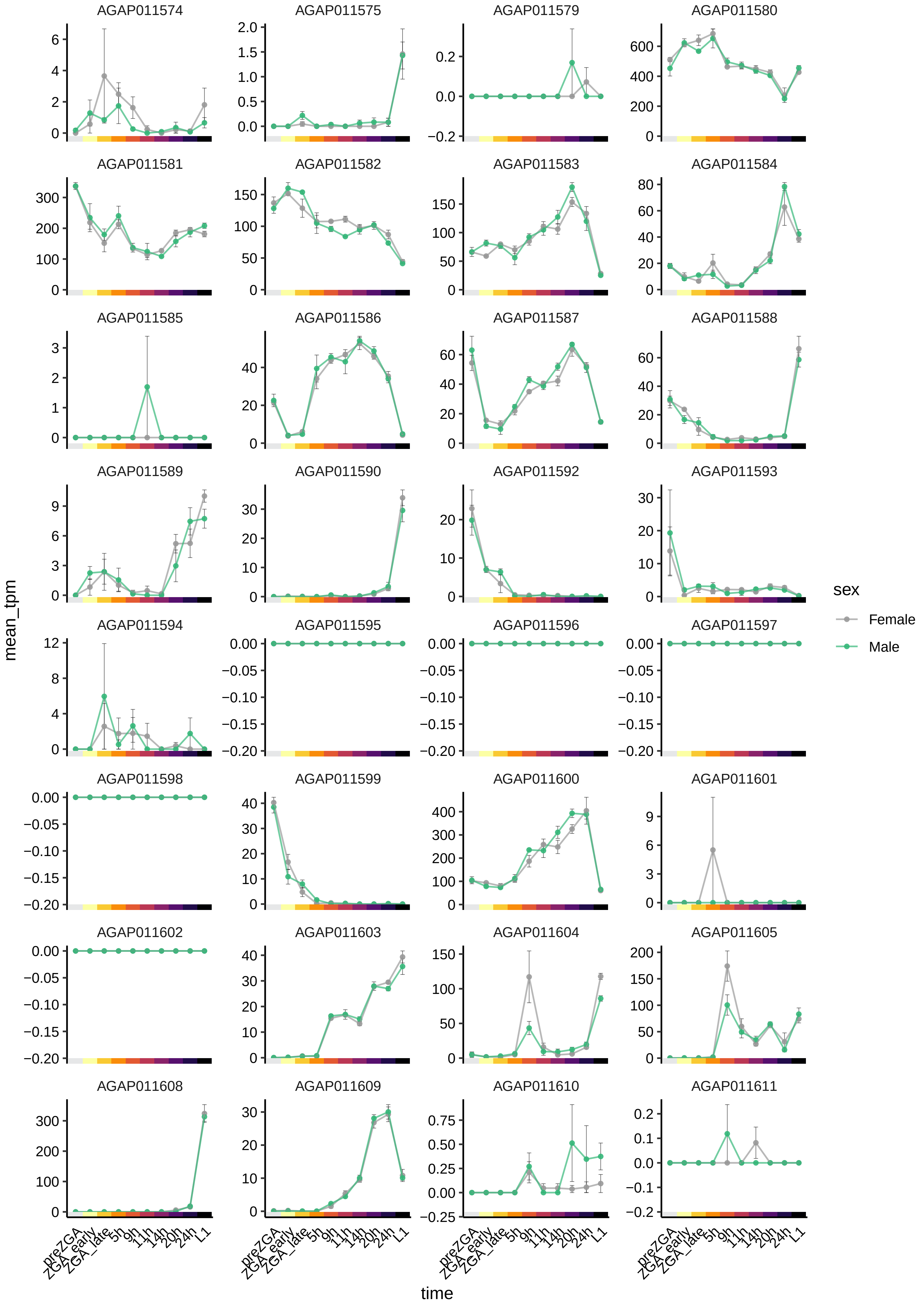

AGAP011574
AGAP011575
AGAP011579
AGAP011580
AGAP011581
AGAP011582
AGAP011583
AGAP011584
AGAP011585
AGAP011586
AGAP011587
AGAP011588
AGAP011589
AGAP011590
AGAP011592
AGAP011593
AGAP011594
AGAP011595
AGAP011596
AGAP011597
AGAP011598
AGAP011599
AGAP011600
AGAP011601
AGAP011602
AGAP011603
AGAP011604
AGAP011605
AGAP011608
AGAP011609
AGAP011610
AGAP011611
mean_tpm
time
preZGA
ZGA_early
ZGA_late
5h
9h
11h
14h
20h
24h
L1
sex
Female
Male

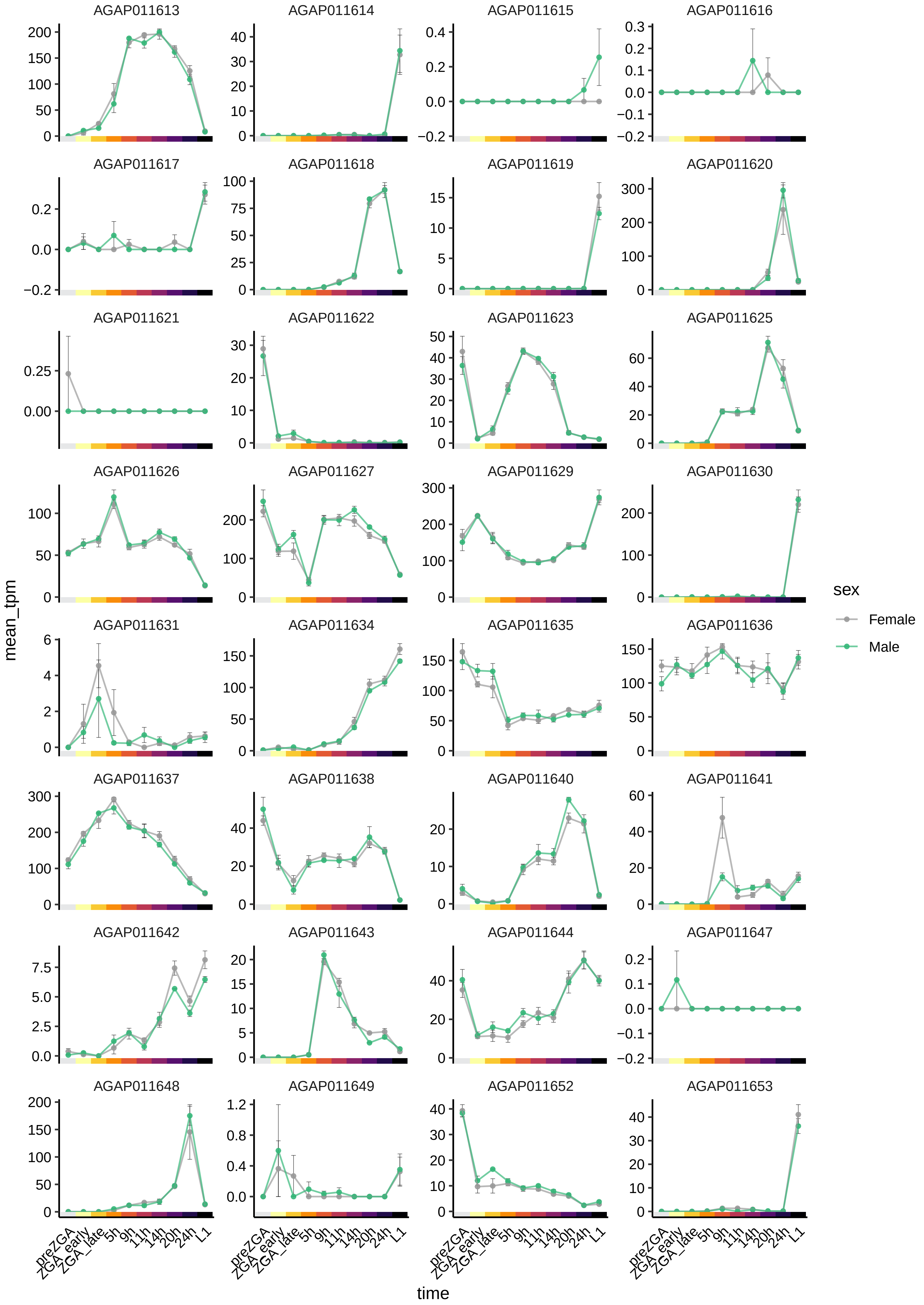

AGAP011613
AGAP011614
AGAP011615
AGAP011616
AGAP011617
AGAP011618
AGAP011619
AGAP011620
AGAP011621
AGAP011622
AGAP011623
AGAP011625
AGAP011626
AGAP011627
AGAP011629
AGAP011630
AGAP011631
AGAP011634
AGAP011635
AGAP011636
AGAP011637
AGAP011638
AGAP011640
AGAP011641
AGAP011642
AGAP011643
AGAP011644
AGAP011647
AGAP011648
AGAP011649
AGAP011652
AGAP011653
mean_tpm
time
preZGA
ZGA_early
ZGA_late
5h
9h
11h
14h
20h
24h
L1
sex
Female
Male

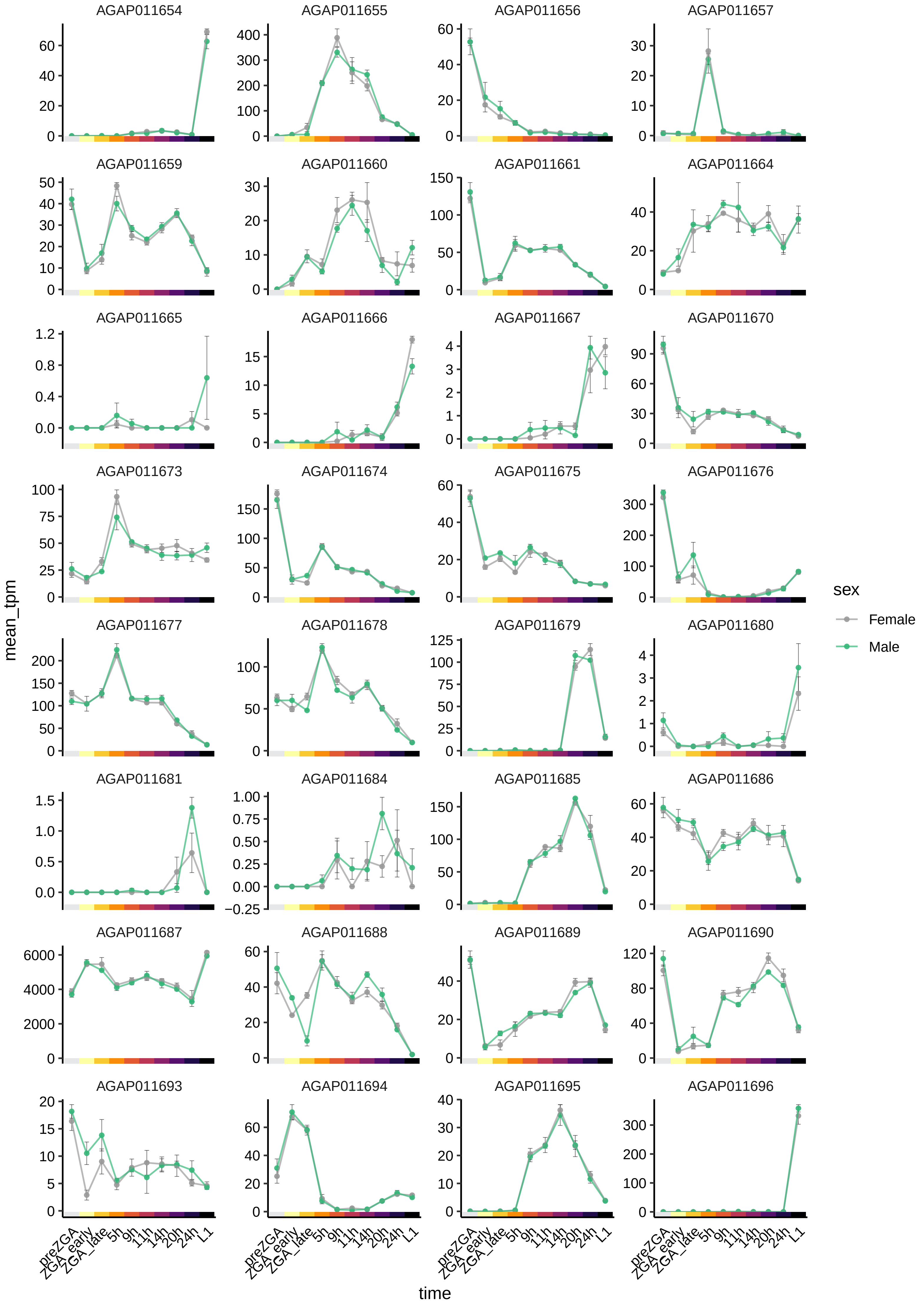

AGAP011654
AGAP011655
AGAP011656
AGAP011657
AGAP011659
AGAP011660
AGAP011661
AGAP011664
AGAP011665
AGAP011666
AGAP011667
AGAP011670
AGAP011673
AGAP011674
AGAP011675
AGAP011676
AGAP011677
AGAP011678
AGAP011679
AGAP011680
AGAP011681
AGAP011684
AGAP011685
AGAP011686
AGAP011687
AGAP011688
AGAP011689
AGAP011690
AGAP011693
AGAP011694
AGAP011695
AGAP011696
mean_tpm
time
preZGA
ZGA_early
ZGA_late
5h
9h
11h
14h
20h
24h
L1
sex
Female
Male

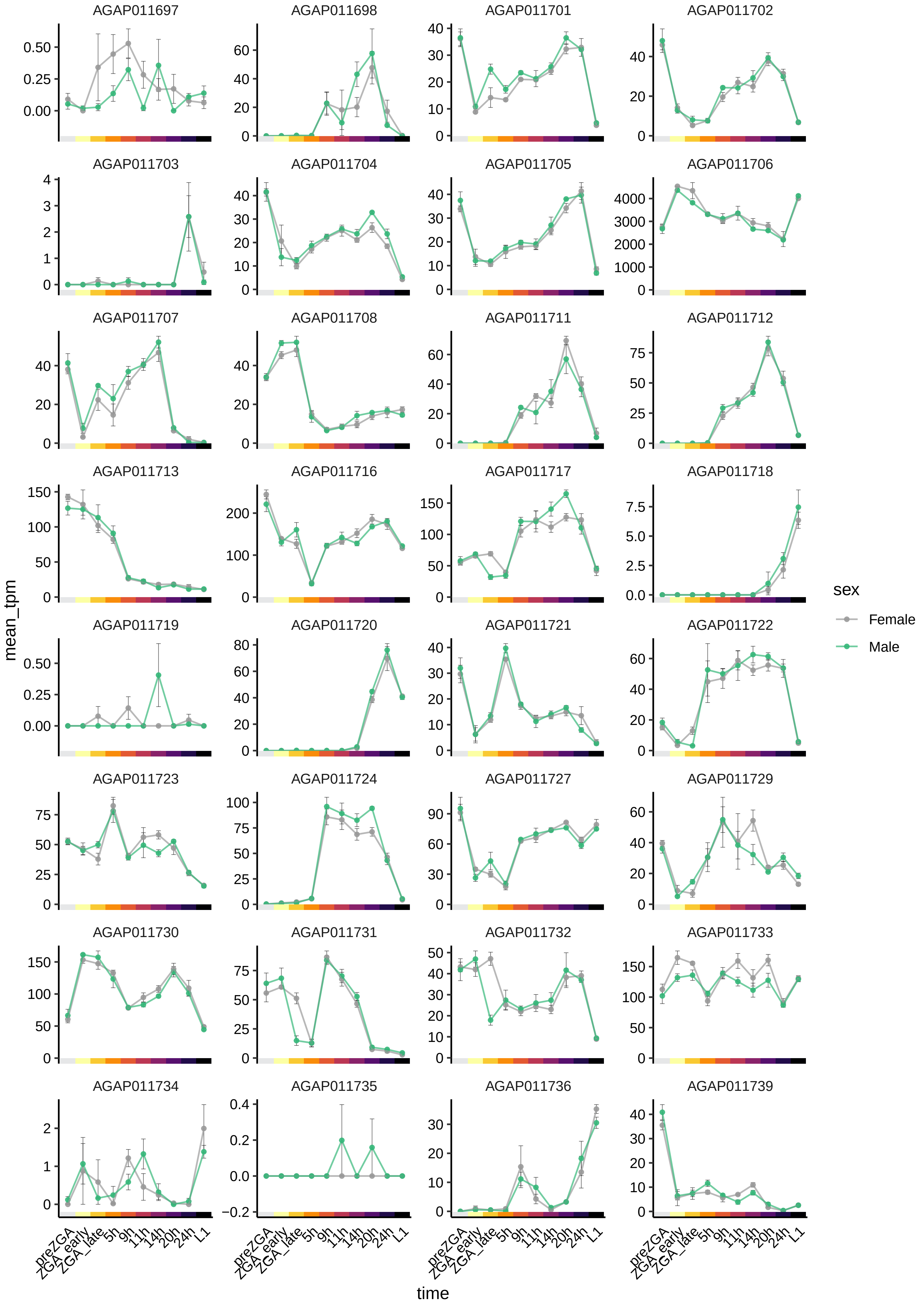

AGAP011697
AGAP011698
AGAP011701
AGAP011702
AGAP011703
AGAP011704
AGAP011705
AGAP011706
AGAP011707
AGAP011708
AGAP011711
AGAP011712
AGAP011713
AGAP011716
AGAP011717
AGAP011718
AGAP011719
AGAP011720
AGAP011721
AGAP011722
AGAP011723
AGAP011724
AGAP011727
AGAP011729
AGAP011730
AGAP011731
AGAP011732
AGAP011733
AGAP011734
AGAP011735
AGAP011736
AGAP011739
mean_tpm
time
preZGA
ZGA_early
ZGA_late
5h
9h
11h
14h
20h
24h
L1
sex
Female
Male

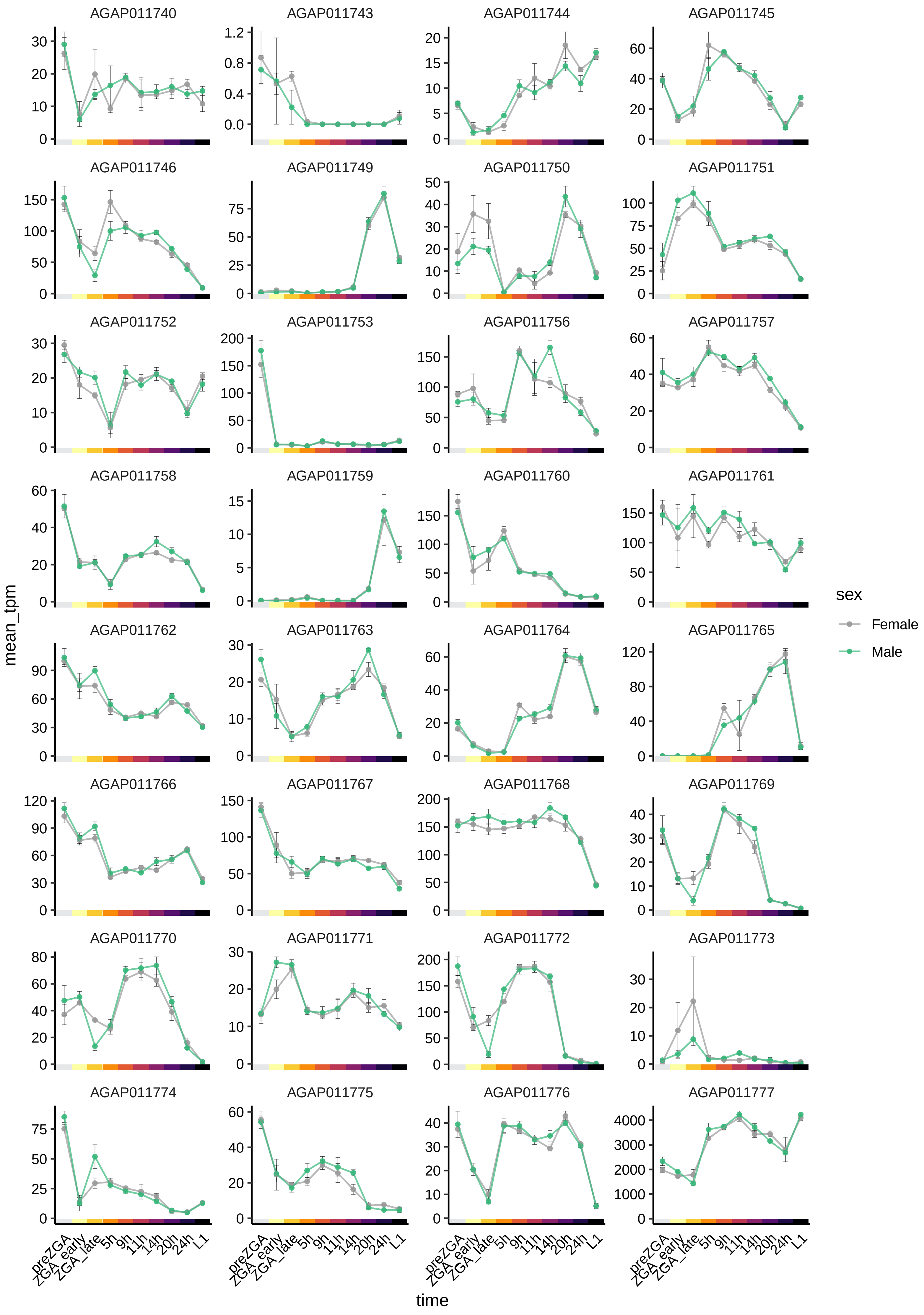

AGAP011740
AGAP011743
AGAP011744
AGAP011745
AGAP011746
AGAP011749
AGAP011750
AGAP011751
AGAP011752
AGAP011753
AGAP011756
AGAP011757
AGAP011758
AGAP011759
AGAP011760
AGAP011761
AGAP011762
AGAP011763
AGAP011764
AGAP011765
AGAP011766
AGAP011767
AGAP011768
AGAP011769
AGAP011770
AGAP011771
AGAP011772
AGAP011773
AGAP011774
AGAP011775
AGAP011776
AGAP011777
mean_tpm
time
preZGA
ZGA_early
ZGA_late
5h
9h
11h
14h
20h
24h
L1
sex
Female
Male

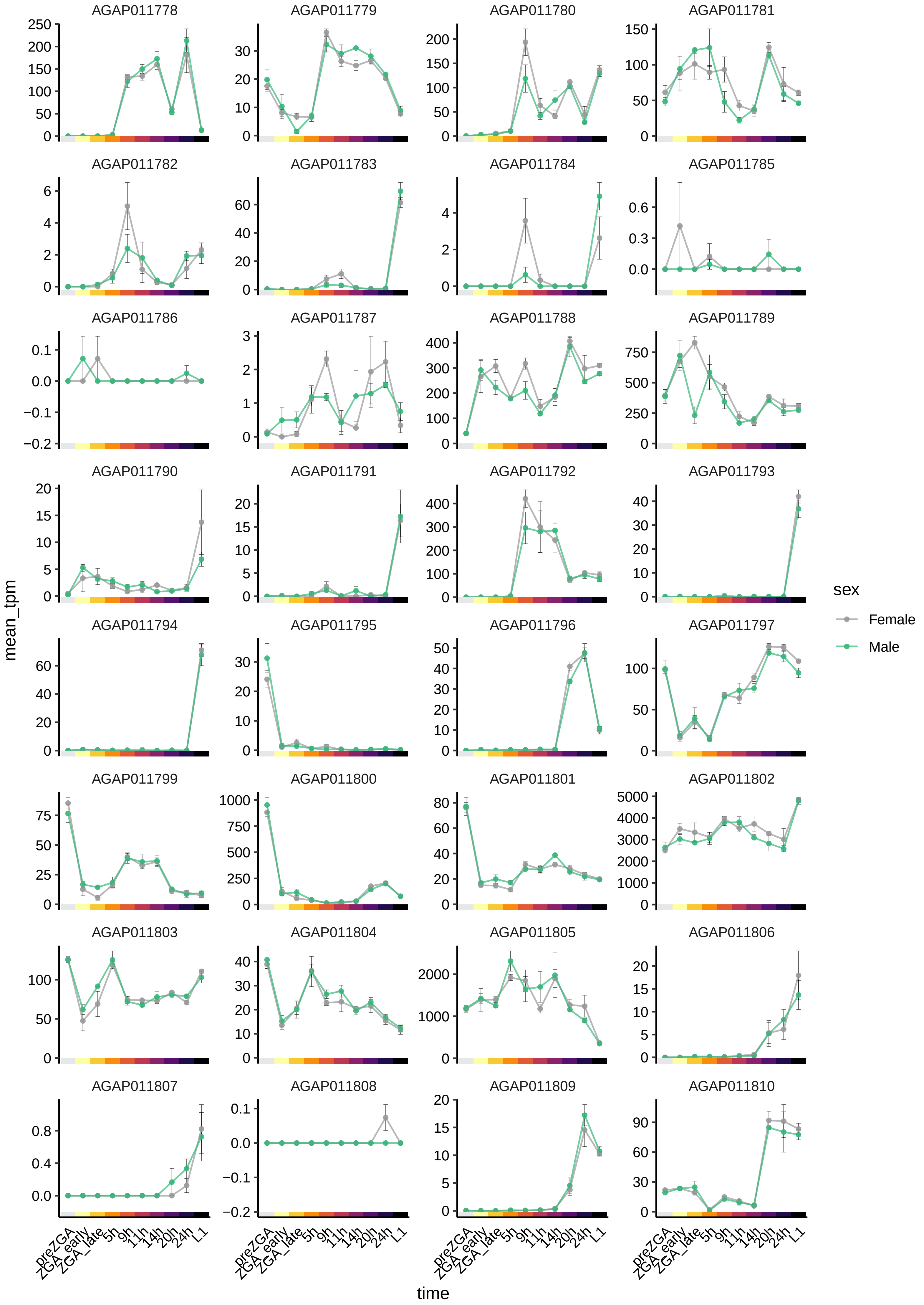

AGAP011778
AGAP011779
AGAP011780
AGAP011781
AGAP011782
AGAP011783
AGAP011784
AGAP011785
AGAP011786
AGAP011787
AGAP011788
AGAP011789
AGAP011790
AGAP011791
AGAP011792
AGAP011793
AGAP011794
AGAP011795
AGAP011796
AGAP011797
AGAP011799
AGAP011800
AGAP011801
AGAP011802
AGAP011803
AGAP011804
AGAP011805
AGAP011806
AGAP011807
AGAP011808
AGAP011809
AGAP011810
mean_tpm
time
preZGA
ZGA_early
ZGA_late
5h
9h
11h
14h
20h
24h
L1
sex
Female
Male

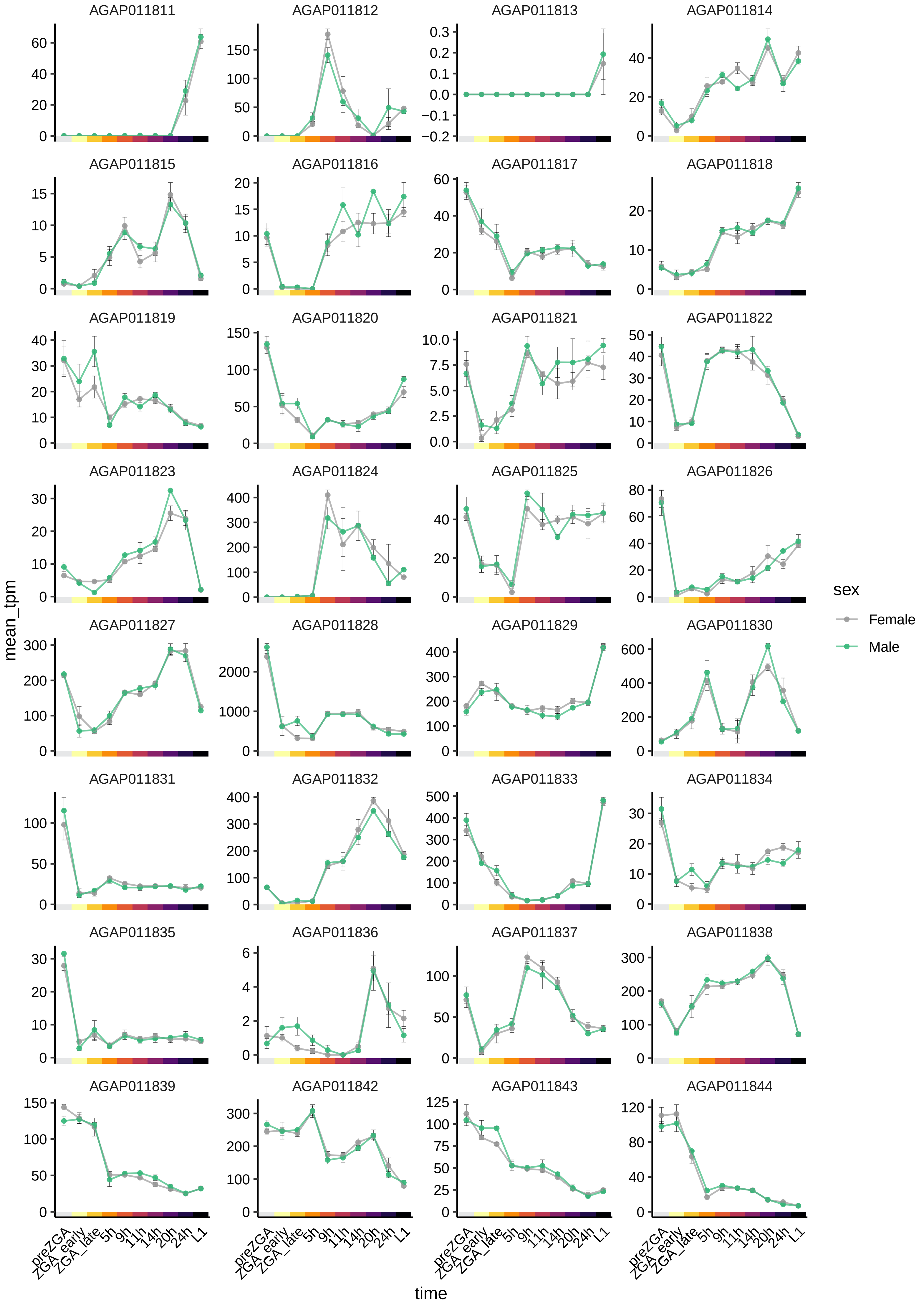

AGAP011811
AGAP011812
AGAP011813
AGAP011814
AGAP011815
AGAP011816
AGAP011817
AGAP011818
AGAP011819
AGAP011820
AGAP011821
AGAP011822
AGAP011823
AGAP011824
AGAP011825
AGAP011826
AGAP011827
AGAP011828
AGAP011829
AGAP011830
AGAP011831
AGAP011832
AGAP011833
AGAP011834
AGAP011835
AGAP011836
AGAP011837
AGAP011838
AGAP011839
AGAP011842
AGAP011843
AGAP011844
mean_tpm
time
preZGA
ZGA_early
ZGA_late
5h
9h
11h
14h
20h
24h
L1
sex
Female
Male

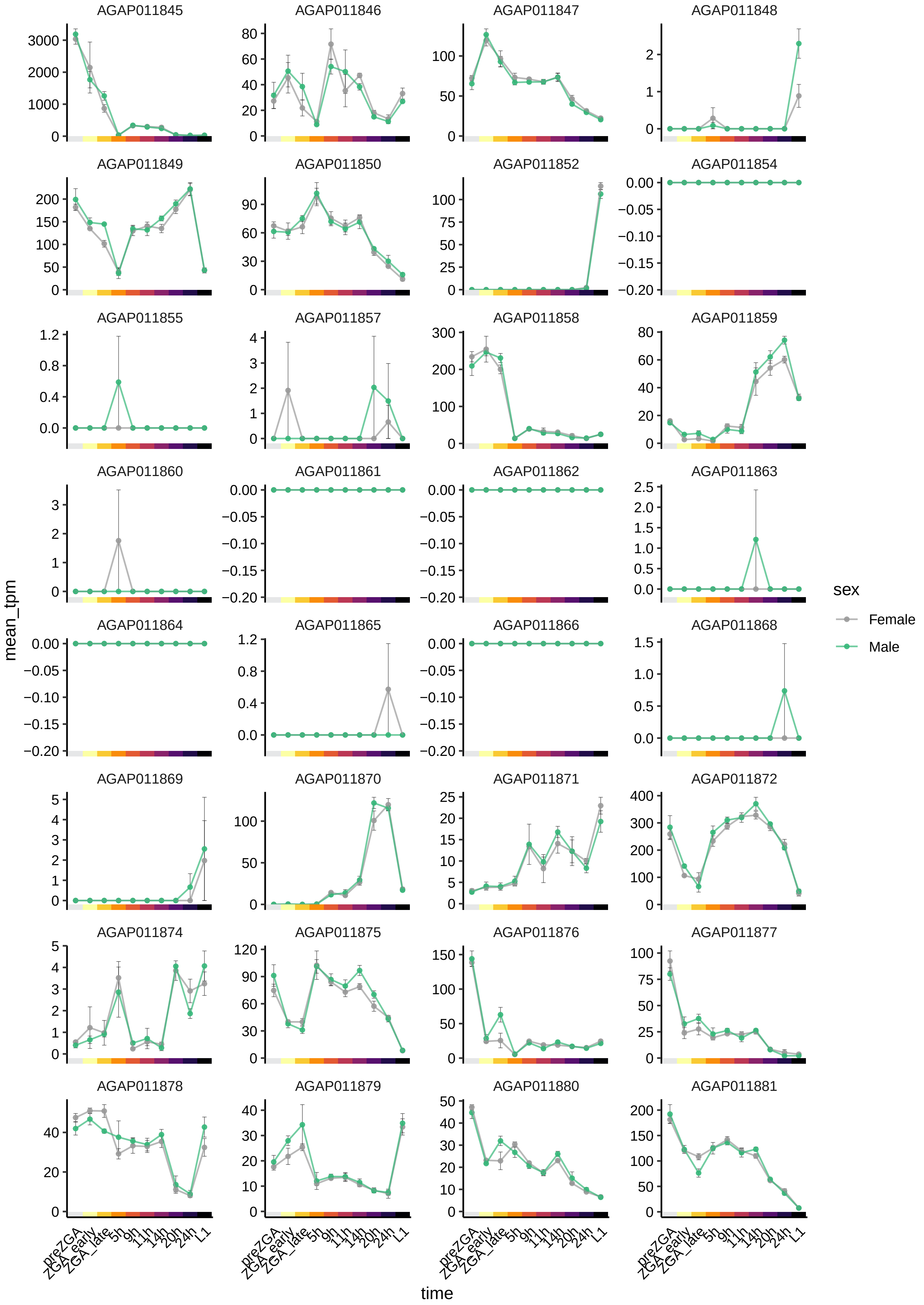
AGAP011845
AGAP011846
AGAP011847
AGAP011848
AGAP011849
AGAP011850
AGAP011852
AGAP011854
AGAP011855
AGAP011857
AGAP011858
AGAP011859
AGAP011860
AGAP011861
AGAP011862
AGAP011863
AGAP011864
AGAP011865
AGAP011866
AGAP011868
AGAP011869
AGAP011870
AGAP011871
AGAP011872
AGAP011874
AGAP011875
AGAP011876
AGAP011877
AGAP011878
AGAP011879
AGAP011880
AGAP011881
mean_tpm
time
preZGA
ZGA_early
ZGA_late
5h
9h
11h
14h
20h
24h
L1
sex
Female
Male

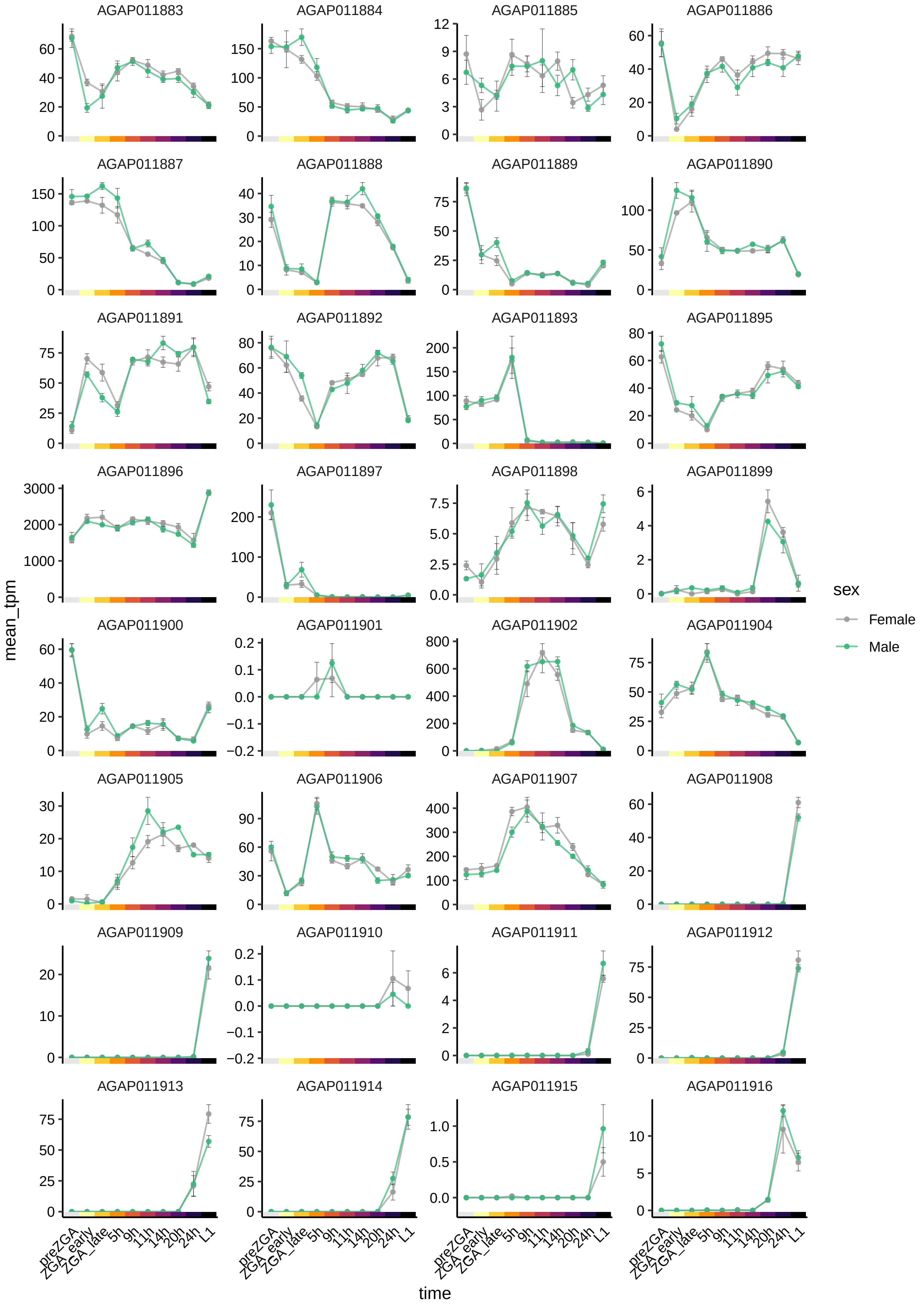

AGAP011883
AGAP011884
AGAP011885
AGAP011886
AGAP011887
AGAP011888
AGAP011889
AGAP011890
AGAP011891
AGAP011892
AGAP011893
AGAP011895
AGAP011896
AGAP011897
AGAP011898
AGAP011899
AGAP011900
AGAP011901
AGAP011902
AGAP011904
AGAP011905
AGAP011906
AGAP011907
AGAP011908
AGAP011909
AGAP011910
AGAP011911
AGAP011912
AGAP011913
AGAP011914
AGAP011915
AGAP011916
mean_tpm
time
preZGA
ZGA_early
ZGA_late
5h
9h
11h
14h
20h
24h
L1
sex
Female
Male

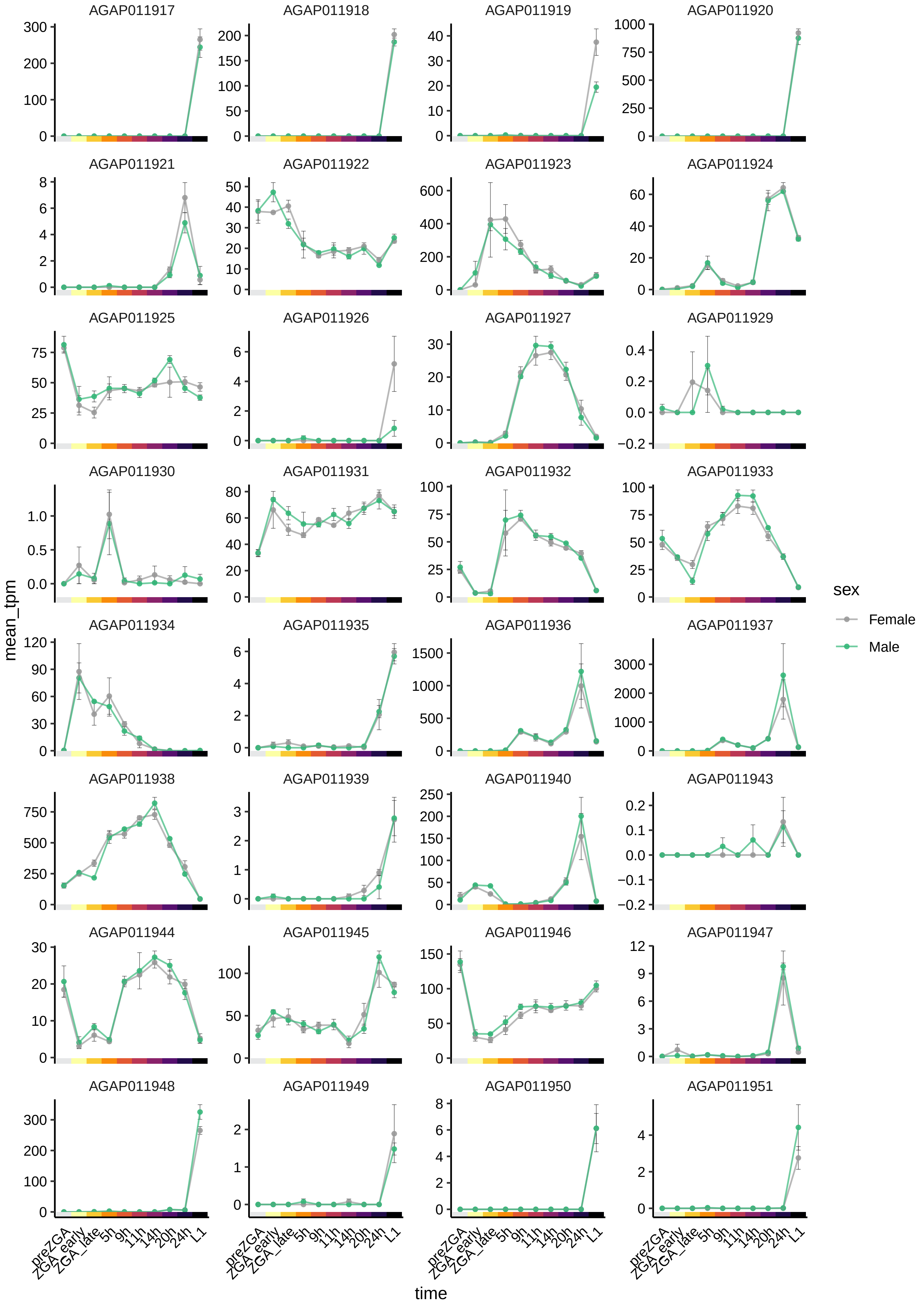

AGAP011917
AGAP011918
AGAP011919
AGAP011920
AGAP011921
AGAP011922
AGAP011923
AGAP011924
AGAP011925
AGAP011926
AGAP011927
AGAP011929
AGAP011930
AGAP011931
AGAP011932
AGAP011933
AGAP011934
AGAP011935
AGAP011936
AGAP011937
AGAP011938
AGAP011939
AGAP011940
AGAP011943
AGAP011944
AGAP011945
AGAP011946
AGAP011947
AGAP011948
AGAP011949
AGAP011950
AGAP011951
mean_tpm
time
preZGA
ZGA_early
ZGA_late
5h
9h
11h
14h
20h
24h
L1
sex
Female
Male

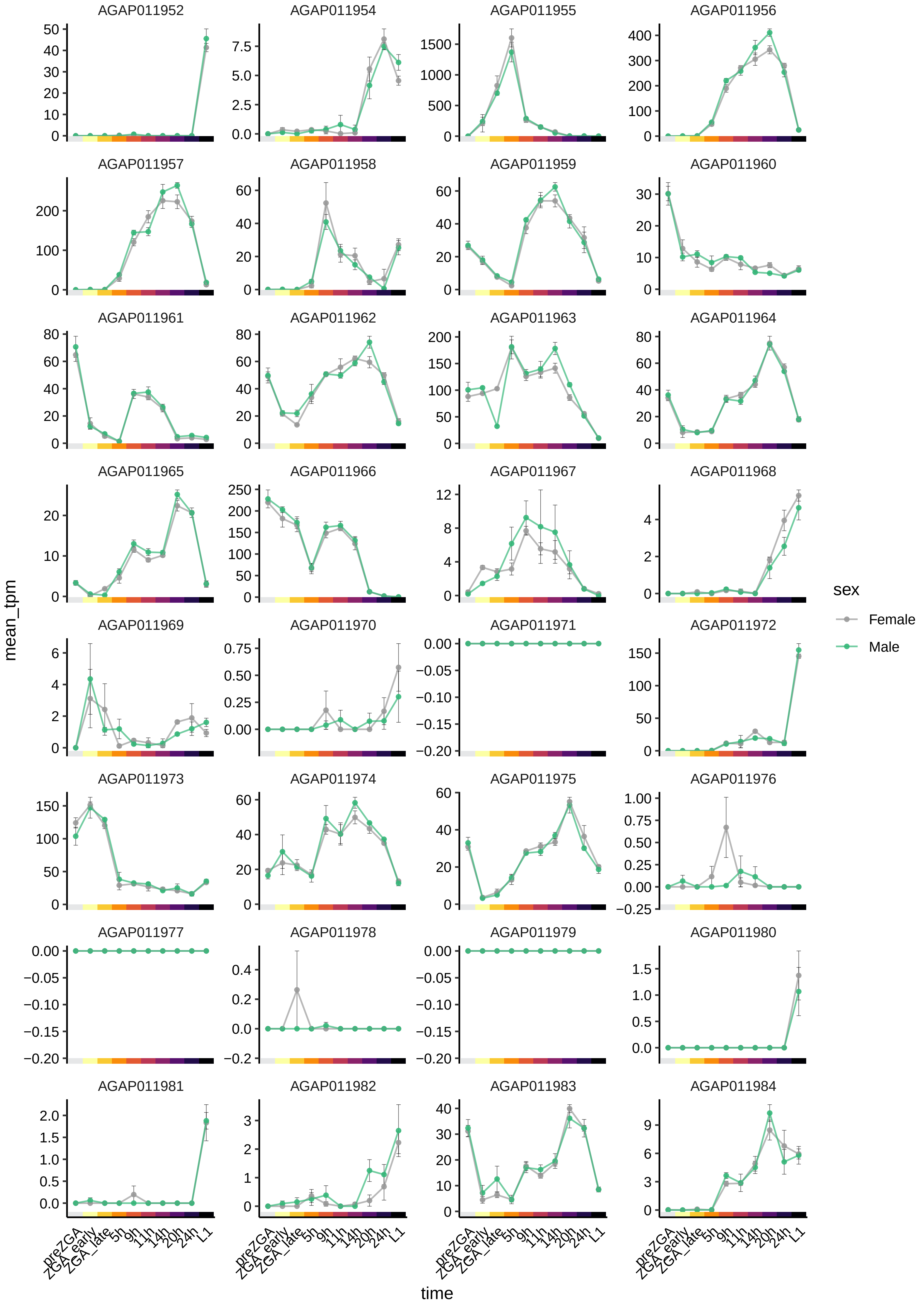

AGAP011952
AGAP011954
AGAP011955
AGAP011956
AGAP011957
AGAP011958
AGAP011959
AGAP011960
AGAP011961
AGAP011962
AGAP011963
AGAP011964
AGAP011965
AGAP011966
AGAP011967
AGAP011968
AGAP011969
AGAP011970
AGAP011971
AGAP011972
AGAP011973
AGAP011974
AGAP011975
AGAP011976
AGAP011977
AGAP011978
AGAP011979
AGAP011980
AGAP011981
AGAP011982
AGAP011983
AGAP011984
mean_tpm
time
preZGA
ZGA_early
ZGA_late
5h
9h
11h
14h
20h
24h
L1
sex
Female
Male

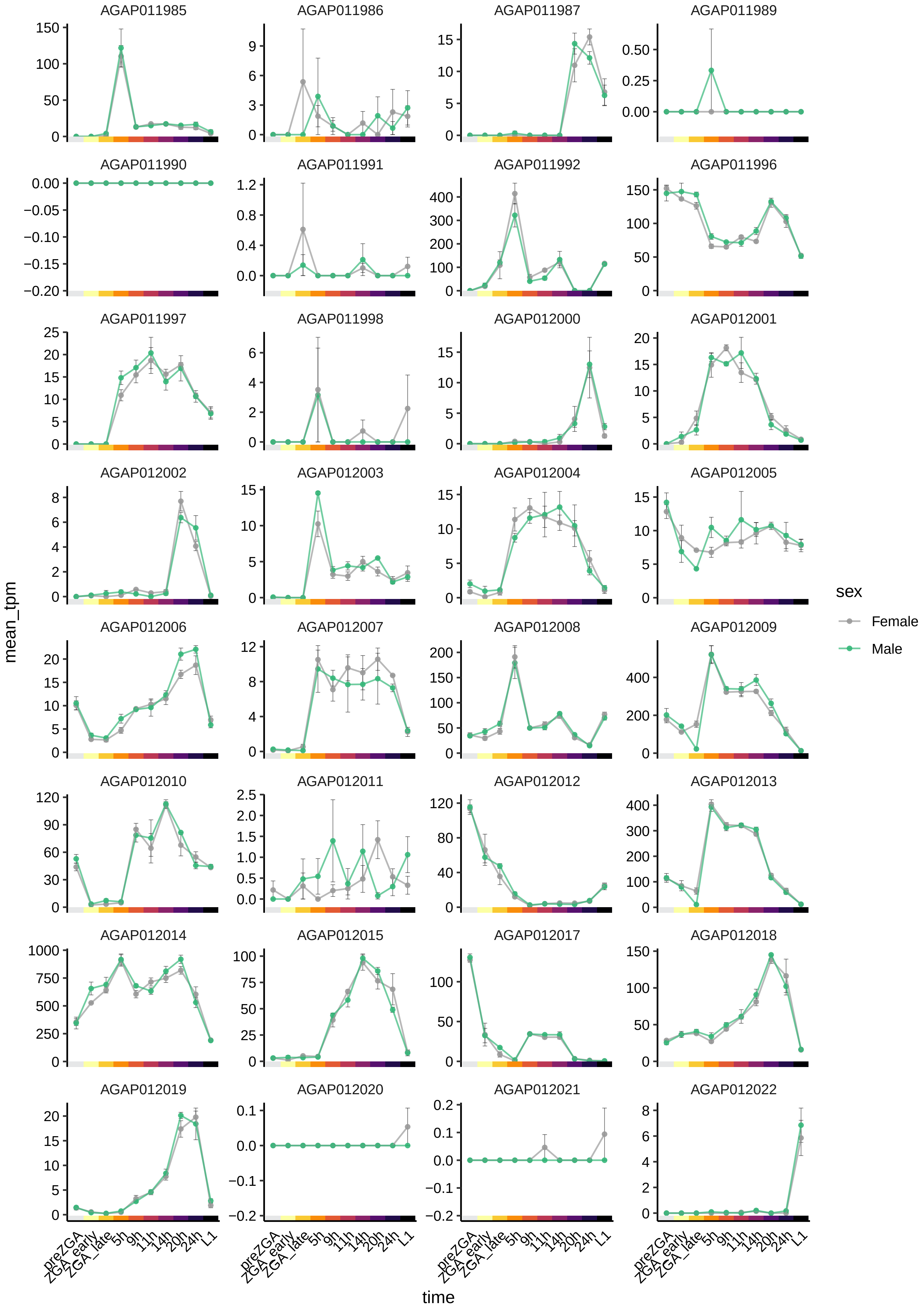

AGAP011985
AGAP011986
AGAP011987
AGAP011989
AGAP011990
AGAP011991
AGAP011992
AGAP011996
AGAP011997
AGAP011998
AGAP012000
AGAP012001
AGAP012002
AGAP012003
AGAP012004
AGAP012005
AGAP012006
AGAP012007
AGAP012008
AGAP012009
AGAP012010
AGAP012011
AGAP012012
AGAP012013
AGAP012014
AGAP012015
AGAP012017
AGAP012018
AGAP012019
AGAP012020
AGAP012021
AGAP012022
mean_tpm
time
preZGA
ZGA_early
ZGA_late
5h
9h
11h
14h
20h
24h
L1
sex
Female
Male

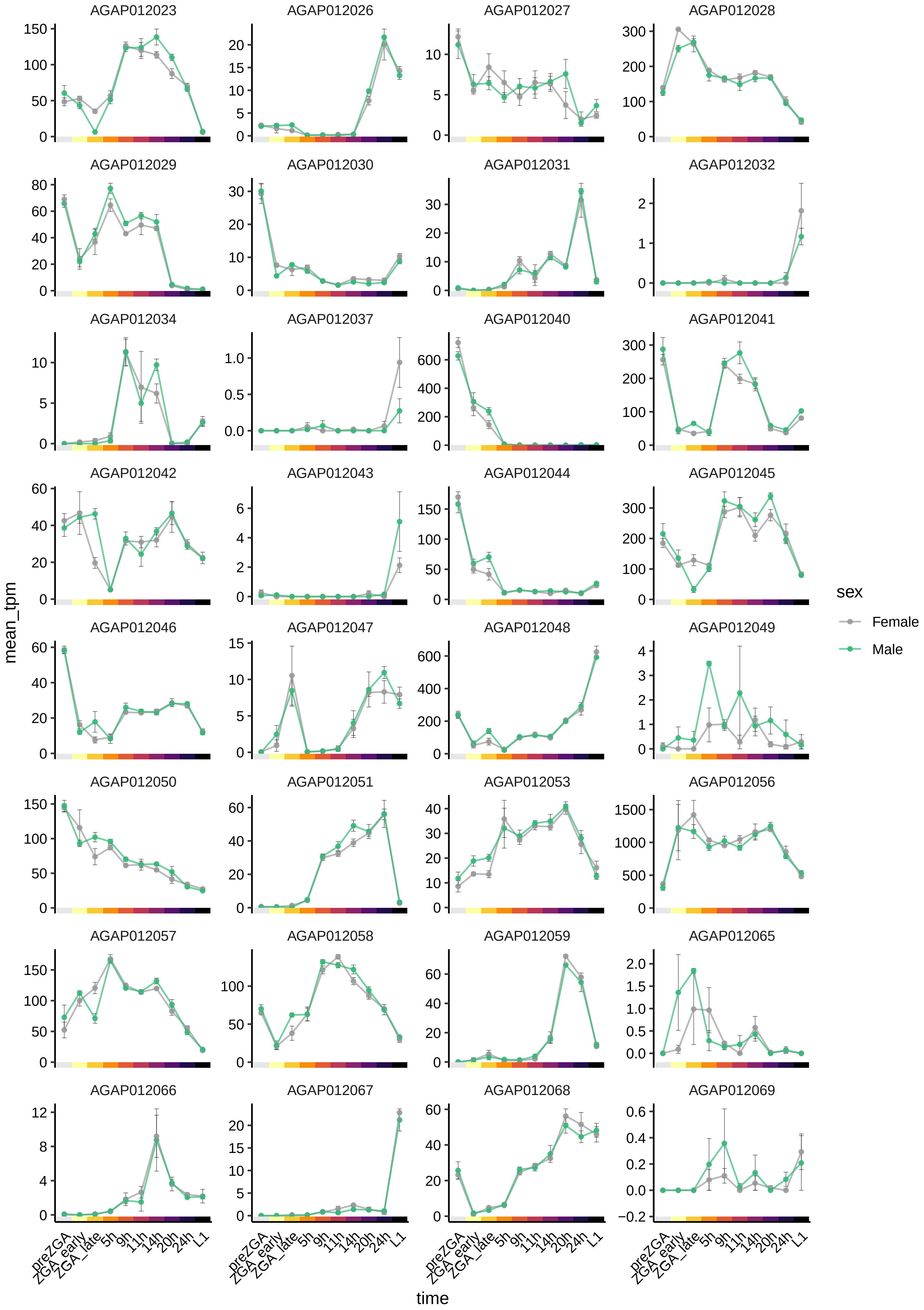
AGAP012023
AGAP012026
AGAP012027
AGAP012028
AGAP012029
AGAP012030
AGAP012031
AGAP012032
AGAP012034
AGAP012037
AGAP012040
AGAP012041
AGAP012042
AGAP012043
AGAP012044
AGAP012045
AGAP012046
AGAP012047
AGAP012048
AGAP012049
AGAP012050
AGAP012051
AGAP012053
AGAP012056
AGAP012057
AGAP012058
AGAP012059
AGAP012065
AGAP012066
AGAP012067
AGAP012068
AGAP012069
mean_tpm
time
preZGA
ZGA_early
ZGA_late
5h
9h
11h
14h
20h
24h
L1
sex
Female
Male

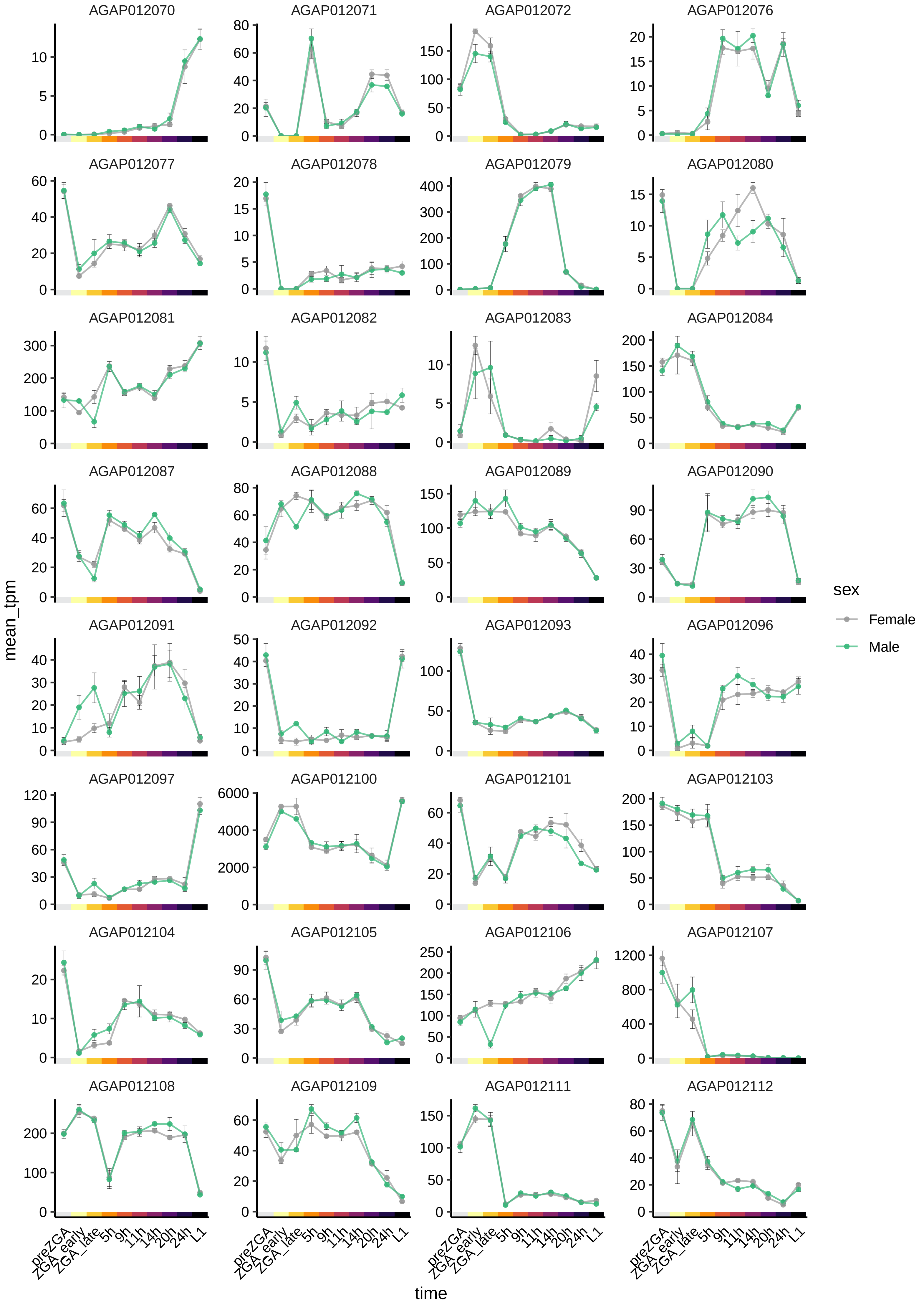

AGAP012070
AGAP012071
AGAP012072
AGAP012076
AGAP012077
AGAP012078
AGAP012079
AGAP012080
AGAP012081
AGAP012082
AGAP012083
AGAP012084
AGAP012087
AGAP012088
AGAP012089
AGAP012090
AGAP012091
AGAP012092
AGAP012093
AGAP012096
AGAP012097
AGAP012100
AGAP012101
AGAP012103
AGAP012104
AGAP012105
AGAP012106
AGAP012107
AGAP012108
AGAP012109
AGAP012111
AGAP012112
mean_tpm
time
preZGA
ZGA_early
ZGA_late
5h
9h
11h
14h
20h
24h
L1
sex
Female
Male

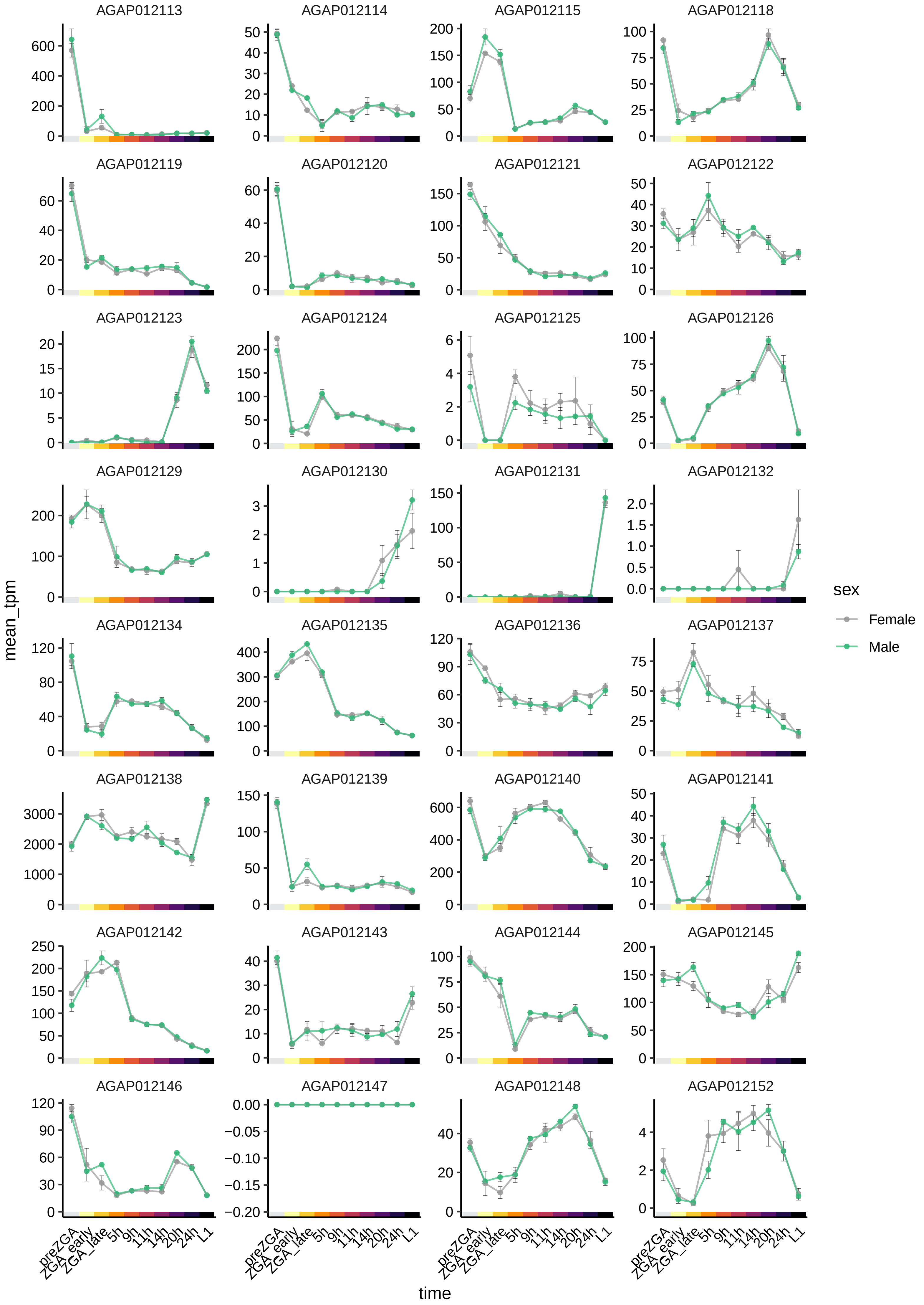

AGAP012113
AGAP012114
AGAP012115
AGAP012118
AGAP012119
AGAP012120
AGAP012121
AGAP012122
AGAP012123
AGAP012124
AGAP012125
AGAP012126
AGAP012129
AGAP012130
AGAP012131
AGAP012132
AGAP012134
AGAP012135
AGAP012136
AGAP012137
AGAP012138
AGAP012139
AGAP012140
AGAP012141
AGAP012142
AGAP012143
AGAP012144
AGAP012145
AGAP012146
AGAP012147
AGAP012148
AGAP012152
mean_tpm
time
preZGA
ZGA_early
ZGA_late
5h
9h
11h
14h
20h
24h
L1
sex
Female
Male

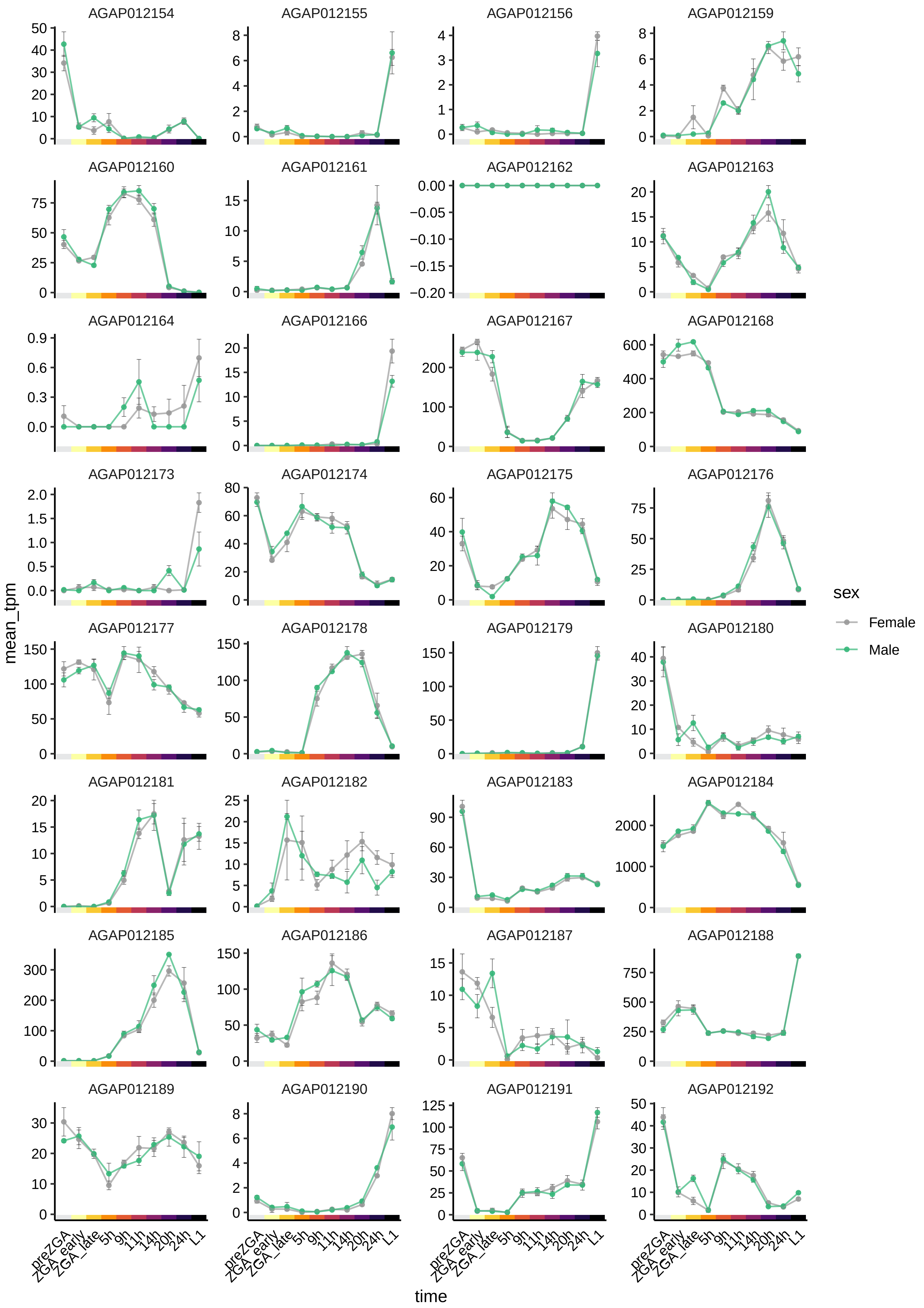

AGAP012154
AGAP012155
AGAP012156
AGAP012159
AGAP012160
AGAP012161
AGAP012162
AGAP012163
AGAP012164
AGAP012166
AGAP012167
AGAP012168
AGAP012173
AGAP012174
AGAP012175
AGAP012176
AGAP012177
AGAP012178
AGAP012179
AGAP012180
AGAP012181
AGAP012182
AGAP012183
AGAP012184
AGAP012185
AGAP012186
AGAP012187
AGAP012188
AGAP012189
AGAP012190
AGAP012191
AGAP012192
mean_tpm
time
preZGA
ZGA_early
ZGA_late
5h
9h
11h
14h
20h
24h
L1
sex
Female
Male

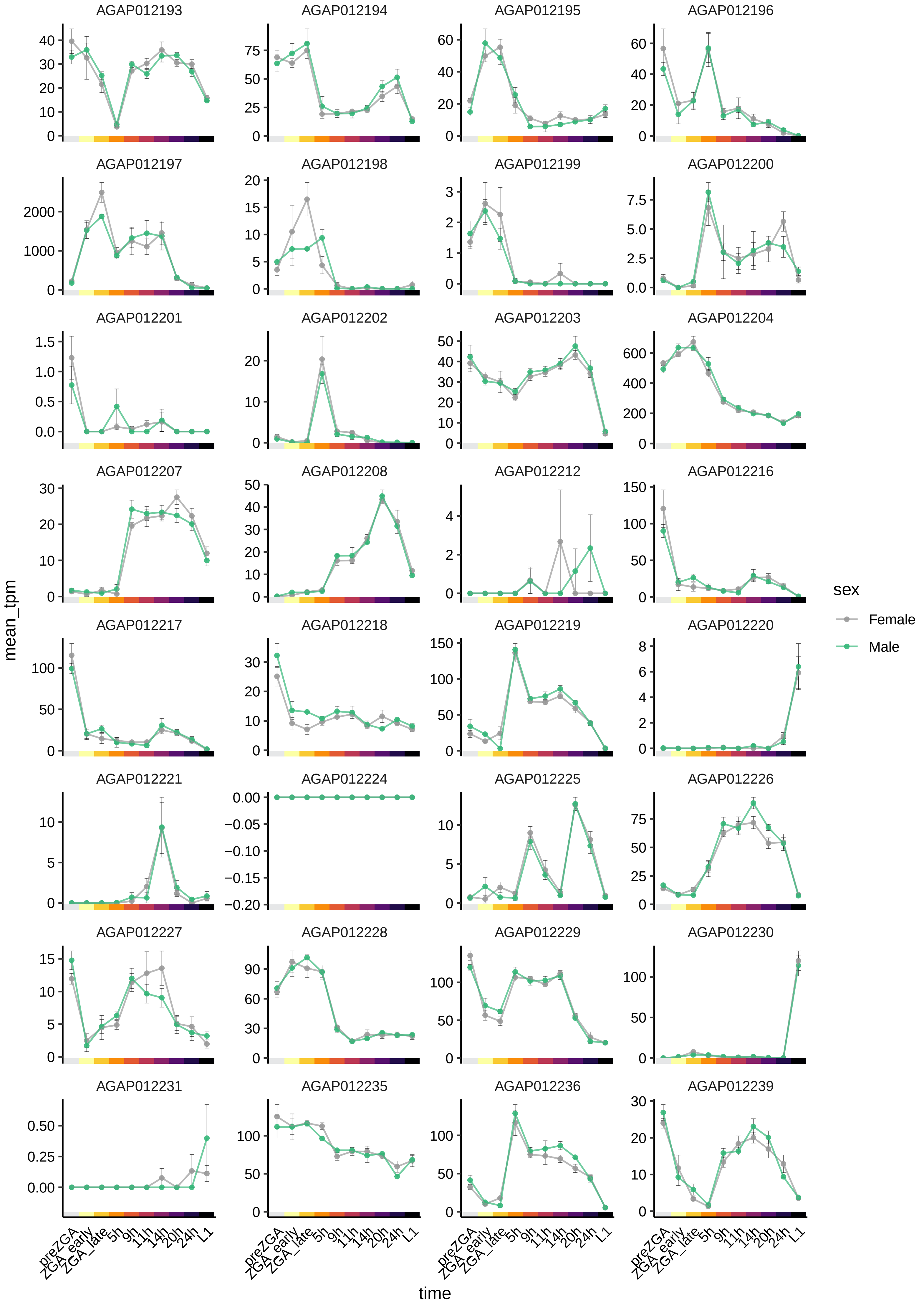

AGAP012193
AGAP012194
AGAP012195
AGAP012196
AGAP012197
AGAP012198
AGAP012199
AGAP012200
AGAP012201
AGAP012202
AGAP012203
AGAP012204
AGAP012207
AGAP012208
AGAP012212
AGAP012216
AGAP012217
AGAP012218
AGAP012219
AGAP012220
AGAP012221
AGAP012224
AGAP012225
AGAP012226
AGAP012227
AGAP012228
AGAP012229
AGAP012230
AGAP012231
AGAP012235
AGAP012236
AGAP012239
mean_tpm
time
preZGA
ZGA_early
ZGA_late
5h
9h
11h
14h
20h
24h
L1
sex
Female
Male

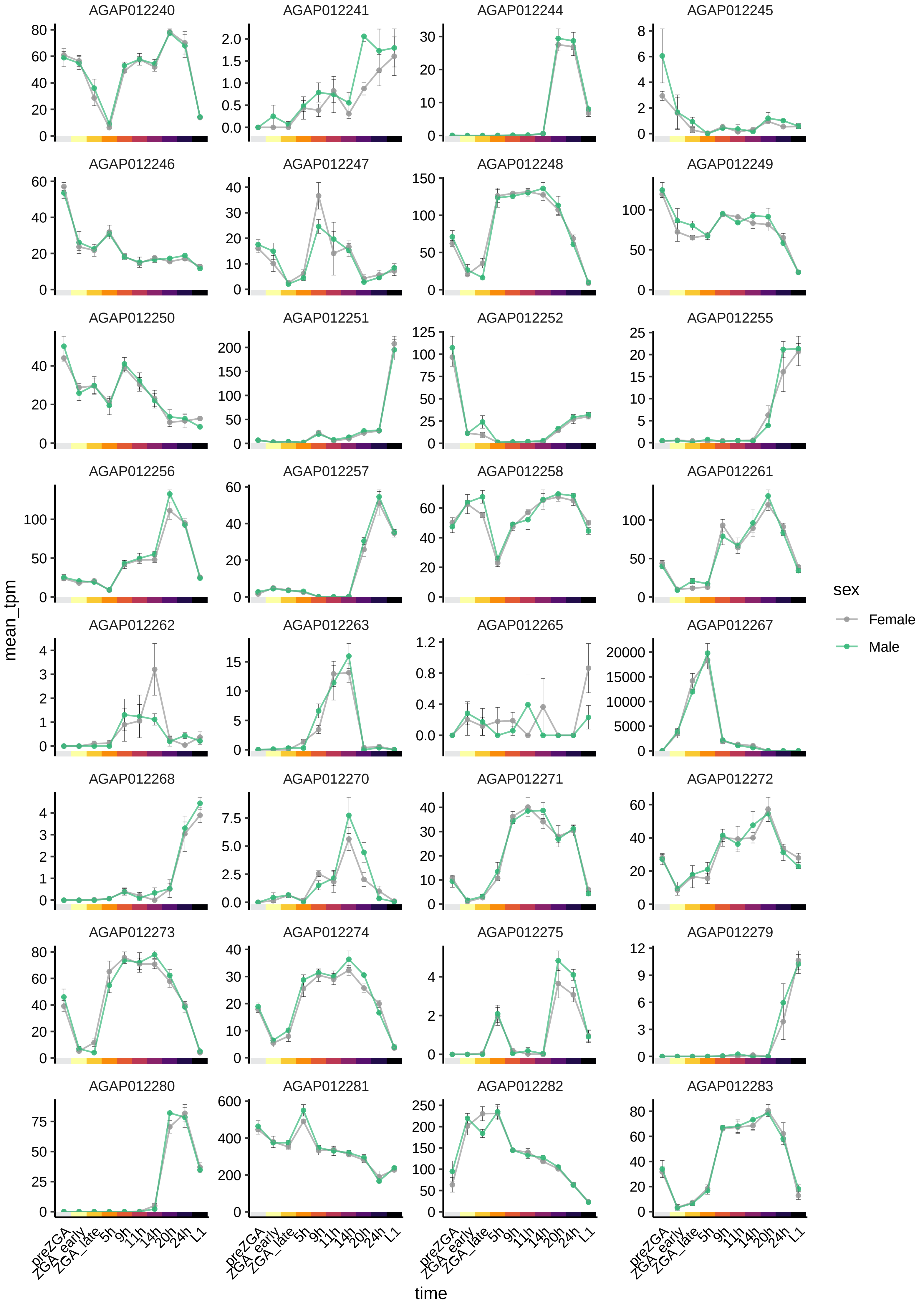
AGAP012240
AGAP012241
AGAP012244
AGAP012245
AGAP012246
AGAP012247
AGAP012248
AGAP012249
AGAP012250
AGAP012251
AGAP012252
AGAP012255
AGAP012256
AGAP012257
AGAP012258
AGAP012261
AGAP012262
AGAP012263
AGAP012265
AGAP012267
AGAP012268
AGAP012270
AGAP012271
AGAP012272
AGAP012273
AGAP012274
AGAP012275
AGAP012279
AGAP012280
AGAP012281
AGAP012282
AGAP012283
mean_tpm
time
preZGA
ZGA_early
ZGA_late
5h
9h
11h
14h
20h
24h
L1
sex
Female
Male

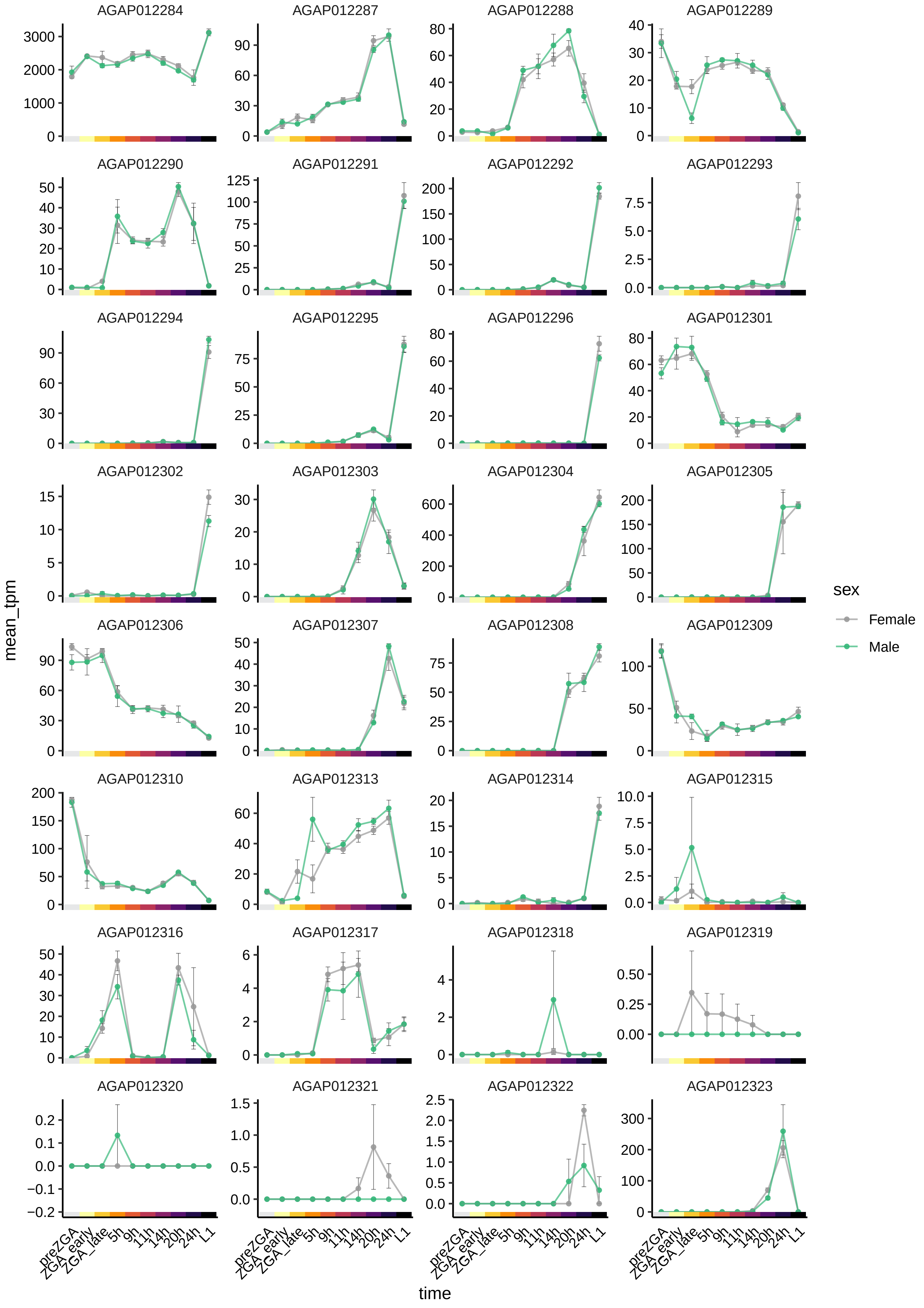
AGAP012284
AGAP012287
AGAP012288
AGAP012289
AGAP012290
AGAP012291
AGAP012292
AGAP012293
AGAP012294
AGAP012295
AGAP012296
AGAP012301
AGAP012302
AGAP012303
AGAP012304
AGAP012305
AGAP012306
AGAP012307
AGAP012308
AGAP012309
AGAP012310
AGAP012313
AGAP012314
AGAP012315
AGAP012316
AGAP012317
AGAP012318
AGAP012319
AGAP012320
AGAP012321
AGAP012322
AGAP012323
mean_tpm
time
preZGA
ZGA_early
ZGA_late
5h
9h
11h
14h
20h
24h
L1
sex
Female
Male

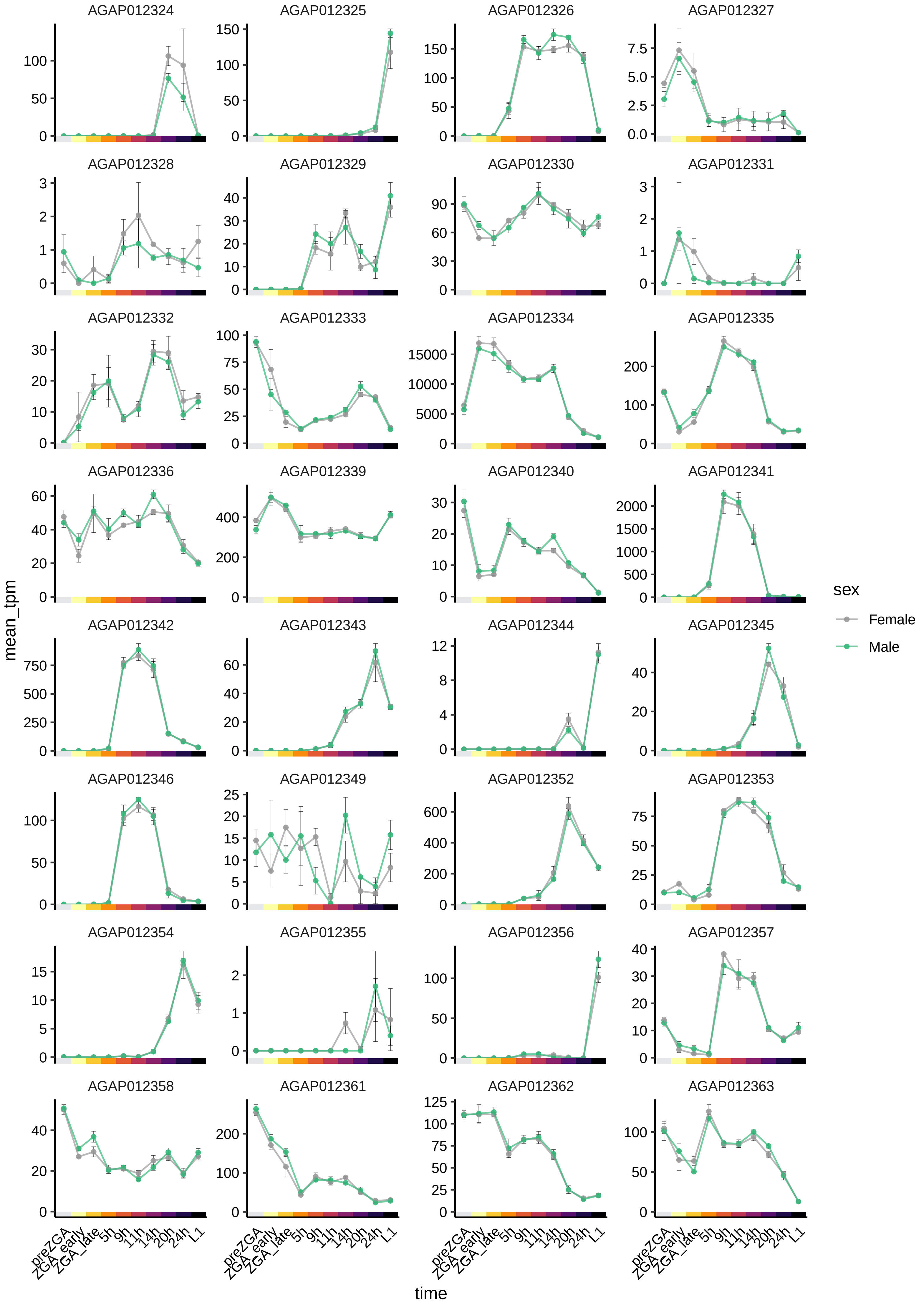
AGAP012324
AGAP012325
AGAP012326
AGAP012327
AGAP012328
AGAP012329
AGAP012330
AGAP012331
AGAP012332
AGAP012333
AGAP012334
AGAP012335
AGAP012336
AGAP012339
AGAP012340
AGAP012341
AGAP012342
AGAP012343
AGAP012344
AGAP012345
AGAP012346
AGAP012349
AGAP012352
AGAP012353
AGAP012354
AGAP012355
AGAP012356
AGAP012357
AGAP012358
AGAP012361
AGAP012362
AGAP012363
mean_tpm
time
preZGA
ZGA_early
ZGA_late
5h
9h
11h
14h
20h
24h
L1
sex
Female
Male

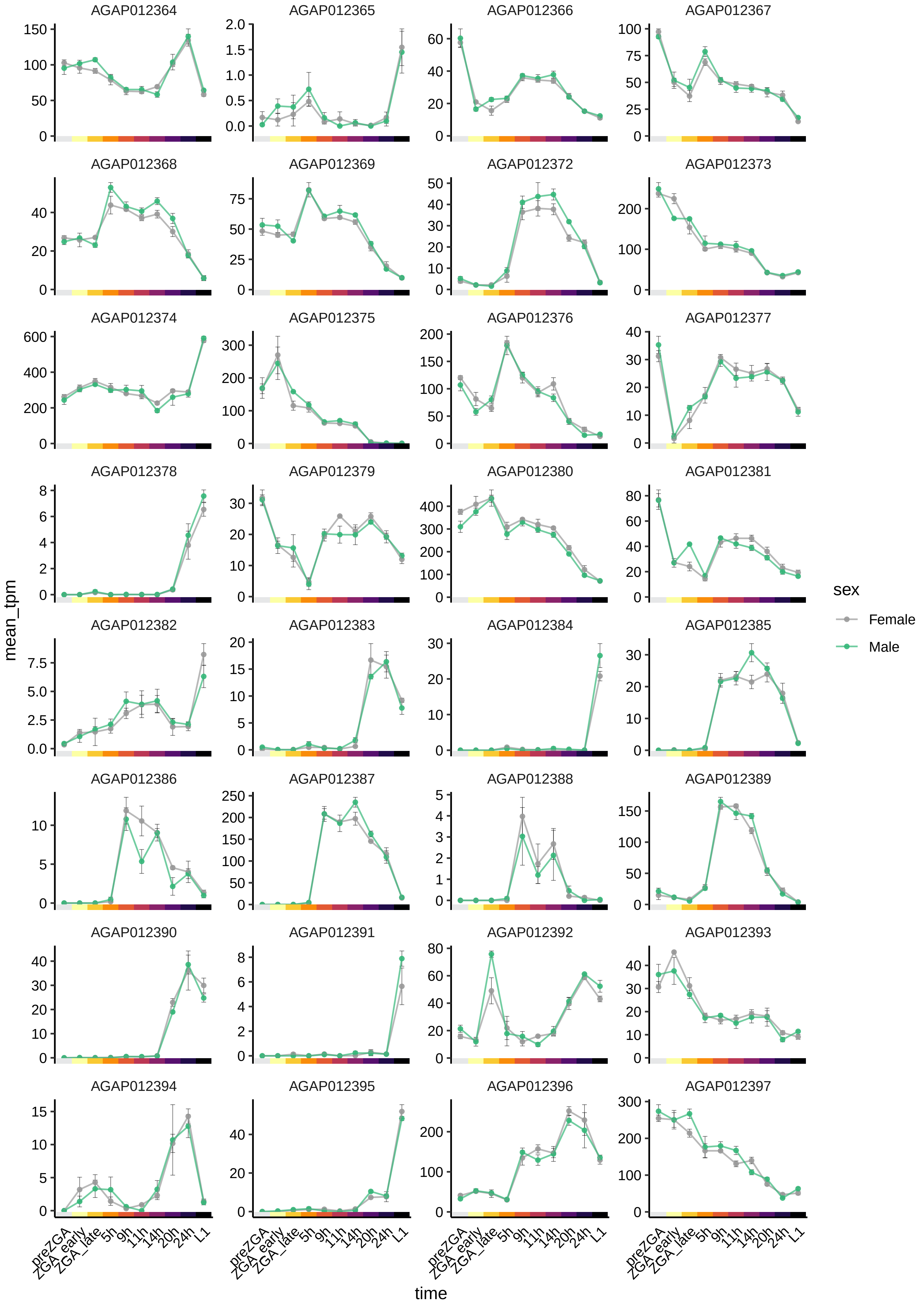
AGAP012364
AGAP012365
AGAP012366
AGAP012367
AGAP012368
AGAP012369
AGAP012372
AGAP012373
AGAP012374
AGAP012375
AGAP012376
AGAP012377
AGAP012378
AGAP012379
AGAP012380
AGAP012381
AGAP012382
AGAP012383
AGAP012384
AGAP012385
AGAP012386
AGAP012387
AGAP012388
AGAP012389
AGAP012390
AGAP012391
AGAP012392
AGAP012393
AGAP012394
AGAP012395
AGAP012396
AGAP012397
mean_tpm
time
preZGA
ZGA_early
ZGA_late
5h
9h
11h
14h
20h
24h
L1
sex
Female
Male

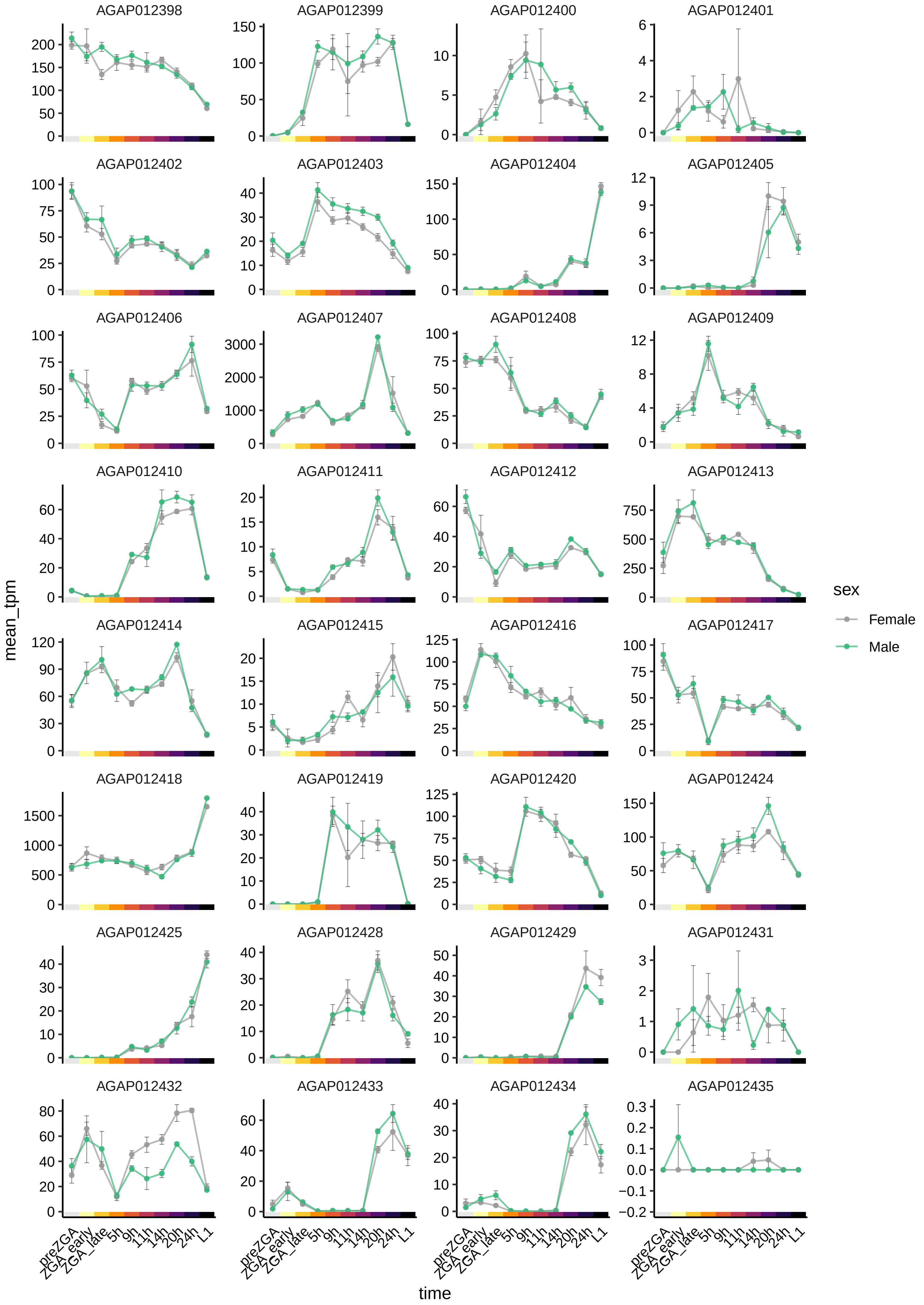

AGAP012398
AGAP012399
AGAP012400
AGAP012401
AGAP012402
AGAP012403
AGAP012404
AGAP012405
AGAP012406
AGAP012407
AGAP012408
AGAP012409
AGAP012410
AGAP012411
AGAP012412
AGAP012413
AGAP012414
AGAP012415
AGAP012416
AGAP012417
AGAP012418
AGAP012419
AGAP012420
AGAP012424
AGAP012425
AGAP012428
AGAP012429
AGAP012431
AGAP012432
AGAP012433
AGAP012434
AGAP012435
mean_tpm
time
preZGA
ZGA_early
ZGA_late
5h
9h
11h
14h
20h
24h
L1
sex
Female
Male

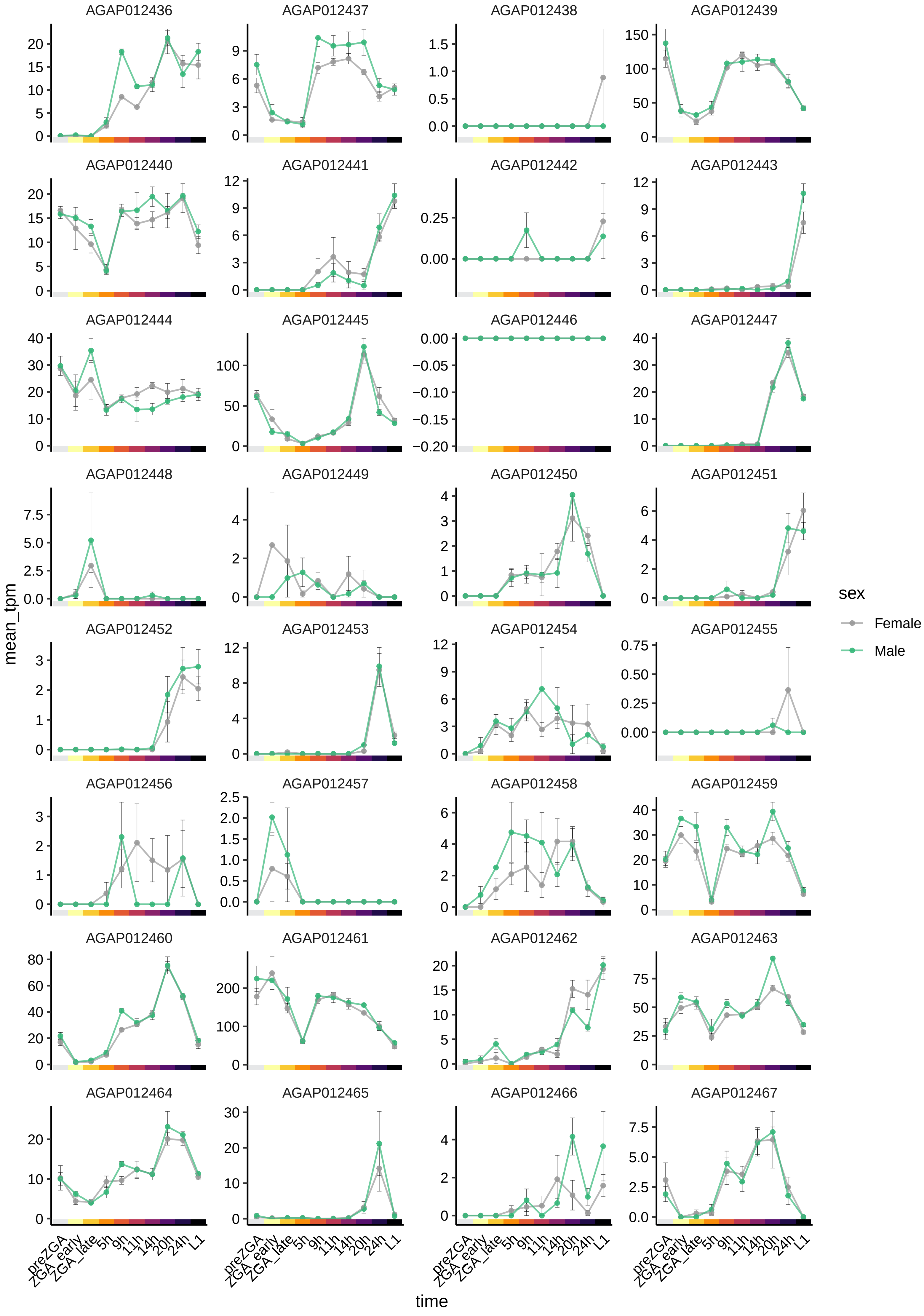

AGAP012436
AGAP012437
AGAP012438
AGAP012439
AGAP012440
AGAP012441
AGAP012442
AGAP012443
AGAP012444
AGAP012445
AGAP012446
AGAP012447
AGAP012448
AGAP012449
AGAP012450
AGAP012451
AGAP012452
AGAP012453
AGAP012454
AGAP012455
AGAP012456
AGAP012457
AGAP012458
AGAP012459
AGAP012460
AGAP012461
AGAP012462
AGAP012463
AGAP012464
AGAP012465
AGAP012466
AGAP012467
mean_tpm
time
preZGA
ZGA_early
ZGA_late
5h
9h
11h
14h
20h
24h
L1
sex
Female
Male

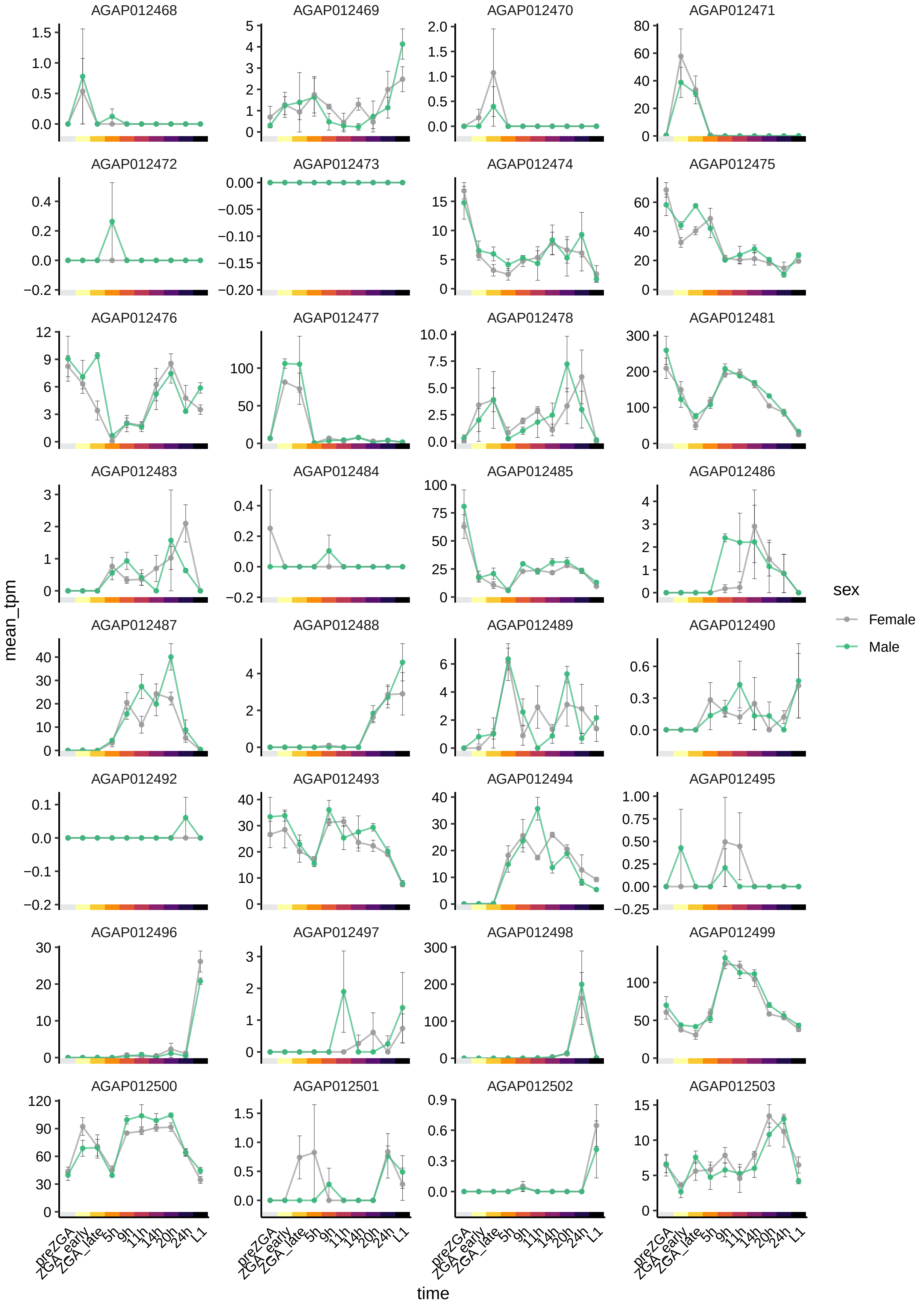
AGAP012468
AGAP012469
AGAP012470
AGAP012471
AGAP012472
AGAP012473
AGAP012474
AGAP012475
AGAP012476
AGAP012477
AGAP012478
AGAP012481
AGAP012483
AGAP012484
AGAP012485
AGAP012486
AGAP012487
AGAP012488
AGAP012489
AGAP012490
AGAP012492
AGAP012493
AGAP012494
AGAP012495
AGAP012496
AGAP012497
AGAP012498
AGAP012499
AGAP012500
AGAP012501
AGAP012502
AGAP012503
mean_tpm
time
preZGA
ZGA_early
ZGA_late
5h
9h
11h
14h
20h
24h
L1
sex
Female
Male

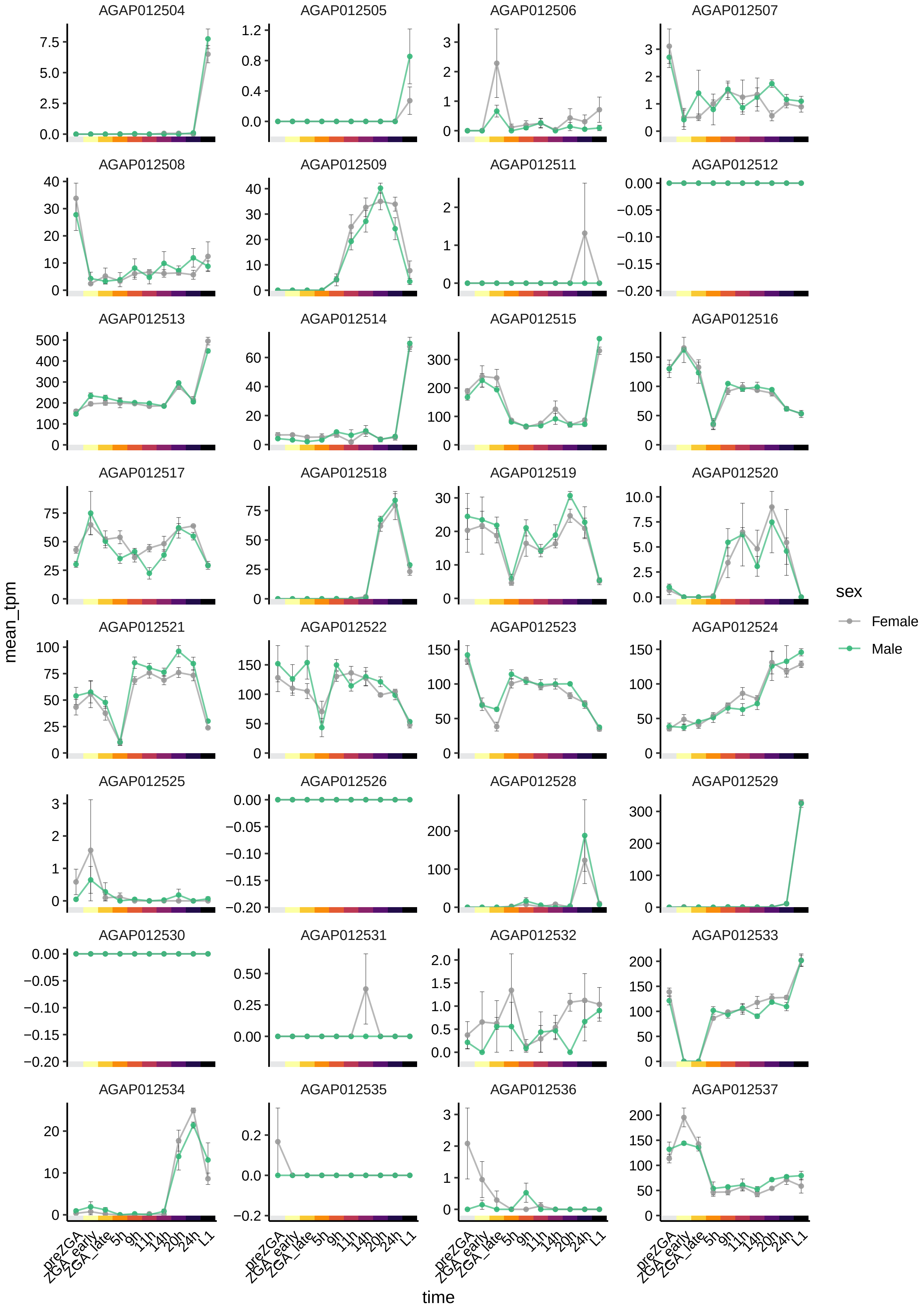
AGAP012504
AGAP012505
AGAP012506
AGAP012507
AGAP012508
AGAP012509
AGAP012511
AGAP012512
AGAP012513
AGAP012514
AGAP012515
AGAP012516
AGAP012517
AGAP012518
AGAP012519
AGAP012520
AGAP012521
AGAP012522
AGAP012523
AGAP012524
AGAP012525
AGAP012526
AGAP012528
AGAP012529
AGAP012530
AGAP012531
AGAP012532
AGAP012533
AGAP012534
AGAP012535
AGAP012536
AGAP012537
mean_tpm
time
preZGA
ZGA_early
ZGA_late
5h
9h
11h
14h
20h
24h
L1
sex
Female
Male

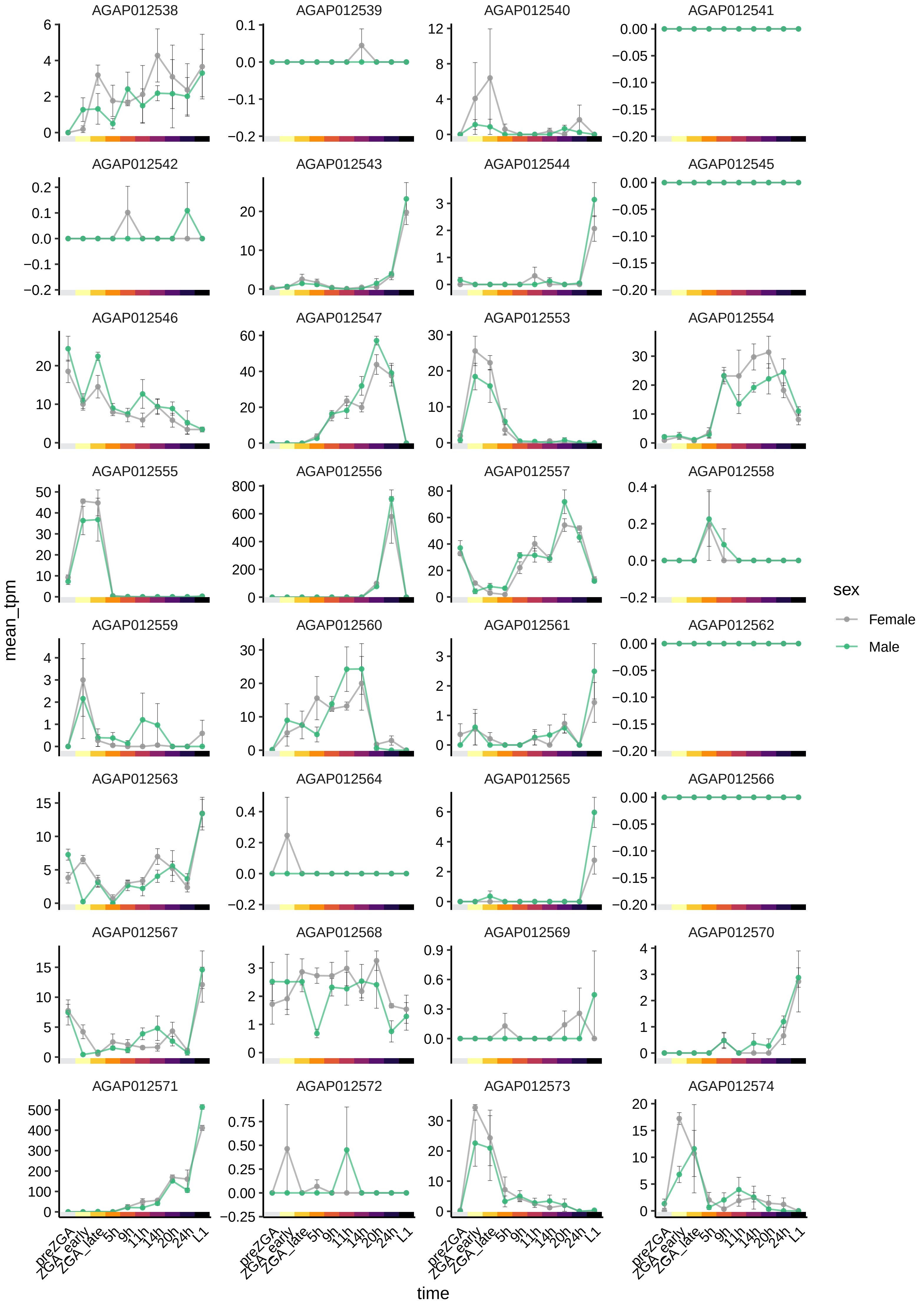

AGAP012538
AGAP012539
AGAP012540
AGAP012541
AGAP012542
AGAP012543
AGAP012544
AGAP012545
AGAP012546
AGAP012547
AGAP012553
AGAP012554
AGAP012555
AGAP012556
AGAP012557
AGAP012558
AGAP012559
AGAP012560
AGAP012561
AGAP012562
AGAP012563
AGAP012564
AGAP012565
AGAP012566
AGAP012567
AGAP012568
AGAP012569
AGAP012570
AGAP012571
AGAP012572
AGAP012573
AGAP012574
mean_tpm
time
preZGA
ZGA_early
ZGA_late
5h
9h
11h
14h
20h
24h
L1
sex
Female
Male

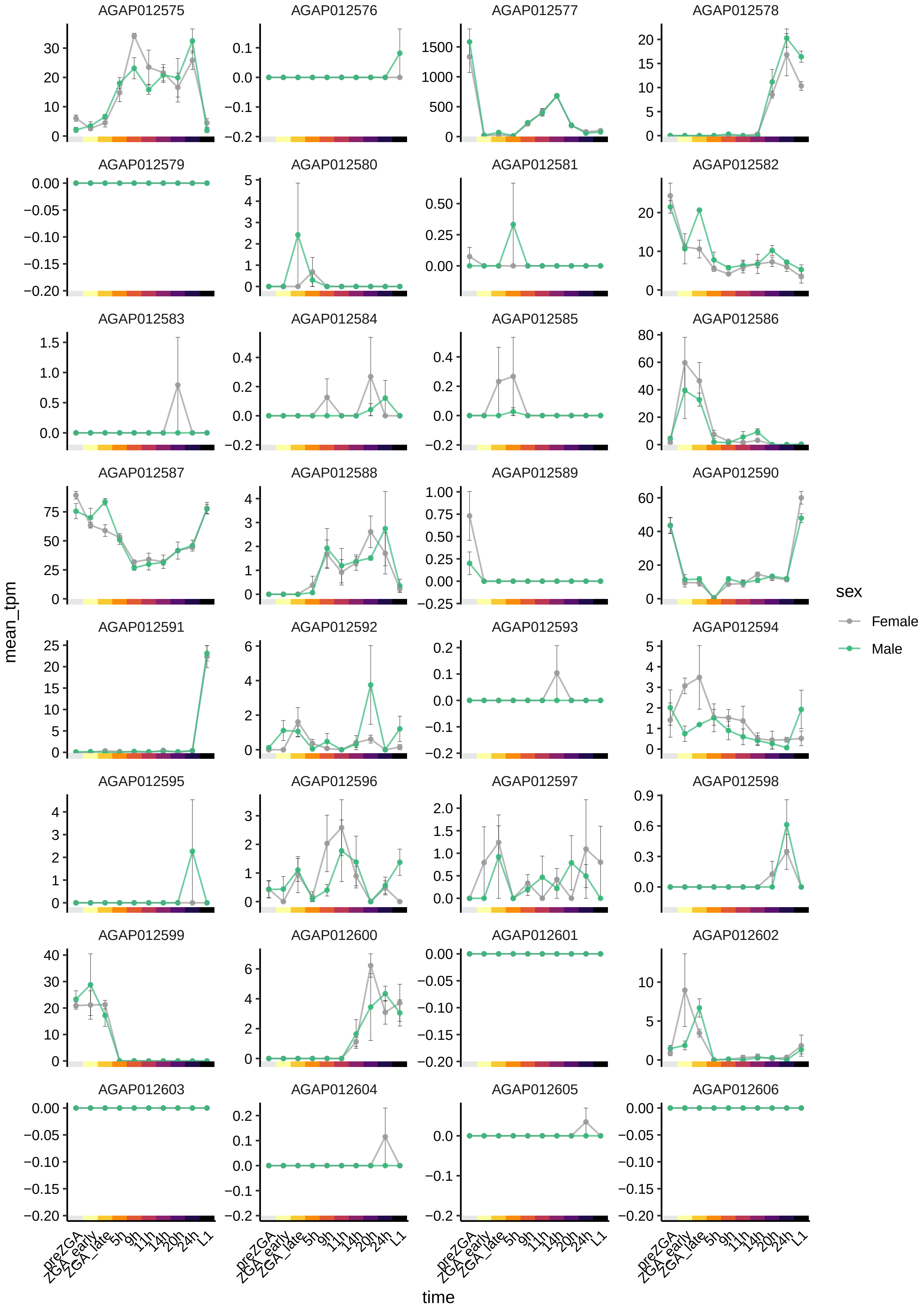
AGAP012575
AGAP012576
AGAP012577
AGAP012578
AGAP012579
AGAP012580
AGAP012581
AGAP012582
AGAP012583
AGAP012584
AGAP012585
AGAP012586
AGAP012587
AGAP012588
AGAP012589
AGAP012590
AGAP012591
AGAP012592
AGAP012593
AGAP012594
AGAP012595
AGAP012596
AGAP012597
AGAP012598
AGAP012599
AGAP012600
AGAP012601
AGAP012602
AGAP012603
AGAP012604
AGAP012605
AGAP012606
mean_tpm
time
preZGA
ZGA_early
ZGA_late
5h
9h
11h
14h
20h
24h
L1
sex
Female
Male

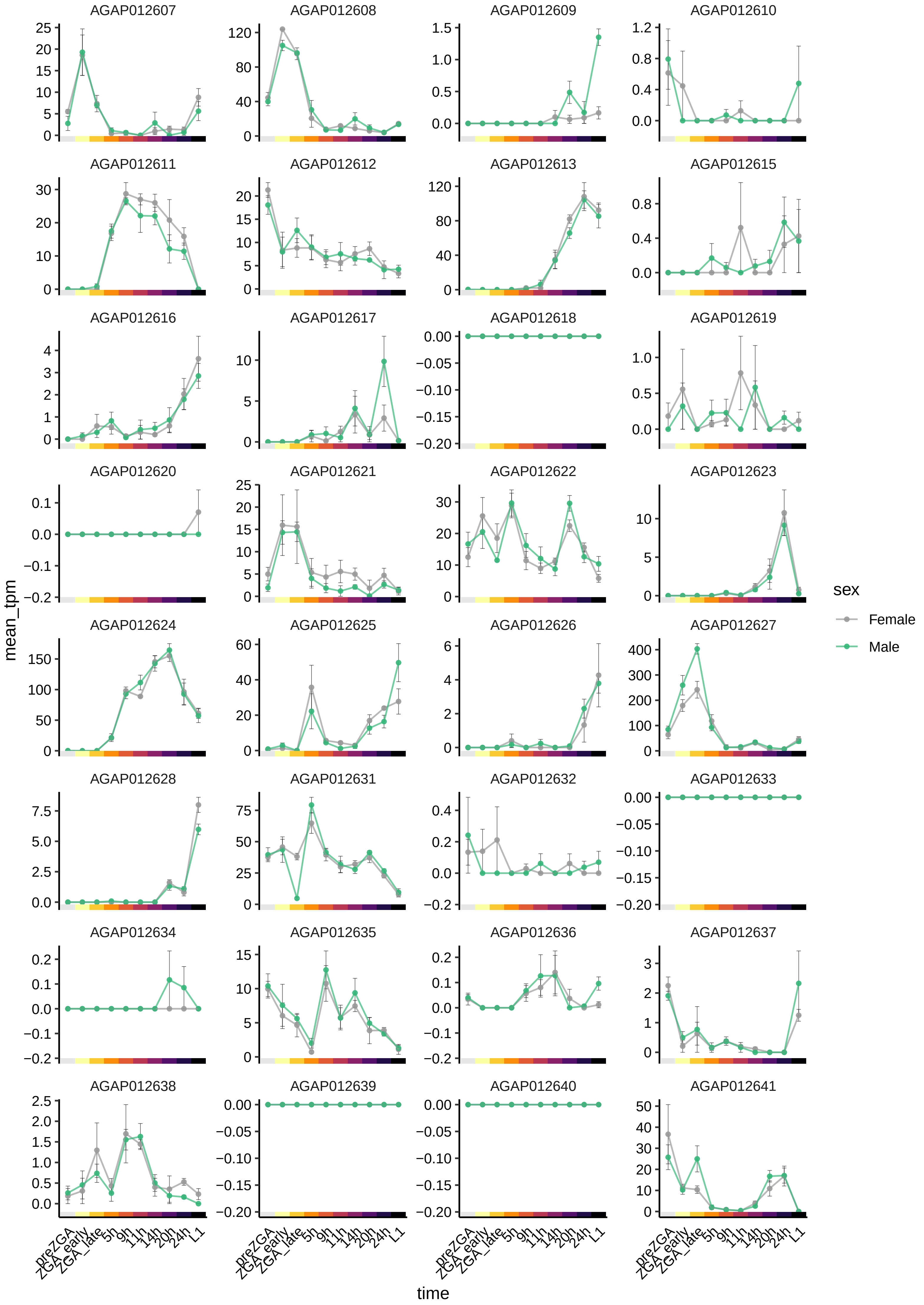

AGAP012607
AGAP012608
AGAP012609
AGAP012610
AGAP012611
AGAP012612
AGAP012613
AGAP012615
AGAP012616
AGAP012617
AGAP012618
AGAP012619
AGAP012620
AGAP012621
AGAP012622
AGAP012623
AGAP012624
AGAP012625
AGAP012626
AGAP012627
AGAP012628
AGAP012631
AGAP012632
AGAP012633
AGAP012634
AGAP012635
AGAP012636
AGAP012637
AGAP012638
AGAP012639
AGAP012640
AGAP012641
mean_tpm
time
preZGA
ZGA_early
ZGA_late
5h
9h
11h
14h
20h
24h
L1
sex
Female
Male

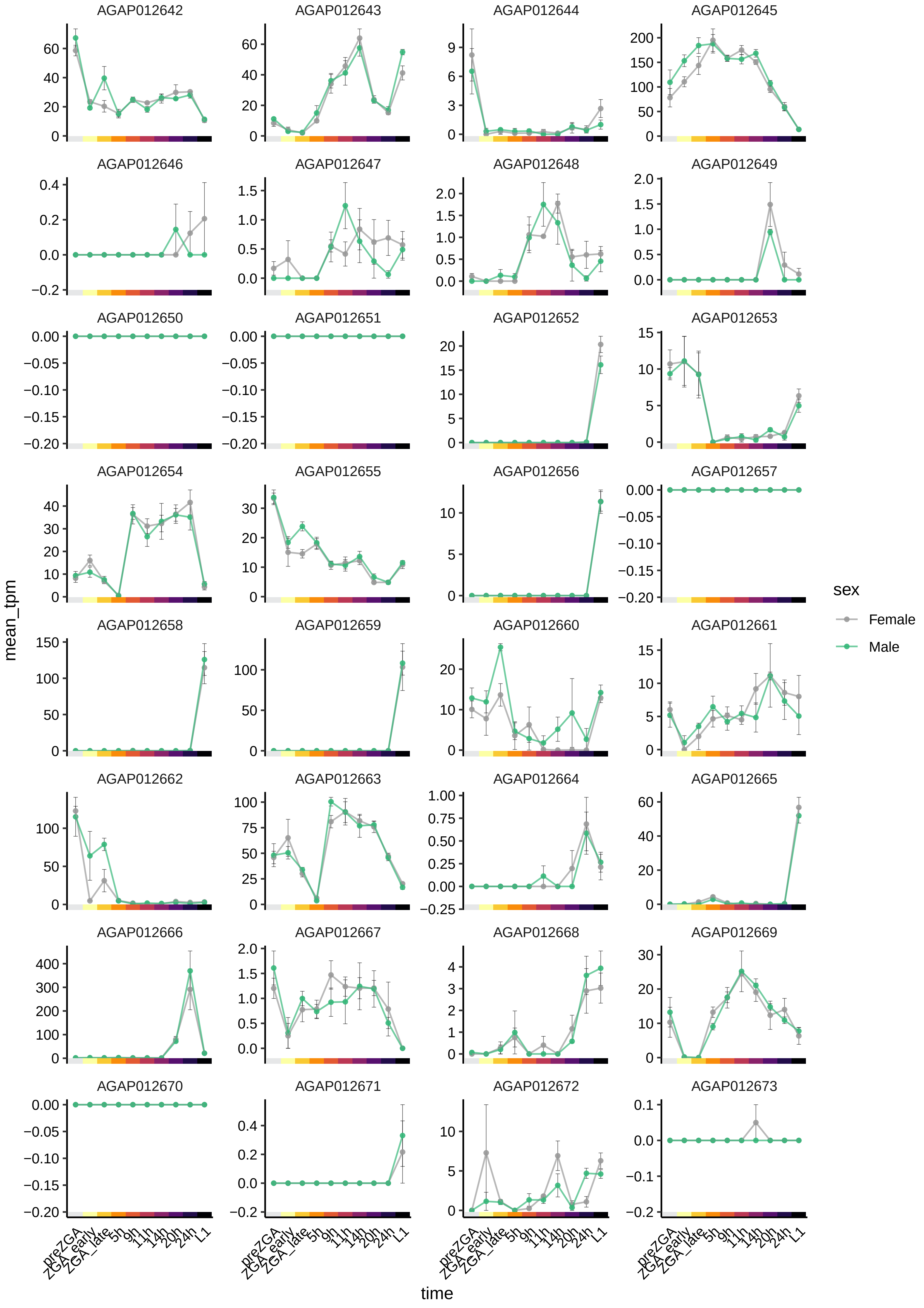

AGAP012642
AGAP012643
AGAP012644
AGAP012645
AGAP012646
AGAP012647
AGAP012648
AGAP012649
AGAP012650
AGAP012651
AGAP012652
AGAP012653
AGAP012654
AGAP012655
AGAP012656
AGAP012657
AGAP012658
AGAP012659
AGAP012660
AGAP012661
AGAP012662
AGAP012663
AGAP012664
AGAP012665
AGAP012666
AGAP012667
AGAP012668
AGAP012669
AGAP012670
AGAP012671
AGAP012672
AGAP012673
mean_tpm
time
preZGA
ZGA_early
ZGA_late
5h
9h
11h
14h
20h
24h
L1
sex
Female
Male

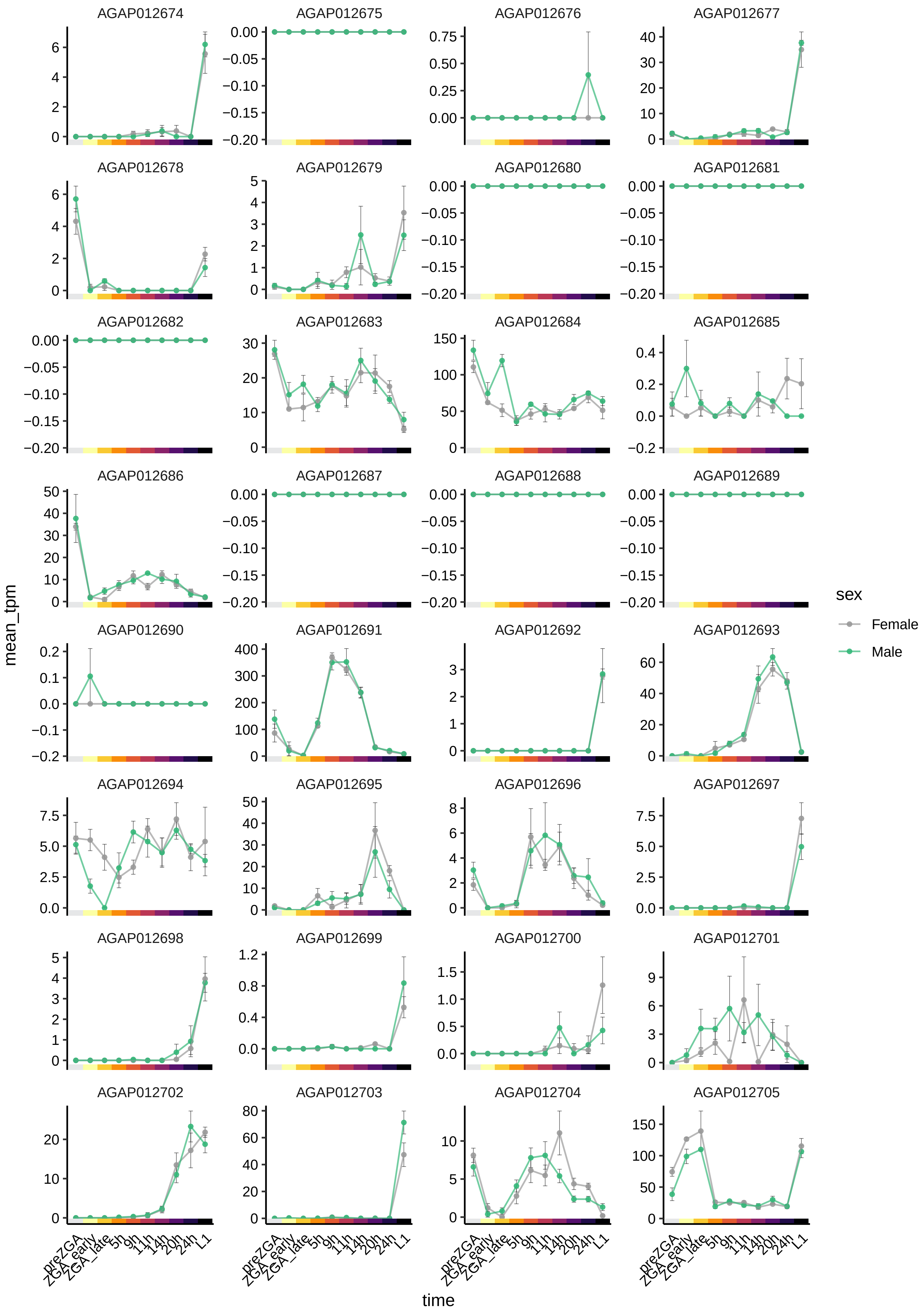

AGAP012674
AGAP012675
AGAP012676
AGAP012677
AGAP012678
AGAP012679
AGAP012680
AGAP012681
AGAP012682
AGAP012683
AGAP012684
AGAP012685
AGAP012686
AGAP012687
AGAP012688
AGAP012689
AGAP012690
AGAP012691
AGAP012692
AGAP012693
AGAP012694
AGAP012695
AGAP012696
AGAP012697
AGAP012698
AGAP012699
AGAP012700
AGAP012701
AGAP012702
AGAP012703
AGAP012704
AGAP012705
mean_tpm
time
preZGA
ZGA_early
ZGA_late
5h
9h
11h
14h
20h
24h
L1
sex
Female
Male

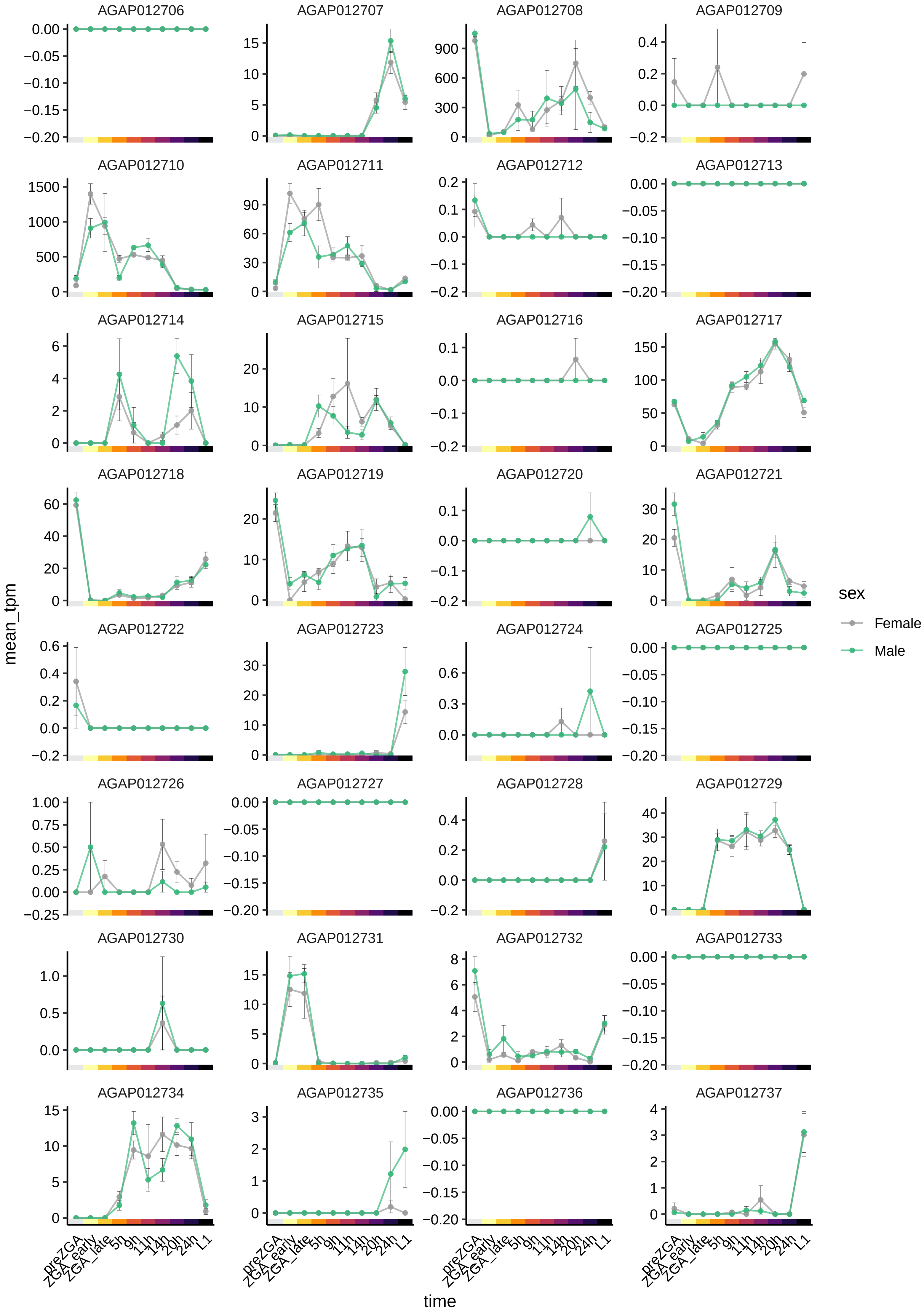
AGAP012706
AGAP012707
AGAP012708
AGAP012709
AGAP012710
AGAP012711
AGAP012712
AGAP012713
AGAP012714
AGAP012715
AGAP012716
AGAP012717
AGAP012718
AGAP012719
AGAP012720
AGAP012721
AGAP012722
AGAP012723
AGAP012724
AGAP012725
AGAP012726
AGAP012727
AGAP012728
AGAP012729
AGAP012730
AGAP012731
AGAP012732
AGAP012733
AGAP012734
AGAP012735
AGAP012736
AGAP012737
mean_tpm
time
preZGA
ZGA_early
ZGA_late
5h
9h
11h
14h
20h
24h
L1
sex
Female
Male

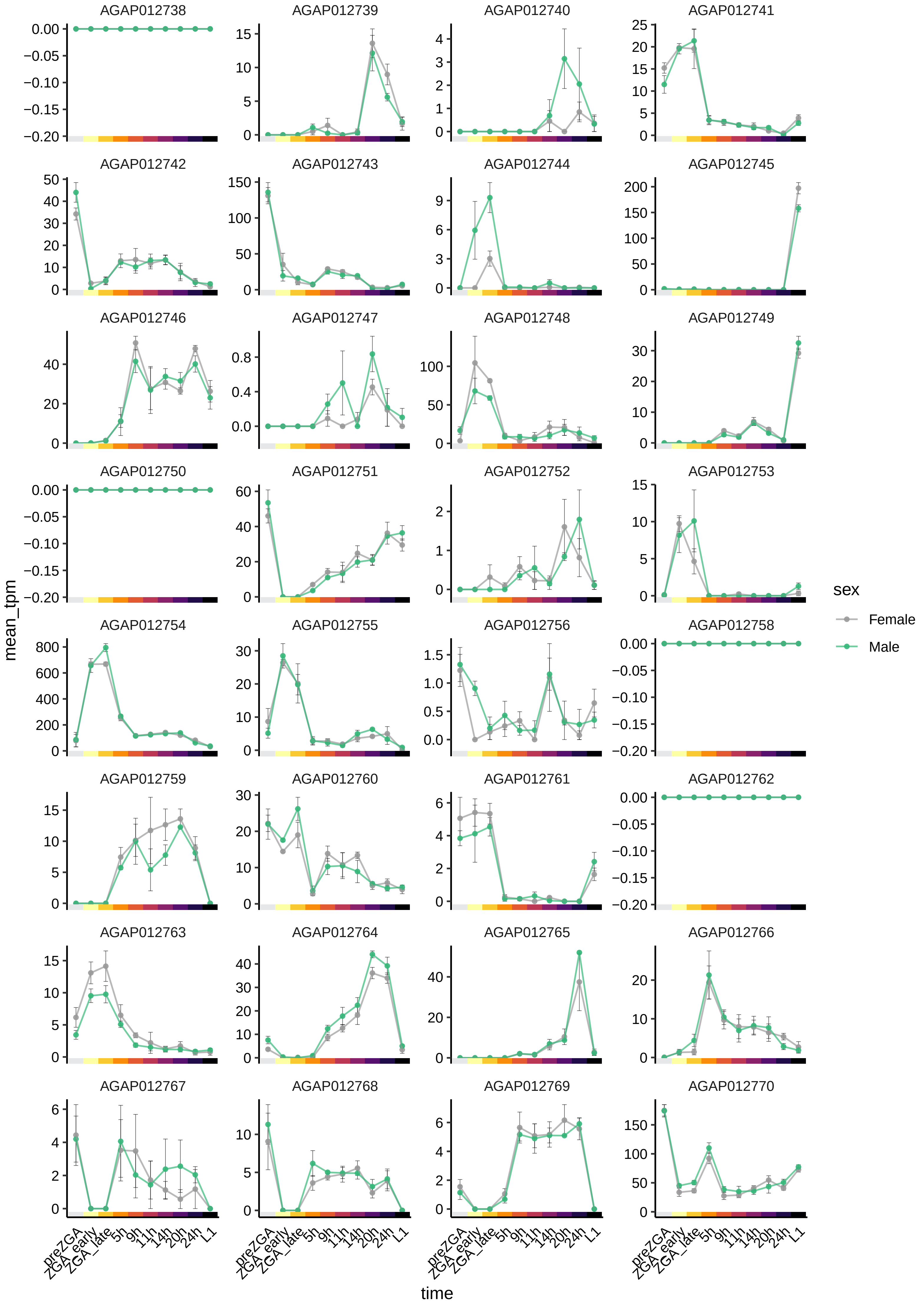

AGAP012738
AGAP012739
AGAP012740
AGAP012741
AGAP012742
AGAP012743
AGAP012744
AGAP012745
AGAP012746
AGAP012747
AGAP012748
AGAP012749
AGAP012750
AGAP012751
AGAP012752
AGAP012753
AGAP012754
AGAP012755
AGAP012756
AGAP012758
AGAP012759
AGAP012760
AGAP012761
AGAP012762
AGAP012763
AGAP012764
AGAP012765
AGAP012766
AGAP012767
AGAP012768
AGAP012769
AGAP012770
mean_tpm
time
preZGA
ZGA_early
ZGA_late
5h
9h
11h
14h
20h
24h
L1
sex
Female
Male

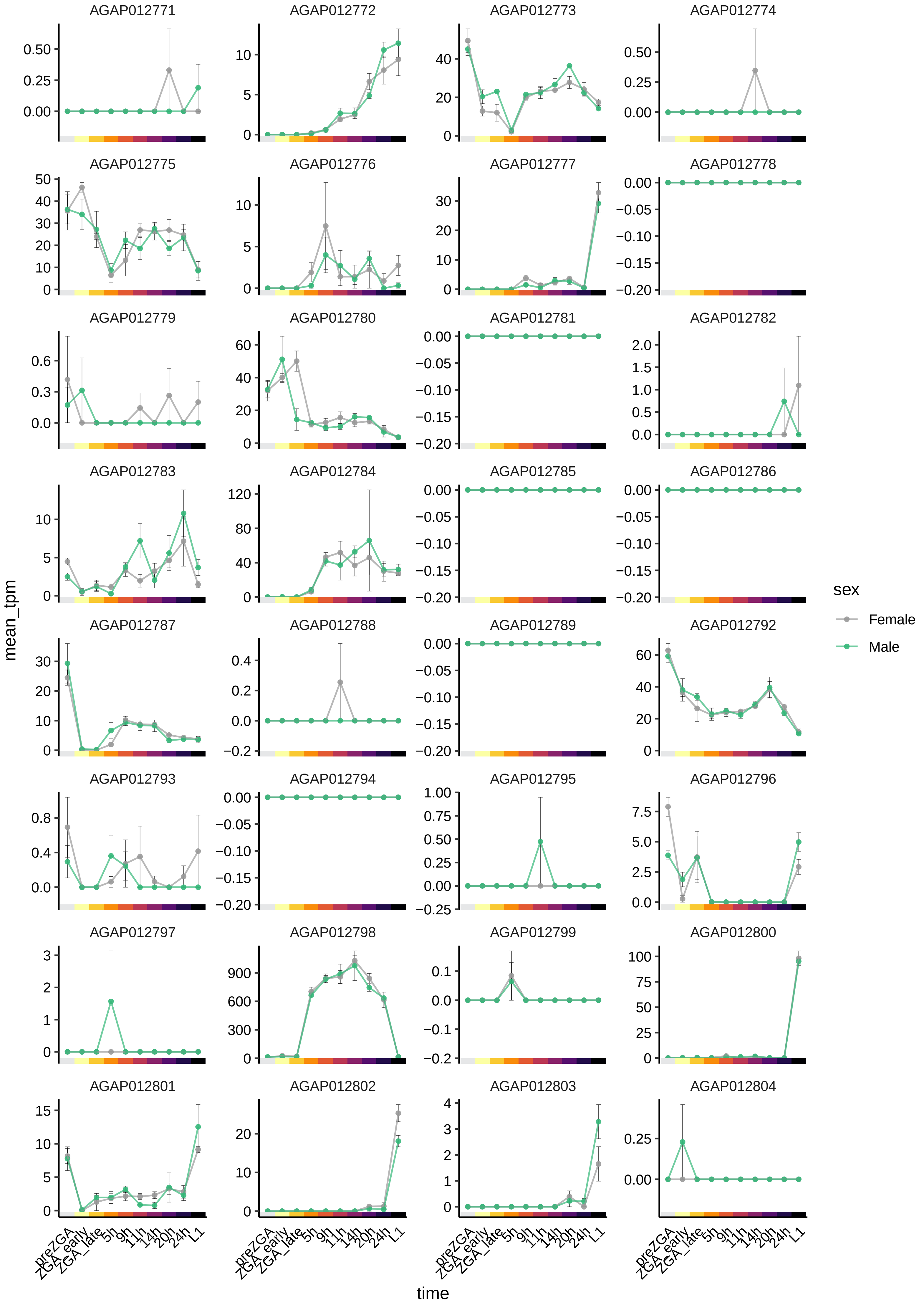

AGAP012771
AGAP012772
AGAP012773
AGAP012774
AGAP012775
AGAP012776
AGAP012777
AGAP012778
AGAP012779
AGAP012780
AGAP012781
AGAP012782
AGAP012783
AGAP012784
AGAP012785
AGAP012786
AGAP012787
AGAP012788
AGAP012789
AGAP012792
AGAP012793
AGAP012794
AGAP012795
AGAP012796
AGAP012797
AGAP012798
AGAP012799
AGAP012800
AGAP012801
AGAP012802
AGAP012803
AGAP012804
mean_tpm
time
preZGA
ZGA_early
ZGA_late
5h
9h
11h
14h
20h
24h
L1
sex
Female
Male

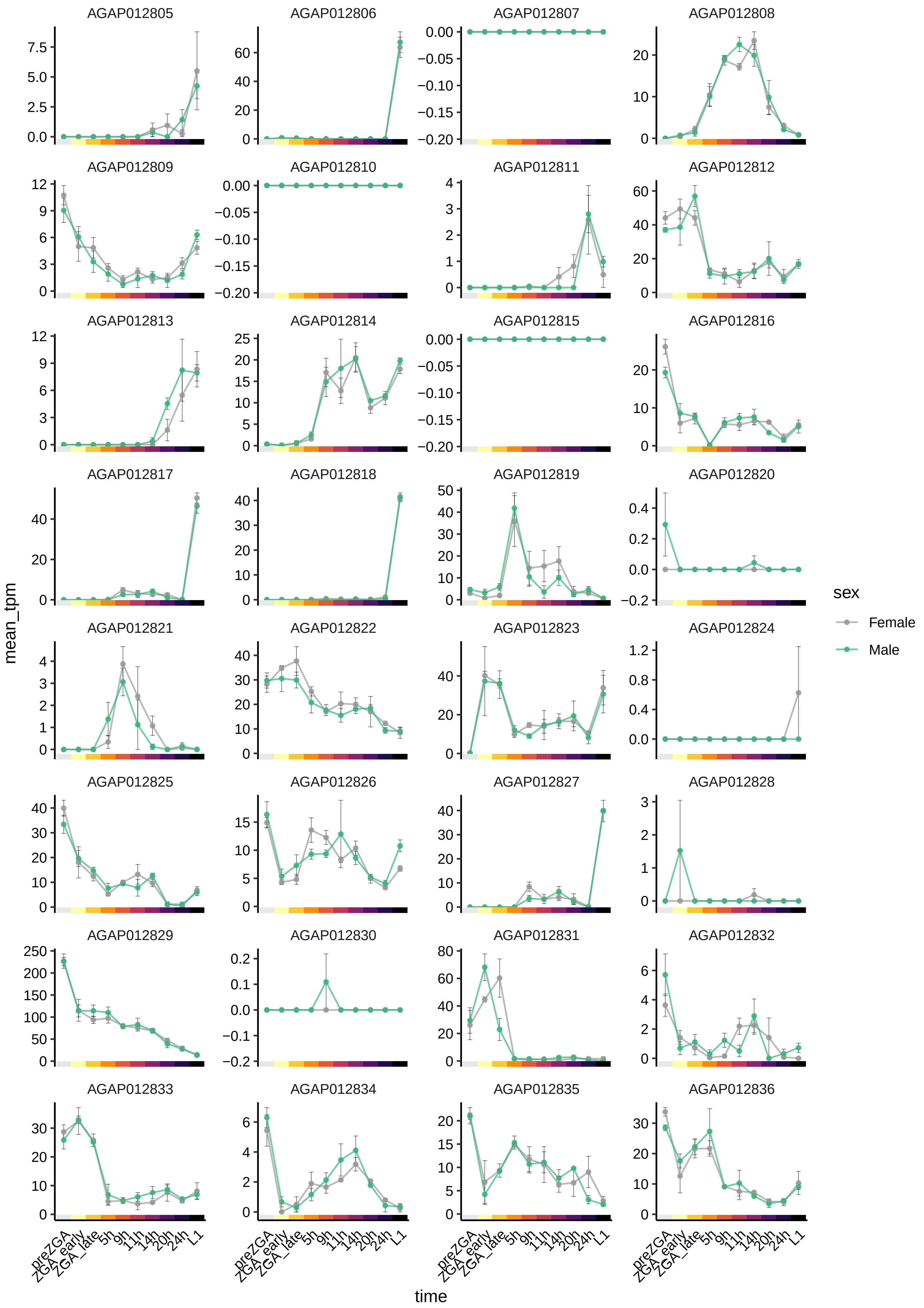
AGAP012805
AGAP012806
AGAP012807
AGAP012808
AGAP012809
AGAP012810
AGAP012811
AGAP012812
AGAP012813
AGAP012814
AGAP012815
AGAP012816
AGAP012817
AGAP012818
AGAP012819
AGAP012820
AGAP012821
AGAP012822
AGAP012823
AGAP012824
AGAP012825
AGAP012826
AGAP012827
AGAP012828
AGAP012829
AGAP012830
AGAP012831
AGAP012832
AGAP012833
AGAP012834
AGAP012835
AGAP012836
mean_tpm
time
preZGA
ZGA_early
ZGA_late
5h
9h
11h
14h
20h
24h
L1
sex
Female
Male

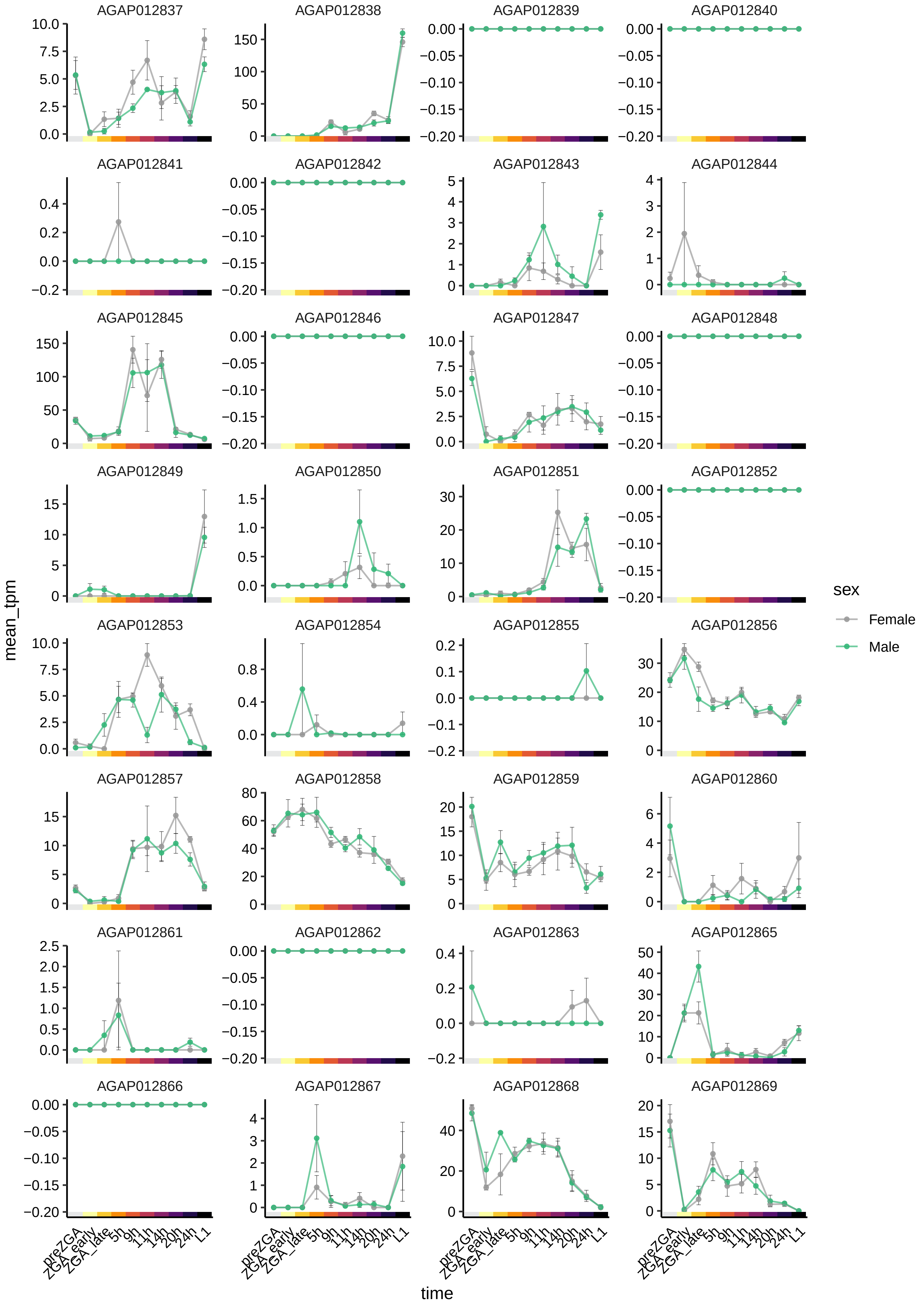

AGAP012837
AGAP012838
AGAP012839
AGAP012840
AGAP012841
AGAP012842
AGAP012843
AGAP012844
AGAP012845
AGAP012846
AGAP012847
AGAP012848
AGAP012849
AGAP012850
AGAP012851
AGAP012852
AGAP012853
AGAP012854
AGAP012855
AGAP012856
AGAP012857
AGAP012858
AGAP012859
AGAP012860
AGAP012861
AGAP012862
AGAP012863
AGAP012865
AGAP012866
AGAP012867
AGAP012868
AGAP012869
mean_tpm
time
preZGA
ZGA_early
ZGA_late
5h
9h
11h
14h
20h
24h
L1
sex
Female
Male

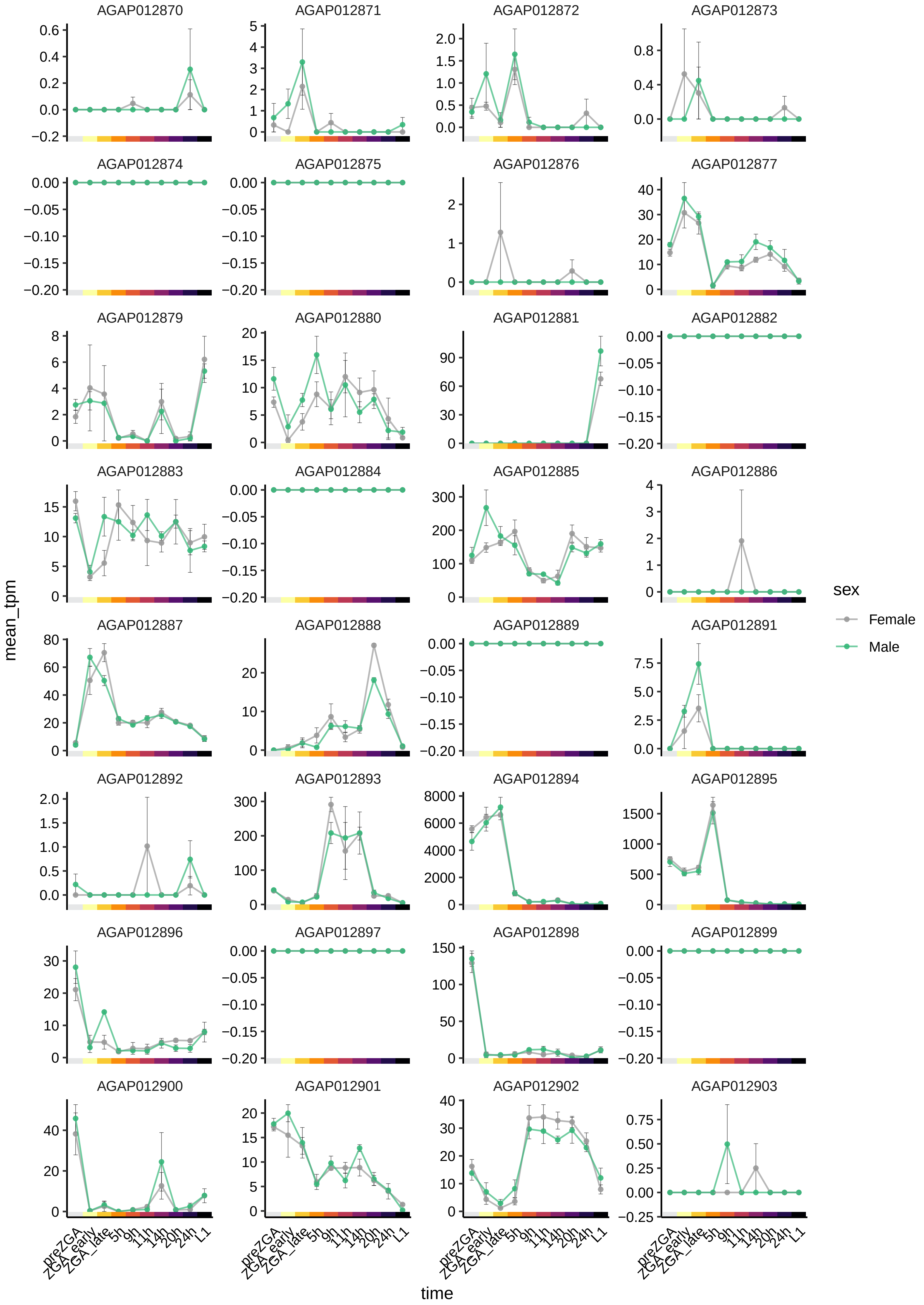

AGAP012870
AGAP012871
AGAP012872
AGAP012873
AGAP012874
AGAP012875
AGAP012876
AGAP012877
AGAP012879
AGAP012880
AGAP012881
AGAP012882
AGAP012883
AGAP012884
AGAP012885
AGAP012886
AGAP012887
AGAP012888
AGAP012889
AGAP012891
AGAP012892
AGAP012893
AGAP012894
AGAP012895
AGAP012896
AGAP012897
AGAP012898
AGAP012899
AGAP012900
AGAP012901
AGAP012902
AGAP012903
mean_tpm
time
preZGA
ZGA_early
ZGA_late
5h
9h
11h
14h
20h
24h
L1
sex
Female
Male

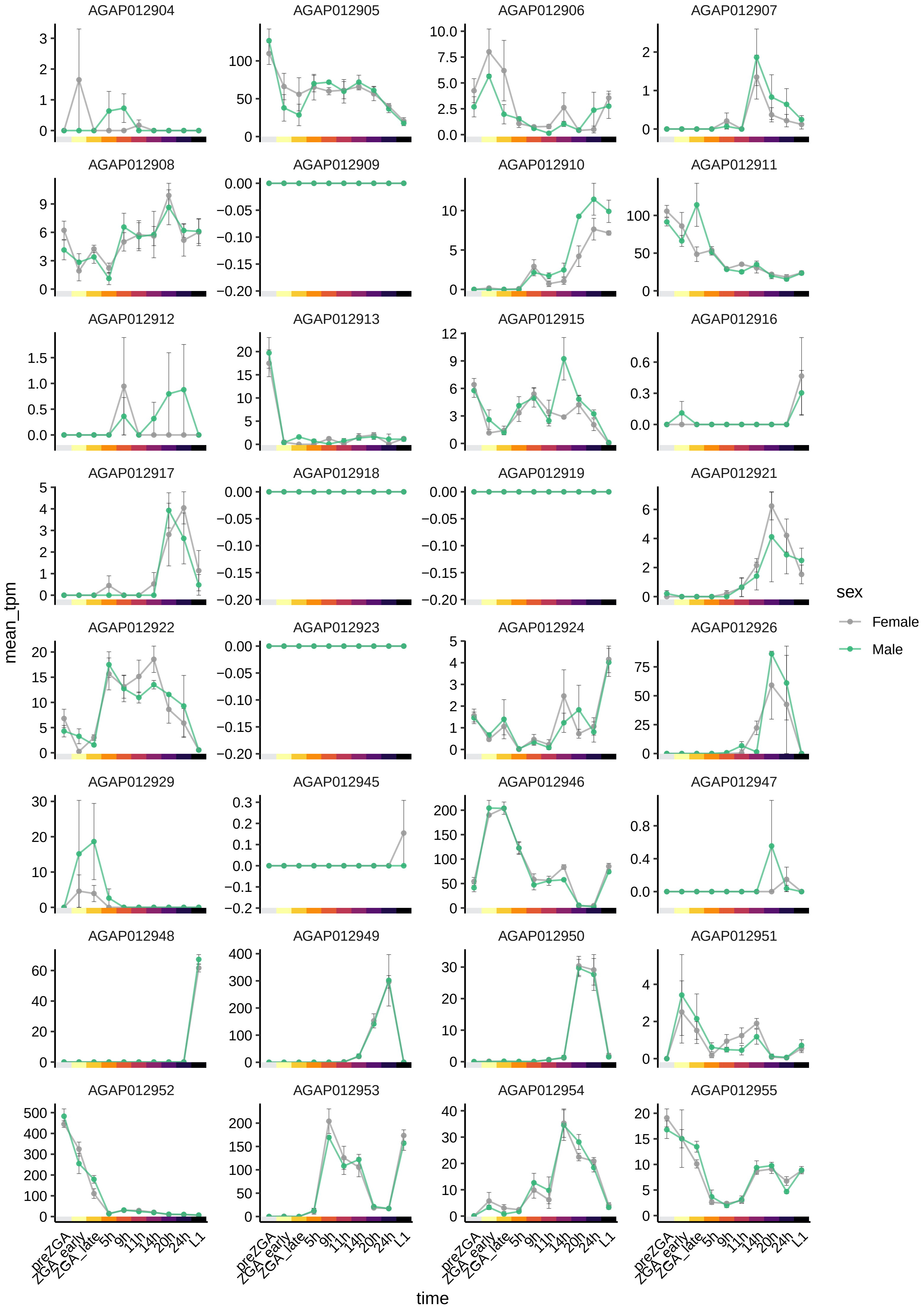
AGAP012904
AGAP012905
AGAP012906
AGAP012907
AGAP012908
AGAP012909
AGAP012910
AGAP012911
AGAP012912
AGAP012913
AGAP012915
AGAP012916
AGAP012917
AGAP012918
AGAP012919
AGAP012921
AGAP012922
AGAP012923
AGAP012924
AGAP012926
AGAP012929
AGAP012945
AGAP012946
AGAP012947
AGAP012948
AGAP012949
AGAP012950
AGAP012951
AGAP012952
AGAP012953
AGAP012954
AGAP012955
mean_tpm
time
preZGA
ZGA_early
ZGA_late
5h
9h
11h
14h
20h
24h
L1
sex
Female
Male

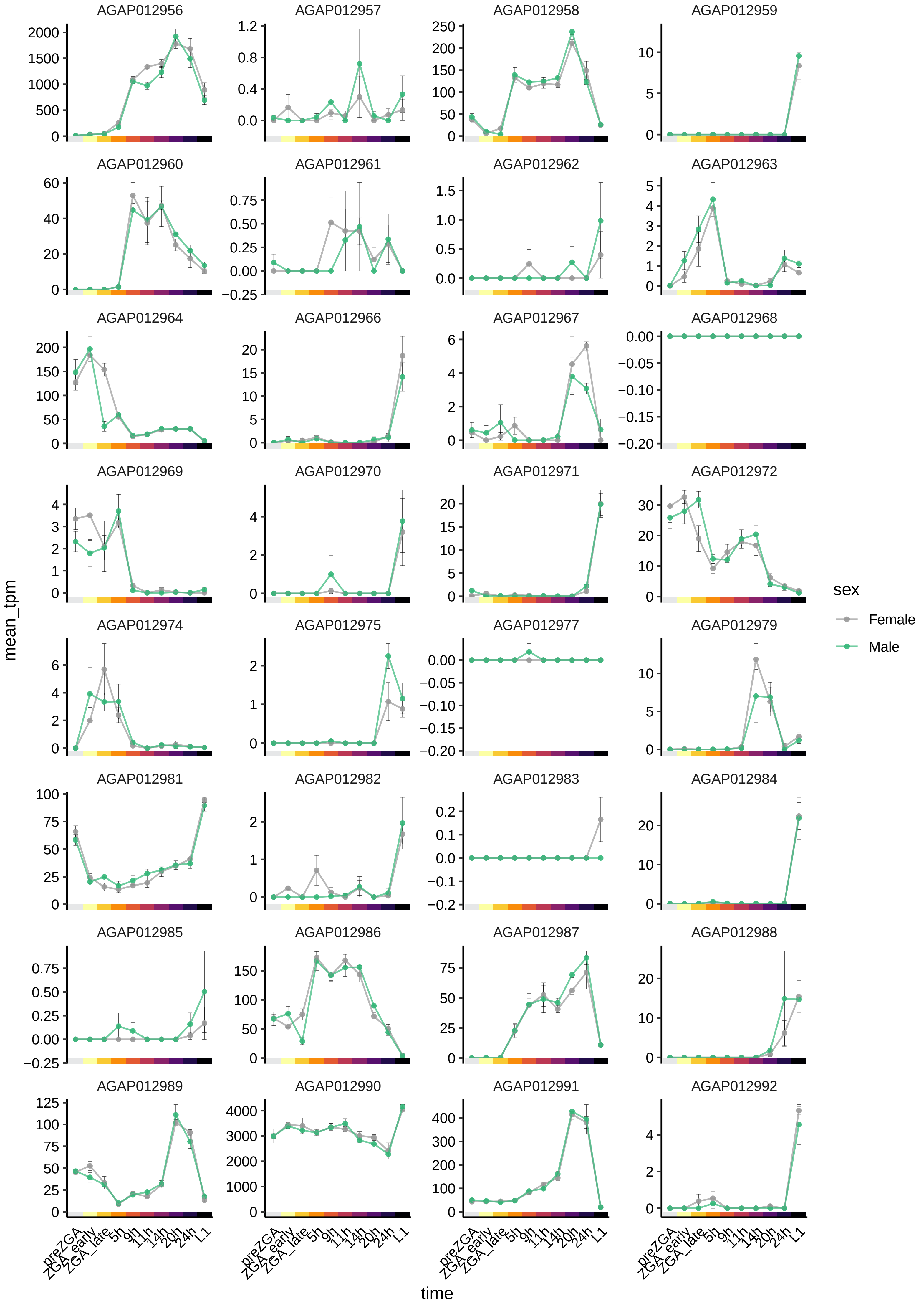
AGAP012956
AGAP012957
AGAP012958
AGAP012959
AGAP012960
AGAP012961
AGAP012962
AGAP012963
AGAP012964
AGAP012966
AGAP012967
AGAP012968
AGAP012969
AGAP012970
AGAP012971
AGAP012972
AGAP012974
AGAP012975
AGAP012977
AGAP012979
AGAP012981
AGAP012982
AGAP012983
AGAP012984
AGAP012985
AGAP012986
AGAP012987
AGAP012988
AGAP012989
AGAP012990
AGAP012991
AGAP012992
mean_tpm
time
preZGA
ZGA_early
ZGA_late
5h
9h
11h
14h
20h
24h
L1
sex
Female
Male

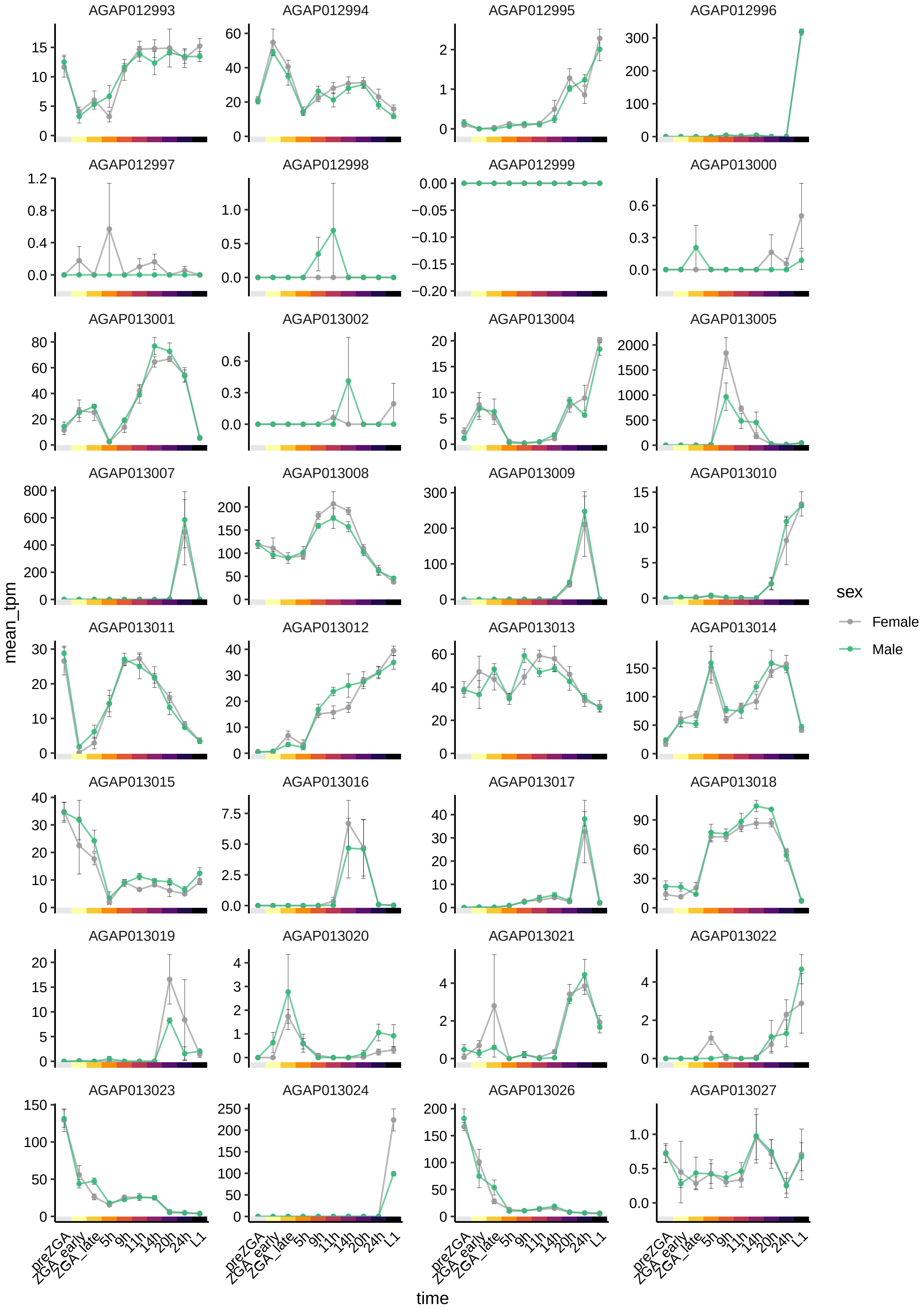
AGAP012993
AGAP012994
AGAP012995
AGAP012996
AGAP012997
AGAP012998
AGAP012999
AGAP013000
AGAP013001
AGAP013002
AGAP013004
AGAP013005
AGAP013007
AGAP013008
AGAP013009
AGAP013010
AGAP013011
AGAP013012
AGAP013013
AGAP013014
AGAP013015
AGAP013016
AGAP013017
AGAP013018
AGAP013019
AGAP013020
AGAP013021
AGAP013022
AGAP013023
AGAP013024
AGAP013026
AGAP013027
mean_tpm
time
preZGA
ZGA_early
ZGA_late
5h
9h
11h
14h
20h
24h
L1
sex
Female
Male

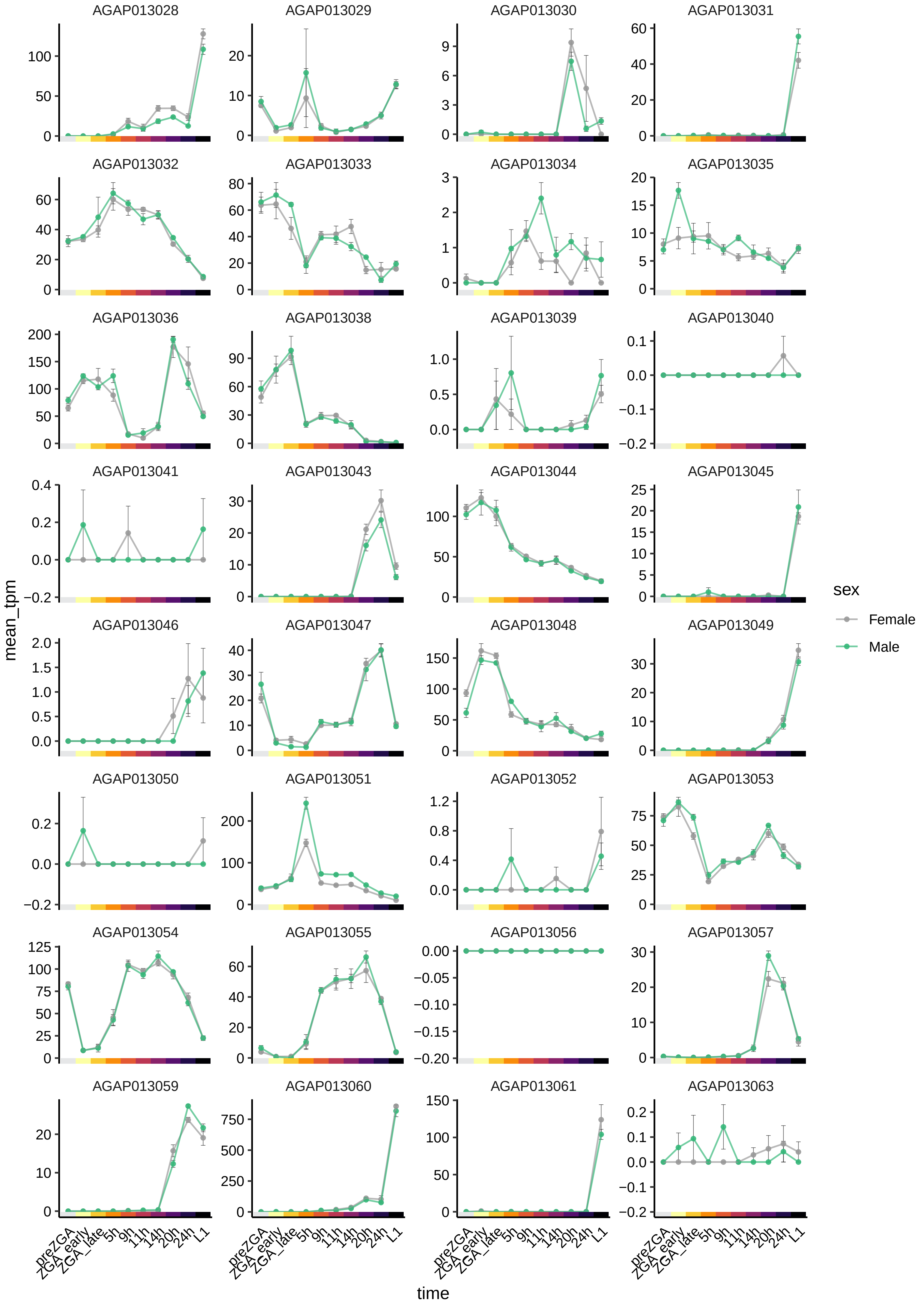
AGAP013028
AGAP013029
AGAP013030
AGAP013031
AGAP013032
AGAP013033
AGAP013034
AGAP013035
AGAP013036
AGAP013038
AGAP013039
AGAP013040
AGAP013041
AGAP013043
AGAP013044
AGAP013045
AGAP013046
AGAP013047
AGAP013048
AGAP013049
AGAP013050
AGAP013051
AGAP013052
AGAP013053
AGAP013054
AGAP013055
AGAP013056
AGAP013057
AGAP013059
AGAP013060
AGAP013061
AGAP013063
mean_tpm
time
preZGA
ZGA_early
ZGA_late
5h
9h
11h
14h
20h
24h
L1
sex
Female
Male

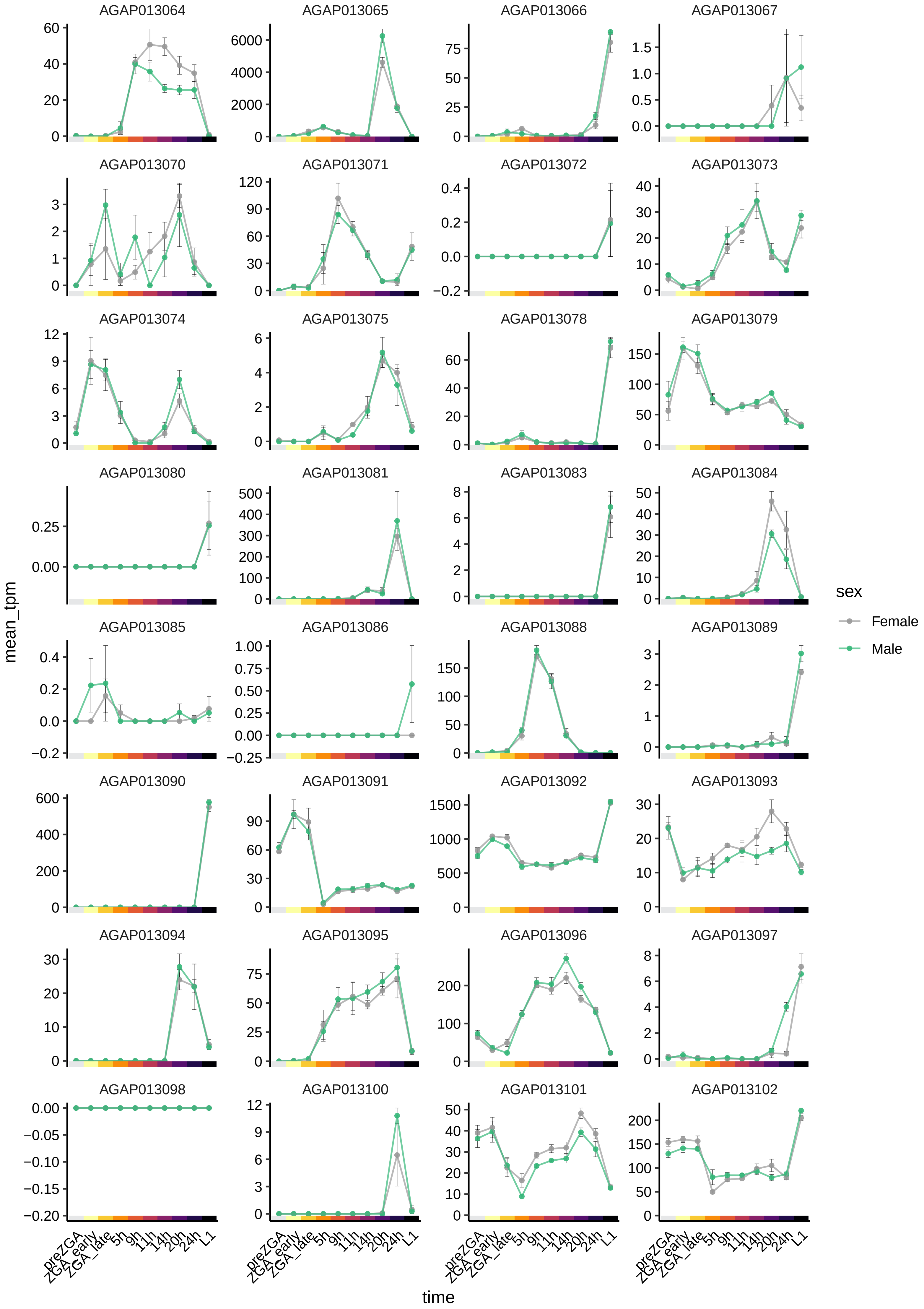

AGAP013064
AGAP013065
AGAP013066
AGAP013067
AGAP013070
AGAP013071
AGAP013072
AGAP013073
AGAP013074
AGAP013075
AGAP013078
AGAP013079
AGAP013080
AGAP013081
AGAP013083
AGAP013084
AGAP013085
AGAP013086
AGAP013088
AGAP013089
AGAP013090
AGAP013091
AGAP013092
AGAP013093
AGAP013094
AGAP013095
AGAP013096
AGAP013097
AGAP013098
AGAP013100
AGAP013101
AGAP013102
mean_tpm
time
preZGA
ZGA_early
ZGA_late
5h
9h
11h
14h
20h
24h
L1
sex
Female
Male

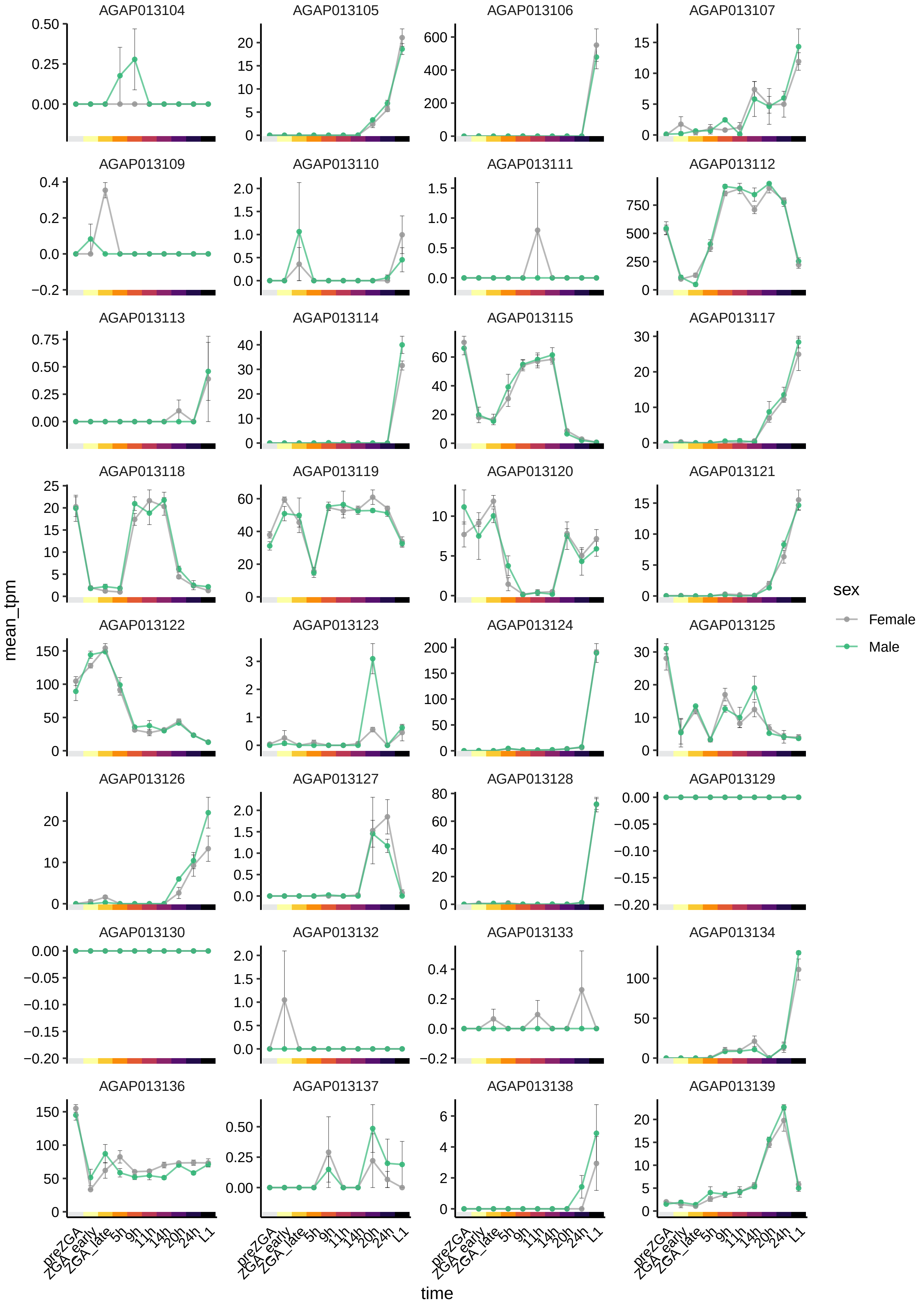

AGAP013104
AGAP013105
AGAP013106
AGAP013107
AGAP013109
AGAP013110
AGAP013111
AGAP013112
AGAP013113
AGAP013114
AGAP013115
AGAP013117
AGAP013118
AGAP013119
AGAP013120
AGAP013121
AGAP013122
AGAP013123
AGAP013124
AGAP013125
AGAP013126
AGAP013127
AGAP013128
AGAP013129
AGAP013130
AGAP013132
AGAP013133
AGAP013134
AGAP013136
AGAP013137
AGAP013138
AGAP013139
mean_tpm
time
preZGA
ZGA_early
ZGA_late
5h
9h
11h
14h
20h
24h
L1
sex
Female
Male

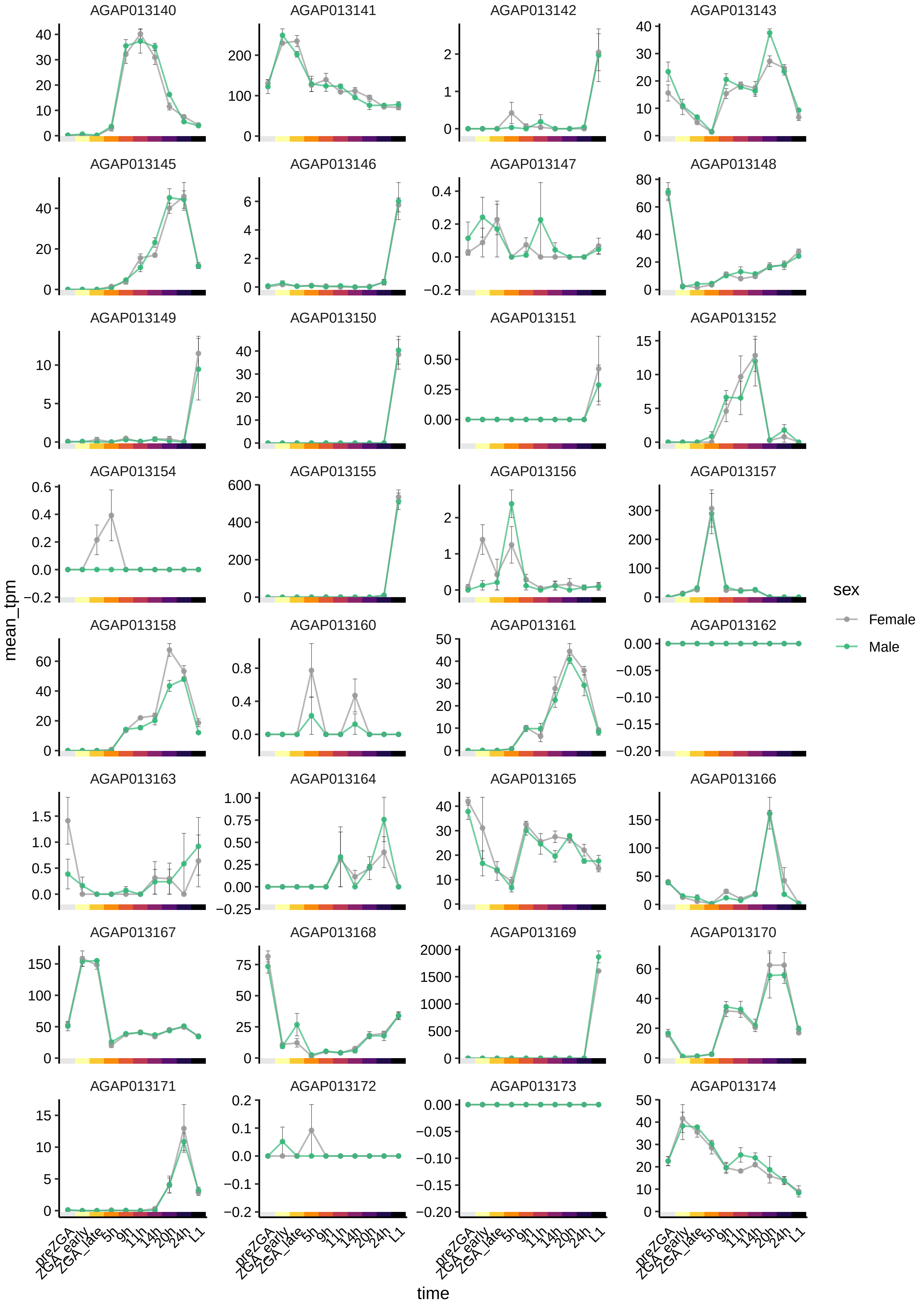
AGAP013140
AGAP013141
AGAP013142
AGAP013143
AGAP013145
AGAP013146
AGAP013147
AGAP013148
AGAP013149
AGAP013150
AGAP013151
AGAP013152
AGAP013154
AGAP013155
AGAP013156
AGAP013157
AGAP013158
AGAP013160
AGAP013161
AGAP013162
AGAP013163
AGAP013164
AGAP013165
AGAP013166
AGAP013167
AGAP013168
AGAP013169
AGAP013170
AGAP013171
AGAP013172
AGAP013173
AGAP013174
mean_tpm
time
preZGA
ZGA_early
ZGA_late
5h
9h
11h
14h
20h
24h
L1
sex
Female
Male

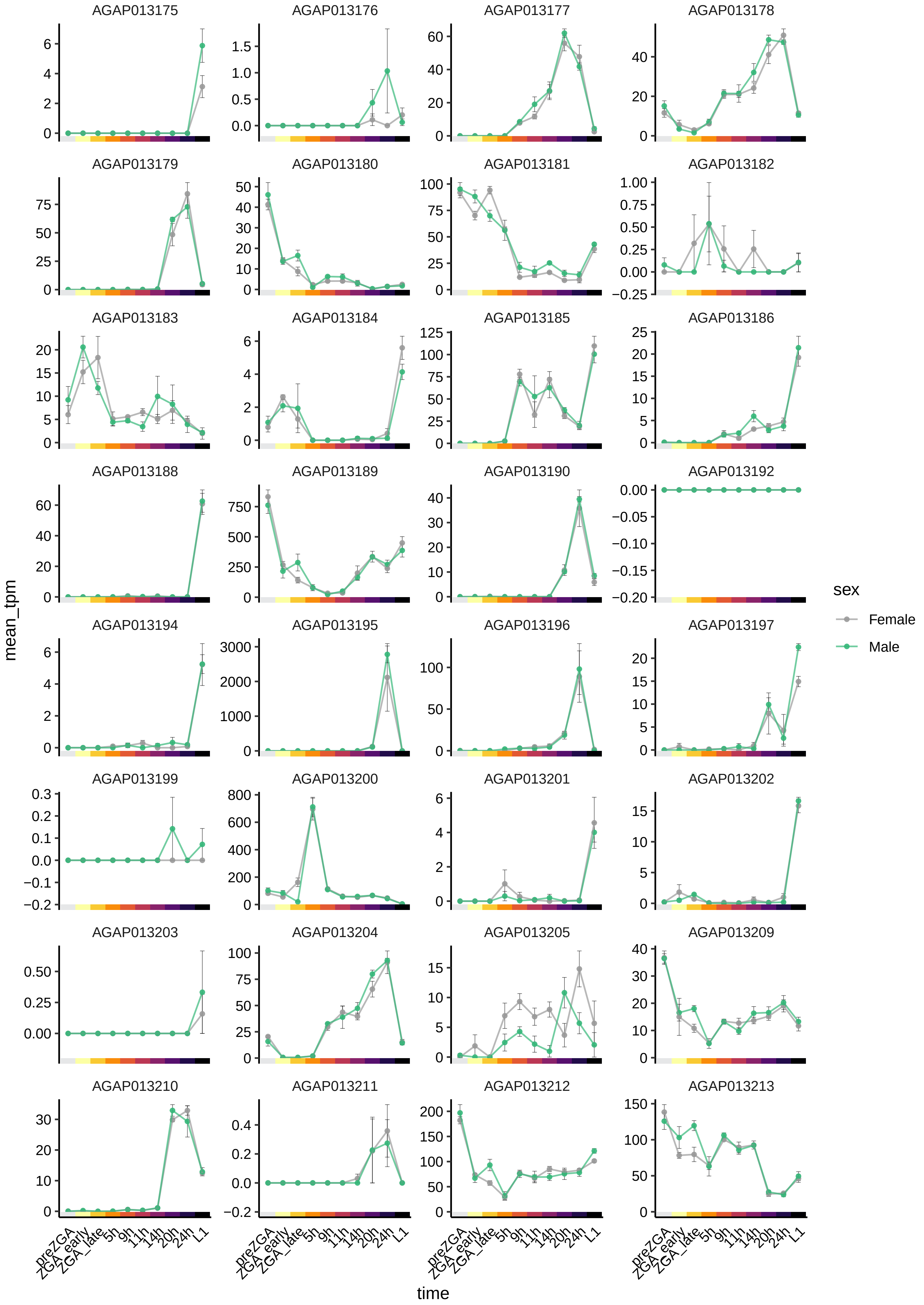

AGAP013175
AGAP013176
AGAP013177
AGAP013178
AGAP013179
AGAP013180
AGAP013181
AGAP013182
AGAP013183
AGAP013184
AGAP013185
AGAP013186
AGAP013188
AGAP013189
AGAP013190
AGAP013192
AGAP013194
AGAP013195
AGAP013196
AGAP013197
AGAP013199
AGAP013200
AGAP013201
AGAP013202
AGAP013203
AGAP013204
AGAP013205
AGAP013209
AGAP013210
AGAP013211
AGAP013212
AGAP013213
mean_tpm
time
preZGA
ZGA_early
ZGA_late
5h
9h
11h
14h
20h
24h
L1
sex
Female
Male

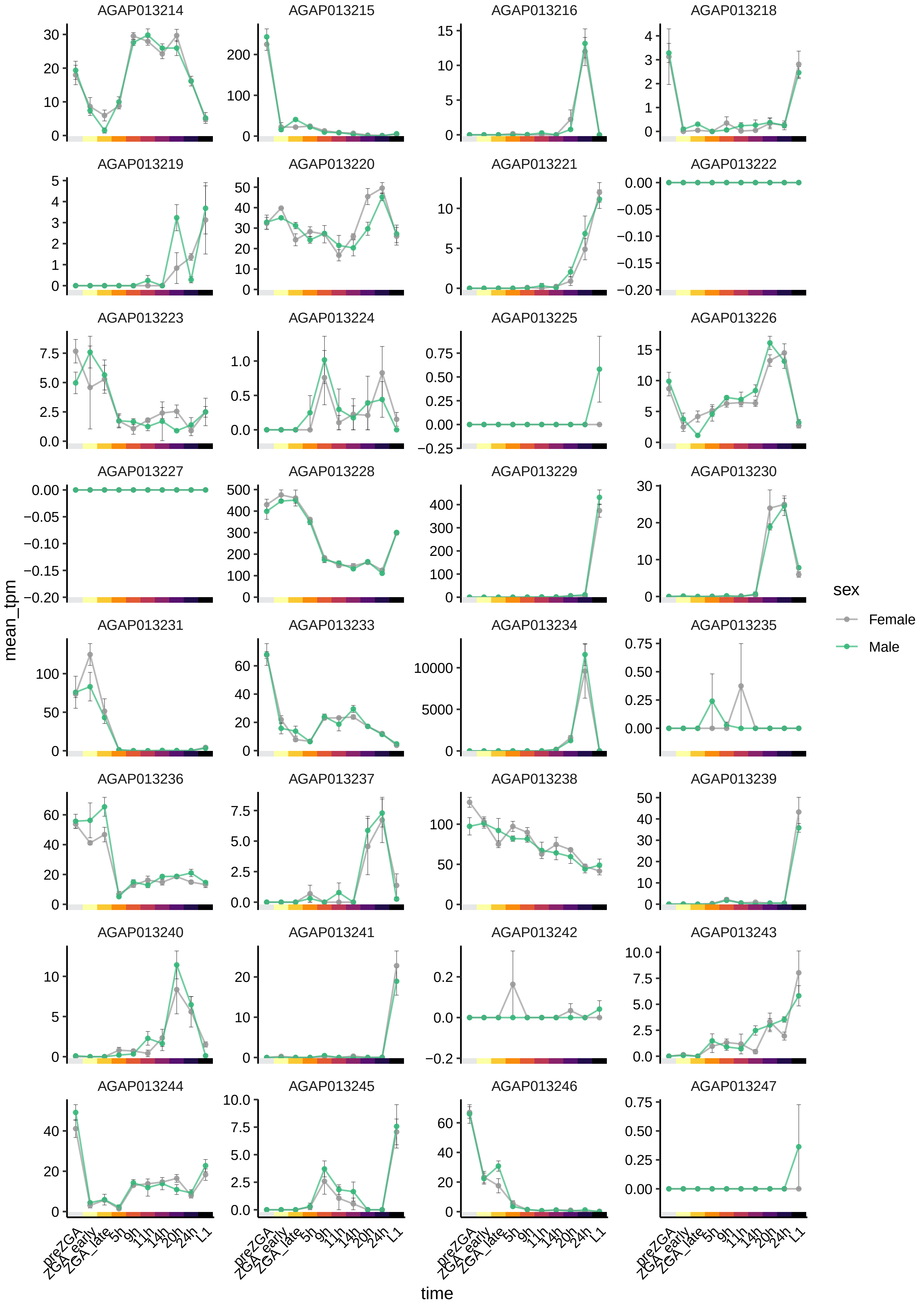

AGAP013214
AGAP013215
AGAP013216
AGAP013218
AGAP013219
AGAP013220
AGAP013221
AGAP013222
AGAP013223
AGAP013224
AGAP013225
AGAP013226
AGAP013227
AGAP013228
AGAP013229
AGAP013230
AGAP013231
AGAP013233
AGAP013234
AGAP013235
AGAP013236
AGAP013237
AGAP013238
AGAP013239
AGAP013240
AGAP013241
AGAP013242
AGAP013243
AGAP013244
AGAP013245
AGAP013246
AGAP013247
mean_tpm
time
preZGA
ZGA_early
ZGA_late
5h
9h
11h
14h
20h
24h
L1
sex
Female
Male

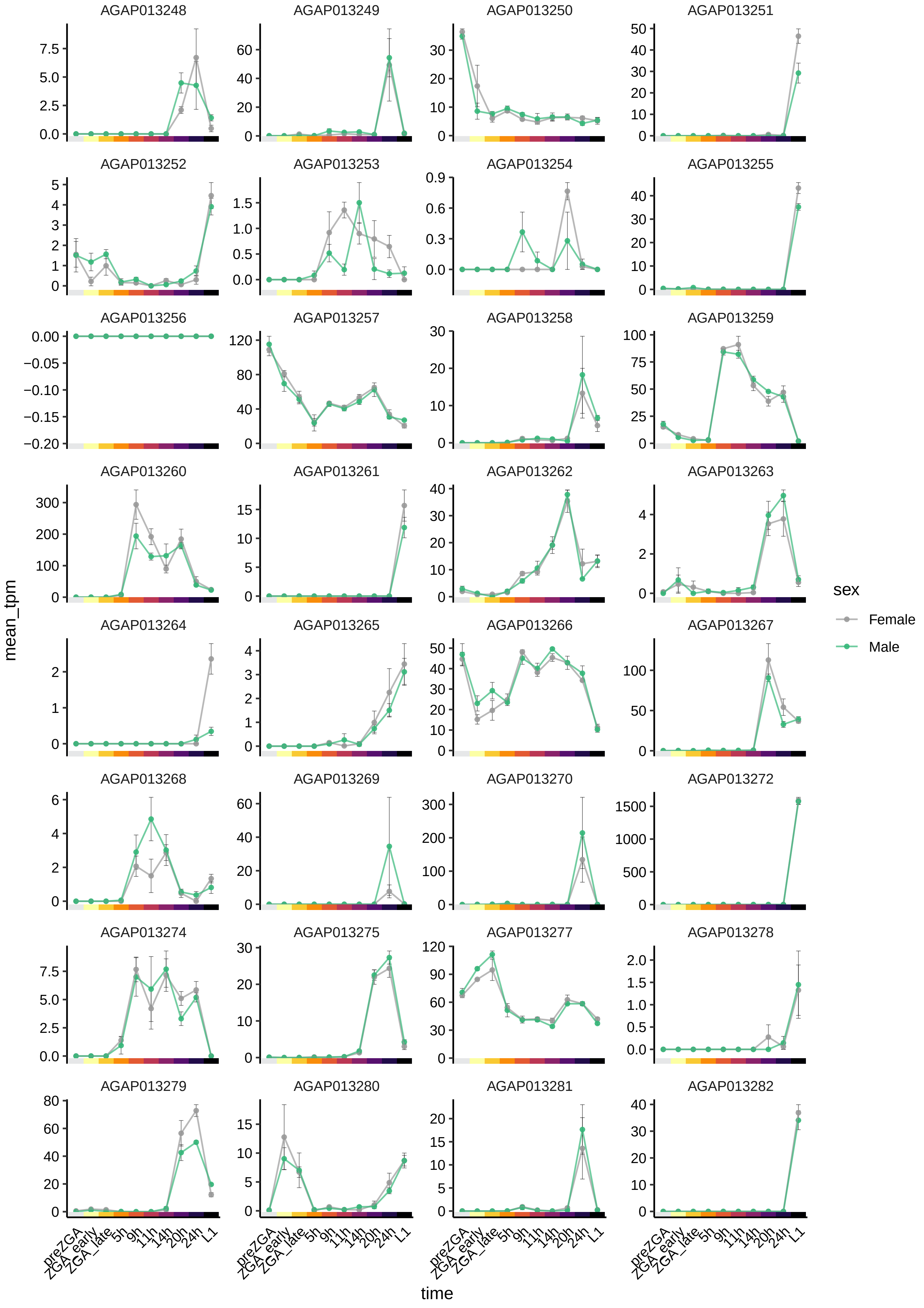

AGAP013248
AGAP013249
AGAP013250
AGAP013251
AGAP013252
AGAP013253
AGAP013254
AGAP013255
AGAP013256
AGAP013257
AGAP013258
AGAP013259
AGAP013260
AGAP013261
AGAP013262
AGAP013263
AGAP013264
AGAP013265
AGAP013266
AGAP013267
AGAP013268
AGAP013269
AGAP013270
AGAP013272
AGAP013274
AGAP013275
AGAP013277
AGAP013278
AGAP013279
AGAP013280
AGAP013281
AGAP013282
mean_tpm
time
preZGA
ZGA_early
ZGA_late
5h
9h
11h
14h
20h
24h
L1
sex
Female
Male

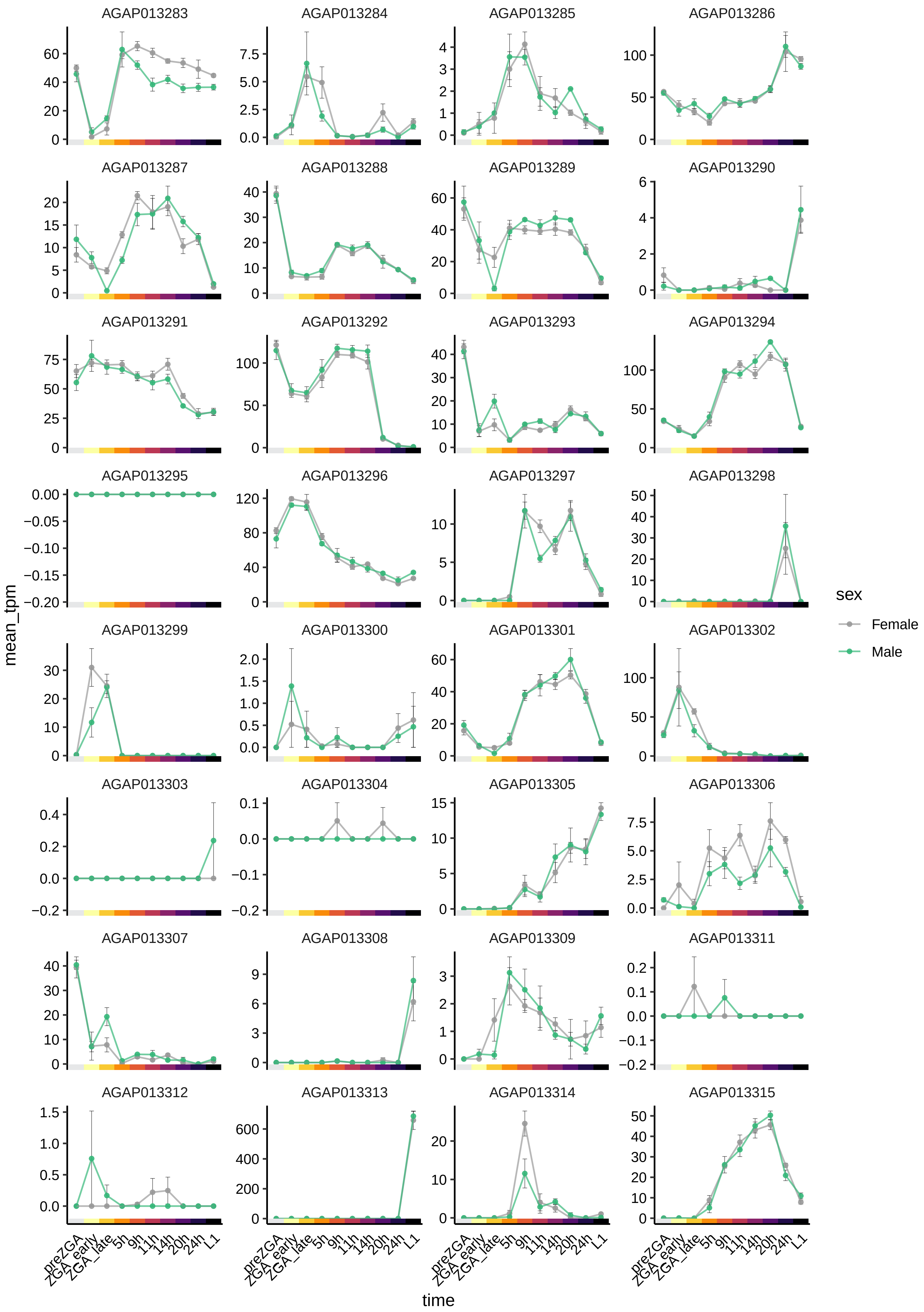
AGAP013283
AGAP013284
AGAP013285
AGAP013286
AGAP013287
AGAP013288
AGAP013289
AGAP013290
AGAP013291
AGAP013292
AGAP013293
AGAP013294
AGAP013295
AGAP013296
AGAP013297
AGAP013298
AGAP013299
AGAP013300
AGAP013301
AGAP013302
AGAP013303
AGAP013304
AGAP013305
AGAP013306
AGAP013307
AGAP013308
AGAP013309
AGAP013311
AGAP013312
AGAP013313
AGAP013314
AGAP013315
mean_tpm
time
preZGA
ZGA_early
ZGA_late
5h
9h
11h
14h
20h
24h
L1
sex
Female
Male

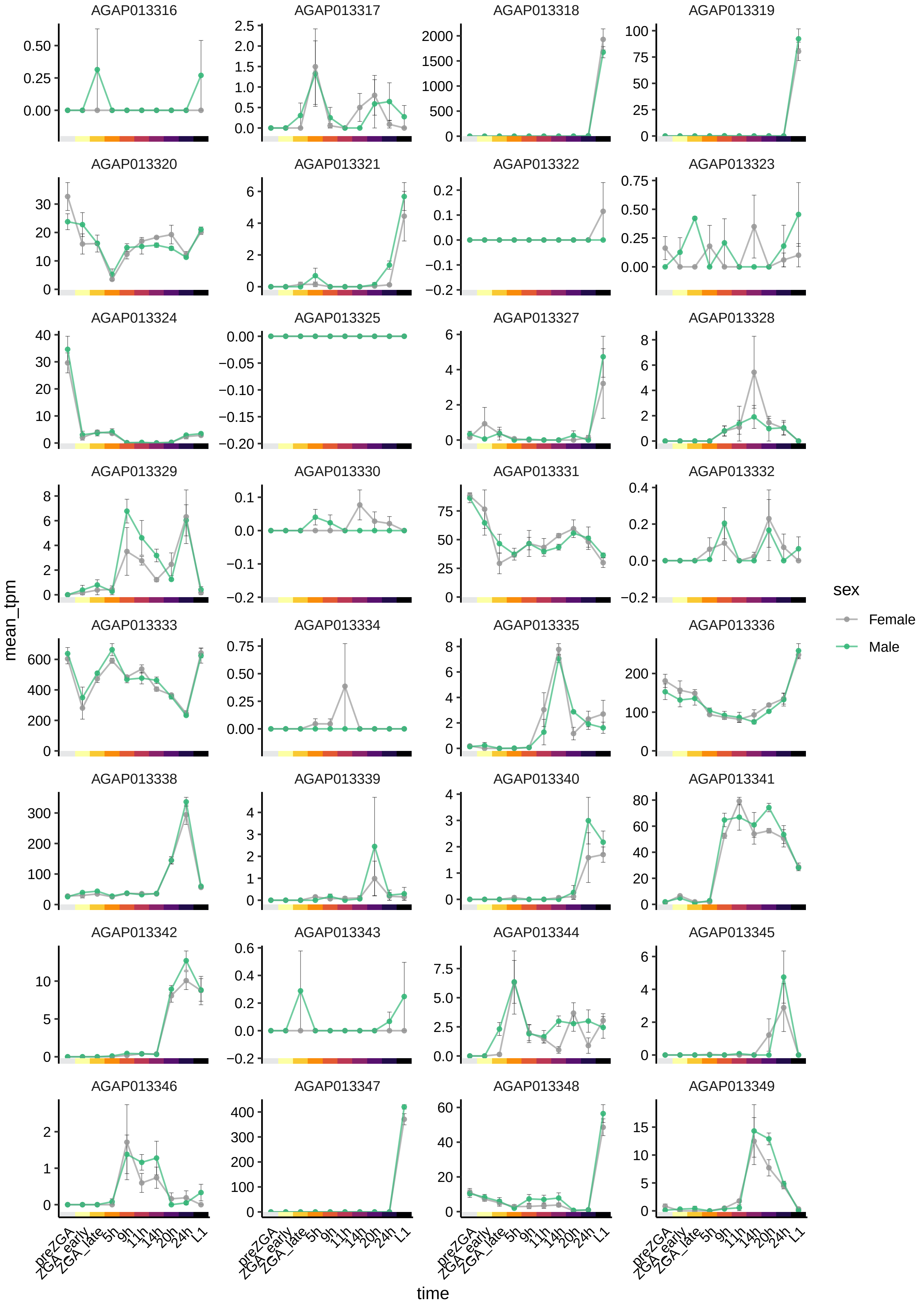

AGAP013316
AGAP013317
AGAP013318
AGAP013319
AGAP013320
AGAP013321
AGAP013322
AGAP013323
AGAP013324
AGAP013325
AGAP013327
AGAP013328
AGAP013329
AGAP013330
AGAP013331
AGAP013332
AGAP013333
AGAP013334
AGAP013335
AGAP013336
AGAP013338
AGAP013339
AGAP013340
AGAP013341
AGAP013342
AGAP013343
AGAP013344
AGAP013345
AGAP013346
AGAP013347
AGAP013348
AGAP013349
mean_tpm
time
preZGA
ZGA_early
ZGA_late
5h
9h
11h
14h
20h
24h
L1
sex
Female
Male

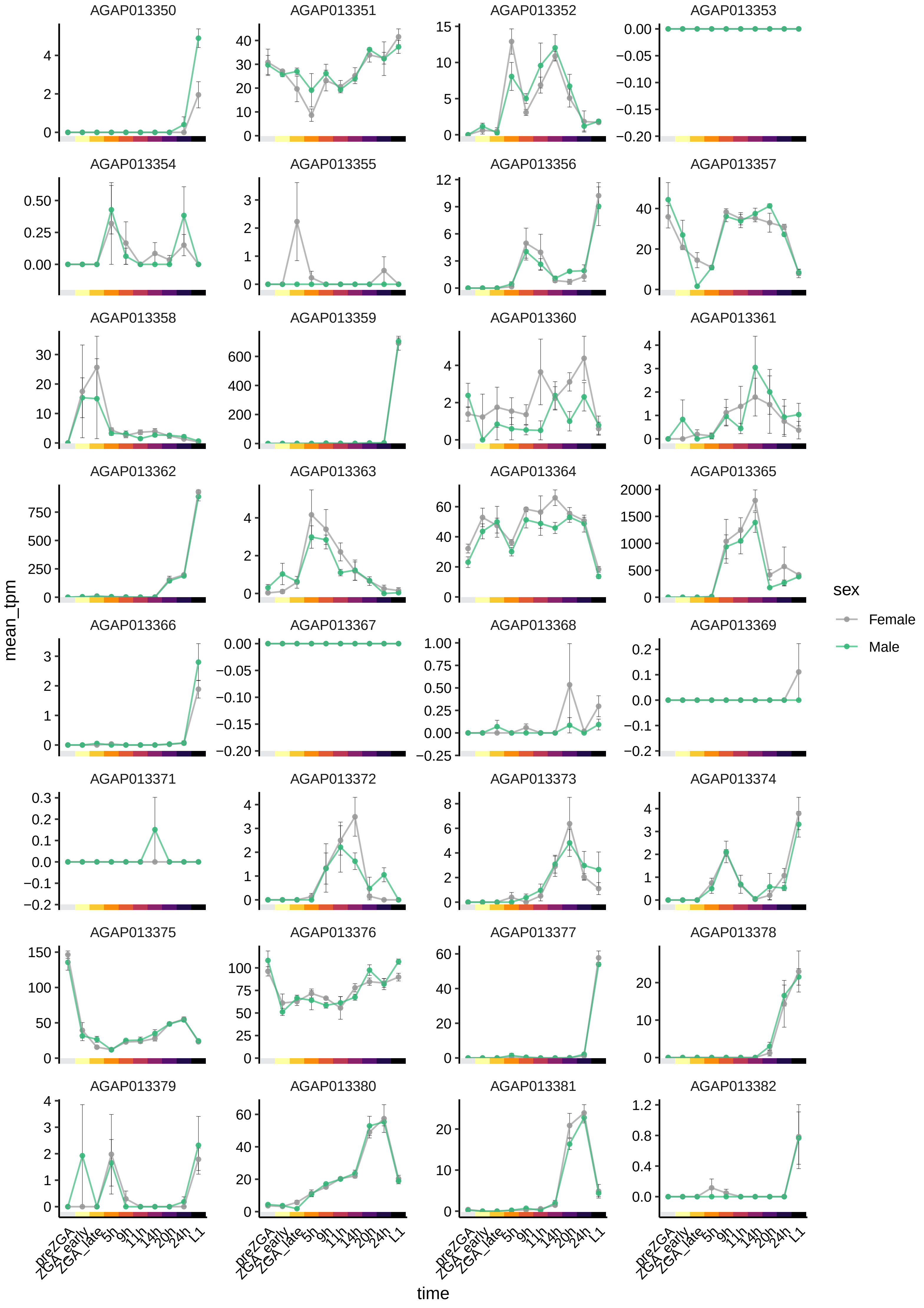

AGAP013350
AGAP013351
AGAP013352
AGAP013353
AGAP013354
AGAP013355
AGAP013356
AGAP013357
AGAP013358
AGAP013359
AGAP013360
AGAP013361
AGAP013362
AGAP013363
AGAP013364
AGAP013365
AGAP013366
AGAP013367
AGAP013368
AGAP013369
AGAP013371
AGAP013372
AGAP013373
AGAP013374
AGAP013375
AGAP013376
AGAP013377
AGAP013378
AGAP013379
AGAP013380
AGAP013381
AGAP013382
mean_tpm
time
preZGA
ZGA_early
ZGA_late
5h
9h
11h
14h
20h
24h
L1
sex
Female
Male

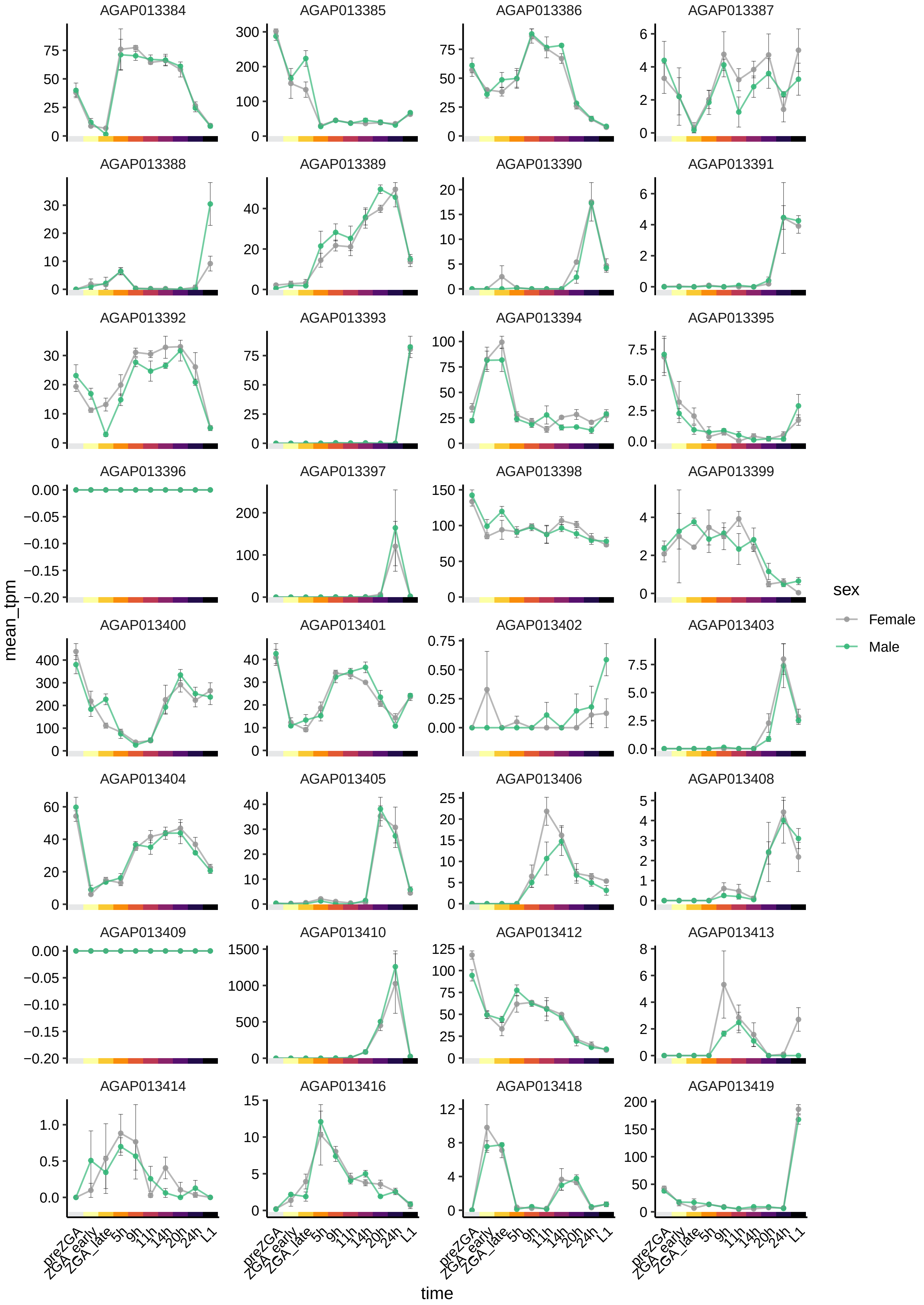

AGAP013384
AGAP013385
AGAP013386
AGAP013387
AGAP013388
AGAP013389
AGAP013390
AGAP013391
AGAP013392
AGAP013393
AGAP013394
AGAP013395
AGAP013396
AGAP013397
AGAP013398
AGAP013399
AGAP013400
AGAP013401
AGAP013402
AGAP013403
AGAP013404
AGAP013405
AGAP013406
AGAP013408
AGAP013409
AGAP013410
AGAP013412
AGAP013413
AGAP013414
AGAP013416
AGAP013418
AGAP013419
mean_tpm
time
preZGA
ZGA_early
ZGA_late
5h
9h
11h
14h
20h
24h
L1
sex
Female
Male

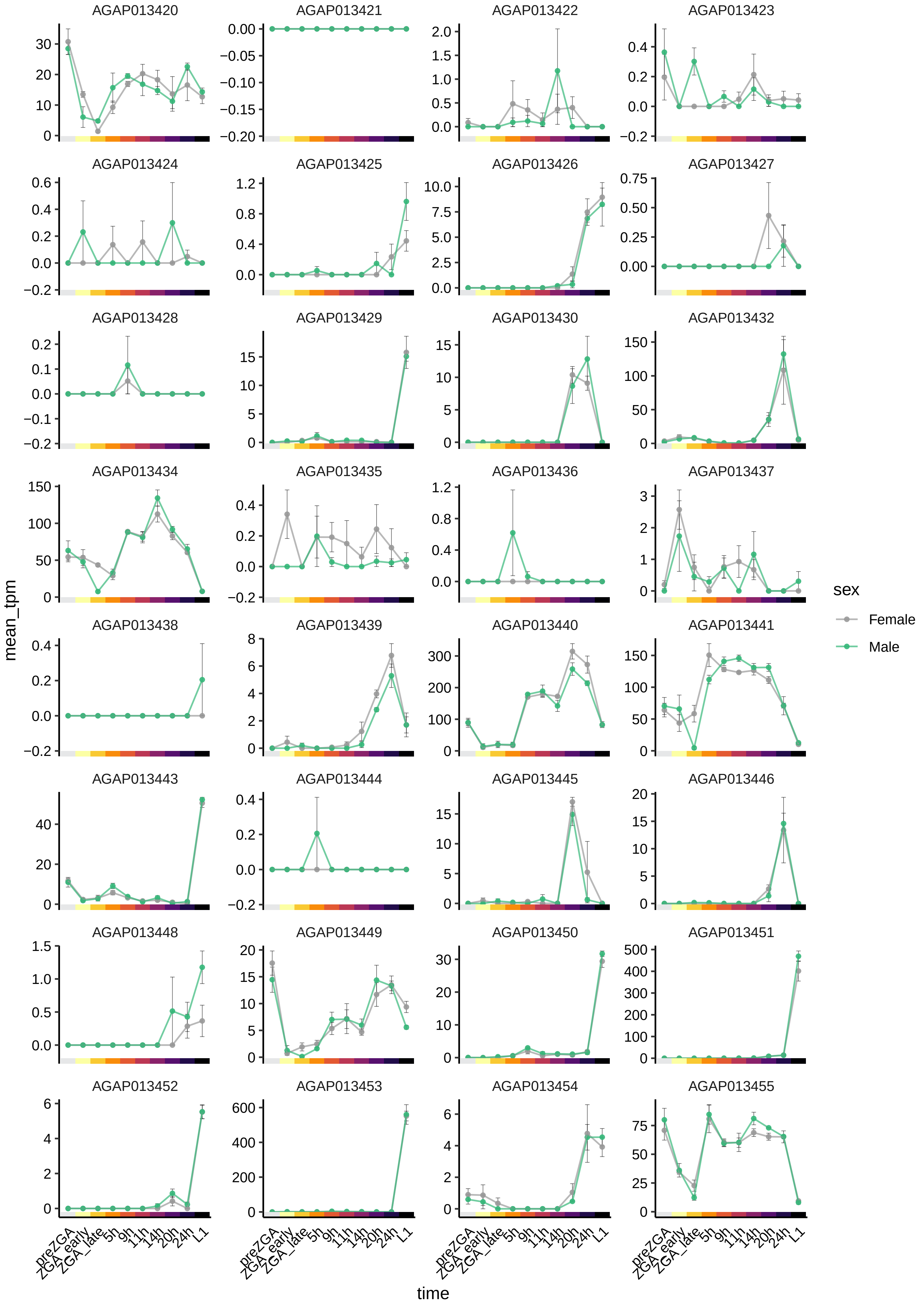

AGAP013420
AGAP013421
AGAP013422
AGAP013423
AGAP013424
AGAP013425
AGAP013426
AGAP013427
AGAP013428
AGAP013429
AGAP013430
AGAP013432
AGAP013434
AGAP013435
AGAP013436
AGAP013437
AGAP013438
AGAP013439
AGAP013440
AGAP013441
AGAP013443
AGAP013444
AGAP013445
AGAP013446
AGAP013448
AGAP013449
AGAP013450
AGAP013451
AGAP013452
AGAP013453
AGAP013454
AGAP013455
mean_tpm
time
preZGA
ZGA_early
ZGA_late
5h
9h
11h
14h
20h
24h
L1
sex
Female
Male

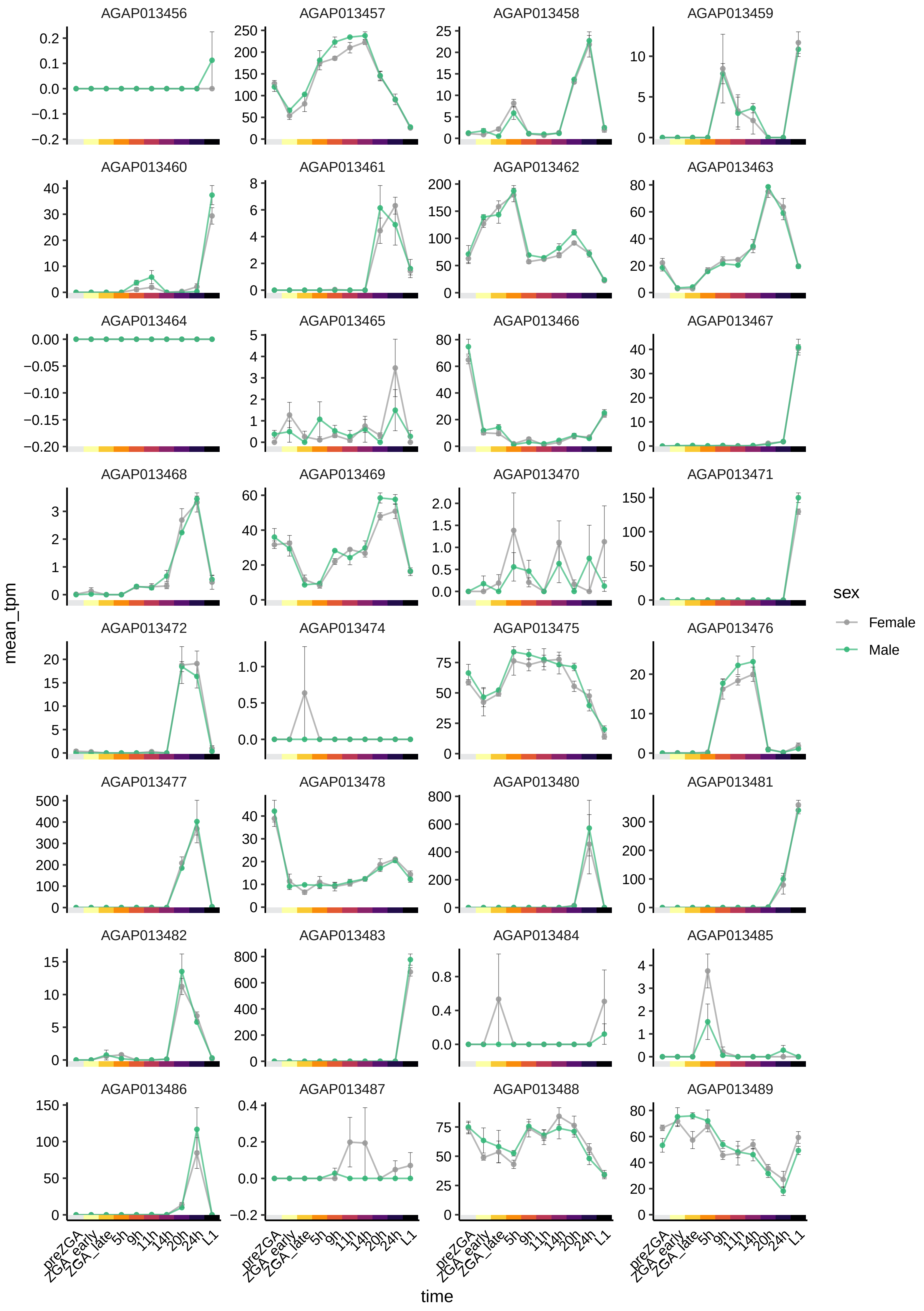

AGAP013456
AGAP013457
AGAP013458
AGAP013459
AGAP013460
AGAP013461
AGAP013462
AGAP013463
AGAP013464
AGAP013465
AGAP013466
AGAP013467
AGAP013468
AGAP013469
AGAP013470
AGAP013471
AGAP013472
AGAP013474
AGAP013475
AGAP013476
AGAP013477
AGAP013478
AGAP013480
AGAP013481
AGAP013482
AGAP013483
AGAP013484
AGAP013485
AGAP013486
AGAP013487
AGAP013488
AGAP013489
mean_tpm
time
preZGA
ZGA_early
ZGA_late
5h
9h
11h
14h
20h
24h
L1
sex
Female
Male

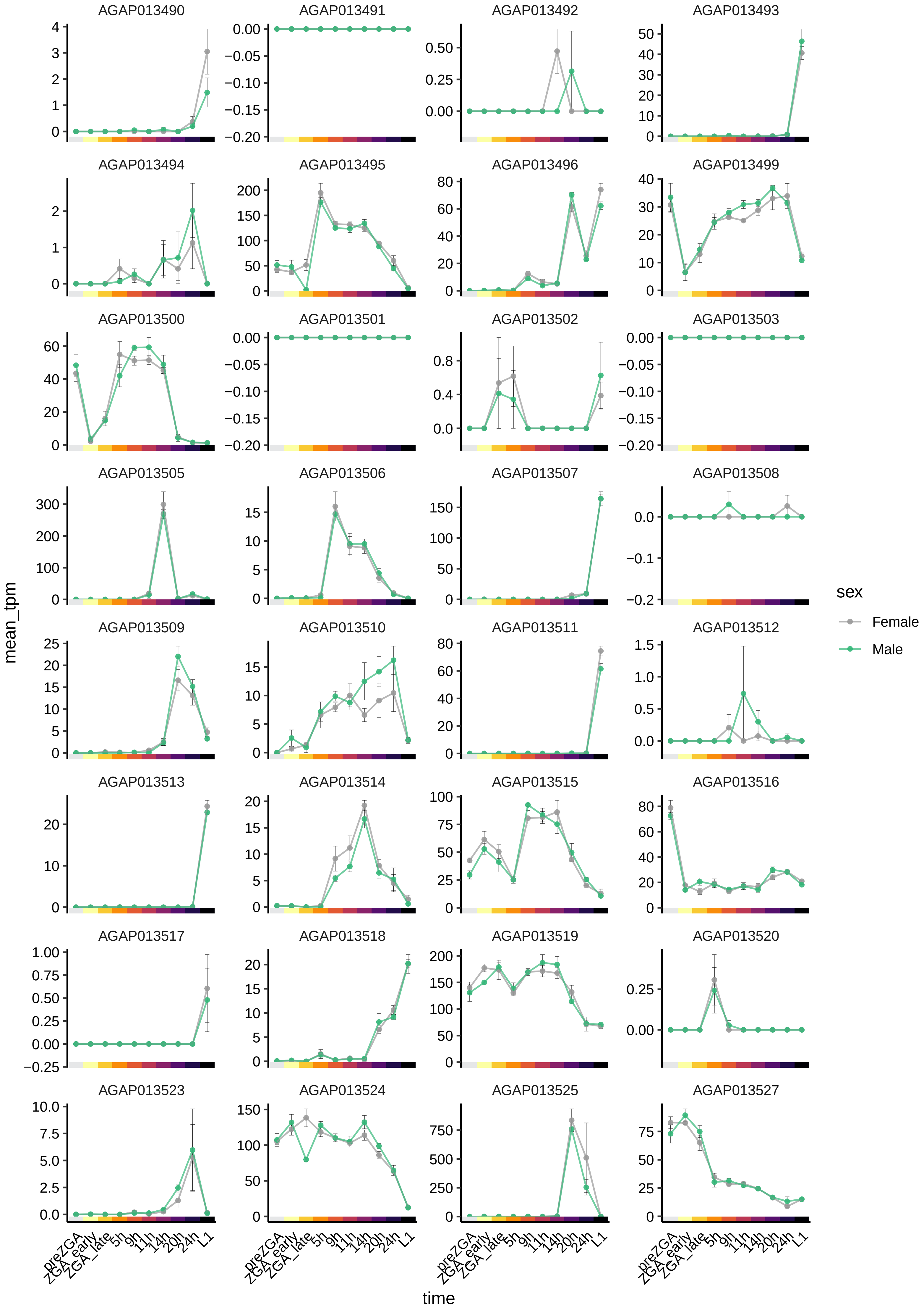
AGAP013490
AGAP013491
AGAP013492
AGAP013493
AGAP013494
AGAP013495
AGAP013496
AGAP013499
AGAP013500
AGAP013501
AGAP013502
AGAP013503
AGAP013505
AGAP013506
AGAP013507
AGAP013508
AGAP013509
AGAP013510
AGAP013511
AGAP013512
AGAP013513
AGAP013514
AGAP013515
AGAP013516
AGAP013517
AGAP013518
AGAP013519
AGAP013520
AGAP013523
AGAP013524
AGAP013525
AGAP013527
mean_tpm
time
preZGA
ZGA_early
ZGA_late
5h
9h
11h
14h
20h
24h
L1
sex
Female
Male

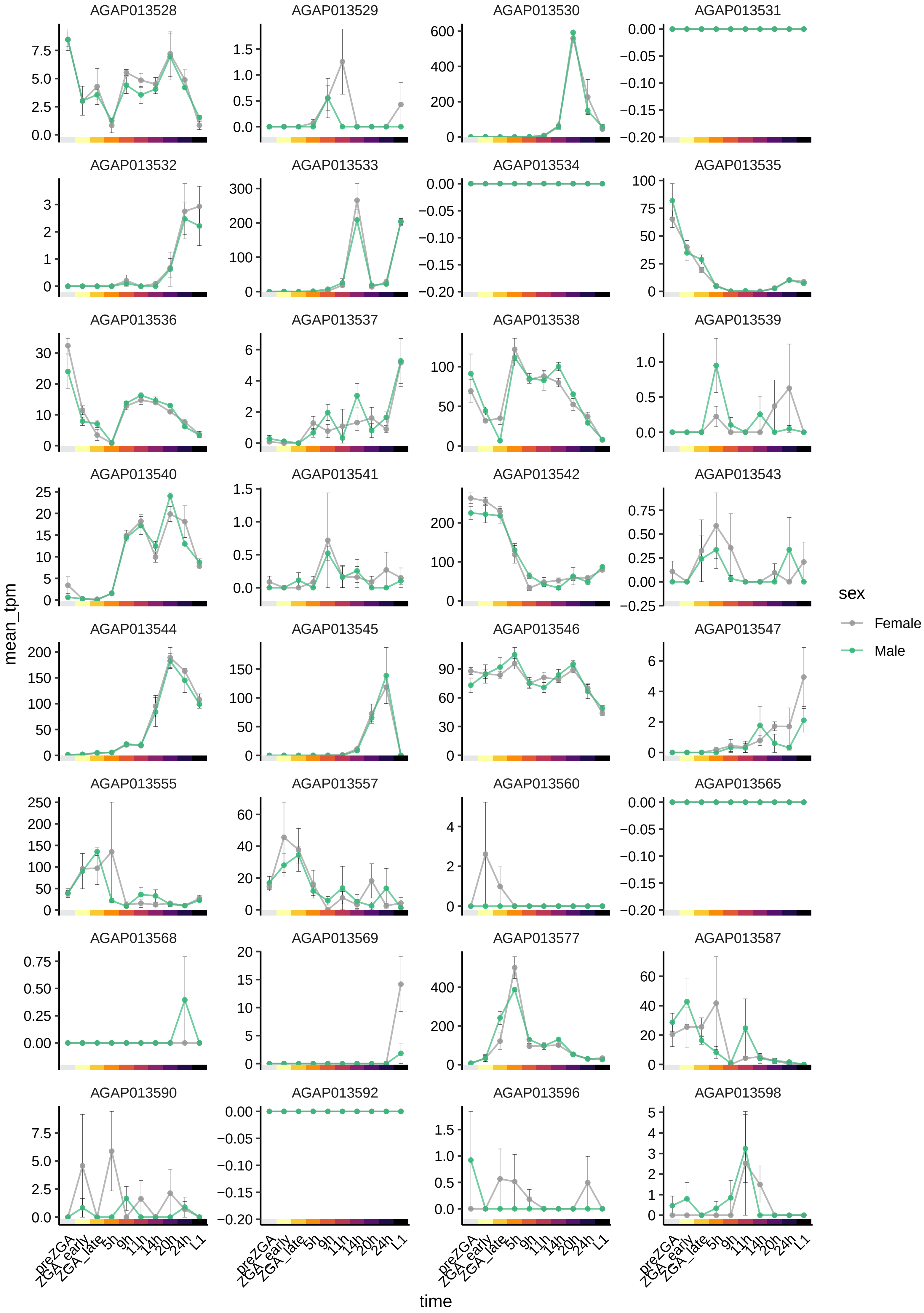
AGAP013528
AGAP013529
AGAP013530
AGAP013531
AGAP013532
AGAP013533
AGAP013534
AGAP013535
AGAP013536
AGAP013537
AGAP013538
AGAP013539
AGAP013540
AGAP013541
AGAP013542
AGAP013543
AGAP013544
AGAP013545
AGAP013546
AGAP013547
AGAP013555
AGAP013557
AGAP013560
AGAP013565
AGAP013568
AGAP013569
AGAP013577
AGAP013587
AGAP013590
AGAP013592
AGAP013596
AGAP013598
mean_tpm
time
preZGA
ZGA_early
ZGA_late
5h
9h
11h
14h
20h
24h
L1
sex
Female
Male

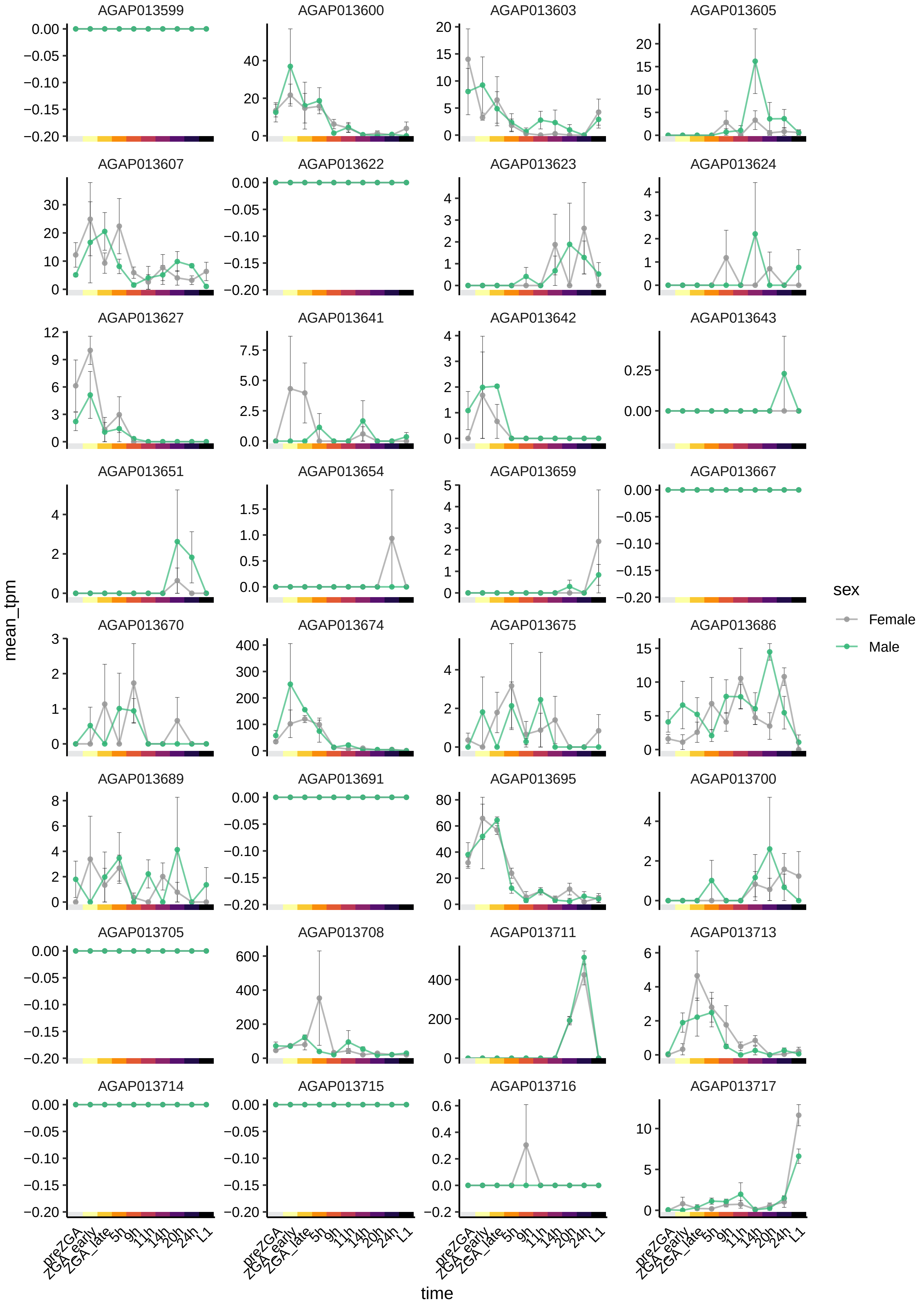
AGAP013599
AGAP013600
AGAP013603
AGAP013605
AGAP013607
AGAP013622
AGAP013623
AGAP013624
AGAP013627
AGAP013641
AGAP013642
AGAP013643
AGAP013651
AGAP013654
AGAP013659
AGAP013667
AGAP013670
AGAP013674
AGAP013675
AGAP013686
AGAP013689
AGAP013691
AGAP013695
AGAP013700
AGAP013705
AGAP013708
AGAP013711
AGAP013713
AGAP013714
AGAP013715
AGAP013716
AGAP013717
mean_tpm
time
preZGA
ZGA_early
ZGA_late
5h
9h
11h
14h
20h
24h
L1
sex
Female
Male

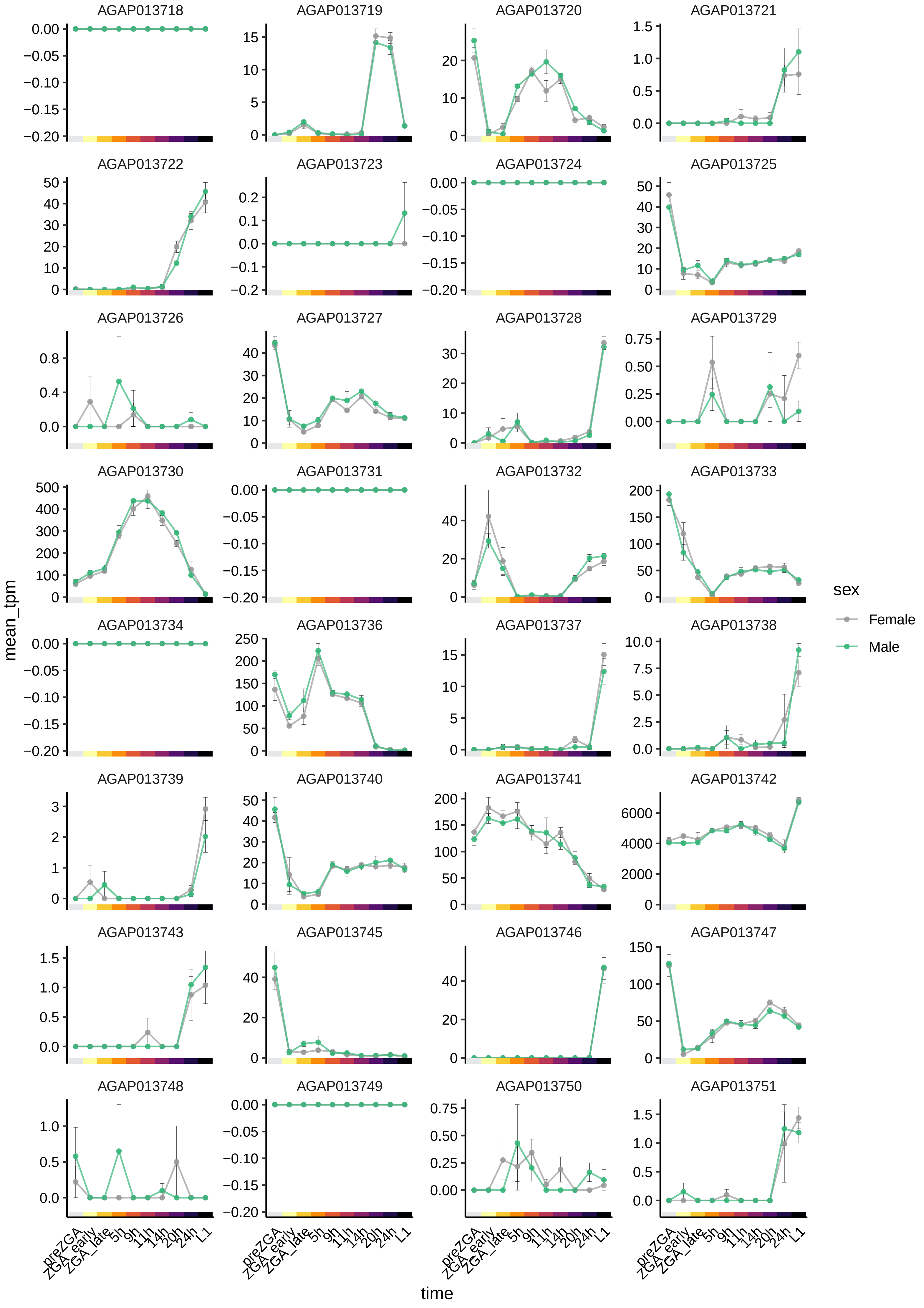
AGAP013718
AGAP013719
AGAP013720
AGAP013721
AGAP013722
AGAP013723
AGAP013724
AGAP013725
AGAP013726
AGAP013727
AGAP013728
AGAP013729
AGAP013730
AGAP013731
AGAP013732
AGAP013733
AGAP013734
AGAP013736
AGAP013737
AGAP013738
AGAP013739
AGAP013740
AGAP013741
AGAP013742
AGAP013743
AGAP013745
AGAP013746
AGAP013747
AGAP013748
AGAP013749
AGAP013750
AGAP013751
mean_tpm
time
preZGA
ZGA_early
ZGA_late
5h
9h
11h
14h
20h
24h
L1
sex
Female
Male

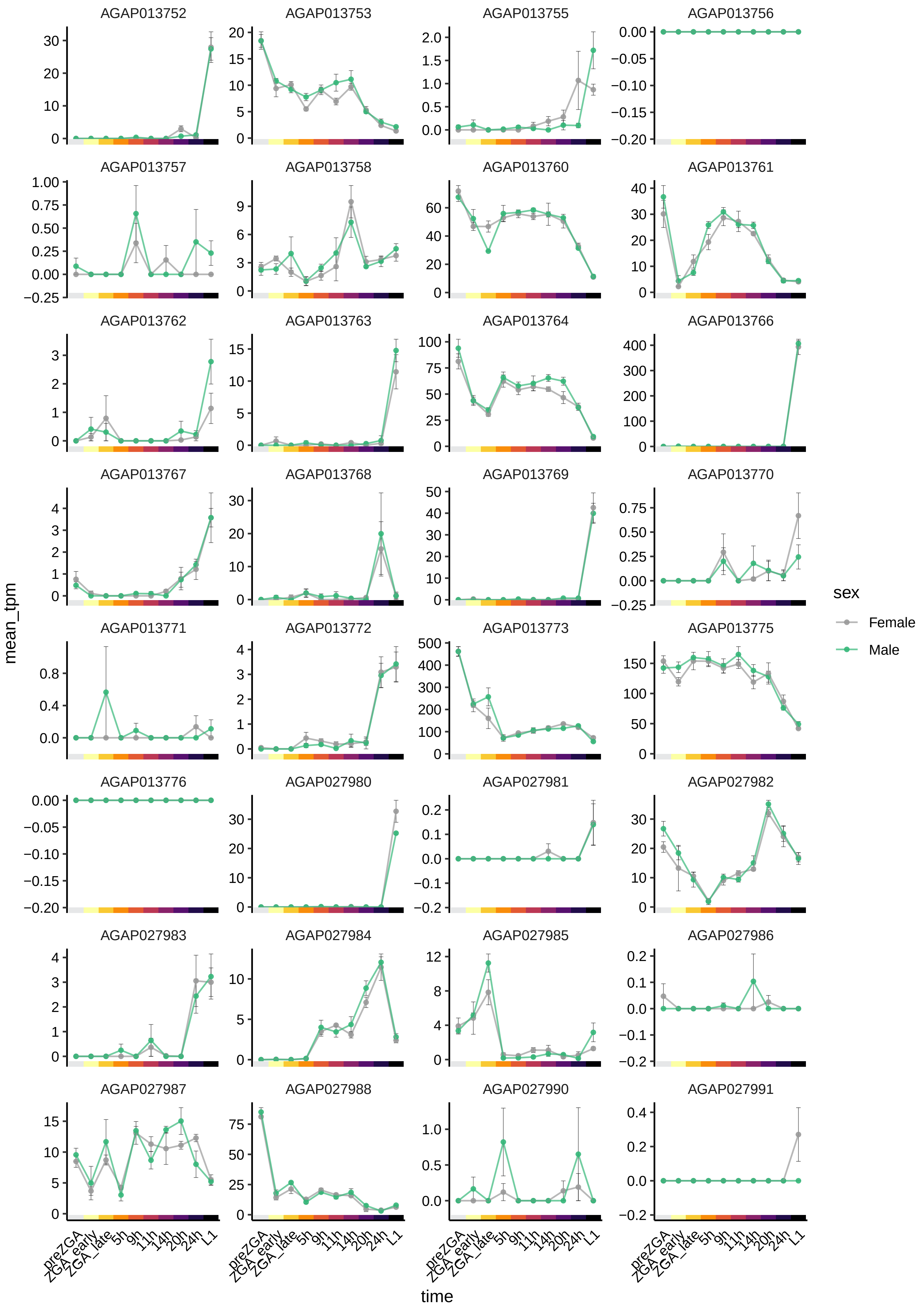
AGAP013752
AGAP013753
AGAP013755
AGAP013756
AGAP013757
AGAP013758
AGAP013760
AGAP013761
AGAP013762
AGAP013763
AGAP013764
AGAP013766
AGAP013767
AGAP013768
AGAP013769
AGAP013770
AGAP013771
AGAP013772
AGAP013773
AGAP013775
AGAP013776
AGAP027980
AGAP027981
AGAP027982
AGAP027983
AGAP027984
AGAP027985
AGAP027986
AGAP027987
AGAP027988
AGAP027990
AGAP027991
mean_tpm
time
preZGA
ZGA_early
ZGA_late
5h
9h
11h
14h
20h
24h
L1
sex
Female
Male

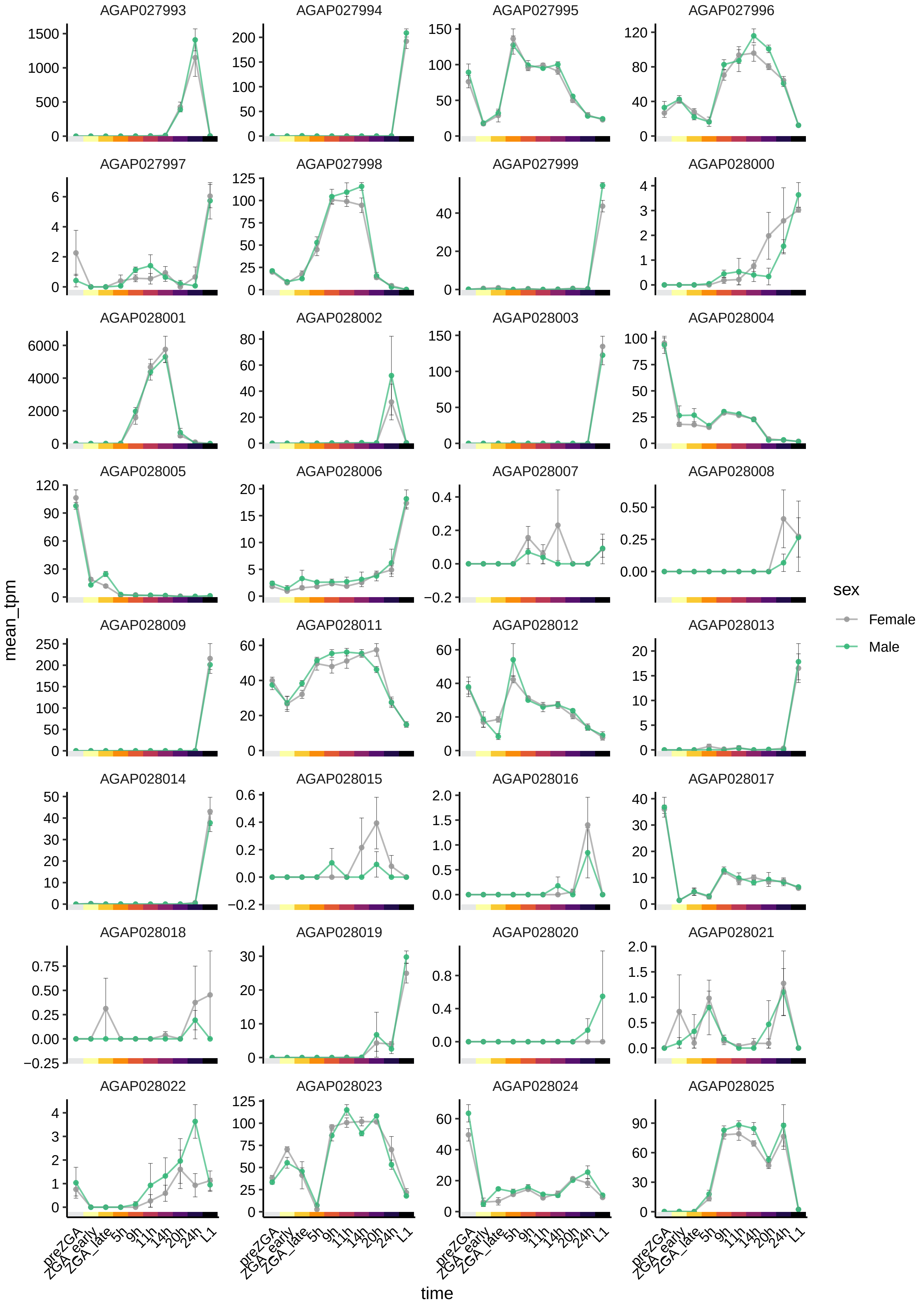

AGAP027993
AGAP027994
AGAP027995
AGAP027996
AGAP027997
AGAP027998
AGAP027999
AGAP028000
AGAP028001
AGAP028002
AGAP028003
AGAP028004
AGAP028005
AGAP028006
AGAP028007
AGAP028008
AGAP028009
AGAP028011
AGAP028012
AGAP028013
AGAP028014
AGAP028015
AGAP028016
AGAP028017
AGAP028018
AGAP028019
AGAP028020
AGAP028021
AGAP028022
AGAP028023
AGAP028024
AGAP028025
mean_tpm
time
preZGA
ZGA_early
ZGA_late
5h
9h
11h
14h
20h
24h
L1
sex
Female
Male

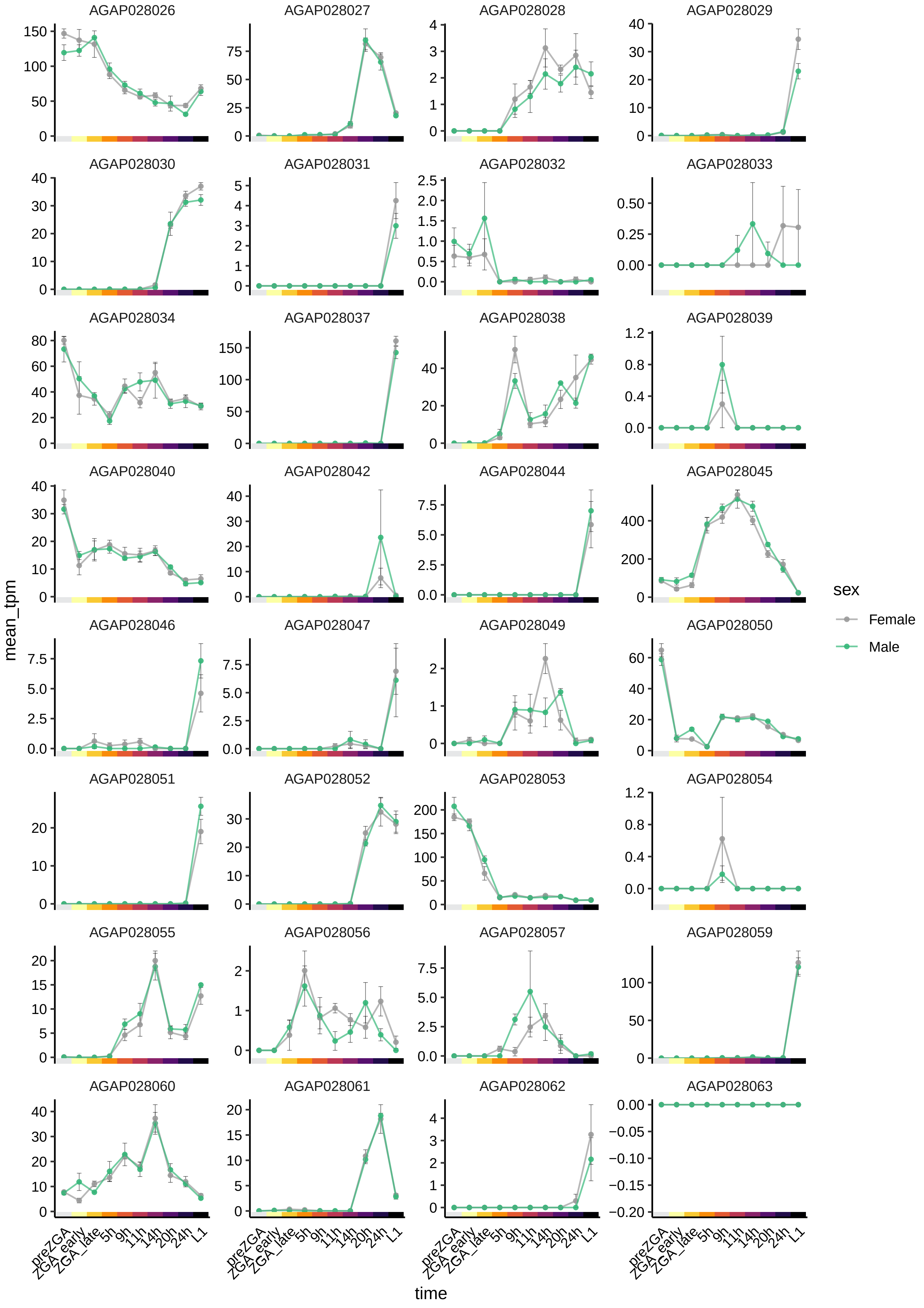

AGAP028026
AGAP028027
AGAP028028
AGAP028029
AGAP028030
AGAP028031
AGAP028032
AGAP028033
AGAP028034
AGAP028037
AGAP028038
AGAP028039
AGAP028040
AGAP028042
AGAP028044
AGAP028045
AGAP028046
AGAP028047
AGAP028049
AGAP028050
AGAP028051
AGAP028052
AGAP028053
AGAP028054
AGAP028055
AGAP028056
AGAP028057
AGAP028059
AGAP028060
AGAP028061
AGAP028062
AGAP028063
mean_tpm
time
preZGA
ZGA_early
ZGA_late
5h
9h
11h
14h
20h
24h
L1
sex
Female
Male

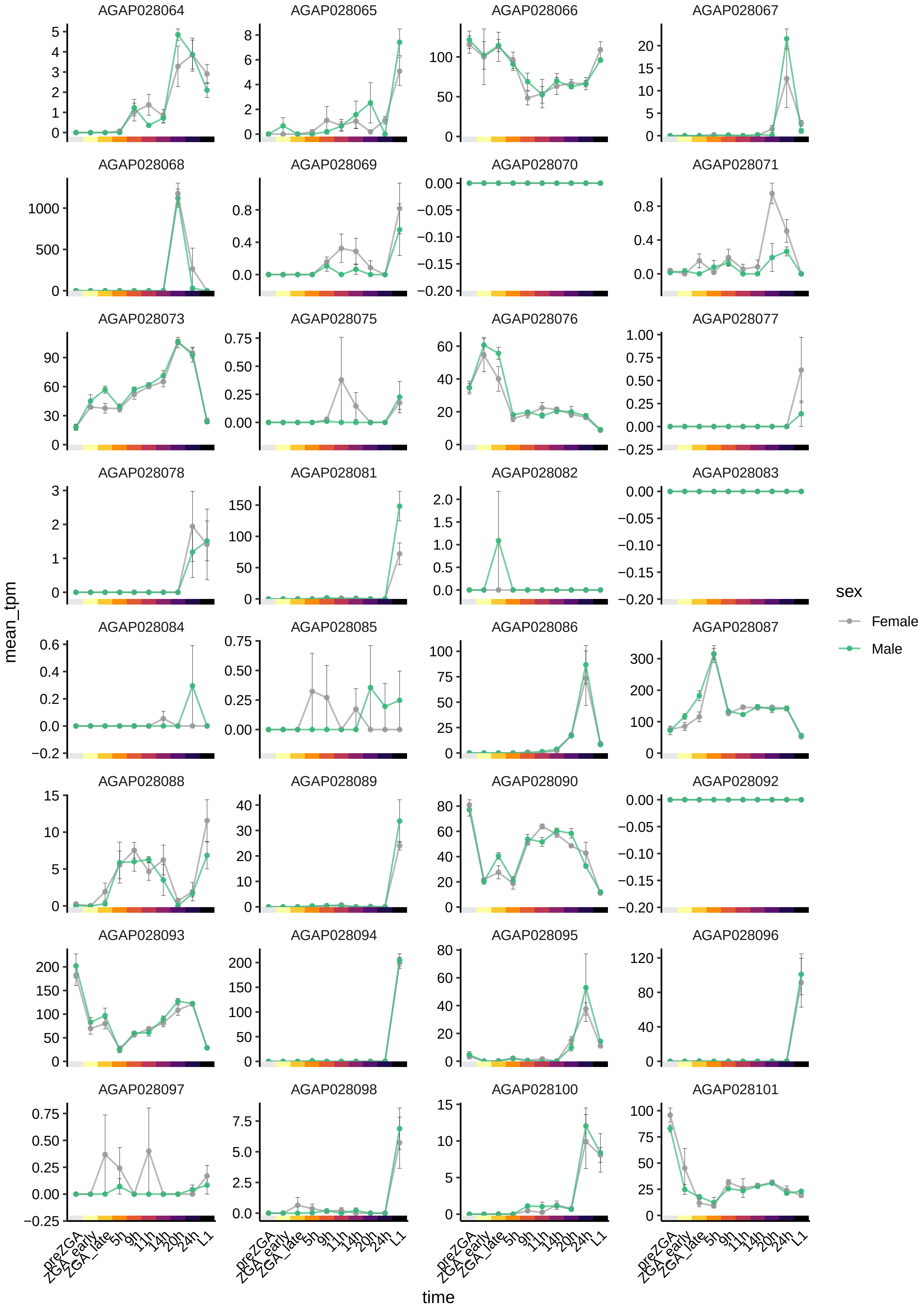
AGAP028064
AGAP028065
AGAP028066
AGAP028067
AGAP028068
AGAP028069
AGAP028070
AGAP028071
AGAP028073
AGAP028075
AGAP028076
AGAP028077
AGAP028078
AGAP028081
AGAP028082
AGAP028083
AGAP028084
AGAP028085
AGAP028086
AGAP028087
AGAP028088
AGAP028089
AGAP028090
AGAP028092
AGAP028093
AGAP028094
AGAP028095
AGAP028096
AGAP028097
AGAP028098
AGAP028100
AGAP028101
mean_tpm
time
preZGA
ZGA_early
ZGA_late
5h
9h
11h
14h
20h
24h
L1
sex
Female
Male

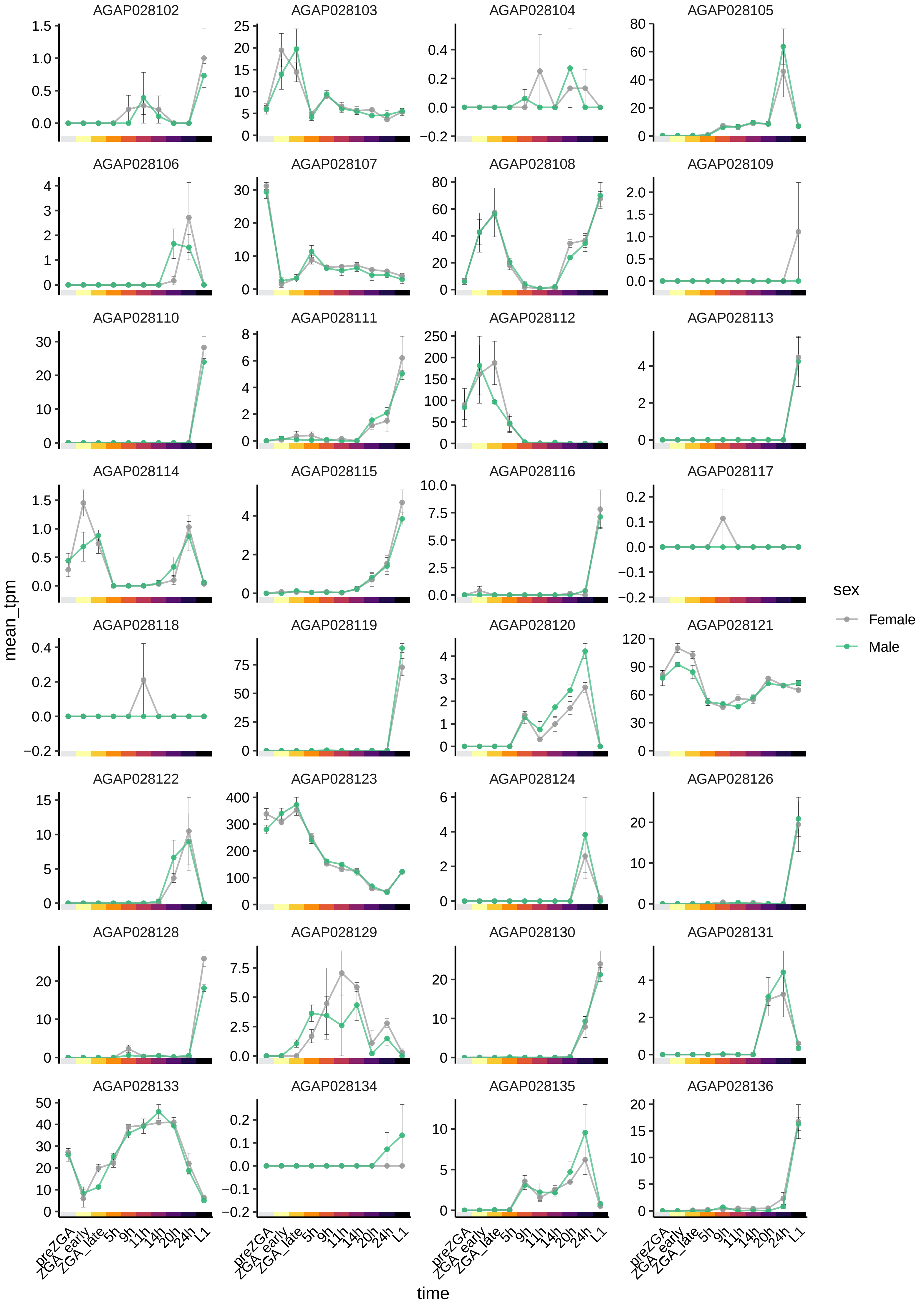

AGAP028102
AGAP028103
AGAP028104
AGAP028105
AGAP028106
AGAP028107
AGAP028108
AGAP028109
AGAP028110
AGAP028111
AGAP028112
AGAP028113
AGAP028114
AGAP028115
AGAP028116
AGAP028117
AGAP028118
AGAP028119
AGAP028120
AGAP028121
AGAP028122
AGAP028123
AGAP028124
AGAP028126
AGAP028128
AGAP028129
AGAP028130
AGAP028131
AGAP028133
AGAP028134
AGAP028135
AGAP028136
mean_tpm
time
preZGA
ZGA_early
ZGA_late
5h
9h
11h
14h
20h
24h
L1
sex
Female
Male

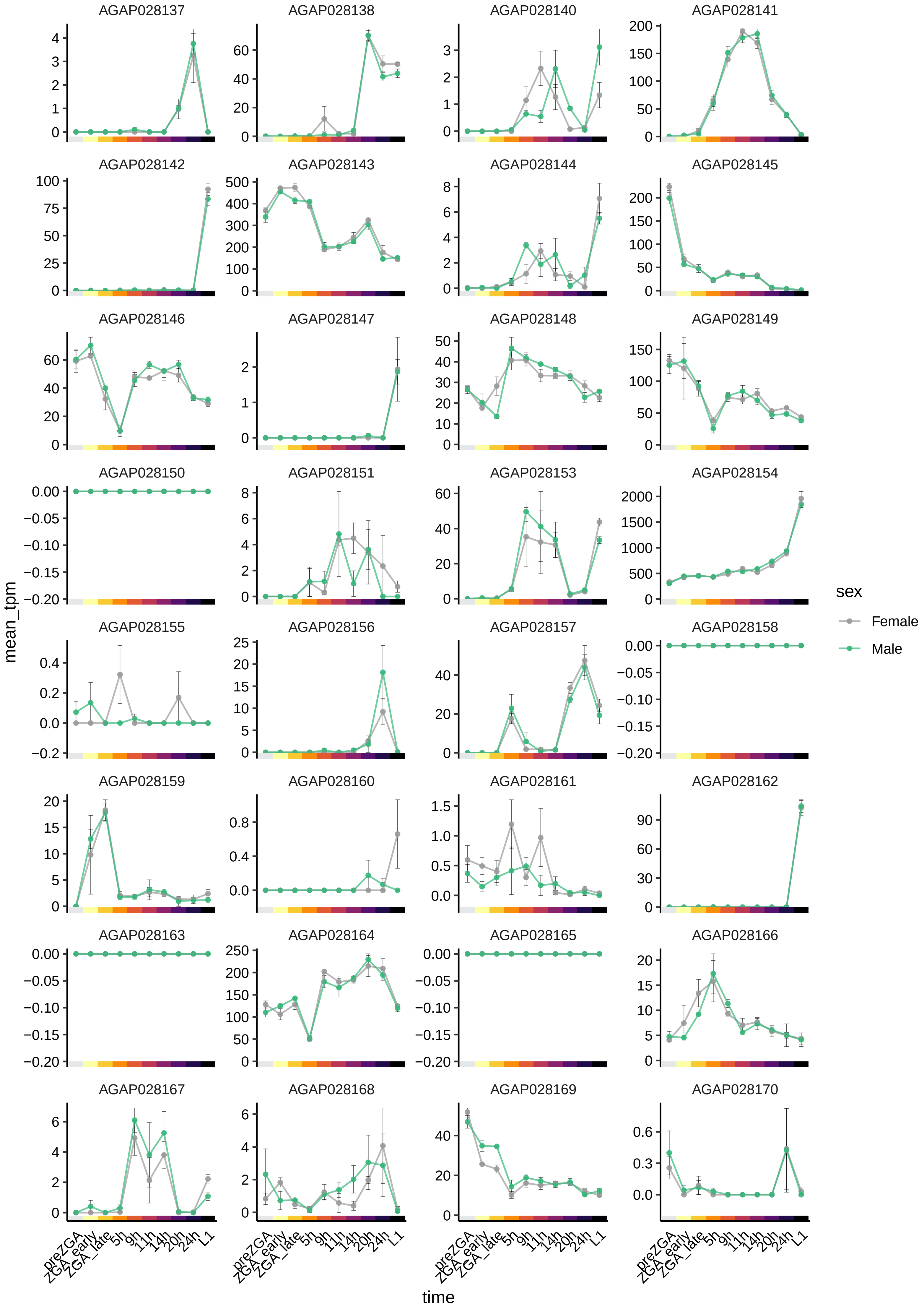

AGAP028137
AGAP028138
AGAP028140
AGAP028141
AGAP028142
AGAP028143
AGAP028144
AGAP028145
AGAP028146
AGAP028147
AGAP028148
AGAP028149
AGAP028150
AGAP028151
AGAP028153
AGAP028154
AGAP028155
AGAP028156
AGAP028157
AGAP028158
AGAP028159
AGAP028160
AGAP028161
AGAP028162
AGAP028163
AGAP028164
AGAP028165
AGAP028166
AGAP028167
AGAP028168
AGAP028169
AGAP028170
mean_tpm
time
preZGA
ZGA_early
ZGA_late
5h
9h
11h
14h
20h
24h
L1
sex
Female
Male

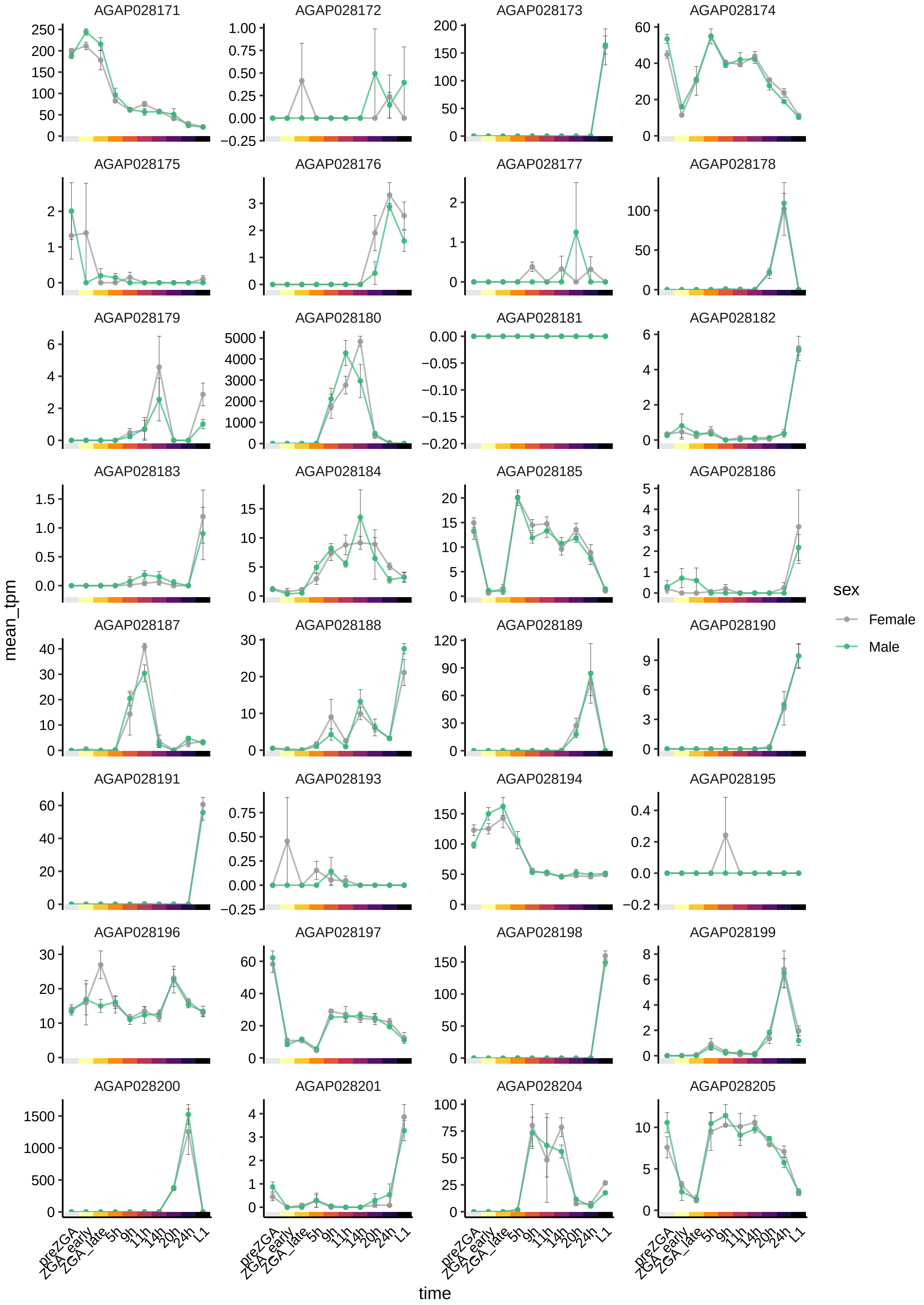
AGAP028171
AGAP028172
AGAP028173
AGAP028174
AGAP028175
AGAP028176
AGAP028177
AGAP028178
AGAP028179
AGAP028180
AGAP028181
AGAP028182
AGAP028183
AGAP028184
AGAP028185
AGAP028186
AGAP028187
AGAP028188
AGAP028189
AGAP028190
AGAP028191
AGAP028193
AGAP028194
AGAP028195
AGAP028196
AGAP028197
AGAP028198
AGAP028199
AGAP028200
AGAP028201
AGAP028204
AGAP028205
mean_tpm
time
preZGA
ZGA_early
ZGA_late
5h
9h
11h
14h
20h
24h
L1
sex
Female
Male

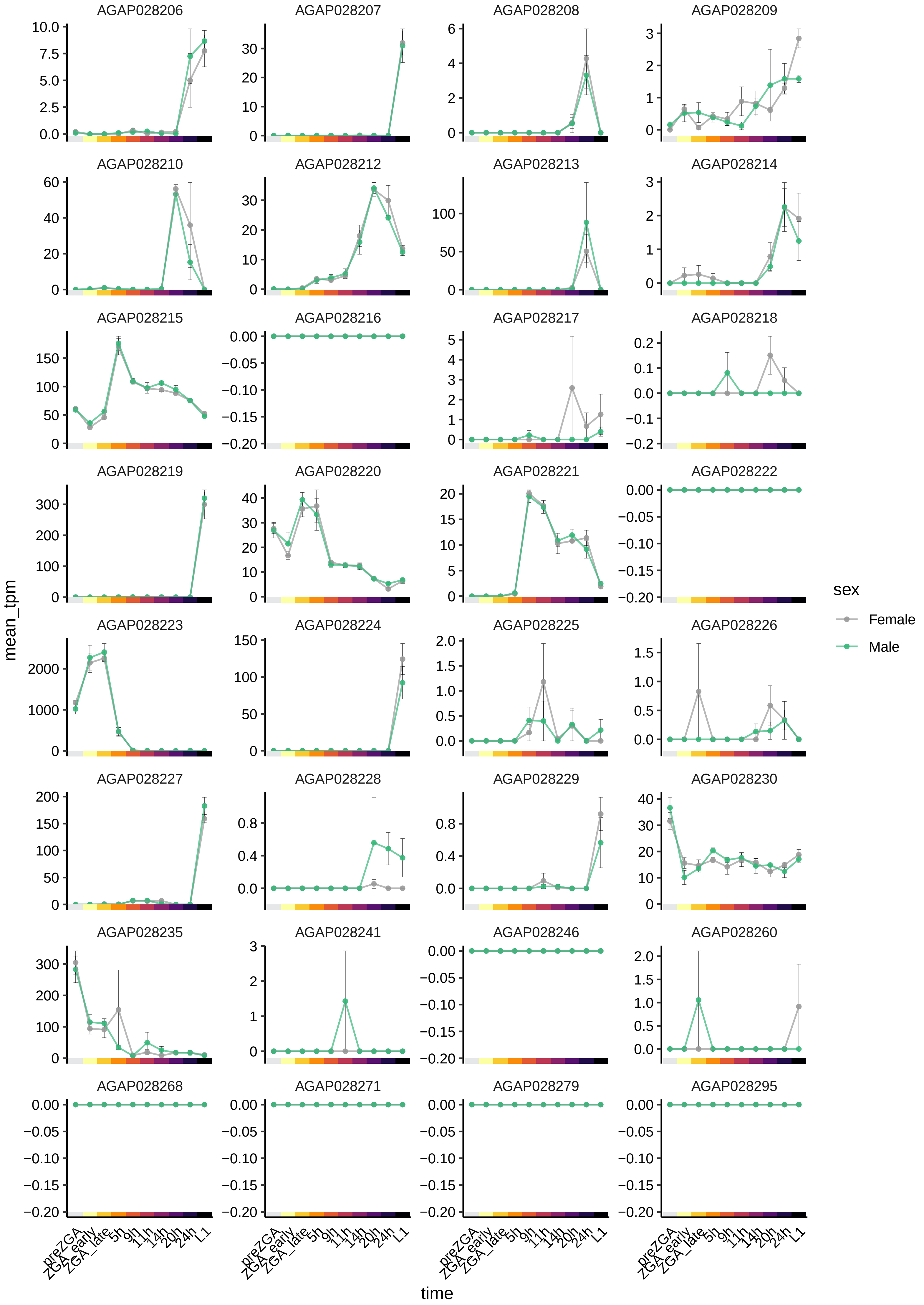

AGAP028206
AGAP028207
AGAP028208
AGAP028209
AGAP028210
AGAP028212
AGAP028213
AGAP028214
AGAP028215
AGAP028216
AGAP028217
AGAP028218
AGAP028219
AGAP028220
AGAP028221
AGAP028222
AGAP028223
AGAP028224
AGAP028225
AGAP028226
AGAP028227
AGAP028228
AGAP028229
AGAP028230
AGAP028235
AGAP028241
AGAP028246
AGAP028260
AGAP028268
AGAP028271
AGAP028279
AGAP028295
mean_tpm
time
preZGA
ZGA_early
ZGA_late
5h
9h
11h
14h
20h
24h
L1
sex
Female
Male

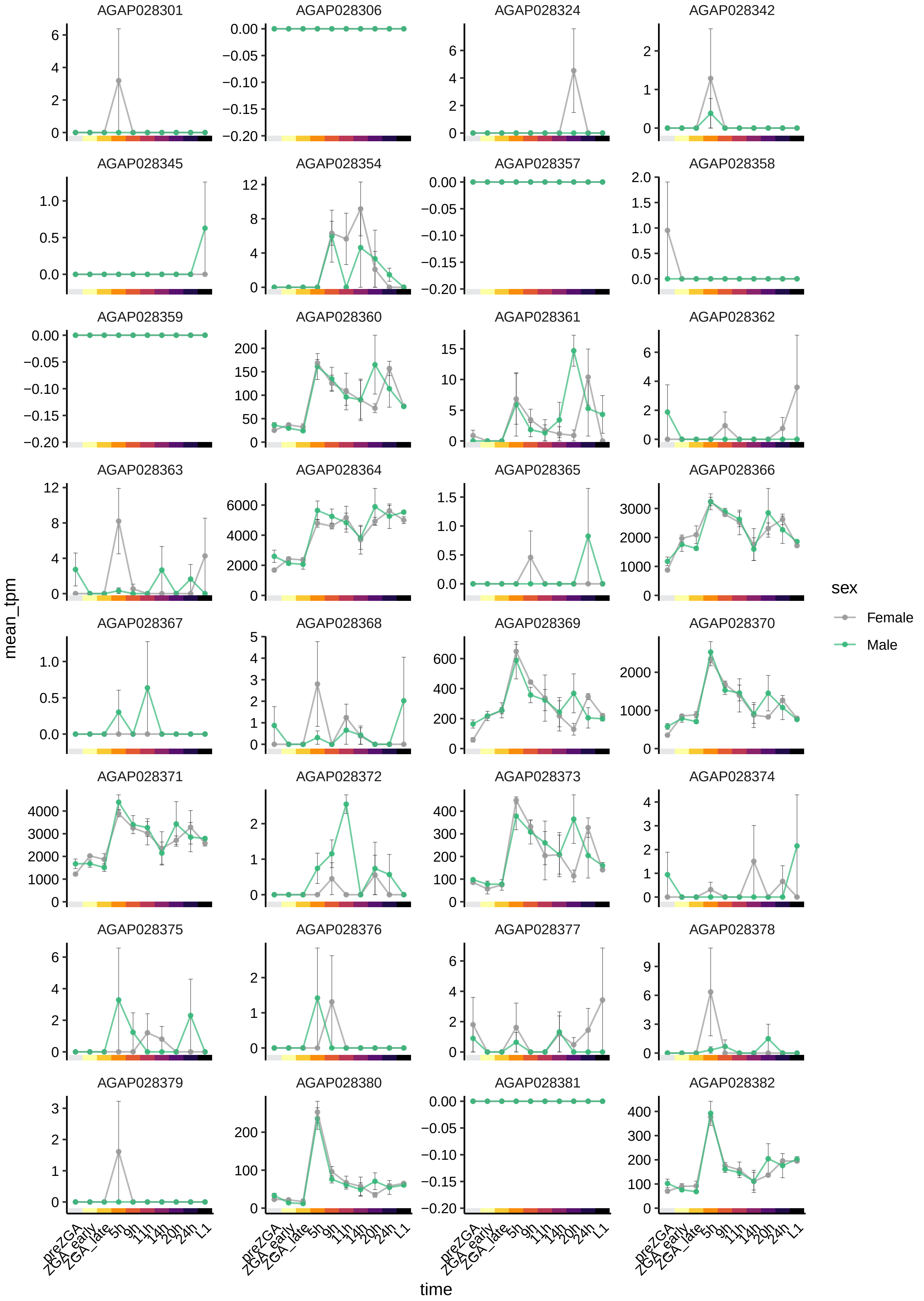

AGAP028301
AGAP028306
AGAP028324
AGAP028342
AGAP028345
AGAP028354
AGAP028357
AGAP028358
AGAP028359
AGAP028360
AGAP028361
AGAP028362
AGAP028363
AGAP028364
AGAP028365
AGAP028366
AGAP028367
AGAP028368
AGAP028369
AGAP028370
AGAP028371
AGAP028372
AGAP028373
AGAP028374
AGAP028375
AGAP028376
AGAP028377
AGAP028378
AGAP028379
AGAP028380
AGAP028381
AGAP028382
mean_tpm
time
preZGA
ZGA_early
ZGA_late
5h
9h
11h
14h
20h
24h
L1
sex
Female
Male

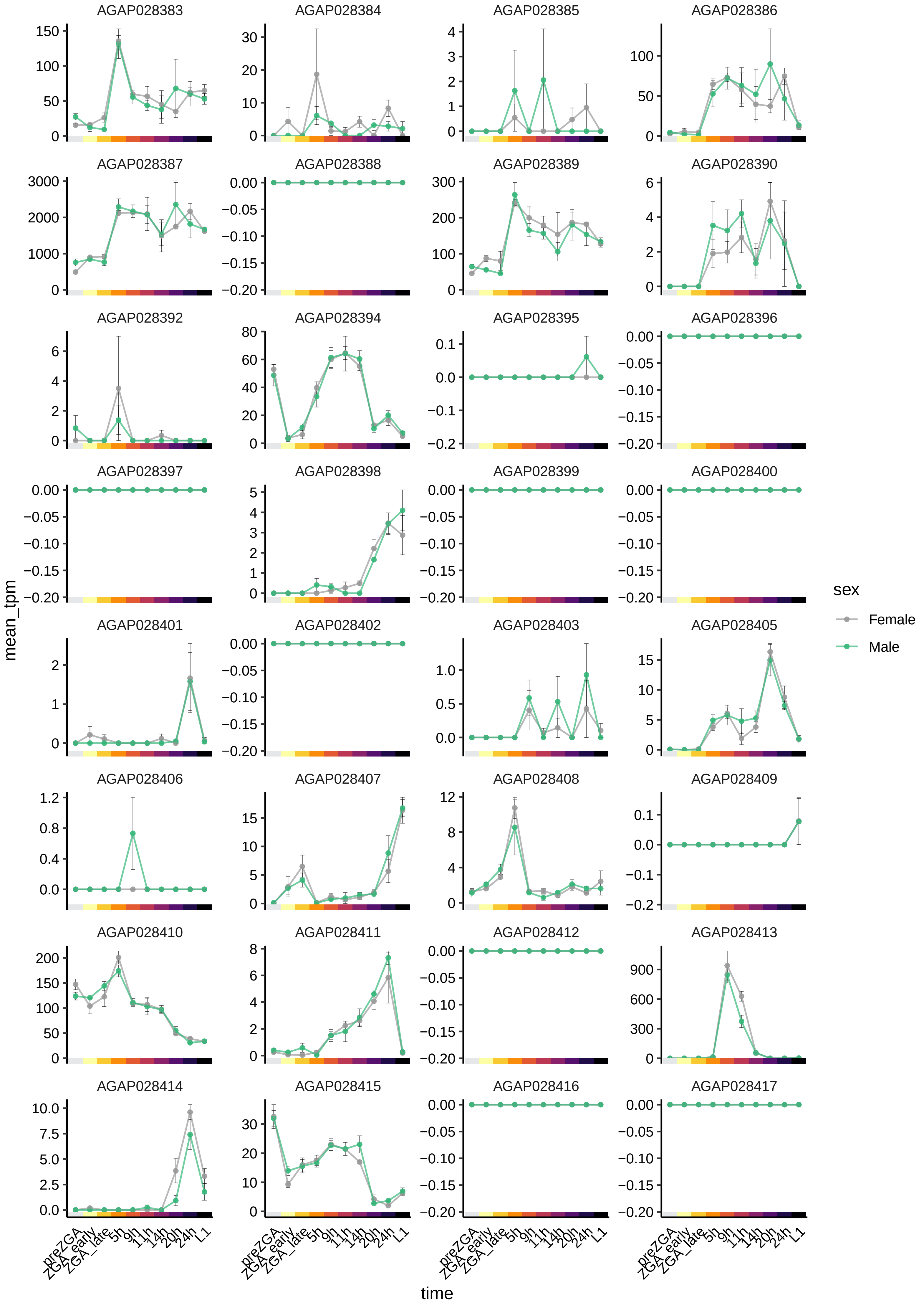
AGAP028383
AGAP028384
AGAP028385
AGAP028386
AGAP028387
AGAP028388
AGAP028389
AGAP028390
AGAP028392
AGAP028394
AGAP028395
AGAP028396
AGAP028397
AGAP028398
AGAP028399
AGAP028400
AGAP028401
AGAP028402
AGAP028403
AGAP028405
AGAP028406
AGAP028407
AGAP028408
AGAP028409
AGAP028410
AGAP028411
AGAP028412
AGAP028413
AGAP028414
AGAP028415
AGAP028416
AGAP028417
mean_tpm
time
preZGA
ZGA_early
ZGA_late
5h
9h
11h
14h
20h
24h
L1
sex
Female
Male

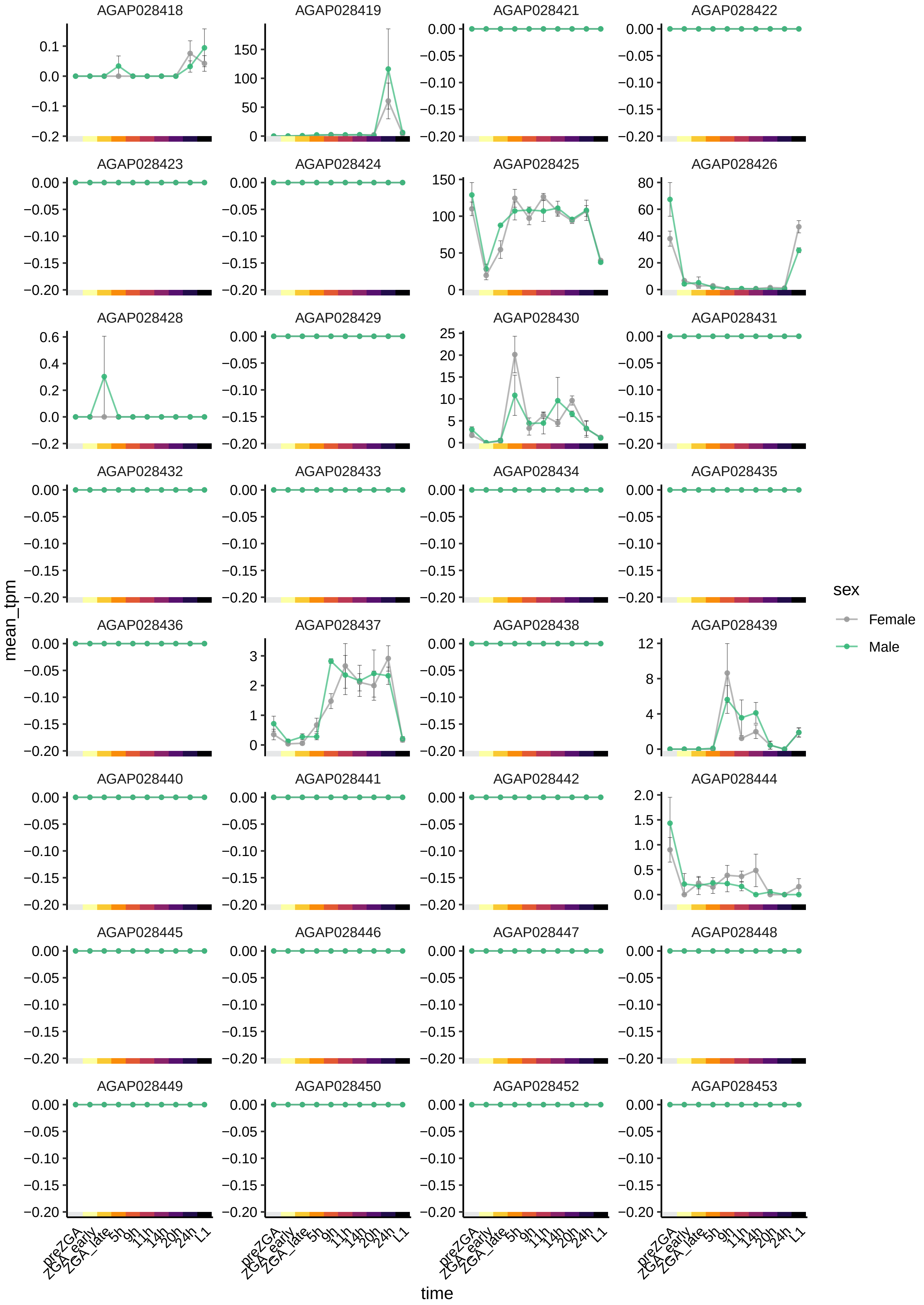

AGAP028418
AGAP028419
AGAP028421
AGAP028422
AGAP028423
AGAP028424
AGAP028425
AGAP028426
AGAP028428
AGAP028429
AGAP028430
AGAP028431
AGAP028432
AGAP028433
AGAP028434
AGAP028435
AGAP028436
AGAP028437
AGAP028438
AGAP028439
AGAP028440
AGAP028441
AGAP028442
AGAP028444
AGAP028445
AGAP028446
AGAP028447
AGAP028448
AGAP028449
AGAP028450
AGAP028452
AGAP028453
mean_tpm
time
preZGA
ZGA_early
ZGA_late
5h
9h
11h
14h
20h
24h
L1
sex
Female
Male

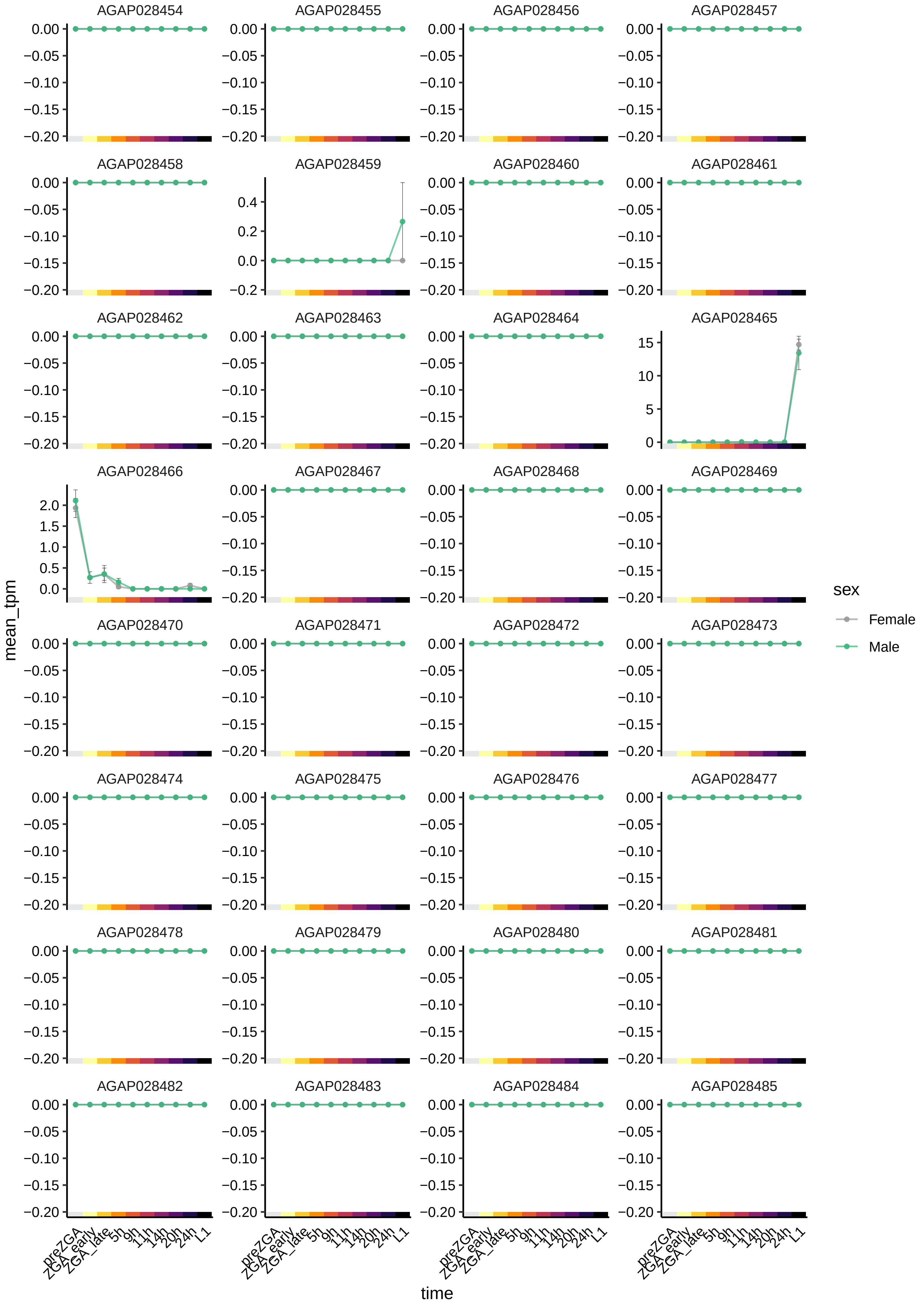

AGAP028454
AGAP028455
AGAP028456
AGAP028457
AGAP028458
AGAP028459
AGAP028460
AGAP028461
AGAP028462
AGAP028463
AGAP028464
AGAP028465
AGAP028466
AGAP028467
AGAP028468
AGAP028469
AGAP028470
AGAP028471
AGAP028472
AGAP028473
AGAP028474
AGAP028475
AGAP028476
AGAP028477
AGAP028478
AGAP028479
AGAP028480
AGAP028481
AGAP028482
AGAP028483
AGAP028484
AGAP028485
mean_tpm
time
preZGA
ZGA_early
ZGA_late
5h
9h
11h
14h
20h
24h
L1
sex
Female
Male

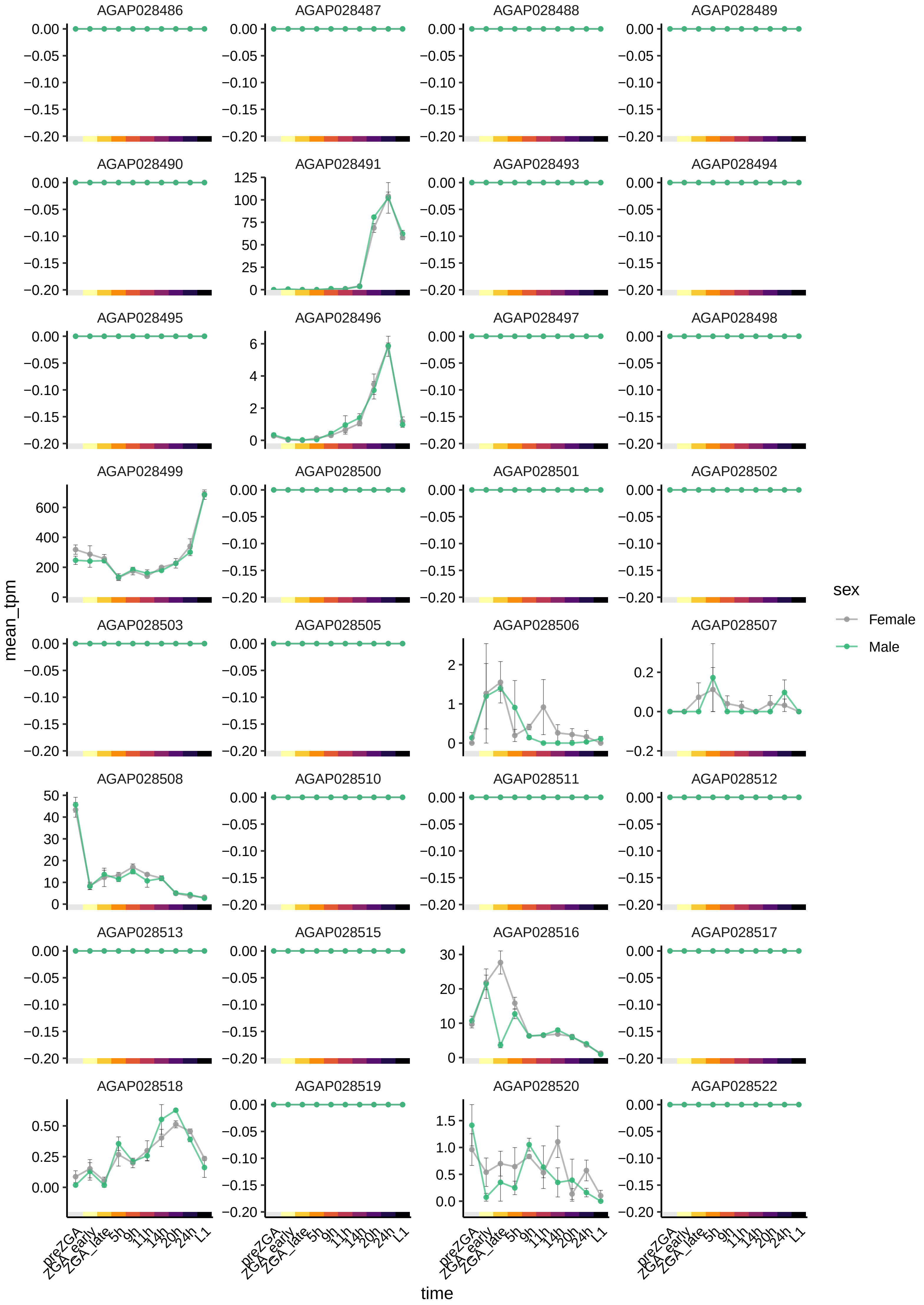

AGAP028486
AGAP028487
AGAP028488
AGAP028489
AGAP028490
AGAP028491
AGAP028493
AGAP028494
AGAP028495
AGAP028496
AGAP028497
AGAP028498
AGAP028499
AGAP028500
AGAP028501
AGAP028502
AGAP028503
AGAP028505
AGAP028506
AGAP028507
AGAP028508
AGAP028510
AGAP028511
AGAP028512
AGAP028513
AGAP028515
AGAP028516
AGAP028517
AGAP028518
AGAP028519
AGAP028520
AGAP028522
mean_tpm
time
preZGA
ZGA_early
ZGA_late
5h
9h
11h
14h
20h
24h
L1
sex
Female
Male

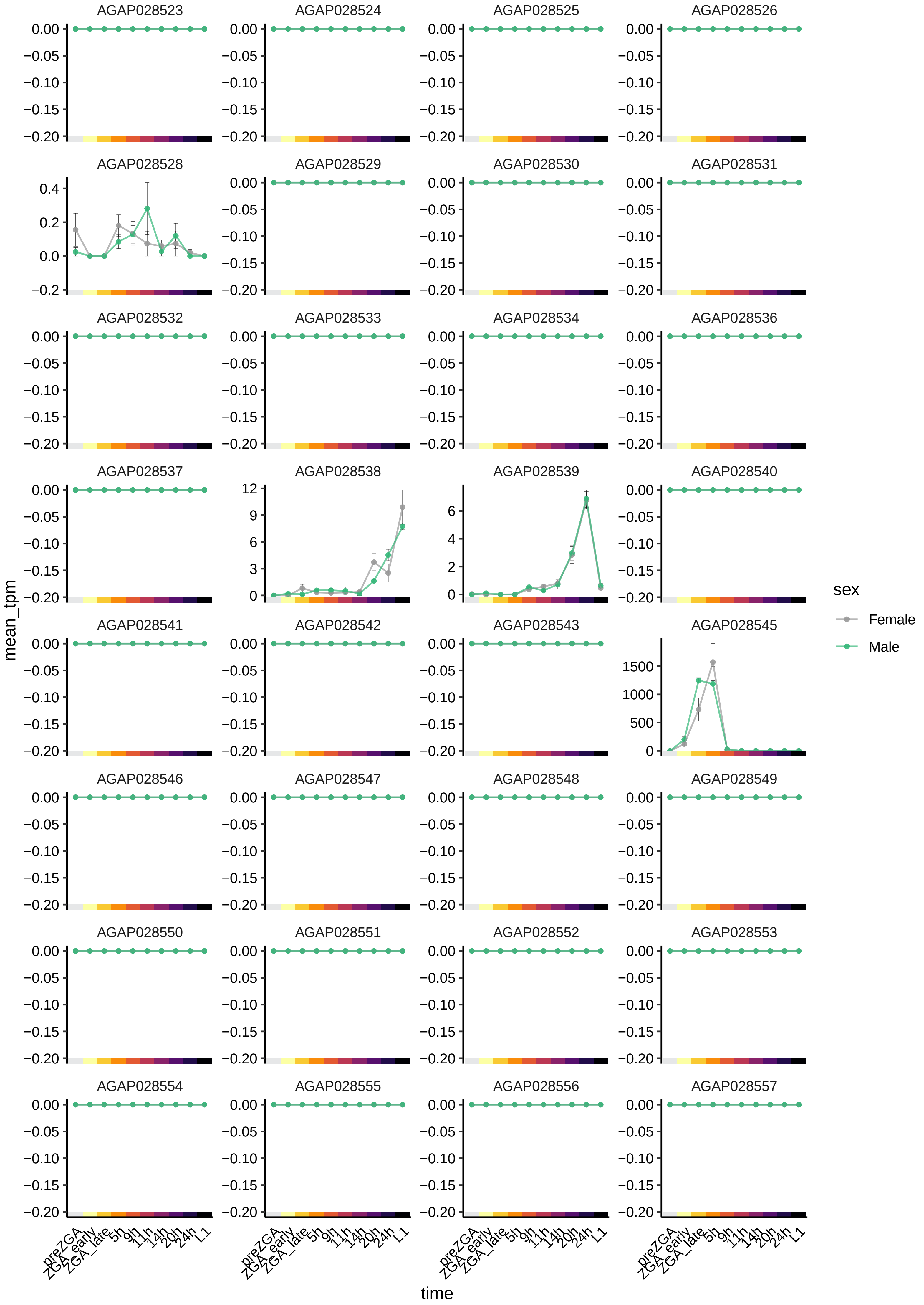

AGAP028523
AGAP028524
AGAP028525
AGAP028526
AGAP028528
AGAP028529
AGAP028530
AGAP028531
AGAP028532
AGAP028533
AGAP028534
AGAP028536
AGAP028537
AGAP028538
AGAP028539
AGAP028540
AGAP028541
AGAP028542
AGAP028543
AGAP028545
AGAP028546
AGAP028547
AGAP028548
AGAP028549
AGAP028550
AGAP028551
AGAP028552
AGAP028553
AGAP028554
AGAP028555
AGAP028556
AGAP028557
mean_tpm
time
preZGA
ZGA_early
ZGA_late
5h
9h
11h
14h
20h
24h
L1
sex
Female
Male

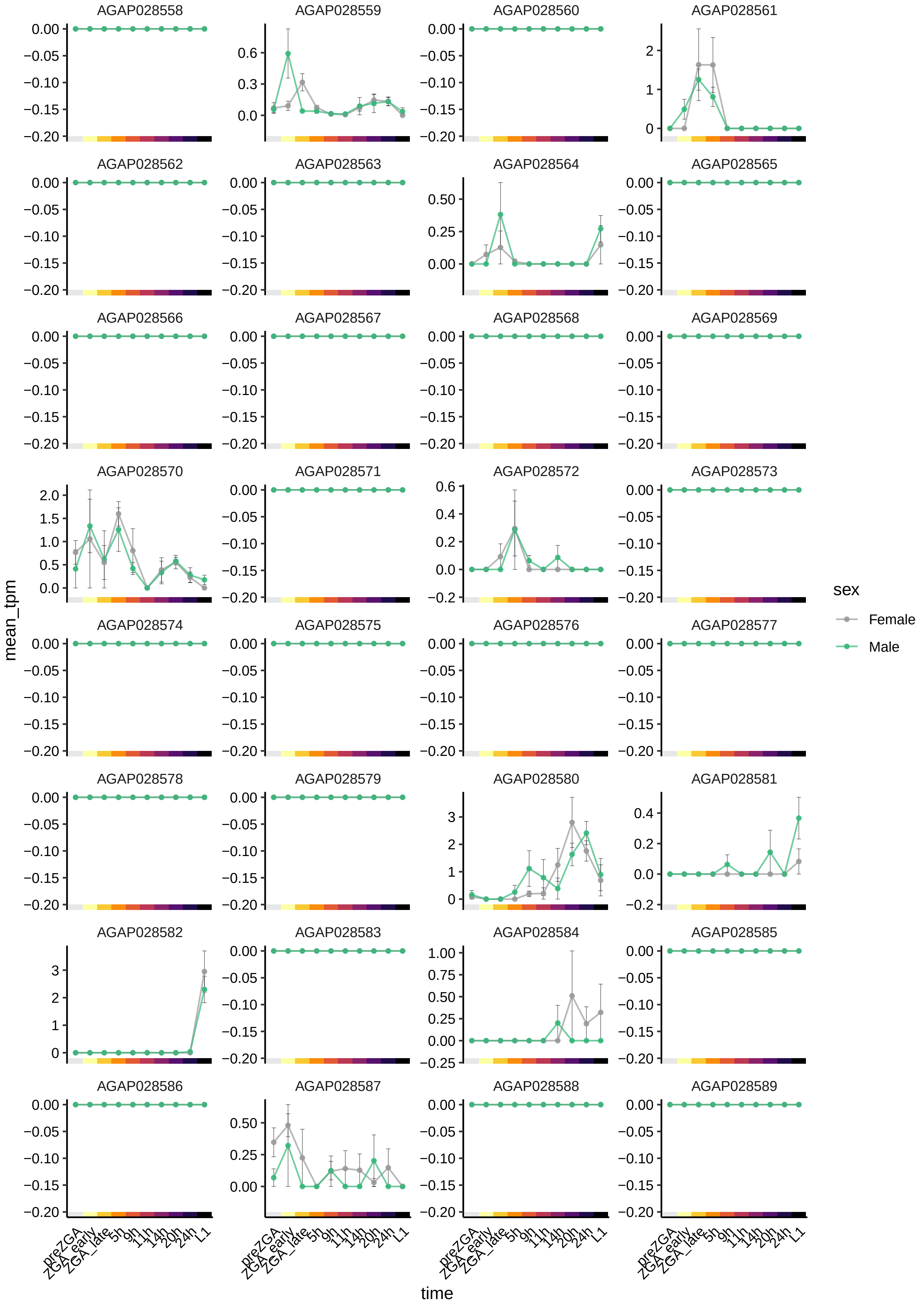

AGAP028558
AGAP028559
AGAP028560
AGAP028561
AGAP028562
AGAP028563
AGAP028564
AGAP028565
AGAP028566
AGAP028567
AGAP028568
AGAP028569
AGAP028570
AGAP028571
AGAP028572
AGAP028573
AGAP028574
AGAP028575
AGAP028576
AGAP028577
AGAP028578
AGAP028579
AGAP028580
AGAP028581
AGAP028582
AGAP028583
AGAP028584
AGAP028585
AGAP028586
AGAP028587
AGAP028588
AGAP028589
mean_tpm
time
preZGA
ZGA_early
ZGA_late
5h
9h
11h
14h
20h
24h
L1
sex
Female
Male

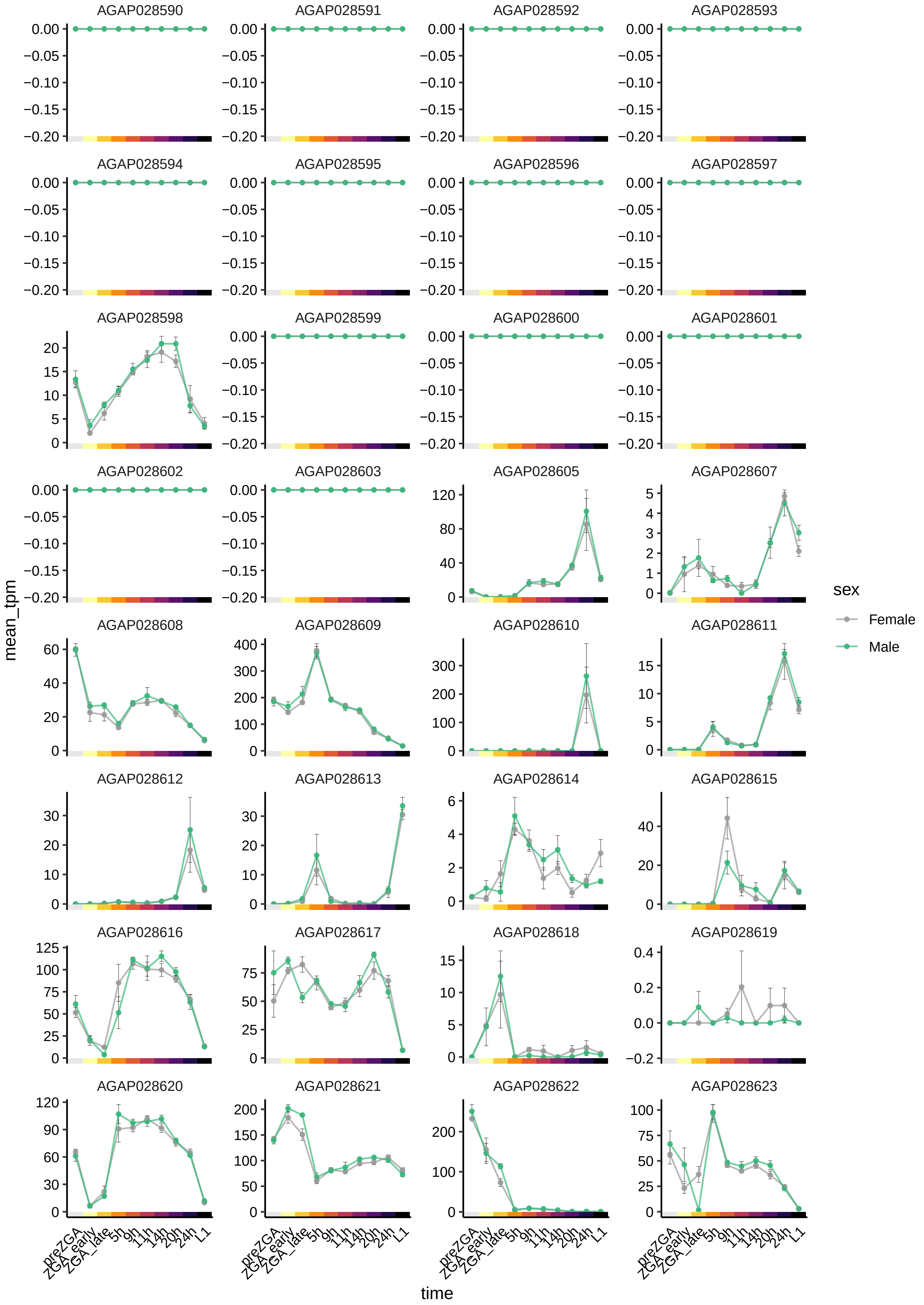
AGAP028590
AGAP028591
AGAP028592
AGAP028593
AGAP028594
AGAP028595
AGAP028596
AGAP028597
AGAP028598
AGAP028599
AGAP028600
AGAP028601
AGAP028602
AGAP028603
AGAP028605
AGAP028607
AGAP028608
AGAP028609
AGAP028610
AGAP028611
AGAP028612
AGAP028613
AGAP028614
AGAP028615
AGAP028616
AGAP028617
AGAP028618
AGAP028619
AGAP028620
AGAP028621
AGAP028622
AGAP028623
mean_tpm
time
preZGA
ZGA_early
ZGA_late
5h
9h
11h
14h
20h
24h
L1
sex
Female
Male

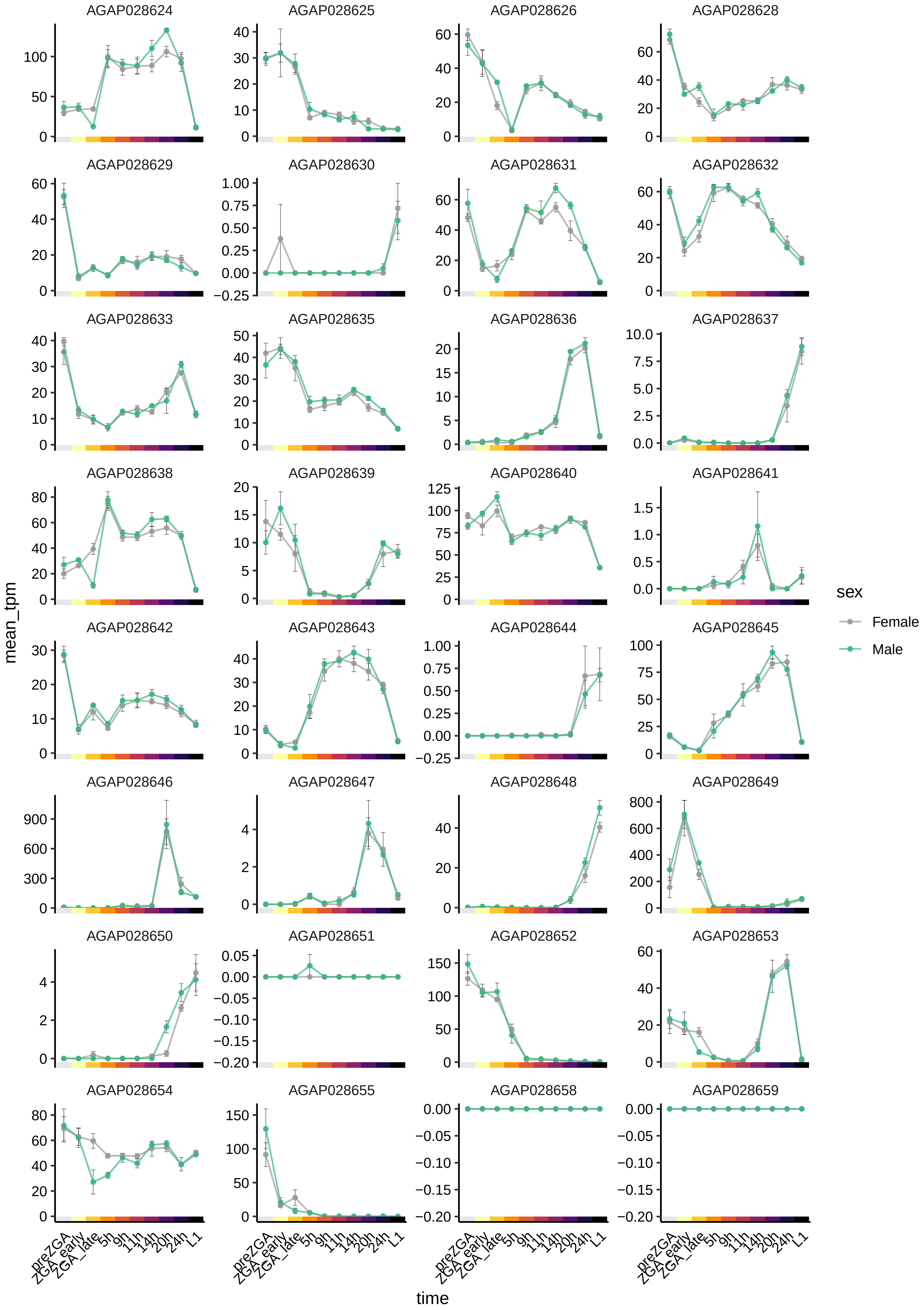

AGAP028624
AGAP028625
AGAP028626
AGAP028628
AGAP028629
AGAP028630
AGAP028631
AGAP028632
AGAP028633
AGAP028635
AGAP028636
AGAP028637
AGAP028638
AGAP028639
AGAP028640
AGAP028641
AGAP028642
AGAP028643
AGAP028644
AGAP028645
AGAP028646
AGAP028647
AGAP028648
AGAP028649
AGAP028650
AGAP028651
AGAP028652
AGAP028653
AGAP028654
AGAP028655
AGAP028658
AGAP028659
mean_tpm
time
preZGA
ZGA_early
ZGA_late
5h
9h
11h
14h
20h
24h
L1
sex
Female
Male

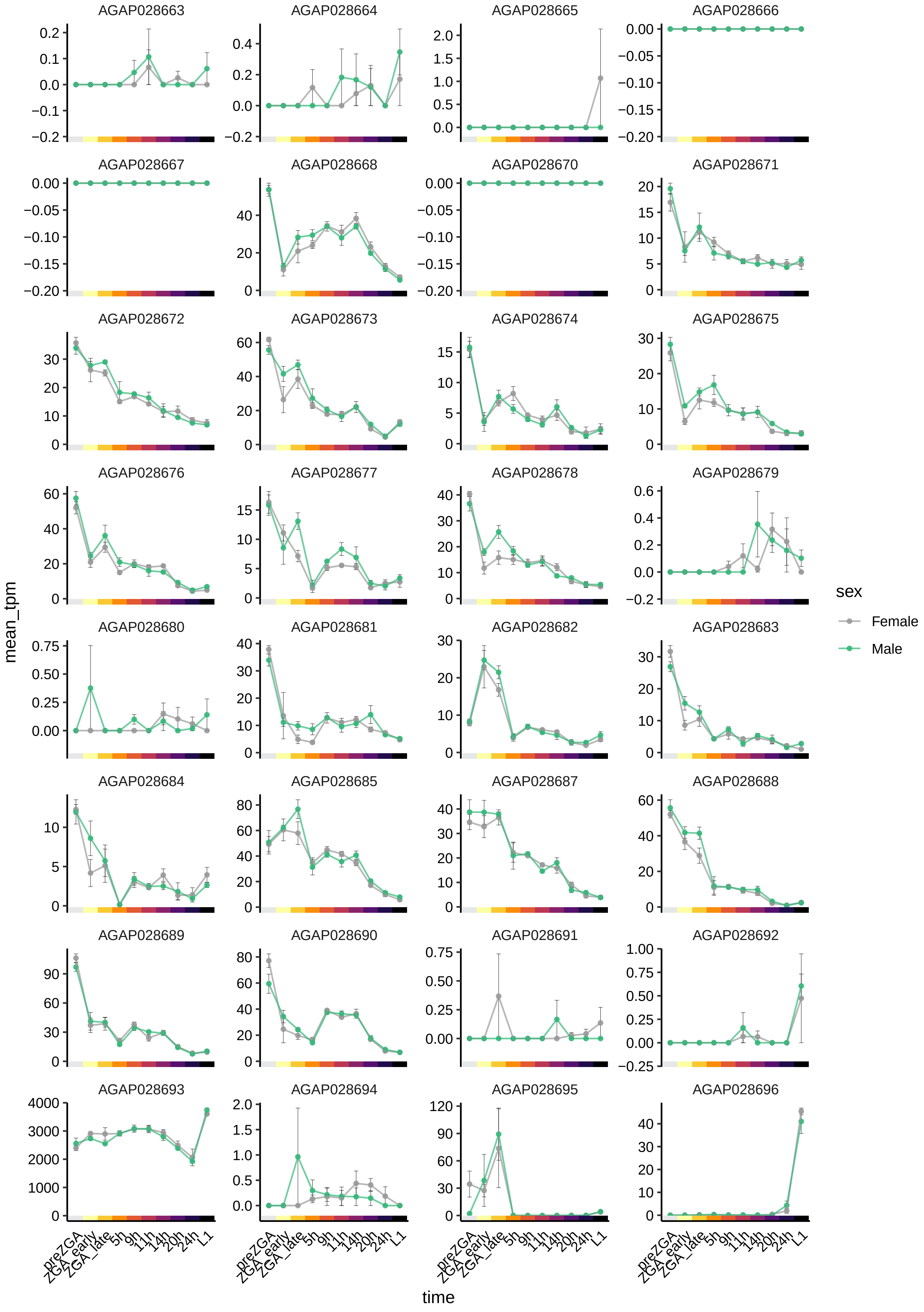
AGAP028663
AGAP028664
AGAP028665
AGAP028666
AGAP028667
AGAP028668
AGAP028670
AGAP028671
AGAP028672
AGAP028673
AGAP028674
AGAP028675
AGAP028676
AGAP028677
AGAP028678
AGAP028679
AGAP028680
AGAP028681
AGAP028682
AGAP028683
AGAP028684
AGAP028685
AGAP028687
AGAP028688
AGAP028689
AGAP028690
AGAP028691
AGAP028692
AGAP028693
AGAP028694
AGAP028695
AGAP028696
mean_tpm
time
preZGA
ZGA_early
ZGA_late
5h
9h
11h
14h
20h
24h
L1
sex
Female
Male

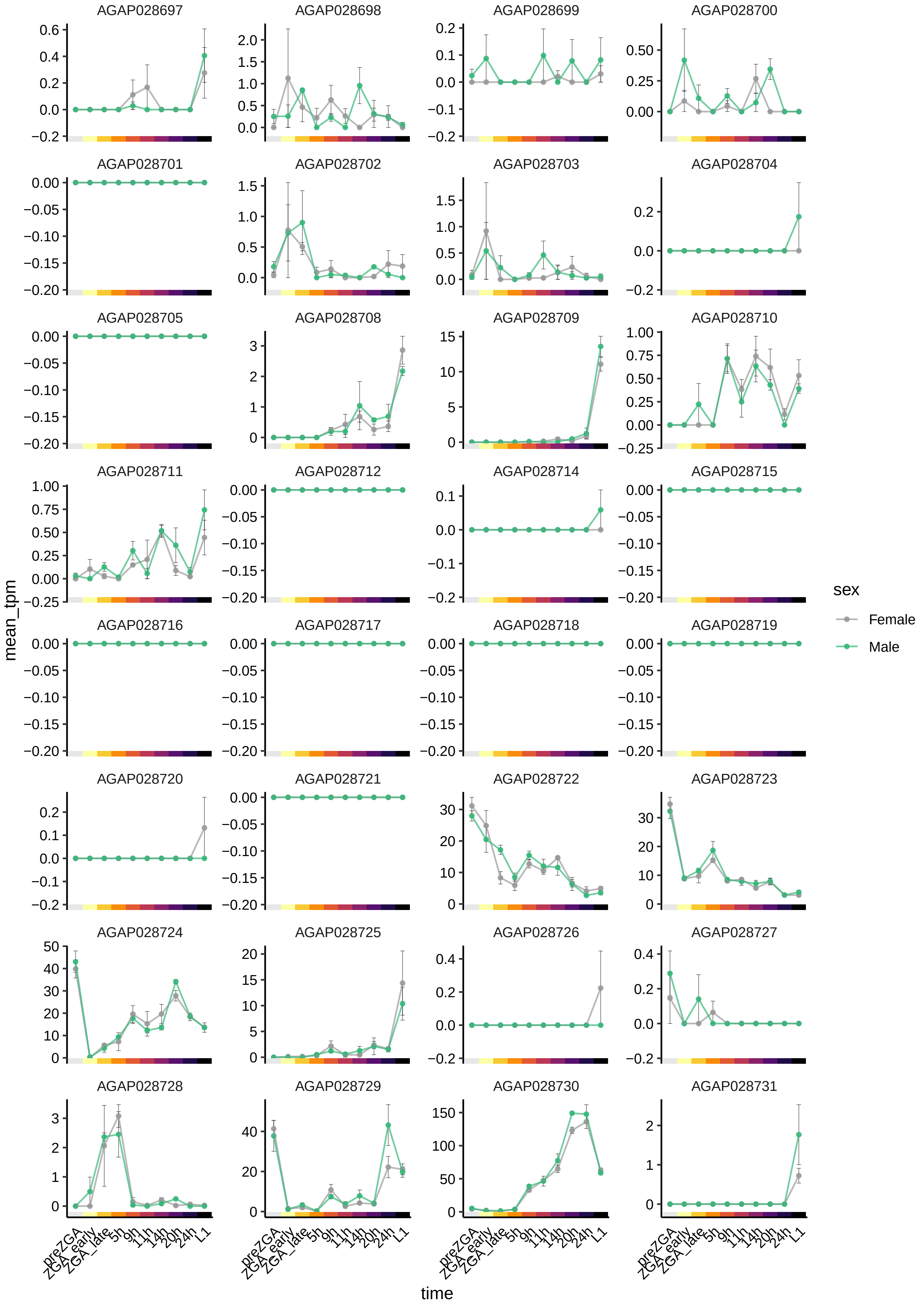

AGAP028697
AGAP028698
AGAP028699
AGAP028700
AGAP028701
AGAP028702
AGAP028703
AGAP028704
AGAP028705
AGAP028708
AGAP028709
AGAP028710
AGAP028711
AGAP028712
AGAP028714
AGAP028715
AGAP028716
AGAP028717
AGAP028718
AGAP028719
AGAP028720
AGAP028721
AGAP028722
AGAP028723
AGAP028724
AGAP028725
AGAP028726
AGAP028727
AGAP028728
AGAP028729
AGAP028730
AGAP028731
mean_tpm
time
preZGA
ZGA_early
ZGA_late
5h
9h
11h
14h
20h
24h
L1
sex
Female
Male

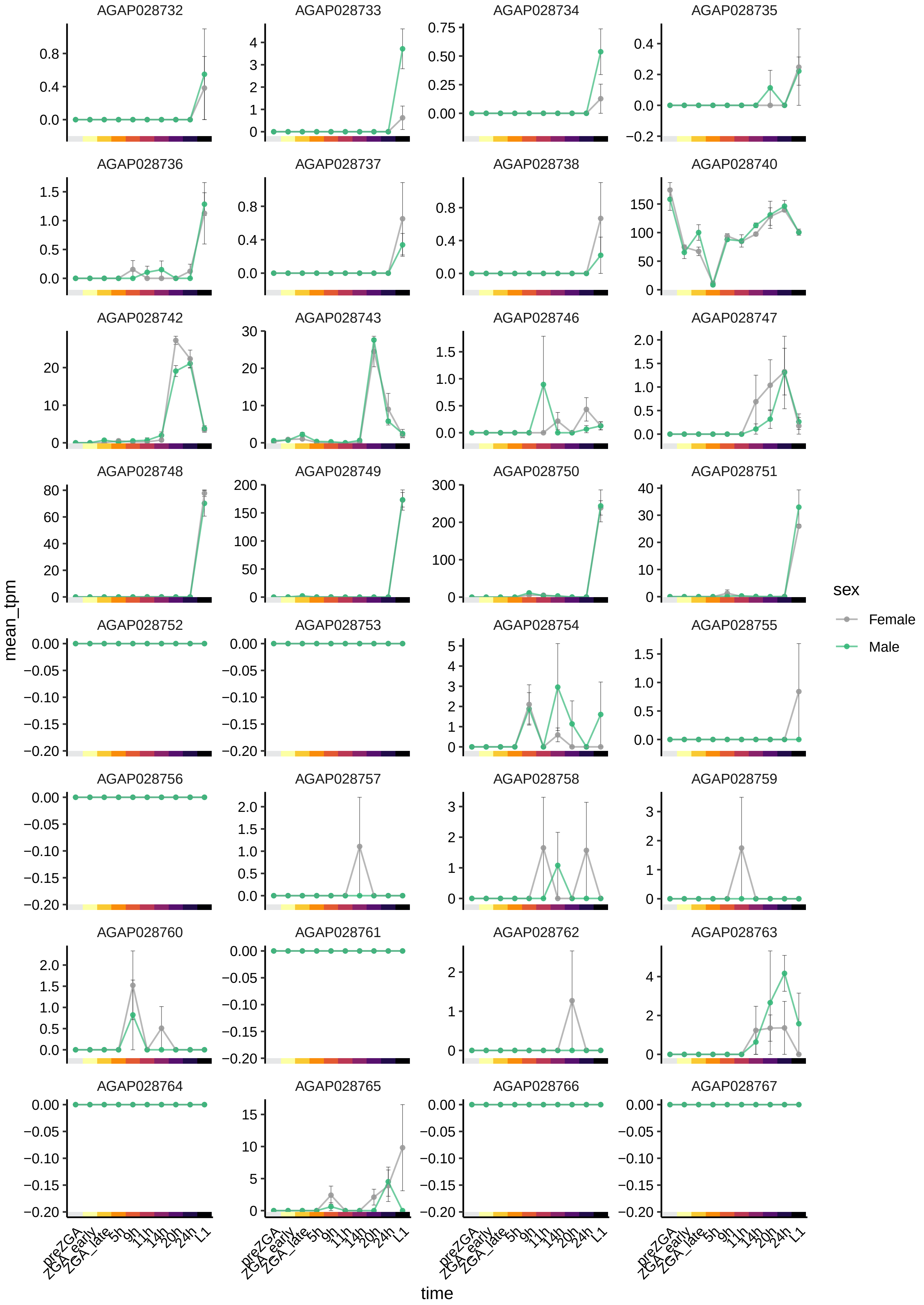

AGAP028732
AGAP028733
AGAP028734
AGAP028735
AGAP028736
AGAP028737
AGAP028738
AGAP028740
AGAP028742
AGAP028743
AGAP028746
AGAP028747
AGAP028748
AGAP028749
AGAP028750
AGAP028751
AGAP028752
AGAP028753
AGAP028754
AGAP028755
AGAP028756
AGAP028757
AGAP028758
AGAP028759
AGAP028760
AGAP028761
AGAP028762
AGAP028763
AGAP028764
AGAP028765
AGAP028766
AGAP028767
mean_tpm
time
preZGA
ZGA_early
ZGA_late
5h
9h
11h
14h
20h
24h
L1
sex
Female
Male

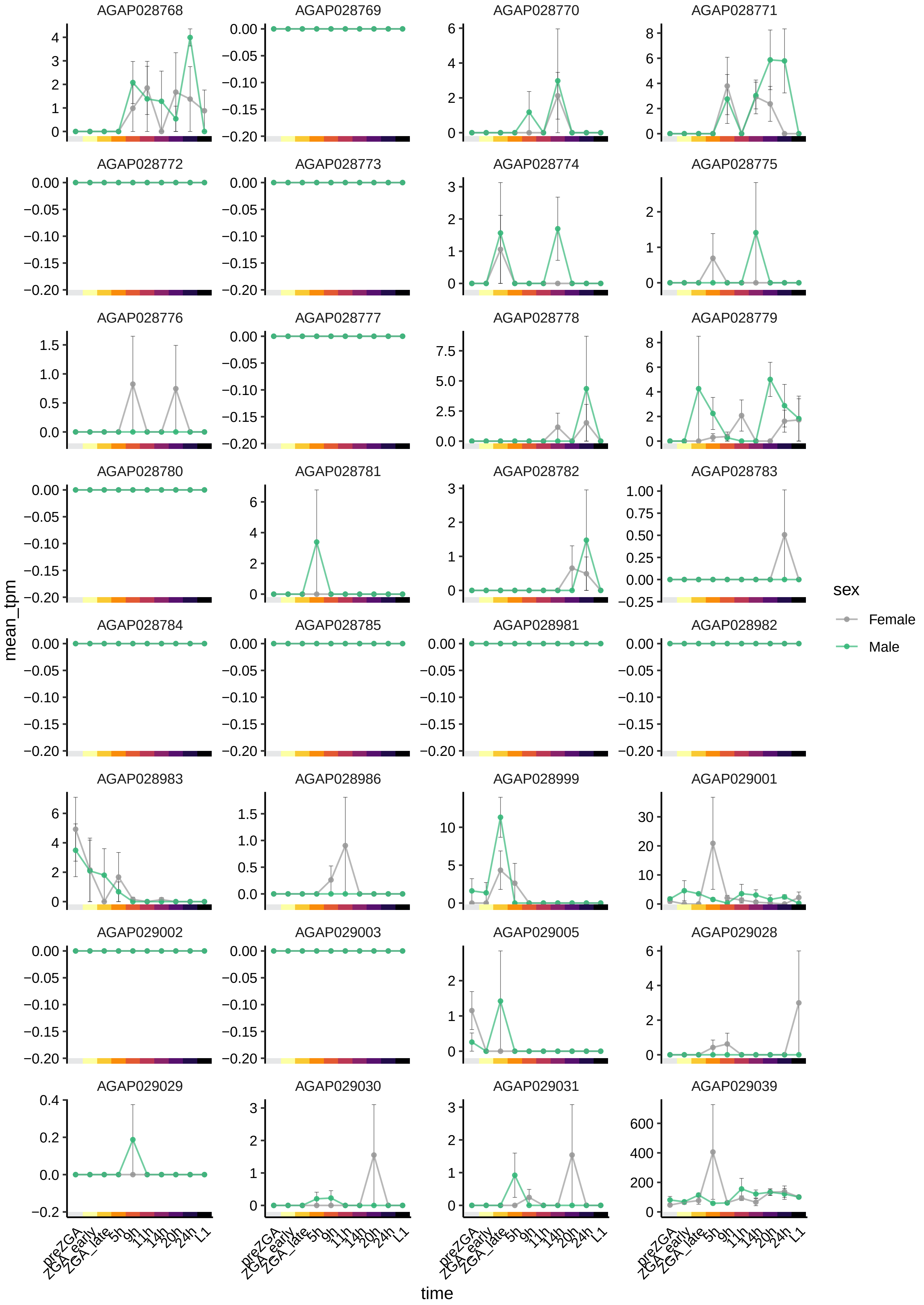

AGAP028768
AGAP028769
AGAP028770
AGAP028771
AGAP028772
AGAP028773
AGAP028774
AGAP028775
AGAP028776
AGAP028777
AGAP028778
AGAP028779
AGAP028780
AGAP028781
AGAP028782
AGAP028783
AGAP028784
AGAP028785
AGAP028981
AGAP028982
AGAP028983
AGAP028986
AGAP028999
AGAP029001
AGAP029002
AGAP029003
AGAP029005
AGAP029028
AGAP029029
AGAP029030
AGAP029031
AGAP029039
mean_tpm
time
preZGA
ZGA_early
ZGA_late
5h
9h
11h
14h
20h
24h
L1
sex
Female
Male

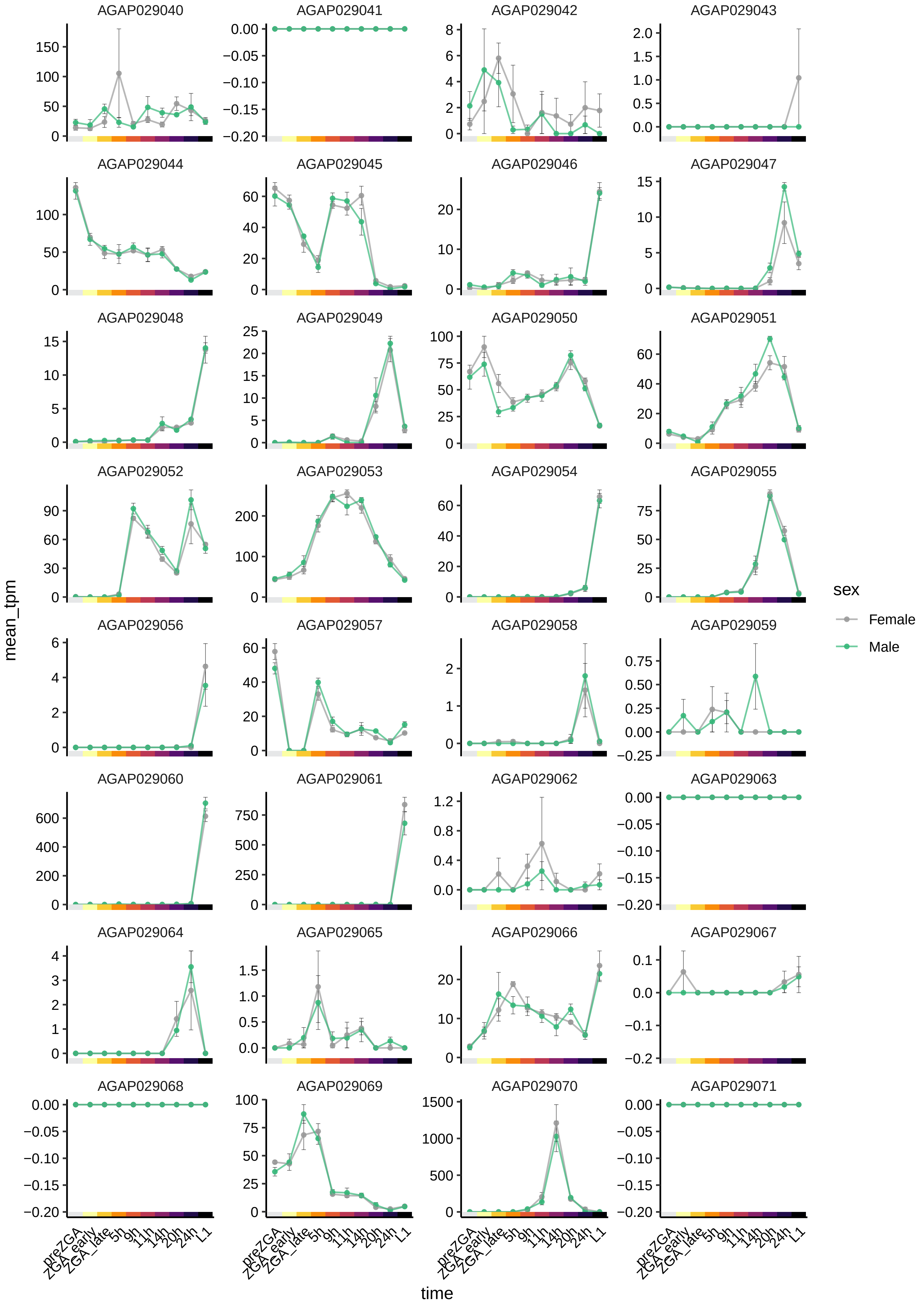

AGAP029040
AGAP029041
AGAP029042
AGAP029043
AGAP029044
AGAP029045
AGAP029046
AGAP029047
AGAP029048
AGAP029049
AGAP029050
AGAP029051
AGAP029052
AGAP029053
AGAP029054
AGAP029055
AGAP029056
AGAP029057
AGAP029058
AGAP029059
AGAP029060
AGAP029061
AGAP029062
AGAP029063
AGAP029064
AGAP029065
AGAP029066
AGAP029067
AGAP029068
AGAP029069
AGAP029070
AGAP029071
mean_tpm
time
preZGA
ZGA_early
ZGA_late
5h
9h
11h
14h
20h
24h
L1
sex
Female
Male

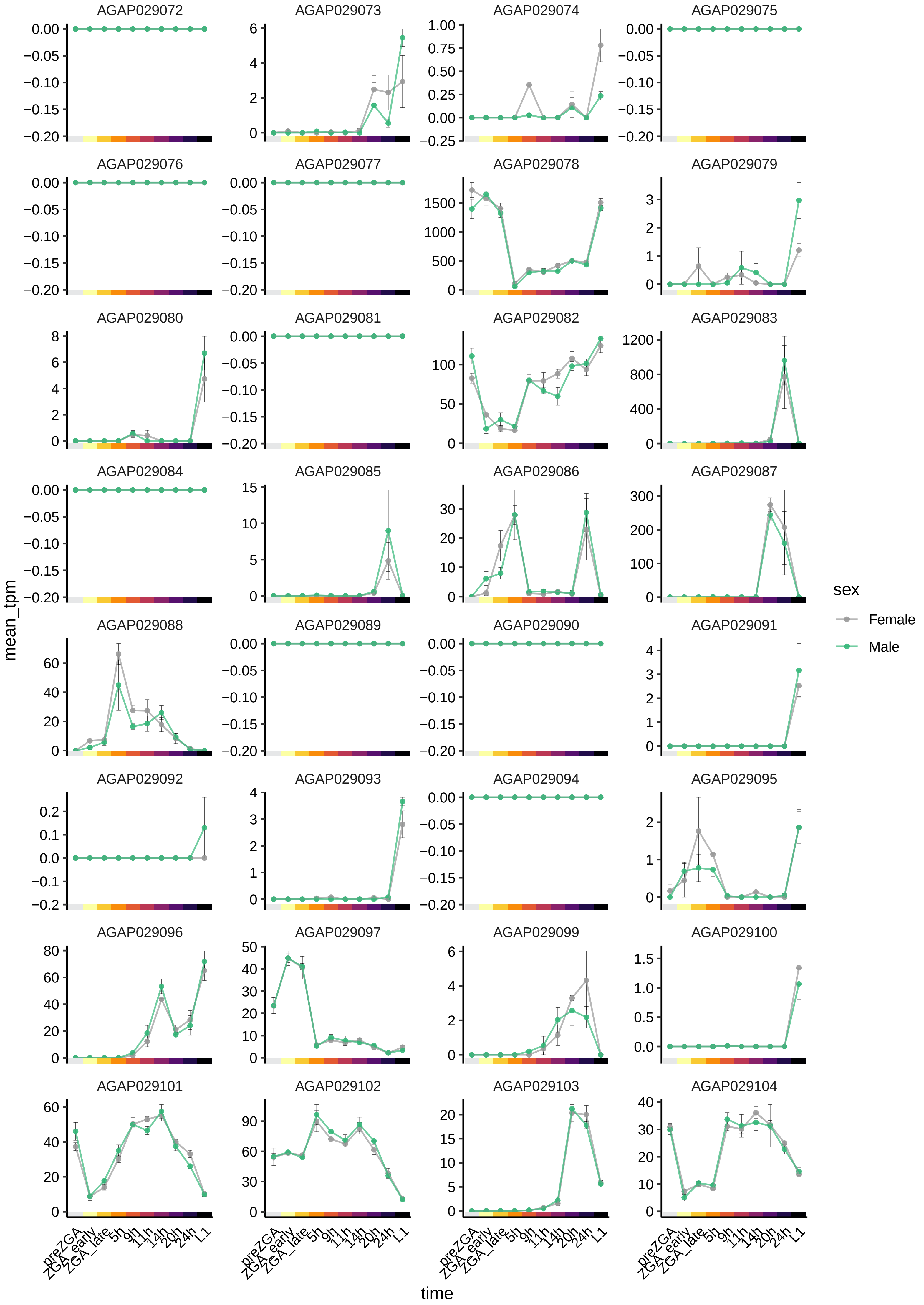

AGAP029072
AGAP029073
AGAP029074
AGAP029075
AGAP029076
AGAP029077
AGAP029078
AGAP029079
AGAP029080
AGAP029081
AGAP029082
AGAP029083
AGAP029084
AGAP029085
AGAP029086
AGAP029087
AGAP029088
AGAP029089
AGAP029090
AGAP029091
AGAP029092
AGAP029093
AGAP029094
AGAP029095
AGAP029096
AGAP029097
AGAP029099
AGAP029100
AGAP029101
AGAP029102
AGAP029103
AGAP029104
mean_tpm
time
preZGA
ZGA_early
ZGA_late
5h
9h
11h
14h
20h
24h
L1
sex
Female
Male

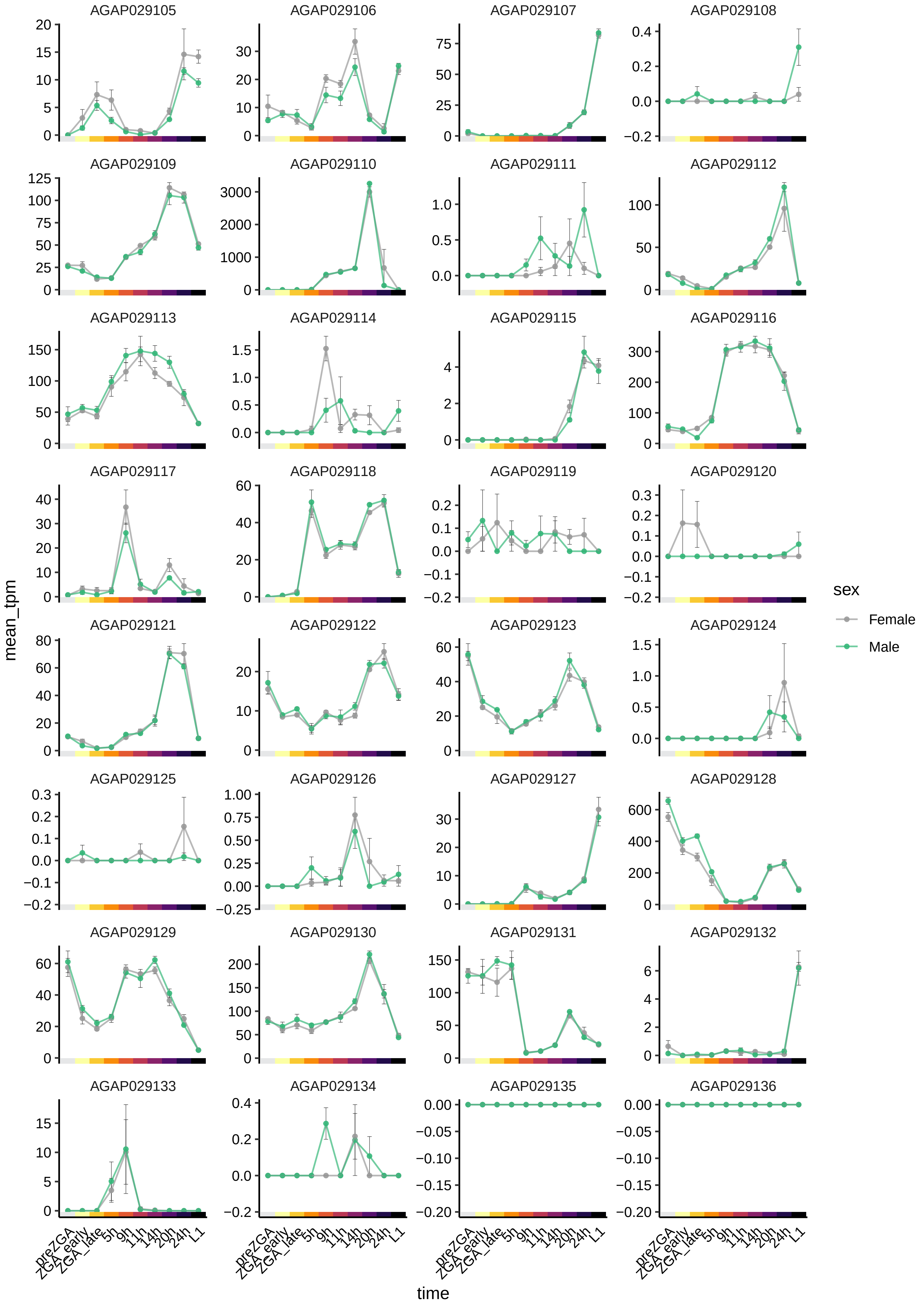
AGAP029105
AGAP029106
AGAP029107
AGAP029108
AGAP029109
AGAP029110
AGAP029111
AGAP029112
AGAP029113
AGAP029114
AGAP029115
AGAP029116
AGAP029117
AGAP029118
AGAP029119
AGAP029120
AGAP029121
AGAP029122
AGAP029123
AGAP029124
AGAP029125
AGAP029126
AGAP029127
AGAP029128
AGAP029129
AGAP029130
AGAP029131
AGAP029132
AGAP029133
AGAP029134
AGAP029135
AGAP029136
mean_tpm
time
preZGA
ZGA_early
ZGA_late
5h
9h
11h
14h
20h
24h
L1
sex
Female
Male

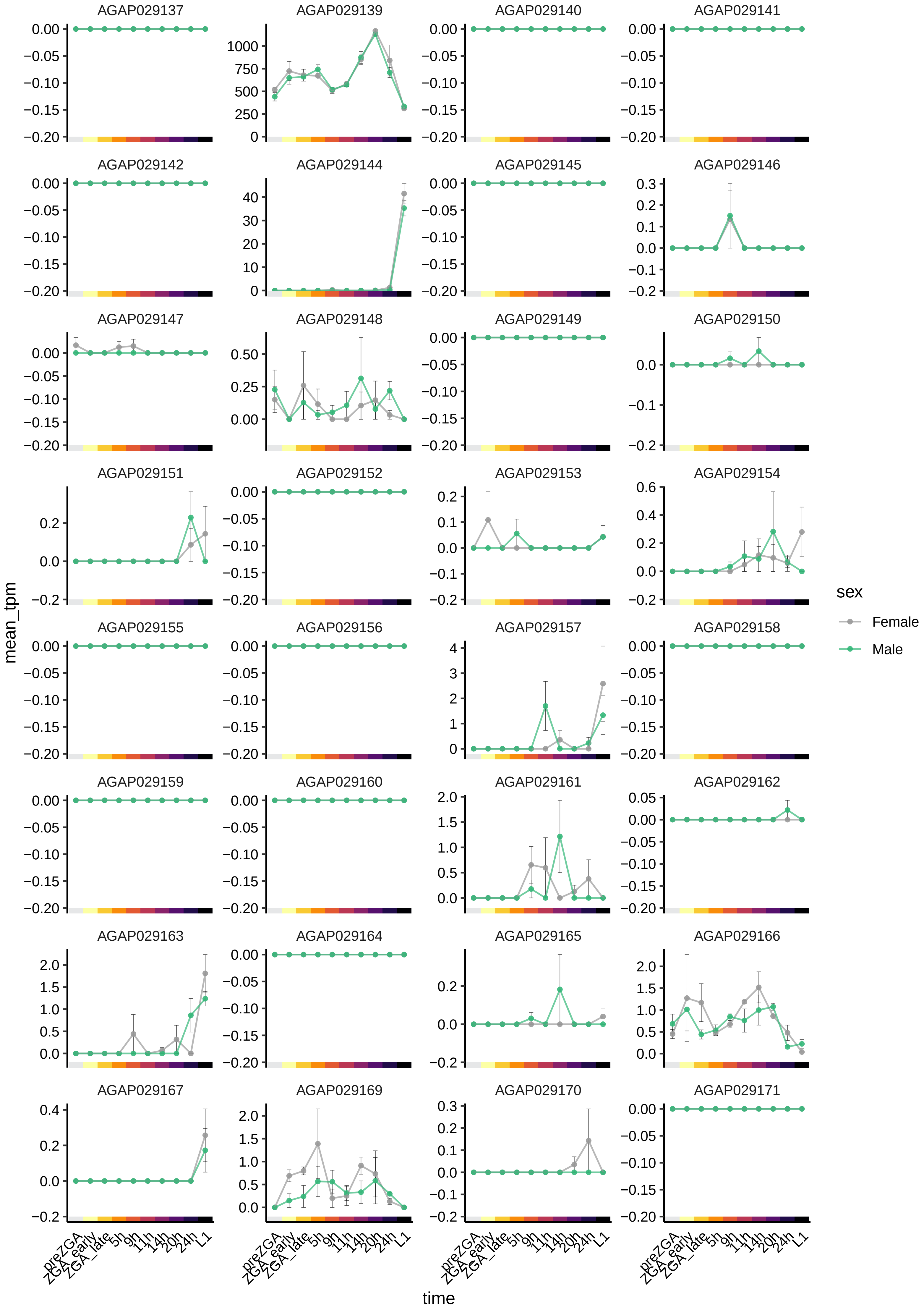

AGAP029137
AGAP029139
AGAP029140
AGAP029141
AGAP029142
AGAP029144
AGAP029145
AGAP029146
AGAP029147
AGAP029148
AGAP029149
AGAP029150
AGAP029151
AGAP029152
AGAP029153
AGAP029154
AGAP029155
AGAP029156
AGAP029157
AGAP029158
AGAP029159
AGAP029160
AGAP029161
AGAP029162
AGAP029163
AGAP029164
AGAP029165
AGAP029166
AGAP029167
AGAP029169
AGAP029170
AGAP029171
mean_tpm
time
preZGA
ZGA_early
ZGA_late
5h
9h
11h
14h
20h
24h
L1
sex
Female
Male

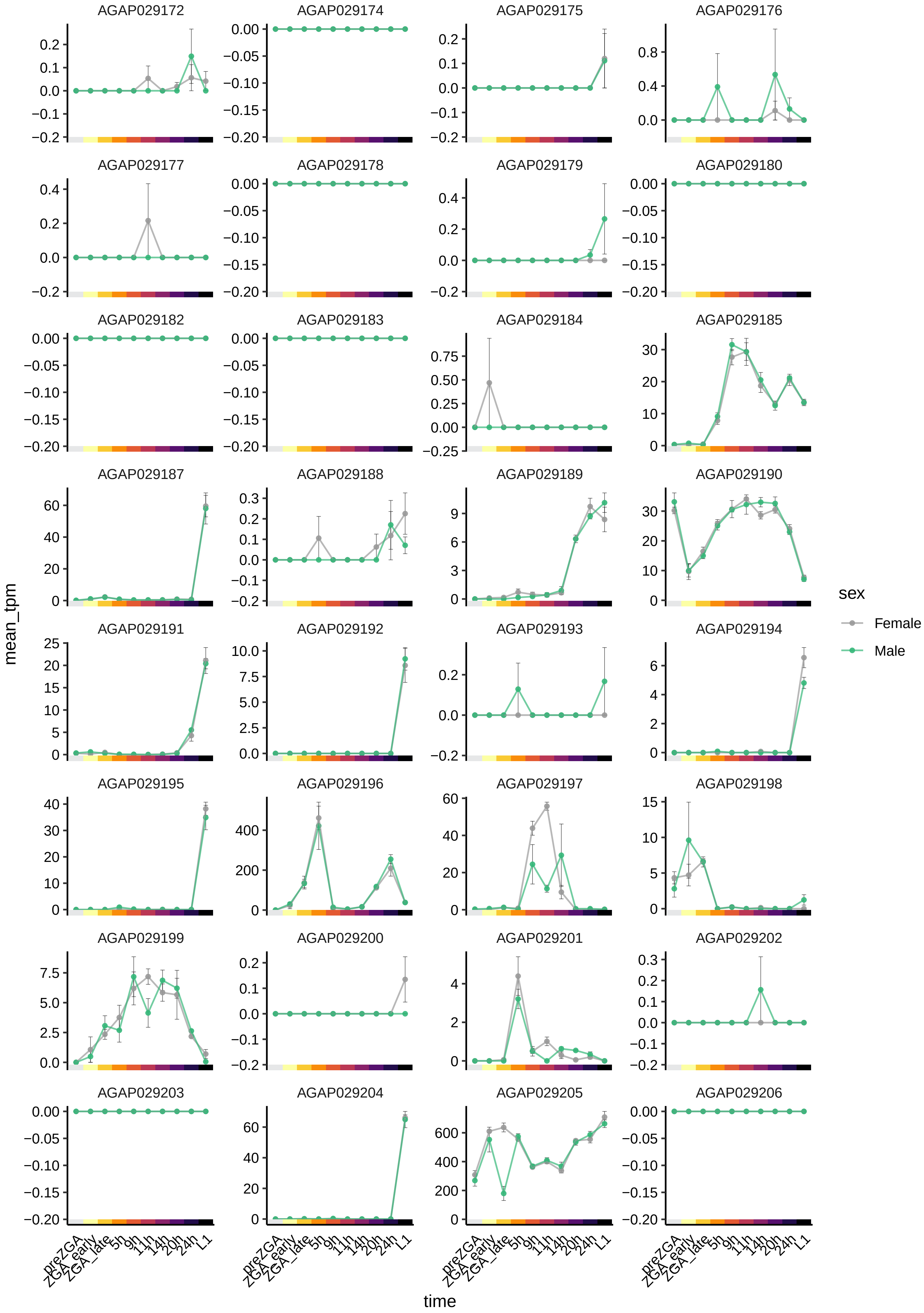

AGAP029172
AGAP029174
AGAP029175
AGAP029176
AGAP029177
AGAP029178
AGAP029179
AGAP029180
AGAP029182
AGAP029183
AGAP029184
AGAP029185
AGAP029187
AGAP029188
AGAP029189
AGAP029190
AGAP029191
AGAP029192
AGAP029193
AGAP029194
AGAP029195
AGAP029196
AGAP029197
AGAP029198
AGAP029199
AGAP029200
AGAP029201
AGAP029202
AGAP029203
AGAP029204
AGAP029205
AGAP029206
mean_tpm
time
preZGA
ZGA_early
ZGA_late
5h
9h
11h
14h
20h
24h
L1
sex
Female
Male

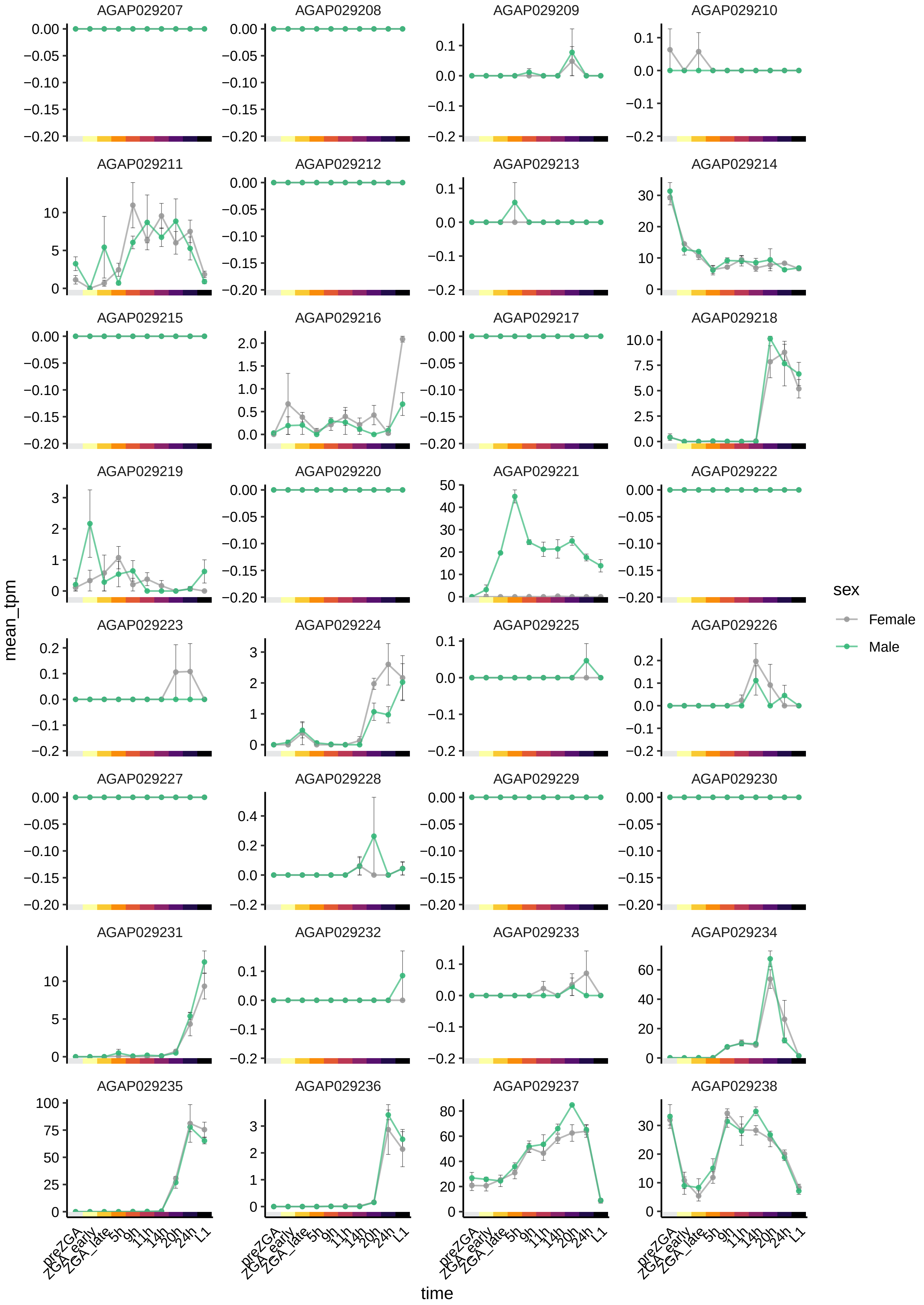
AGAP029207
AGAP029208
AGAP029209
AGAP029210
AGAP029211
AGAP029212
AGAP029213
AGAP029214
AGAP029215
AGAP029216
AGAP029217
AGAP029218
AGAP029219
AGAP029220
AGAP029221
AGAP029222
AGAP029223
AGAP029224
AGAP029225
AGAP029226
AGAP029227
AGAP029228
AGAP029229
AGAP029230
AGAP029231
AGAP029232
AGAP029233
AGAP029234
AGAP029235
AGAP029236
AGAP029237
AGAP029238
mean_tpm
time
preZGA
ZGA_early
ZGA_late
5h
9h
11h
14h
20h
24h
L1
sex
Female
Male

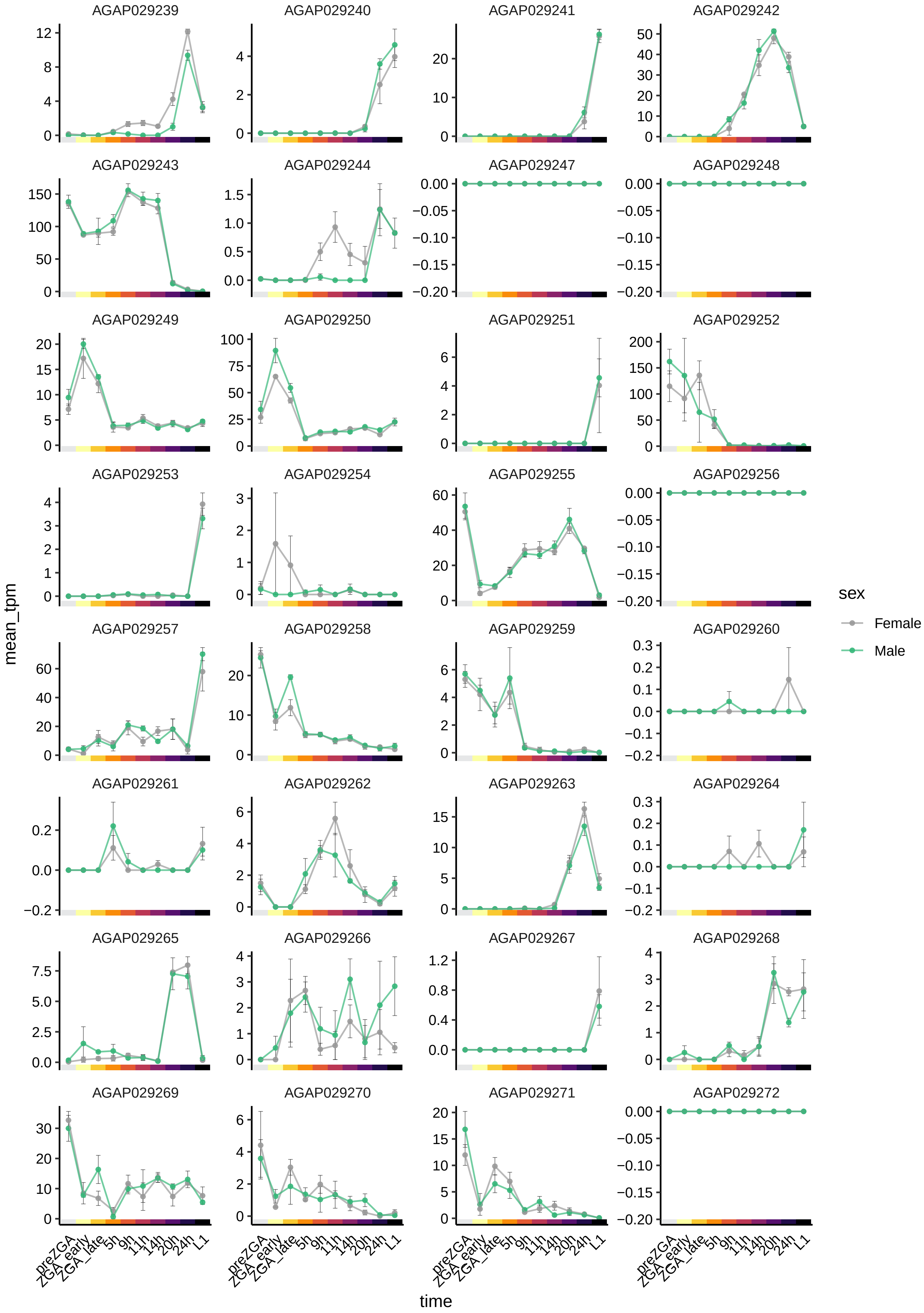

AGAP029239
AGAP029240
AGAP029241
AGAP029242
AGAP029243
AGAP029244
AGAP029247
AGAP029248
AGAP029249
AGAP029250
AGAP029251
AGAP029252
AGAP029253
AGAP029254
AGAP029255
AGAP029256
AGAP029257
AGAP029258
AGAP029259
AGAP029260
AGAP029261
AGAP029262
AGAP029263
AGAP029264
AGAP029265
AGAP029266
AGAP029267
AGAP029268
AGAP029269
AGAP029270
AGAP029271
AGAP029272
mean_tpm
time
preZGA
ZGA_early
ZGA_late
5h
9h
11h
14h
20h
24h
L1
sex
Female
Male

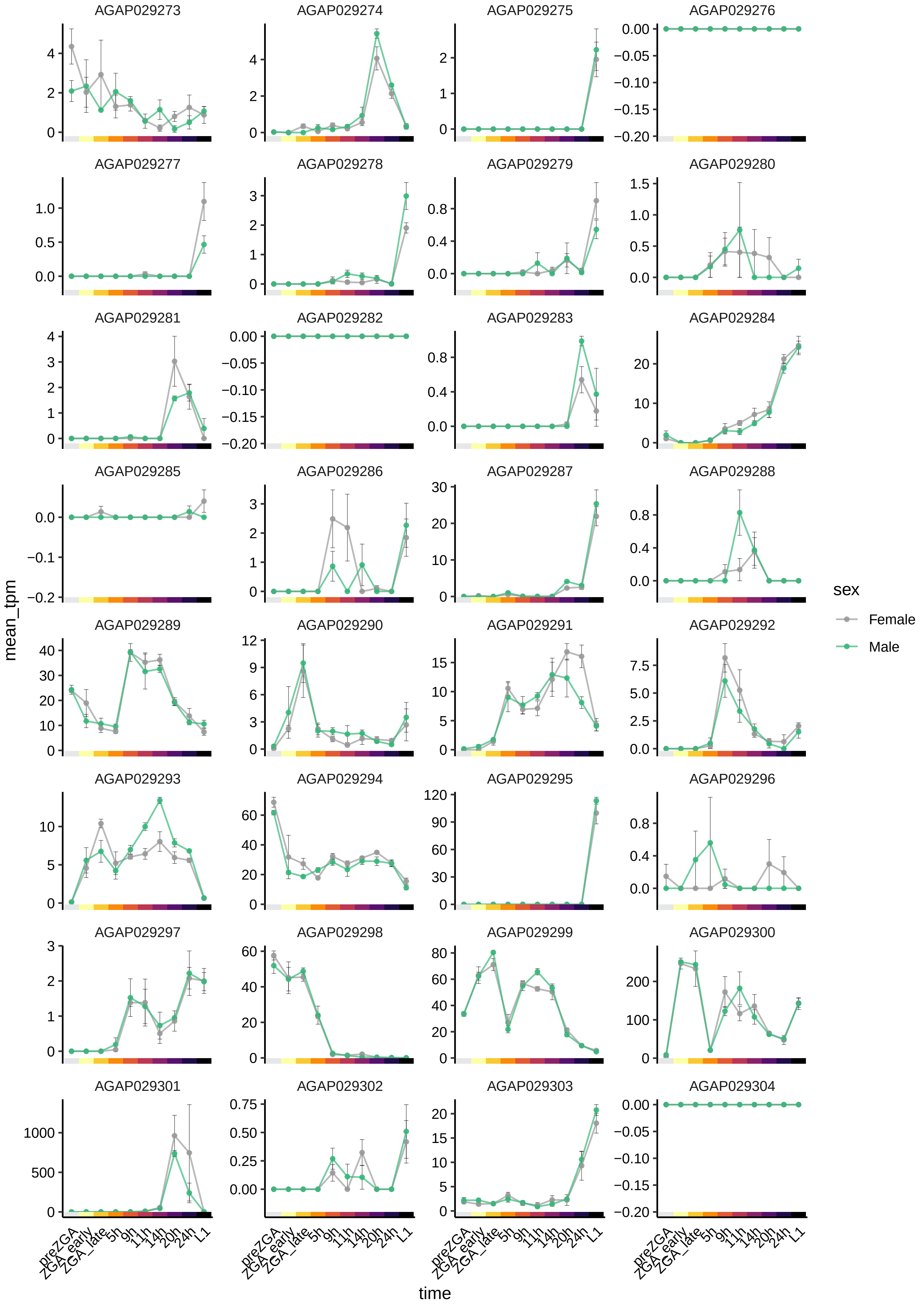

AGAP029273
AGAP029274
AGAP029275
AGAP029276
AGAP029277
AGAP029278
AGAP029279
AGAP029280
AGAP029281
AGAP029282
AGAP029283
AGAP029284
AGAP029285
AGAP029286
AGAP029287
AGAP029288
AGAP029289
AGAP029290
AGAP029291
AGAP029292
AGAP029293
AGAP029294
AGAP029295
AGAP029296
AGAP029297
AGAP029298
AGAP029299
AGAP029300
AGAP029301
AGAP029302
AGAP029303
AGAP029304
mean_tpm
time
preZGA
ZGA_early
ZGA_late
5h
9h
11h
14h
20h
24h
L1
sex
Female
Male

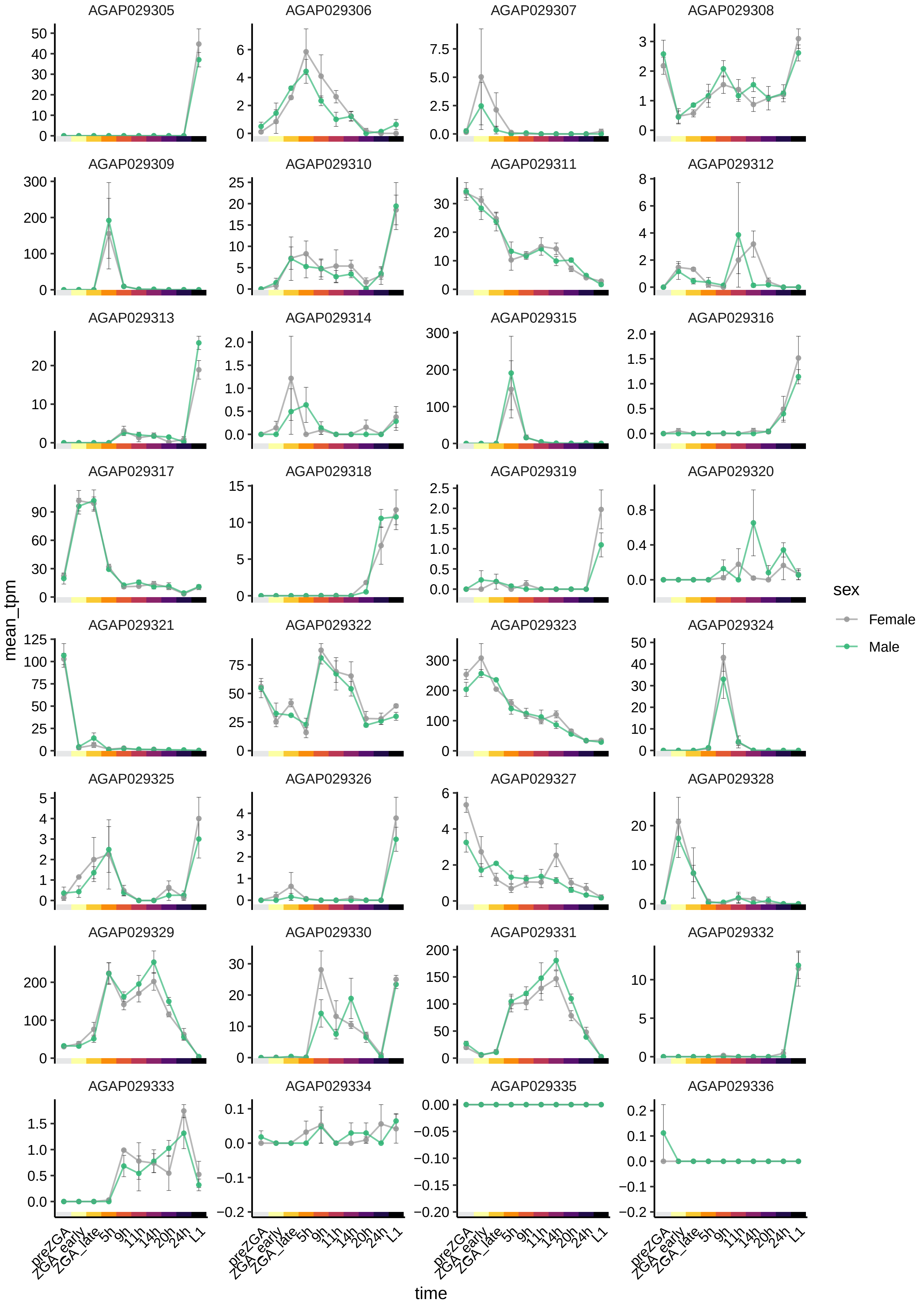
AGAP029305
AGAP029306
AGAP029307
AGAP029308
AGAP029309
AGAP029310
AGAP029311
AGAP029312
AGAP029313
AGAP029314
AGAP029315
AGAP029316
AGAP029317
AGAP029318
AGAP029319
AGAP029320
AGAP029321
AGAP029322
AGAP029323
AGAP029324
AGAP029325
AGAP029326
AGAP029327
AGAP029328
AGAP029329
AGAP029330
AGAP029331
AGAP029332
AGAP029333
AGAP029334
AGAP029335
AGAP029336
mean_tpm
time
preZGA
ZGA_early
ZGA_late
5h
9h
11h
14h
20h
24h
L1
sex
Female
Male

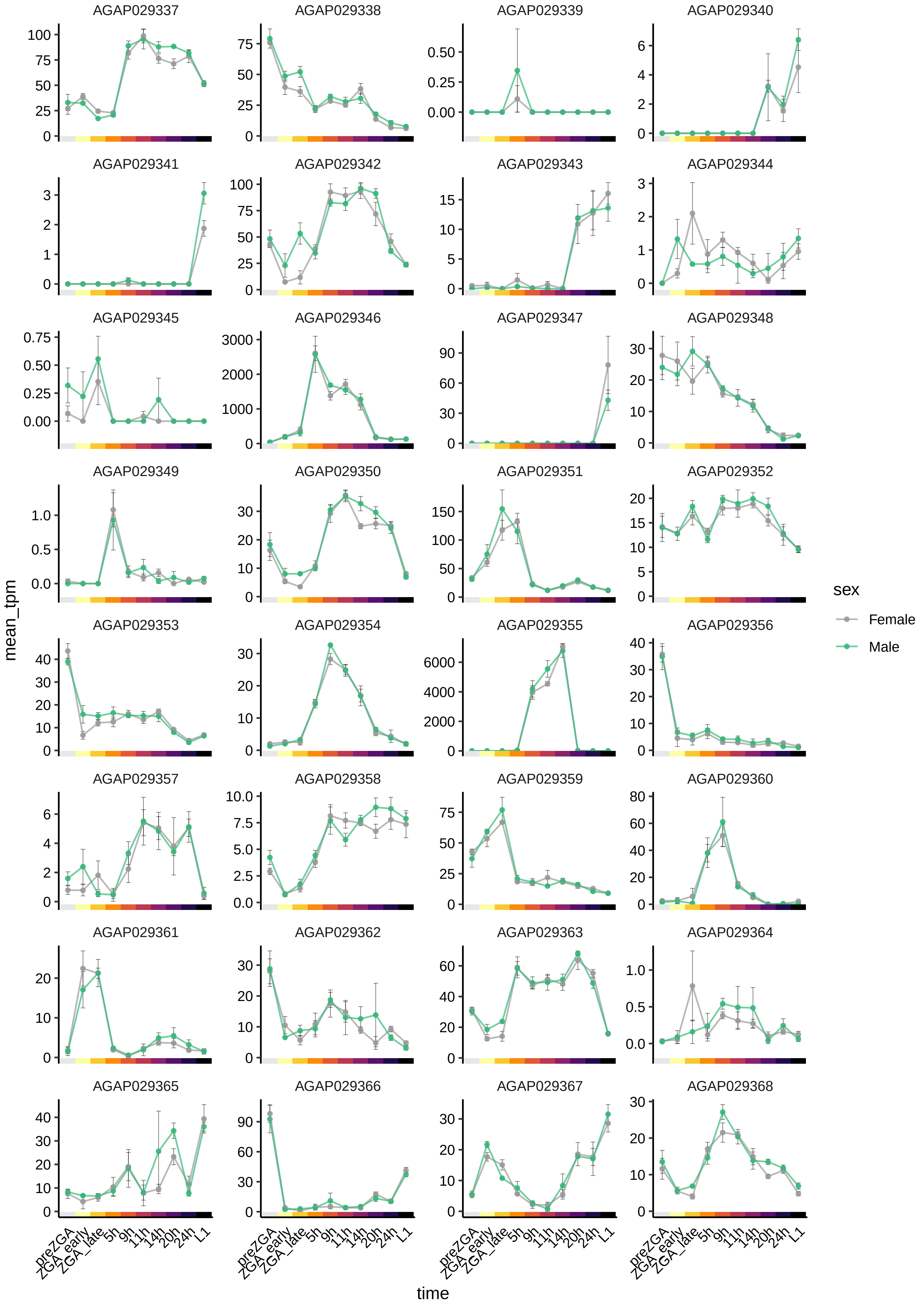

AGAP029337
AGAP029338
AGAP029339
AGAP029340
AGAP029341
AGAP029342
AGAP029343
AGAP029344
AGAP029345
AGAP029346
AGAP029347
AGAP029348
AGAP029349
AGAP029350
AGAP029351
AGAP029352
AGAP029353
AGAP029354
AGAP029355
AGAP029356
AGAP029357
AGAP029358
AGAP029359
AGAP029360
AGAP029361
AGAP029362
AGAP029363
AGAP029364
AGAP029365
AGAP029366
AGAP029367
AGAP029368
mean_tpm
time
preZGA
ZGA_early
ZGA_late
5h
9h
11h
14h
20h
24h
L1
sex
Female
Male

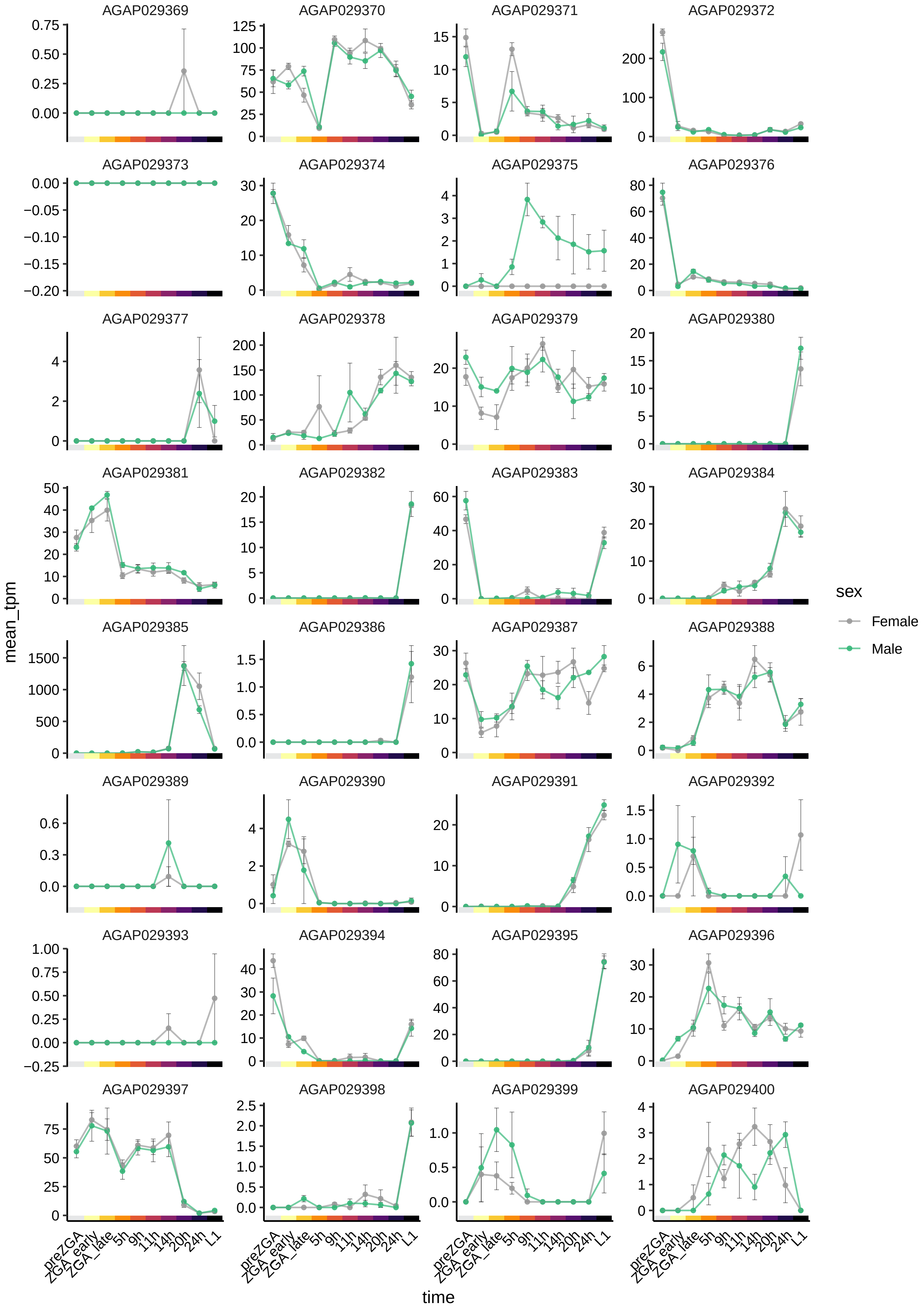
AGAP029369
AGAP029370
AGAP029371
AGAP029372
AGAP029373
AGAP029374
AGAP029375
AGAP029376
AGAP029377
AGAP029378
AGAP029379
AGAP029380
AGAP029381
AGAP029382
AGAP029383
AGAP029384
AGAP029385
AGAP029386
AGAP029387
AGAP029388
AGAP029389
AGAP029390
AGAP029391
AGAP029392
AGAP029393
AGAP029394
AGAP029395
AGAP029396
AGAP029397
AGAP029398
AGAP029399
AGAP029400
mean_tpm
time
preZGA
ZGA_early
ZGA_late
5h
9h
11h
14h
20h
24h
L1
sex
Female
Male

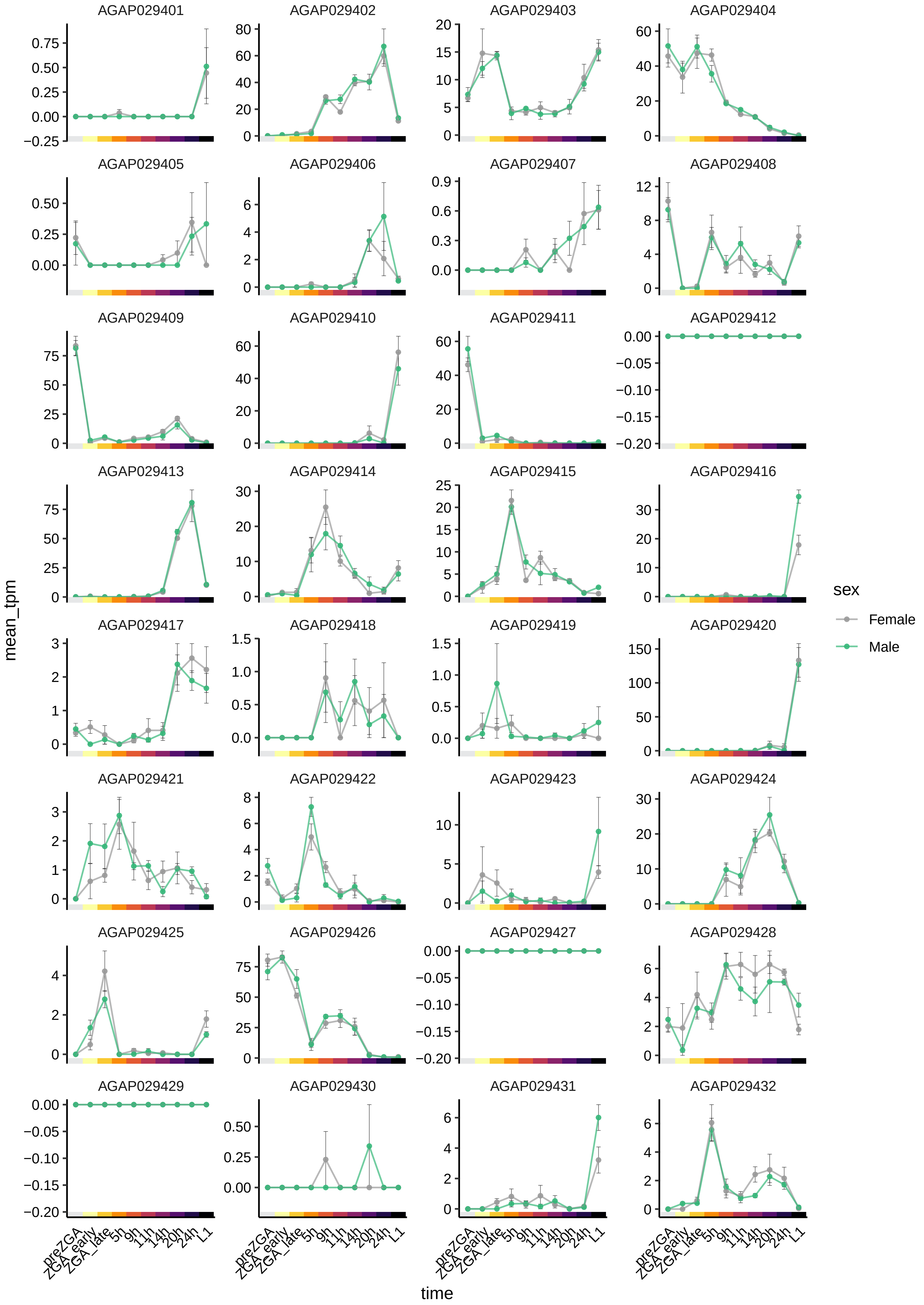
AGAP029401
AGAP029402
AGAP029403
AGAP029404
AGAP029405
AGAP029406
AGAP029407
AGAP029408
AGAP029409
AGAP029410
AGAP029411
AGAP029412
AGAP029413
AGAP029414
AGAP029415
AGAP029416
AGAP029417
AGAP029418
AGAP029419
AGAP029420
AGAP029421
AGAP029422
AGAP029423
AGAP029424
AGAP029425
AGAP029426
AGAP029427
AGAP029428
AGAP029429
AGAP029430
AGAP029431
AGAP029432
mean_tpm
time
preZGA
ZGA_early
ZGA_late
5h
9h
11h
14h
20h
24h
L1
sex
Female
Male

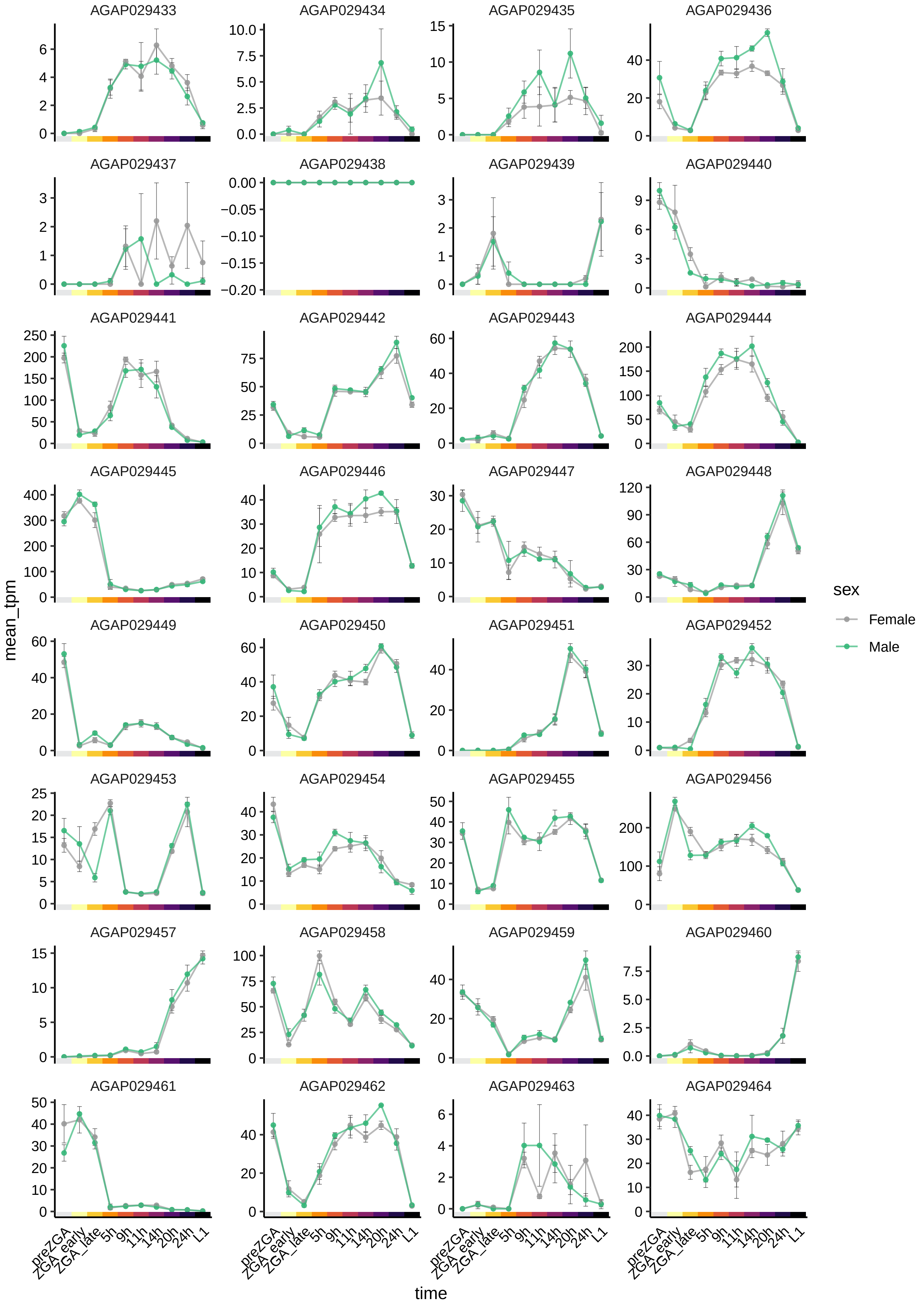
AGAP029433
AGAP029434
AGAP029435
AGAP029436
AGAP029437
AGAP029438
AGAP029439
AGAP029440
AGAP029441
AGAP029442
AGAP029443
AGAP029444
AGAP029445
AGAP029446
AGAP029447
AGAP029448
AGAP029449
AGAP029450
AGAP029451
AGAP029452
AGAP029453
AGAP029454
AGAP029455
AGAP029456
AGAP029457
AGAP029458
AGAP029459
AGAP029460
AGAP029461
AGAP029462
AGAP029463
AGAP029464
mean_tpm
time
preZGA
ZGA_early
ZGA_late
5h
9h
11h
14h
20h
24h
L1
sex
Female
Male

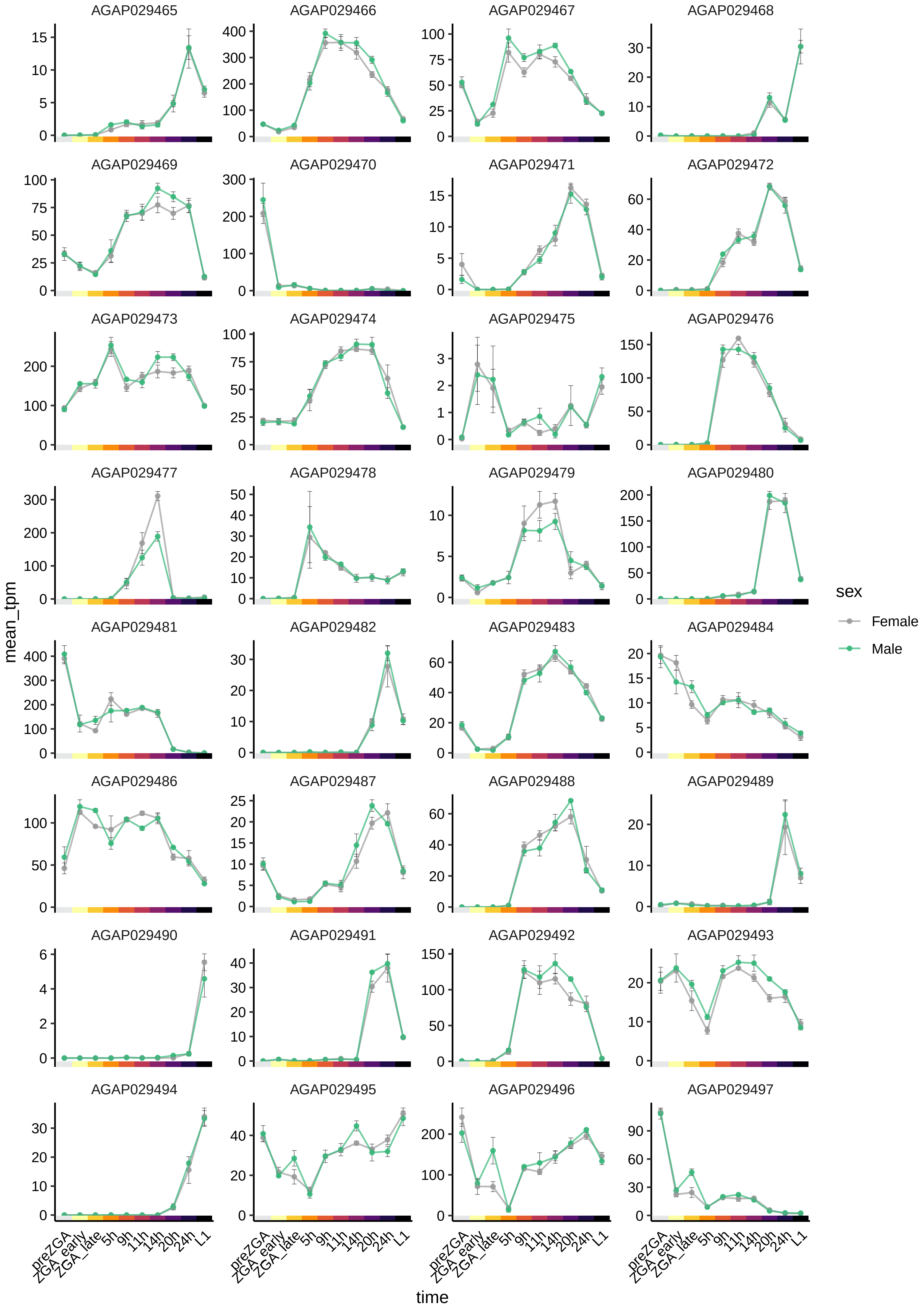

AGAP029465
AGAP029466
AGAP029467
AGAP029468
AGAP029469
AGAP029470
AGAP029471
AGAP029472
AGAP029473
AGAP029474
AGAP029475
AGAP029476
AGAP029477
AGAP029478
AGAP029479
AGAP029480
AGAP029481
AGAP029482
AGAP029483
AGAP029484
AGAP029486
AGAP029487
AGAP029488
AGAP029489
AGAP029490
AGAP029491
AGAP029492
AGAP029493
AGAP029494
AGAP029495
AGAP029496
AGAP029497
mean_tpm
time
preZGA
ZGA_early
ZGA_late
5h
9h
11h
14h
20h
24h
L1
sex
Female
Male

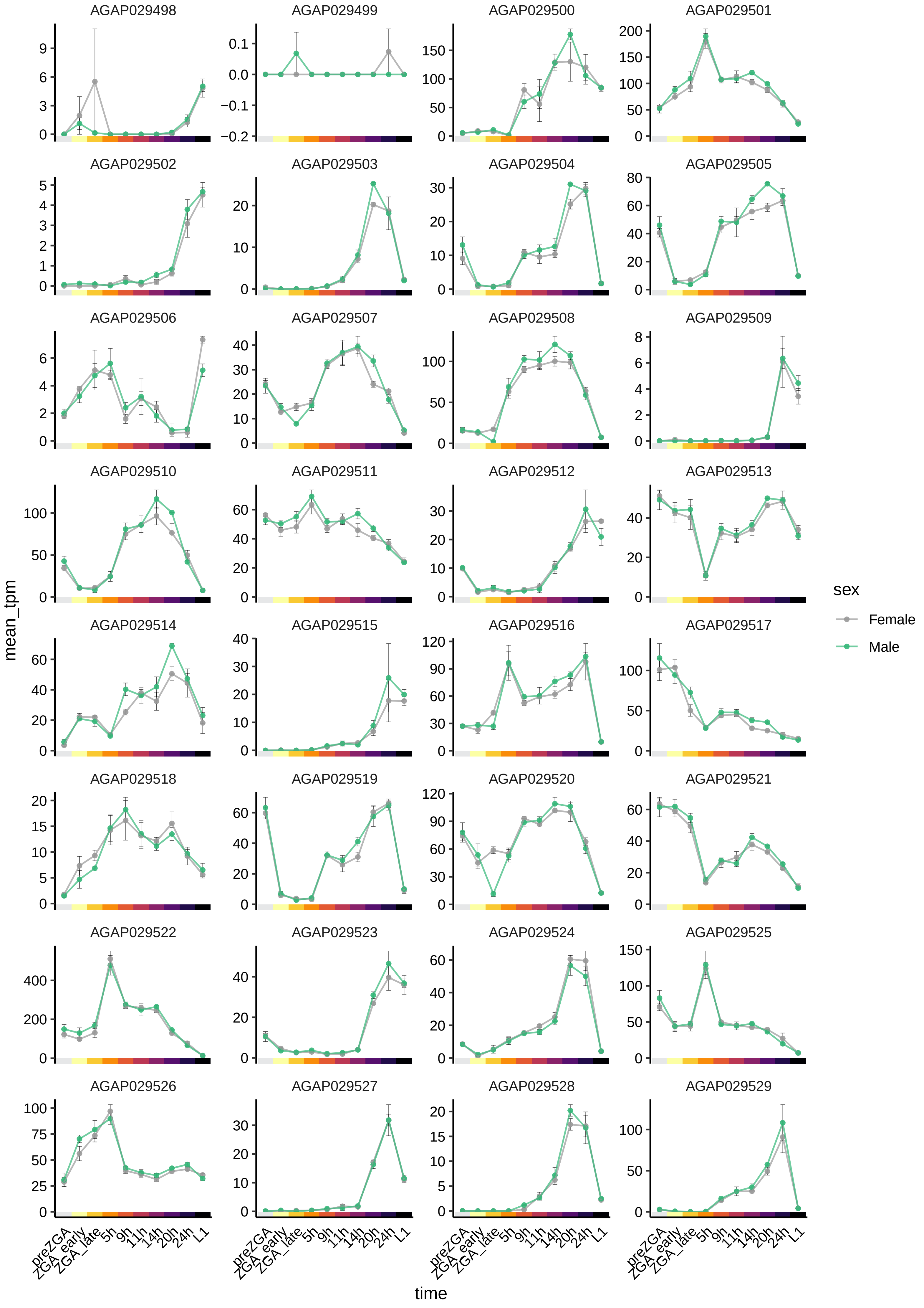
AGAP029498
AGAP029499
AGAP029500
AGAP029501
AGAP029502
AGAP029503
AGAP029504
AGAP029505
AGAP029506
AGAP029507
AGAP029508
AGAP029509
AGAP029510
AGAP029511
AGAP029512
AGAP029513
AGAP029514
AGAP029515
AGAP029516
AGAP029517
AGAP029518
AGAP029519
AGAP029520
AGAP029521
AGAP029522
AGAP029523
AGAP029524
AGAP029525
AGAP029526
AGAP029527
AGAP029528
AGAP029529
mean_tpm
time
preZGA
ZGA_early
ZGA_late
5h
9h
11h
14h
20h
24h
L1
sex
Female
Male

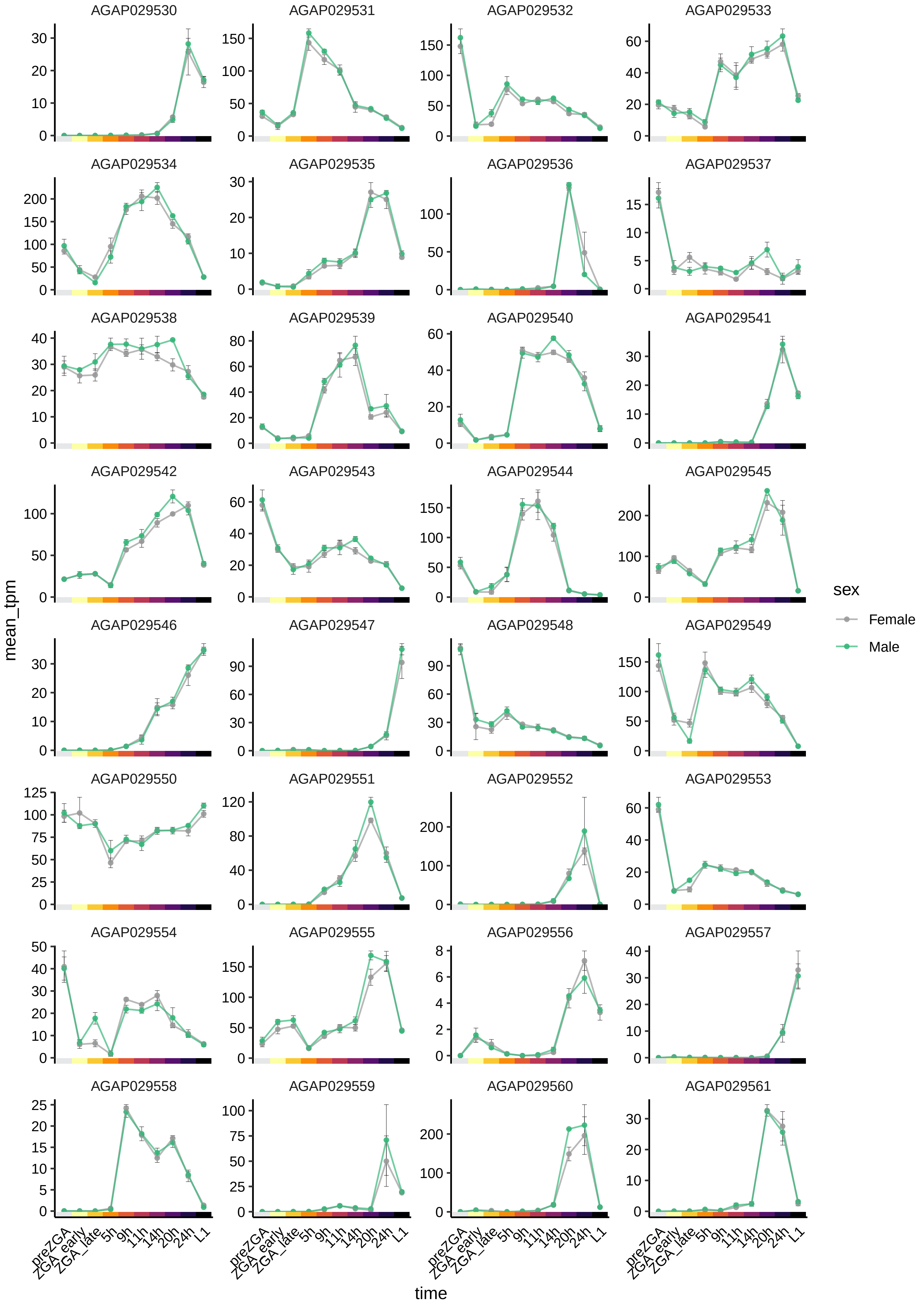
AGAP029530
AGAP029531
AGAP029532
AGAP029533
AGAP029534
AGAP029535
AGAP029536
AGAP029537
AGAP029538
AGAP029539
AGAP029540
AGAP029541
AGAP029542
AGAP029543
AGAP029544
AGAP029545
AGAP029546
AGAP029547
AGAP029548
AGAP029549
AGAP029550
AGAP029551
AGAP029552
AGAP029553
AGAP029554
AGAP029555
AGAP029556
AGAP029557
AGAP029558
AGAP029559
AGAP029560
AGAP029561
mean_tpm
time
preZGA
ZGA_early
ZGA_late
5h
9h
11h
14h
20h
24h
L1
sex
Female
Male

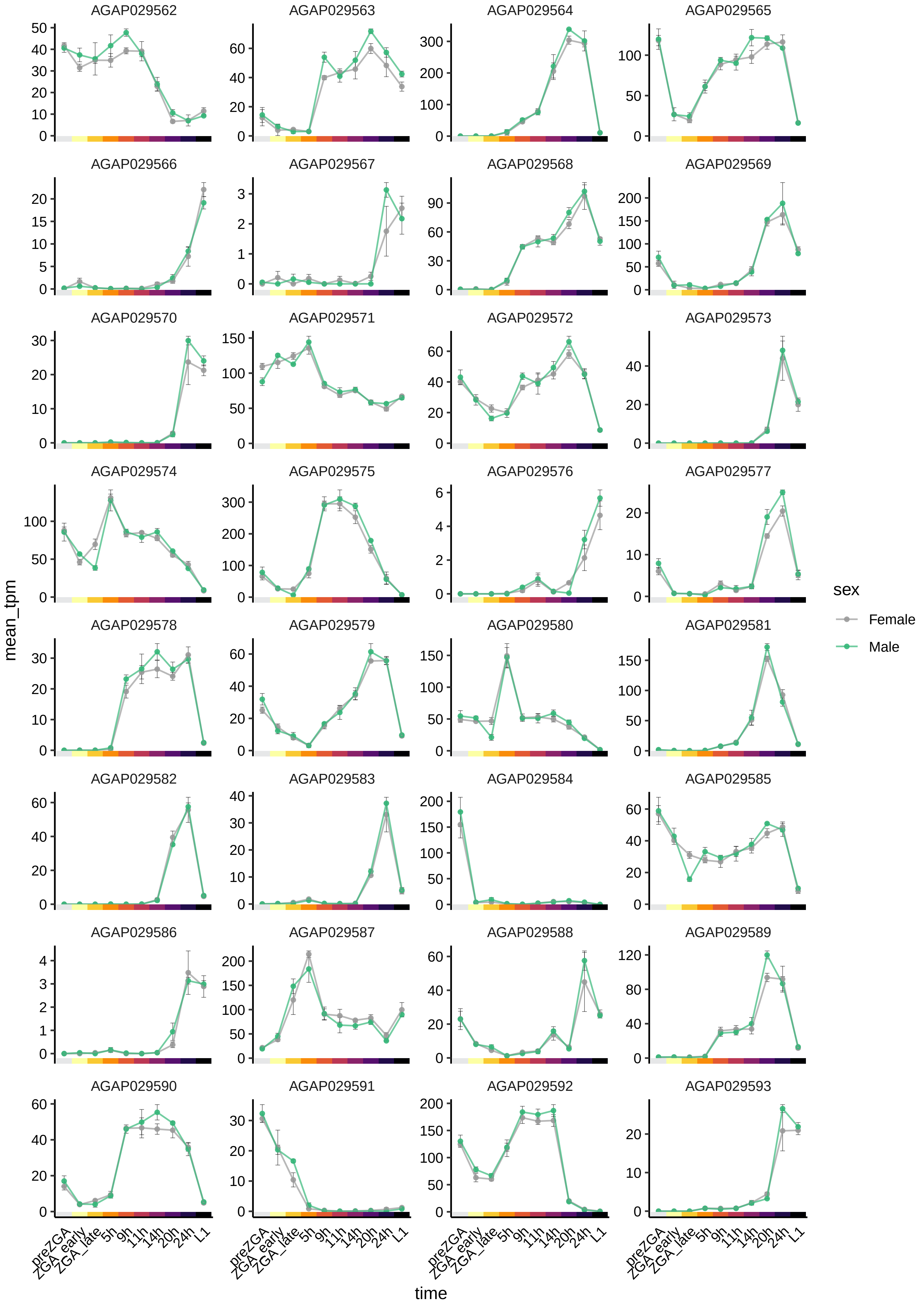
AGAP029562
AGAP029563
AGAP029564
AGAP029565
AGAP029566
AGAP029567
AGAP029568
AGAP029569
AGAP029570
AGAP029571
AGAP029572
AGAP029573
AGAP029574
AGAP029575
AGAP029576
AGAP029577
AGAP029578
AGAP029579
AGAP029580
AGAP029581
AGAP029582
AGAP029583
AGAP029584
AGAP029585
AGAP029586
AGAP029587
AGAP029588
AGAP029589
AGAP029590
AGAP029591
AGAP029592
AGAP029593
mean_tpm
time
preZGA
ZGA_early
ZGA_late
5h
9h
11h
14h
20h
24h
L1
sex
Female
Male

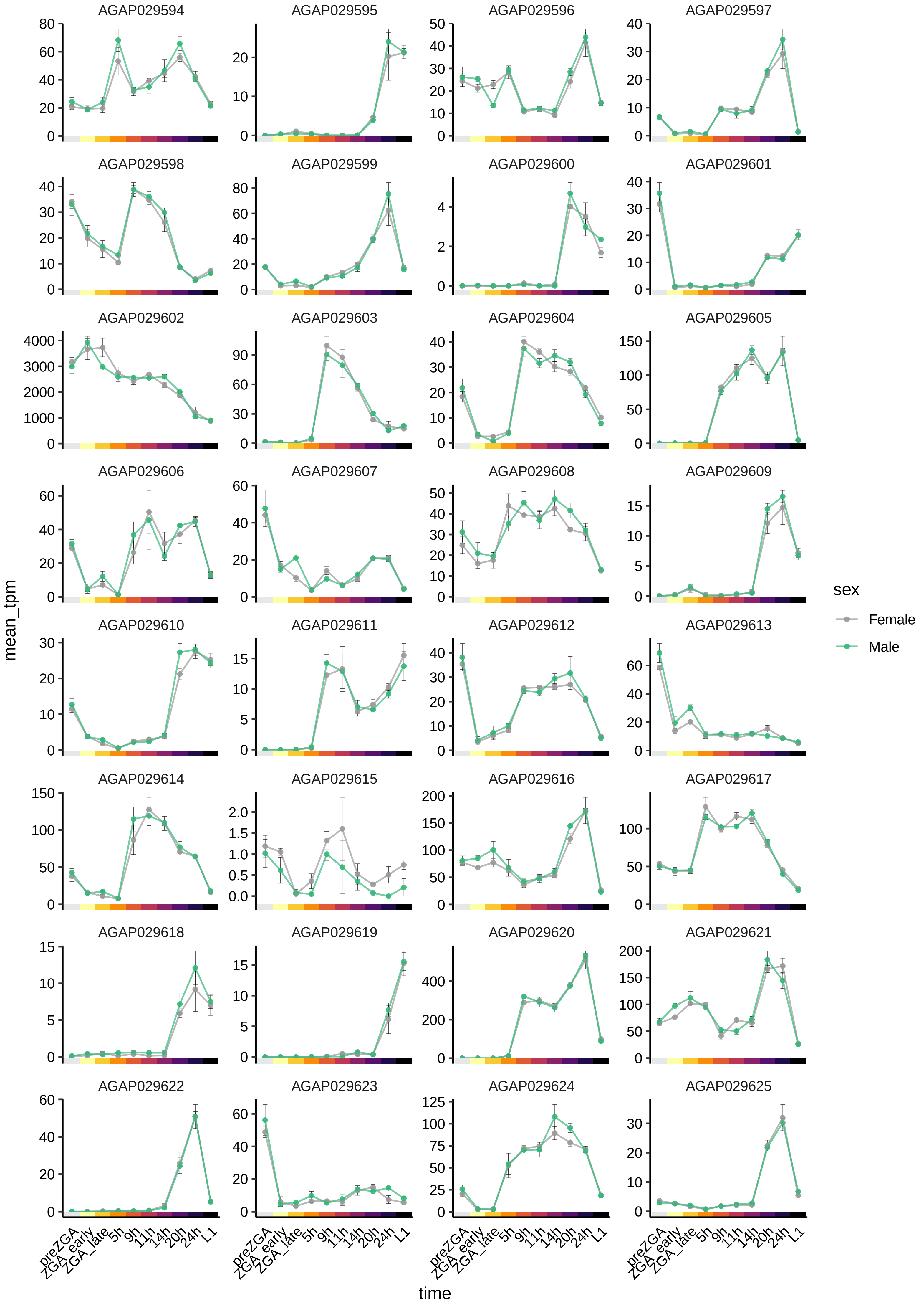
AGAP029594
AGAP029595
AGAP029596
AGAP029597
AGAP029598
AGAP029599
AGAP029600
AGAP029601
AGAP029602
AGAP029603
AGAP029604
AGAP029605
AGAP029606
AGAP029607
AGAP029608
AGAP029609
AGAP029610
AGAP029611
AGAP029612
AGAP029613
AGAP029614
AGAP029615
AGAP029616
AGAP029617
AGAP029618
AGAP029619
AGAP029620
AGAP029621
AGAP029622
AGAP029623
AGAP029624
AGAP029625
mean_tpm
time
preZGA
ZGA_early
ZGA_late
5h
9h
11h
14h
20h
24h
L1
sex
Female
Male

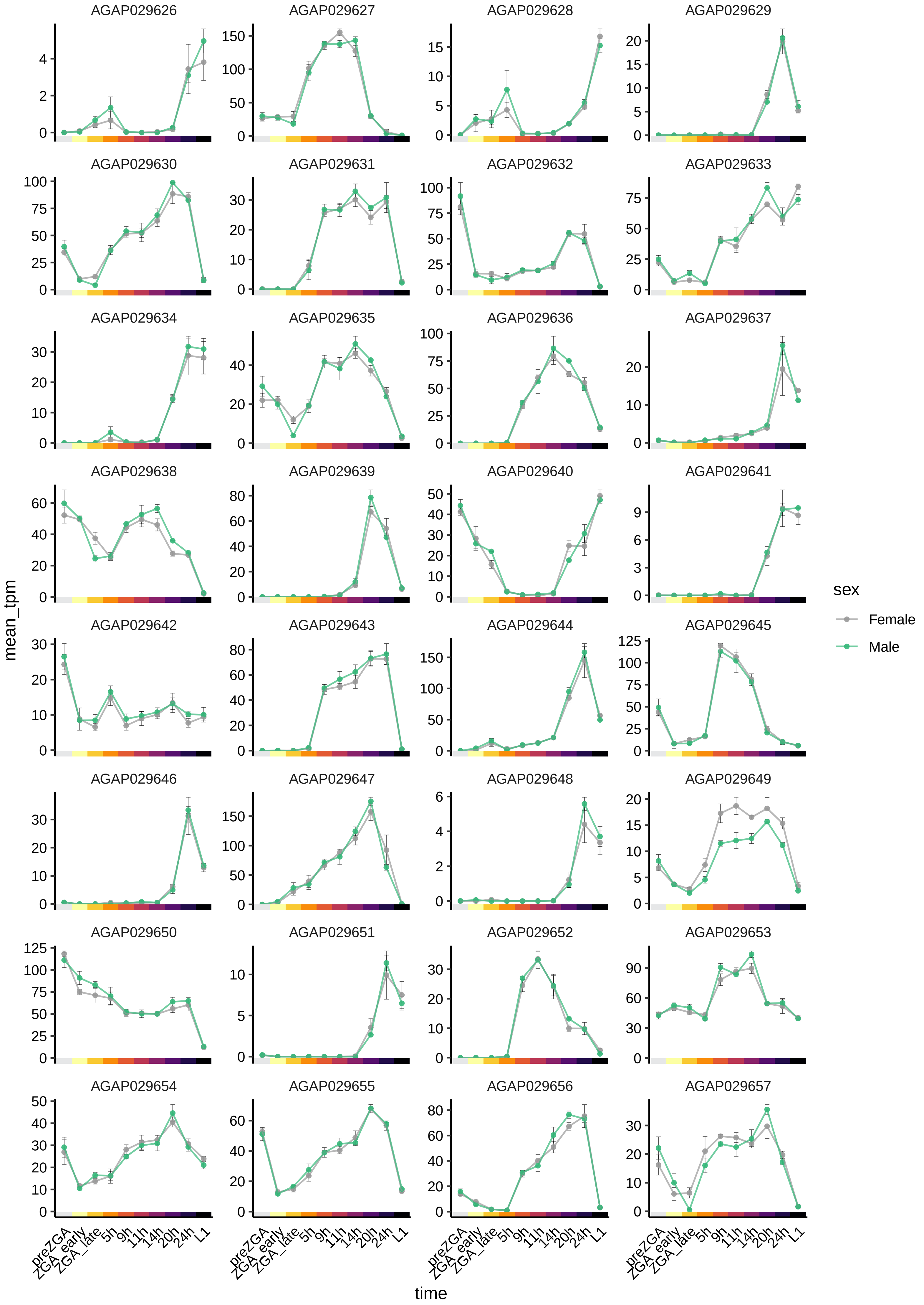

AGAP029626
AGAP029627
AGAP029628
AGAP029629
AGAP029630
AGAP029631
AGAP029632
AGAP029633
AGAP029634
AGAP029635
AGAP029636
AGAP029637
AGAP029638
AGAP029639
AGAP029640
AGAP029641
AGAP029642
AGAP029643
AGAP029644
AGAP029645
AGAP029646
AGAP029647
AGAP029648
AGAP029649
AGAP029650
AGAP029651
AGAP029652
AGAP029653
AGAP029654
AGAP029655
AGAP029656
AGAP029657
mean_tpm
time
preZGA
ZGA_early
ZGA_late
5h
9h
11h
14h
20h
24h
L1
sex
Female
Male

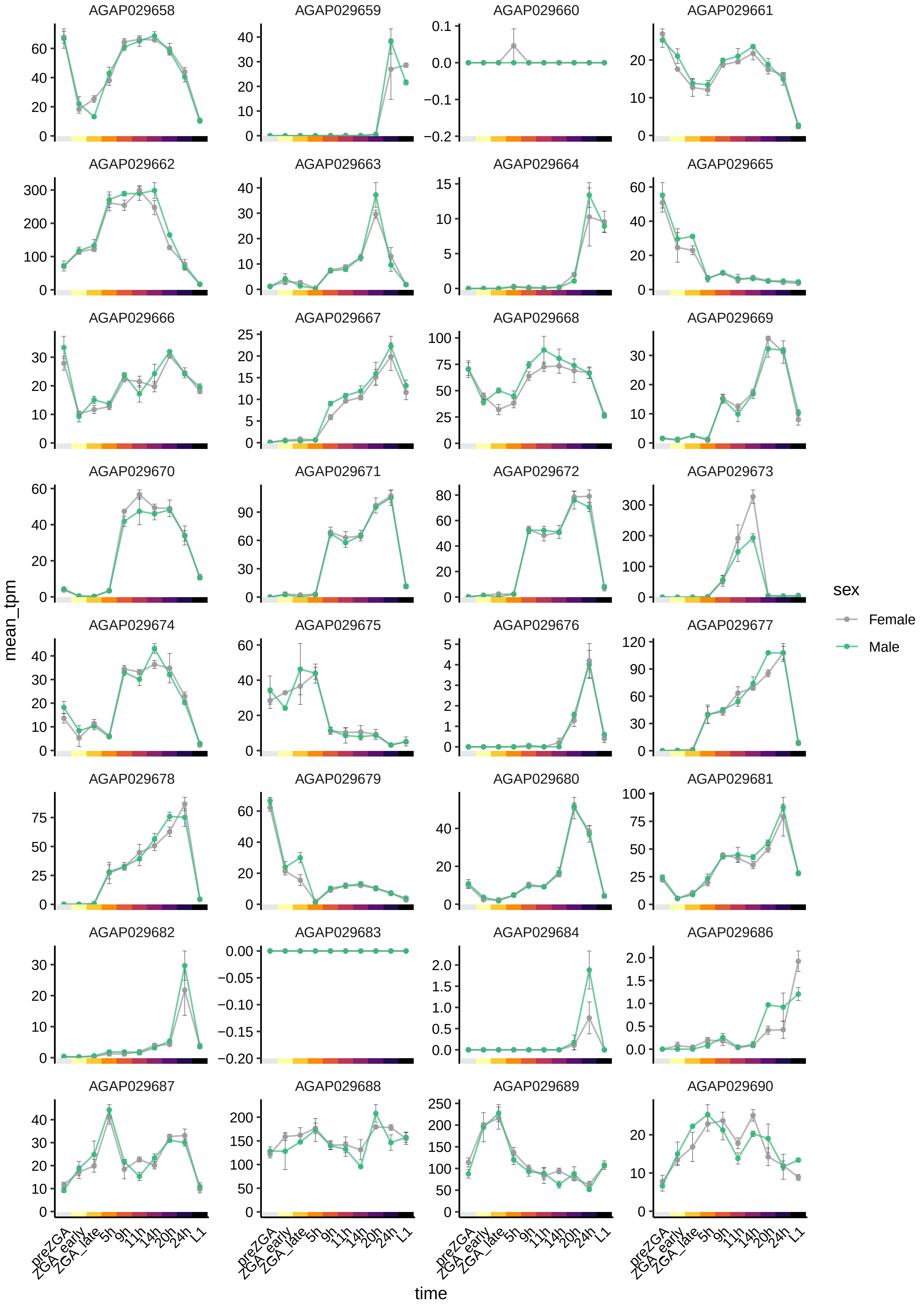
AGAP029658
AGAP029659
AGAP029660
AGAP029661
AGAP029662
AGAP029663
AGAP029664
AGAP029665
AGAP029666
AGAP029667
AGAP029668
AGAP029669
AGAP029670
AGAP029671
AGAP029672
AGAP029673
AGAP029674
AGAP029675
AGAP029676
AGAP029677
AGAP029678
AGAP029679
AGAP029680
AGAP029681
AGAP029682
AGAP029683
AGAP029684
AGAP029686
AGAP029687
AGAP029688
AGAP029689
AGAP029690
mean_tpm
time
preZGA
ZGA_early
ZGA_late
5h
9h
11h
14h
20h
24h
L1
sex
Female
Male

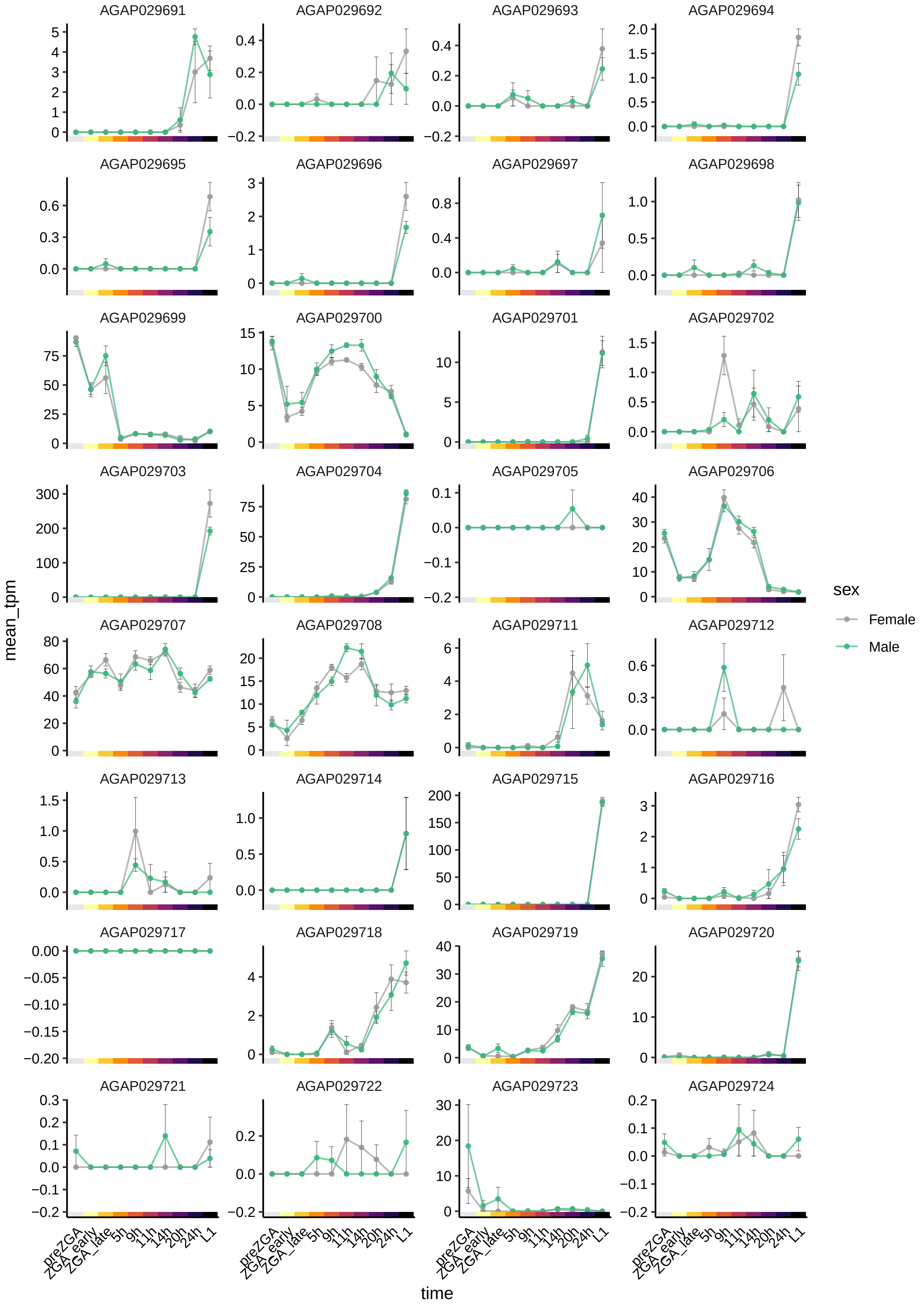

AGAP029691
AGAP029692
AGAP029693
AGAP029694
AGAP029695
AGAP029696
AGAP029697
AGAP029698
AGAP029699
AGAP029700
AGAP029701
AGAP029702
AGAP029703
AGAP029704
AGAP029705
AGAP029706
AGAP029707
AGAP029708
AGAP029711
AGAP029712
AGAP029713
AGAP029714
AGAP029715
AGAP029716
AGAP029717
AGAP029718
AGAP029719
AGAP029720
AGAP029721
AGAP029722
AGAP029723
AGAP029724
mean_tpm
time
preZGA
ZGA_early
ZGA_late
5h
9h
11h
14h
20h
24h
L1
sex
Female
Male

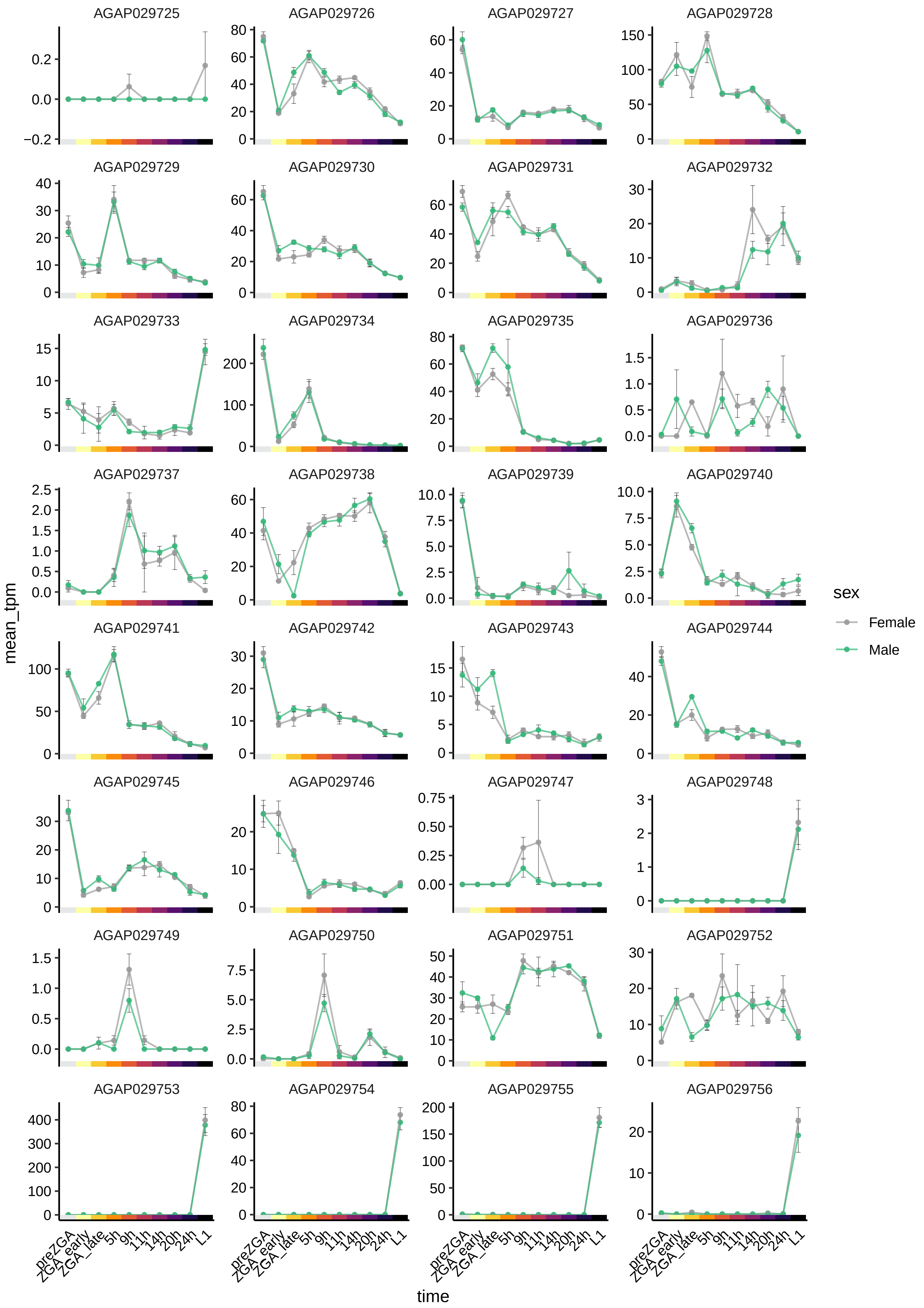

AGAP029725
AGAP029726
AGAP029727
AGAP029728
AGAP029729
AGAP029730
AGAP029731
AGAP029732
AGAP029733
AGAP029734
AGAP029735
AGAP029736
AGAP029737
AGAP029738
AGAP029739
AGAP029740
AGAP029741
AGAP029742
AGAP029743
AGAP029744
AGAP029745
AGAP029746
AGAP029747
AGAP029748
AGAP029749
AGAP029750
AGAP029751
AGAP029752
AGAP029753
AGAP029754
AGAP029755
AGAP029756
mean_tpm
time
preZGA
ZGA_early
ZGA_late
5h
9h
11h
14h
20h
24h
L1
sex
Female
Male

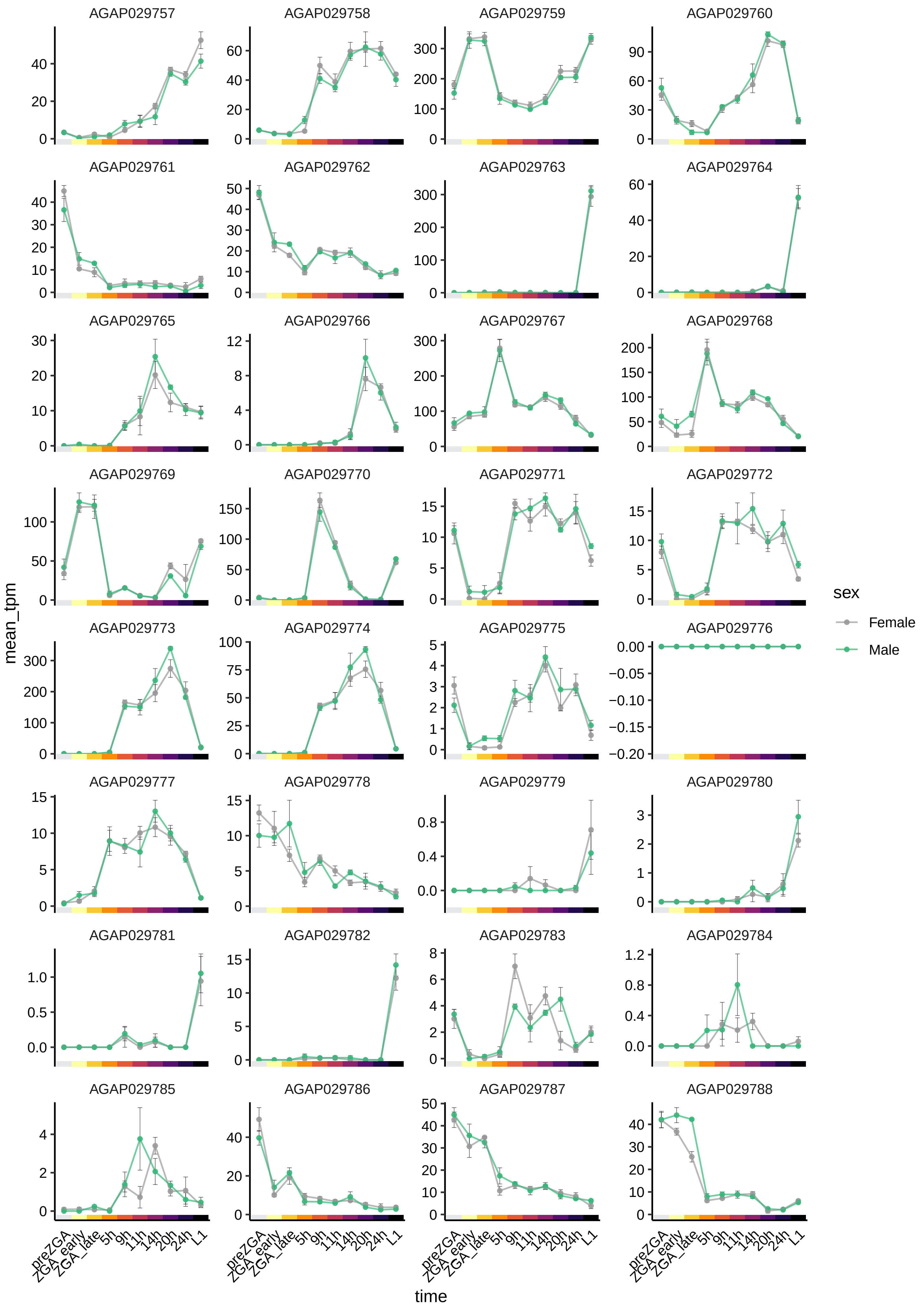
AGAP029757
AGAP029758
AGAP029759
AGAP029760
AGAP029761
AGAP029762
AGAP029763
AGAP029764
AGAP029765
AGAP029766
AGAP029767
AGAP029768
AGAP029769
AGAP029770
AGAP029771
AGAP029772
AGAP029773
AGAP029774
AGAP029775
AGAP029776
AGAP029777
AGAP029778
AGAP029779
AGAP029780
AGAP029781
AGAP029782
AGAP029783
AGAP029784
AGAP029785
AGAP029786
AGAP029787
AGAP029788
mean_tpm
time
preZGA
ZGA_early
ZGA_late
5h
9h
11h
14h
20h
24h
L1
sex
Female
Male

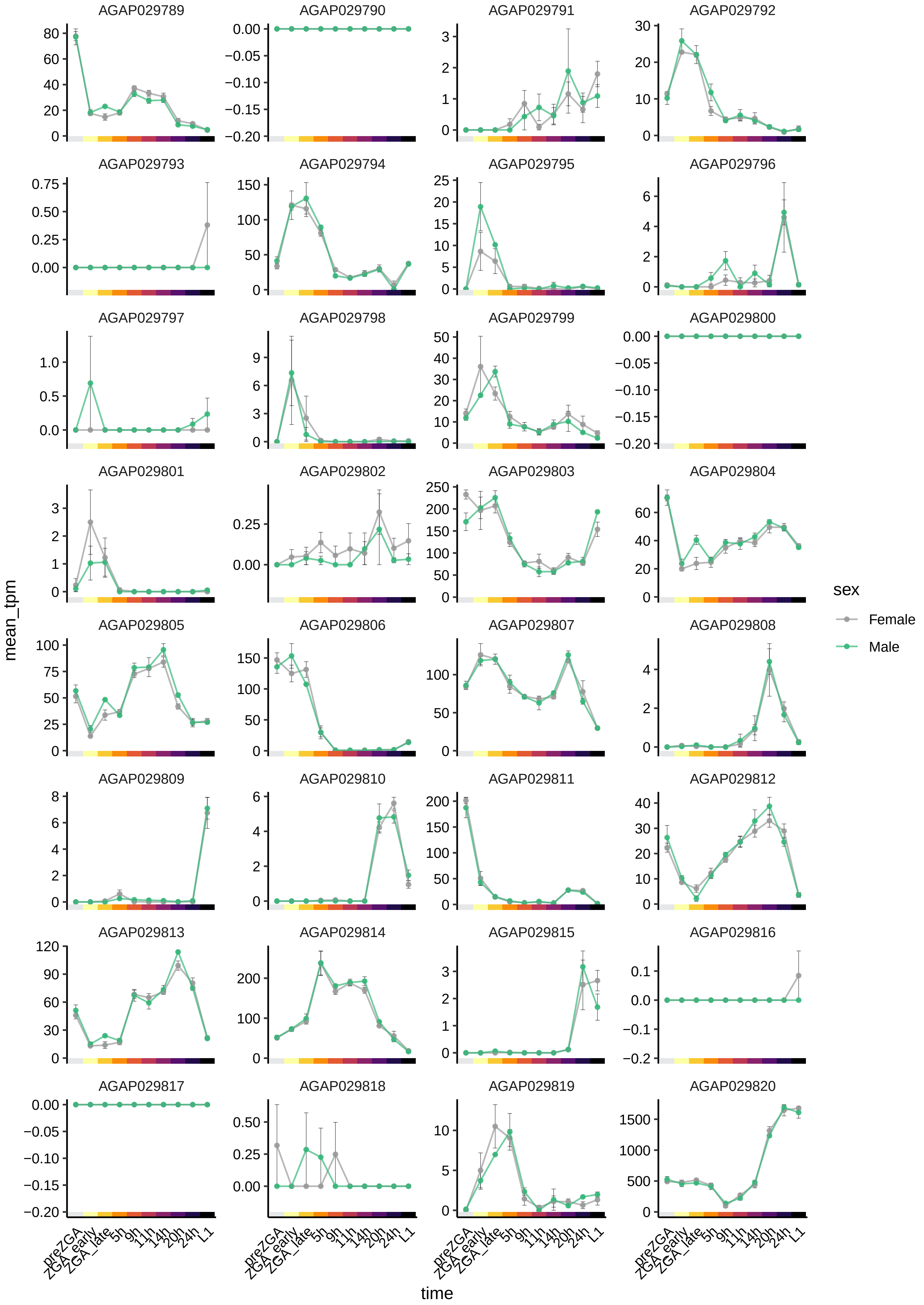
AGAP029789
AGAP029790
AGAP029791
AGAP029792
AGAP029793
AGAP029794
AGAP029795
AGAP029796
AGAP029797
AGAP029798
AGAP029799
AGAP029800
AGAP029801
AGAP029802
AGAP029803
AGAP029804
AGAP029805
AGAP029806
AGAP029807
AGAP029808
AGAP029809
AGAP029810
AGAP029811
AGAP029812
AGAP029813
AGAP029814
AGAP029815
AGAP029816
AGAP029817
AGAP029818
AGAP029819
AGAP029820
mean_tpm
time
preZGA
ZGA_early
ZGA_late
5h
9h
11h
14h
20h
24h
L1
sex
Female
Male

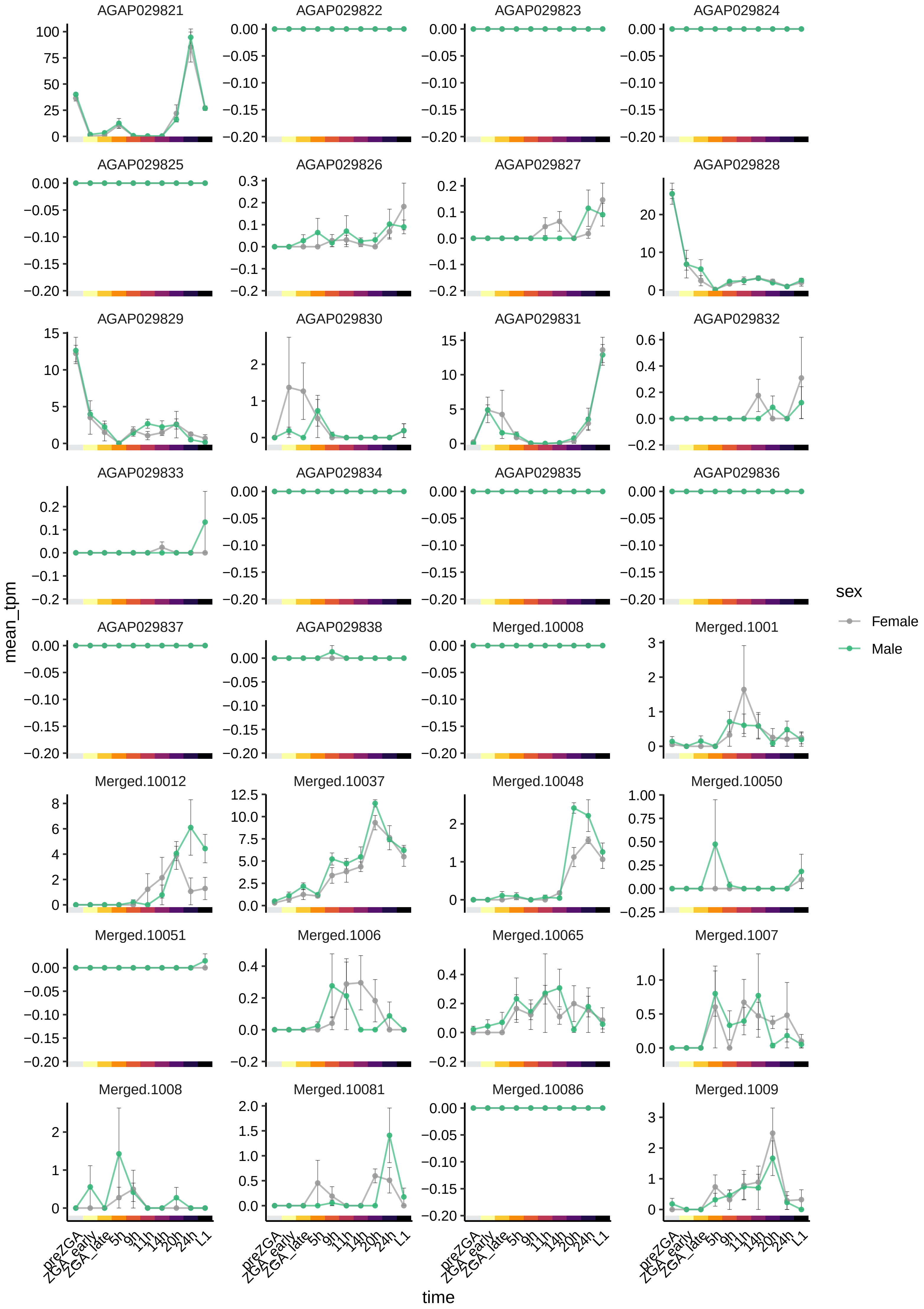
AGAP029821
AGAP029822
AGAP029823
AGAP029824
AGAP029825
AGAP029826
AGAP029827
AGAP029828
AGAP029829
AGAP029830
AGAP029831
AGAP029832
AGAP029833
AGAP029834
AGAP029835
AGAP029836
AGAP029837
AGAP029838
Merged.10008
Merged.1001
Merged.10012
Merged.10037
Merged.10048
Merged.10050
Merged.10051
Merged.1006
Merged.10065
Merged.1007
Merged.1008
Merged.10081
Merged.10086
Merged.1009
mean_tpm
time
preZGA
ZGA_early
ZGA_late
5h
9h
11h
14h
20h
24h
L1
sex
Female
Male

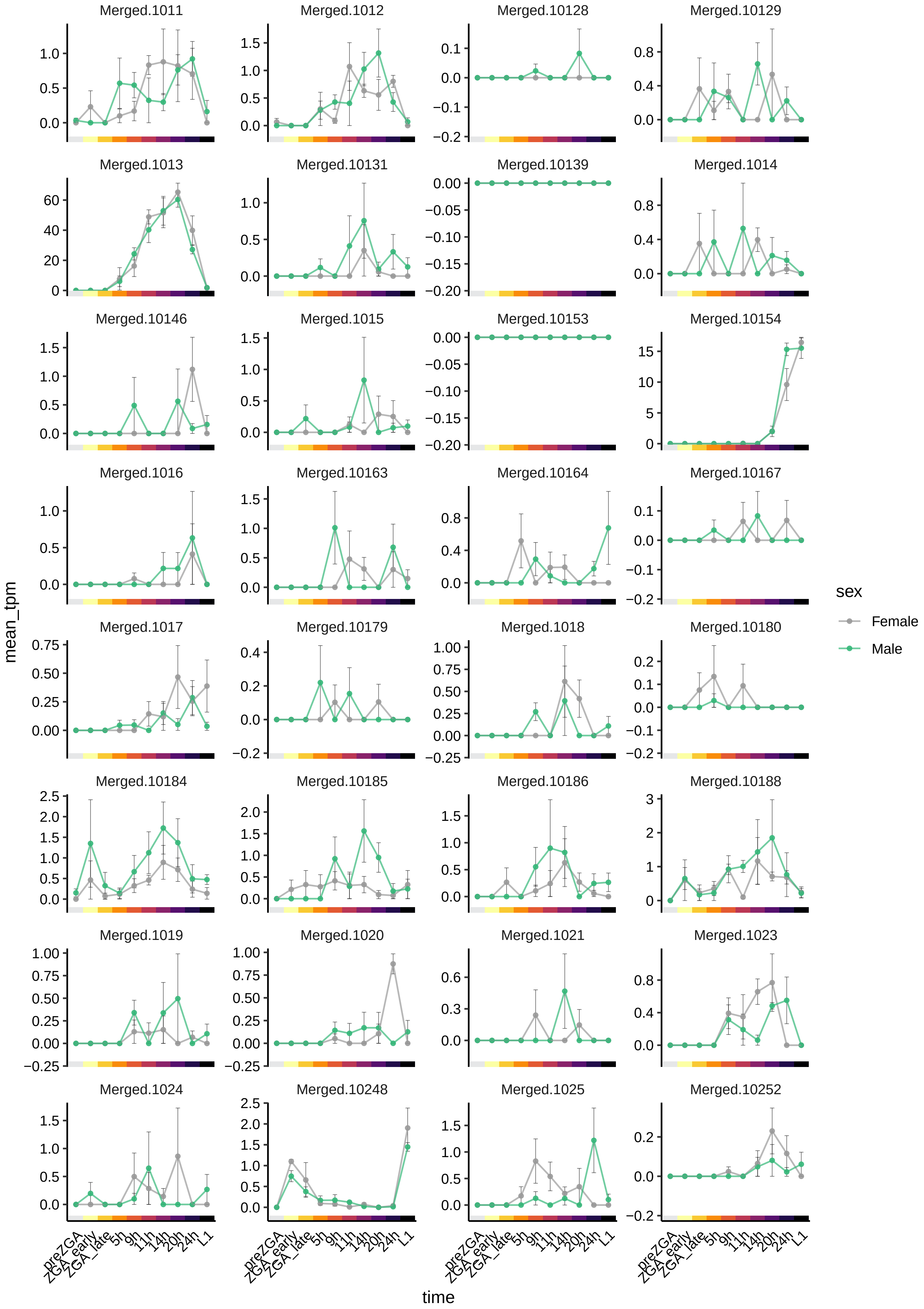

Merged.1011
Merged.1012
Merged.10128
Merged.10129
Merged.1013
Merged.10131
Merged.10139
Merged.1014
Merged.10146
Merged.1015
Merged.10153
Merged.10154
Merged.1016
Merged.10163
Merged.10164
Merged.10167
Merged.1017
Merged.10179
Merged.1018
Merged.10180
Merged.10184
Merged.10185
Merged.10186
Merged.10188
Merged.1019
Merged.1020
Merged.1021
Merged.1023
Merged.1024
Merged.10248
Merged.1025
Merged.10252
mean_tpm
time
preZGA
ZGA_early
ZGA_late
5h
9h
11h
14h
20h
24h
L1
sex
Female
Male

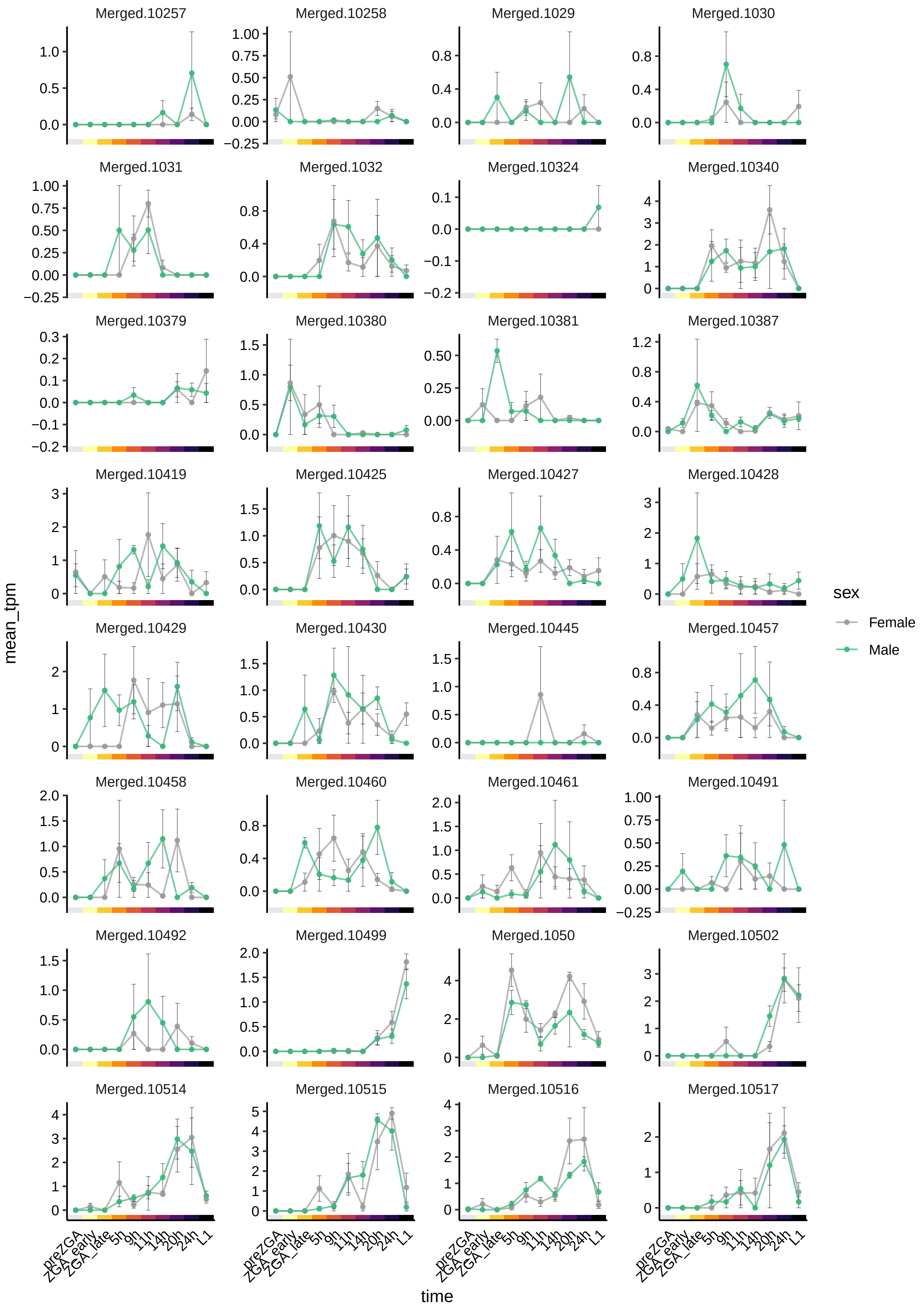

Merged.10257
Merged.10258
Merged.1029
Merged.1030
Merged.1031
Merged.1032
Merged.10324
Merged.10340
Merged.10379
Merged.10380
Merged.10381
Merged.10387
Merged.10419
Merged.10425
Merged.10427
Merged.10428
Merged.10429
Merged.10430
Merged.10445
Merged.10457
Merged.10458
Merged.10460
Merged.10461
Merged.10491
Merged.10492
Merged.10499
Merged.1050
Merged.10502
Merged.10514
Merged.10515
Merged.10516
Merged.10517
mean_tpm
time
preZGA
ZGA_early
ZGA_late
5h
9h
11h
14h
20h
24h
L1
sex
Female
Male

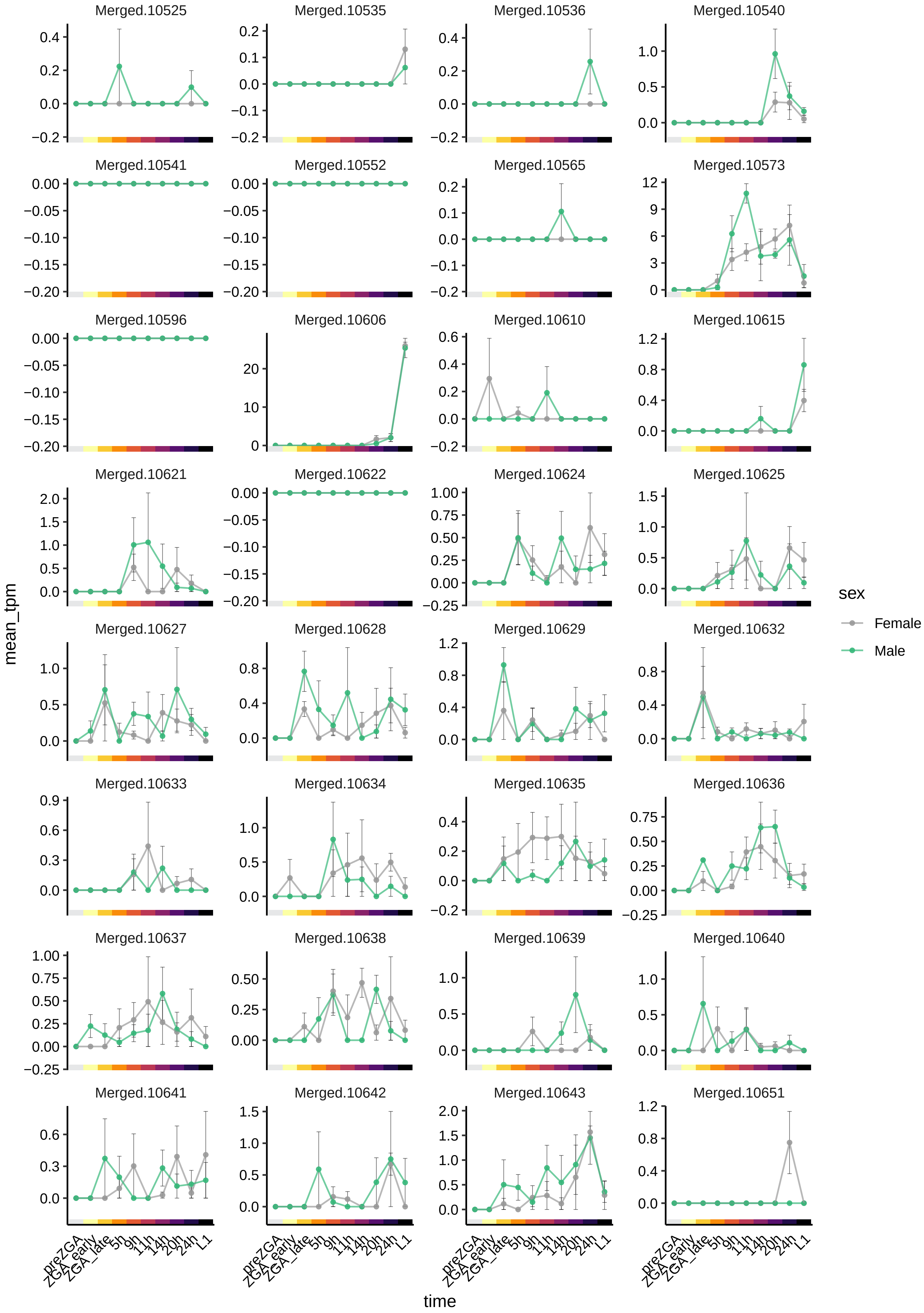

Merged.10525
Merged.10535
Merged.10536
Merged.10540
Merged.10541
Merged.10552
Merged.10565
Merged.10573
Merged.10596
Merged.10606
Merged.10610
Merged.10615
Merged.10621
Merged.10622
Merged.10624
Merged.10625
Merged.10627
Merged.10628
Merged.10629
Merged.10632
Merged.10633
Merged.10634
Merged.10635
Merged.10636
Merged.10637
Merged.10638
Merged.10639
Merged.10640
Merged.10641
Merged.10642
Merged.10643
Merged.10651
mean_tpm
time
preZGA
ZGA_early
ZGA_late
5h
9h
11h
14h
20h
24h
L1
sex
Female
Male

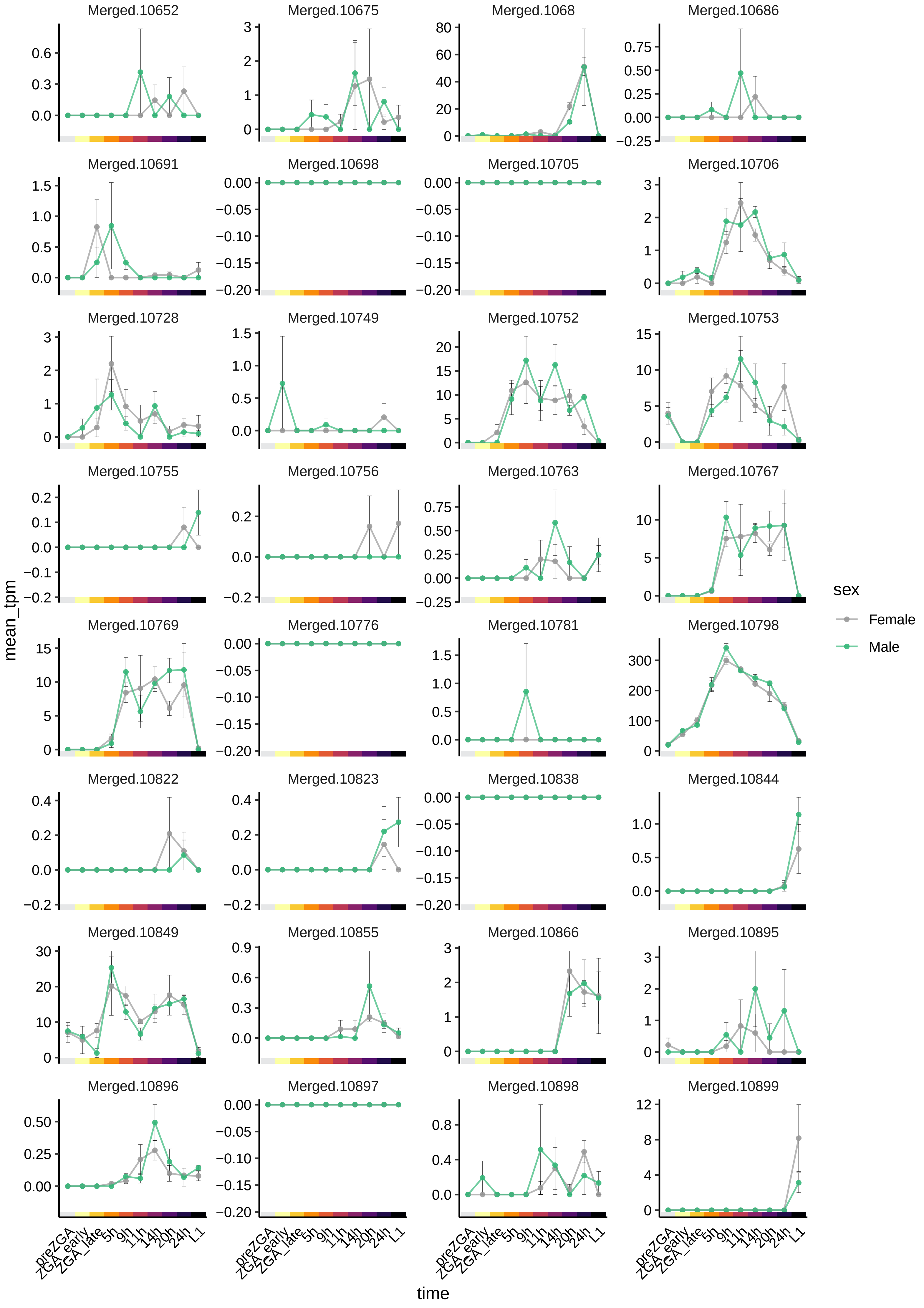

Merged.10652
Merged.10675
Merged.1068
Merged.10686
Merged.10691
Merged.10698
Merged.10705
Merged.10706
Merged.10728
Merged.10749
Merged.10752
Merged.10753
Merged.10755
Merged.10756
Merged.10763
Merged.10767
Merged.10769
Merged.10776
Merged.10781
Merged.10798
Merged.10822
Merged.10823
Merged.10838
Merged.10844
Merged.10849
Merged.10855
Merged.10866
Merged.10895
Merged.10896
Merged.10897
Merged.10898
Merged.10899
mean_tpm
time
preZGA
ZGA_early
ZGA_late
5h
9h
11h
14h
20h
24h
L1
sex
Female
Male

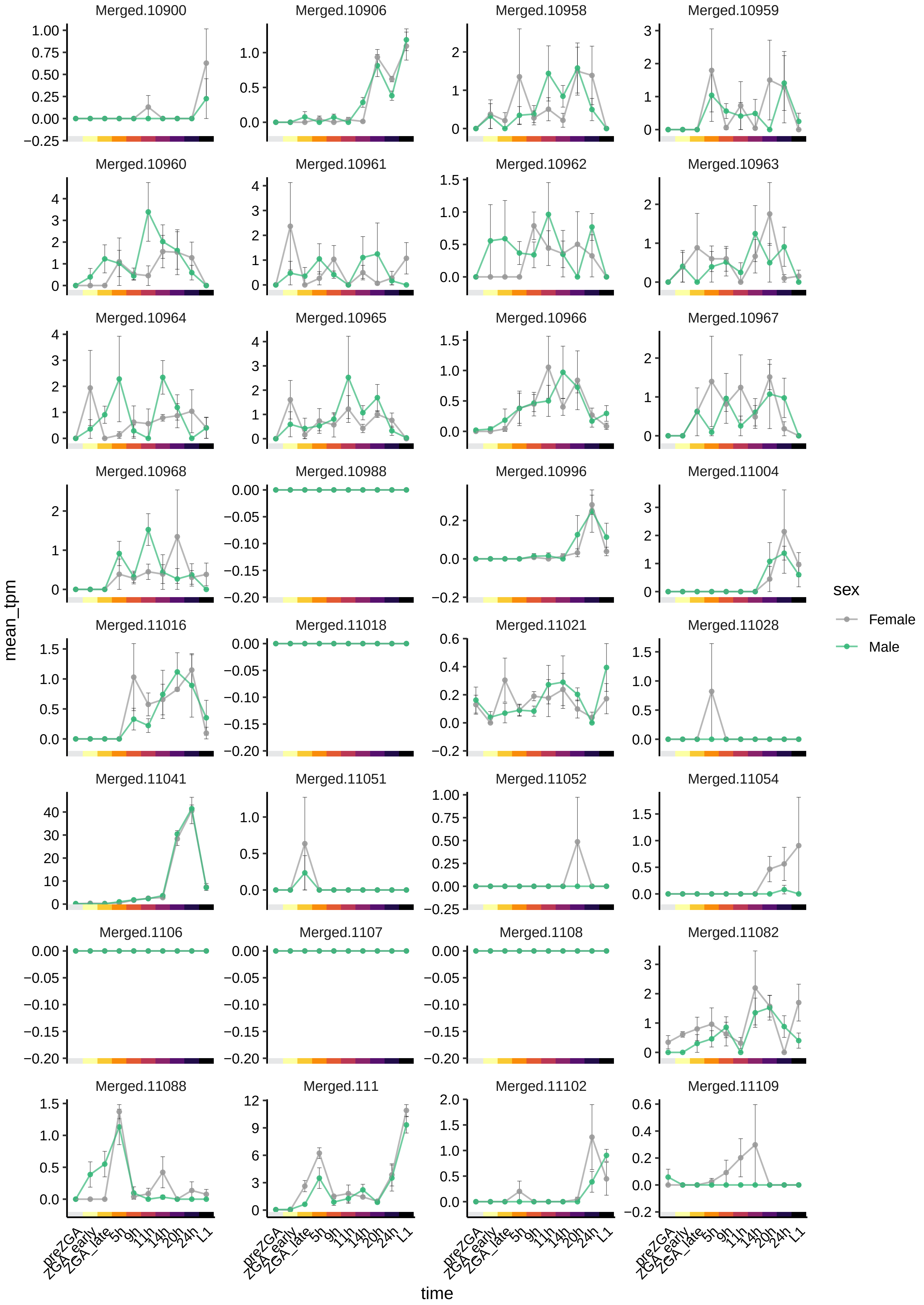

Merged.10900
Merged.10906
Merged.10958
Merged.10959
Merged.10960
Merged.10961
Merged.10962
Merged.10963
Merged.10964
Merged.10965
Merged.10966
Merged.10967
Merged.10968
Merged.10988
Merged.10996
Merged.11004
Merged.11016
Merged.11018
Merged.11021
Merged.11028
Merged.11041
Merged.11051
Merged.11052
Merged.11054
Merged.1106
Merged.1107
Merged.1108
Merged.11082
Merged.11088
Merged.111
Merged.11102
Merged.11109
mean_tpm
time
preZGA
ZGA_early
ZGA_late
5h
9h
11h
14h
20h
24h
L1
sex
Female
Male

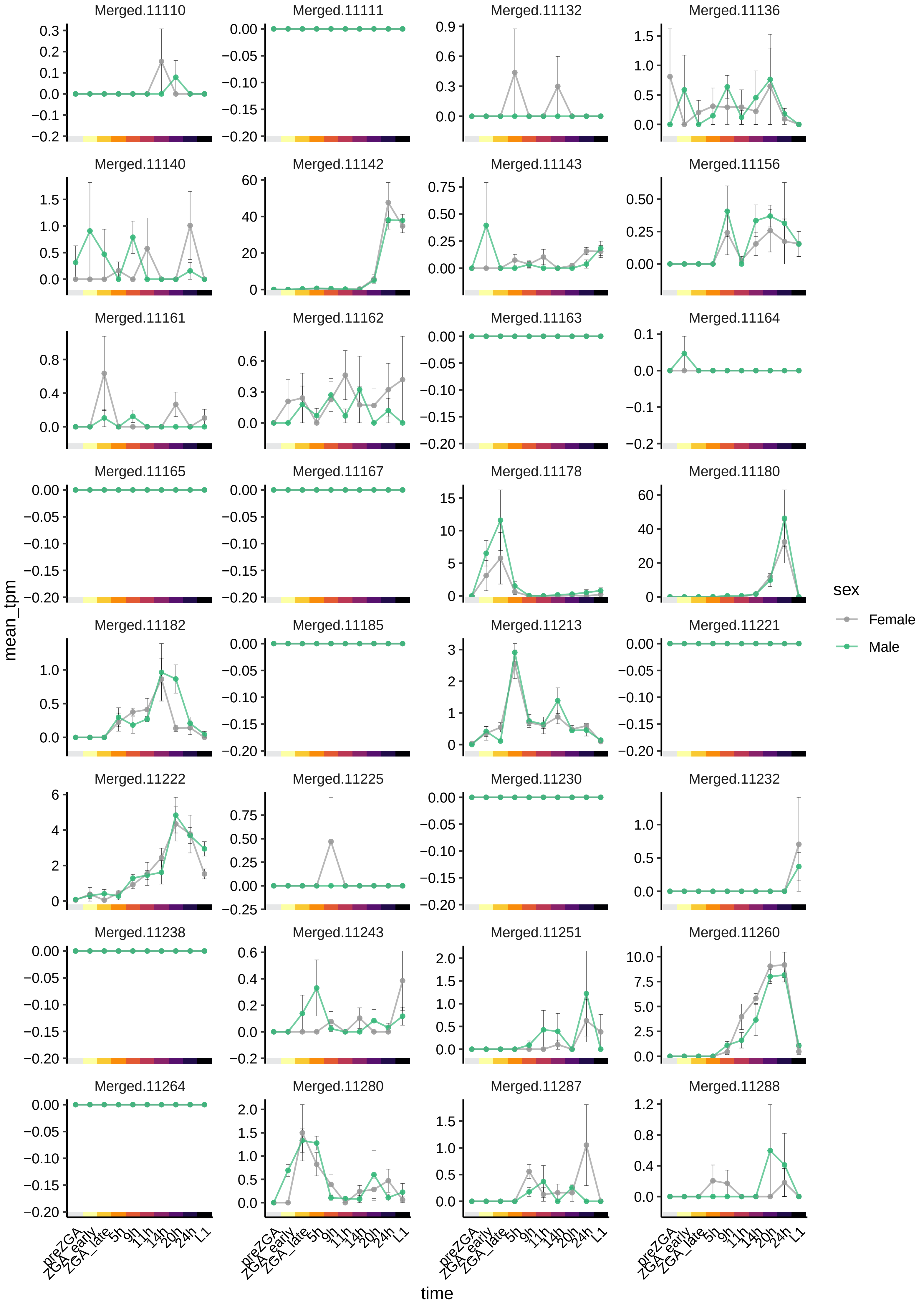
Merged.11110
Merged.11111
Merged.11132
Merged.11136
Merged.11140
Merged.11142
Merged.11143
Merged.11156
Merged.11161
Merged.11162
Merged.11163
Merged.11164
Merged.11165
Merged.11167
Merged.11178
Merged.11180
Merged.11182
Merged.11185
Merged.11213
Merged.11221
Merged.11222
Merged.11225
Merged.11230
Merged.11232
Merged.11238
Merged.11243
Merged.11251
Merged.11260
Merged.11264
Merged.11280
Merged.11287
Merged.11288
mean_tpm
time
preZGA
ZGA_early
ZGA_late
5h
9h
11h
14h
20h
24h
L1
sex
Female
Male

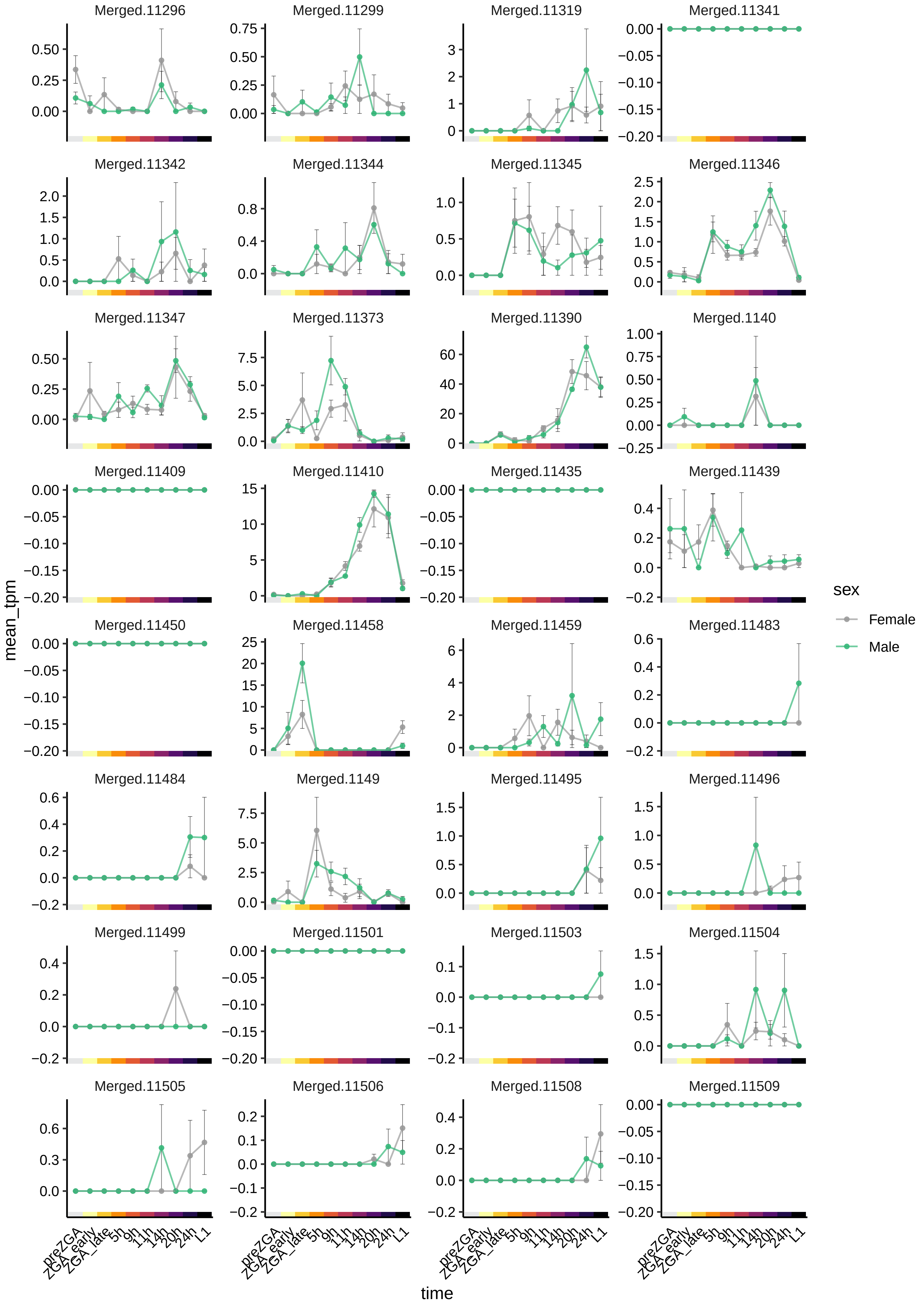

Merged.11296
Merged.11299
Merged.11319
Merged.11341
Merged.11342
Merged.11344
Merged.11345
Merged.11346
Merged.11347
Merged.11373
Merged.11390
Merged.1140
Merged.11409
Merged.11410
Merged.11435
Merged.11439
Merged.11450
Merged.11458
Merged.11459
Merged.11483
Merged.11484
Merged.1149
Merged.11495
Merged.11496
Merged.11499
Merged.11501
Merged.11503
Merged.11504
Merged.11505
Merged.11506
Merged.11508
Merged.11509
mean_tpm
time
preZGA
ZGA_early
ZGA_late
5h
9h
11h
14h
20h
24h
L1
sex
Female
Male

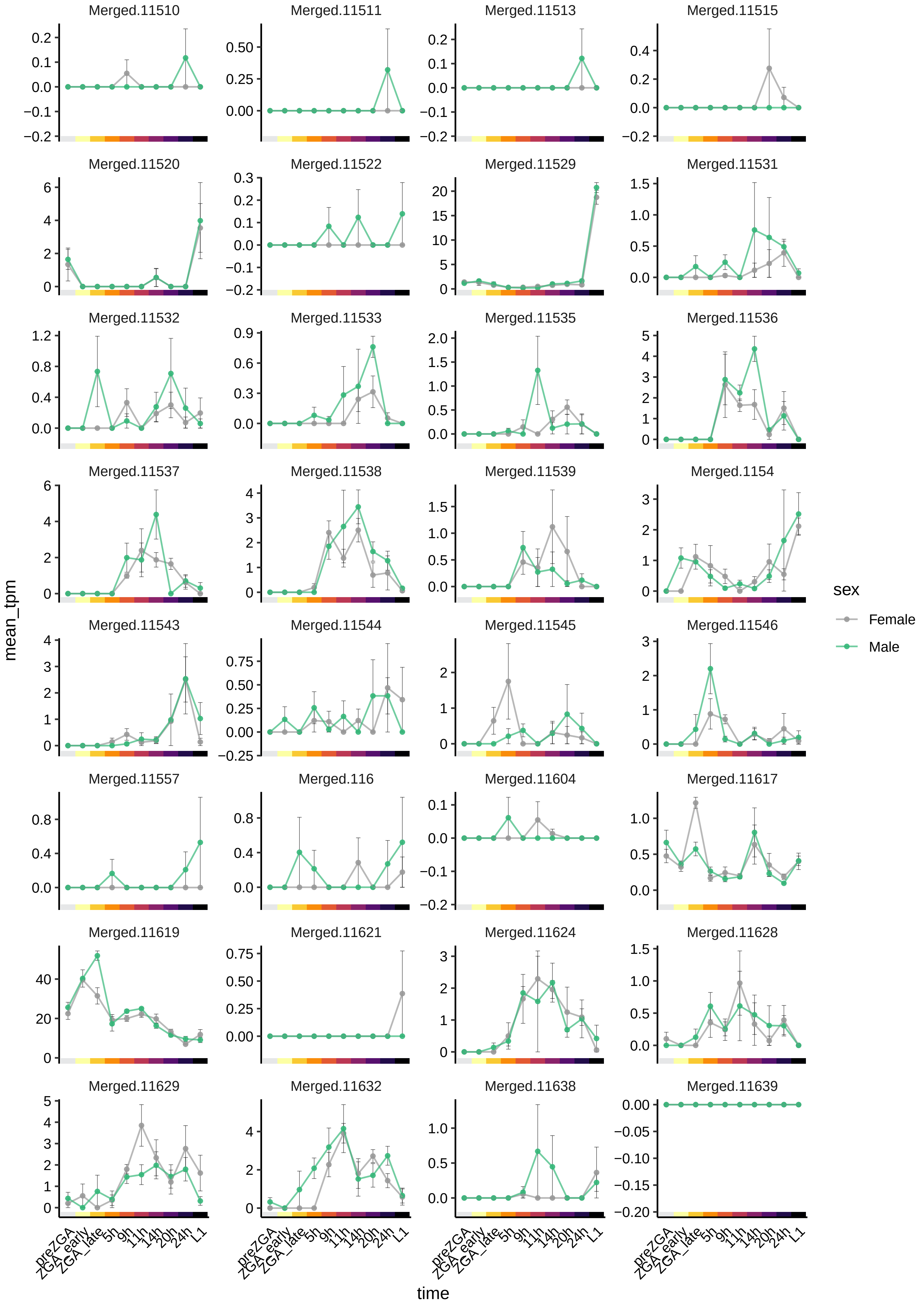

Merged.11510
Merged.11511
Merged.11513
Merged.11515
Merged.11520
Merged.11522
Merged.11529
Merged.11531
Merged.11532
Merged.11533
Merged.11535
Merged.11536
Merged.11537
Merged.11538
Merged.11539
Merged.1154
Merged.11543
Merged.11544
Merged.11545
Merged.11546
Merged.11557
Merged.116
Merged.11604
Merged.11617
Merged.11619
Merged.11621
Merged.11624
Merged.11628
Merged.11629
Merged.11632
Merged.11638
Merged.11639
mean_tpm
time
preZGA
ZGA_early
ZGA_late
5h
9h
11h
14h
20h
24h
L1
sex
Female
Male

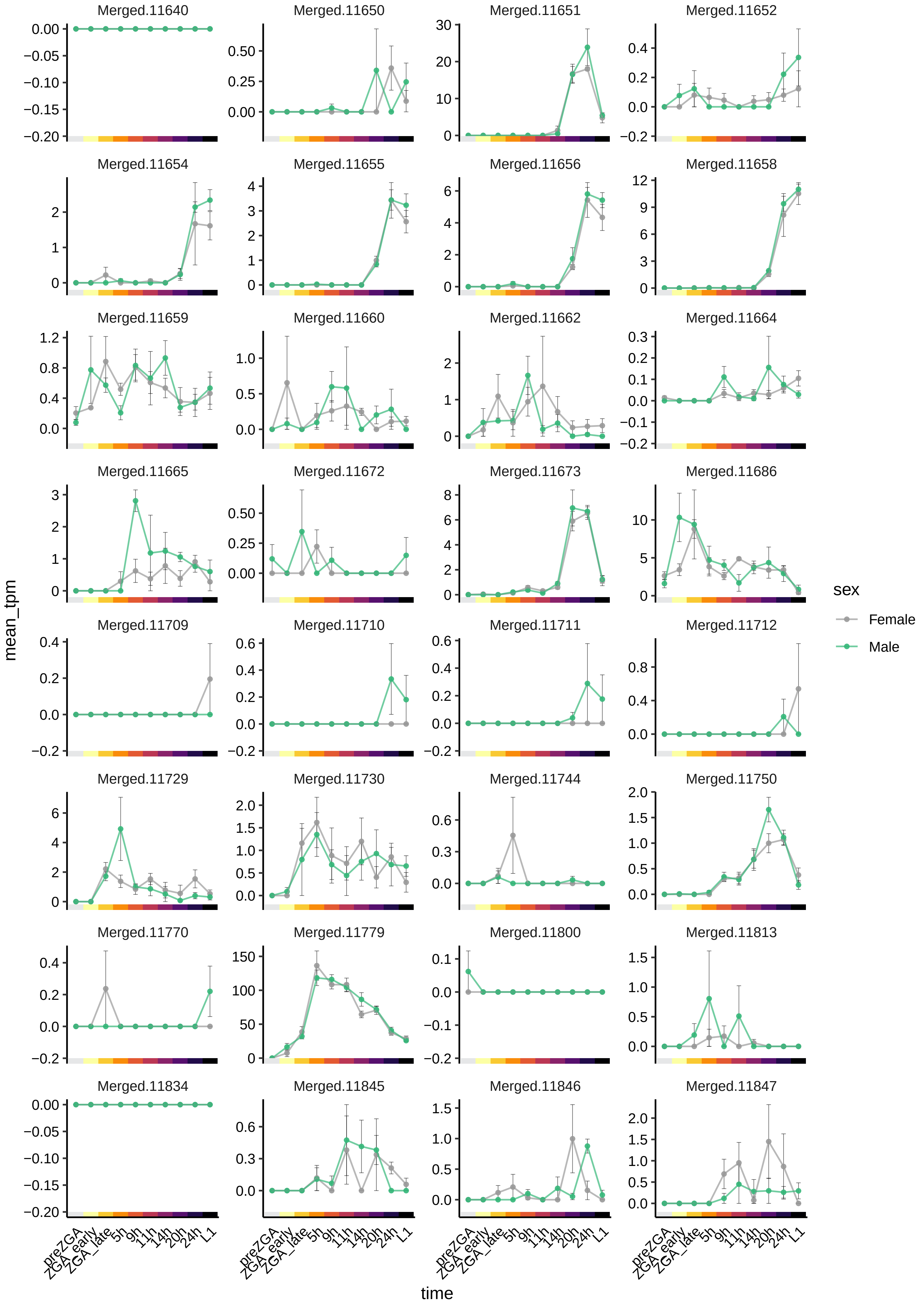
Merged.11640
Merged.11650
Merged.11651
Merged.11652
Merged.11654
Merged.11655
Merged.11656
Merged.11658
Merged.11659
Merged.11660
Merged.11662
Merged.11664
Merged.11665
Merged.11672
Merged.11673
Merged.11686
Merged.11709
Merged.11710
Merged.11711
Merged.11712
Merged.11729
Merged.11730
Merged.11744
Merged.11750
Merged.11770
Merged.11779
Merged.11800
Merged.11813
Merged.11834
Merged.11845
Merged.11846
Merged.11847
mean_tpm
time
preZGA
ZGA_early
ZGA_late
5h
9h
11h
14h
20h
24h
L1
sex
Female
Male

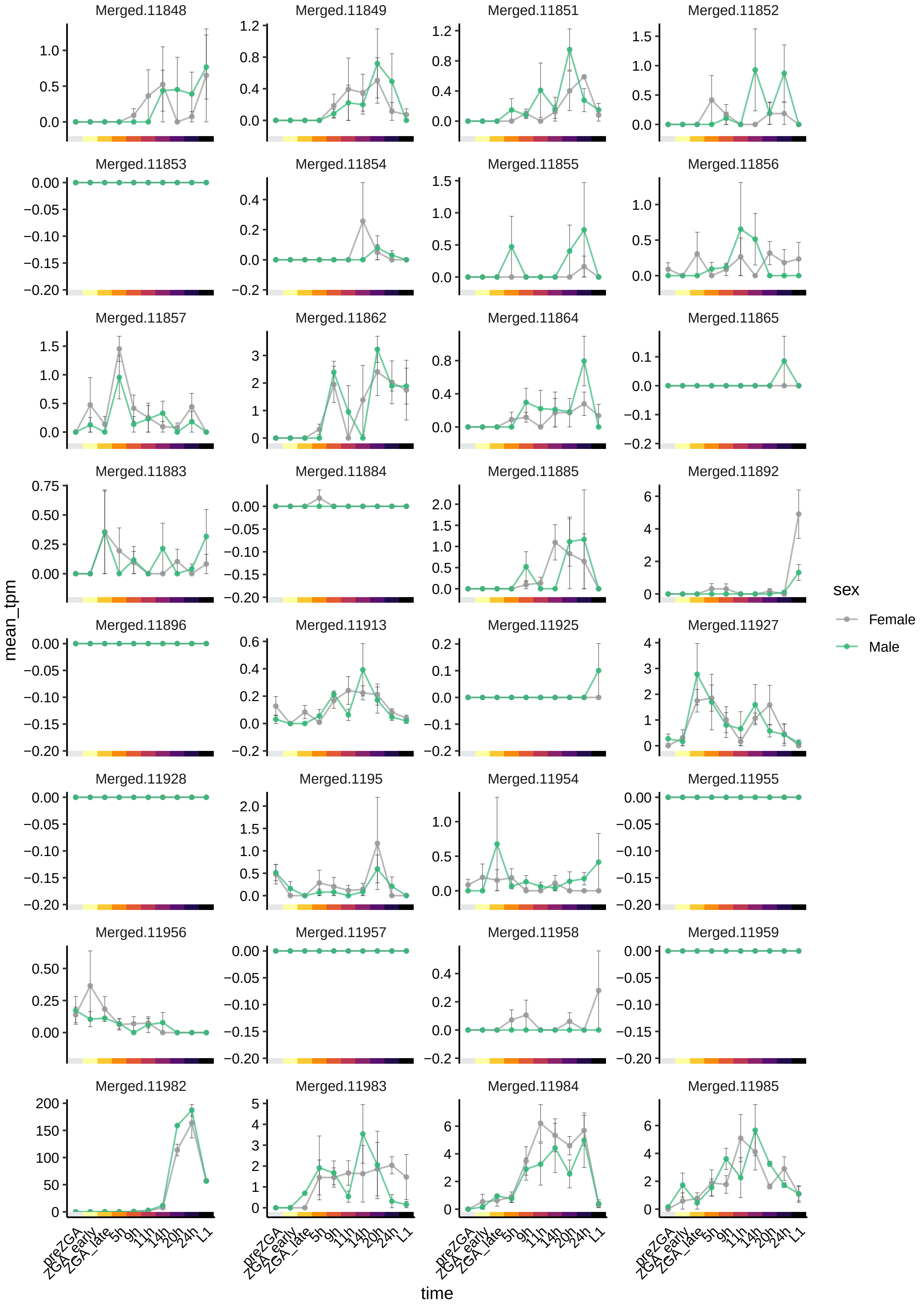

Merged.11848
Merged.11849
Merged.11851
Merged.11852
Merged.11853
Merged.11854
Merged.11855
Merged.11856
Merged.11857
Merged.11862
Merged.11864
Merged.11865
Merged.11883
Merged.11884
Merged.11885
Merged.11892
Merged.11896
Merged.11913
Merged.11925
Merged.11927
Merged.11928
Merged.1195
Merged.11954
Merged.11955
Merged.11956
Merged.11957
Merged.11958
Merged.11959
Merged.11982
Merged.11983
Merged.11984
Merged.11985
mean_tpm
time
preZGA
ZGA_early
ZGA_late
5h
9h
11h
14h
20h
24h
L1
sex
Female
Male

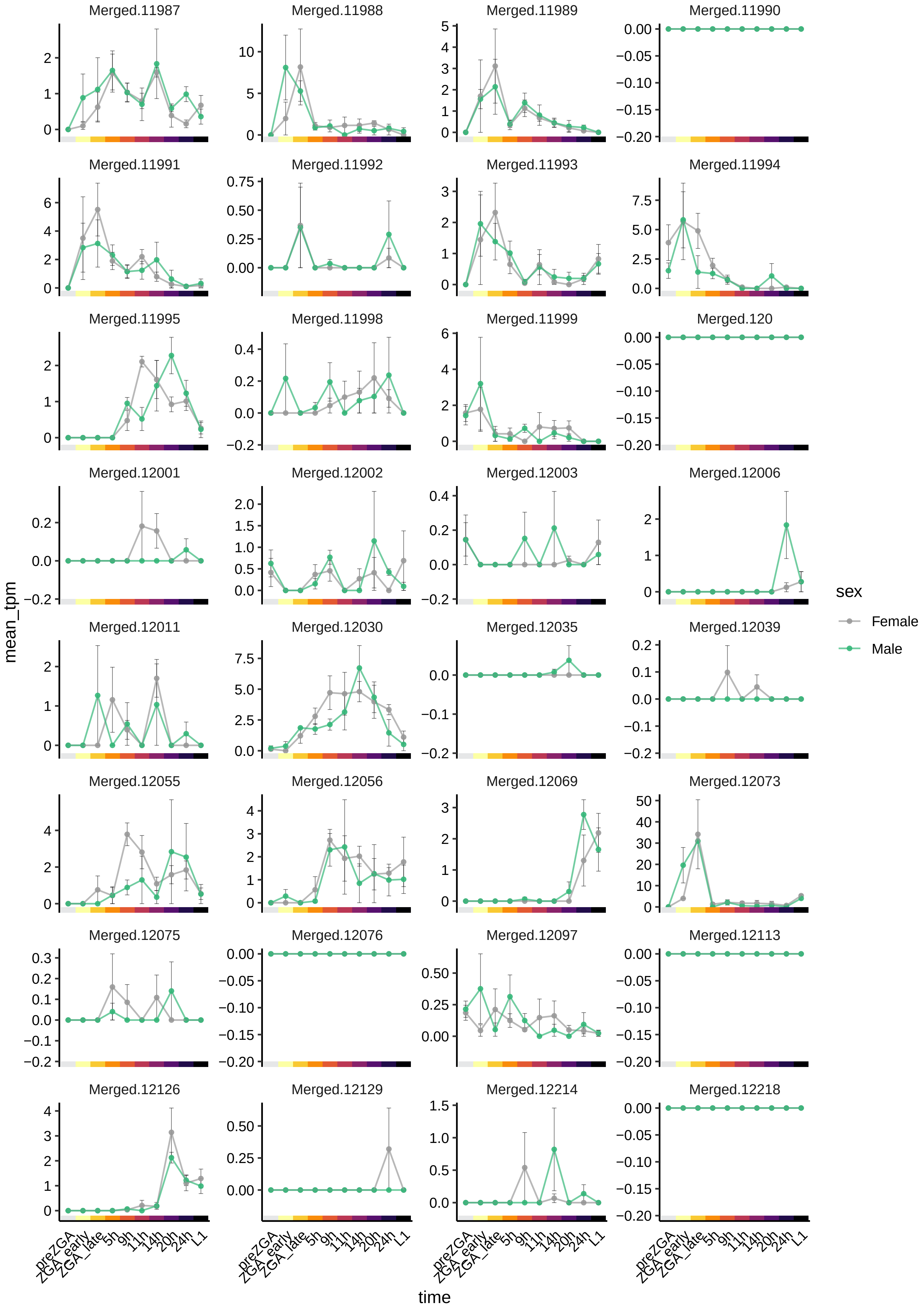
Merged.11987
Merged.11988
Merged.11989
Merged.11990
Merged.11991
Merged.11992
Merged.11993
Merged.11994
Merged.11995
Merged.11998
Merged.11999
Merged.120
Merged.12001
Merged.12002
Merged.12003
Merged.12006
Merged.12011
Merged.12030
Merged.12035
Merged.12039
Merged.12055
Merged.12056
Merged.12069
Merged.12073
Merged.12075
Merged.12076
Merged.12097
Merged.12113
Merged.12126
Merged.12129
Merged.12214
Merged.12218
mean_tpm
time
preZGA
ZGA_early
ZGA_late
5h
9h
11h
14h
20h
24h
L1
sex
Female
Male

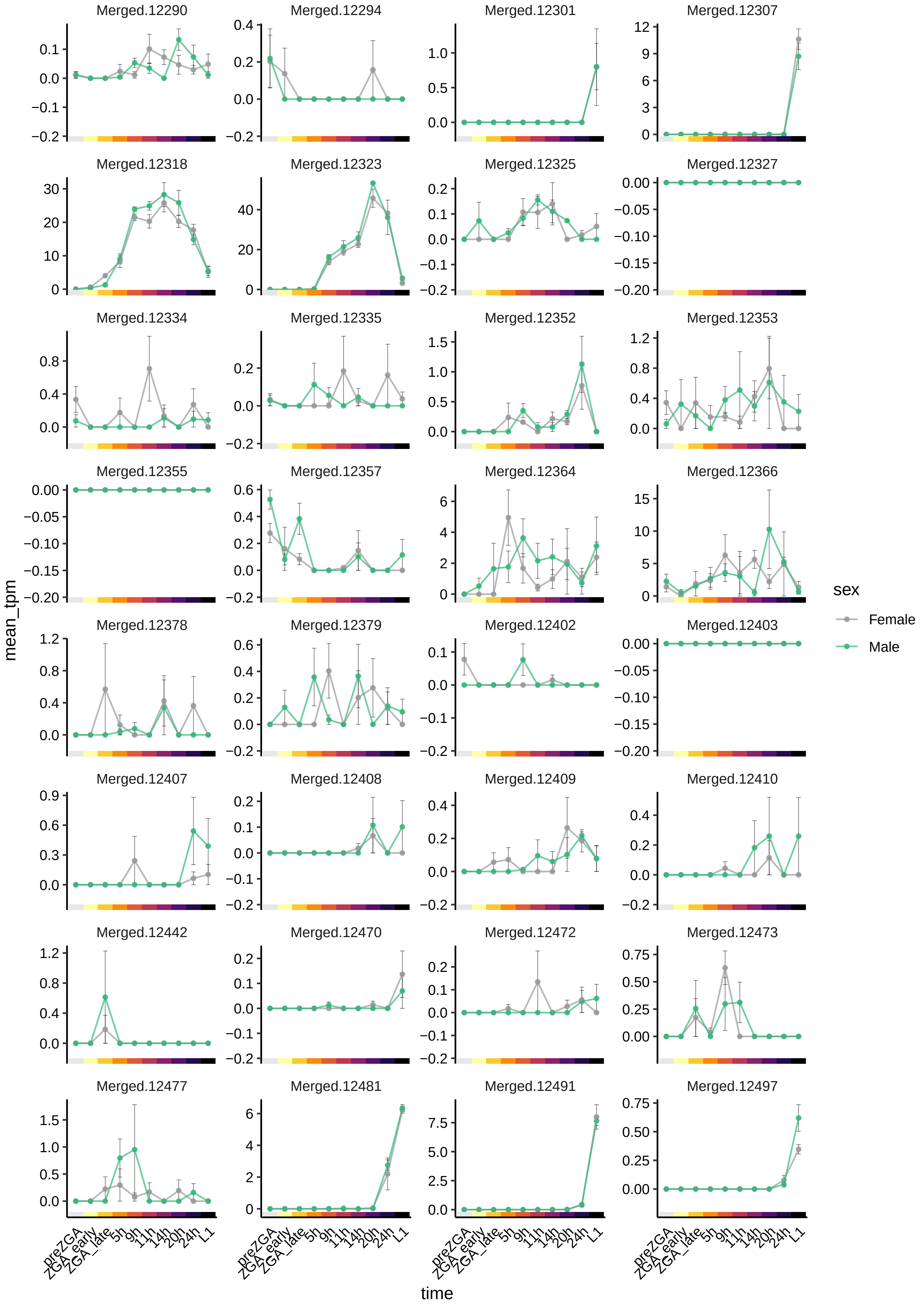

Merged.12290
Merged.12294
Merged.12301
Merged.12307
Merged.12318
Merged.12323
Merged.12325
Merged.12327
Merged.12334
Merged.12335
Merged.12352
Merged.12353
Merged.12355
Merged.12357
Merged.12364
Merged.12366
Merged.12378
Merged.12379
Merged.12402
Merged.12403
Merged.12407
Merged.12408
Merged.12409
Merged.12410
Merged.12442
Merged.12470
Merged.12472
Merged.12473
Merged.12477
Merged.12481
Merged.12491
Merged.12497
mean_tpm
time
preZGA
ZGA_early
ZGA_late
5h
9h
11h
14h
20h
24h
L1
sex
Female
Male

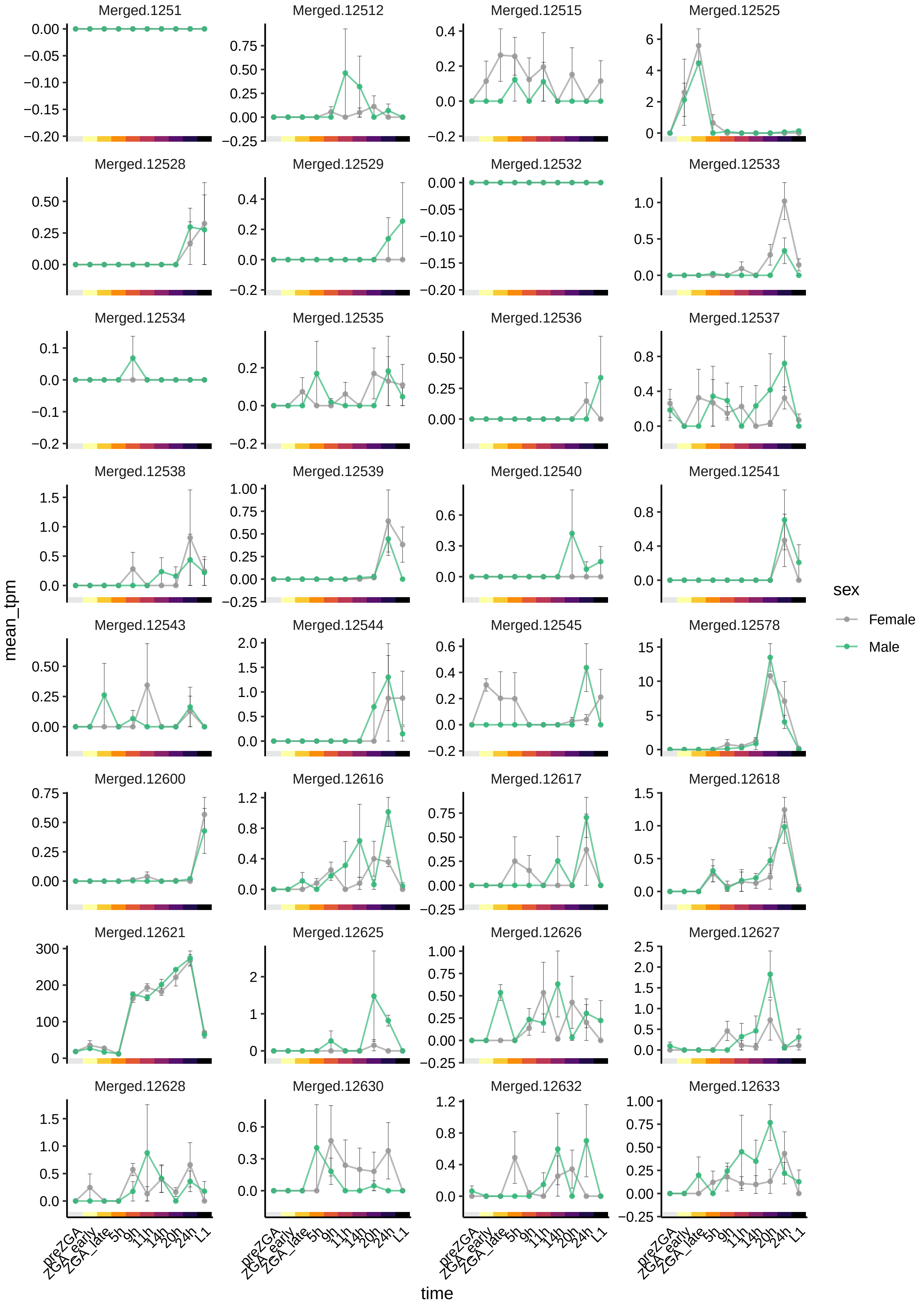

Merged.1251
Merged.12512
Merged.12515
Merged.12525
Merged.12528
Merged.12529
Merged.12532
Merged.12533
Merged.12534
Merged.12535
Merged.12536
Merged.12537
Merged.12538
Merged.12539
Merged.12540
Merged.12541
Merged.12543
Merged.12544
Merged.12545
Merged.12578
Merged.12600
Merged.12616
Merged.12617
Merged.12618
Merged.12621
Merged.12625
Merged.12626
Merged.12627
Merged.12628
Merged.12630
Merged.12632
Merged.12633
mean_tpm
time
preZGA
ZGA_early
ZGA_late
5h
9h
11h
14h
20h
24h
L1
sex
Female
Male

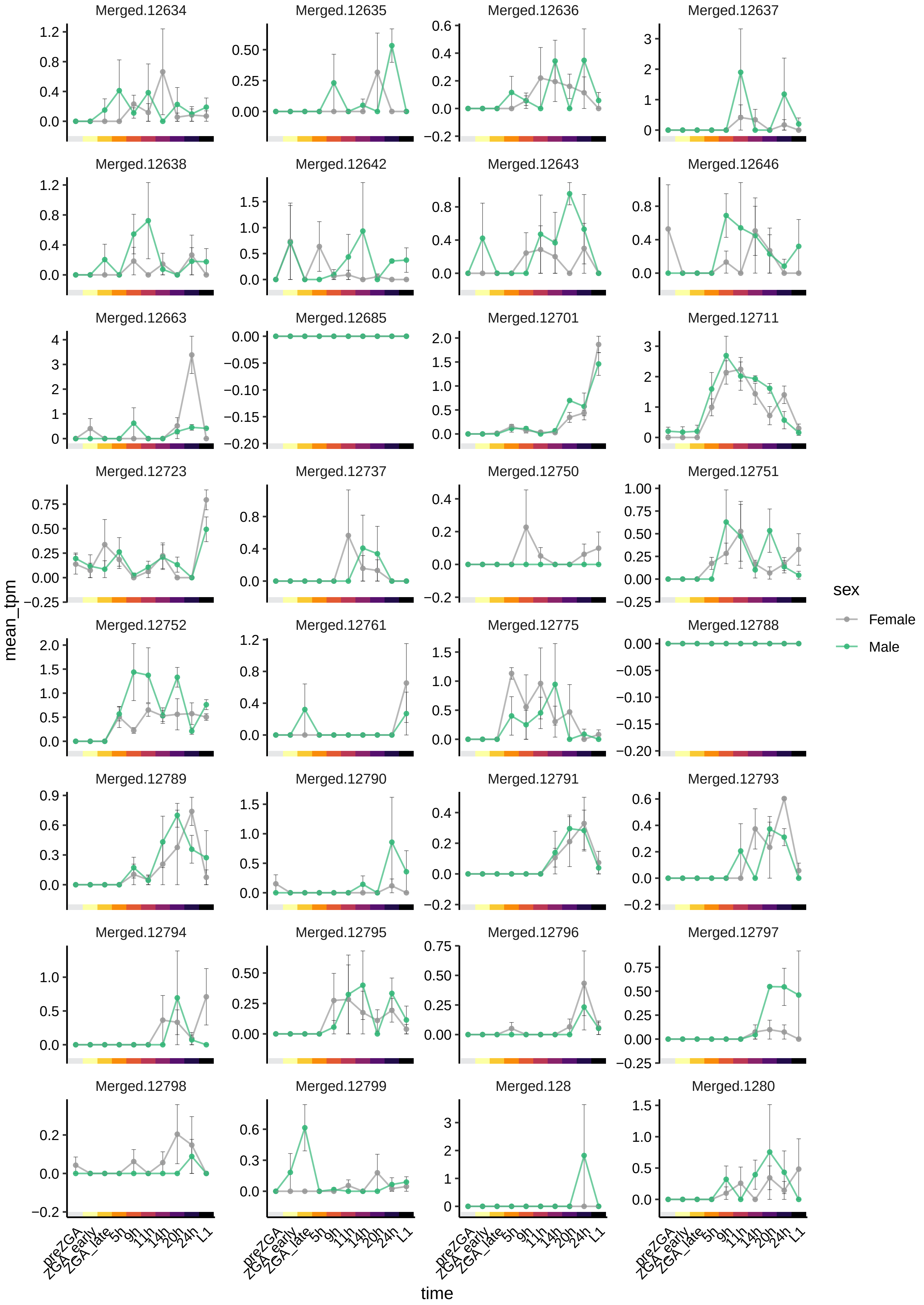
Merged.12634
Merged.12635
Merged.12636
Merged.12637
Merged.12638
Merged.12642
Merged.12643
Merged.12646
Merged.12663
Merged.12685
Merged.12701
Merged.12711
Merged.12723
Merged.12737
Merged.12750
Merged.12751
Merged.12752
Merged.12761
Merged.12775
Merged.12788
Merged.12789
Merged.12790
Merged.12791
Merged.12793
Merged.12794
Merged.12795
Merged.12796
Merged.12797
Merged.12798
Merged.12799
Merged.128
Merged.1280
mean_tpm
time
preZGA
ZGA_early
ZGA_late
5h
9h
11h
14h
20h
24h
L1
sex
Female
Male

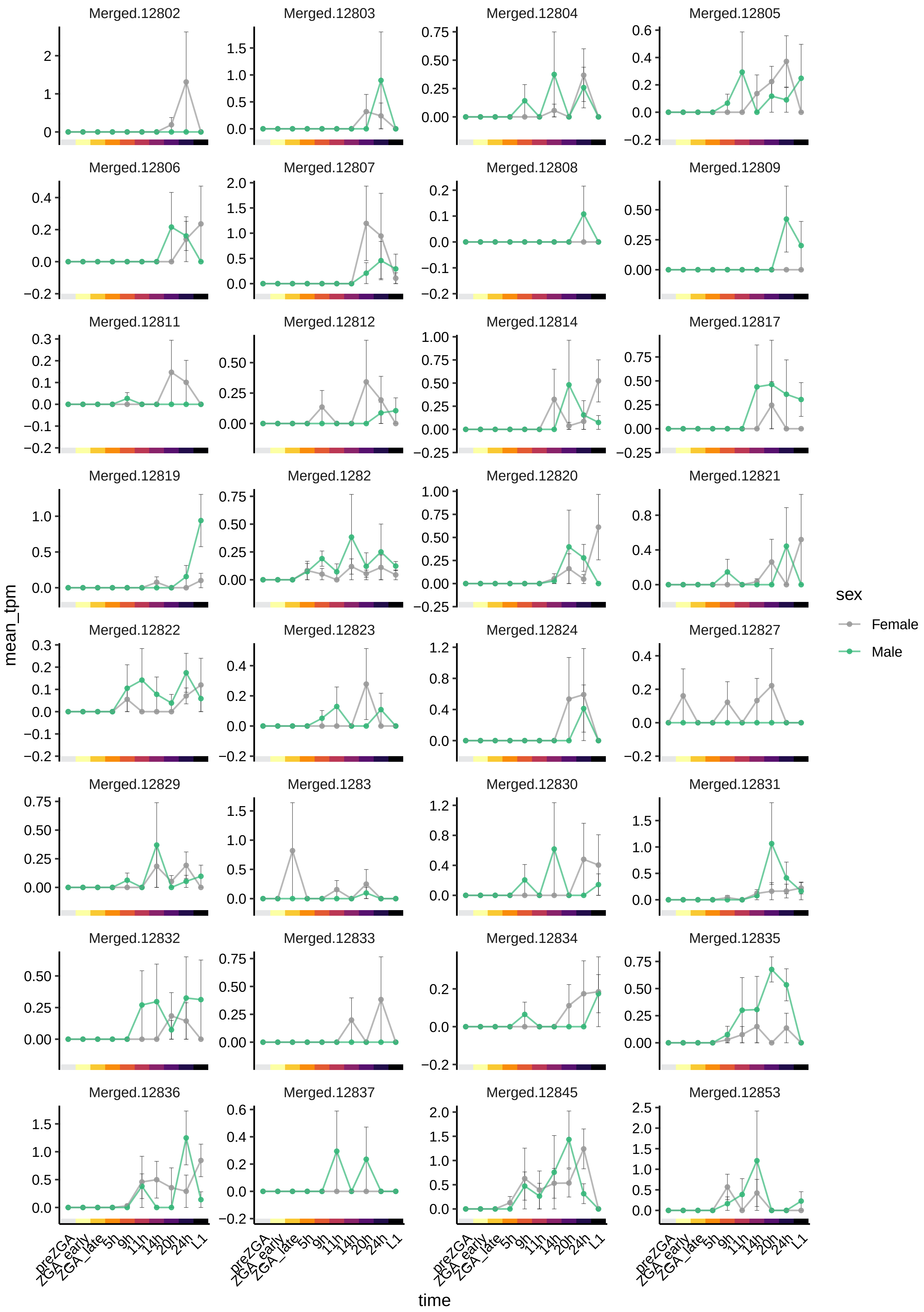

Merged.12802
Merged.12803
Merged.12804
Merged.12805
Merged.12806
Merged.12807
Merged.12808
Merged.12809
Merged.12811
Merged.12812
Merged.12814
Merged.12817
Merged.12819
Merged.1282
Merged.12820
Merged.12821
Merged.12822
Merged.12823
Merged.12824
Merged.12827
Merged.12829
Merged.1283
Merged.12830
Merged.12831
Merged.12832
Merged.12833
Merged.12834
Merged.12835
Merged.12836
Merged.12837
Merged.12845
Merged.12853
mean_tpm
time
preZGA
ZGA_early
ZGA_late
5h
9h
11h
14h
20h
24h
L1
sex
Female
Male

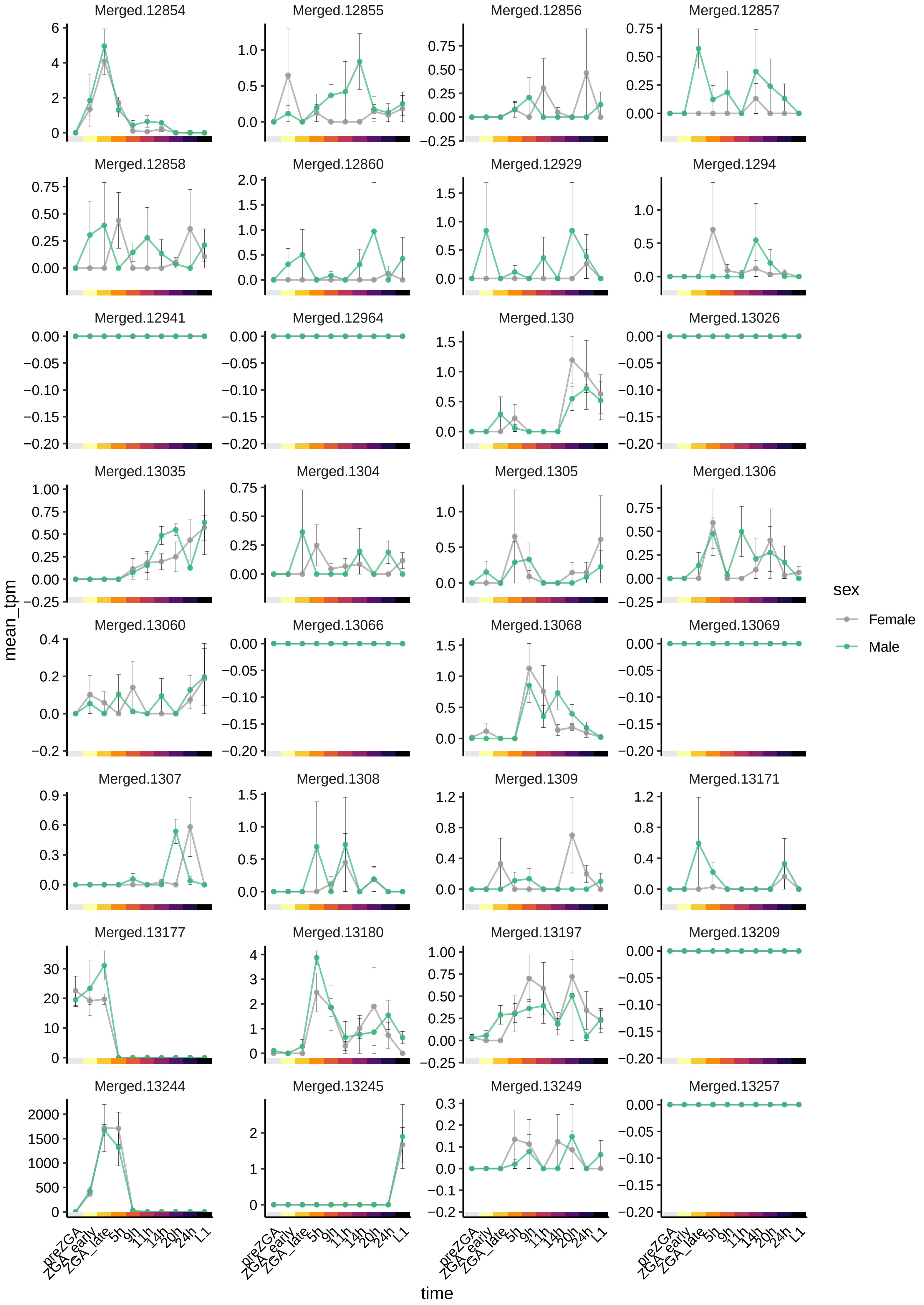

Merged.12854
Merged.12855
Merged.12856
Merged.12857
Merged.12858
Merged.12860
Merged.12929
Merged.1294
Merged.12941
Merged.12964
Merged.130
Merged.13026
Merged.13035
Merged.1304
Merged.1305
Merged.1306
Merged.13060
Merged.13066
Merged.13068
Merged.13069
Merged.1307
Merged.1308
Merged.1309
Merged.13171
Merged.13177
Merged.13180
Merged.13197
Merged.13209
Merged.13244
Merged.13245
Merged.13249
Merged.13257
mean_tpm
time
preZGA
ZGA_early
ZGA_late
5h
9h
11h
14h
20h
24h
L1
sex
Female
Male

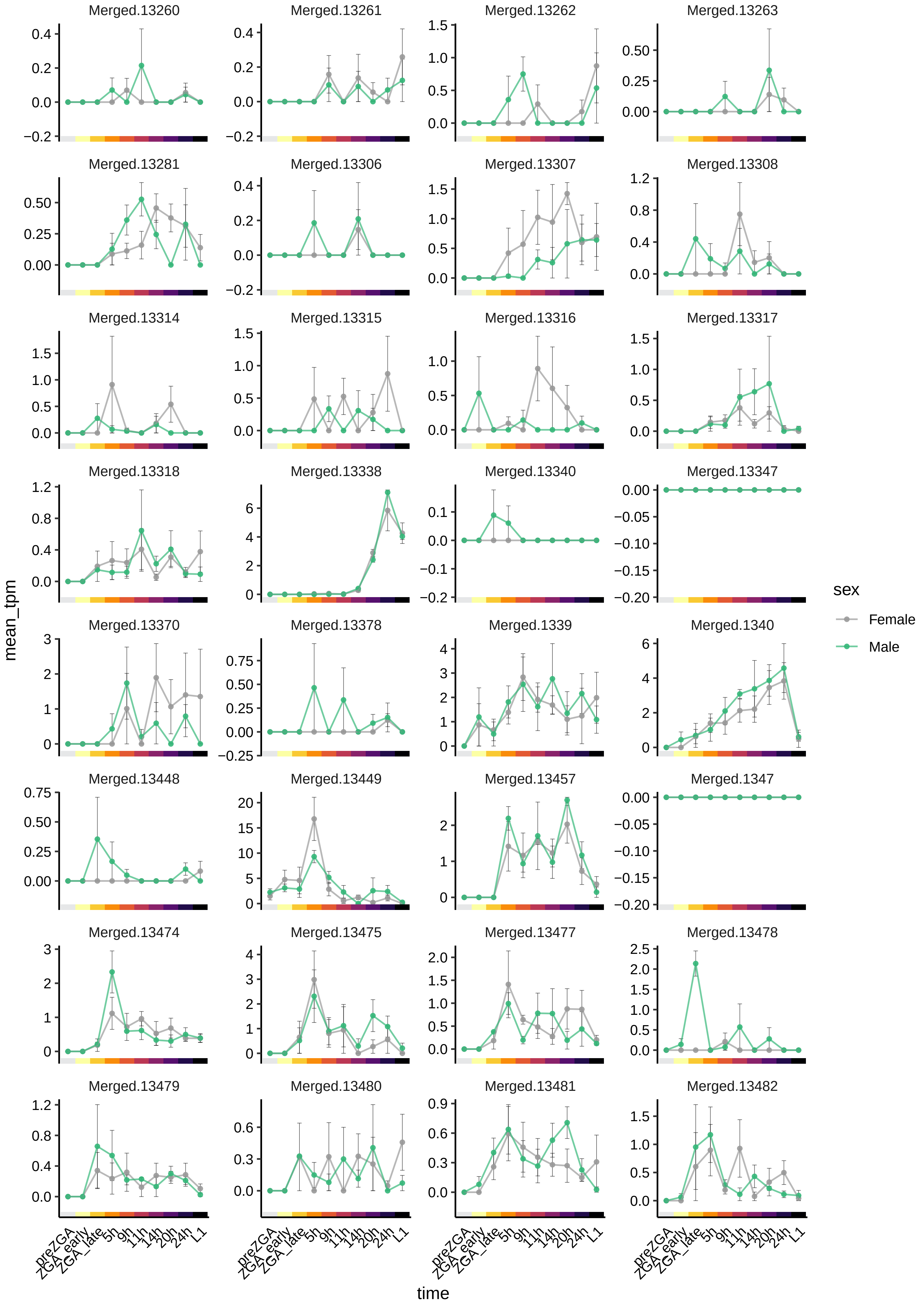
Merged.13260
Merged.13261
Merged.13262
Merged.13263
Merged.13281
Merged.13306
Merged.13307
Merged.13308
Merged.13314
Merged.13315
Merged.13316
Merged.13317
Merged.13318
Merged.13338
Merged.13340
Merged.13347
Merged.13370
Merged.13378
Merged.1339
Merged.1340
Merged.13448
Merged.13449
Merged.13457
Merged.1347
Merged.13474
Merged.13475
Merged.13477
Merged.13478
Merged.13479
Merged.13480
Merged.13481
Merged.13482
mean_tpm
time
preZGA
ZGA_early
ZGA_late
5h
9h
11h
14h
20h
24h
L1
sex
Female
Male

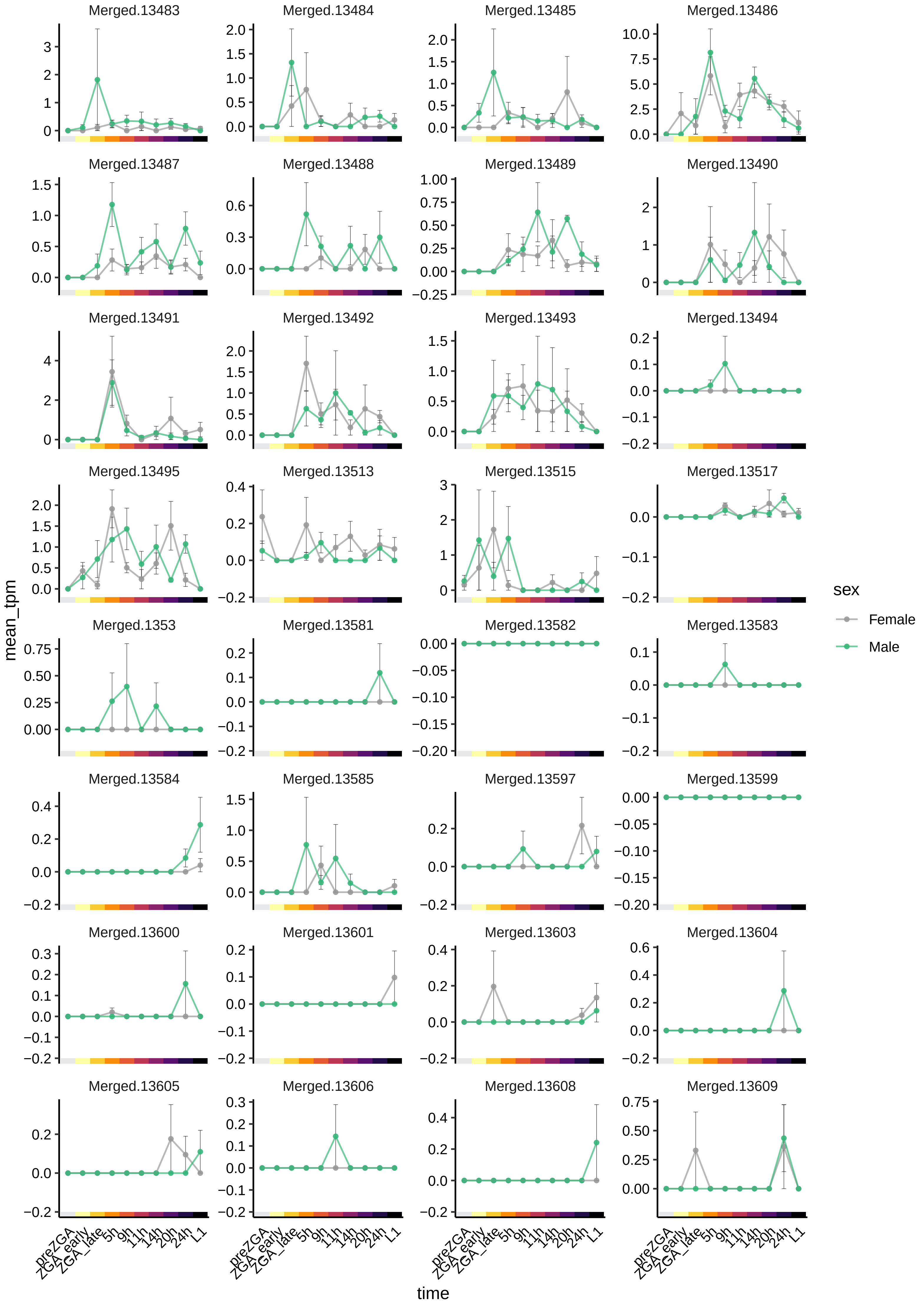

Merged.13483
Merged.13484
Merged.13485
Merged.13486
Merged.13487
Merged.13488
Merged.13489
Merged.13490
Merged.13491
Merged.13492
Merged.13493
Merged.13494
Merged.13495
Merged.13513
Merged.13515
Merged.13517
Merged.1353
Merged.13581
Merged.13582
Merged.13583
Merged.13584
Merged.13585
Merged.13597
Merged.13599
Merged.13600
Merged.13601
Merged.13603
Merged.13604
Merged.13605
Merged.13606
Merged.13608
Merged.13609
mean_tpm
time
preZGA
ZGA_early
ZGA_late
5h
9h
11h
14h
20h
24h
L1
sex
Female
Male

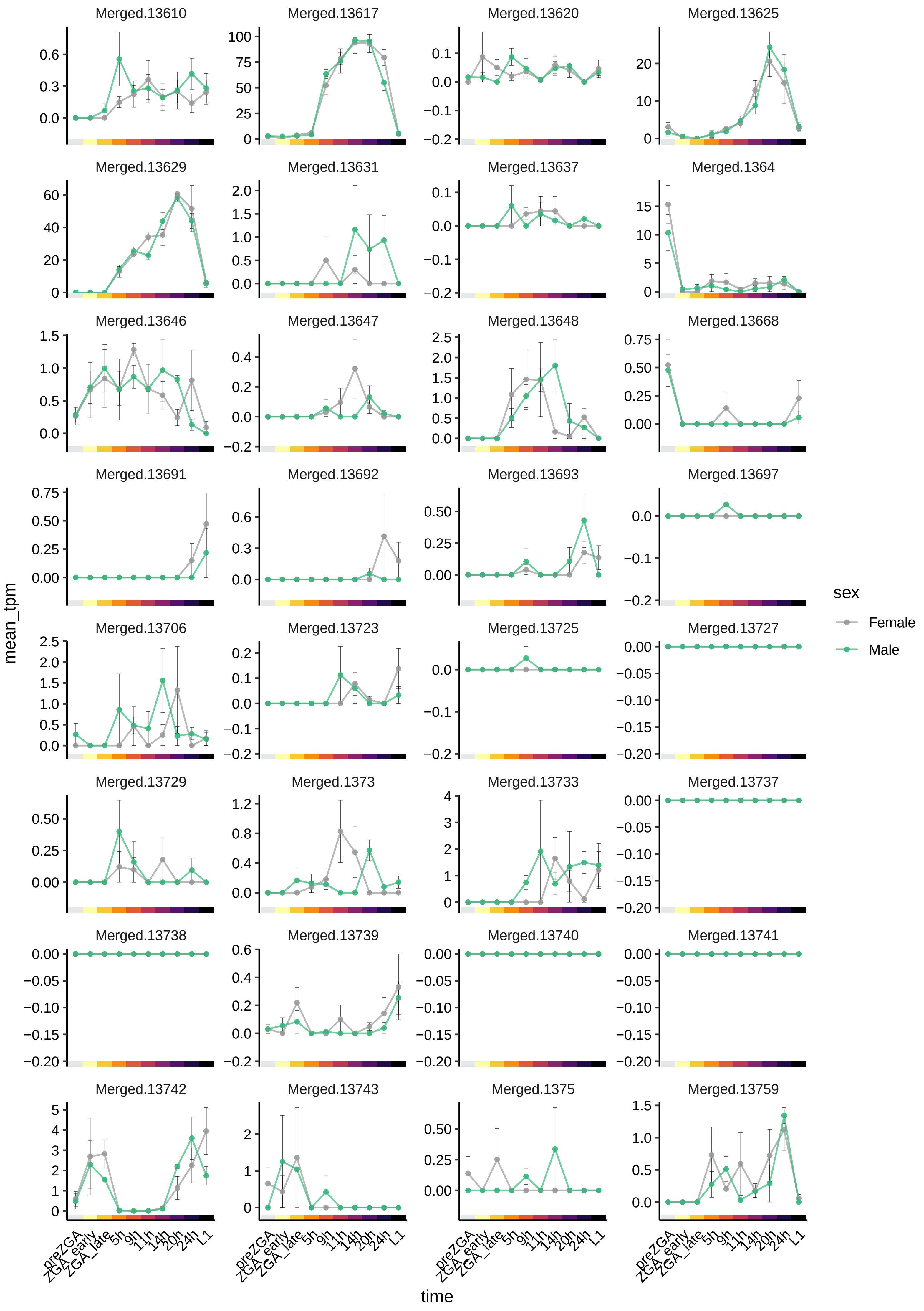

Merged.13610
Merged.13617
Merged.13620
Merged.13625
Merged.13629
Merged.13631
Merged.13637
Merged.1364
Merged.13646
Merged.13647
Merged.13648
Merged.13668
Merged.13691
Merged.13692
Merged.13693
Merged.13697
Merged.13706
Merged.13723
Merged.13725
Merged.13727
Merged.13729
Merged.1373
Merged.13733
Merged.13737
Merged.13738
Merged.13739
Merged.13740
Merged.13741
Merged.13742
Merged.13743
Merged.1375
Merged.13759
mean_tpm
time
preZGA
ZGA_early
ZGA_late
5h
9h
11h
14h
20h
24h
L1
sex
Female
Male

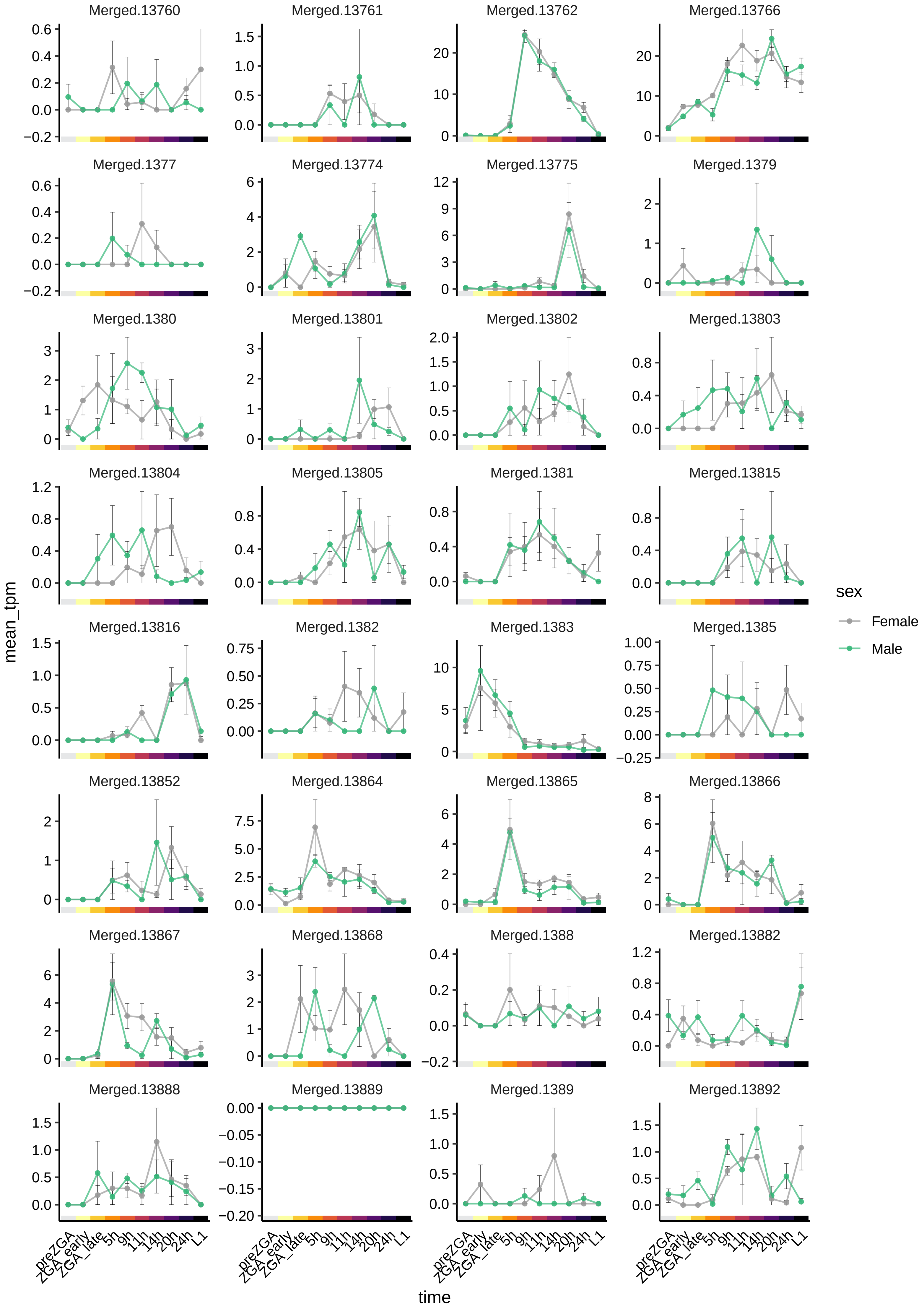

Merged.13760
Merged.13761
Merged.13762
Merged.13766
Merged.1377
Merged.13774
Merged.13775
Merged.1379
Merged.1380
Merged.13801
Merged.13802
Merged.13803
Merged.13804
Merged.13805
Merged.1381
Merged.13815
Merged.13816
Merged.1382
Merged.1383
Merged.1385
Merged.13852
Merged.13864
Merged.13865
Merged.13866
Merged.13867
Merged.13868
Merged.1388
Merged.13882
Merged.13888
Merged.13889
Merged.1389
Merged.13892
mean_tpm
time
preZGA
ZGA_early
ZGA_late
5h
9h
11h
14h
20h
24h
L1
sex
Female
Male

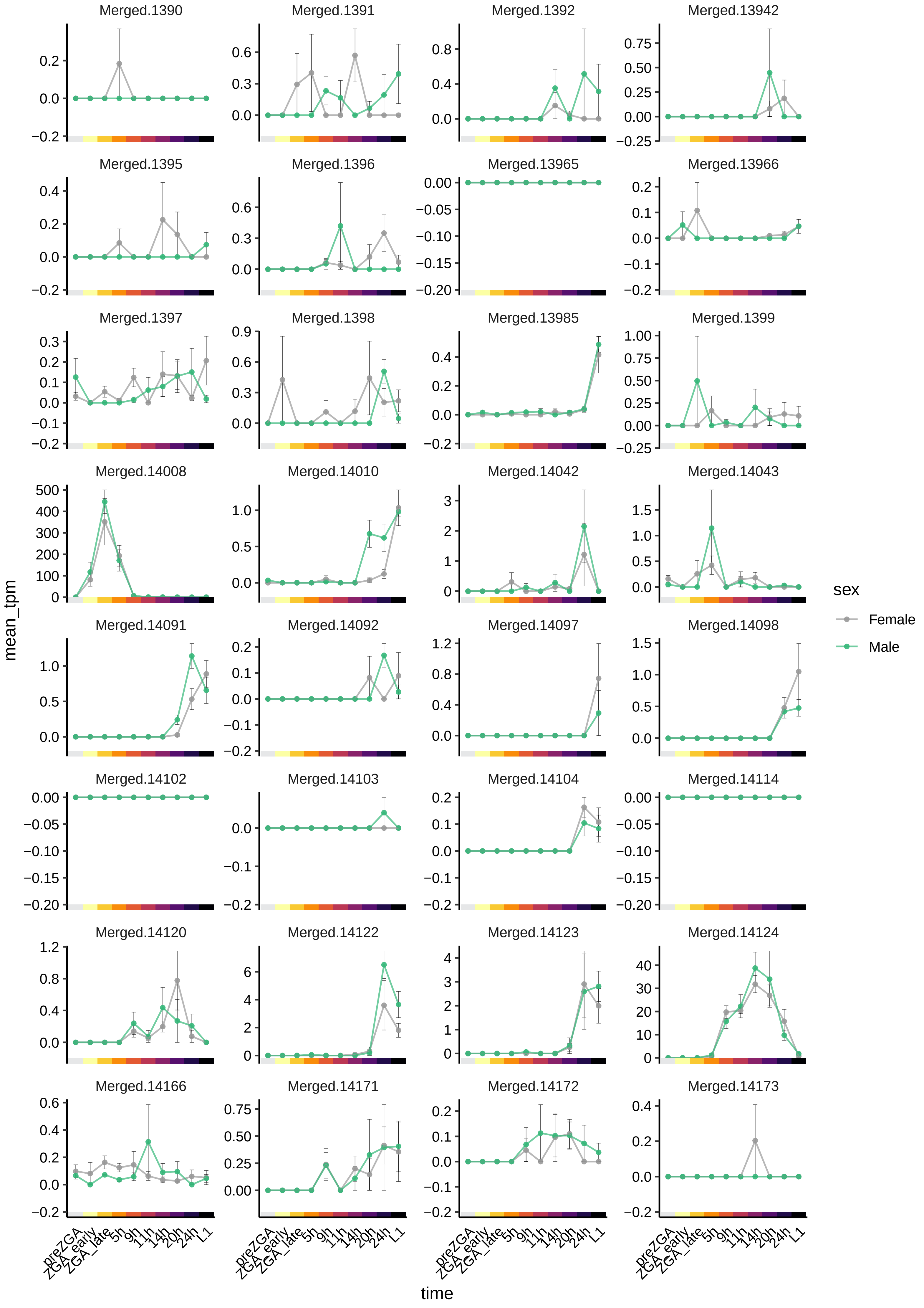

Merged.1390
Merged.1391
Merged.1392
Merged.13942
Merged.1395
Merged.1396
Merged.13965
Merged.13966
Merged.1397
Merged.1398
Merged.13985
Merged.1399
Merged.14008
Merged.14010
Merged.14042
Merged.14043
Merged.14091
Merged.14092
Merged.14097
Merged.14098
Merged.14102
Merged.14103
Merged.14104
Merged.14114
Merged.14120
Merged.14122
Merged.14123
Merged.14124
Merged.14166
Merged.14171
Merged.14172
Merged.14173
mean_tpm
time
preZGA
ZGA_early
ZGA_late
5h
9h
11h
14h
20h
24h
L1
sex
Female
Male

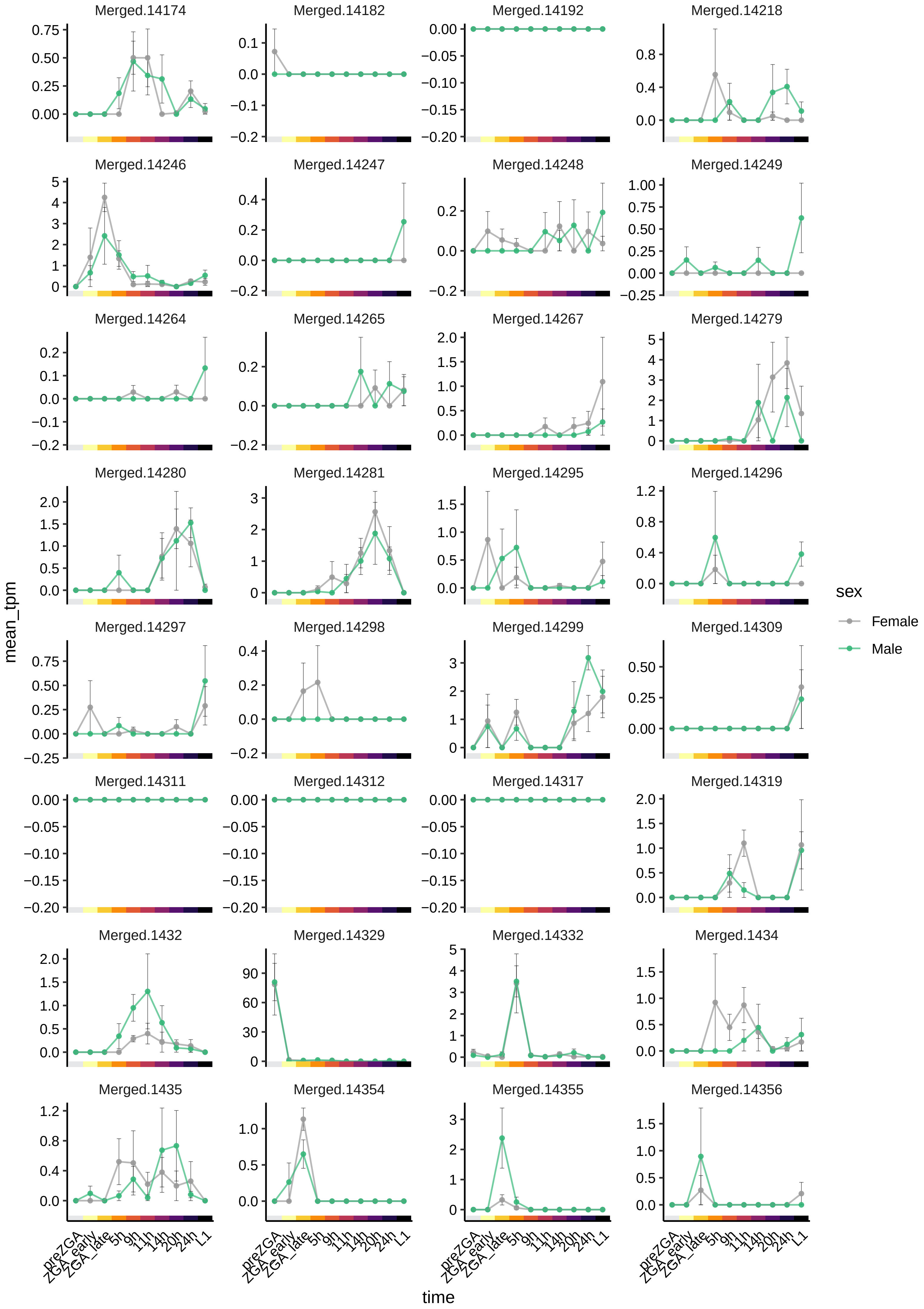

Merged.14174
Merged.14182
Merged.14192
Merged.14218
Merged.14246
Merged.14247
Merged.14248
Merged.14249
Merged.14264
Merged.14265
Merged.14267
Merged.14279
Merged.14280
Merged.14281
Merged.14295
Merged.14296
Merged.14297
Merged.14298
Merged.14299
Merged.14309
Merged.14311
Merged.14312
Merged.14317
Merged.14319
Merged.1432
Merged.14329
Merged.14332
Merged.1434
Merged.1435
Merged.14354
Merged.14355
Merged.14356
mean_tpm
time
preZGA
ZGA_early
ZGA_late
5h
9h
11h
14h
20h
24h
L1
sex
Female
Male

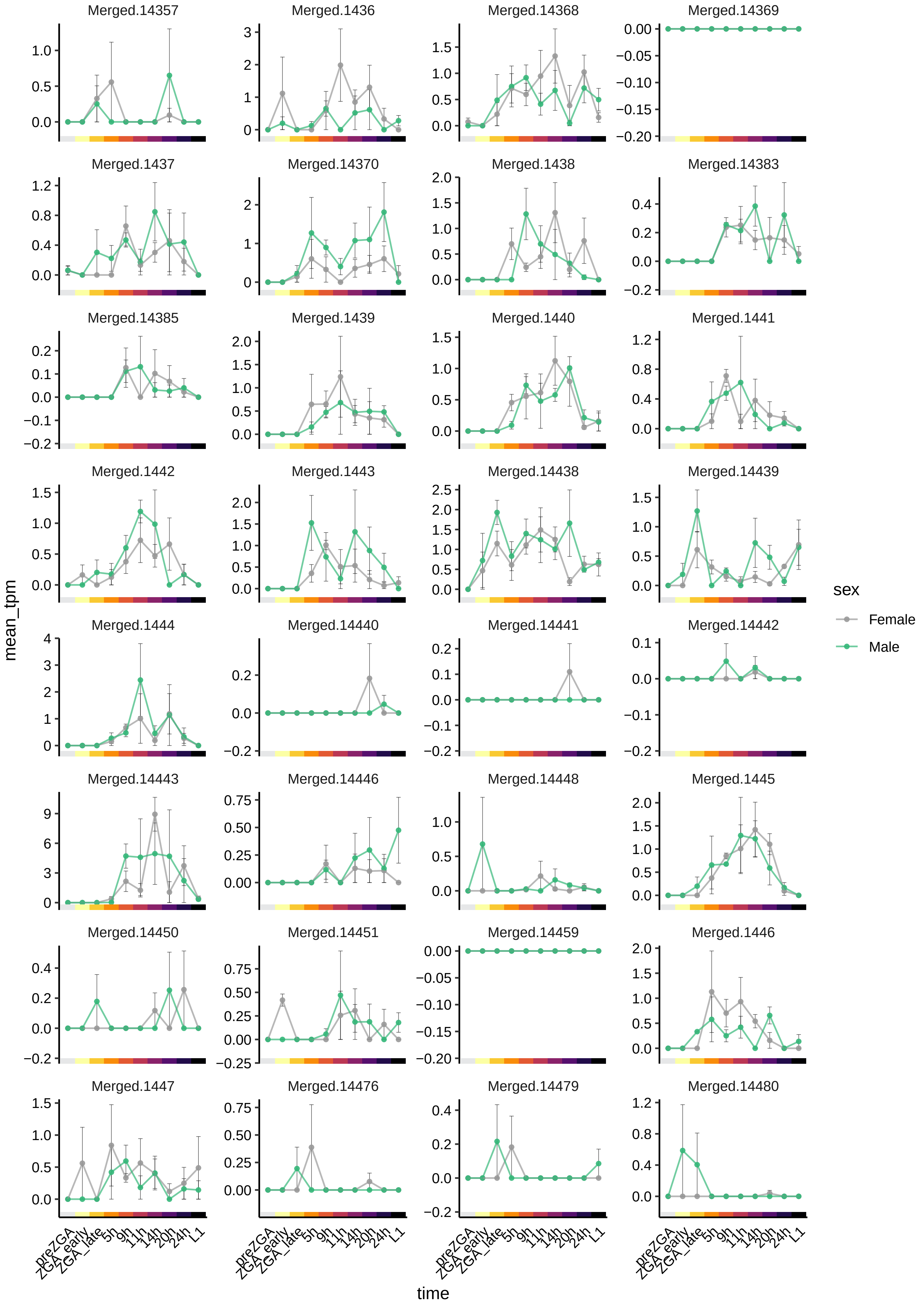

Merged.14357
Merged.1436
Merged.14368
Merged.14369
Merged.1437
Merged.14370
Merged.1438
Merged.14383
Merged.14385
Merged.1439
Merged.1440
Merged.1441
Merged.1442
Merged.1443
Merged.14438
Merged.14439
Merged.1444
Merged.14440
Merged.14441
Merged.14442
Merged.14443
Merged.14446
Merged.14448
Merged.1445
Merged.14450
Merged.14451
Merged.14459
Merged.1446
Merged.1447
Merged.14476
Merged.14479
Merged.14480
mean_tpm
time
preZGA
ZGA_early
ZGA_late
5h
9h
11h
14h
20h
24h
L1
sex
Female
Male

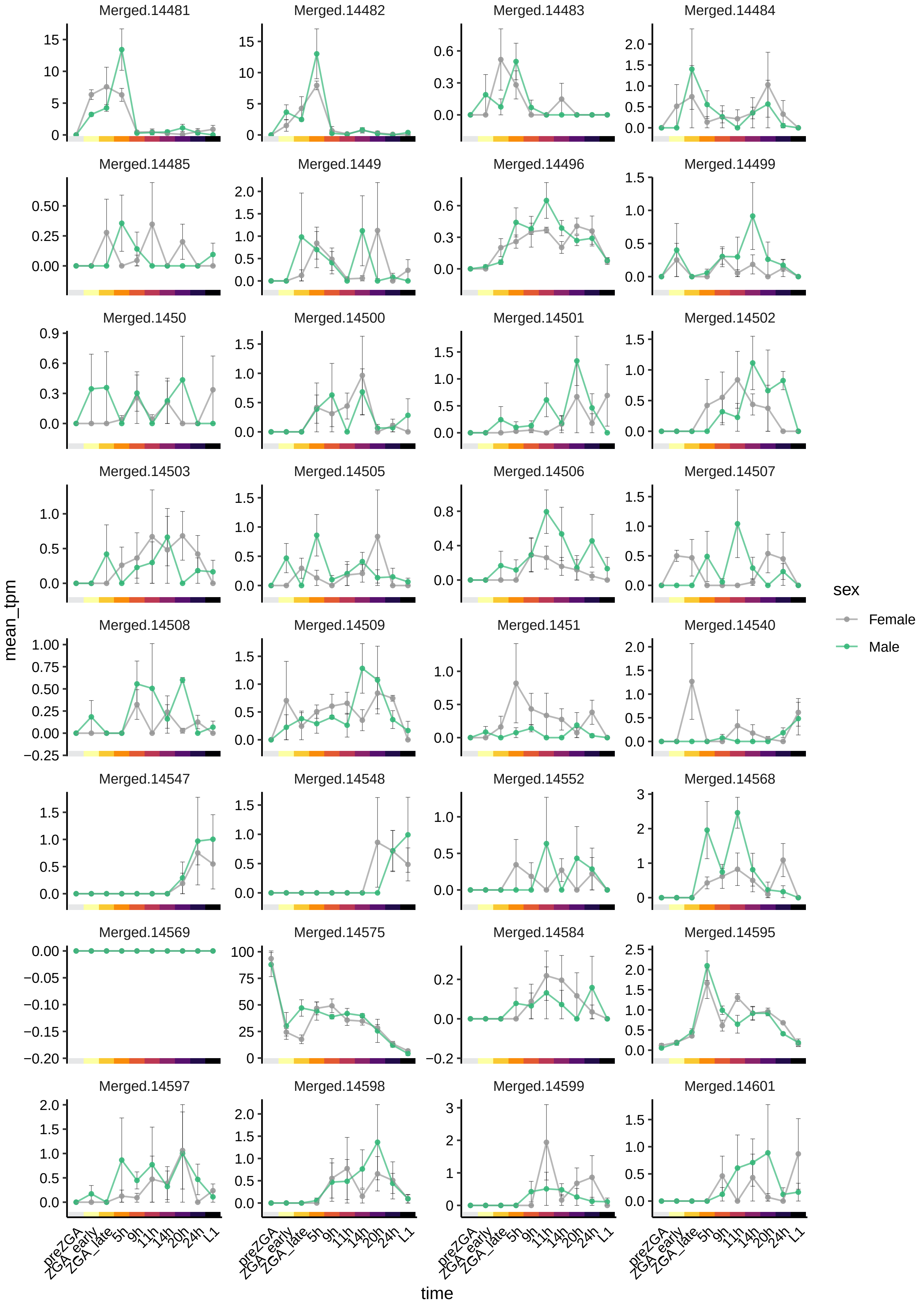

Merged.14481
Merged.14482
Merged.14483
Merged.14484
Merged.14485
Merged.1449
Merged.14496
Merged.14499
Merged.1450
Merged.14500
Merged.14501
Merged.14502
Merged.14503
Merged.14505
Merged.14506
Merged.14507
Merged.14508
Merged.14509
Merged.1451
Merged.14540
Merged.14547
Merged.14548
Merged.14552
Merged.14568
Merged.14569
Merged.14575
Merged.14584
Merged.14595
Merged.14597
Merged.14598
Merged.14599
Merged.14601
mean_tpm
time
preZGA
ZGA_early
ZGA_late
5h
9h
11h
14h
20h
24h
L1
sex
Female
Male

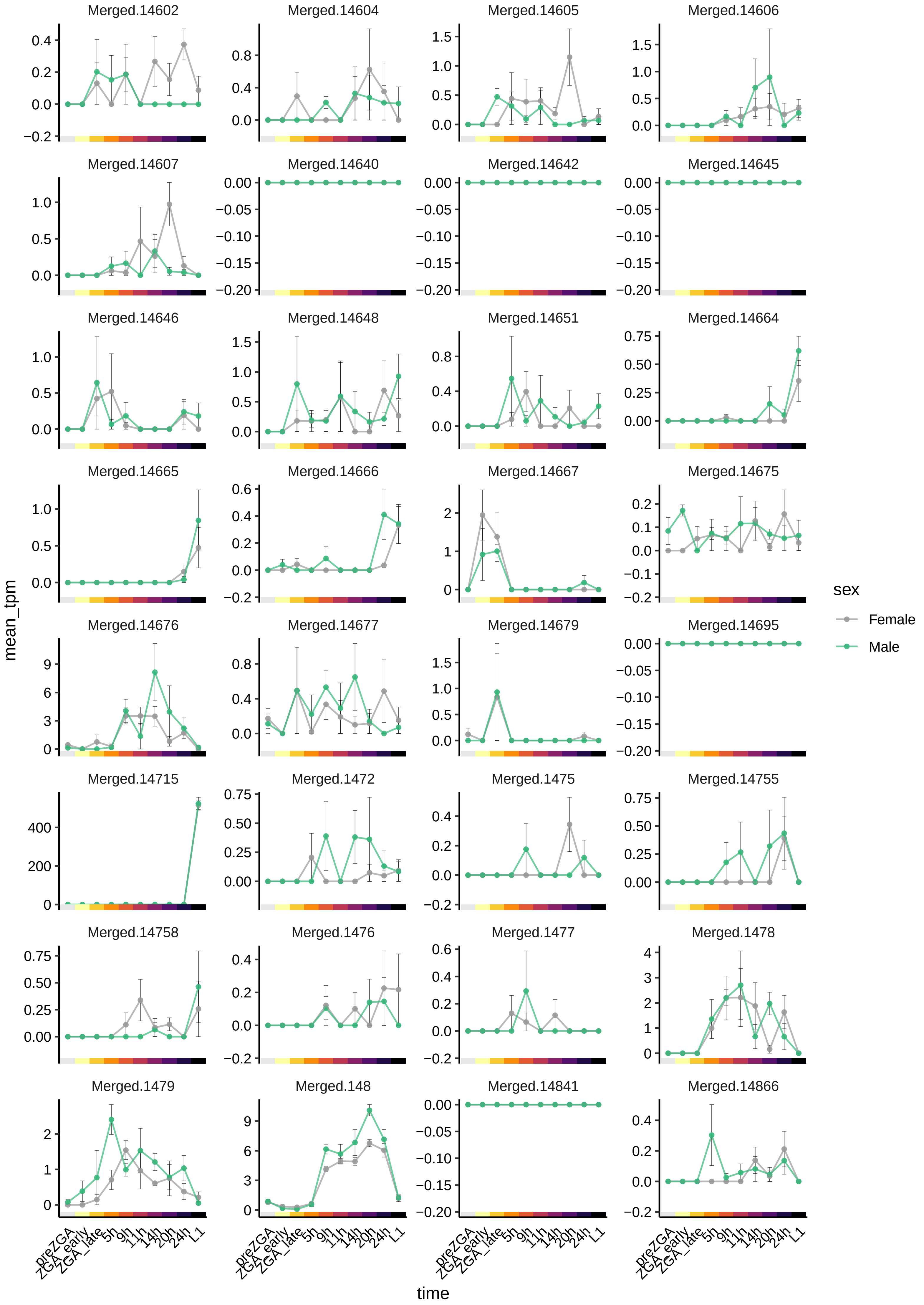
Merged.14602
Merged.14604
Merged.14605
Merged.14606
Merged.14607
Merged.14640
Merged.14642
Merged.14645
Merged.14646
Merged.14648
Merged.14651
Merged.14664
Merged.14665
Merged.14666
Merged.14667
Merged.14675
Merged.14676
Merged.14677
Merged.14679
Merged.14695
Merged.14715
Merged.1472
Merged.1475
Merged.14755
Merged.14758
Merged.1476
Merged.1477
Merged.1478
Merged.1479
Merged.148
Merged.14841
Merged.14866
mean_tpm
time
preZGA
ZGA_early
ZGA_late
5h
9h
11h
14h
20h
24h
L1
sex
Female
Male

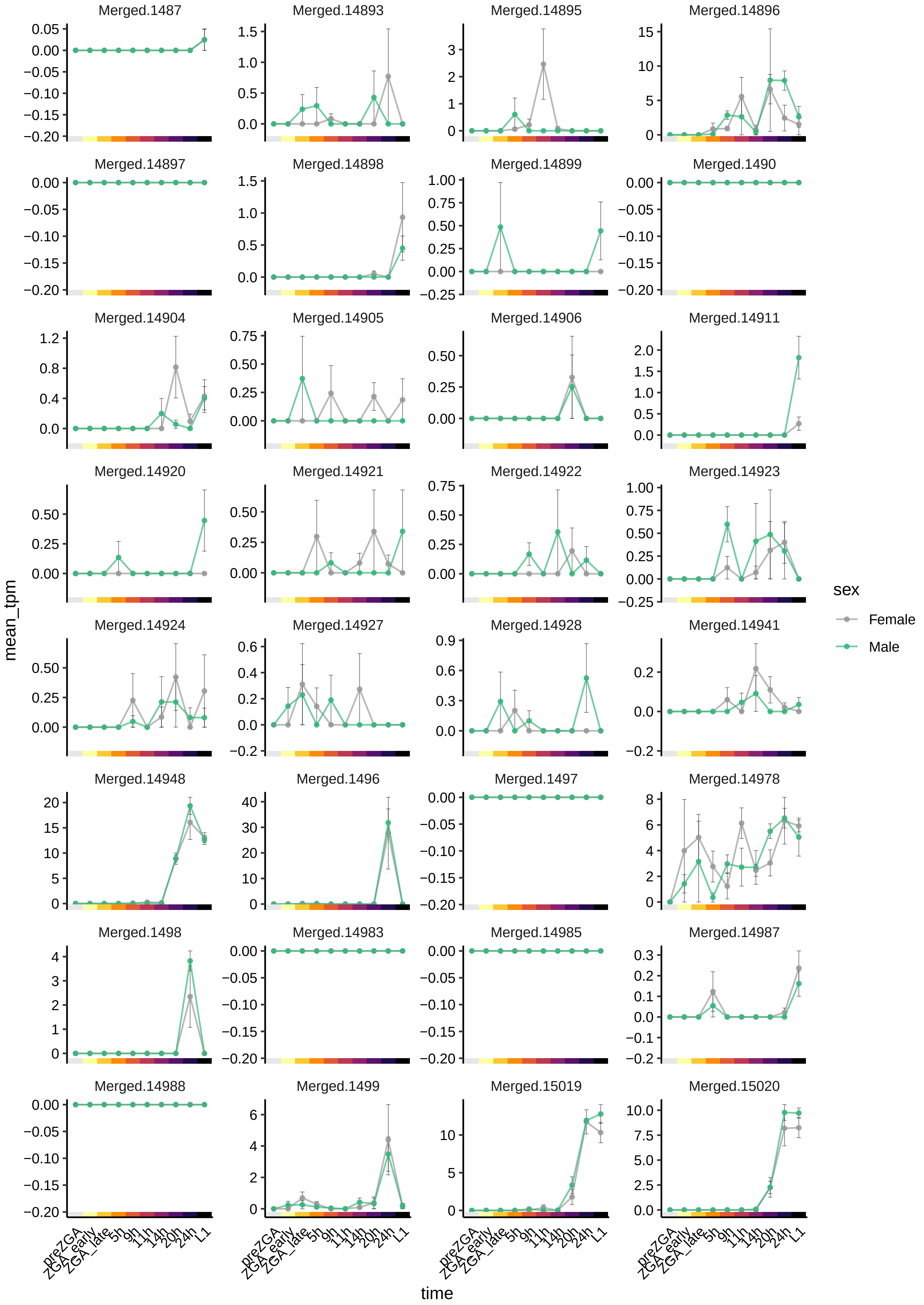
Merged.1487
Merged.14893
Merged.14895
Merged.14896
Merged.14897
Merged.14898
Merged.14899
Merged.1490
Merged.14904
Merged.14905
Merged.14906
Merged.14911
Merged.14920
Merged.14921
Merged.14922
Merged.14923
Merged.14924
Merged.14927
Merged.14928
Merged.14941
Merged.14948
Merged.1496
Merged.1497
Merged.14978
Merged.1498
Merged.14983
Merged.14985
Merged.14987
Merged.14988
Merged.1499
Merged.15019
Merged.15020
mean_tpm
time
preZGA
ZGA_early
ZGA_late
5h
9h
11h
14h
20h
24h
L1
sex
Female
Male

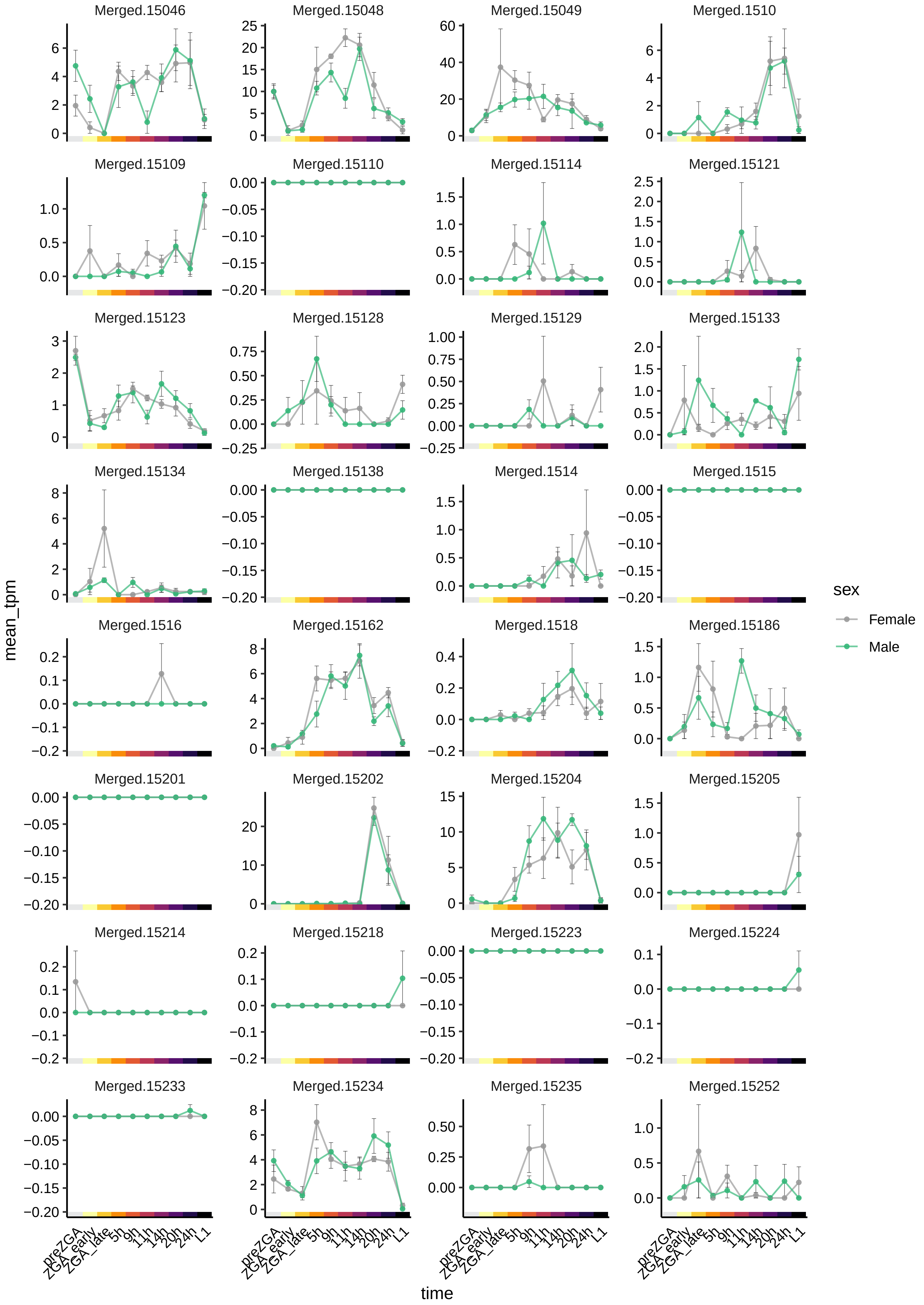
Merged.15046
Merged.15048
Merged.15049
Merged.1510
Merged.15109
Merged.15110
Merged.15114
Merged.15121
Merged.15123
Merged.15128
Merged.15129
Merged.15133
Merged.15134
Merged.15138
Merged.1514
Merged.1515
Merged.1516
Merged.15162
Merged.1518
Merged.15186
Merged.15201
Merged.15202
Merged.15204
Merged.15205
Merged.15214
Merged.15218
Merged.15223
Merged.15224
Merged.15233
Merged.15234
Merged.15235
Merged.15252
mean_tpm
time
preZGA
ZGA_early
ZGA_late
5h
9h
11h
14h
20h
24h
L1
sex
Female
Male

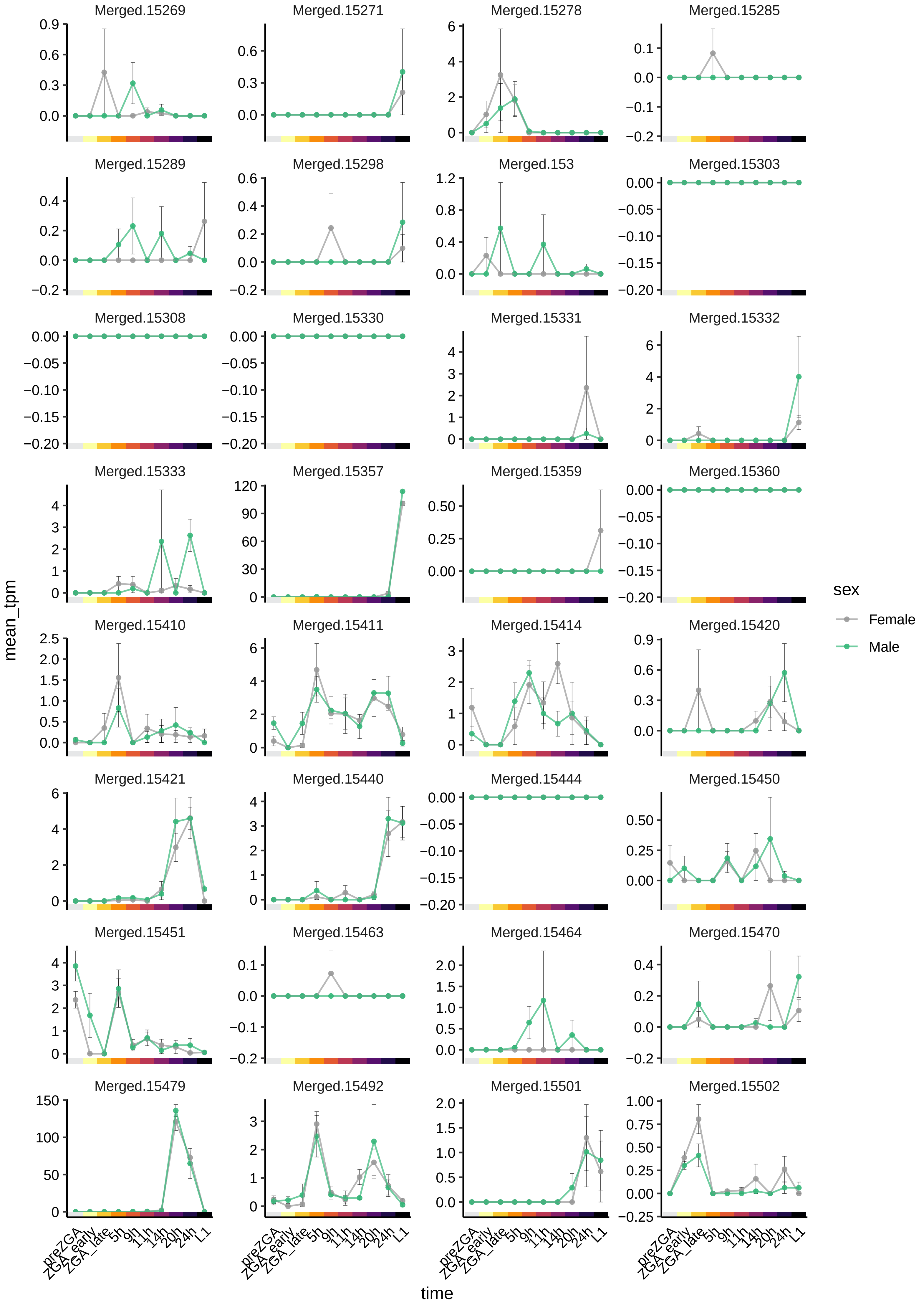

Merged.15269
Merged.15271
Merged.15278
Merged.15285
Merged.15289
Merged.15298
Merged.153
Merged.15303
Merged.15308
Merged.15330
Merged.15331
Merged.15332
Merged.15333
Merged.15357
Merged.15359
Merged.15360
Merged.15410
Merged.15411
Merged.15414
Merged.15420
Merged.15421
Merged.15440
Merged.15444
Merged.15450
Merged.15451
Merged.15463
Merged.15464
Merged.15470
Merged.15479
Merged.15492
Merged.15501
Merged.15502
mean_tpm
time
preZGA
ZGA_early
ZGA_late
5h
9h
11h
14h
20h
24h
L1
sex
Female
Male

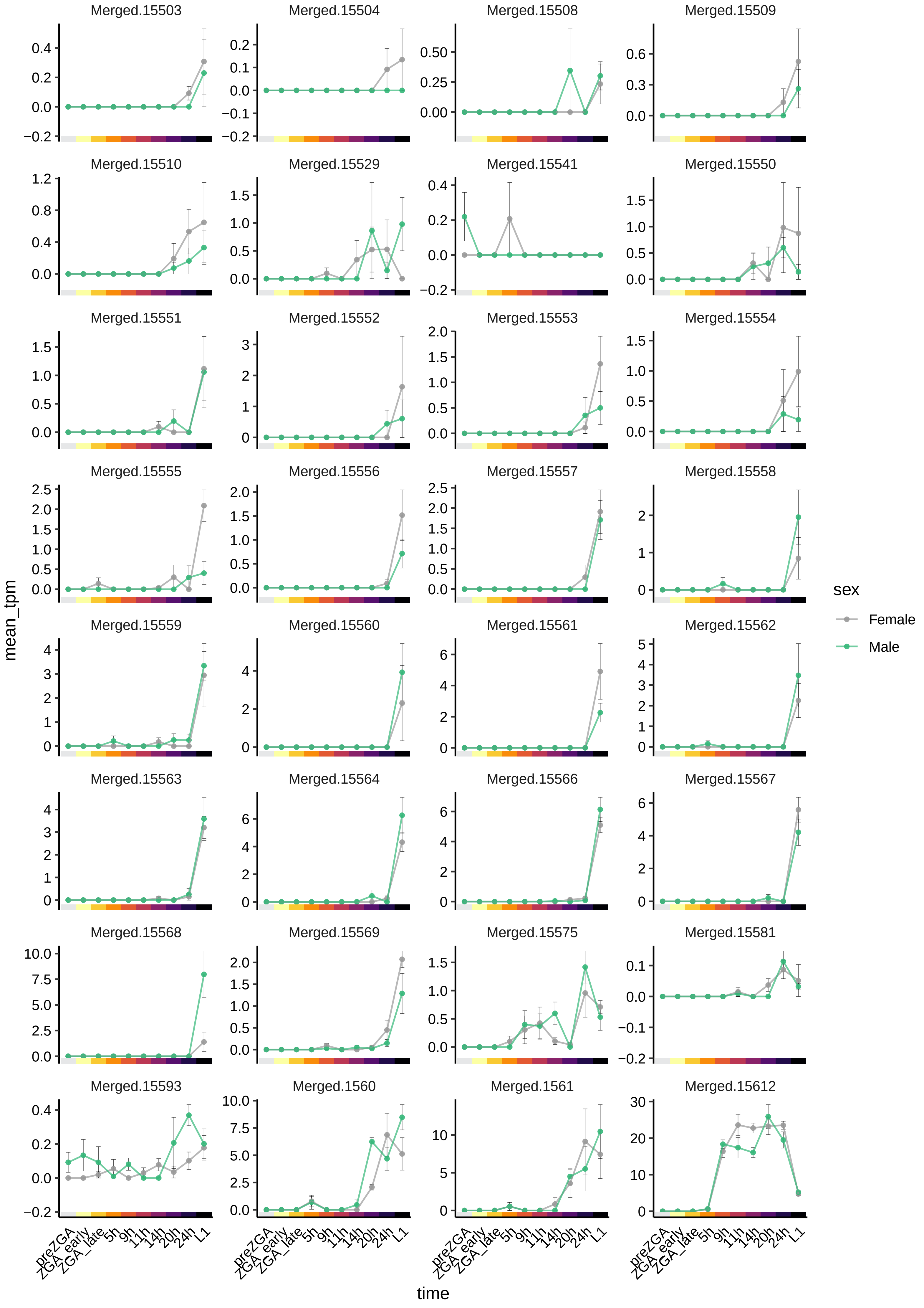
Merged.15503
Merged.15504
Merged.15508
Merged.15509
Merged.15510
Merged.15529
Merged.15541
Merged.15550
Merged.15551
Merged.15552
Merged.15553
Merged.15554
Merged.15555
Merged.15556
Merged.15557
Merged.15558
Merged.15559
Merged.15560
Merged.15561
Merged.15562
Merged.15563
Merged.15564
Merged.15566
Merged.15567
Merged.15568
Merged.15569
Merged.15575
Merged.15581
Merged.15593
Merged.1560
Merged.1561
Merged.15612
mean_tpm
time
preZGA
ZGA_early
ZGA_late
5h
9h
11h
14h
20h
24h
L1
sex
Female
Male

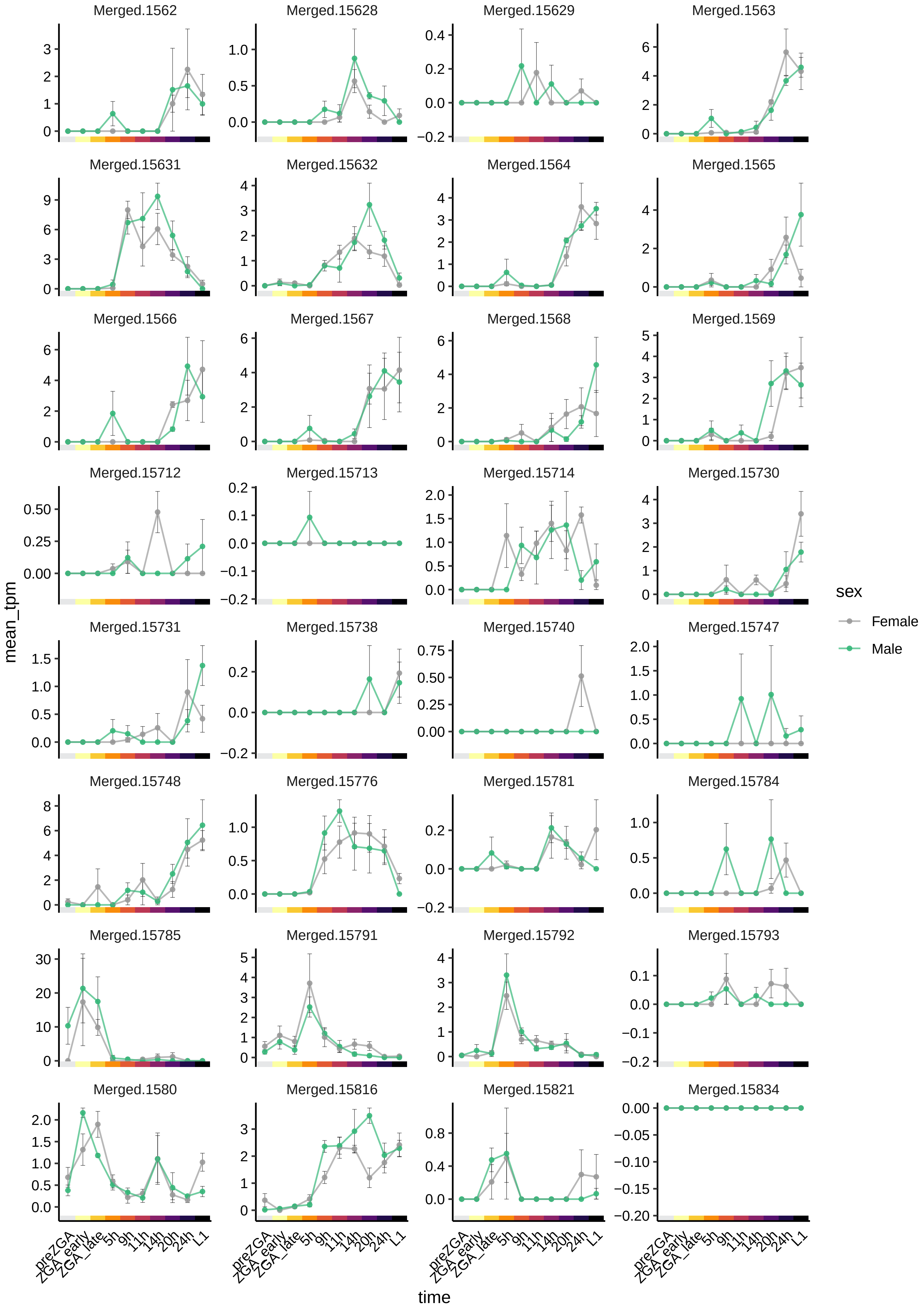

Merged.1562
Merged.15628
Merged.15629
Merged.1563
Merged.15631
Merged.15632
Merged.1564
Merged.1565
Merged.1566
Merged.1567
Merged.1568
Merged.1569
Merged.15712
Merged.15713
Merged.15714
Merged.15730
Merged.15731
Merged.15738
Merged.15740
Merged.15747
Merged.15748
Merged.15776
Merged.15781
Merged.15784
Merged.15785
Merged.15791
Merged.15792
Merged.15793
Merged.1580
Merged.15816
Merged.15821
Merged.15834
mean_tpm
time
preZGA
ZGA_early
ZGA_late
5h
9h
11h
14h
20h
24h
L1
sex
Female
Male

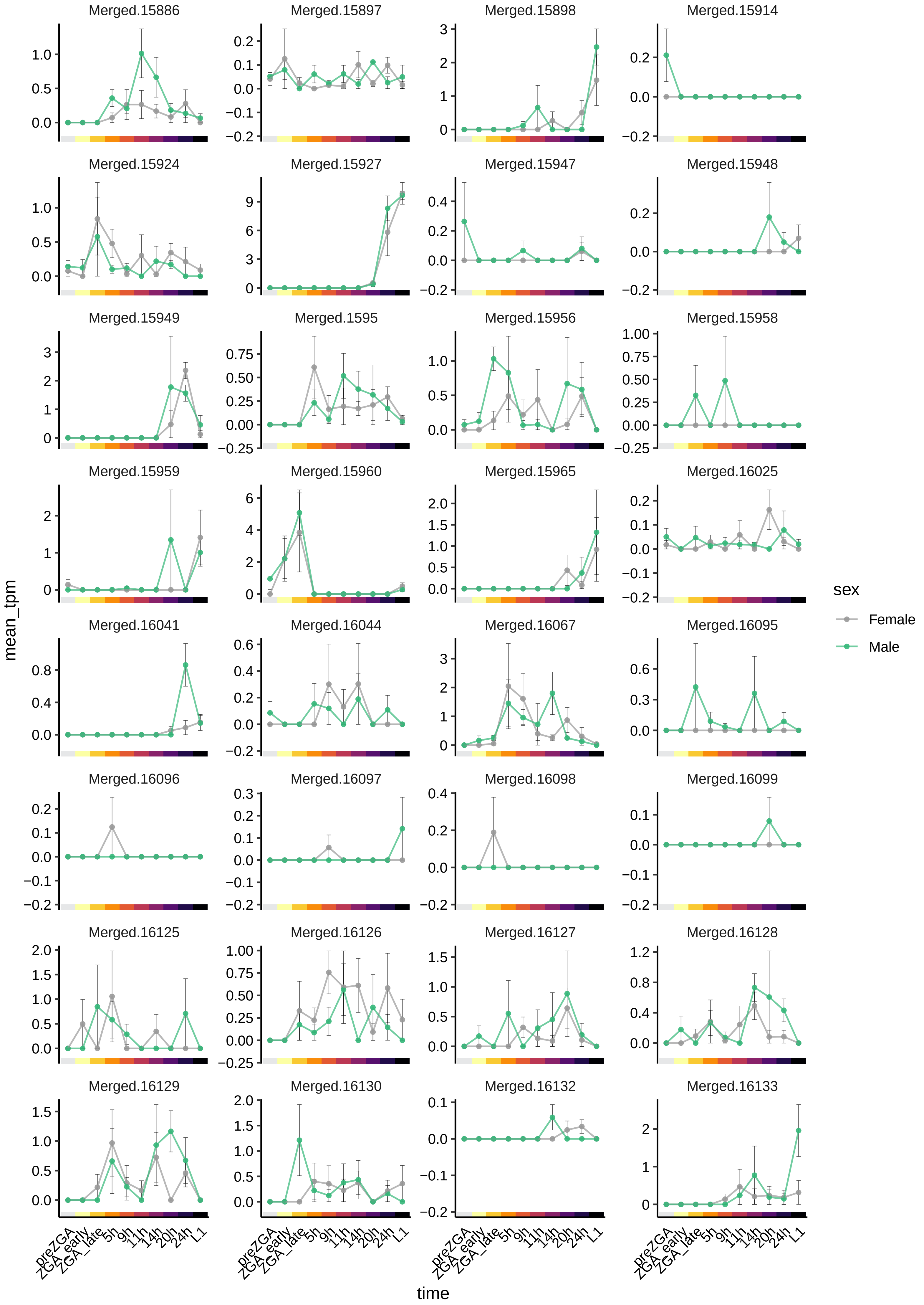

Merged.15886
Merged.15897
Merged.15898
Merged.15914
Merged.15924
Merged.15927
Merged.15947
Merged.15948
Merged.15949
Merged.1595
Merged.15956
Merged.15958
Merged.15959
Merged.15960
Merged.15965
Merged.16025
Merged.16041
Merged.16044
Merged.16067
Merged.16095
Merged.16096
Merged.16097
Merged.16098
Merged.16099
Merged.16125
Merged.16126
Merged.16127
Merged.16128
Merged.16129
Merged.16130
Merged.16132
Merged.16133
mean_tpm
time
preZGA
ZGA_early
ZGA_late
5h
9h
11h
14h
20h
24h
L1
sex
Female
Male

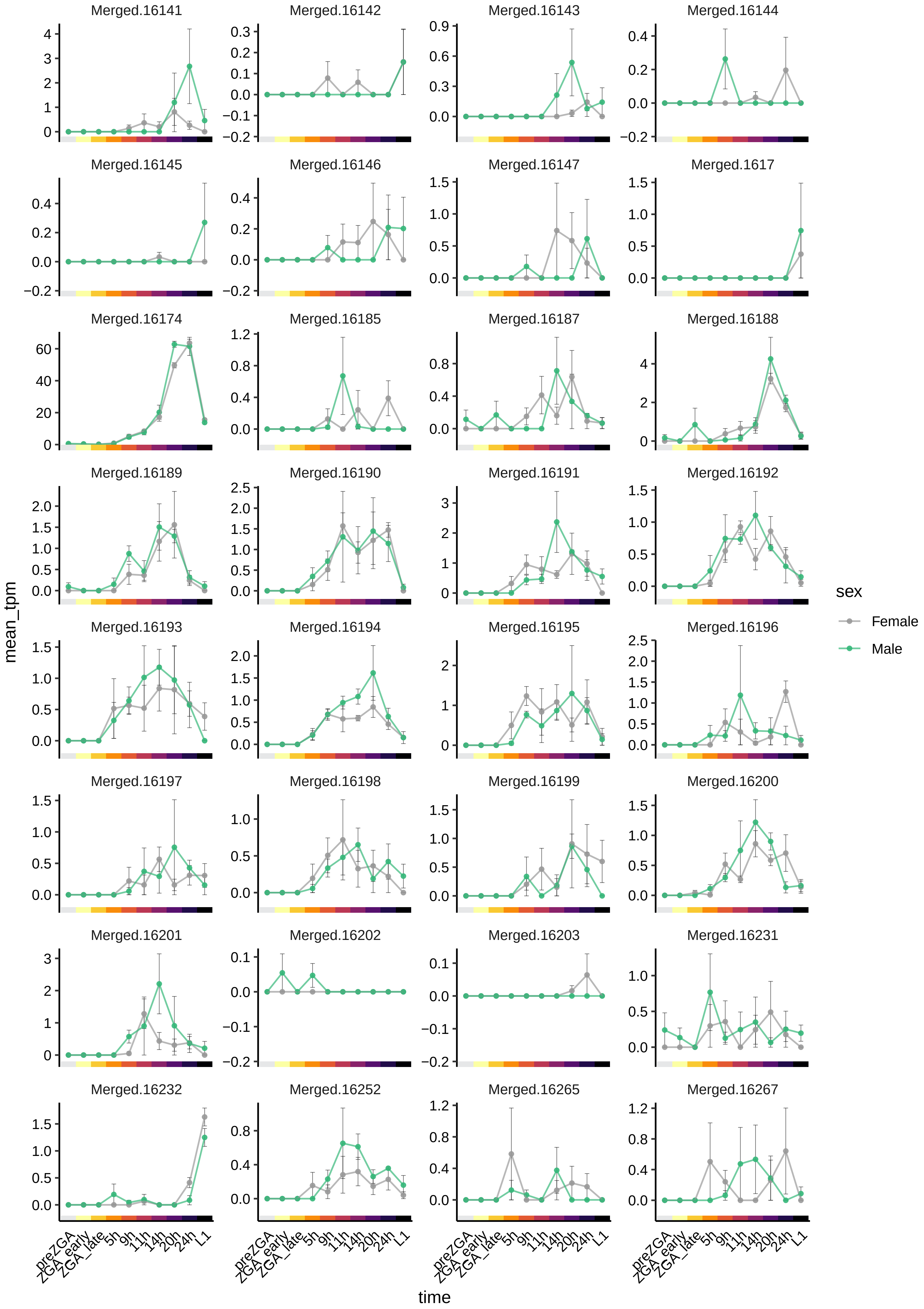

Merged.16141
Merged.16142
Merged.16143
Merged.16144
Merged.16145
Merged.16146
Merged.16147
Merged.1617
Merged.16174
Merged.16185
Merged.16187
Merged.16188
Merged.16189
Merged.16190
Merged.16191
Merged.16192
Merged.16193
Merged.16194
Merged.16195
Merged.16196
Merged.16197
Merged.16198
Merged.16199
Merged.16200
Merged.16201
Merged.16202
Merged.16203
Merged.16231
Merged.16232
Merged.16252
Merged.16265
Merged.16267
mean_tpm
time
preZGA
ZGA_early
ZGA_late
5h
9h
11h
14h
20h
24h
L1
sex
Female
Male

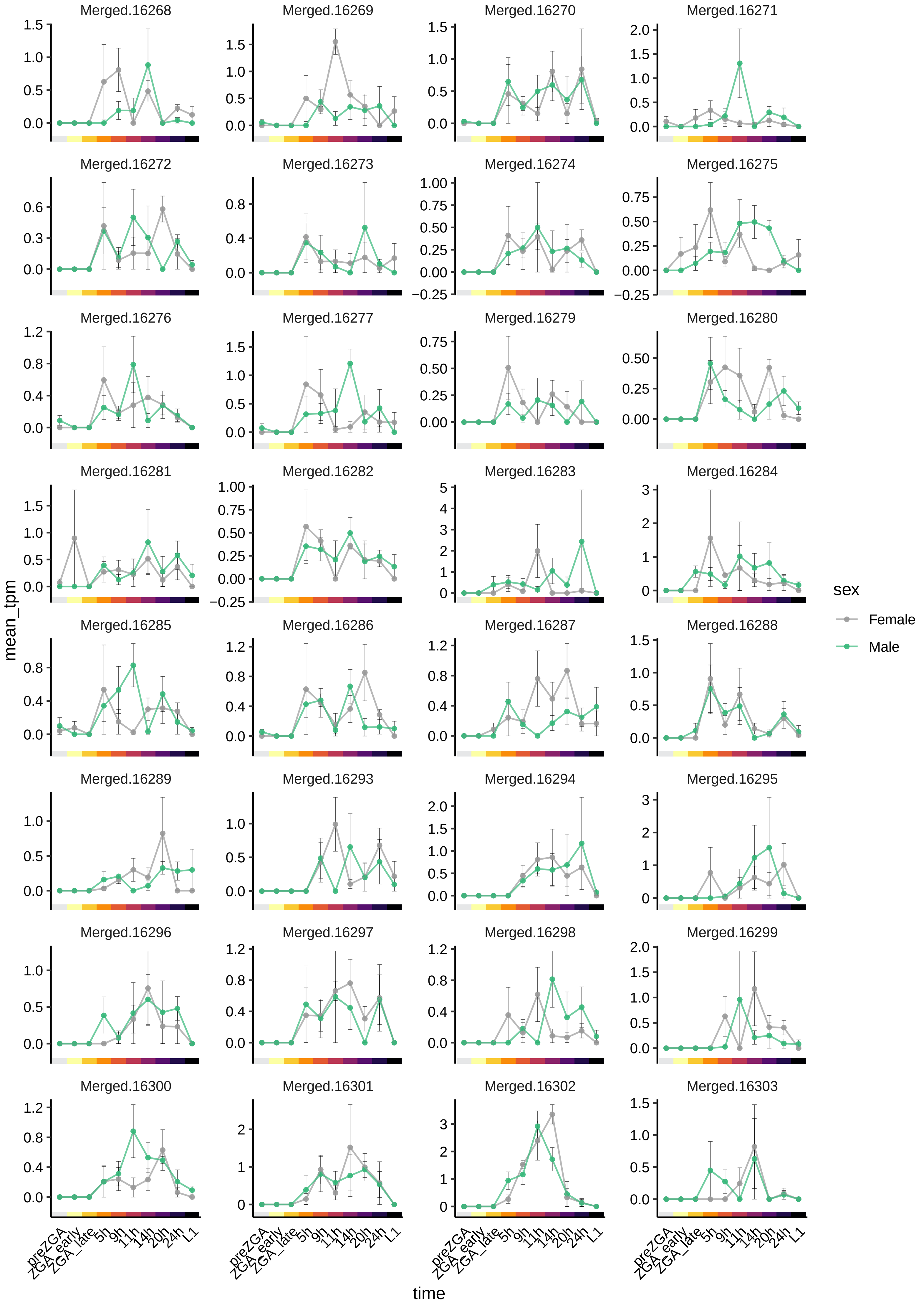

Merged.16268
Merged.16269
Merged.16270
Merged.16271
Merged.16272
Merged.16273
Merged.16274
Merged.16275
Merged.16276
Merged.16277
Merged.16279
Merged.16280
Merged.16281
Merged.16282
Merged.16283
Merged.16284
Merged.16285
Merged.16286
Merged.16287
Merged.16288
Merged.16289
Merged.16293
Merged.16294
Merged.16295
Merged.16296
Merged.16297
Merged.16298
Merged.16299
Merged.16300
Merged.16301
Merged.16302
Merged.16303
mean_tpm
time
preZGA
ZGA_early
ZGA_late
5h
9h
11h
14h
20h
24h
L1
sex
Female
Male

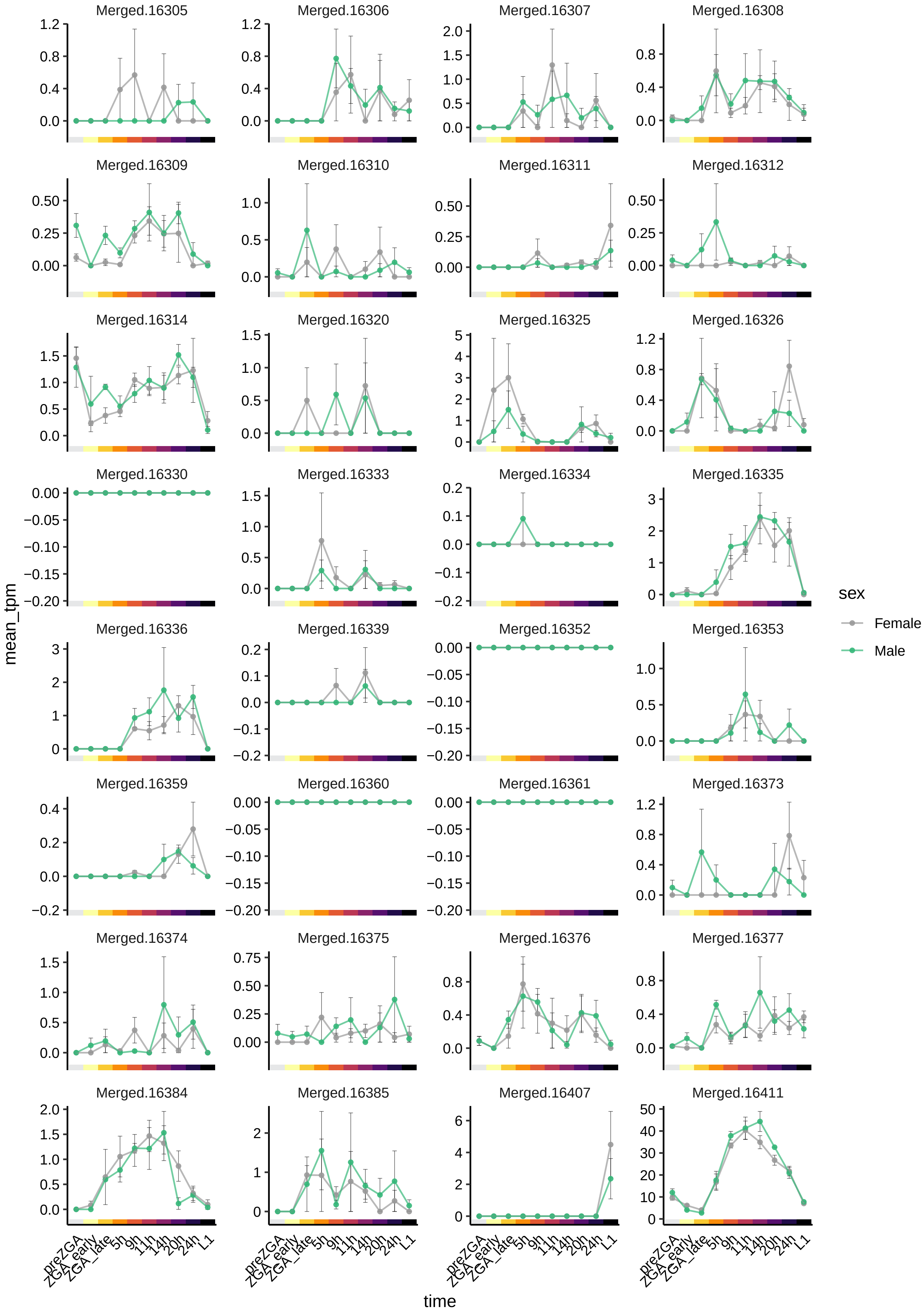

Merged.16305
Merged.16306
Merged.16307
Merged.16308
Merged.16309
Merged.16310
Merged.16311
Merged.16312
Merged.16314
Merged.16320
Merged.16325
Merged.16326
Merged.16330
Merged.16333
Merged.16334
Merged.16335
Merged.16336
Merged.16339
Merged.16352
Merged.16353
Merged.16359
Merged.16360
Merged.16361
Merged.16373
Merged.16374
Merged.16375
Merged.16376
Merged.16377
Merged.16384
Merged.16385
Merged.16407
Merged.16411
mean_tpm
time
preZGA
ZGA_early
ZGA_late
5h
9h
11h
14h
20h
24h
L1
sex
Female
Male

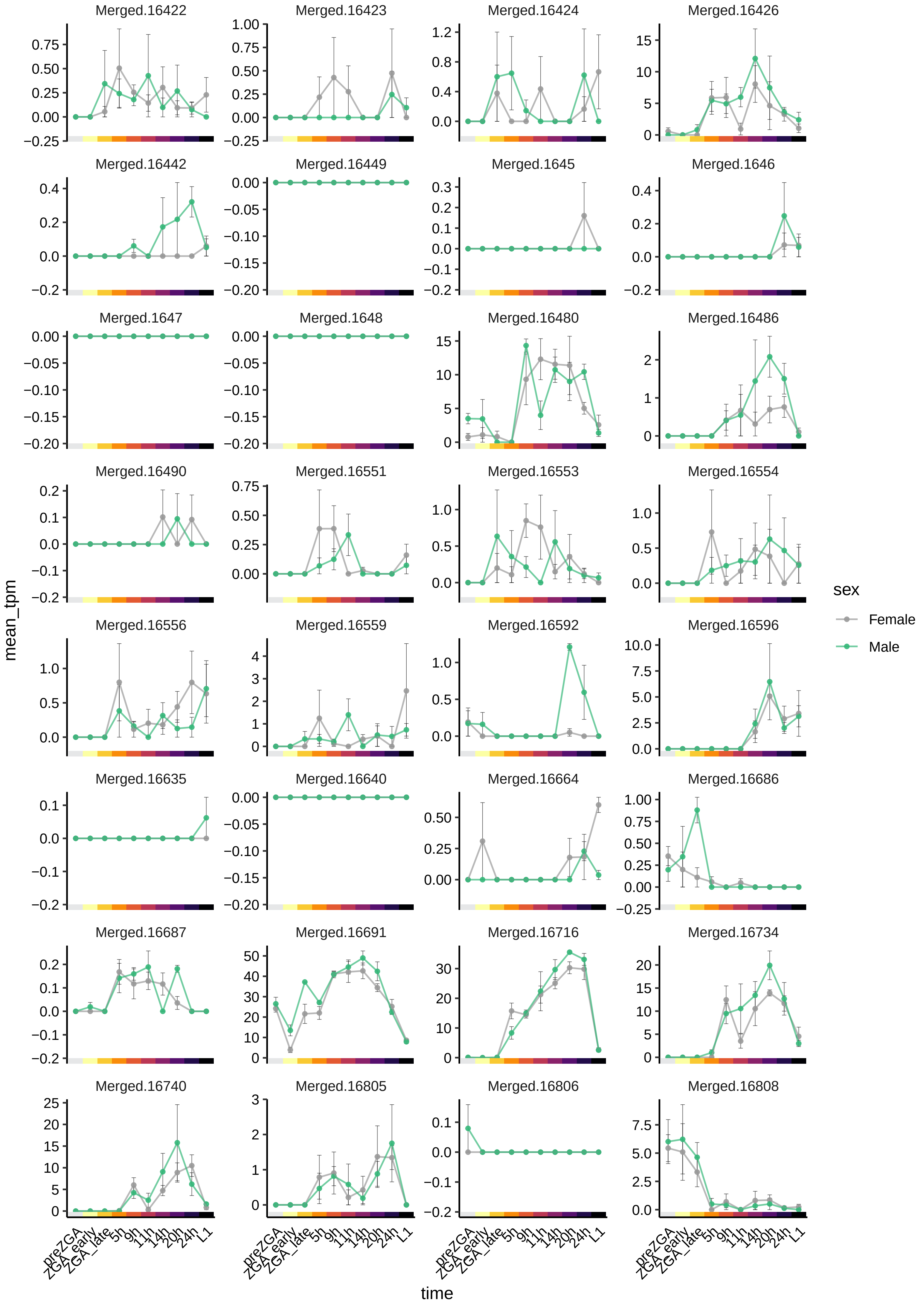
Merged.16422
Merged.16423
Merged.16424
Merged.16426
Merged.16442
Merged.16449
Merged.1645
Merged.1646
Merged.1647
Merged.1648
Merged.16480
Merged.16486
Merged.16490
Merged.16551
Merged.16553
Merged.16554
Merged.16556
Merged.16559
Merged.16592
Merged.16596
Merged.16635
Merged.16640
Merged.16664
Merged.16686
Merged.16687
Merged.16691
Merged.16716
Merged.16734
Merged.16740
Merged.16805
Merged.16806
Merged.16808
mean_tpm
time
preZGA
ZGA_early
ZGA_late
5h
9h
11h
14h
20h
24h
L1
sex
Female
Male

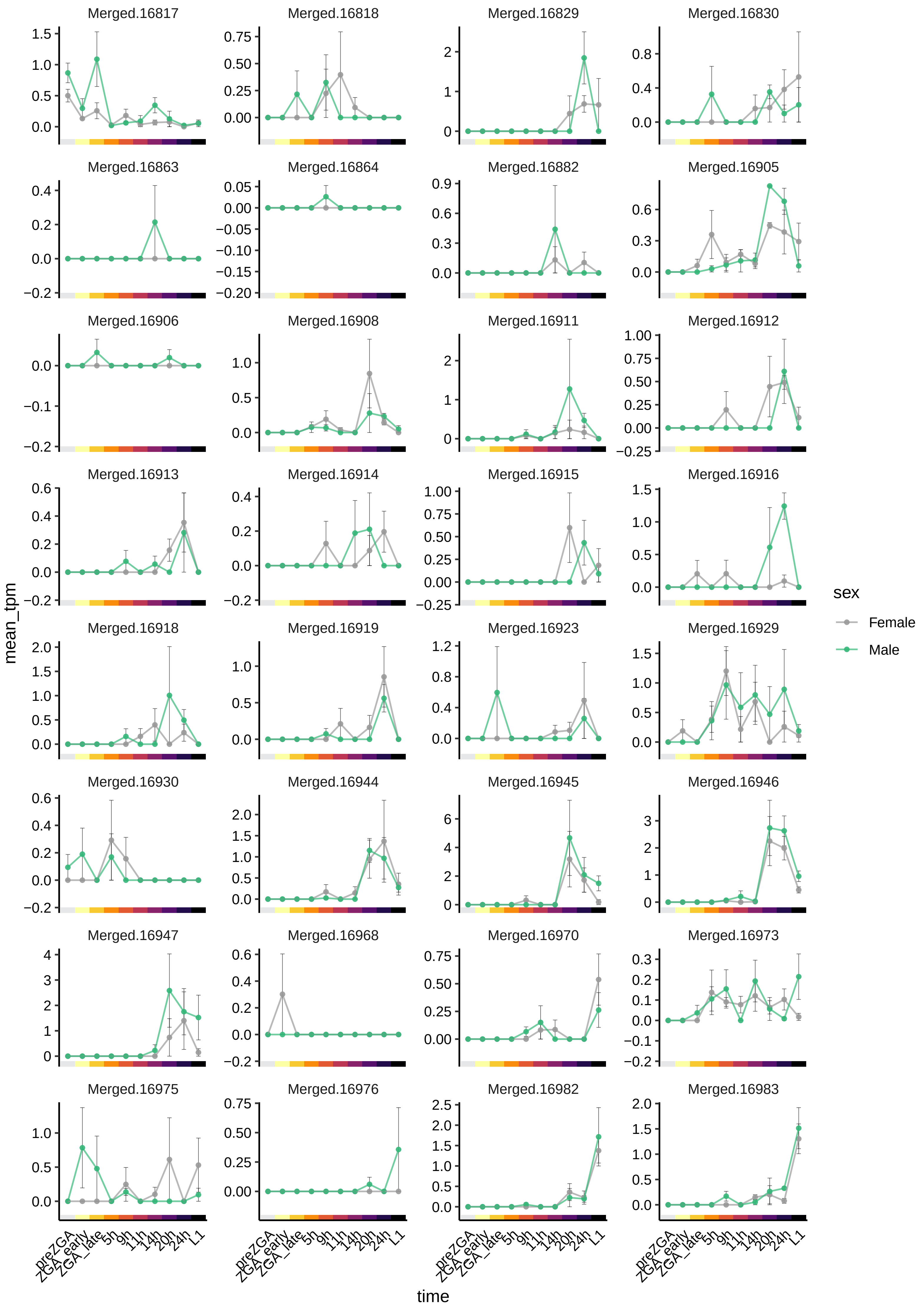
Merged.16817
Merged.16818
Merged.16829
Merged.16830
Merged.16863
Merged.16864
Merged.16882
Merged.16905
Merged.16906
Merged.16908
Merged.16911
Merged.16912
Merged.16913
Merged.16914
Merged.16915
Merged.16916
Merged.16918
Merged.16919
Merged.16923
Merged.16929
Merged.16930
Merged.16944
Merged.16945
Merged.16946
Merged.16947
Merged.16968
Merged.16970
Merged.16973
Merged.16975
Merged.16976
Merged.16982
Merged.16983
mean_tpm
time
preZGA
ZGA_early
ZGA_late
5h
9h
11h
14h
20h
24h
L1
sex
Female
Male

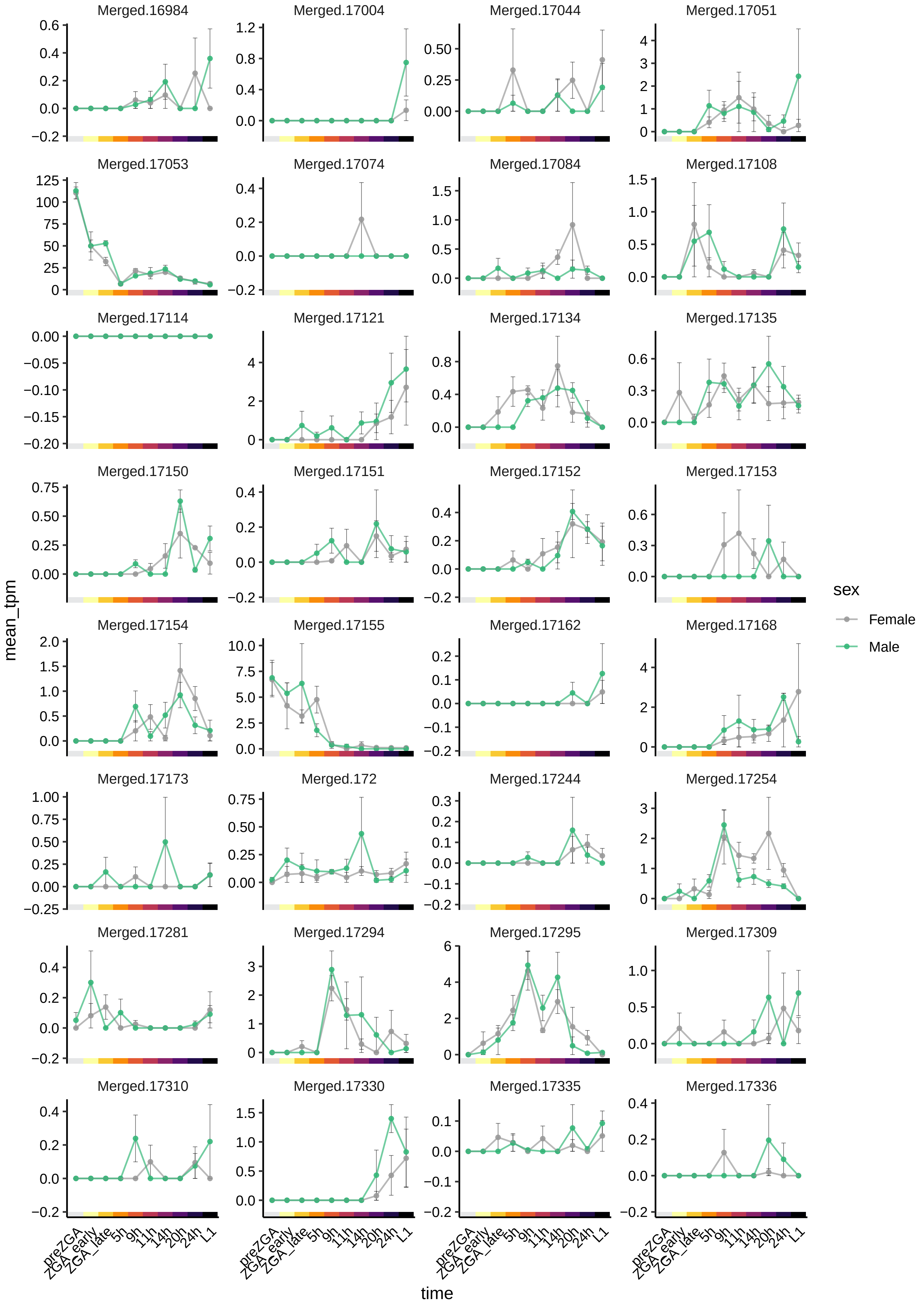
Merged.16984
Merged.17004
Merged.17044
Merged.17051
Merged.17053
Merged.17074
Merged.17084
Merged.17108
Merged.17114
Merged.17121
Merged.17134
Merged.17135
Merged.17150
Merged.17151
Merged.17152
Merged.17153
Merged.17154
Merged.17155
Merged.17162
Merged.17168
Merged.17173
Merged.172
Merged.17244
Merged.17254
Merged.17281
Merged.17294
Merged.17295
Merged.17309
Merged.17310
Merged.17330
Merged.17335
Merged.17336
mean_tpm
time
preZGA
ZGA_early
ZGA_late
5h
9h
11h
14h
20h
24h
L1
sex
Female
Male

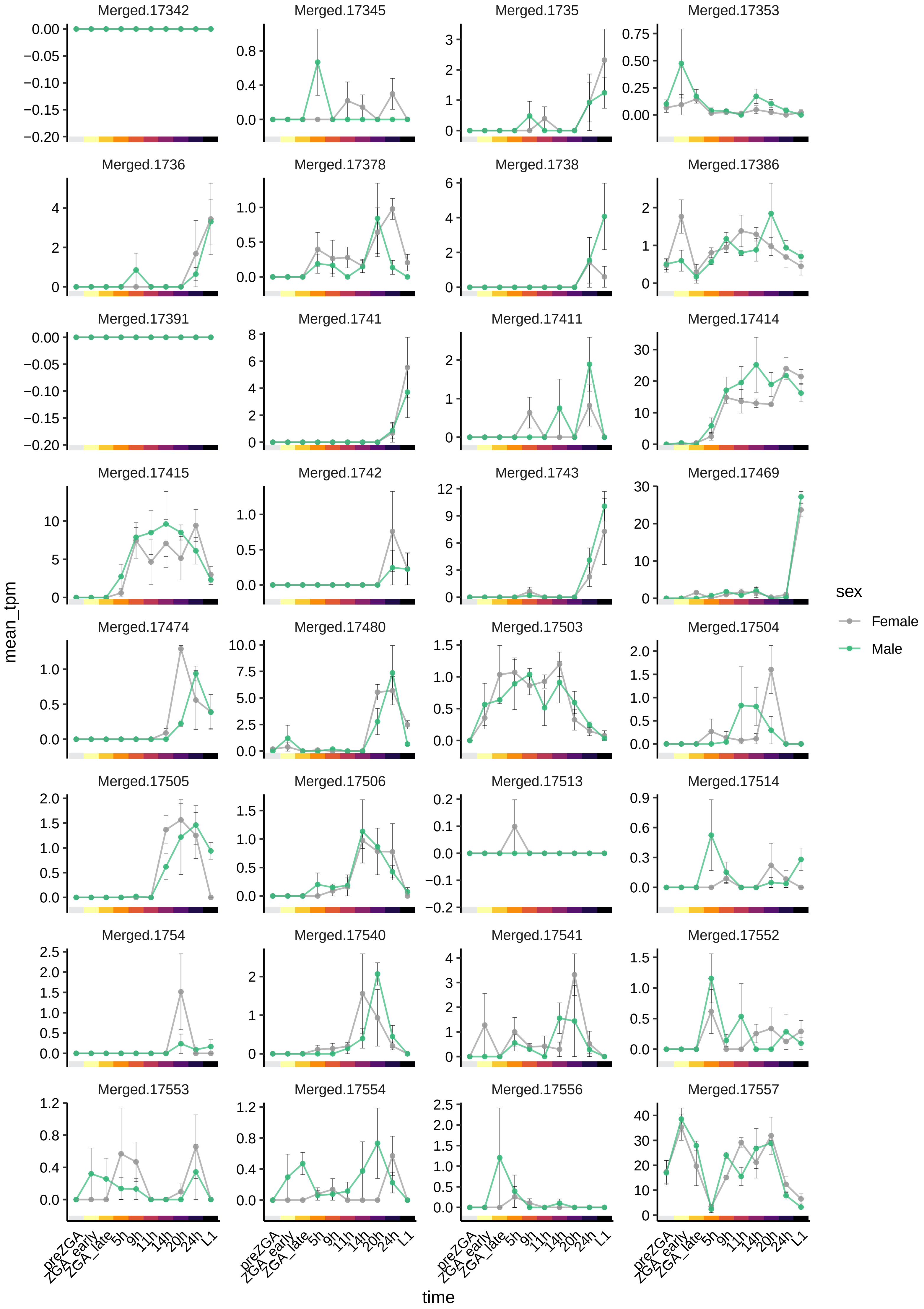

Merged.17342
Merged.17345
Merged.1735
Merged.17353
Merged.1736
Merged.17378
Merged.1738
Merged.17386
Merged.17391
Merged.1741
Merged.17411
Merged.17414
Merged.17415
Merged.1742
Merged.1743
Merged.17469
Merged.17474
Merged.17480
Merged.17503
Merged.17504
Merged.17505
Merged.17506
Merged.17513
Merged.17514
Merged.1754
Merged.17540
Merged.17541
Merged.17552
Merged.17553
Merged.17554
Merged.17556
Merged.17557
mean_tpm
time
preZGA
ZGA_early
ZGA_late
5h
9h
11h
14h
20h
24h
L1
sex
Female
Male

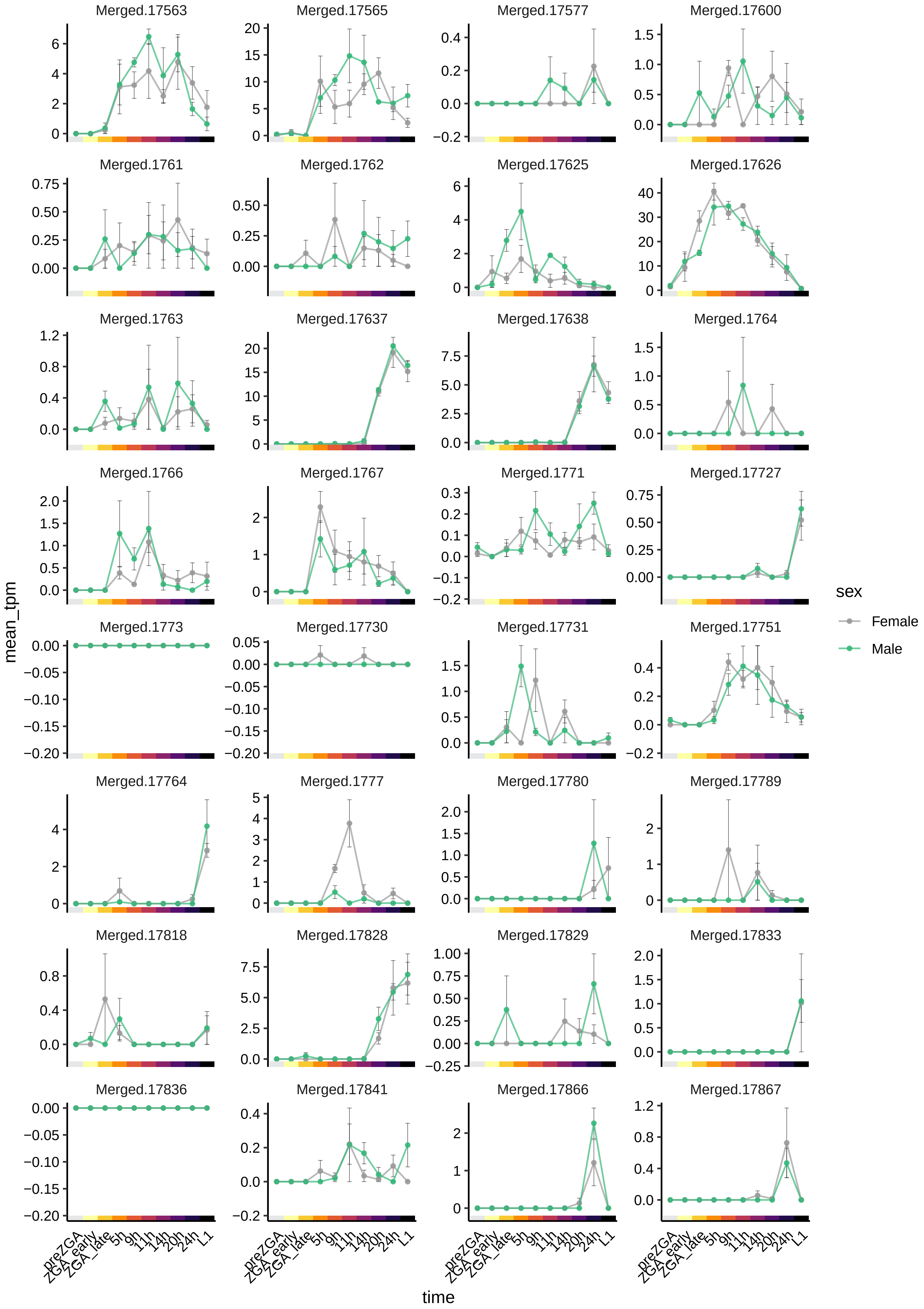

Merged.17563
Merged.17565
Merged.17577
Merged.17600
Merged.1761
Merged.1762
Merged.17625
Merged.17626
Merged.1763
Merged.17637
Merged.17638
Merged.1764
Merged.1766
Merged.1767
Merged.1771
Merged.17727
Merged.1773
Merged.17730
Merged.17731
Merged.17751
Merged.17764
Merged.1777
Merged.17780
Merged.17789
Merged.17818
Merged.17828
Merged.17829
Merged.17833
Merged.17836
Merged.17841
Merged.17866
Merged.17867
mean_tpm
time
preZGA
ZGA_early
ZGA_late
5h
9h
11h
14h
20h
24h
L1
sex
Female
Male

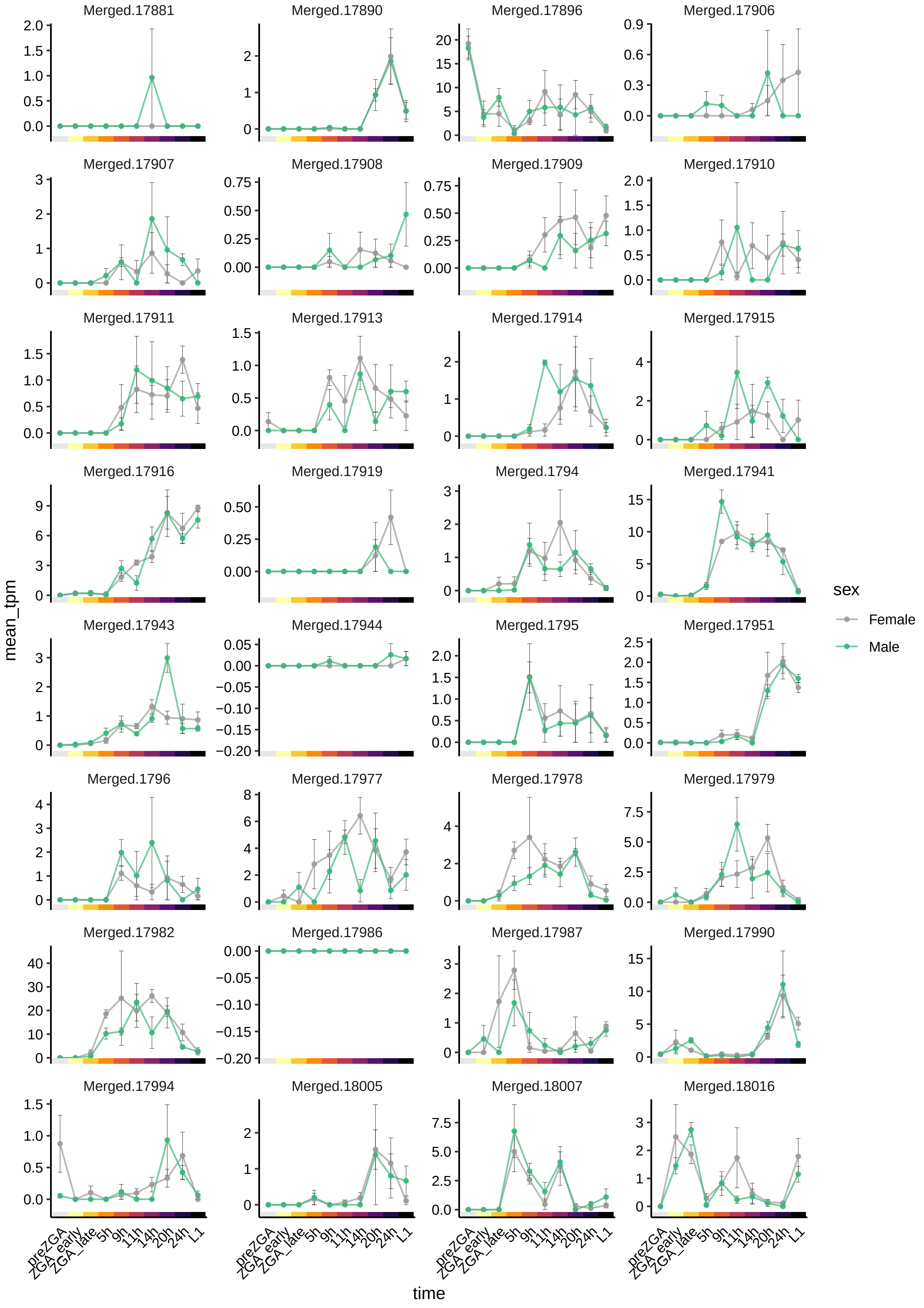
Merged.17881
Merged.17890
Merged.17896
Merged.17906
Merged.17907
Merged.17908
Merged.17909
Merged.17910
Merged.17911
Merged.17913
Merged.17914
Merged.17915
Merged.17916
Merged.17919
Merged.1794
Merged.17941
Merged.17943
Merged.17944
Merged.1795
Merged.17951
Merged.1796
Merged.17977
Merged.17978
Merged.17979
Merged.17982
Merged.17986
Merged.17987
Merged.17990
Merged.17994
Merged.18005
Merged.18007
Merged.18016
mean_tpm
time
preZGA
ZGA_early
ZGA_late
5h
9h
11h
14h
20h
24h
L1
sex
Female
Male

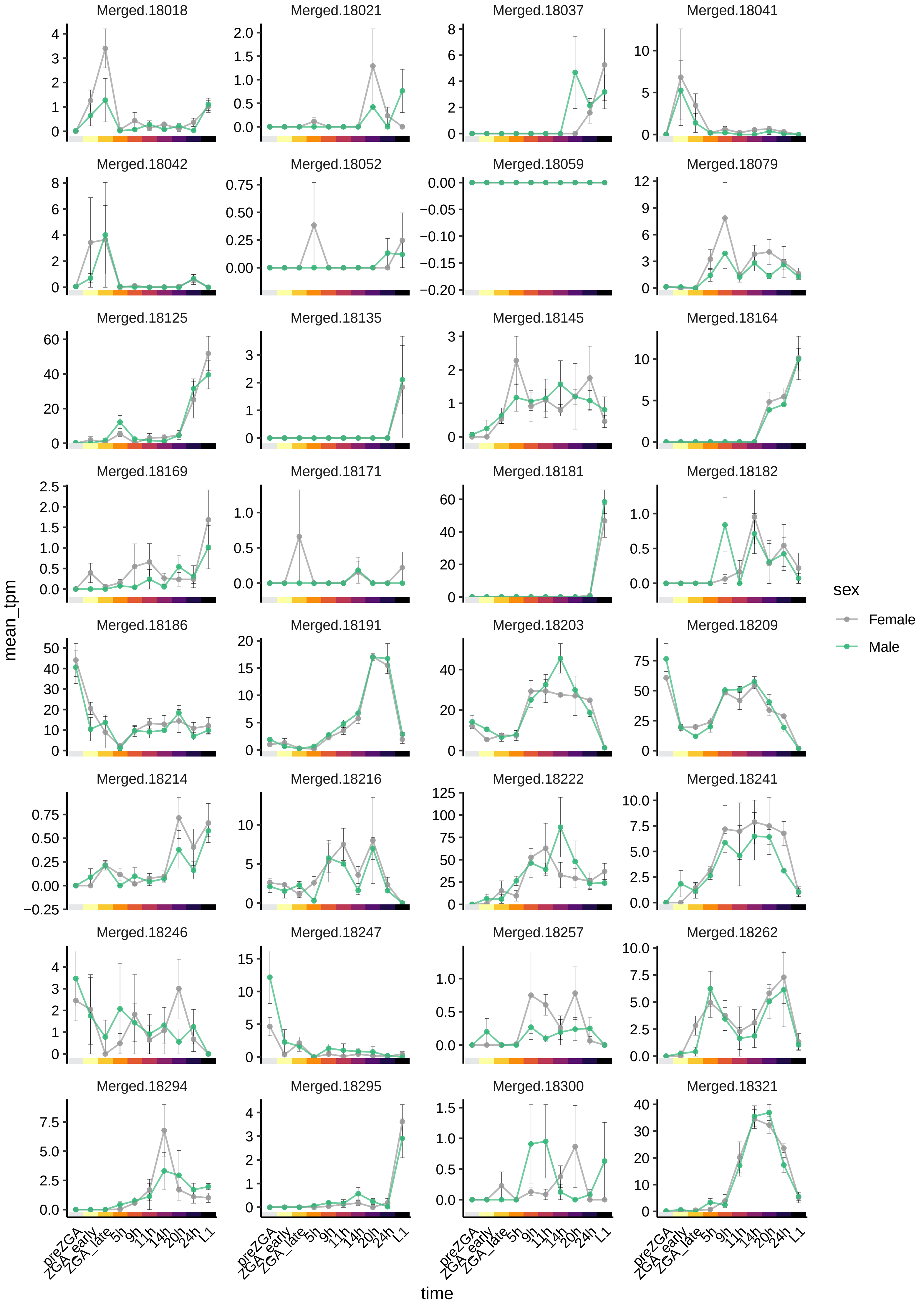

Merged.18018
Merged.18021
Merged.18037
Merged.18041
Merged.18042
Merged.18052
Merged.18059
Merged.18079
Merged.18125
Merged.18135
Merged.18145
Merged.18164
Merged.18169
Merged.18171
Merged.18181
Merged.18182
Merged.18186
Merged.18191
Merged.18203
Merged.18209
Merged.18214
Merged.18216
Merged.18222
Merged.18241
Merged.18246
Merged.18247
Merged.18257
Merged.18262
Merged.18294
Merged.18295
Merged.18300
Merged.18321
mean_tpm
time
preZGA
ZGA_early
ZGA_late
5h
9h
11h
14h
20h
24h
L1
sex
Female
Male

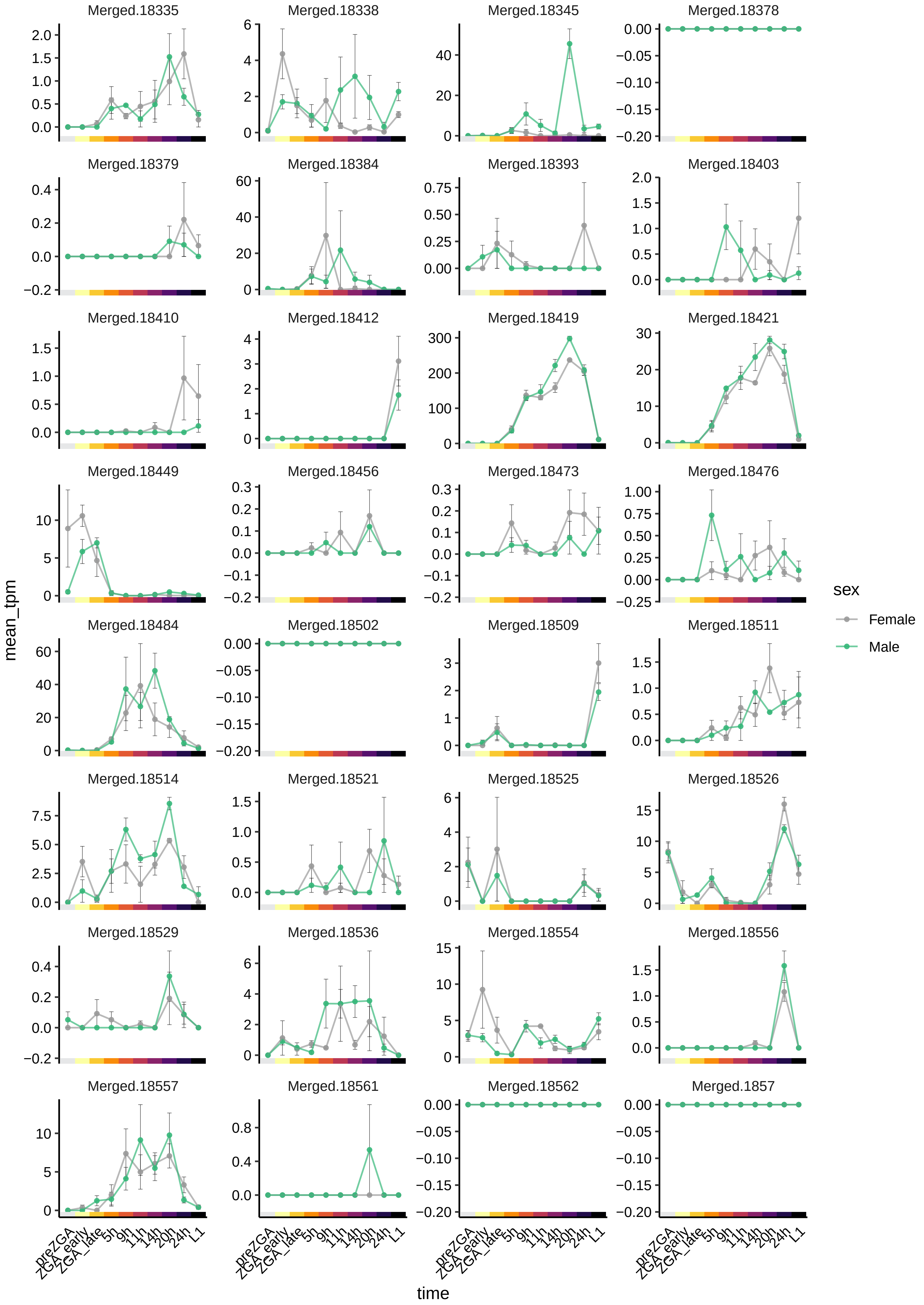
Merged.18335
Merged.18338
Merged.18345
Merged.18378
Merged.18379
Merged.18384
Merged.18393
Merged.18403
Merged.18410
Merged.18412
Merged.18419
Merged.18421
Merged.18449
Merged.18456
Merged.18473
Merged.18476
Merged.18484
Merged.18502
Merged.18509
Merged.18511
Merged.18514
Merged.18521
Merged.18525
Merged.18526
Merged.18529
Merged.18536
Merged.18554
Merged.18556
Merged.18557
Merged.18561
Merged.18562
Merged.1857
mean_tpm
time
preZGA
ZGA_early
ZGA_late
5h
9h
11h
14h
20h
24h
L1
sex
Female
Male

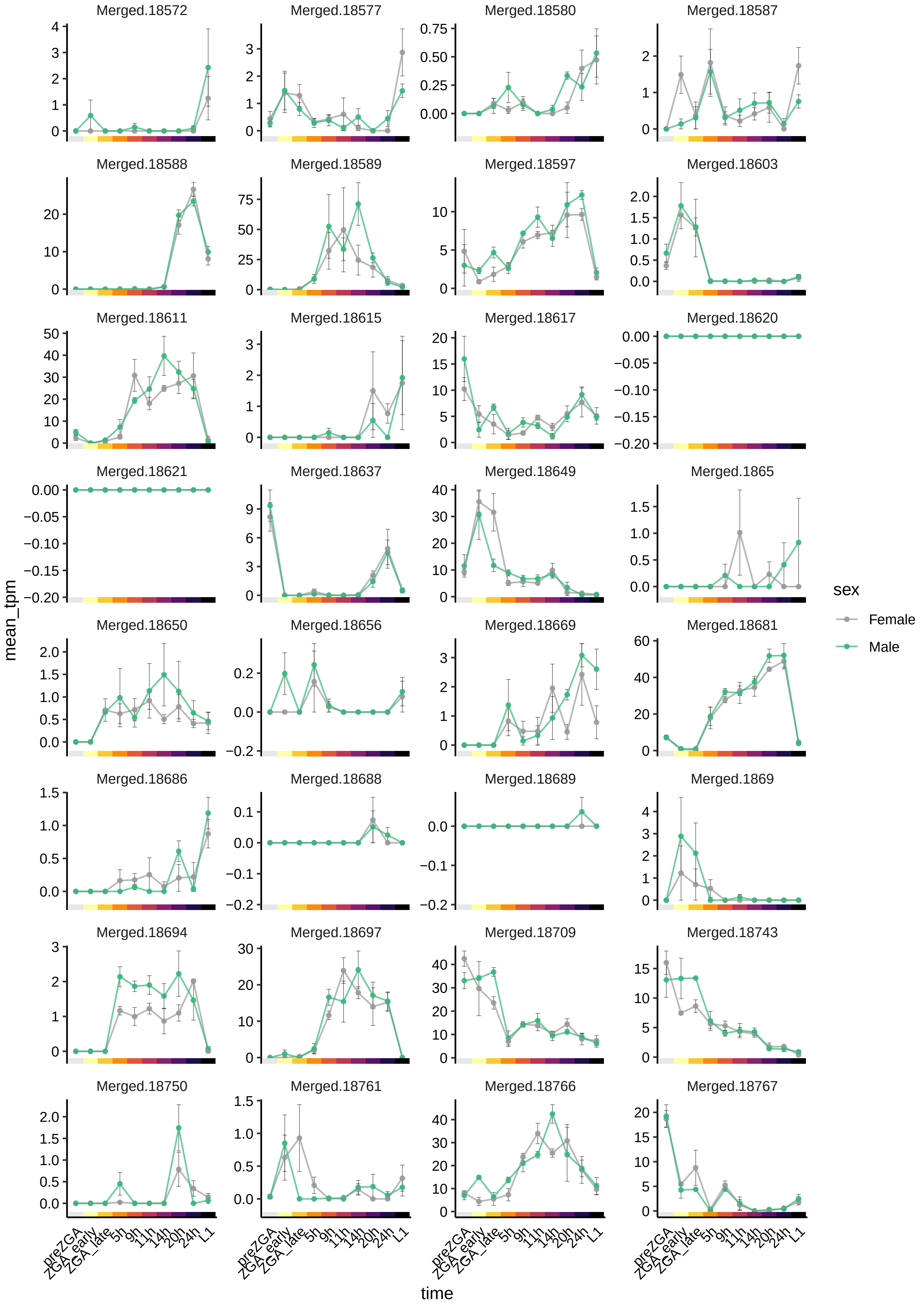

Merged.18572
Merged.18577
Merged.18580
Merged.18587
Merged.18588
Merged.18589
Merged.18597
Merged.18603
Merged.18611
Merged.18615
Merged.18617
Merged.18620
Merged.18621
Merged.18637
Merged.18649
Merged.1865
Merged.18650
Merged.18656
Merged.18669
Merged.18681
Merged.18686
Merged.18688
Merged.18689
Merged.1869
Merged.18694
Merged.18697
Merged.18709
Merged.18743
Merged.18750
Merged.18761
Merged.18766
Merged.18767
mean_tpm
time
preZGA
ZGA_early
ZGA_late
5h
9h
11h
14h
20h
24h
L1
sex
Female
Male

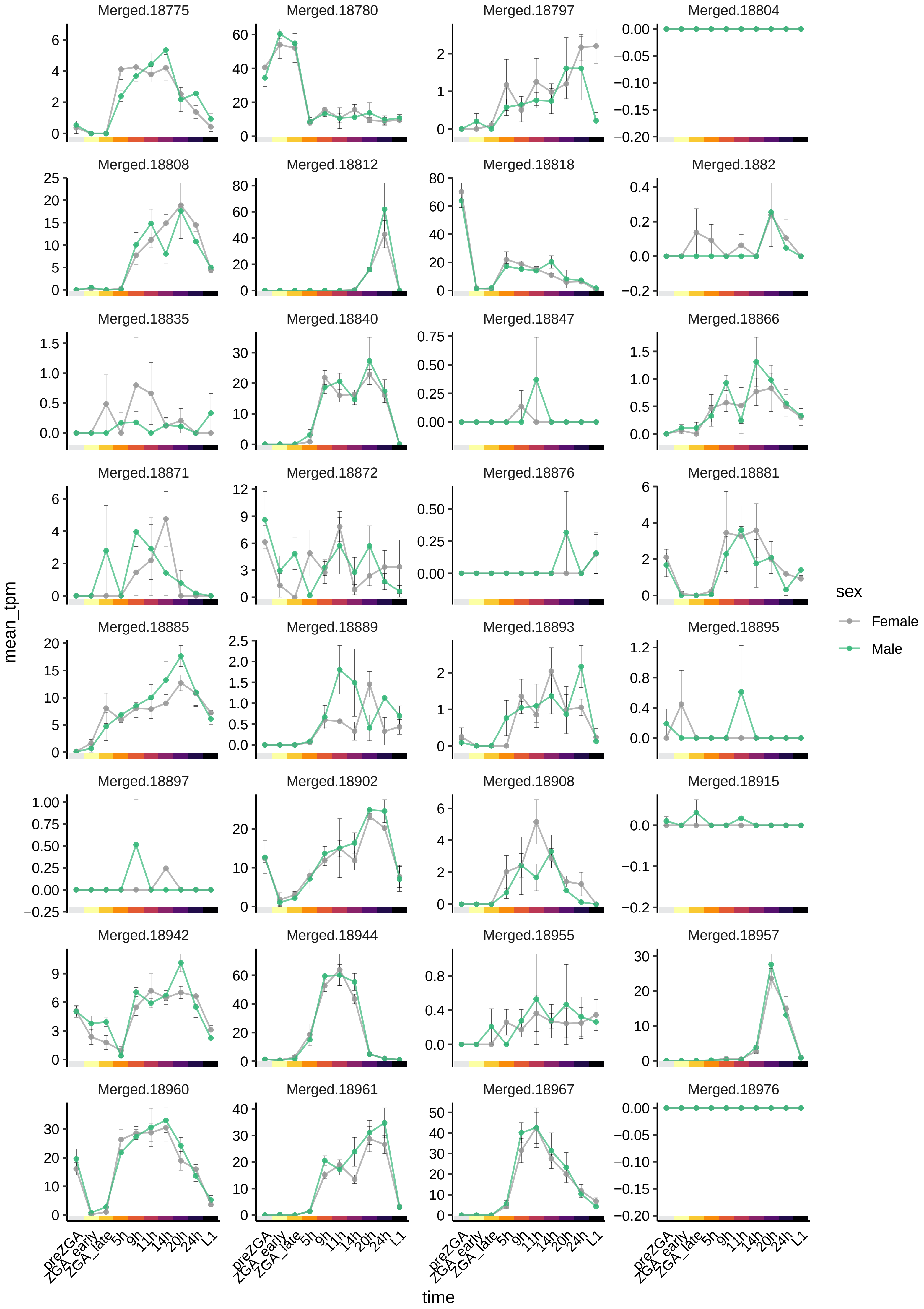

Merged.18775
Merged.18780
Merged.18797
Merged.18804
Merged.18808
Merged.18812
Merged.18818
Merged.1882
Merged.18835
Merged.18840
Merged.18847
Merged.18866
Merged.18871
Merged.18872
Merged.18876
Merged.18881
Merged.18885
Merged.18889
Merged.18893
Merged.18895
Merged.18897
Merged.18902
Merged.18908
Merged.18915
Merged.18942
Merged.18944
Merged.18955
Merged.18957
Merged.18960
Merged.18961
Merged.18967
Merged.18976
mean_tpm
time
preZGA
ZGA_early
ZGA_late
5h
9h
11h
14h
20h
24h
L1
sex
Female
Male

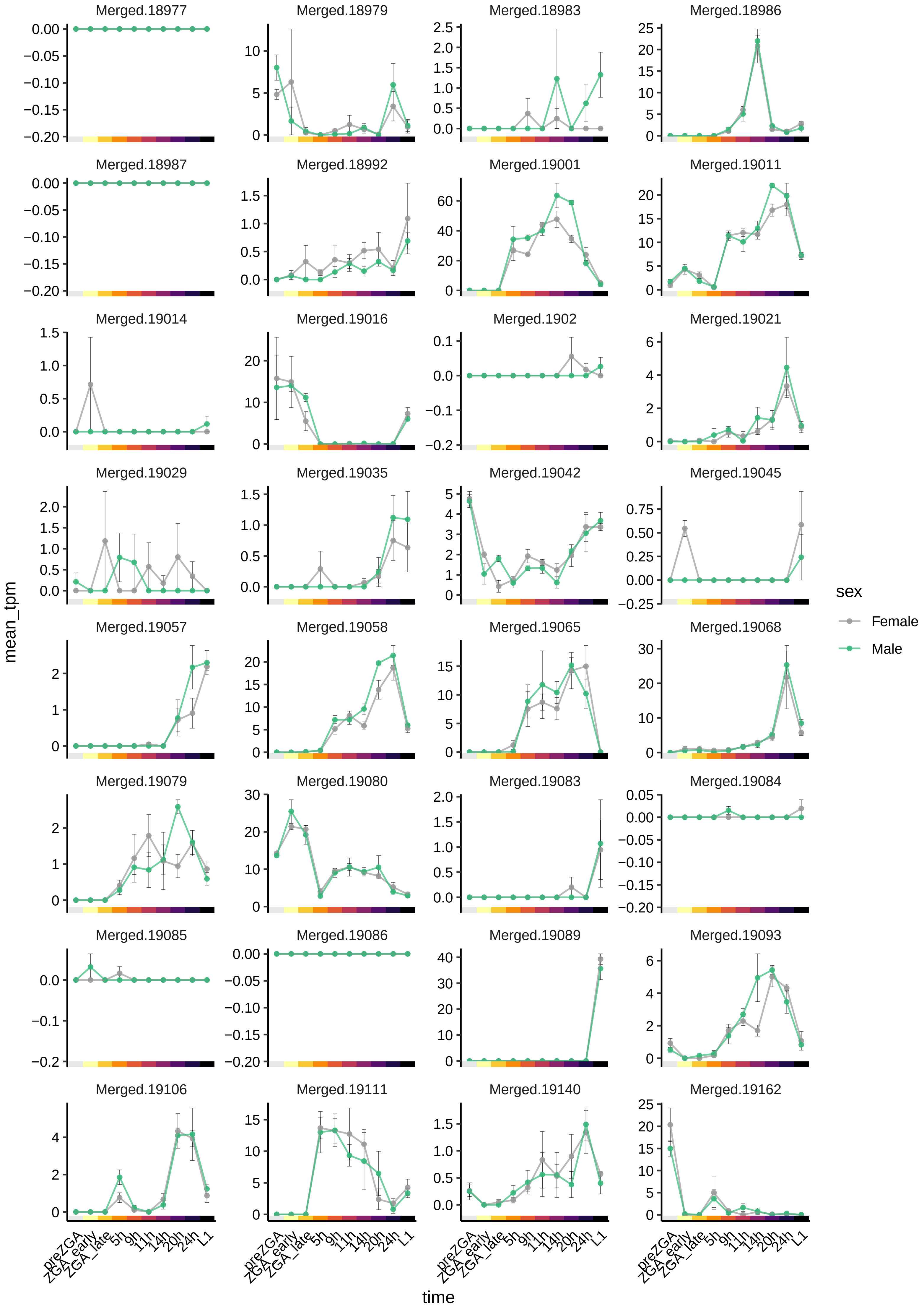

Merged.18977
Merged.18979
Merged.18983
Merged.18986
Merged.18987
Merged.18992
Merged.19001
Merged.19011
Merged.19014
Merged.19016
Merged.1902
Merged.19021
Merged.19029
Merged.19035
Merged.19042
Merged.19045
Merged.19057
Merged.19058
Merged.19065
Merged.19068
Merged.19079
Merged.19080
Merged.19083
Merged.19084
Merged.19085
Merged.19086
Merged.19089
Merged.19093
Merged.19106
Merged.19111
Merged.19140
Merged.19162
mean_tpm
time
preZGA
ZGA_early
ZGA_late
5h
9h
11h
14h
20h
24h
L1
sex
Female
Male

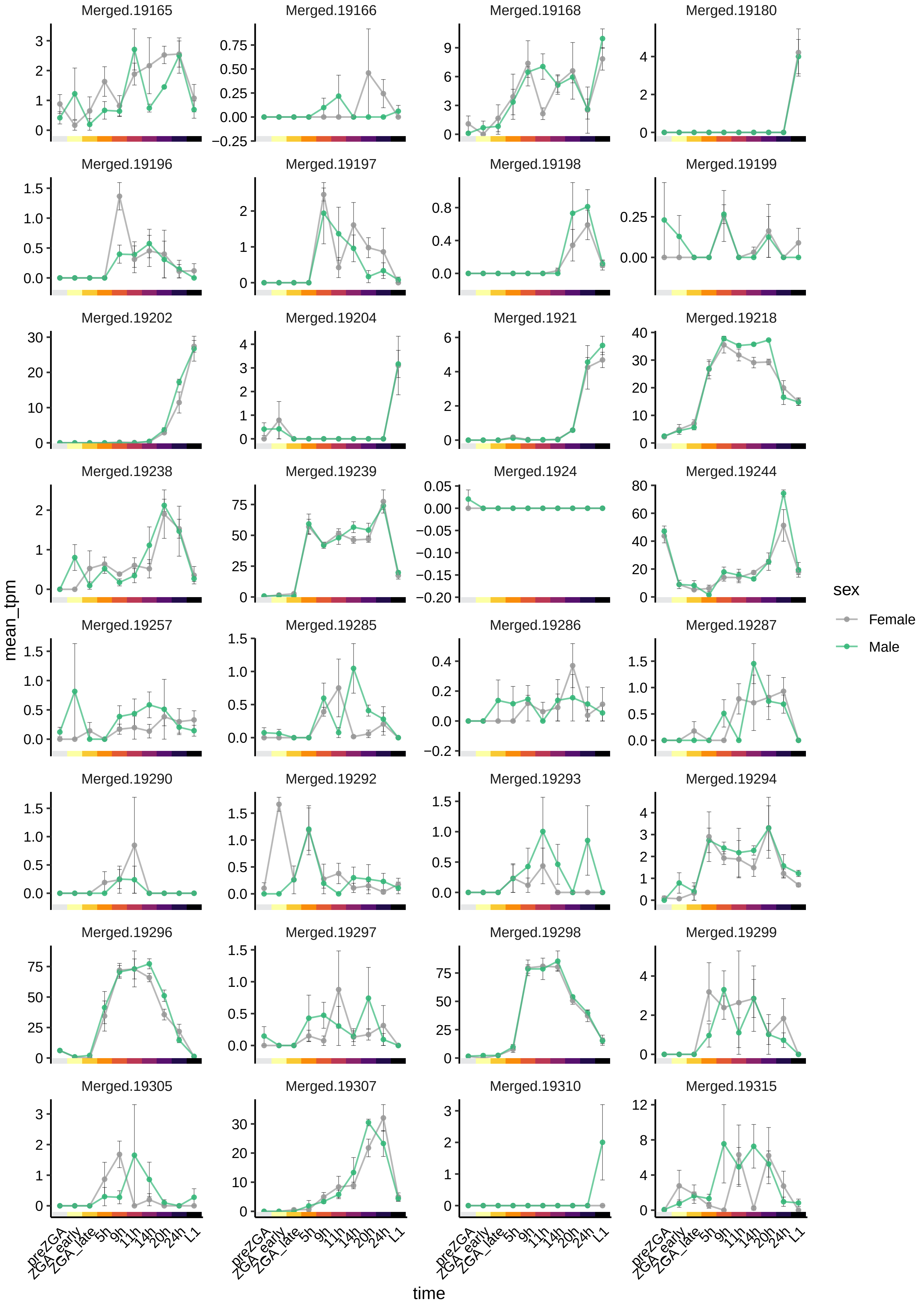

Merged.19165
Merged.19166
Merged.19168
Merged.19180
Merged.19196
Merged.19197
Merged.19198
Merged.19199
Merged.19202
Merged.19204
Merged.1921
Merged.19218
Merged.19238
Merged.19239
Merged.1924
Merged.19244
Merged.19257
Merged.19285
Merged.19286
Merged.19287
Merged.19290
Merged.19292
Merged.19293
Merged.19294
Merged.19296
Merged.19297
Merged.19298
Merged.19299
Merged.19305
Merged.19307
Merged.19310
Merged.19315
mean_tpm
time
preZGA
ZGA_early
ZGA_late
5h
9h
11h
14h
20h
24h
L1
sex
Female
Male

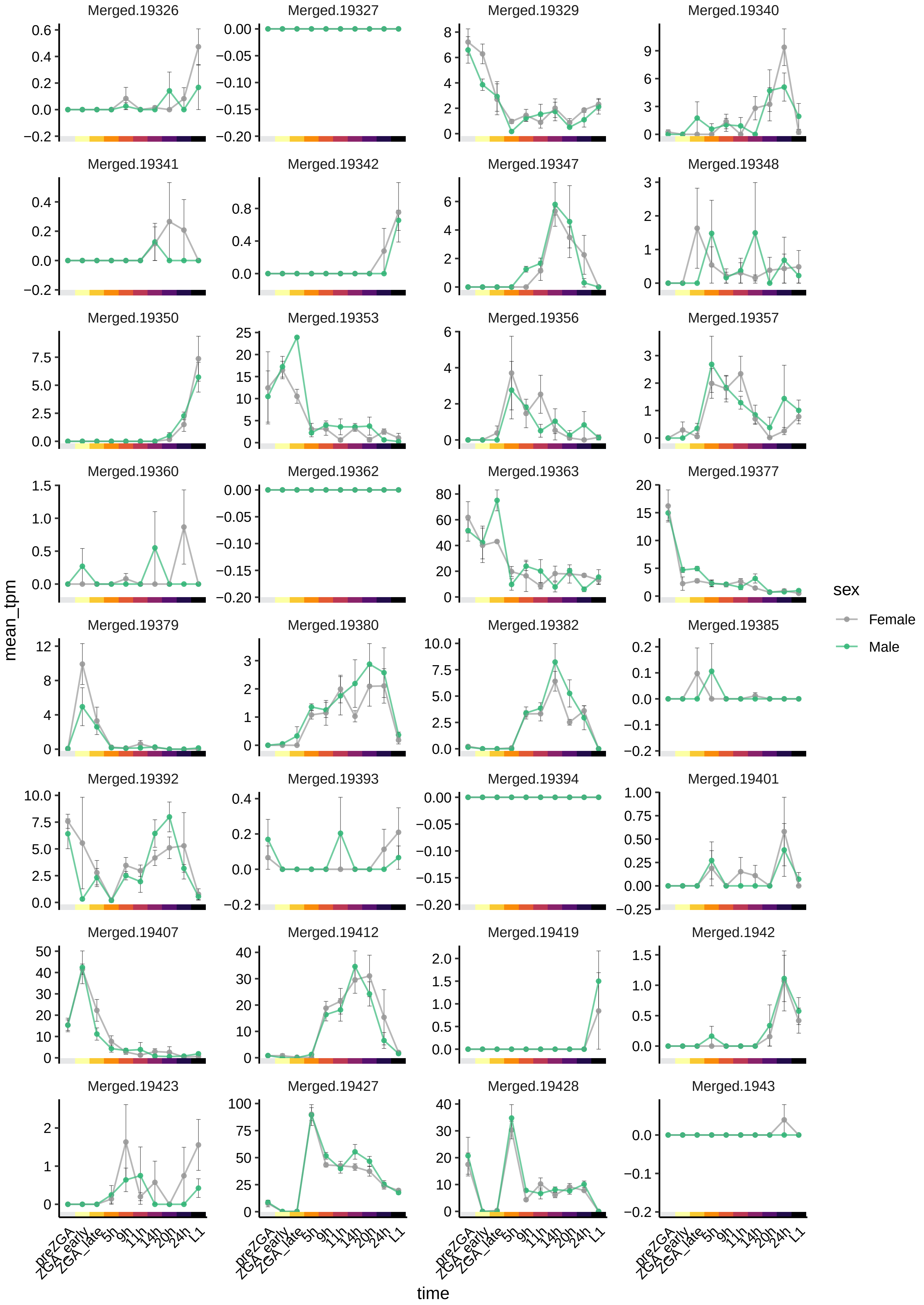

Merged.19326
Merged.19327
Merged.19329
Merged.19340
Merged.19341
Merged.19342
Merged.19347
Merged.19348
Merged.19350
Merged.19353
Merged.19356
Merged.19357
Merged.19360
Merged.19362
Merged.19363
Merged.19377
Merged.19379
Merged.19380
Merged.19382
Merged.19385
Merged.19392
Merged.19393
Merged.19394
Merged.19401
Merged.19407
Merged.19412
Merged.19419
Merged.1942
Merged.19423
Merged.19427
Merged.19428
Merged.1943
mean_tpm
time
preZGA
ZGA_early
ZGA_late
5h
9h
11h
14h
20h
24h
L1
sex
Female
Male

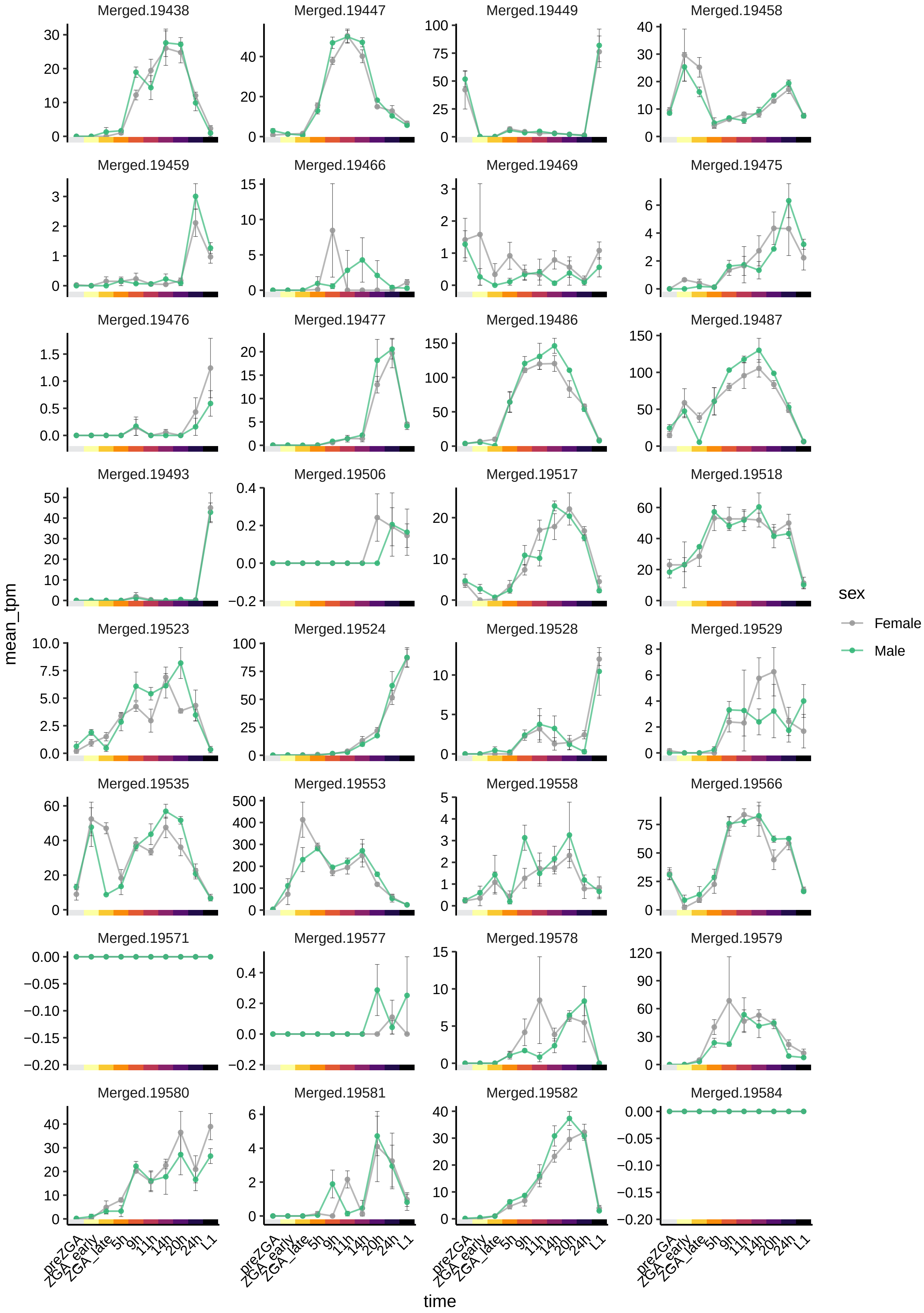

Merged.19438
Merged.19447
Merged.19449
Merged.19458
Merged.19459
Merged.19466
Merged.19469
Merged.19475
Merged.19476
Merged.19477
Merged.19486
Merged.19487
Merged.19493
Merged.19506
Merged.19517
Merged.19518
Merged.19523
Merged.19524
Merged.19528
Merged.19529
Merged.19535
Merged.19553
Merged.19558
Merged.19566
Merged.19571
Merged.19577
Merged.19578
Merged.19579
Merged.19580
Merged.19581
Merged.19582
Merged.19584
mean_tpm
time
preZGA
ZGA_early
ZGA_late
5h
9h
11h
14h
20h
24h
L1
sex
Female
Male

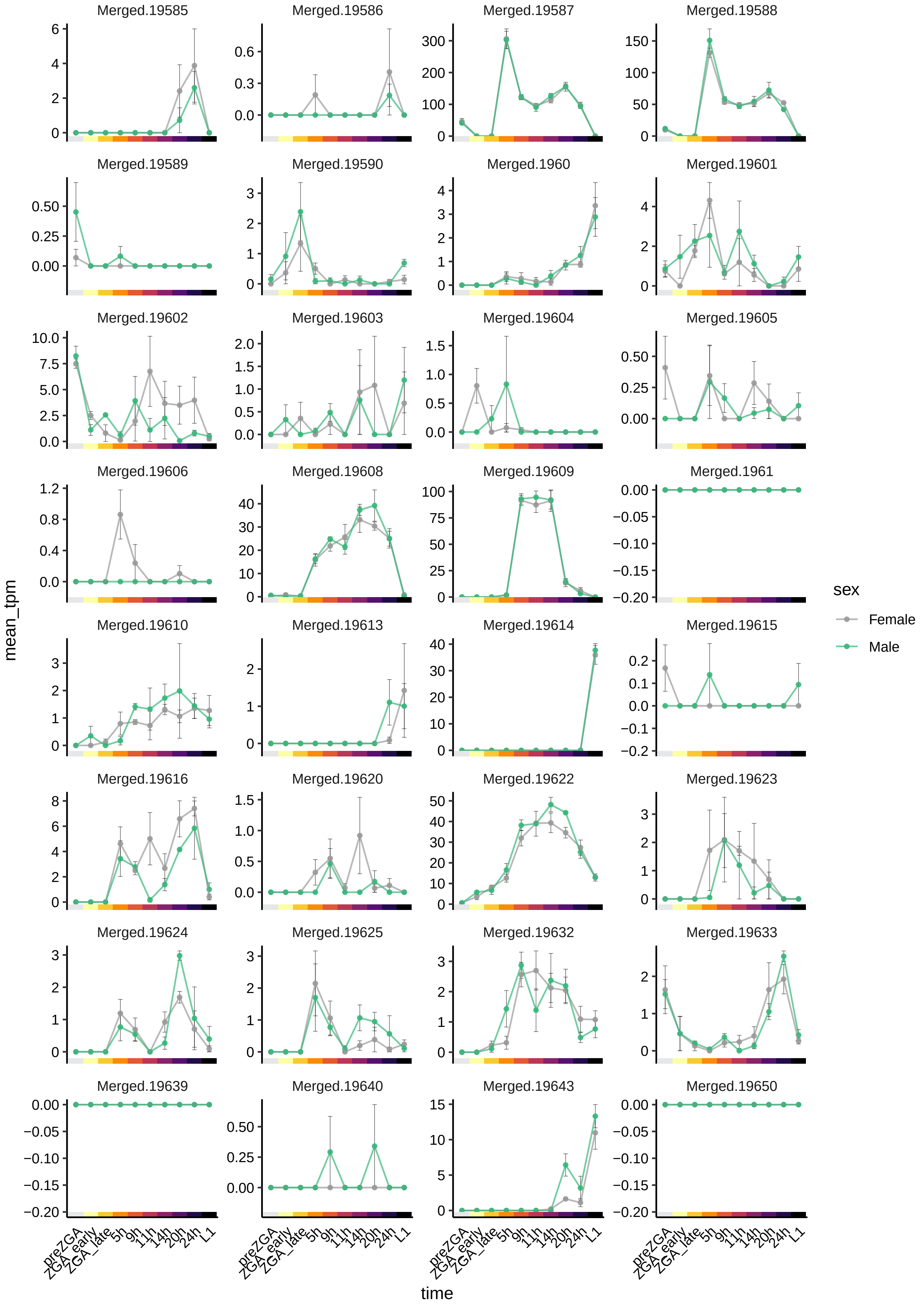

Merged.19585
Merged.19586
Merged.19587
Merged.19588
Merged.19589
Merged.19590
Merged.1960
Merged.19601
Merged.19602
Merged.19603
Merged.19604
Merged.19605
Merged.19606
Merged.19608
Merged.19609
Merged.1961
Merged.19610
Merged.19613
Merged.19614
Merged.19615
Merged.19616
Merged.19620
Merged.19622
Merged.19623
Merged.19624
Merged.19625
Merged.19632
Merged.19633
Merged.19639
Merged.19640
Merged.19643
Merged.19650
mean_tpm
time
preZGA
ZGA_early
ZGA_late
5h
9h
11h
14h
20h
24h
L1
sex
Female
Male

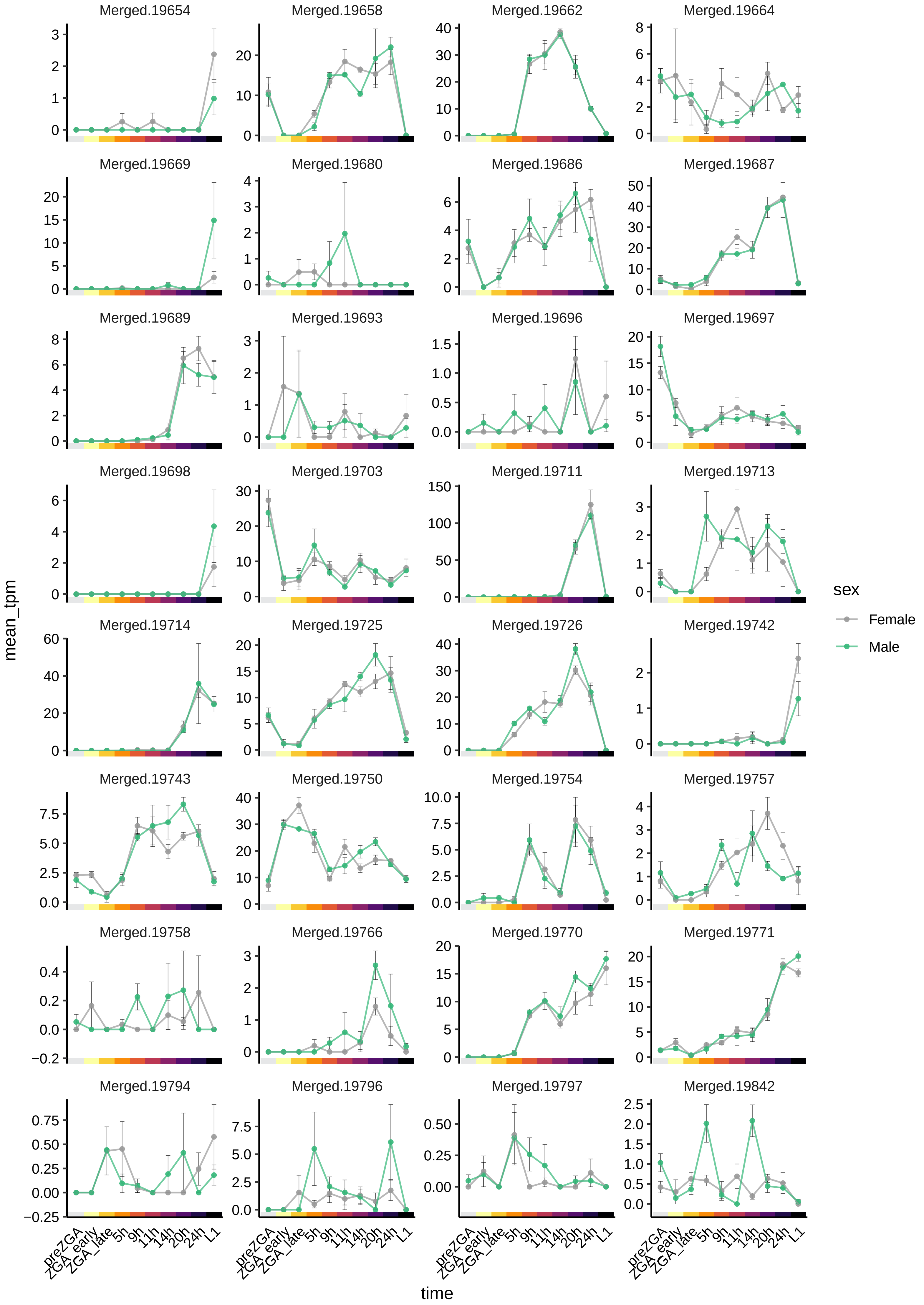

Merged.19654
Merged.19658
Merged.19662
Merged.19664
Merged.19669
Merged.19680
Merged.19686
Merged.19687
Merged.19689
Merged.19693
Merged.19696
Merged.19697
Merged.19698
Merged.19703
Merged.19711
Merged.19713
Merged.19714
Merged.19725
Merged.19726
Merged.19742
Merged.19743
Merged.19750
Merged.19754
Merged.19757
Merged.19758
Merged.19766
Merged.19770
Merged.19771
Merged.19794
Merged.19796
Merged.19797
Merged.19842
mean_tpm
time
preZGA
ZGA_early
ZGA_late
5h
9h
11h
14h
20h
24h
L1
sex
Female
Male

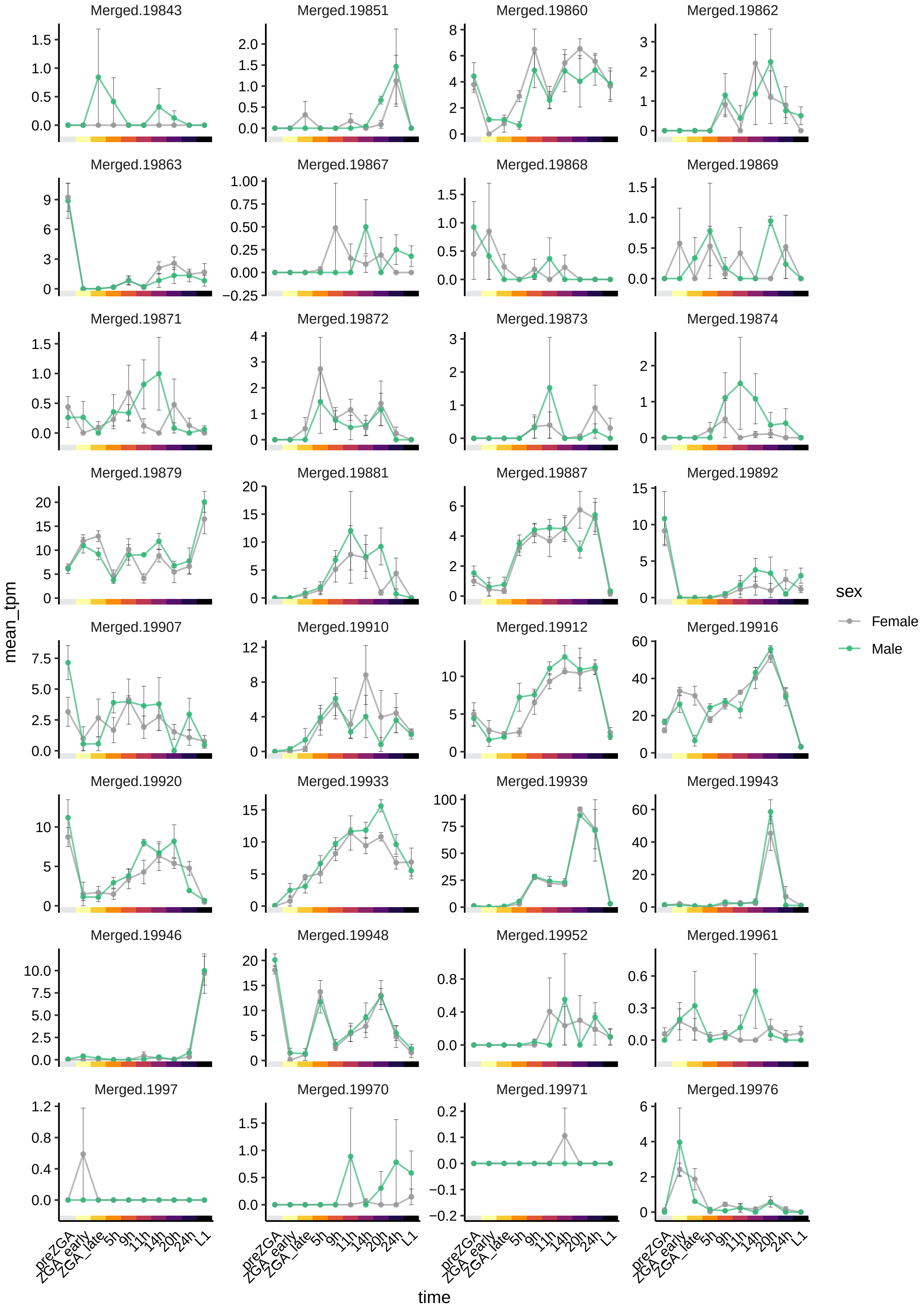
Merged.19843
Merged.19851
Merged.19860
Merged.19862
Merged.19863
Merged.19867
Merged.19868
Merged.19869
Merged.19871
Merged.19872
Merged.19873
Merged.19874
Merged.19879
Merged.19881
Merged.19887
Merged.19892
Merged.19907
Merged.19910
Merged.19912
Merged.19916
Merged.19920
Merged.19933
Merged.19939
Merged.19943
Merged.19946
Merged.19948
Merged.19952
Merged.19961
Merged.1997
Merged.19970
Merged.19971
Merged.19976
mean_tpm
time
preZGA ZGA_early ZGA_late 5h 9h 11h 14h 20h 24h L1
sex
Female
Male

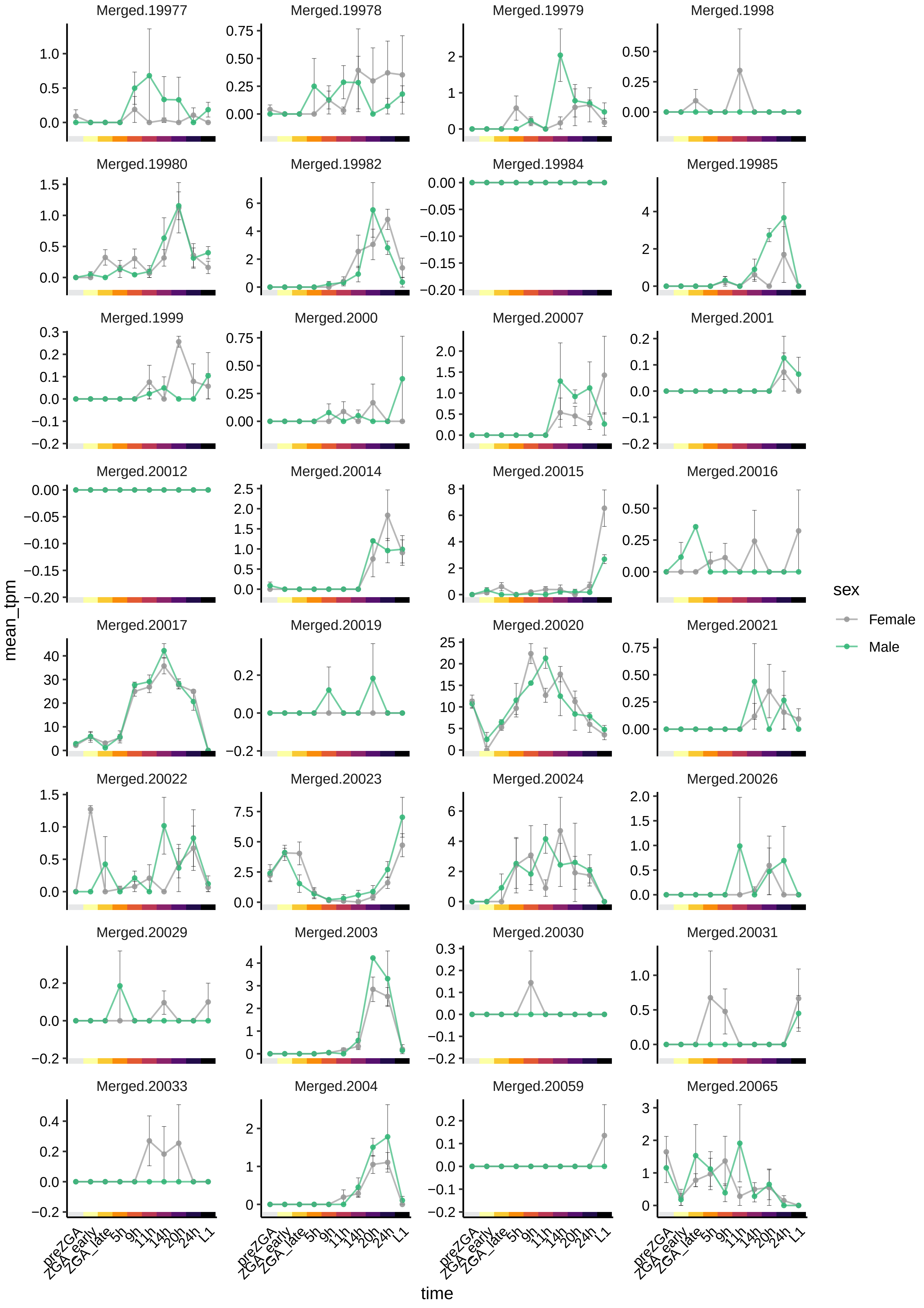
Merged.19977
Merged.19978
Merged.19979
Merged.1998
Merged.19980
Merged.19982
Merged.19984
Merged.19985
Merged.1999
Merged.2000
Merged.20007
Merged.2001
Merged.20012
Merged.20014
Merged.20015
Merged.20016
Merged.20017
Merged.20019
Merged.20020
Merged.20021
Merged.20022
Merged.20023
Merged.20024
Merged.20026
Merged.20029
Merged.2003
Merged.20030
Merged.20031
Merged.20033
Merged.2004
Merged.20059
Merged.20065
mean_tpm
time
preZGA
ZGA_early
ZGA_late
5h
9h
11h
14h
20h
24h
L1
sex
Female
Male

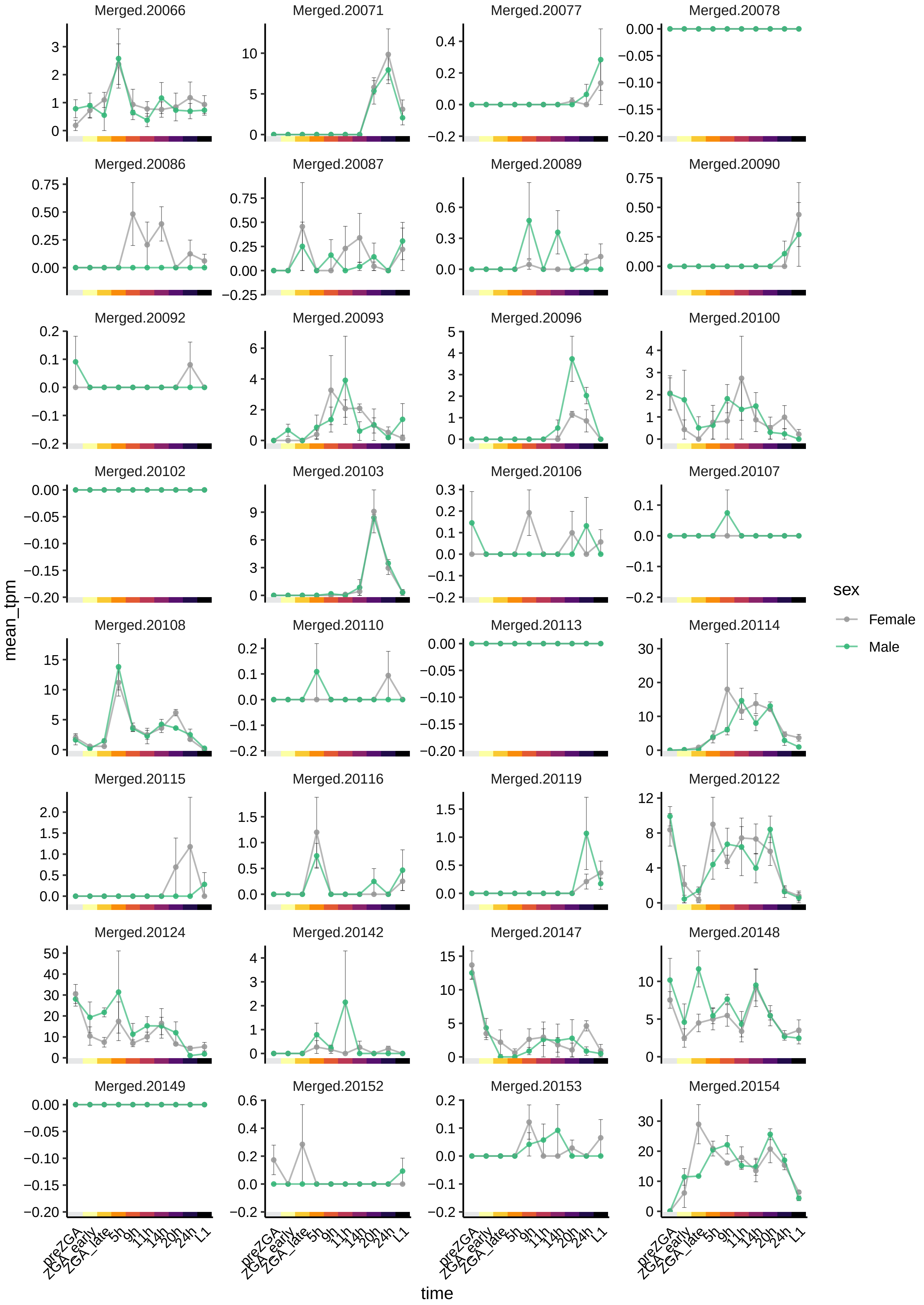

Merged.20066
Merged.20071
Merged.20077
Merged.20078
Merged.20086
Merged.20087
Merged.20089
Merged.20090
Merged.20092
Merged.20093
Merged.20096
Merged.20100
Merged.20102
Merged.20103
Merged.20106
Merged.20107
Merged.20108
Merged.20110
Merged.20113
Merged.20114
Merged.20115
Merged.20116
Merged.20119
Merged.20122
Merged.20124
Merged.20142
Merged.20147
Merged.20148
Merged.20149
Merged.20152
Merged.20153
Merged.20154
mean_tpm
time
preZGA
ZGA_early
ZGA_late
5h
9h
11h
14h
20h
24h
L1
sex
Female
Male

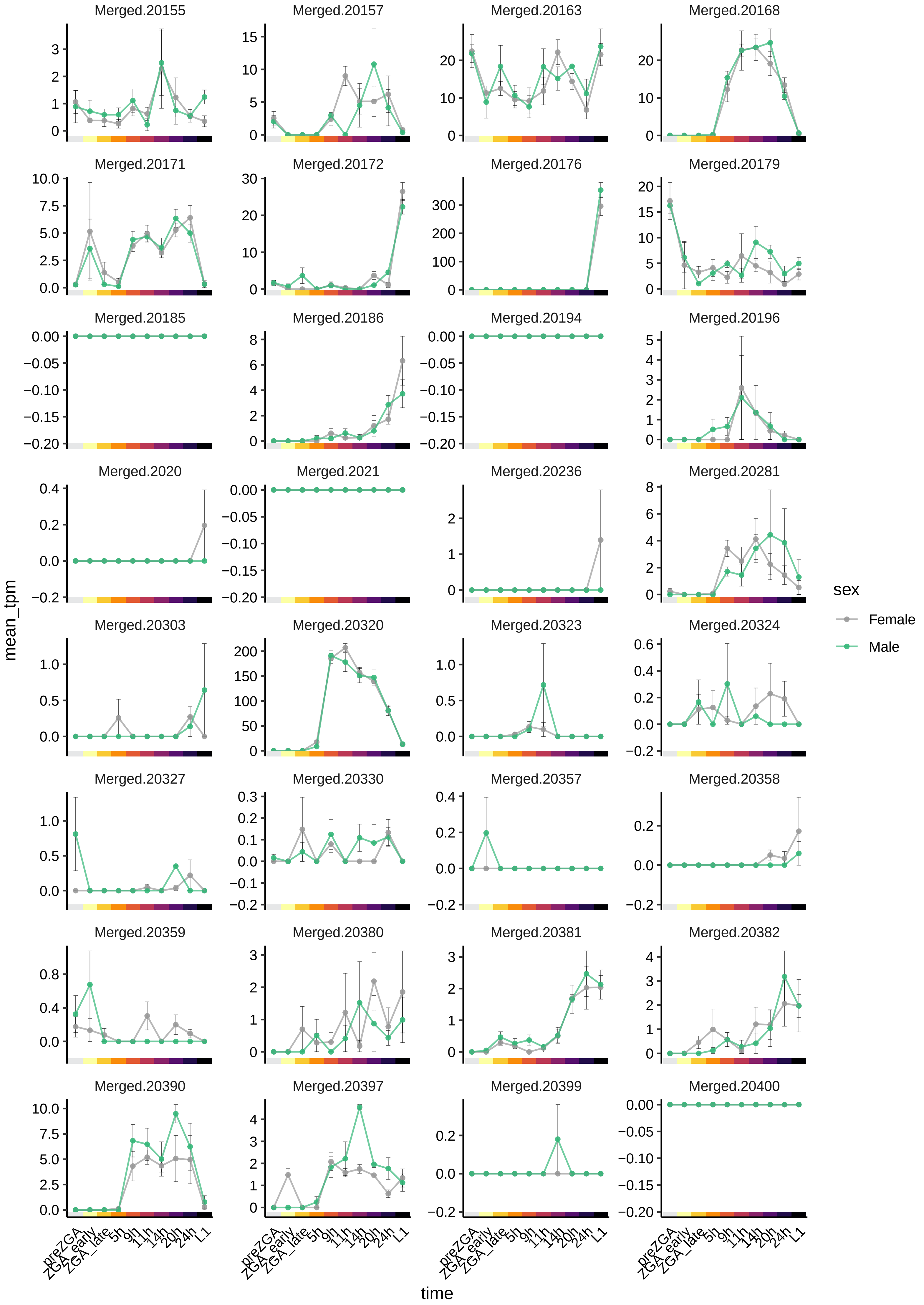

Merged.20155
Merged.20157
Merged.20163
Merged.20168
Merged.20171
Merged.20172
Merged.20176
Merged.20179
Merged.20185
Merged.20186
Merged.20194
Merged.20196
Merged.2020
Merged.2021
Merged.20236
Merged.20281
Merged.20303
Merged.20320
Merged.20323
Merged.20324
Merged.20327
Merged.20330
Merged.20357
Merged.20358
Merged.20359
Merged.20380
Merged.20381
Merged.20382
Merged.20390
Merged.20397
Merged.20399
Merged.20400
mean_tpm
time
preZGA
ZGA_early
ZGA_late
5h
9h
11h
14h
20h
24h
L1
sex
Female
Male

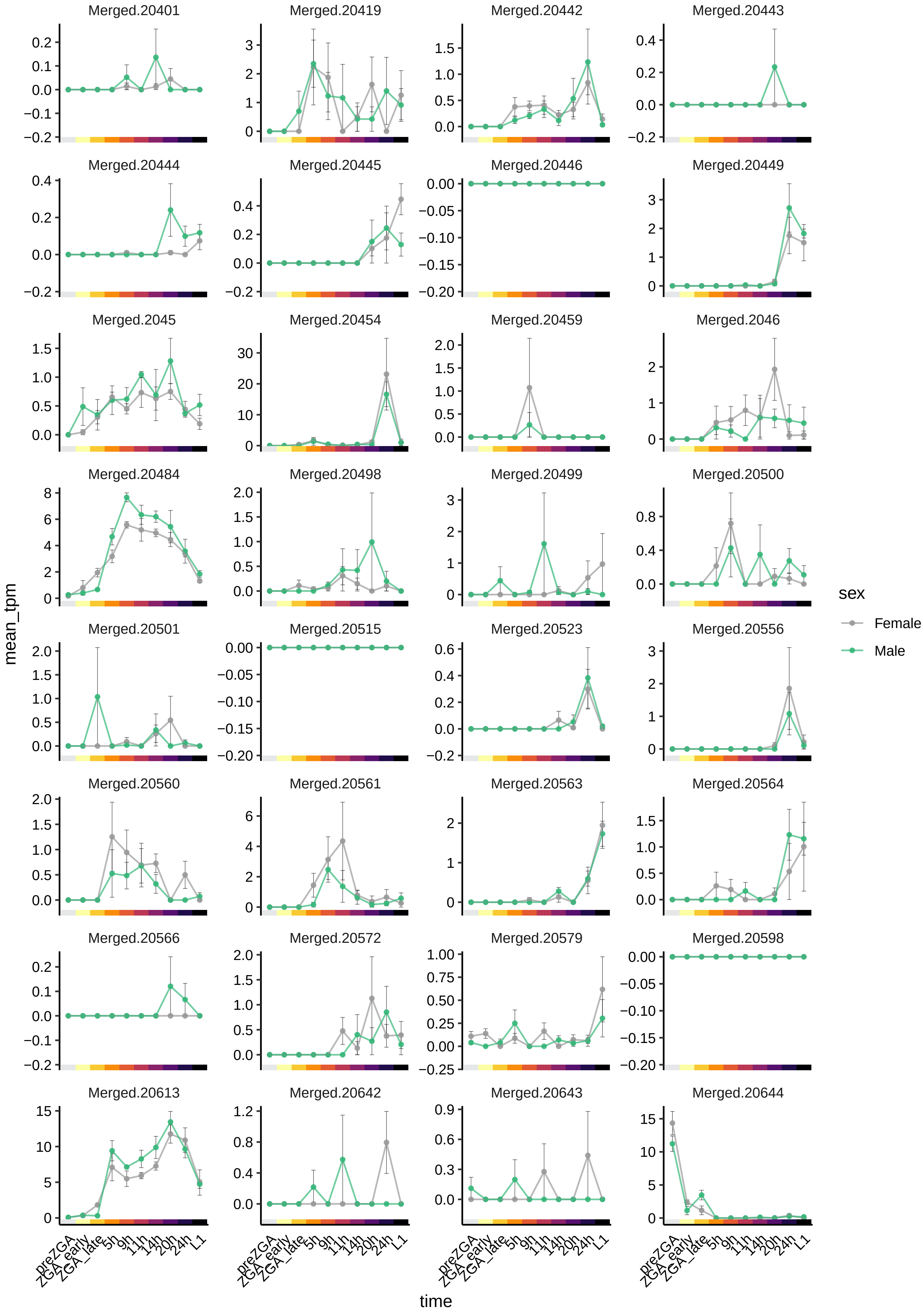
Merged.20401
Merged.20419
Merged.20442
Merged.20443
Merged.20444
Merged.20445
Merged.20446
Merged.20449
Merged.2045
Merged.20454
Merged.20459
Merged.2046
Merged.20484
Merged.20498
Merged.20499
Merged.20500
Merged.20501
Merged.20515
Merged.20523
Merged.20556
Merged.20560
Merged.20561
Merged.20563
Merged.20564
Merged.20566
Merged.20572
Merged.20579
Merged.20598
Merged.20613
Merged.20642
Merged.20643
Merged.20644
mean_tpm
time
preZGA
ZGA_early
ZGA_late
5h
9h
11h
14h
20h
24h
L1
sex
Female
Male

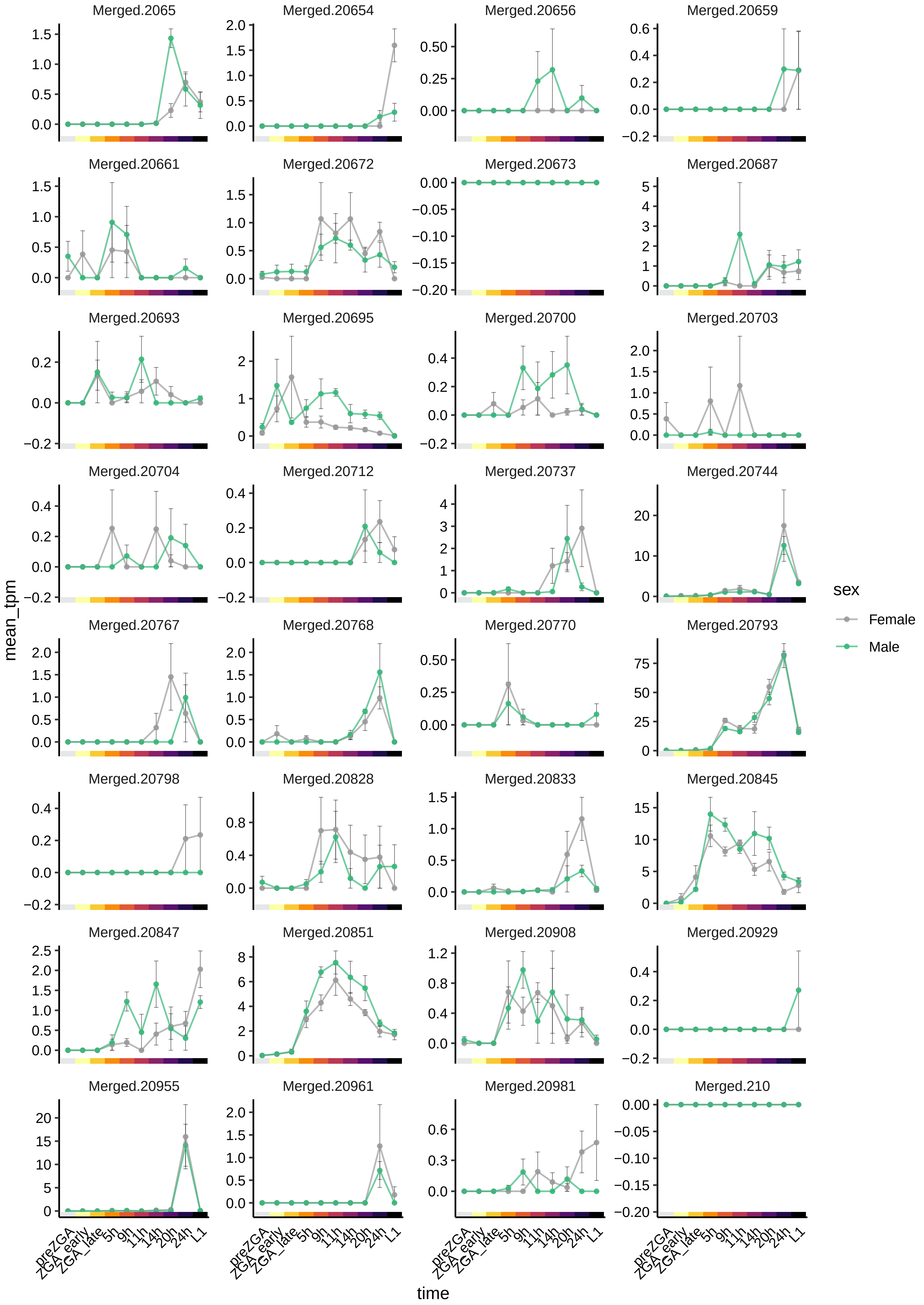

Merged.2065
Merged.20654
Merged.20656
Merged.20659
Merged.20661
Merged.20672
Merged.20673
Merged.20687
Merged.20693
Merged.20695
Merged.20700
Merged.20703
Merged.20704
Merged.20712
Merged.20737
Merged.20744
Merged.20767
Merged.20768
Merged.20770
Merged.20793
Merged.20798
Merged.20828
Merged.20833
Merged.20845
Merged.20847
Merged.20851
Merged.20908
Merged.20929
Merged.20955
Merged.20961
Merged.20981
Merged.210
mean_tpm
time
preZGA
ZGA_early
ZGA_late
5h
9h
11h
14h
20h
24h
L1
sex
Female
Male

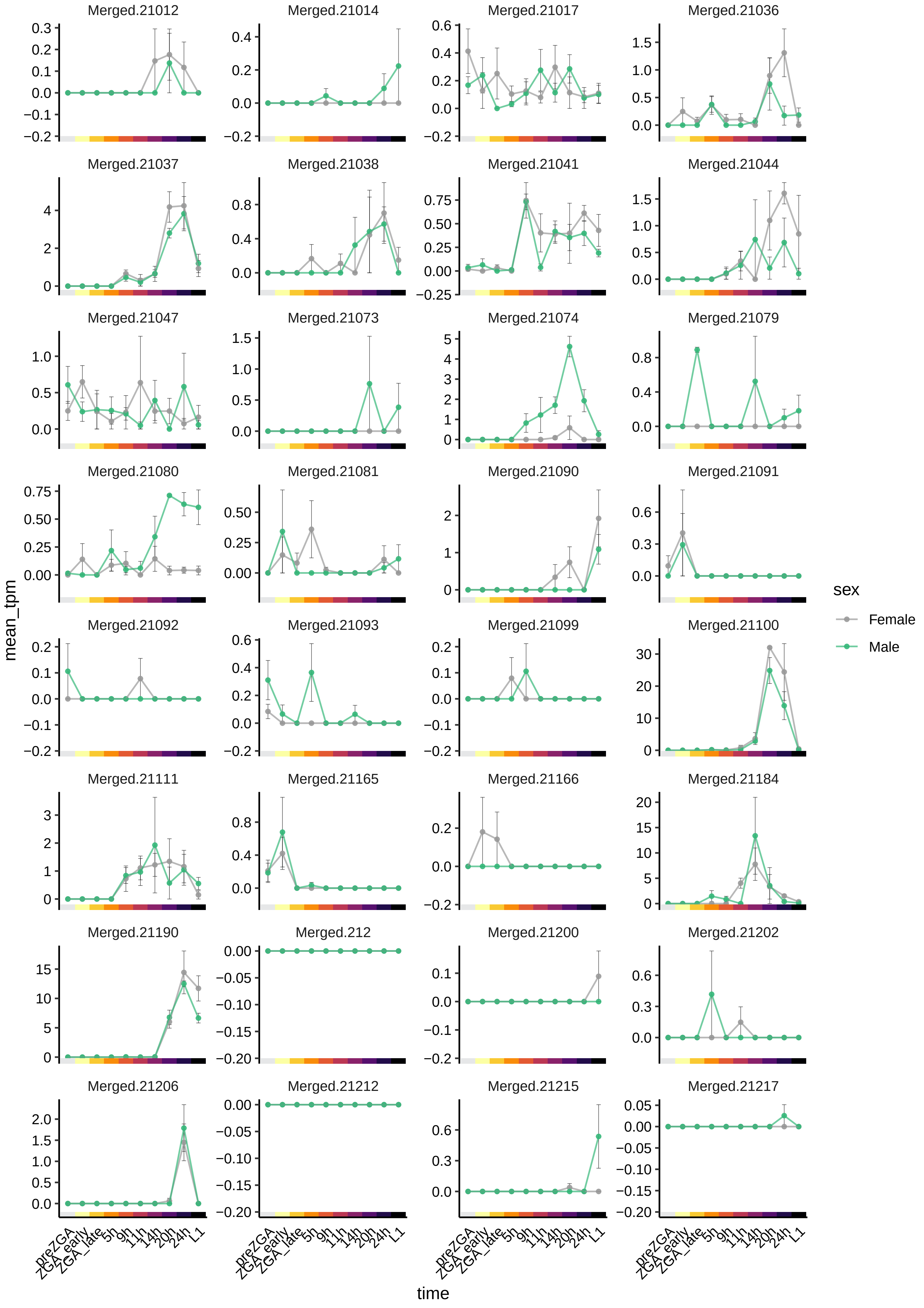

Merged.21012
Merged.21014
Merged.21017
Merged.21036
Merged.21037
Merged.21038
Merged.21041
Merged.21044
Merged.21047
Merged.21073
Merged.21074
Merged.21079
Merged.21080
Merged.21081
Merged.21090
Merged.21091
Merged.21092
Merged.21093
Merged.21099
Merged.21100
Merged.21111
Merged.21165
Merged.21166
Merged.21184
Merged.21190
Merged.212
Merged.21200
Merged.21202
Merged.21206
Merged.21212
Merged.21215
Merged.21217
mean_tpm
time
preZGA
ZGA_early
ZGA_late
5h
9h
11h
14h
20h
24h
L1
sex
Female
Male

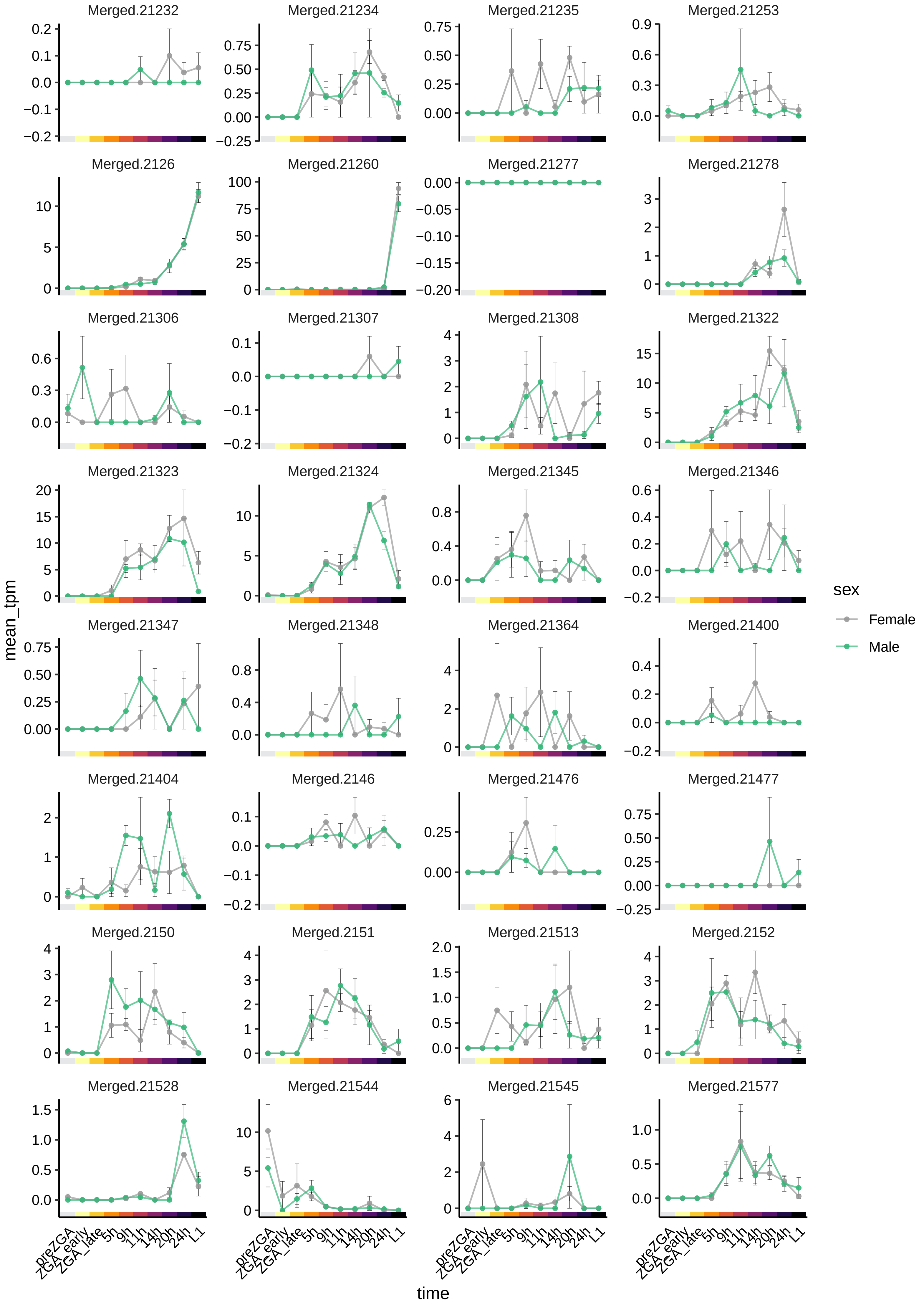

Merged.21232
Merged.21234
Merged.21235
Merged.21253
Merged.2126
Merged.21260
Merged.21277
Merged.21278
Merged.21306
Merged.21307
Merged.21308
Merged.21322
Merged.21323
Merged.21324
Merged.21345
Merged.21346
Merged.21347
Merged.21348
Merged.21364
Merged.21400
Merged.21404
Merged.2146
Merged.21476
Merged.21477
Merged.2150
Merged.2151
Merged.21513
Merged.2152
Merged.21528
Merged.21544
Merged.21545
Merged.21577
mean_tpm
time
preZGA
ZGA_early
ZGA_late
5h
9h
11h
14h
20h
24h
L1
sex
Female
Male

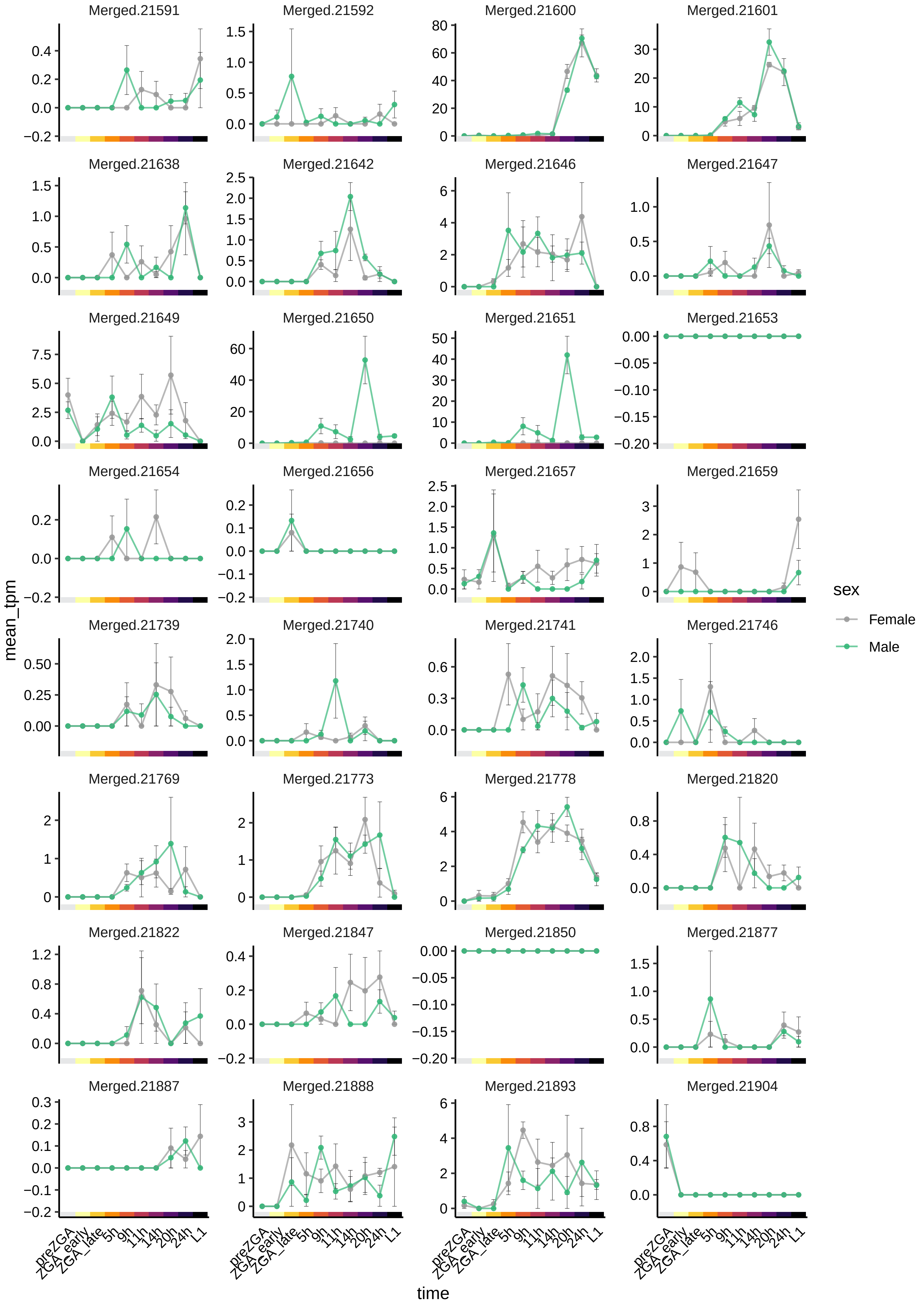
Merged.21591
Merged.21592
Merged.21600
Merged.21601
Merged.21638
Merged.21642
Merged.21646
Merged.21647
Merged.21649
Merged.21650
Merged.21651
Merged.21653
Merged.21654
Merged.21656
Merged.21657
Merged.21659
Merged.21739
Merged.21740
Merged.21741
Merged.21746
Merged.21769
Merged.21773
Merged.21778
Merged.21820
Merged.21822
Merged.21847
Merged.21850
Merged.21877
Merged.21887
Merged.21888
Merged.21893
Merged.21904
mean_tpm
time
preZGA
ZGA_early
ZGA_late
5h
9h
11h
14h
20h
24h
L1
sex
Female
Male

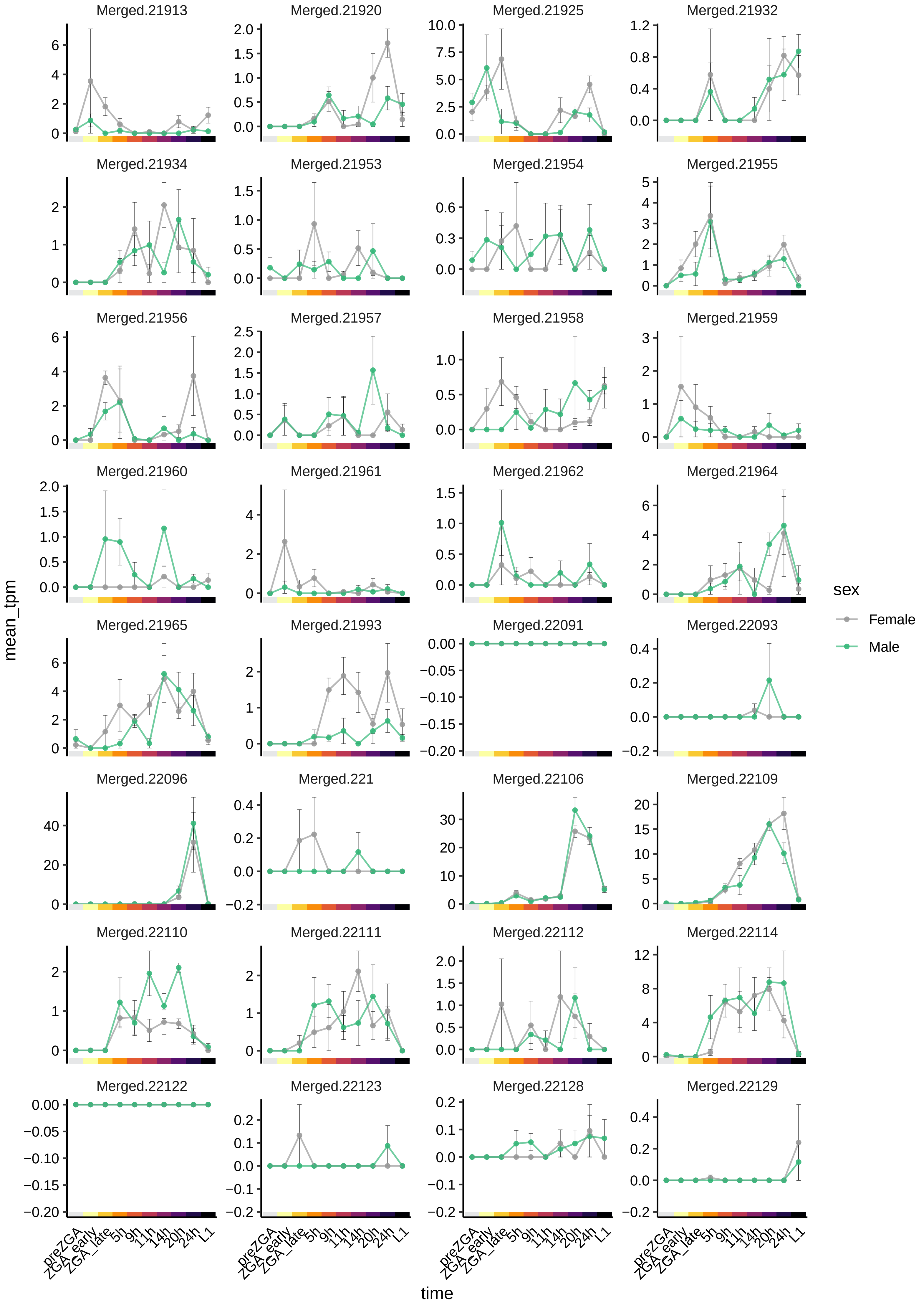

Merged.21913
Merged.21920
Merged.21925
Merged.21932
Merged.21934
Merged.21953
Merged.21954
Merged.21955
Merged.21956
Merged.21957
Merged.21958
Merged.21959
Merged.21960
Merged.21961
Merged.21962
Merged.21964
Merged.21965
Merged.21993
Merged.22091
Merged.22093
Merged.22096
Merged.221
Merged.22106
Merged.22109
Merged.22110
Merged.22111
Merged.22112
Merged.22114
Merged.22122
Merged.22123
Merged.22128
Merged.22129
mean_tpm
time
preZGA
ZGA_early
ZGA_late
5h
9h
11h
14h
20h
24h
L1
sex
Female
Male

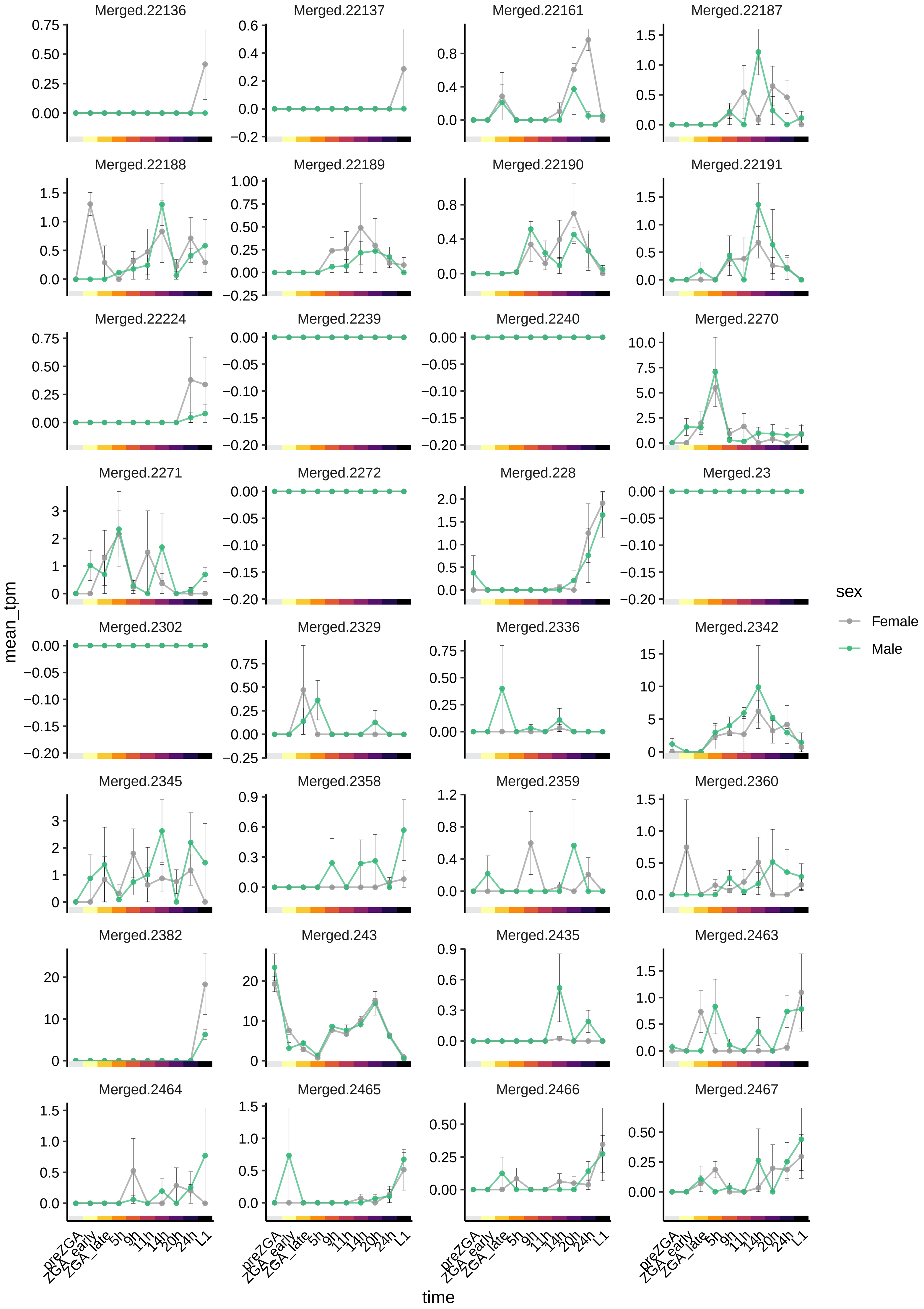

Merged.22136
Merged.22137
Merged.22161
Merged.22187
Merged.22188
Merged.22189
Merged.22190
Merged.22191
Merged.22224
Merged.2239
Merged.2240
Merged.2270
Merged.2271
Merged.2272
Merged.228
Merged.23
Merged.2302
Merged.2329
Merged.2336
Merged.2342
Merged.2345
Merged.2358
Merged.2359
Merged.2360
Merged.2382
Merged.243
Merged.2435
Merged.2463
Merged.2464
Merged.2465
Merged.2466
Merged.2467
mean_tpm
time
preZGA
ZGA_early
ZGA_late
5h
9h
11h
14h
20h
24h
L1
sex
Female
Male

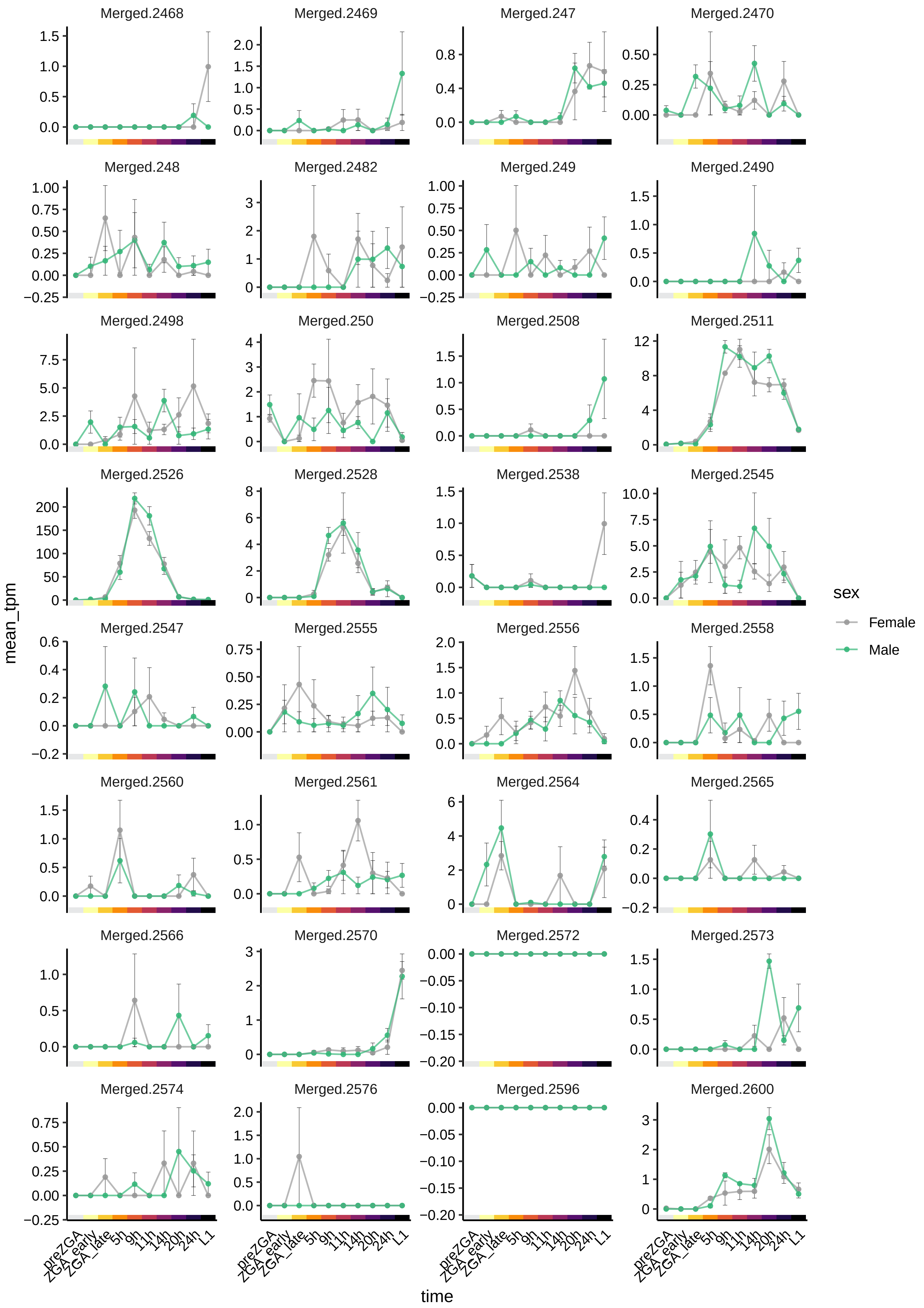

Merged.2468
Merged.2469
Merged.247
Merged.2470
Merged.248
Merged.2482
Merged.249
Merged.2490
Merged.2498
Merged.250
Merged.2508
Merged.2511
Merged.2526
Merged.2528
Merged.2538
Merged.2545
Merged.2547
Merged.2555
Merged.2556
Merged.2558
Merged.2560
Merged.2561
Merged.2564
Merged.2565
Merged.2566
Merged.2570
Merged.2572
Merged.2573
Merged.2574
Merged.2576
Merged.2596
Merged.2600
mean_tpm
time
preZGA
ZGA_early
ZGA_late
5h
9h
11h
14h
20h
24h
L1
sex
Female
Male

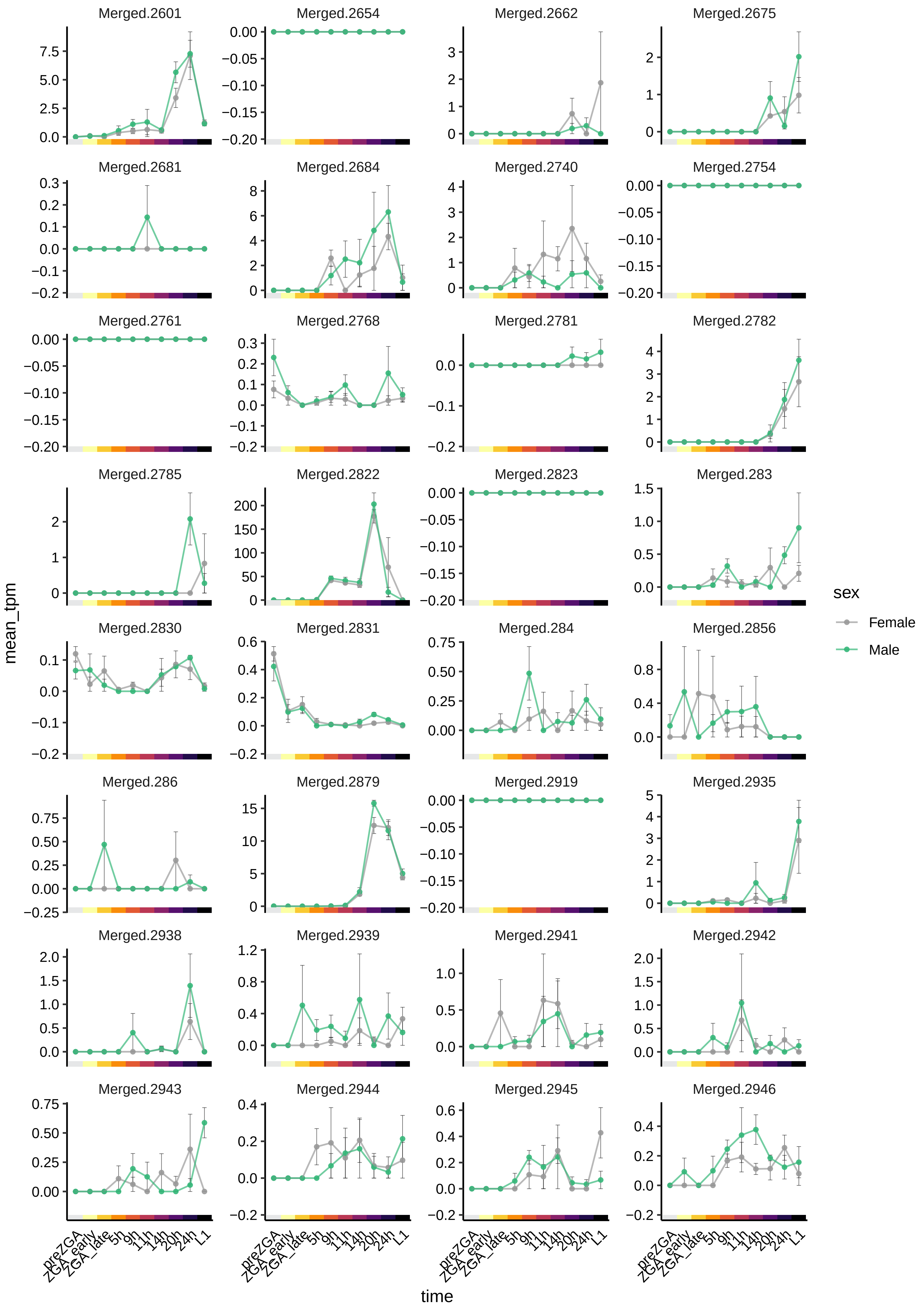
Merged.2601
Merged.2654
Merged.2662
Merged.2675
Merged.2681
Merged.2684
Merged.2740
Merged.2754
Merged.2761
Merged.2768
Merged.2781
Merged.2782
Merged.2785
Merged.2822
Merged.2823
Merged.283
Merged.2830
Merged.2831
Merged.284
Merged.2856
Merged.286
Merged.2879
Merged.2919
Merged.2935
Merged.2938
Merged.2939
Merged.2941
Merged.2942
Merged.2943
Merged.2944
Merged.2945
Merged.2946
mean_tpm
time
preZGA
ZGA_early
ZGA_late
5h
9h
11h
14h
20h
24h
L1
sex
Female
Male

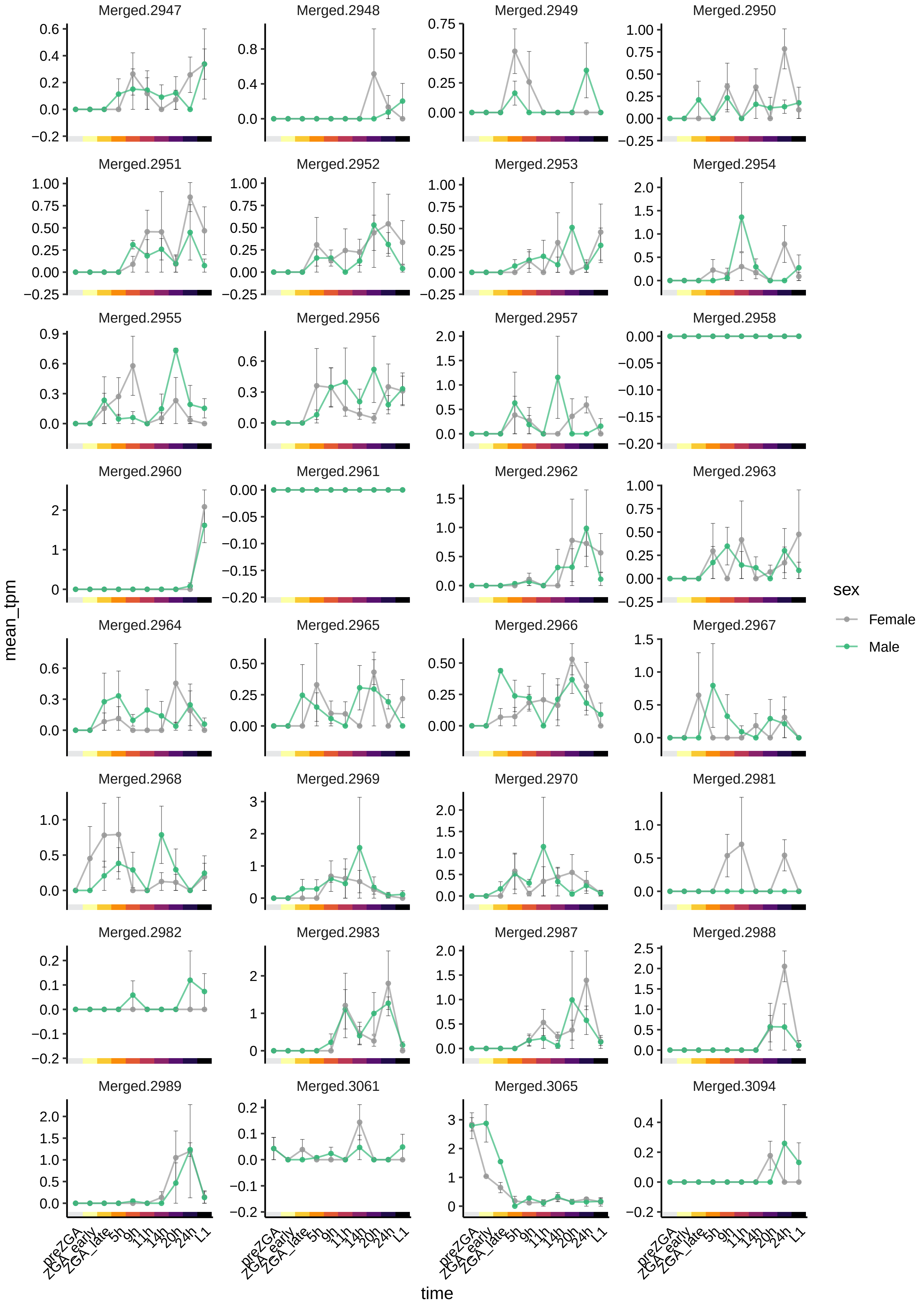

Merged.2947
Merged.2948
Merged.2949
Merged.2950
Merged.2951
Merged.2952
Merged.2953
Merged.2954
Merged.2955
Merged.2956
Merged.2957
Merged.2958
Merged.2960
Merged.2961
Merged.2962
Merged.2963
Merged.2964
Merged.2965
Merged.2966
Merged.2967
Merged.2968
Merged.2969
Merged.2970
Merged.2981
Merged.2982
Merged.2983
Merged.2987
Merged.2988
Merged.2989
Merged.3061
Merged.3065
Merged.3094
mean_tpm
time
preZGA
ZGA_early
ZGA_late
5h
9h
11h
14h
20h
24h
L1
sex
Female
Male

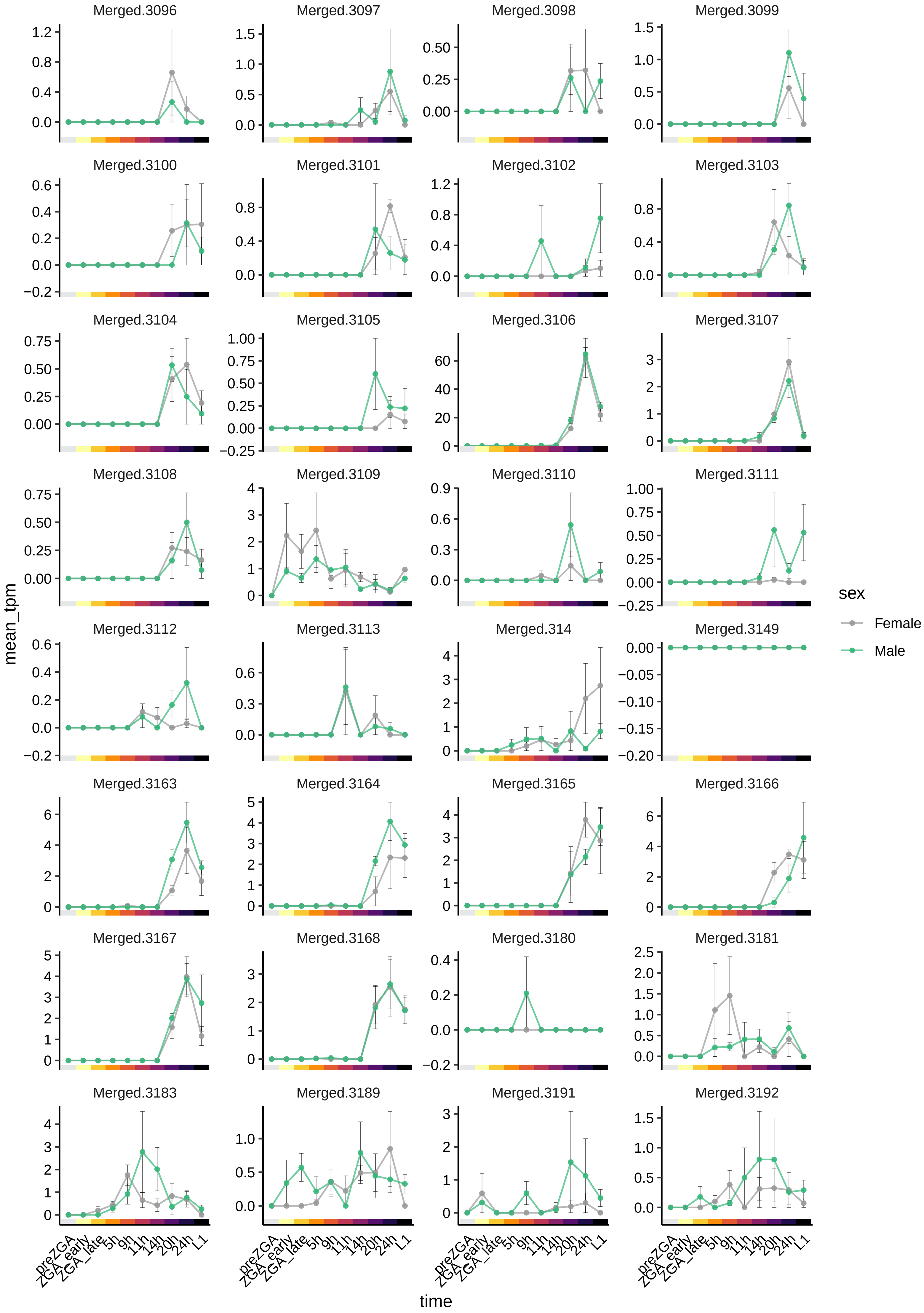

Merged.3096
Merged.3097
Merged.3098
Merged.3099
Merged.3100
Merged.3101
Merged.3102
Merged.3103
Merged.3104
Merged.3105
Merged.3106
Merged.3107
Merged.3108
Merged.3109
Merged.3110
Merged.3111
Merged.3112
Merged.3113
Merged.314
Merged.3149
Merged.3163
Merged.3164
Merged.3165
Merged.3166
Merged.3167
Merged.3168
Merged.3180
Merged.3181
Merged.3183
Merged.3189
Merged.3191
Merged.3192
mean_tpm
time
preZGA
ZGA_early
ZGA_late
5h
9h
11h
14h
20h
24h
L1
sex
Female
Male

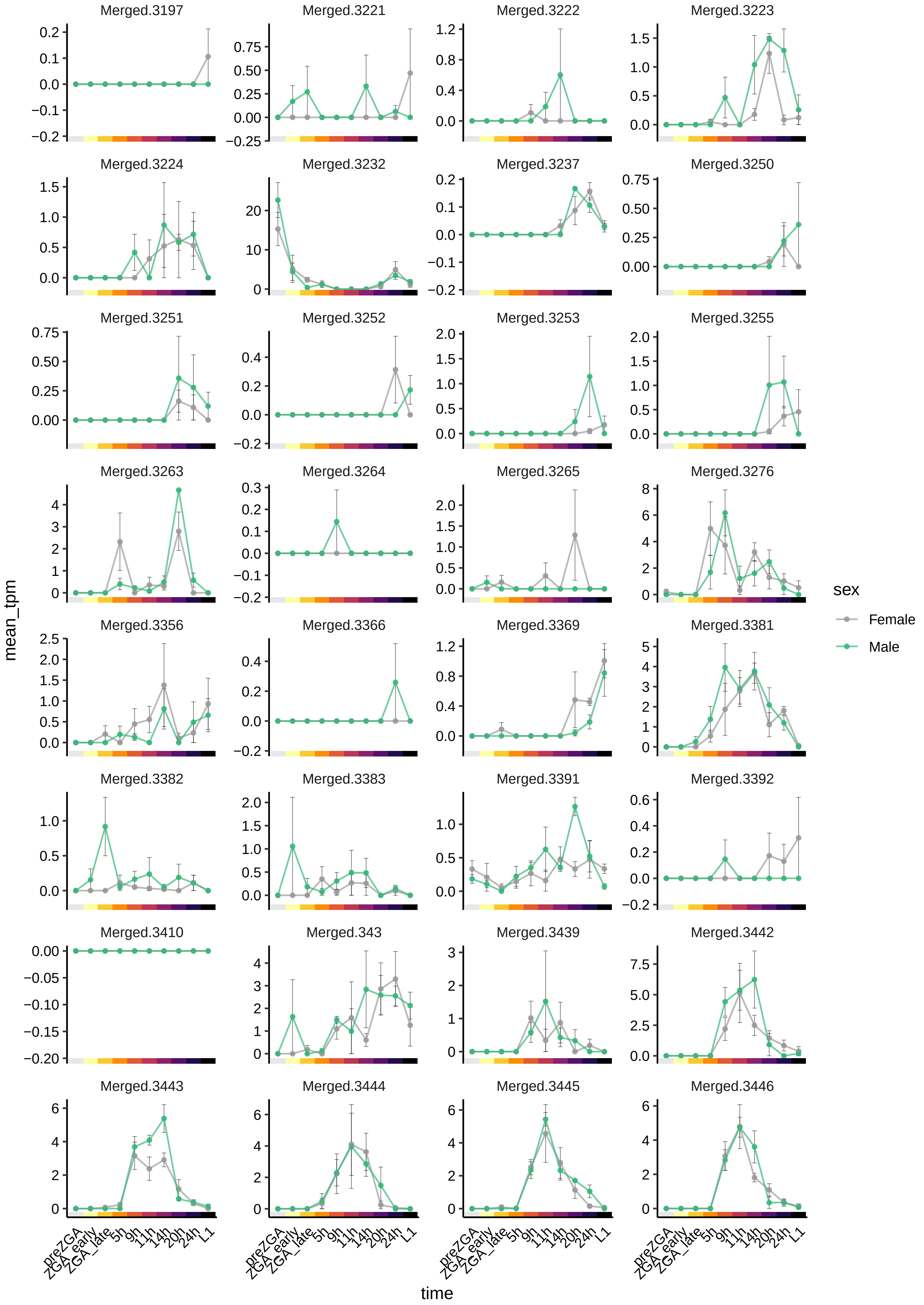
Merged.3197
Merged.3221
Merged.3222
Merged.3223
Merged.3224
Merged.3232
Merged.3237
Merged.3250
Merged.3251
Merged.3252
Merged.3253
Merged.3255
Merged.3263
Merged.3264
Merged.3265
Merged.3276
Merged.3356
Merged.3366
Merged.3369
Merged.3381
Merged.3382
Merged.3383
Merged.3391
Merged.3392
Merged.3410
Merged.343
Merged.3439
Merged.3442
Merged.3443
Merged.3444
Merged.3445
Merged.3446
mean_tpm
time
preZGA
ZGA_early
ZGA_late
5h
9h
11h
14h
20h
24h
L1
sex
Female
Male

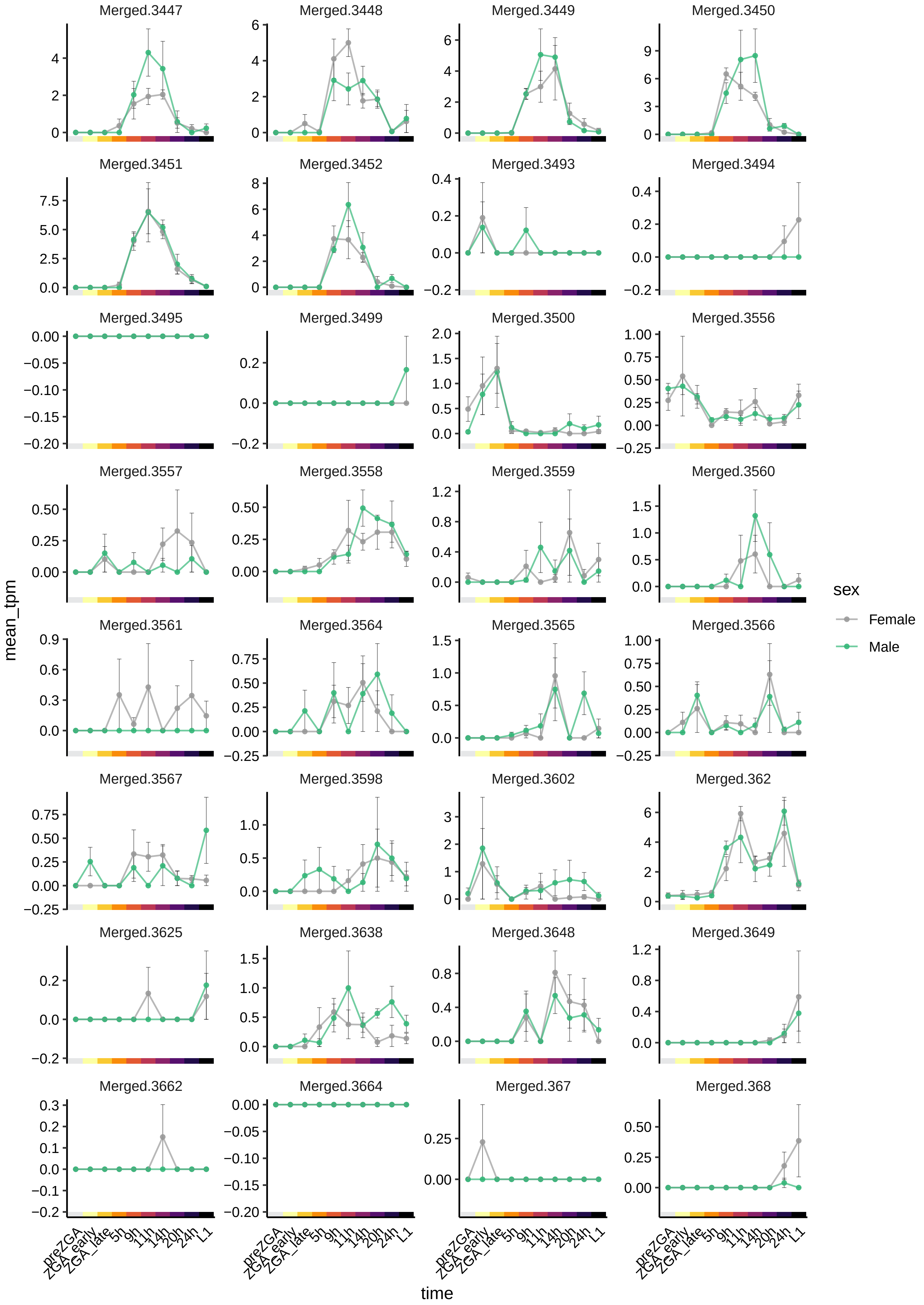

Merged.3447
Merged.3448
Merged.3449
Merged.3450
Merged.3451
Merged.3452
Merged.3493
Merged.3494
Merged.3495
Merged.3499
Merged.3500
Merged.3556
Merged.3557
Merged.3558
Merged.3559
Merged.3560
Merged.3561
Merged.3564
Merged.3565
Merged.3566
Merged.3567
Merged.3598
Merged.3602
Merged.362
Merged.3625
Merged.3638
Merged.3648
Merged.3649
Merged.3662
Merged.3664
Merged.367
Merged.368
mean_tpm
time
preZGA
ZGA_early
ZGA_late
5h
9h
11h
14h
20h
24h
L1
sex
Female
Male

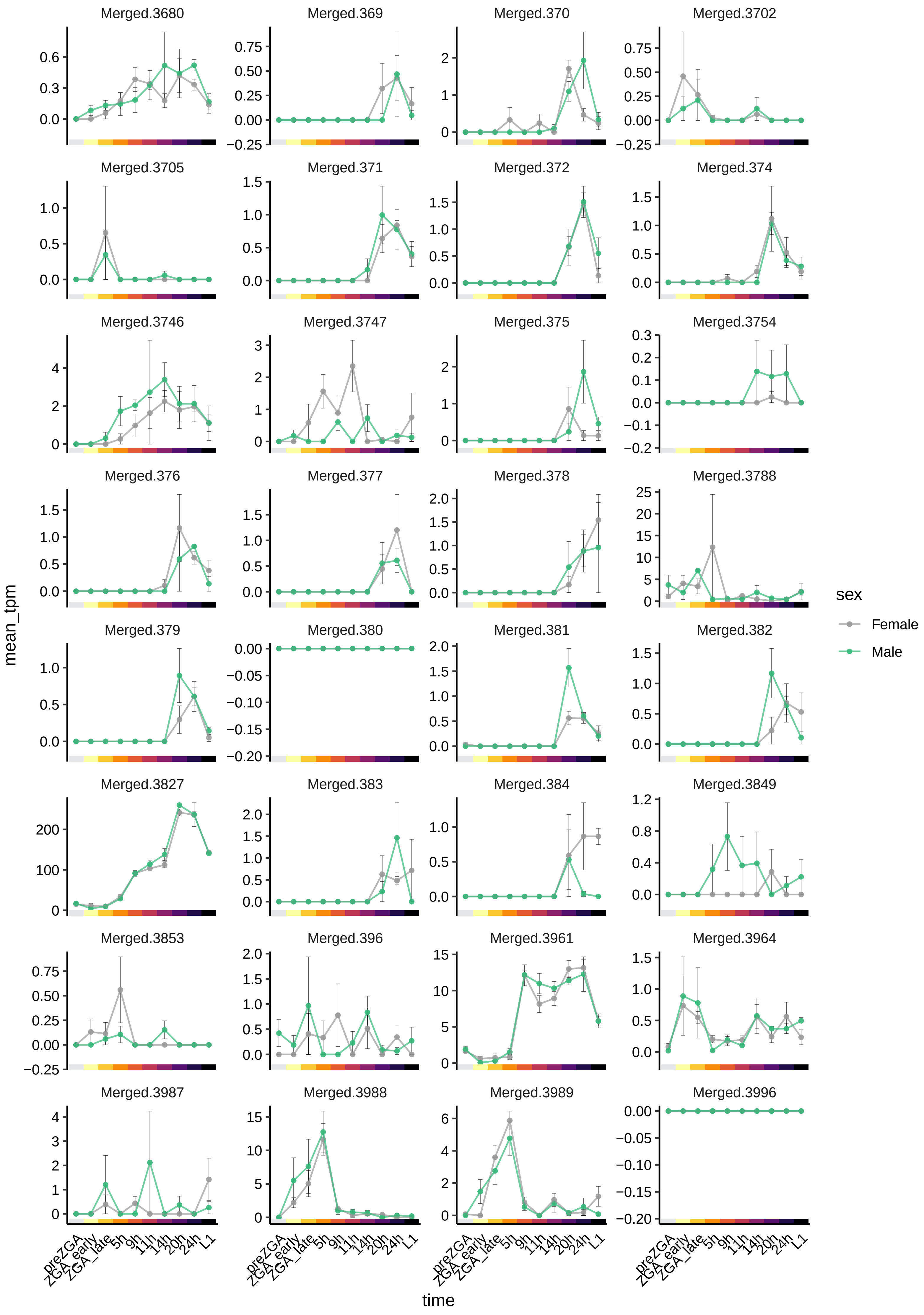

Merged.3680
Merged.369
Merged.370
Merged.3702
Merged.3705
Merged.371
Merged.372
Merged.374
Merged.3746
Merged.3747
Merged.375
Merged.3754
Merged.376
Merged.377
Merged.378
Merged.3788
Merged.379
Merged.380
Merged.381
Merged.382
Merged.3827
Merged.383
Merged.384
Merged.3849
Merged.3853
Merged.396
Merged.3961
Merged.3964
Merged.3987
Merged.3988
Merged.3989
Merged.3996
mean_tpm
time
preZGA
ZGA_early
ZGA_late
5h
9h
11h
14h
20h
24h
L1
sex
Female
Male

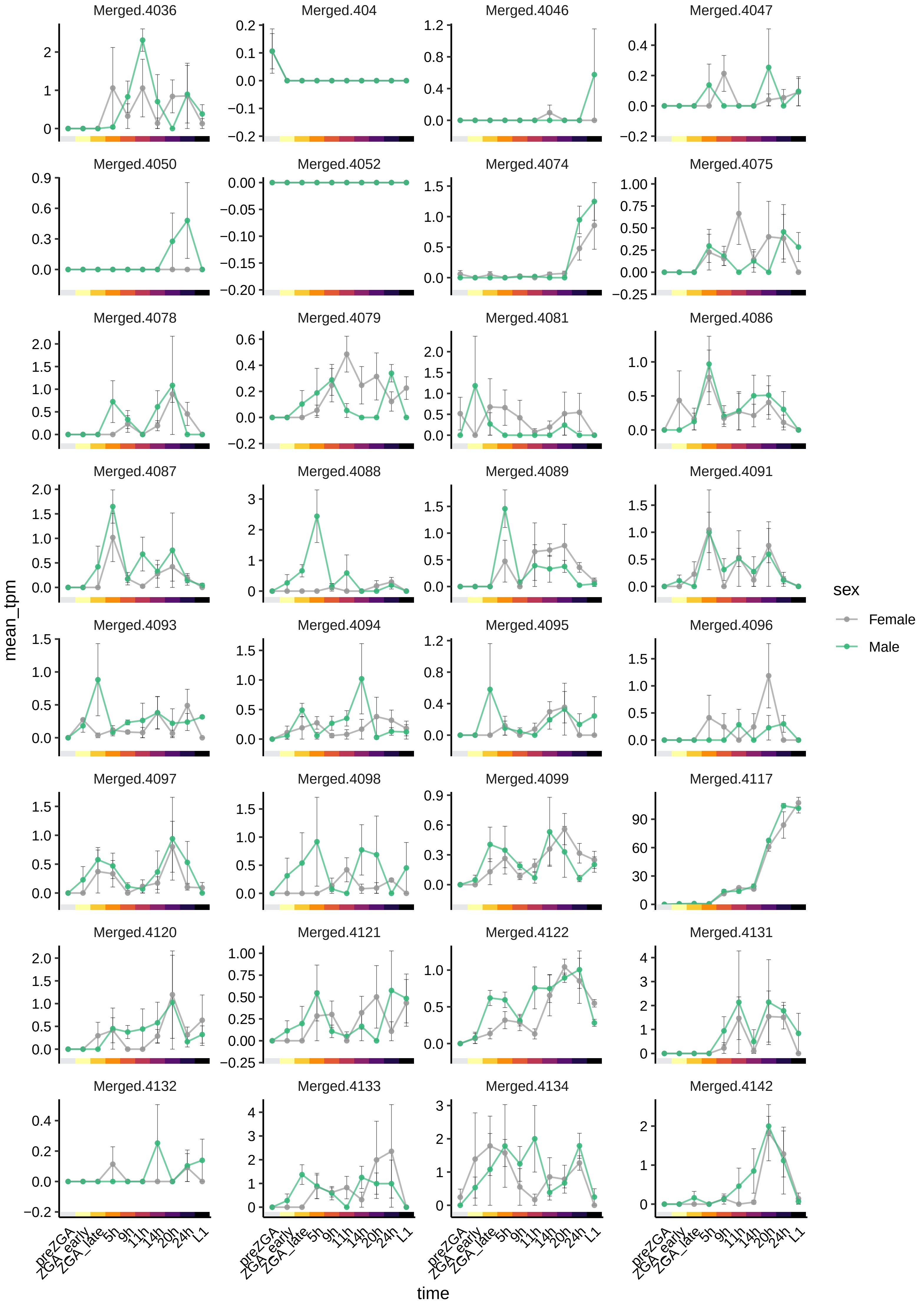
Merged.4036
Merged.404
Merged.4046
Merged.4047
Merged.4050
Merged.4052
Merged.4074
Merged.4075
Merged.4078
Merged.4079
Merged.4081
Merged.4086
Merged.4087
Merged.4088
Merged.4089
Merged.4091
Merged.4093
Merged.4094
Merged.4095
Merged.4096
Merged.4097
Merged.4098
Merged.4099
Merged.4117
Merged.4120
Merged.4121
Merged.4122
Merged.4131
Merged.4132
Merged.4133
Merged.4134
Merged.4142
mean_tpm
time
preZGA
ZGA_early
ZGA_late
5h
9h
11h
14h
20h
24h
L1
sex
Female
Male

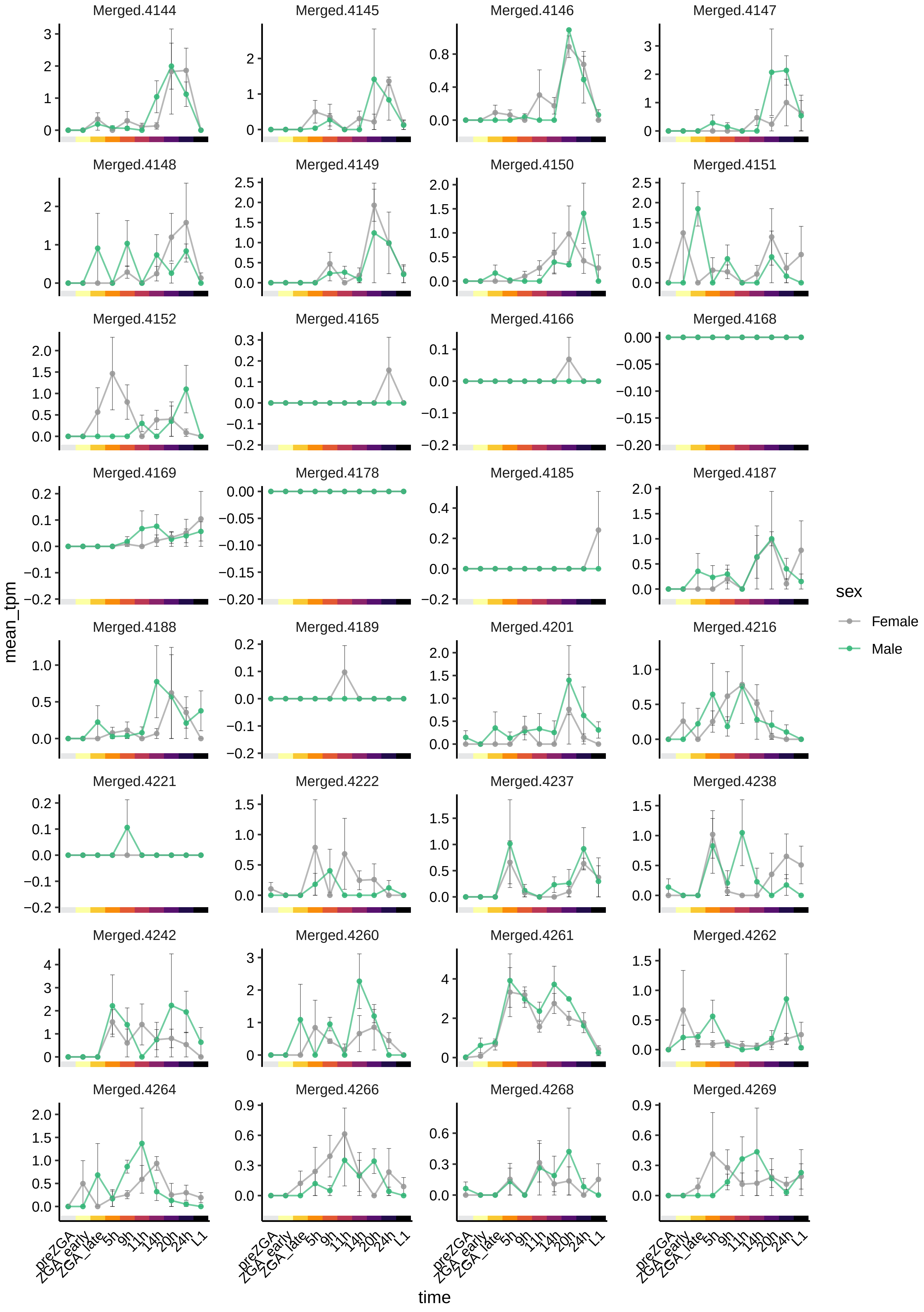

Merged.4144
Merged.4145
Merged.4146
Merged.4147
Merged.4148
Merged.4149
Merged.4150
Merged.4151
Merged.4152
Merged.4165
Merged.4166
Merged.4168
Merged.4169
Merged.4178
Merged.4185
Merged.4187
Merged.4188
Merged.4189
Merged.4201
Merged.4216
Merged.4221
Merged.4222
Merged.4237
Merged.4238
Merged.4242
Merged.4260
Merged.4261
Merged.4262
Merged.4264
Merged.4266
Merged.4268
Merged.4269
mean_tpm
time
preZGA
ZGA_early
ZGA_late
5h
9h
11h
14h
20h
24h
L1
sex
Female
Male

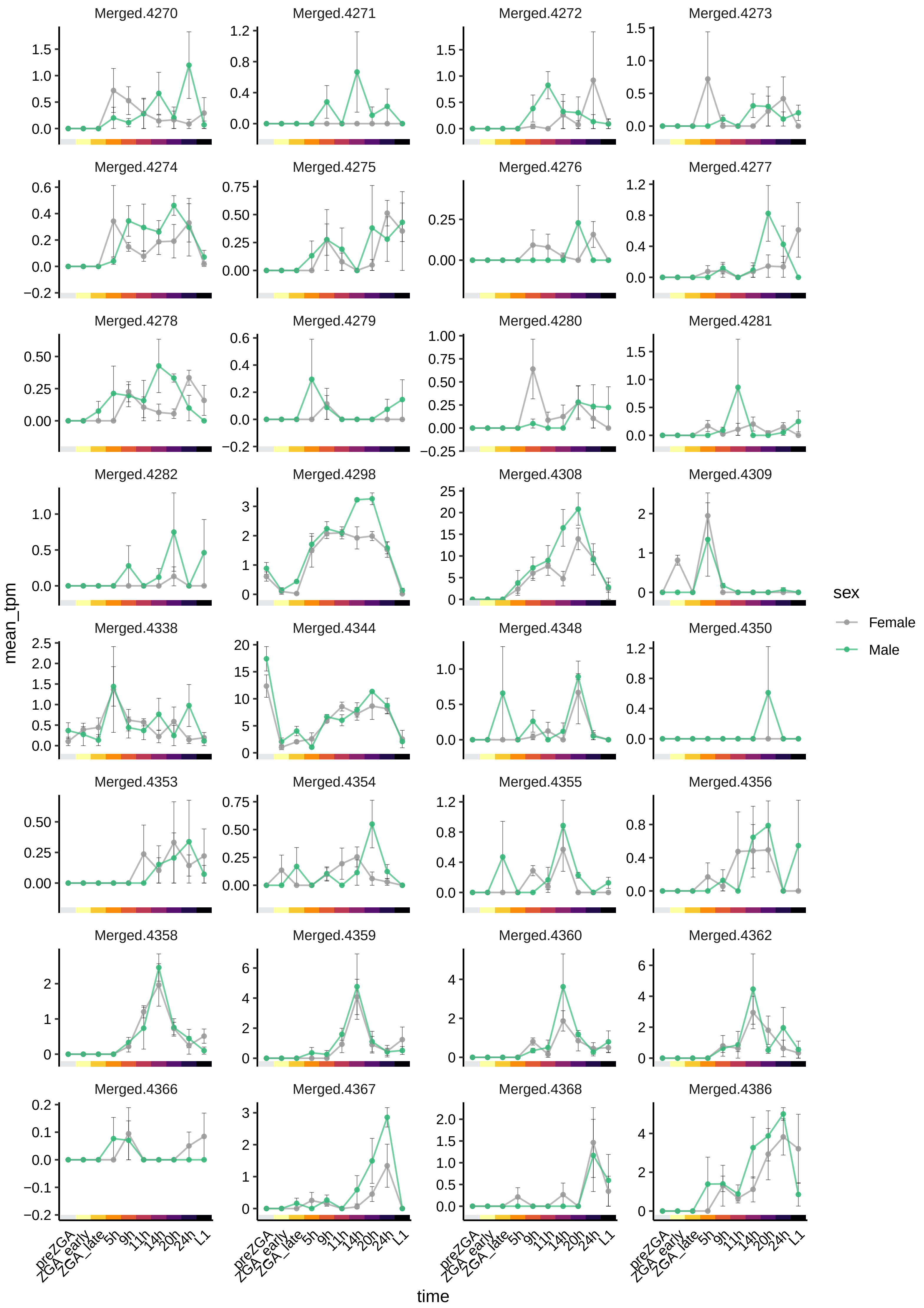

Merged.4270
Merged.4271
Merged.4272
Merged.4273
Merged.4274
Merged.4275
Merged.4276
Merged.4277
Merged.4278
Merged.4279
Merged.4280
Merged.4281
Merged.4282
Merged.4298
Merged.4308
Merged.4309
Merged.4338
Merged.4344
Merged.4348
Merged.4350
Merged.4353
Merged.4354
Merged.4355
Merged.4356
Merged.4358
Merged.4359
Merged.4360
Merged.4362
Merged.4366
Merged.4367
Merged.4368
Merged.4386
mean_tpm
time
preZGA
ZGA_early
ZGA_late
5h
9h
11h
14h
20h
24h
L1
sex
Female
Male

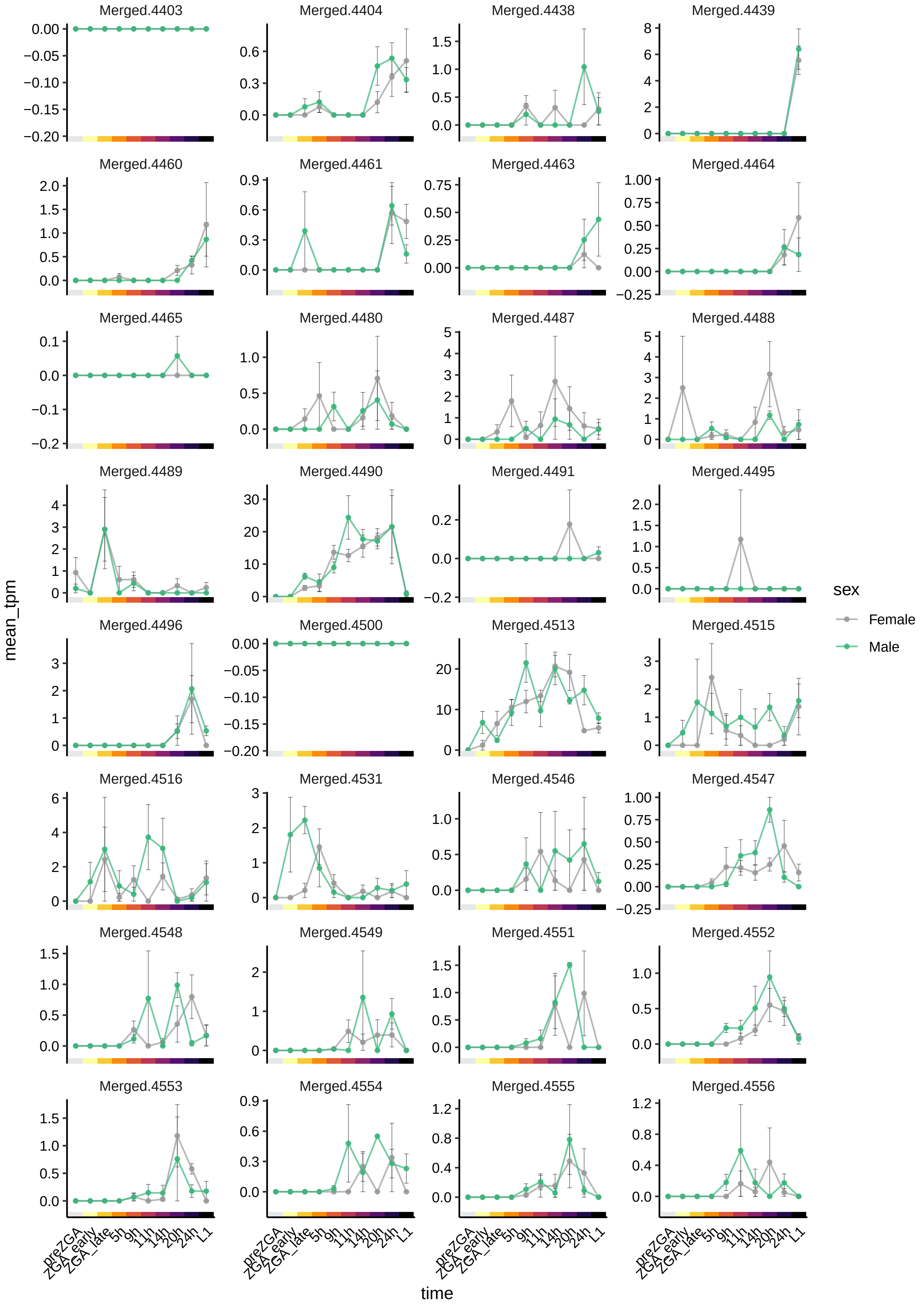
Merged.4403
Merged.4404
Merged.4438
Merged.4439
Merged.4460
Merged.4461
Merged.4463
Merged.4464
Merged.4465
Merged.4480
Merged.4487
Merged.4488
Merged.4489
Merged.4490
Merged.4491
Merged.4495
Merged.4496
Merged.4500
Merged.4513
Merged.4515
Merged.4516
Merged.4531
Merged.4546
Merged.4547
Merged.4548
Merged.4549
Merged.4551
Merged.4552
Merged.4553
Merged.4554
Merged.4555
Merged.4556
mean_tpm
time
preZGA
ZGA_early
ZGA_late
5h
9h
11h
14h
20h
24h
L1
sex
Female
Male

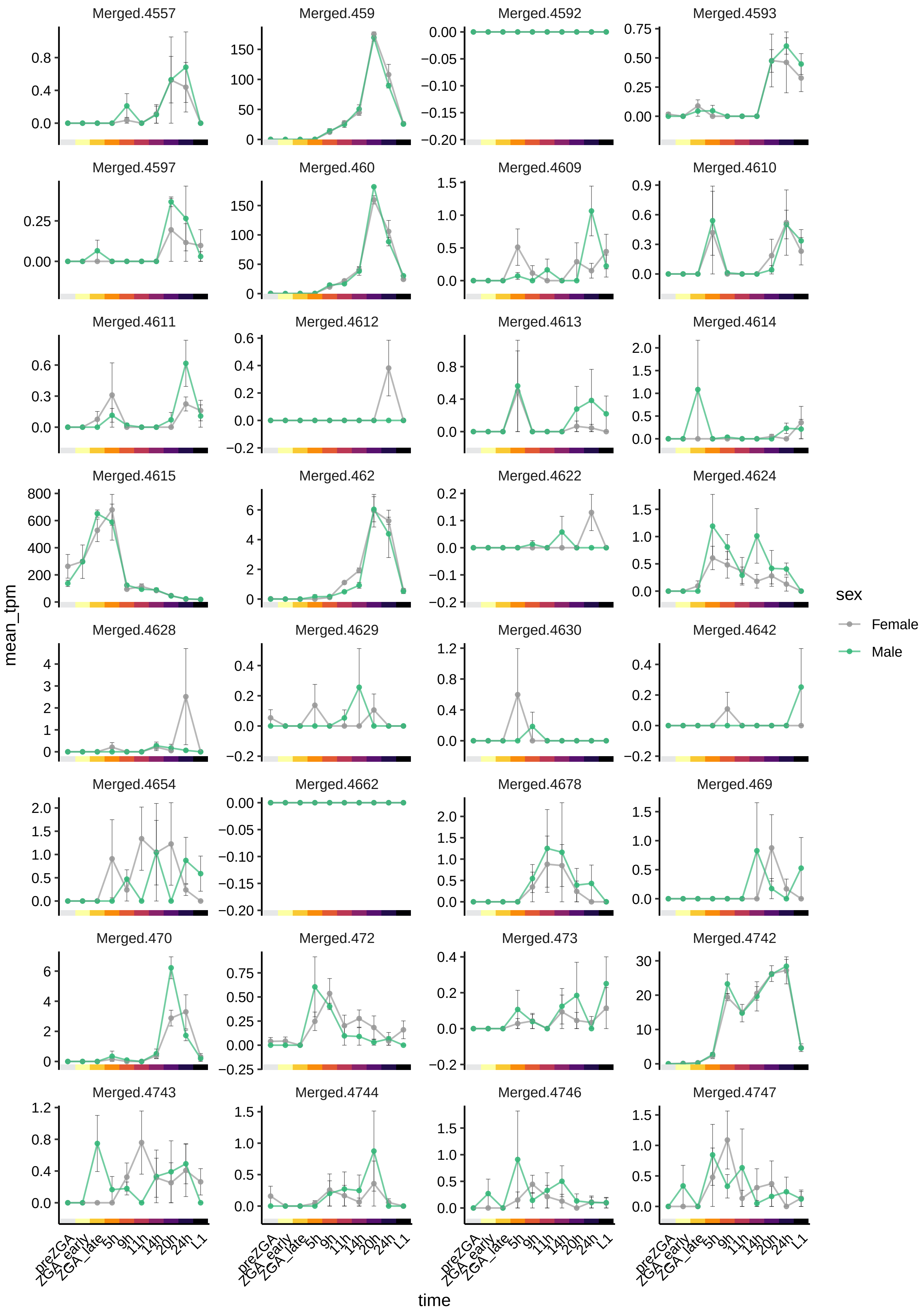

Merged.4557
Merged.459
Merged.4592
Merged.4593
Merged.4597
Merged.460
Merged.4609
Merged.4610
Merged.4611
Merged.4612
Merged.4613
Merged.4614
Merged.4615
Merged.462
Merged.4622
Merged.4624
Merged.4628
Merged.4629
Merged.4630
Merged.4642
Merged.4654
Merged.4662
Merged.4678
Merged.469
Merged.470
Merged.472
Merged.473
Merged.4742
Merged.4743
Merged.4744
Merged.4746
Merged.4747
mean_tpm
time
preZGA
ZGA_early
ZGA_late
5h
9h
11h
14h
20h
24h
L1
sex
Female
Male

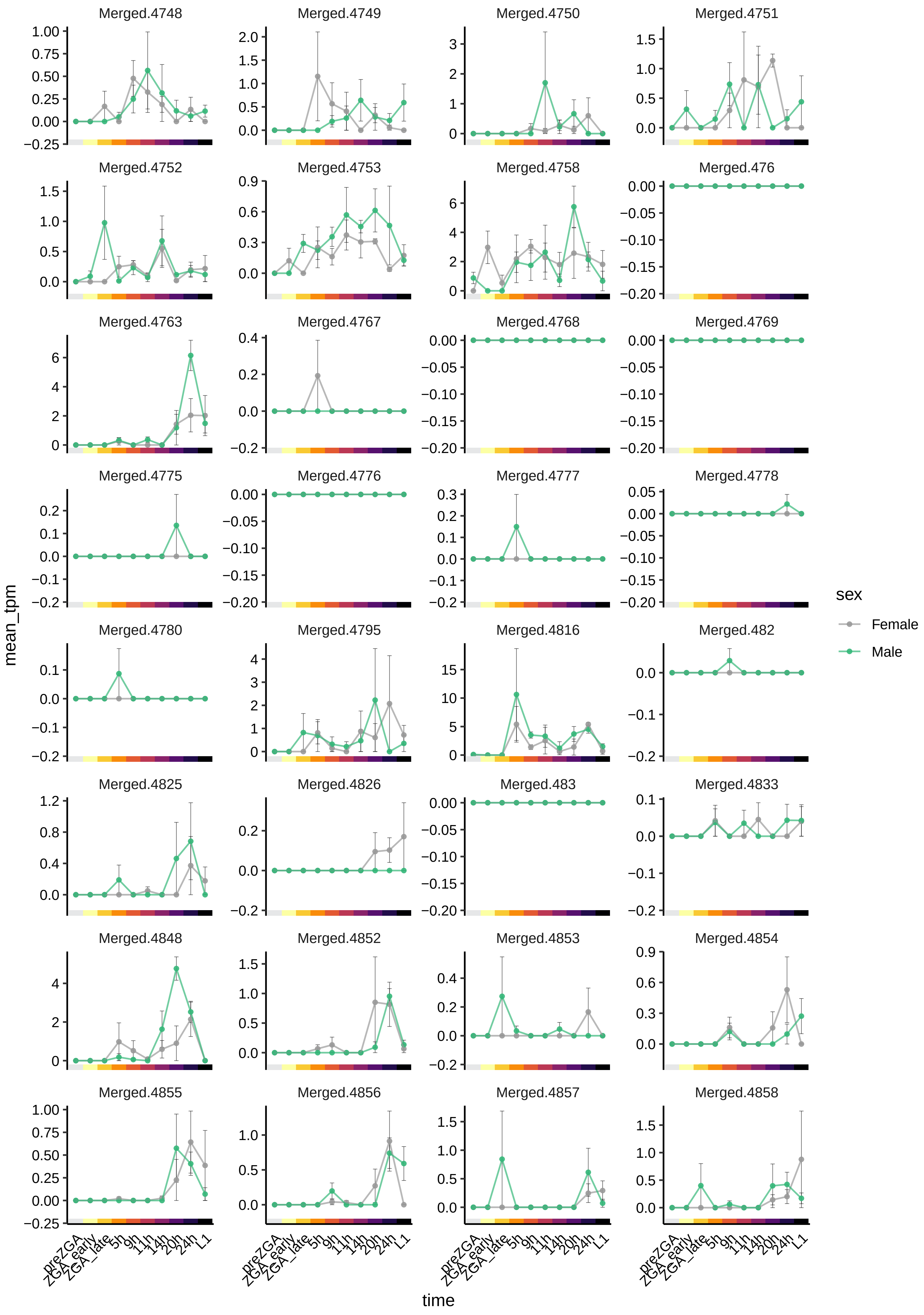

Merged.4748
Merged.4749
Merged.4750
Merged.4751
Merged.4752
Merged.4753
Merged.4758
Merged.476
Merged.4763
Merged.4767
Merged.4768
Merged.4769
Merged.4775
Merged.4776
Merged.4777
Merged.4778
Merged.4780
Merged.4795
Merged.4816
Merged.482
Merged.4825
Merged.4826
Merged.483
Merged.4833
Merged.4848
Merged.4852
Merged.4853
Merged.4854
Merged.4855
Merged.4856
Merged.4857
Merged.4858
mean_tpm
time
preZGA
ZGA_early
ZGA_late
5h
9h
11h
14h
20h
24h
L1
sex
Female
Male

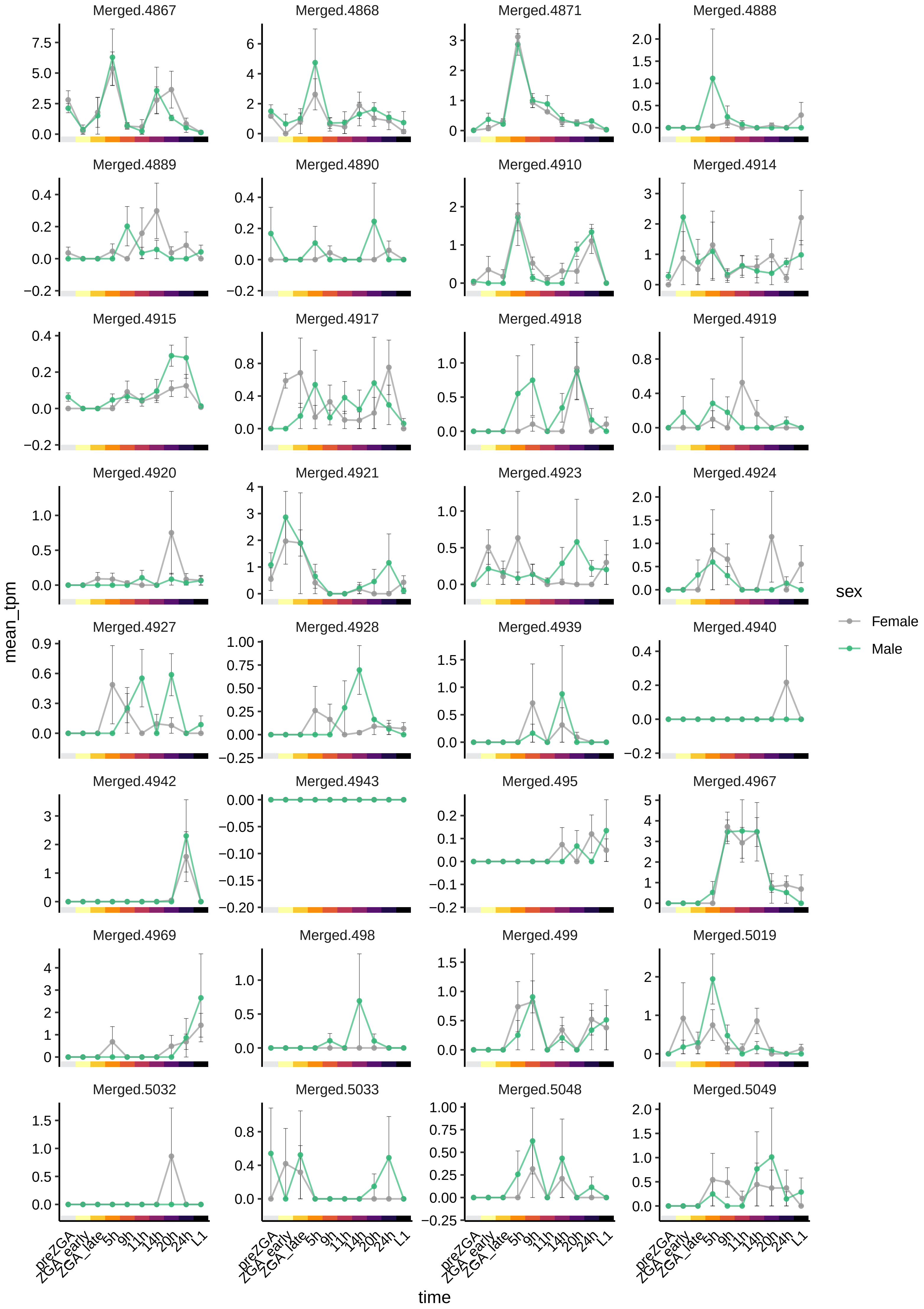
Merged.4867
Merged.4868
Merged.4871
Merged.4888
Merged.4889
Merged.4890
Merged.4910
Merged.4914
Merged.4915
Merged.4917
Merged.4918
Merged.4919
Merged.4920
Merged.4921
Merged.4923
Merged.4924
Merged.4927
Merged.4928
Merged.4939
Merged.4940
Merged.4942
Merged.4943
Merged.495
Merged.4967
Merged.4969
Merged.498
Merged.499
Merged.5019
Merged.5032
Merged.5033
Merged.5048
Merged.5049
mean_tpm
time
preZGA
ZGA_early
ZGA_late
5h
9h
11h
14h
20h
24h
L1
sex
Female
Male

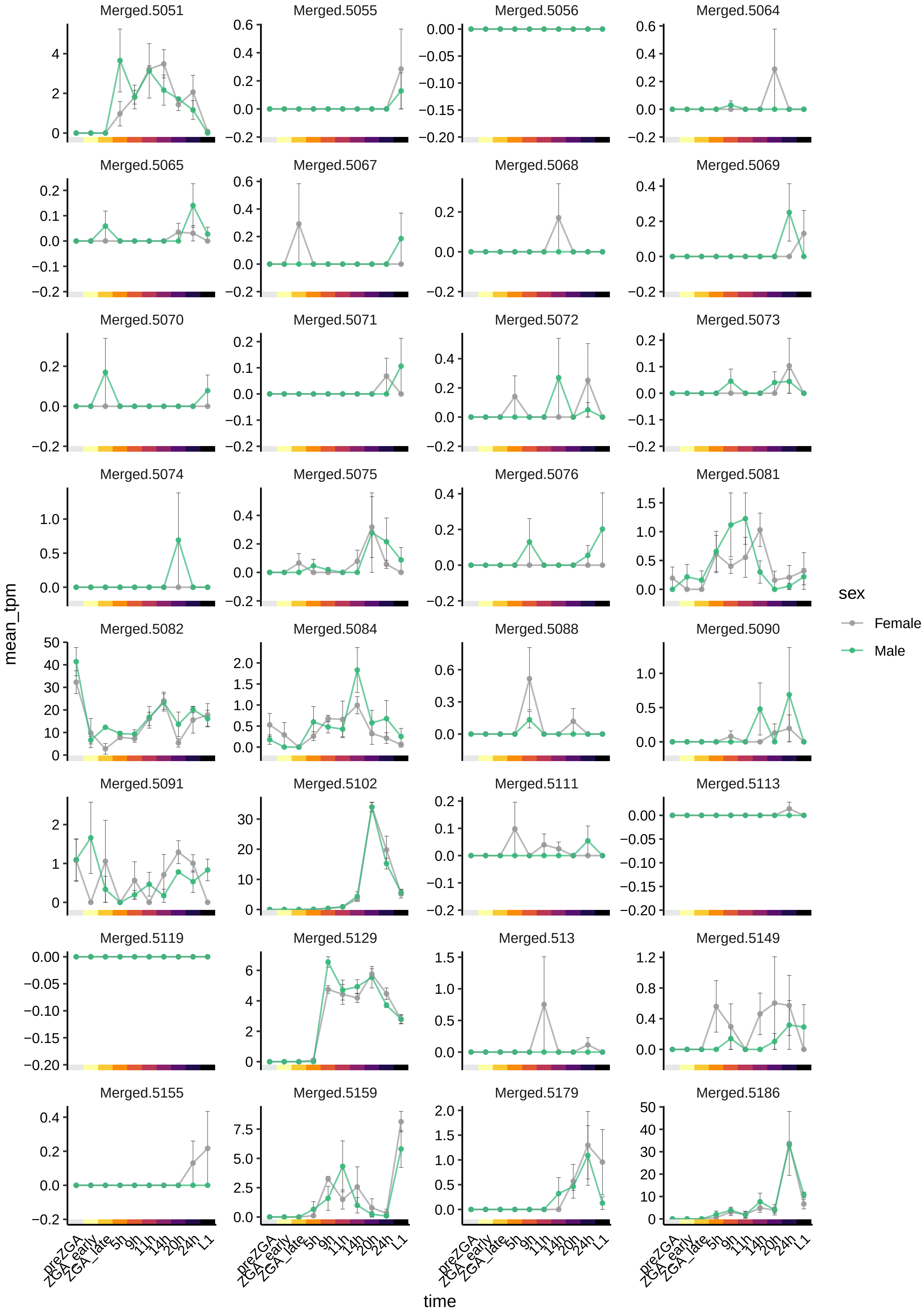
Merged.5051
Merged.5055
Merged.5056
Merged.5064
Merged.5065
Merged.5067
Merged.5068
Merged.5069
Merged.5070
Merged.5071
Merged.5072
Merged.5073
Merged.5074
Merged.5075
Merged.5076
Merged.5081
Merged.5082
Merged.5084
Merged.5088
Merged.5090
Merged.5091
Merged.5102
Merged.5111
Merged.5113
Merged.5119
Merged.5129
Merged.513
Merged.5149
Merged.5155
Merged.5159
Merged.5179
Merged.5186
mean_tpm
time
preZGA
ZGA_early
ZGA_late
5h
9h
11h
14h
20h
24h
L1
sex
Female
Male

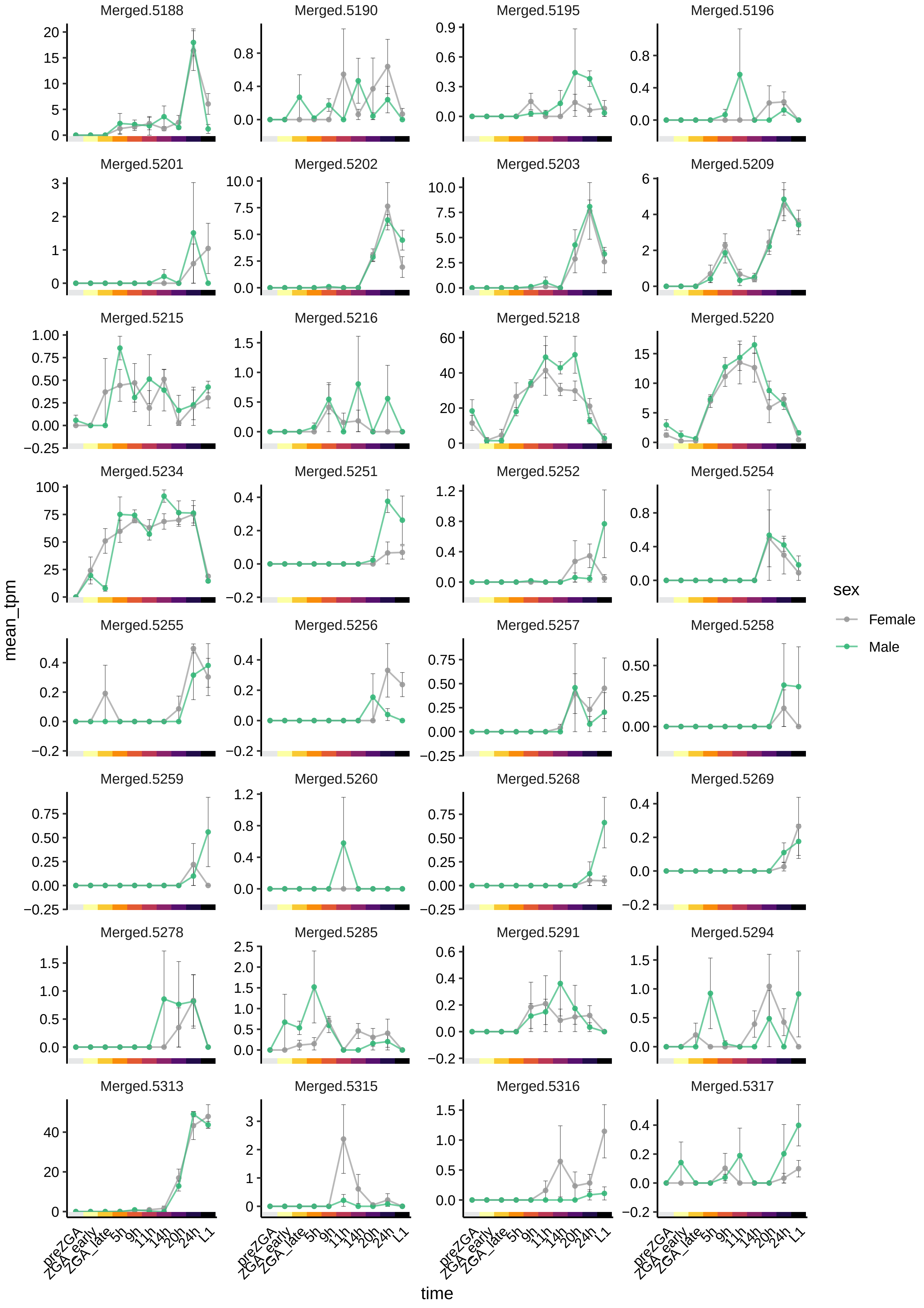

Merged.5188
Merged.5190
Merged.5195
Merged.5196
Merged.5201
Merged.5202
Merged.5203
Merged.5209
Merged.5215
Merged.5216
Merged.5218
Merged.5220
Merged.5234
Merged.5251
Merged.5252
Merged.5254
Merged.5255
Merged.5256
Merged.5257
Merged.5258
Merged.5259
Merged.5260
Merged.5268
Merged.5269
Merged.5278
Merged.5285
Merged.5291
Merged.5294
Merged.5313
Merged.5315
Merged.5316
Merged.5317
mean_tpm
time
preZGA
ZGA_early
ZGA_late
5h
9h
11h
14h
20h
24h
L1
sex
Female
Male

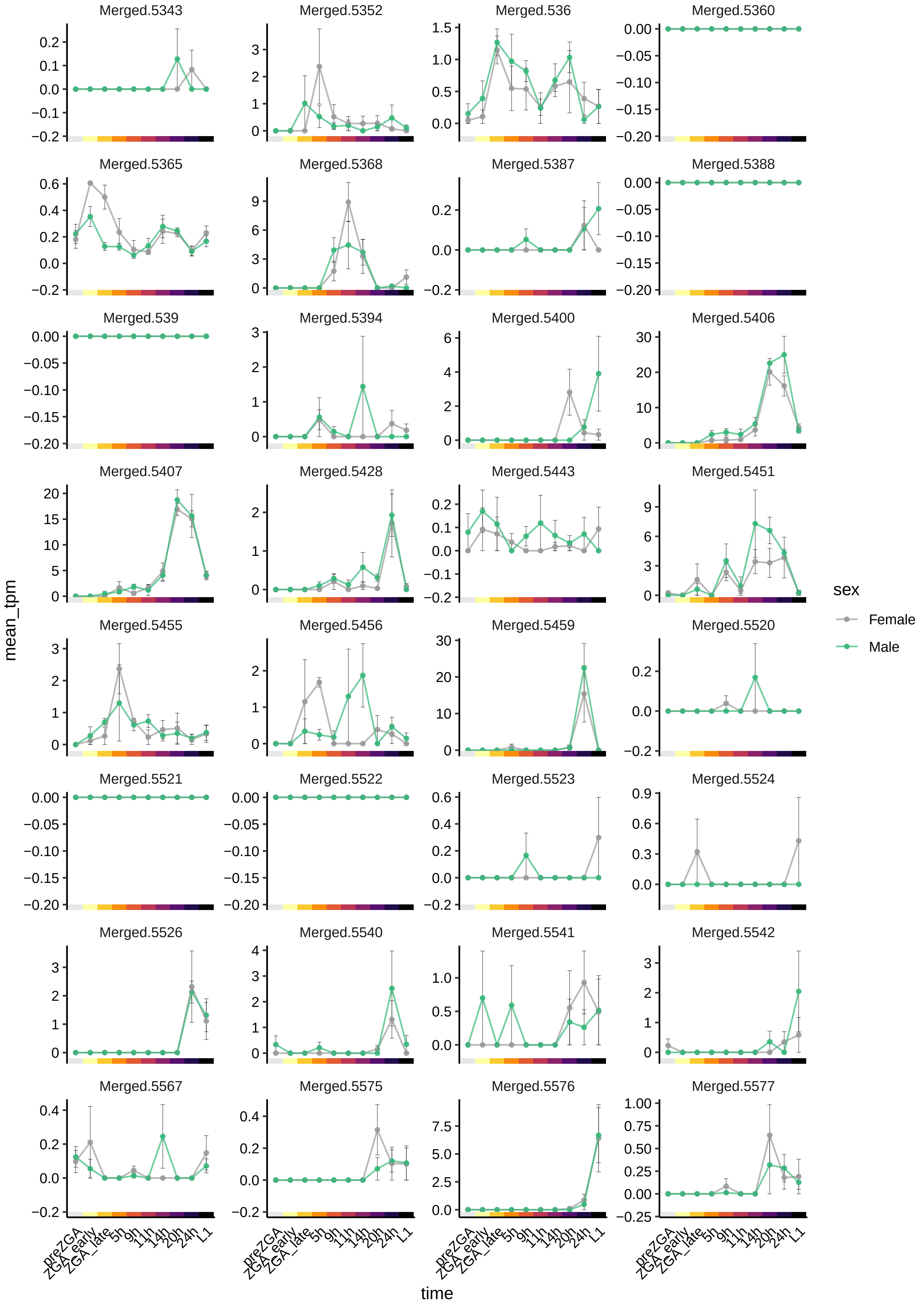

Merged.5343
Merged.5352
Merged.536
Merged.5360
Merged.5365
Merged.5368
Merged.5387
Merged.5388
Merged.539
Merged.5394
Merged.5400
Merged.5406
Merged.5407
Merged.5428
Merged.5443
Merged.5451
Merged.5455
Merged.5456
Merged.5459
Merged.5520
Merged.5521
Merged.5522
Merged.5523
Merged.5524
Merged.5526
Merged.5540
Merged.5541
Merged.5542
Merged.5567
Merged.5575
Merged.5576
Merged.5577
mean_tpm
time
preZGA
ZGA_early
ZGA_late
5h
9h
11h
14h
20h
24h
L1
sex
Female
Male

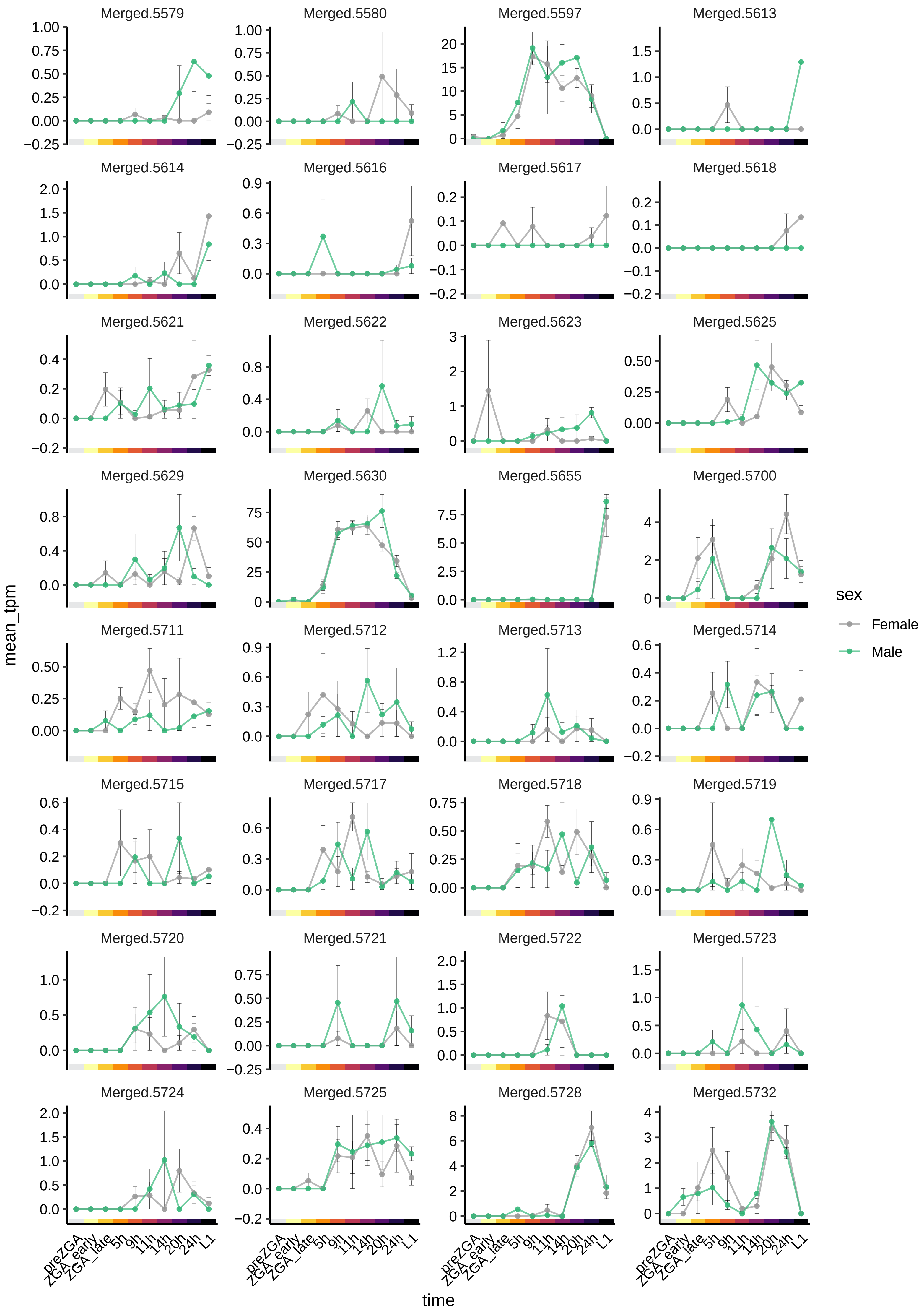

Merged.5579
Merged.5580
Merged.5597
Merged.5613
Merged.5614
Merged.5616
Merged.5617
Merged.5618
Merged.5621
Merged.5622
Merged.5623
Merged.5625
Merged.5629
Merged.5630
Merged.5655
Merged.5700
Merged.5711
Merged.5712
Merged.5713
Merged.5714
Merged.5715
Merged.5717
Merged.5718
Merged.5719
Merged.5720
Merged.5721
Merged.5722
Merged.5723
Merged.5724
Merged.5725
Merged.5728
Merged.5732
mean_tpm
time
preZGA
ZGA_early
ZGA_late
5h
9h
11h
14h
20h
24h
L1
sex
Female
Male

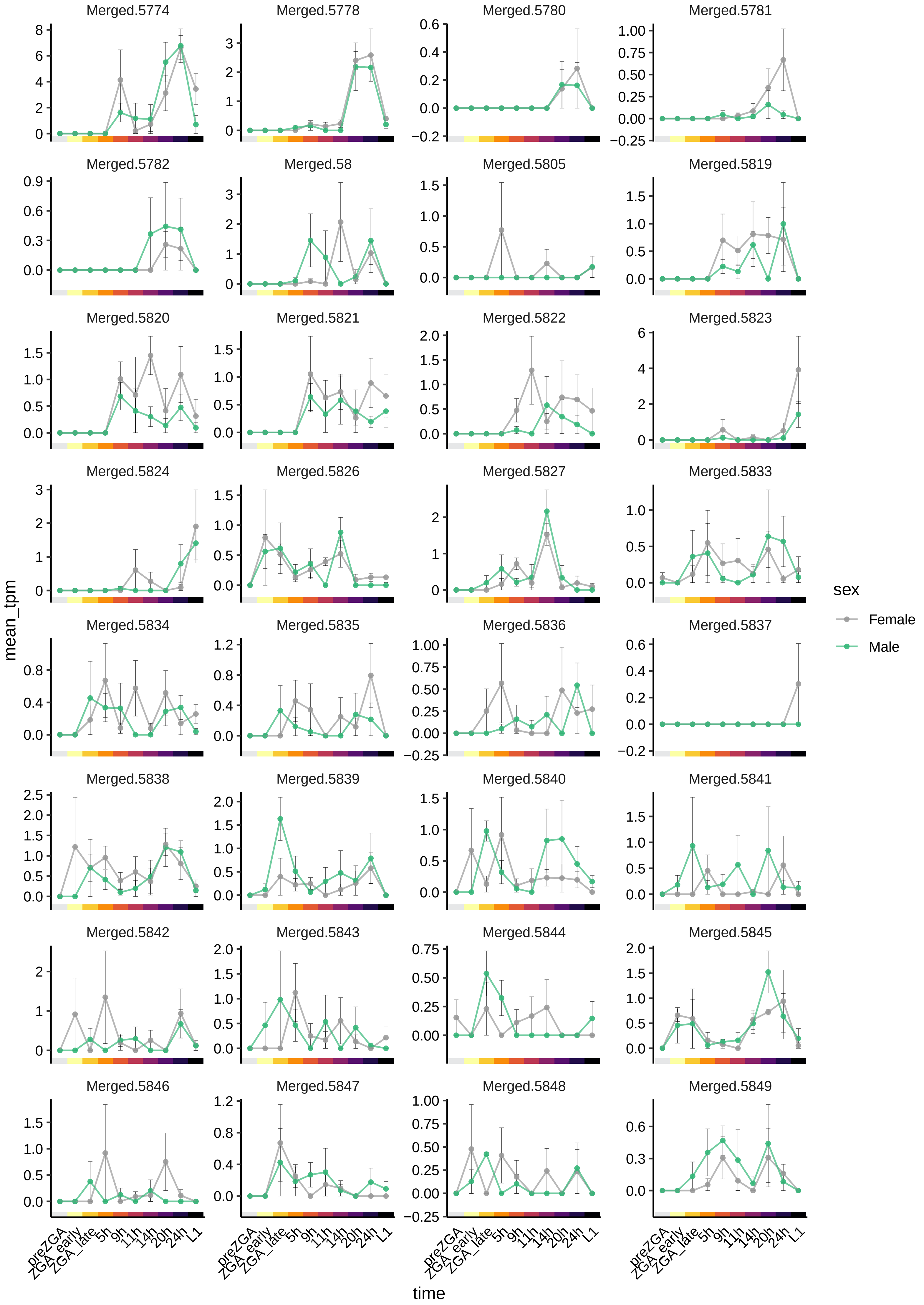

Merged.5774
Merged.5778
Merged.5780
Merged.5781
Merged.5782
Merged.58
Merged.5805
Merged.5819
Merged.5820
Merged.5821
Merged.5822
Merged.5823
Merged.5824
Merged.5826
Merged.5827
Merged.5833
Merged.5834
Merged.5835
Merged.5836
Merged.5837
Merged.5838
Merged.5839
Merged.5840
Merged.5841
Merged.5842
Merged.5843
Merged.5844
Merged.5845
Merged.5846
Merged.5847
Merged.5848
Merged.5849
mean_tpm
time
preZGA
ZGA_early
ZGA_late
5h
9h
11h
14h
20h
24h
L1
sex
Female
Male

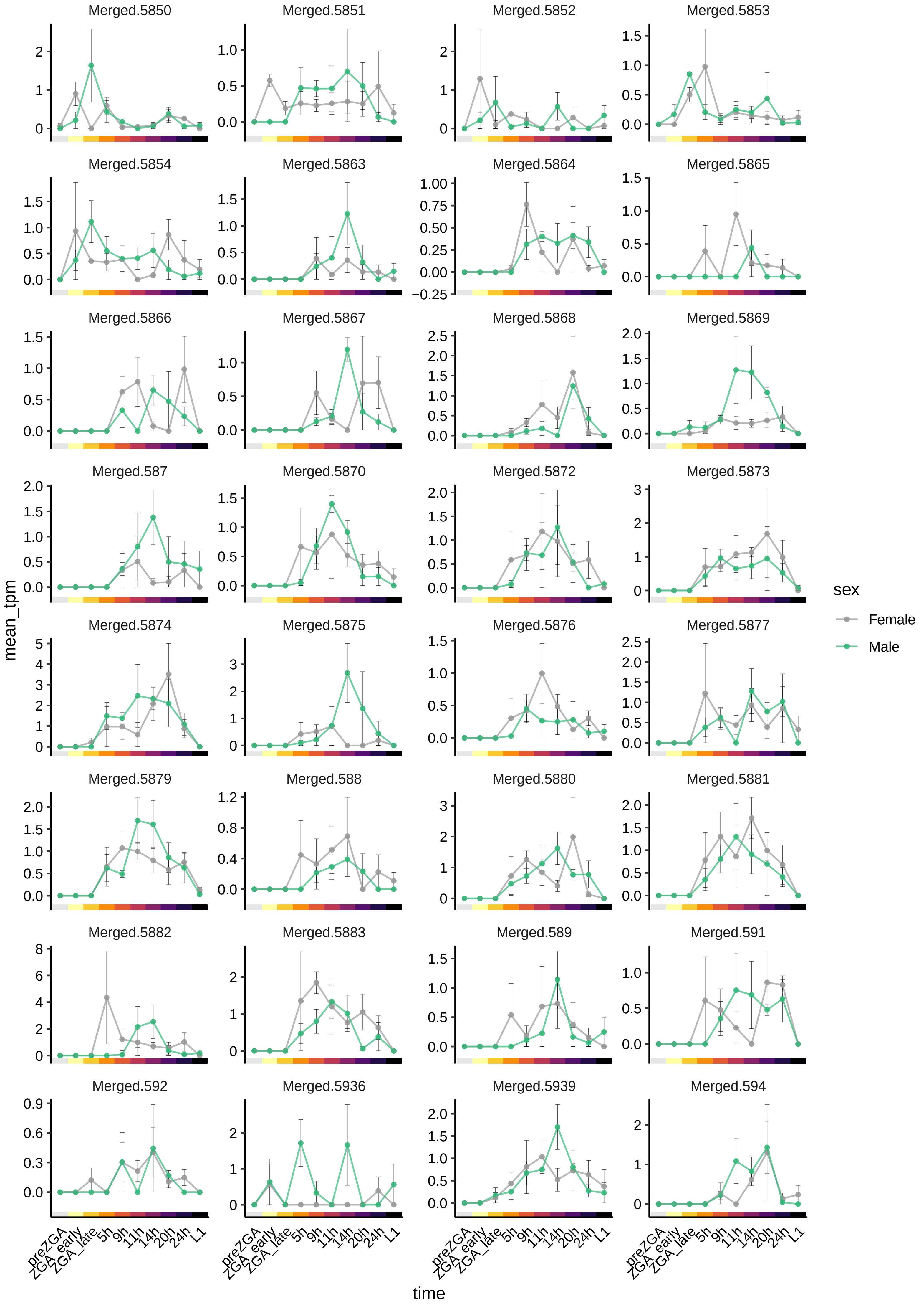

Merged.5850
Merged.5851
Merged.5852
Merged.5853
Merged.5854
Merged.5863
Merged.5864
Merged.5865
Merged.5866
Merged.5867
Merged.5868
Merged.5869
Merged.587
Merged.5870
Merged.5872
Merged.5873
Merged.5874
Merged.5875
Merged.5876
Merged.5877
Merged.5879
Merged.588
Merged.5880
Merged.5881
Merged.5882
Merged.5883
Merged.589
Merged.591
Merged.592
Merged.5936
Merged.5939
Merged.594
mean_tpm
time
preZGA
ZGA_early
ZGA_late
5h
9h
11h
14h
20h
24h
L1
sex
Female
Male

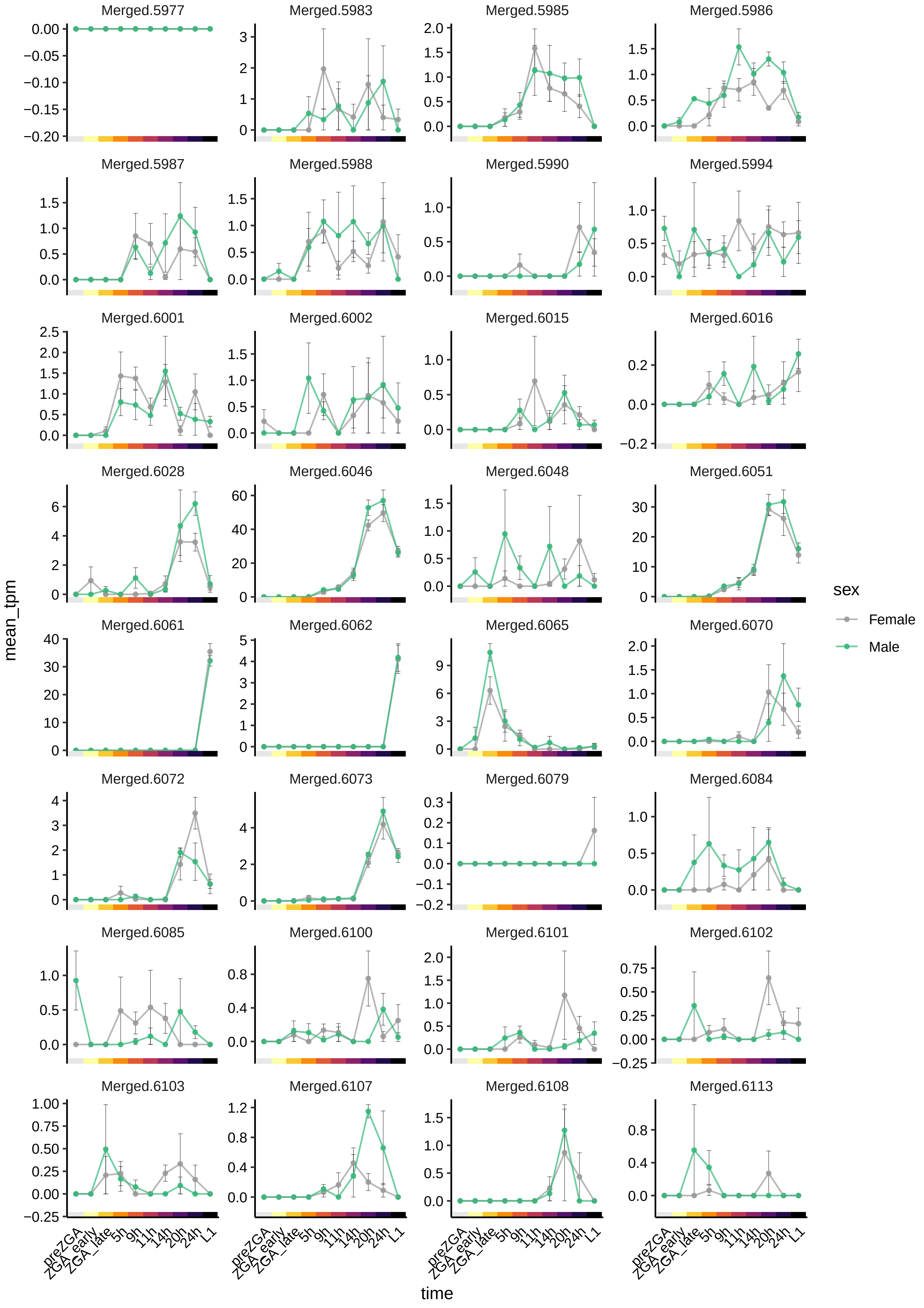
Merged.5977
Merged.5983
Merged.5985
Merged.5986
Merged.5987
Merged.5988
Merged.5990
Merged.5994
Merged.6001
Merged.6002
Merged.6015
Merged.6016
Merged.6028
Merged.6046
Merged.6048
Merged.6051
Merged.6061
Merged.6062
Merged.6065
Merged.6070
Merged.6072
Merged.6073
Merged.6079
Merged.6084
Merged.6085
Merged.6100
Merged.6101
Merged.6102
Merged.6103
Merged.6107
Merged.6108
Merged.6113
mean_tpm
time
preZGA
ZGA_early
ZGA_late
5h
9h
11h
14h
20h
24h
L1
sex
Female
Male

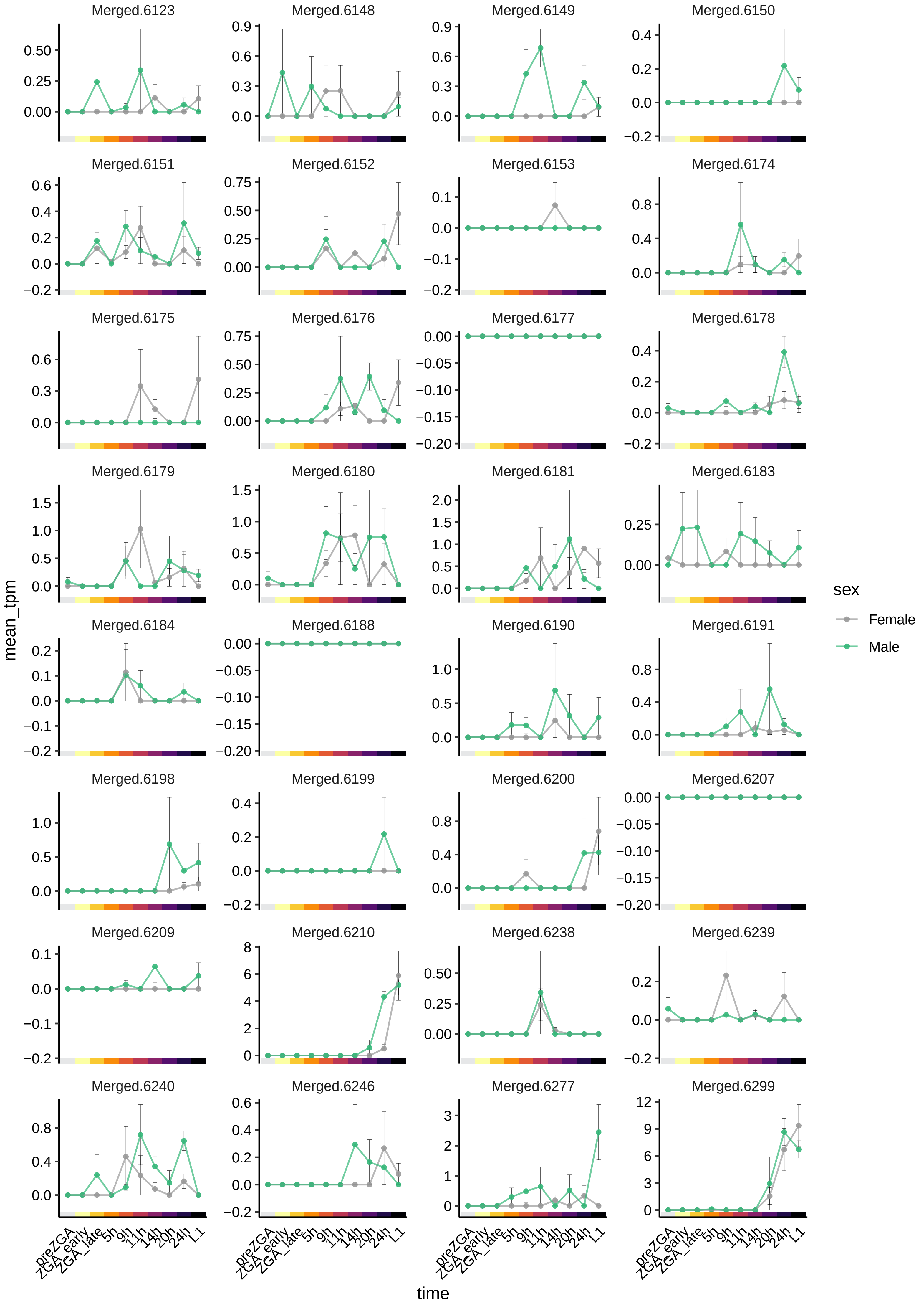

Merged.6123
Merged.6148
Merged.6149
Merged.6150
Merged.6151
Merged.6152
Merged.6153
Merged.6174
Merged.6175
Merged.6176
Merged.6177
Merged.6178
Merged.6179
Merged.6180
Merged.6181
Merged.6183
Merged.6184
Merged.6188
Merged.6190
Merged.6191
Merged.6198
Merged.6199
Merged.6200
Merged.6207
Merged.6209
Merged.6210
Merged.6238
Merged.6239
Merged.6240
Merged.6246
Merged.6277
Merged.6299
mean_tpm
time
preZGA
ZGA_early
ZGA_late
5h
9h
11h
14h
20h
24h
L1
sex
Female
Male

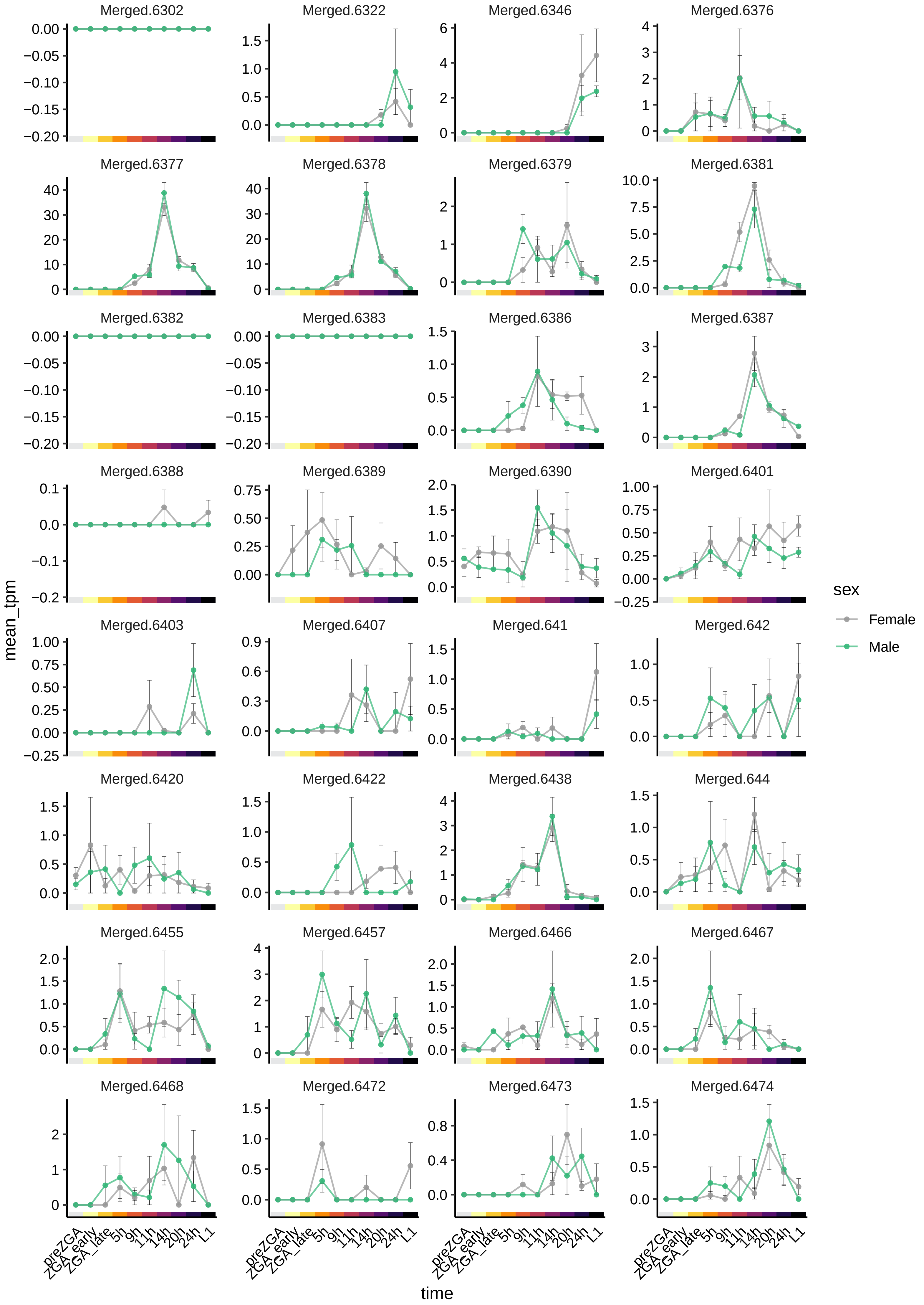

Merged.6302
Merged.6322
Merged.6346
Merged.6376
Merged.6377
Merged.6378
Merged.6379
Merged.6381
Merged.6382
Merged.6383
Merged.6386
Merged.6387
Merged.6388
Merged.6389
Merged.6390
Merged.6401
Merged.6403
Merged.6407
Merged.641
Merged.642
Merged.6420
Merged.6422
Merged.6438
Merged.644
Merged.6455
Merged.6457
Merged.6466
Merged.6467
Merged.6468
Merged.6472
Merged.6473
Merged.6474
mean_tpm
time
preZGA
ZGA_early
ZGA_late
5h
9h
11h
14h
20h
24h
L1
sex
Female
Male

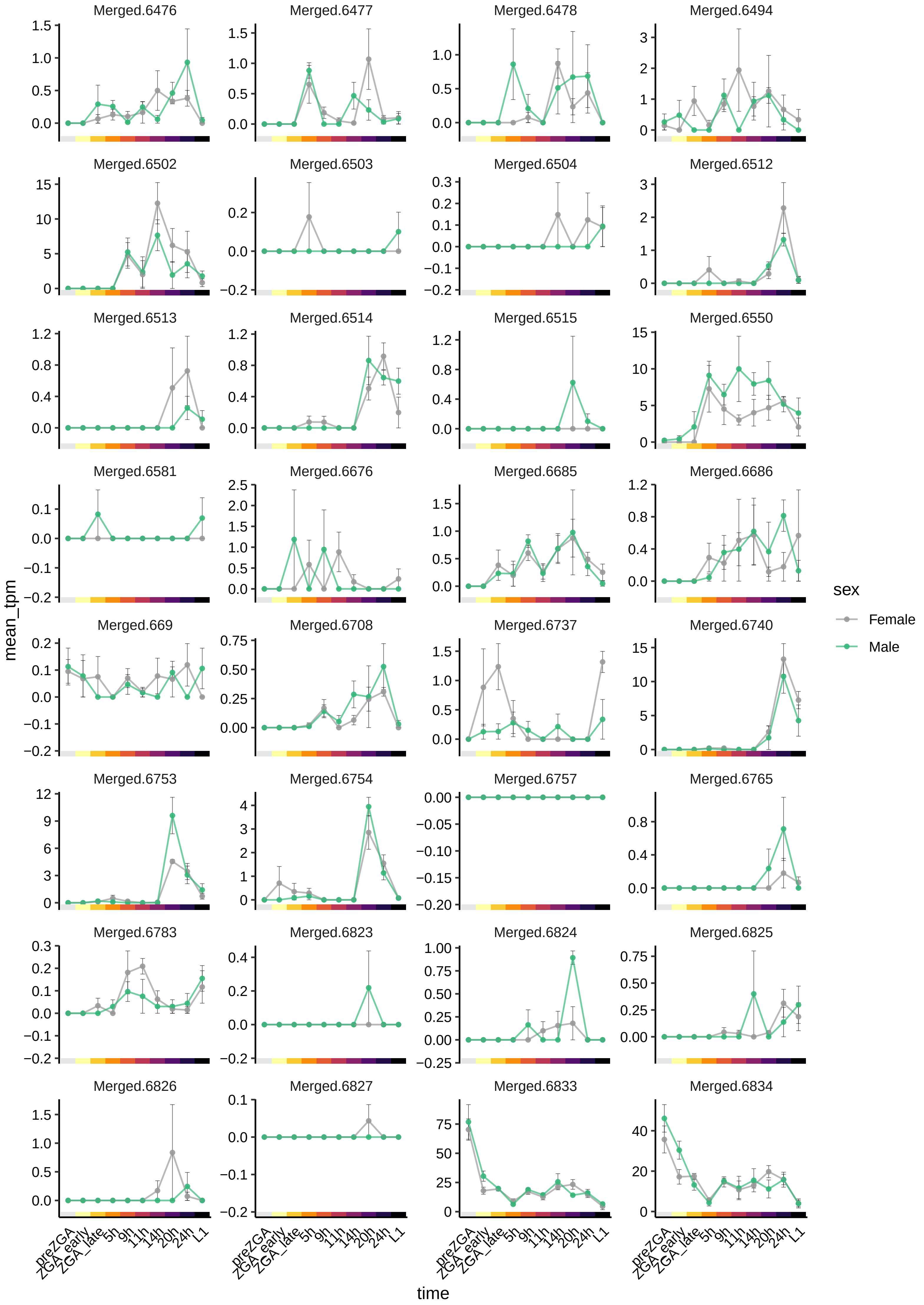
Merged.6476
Merged.6477
Merged.6478
Merged.6494
Merged.6502
Merged.6503
Merged.6504
Merged.6512
Merged.6513
Merged.6514
Merged.6515
Merged.6550
Merged.6581
Merged.6676
Merged.6685
Merged.6686
Merged.669
Merged.6708
Merged.6737
Merged.6740
Merged.6753
Merged.6754
Merged.6757
Merged.6765
Merged.6783
Merged.6823
Merged.6824
Merged.6825
Merged.6826
Merged.6827
Merged.6833
Merged.6834
mean_tpm
time
preZGA
ZGA_early
ZGA_late
5h
9h
11h
14h
20h
24h
L1
sex
Female
Male

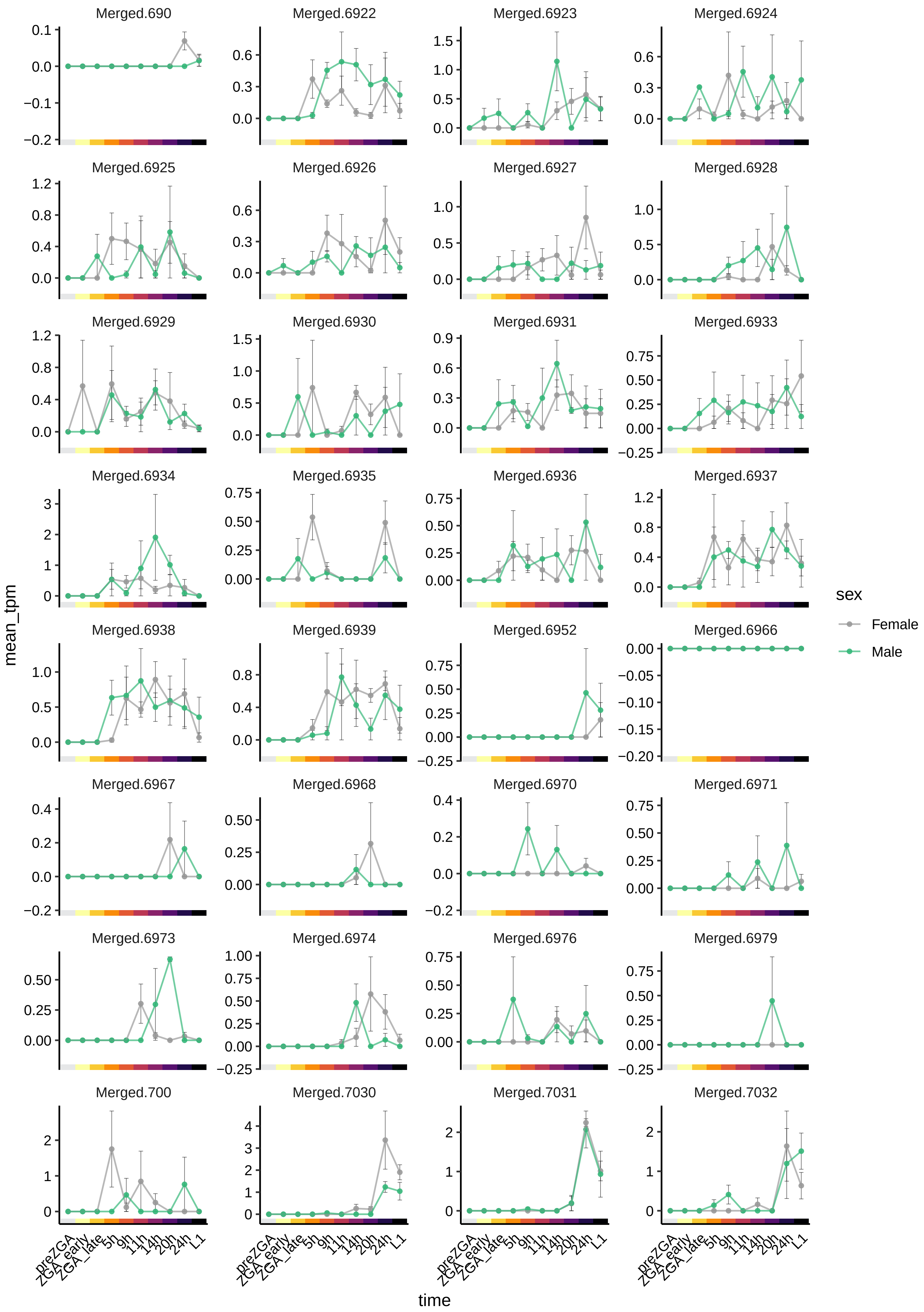

Merged.690
Merged.6922
Merged.6923
Merged.6924
Merged.6925
Merged.6926
Merged.6927
Merged.6928
Merged.6929
Merged.6930
Merged.6931
Merged.6933
Merged.6934
Merged.6935
Merged.6936
Merged.6937
Merged.6938
Merged.6939
Merged.6952
Merged.6966
Merged.6967
Merged.6968
Merged.6970
Merged.6971
Merged.6973
Merged.6974
Merged.6976
Merged.6979
Merged.700
Merged.7030
Merged.7031
Merged.7032
mean_tpm
time
preZGA
ZGA_early
ZGA_late
5h
9h
11h
14h
20h
24h
L1
sex
Female
Male

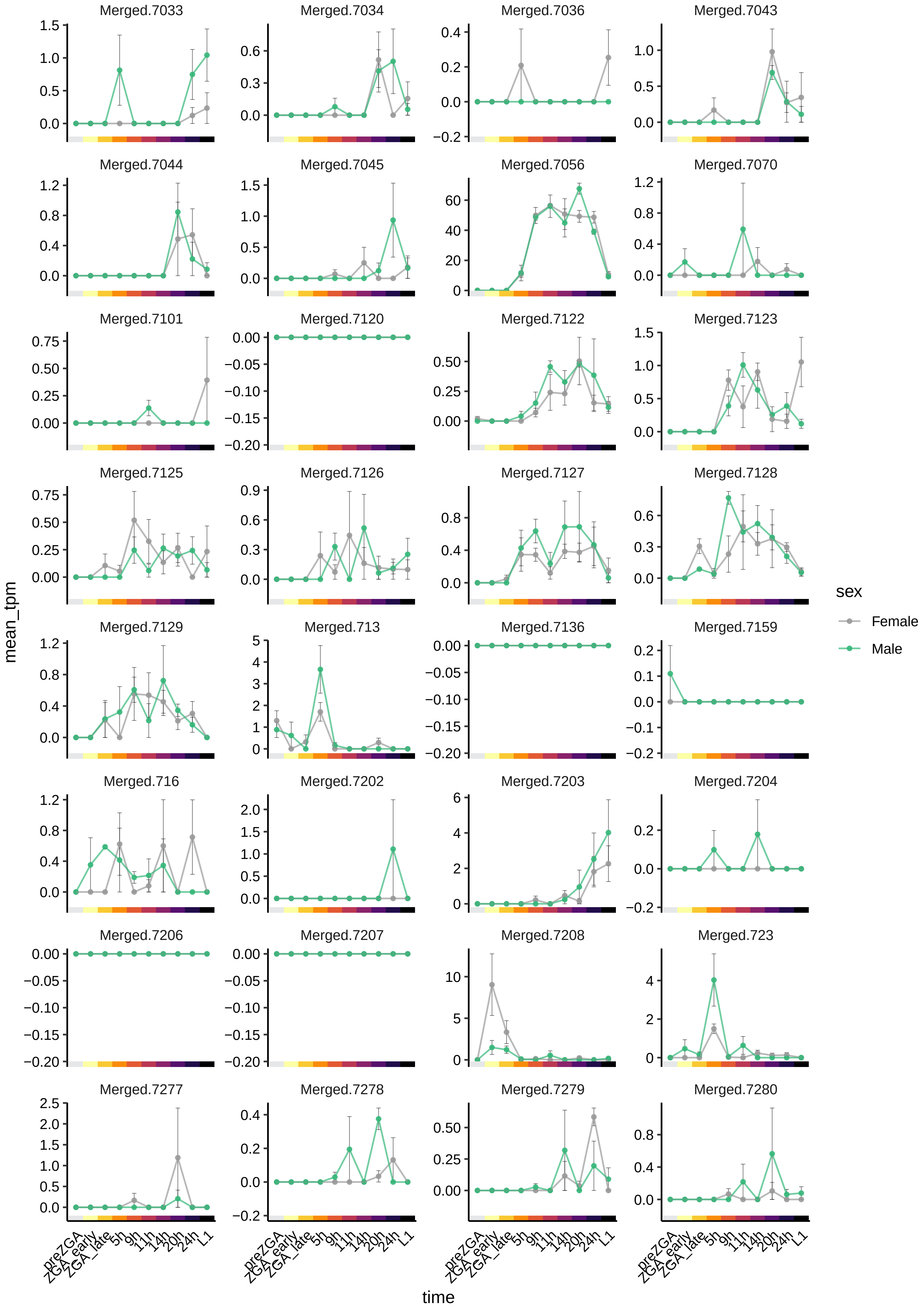

Merged.7033
Merged.7034
Merged.7036
Merged.7043
Merged.7044
Merged.7045
Merged.7056
Merged.7070
Merged.7101
Merged.7120
Merged.7122
Merged.7123
Merged.7125
Merged.7126
Merged.7127
Merged.7128
Merged.7129
Merged.713
Merged.7136
Merged.7159
Merged.716
Merged.7202
Merged.7203
Merged.7204
Merged.7206
Merged.7207
Merged.7208
Merged.723
Merged.7277
Merged.7278
Merged.7279
Merged.7280
mean_tpm
preZGA
ZGA_early
ZGA_late
5h
9h
11h
14h
20h
24h
L1
time
sex
Female
Male

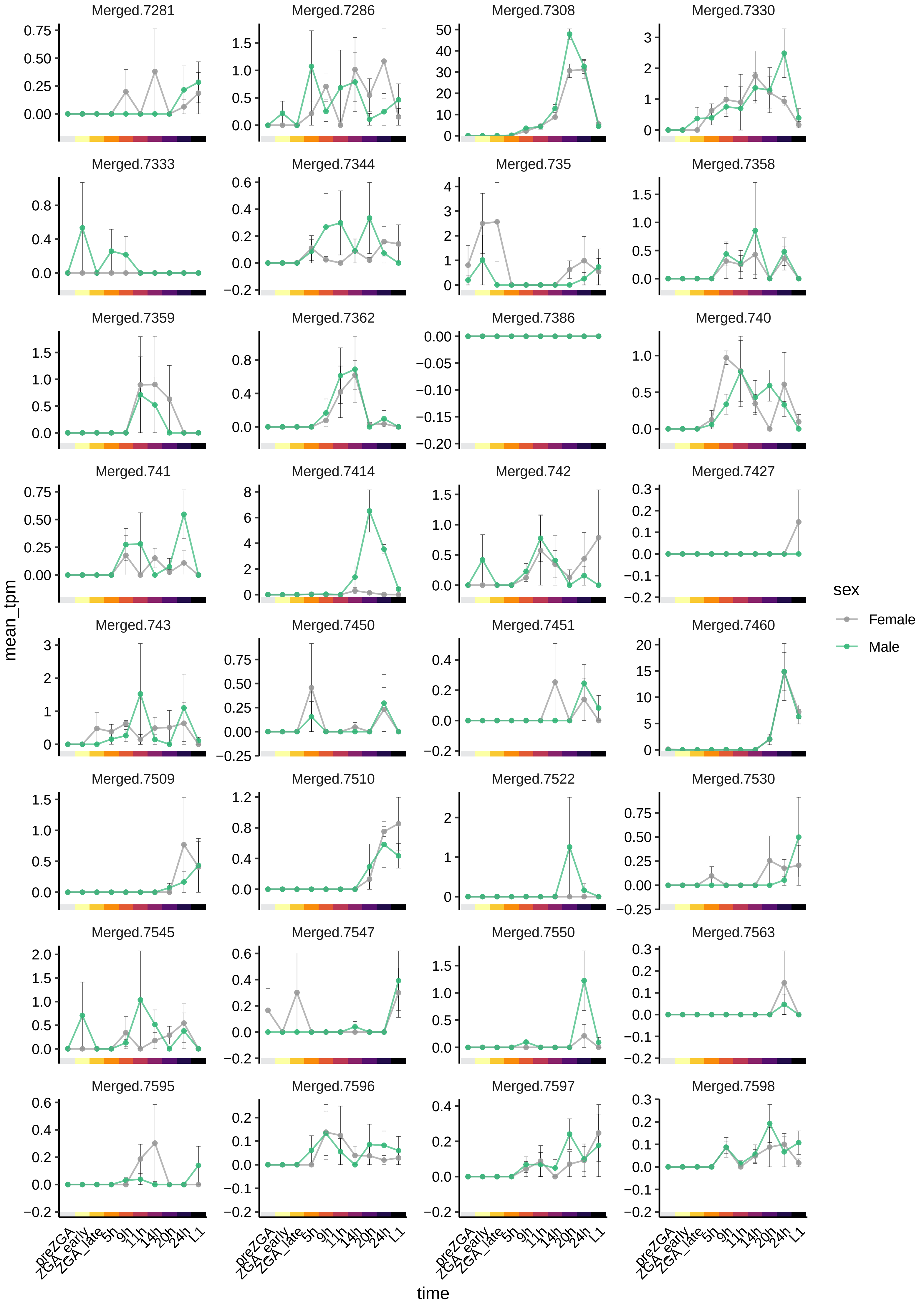

Merged.7281
Merged.7286
Merged.7308
Merged.7330
Merged.7333
Merged.7344
Merged.735
Merged.7358
Merged.7359
Merged.7362
Merged.7386
Merged.740
Merged.741
Merged.7414
Merged.742
Merged.7427
Merged.743
Merged.7450
Merged.7451
Merged.7460
Merged.7509
Merged.7510
Merged.7522
Merged.7530
Merged.7545
Merged.7547
Merged.7550
Merged.7563
Merged.7595
Merged.7596
Merged.7597
Merged.7598
mean_tpm
time
preZGA
ZGA_early
ZGA_late
5h
9h
11h
14h
20h
24h
L1
sex
Female
Male

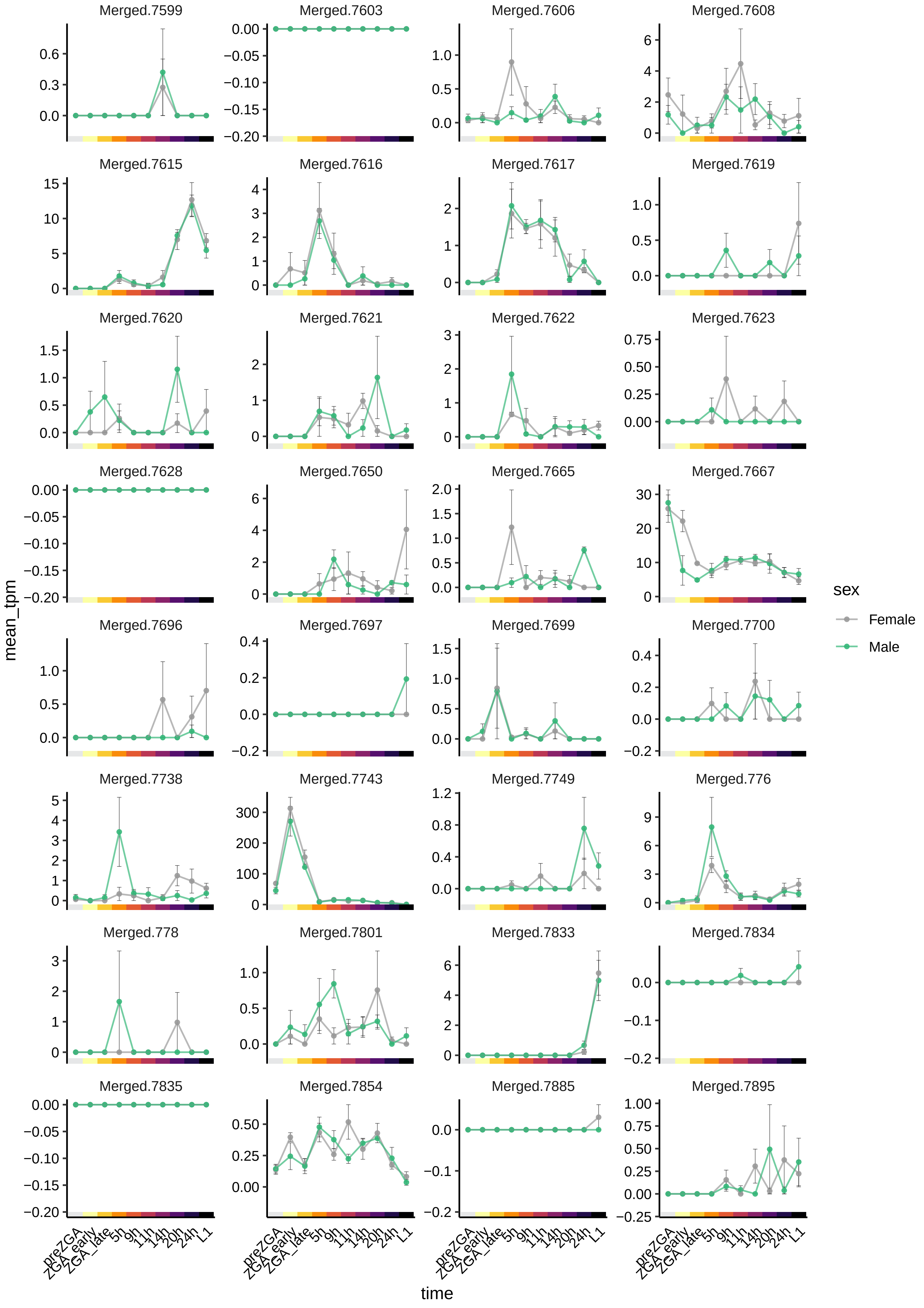
Merged.7599
Merged.7603
Merged.7606
Merged.7608
Merged.7615
Merged.7616
Merged.7617
Merged.7619
Merged.7620
Merged.7621
Merged.7622
Merged.7623
Merged.7628
Merged.7650
Merged.7665
Merged.7667
Merged.7696
Merged.7697
Merged.7699
Merged.7700
Merged.7738
Merged.7743
Merged.7749
Merged.776
Merged.778
Merged.7801
Merged.7833
Merged.7834
Merged.7835
Merged.7854
Merged.7885
Merged.7895
mean_tpm
time
preZGA
ZGA_early
ZGA_late
5h
9h
11h
14h
20h
24h
L1
sex
Female
Male

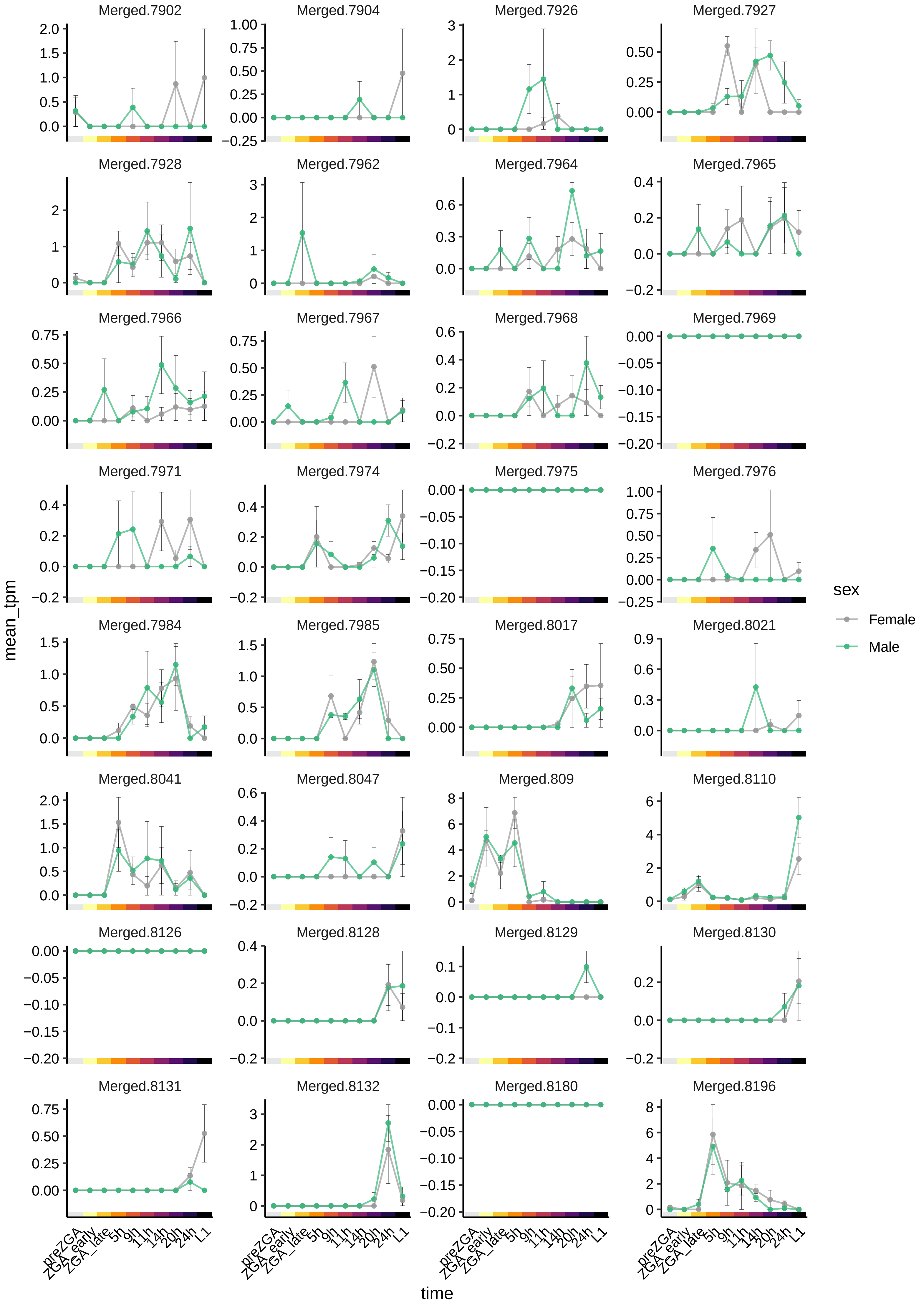
Merged.7902
Merged.7904
Merged.7926
Merged.7927
Merged.7928
Merged.7962
Merged.7964
Merged.7965
Merged.7966
Merged.7967
Merged.7968
Merged.7969
Merged.7971
Merged.7974
Merged.7975
Merged.7976
Merged.7984
Merged.7985
Merged.8017
Merged.8021
Merged.8041
Merged.8047
Merged.809
Merged.8110
Merged.8126
Merged.8128
Merged.8129
Merged.8130
Merged.8131
Merged.8132
Merged.8180
Merged.8196
mean_tpm
time
preZGA
ZGA_early
ZGA_late
5h
9h
11h
14h
20h
24h
L1
sex
Female
Male

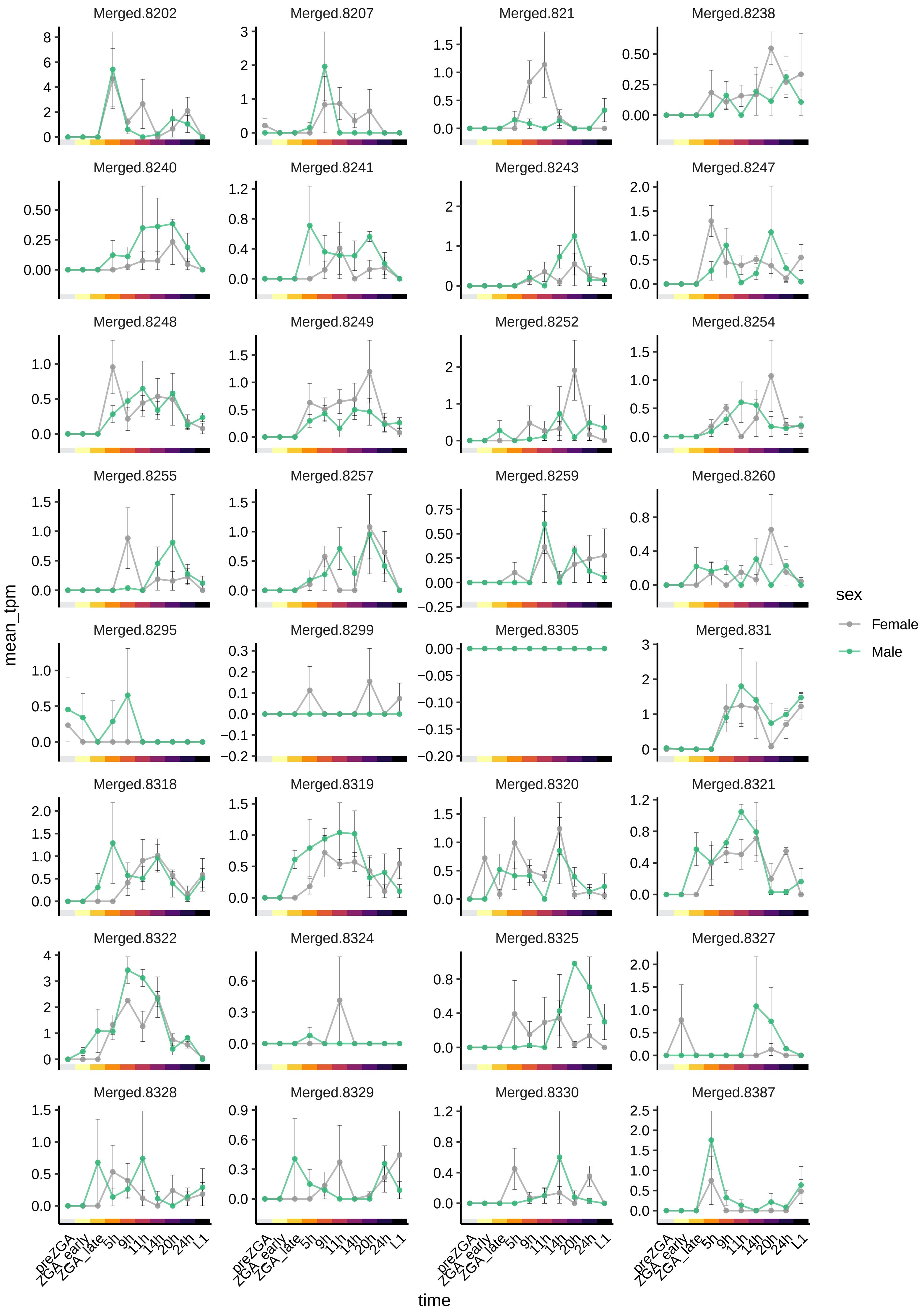

Merged.8202
Merged.8207
Merged.821
Merged.8238
Merged.8240
Merged.8241
Merged.8243
Merged.8247
Merged.8248
Merged.8249
Merged.8252
Merged.8254
Merged.8255
Merged.8257
Merged.8259
Merged.8260
Merged.8295
Merged.8299
Merged.8305
Merged.831
Merged.8318
Merged.8319
Merged.8320
Merged.8321
Merged.8322
Merged.8324
Merged.8325
Merged.8327
Merged.8328
Merged.8329
Merged.8330
Merged.8387
mean_tpm
time
preZGA
ZGA_early
ZGA_late
5h
9h
11h
14h
20h
24h
L1
sex
Female
Male

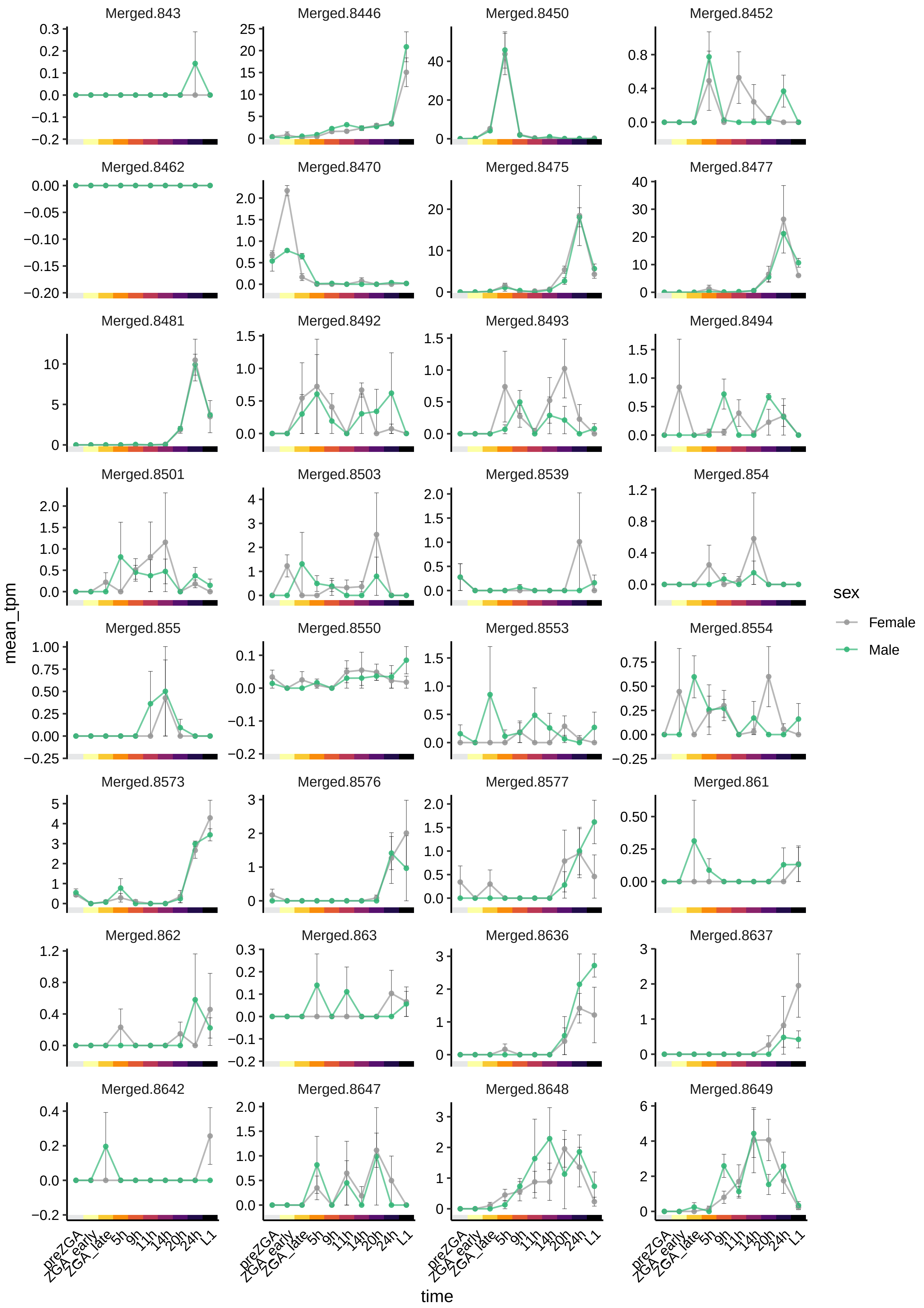
Merged.843
Merged.8446
Merged.8450
Merged.8452
Merged.8462
Merged.8470
Merged.8475
Merged.8477
Merged.8481
Merged.8492
Merged.8493
Merged.8494
Merged.8501
Merged.8503
Merged.8539
Merged.854
Merged.855
Merged.8550
Merged.8553
Merged.8554
Merged.8573
Merged.8576
Merged.8577
Merged.861
Merged.862
Merged.863
Merged.8636
Merged.8637
Merged.8642
Merged.8647
Merged.8648
Merged.8649
mean_tpm
time
preZGA
ZGA_early
ZGA_late
5h
9h
11h
14h
20h
24h
L1
sex
Female
Male

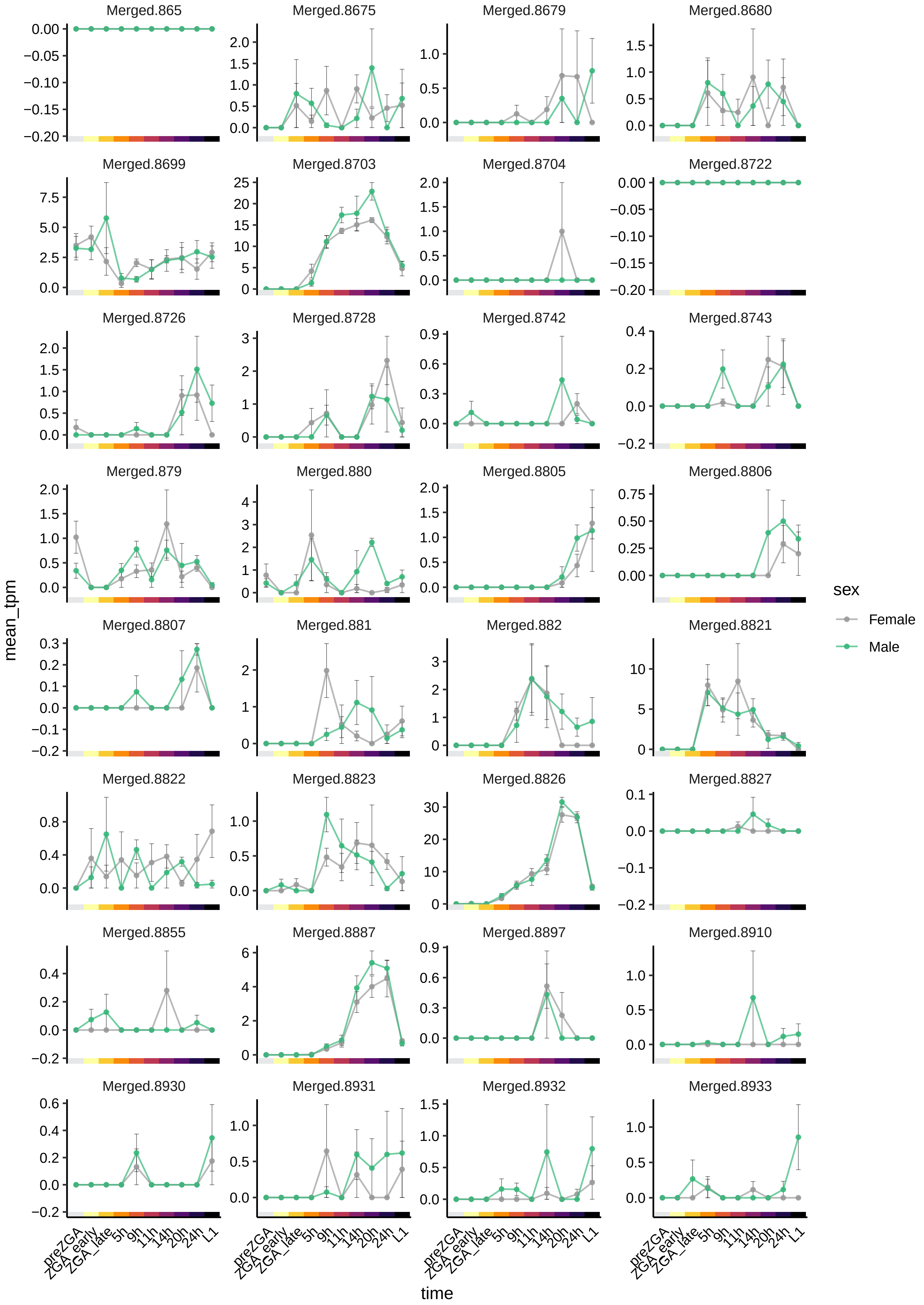

Merged.865
Merged.8675
Merged.8679
Merged.8680
Merged.8699
Merged.8703
Merged.8704
Merged.8722
Merged.8726
Merged.8728
Merged.8742
Merged.8743
Merged.879
Merged.880
Merged.8805
Merged.8806
Merged.8807
Merged.881
Merged.882
Merged.8821
Merged.8822
Merged.8823
Merged.8826
Merged.8827
Merged.8855
Merged.8887
Merged.8897
Merged.8910
Merged.8930
Merged.8931
Merged.8932
Merged.8933
mean_tpm
time
preZGA
ZGA_early
ZGA_late
5h
9h
11h
14h
20h
24h
L1
sex
Female
Male

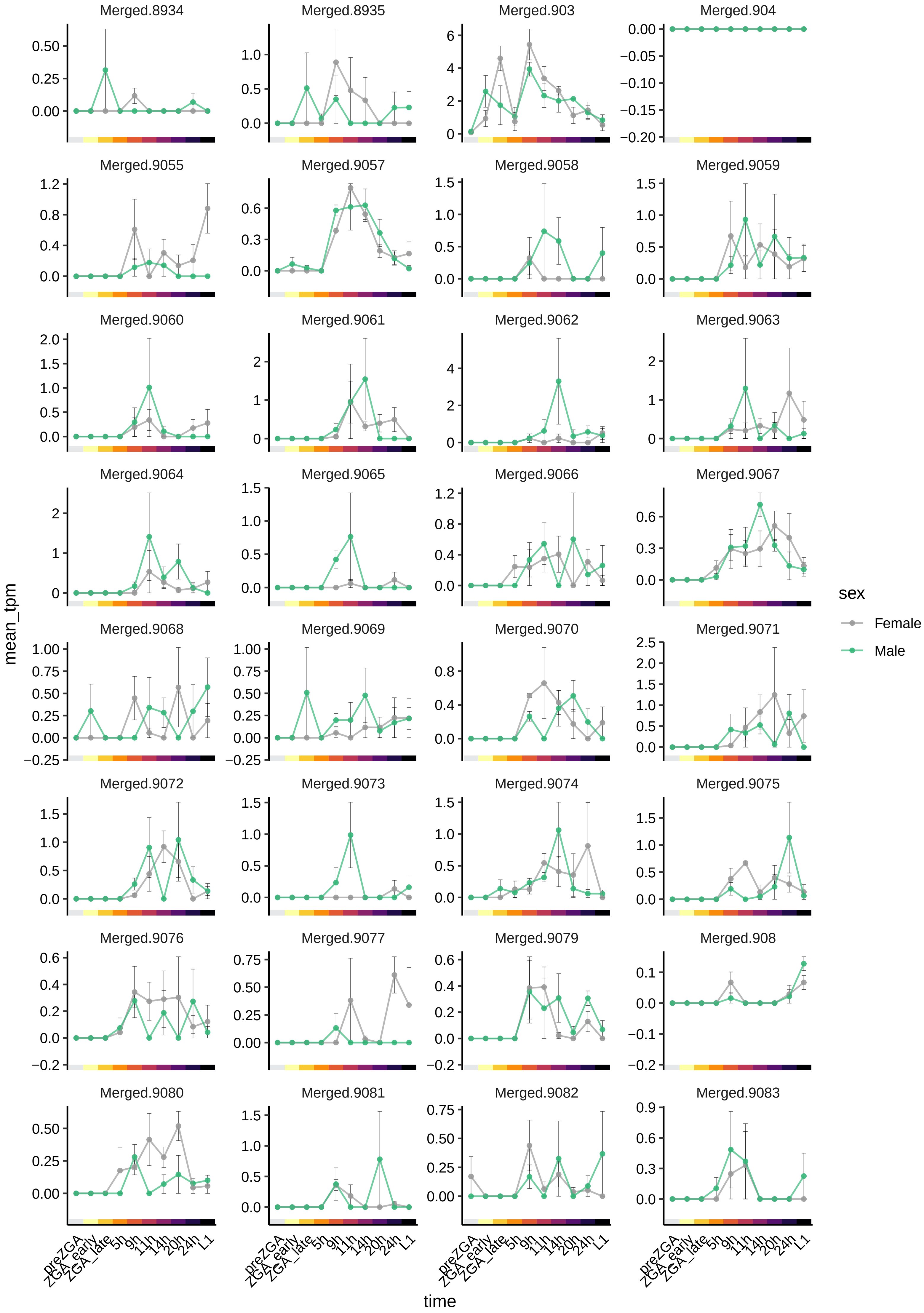

Merged.8934
Merged.8935
Merged.903
Merged.904
Merged.9055
Merged.9057
Merged.9058
Merged.9059
Merged.9060
Merged.9061
Merged.9062
Merged.9063
Merged.9064
Merged.9065
Merged.9066
Merged.9067
Merged.9068
Merged.9069
Merged.9070
Merged.9071
Merged.9072
Merged.9073
Merged.9074
Merged.9075
Merged.9076
Merged.9077
Merged.9079
Merged.908
Merged.9080
Merged.9081
Merged.9082
Merged.9083
mean_tpm
time
preZGA
ZGA_early
ZGA_late
5h
9h
11h
14h
20h
24h
L1
sex
Female
Male

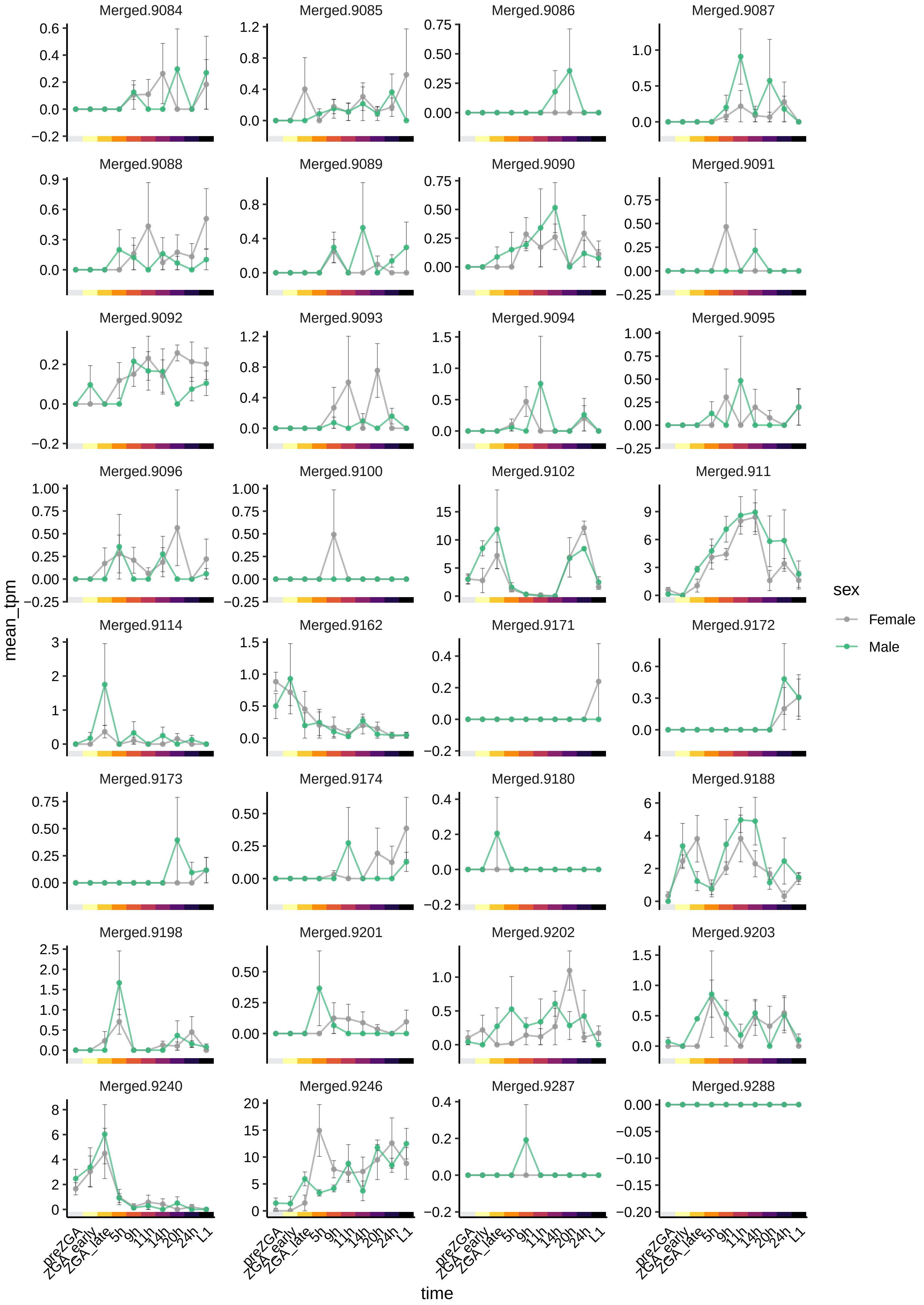
Merged.9084
Merged.9085
Merged.9086
Merged.9087
Merged.9088
Merged.9089
Merged.9090
Merged.9091
Merged.9092
Merged.9093
Merged.9094
Merged.9095
Merged.9096
Merged.9100
Merged.9102
Merged.911
Merged.9114
Merged.9162
Merged.9171
Merged.9172
Merged.9173
Merged.9174
Merged.9180
Merged.9188
Merged.9198
Merged.9201
Merged.9202
Merged.9203
Merged.9240
Merged.9246
Merged.9287
Merged.9288
mean_tpm
time
preZGA
ZGA_early
ZGA_late
5h
9h
11h
14h
20h
24h
L1
sex
Female
Male

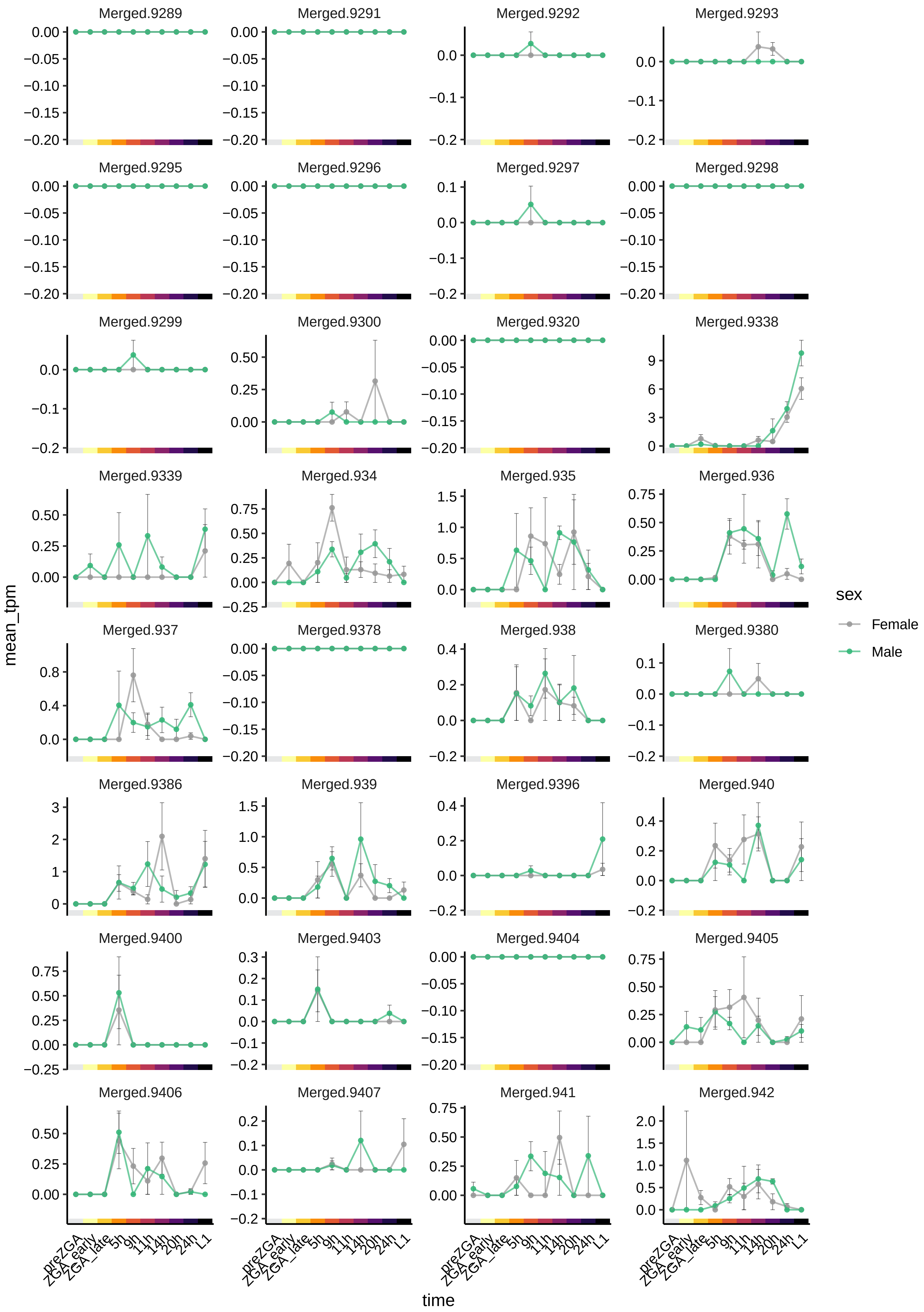

Merged.9289
Merged.9291
Merged.9292
Merged.9293
Merged.9295
Merged.9296
Merged.9297
Merged.9298
Merged.9299
Merged.9300
Merged.9320
Merged.9338
Merged.9339
Merged.934
Merged.935
Merged.936
Merged.937
Merged.9378
Merged.938
Merged.9380
Merged.9386
Merged.939
Merged.9396
Merged.940
Merged.9400
Merged.9403
Merged.9404
Merged.9405
Merged.9406
Merged.9407
Merged.941
Merged.942
mean_tpm
time
preZGA
ZGA_early
ZGA_late
5h
9h
11h
14h
20h
24h
L1
sex
Female
Male

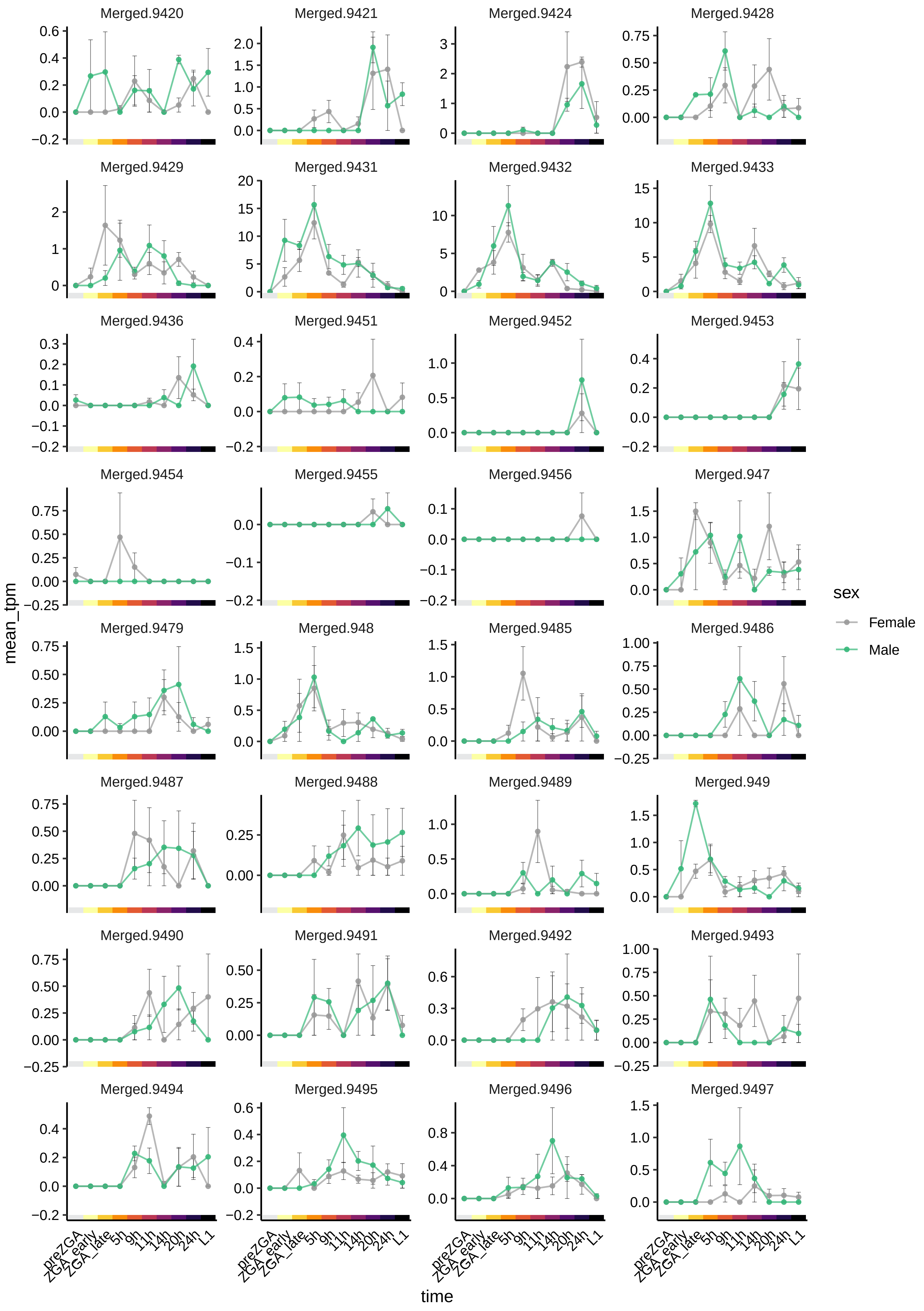

Merged.9420
Merged.9421
Merged.9424
Merged.9428
Merged.9429
Merged.9431
Merged.9432
Merged.9433
Merged.9436
Merged.9451
Merged.9452
Merged.9453
Merged.9454
Merged.9455
Merged.9456
Merged.947
Merged.9479
Merged.948
Merged.9485
Merged.9486
Merged.9487
Merged.9488
Merged.9489
Merged.949
Merged.9490
Merged.9491
Merged.9492
Merged.9493
Merged.9494
Merged.9495
Merged.9496
Merged.9497
mean_tpm
time
preZGA
ZGA_early
ZGA_late
5h
9h
11h
14h
20h
24h
L1
sex
Female
Male

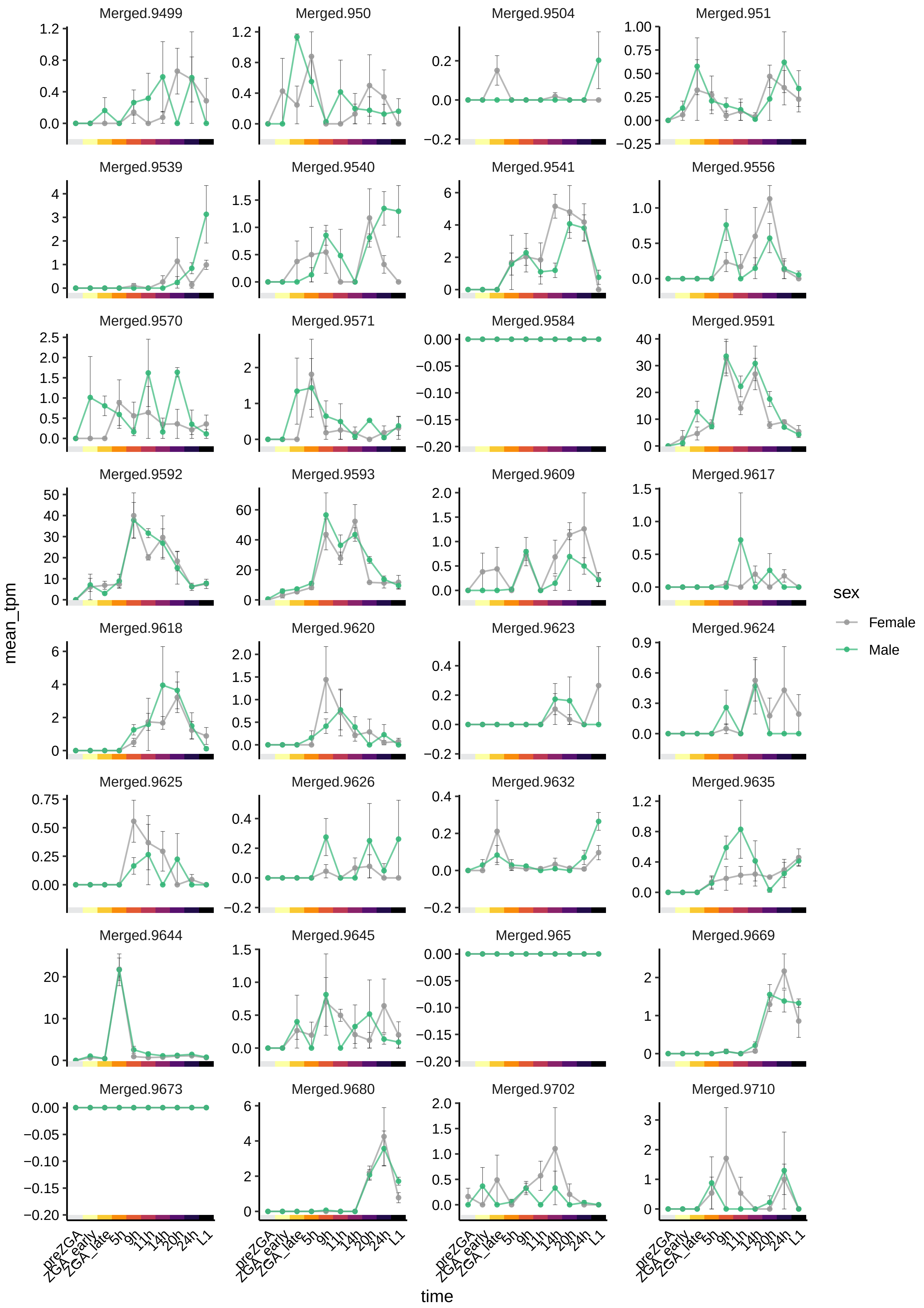
Merged.9499
Merged.950
Merged.9504
Merged.951
Merged.9539
Merged.9540
Merged.9541
Merged.9556
Merged.9570
Merged.9571
Merged.9584
Merged.9591
Merged.9592
Merged.9593
Merged.9609
Merged.9617
Merged.9618
Merged.9620
Merged.9623
Merged.9624
Merged.9625
Merged.9626
Merged.9632
Merged.9635
Merged.9644
Merged.9645
Merged.965
Merged.9669
Merged.9673
Merged.9680
Merged.9702
Merged.9710
mean_tpm
time
preZGA
ZGA_early
ZGA_late
5h
9h
11h
14h
20h
24h
L1
sex
Female
Male

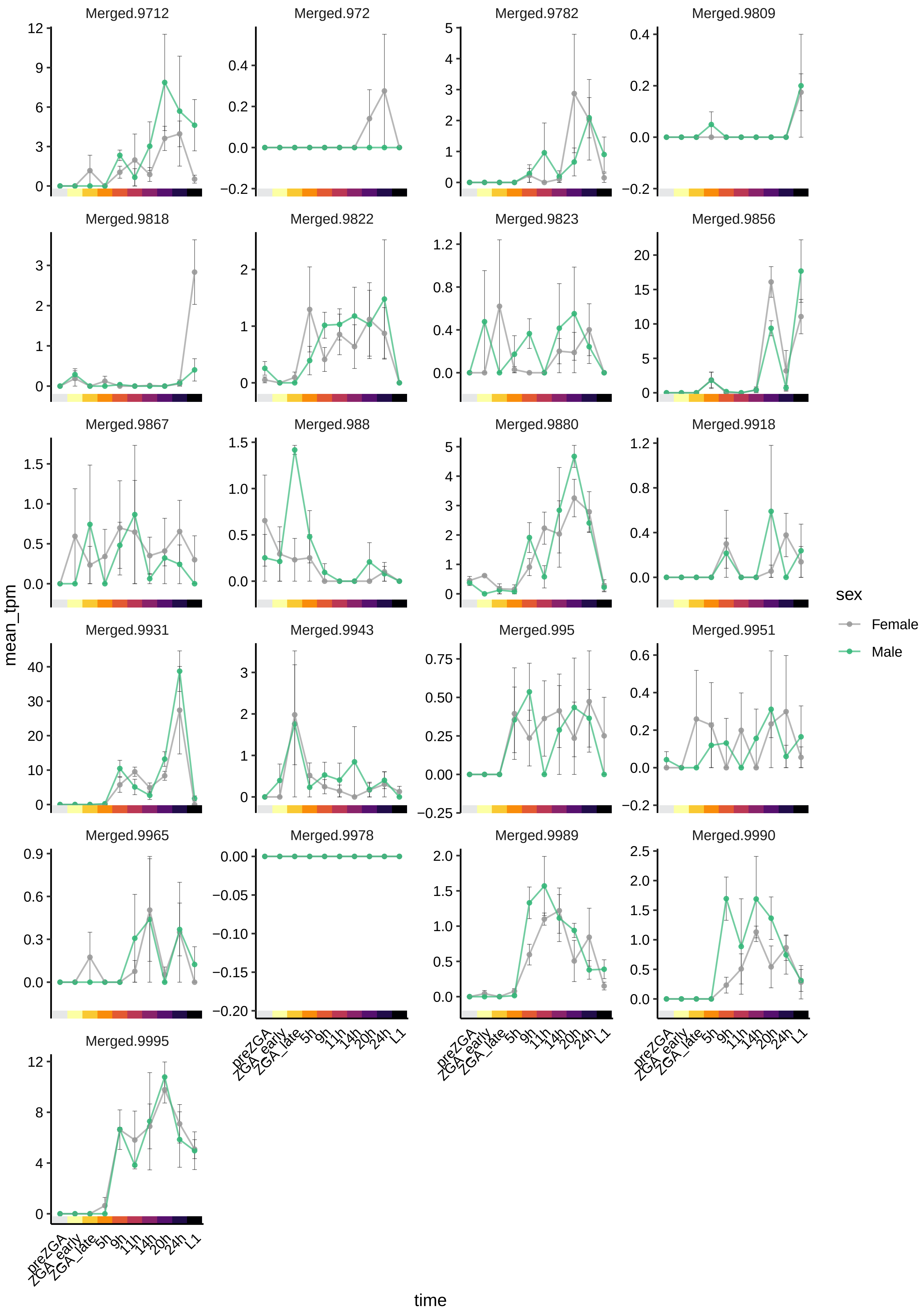
Merged.9712
Merged.972
Merged.9782
Merged.9809
Merged.9818
Merged.9822
Merged.9823
Merged.9856
Merged.9867
Merged.988
Merged.9880
Merged.9918
Merged.9931
Merged.9943
Merged.995
Merged.9951
Merged.9965
Merged.9978
Merged.9989
Merged.9990
Merged.9995
mean_tpm
time
preZGA
ZGA_early
ZGA_late
5h
9h
11h
14h
20h
24h
L1
sex
Female
Male
